# LysM Receptor Proteins are Required for Ectomycorrhizal Symbiosis in Poplar

**DOI:** 10.64898/2025.12.17.694584

**Authors:** K.M. Omenge, K. Deecke, T.B. Irving, T.A. Rush, E.S. Wallner, J. Keilwagen, M. Tiwari, S. Nagalla, S. Fort, C. Hallmann, M. Fladung, J.-M. Ané, S. Werner

## Abstract

Successful symbiotic associations often rely on the perception of specific microbial signals by host plants through pattern recognition receptors. LysM receptor-like kinases are key receptors in both rhizobial and arbuscular mycorrhizal symbioses. In this study, we provide the first evidence that such LysM receptor-like kinases also control the establishment of ectomycorrhizal associations in poplar. Using CRISPR/Cas9-mediated genome editing, we knocked out three NFP-like candidate genes in various combinations. We examined their roles in the perception of the ectomycorrhizal fungus *Laccaria bicolor*, as well as chitin oligomers and lipo-chitooligosaccharides. We demonstrate that all three receptors are necessary for the recognition of symbiosis-related signals, calcium spiking, symbiotic gene expression, and ectomycorrhizal root phenotypes.

## Introduction

Most terrestrial angiosperms engage in a symbiotic association with mycorrhizal fungi forming a mutually beneficial endosymbiosis that has persisted for over 400 million years (Selosse et al., 2015). The colonization of plant roots by mycorrhizal fungi enhances host water and nutrient acquisition, particularly phosphorus and nitrogen, while simultaneously bolstering plant resilience to biotic and abiotic stresses (Parniske, 2008; Smith and Smith, 2011). Reciprocally, the host plant supplies fungi with photosynthetically derived organic compounds, primarily lipids and carbohydrates that are necessary for maintaining this symbiotic relationship (Solaimanand and Saito, 1997; Jiang et al., 2017). Ectomycorrhizal (EM) and arbuscular mycorrhizal (AM) fungi produce cell wall-derived signals which function as microbial-associated molecular patterns (MAMPs). MAMPs are recognized by the plant via plasma membrane-localized pattern recognition receptors (PRRs), which initiate downstream signaling pathways (Sánchez-Vallet et al., 2015; Madsen et al., 2003; Kaku et al., 2006). Among these PRRs, lysin motif (LysM) domain-containing receptor proteins perceive chitin-derived oligosaccharides, mediating both host immune responses and symbiotic signaling during plant-microbe interactions (Kaku et al., 2006; Gough and Cullimore, 2011; Khokhani et al., 2021). LysM domain containing receptor proteins can be divided in 3 classes: LysM-Receptor Like Kinases (LysM-RLKs) with functional kinase domain like the *Arabidopsis thaliana* CERK1 or *Medicago truncatula* LYK3 receptors belong to the LYK subfamily, whilst the LYR subfamily, like *M. truncatula* NFP, is characterized by phosphorylation-inactive kinase domains missing the P-rich loop (Petutschnig et al., 2010; Buendia et al., 2018). The third group (LYMs) are GPI-anchored LysM-Receptor Like Proteins. LYRs and LYMs are dependent on co-receptor formation with LYK proteins and it is hypothesized that LYRs and LYMs signaling is dependent on ligand triggered binding of a LYR or LYM to a LYK activating the LYK kinase domain (Buendia et al., 2018). Decorated chitin molecules of different length act as ligands for LysM receptors. The so-called nodulation (Nod) factors produced by rhizobia to initiate symbiosis with legumes are lipo-chitooligosaccharides (LCOs). Both LCOs and non-acylated COs have been shown to initiate Ca^2+^ spiking as part of the AM symbiosis (He et al., 2019). Activation of heteromeric receptor complexes by LCOs and/or COs in combination with nutrient status signals leads to the activation of the Common Symbiosis Signaling Pathway (CSSP) (Li et al., 2022), characterized by nuclear calcium spiking in root cells that is mediated by the CASTOR, POLLUX/DMI1 and cyclic nucleotide-gated channels CNGC15 cation channels and the MCA8 calcium pump (MacLean et al., 2017; Capoen et al., 2011; Kim et al., 2019). Ca^2+^ spiking within the nucleoplasm activates the calcium- and calmodulin-dependent protein kinase (CCaMK) which phosphorylates the transcription factor IPD3/CYCLOPS (Singh and Parniske, 2012; Messinese et al., 2007) to initiate changes in downstream gene expression needed for symbiosis. For rhizobia and AM fungi, this pathway is well described. For EM symbiosis a potential involvement of the CSSP was not known until Cope and co-workers demonstrated that the EM fungus *Laccaria bicolor* produces a diverse array of sulfated and non-sulfated LCOs that trigger Ca^2+^ spiking in the *Populus tremula x alba* hybrid also known as *Populus x canescens* (poplar) in a CASTOR/POLLUX dependent manner (Cope et al., 2019). Contrasting to that, gymnosperm *Pinus sp.* forms EM symbiosis but lacks crucial CSSP components (Garcia et al., 2015) while other EM fungi such as *Hebeloma cylindrosporum* produce LCOs capable of eliciting calcium spiking in poplar despite not being able to form EM associations with this host (R Cope et al., 2021) indicating that other MAMPs and PRRs such as the poplar G-type lectin receptor-like kinase PtLec1 also control the establishment of these EM associations (Labbé et al., 2019). The production of COs and LCOs by EM fungi and the role of CSSP genes in the establishment of EM associations (Cope et al., 2019; R Cope et al., 2021; Rush et al., 2020), led to the hypothesis that EM formation in woody plants might also rely on LysM receptors as crucial facilitators of symbiosis. To investigate this, we focused our experiments on three specific poplar LysM genes, which were named NFP-like (*NFP-like 1*, *NFP-like 3*, and *NFP-like 4*), due to their similarity to the *M. truncatula* NFP, a key player in AM and root nodule symbiosis (Bruegmann et al., 2019). Knockouts using CRISPR/Cas9 mediated gene editing of *NFP-like 1*, *NFP-like 3*, and *NFP-like 4* in poplar resulted in a significant impairment in the induction of calcium spiking, altered downstream gene expression as well as EM associations.

## Results

### NFP-like receptors are required for induction of the CSSP in poplar

Calcium oscillations were monitored in ∼ 1 cm roots of wild-type plants, *nfp-like 1*, *nfp-like 3,4* and *nfp-like 1,3,4* mutants that were transformed with a nuclear-localized calcium sensor gGECO after treatment with LCOs from *Rhizobium sp.* IRBG74 as described in (Cope et al., 2019) and (Irving et al., 2022). Additionally, wild-type plants as well as the *nfp-like 1,3,4* line were treated with synthetic COs, sLCOs and nsLCOs. Wild-type roots show consistent nuclear calcium oscillations as already published (Cope et al., 2019). For the single *nfp-like 1* and the double *nfp-like 3,4* mutants, two independent genome edited lines were used. The *nfp-like 1* mutant lines displayed a calcium spiking response (Fig. 1A and 1C). The number of nuclei spiking more than three times was comparable to wild-type, however the average number of flashes per nucleus was lower in the *nfp-like 1* mutant (Fig. 1B). Contrasting to that, the *nfp-like 3,4* double and *nfp-like 1,3,4* triple mutants display no nuclear spiking, indicating that either *NFP-like 3*, *NFP-like 4* or both of them are crucial for calcium oscillations via CSSP but not *NFP-like 1* (Fig. 1A, 1B and 1C). Given that the CSSP can be activated in different plant species in response to diverse chitin molecules (Cope et al., 2019; Genre et al., 2013; Sun et al., 2015), we treated the *nfp-like 1,3,4* triple mutant alongside with wild-type roots with CO4, CO8, synthesized C16:0 sLCO or C16:0 nsLCOs and monitored Ca^2+^ spiking (Fig. 1D). The *nfp-like 1,3,4* mutant was also severely affected in its response, with no or drastically reduced spiking after treatment with the purified LCO and CO species (Fig. 1D). Therefore, NFP-like receptors of poplar are required for the induction of nuclear calcium oscillations.

**Figure 1:**
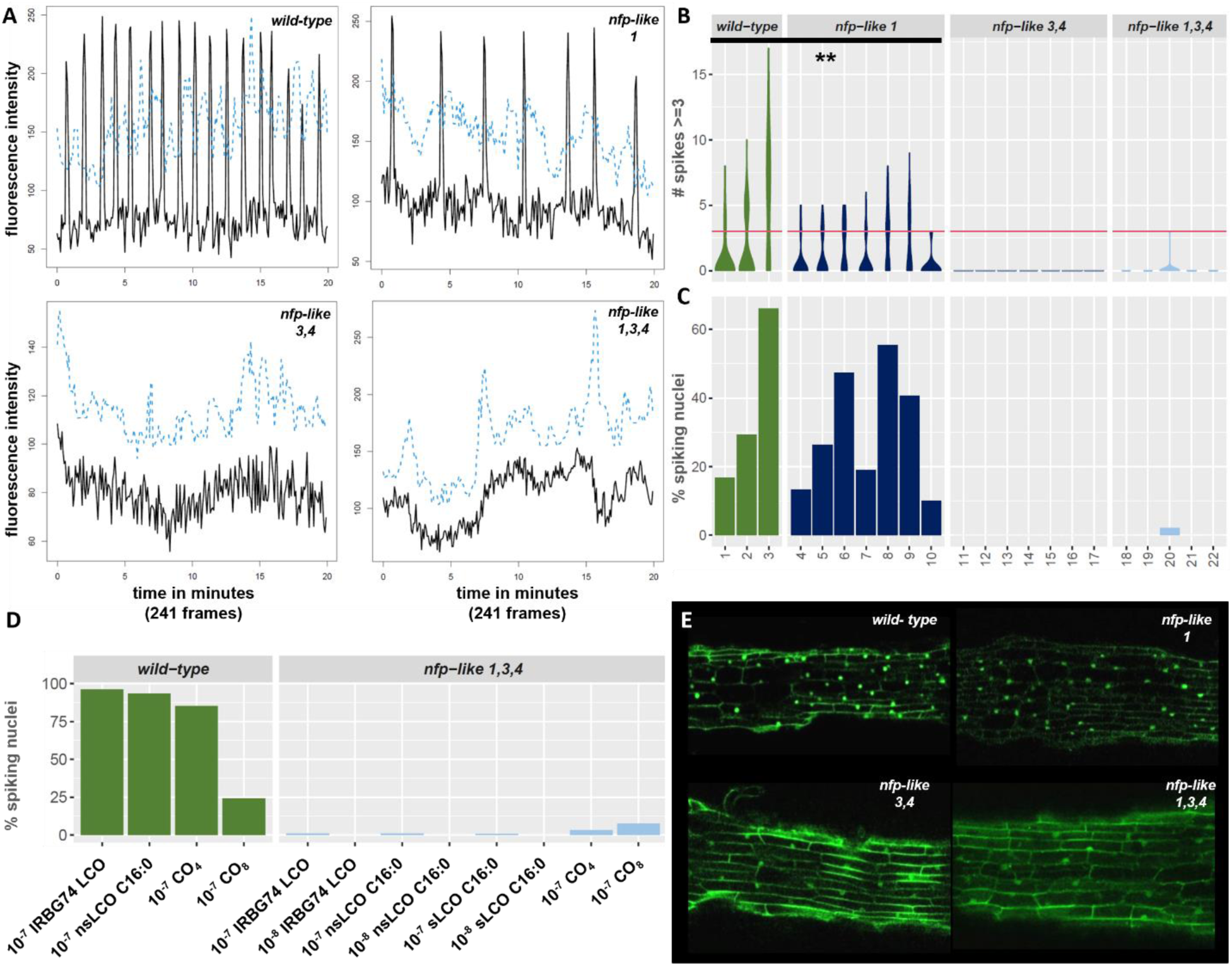
Wild-type plants and *nfp-like* mutants expressing a gGECO reporter construct were treated with LCOs ad COs to monitor nuclear calcium spiking. (A) Representative Ca^2+^-spiking patterns of root cell nuclei of wild**-**type, *nfp-like 1*, *nfp-like 3,4* and *nfp-like 1,3,4* treated with *Rhizobium sp*. IRBG74 LCO (10^-7^ M) over 20 minutes. Pictures were taken every 5 seconds resulting in 241 frames in total. (B) Number of nuclear spikes in every root sample. The red line indicates the threshold of 3 spikes/nucleus that was set as minimum spiking number a nucleus must have to be counted in (C). In each sample the maximum number of nuclei visible in the overlay picture (see methods) was counted. (C) Percentage of spiking nuclei in wild-type plants and the different mutant lines. For analysis, visible nuclei were labeled as region of interest in 20 min videos of roots (E) and spikes were counted. For *nfp-like 1* and nfp-like 3,4 two independent single mutant lines (independent CRISPR events) were recorded. For *nfp-like 1* sample 4-7 represent CRISPR line 1, 8-10 represent CRISPR line 2, For *nfp-like 3,4*, 11-14 are replicates of a CRISPR line, while 15-17 are replicates for another independent double mutant line. For the *nfp-like 1,3,4* triple mutant 5 replicates were done for the same line (18-22). (D) Wild-type plants roots were treated with the indicated LCOs/COs with the concentration 10^-7^ M once. The *nfp-like1,3,4* mutant treatments with different 10^-7^ M LCOs/COs were repeated 3 times with independent replicates while LCO treatments with the concentration of 10^-8^ M were repeated twice. For representation of repetitions, average of % spiking nuclei were calculated. (E) Representative confocal images of transformed roots during induced Ca^2+^response with LCOs isolated from *Rhizobium sp*. IRBG74.

### The amino acid residues forming the hydrophobic patch in the LysM2 domain in legume NFPs are conserved in poplar NFP-like proteins

To investigate the mode of action of NFP-like candidates, we compared the sequence and structure of LysM2 domains that are critical for LCO binding in already characterized NFP proteins like of *M. truncatula* with the domains present in NFP-like proteins in *P. x canescens*. Sequence alignments of the *P. tremula* and *P. alba* alleles of the *canescens* hybrid with domains of LCO binding proteins of other species confirmed the presence of the amino acids giving LysM2 its hydrophobic character at the binding site (Gysel et al., 2021; Cope et al., 2023) (Fig. 2A). However, NFP-like 1 and 3 harbor a negative patch that is predicted to impair binding of LCOs in this region that relies on positioning of the fatty acid chain in the hydrophobic area (Gysel et al., 2021) (Fig. 2B). In the NFP-like 1 protein of the *P. tremula* allele this patch is formed by two negative aspartate residues in position 153 and 156 (Fig. 2B), in the *P. alba* allele only by D156 that is located in the middle of the hydrophobic area formed by the leucine and alanine in position 147 and 154 respectively (Fig. 2 A). For both, the *P. alba* and *P. tremula* variant of NFP-like 3, negative charges resulting from D153 and E157 are present directly next to the predicted hydrophobic area that might be in consequence only moderately hydrophobic (Fig. 2 B). In contrast, the groove of both NFP-like 4 variants (*P. alba* and *P. tremula*) is strongly hydrophobic, similar to that of *M. truncatula* NFP (Fig. 2 B). We also analyzed the intracellular kinase domain of the poplar NFP-like candidates and compared them to AtCERK1 and MtLYK3 and MtNFP and found that they lack the P-loop as well have an altered DFG motive like MtNFP in contrast to AtCERK1 and MtLYK3 (Suppl. Fig. S1).

**Figure 2:**
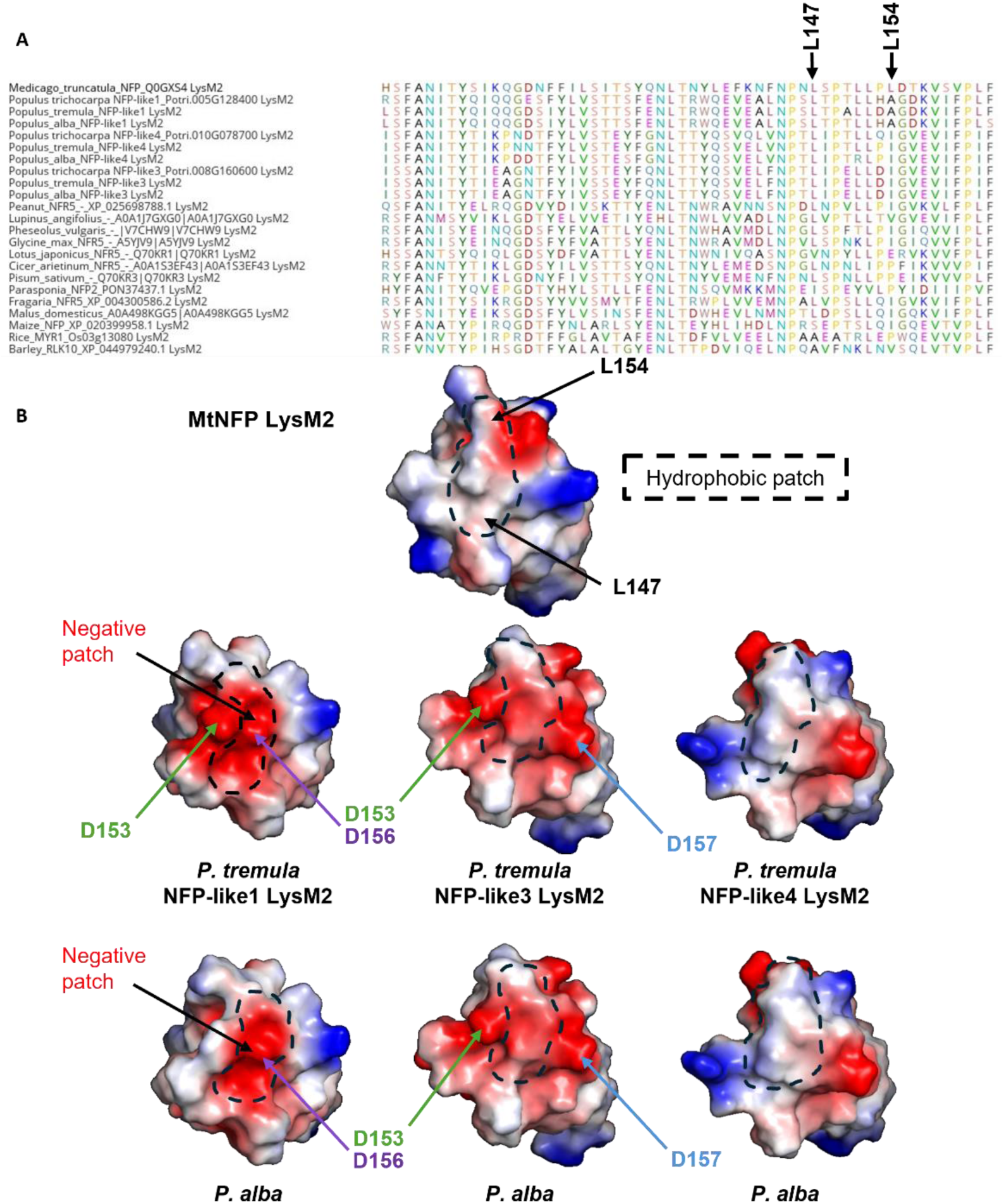
Protein alignment of LysM2 domains of Medicago, Poplar, legumes and monocots and predictions of hydrophobic patches on LysM2 domains of Medicago and Poplar. (A) The lysine residues L147 and L154 important for LCO binding in legumes have been highlighted. These hydrophobic residues are conserved as leucine, isoleucine or alanine in *Populus trichocarpa*, *Populus tremula* and *Populus alba* NFP-like 1, NFP-like 3 and NFP-like 4. (B) Structures of LysM2 domains have been predicted using Alphafold2. Red, blue and white colors indicate acidic (negatively charged), basic (positively charged) and hydrophobic residues respectively. In green, purple and blue negatively charged aspartate residues (D153 and D156) and glutamate (E157) are indicated. A negative patch is present in the LysM2 domain of *P. tremula* NFP-like 1 enhancing the negative charge in this region. In LysM2 of *P. alba* NFP-like 1 D156 is as well present in the middle of the hydrophobic patch. In LysM 2 of NFP-like 3 the hydrophobic area is framed by two negative patches (D153 and E157) in both, *P. alba* and *P. tremula*.

### NFP-like receptors control LCO-induced gene expression

To determine, if *NFP-like* genes are part of a positive feedback loop downstream of CSSP, we monitored their expression upon LCO treatment in wild-type plants. Roots were treated with 18:1 nsLCOs and then incubated for another five hours before analysis. These five hours were chosen as that period was previously described for LCO triggered gene expression (Irving et al., 2022). For all *NFP-like* genes we saw a significantly enhanced gene expression upon LCO treatment in wild-type plants (Fig. 3A) confirming our hypothesis of a LCO induced positive feedback loop. A recent study showed that poplar putative sulfotransferase *PtSS1* and a lateral root transcription factor Nodule Inception 2b (*PtNIN2b*) are rapidly induced by LCO treatment in wild-type plants (Irving et al., 2022). *PtSS1* is a strong marker for LCO induced gene expression in the outer cell layers of mature roots with a maximal expression after four to five hours upon treatment. *PtNIN2b* expression is also significantly induced after LCO treatment and is required for LCO-induced lateral root development in poplar. Building upon these findings, we replicated the LCO treatment assay using *nfp-like* knockout lines. For wild-type plants we were able to replicate the results from Irving et al. where LCO treatment significantly enhances gene expression of both *PtSS1* and Pt*NIN2b* (Fig 3B and C). For *nfp-like 1*, *nfp-like 3,4* and *nfp-like 1,3,4,* we did not see induction of gene expression for both genes. This indicates that for proper LCO mediated transcriptional changes either *NFP-like 1* and/or *NFP-like 3* and/or *4* or all of them are required, even though for nuclear calcium spiking *NFP-like 1* seemed dispensable. To determine which of the receptors are the key players for LCO mediated signaling, we designed complementation constructs for all the *NFP-like* genes and complemented the mutants in different combinations. With these lines, LCO assays were either performed with stable complementation lines (for *nfp-like 1* mutants) or with transformed roots emerging from leaves after *A. rhizogenes* transformations with complementation constructs. To exclude the possibility that roots directly used after leaf transformation exhibit another transcriptional response upon LCO treatment than whole plants, we also transformed wild-type leaves with *A. rhizogenes* control strains to monitor the expression of the same genes as in the complemented mutant lines for comparison. Leaves of the *nfp-like 3,4* double mutant were transformed with complementation constructs for *NFP-like 3*, *NFP-like 4* or *NFP-like 3,4* to separate their mediated signaling responses from each other. After treating the roots for one hour with 18:1 nsLCOs and five hours incubation time, the expression of Pt*NIN2b* and *PtSS1* was monitored in the different combinations. Complementation of *nfp-like 1* mutants restored expression induction of both genes after LCO treatment (Figure 3D and E). Complementation of the *nfp-like 3,4* double mutant with *NFP-like 3* alone restored LCO dependent expression. Significant induction of gene expression was visible when restoring the function of *NFP-like 4,* but much weaker compared to the complementation with *NFP-like 3* (Fig. 3D and E). Restoring both genes in the double mutant background also leads to significantly enhanced gene expression after LCO treatment compared to the ethanol control (Fig. 3D and E). In all lines, we also see higher relative baseline expression of *PtSS1* and *PtNIN2b* compared to wild-type conditions. One explanation for that could be that all complementation constructs are under control of very strong constitutive promoters that might lead to a significantly higher abundance of NFP-like proteins in the cell, influencing signaling within their host. In general, this effect might be neglectable since LCOs still lead to significant downstream gene expression inductions compared to control situations. In conclusion, our experiments demonstrate that restoring the function of the single *NFP-like* genes in the single and the double mutant also restores LCO induced expression of genes upregulated during symbiosis signaling cascades. In the *nfp-like 3,4* double mutant, specifically complementation with the *NFP-like 3* gene leads to a significantly enhanced gene expression of *PtSS1* as well as *PtNIN2b* indicating that *NFP-like 3* seems to be more important for downstream signaling than *NFP-like 4*.

**Figure 3:**
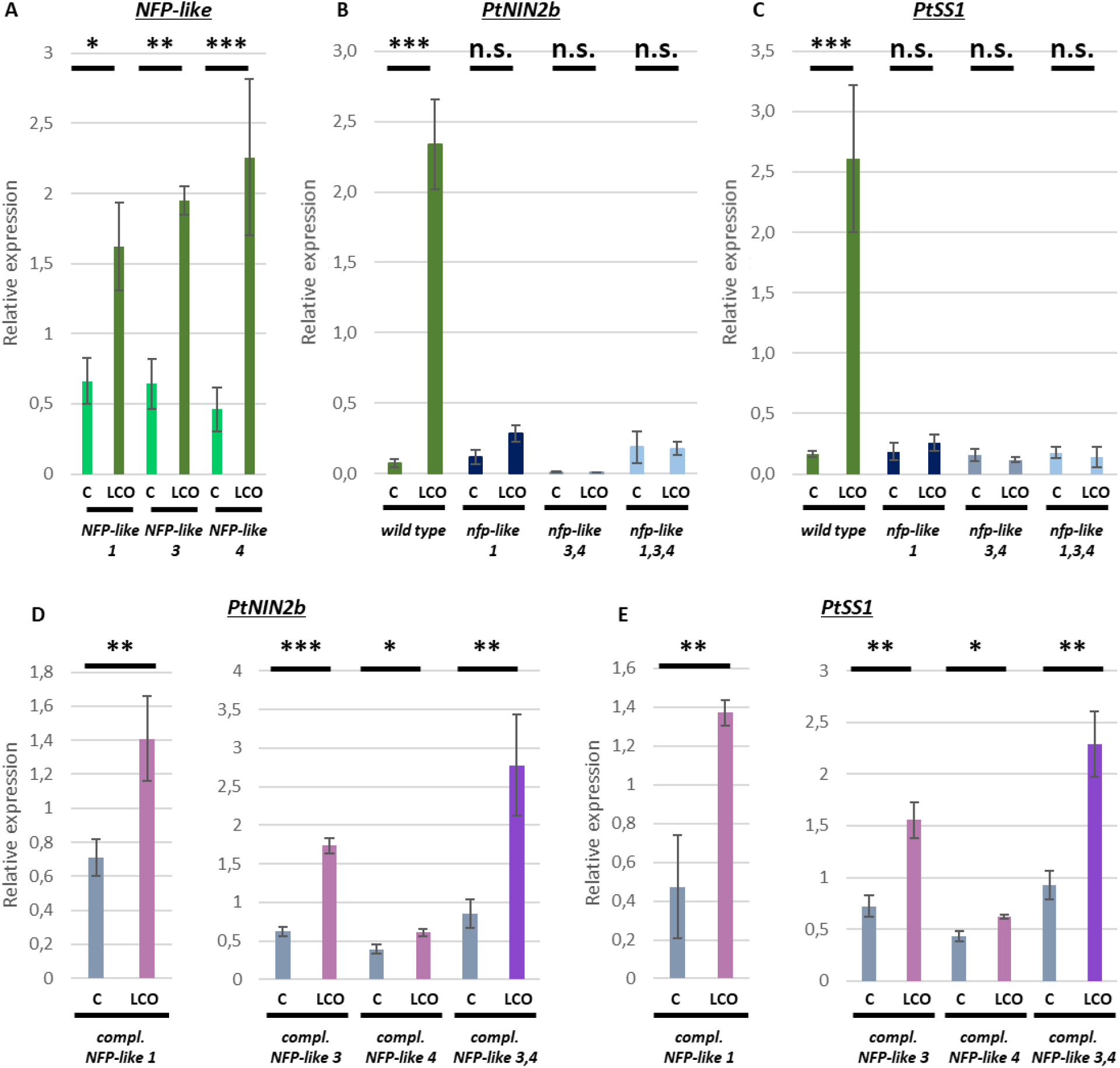
Expression of NFP-like receptors is enhanced upon LCO treatment and receptors themselves are crucial for CSSP mediated gene expression. (A) Relative gene expression of *NFP-like 1*, *NFP-like 3* and *NFP-like 4* genes in roots 5 hours after inoculation with LCO as determined by qRT-PCR. (B, C) Expression levels of CSSP inducible genes, *PtNIN2b* and *PtSS1* in the roots of wild-type plants and the single (*nfp-like 1*) double (*nfp-like 3,4*) and triple *(nfp-like 1,3,4)* mutants. Wild-type and *nfp-like* mutants were harvested after 5 hours after LCO treatment for quantifying gene expression of *PtNIN2b* (B) and *PtSS1* (C). (D-E) Restored expression of *PtNIN2b* of *PtSS1* in genetically complemented transgenic roots of *nfp-like 1/NFP-like 1*, *nfp-like 3,4/NFP-like 3*, *nfp-like 3,4/NFP-like 4* and *nfp-like 3,4/NFP-like 3,4*. Relative expression was measured in every trial with at least three biological replicates each trial (n=3). Significance was tested for all samples via ANOVA with posthoc Tukey test (Significance code: p ≤ 0.0001 ‘***’; p ≤ 0.01 ‘**’; p ≤ 0.05 ‘*’; p ≤ 0.1 ‘.’; p ≥ 0.1 not significant (n.s.)).

### NFP-like 3 is the main receptor for downstream gene induction

We observed that expression of *NFP-like* genes is also induced by LCO treatment as *PtSS1* and *PtNIN2b*. To understand potential positive feedback loops and to get a clearer picture of the role of the different receptor candidates in triggering downstream responses, we analyzed the expression of *NFP-like 1*, *3* and *4* in the different complementation lines. Due to the results of the complementation of the *nfp-like 3,4* double mutant (Fig. 3D and E), we hypothesized that the expression of the different receptor genes might be induced by the same signaling pathway as *PtSS1* and *PtNIN2b.* We performed expression analysis of *NFP-like 1* in the *nfp-like 3,4* complemented mutant lines that were used before with the described treatment (Fig. 3D and E) and observed a significant increase in expression when *NFP-like 3* (Fig. 3 A) was restored but not with *NFP-like 4* (Fig. 3 A). Complementing the mutant with both*, NFP-like 3* and *4*, resulted as well in a significant increase in gene expression of *NFP-like 1* after LCO treatment (Fig. 4A). These results strongly support the idea that *NFP-like 1* gene expression is regulated mainly by the presence of the NFP-like 3 receptor when LCOs are perceived in a positive feedback look. To determine if the expression of *NFP-like 3* and *NFP-like 4* genes is dependent on the presence of the NFP-like 1 receptor, we analyzed their expression in the *nfp-like 1* single mutant background as we did for *PtSS1* and *PtNIN2b*. In contrast to them, expression of *NFP-like 3* as well as *NFP-like 4* was significantly induced by LCO treatment (Fig. 4B) indicating that they are as well part of a positive feedback loop with themselves but not dependent on the presence of the NFP-like 1 receptor for being induced by LCOs. These results suggest that there might be different sets of genes for LCO/EM-fungus perception and development regulation needed for mycorrhiza formation that are induced in dependency of specific or all receptors being functional.

**Figure 4:**
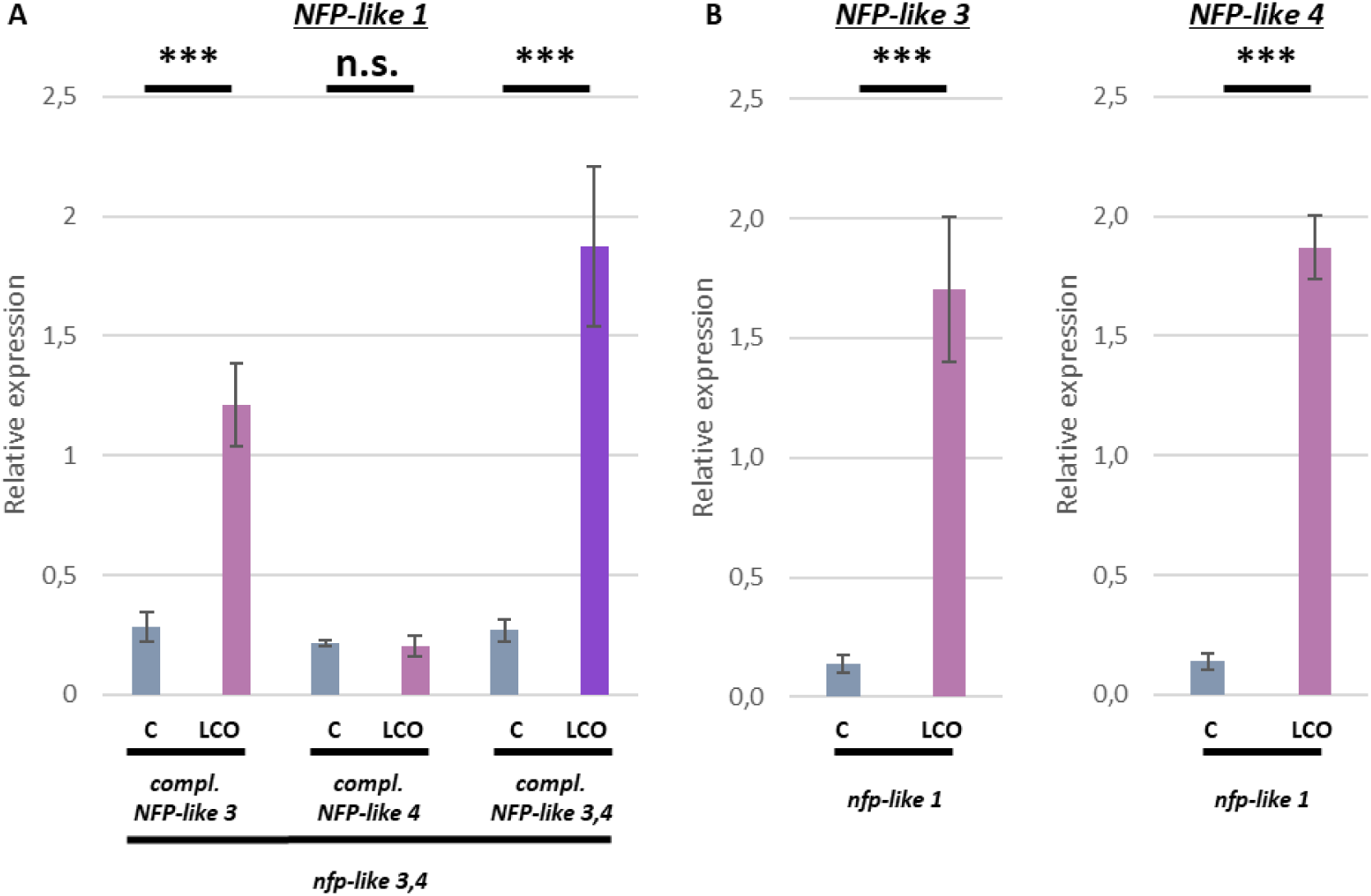
LCO-induced expression of *NFP-like 1* is dependent on NFP-like 3 but not vice versa. (A) Relative gene expression of *NFP-like 1* was analyzed 5 hours after LCO treatment in the *nfp-like 3,4* double mutant complemented with *NFP-like 3*, *NFP-like 4* or both by qRT-PCR. (B) Relative expression of *NFP-like 3* and *NFP-like 4* was analyzed 5 hours after LCO treatment in the *nfp-like 1* single mutant by qRT-PCR. Relative expression was measured in every trial with at least three biological replicates each trial (n=3). Significance was tested for all samples via ANOVA with posthoc Tukey test (Significance code: p ≤ 0.0001 ‘***’; p ≤ 0.01 ‘**’; p ≤ 0.05 ‘*’; p ≤ 0.1 ‘.’; p ≥ 0.1 n.s.).

### NFP-like receptors are crucial for the establishment of EM associations

To investigate if the differences of signal transduction as well as on the level of gene expression also result in altered mycorrhization, we cultivated wild-type plants and *nfp-like* mutants for four weeks with *L. bicolor.* In Cope et al. 2019, disturbance of signaling via CSSP by knocking down *CASTOR* and *POLLUX* or *CCaMK* led to reduced colorization with *L. bicolor* (Cope et al., 2019). Our assumption is that if the three NFP-like proteins localizing to the plasma membrane (Suppl. Fig. S2) are the starting point of signaling by perceiving the fungus via its LCOs and COs, mycorrhization also has to be altered as consequence of their knockout. After four weeks of co-cultivation between plants and *L. bicolor,* lateral root tips were investigated for the presence of the typical EM-mantle as well as tip swelling and arrested root growth associated with this symbiosis (Vayssières et al., 2015) (Fig. 5A) compared to tips without signs of mycorrhization in presence of the fungus (Fig. 5A). For all mutant lines, we found a significant reduction of the mycorrhizal phenotype (fewer swollen root tips/complete mantle vs. without). In wild-type plants the average mycorrhization rate was 66% of all observed tips (Fig. 5B). The *nfp-like 1* single mutant shows only 46 % mycorrhizal like tips, i.e. significantly less than in the wild-type plants. Reduction in receptor availability enhanced the severity, with the *nfp-like 3,4* double mutant at 31% and the triple mutant with 28% mycorrhizal like tips (Fig. 5B). These results show that all three receptors are important for mycorrhiza inducing signaling processes. However, since mycorrhization was not completely impaired, other receptors could be involved in perception of symbiotic signals.

**Figure 5:**
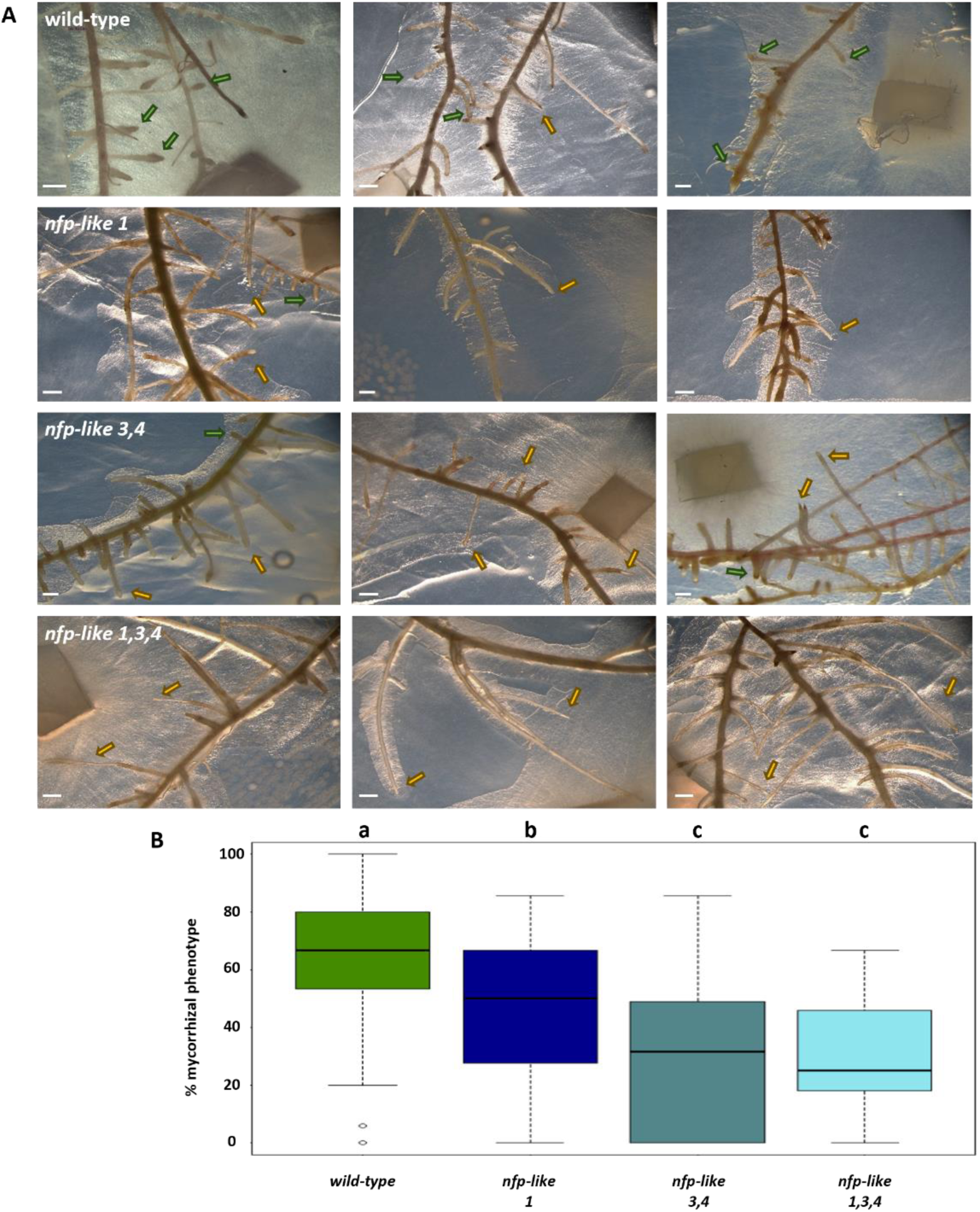
Colonization of wild-type plants and mutant poplar lines by *Laccaria bicolor*. Wild-type plants and *nfp-like* mutants were co-cultivated with *Laccaria bicolor* for four weeks. Pictures were evaluated from 3 experiments with at least 10 roots each line. (A) Example images show observed root phenotypes. Wild-type roots form the typical EM phenotype in contact with *L. bicolor* with mantled tips in different stadia of swelling (green arrows) and less often non-colonized lateral roots or tips that are breaking through the mycorrhizal mantle structure (yellow arrows). For the *nfp-like* mutant lines more tips were observed that are not colonized or outgrow the fungus by breaking the EM mantle (yellow arrows) compared to wild-type roots. Scale bar = 1mm. (B) From all tips in each picture the percentage of EM phenotypic root tips in relation to formed lateral roots in total was calculated. Evaluation was done with R-studio testing significance via Poisson distribution followed by adapted Tukey posthoc test for pairwise comparison. P-values for a-b and a-c are <0.001 (***).

### Gene expression in wild-type plants and *nfp-like 1,3,4* mutants in response to *L. bicolor*

To characterize the differential response of wild-type plants and *nfp-like 1,3,4* triple mutants not only to LCOs, plants were co-cultivated with *L. bicolor* and differential gene expression was studied via RNAseq. To investigate the beginning mycorrhization, we co-cultivated the plants for one week in presence of the fungus, which is enough for early attachment for mycorrhiza formation but after perception-signaling as investigated and described in the paragraphs before. A total of 197 differentially expressed genes (DEGs) were identified in wild type plants co-cultivated with *L. bicolor* compared to the mock treatment. Similarly, 107 DEGs were present in the *nfp-like 1,3,4* triple mutant treated with *L. bicolor* compared to mock. Among these DEGs, 129 and 39 were exclusively present in wild-type plants and *nfp-like 1,3,4,* respectively. Several genes known as key players in symbiosis, such as LecRLK, LRR-RLK, sulfate transporters and chitinase etc., and transcription factors such as WRKY, NAC domain-containing proteins, and AP2/ERF showed high expression in wild-type plants treated with *L. bicolor* than in mock (Fig. 6 C). Among those several genes were not or even downregulated in the *nfp-like 1,3,4* mutant indicating that its reduced perception of the fungus also results in a misregulation of genes required for symbiosis establishment.

**Figure 6.**
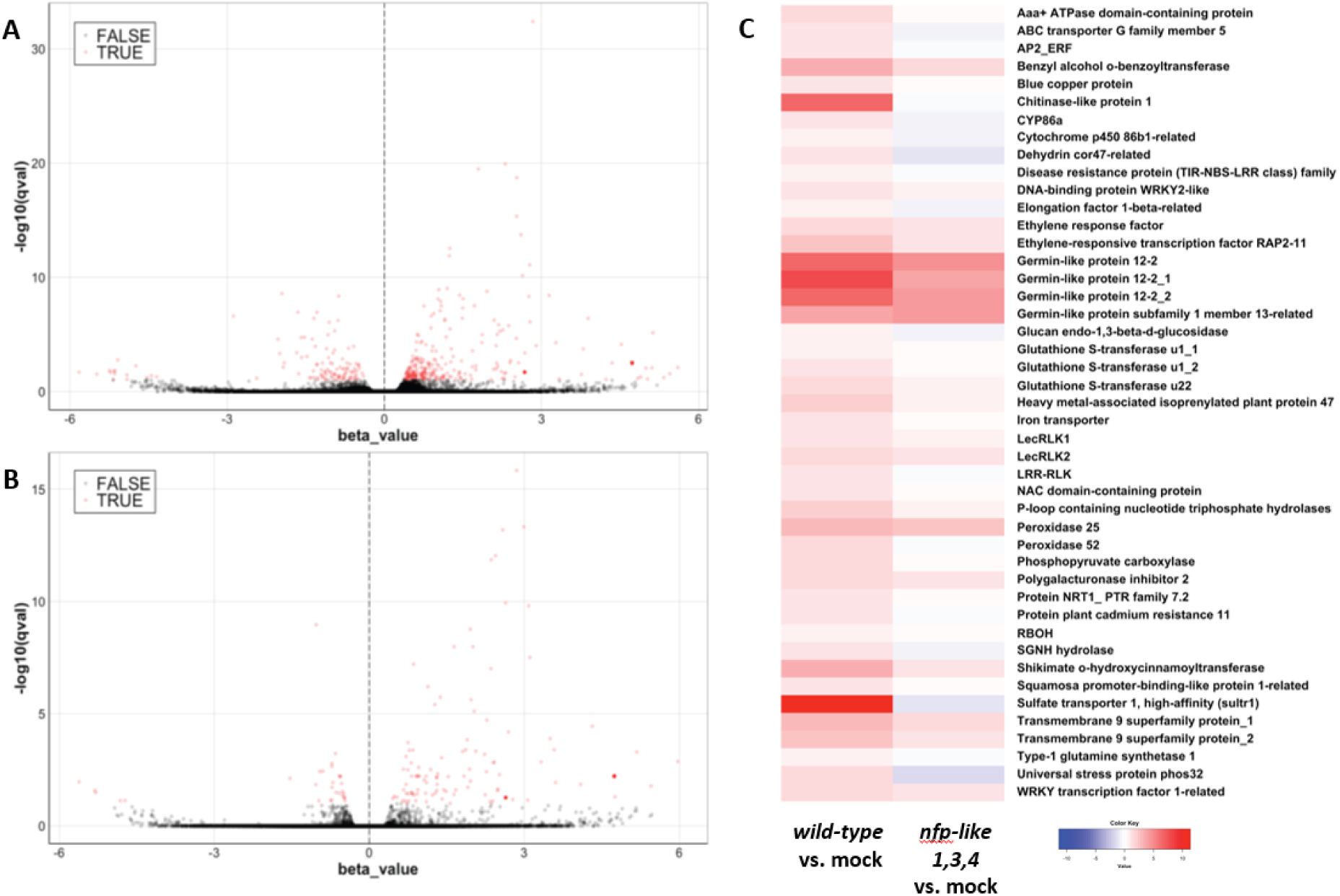
RNAseq based analysis of response of poplar wild type (WT) plants and *nfp-like 1,3,4* mutants upon *L. bicolor* inoculation. (A) Volcano plot representing the DEGs in wild-type plants treated with *L. bicolor* compared to mock. (B) DEGs in *nfp-like 1,3,4* plants treated with *L. bicolor* compared to mock. (C) Heatmap of representative DEGs in WT and *nfp-like 1,3,4* genotypes treated with *L. bicolor* compared to mock. Expressions are log2 transformed values of fold change of TPM between respective samples. Red indicates higher expression and blue indicates lower expression.

## Discussion

LysM receptor proteins that are crucial for the perception of fungal or bacterial chitin-derived signaling molecules are well described in herbaceous plants that form symbiotic interactions with AM fungi or rhizobia. Here we describe for the first time LysM receptor candidates involved in ectomycorrhiza symbiosis of the woody plant *Populus x canescens* with the EM fungus *L. bicolor* as being crucial for the activation of CSSP, downstream gene expression as well as wild type EM phenotypes. Amino acid sequence comparisons of NFP-like 1, 3 and 4 with symbiosis related genes of *M. truncatula*, *L. japonicus* and others demonstrate that our identified candidates share characteristic features crucial for LCO binding in the extracellular LysM2 domain (Fig. 2) as well as the absence of a functional kinase domain typical for LYR class proteins (Suppl. Fig. S1) (Ding et al., 2025). Thus, they are typical co-receptor candidates dependent on the interaction with LYK-class proteins to channel downstream signaling into the CSSP (Buendia et al., 2018). We also describe a negative patch disrupting the hydrophobic patch of LysM2 in NFP-like 1 and one framing it in NFP-like 3 proteins on both *alba* and *tremula* alleles in *P. x canescens,* that is absent in *NFP-like 4* as well as NFP homologs of other species (Fig. 2). These additional charges could reduce binding specificity and affinity to LCOs which might explain why NFP-like 1 does not have a central role in LCO triggered CSSP induction as the *nfp-like 1* knockout mutant still responds with nuclear calcium spiking, even with a lower frequency, while the *nfp-like 3,4* double mutant does not (Fig. 1). For NFP-like 3 the negative charges close to the hydrophobic patch might also interfere with LCO binding. Nevertheless, when restoring its expression in double mutant background, downstream gene expression after 18:1 nsLCO is inducible again showing that the negative patch does not interfere in such a way that interaction with a LCO decorated with a fatty acid is impossible.

Although NFP-like 1 might not be crucial for CSSP activation per se, it is needed for downstream LCO induced gene expression of *PtNIN2b* and *PtSS1*. Both were shown to be significantly upregulated after 18:1 nsLCO treatment in wild type *P. x canescens* ((Irving et al., 2022) and this study) but do not show any transcriptional response in all mutant lines including *nfp-like 1* (Fig. 3 B and C). An explanation might be that downstream signaling after calcium spiking is dependent on a certain level of spiking frequency to induce further signal transduction. Calcium oscillations activate the calcium decoder CCaMK to initiate downstream gene expression and it was shown that its activation is dependent on high levels of calcium input provided by a high calcium oscillation frequency (Singh and Parniske, 2012). Del Cerro and co-workers demonstrated that enhancing oscillation frequency by modifying the calcium-bound form of calmodulin 2 (holo-CaM2) results in accelerated early endosymbiosis signaling and enhanced root nodule symbiosis over time in *M. truncatula* with rhizobia (Del Cerro et al., 2022). A reduced frequency as observed in the *nfp-like 1* mutant might lead to the opposite effect affecting downstream gene expression as well as mycorrhizal root tip formation (Fig. 5). Another reason for the loss of downstream gene induction in calcium spiking *nfp-like 1* mutants could be a CSSP independent signaling pathway as described for EM forming plants which lack critical components of the CSSP (Garcia et al., 2015). Several gymnosperm plants that are able to form EM associations seem to circumvent this missing pathway by so far unknown signaling routes to initiate symbiosis. The contradictory responses in *nfp-like 1* mutants could be another hint towards alternative pathways in poplar which are, next to CSSP, crucial for symbiotic signaling and symbiosis related organ formation. In contrast to *nfp-like 1* plants, the *nfp-like 3,4* and *nfp-like 1,3,4* mutants are severely impaired in CSSP related calcium oscillations (Fig. 1) and downstream gene expression (Fig. 3). The calcium spiking response which is inducible by different LCO and CO species was significantly reduced or absent in the triple mutant after treatment with a diverse set of these signals proving the importance of these receptors for downstream signaling processes (Fig. 1D). Because NFP-like 3 and 4 are as paralogs very similar in their DNA sequence, it was impossible to knock them out separately and all genome editing trials ended up in mutations in both genes (Bruegmann et al., 2019). To circumvent this, we created complemented roots of each gene in the double mutant to identify potential unique functions of the two receptors. By this we were able to show that specifically NFP-like 3 is important as receptor for LCO mediated gene expression. This holds true for genes involved in plant development crucial for symbiotic interactions like *PtNIN2b*, as well as the receptor genes themselves (Fig 3 and 4). Interestingly NFP-like 1 does not have any influence on LCO induced expression of *NFP-like 3* and *4* genes since their expression is still upregulated in *nfp-like 1* mutants upon treatment, in contrast to *PtNIN2b* and *PtSS1*. This indicates that there might be different pools of genes that are activated by unique and separated combinations of NFP-like proteins with their so far unidentified co-receptors. One set of genes consisting of the NFP-like receptors themselves seem to be dependent on the LCO induced signal mediation by NFP-like 3 and to a smaller extend also NFP-like 4. They are specifically needed for early fungus perception and symbiotic partner recognition and their expression is in a positive feedback loop downstream of NFP-like 3 that enhances their LCO induced expression (Fig. 4). Other genes like *PtNIN2b* or *PtSS1* might represent a pool of genes needed for plant development (enhanced lateral root formation for increased target surface for fungus attachment) and symbiotic tissue formation and is dependent on the interplay of all three receptors analyzed in this study. NFP-like 1 could be also a CO receptor in this model. This is supported by the fact that the negative patch disrupting the hydrophobic area in LysM2 domain makes the binding of a LCO with its long fatty acid chain difficult (Gysel et al., 2021). Nevertheless, NFP-like 1 is needed for LCO triggered downstream gene expression associated with symbiosis formation but not when it comes to signal perception (NFP-like expression). An explanation could be that for gene expression activation of these different sets of genes different signaling routes with unique prerequisites have to be fulfilled. These can be specific but simultaneous LCO/CO binding events by different NFP-like proteins (expression of *PtNIN2b* and *PtSS1*) or separate activations (expression of NFP-like receptors). This is also supported by the reduced capacity to form mycorrhizal tip structures (Fig. 5) which can be observed in all mutants. Mature EM tips are characterized by a dense mycelium mantle, no root hairs, swelling, arrested root elongation as well as the formation of the Hartig net. The development of this structure follows a gradient. After attaching to the surface the fungus forms a sheath at the lateral sides of the root tip (colonization stage) that progresses into a full mantle covering the whole root with a visible Hartig net structure invading intercellular spaces in the root epidermal and cortex cell layer (mature EM) (Vayssières et al., 2015). Regulation of these processes seem to be mainly mediated by fungus induced changes in auxin signaling, homeostasis as well as transport leading to the arrest and deactivation of the root apical meristem (Vayssières et al., 2015). In the *nfp-like* mutants the fungus is not as efficient as in wild-type plants to attach to the root, arrest root growth and to form a mantle. Root tips of the mutants more often continue to grow, breaking through the forming mycelium sheath because missing NFP-like mediated signaling seems to alter recognition of the symbiotic partner (Fig. 5). Even though we did not look into Hartig net formation due to the high amounts of samples that are needed for proper statistics to compare the depth of the hyphal network penetrating the intercellular space, we assume from the data we gathered, that the other criteria (growth arrest, tip swelling and mantle formation) are enough to compare potential EM formation between wild-type plants and the mutants. That knocking out all three receptors heavily impact signaling, became also clear in our transcriptomic approach of an early time point of plant-fungus interaction (Fig. 6). After one week of co-cultivation, *L. bicolor* starts to attach to the plant root surface and friend-foe distinction signaling is crucial for further steps of EM formation. In wild-type plants, most of the DEGs showed increased expression upon *L. bicolor* inoculation compared to the mock treatment; however, the *nfp-like 1,3,4* triple mutant appears to be less responsive to *L. bicolor* inoculation and exhibits reduced induction of these genes compared to the mock treatment. Lectin receptor-like kinase PtLecRLK1 mediates the interaction between Populus and the ectomycorrhizal fungi *L. bicolor (Labbé et al., 2019)*. PtLecRLK1 contributes to mantle formation, and higher expression of its homologs (PtXaTreH.11G103000.1 and PtXaTreH.11G102400.1) was observed in wild-type plants inoculated with *L. bicolor* compared to mock-treated plants. Fungal recognition mediated by LecRLK is accompanied by the activation of downstream effectors such as ROS-detoxifying enzymes (glutathione S-transferases, peroxidases), and transcription factors (WRKY, NAC, AP2/ERF). The activation of these genes suggests that symbiotic association with *L. bicolor* involves a molecular orchestration that balances defense and protective responses to facilitate fungal accommodation. The absence of transcriptional rewiring in the *nfp-like 1,3,4* mutant clearly indicates that NFP-like-dependent signaling is indispensable for poplar’s interaction with *L. bicolor* and its colonization. Therefore, similar to the role of NFP in LCOs/COs perception in rhizobia or AMF associations (Choi et al., 2018; Finegan et al., 2025), NFP-like proteins are also required for ectomycorrhizal symbiosis in poplar. Conclusively, the NFP-like proteins we characterized in this study are the first early-signaling receptors described for ectomycorrhiza formation that mediate signaling via CSSP, mediate downstream gene expression as well as mycorrhiza formation. Because EM formation is not completely dysfunctional, the next step could be to focus on other receptors involved in symbiosis formation. Our data supports that there might be other pathways apart from CSSP mediated signal transductions which could also include other ways of perception. In AM symbiosis glucan binding EPR3a receptor complexes are involved in fungal infection as well as arbuscle formation (Kelly et al., 2023). EPR3 receptors are conserved in dicot and monocot plant genomes and if they play a role in EM interactions is completely unknown.

## Materials and Methods

### Plant material, culture and generation of knockout mutants

*Populus* x *canescens* (*Populus tremula x alba*) hybrid clone INRA 717-1B4 was grown and propagated on ½ MS medium (Duchefa Biochemie, Haarlem, Netherlands) for 16-hour day/8-hour night photoperiod and ∼50 μmol.m^-2^.s^-1^ of photon flux. Transgenic lines were obtained by generation of CRISPR/Cas9 constructs to knock out Nod Factor Perception protein like 1 (NFP-like 1) (referring to Potri.005G128400), NFP-like 3 (Potri.008G160600) and NFP-like 4 (Potri.010G078700) proteins followed by leaf-disc transformation protocol (Bruegmann et al., 2019). Candidate selection of genes and generation and verification of *nfp-like 1* and *nfp-like 3,4* knock out lines are described by Bruegmann and co-workers (Bruegmann et al., 2019). Since *NFP-like 3* and *NFP-like 4* are a paralog pair with high sequence similarity, it was impossible to produce single mutant lines for them. To generate *nfp-like 1,3,4* knock out mutants, plants were co-transformed with the vectors used for generating *nfp-like 1* and *nfp-like 3,4* lines. Vectors and sgRNAs used are also described in detail in (Bruegmann et al., 2019). Mutations induced in the two *nfp-like 1,3,4* triple mutant used in this study are shown in suppl. table T1. Off-target prediction was performed using CRISPR-P 2.0 (http://crispr.hzau.edu.cn/cgi-bin/CRISPR2/CRISPR). Potential off-target sequences were amplified, sequenced and modifications excluded. The complementation line for *nfp-like 1* was generated by stably transforming *nfp-like 1* mutants with a construct harboring the coding sequence of *NFP-like 1* under control of an Ubi10 promoter of *A. thaliana* after deletion. Transformation was performed as described in (Bruegmann et al., 2019) .

### Structure prediction of LysM2 domain in NFP-like proteins

Structures were predicted using AlphaFold2 (version 2.2.2) (Jumper et al., 2021) . Electrostatic surfaces were calculated using PDB2PQR and APBS webservers and visualized in PyMOL using APBS tools 2.1 (Jurrus et al., 2018) .

### Construct design for localization and complementation studies

The Greengate method (Addgene Kit #1000000036) (Lampropoulos et al., 2013) was used to clone complementation constructs as well as fluorescent protein fusions for localization studies of NFP-like 3 and NFP-like 4. To complement CRISPR mutants with active Cas9, coding sequences of *NFP-like 3* (1866 bp) and *NFP-like 4* (1857 bp) were mutagenized by introducing five sense mutations within each gRNA binding site. Additionally, the stop codons alongside internal *Bsa*I restriction sites that would interfere with Greengate cloning were removed. The codon-optimized *NFP-like 3* and *NFP-like 4* coding sequences were synthesized by Azenta Life Sciences and PCR-amplified with the CloneAmp HiFi PCR Premix (Takara, Cat#639298) using the following primer pairs with respective Greengate-specific overhangs. The respective PCR products were cloned into the *pUC19* based Greengate module pGGC000 using the T4 DNA Ligase (NEB, Cat#M0202). To generate *UBQ10p:NFP-like 3-VENUS* (7870 bp) and *UBQ10p:NFP-like 4-VENUS* (7861 bp), Greengate modules containing the previously published *UBIQUITIN 10* (*UBQ10*) promoter and terminator (Lampropoulos et al., 2013), *BASTA* plant resistance cassette (Lampropoulos et al., 2013) and linker-VENUS (Wallner et al., 2023) were assembled together with *NFP-like 3* or *NFP-like 4* modules into the *pGreen-IIS* based vector backbone pGGZ003. The single complementation constructs *UBQ10p:NFP-like 3* (7095 bp) and *UBQ10p:NFP-like 4* (7086 bp) were assembled by replacing the coding sequence for *VENUS* with the Greengate d-dummy module that contained a stop codon. To generate the dual expression cassette construct *UBQ10p:NFP-like 3;UBQ10p:NFP-like 4* (10353 bp), the Greengate intermediate vectors pGGM000 and pGGN000 (Lampropoulos et al., 2013) served as backbones for *UBQ10p:NFP-like 3* and *UBQ10p:NFP-like 4*, respectively, and were combined in a final Greengate reaction into destination vector pGGZ003. All Greengate assemblies used the BsaI-HF_v2 (NEB, Cat#R3733) restriction enzyme and the T4 DNA Ligase (NEB, Cat#M0202) and were performed according to the manual (Lampropoulos et al., 2013). The ligation products were transformed into *E. coli* strain DH5α and positive clones were selected by spectinomycin resistance conferred by pGGZ003. To select the correctly assembled plasmids, test digestions were performed on several plasmid preps using the *Kpn*I (NEB, Cat#R3142) restriction enzyme and plasmid sequences were verified by full plasmid sequencing at the Microsynth AG.

### Generation reporter and complementation lines using *A. rhizogenes* leaf transformation

For visualization of nuclear calcium spiking, *Agrobacterium rhizogenes* strain ARqua1 carrying binary vector pEC11579 encoding both a nucleus-localized version of the G-GECO calcium sensor and the fluorescent protein DsRed was used (Cope et al., 2019). For complementation of *nfp-like 3,4* double mutants *A. rhizogenes* ARqua1 was co-transformed with pSOUP helper plasmid (Hayta et al., 2021) (pSOUP was a gift from Wendy Harwood & John Innes Centre - Wheat Transformation and Genome Editing (Addgene plasmid # 165419 ; http://n2t.net/addgene:165419 ; RRID:Addgene_165419)) as well as the before described greenGate complementation constructs. *NFP-like 3* and *NFP-like 4-VENUS* reporter constructs were, as well, co-transformed with pSOUP into *A. rhizogenes* ARqua1. For generation of transgenic roots harboring the different constructs, poplar leaves were transformed with the transgenic *A. rhizogenes* using a protocol published in (Cope et al., 2019). Briefly, poplar leaves were mechanically injured via multiple perpendicular incisions along the midrib on the abaxial leaf surface using a sterile scalpel. The wounded leaves were inoculated with transgenic *A. rhizogenes*, diluted to an optical density at 600 nm (OD_600_) of 0.8. Post-inoculation, the leaves were incubated on co-cultivation medium for 48 hours, followed by transfer to selection medium containing cefotaxime at 200 mg·L⁻¹ and Timentin® (mixture of ticarcillin and clavulanic acid) at 300 mg·L⁻¹ to eliminate residual bacteria. The explants were then maintained in dark conditions at 25 °C for two weeks. In case of lines harboring fluorescent reporter constructs, root development was monitored using a fluorescence stereomicroscope (Leica MZ10F), to identify positive transformants. In all cases, roots generated by this method were directly used for follow-up experiments using qRT-PCR or confocal microscopy as described in the next paragraphs.

### Ca^2+^ spiking assays

Transformed roots showing DsRed expression were extracted from the leaves and incubated for at least 2 hours in liquid woody plant medium with reduced phosphorus (2.5 mM KH_2_PO_4_), diminished nitrogen (no NH_4_NO_3_) and 0.1 mM 2-aminoethoxyvinyl glycine (AVG) (Cope et al., 2019). For monitoring Ca^2+^ spiking, we followed the protocol of Cope and co-workers (Cope et al., 2019). Roots were separately mounted on a slide in a well formed with vacuum grease (Dow Corning) in a solution of water containing LCOs isolated from *Rhizobium sp.* IRBG74 at a 10^-7^ M concentration or in 0.005% ethanol (mock condition) (R Cope et al., 2021). Wild type and *nfp-like 1,3,4* roots were furthermore monitored after treatment with C16:0 sulfated LCO (C16:0 sLCO), C16:0 non-sulfated LCO (C16:0 nsLCOs) and chitotetraose, tetra-N-acetyl (CO4) (IsoSep, Tullinge, Sweden) and chitooctaose, octa-N-acetyl (CO8) (IsoSep, Tullinge, Sweden). After 20 min of incubation, roots were monitored for G-GECO signals using a Leica SP8 confocal microscope with a HC PL APO CS2 20x/0.75 IMM objective (at JKI) or a cLSM780 confocal laser scanning microscope (Carl Zeiss) with an LD C-Apochromat 403/1.1-W Korr M27 objective (University of Wisconsin - Madison). G-GECO was excited with the 488 nm line of an argon laser, and green fluorescent protein emission was detected at 500 to 535 nm. DsRed was excited with a 561 nm line argon laser and emission was detected at 579 to 624 nm wavelength. Pictures were taken in 5 second intervals for 20 min in total with a format of xy: 1024 x 1024 scan resolution with Leica SP8 (or 512 x 512 at Zeiss cLSM780). Settings for all plant lines and treatments were the same.

Microscopy movies made at JKI were analyzed with a semi-automated pipeline using javacv (https://github.com/bytedeco/javacv). For each video, all frames were extracted and the color distribution was normalized to the first frame. Then, for each pixel, the 97.5% quantile across all frames was determined to create a stacked image. Next, for each pixel of the stacked image, the background signal was determined as the 50% quantile of all pixels with a maximum distance of 20, resulting in a background image. The corrected stacked image was obtained by subtracting the background image from the stacked image. Visible nuclei of each cell were manually labeled in the corrected stacked images. A profile for each labeled nucleus was then extracted from the normalized video. Visualization and counting of spikes were done in R (Version 4.4.2). Spikes were determined as values exceeding a dynamic threshold. This threshold was determined for each point independently as the median plus five times the MAD (median absolute deviation) over a centered window of size 21 frames. The corresponding script for the described workflow can be found under https://github.com/JensKeilwagen/nuclei. Videos made at University Wisconsin – Madison (Fig. 1B and C 20-22 and Fig. 1D) were evaluated as described by Cope and co-workers (Cope et al., 2019).

### Evaluation of poplar responses to LCOs

The assay was performed according to (Irving et al., 2022). Poplar wild-type plants and *nfp-like* mutant plants were cultivated on ½ MS-medium until roots reached a length of approximately 25 mm. Young plants were aseptically transferred to square culture plates containing woody plant medium broth with reduced phosphorus (2.5 mM KH_2_PO_4_) and diminished nitrogen (no NH_4_NO_3_) for another 10 days. Plant root systems were incubated in 10^-8^ M synthesized 18:1 non sulfated (ns) LCO (synthesized LCOs provided by S. Fort; CERMAV) or a control solution (0.005% (v/v) solvent for one hour. Following that, the treatment and control solutions were removed, and the plants were returned to the described cultivation conditions for five additional hours. Roots were harvested five hours post-LCO treatment. Excised root samples were immediately snap-frozen in liquid nitrogen to preserve RNA integrity for subsequent RNA isolation and quantitative reverse transcription polymerase chain reaction (qRT-PCR) analysis. In case of the *nfp-like 3,4* complementation lines, roots from transformed leaves were directly used for LCO assays and treated in the same way as for microscopy: roots were incubated for at least two hours in woody plant medium with reduced amounts of phosphorus, diminished nitrogen, and 0.1 mM AVG to block ethylene production and then treated for one hour in LCO-solution. Afterwards they were transferred back to the above-mentioned WPM for another five hours. For RNA isolation 5-10 roots were pooled and snap-frozen in liquid nitrogen for following gene expression analysis. To make sure that this method gives comparable results to whole plant treatment, wild type roots generated by transformation with *A. rhizogenes* containing only pSOUP were treated with LCOs and used for gene expression analysis. To compare the stable *nfp-like 1* complementation with the n*fp-like 3,4* complementation lines, roots were excised from grown plants and treated as roots resulting from *A. rhizogenes* transformation.

### RNA isolation and quantification of gene expression by qRT-PCR

RNA was extracted using the NucleoSpin RNA plant and fungi kit (Macherey-Nagel ref 740120.250) following manufacturer’s instructions. Total RNA quantity and quality were analyzed using a nanodrop spectrophotometer. After DNase I treatment (DNA-free, Thermo Fisher Scientific catalog number ENO521), a test PCR using one of the reference primers followed by gel electrophoresis was performed to check for residual genomic DNA contaminations. The following PCR program was used: denaturation step 99 °C for 2 min followed by repeated cycles of denaturation, annealing, and extension as follows: 35 cycles of 95 °C for 30 s, 65 °C for 30 s and 72 °C for 30 s and a final extension at 72 °C for 5 min. RNA quality and quantity was again determined by nanodrop spectrophotometer before 250 ng RNA were used for cDNA transcription using the RevertAid H Minus reverse transcriptase kit (Thermo Scientific catalog number EP0451) according to the manufacturers protocol. To quantify the expression of LCO responsive genes we performed qRT-PCR for genes encoding for *PtNIN2b* and *PtSS1* (Irving et al., 2022). qRT-PCR runs for the house-keeping genes, actin2 and elongation factor 1B, were run together with the genes of interest for internal normalization (Irving et al., 2022). The reaction was performed in 96 well plates using the qPCR Mastermix EvaGreen® No Rox kit (Bio&Sell item number BS76.590.0200) according to the manufacturer’s protocol. For each sample cDNA was diluted 1:5 (v/v) with ddH_2_O. The qRT-PCR was performed on the BioRad C1000Touch CFX96 system with the following program: initial denaturation step 95 °C for 15 min, followed by repeated cycles of denaturation, annealing, and extension as follows; 39 cycles of 95 °C for 15 s, 58 °C for 30 s. A melt curve analysis was performed to assess the specificity of the PCR product at 95 °C for 10 s, 90 °C for 5s. Primers used in this study (Table S1) were designed with the Primer-BLAST software (National Center for Biotechnology Information) Analysis of the resulting data was implemented by delta-Ct (2^−ΔCT) method (Livak and Schmittgen, 2001). Statistical analyses were performed with R-Studio (version 3.2.4) Significance of results was determined by ANOVA and posthoc Tukey test.

### Evaluation of ectomycorrhizal associations

Cocultures of *Laccaria bicolor* S238N (Felten et al., 2009) and poplar wild-type, *nfp-like1*, *nfp-like3,4* or *nfp-like1,3,4* knock out mutants were conducted using a well-established sandwich system (Cope et al., 2019). Briefly, *L. bicolor* was pre-cultured in the dark at 25 ^◦^C for two weeks on sugar-rich Pachlewski agar medium (P5) (Müller et al., 2013). Poplar micro-cuttings were allowed to root on solid ½ Murashige and Skoog (MS) medium and then transferred to already prepared, cellophane covered plates containing P20 medium supplemented with 0,1% MES (Müller et al., 2013). On the same plates eight plugs of pre-grown *L. bicolor* were arranged one week in advance to give the fungus the chance to establish already. After adding the plants, roots plus fungi were covered with another piece of cellophane membrane. Control plates containing plants without *L. bicolor* were prepared and handled the same way as co-culture plates. Cultivation was performed as described in first paragraph for another four weeks for mycorrhiza establishment. For colonization assays, undisturbed roots were studied using a stereomicroscope and subsequently imaged using a Leica MZ 10F. To evaluate mycorrhization efficiency, the mycorrhizal phenotype (swollen root tip, arrested root growth, hyphal mantle) was compared between wild-type plants and mutant lines in each picture made from the root system of individual plants. Root tips breaking through the hyphal mantle that continued to grow undisturbed were counted as non-mycorrhizal.

### Analysis of gene expression using RNA-Sequencing

Plant cultivation was performed as already described. Briefly, plants were grown with pre-cultured *L. bicolor* on P20 medium covered with cellophane membrane in square plates. Roots were harvested and snap frozen in liquid nitrogen and RNA isolation was done with the NucleoSpin RNA plant and fungi kit (Macherey-Nagel ref 740120.250) according to the manufacturer’s manual. Quality of samples and good RNA integrity numbers (RIN) were confirmed by Bioanalyzer measurement. Library preparation and RNA sequencing was done by Novogene using an Illumina Hiseq pe150 sequencer. Data of the paired end reads were filtered using the FastP program. Filtered paired-end reads were aligned using kallisto version 0.51.1, a k-mer-based pseudoalignment program (Pimentel et al., 2017) with a bootstrap value of 30. For alignment, the transcript sequence from *Populus tremula* x *Populus alba* HAP1 v5.1 genome was downloaded from Phytozome (https://phytozome-next.jgi.doe.gov/info/PtremulaxPopulusalbaHAP1_v5_1). The differentially expressed genes were identified using the Sleuth 0.30.1 package, with a q-value threshold of 0.05.

## Supporting information

Supplementary data Calcium spiking

Supplementary data qRT-PCR & Primers

## 5. Supplementary figures

**Supplementary Fig. S1.**
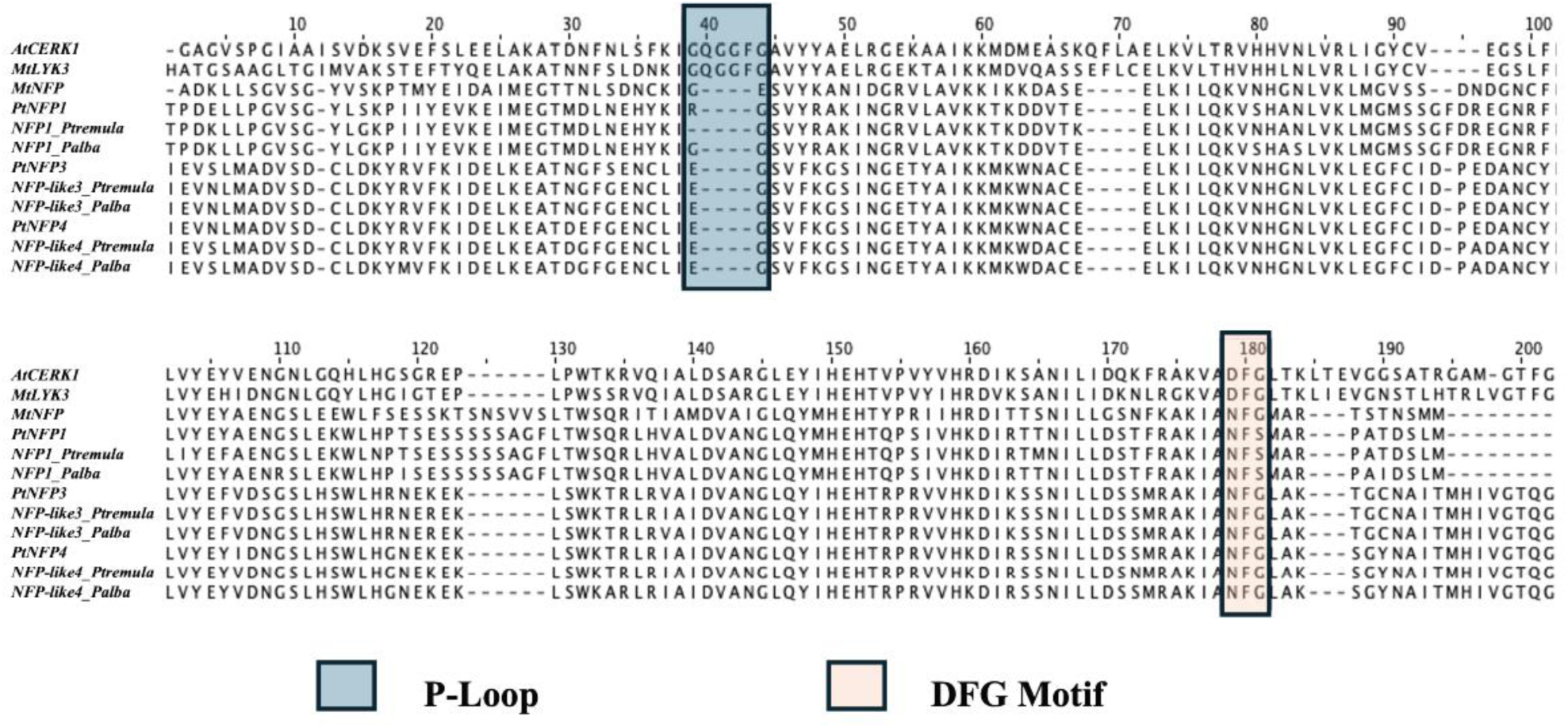
Comparison of the kinase domains in NFP, CERK1, and LYK3 proteins. Multiple sequence alignment of the N-terminal kinase domains of the *A. thaliana* CERK1, *M. truncatula* LYK3 and NFP, and *Populus* sp. NFP-like proteins. The canonical P-loop (GxGxxG) motif required for ATP binding is present in CERK1 and LYK3 but absent in *M. truncatula* and *Populus* sp. NFP-like proteins. Similarly, the DFG motif within the activation loop is altered in *M. truncatula* and *Populus* sp. NFPs, consistent with their lack of kinase activity.

**Supplementary Fig. S2.**
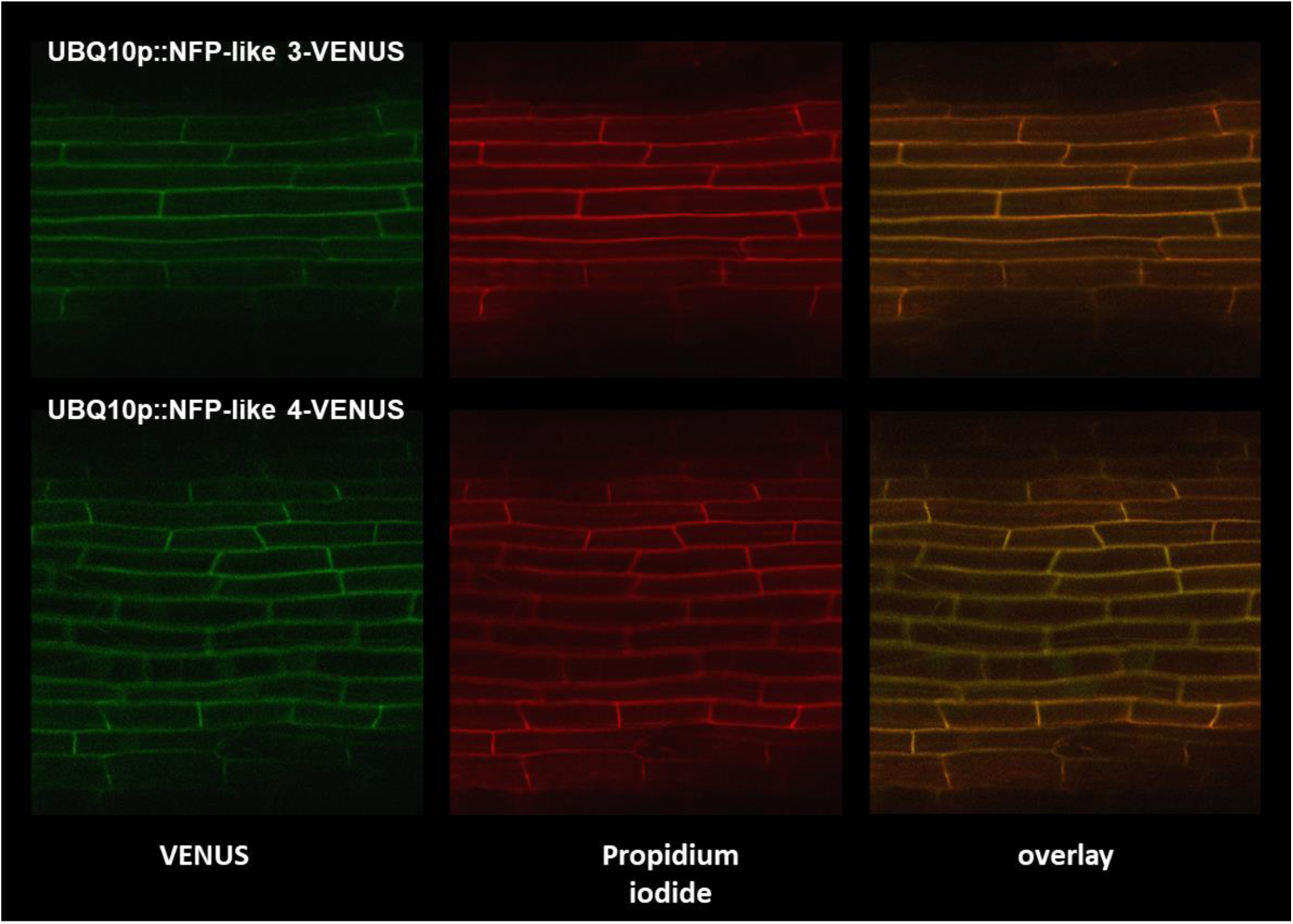
Subcellular localization of NFP-like 3 and NFP-like 4 receptors in poplar roots. Poplar leaves were transformed with *A. rhizogenes* carrying translational fusion constructs of NFP-like 3 and 4 fused to the VENUS fluorescent reporter under control of the *Arabidopsis* Ubiquitin 10 promoter. Protein localization was monitored in transformed roots emerging from poplar leaves. Both NFP-like proteins localize to the plasma membrane as indicated by the VENUS signal. To confirm localization, roots were stained with propidium iodide to highlight the plasma membrane.

**Supplementary Table T1.** Single mutations in the *P. alba* and *P. tremula* alleles of *P. x canescens* in genome edited *nfp-like 1,3,4* mutant lines. Shown are the modifications in the independent *nfp-like 1,3,4* mutant lines N483-8 and N484-3 depending on the modified candidate gene on cDNA level. g1-g4 are the guide RNAs used (see (Bruegmann et al., 2019) for details). For *NFP-like 1* a deletion was identified on the *alba* and *tremula* alleles leading to a stop codon at position 142 in the protein sequence reducing its length to only 22,9% of the full sequence. For *NFP-like 3* we found 2 transcript variants. In N483-8 and N848-3 *NFP-like 3* showed different modifications in the *alba* and *tremula* alleles. Analysis of the sequences demonstrated that insertions and deletions resulted in shortened amino acid sequences as well as changed reading frames that do not result in functional proteins. For *NFP-like 4* the homozygous insertion of thymine resulted in a stop codon at amino acid position 64 which represents only 10.3% of the total protein length.

| Line | DNA used for sequencing | *NFP-like 1* | *NFP-like 3* | *NFP-like 4* |
| --- | --- | --- | --- | --- |
| N483-8 | cDNA | **g1**: Homozygous deletion A on both alleles<br>**g2**: Wild type | **alba**<br>**g3**: Insertion A<br>**g4**: Deletion 74 nt<br>**tremula** (1):<br>**g3**: Insertion T<br>**g4**: Insertion T<br>**tremula** (2)<br>**g3**: Insertion A<br>**g4**: Deletion 74 nt | **g3**: Homozygous insertion A on both alleles<br>**g4**: Heterozygous monoallelic deletion ATAAG |
| N484-3 | cDNA | **g1**: Homozygous deletion A on both alleles<br>**g2**: Wild type | **alba**<br>**g3**: Insertion A<br>**g4**: Insertion A<br>**tremula** (1)<br>**g3**: Insertion T<br>**g4**: Deletion 52 nt<br>**tremula** (2)<br>**g3**: Insertion TT<br>**g4**: Deletion 52 nt | **g3**: Homozygous insertion T on both alleles<br>**g4**: Heterozygous monoallelic deletion CATAAGG |

## Acknowledgments

K.D., M. F., C.H. and S.W. were supported by a grant for the project CHITOPOP (2016-2019) from the BMFTR to the Thuenen Institute (Project ID: 031B0203B) and the Leibniz Institute for Plant Biochemistry (IPB) (Project ID: 031B0203C and D). M.F. and S.W. thank the CHITOPOP consortium partners from the University of Göttingen, Department of Plant Cell Biology, especially Thomas Teichmann for coordination. T.B.I. and M.T. were funded by the Department of Energy (DOE) grant DE-SC0018247 to J.M.A. S.N. was supported by the Hatch grant (#WIS05052) to J.M.A. T.A.R. was supported by the Genomic System Sciences Program, U.S. Department of Energy, Office of Science, Biological and Environmental Research, as part of the Plant-Microbe Interfaces Scientific Focus Area at the Oak Ridge National Laboratory (http://pmi.ornl.gov), which is supported by the Office of Science of the U.S. Department of Energy under Contract No. DE-AC05-00OR22725. E.S.W was funded by a grant from the Austrian Academy of Sciences to the Gregor Mendel Institute and a European Research Council Advanced Grant DENOVO-P (project no. 787613) to Liam Dolan. S.F. acknowledges NanoBio ICMG (UAR 2607) for providing access to mass spectrometry and NMR facilities and the French National Research Agency-ANR through LABEX ARCANE/EUR CBH-EUR-GS(ANR-17-EURE-0003), Glyc@Alps (ANR-15-IDEX-02), and Carnot Polynat (CARN-025-01) for partial financial support. We also thank Francis Martin for providing fungus material (*L. bicolor*). S.W. also thanks Andrea Polle and Bettina Hause for critical discussions and hosting the CHITOPOP project at University Göttingen and IPB as well as Melissa Reimann and Annett van Lengen for sampling and lab support.

## Contributions

S.W., M.F. and J.M.A conceptualized the project. K.O.M., K.D., T.B.I. and S.W. carried out the experiments. T.A.R. conducted the phenotypic analysis of root tips. E.S.W. did the cloning of constructs. J.K. developed and wrote the program for semi-automatic nuclear spike counting. M.T. carried out RNASeq evaluation. S.F. synthesized LCOs. C.H. developed the setup and carried out sampling for roots for RNASeq. J. M.A., S.N. and M.T. conducted domain modeling and sequence analysis. K.O.M., J. M.A., T.B.I., T.A.R. and S.W. wrote the manuscript.

## Data availability

Calcium spiking videos are stored in the open repository Zenodo under https://zenodo.org/records/17776475 and will be openly available after manuscript acceptance.

