## Supplementary data Calcium spiking for "LysM Receptor Proteins are Required for Ectomycorrhizal Symbiosis in Poplar": profiles-spiking-1.pdf

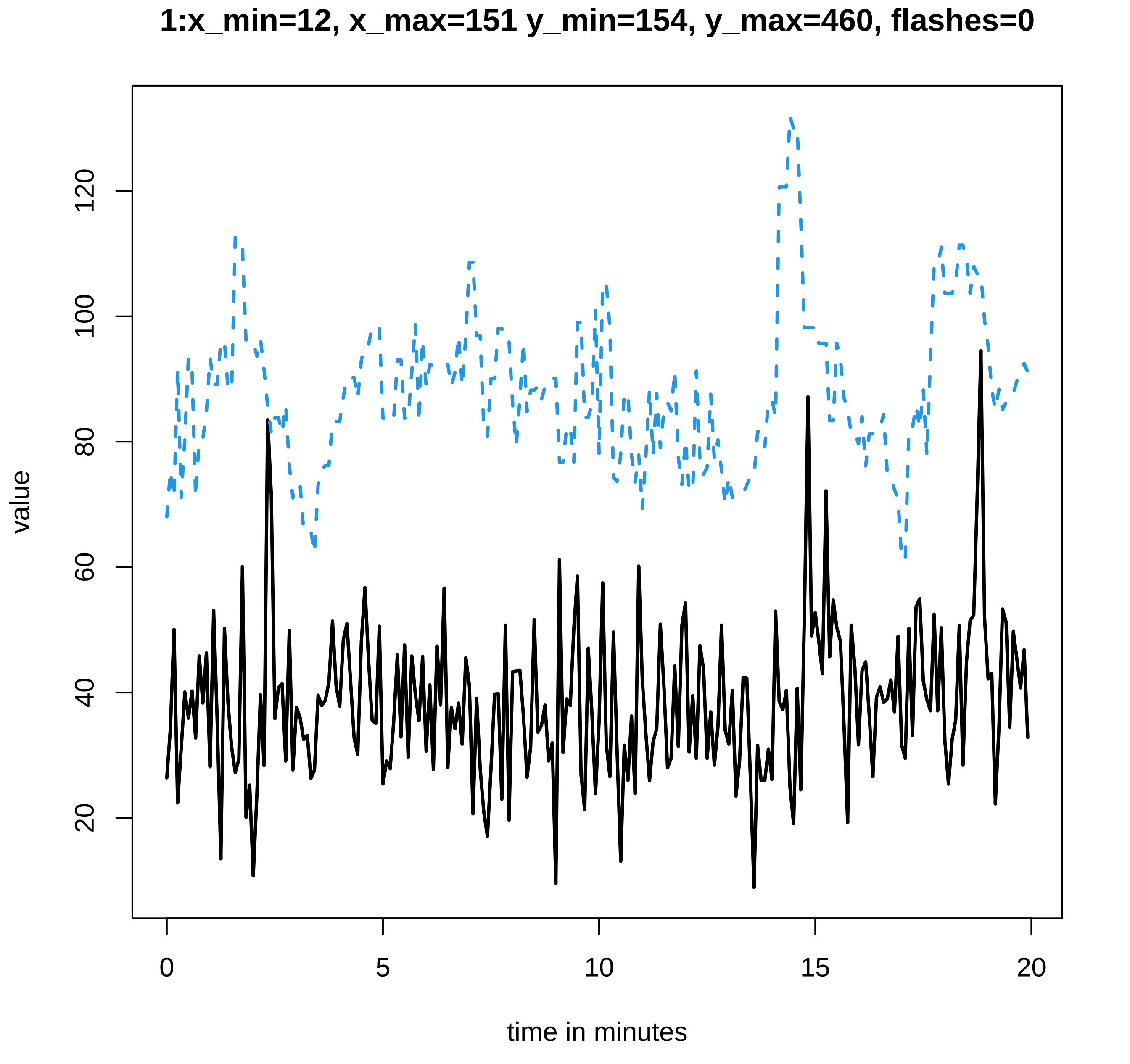
1:x_min=12, x_max=151 y_min=154, y_max=460, flashes=0
value
20
40
60
80
100
120
0
5
10
15
20
time in minutes

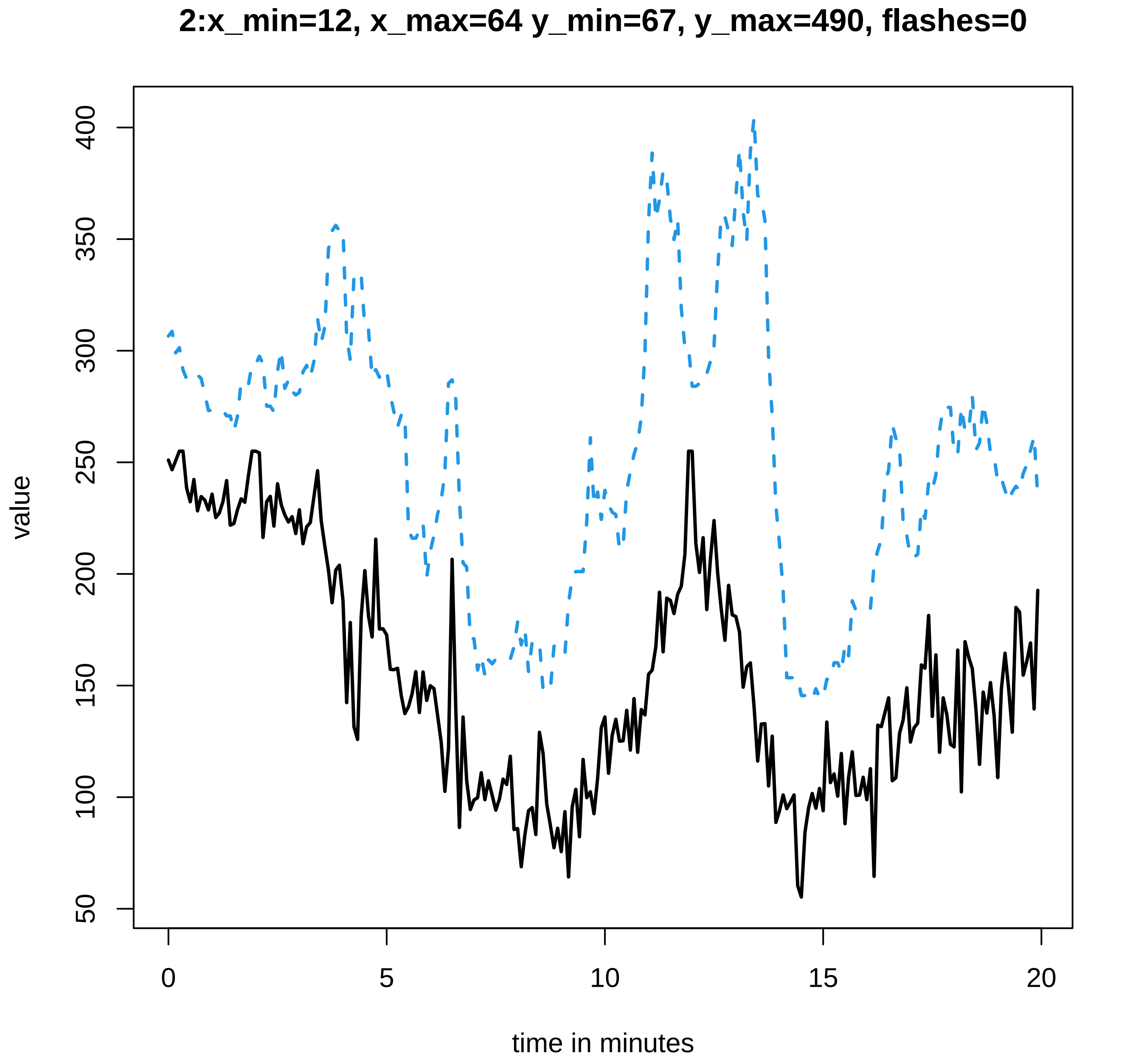
2:x_min=12, x_max=64 y_min=67, y_max=490, flashes=0
value
50
100
150
200
250
300
350
400
0
5
10
15
20
time in minutes

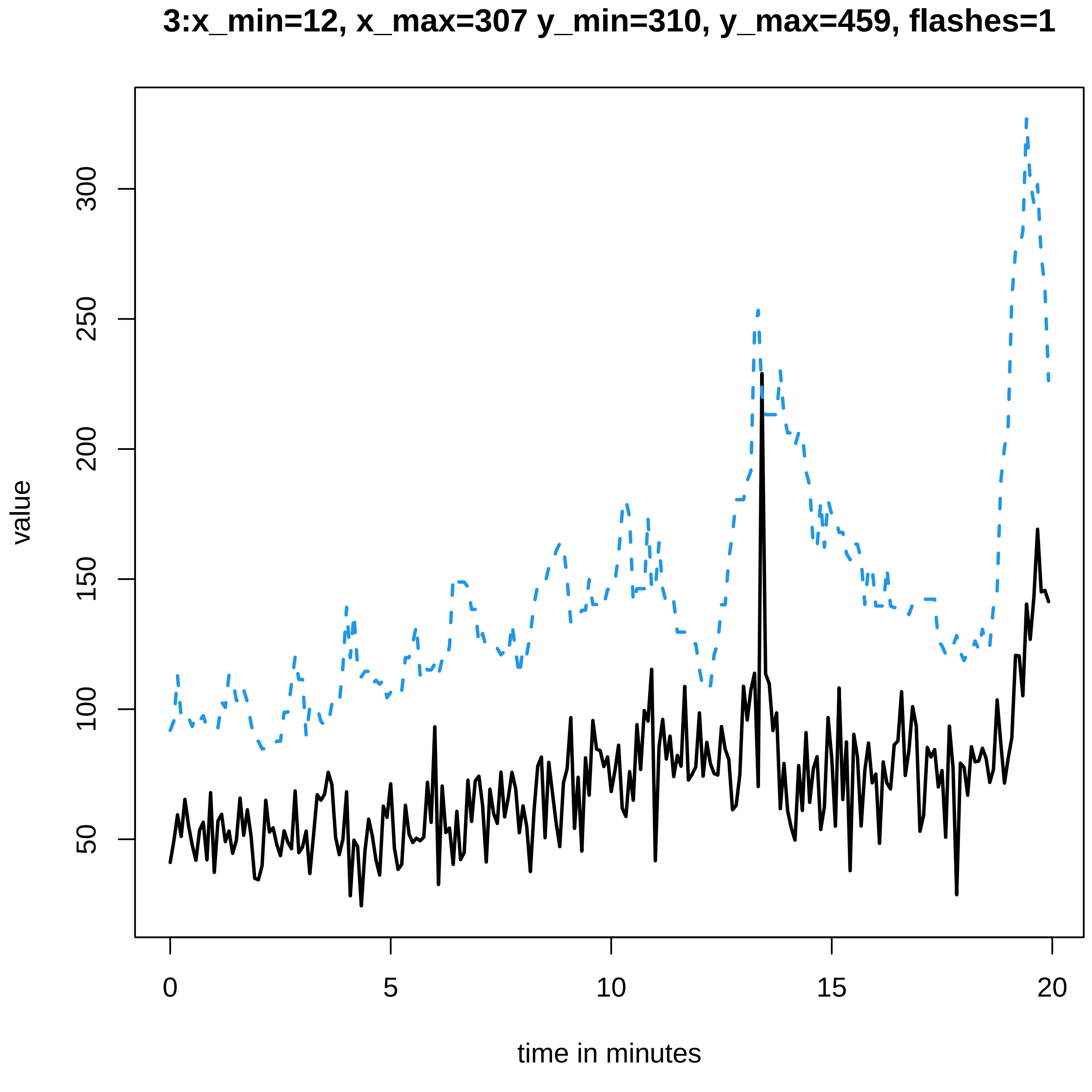
3:x_min=12, x_max=307 y_min=310, y_max=459, flashes=1
value
50
100
150
200
250
300
0
5
10
15
20
time in minutes

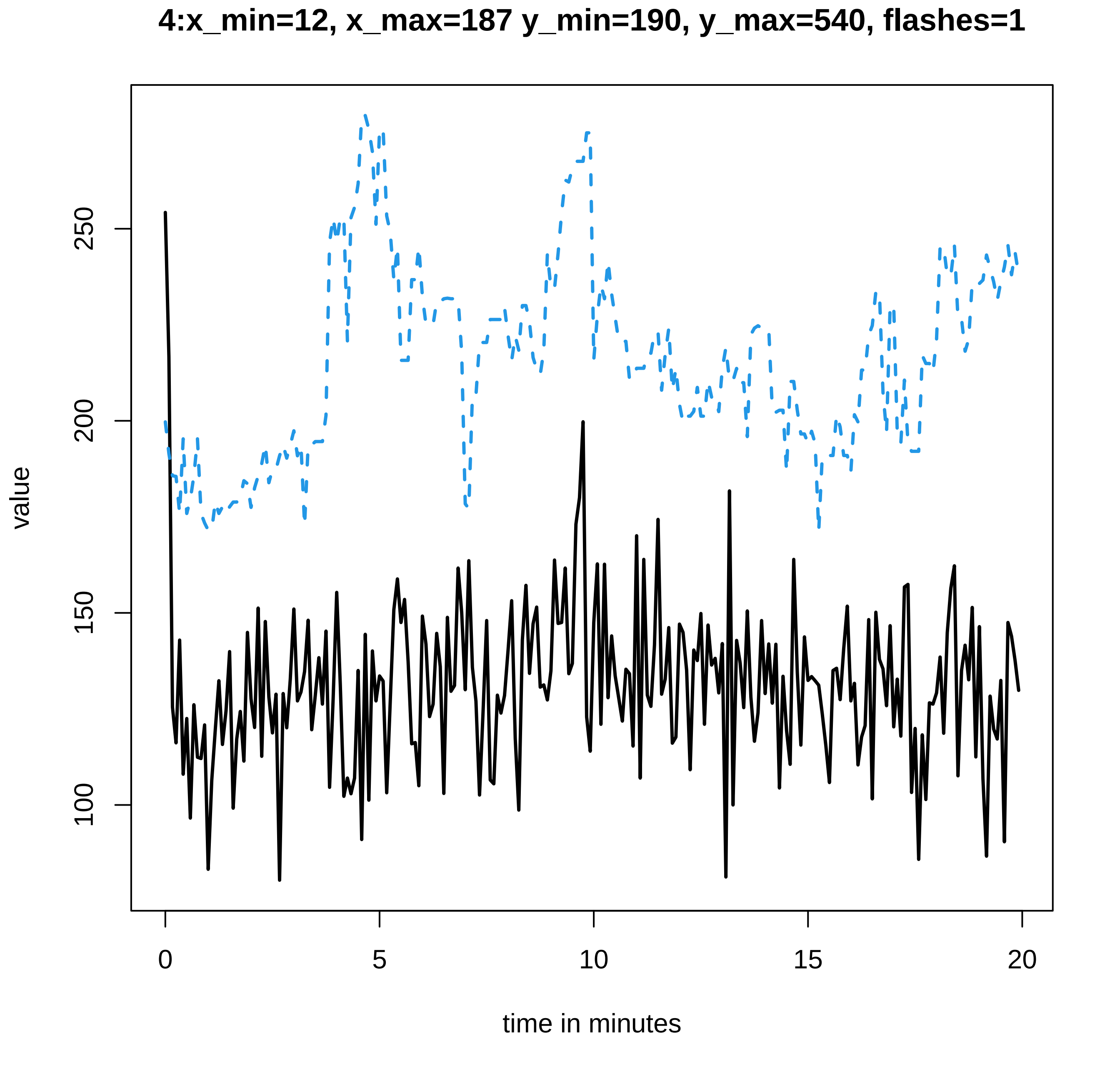

4:x_min=12, x_max=187 y_min=190, y_max=540, flashes=1
value
250
200
150
100
0
5
10
15
20
time in minutes

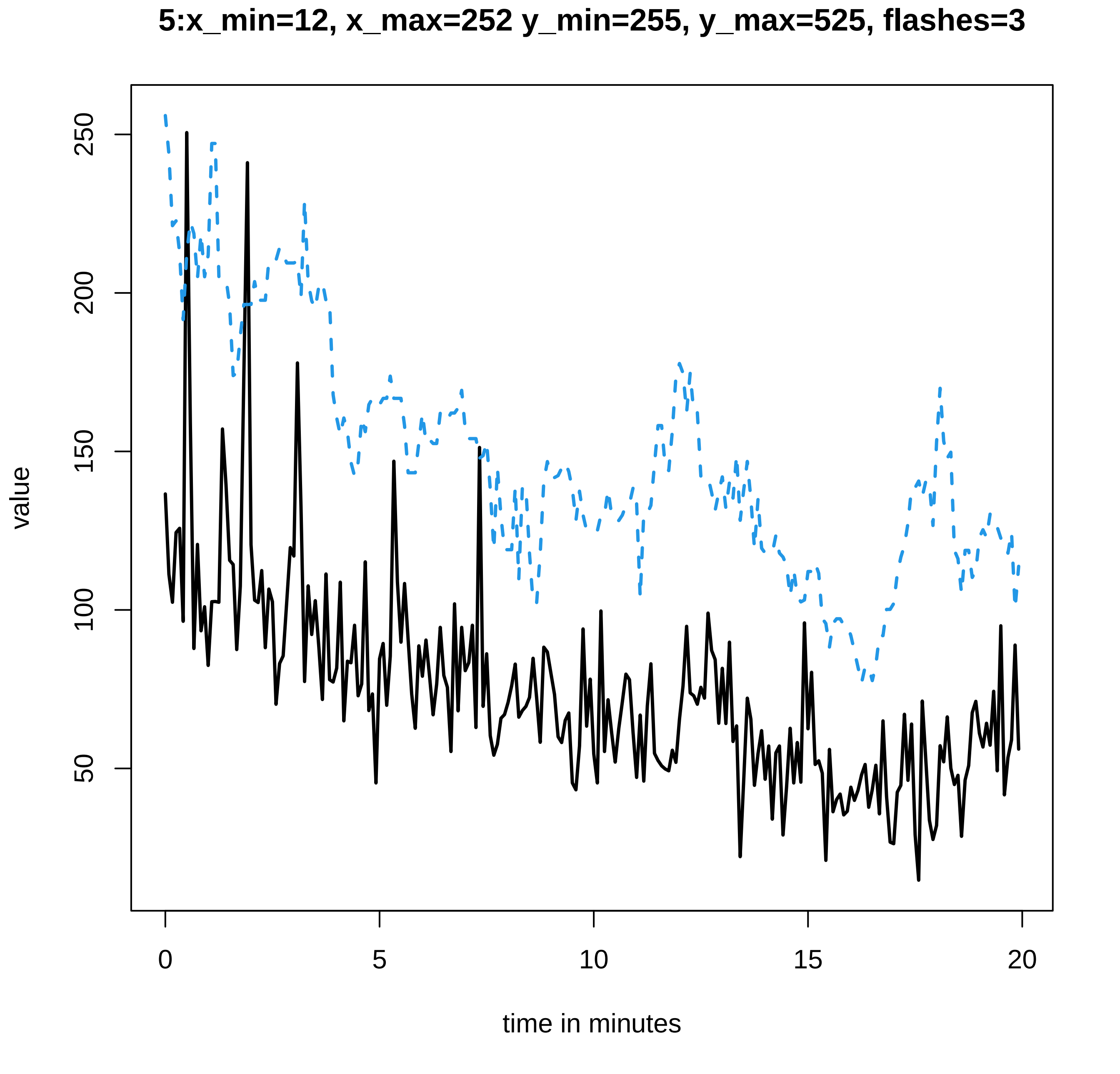
5:x_min=12, x_max=252 y_min=255, y_max=525, flashes=3
value
250
200
150
100
50
0
5
10
15
20
time in minutes

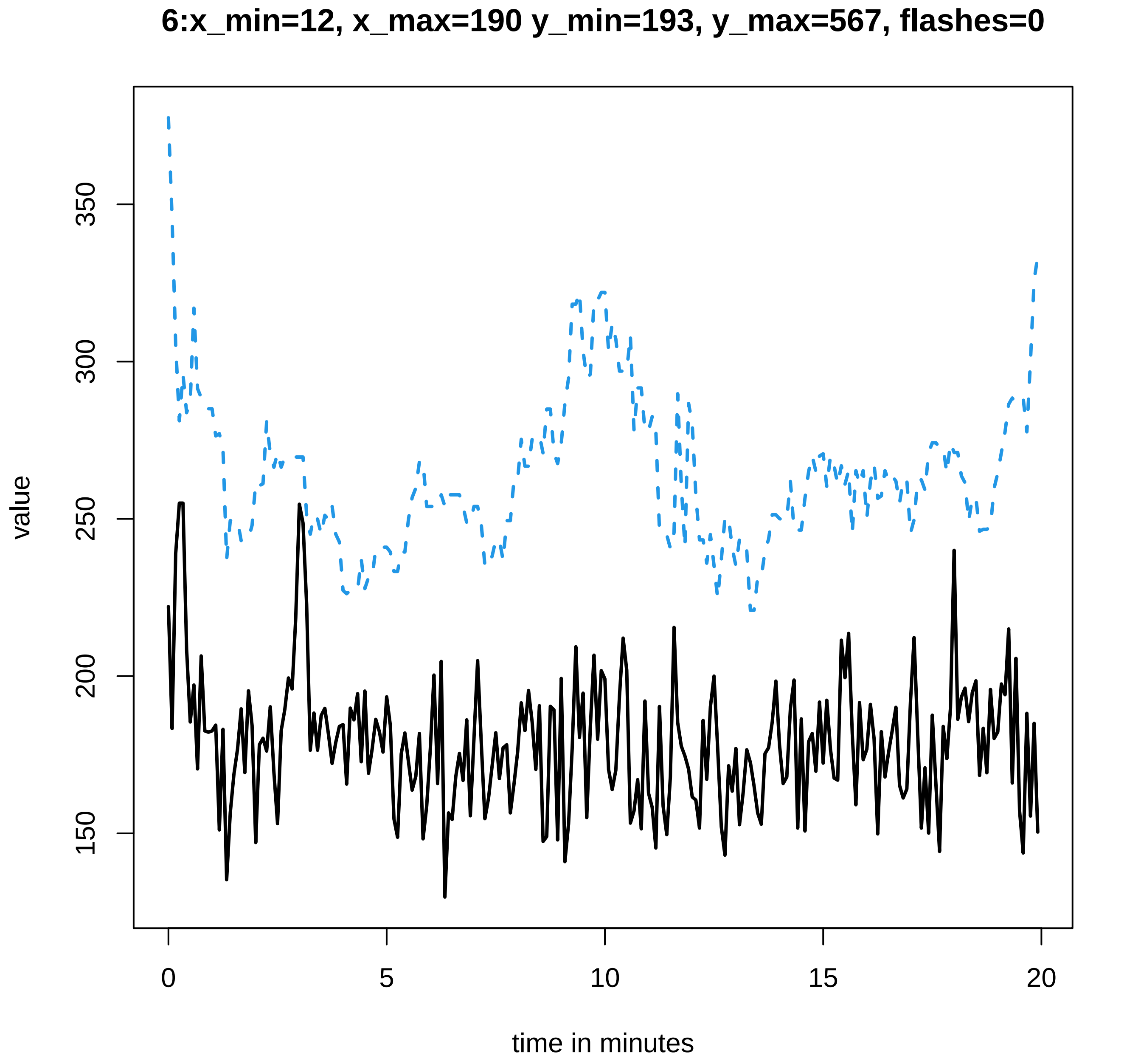
6:x_min=12, x_max=190 y_min=193, y_max=567, flashes=0
value
150
200
250
300
350
0
5
10
15
20
time in minutes

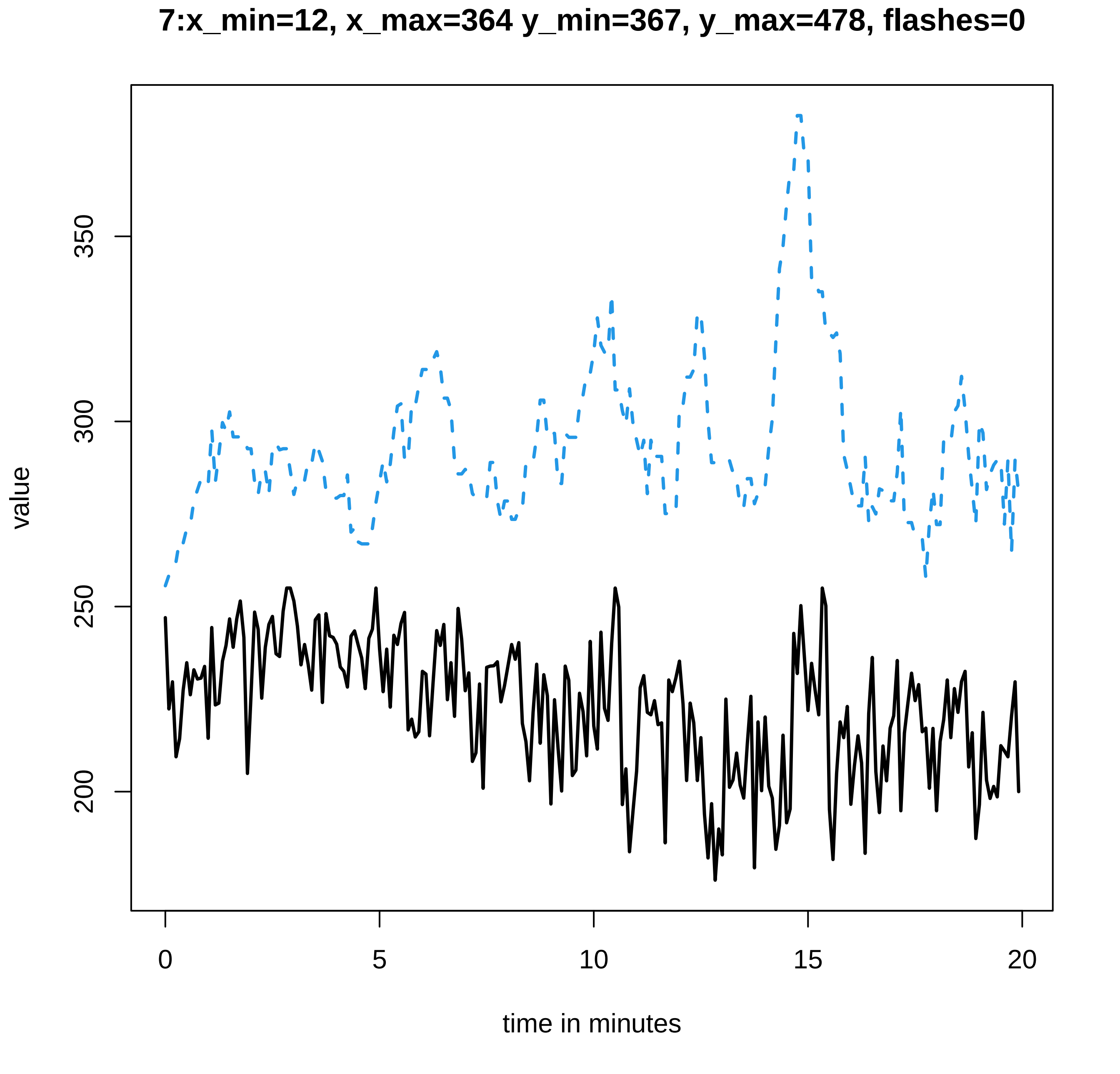
7:x_min=12, x_max=364 y_min=367, y_max=478, flashes=0
value
200
250
300
350
0
5
10
15
20
time in minutes

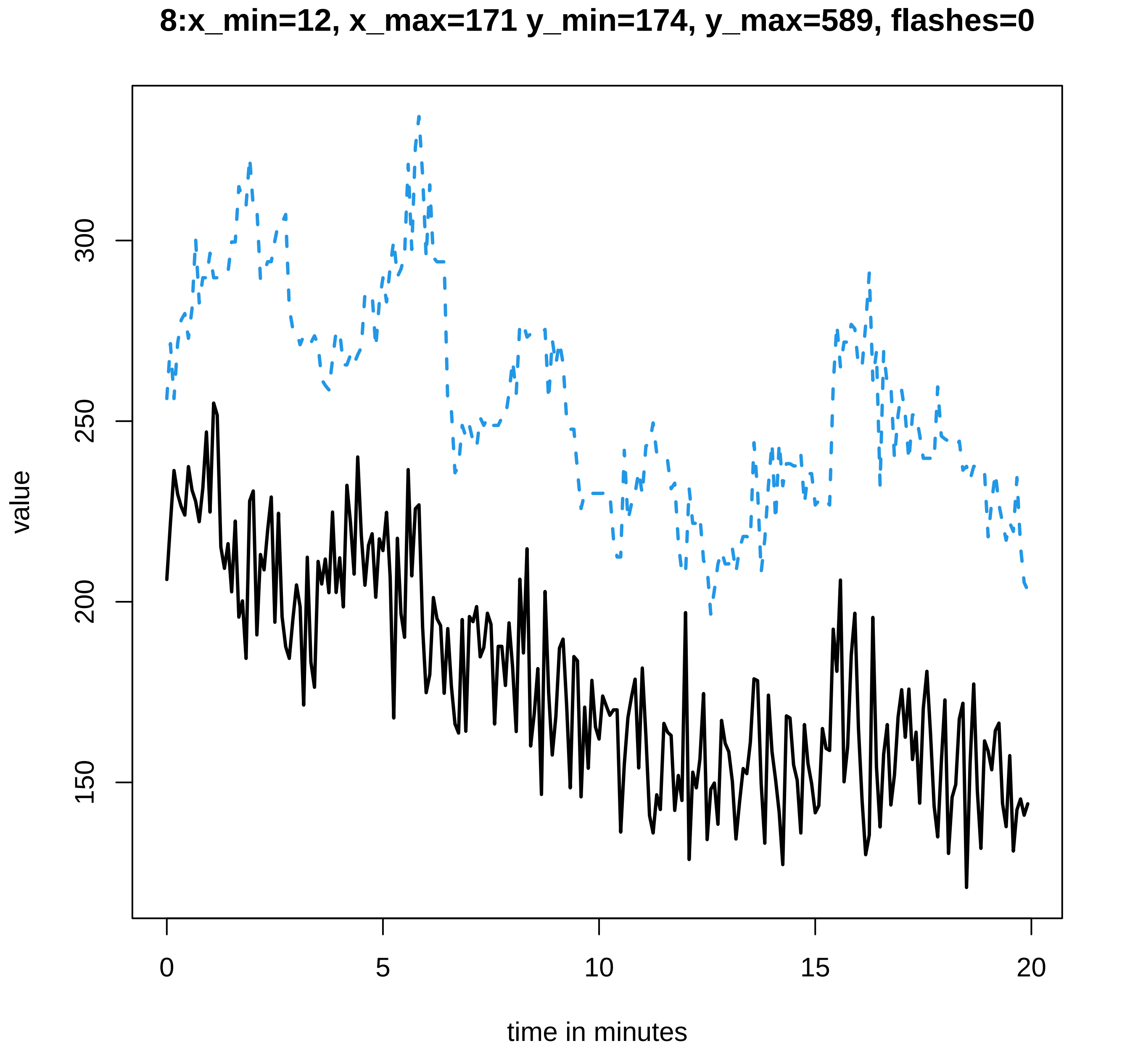
8:x_min=12, x_max=171 y_min=174, y_max=589, flashes=0
value
150
200
250
300
0
5
10
15
20
time in minutes

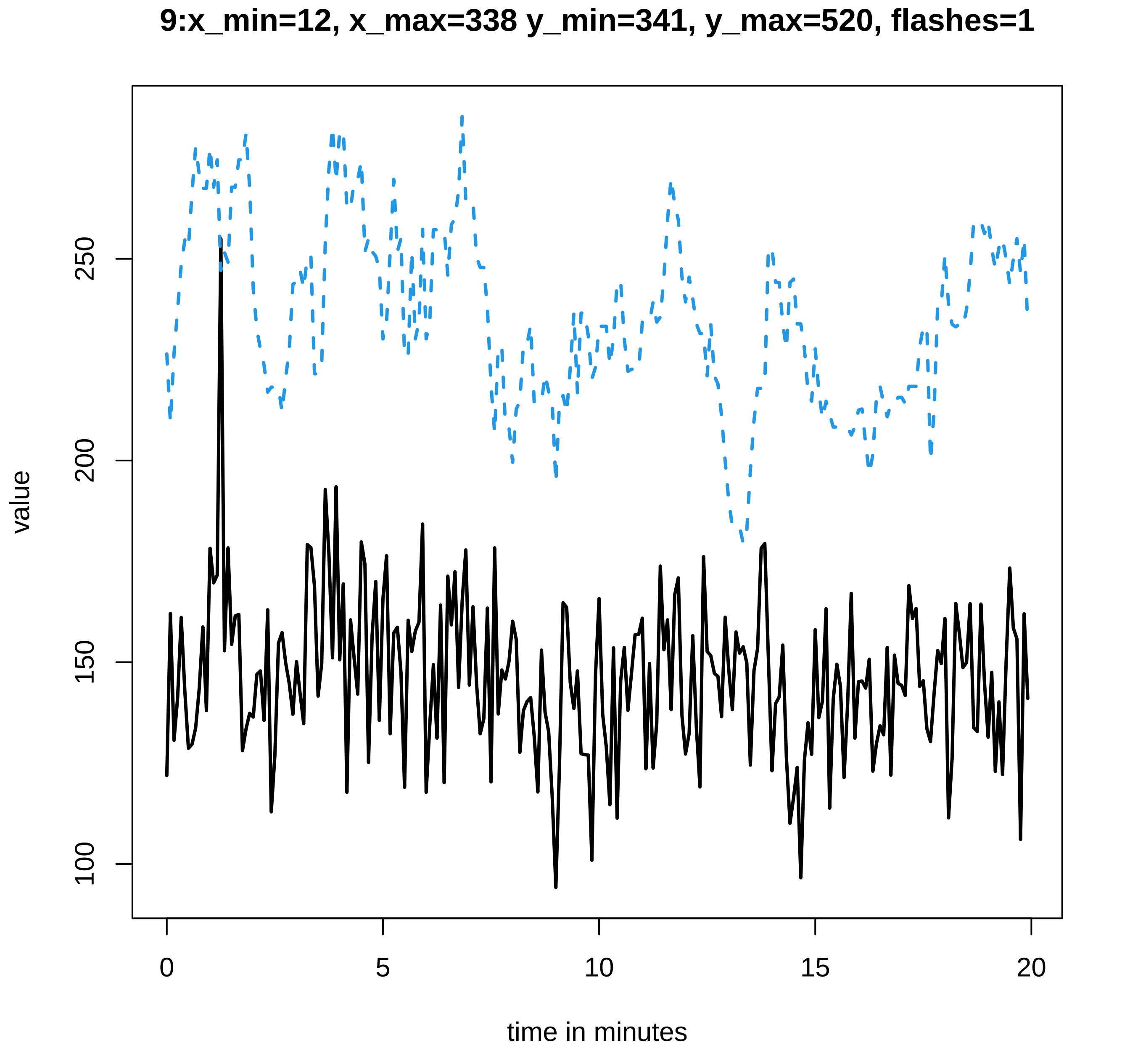

9:x_min=12, x_max=338 y_min=341, y_max=520, flashes=1
value
250
200
150
100
0
5
10
15
20
time in minutes

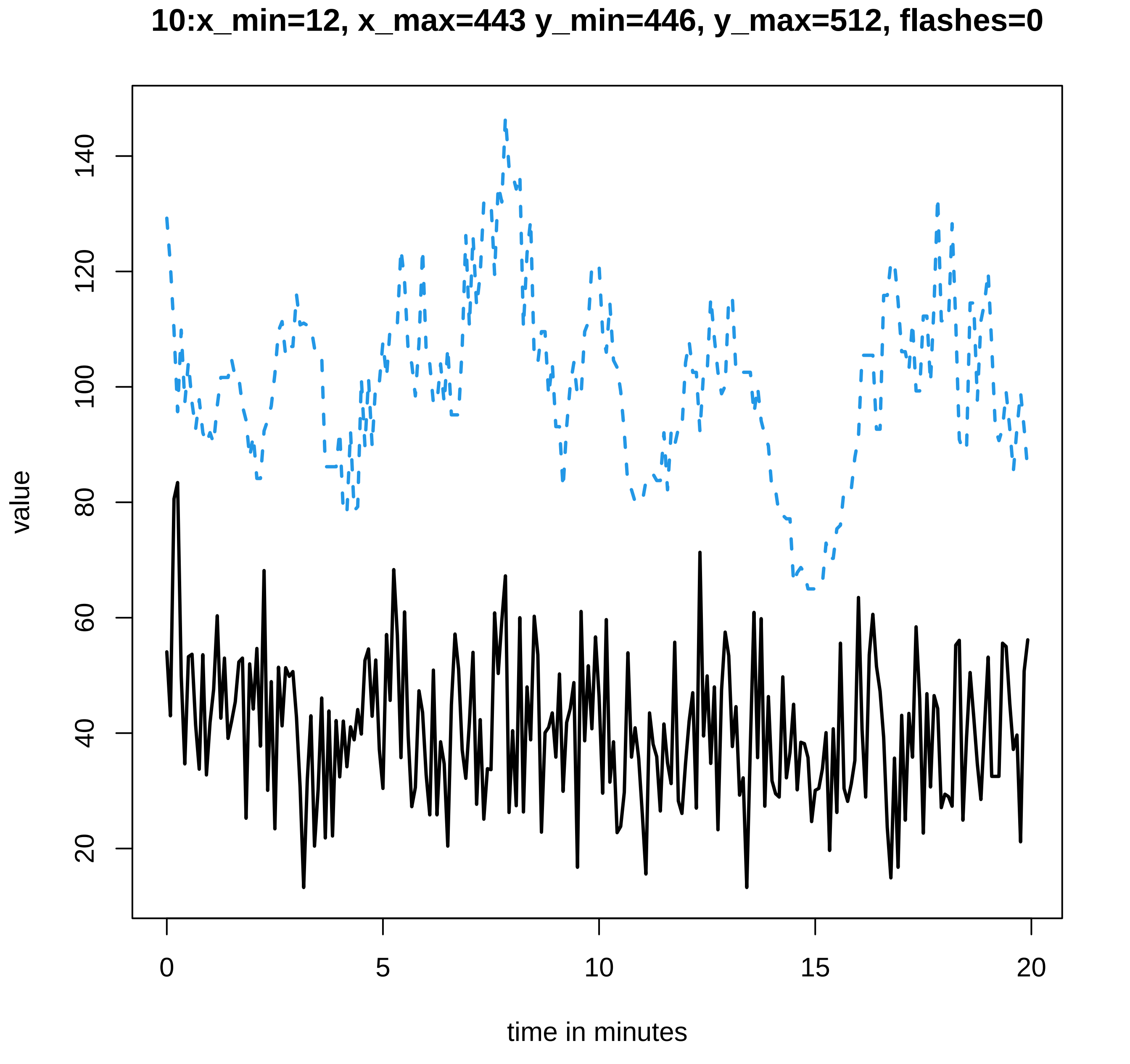
10:x_min=12, x_max=443 y_min=446, y_max=512, flashes=0
value
20
40
60
80
100
120
140
0
5
10
15
20
time in minutes

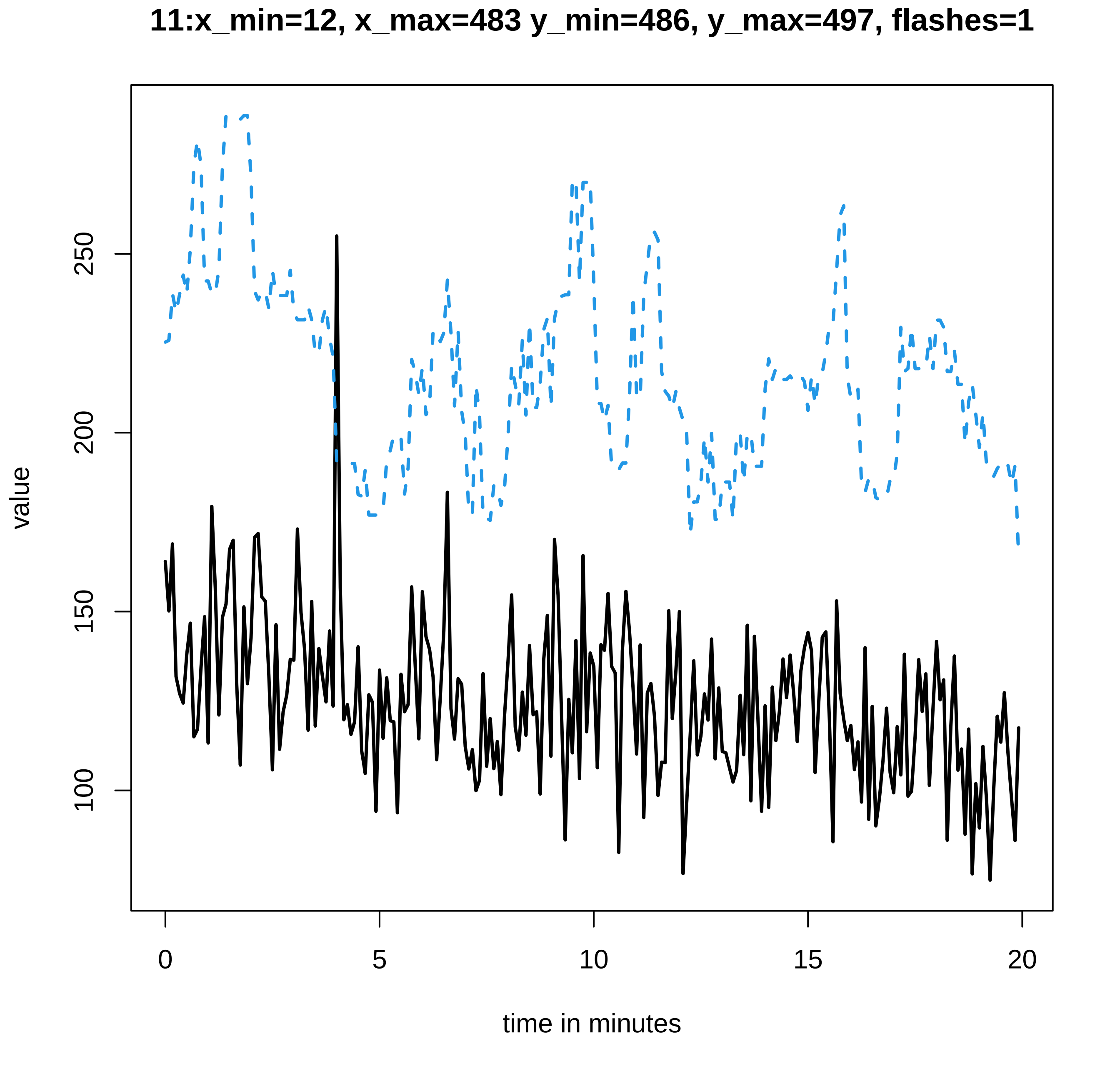

11:x_min=12, x_max=483 y_min=486, y_max=497, flashes=1
value
100
150
200
250
0
5
10
15
20
time in minutes

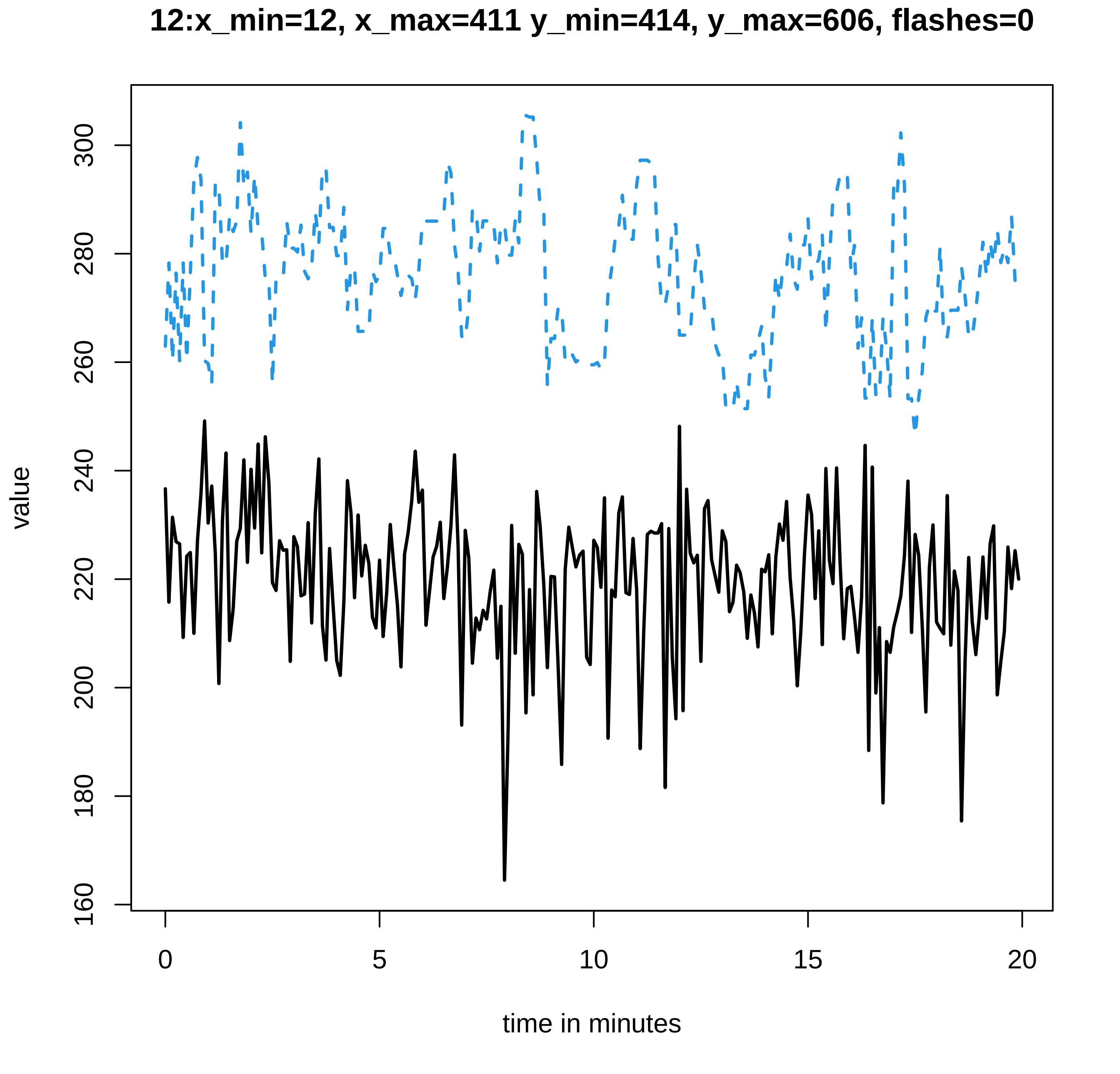

12:x_min=12, x_max=411 y_min=414, y_max=606, flashes=0
value
160
180
200
220
240
260
280
300
0
5
10
15
20
time in minutes

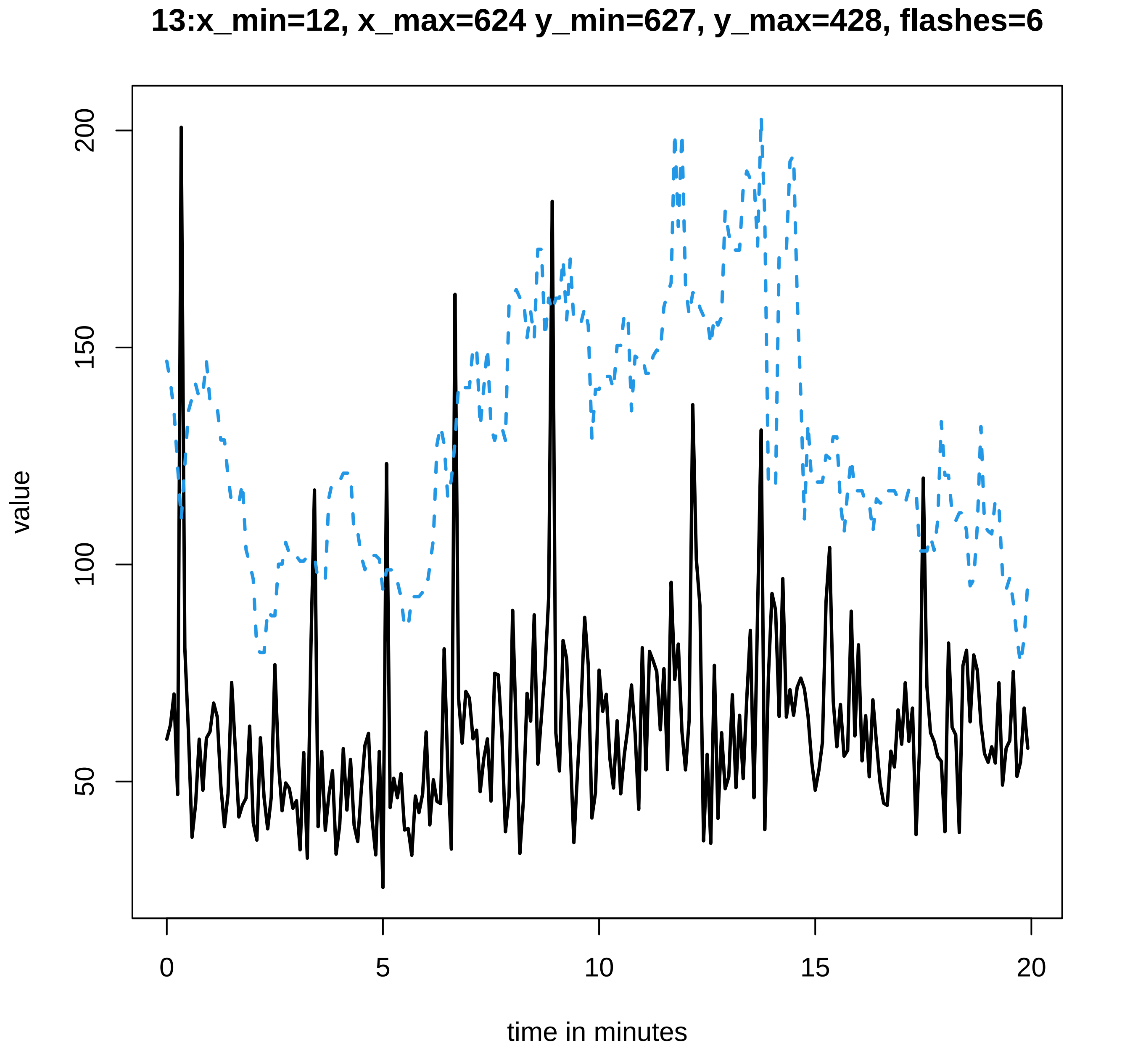
13:x_min=12, x_max=624 y_min=627, y_max=428, flashes=6
value
200
150
100
50
0
5
10
15
20
time in minutes

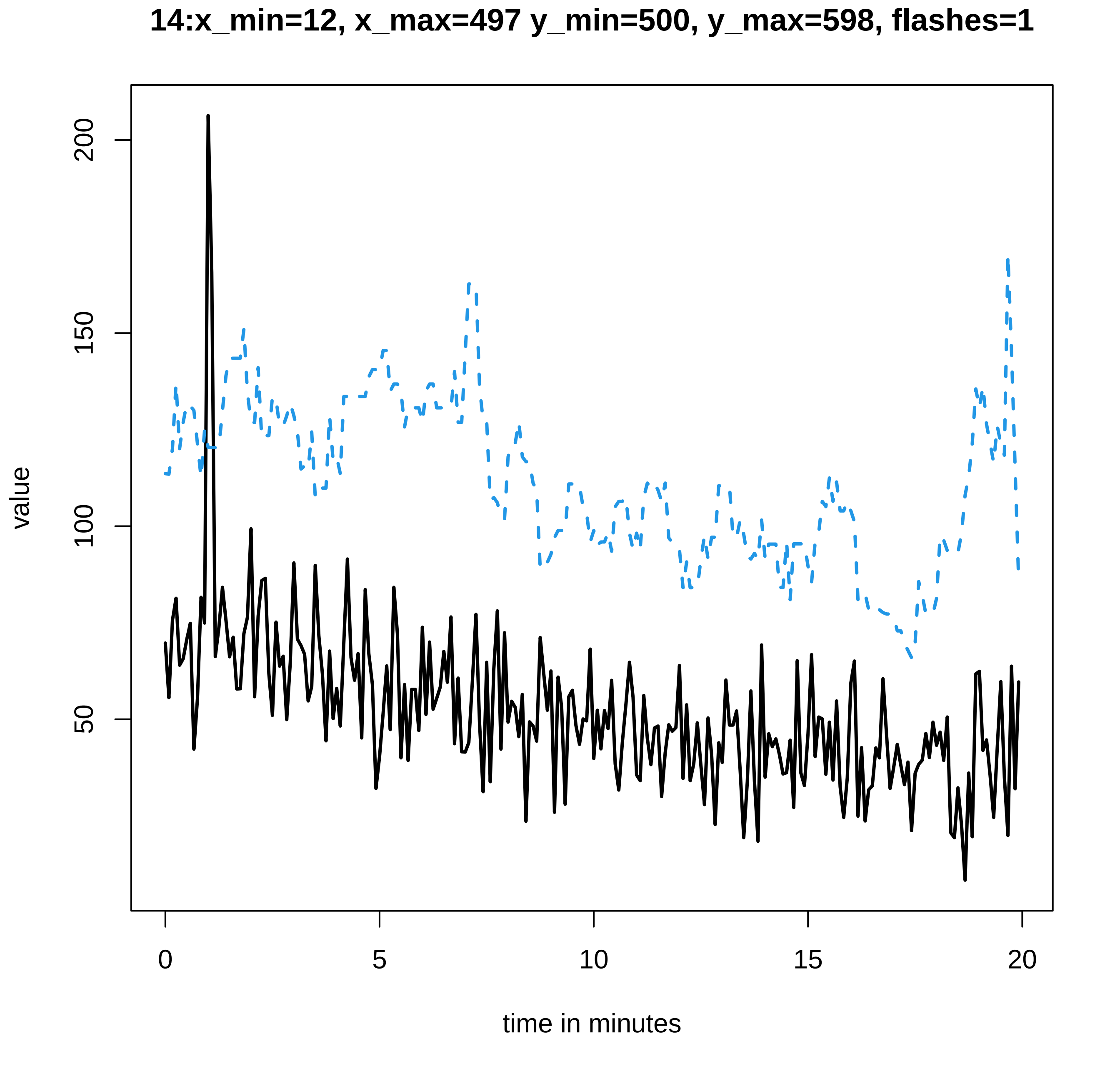

14:x_min=12, x_max=497 y_min=500, y_max=598, flashes=1
value
200
150
100
50
0
5
10
15
20
time in minutes

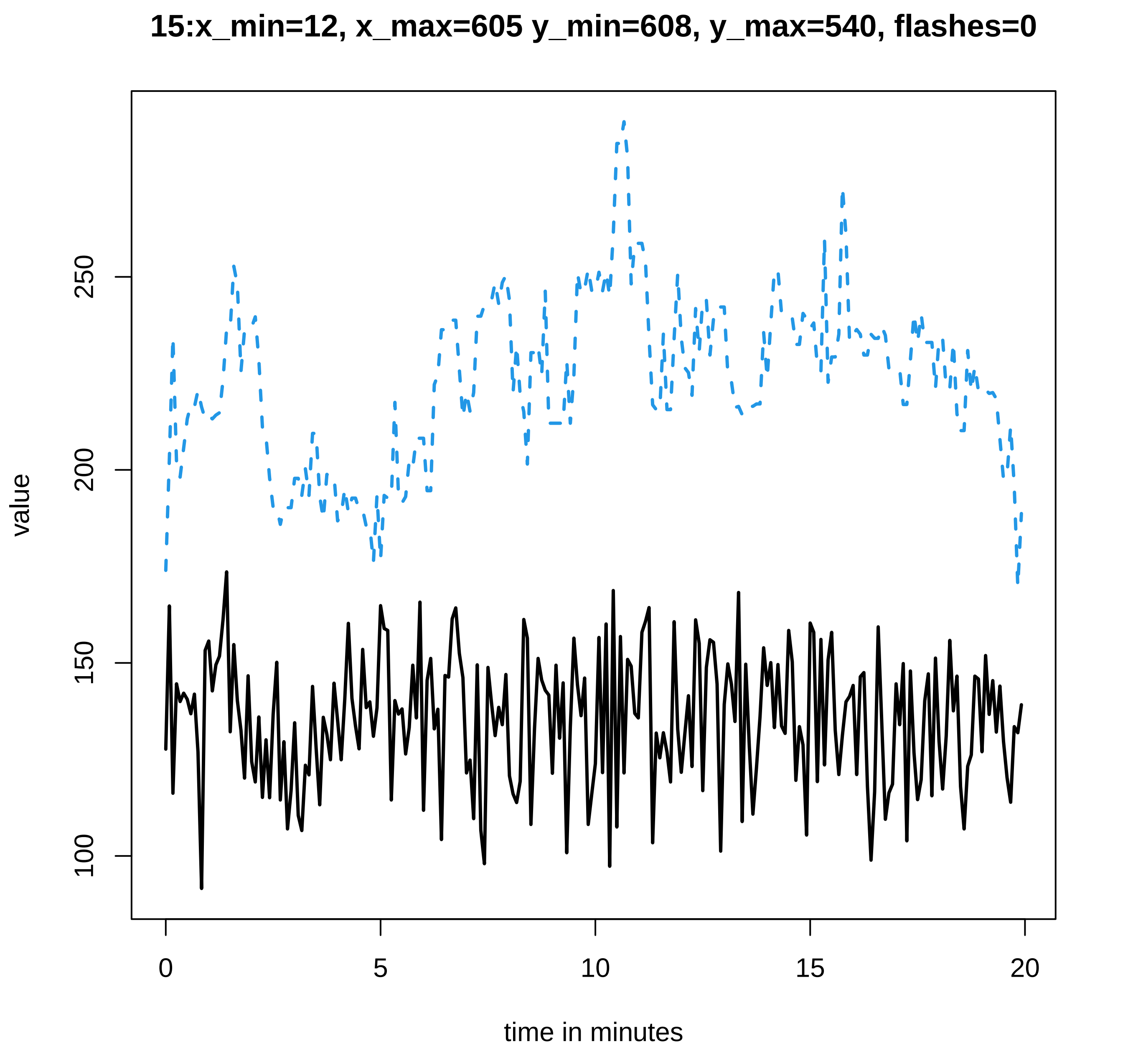
15:x_min=12, x_max=605 y_min=608, y_max=540, flashes=0
value
100
150
200
250
0
5
10
15
20
time in minutes

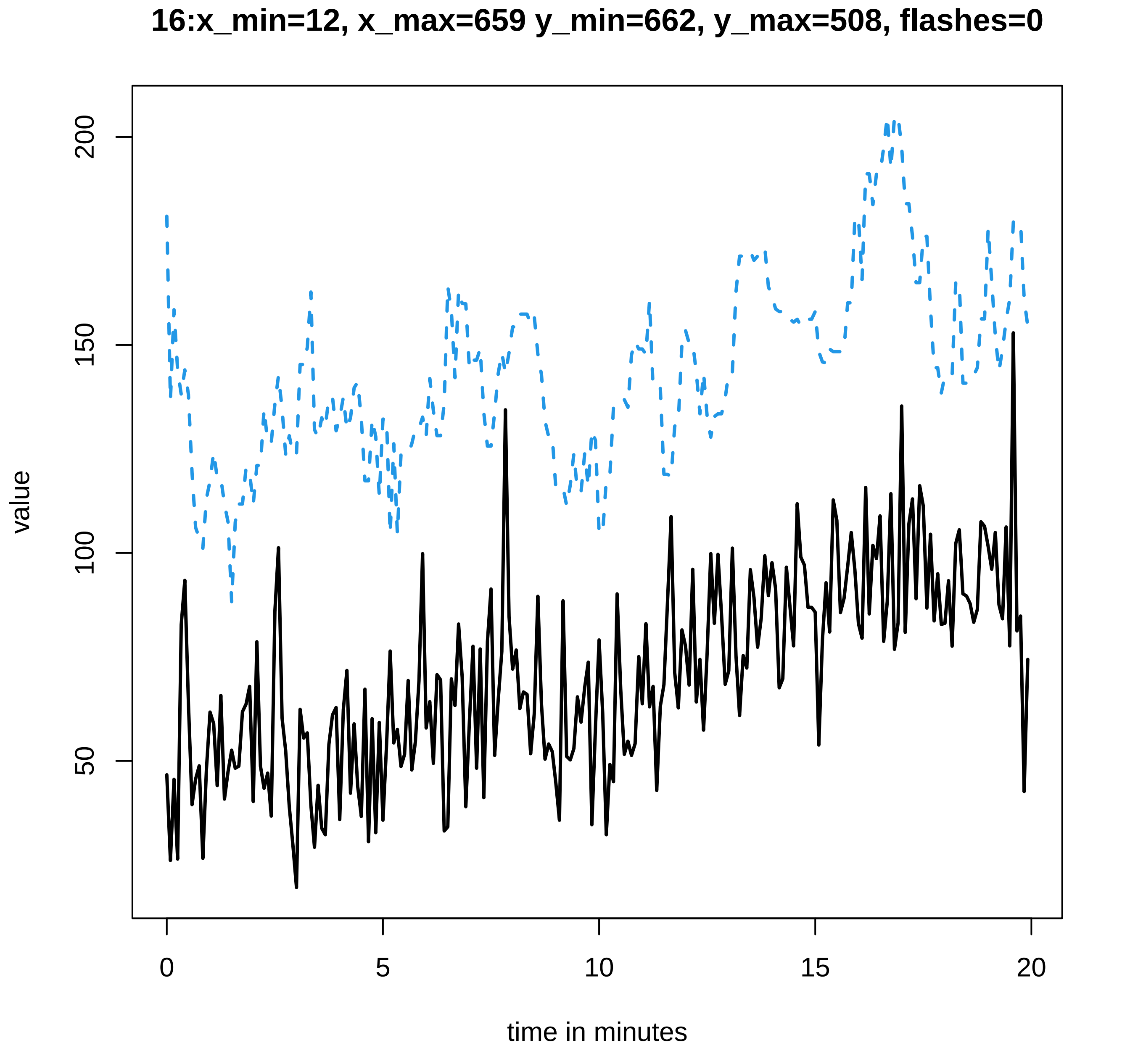

16:x_min=12, x_max=659 y_min=662, y_max=508, flashes=0
value
200
150
100
50
0
5
10
15
20
time in minutes

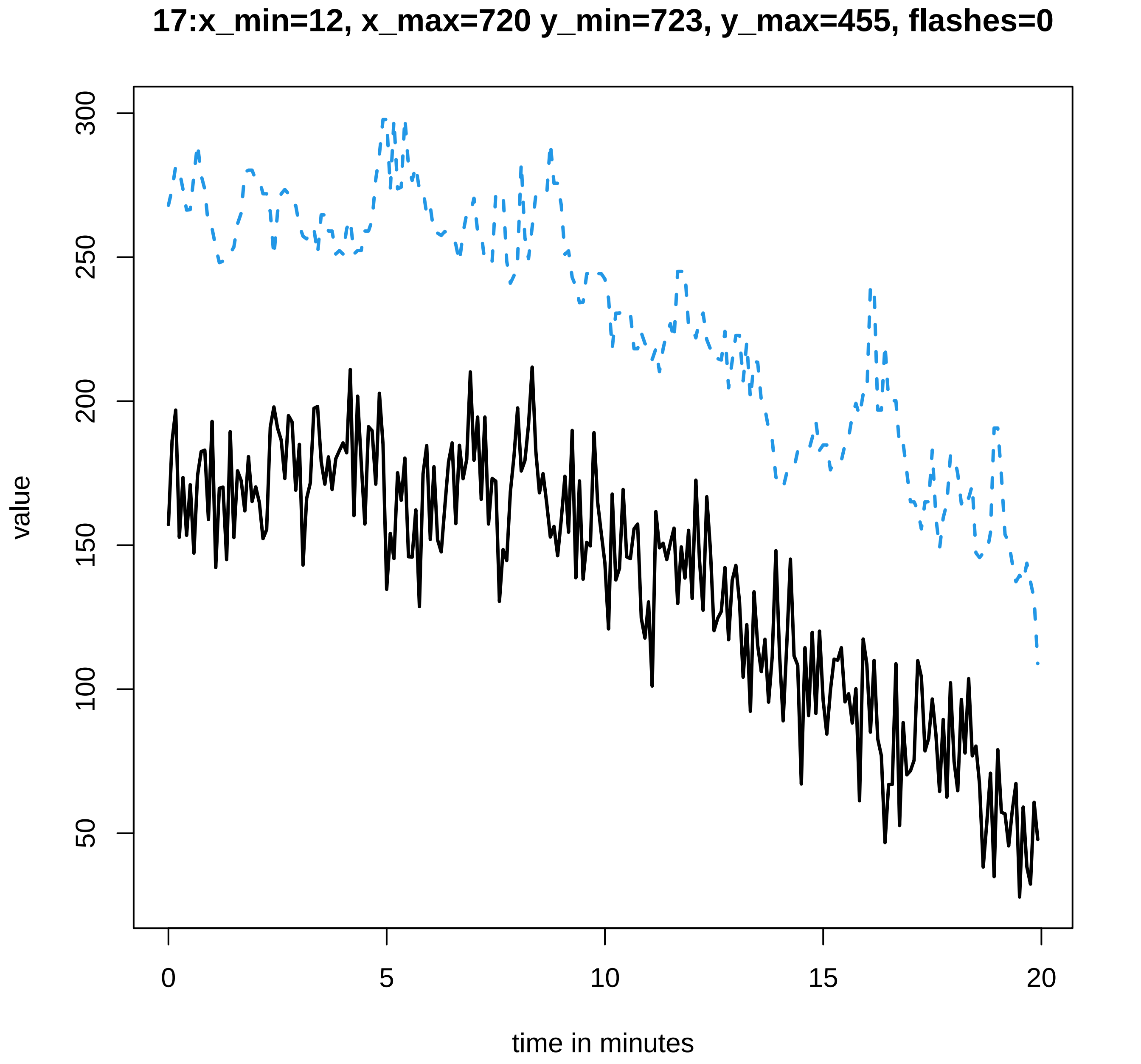
17:x_min=12, x_max=720 y_min=723, y_max=455, flashes=0
value
300
250
200
150
100
50
0
5
10
15
20
time in minutes

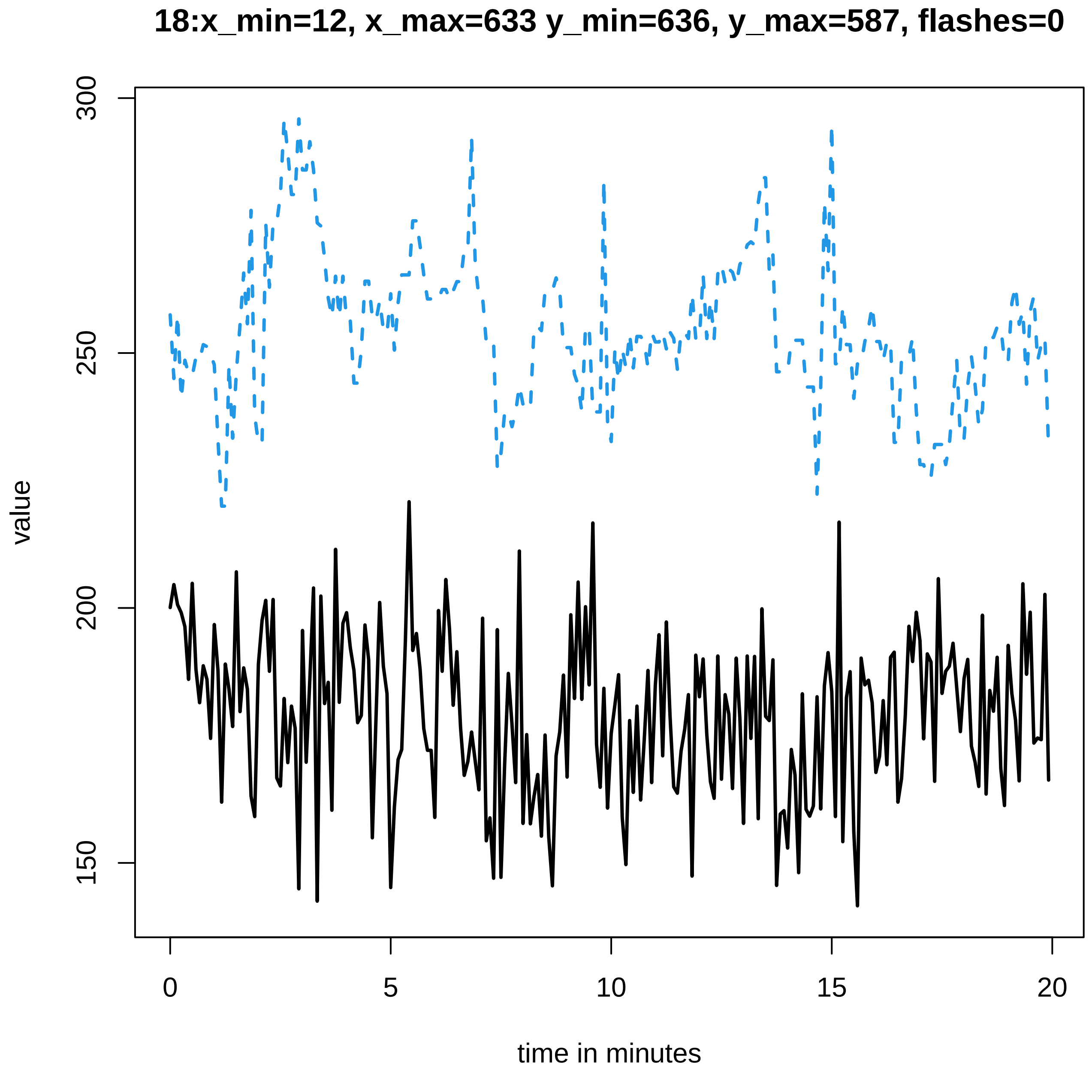

18:x_min=12, x_max=633 y_min=636, y_max=587, flashes=0
value
300
250
200
150
0
5
10
15
20
time in minutes

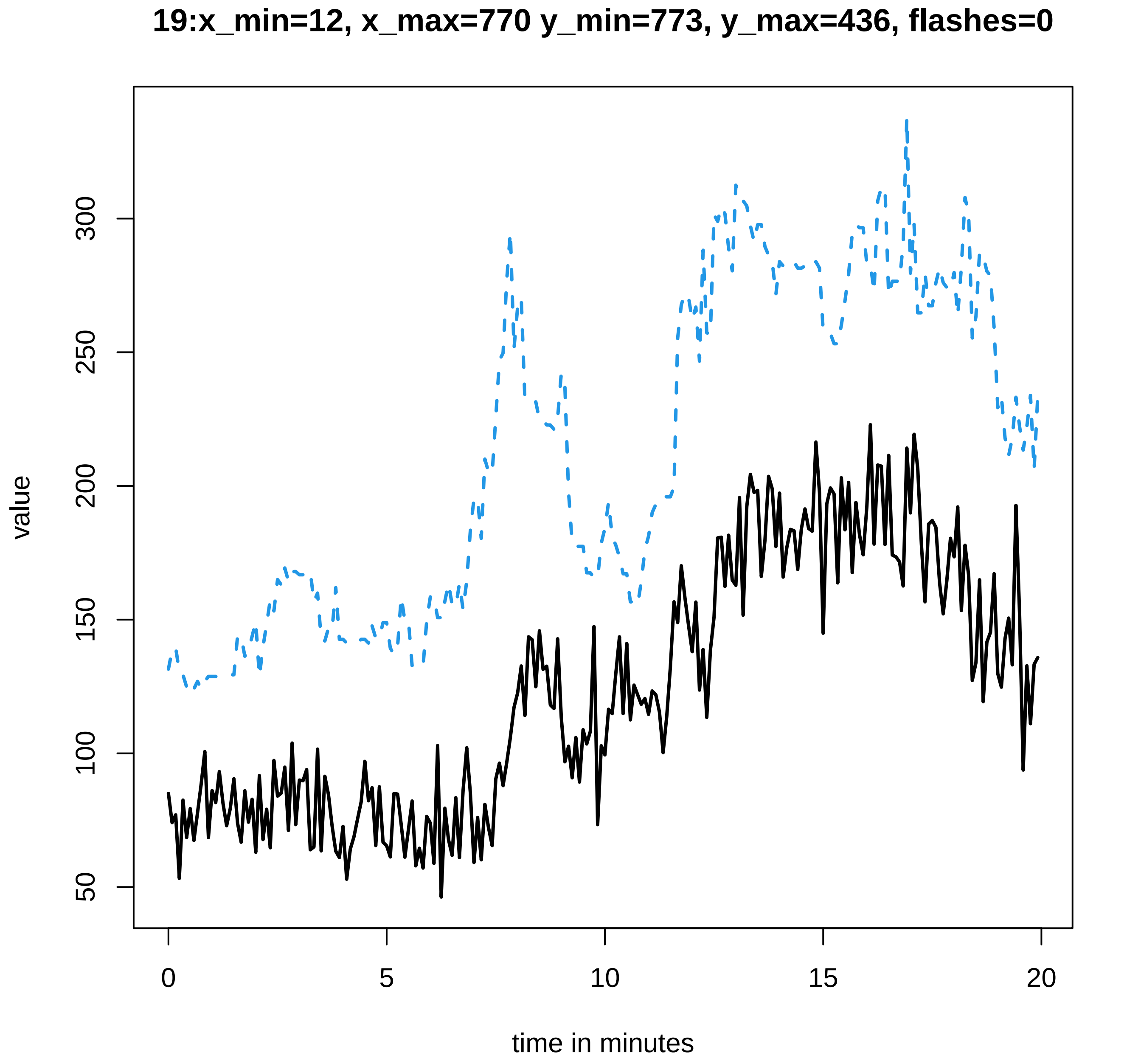
19:x_min=12, x_max=770 y_min=773, y_max=436, flashes=0
value
300
250
200
150
100
50
0
5
10
15
20
time in minutes

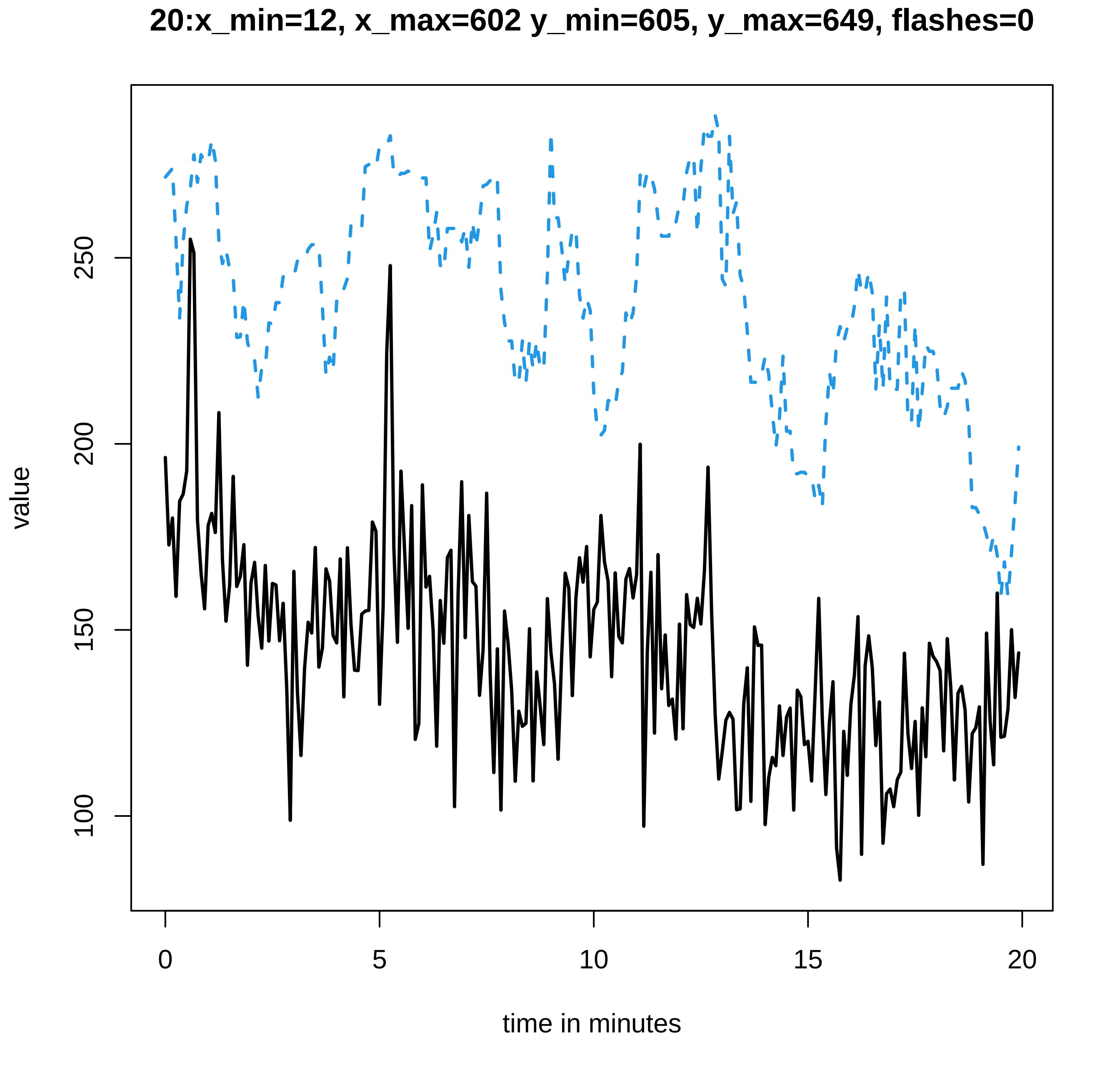

20:x_min=12, x_max=602 y_min=605, y_max=649, flashes=0
value
100
150
200
250
0
5
10
15
20
time in minutes

#### 21:x_min=12, x_max=649 y_min=652, y_max=621, flashes=1

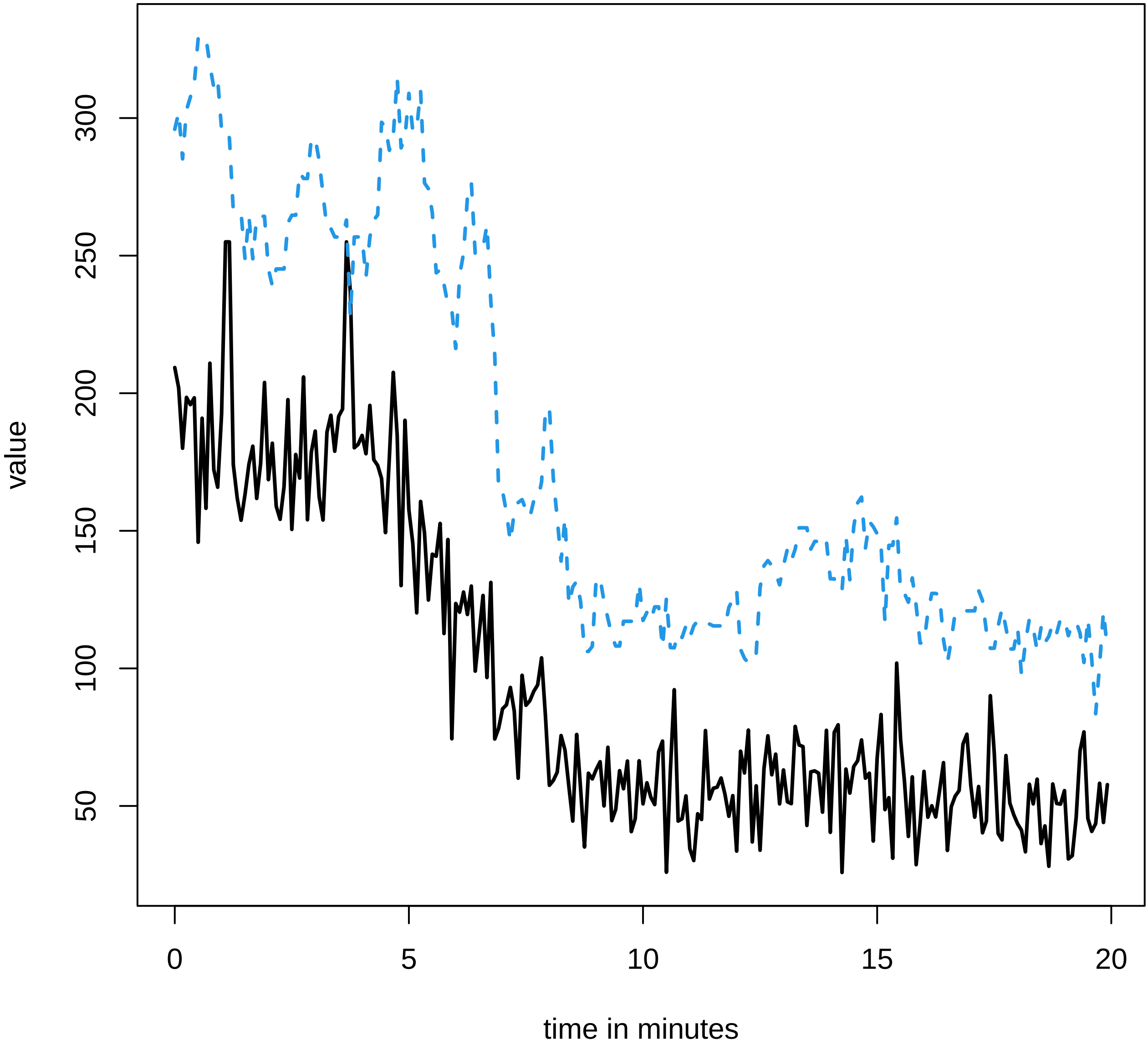

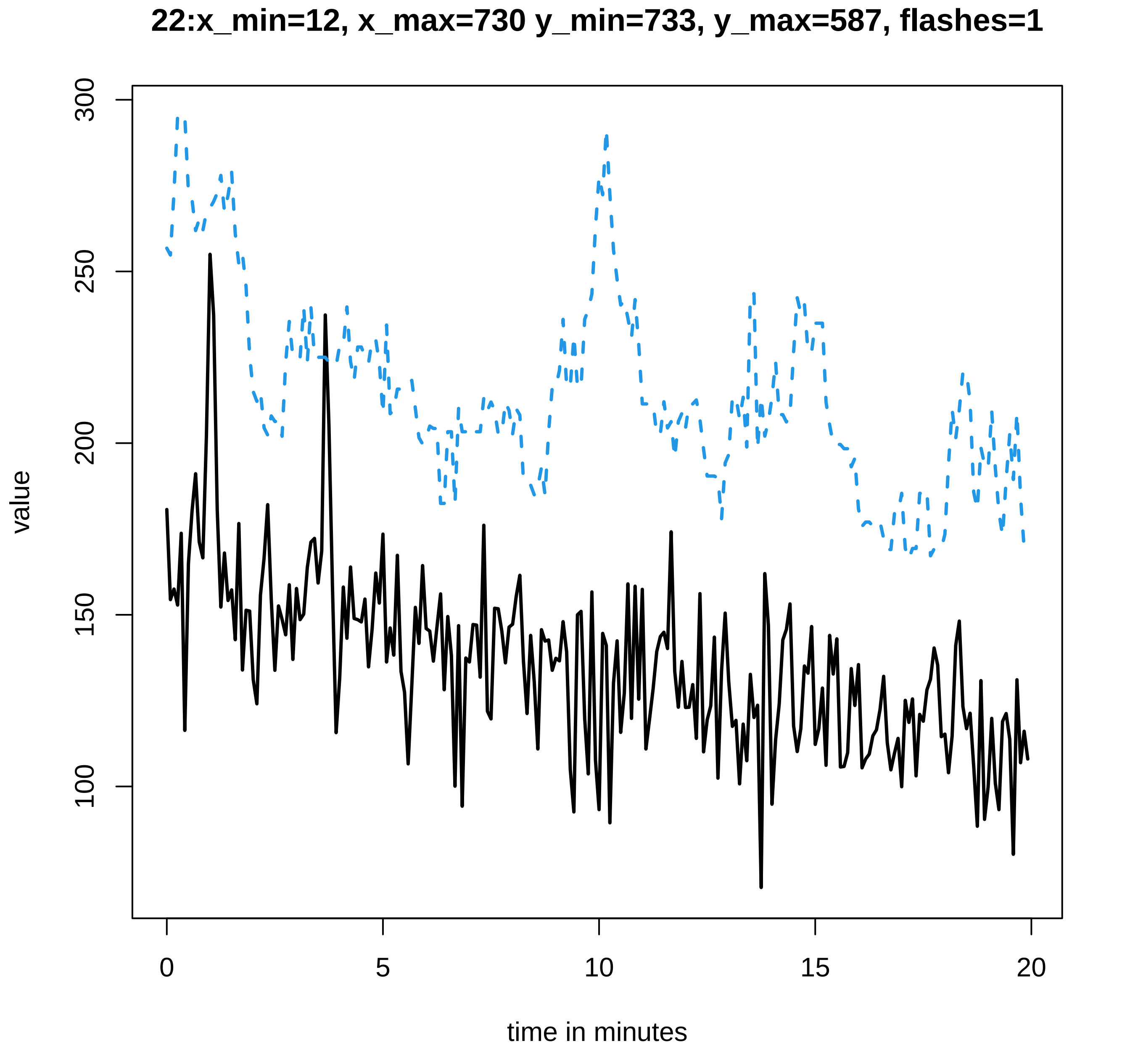

22:x_min=12, x_max=730 y_min=733, y_max=587, flashes=1
300
250
200
150
100
value
0
5
10
15
20
time in minutes

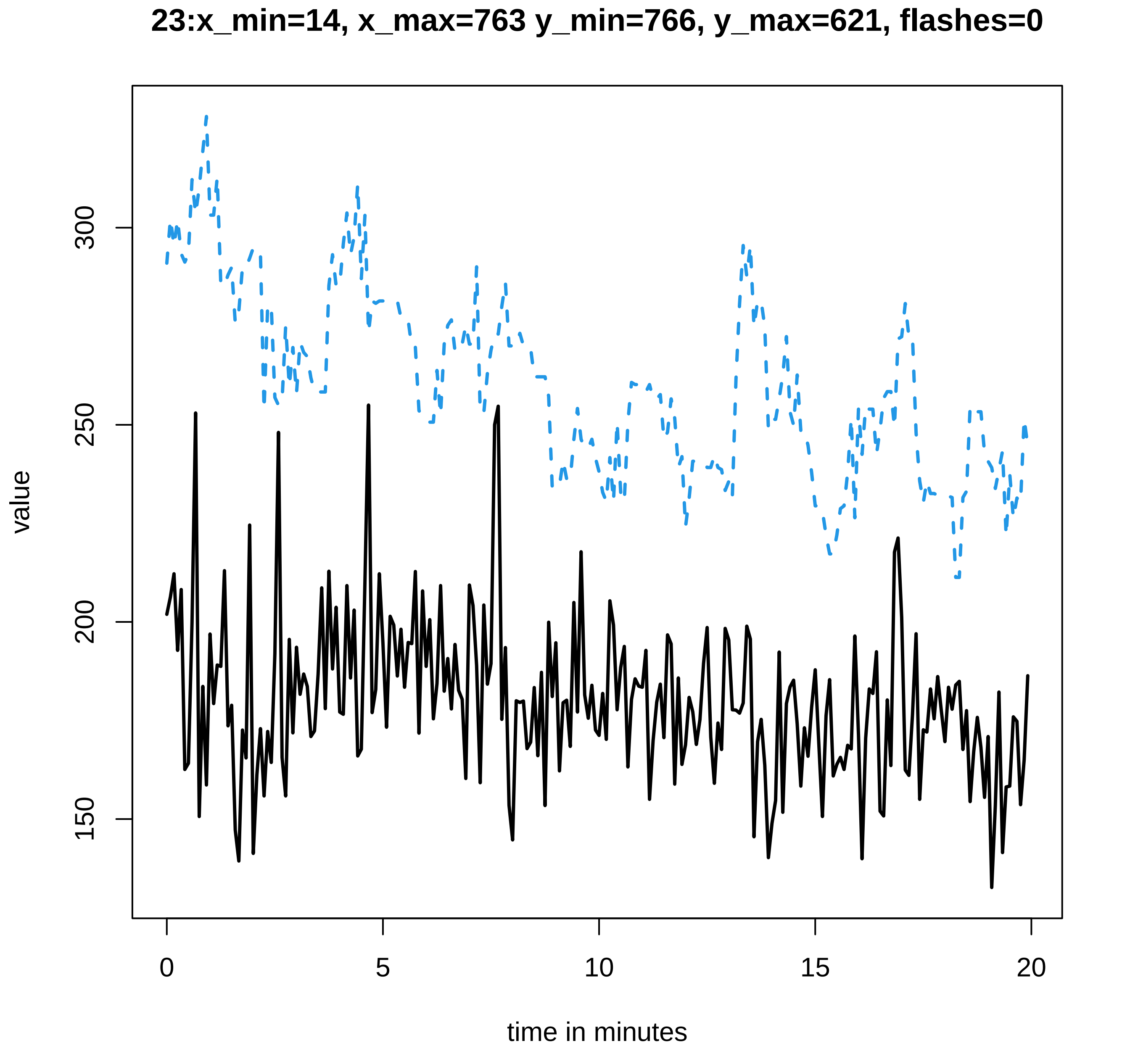
23:x_min=14, x_max=763 y_min=766, y_max=621, flashes=0
value
300
250
200
150
0
5
10
15
20
time in minutes

### 24:x_min=12, x_max=852 y_min=855, y_max=504, flashes=1

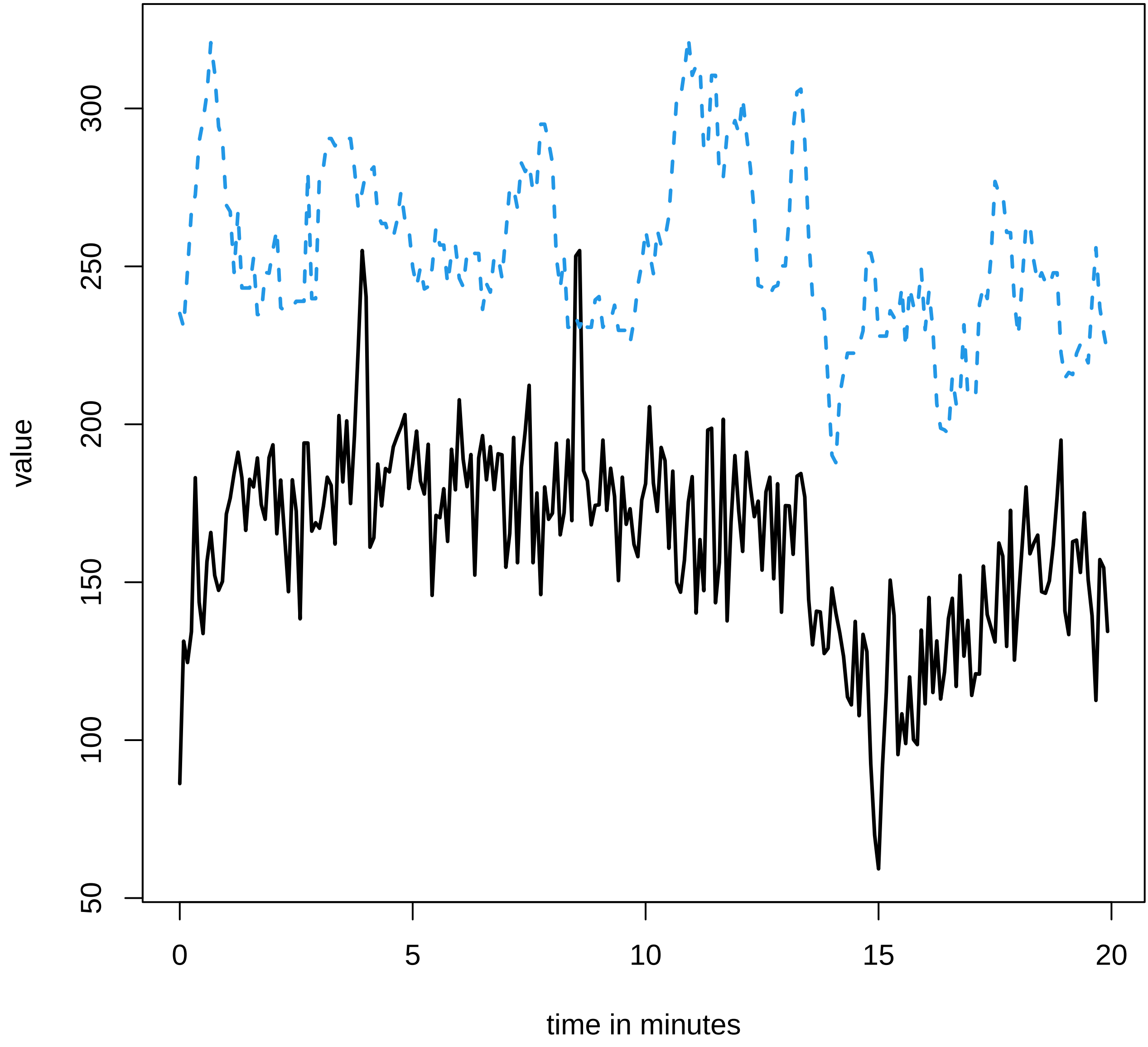

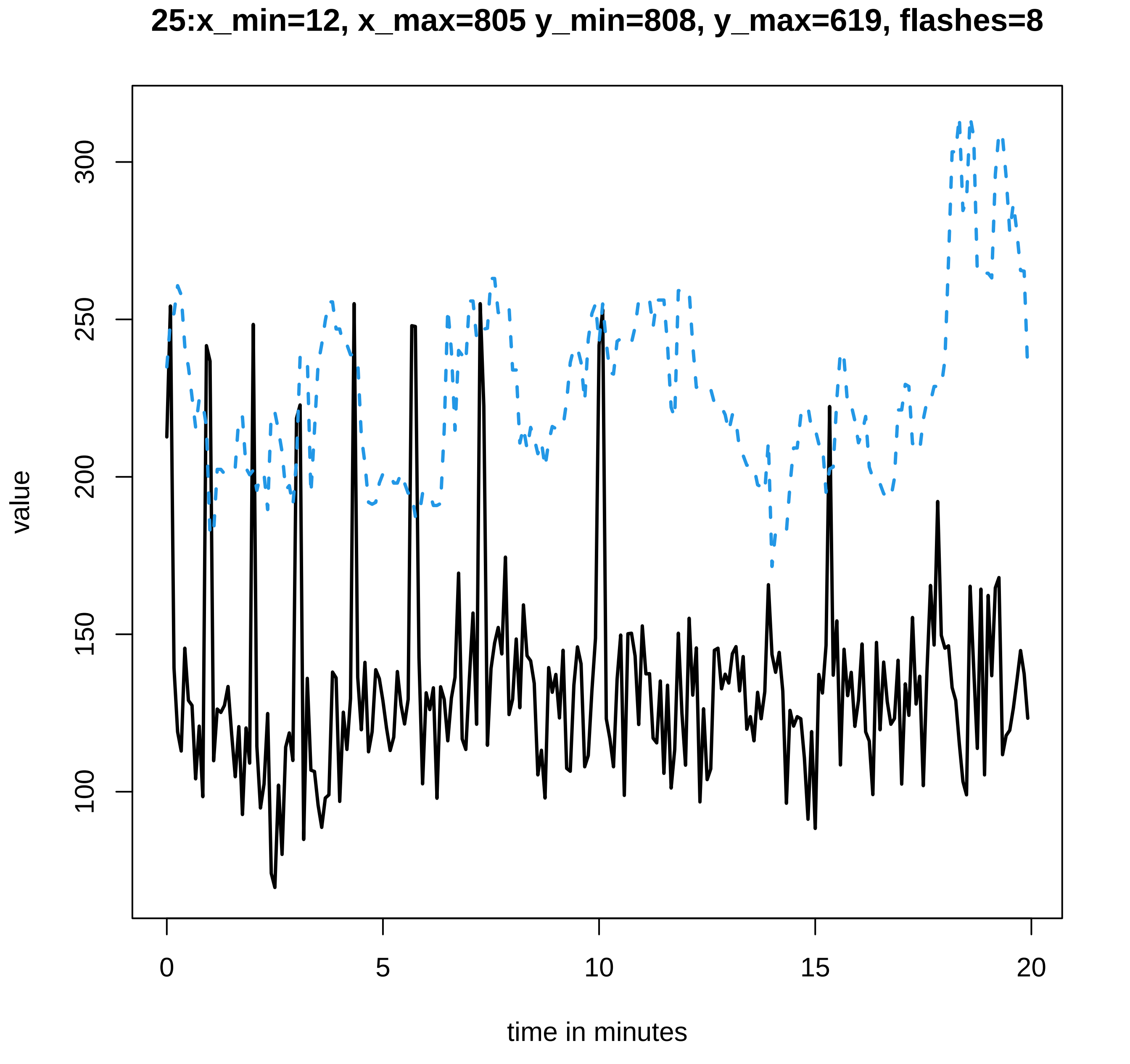
25:x_min=12, x_max=805 y_min=808, y_max=619, flashes=8
value
100
150
200
250
300
0
5
10
15
20
time in minutes

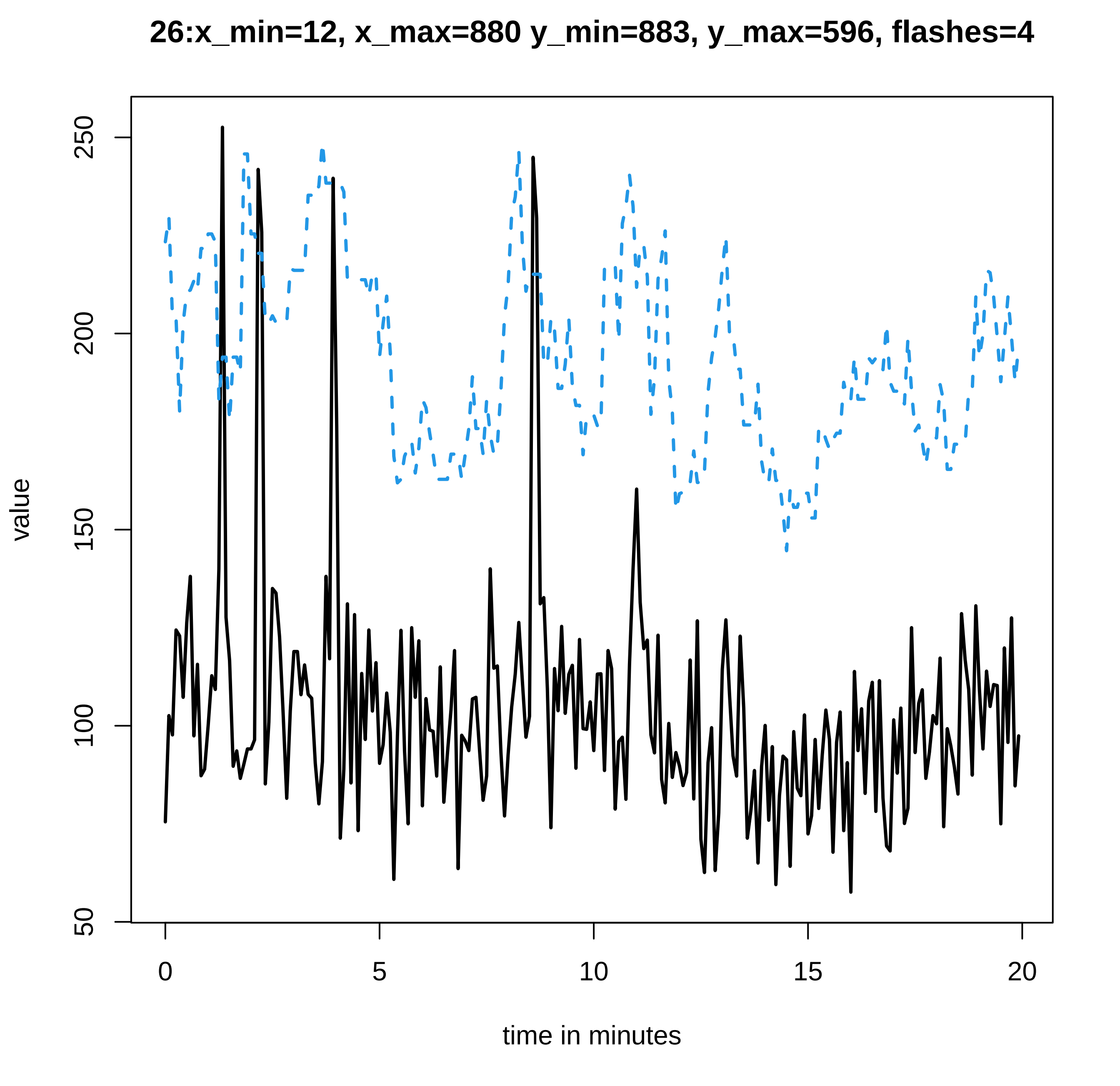
26:x_min=12, x_max=880 y_min=883, y_max=596, flashes=4
value
time in minutes
50
100
150
200
250
0
5
10
15
20

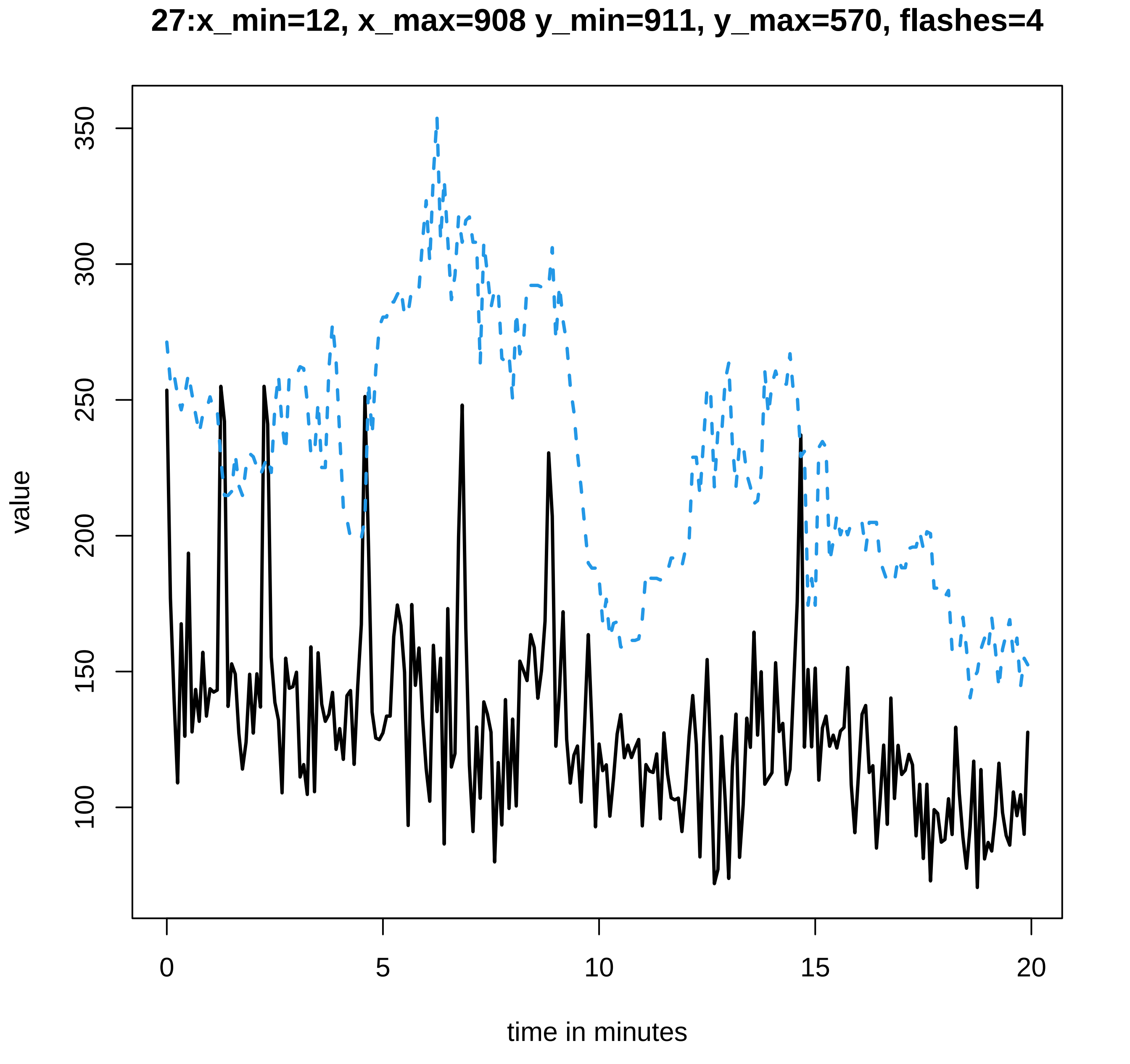
27:x_min=12, x_max=908 y_min=911, y_max=570, flashes=4
value
100
150
200
250
300
350
0
5
10
15
20
time in minutes

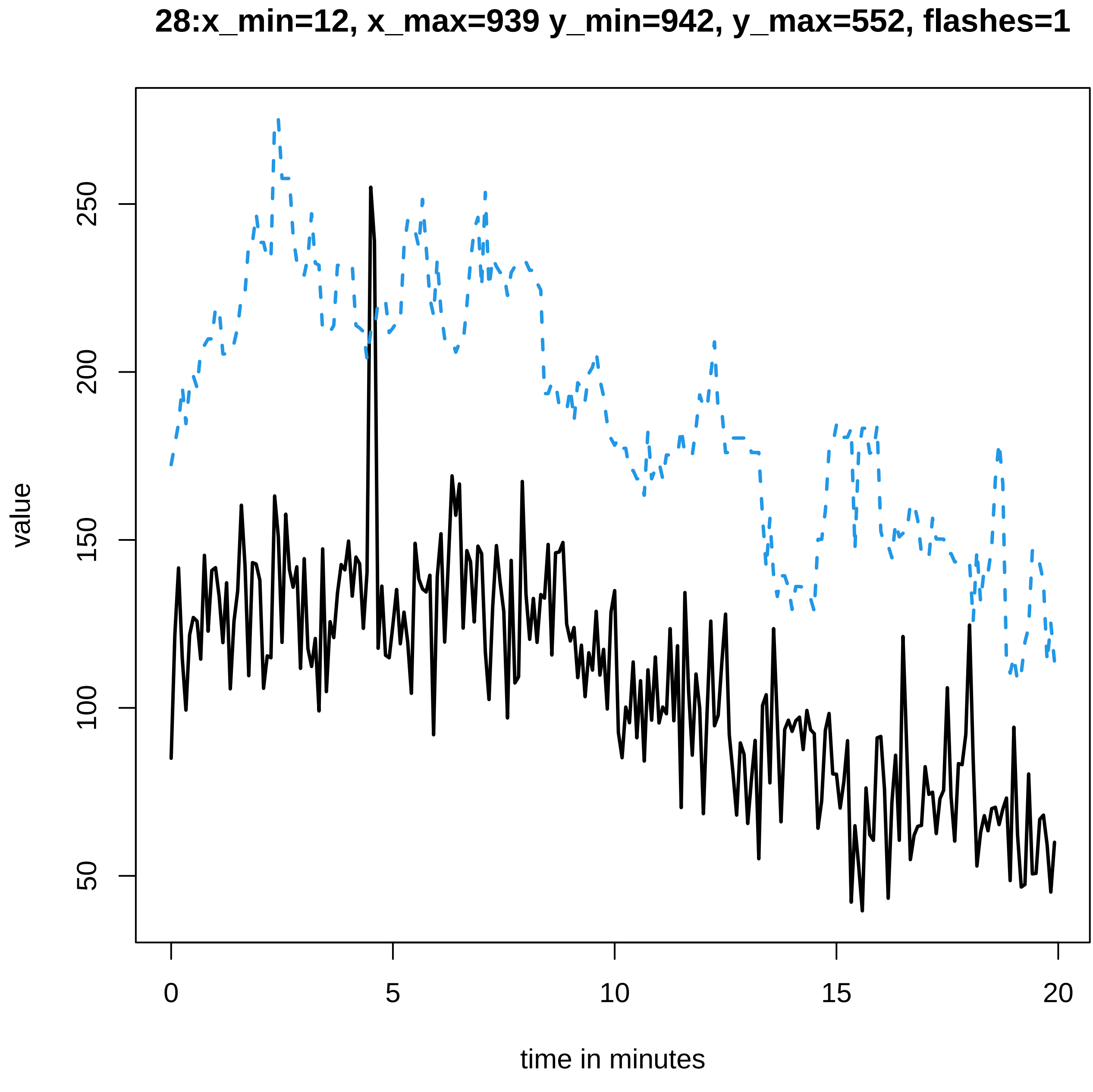

28:x_min=12, x_max=939 y_min=942, y_max=552, flashes=1
value
50
100
150
200
250
0
5
10
15
20
time in minutes

**29:x_min=12, x_max=910 y_min=913, y_max=604, flashes=2**

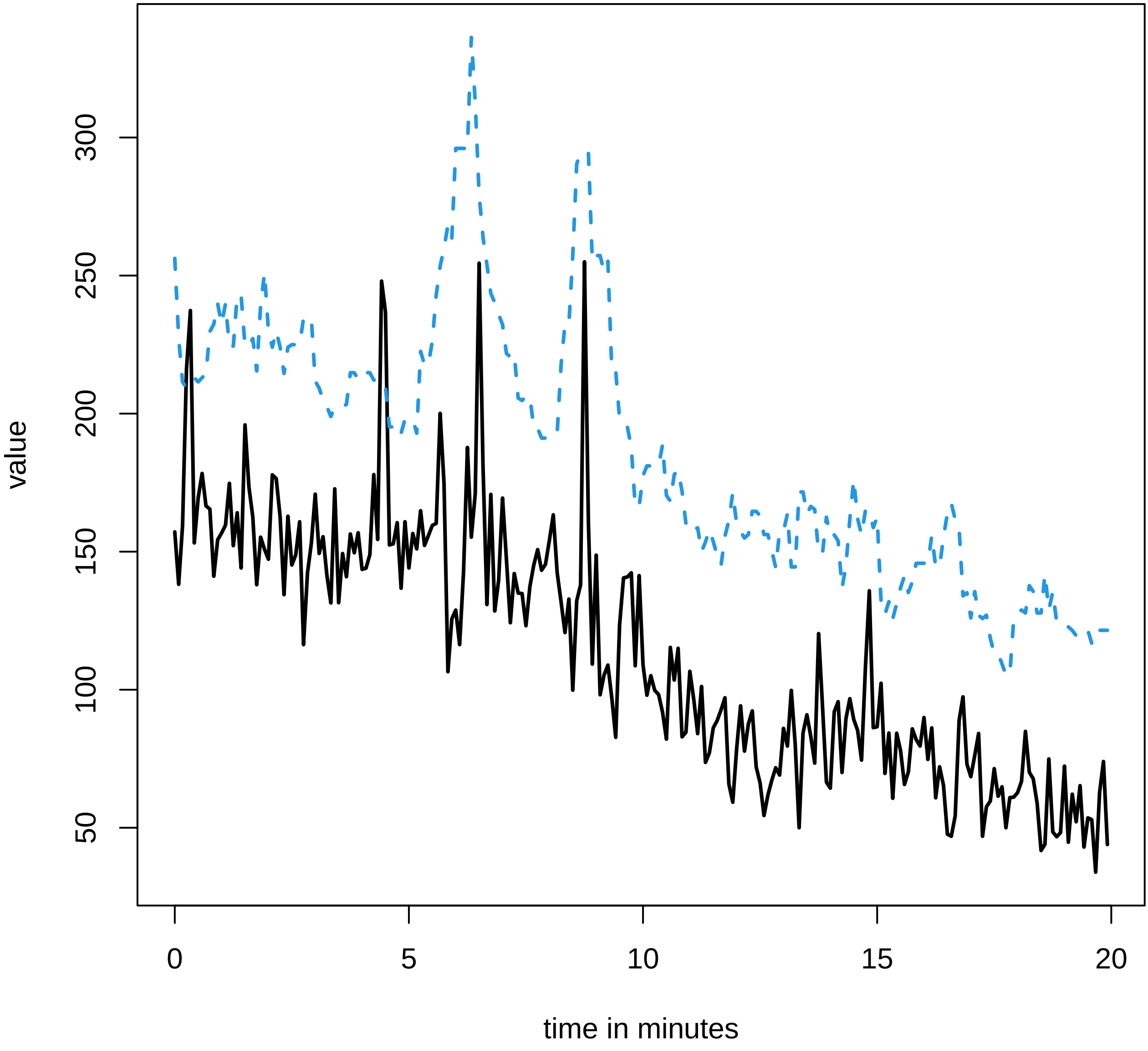

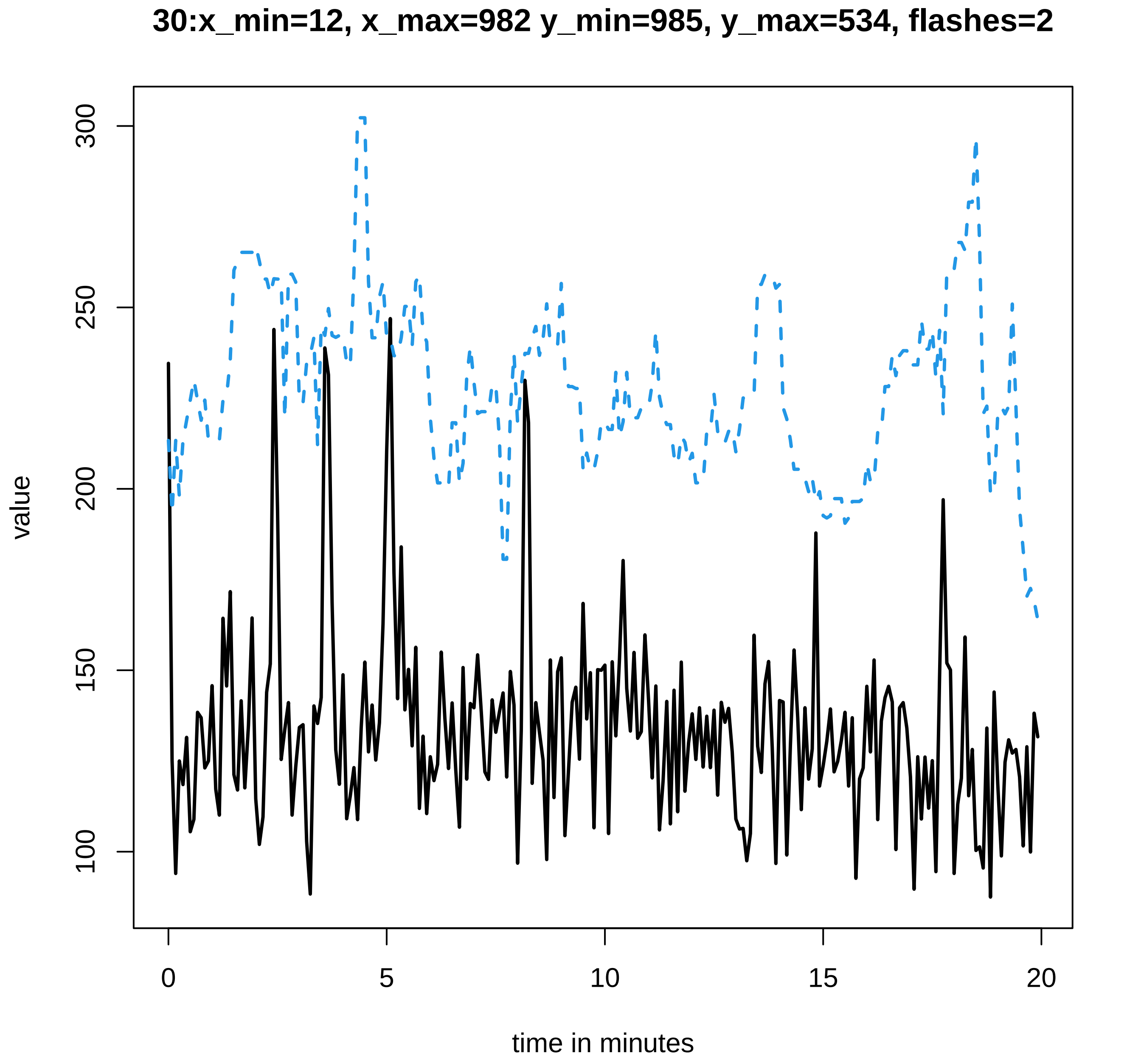
30:x_min=12, x_max=982 y_min=985, y_max=534, flashes=2
value
300
250
200
150
100
0
5
10
15
20
time in minutes
