## Supplementary data Calcium spiking for "LysM Receptor Proteins are Required for Ectomycorrhizal Symbiosis in Poplar": profiles-spiking-2.pdf

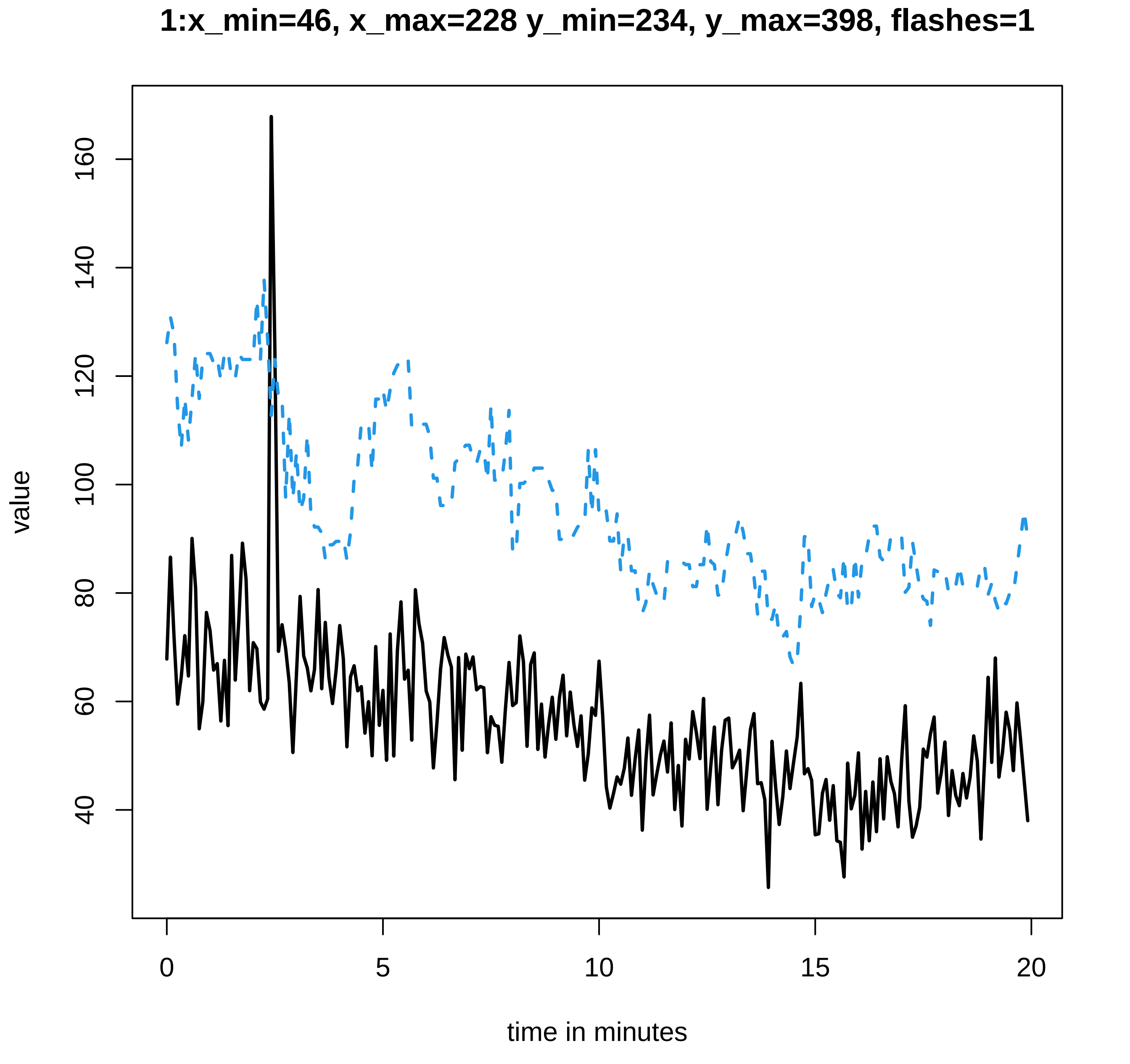
1:x_min=46, x_max=228 y_min=234, y_max=398, flashes=1
value
160
140
120
100
80
60
40
0
5
10
15
20
time in minutes

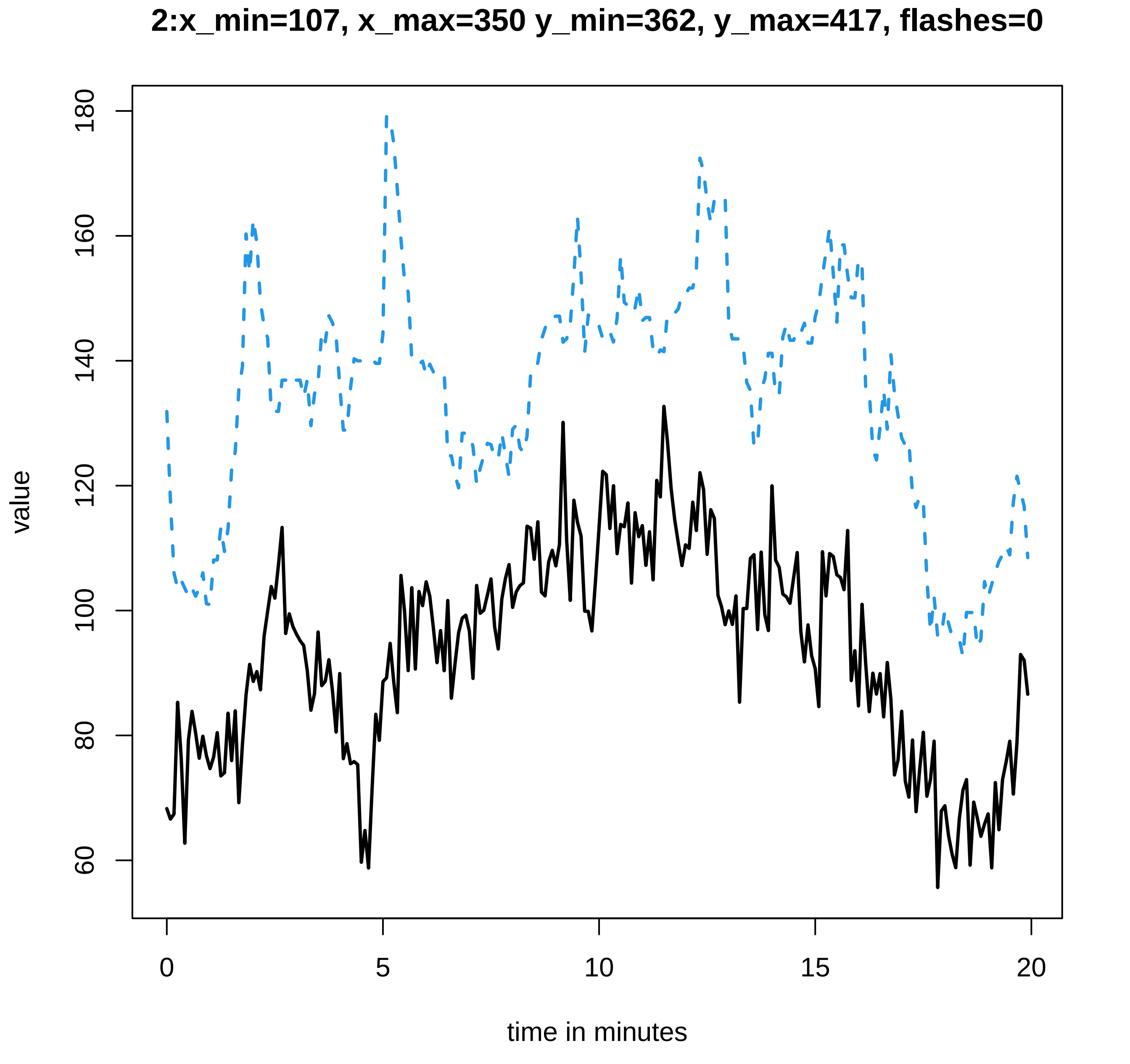

2:x_min=107, x_max=350 y_min=362, y_max=417, flashes=0
value
180
160
140
120
100
80
60
0
5
10
15
20
time in minutes

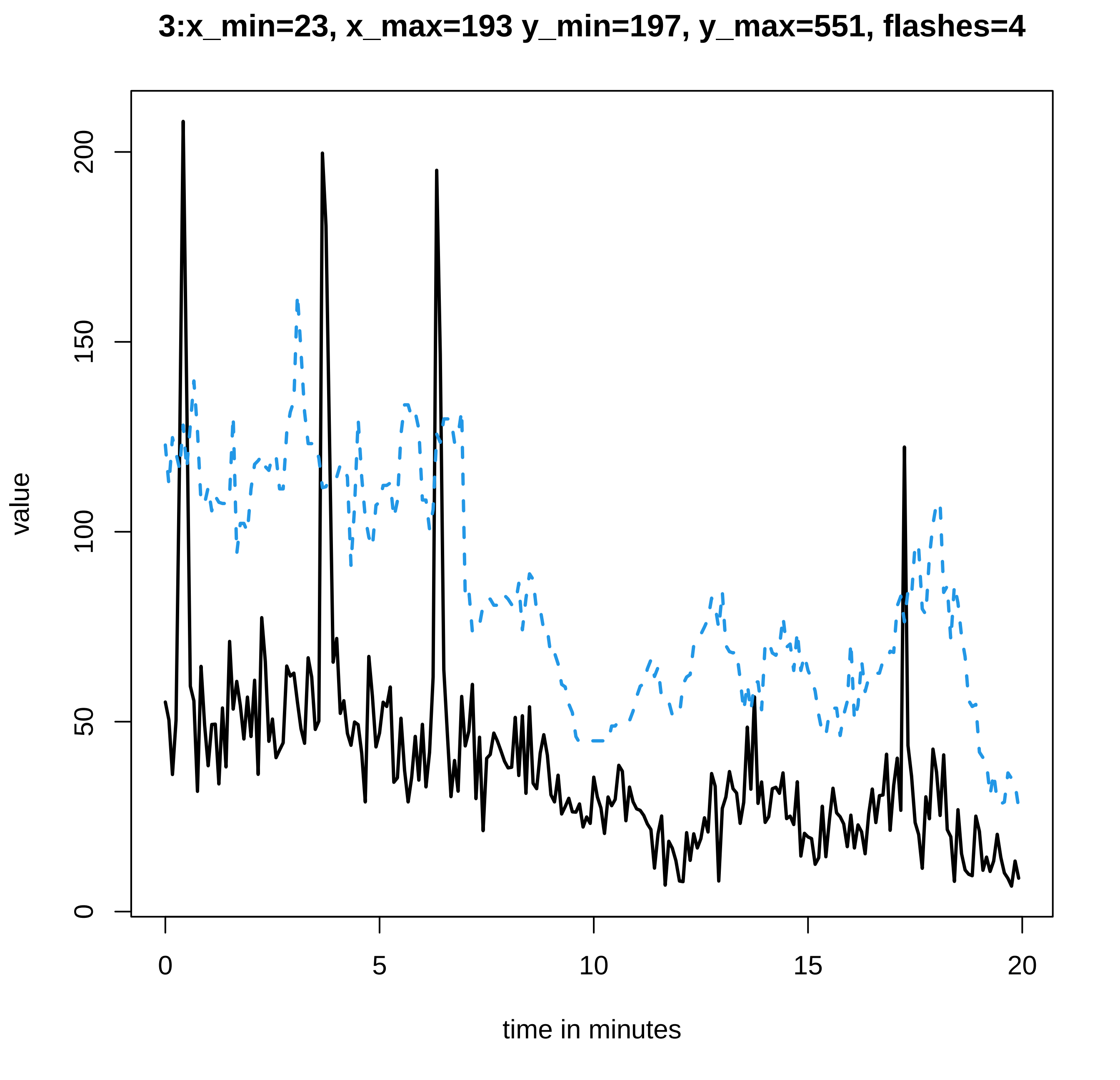

3:x_min=23, x_max=193 y_min=197, y_max=551, flashes=4
value
0
50
100
150
200
0
5
10
15
20
time in minutes

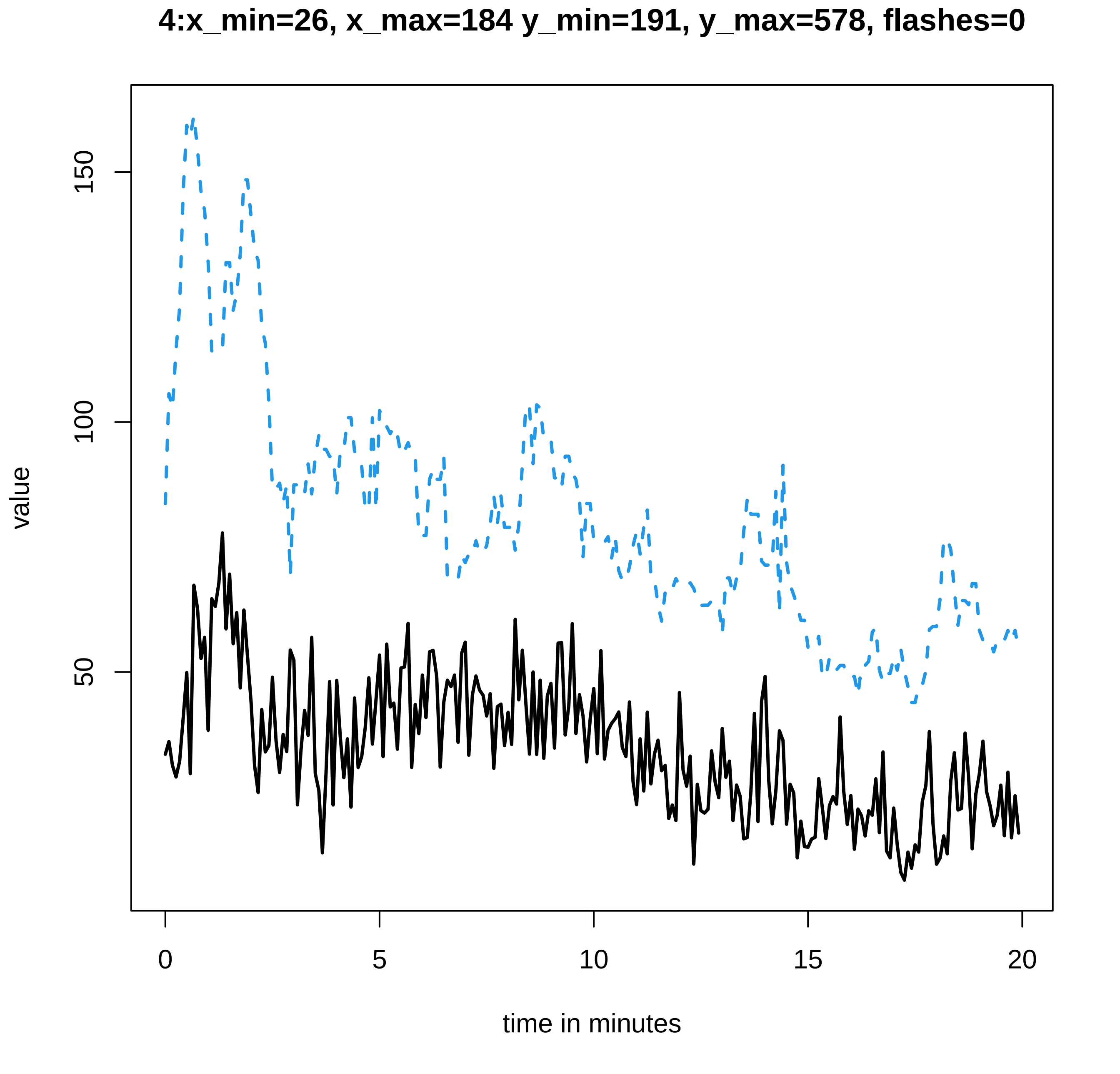
4:x_min=26, x_max=184 y_min=191, y_max=578, flashes=0
value
150
100
50
0
5
10
15
20
time in minutes

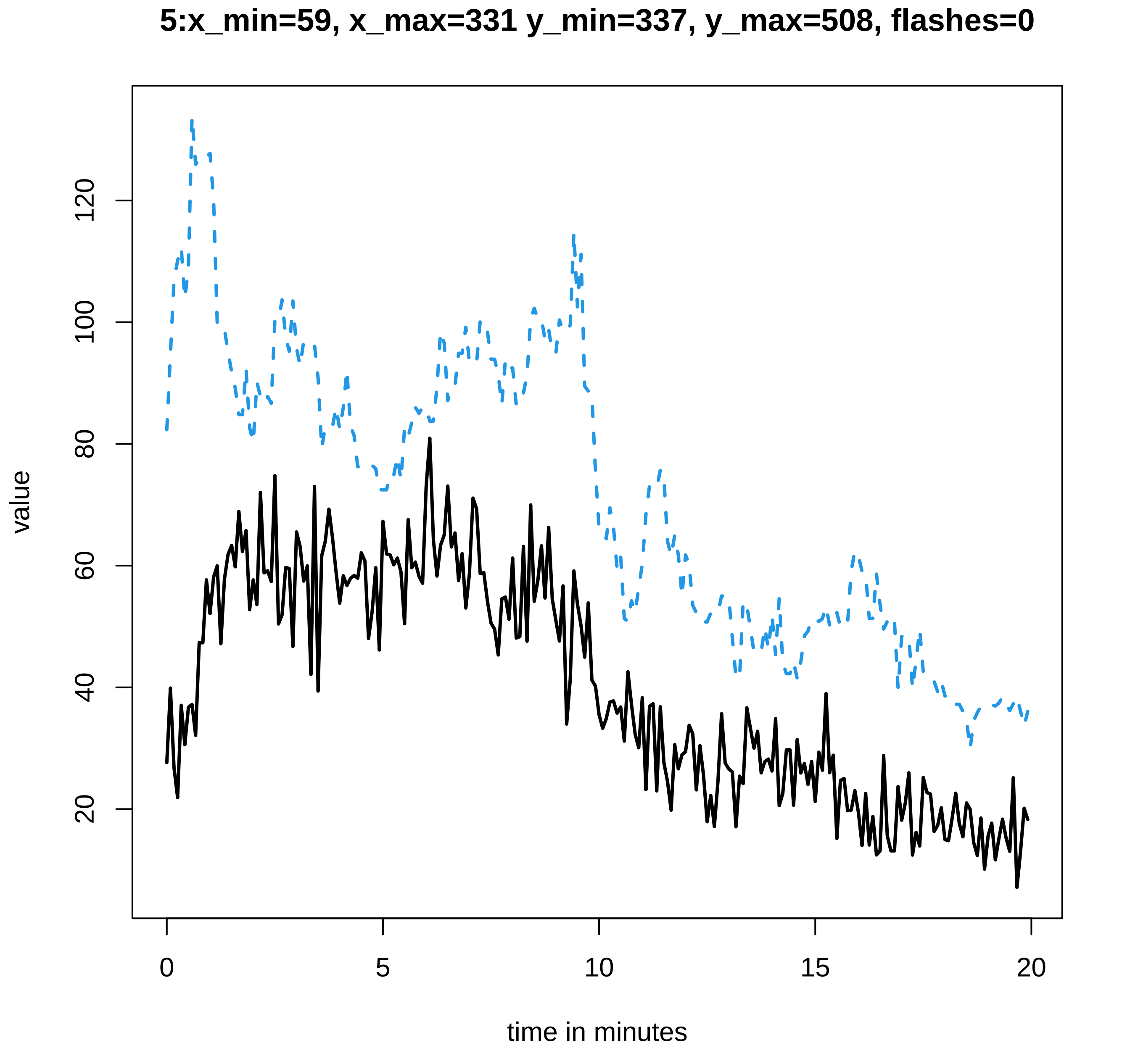
5:x_min=59, x_max=331 y_min=337, y_max=508, flashes=0
value
20
40
60
80
100
120
0
5
10
15
20
time in minutes

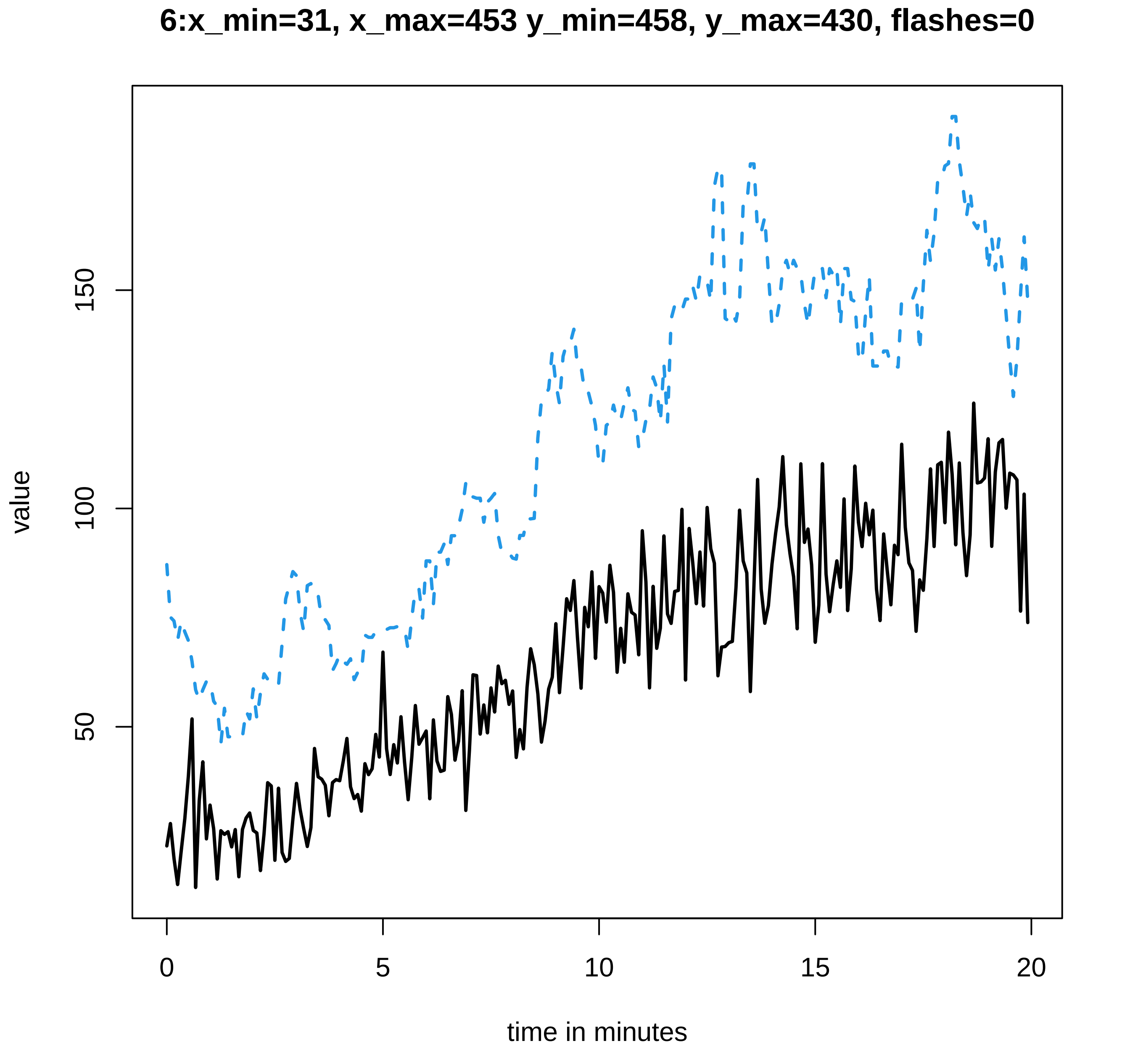
6:x_min=31, x_max=453 y_min=458, y_max=430, flashes=0
value
50
100
150
0
5
10
15
20
time in minutes

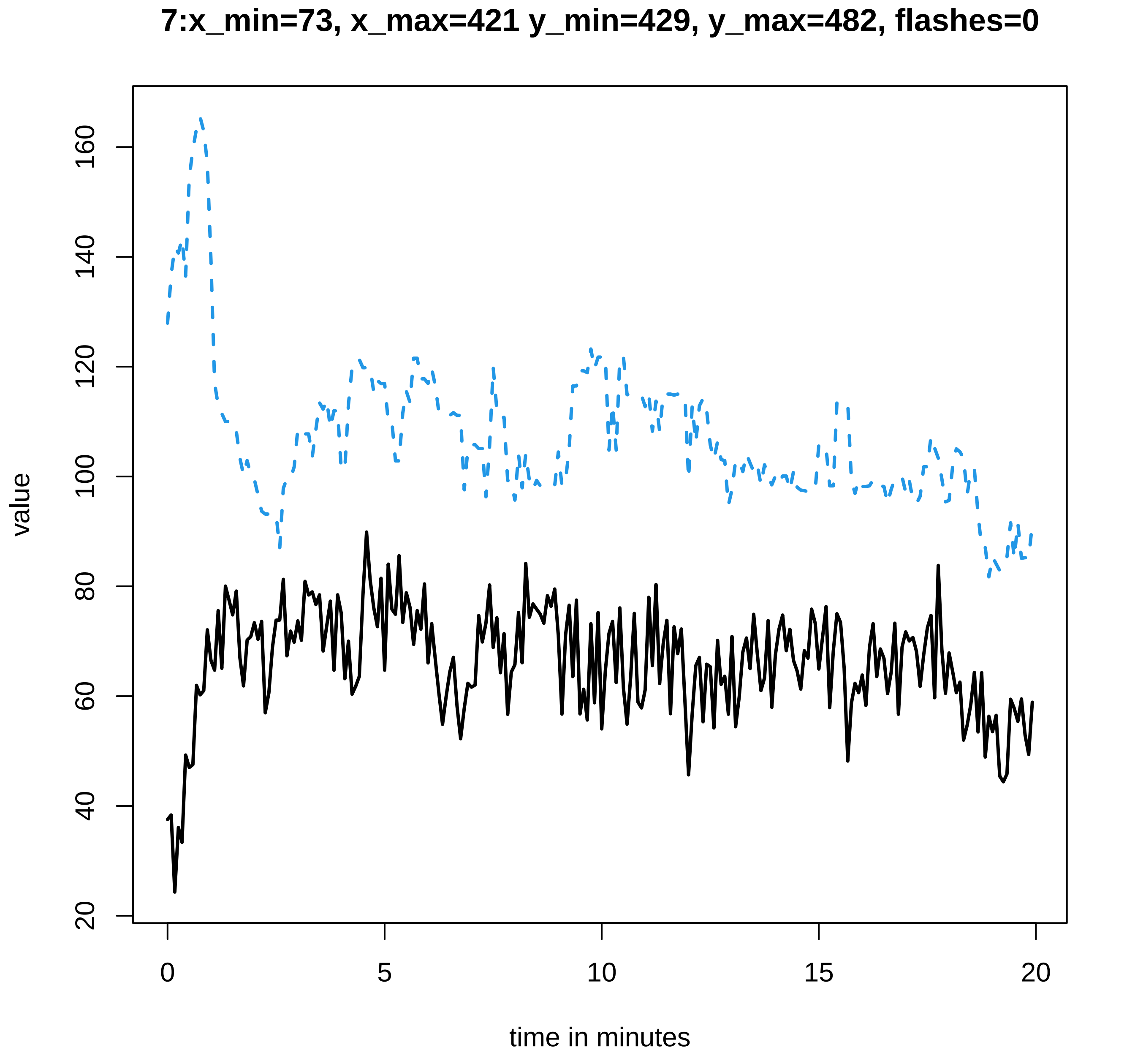
7:x_min=73, x_max=421 y_min=429, y_max=482, flashes=0
value
time in minutes
20
40
60
80
100
120
140
160
0
5
10
15
20

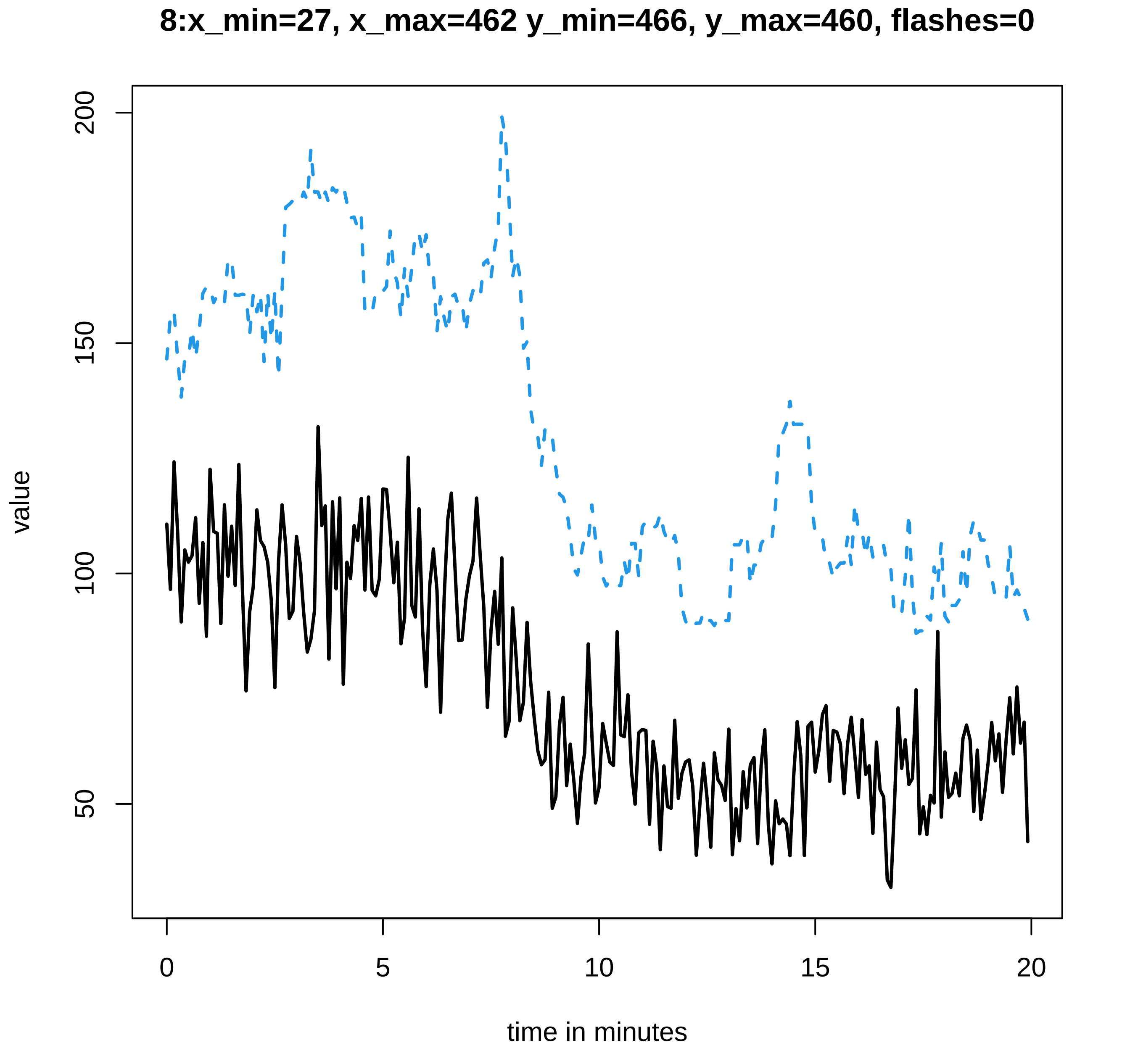
8:x_min=27, x_max=462 y_min=466, y_max=460, flashes=0
value
200
150
100
50
time in minutes
0
5
10
15
20

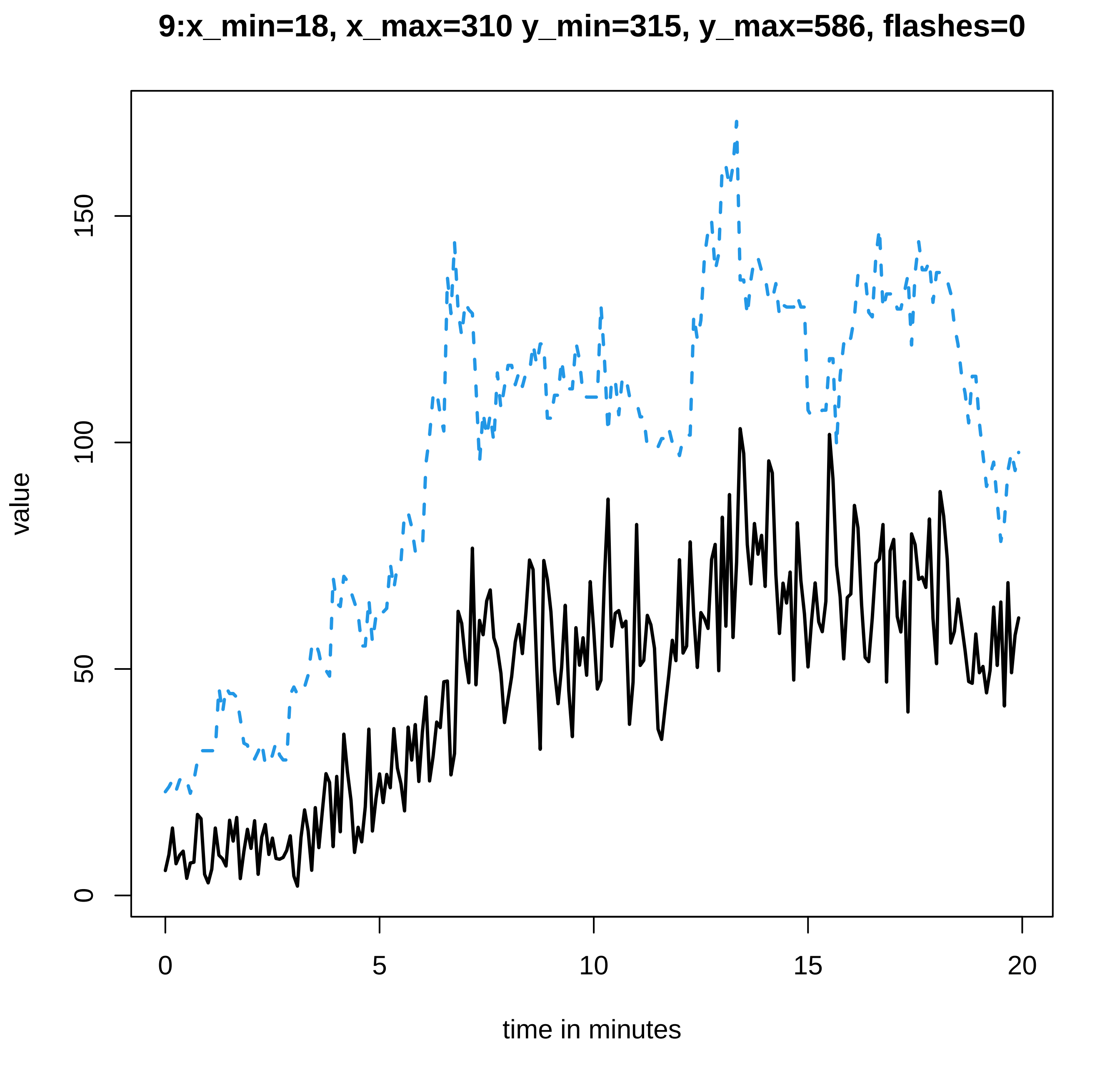

9:x_min=18, x_max=310 y_min=315, y_max=586, flashes=0
value
0
50
100
150
0
5
10
15
20
time in minutes

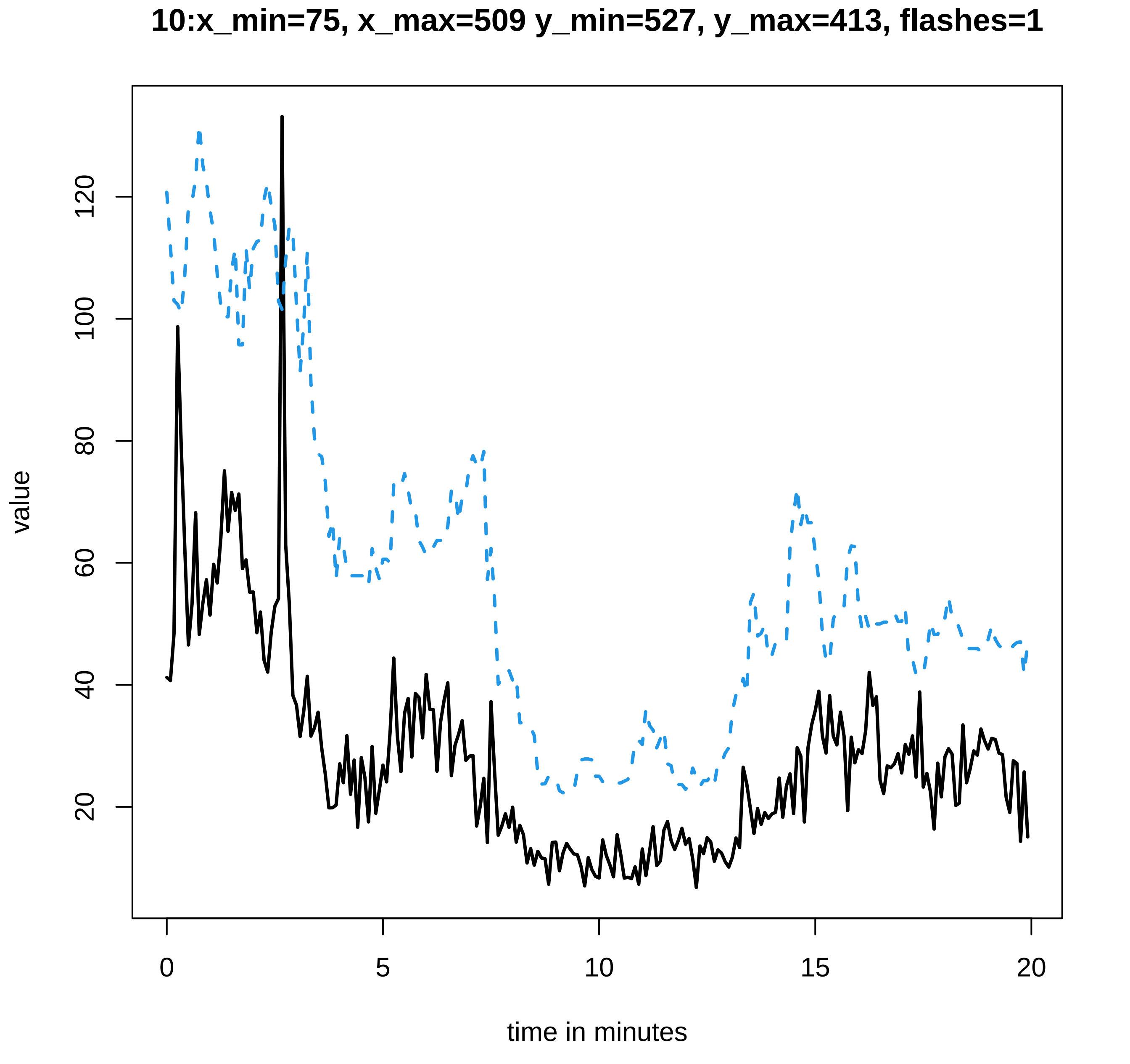
10:x_min=75, x_max=509 y_min=527, y_max=413, flashes=1
value
20
40
60
80
100
120
0
5
10
15
20
time in minutes

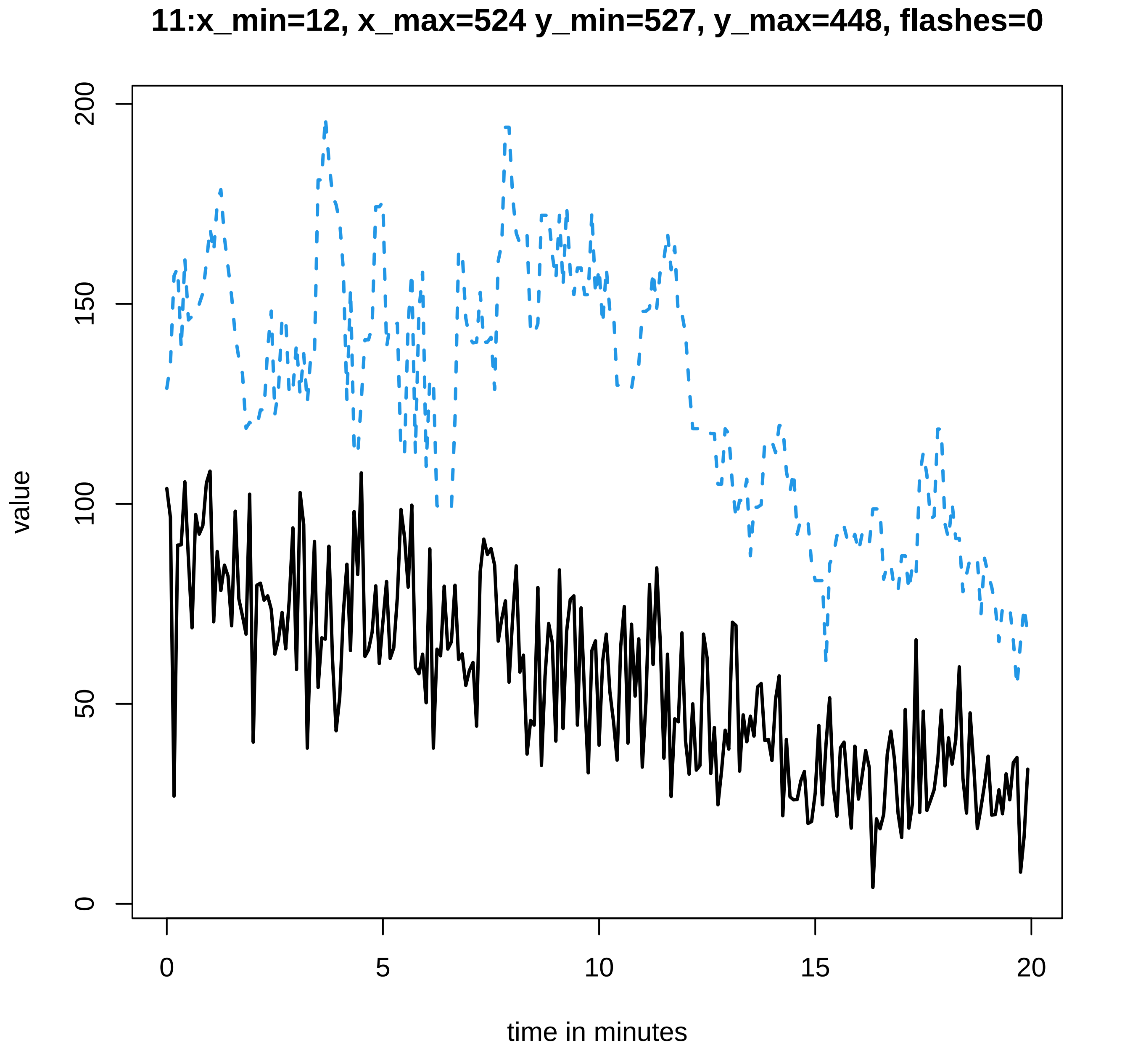

11:x_min=12, x_max=524 y_min=527, y_max=448, flashes=0
value
time in minutes
0
50
100
150
200
0
5
10
15
20

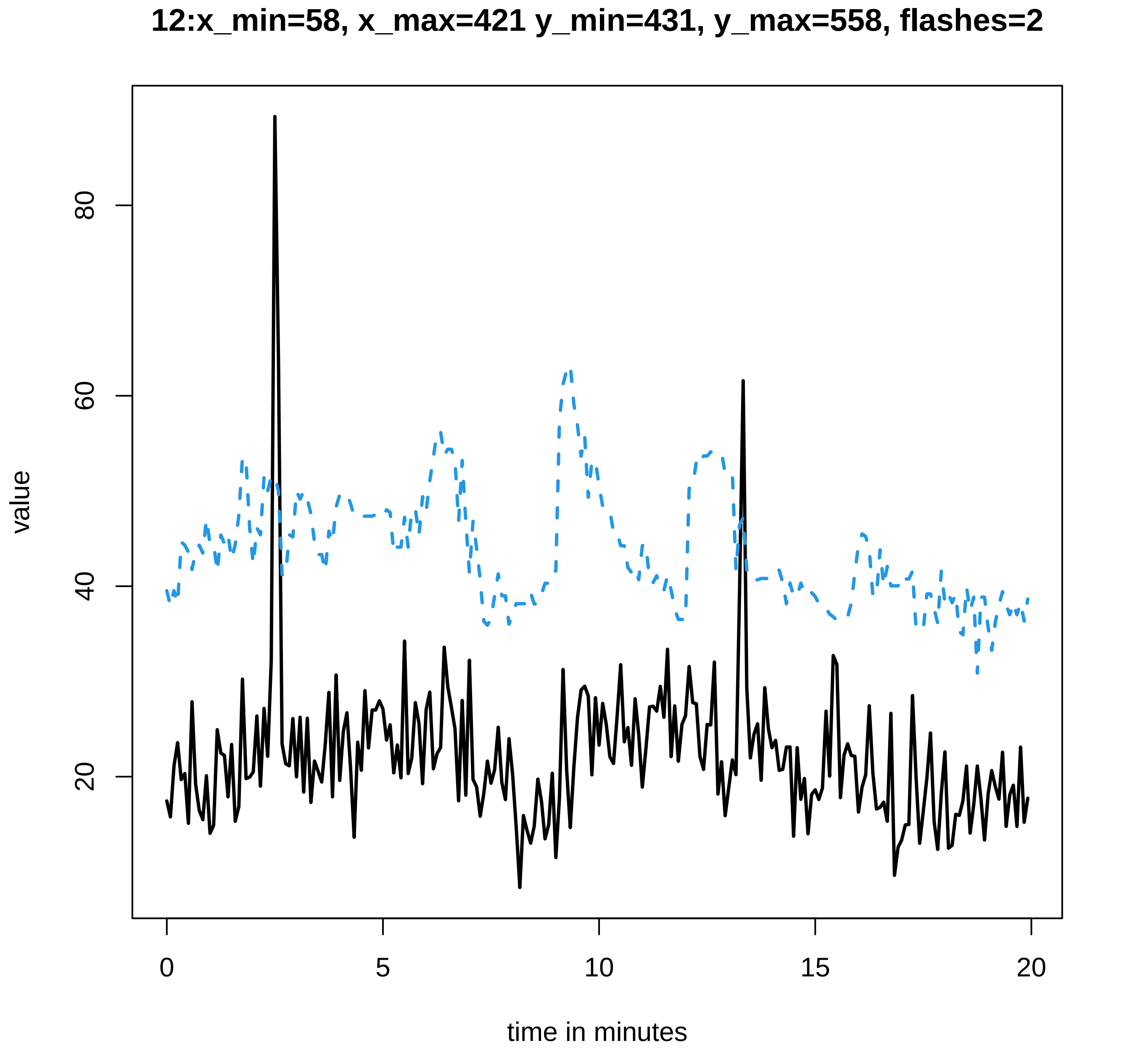

12:x_min=58, x_max=421 y_min=431, y_max=558, flashes=2
value
20
40
60
80
0
5
10
15
20
time in minutes

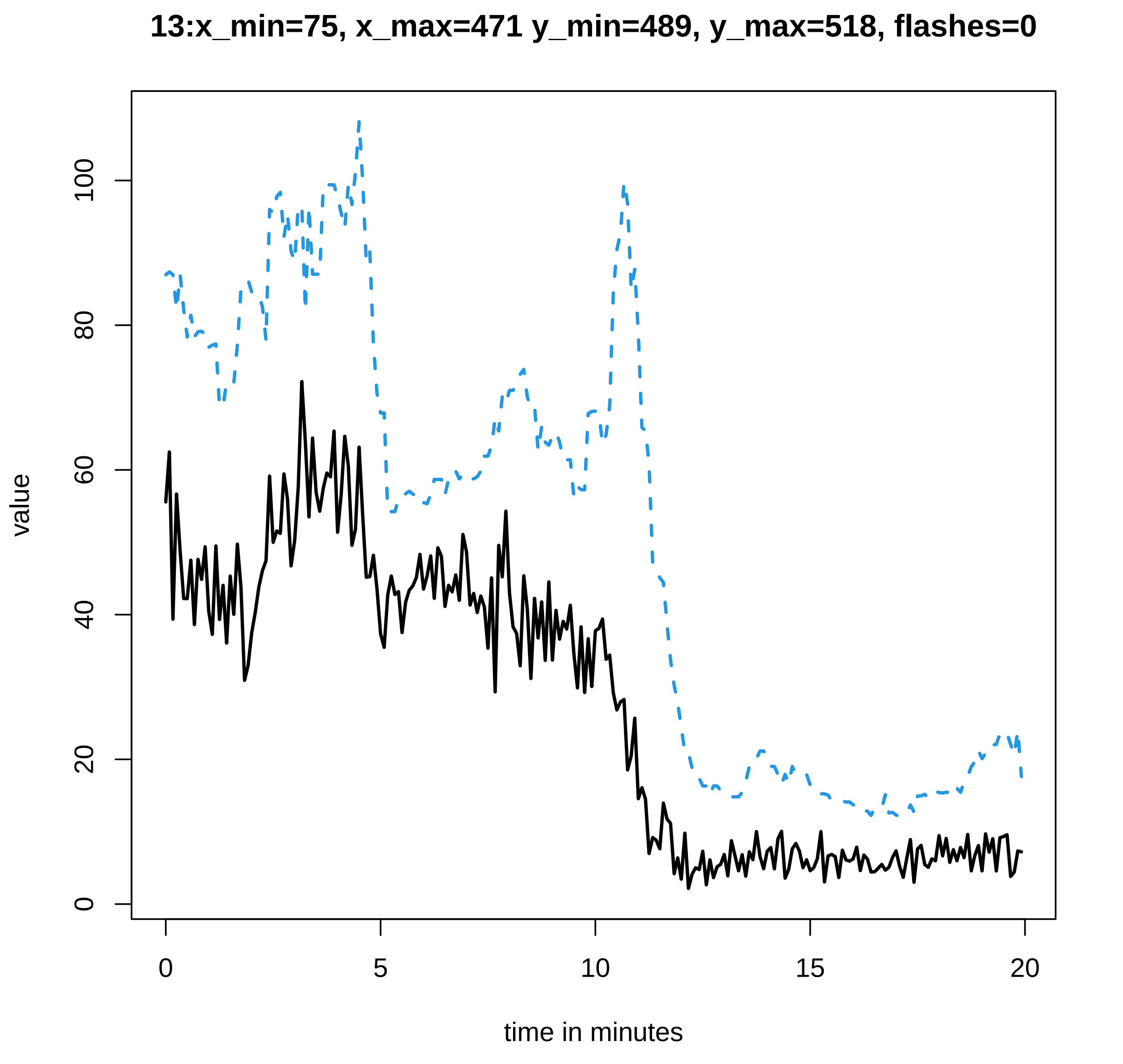
13:x_min=75, x_max=471 y_min=489, y_max=518, flashes=0
value
0
20
40
60
80
100
0
5
10
15
20
time in minutes

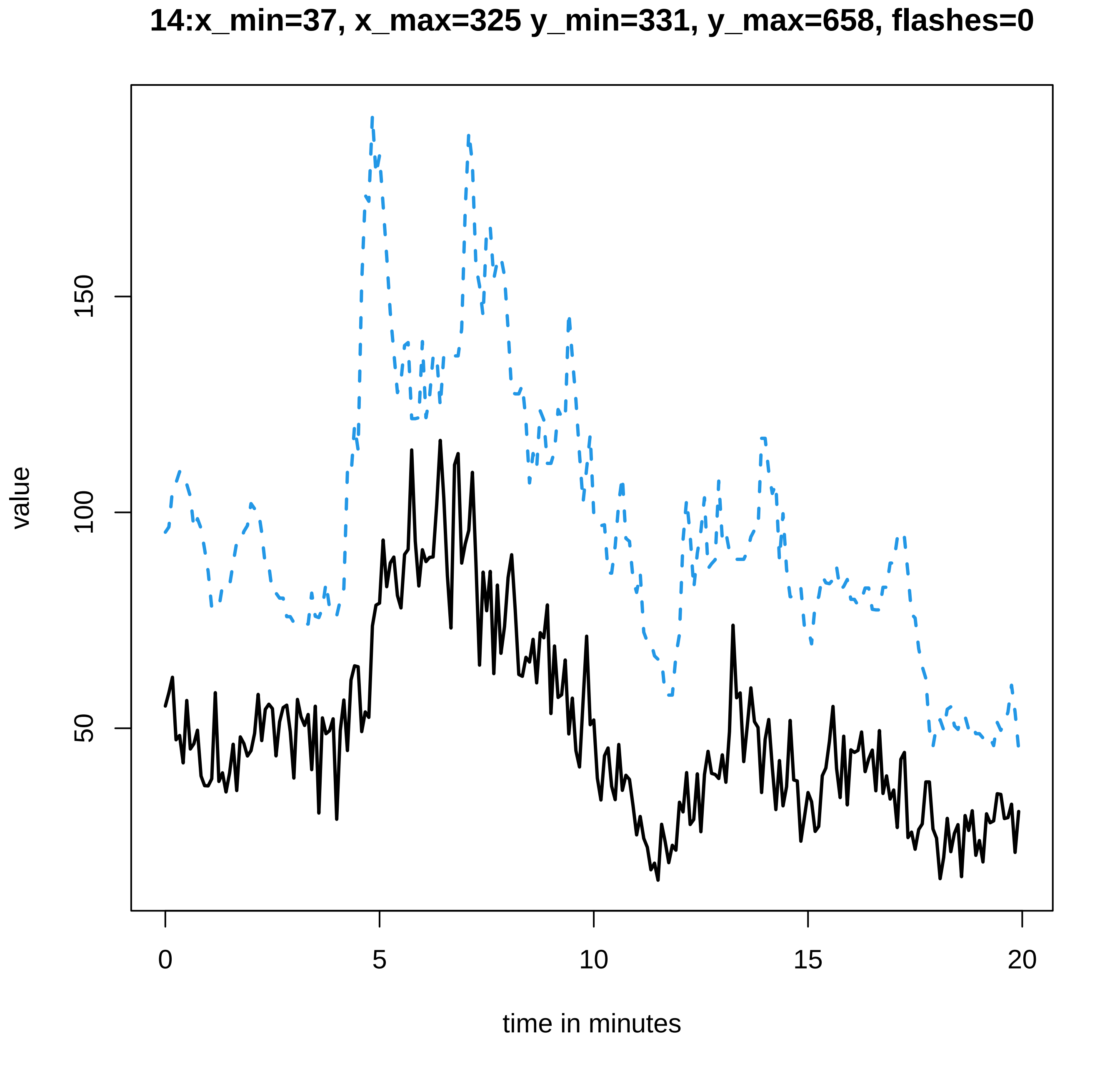
14:x_min=37, x_max=325 y_min=331, y_max=658, flashes=0
value
50
100
150
0
5
10
15
20
time in minutes

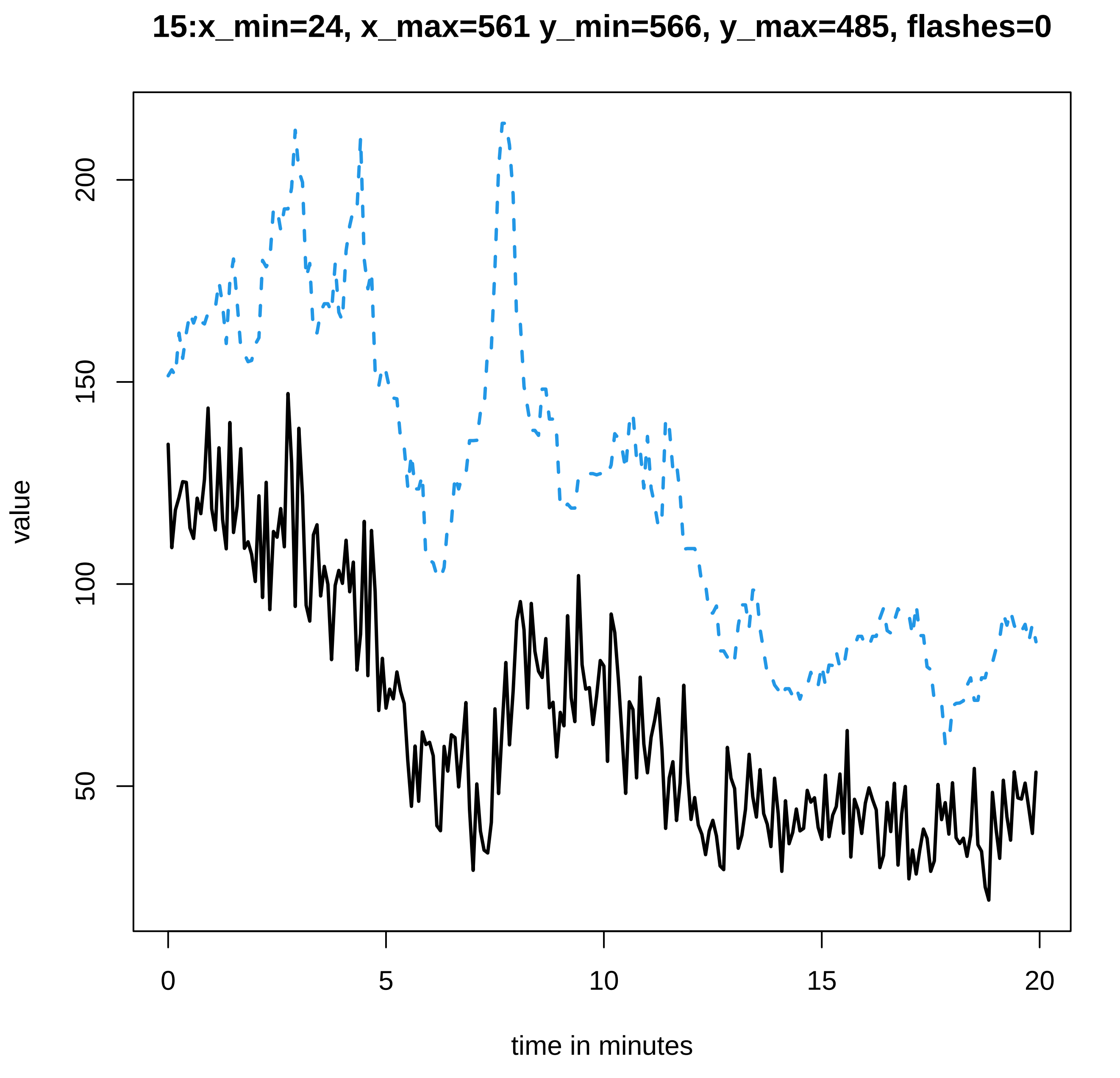
15:x_min=24, x_max=561 y_min=566, y_max=485, flashes=0
value
200
150
100
50
0
5
10
15
20
time in minutes

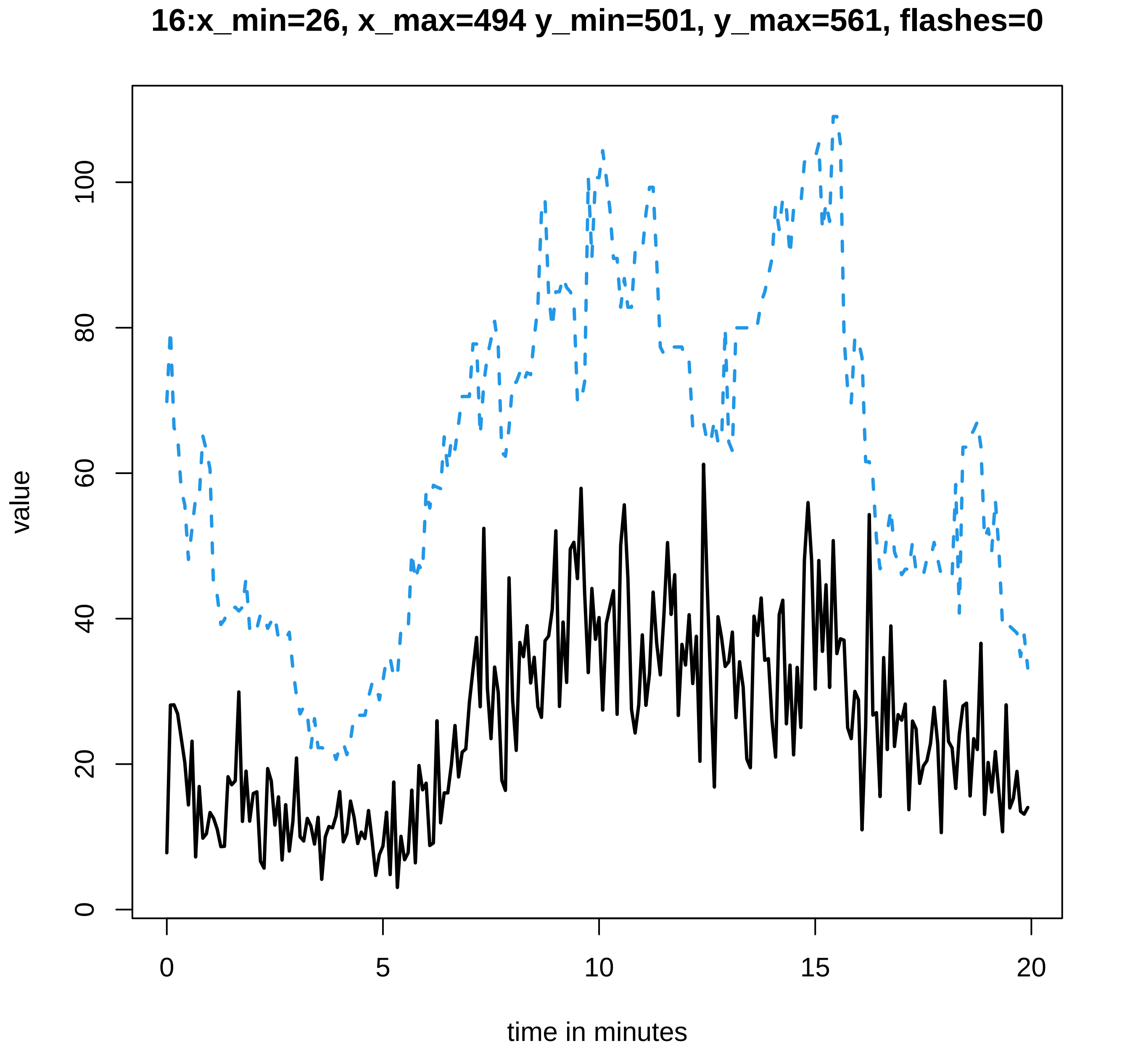
16:x_min=26, x_max=494 y_min=501, y_max=561, flashes=0
value
0
20
40
60
80
100
0
5
10
15
20
time in minutes

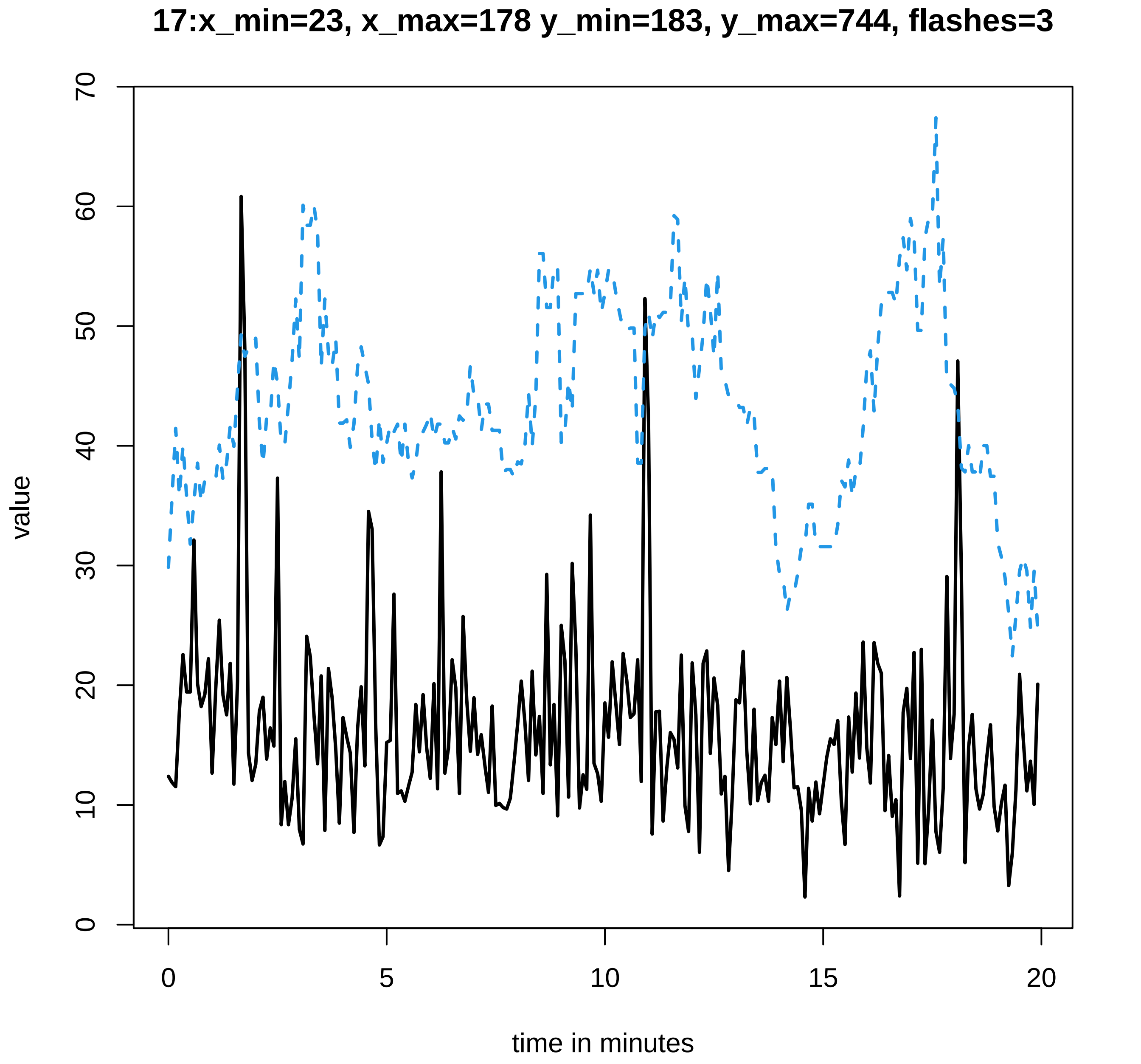
17:x_min=23, x_max=178 y_min=183, y_max=744, flashes=3
value
0
10
20
30
40
50
60
70
0
5
10
15
20
time in minutes

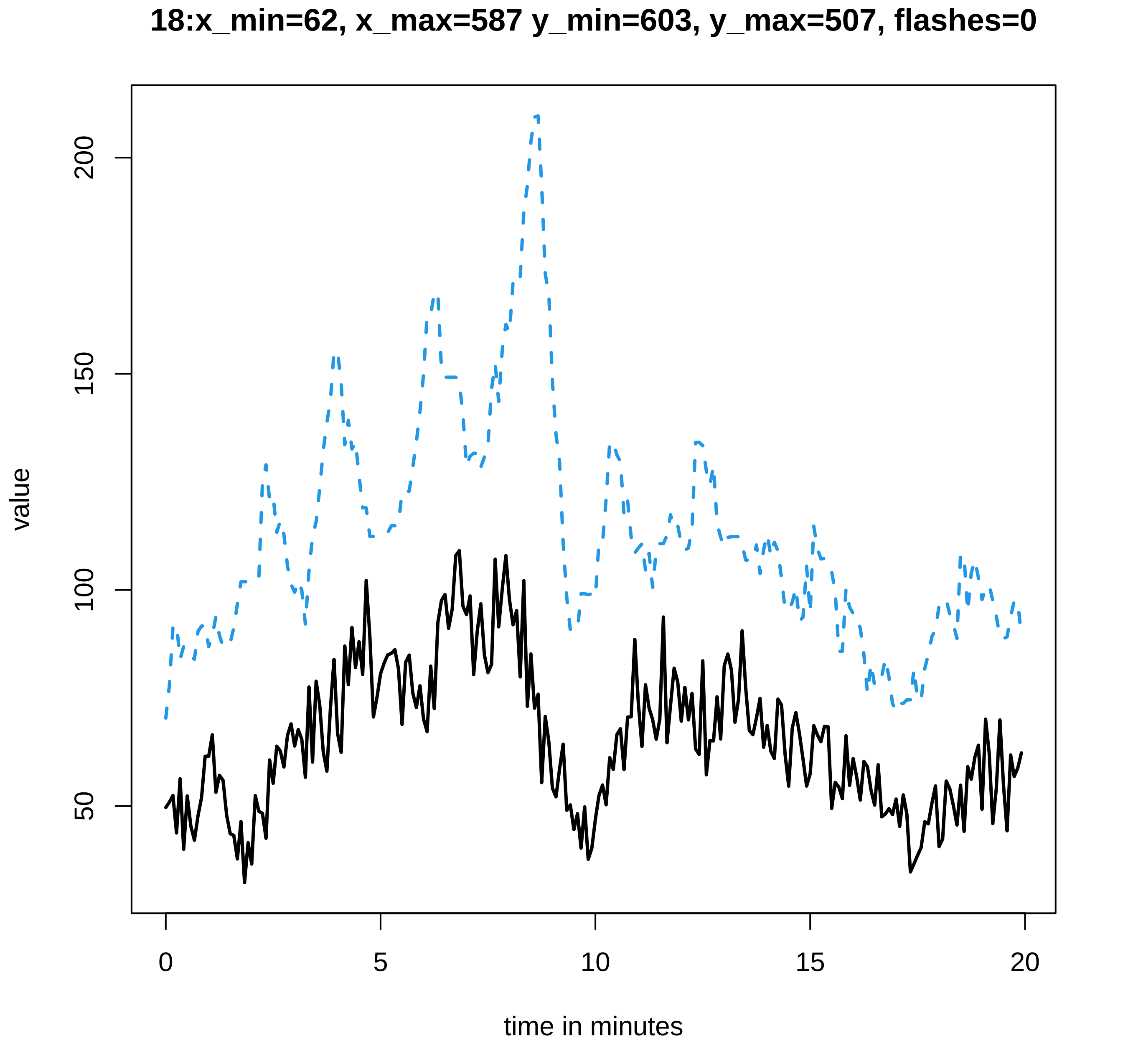
18:x_min=62, x_max=587 y_min=603, y_max=507, flashes=0
value
200
150
100
50
0
5
10
15
20
time in minutes

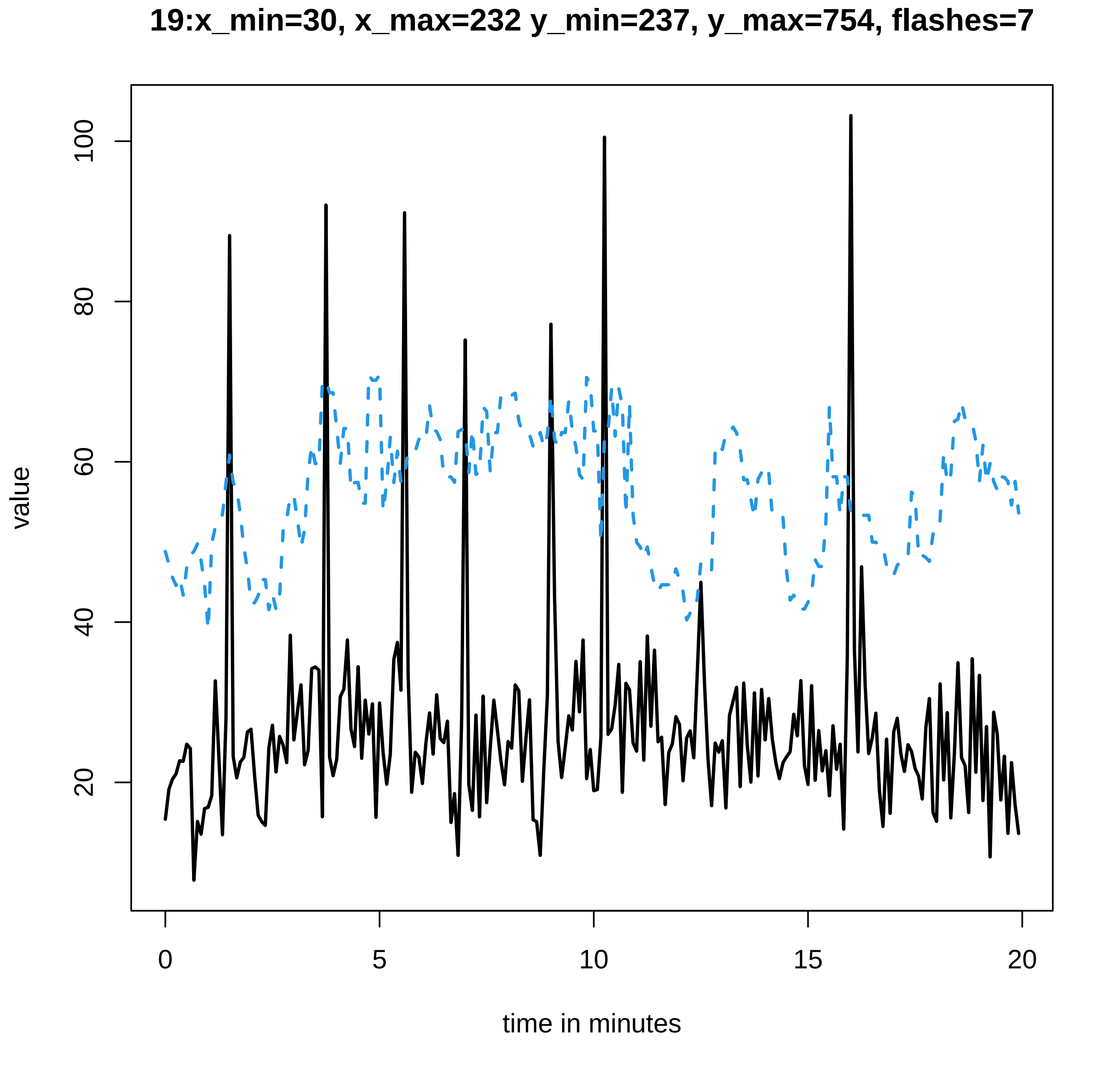
19:x_min=30, x_max=232 y_min=237, y_max=754, flashes=7
value
100
80
60
40
20
0
5
10
15
20
time in minutes

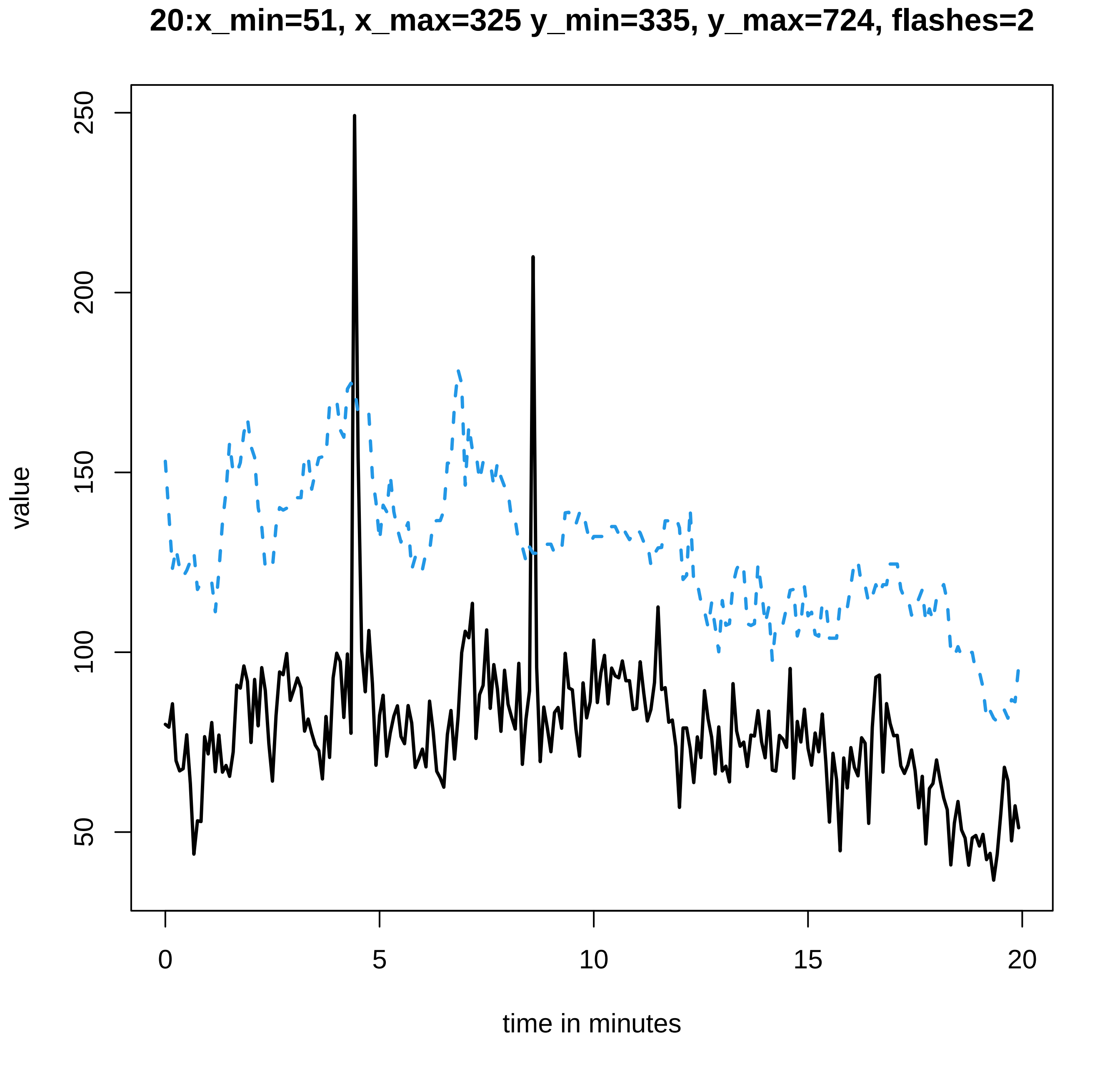
20:x_min=51, x_max=325 y_min=335, y_max=724, flashes=2
value
250
200
150
100
50
0
5
10
15
20
time in minutes

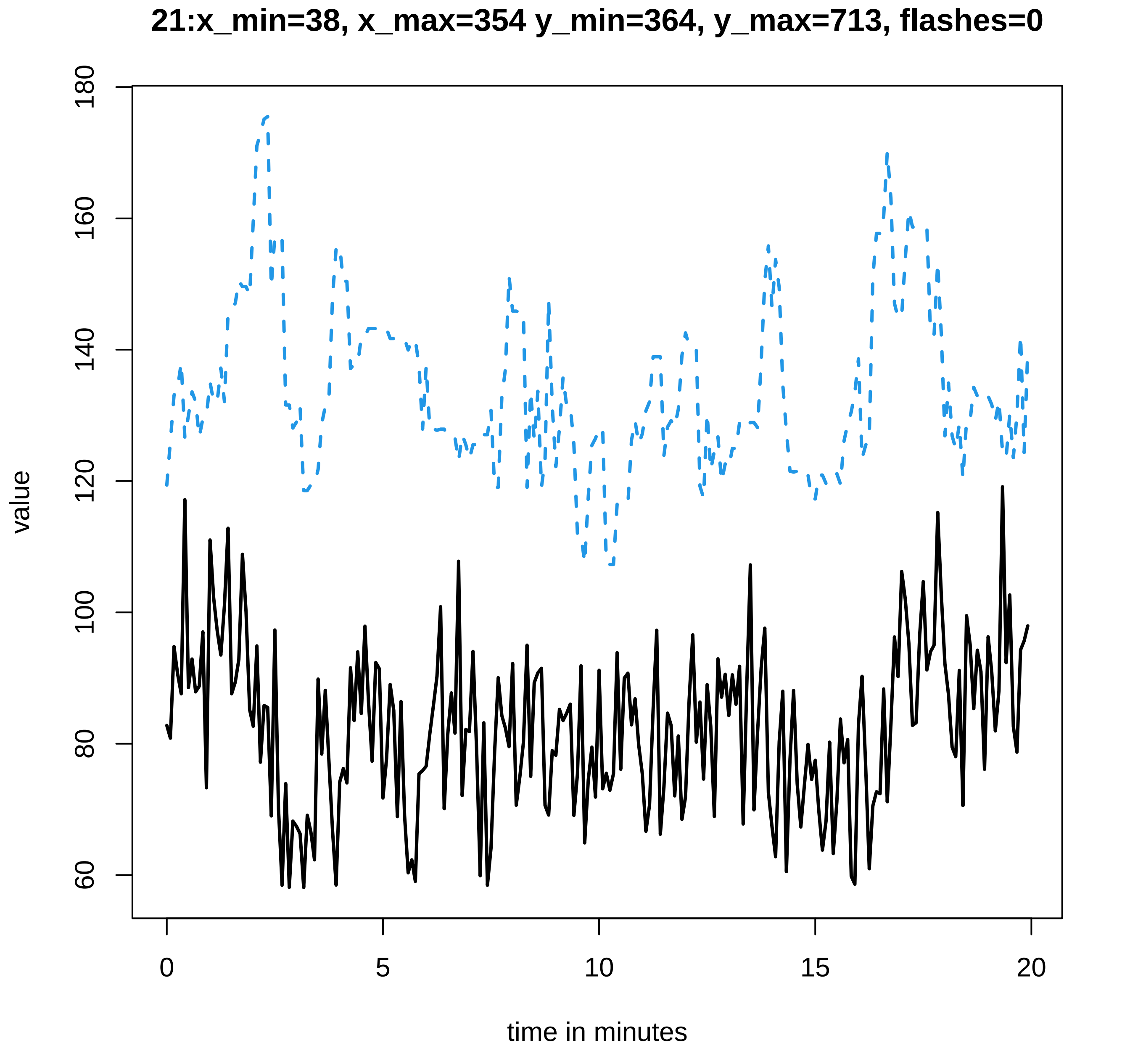
21:x_min=38, x_max=354 y_min=364, y_max=713, flashes=0
value
180
160
140
120
100
80
60
0
5
10
15
20
time in minutes

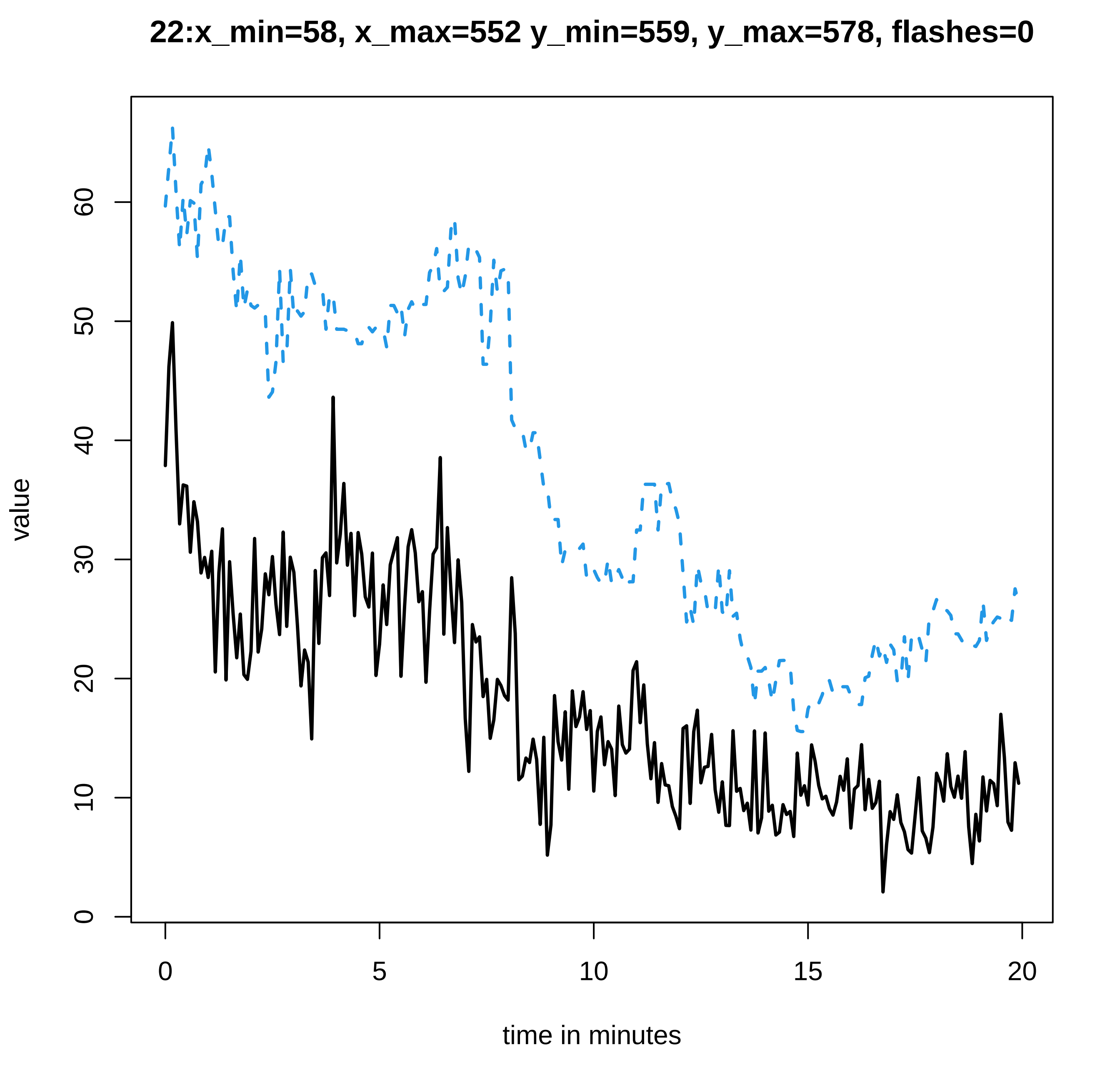
22:x_min=58, x_max=552 y_min=559, y_max=578, flashes=0
value
0
10
20
30
40
50
60
0
5
10
15
20
time in minutes

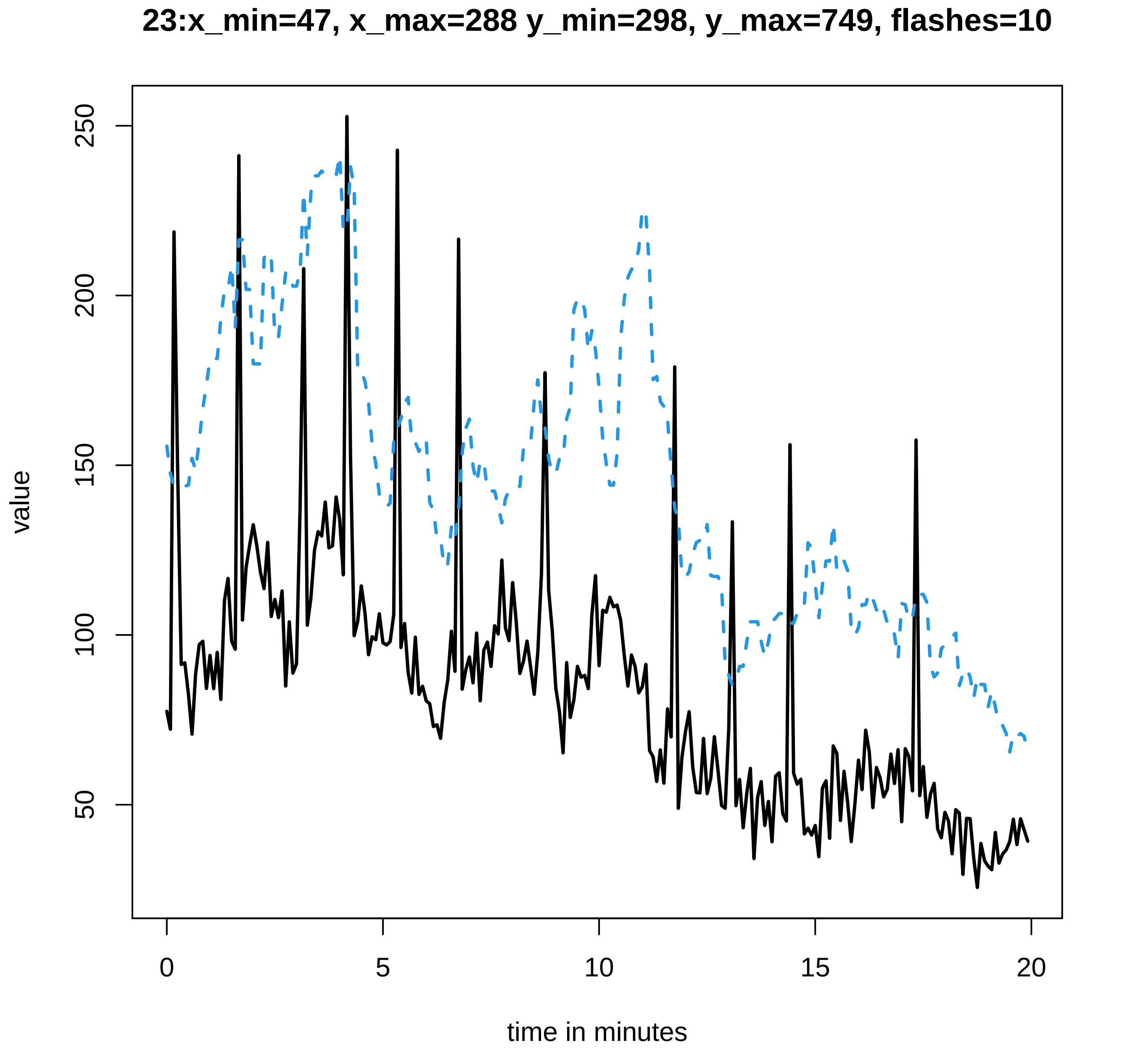

23:x_min=47, x_max=288 y_min=298, y_max=749, flashes=10
value
50
100
150
200
250
0
5
10
15
20
time in minutes

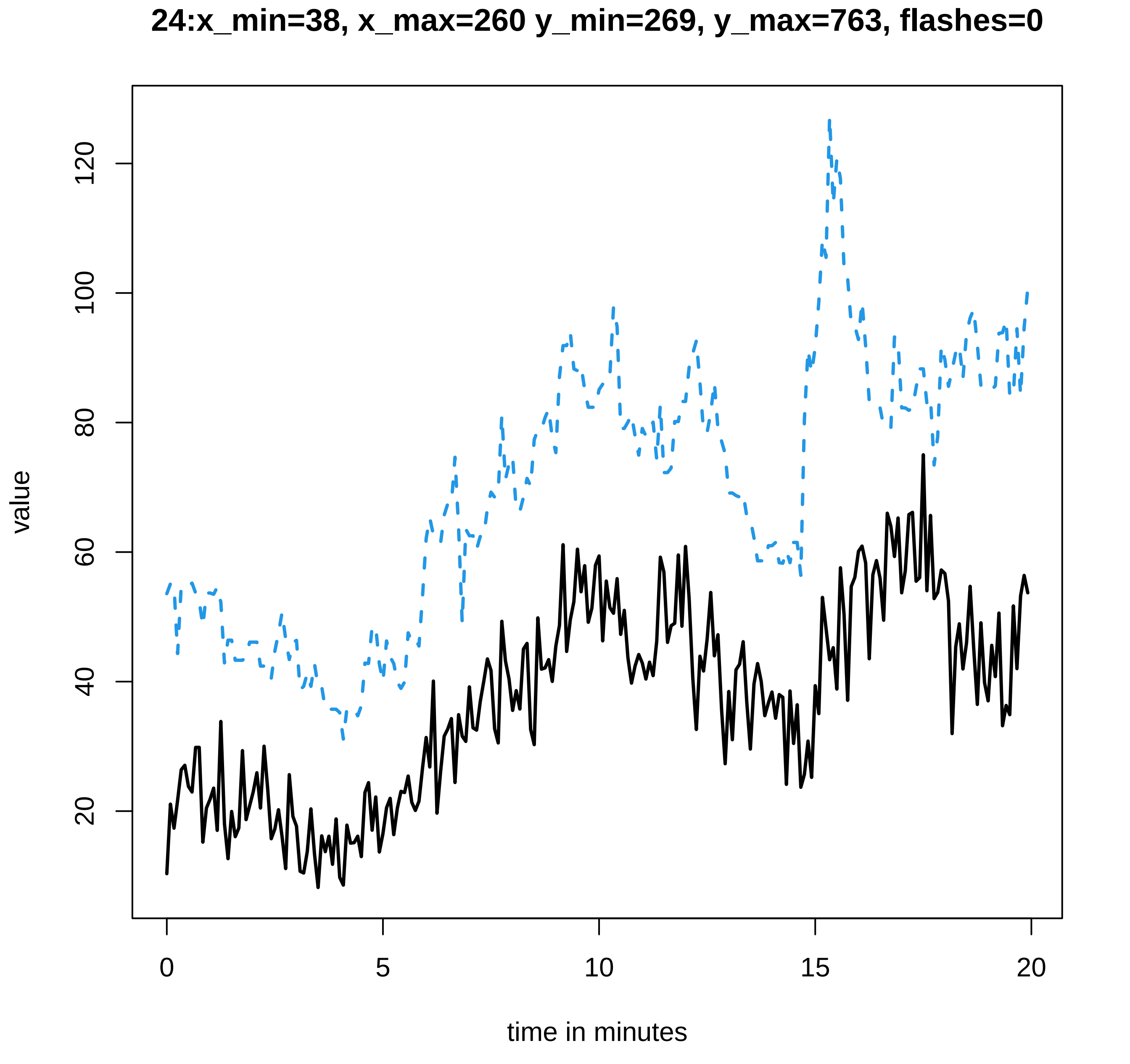
24:x_min=38, x_max=260 y_min=269, y_max=763, flashes=0
value
20
40
60
80
100
120
0
5
10
15
20
time in minutes

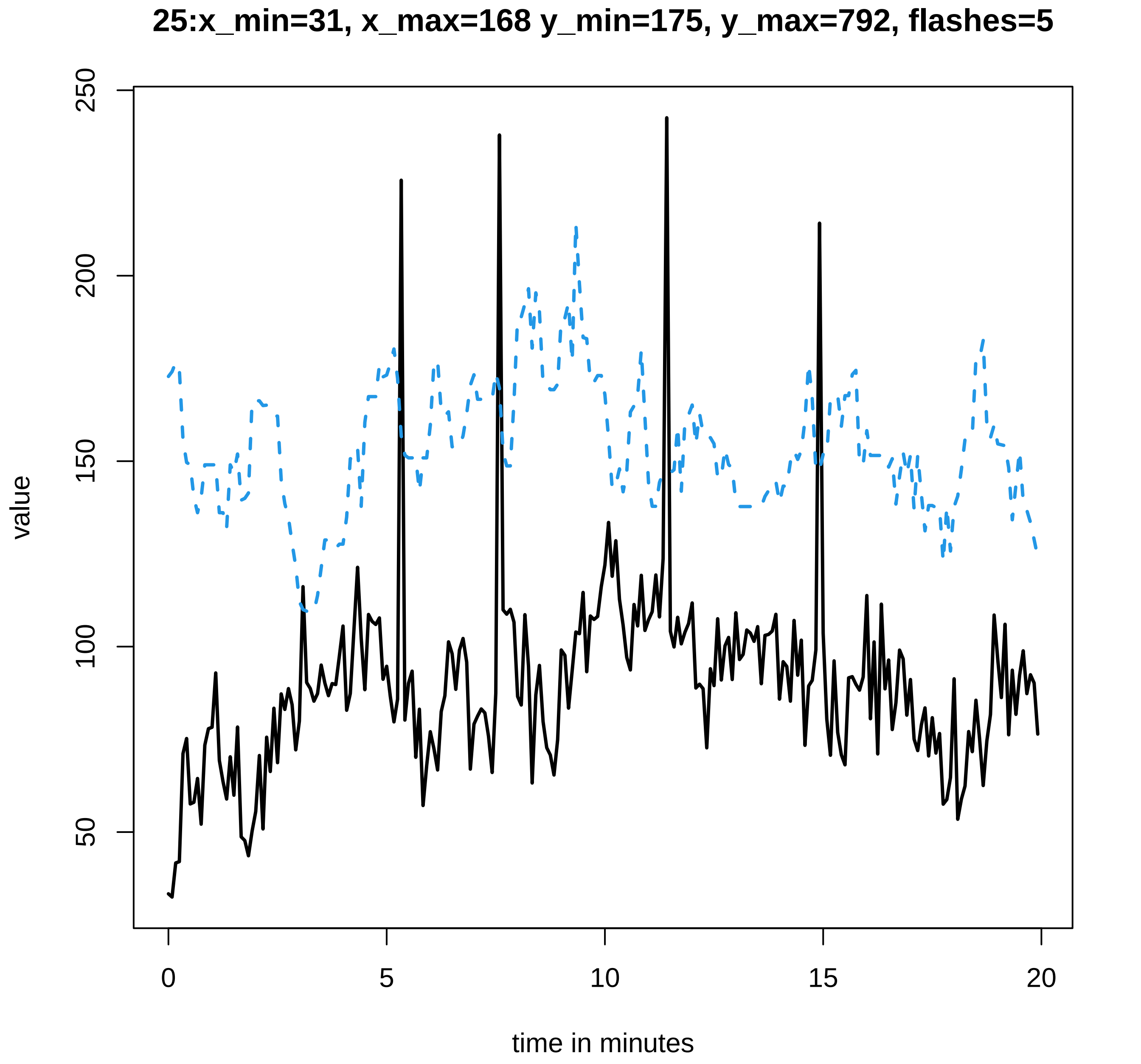

25:x_min=31, x_max=168 y_min=175, y_max=792, flashes=5
value
50
100
150
200
250
0
5
10
15
20
time in minutes

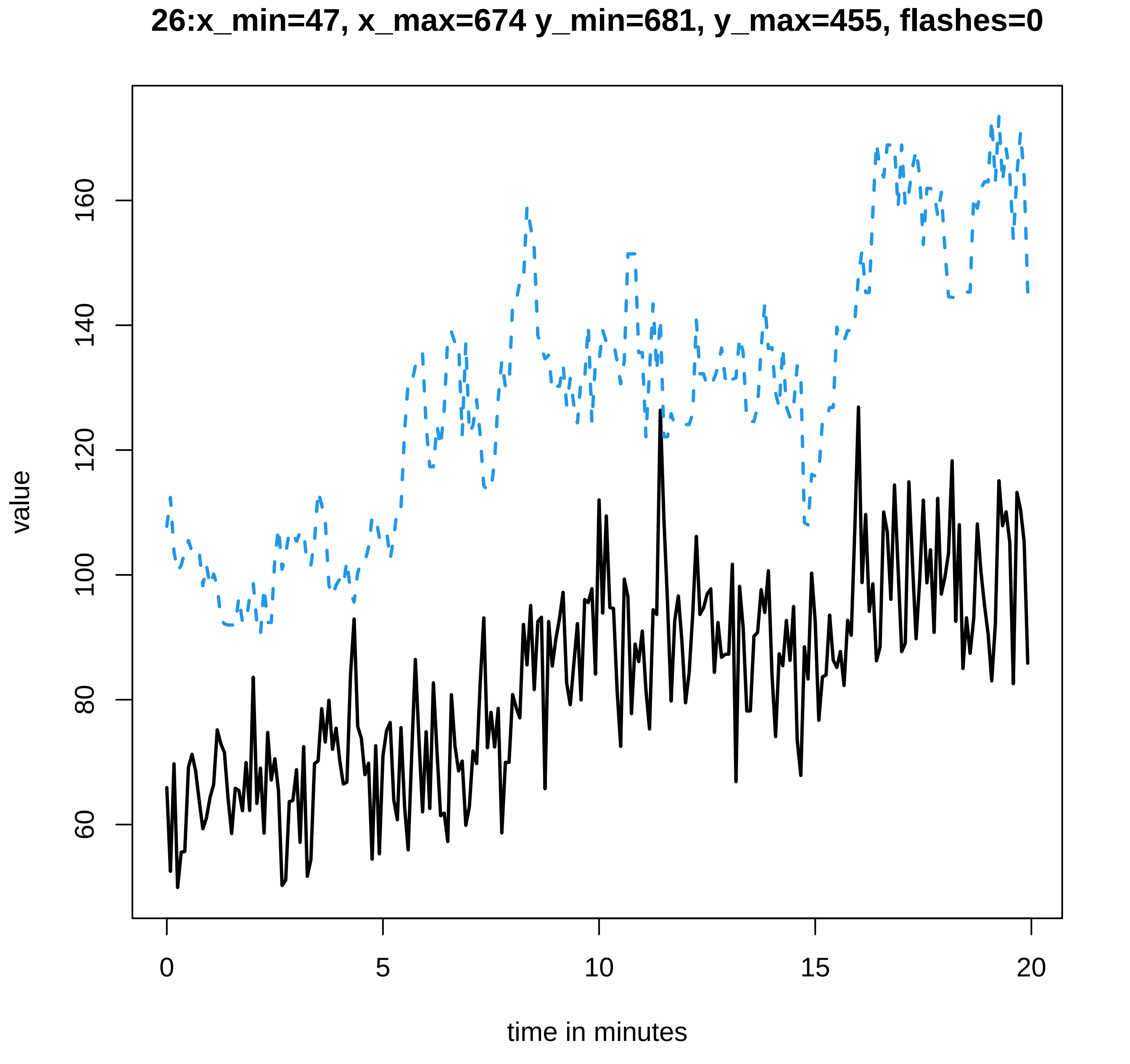

26:x_min=47, x_max=674 y_min=681, y_max=455, flashes=0
value
60
80
100
120
140
160
0
5
10
15
20
time in minutes

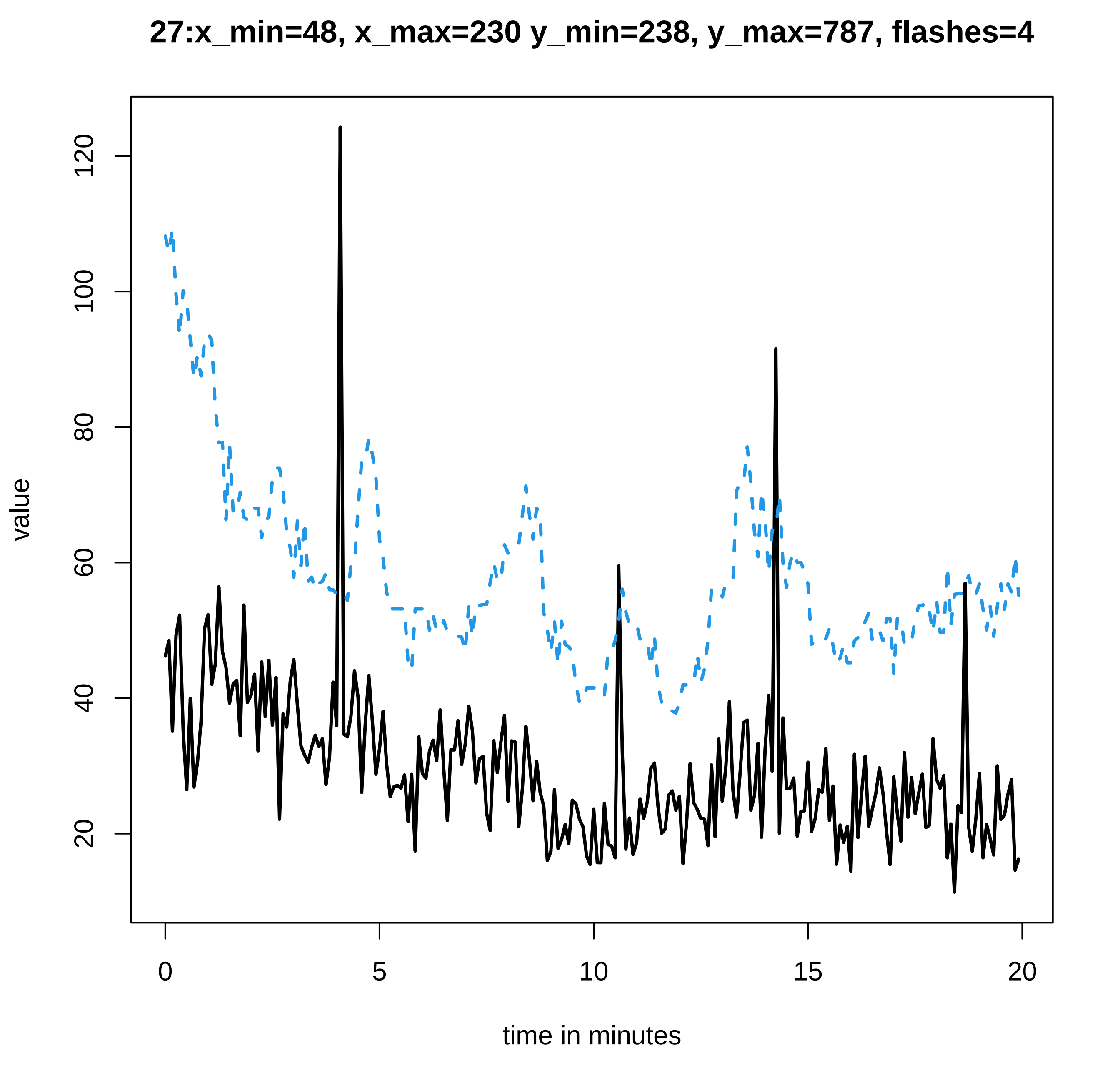
27:x_min=48, x_max=230 y_min=238, y_max=787, flashes=4
value
120
100
80
60
40
20
0
5
10
15
20
time in minutes

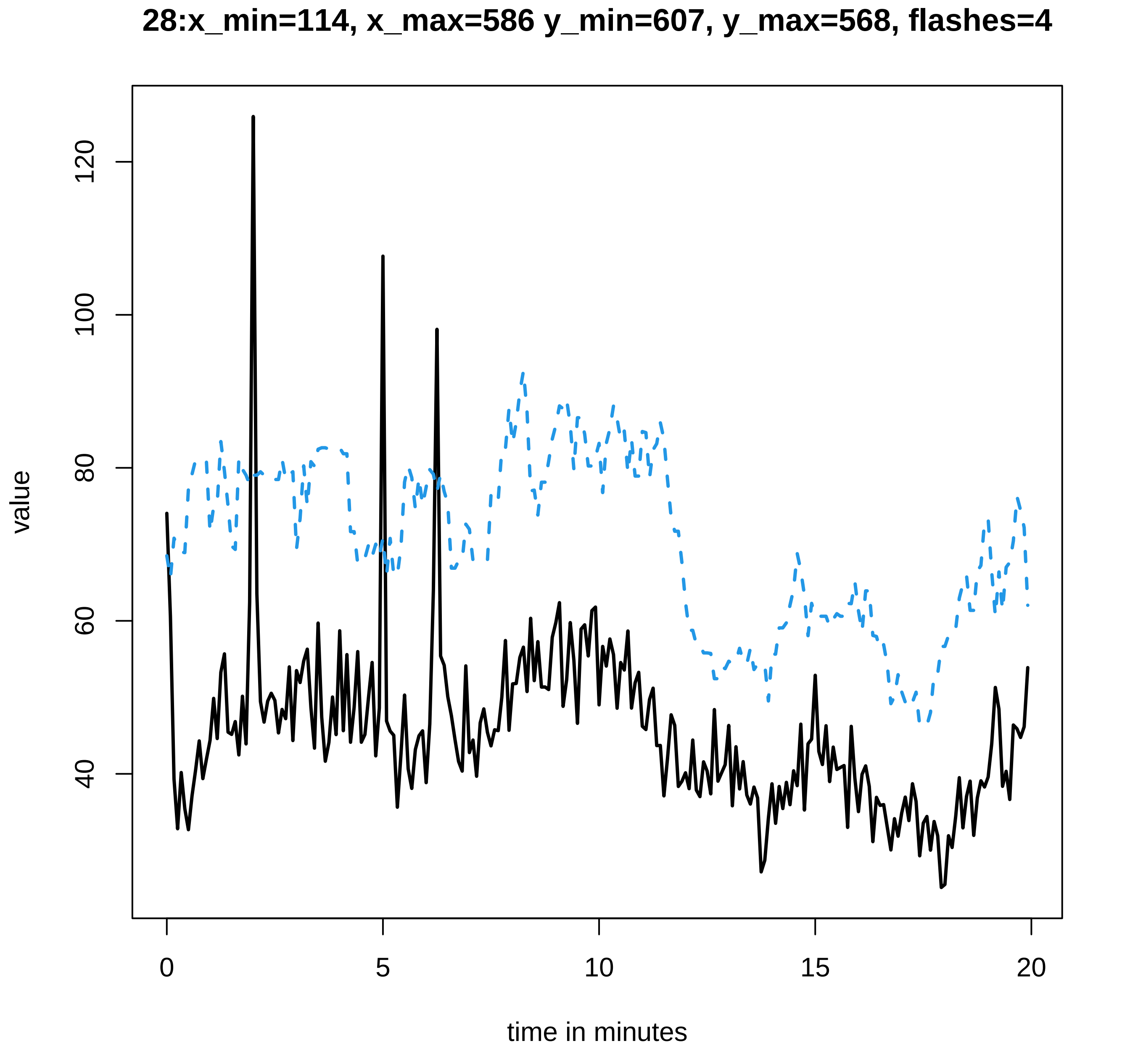

28:x_min=114, x_max=586 y_min=607, y_max=568, flashes=4
value
120
100
80
60
40
0
5
10
15
20
time in minutes

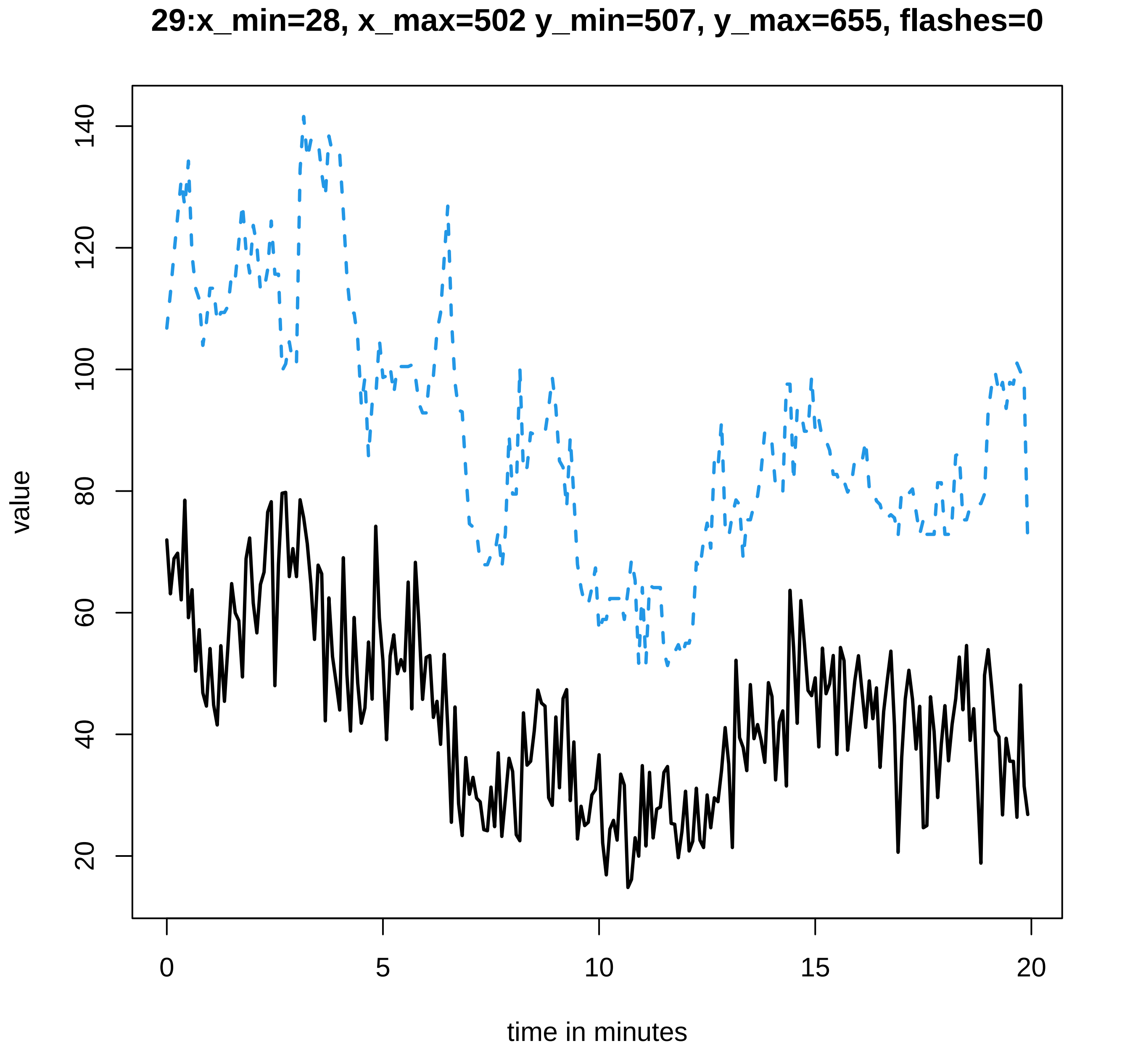
29:x_min=28, x_max=502 y_min=507, y_max=655, flashes=0
value
20
40
60
80
100
120
140
0
5
10
15
20
time in minutes

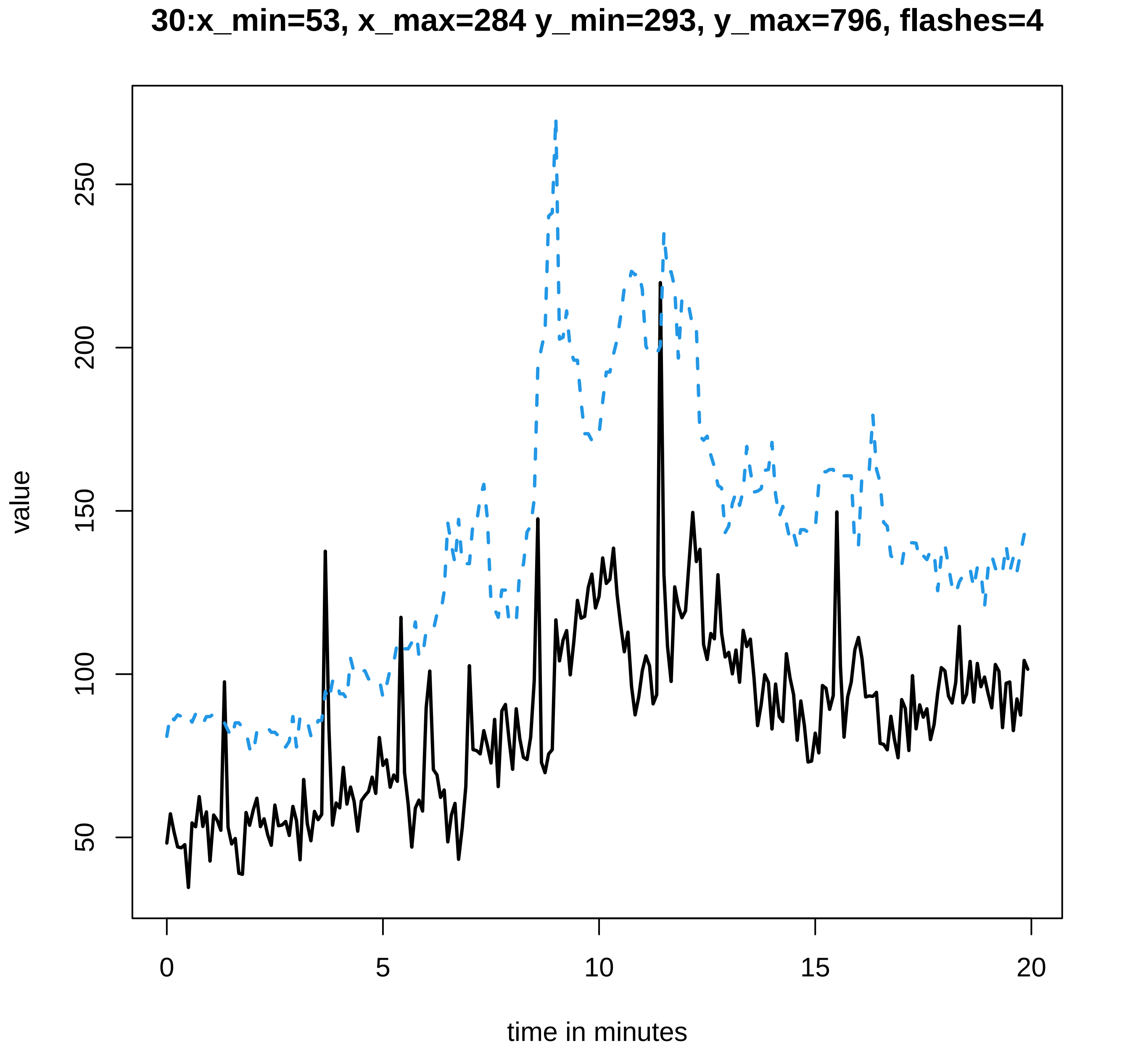

30:x_min=53, x_max=284 y_min=293, y_max=796, flashes=4
value
time in minutes
0
5
10
15
20
50
100
150
200
250

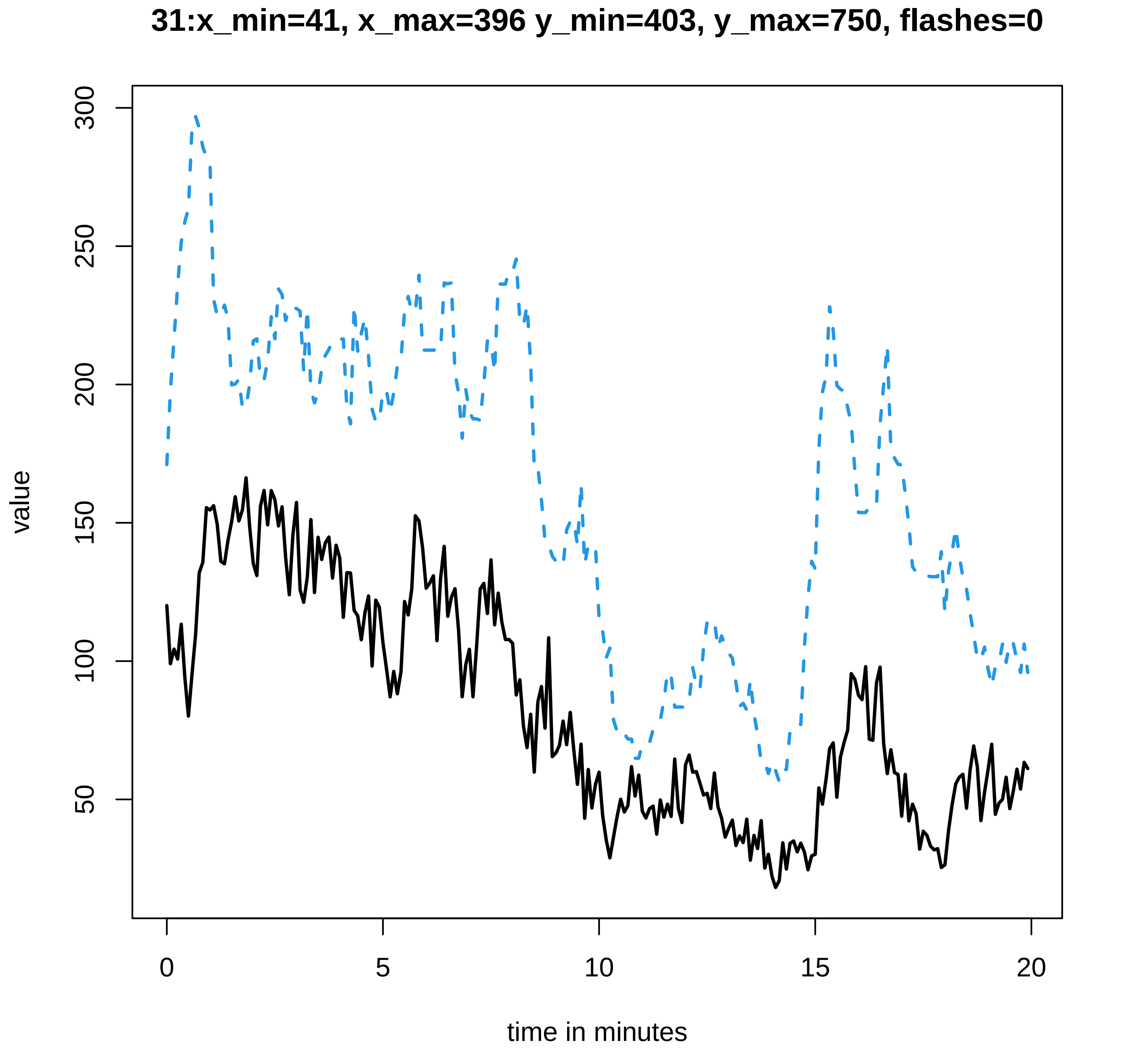

31:x_min=41, x_max=396 y_min=403, y_max=750, flashes=0
value
time in minutes
0
5
10
15
20
50
100
150
200
250
300

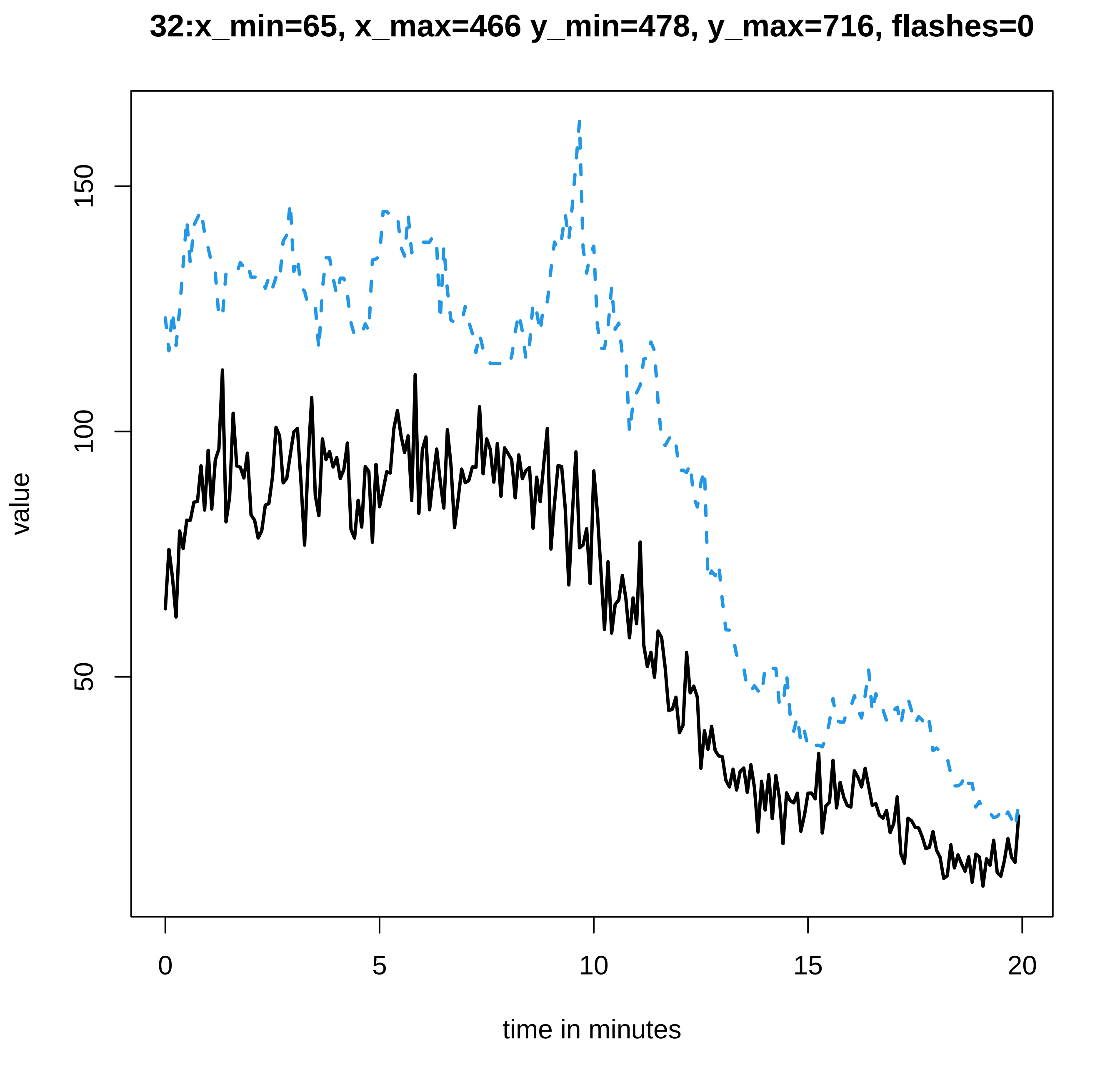
32:x_min=65, x_max=466 y_min=478, y_max=716, flashes=0
value
150
100
50
0
5
10
15
20
time in minutes

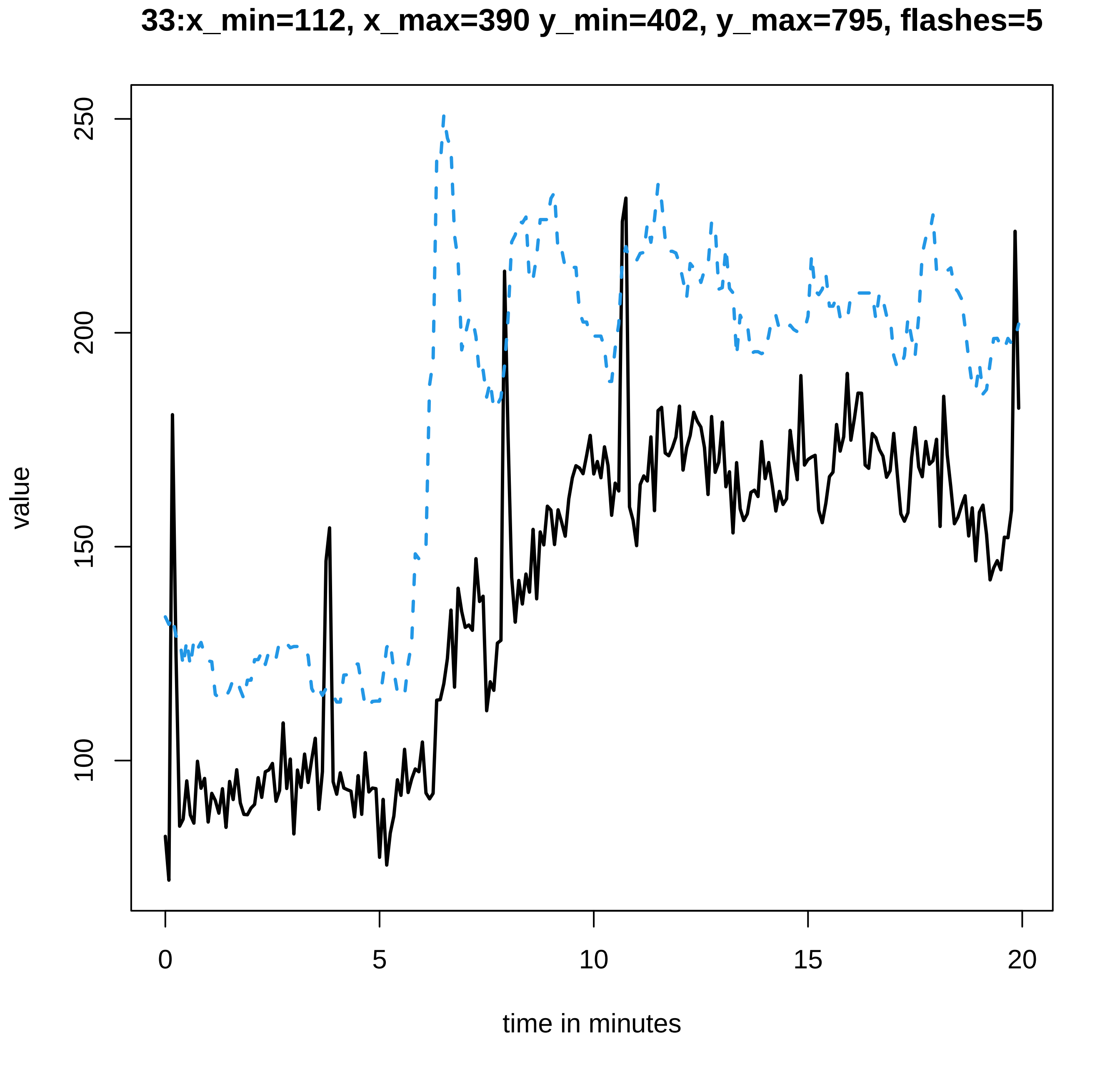
33:x_min=112, x_max=390 y_min=402, y_max=795, flashes=5
value
250
200
150
100
0
5
10
15
20
time in minutes

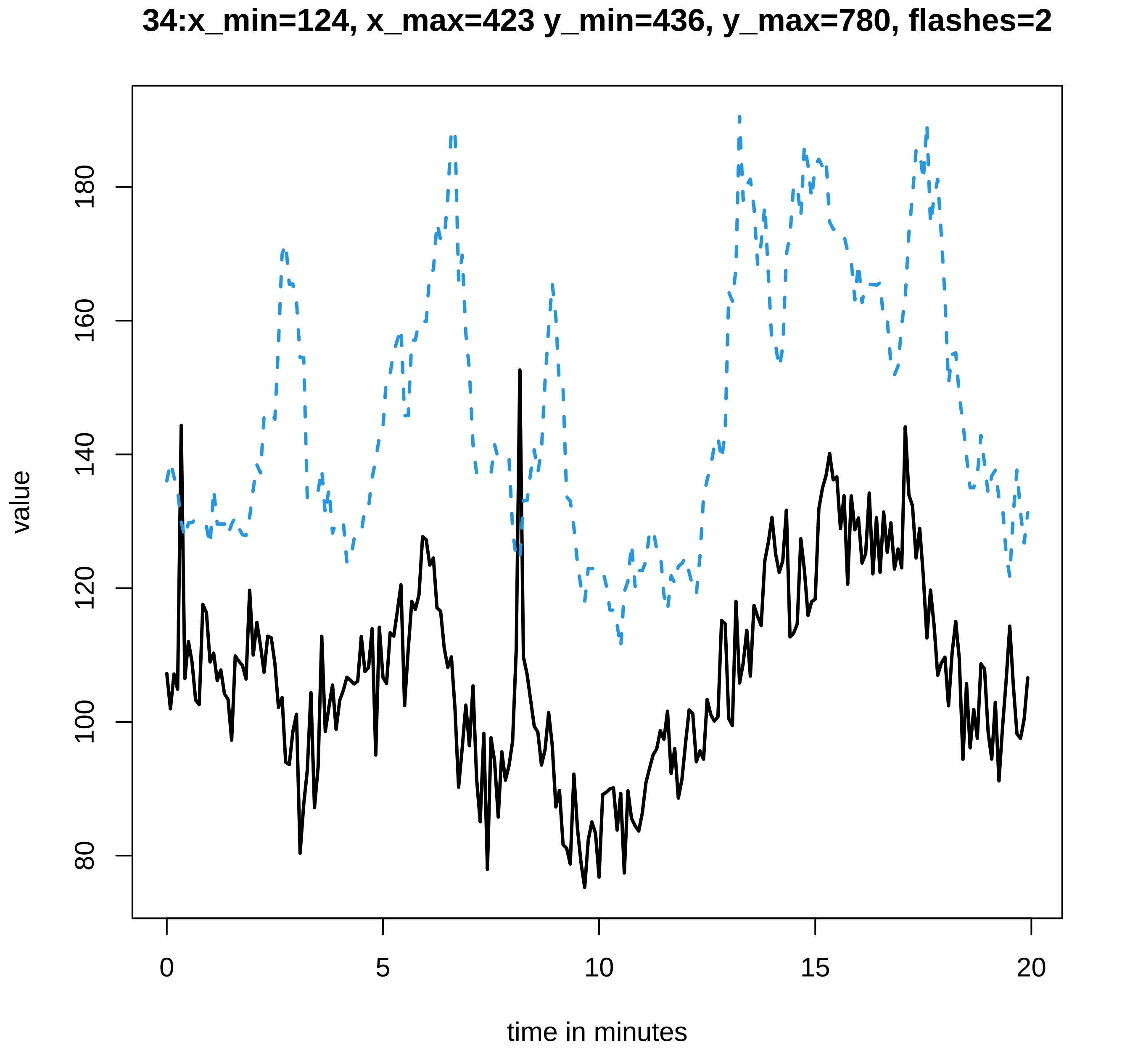
34:x_min=124, x_max=423 y_min=436, y_max=780, flashes=2
value
80
100
120
140
160
180
0
5
10
15
20
time in minutes

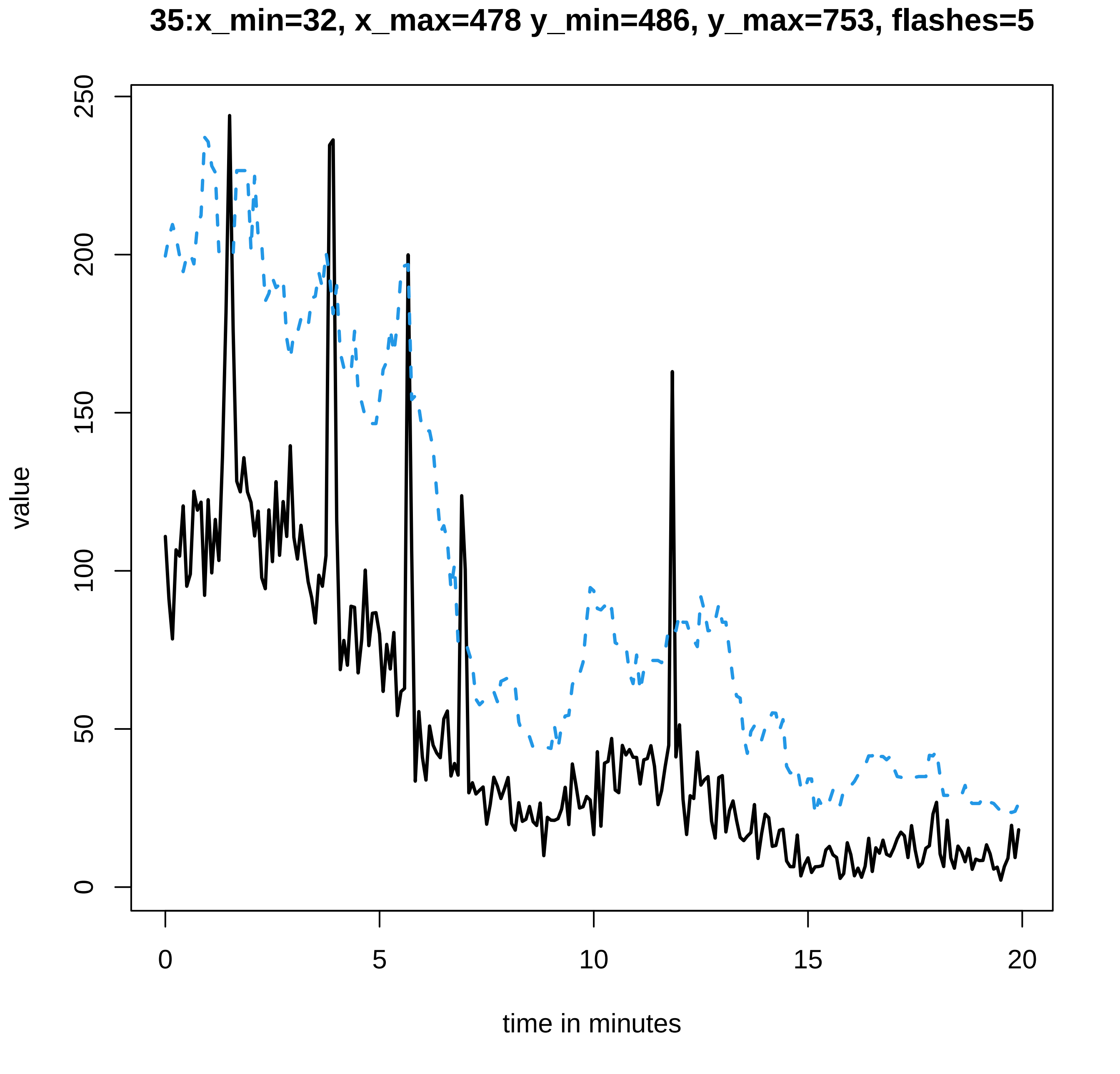

35:x_min=32, x_max=478 y_min=486, y_max=753, flashes=5
value
0
50
100
150
200
250
0
5
10
15
20
time in minutes

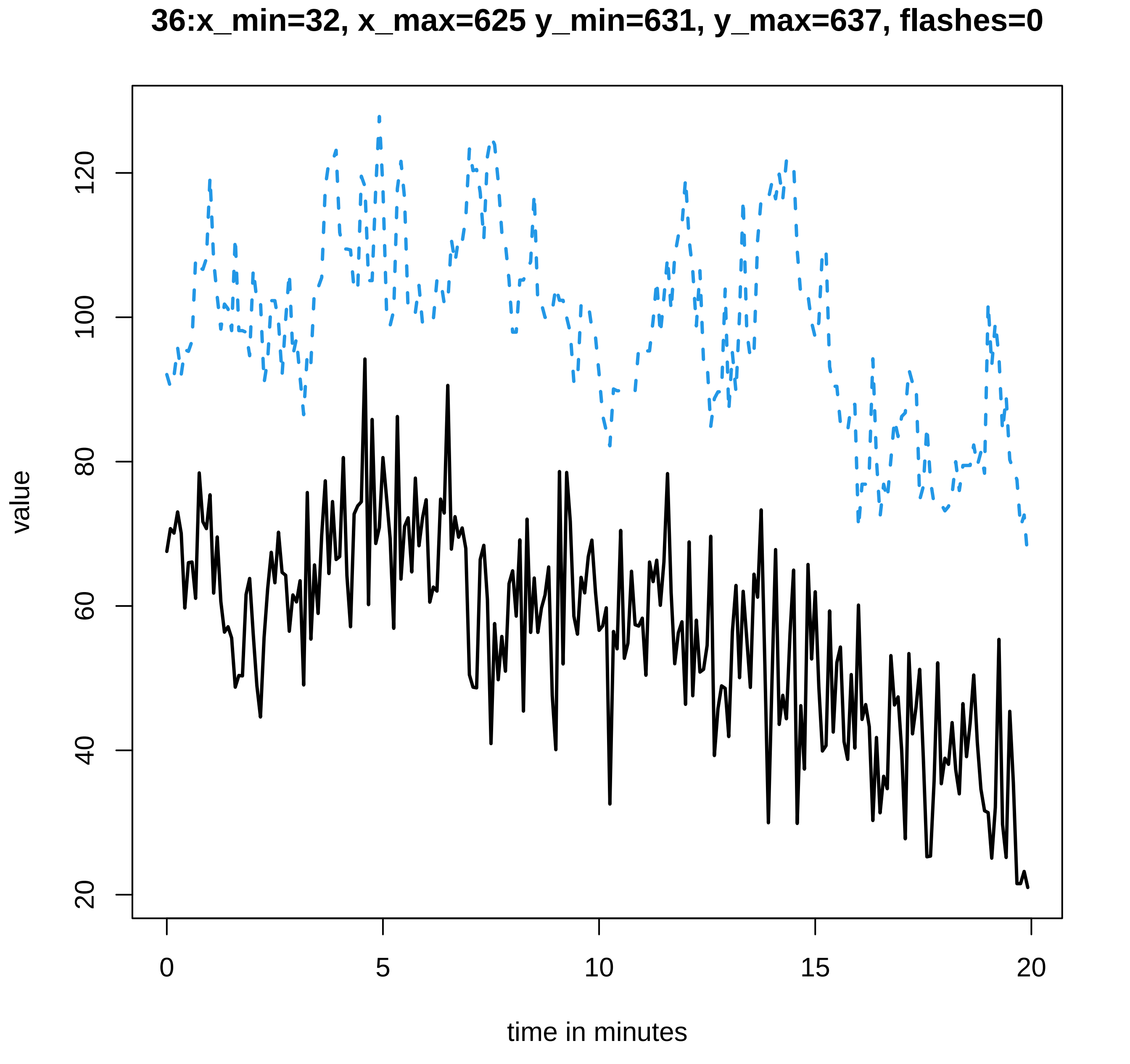
36:x_min=32, x_max=625 y_min=631, y_max=637, flashes=0
value
time in minutes
0
5
10
15
20
20
40
60
80
100
120

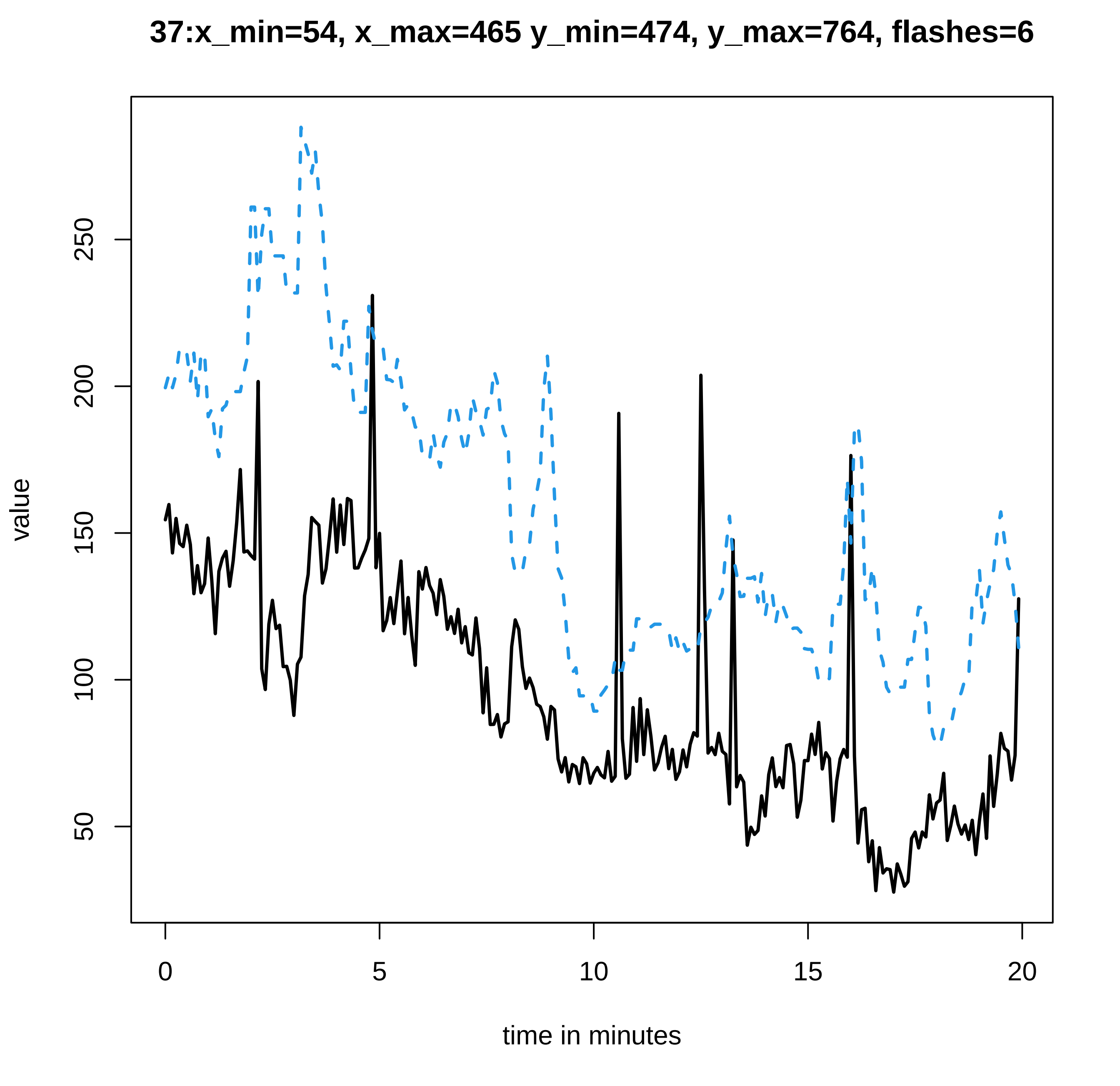

37:x_min=54, x_max=465 y_min=474, y_max=764, flashes=6
value
50
100
150
200
250
0
5
10
15
20
time in minutes

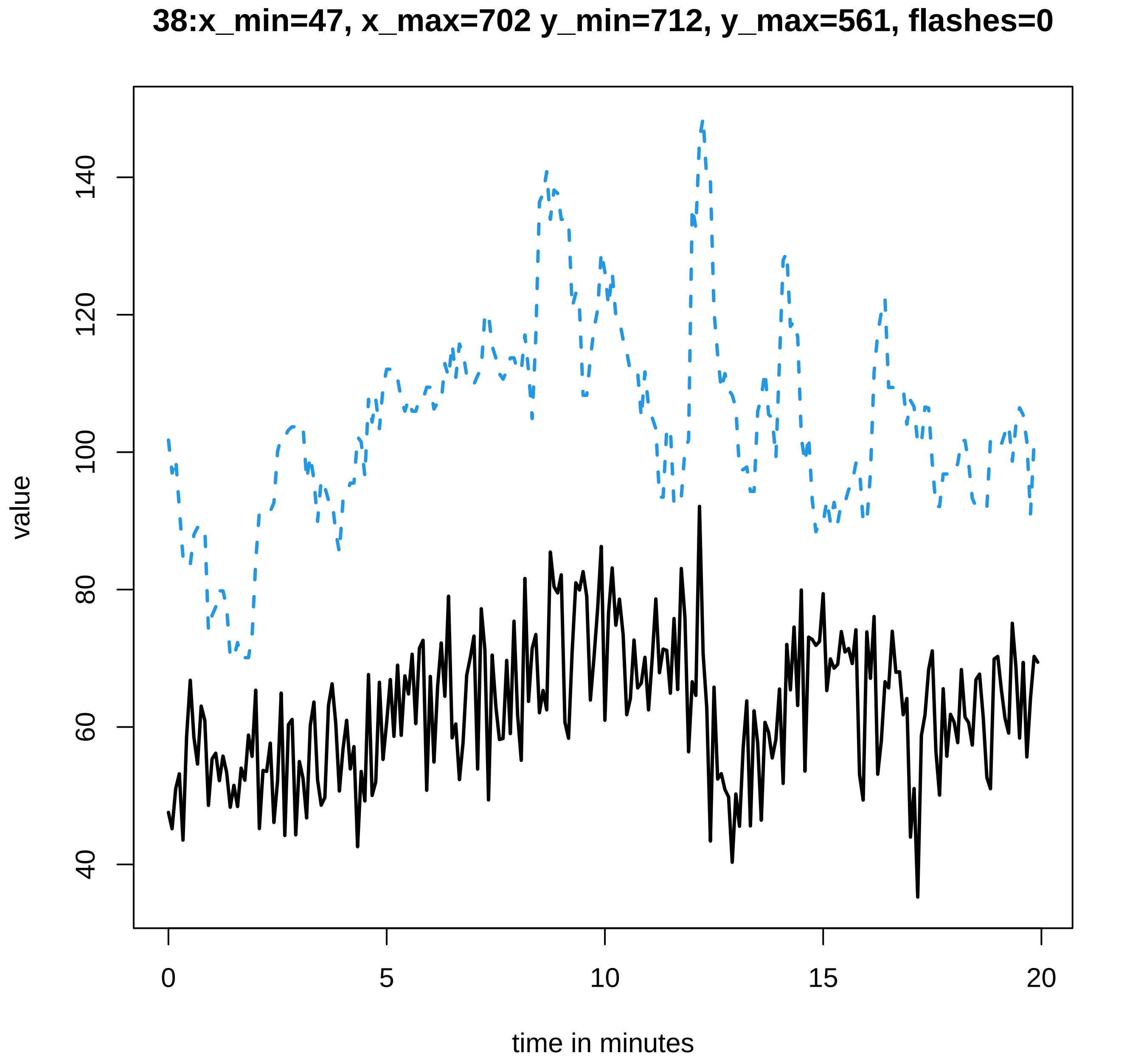
38:x_min=47, x_max=702 y_min=712, y_max=561, flashes=0
value
40
60
80
100
120
140
0
5
10
15
20
time in minutes

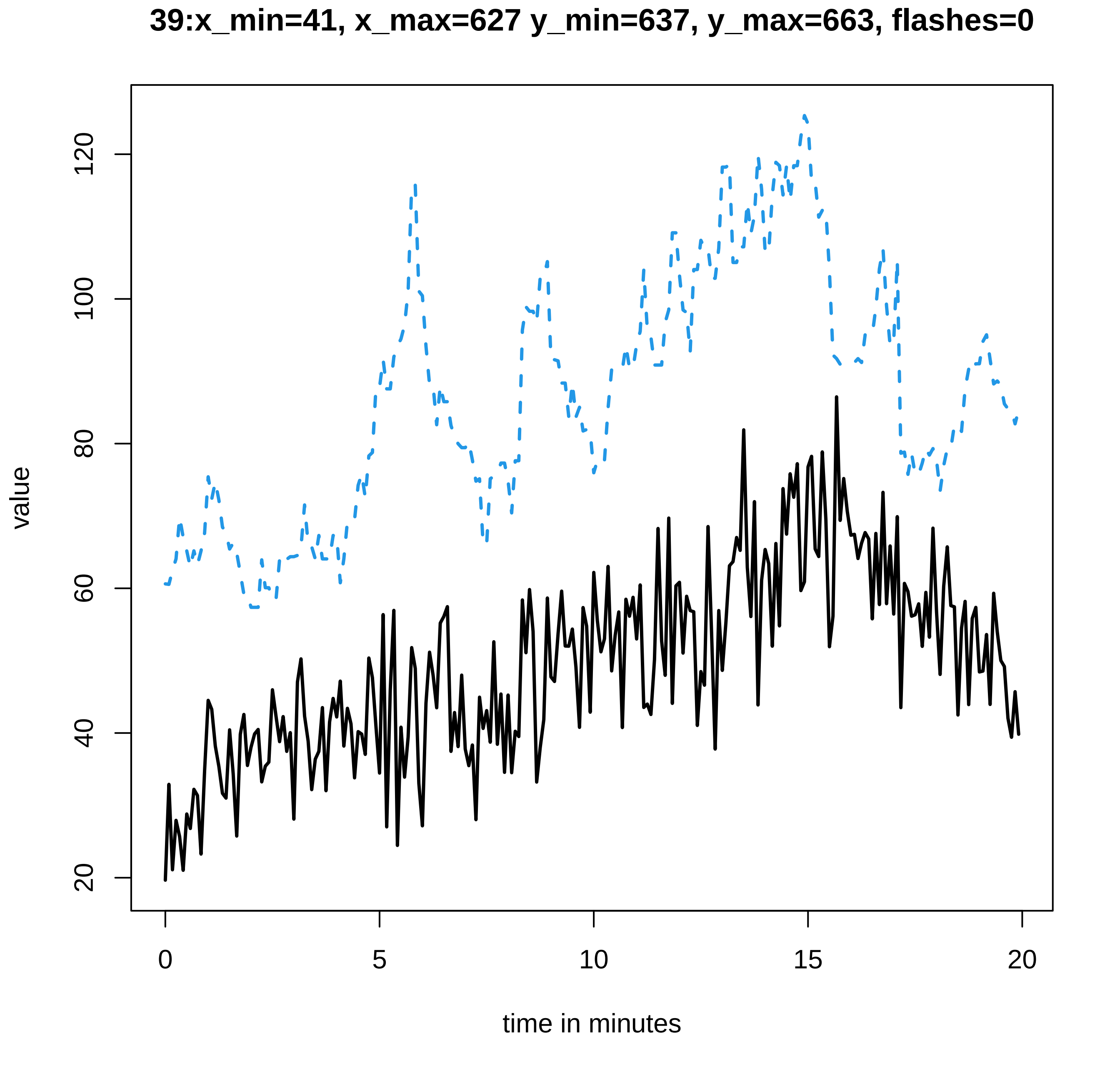

39:x_min=41, x_max=627 y_min=637, y_max=663, flashes=0
value
20
40
60
80
100
120
0
5
10
15
20
time in minutes

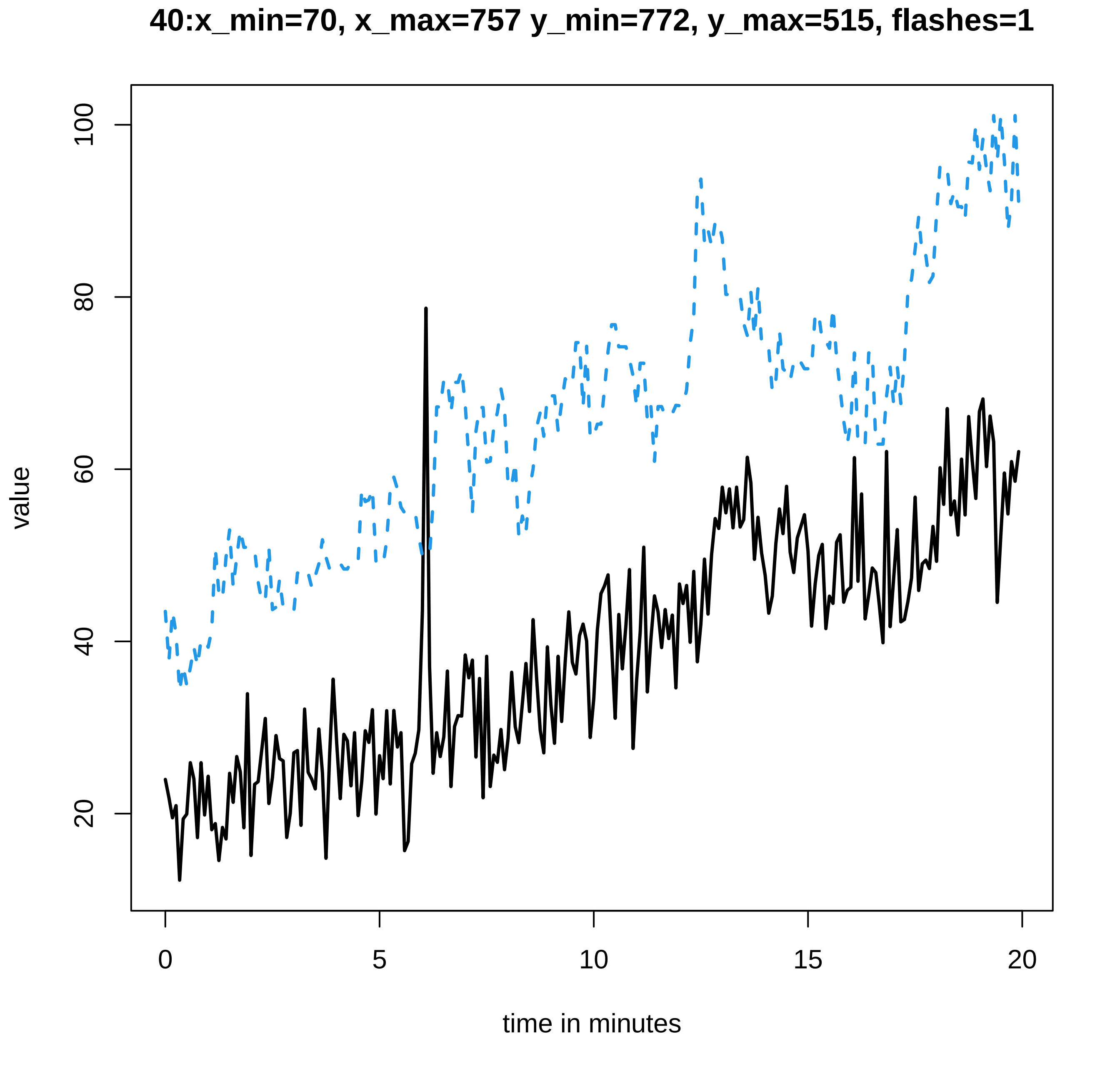
40:x_min=70, x_max=757 y_min=772, y_max=515, flashes=1
value
20
40
60
80
100
0
5
10
15
20
time in minutes

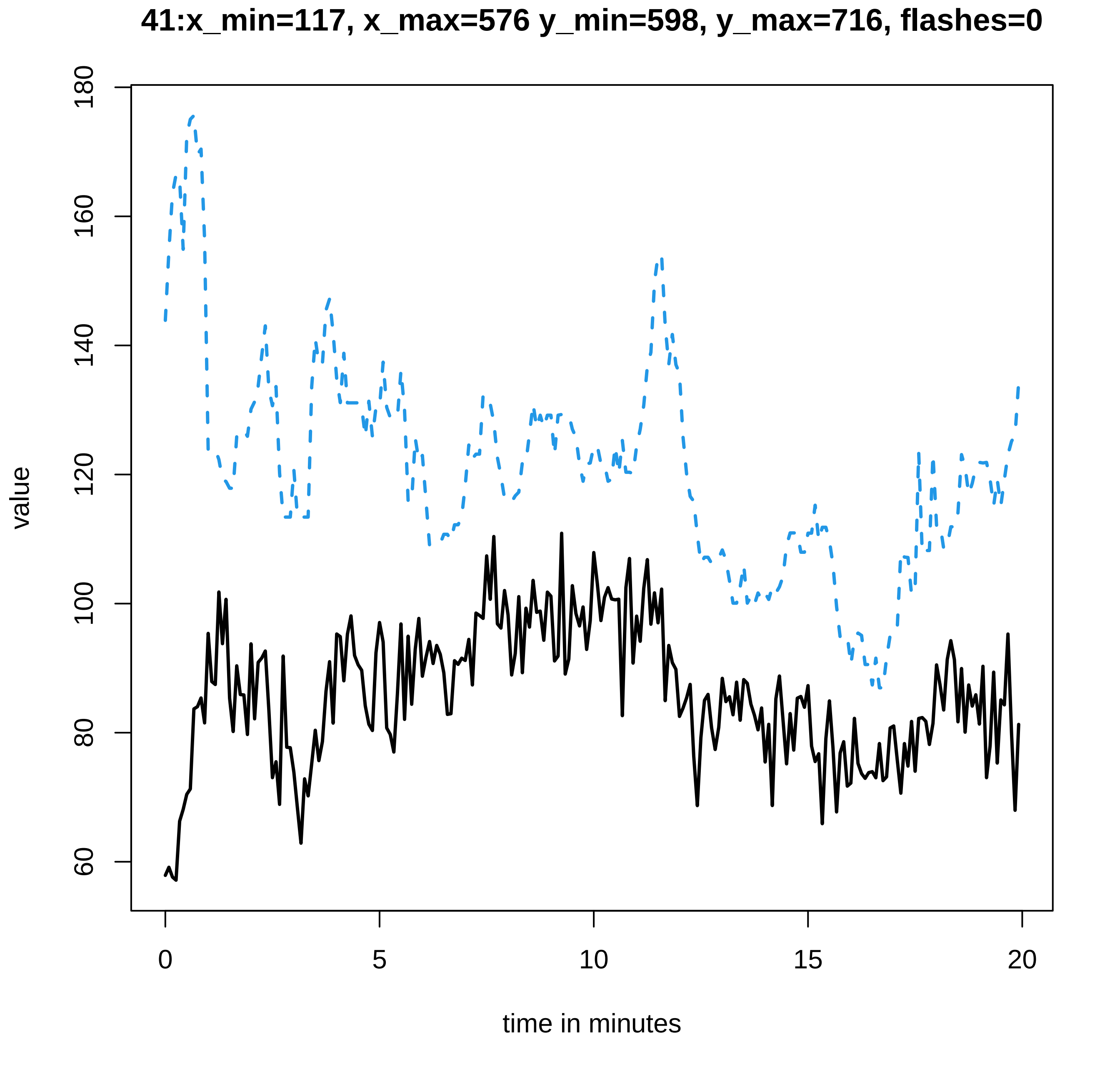
41:x_min=117, x_max=576 y_min=598, y_max=716, flashes=0
value
180
160
140
120
100
80
60
0
5
10
15
20
time in minutes

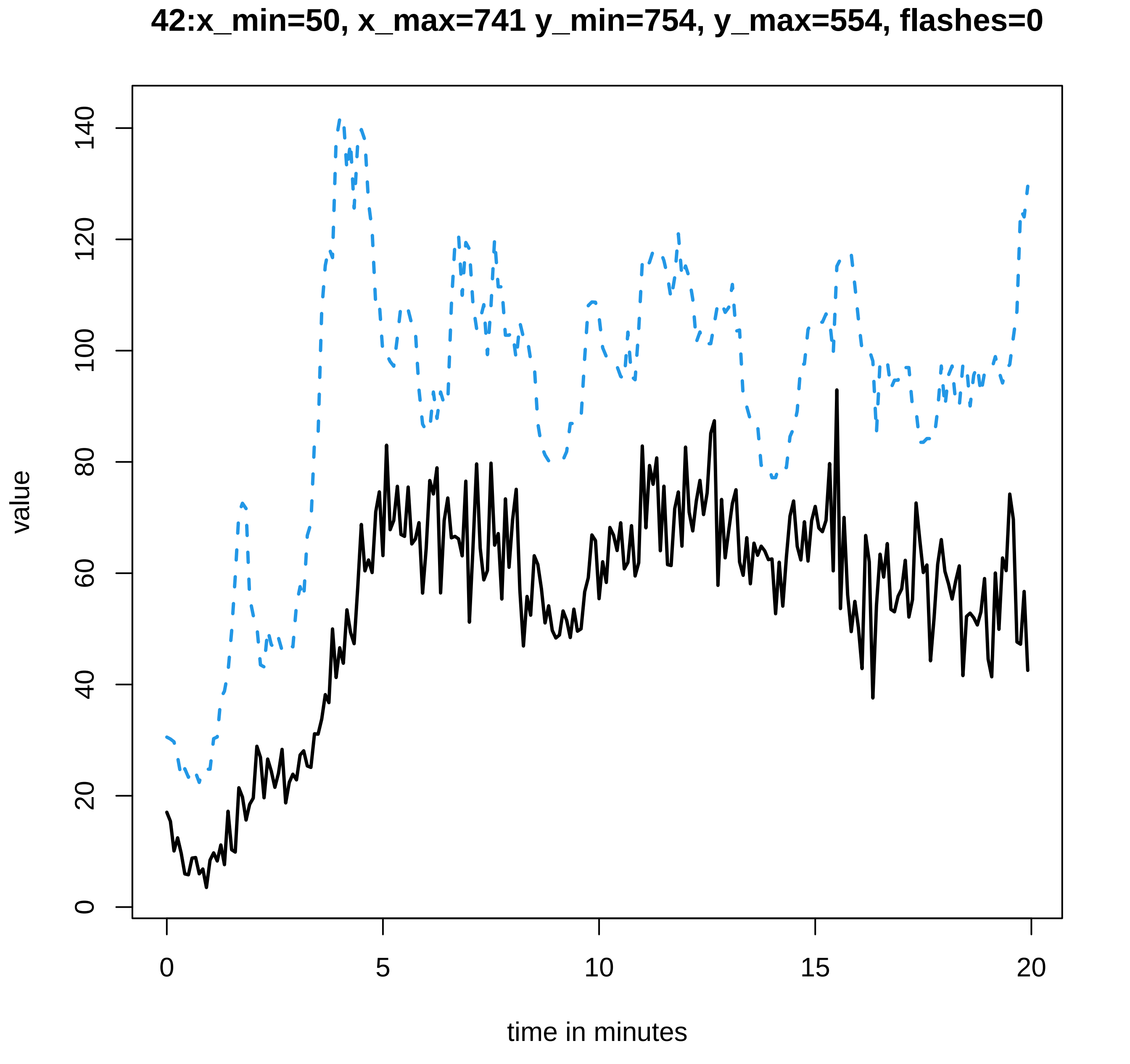

42:x_min=50, x_max=741 y_min=754, y_max=554, flashes=0
value
0
20
40
60
80
100
120
140
0
5
10
15
20
time in minutes

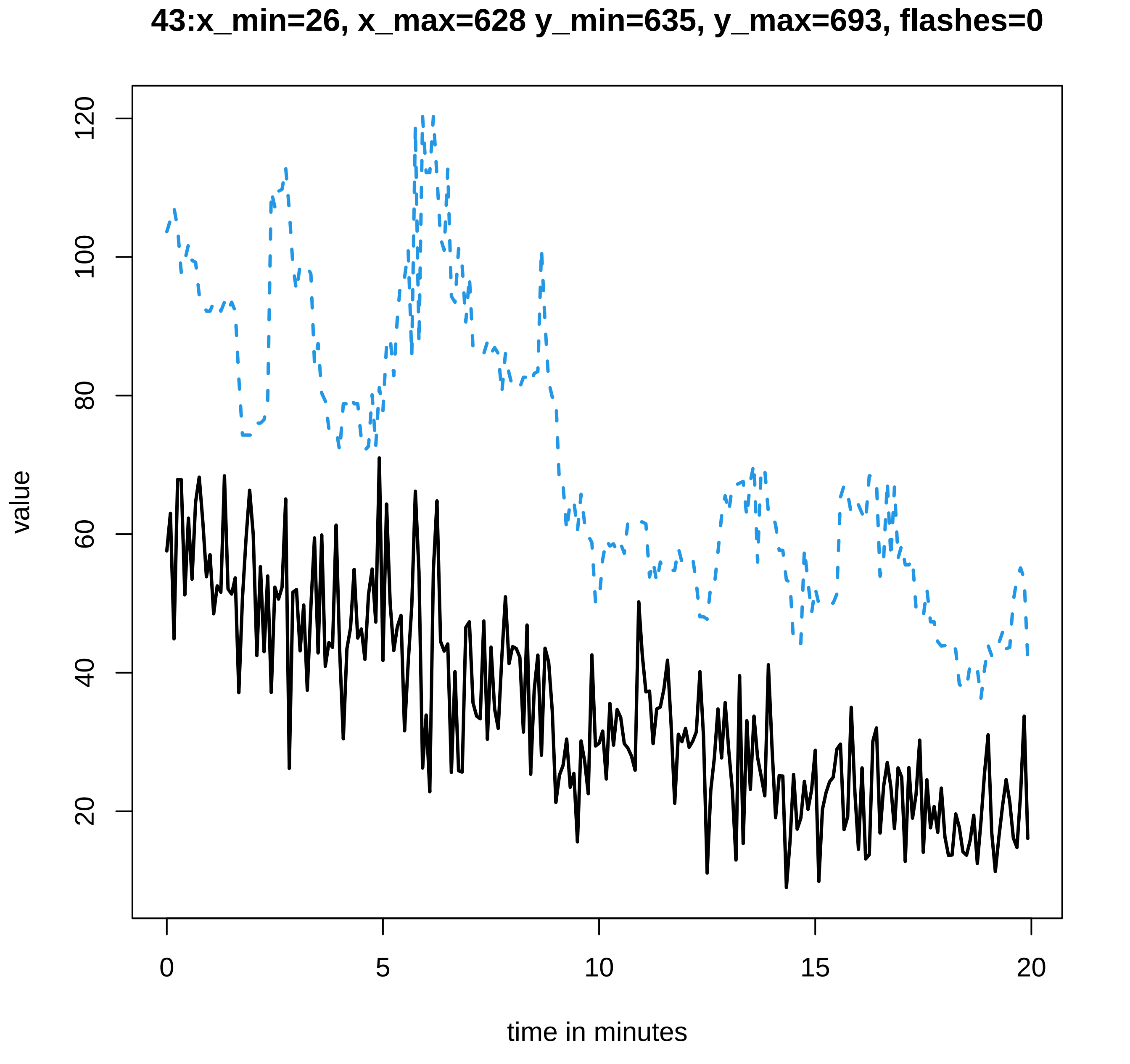
43:x_min=26, x_max=628 y_min=635, y_max=693, flashes=0
value
20
40
60
80
100
120
0
5
10
15
20
time in minutes

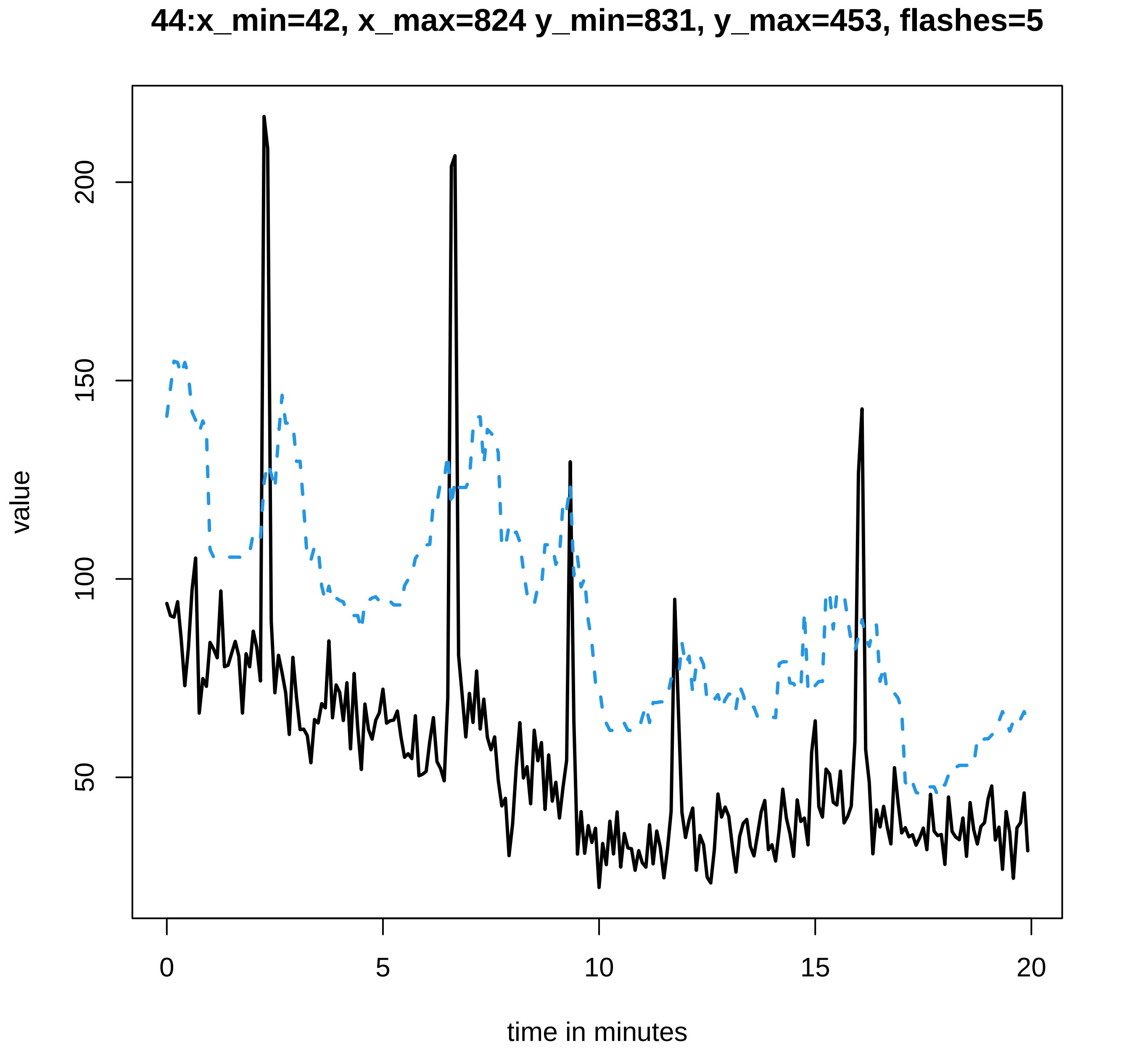
44:x_min=42, x_max=824 y_min=831, y_max=453, flashes=5
value
200
150
100
50
0
5
10
15
20
time in minutes

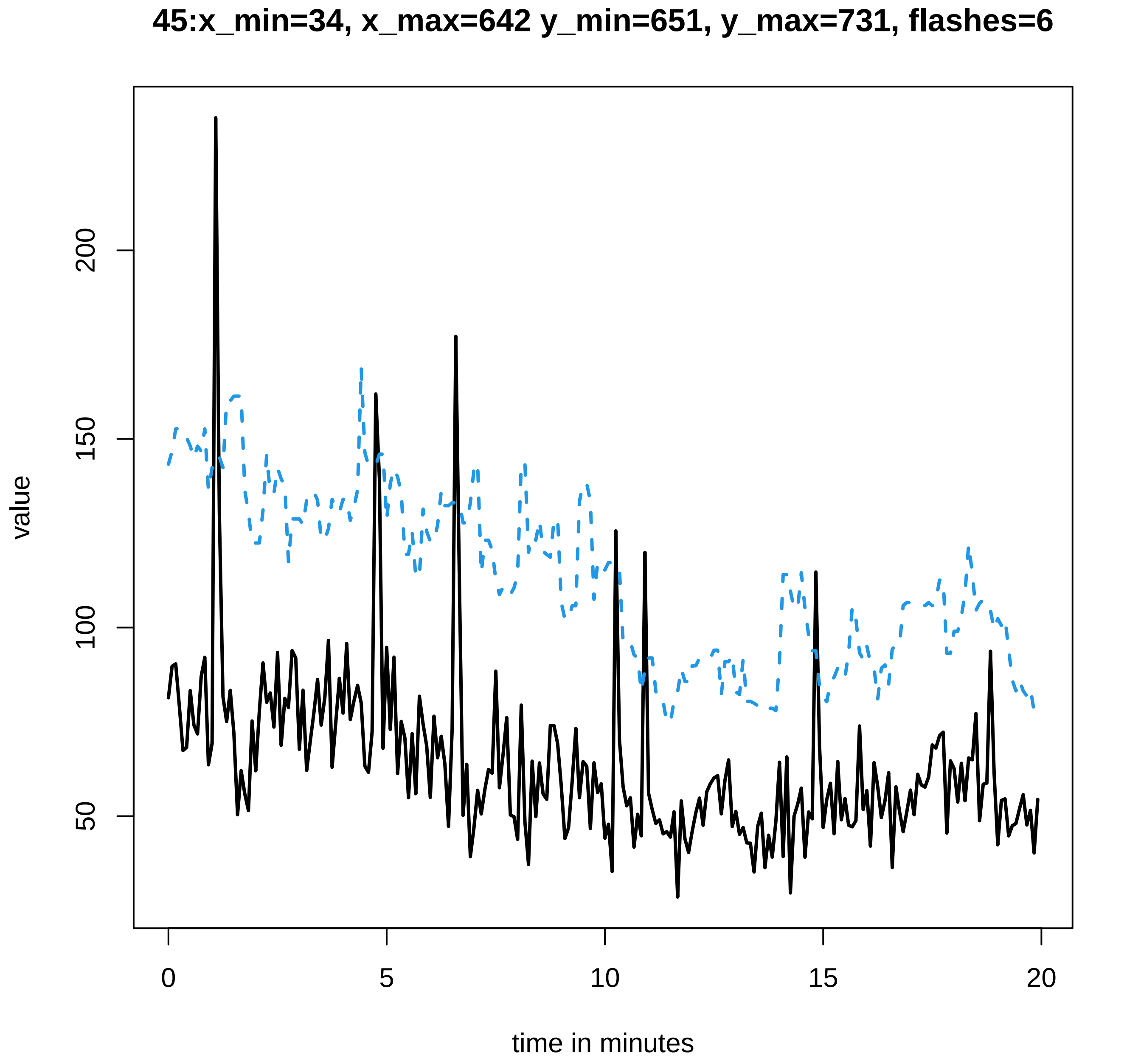
45:x_min=34, x_max=642 y_min=651, y_max=731, flashes=6
value
50
100
150
200
0
5
10
15
20
time in minutes

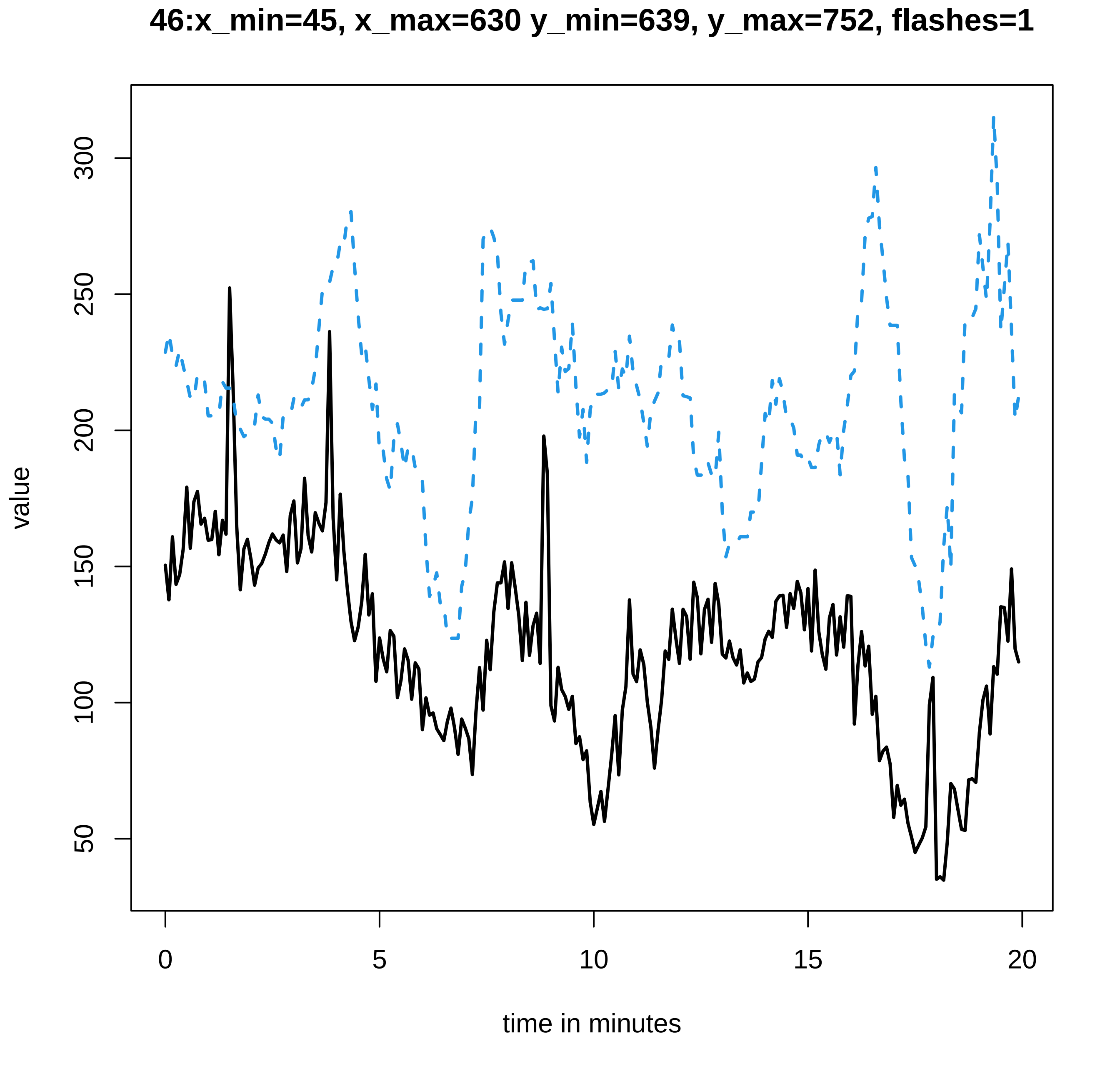

46:x_min=45, x_max=630 y_min=639, y_max=752, flashes=1
value
300
250
200
150
100
50
0
5
10
15
20
time in minutes

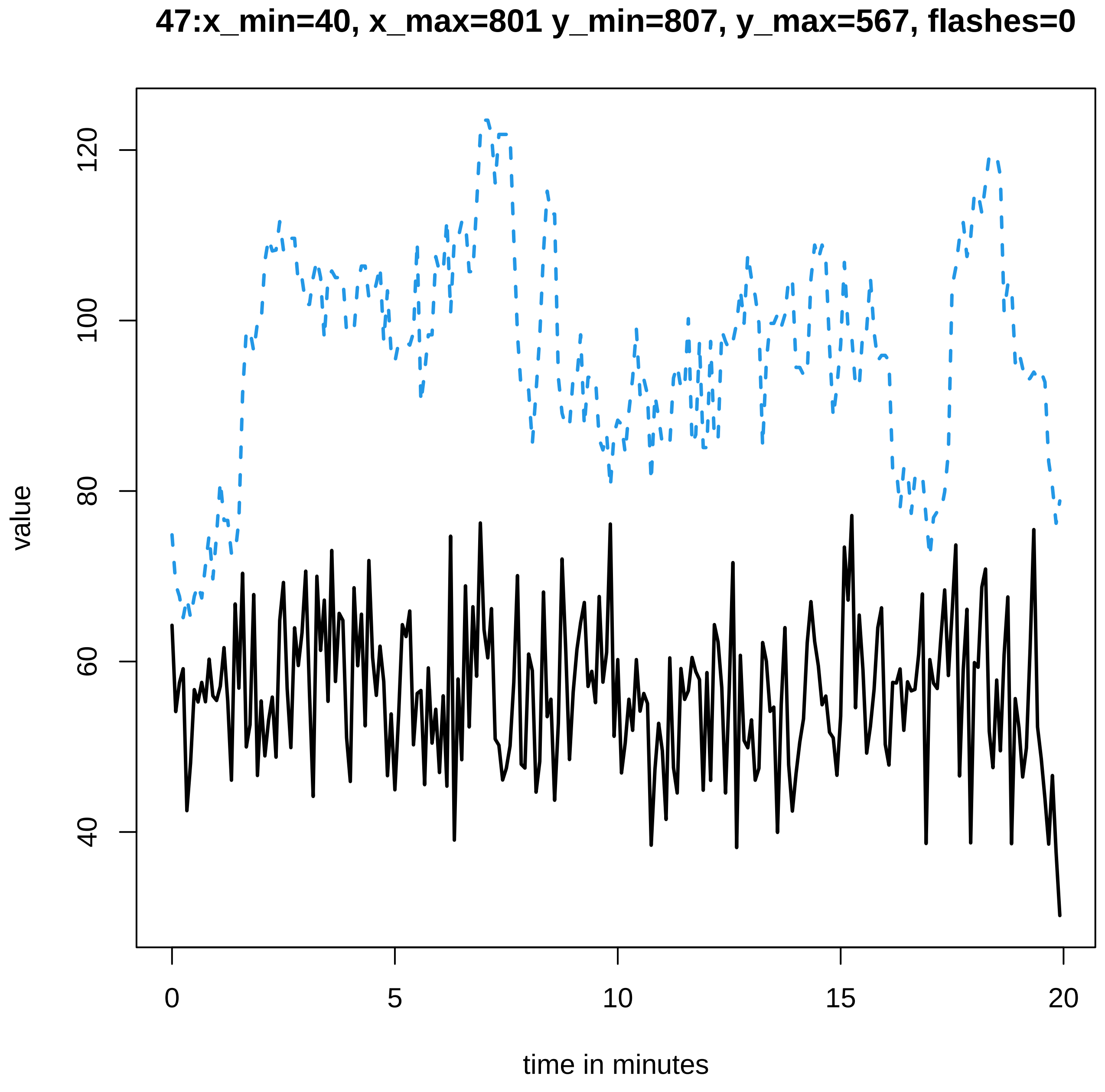
47:x_min=40, x_max=801 y_min=807, y_max=567, flashes=0
value
time in minutes
120
100
80
60
40
0
5
10
15
20

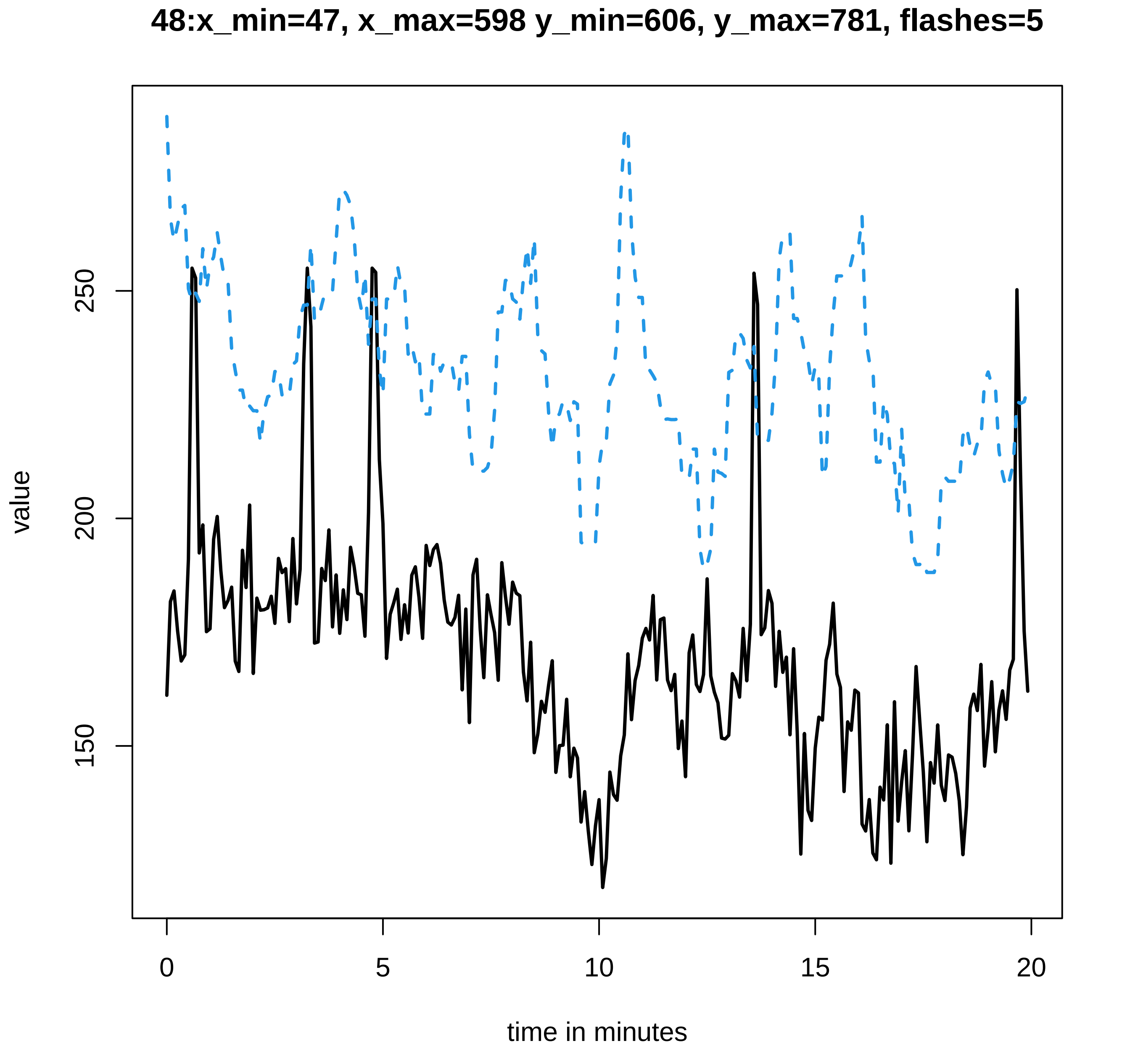
48:x_min=47, x_max=598 y_min=606, y_max=781, flashes=5
value
150
200
250
0
5
10
15
20
time in minutes

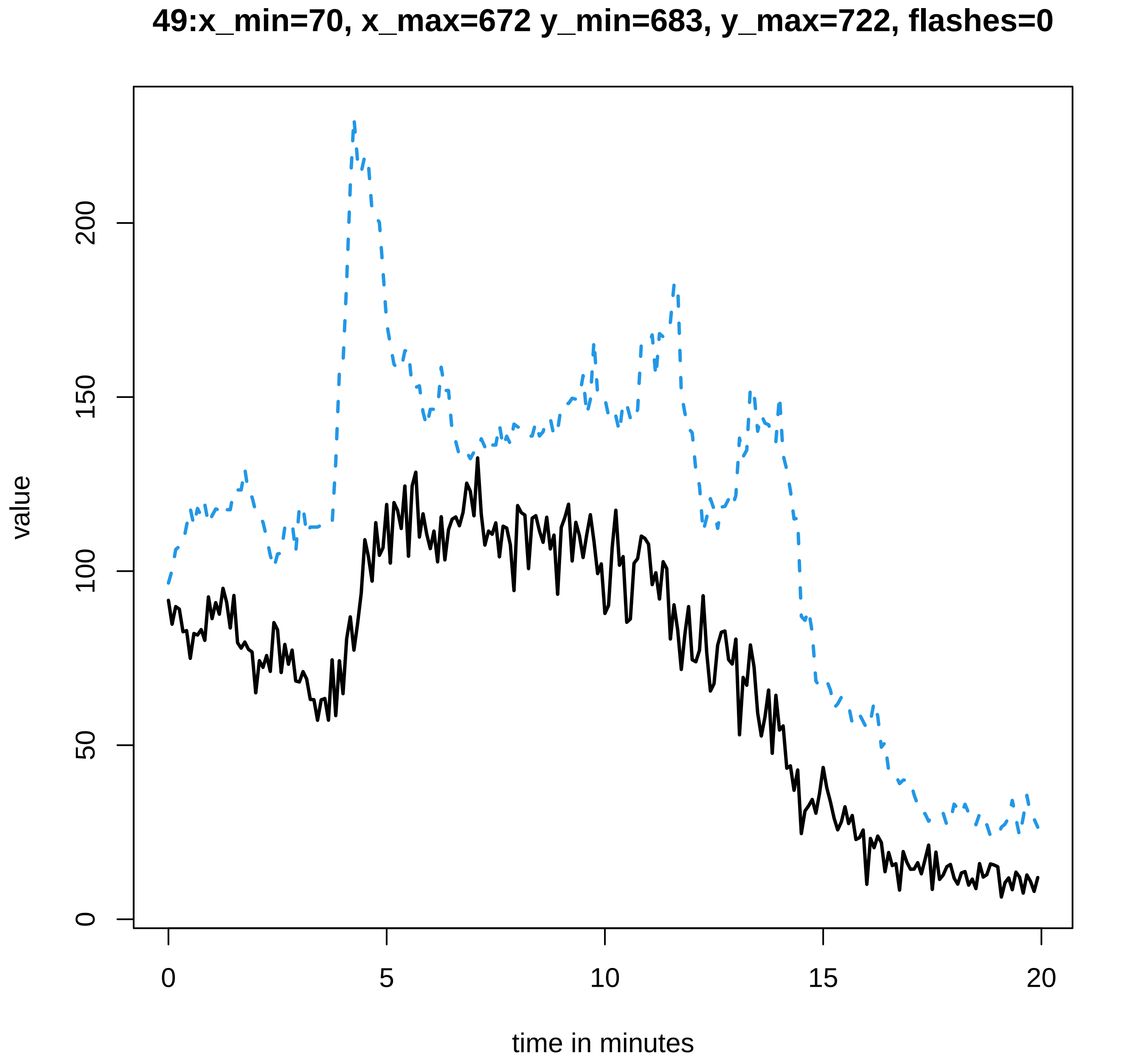

49:x_min=70, x_max=672 y_min=683, y_max=722, flashes=0
value
0
50
100
150
200
0
5
10
15
20
time in minutes

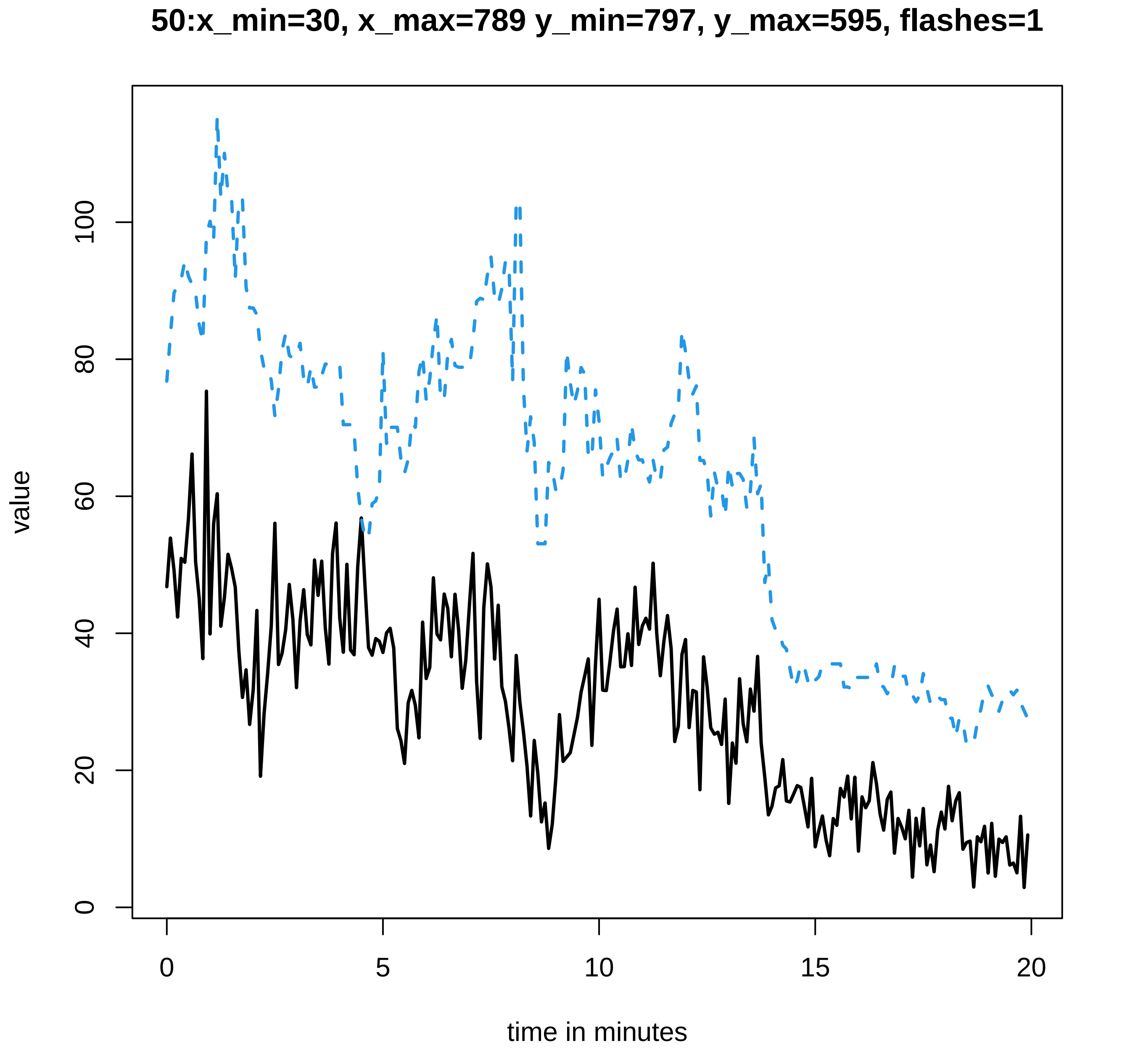
50:x_min=30, x_max=789 y_min=797, y_max=595, flashes=1
value
0
20
40
60
80
100
0
5
10
15
20
time in minutes

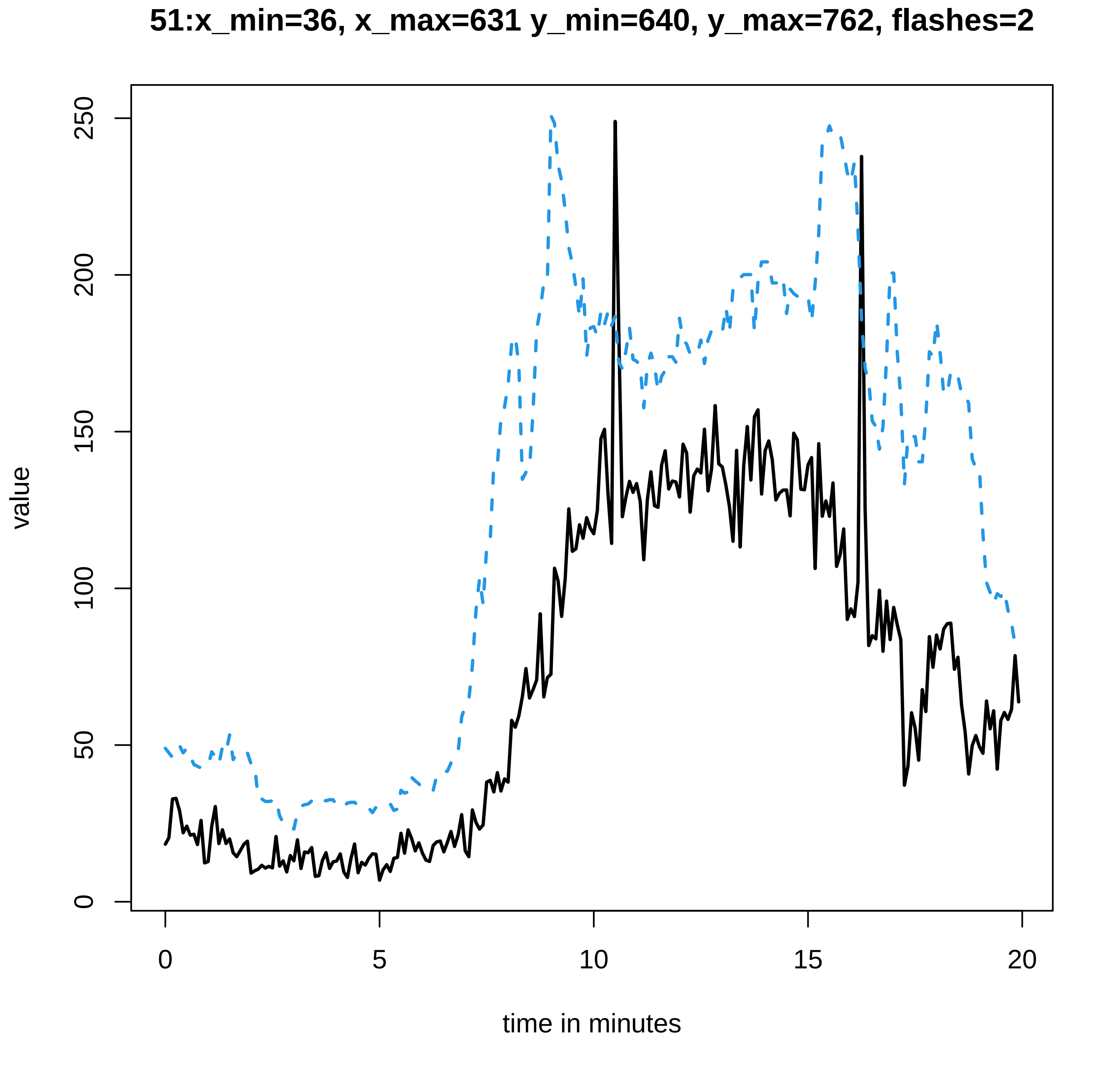
51:x_min=36, x_max=631 y_min=640, y_max=762, flashes=2
value
0
50
100
150
200
250
time in minutes
0
5
10
15
20

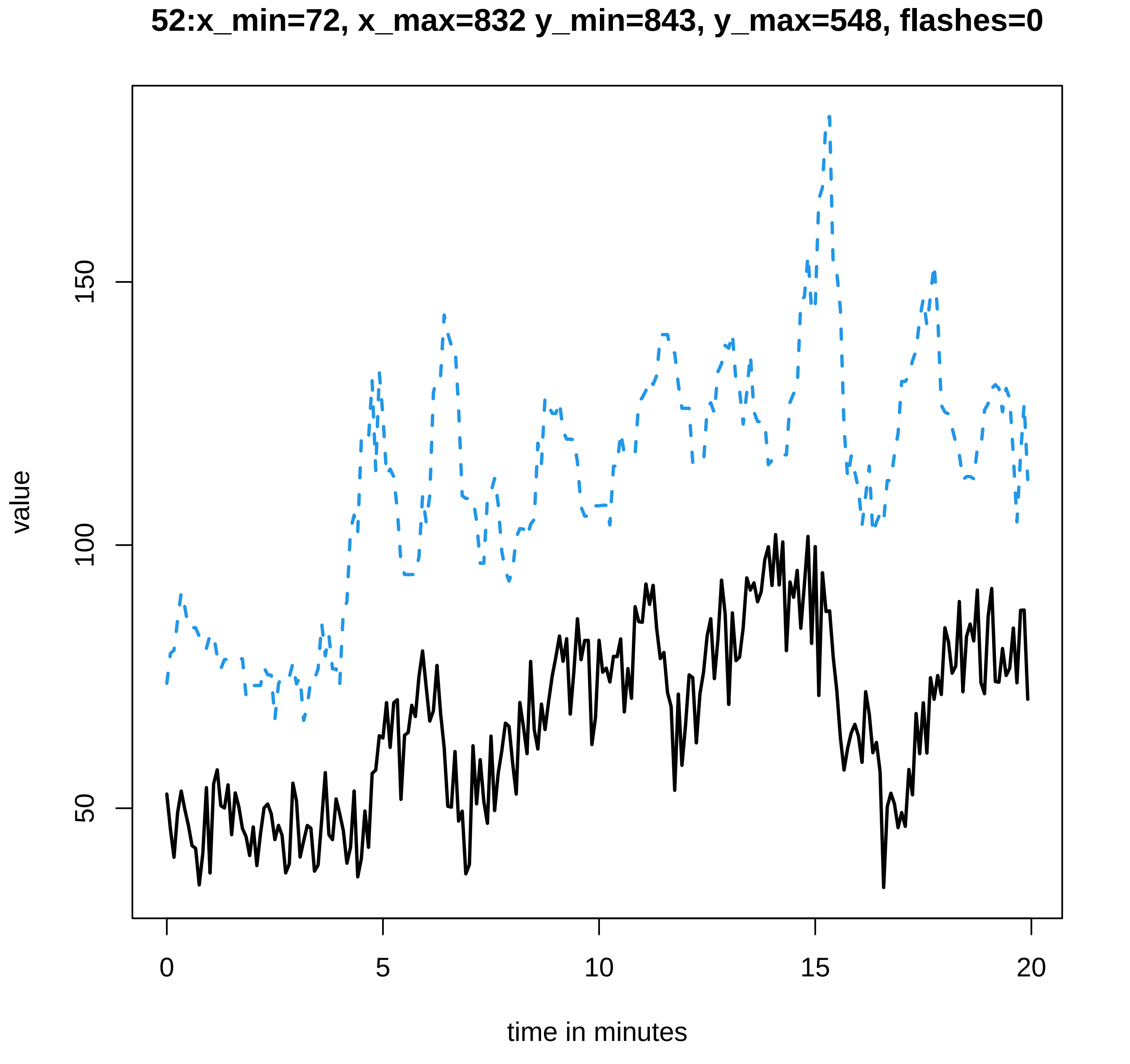

52:x_min=72, x_max=832 y_min=843, y_max=548, flashes=0
value
150
100
50
0
5
10
15
20
time in minutes

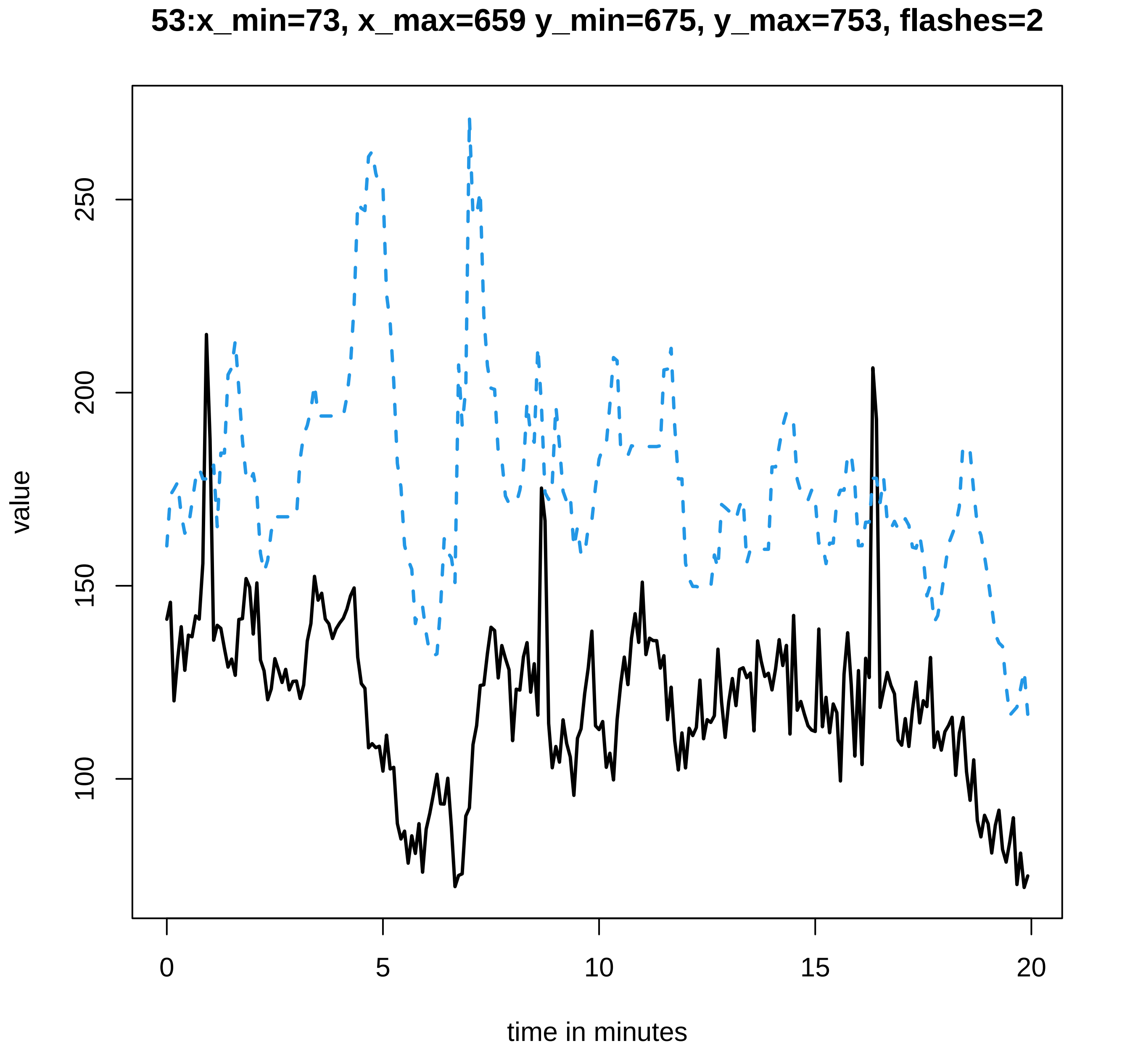

53:x_min=73, x_max=659 y_min=675, y_max=753, flashes=2
value
100
150
200
250
0
5
10
15
20
time in minutes

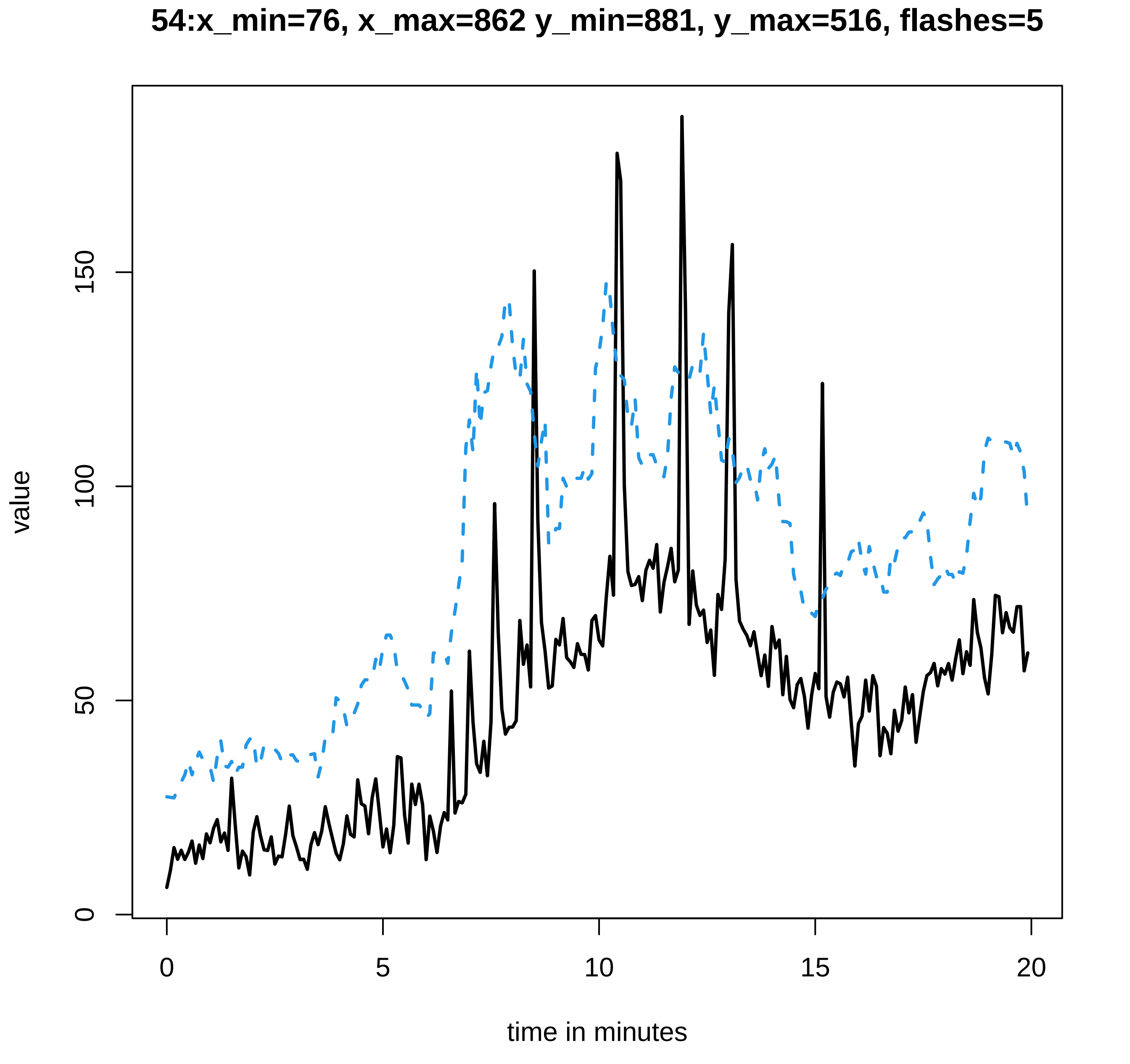

54:x_min=76, x_max=862 y_min=881, y_max=516, flashes=5
value
0
50
100
150
0
5
10
15
20
time in minutes

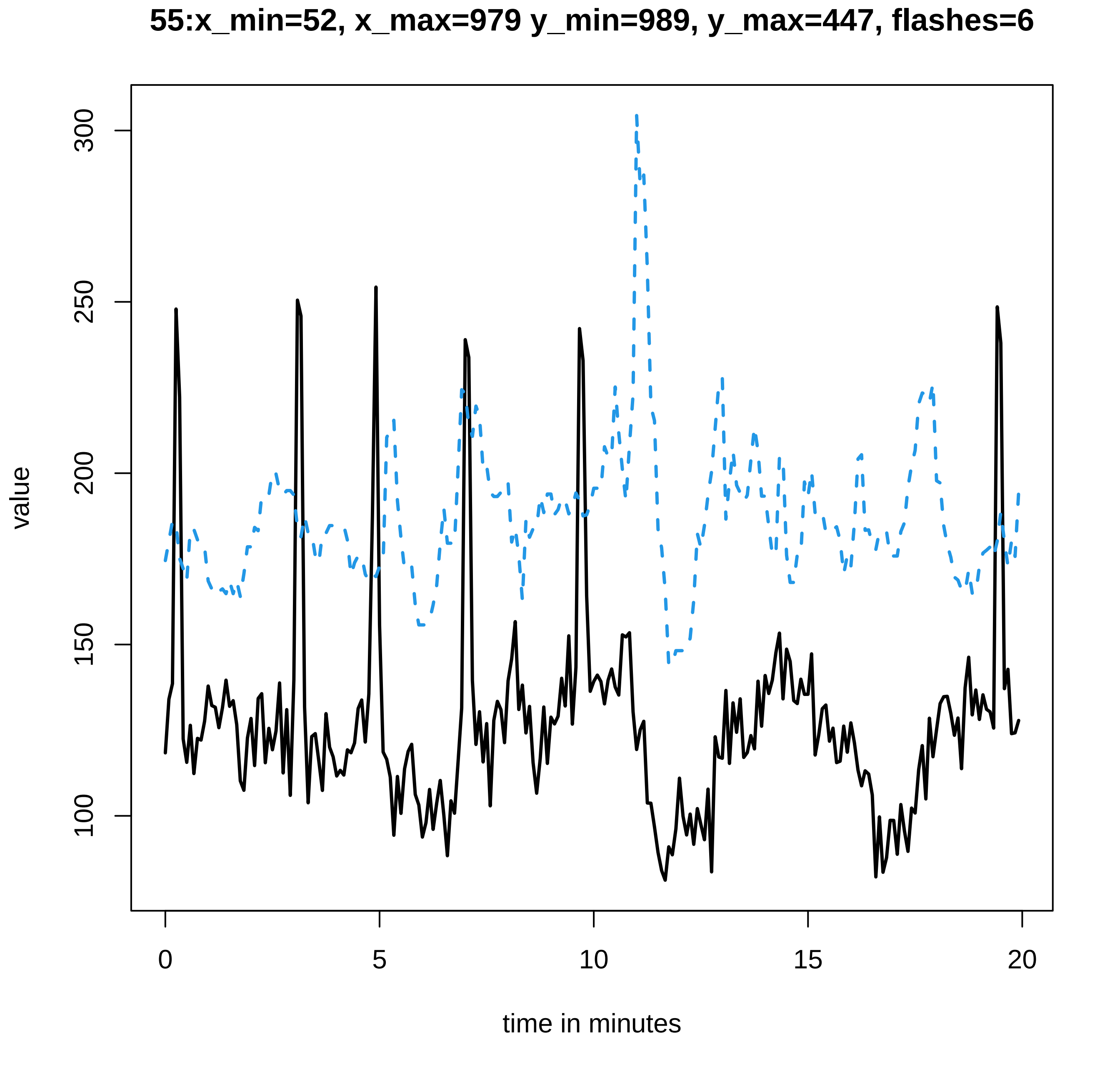
55:x_min=52, x_max=979 y_min=989, y_max=447, flashes=6
value
300
250
200
150
100
0
5
10
15
20
time in minutes

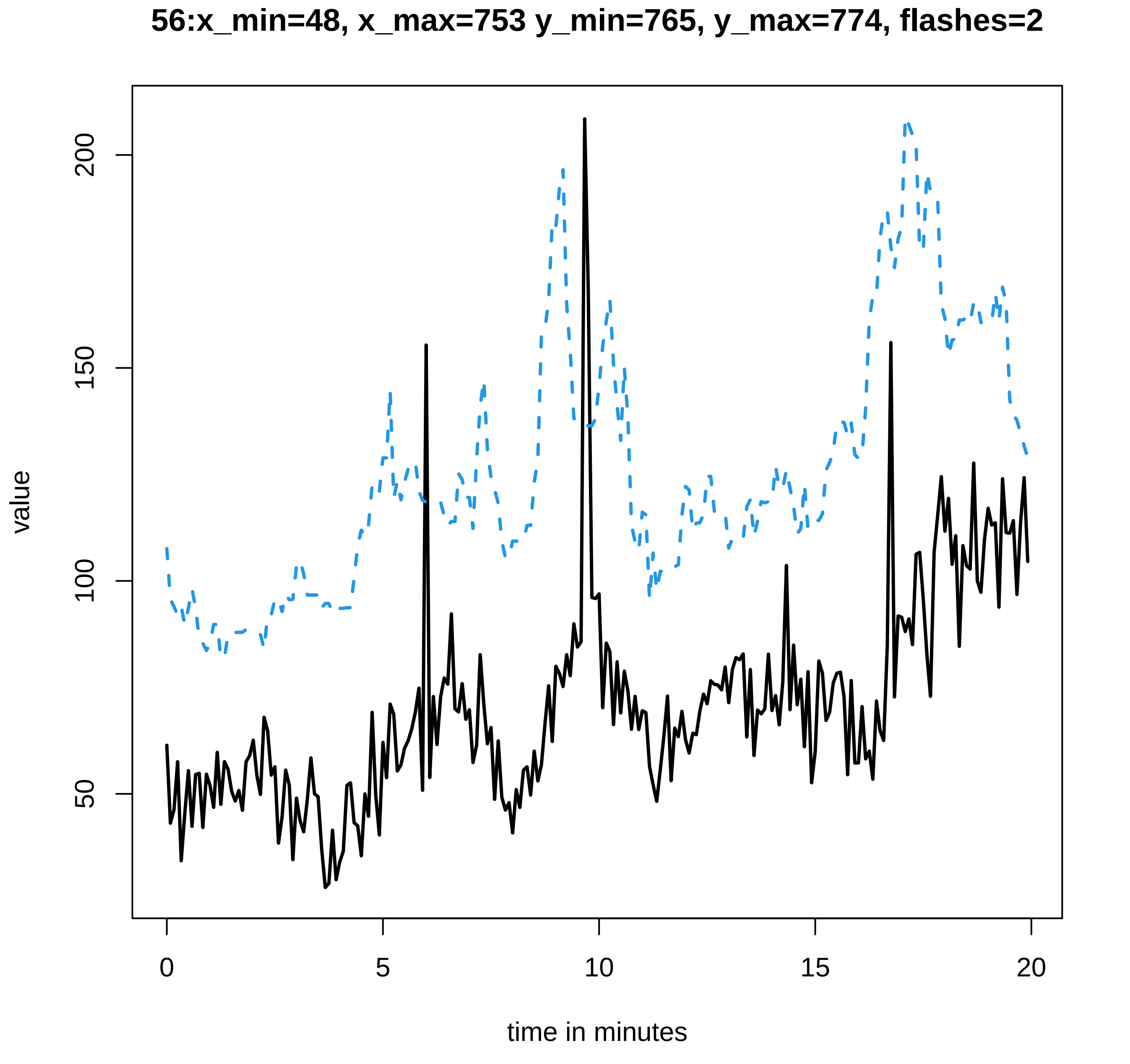
56:x_min=48, x_max=753 y_min=765, y_max=774, flashes=2
value
200
150
100
50
0
5
10
15
20
time in minutes

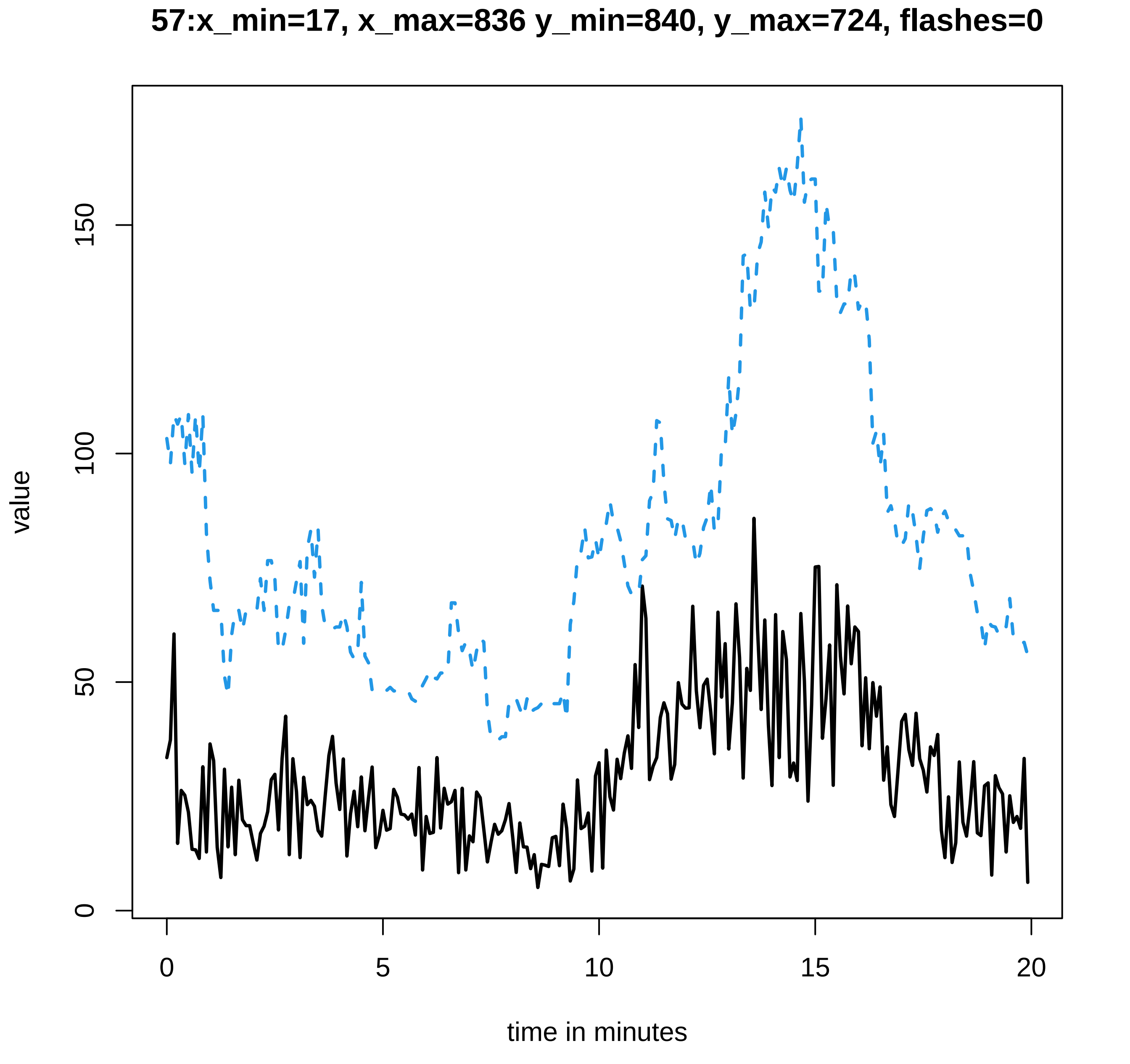

57:x_min=17, x_max=836 y_min=840, y_max=724, flashes=0
value
0
50
100
150
0
5
10
15
20
time in minutes

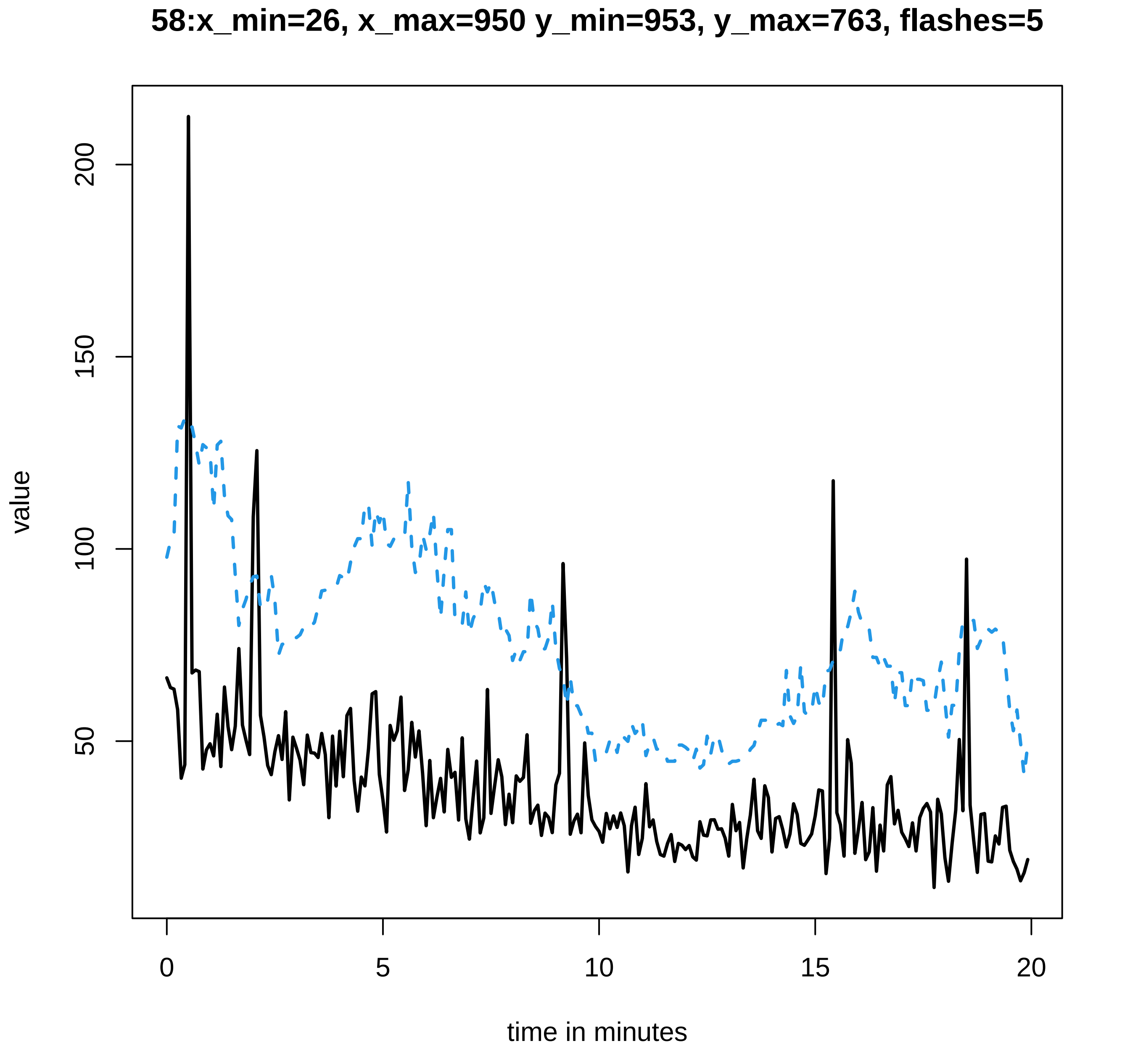
58:x_min=26, x_max=950 y_min=953, y_max=763, flashes=5
value
200
150
100
50
0
5
10
15
20
time in minutes
