## Supplementary data Calcium spiking for "LysM Receptor Proteins are Required for Ectomycorrhizal Symbiosis in Poplar": profiles-spiking-3.pdf

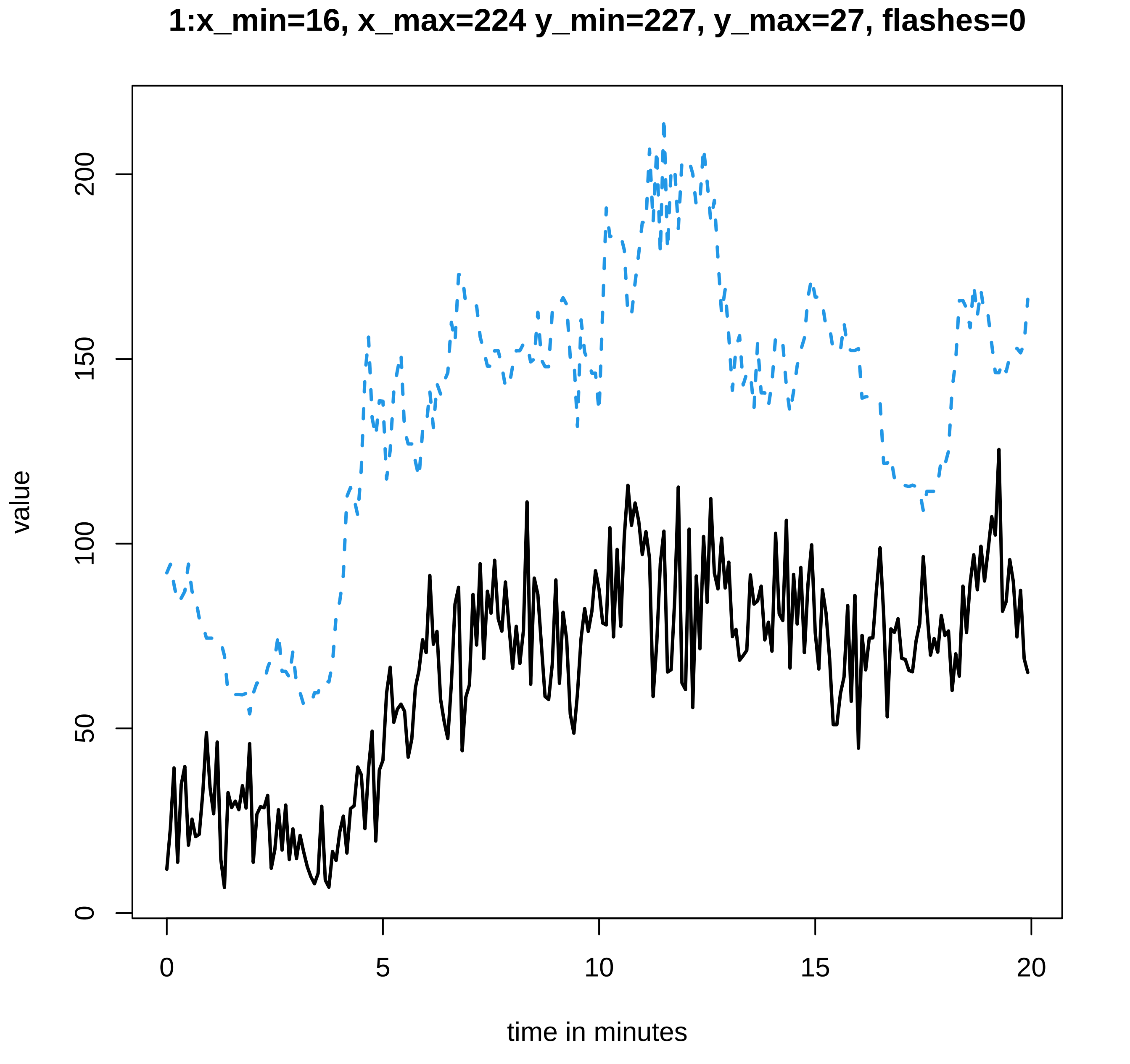

1:x_min=16, x_max=224 y_min=227, y_max=27, flashes=0
value
0
50
100
150
200
0
5
10
15
20
time in minutes

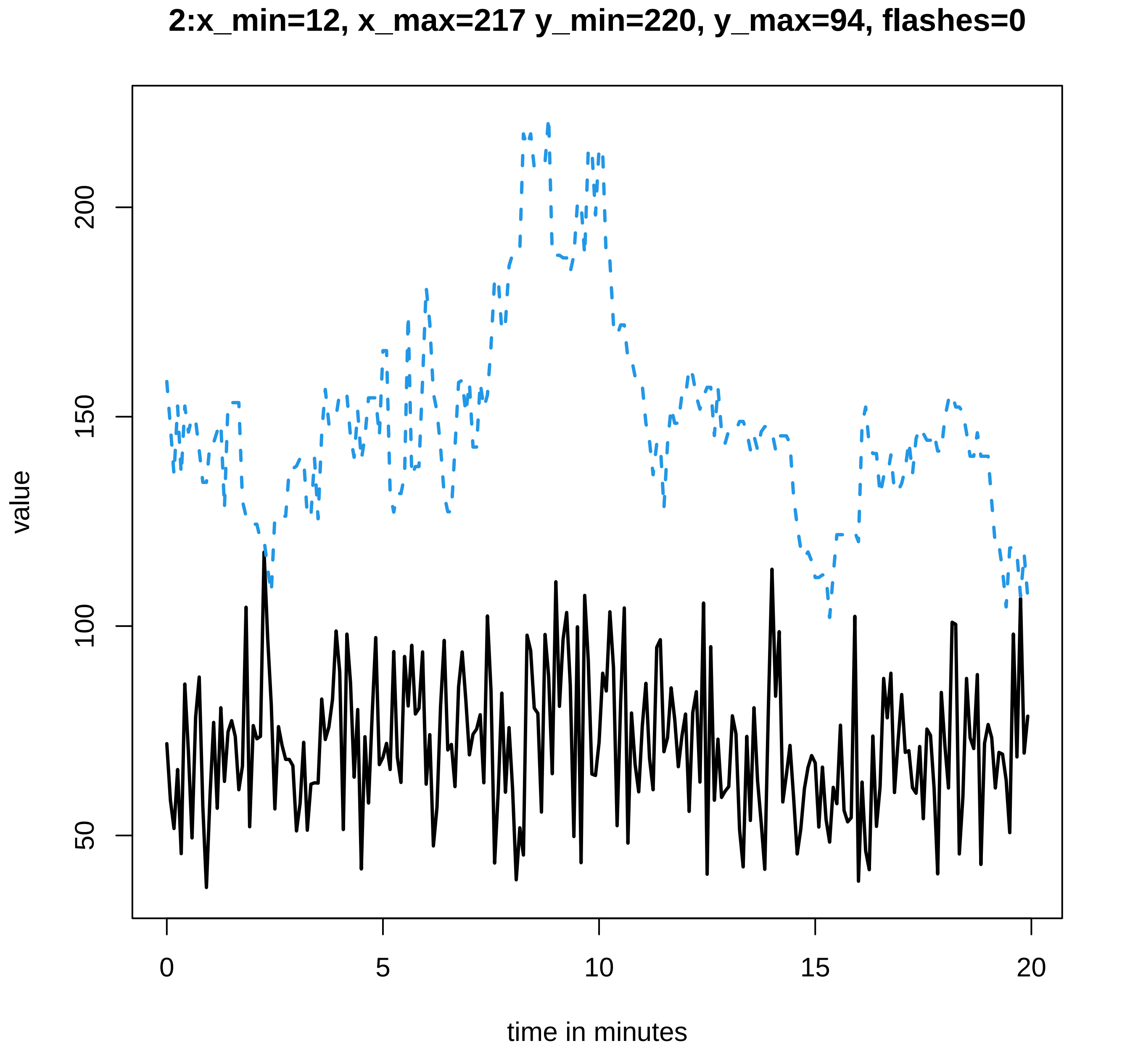

2:x_min=12, x_max=217 y_min=220, y_max=94, flashes=0
value
200
150
100
50
0
5
10
15
20
time in minutes

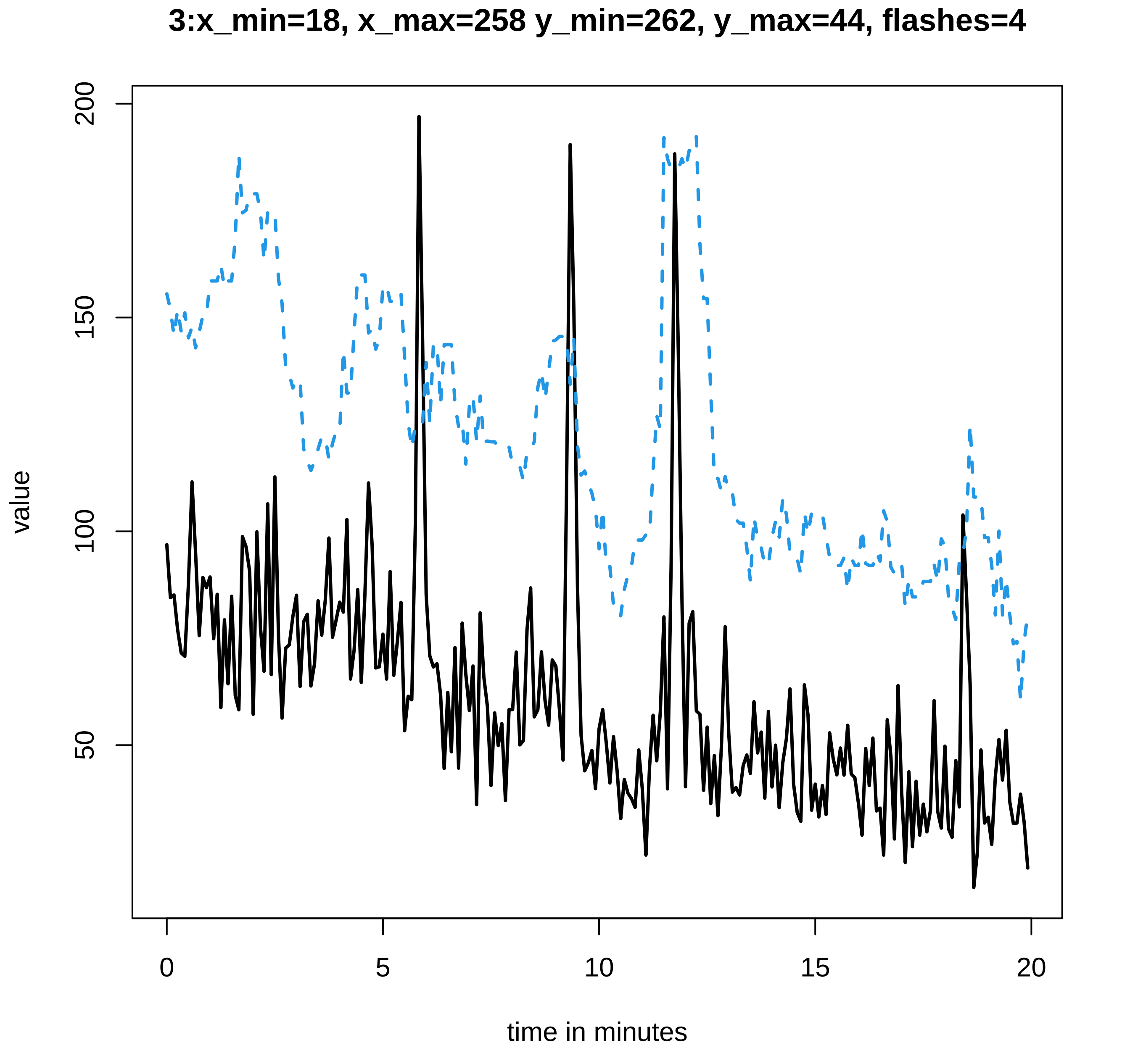
3:x_min=18, x_max=258 y_min=262, y_max=44, flashes=4
value
200
150
100
50
0
5
10
15
20
time in minutes

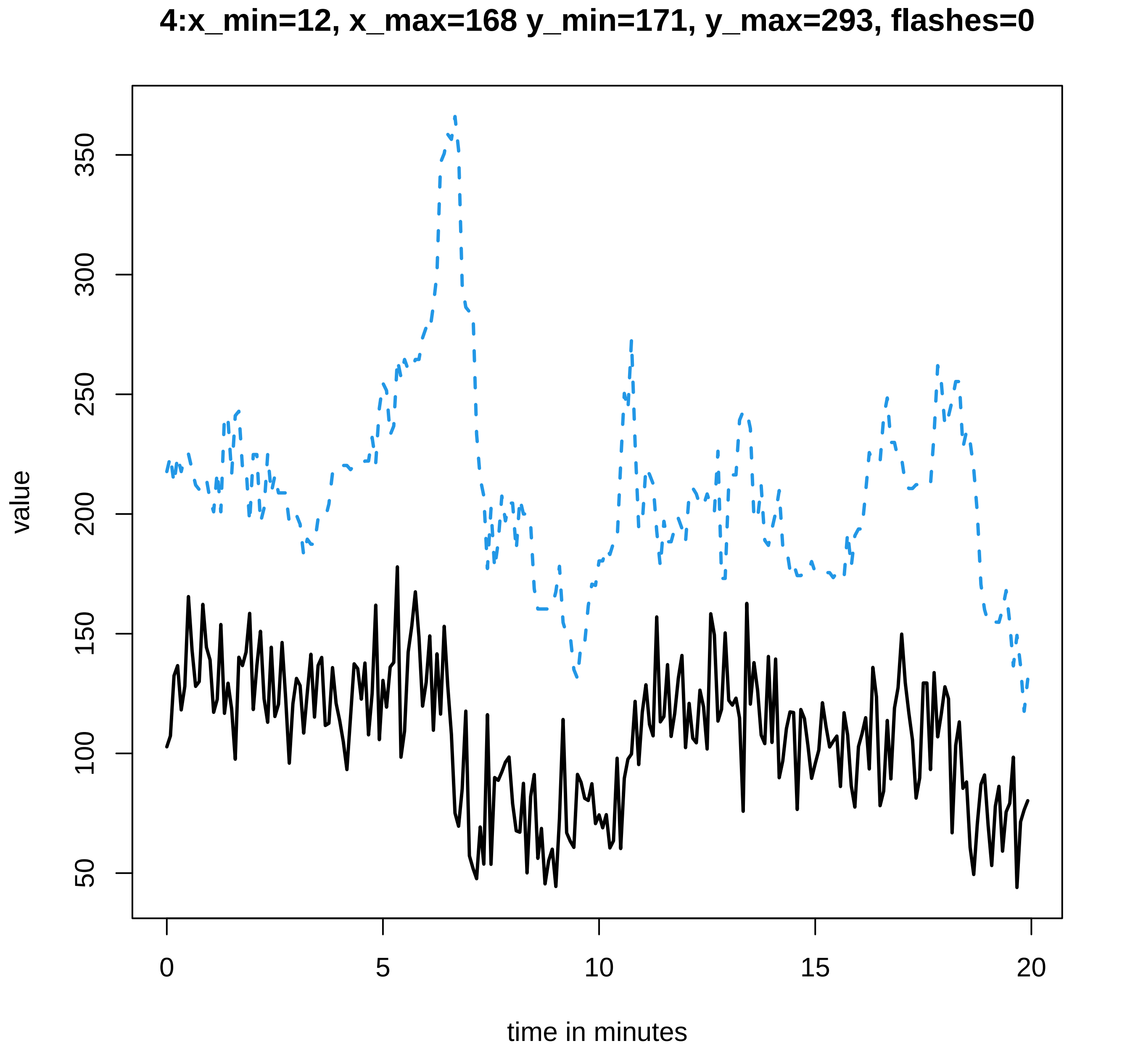
4:x_min=12, x_max=168 y_min=171, y_max=293, flashes=0
value
50
100
150
200
250
300
350
0
5
10
15
20
time in minutes

**5:x_min=32, x_max=128 y_min=133, y_max=376, flashes=11**

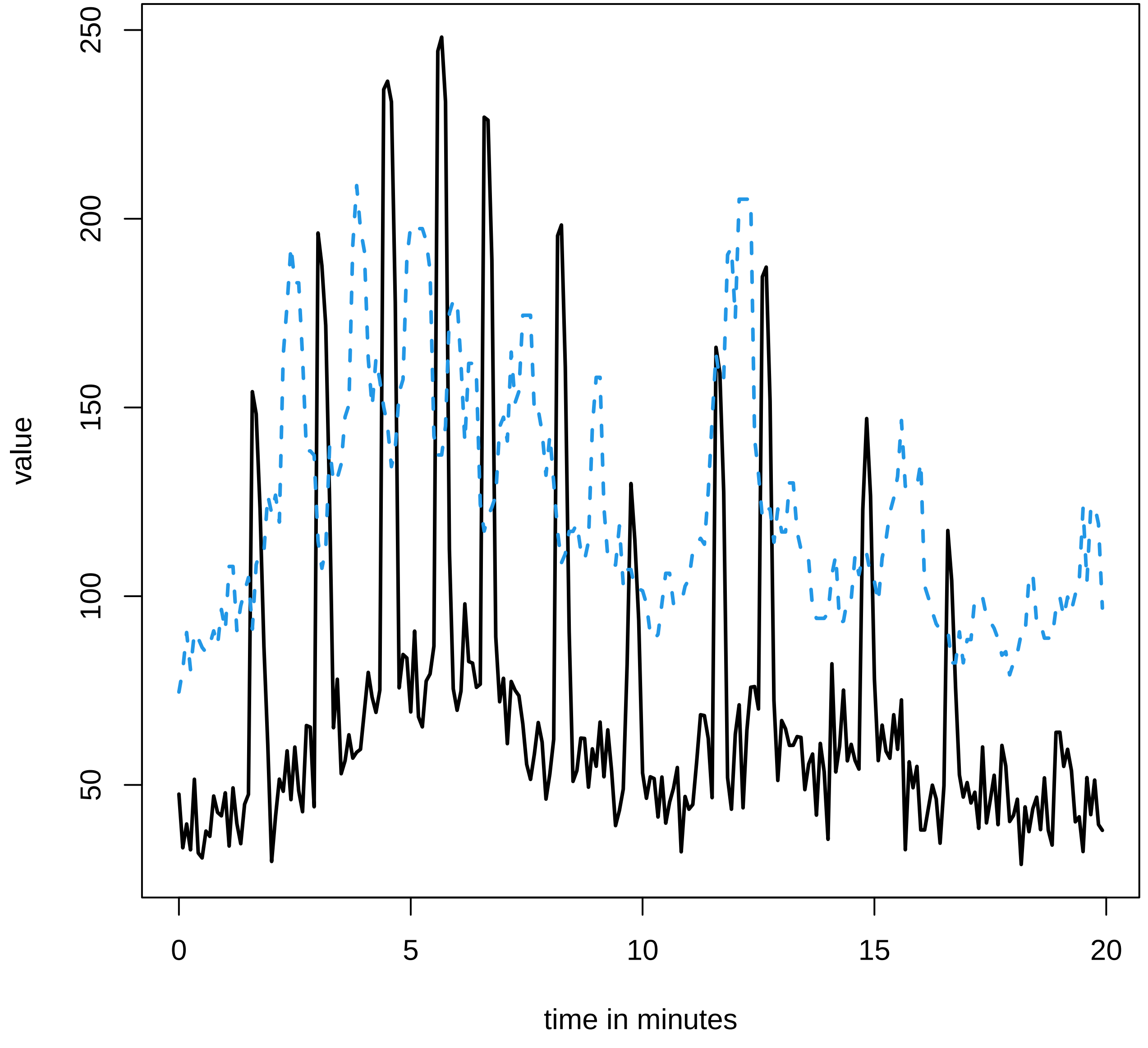

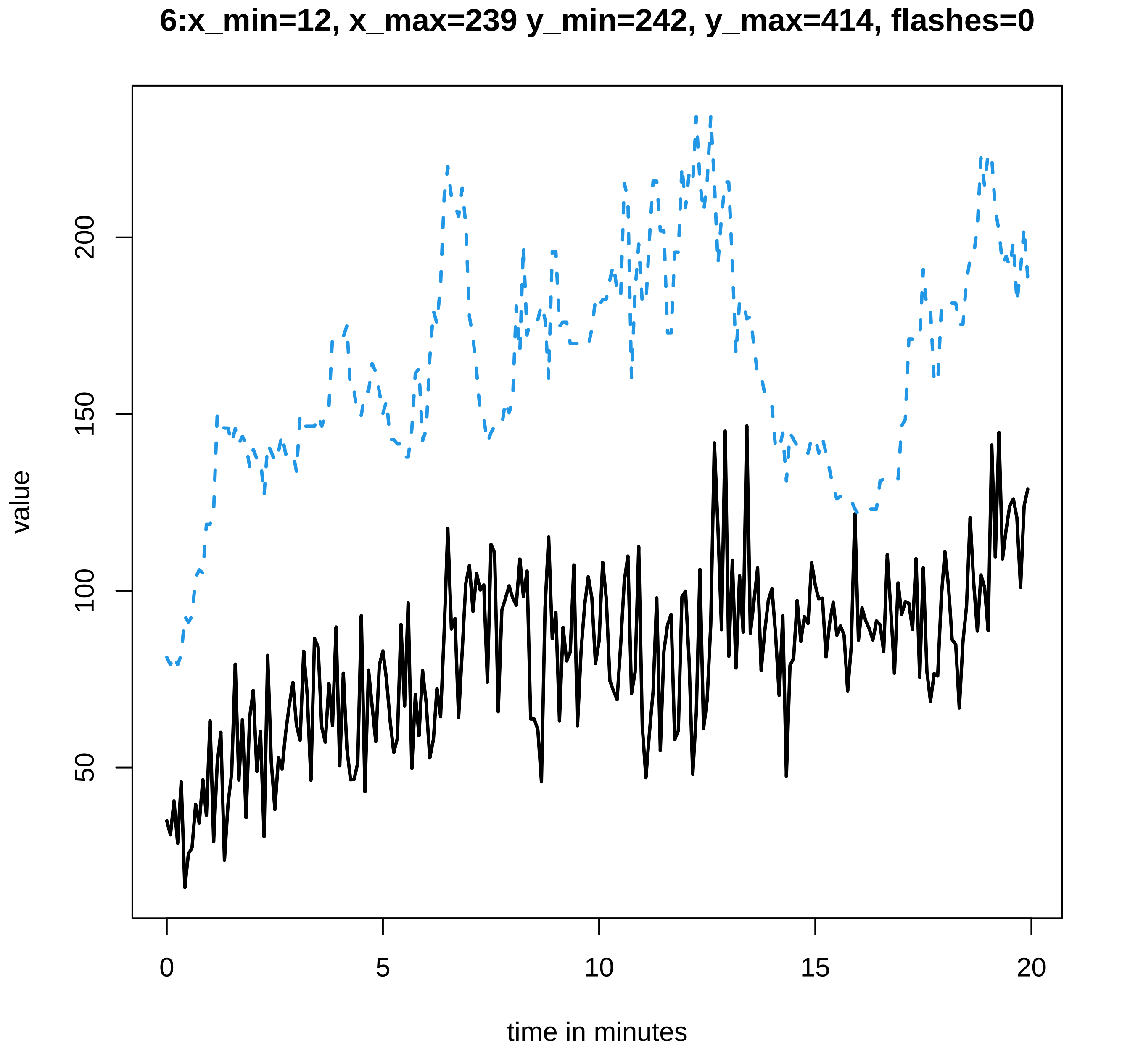

6:x_min=12, x_max=239 y_min=242, y_max=414, flashes=0
value
50
100
150
200
0
5
10
15
20
time in minutes

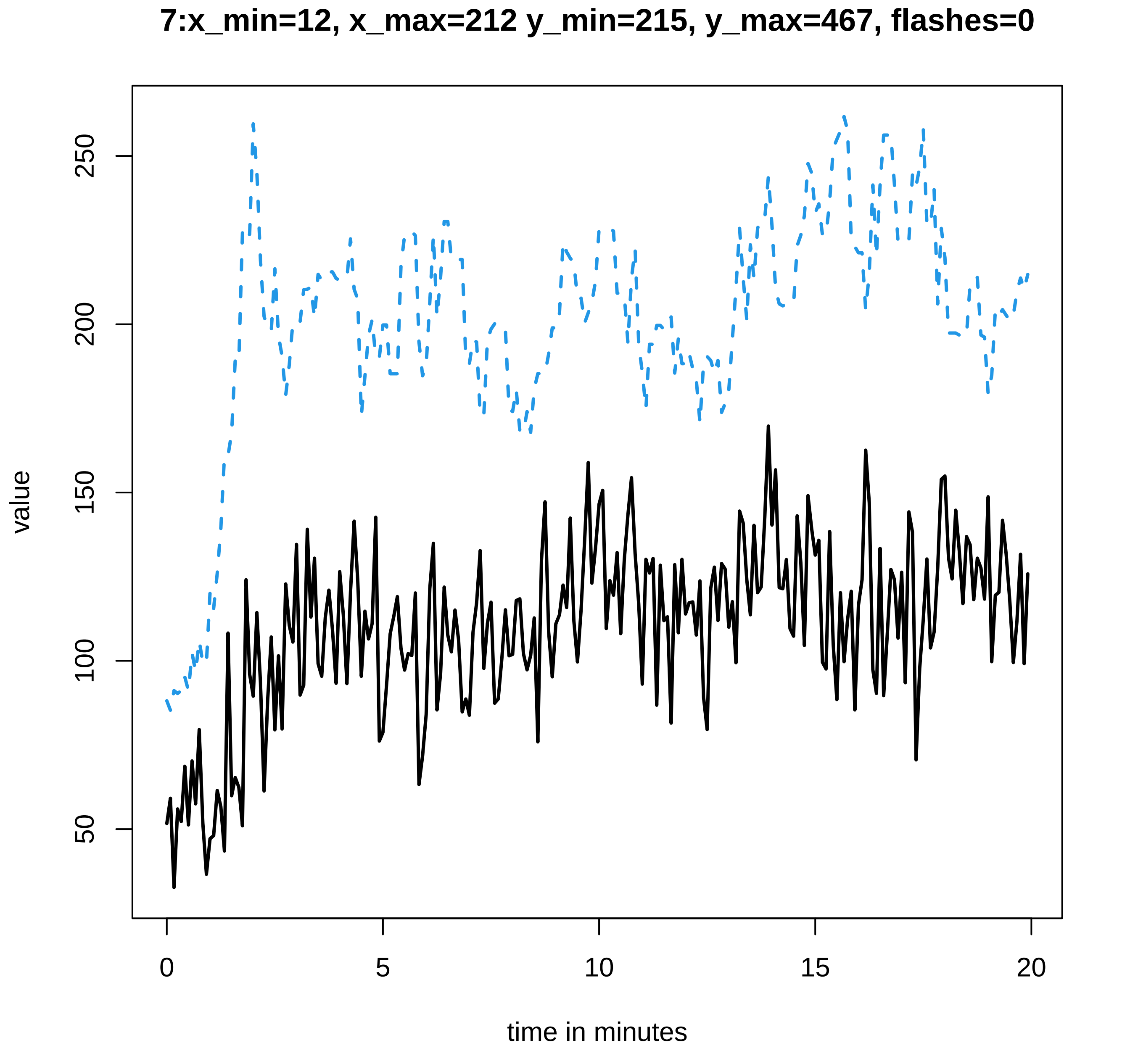

7:x_min=12, x_max=212 y_min=215, y_max=467, flashes=0
value
50
100
150
200
250
0
5
10
15
20
time in minutes

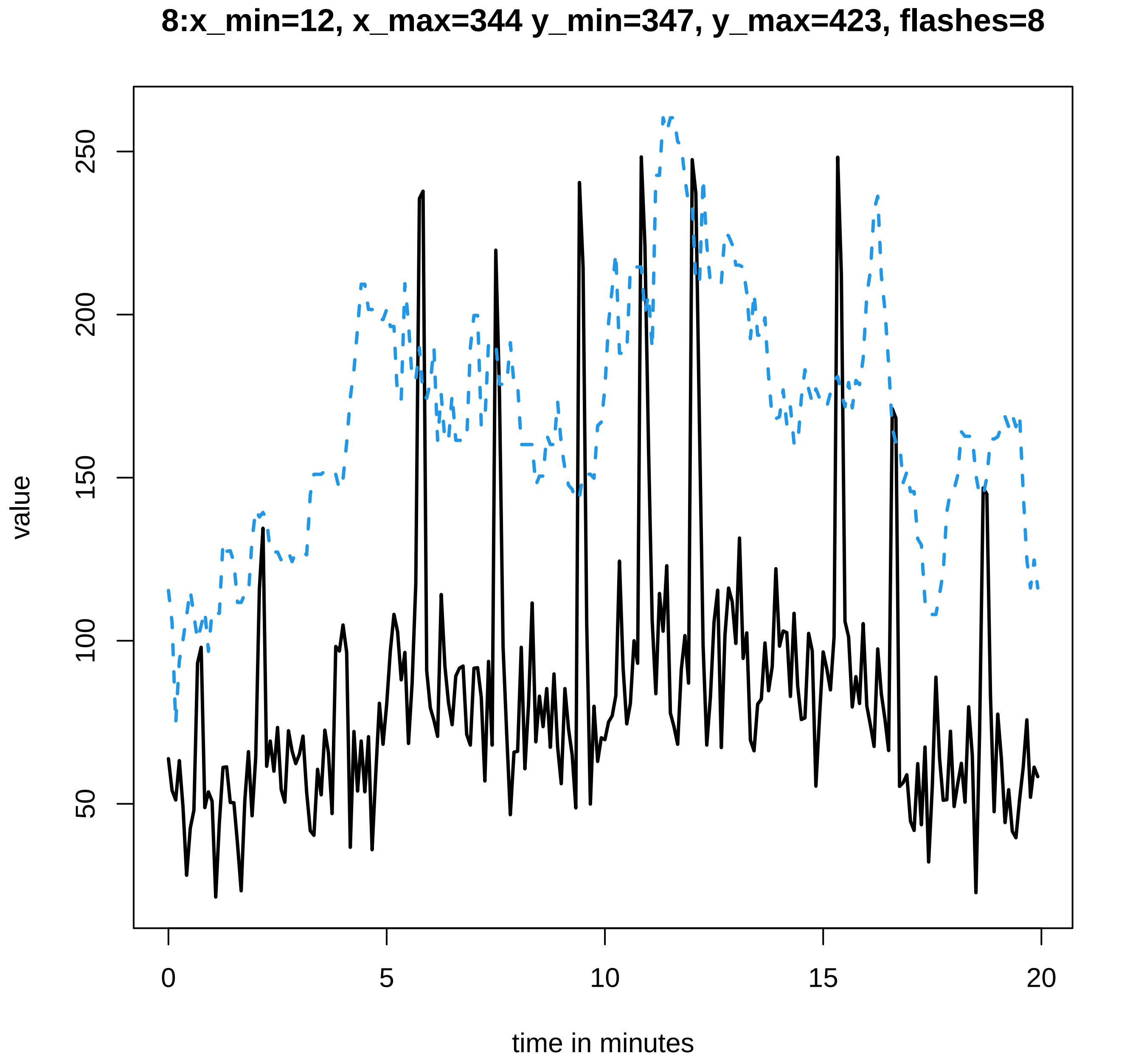

8:x_min=12, x_max=344 y_min=347, y_max=423, flashes=8
value
50
100
150
200
250
0
5
10
15
20
time in minutes

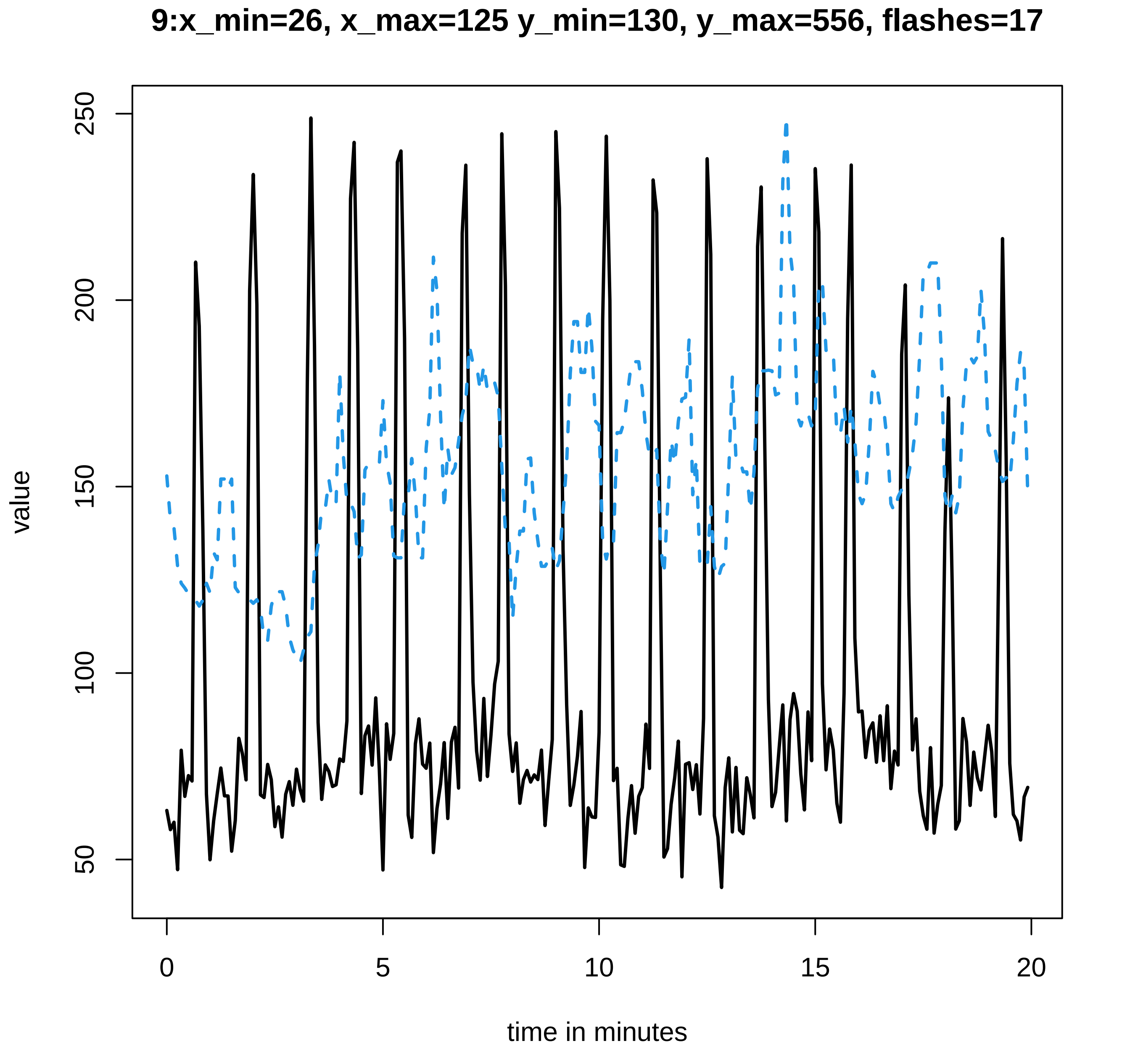
9:x_min=26, x_max=125 y_min=130, y_max=556, flashes=17
value
50
100
150
200
250
0
5
10
15
20
time in minutes

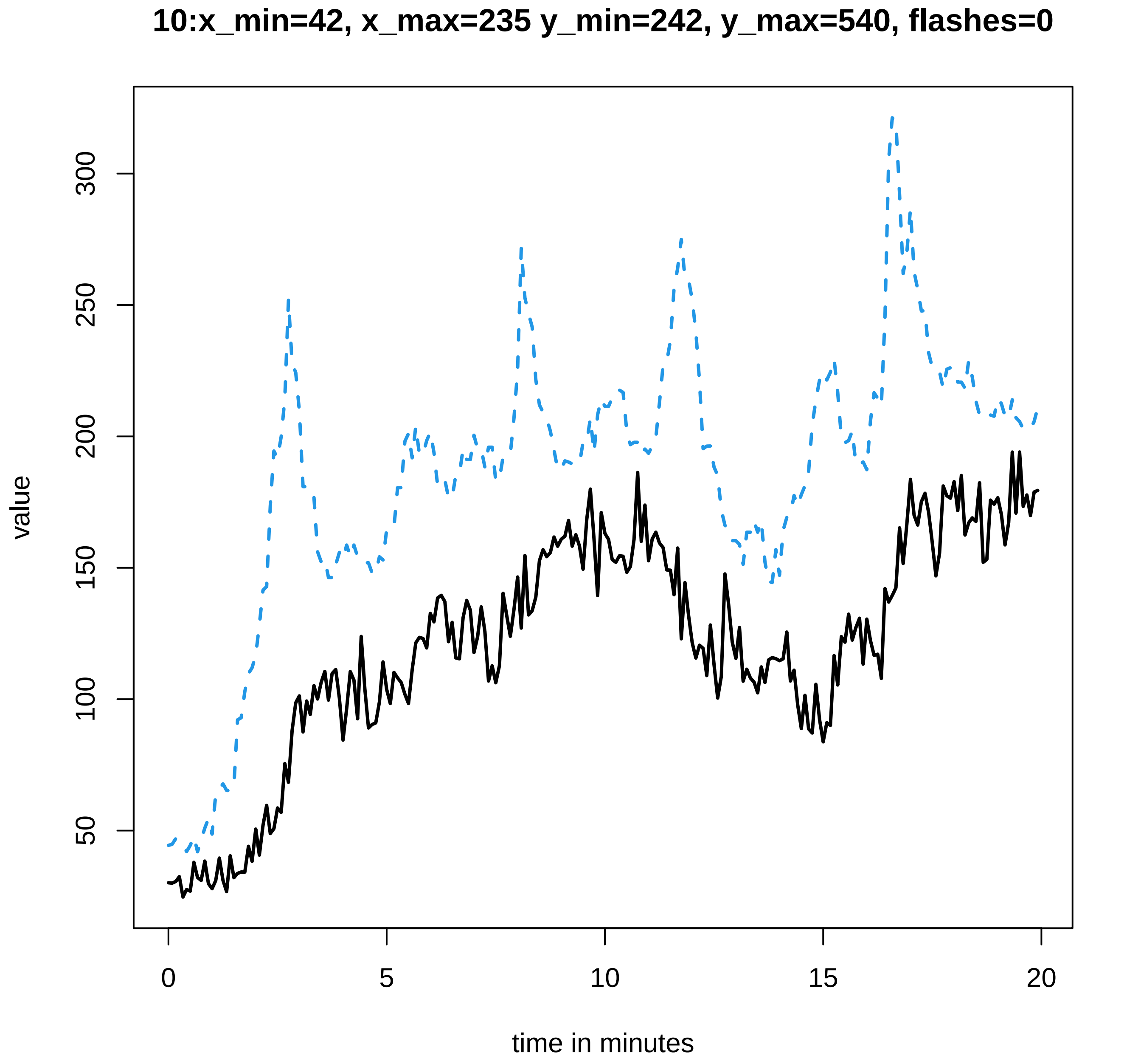

10:x_min=42, x_max=235 y_min=242, y_max=540, flashes=0
value
300
250
200
150
100
50
0
5
10
15
20
time in minutes

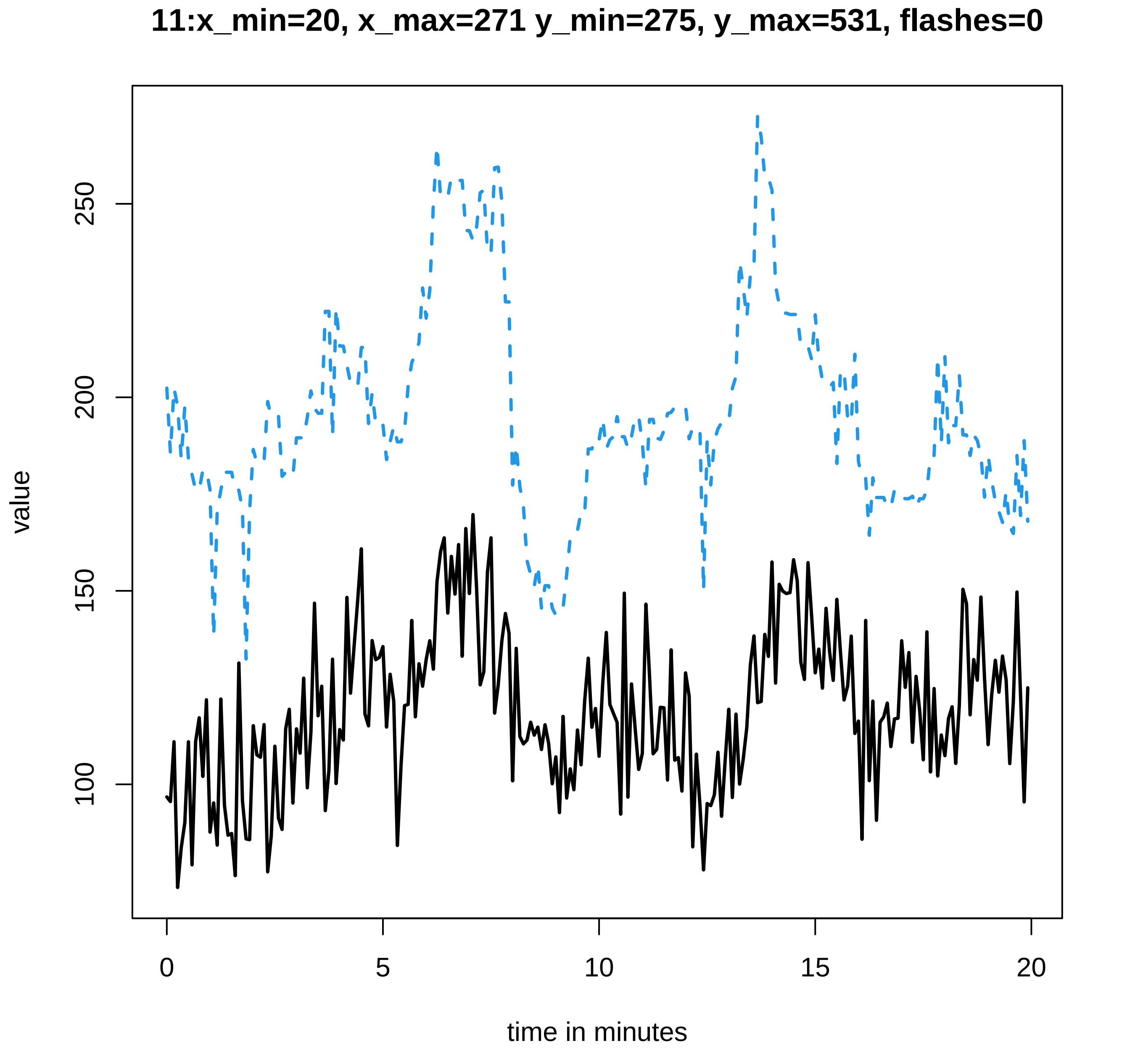

11:x_min=20, x_max=271 y_min=275, y_max=531, flashes=0
value
250
200
150
100
0
5
10
15
20
time in minutes

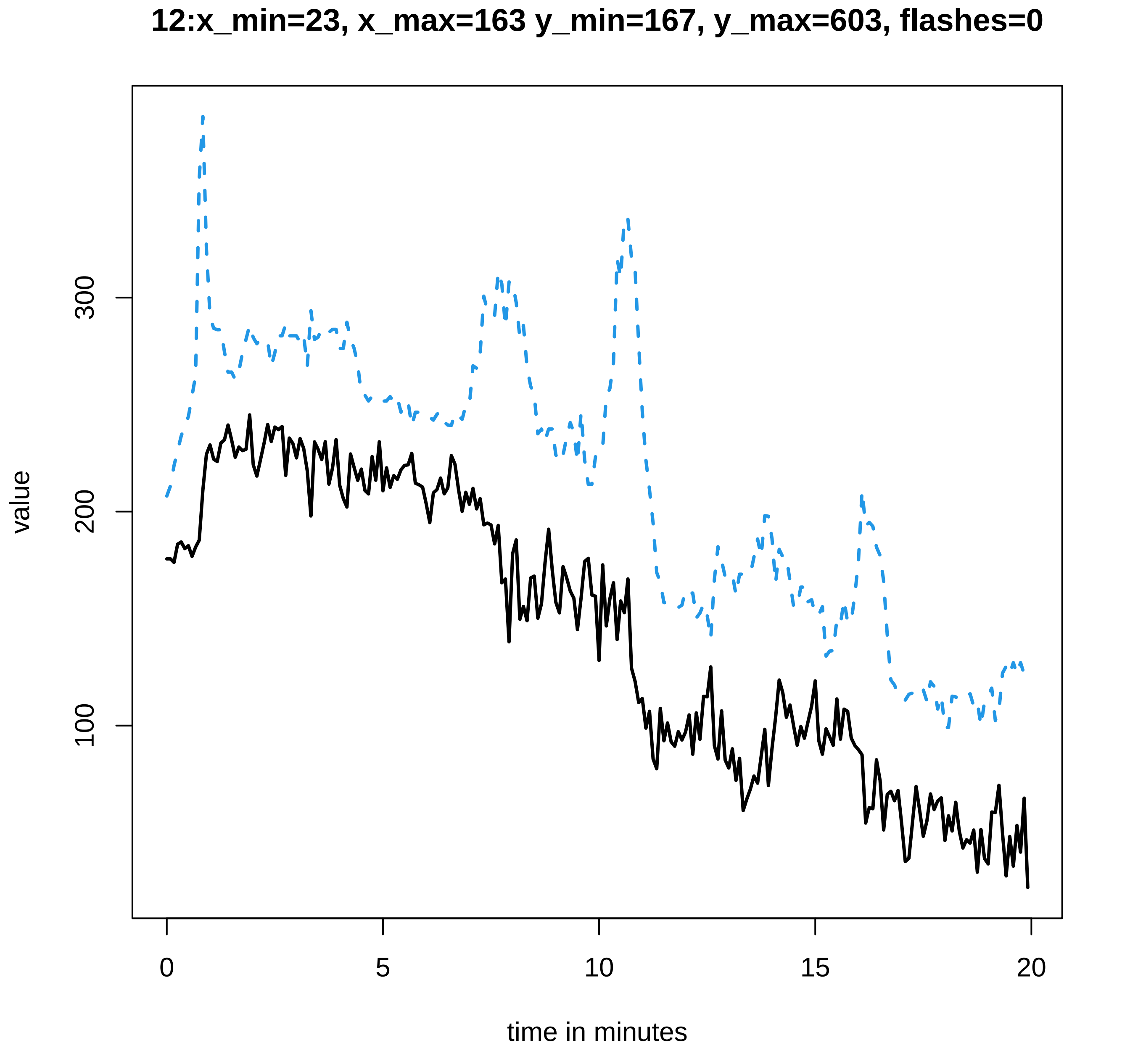

12:x_min=23, x_max=163 y_min=167, y_max=603, flashes=0
value
100
200
300
0
5
10
15
20
time in minutes

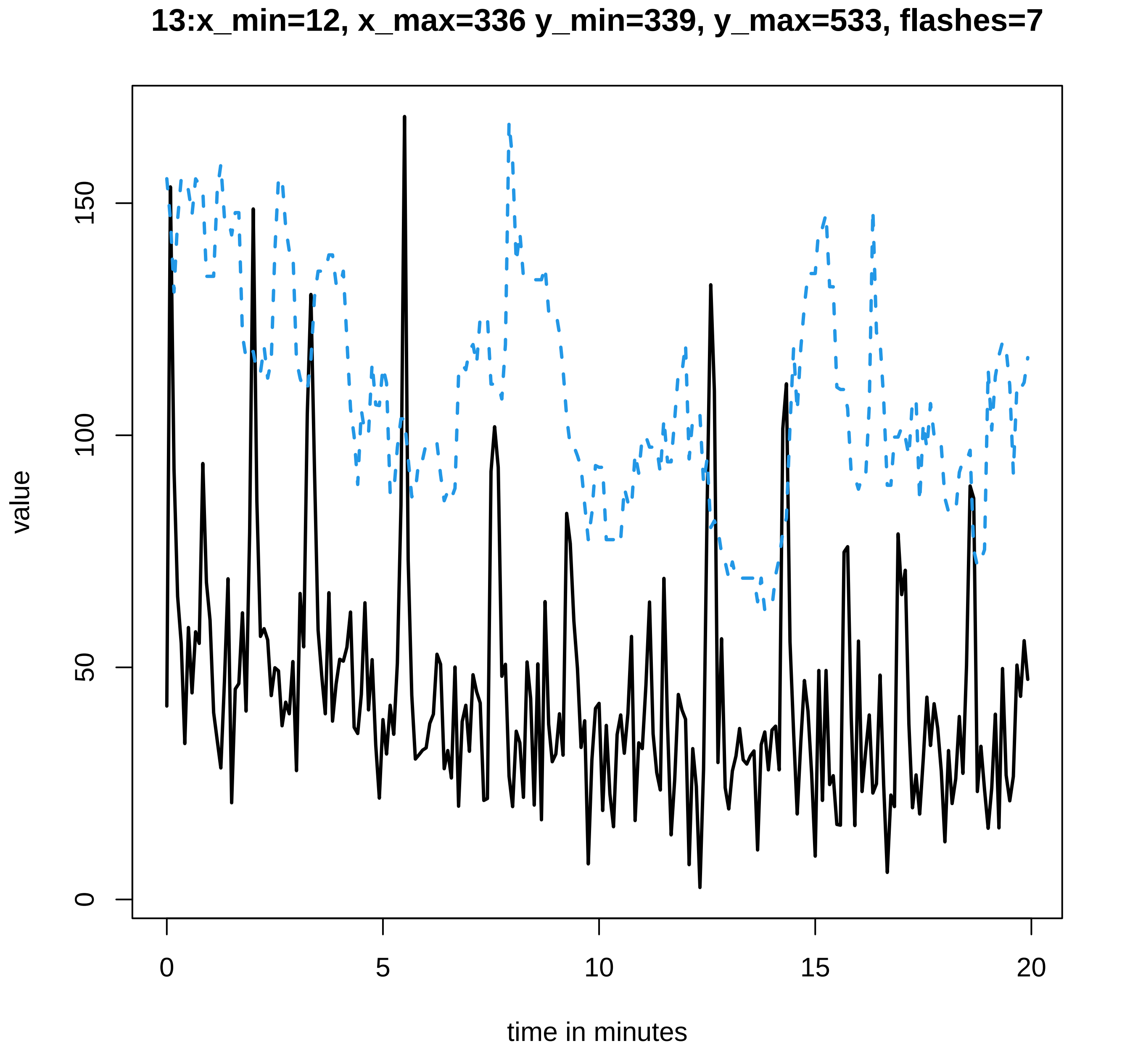
13:x_min=12, x_max=336 y_min=339, y_max=533, flashes=7
value
0
50
100
150
0
5
10
15
20
time in minutes

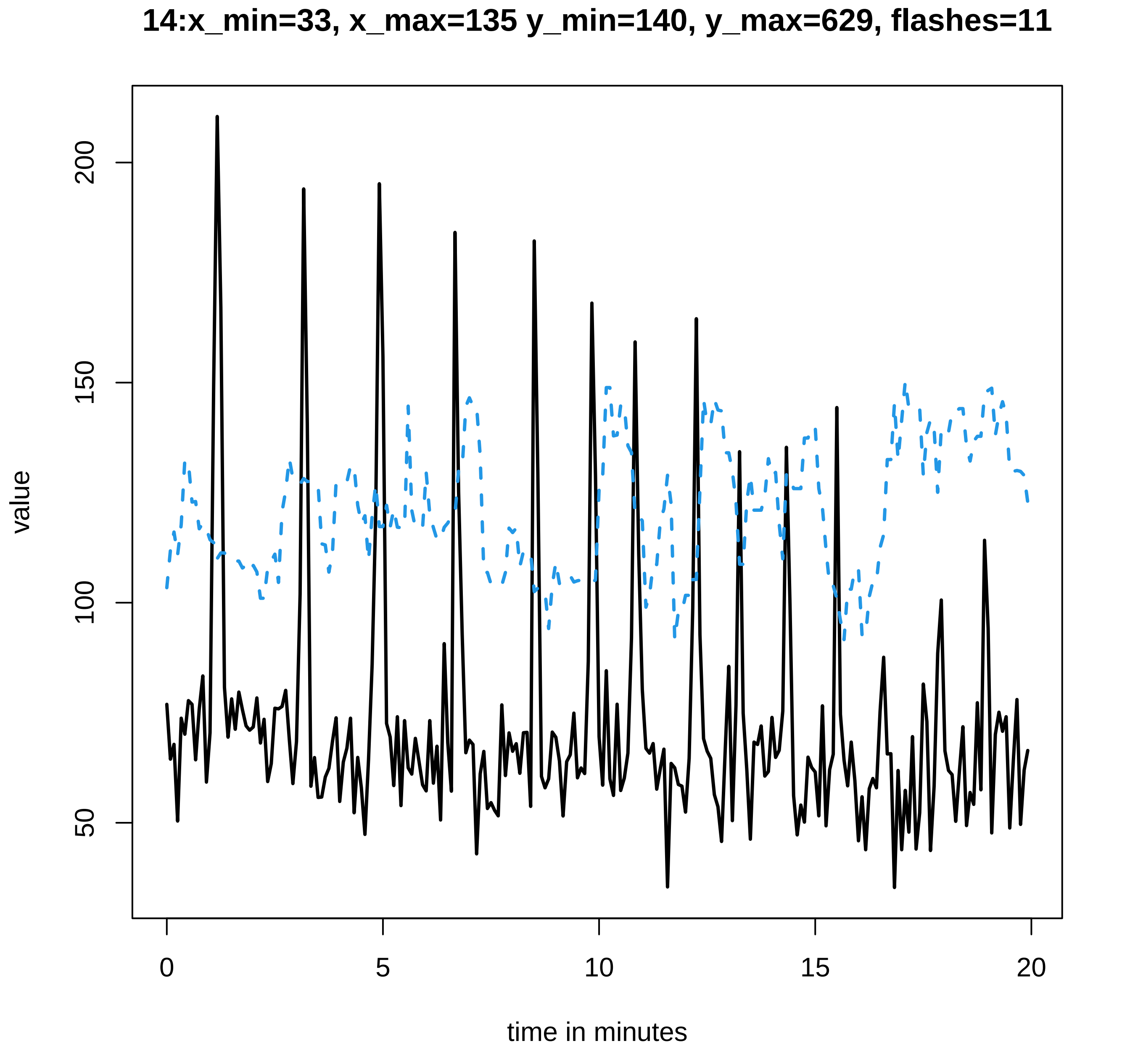
14:x_min=33, x_max=135 y_min=140, y_max=629, flashes=11
value
50
100
150
200
0
5
10
15
20
time in minutes

#### 15:x_min=20, x_max=444 y_min=449, y_max=472, flashes=2

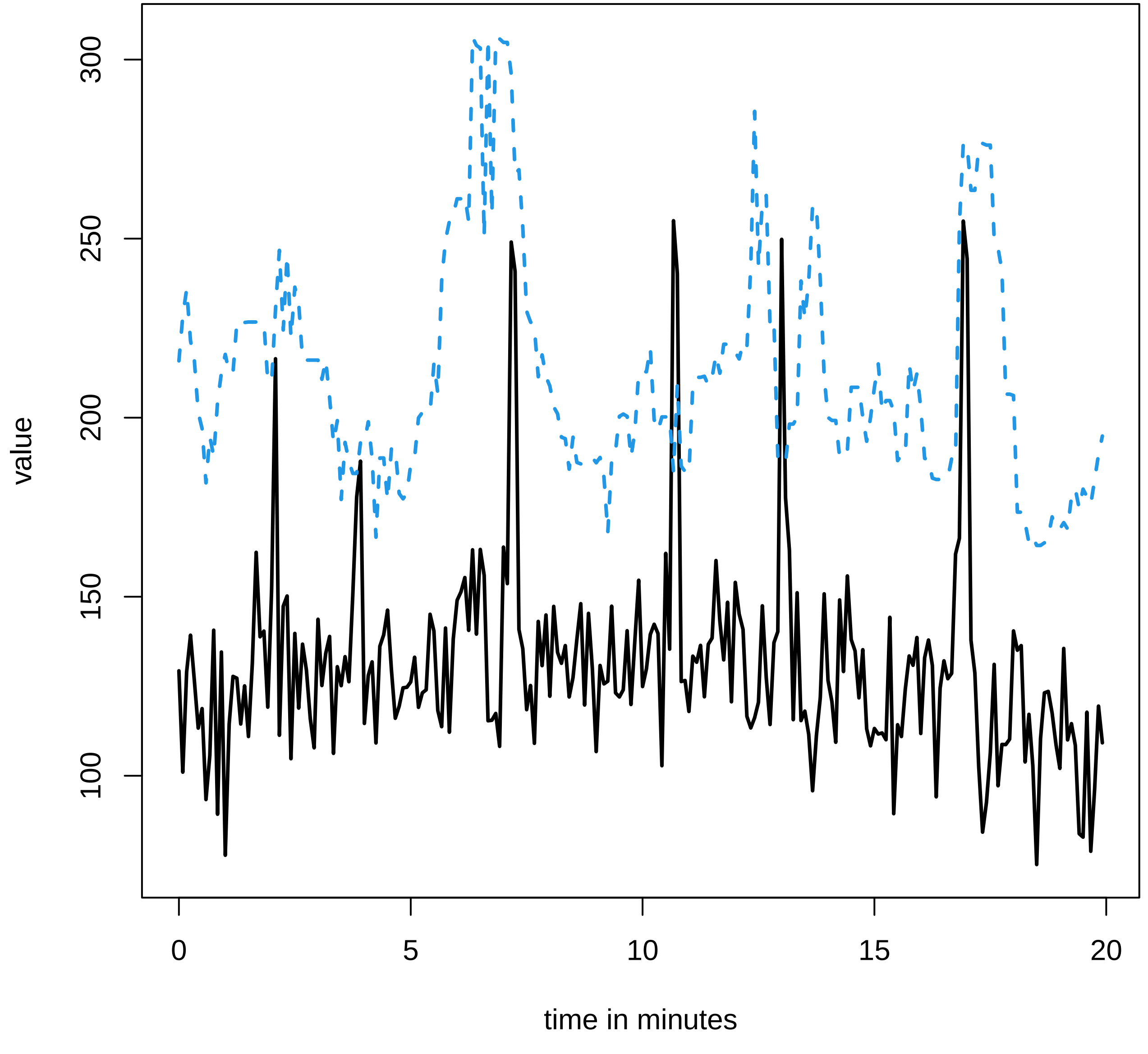

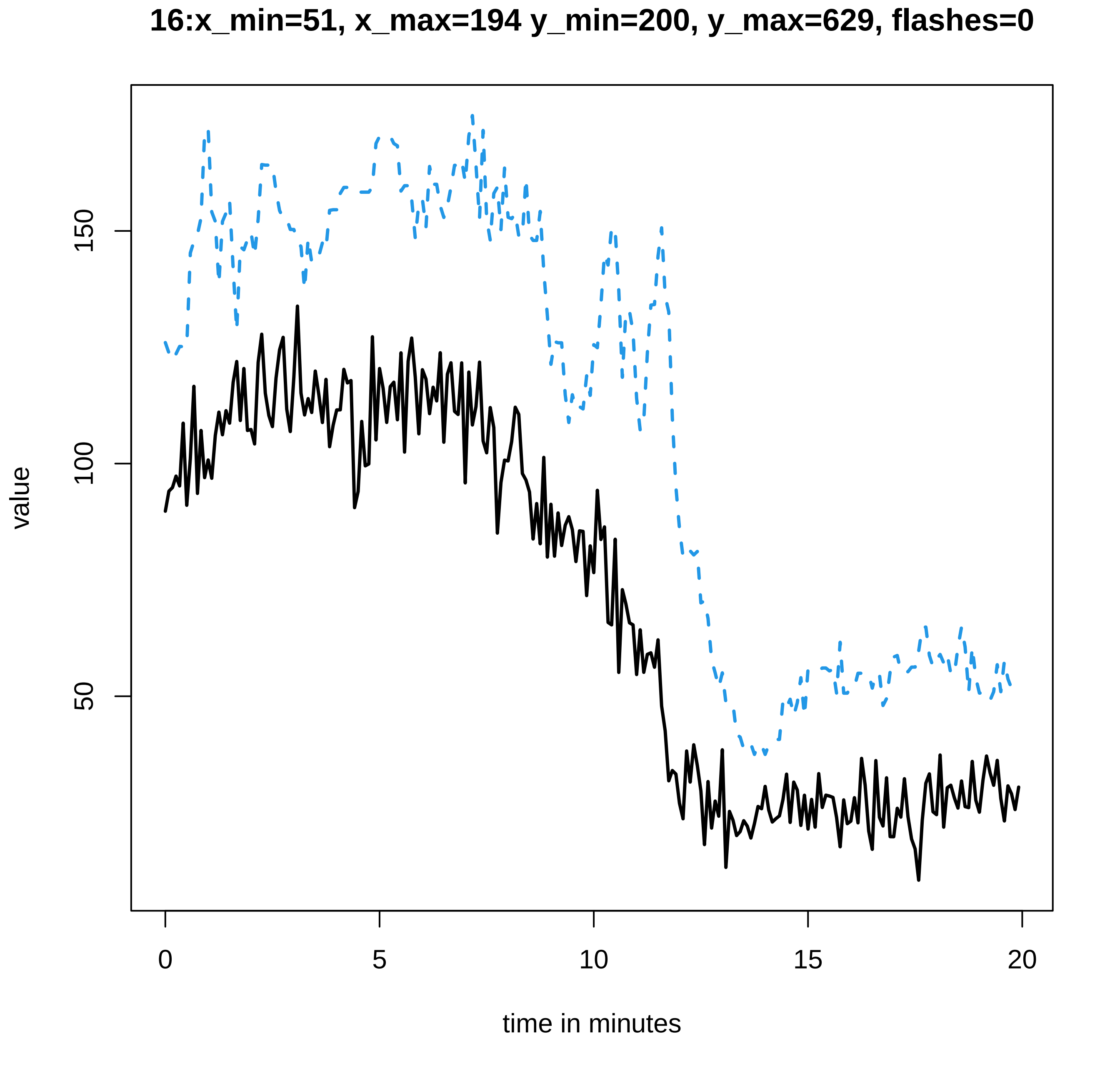

16:x_min=51, x_max=194 y_min=200, y_max=629, flashes=0
value
150
100
50
0
5
10
15
20
time in minutes

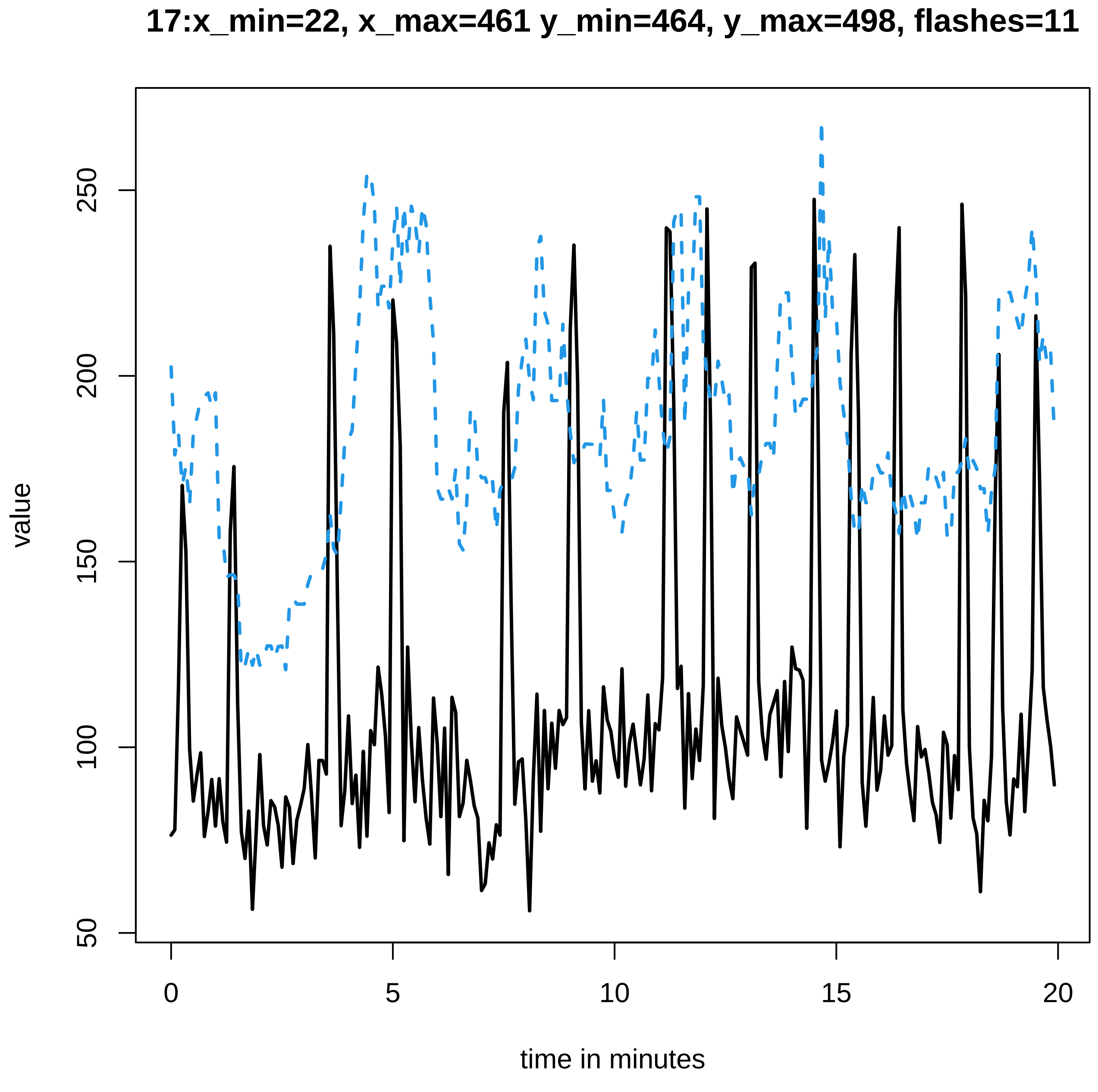
17:x_min=22, x_max=461 y_min=464, y_max=498, flashes=11
value
250
200
150
100
50
0
5
10
15
20
time in minutes

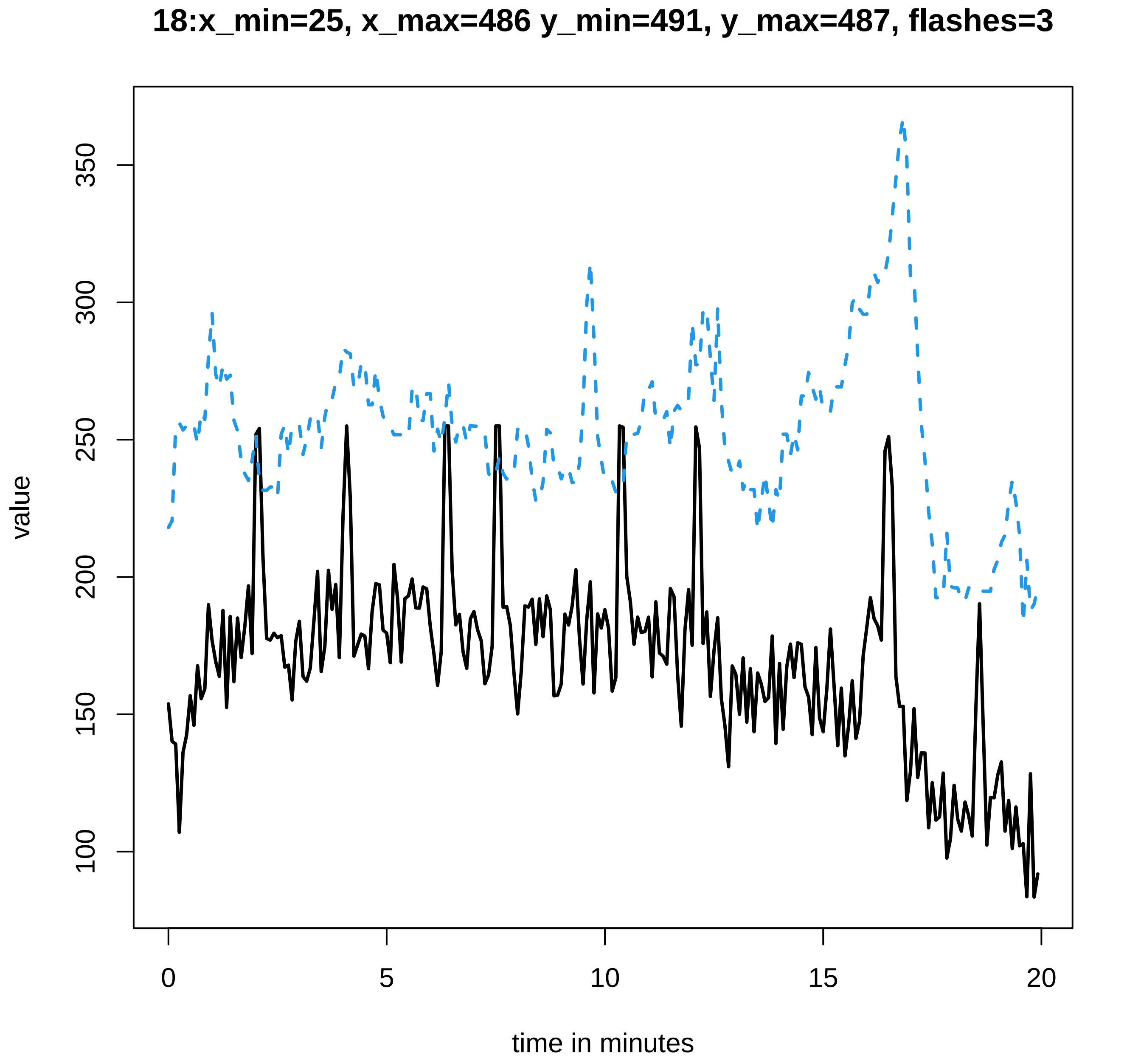
18:x_min=25, x_max=486 y_min=491, y_max=487, flashes=3
value
100
150
200
250
300
350
0
5
10
15
20
time in minutes

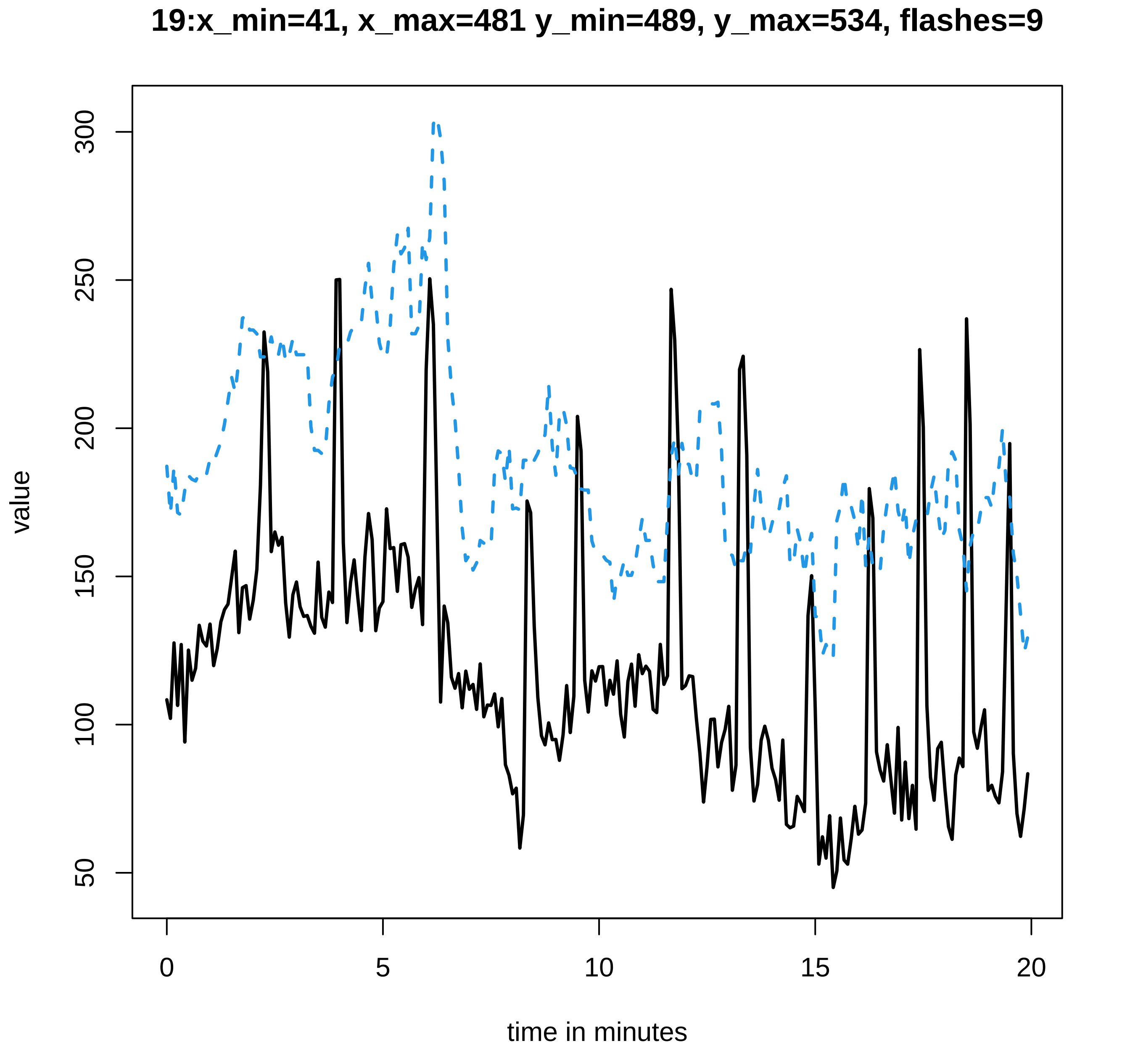
19:x_min=41, x_max=481 y_min=489, y_max=534, flashes=9
value
300
250
200
150
100
50
0
5
10
15
20
time in minutes

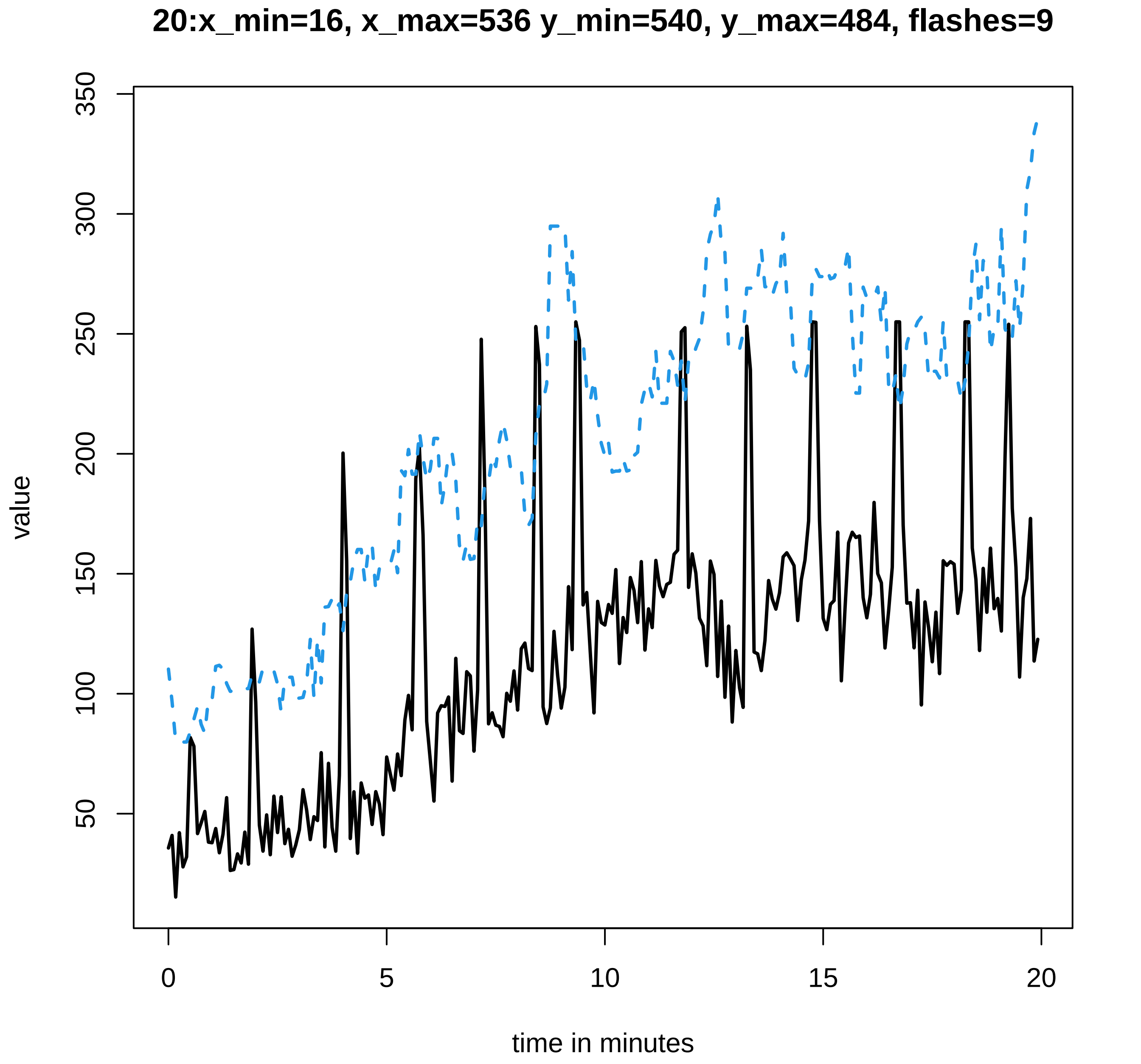

20:x_min=16, x_max=536 y_min=540, y_max=484, flashes=9
value
50
100
150
200
250
300
350
0
5
10
15
20
time in minutes

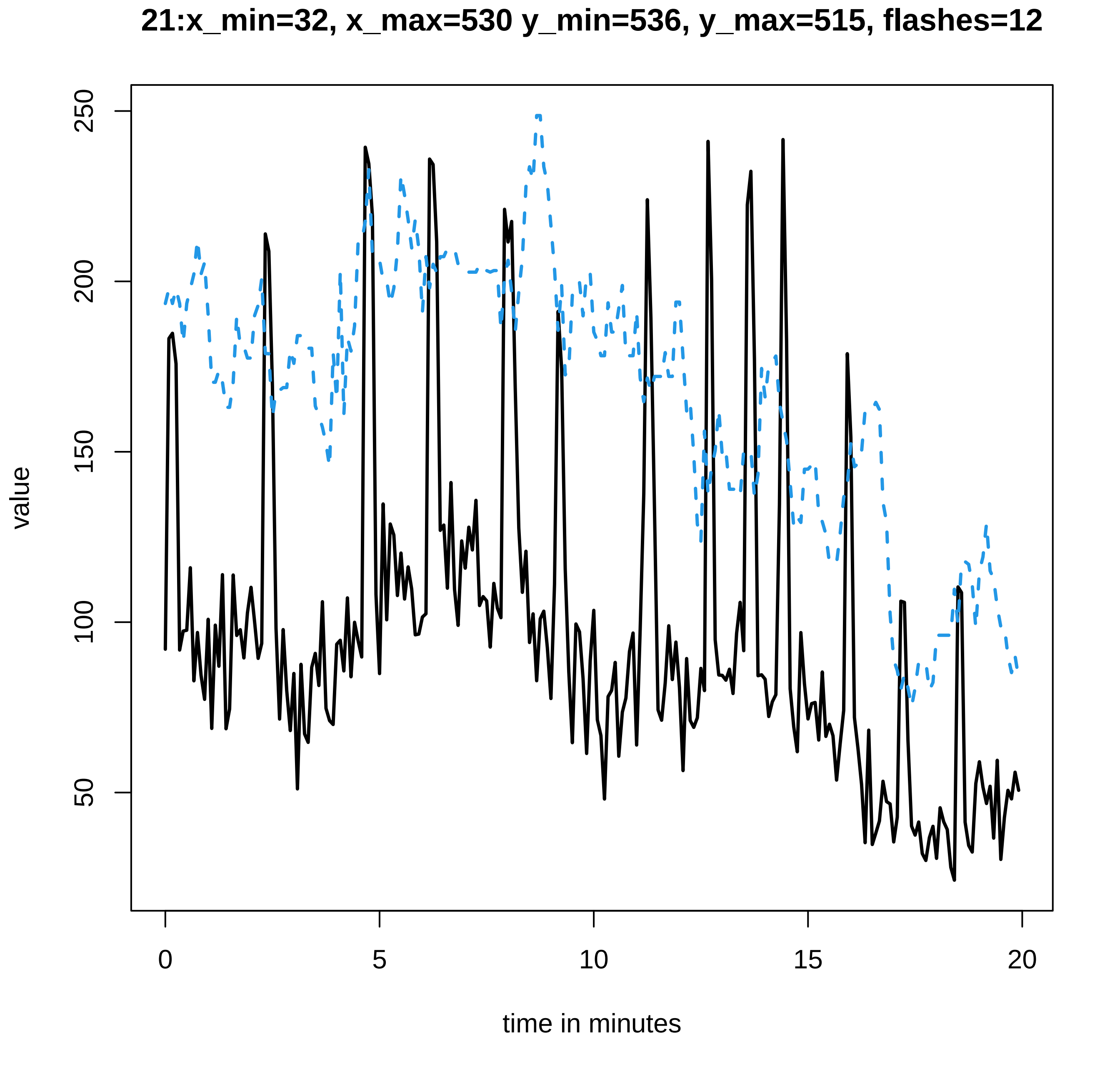
21:x_min=32, x_max=530 y_min=536, y_max=515, flashes=12
value
250
200
150
100
50
0
5
10
15
20
time in minutes

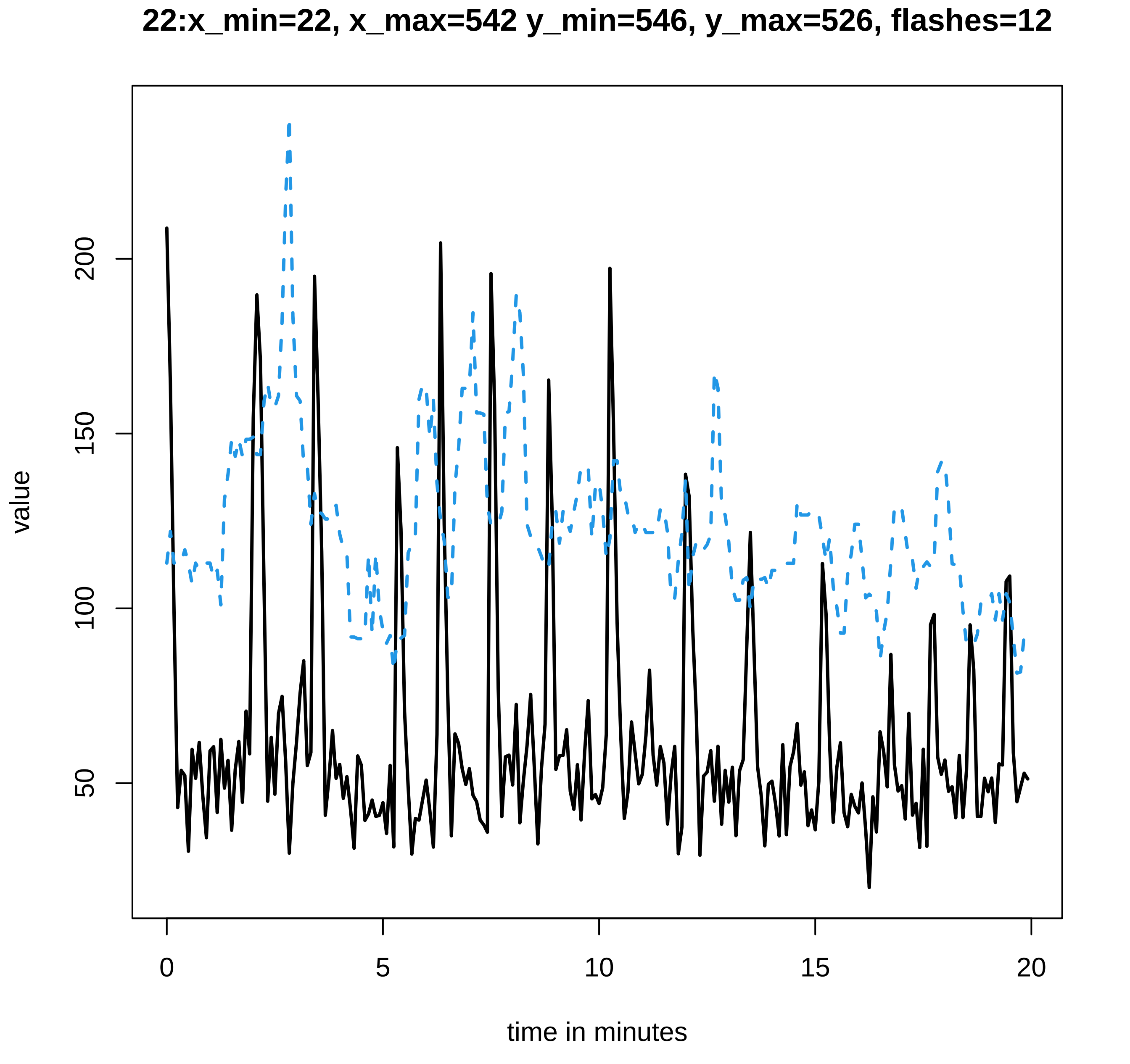
22:x_min=22, x_max=542 y_min=546, y_max=526, flashes=12
value
50
100
150
200
0
5
10
15
20
time in minutes

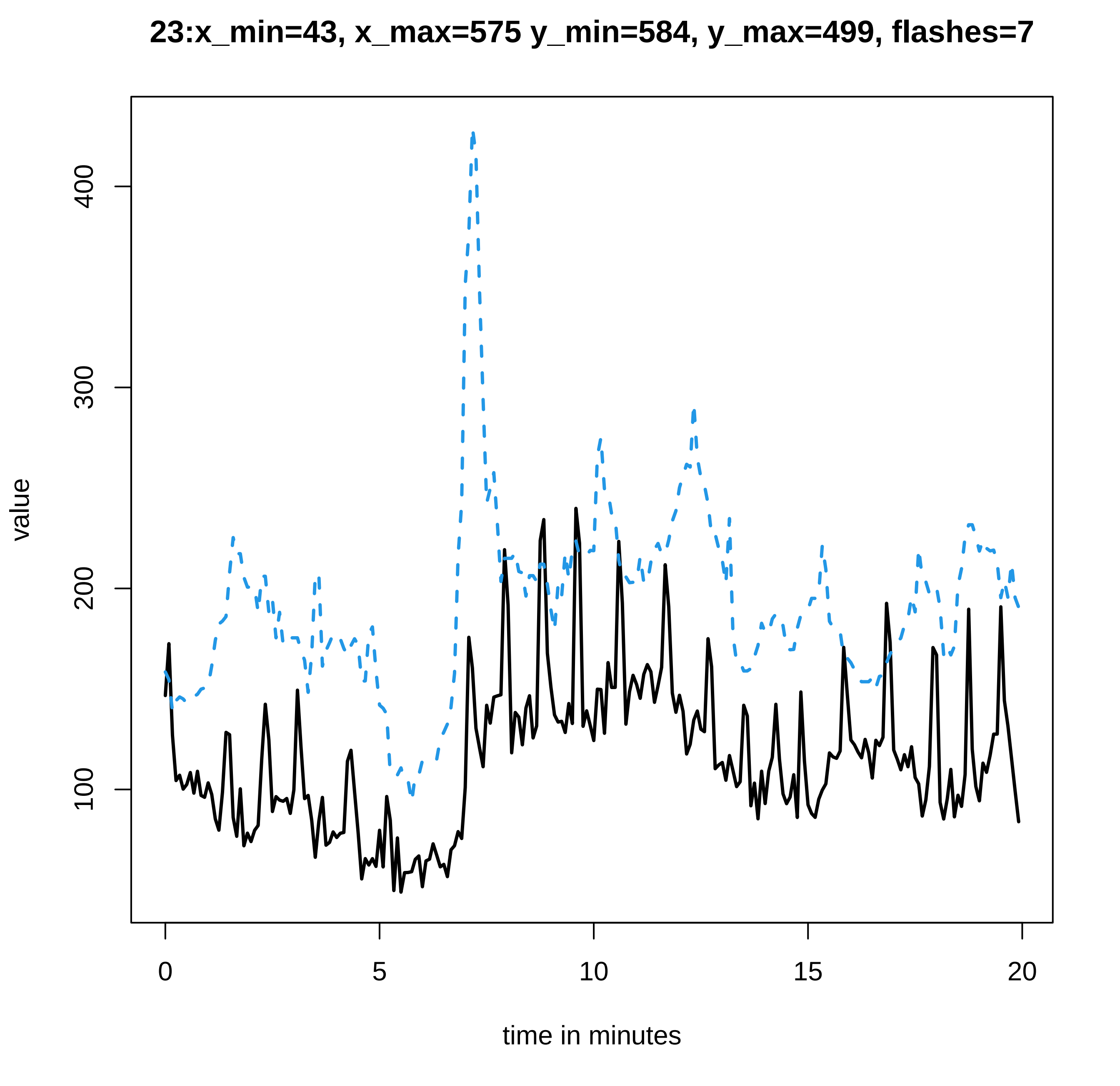

23:x_min=43, x_max=575 y_min=584, y_max=499, flashes=7
value
100
200
300
400
0
5
10
15
20
time in minutes

### 24:x_min=16, x_max=512 y_min=515, y_max=572, flashes=6

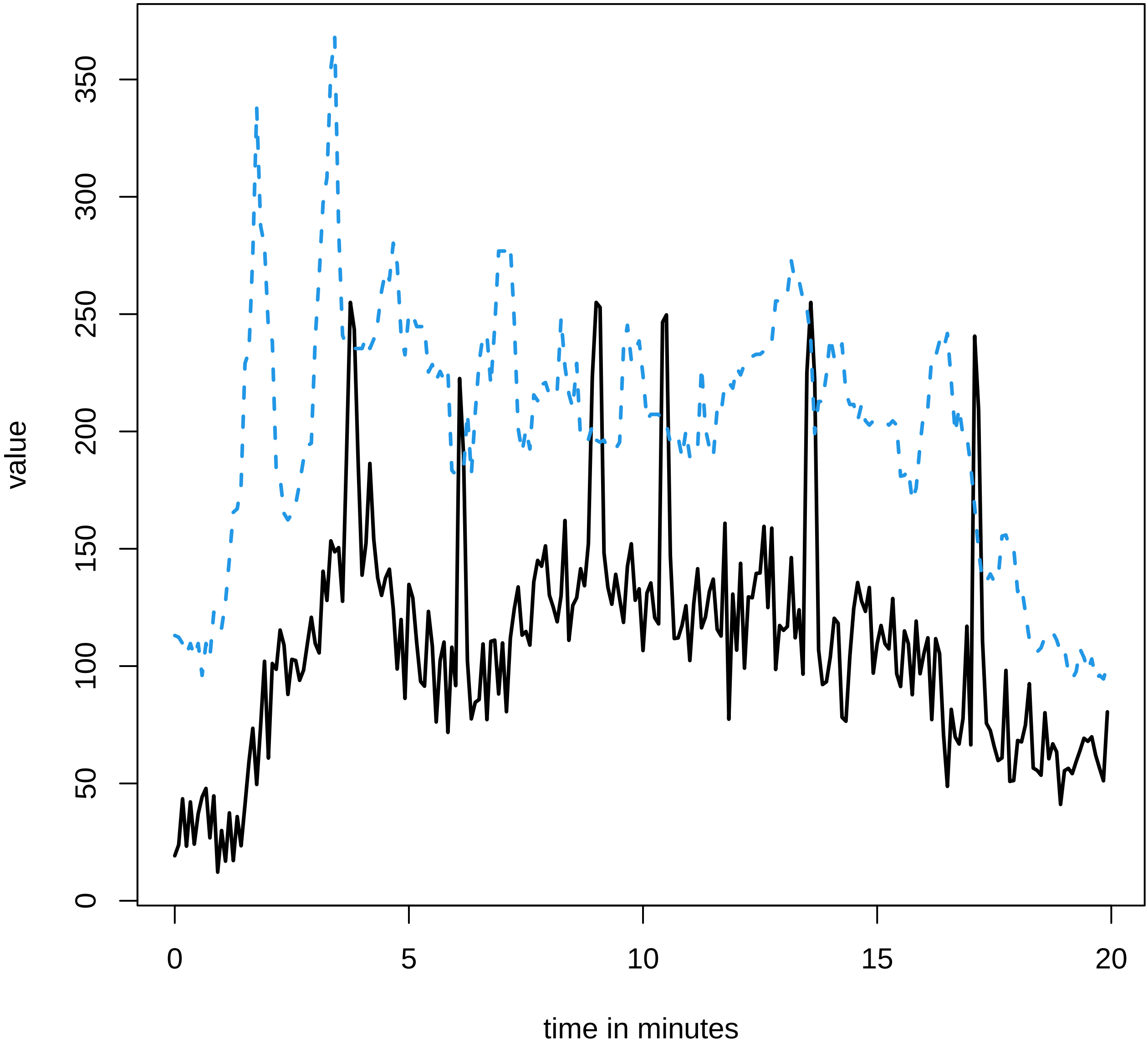

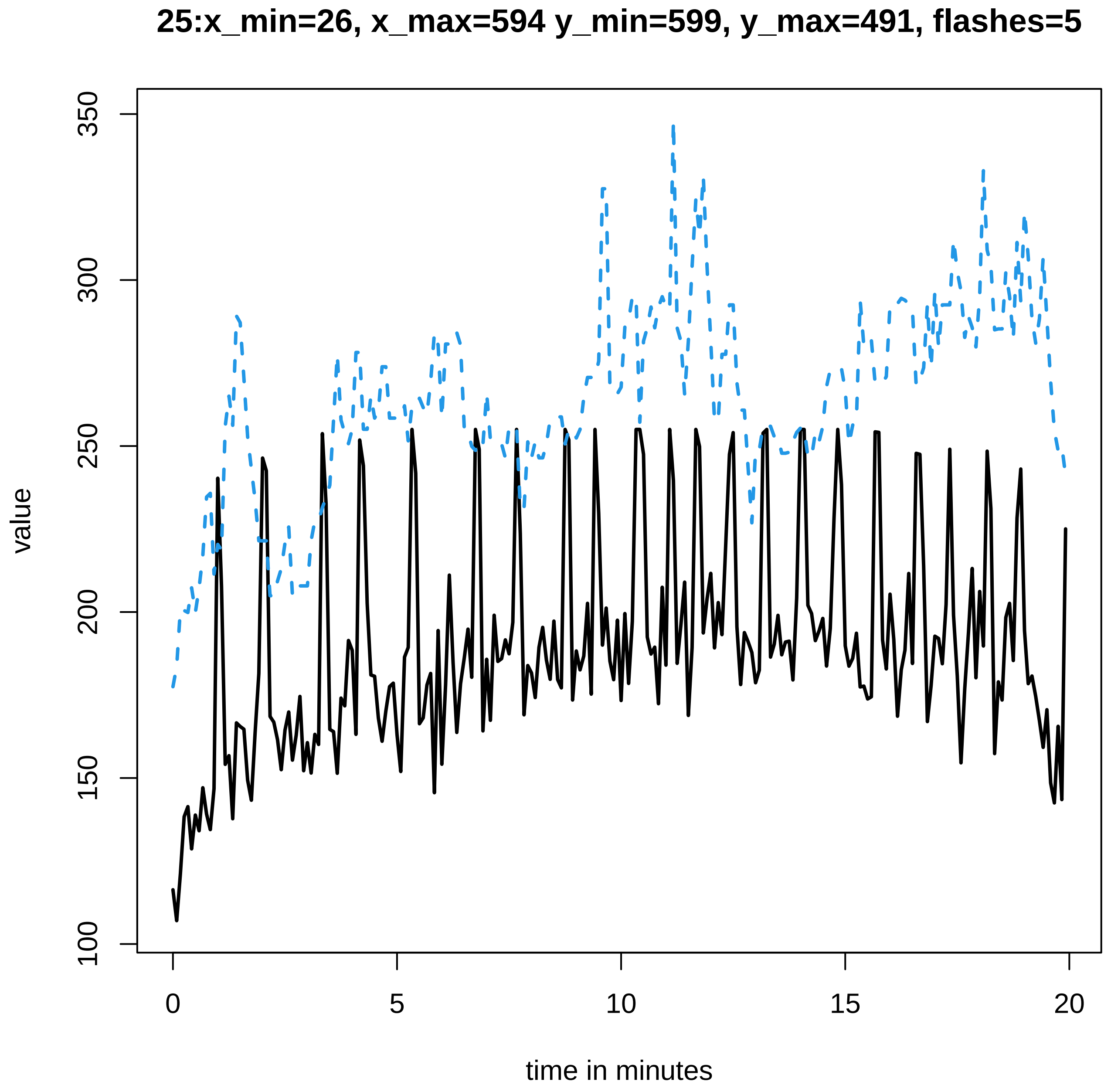

25:x_min=26, x_max=594 y_min=599, y_max=491, flashes=5
value
time in minutes
350
300
250
200
150
100
0
5
10
15
20

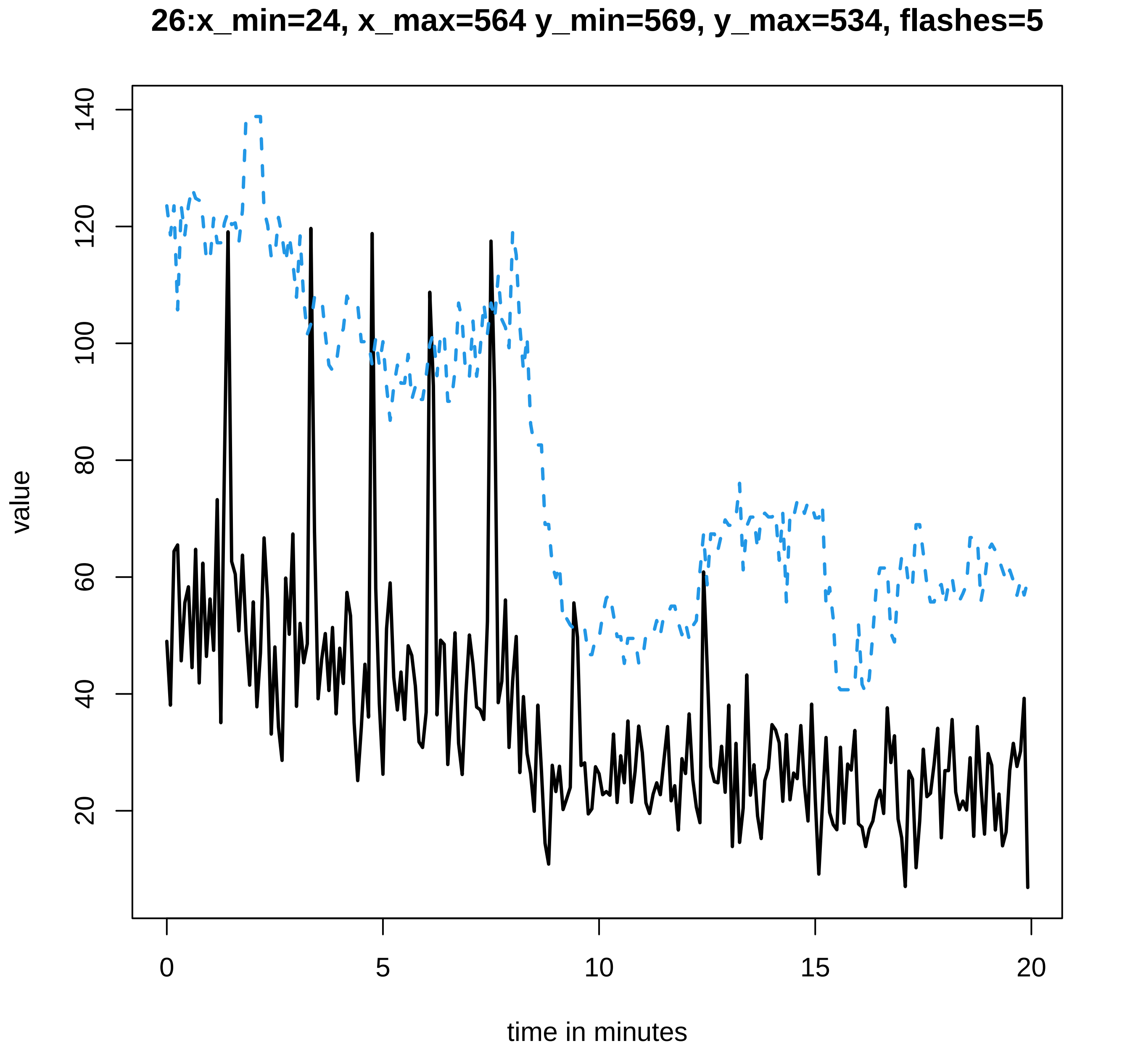

26:x_min=24, x_max=564 y_min=569, y_max=534, flashes=5
value
time in minutes
0
5
10
15
20
20
40
60
80
100
120
140

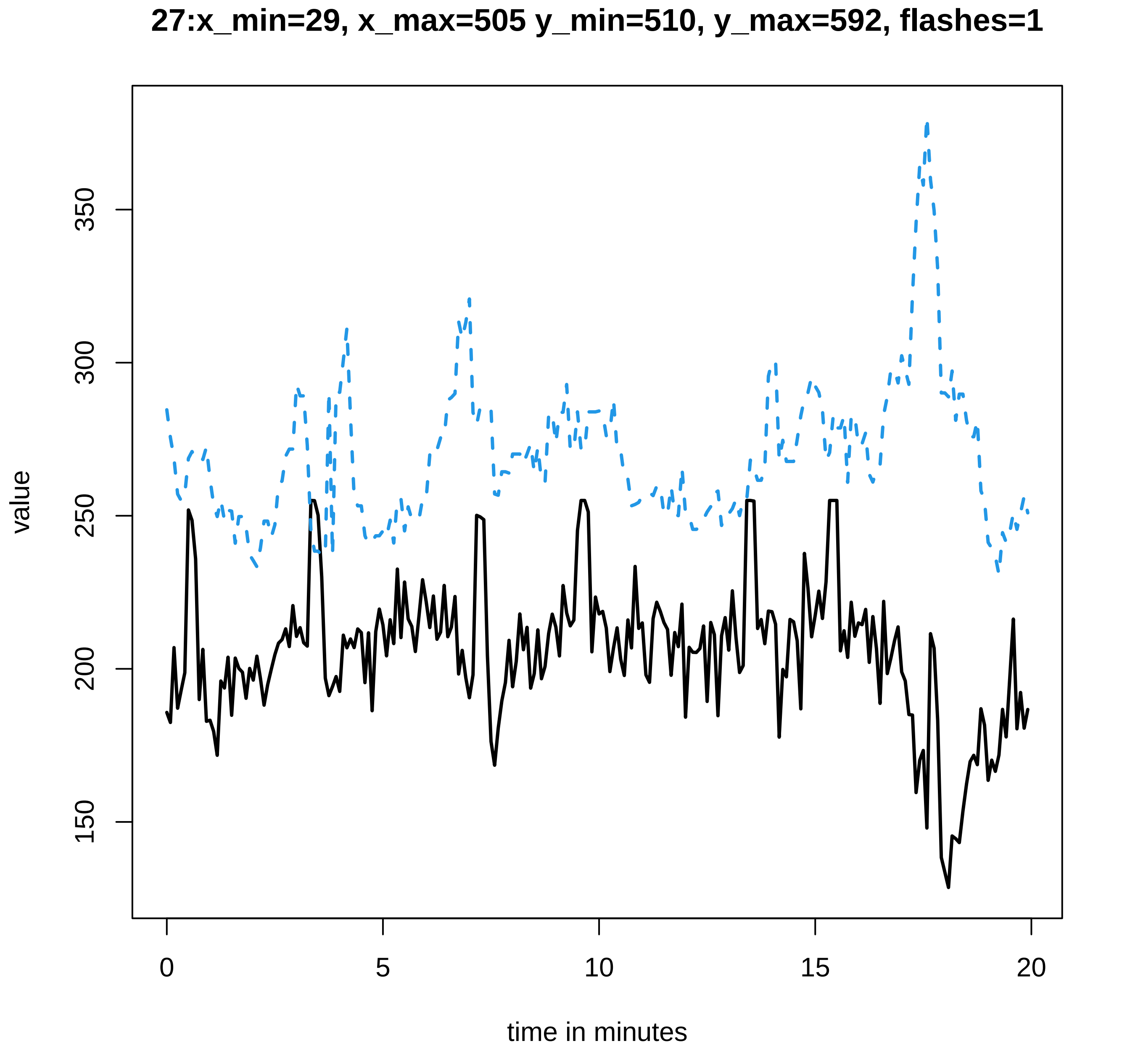
27:x_min=29, x_max=505 y_min=510, y_max=592, flashes=1
value
150
200
250
300
350
0
5
10
15
20
time in minutes

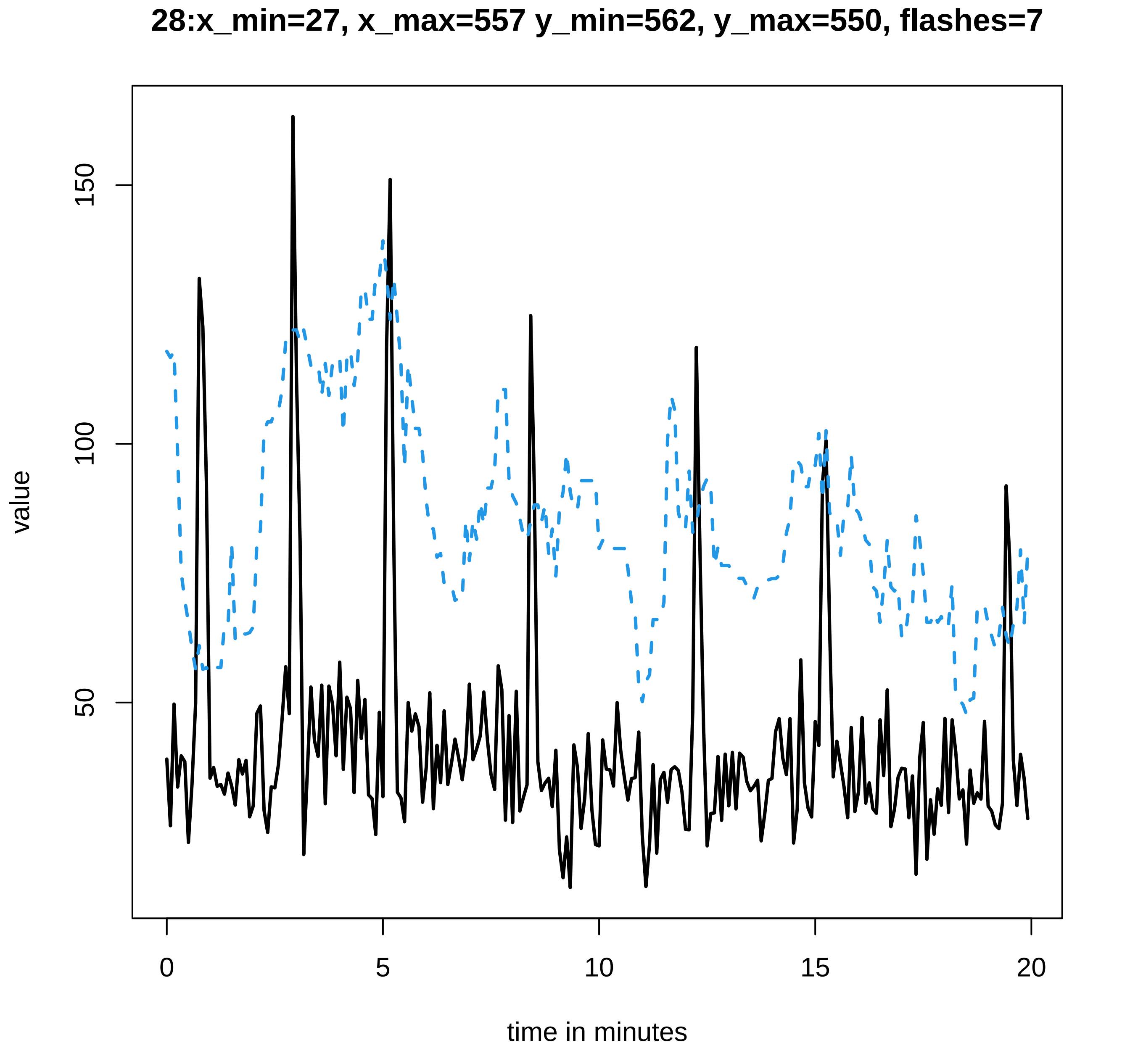

28:x_min=27, x_max=557 y_min=562, y_max=550, flashes=7
value
150
100
50
0
5
10
15
20
time in minutes

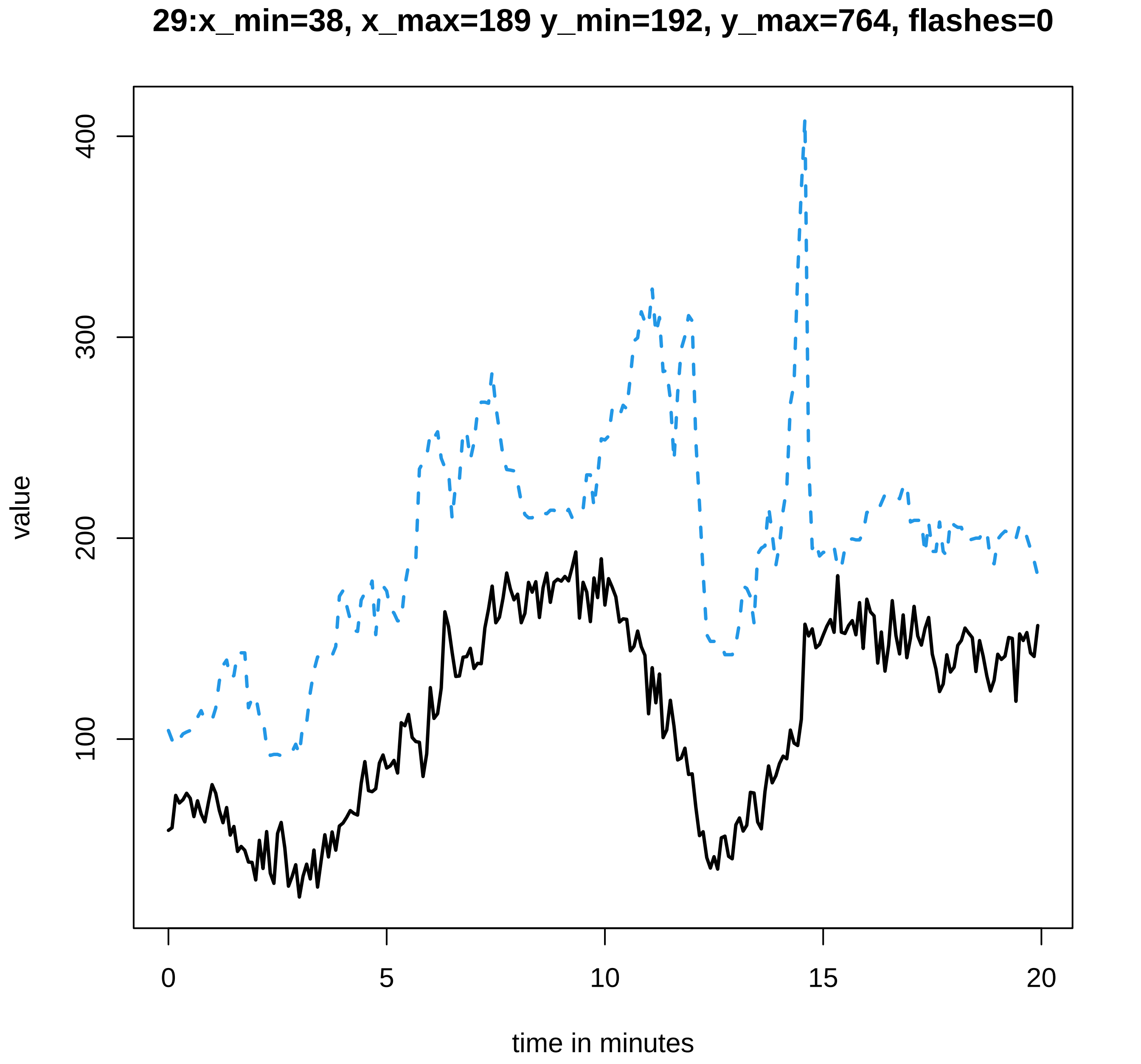

29:x_min=38, x_max=189 y_min=192, y_max=764, flashes=0
value
100
200
300
400
0
5
10
15
20
time in minutes

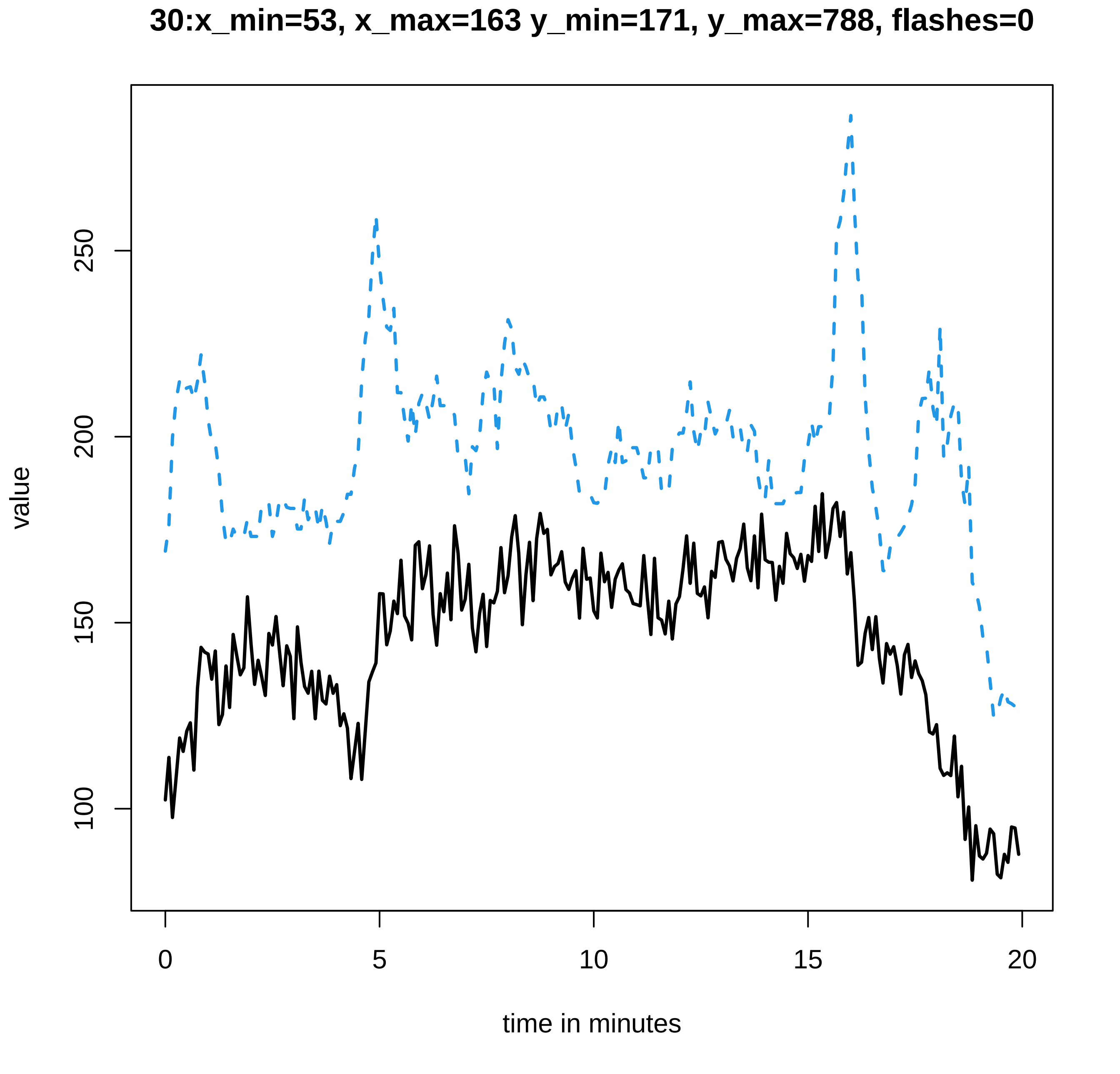

30:x_min=53, x_max=163 y_min=171, y_max=788, flashes=0
value
100
150
200
250
0
5
10
15
20
time in minutes

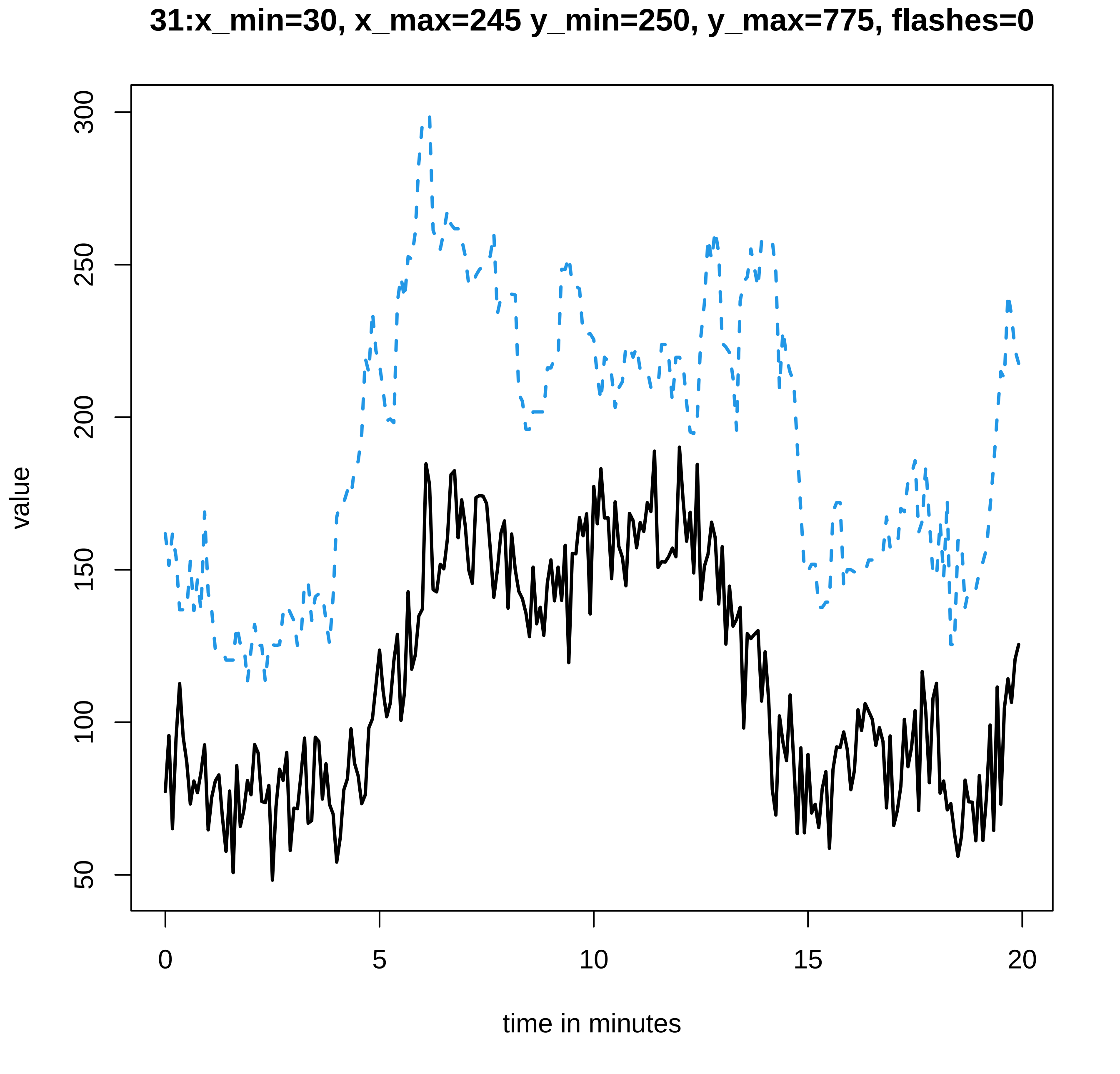
31:x_min=30, x_max=245 y_min=250, y_max=775, flashes=0
value
50
100
150
200
250
300
0
5
10
15
20
time in minutes

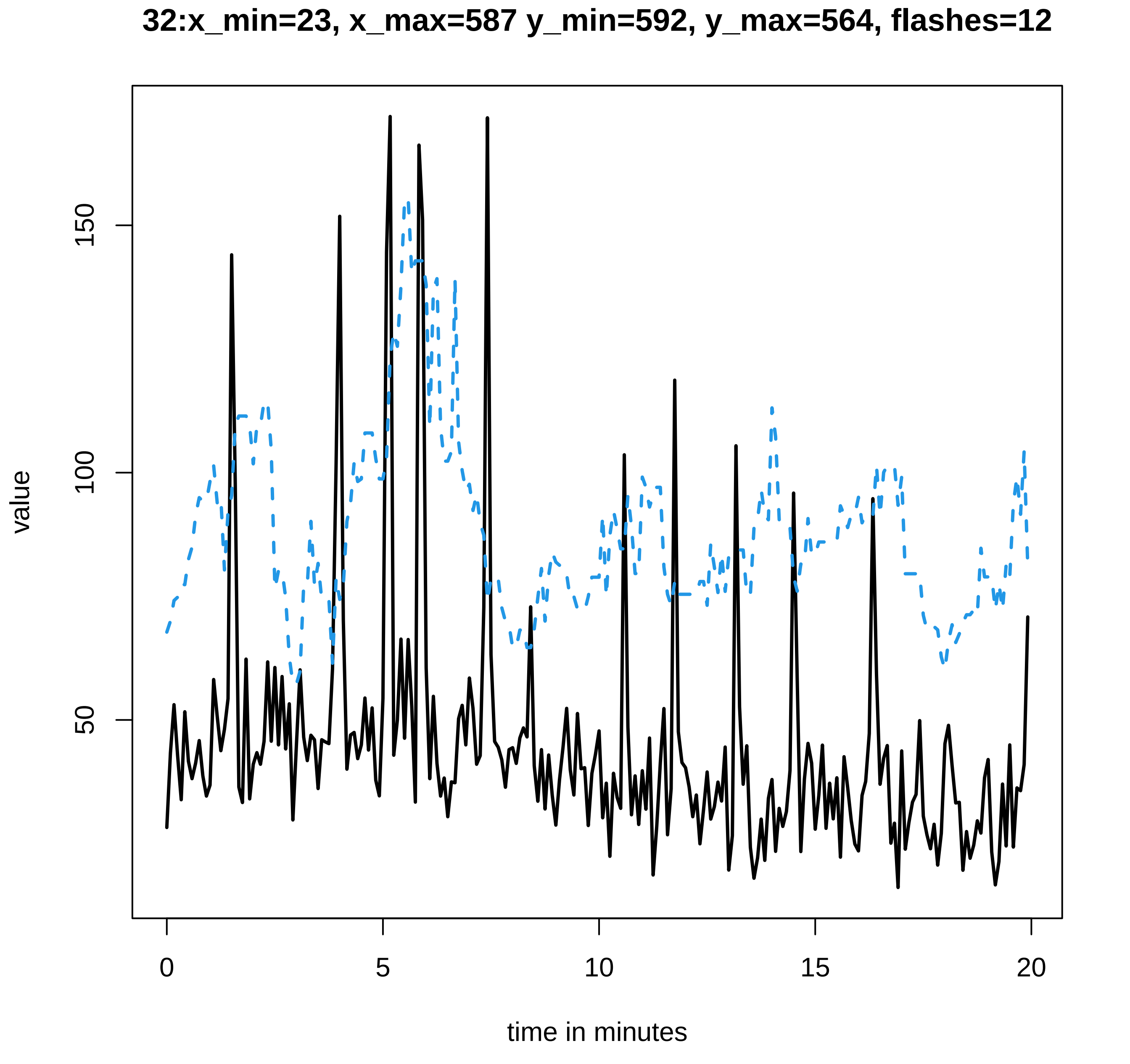

32:x_min=23, x_max=587 y_min=592, y_max=564, flashes=12
value
150
100
50
0
5
10
15
20
time in minutes

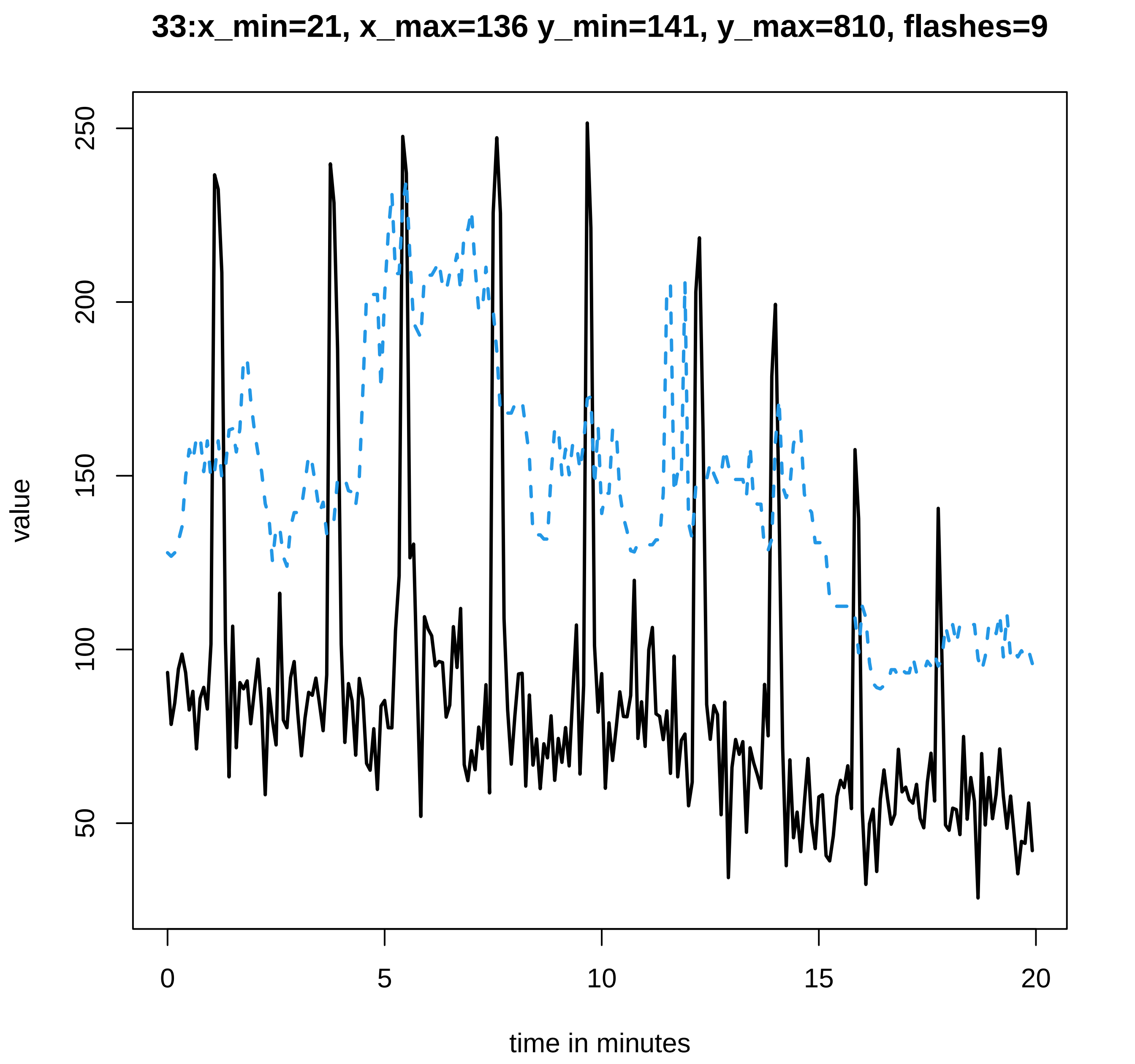
33:x_min=21, x_max=136 y_min=141, y_max=810, flashes=9
value
250
200
150
100
50
0
5
10
15
20
time in minutes

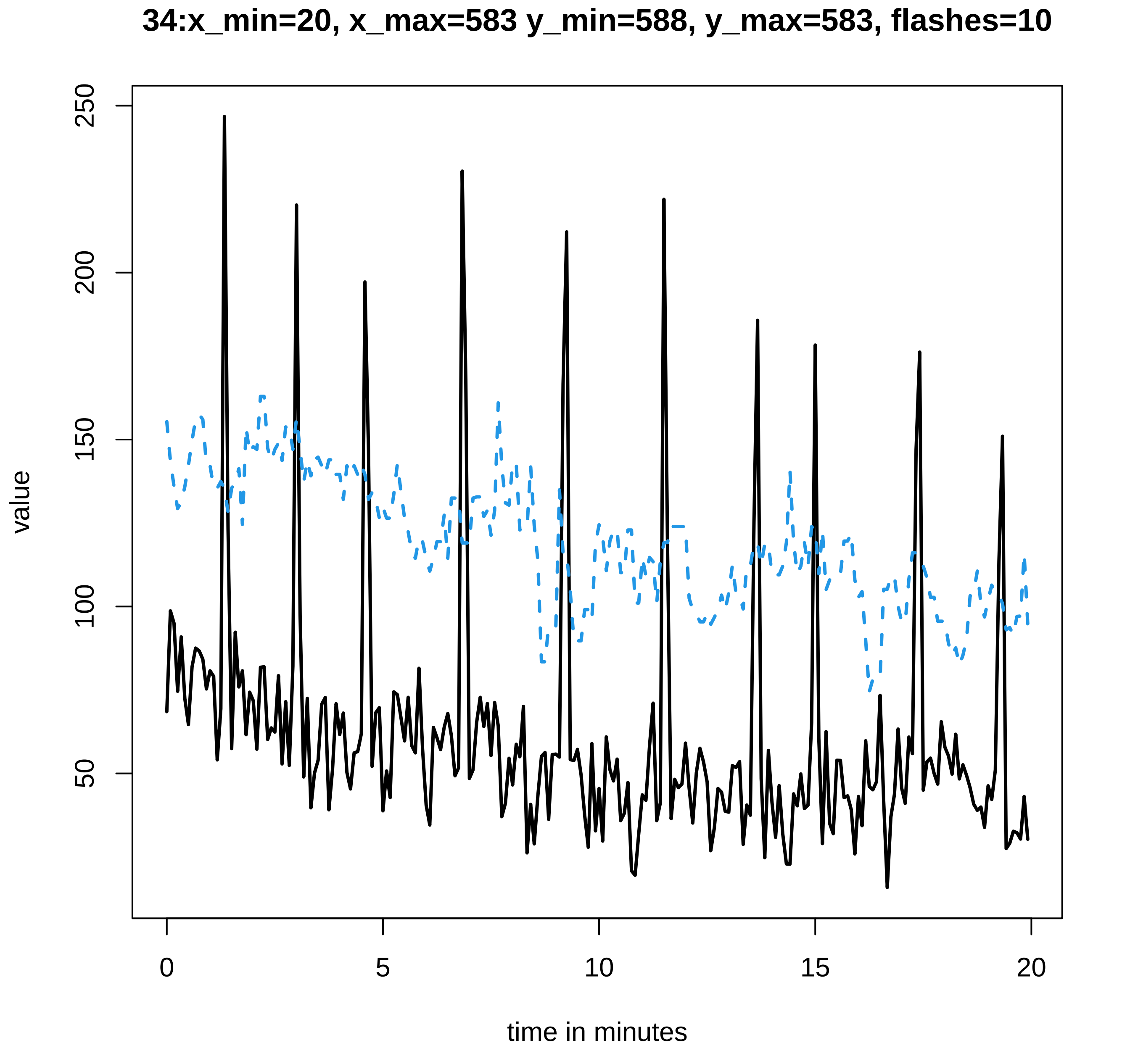
34:x_min=20, x_max=583 y_min=588, y_max=583, flashes=10
value
250
200
150
100
50
0
5
10
15
20
time in minutes

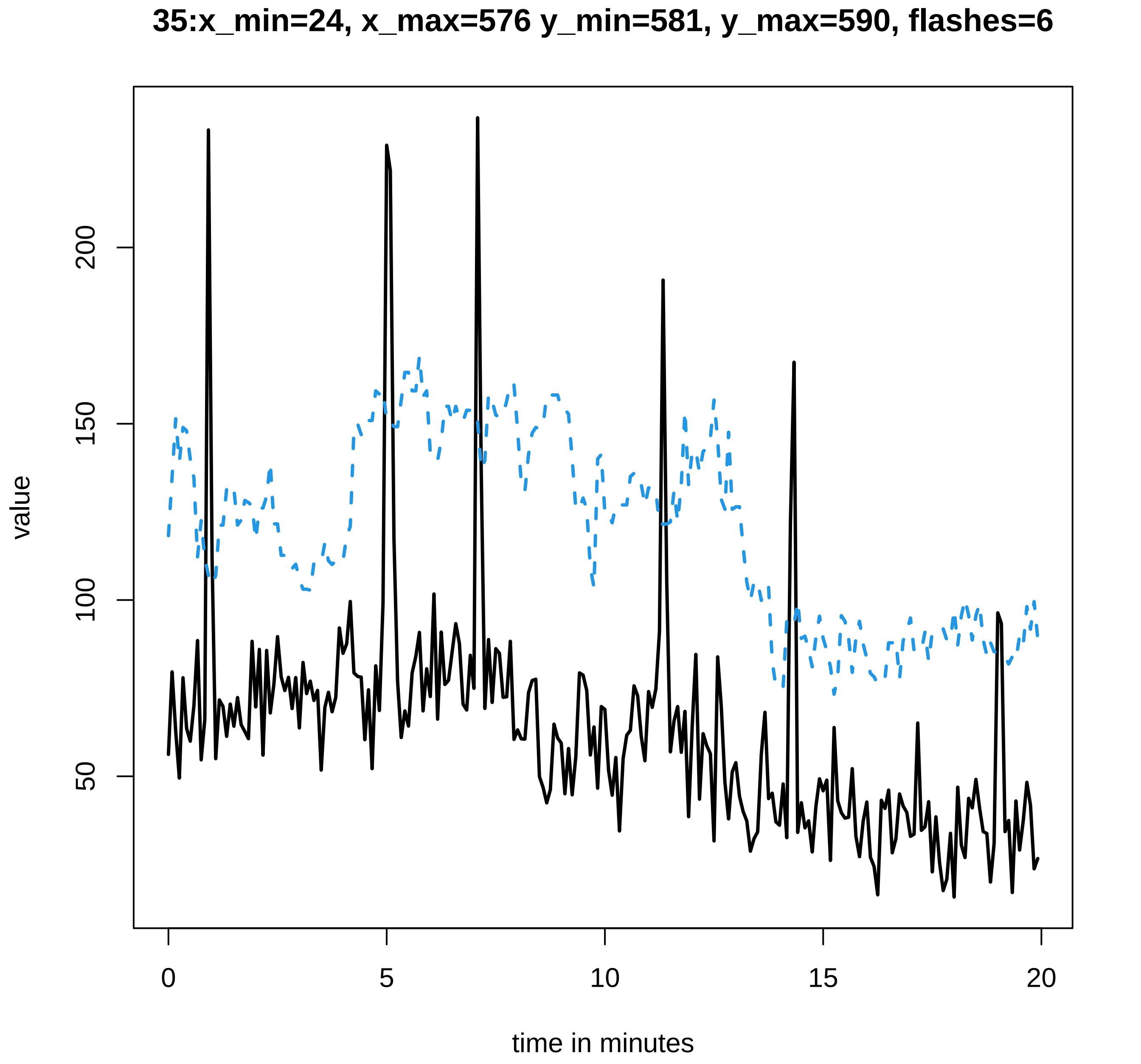

35:x_min=24, x_max=576 y_min=581, y_max=590, flashes=6
value
200
150
100
50
0
5
10
15
20
time in minutes

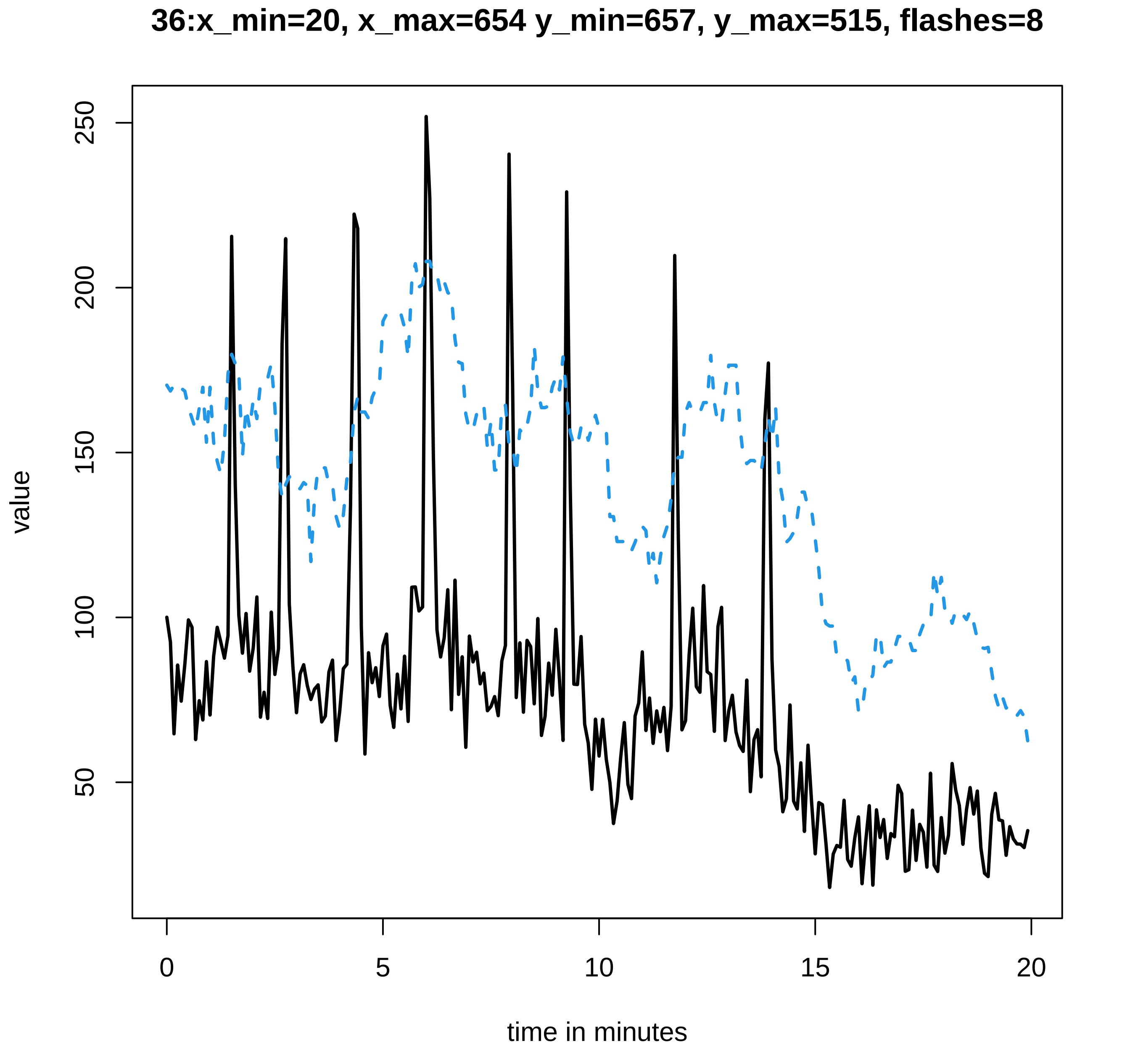
36:x_min=20, x_max=654 y_min=657, y_max=515, flashes=8
value
250
200
150
100
50
0
5
10
15
20
time in minutes

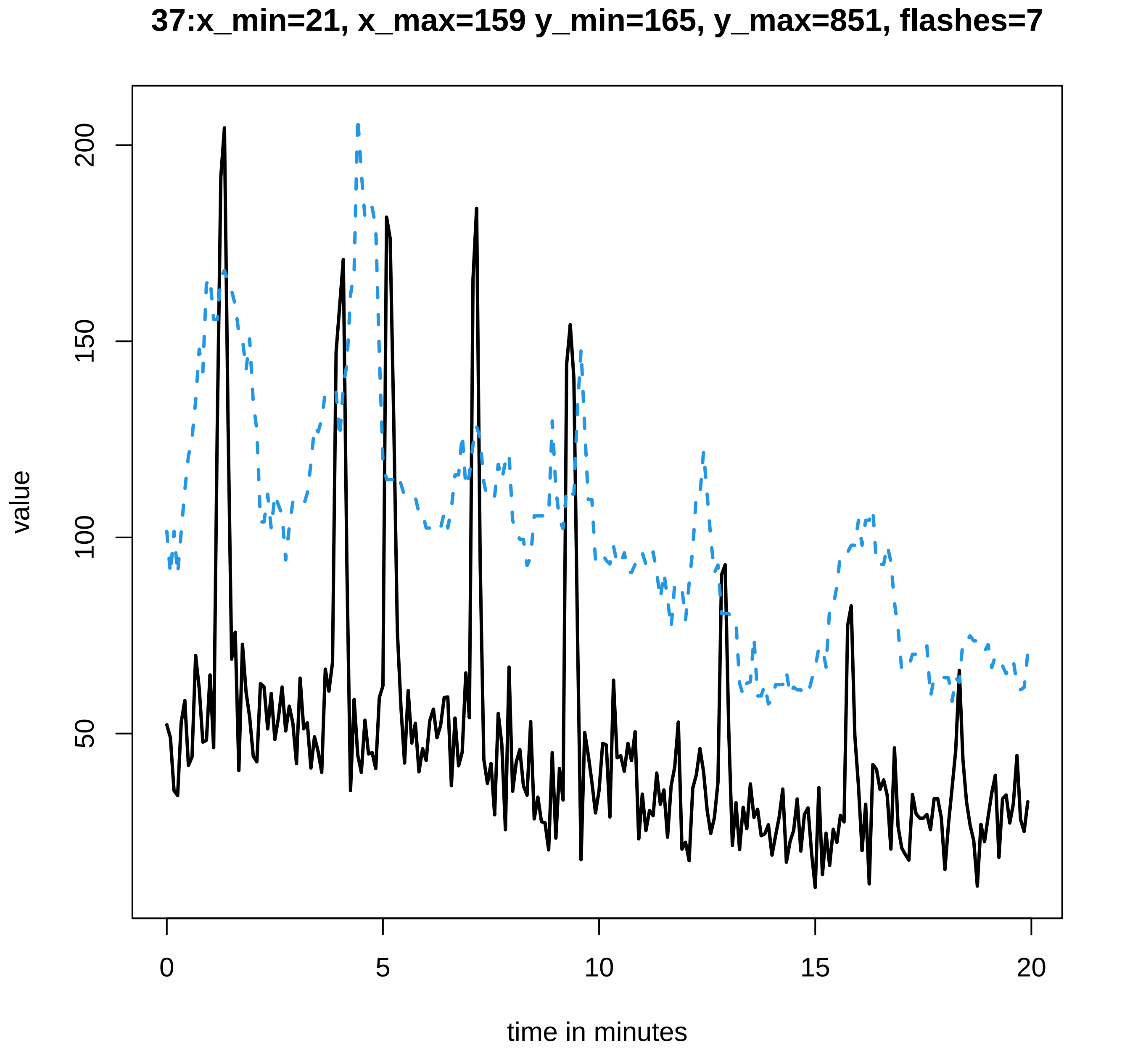

37:x_min=21, x_max=159 y_min=165, y_max=851, flashes=7
value
200
150
100
50
0
5
10
15
20
time in minutes

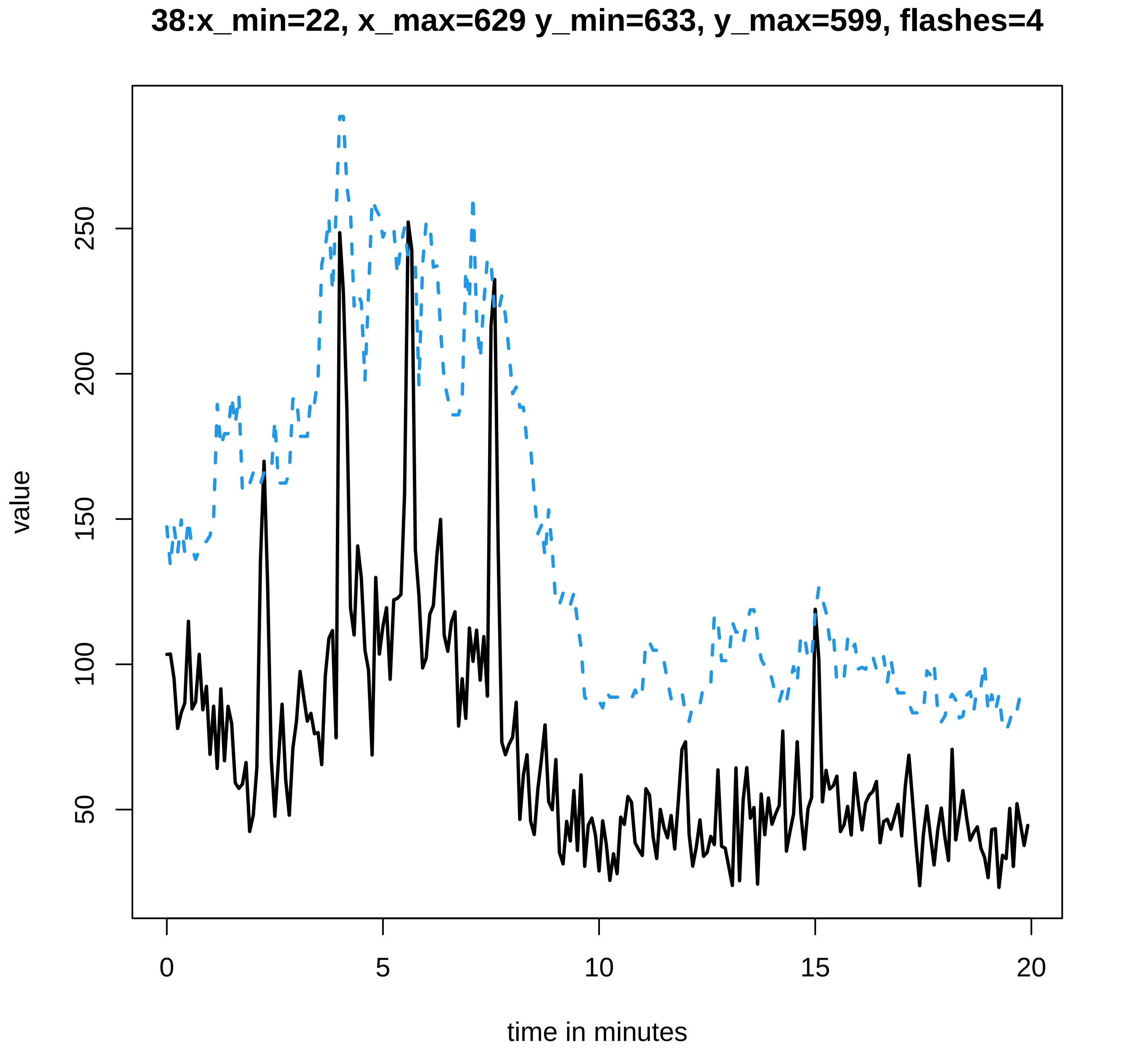

38:x_min=22, x_max=629 y_min=633, y_max=599, flashes=4
value
250
200
150
100
50
0
5
10
15
20
time in minutes

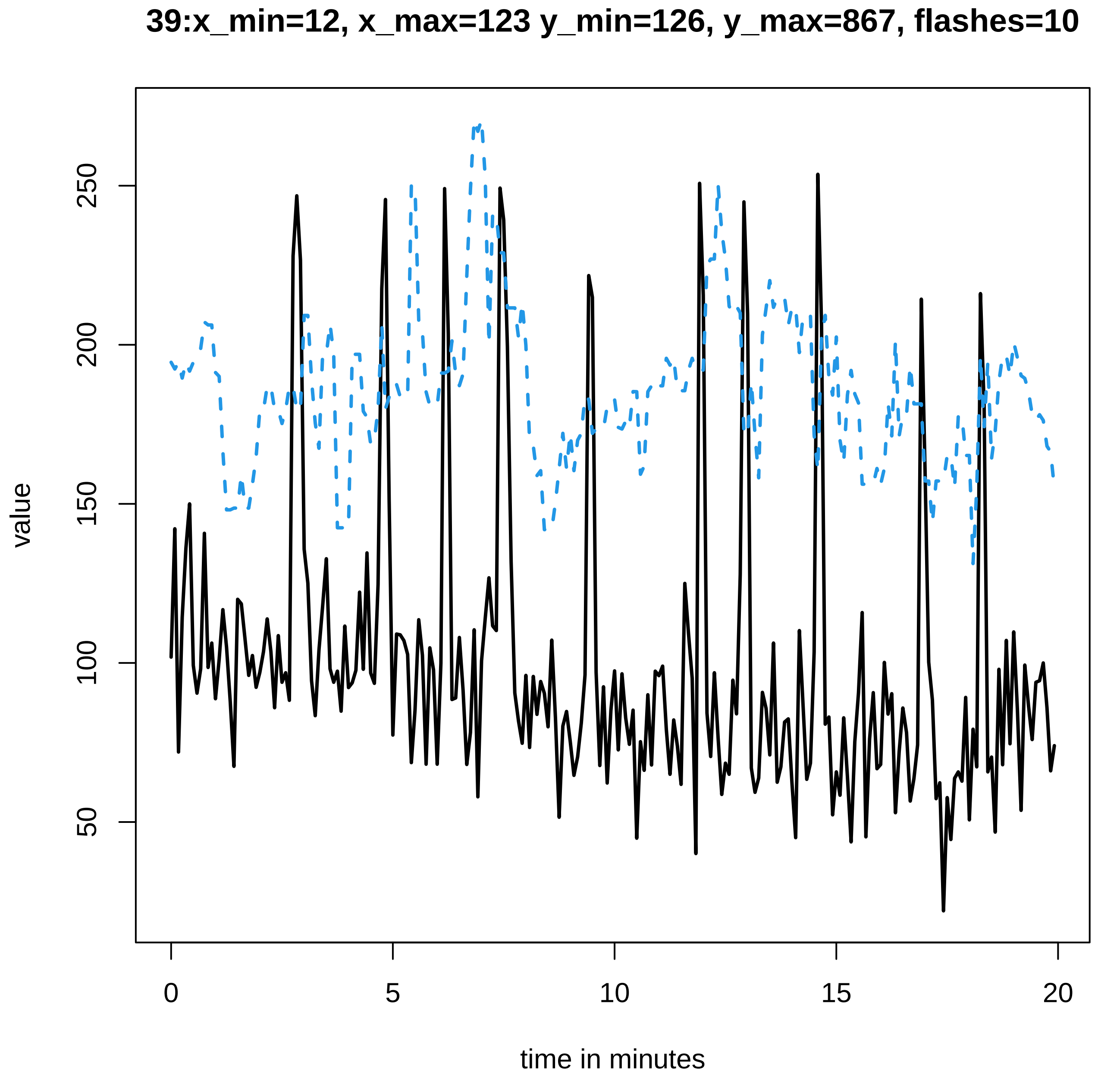
39:x_min=12, x_max=123 y_min=126, y_max=867, flashes=10
value
50
100
150
200
250
0
5
10
15
20
time in minutes

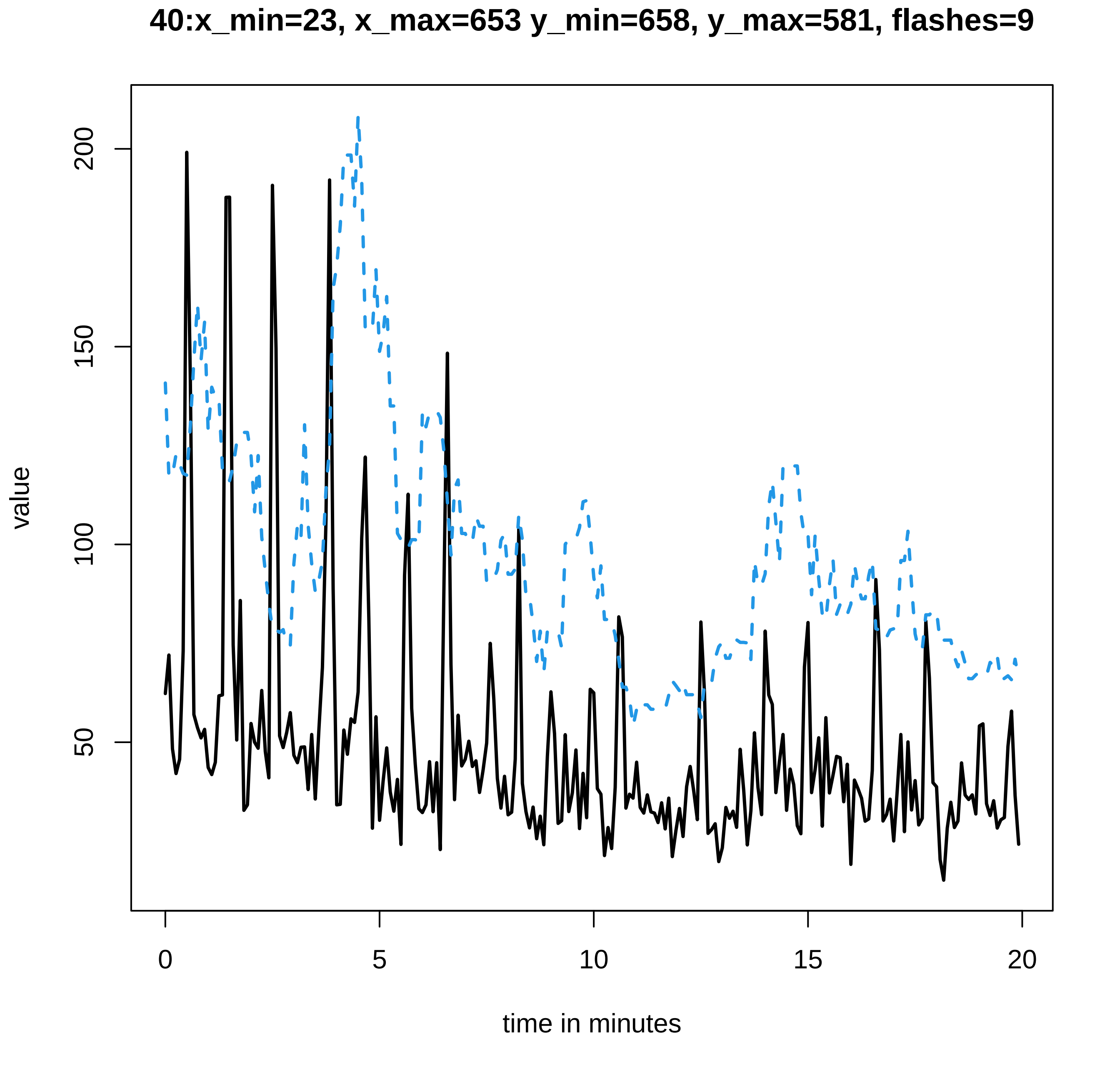

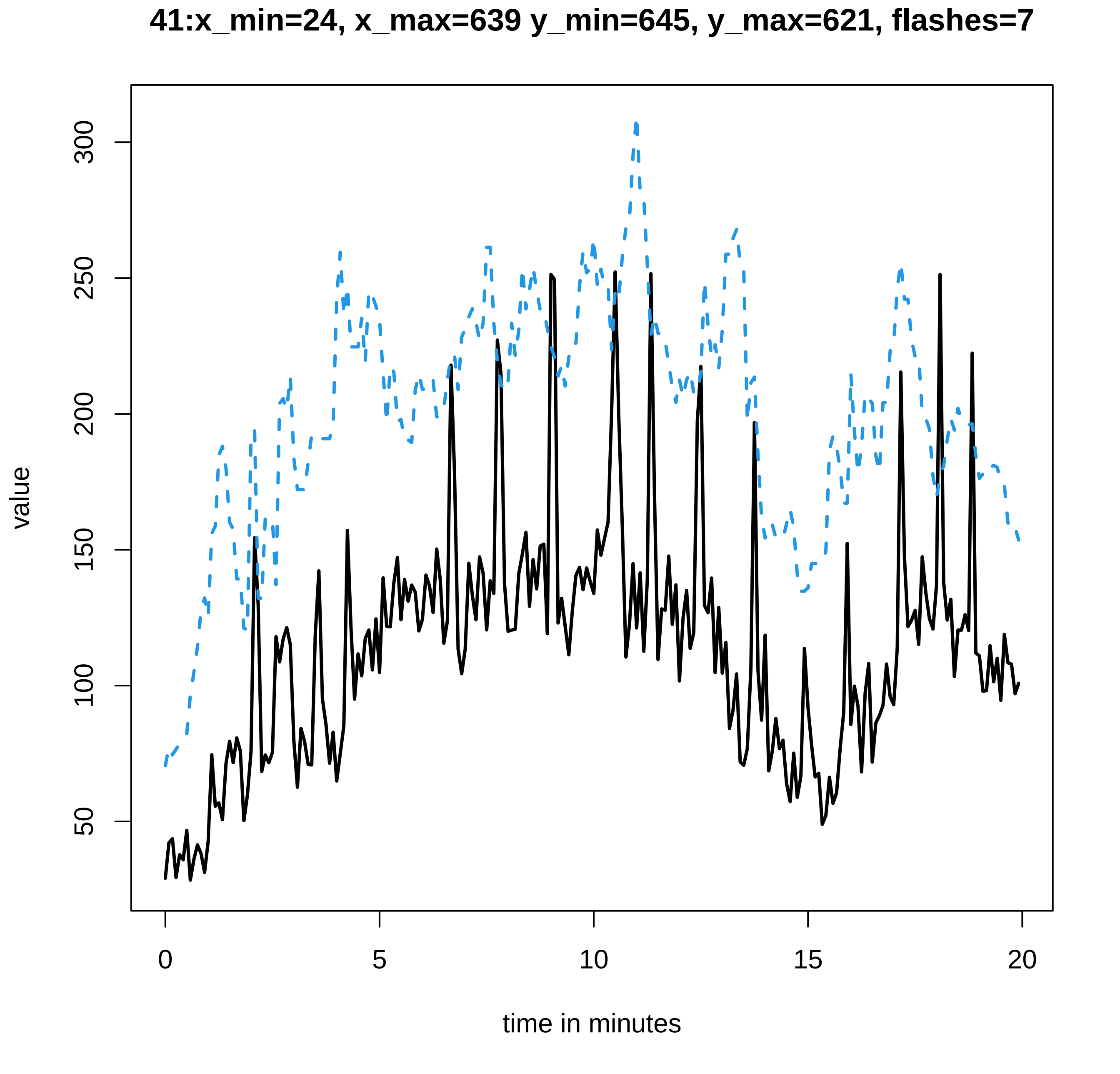
41:x_min=24, x_max=639 y_min=645, y_max=621, flashes=7
value
time in minutes
0
5
10
15
20
50
100
150
200
250
300

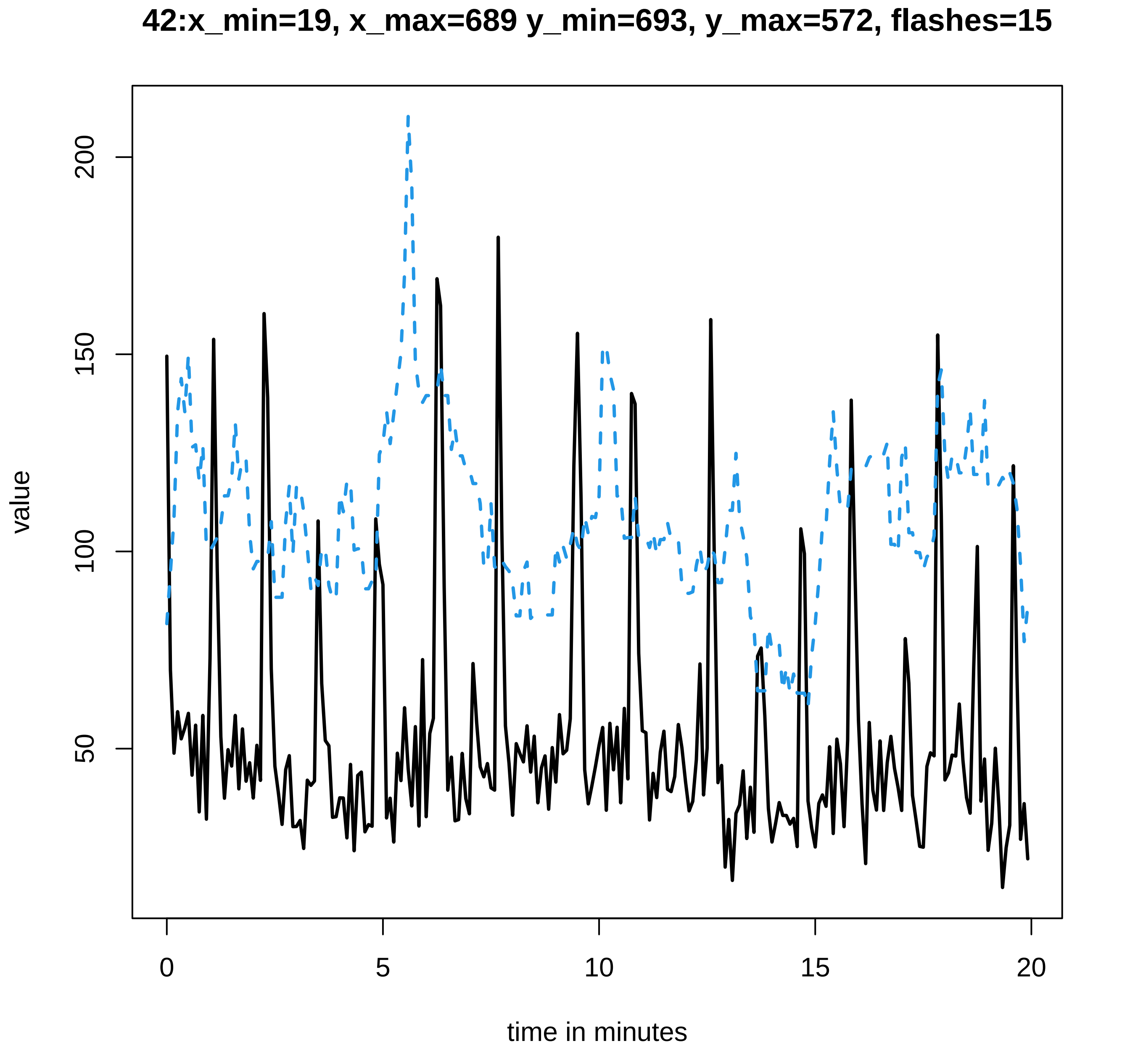
42:x_min=19, x_max=689 y_min=693, y_max=572, flashes=15
value
200
150
100
50
0
5
10
15
20
time in minutes

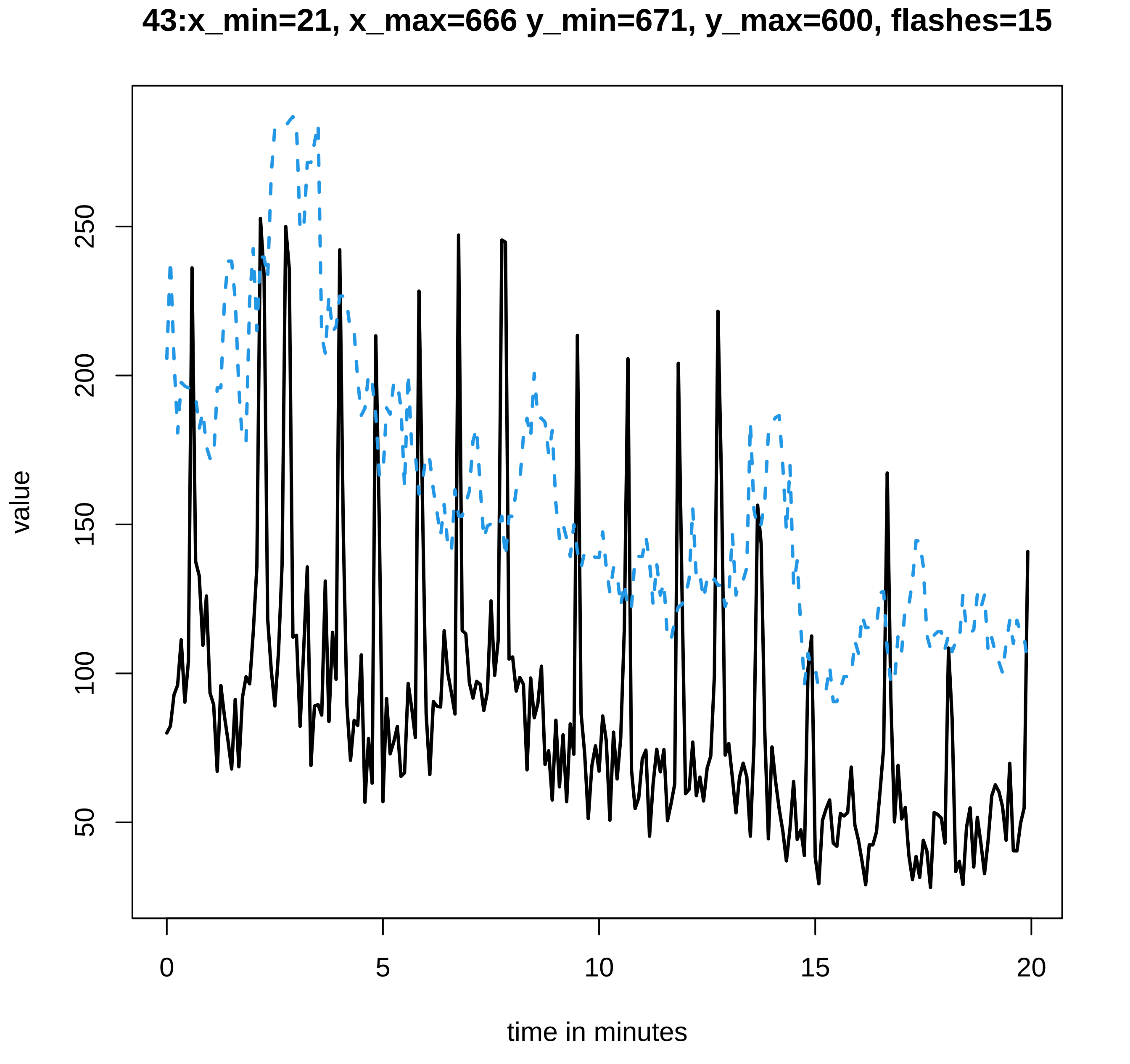
43:x_min=21, x_max=666 y_min=671, y_max=600, flashes=15
value
50
100
150
200
250
0
5
10
15
20
time in minutes

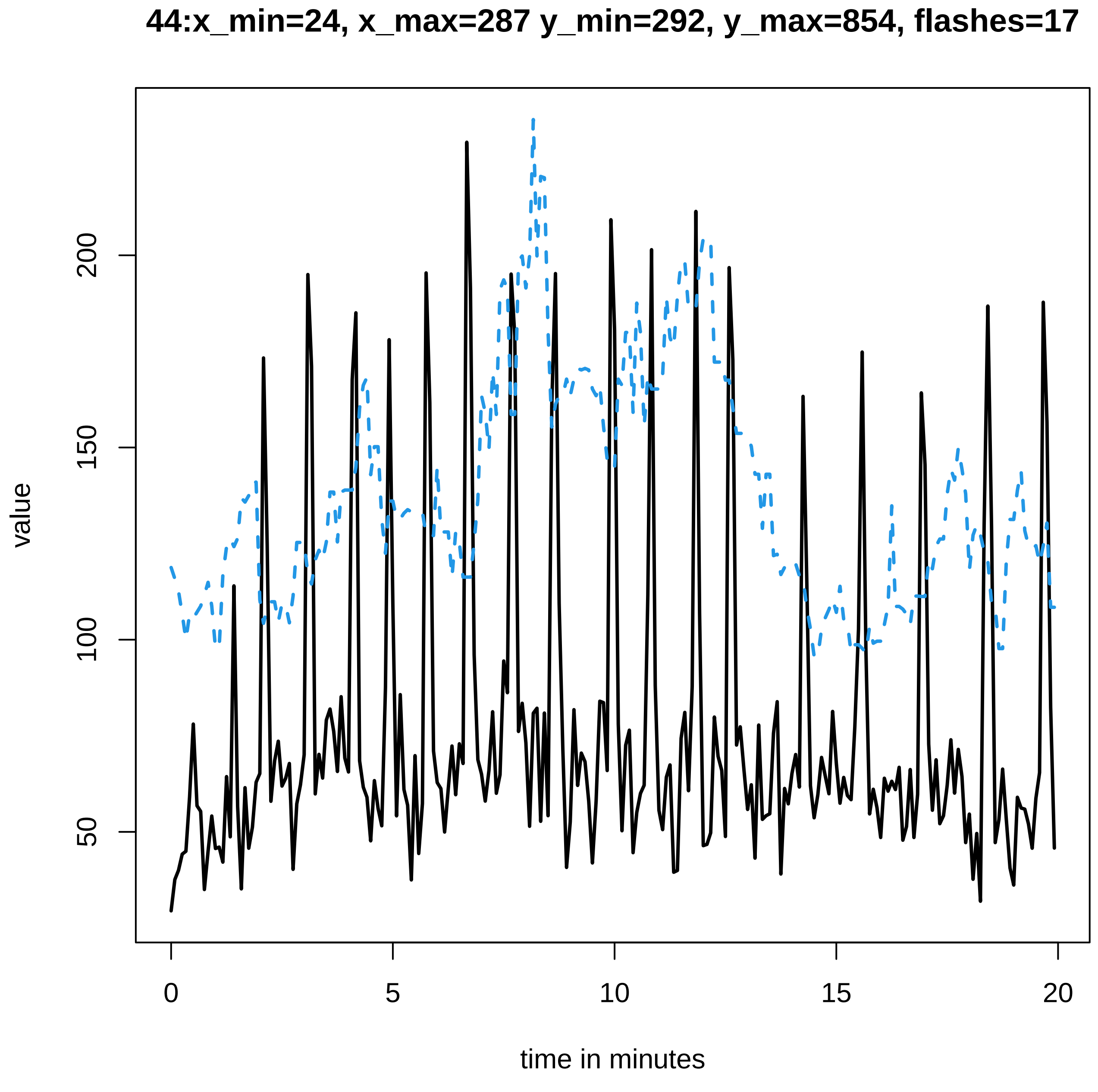

44:x_min=24, x_max=287 y_min=292, y_max=854, flashes=17
value
200
150
100
50
0
5
10
15
20
time in minutes

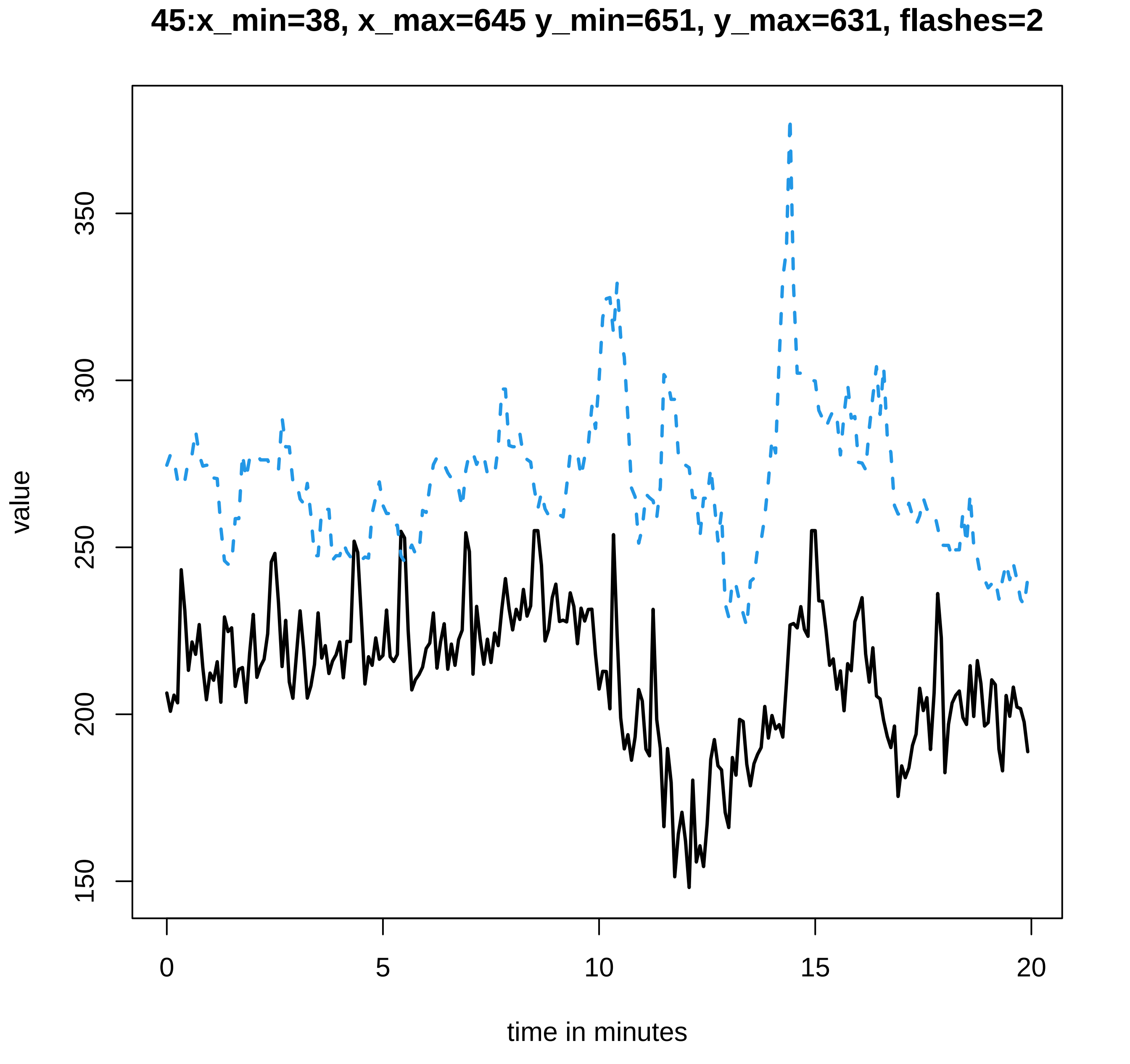
45:x_min=38, x_max=645 y_min=651, y_max=631, flashes=2
value
350
300
250
200
150
0
5
10
15
20
time in minutes

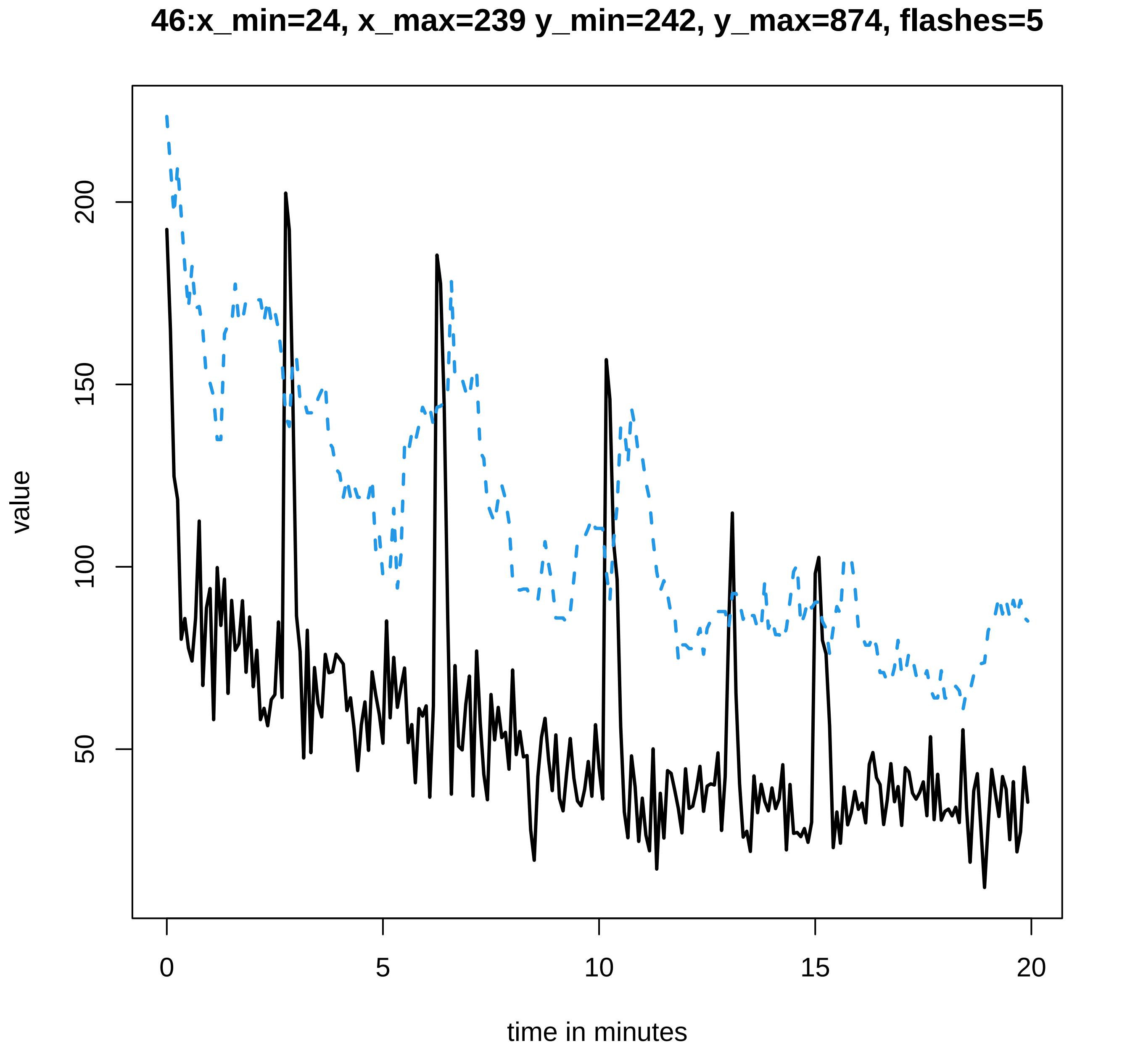
46:x_min=24, x_max=239 y_min=242, y_max=874, flashes=5
value
200
150
100
50
0
5
10
15
20
time in minutes

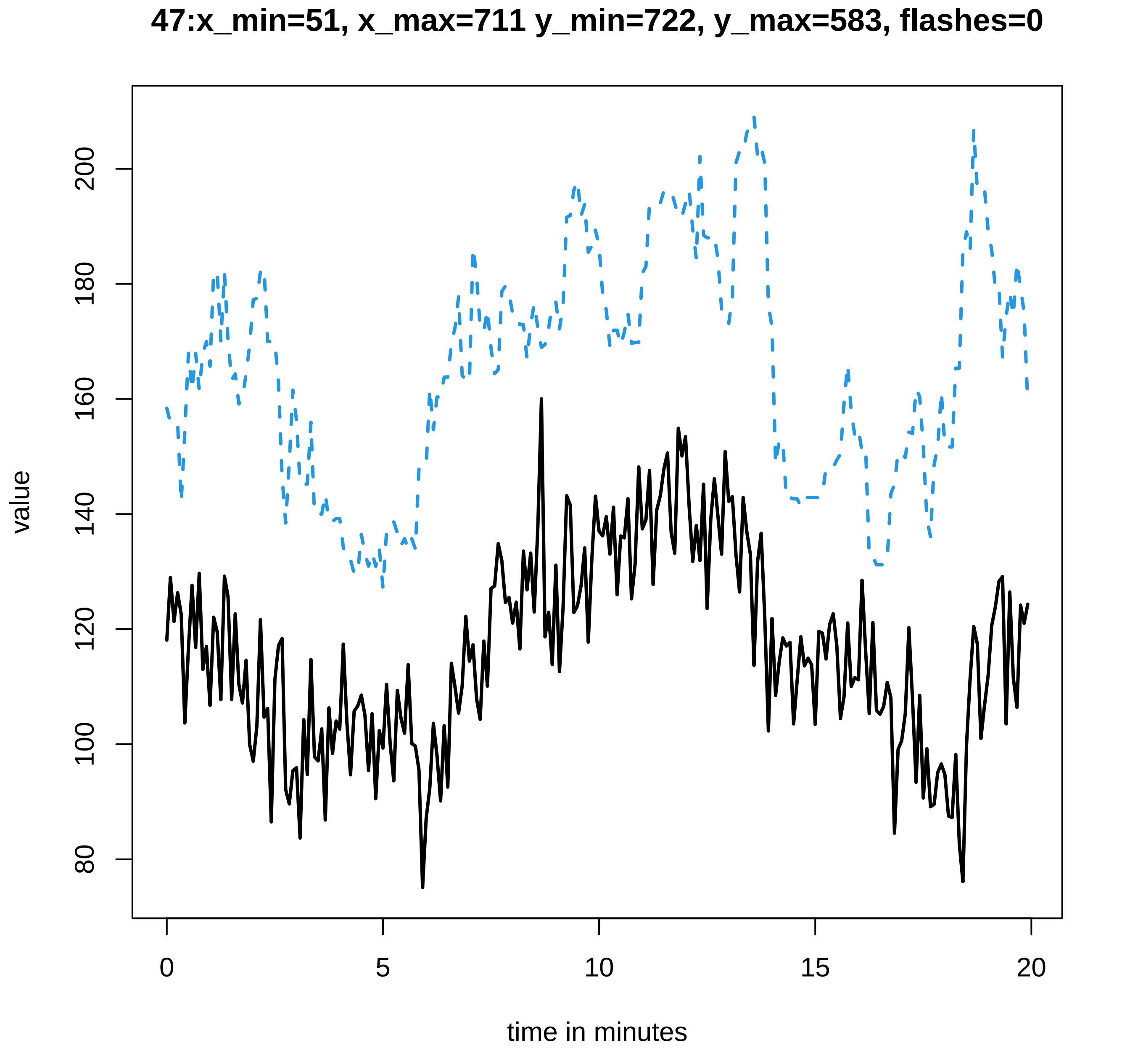

47:x_min=51, x_max=711 y_min=722, y_max=583, flashes=0
value
80
100
120
140
160
180
200
0
5
10
15
20
time in minutes

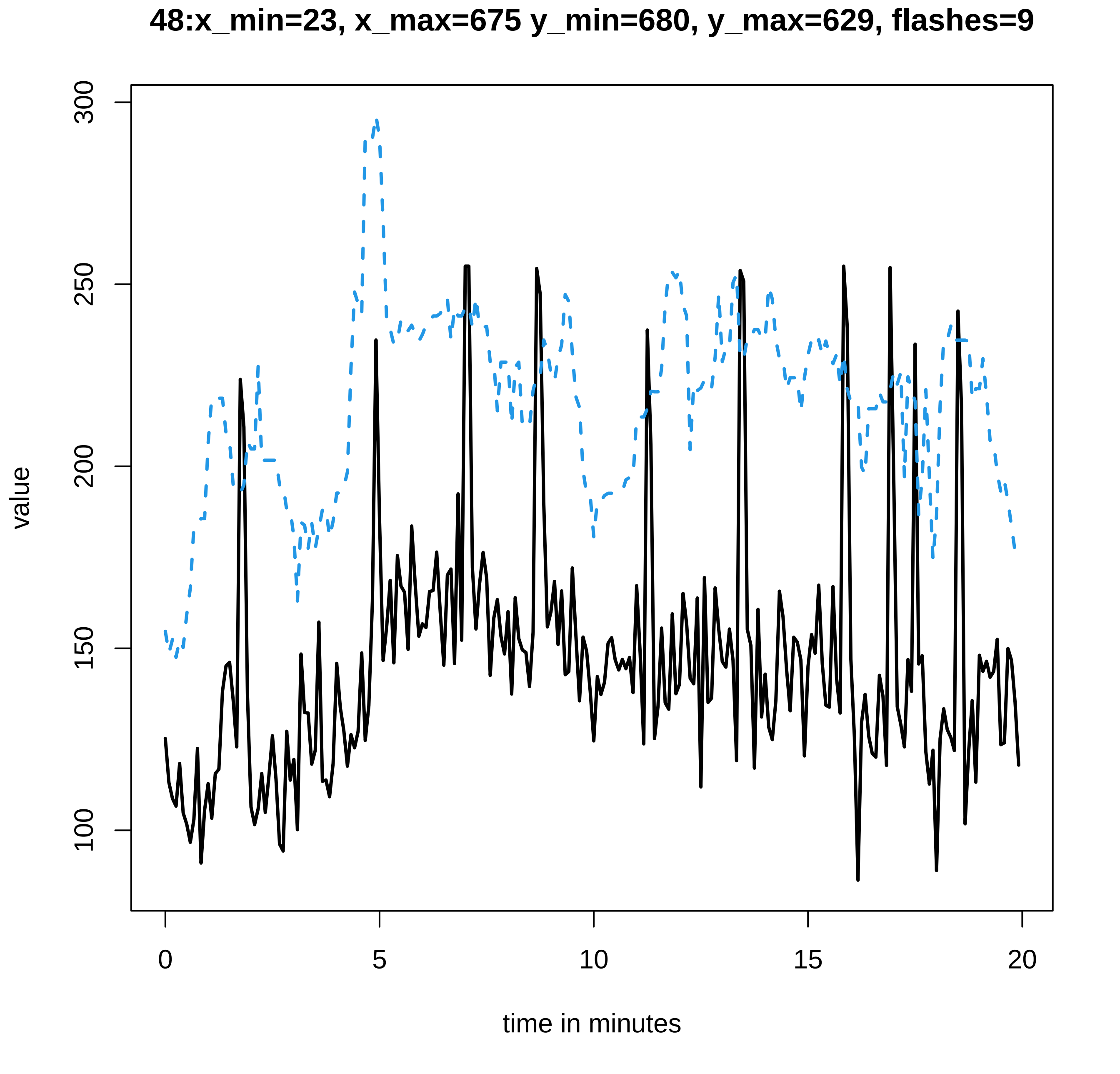
48:x_min=23, x_max=675 y_min=680, y_max=629, flashes=9
value
300
250
200
150
100
0
5
10
15
20
time in minutes

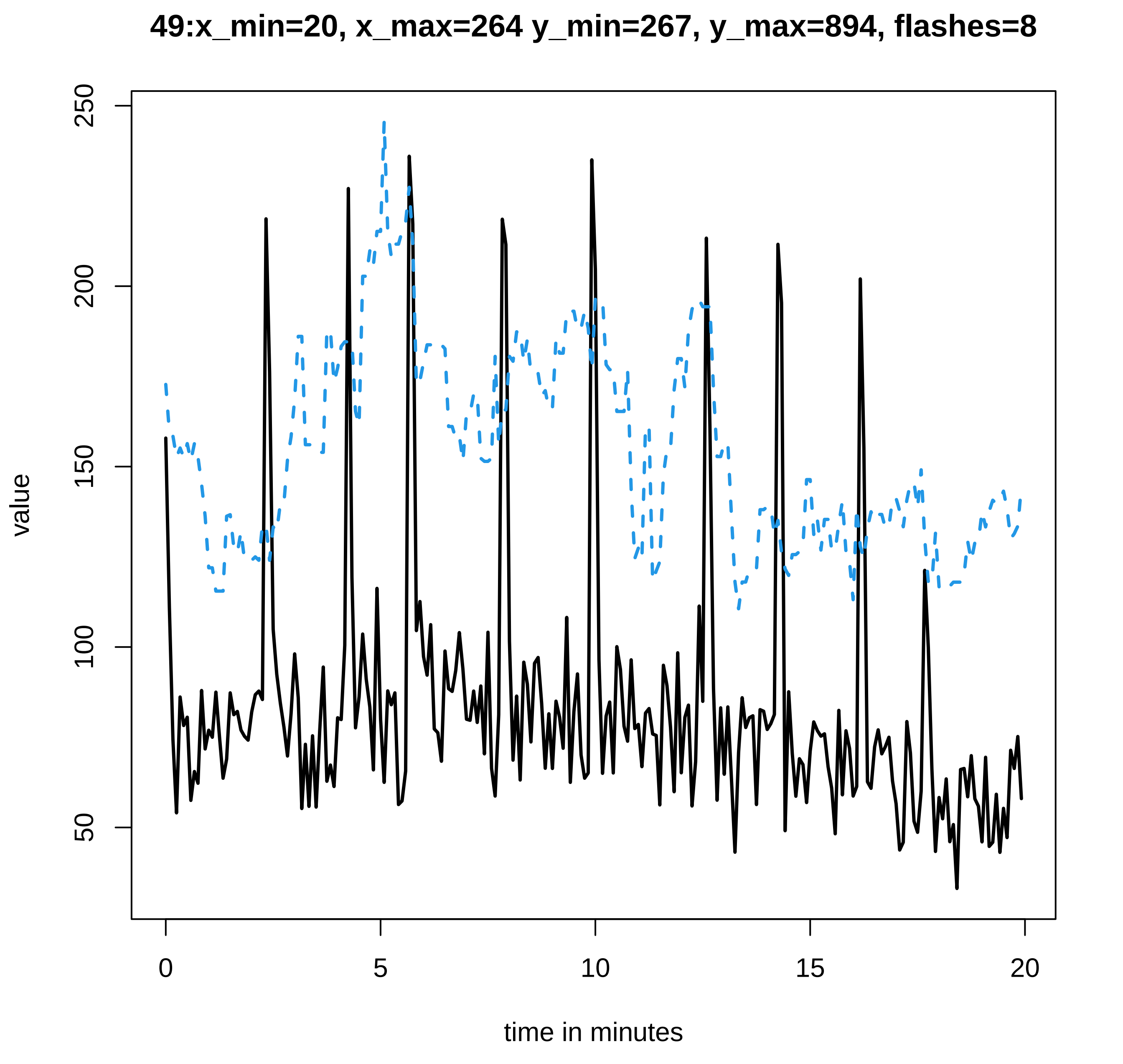

49:x_min=20, x_max=264 y_min=267, y_max=894, flashes=8
value
time in minutes
50
100
150
200
250
0
5
10
15
20

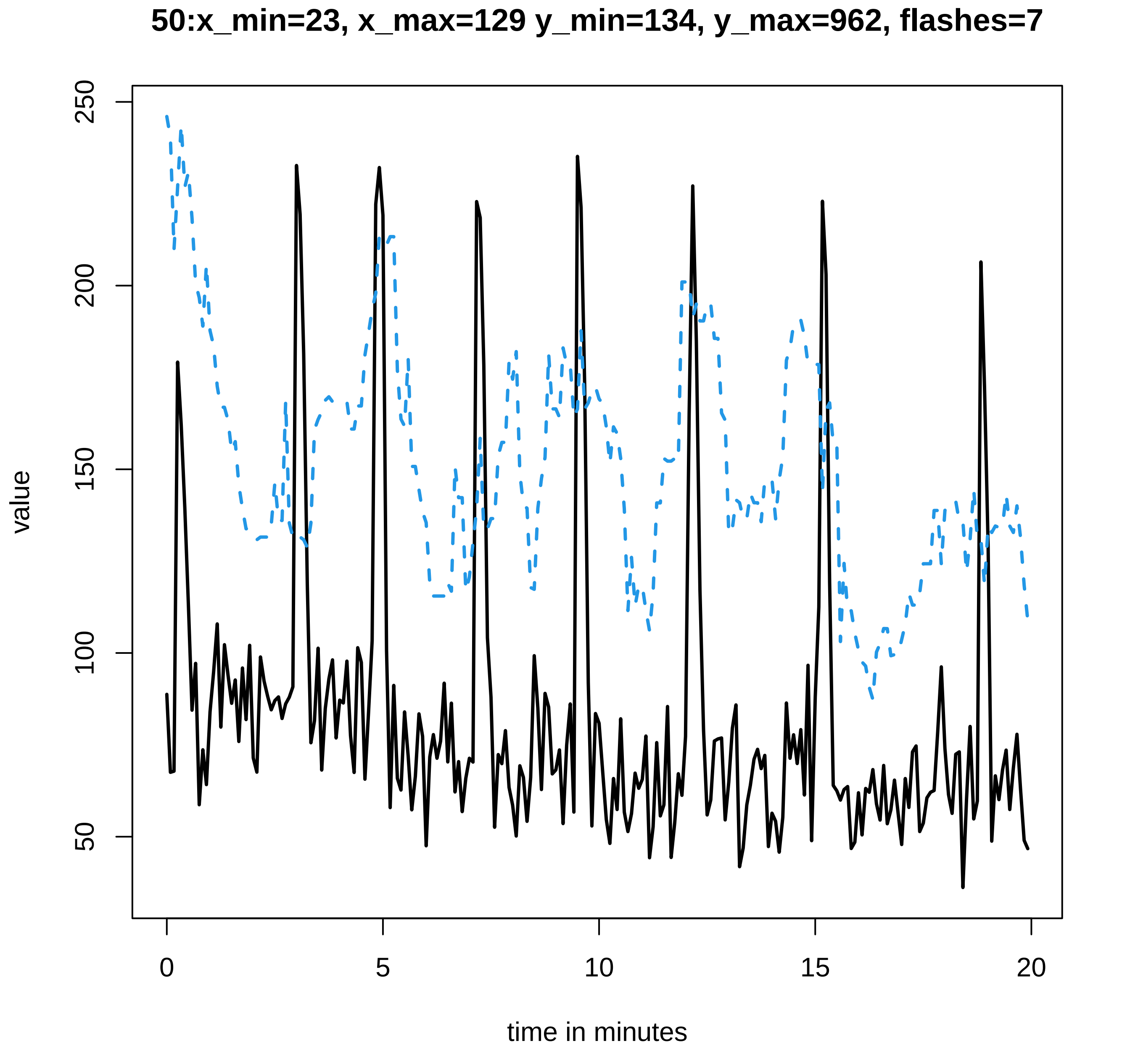

50:x_min=23, x_max=129 y_min=134, y_max=962, flashes=7
value
250
200
150
100
50
0
5
10
15
20
time in minutes

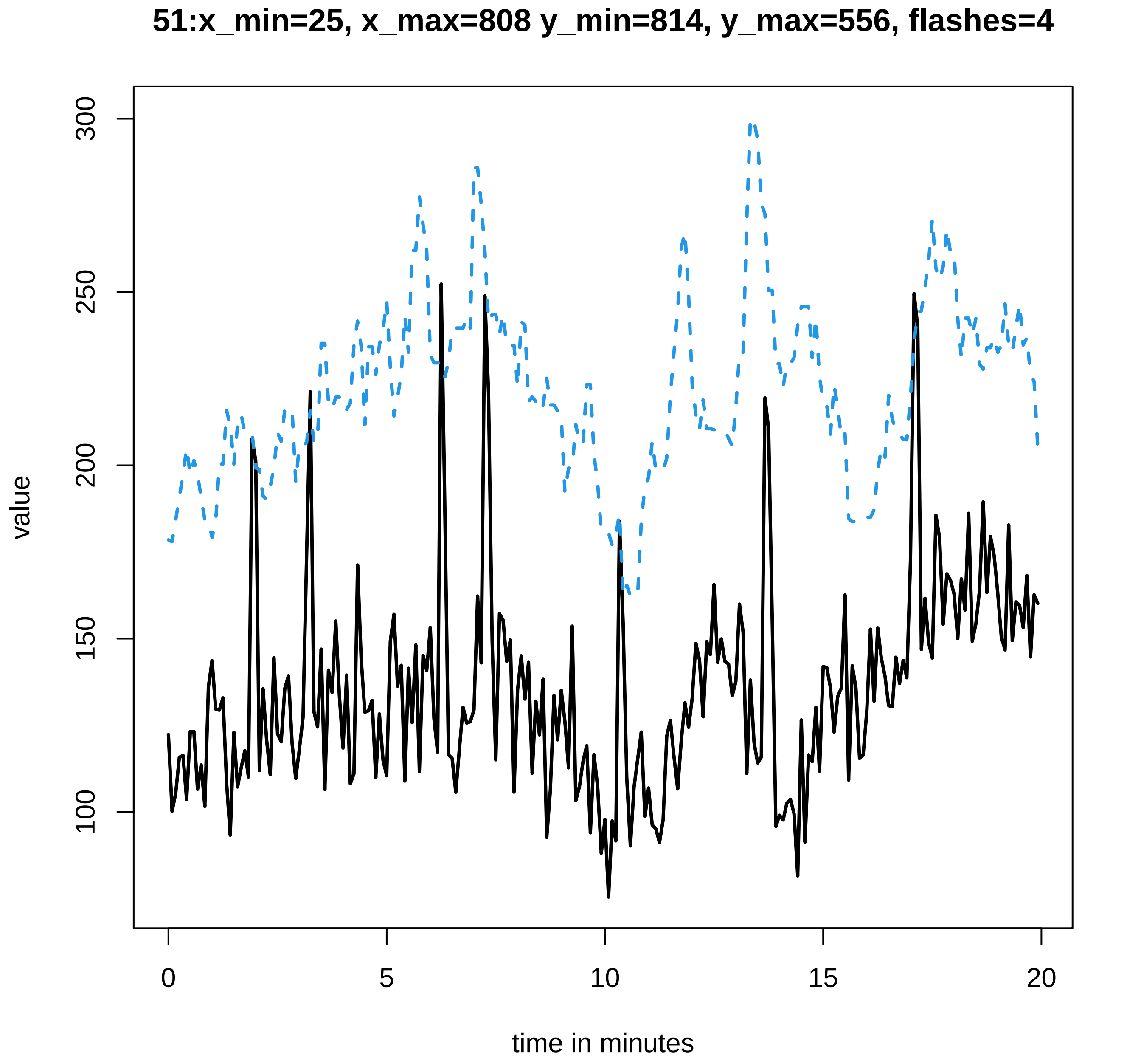
51:x_min=25, x_max=808 y_min=814, y_max=556, flashes=4
value
100
150
200
250
300
0
5
10
15
20
time in minutes

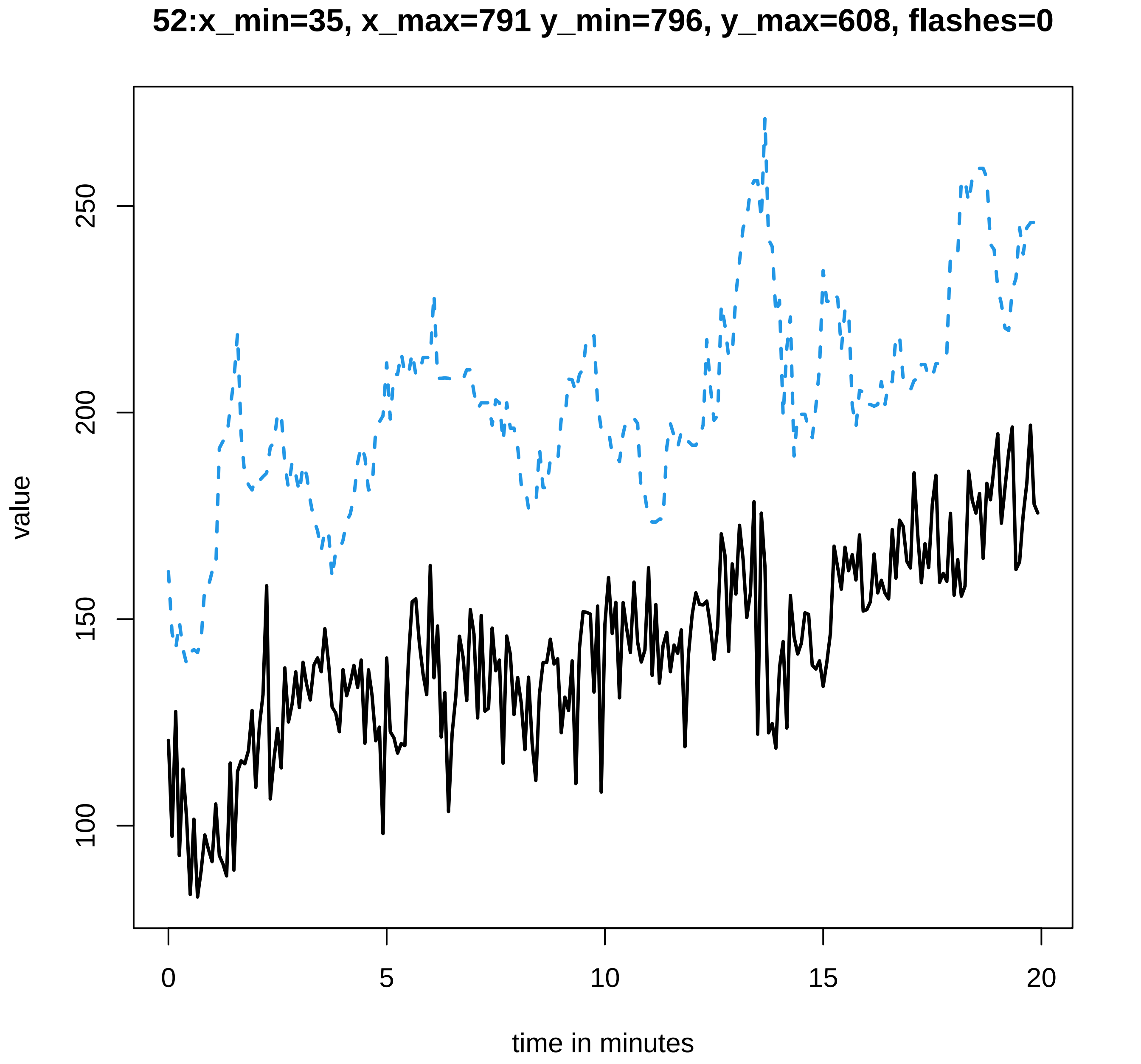
52:x_min=35, x_max=791 y_min=796, y_max=608, flashes=0
value
100
150
200
250
0
5
10
15
20
time in minutes

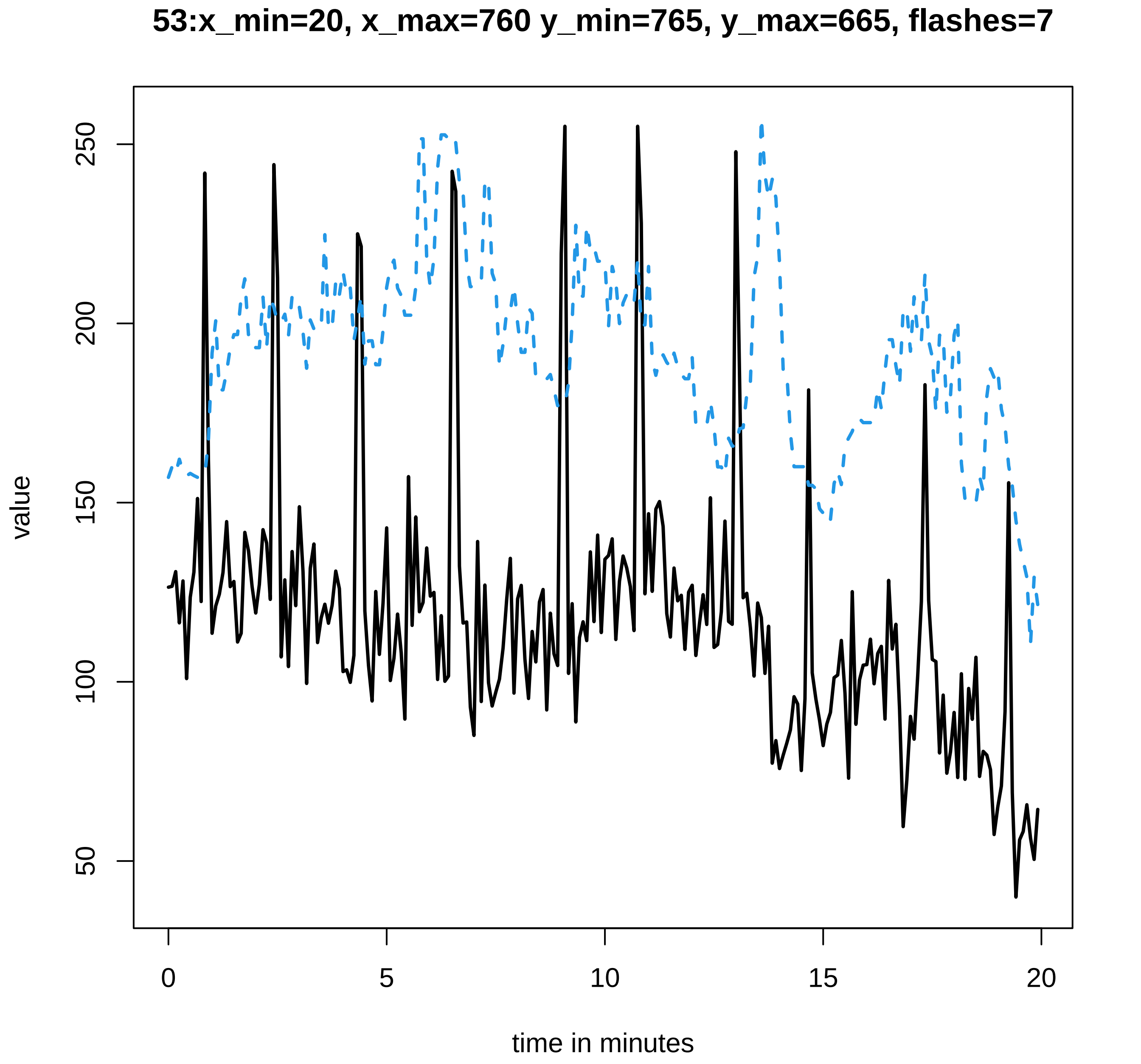

53:x_min=20, x_max=760 y_min=765, y_max=665, flashes=7
value
50
100
150
200
250
0
5
10
15
20
time in minutes

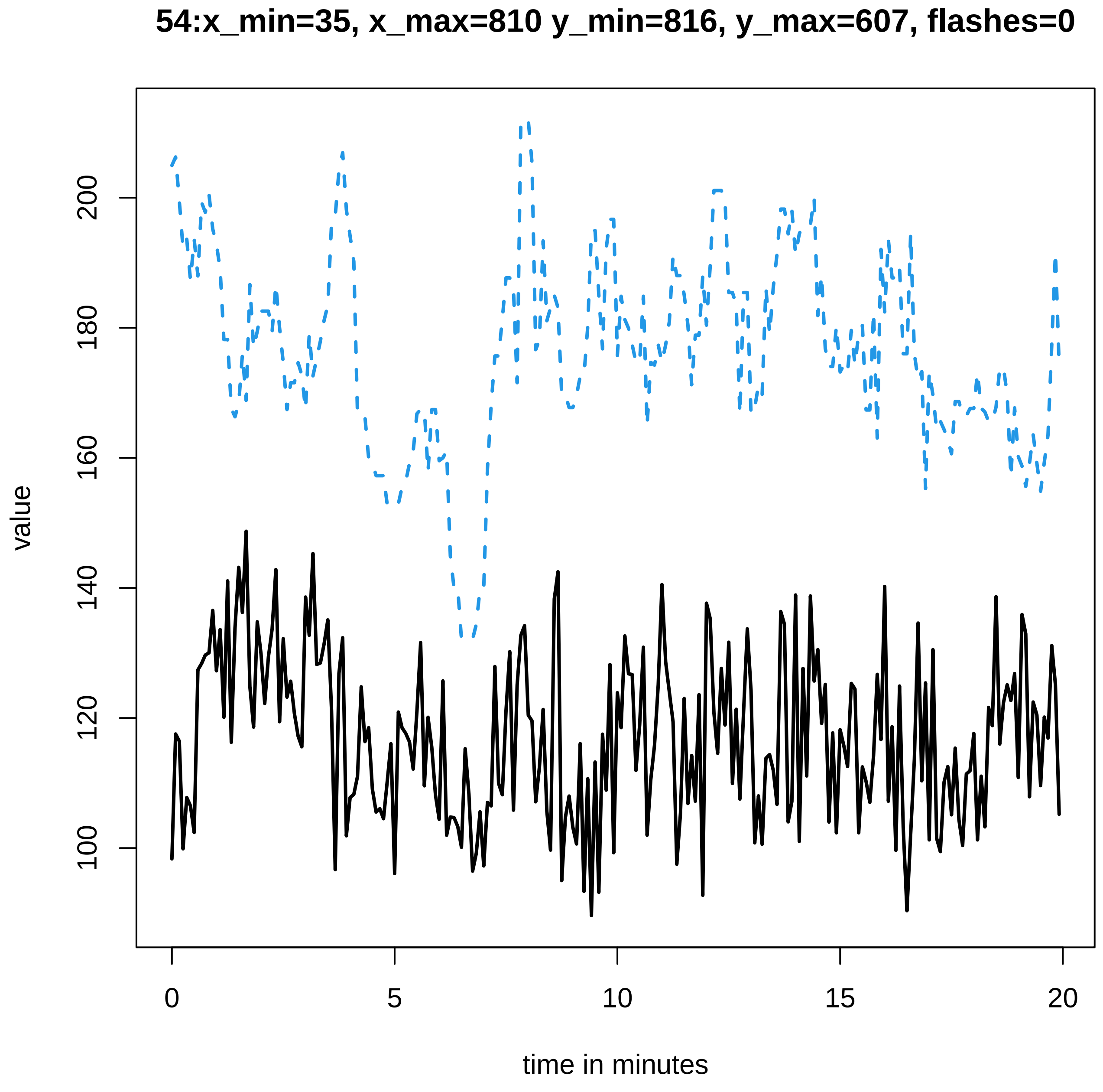
54:x_min=35, x_max=810 y_min=816, y_max=607, flashes=0
value
100
120
140
160
180
200
0
5
10
15
20
time in minutes

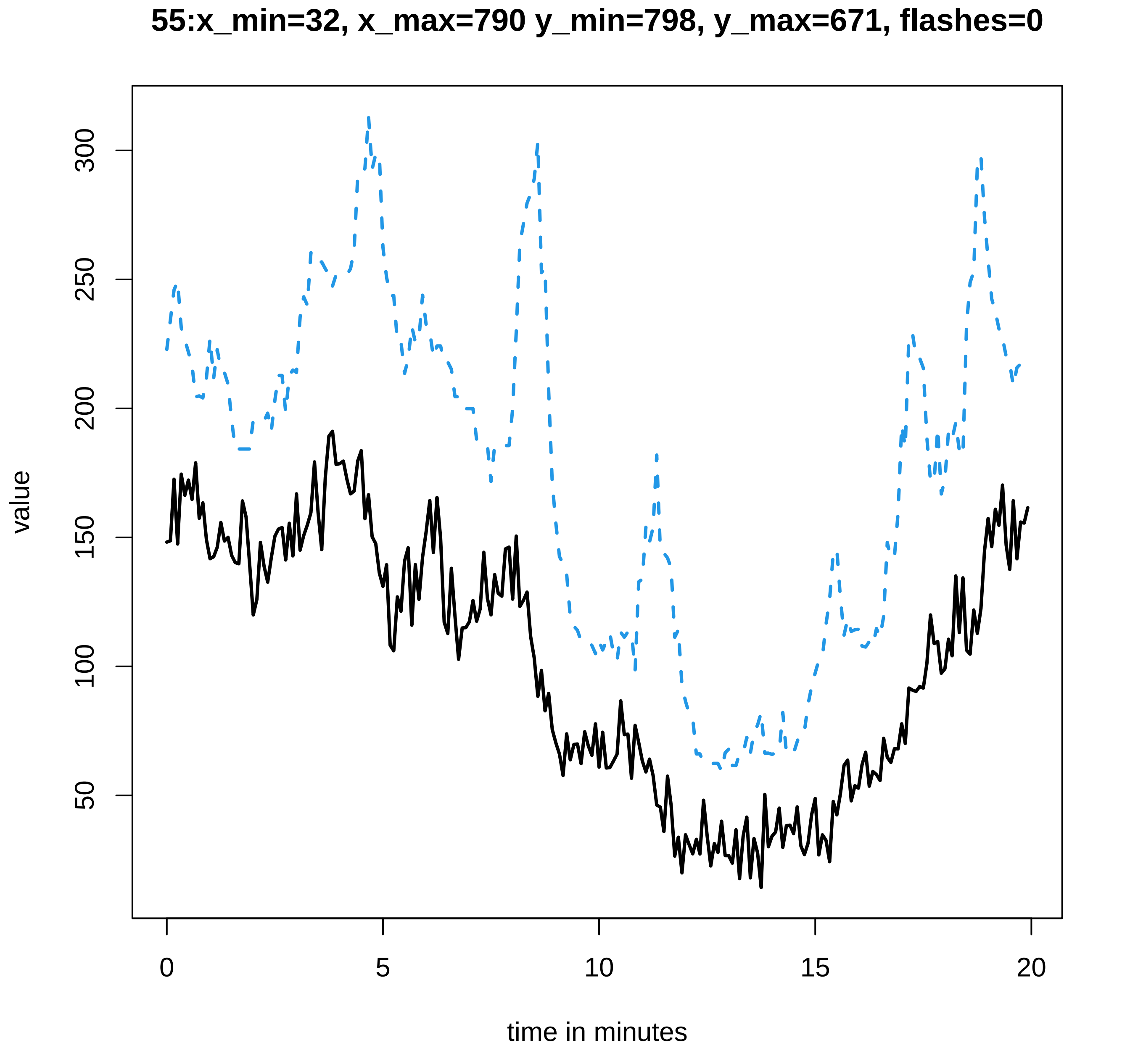

55:x_min=32, x_max=790 y_min=798, y_max=671, flashes=0
value
300
250
200
150
100
50
0
5
10
15
20
time in minutes

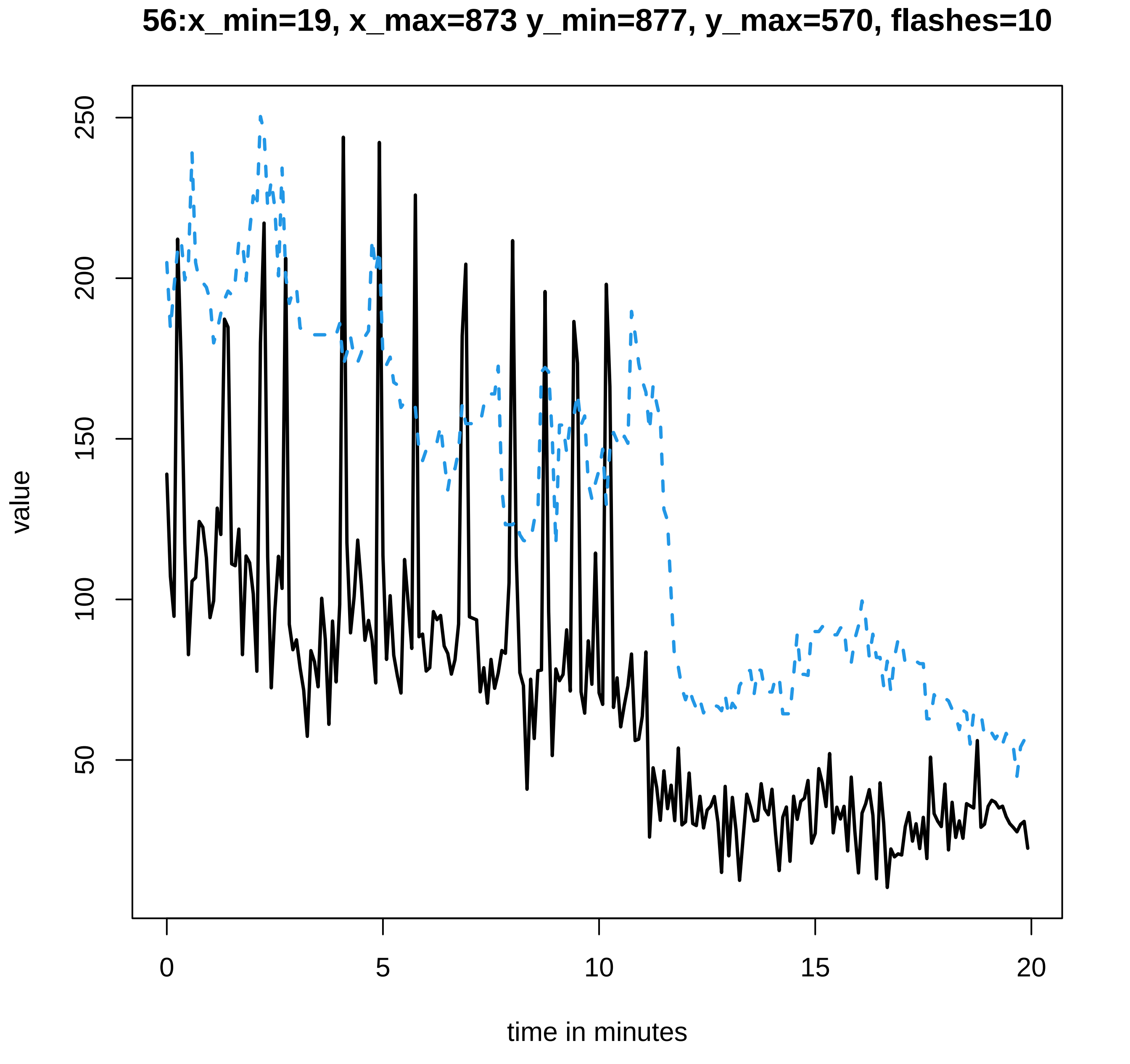

56:x_min=19, x_max=873 y_min=877, y_max=570, flashes=10
value
250
200
150
100
50
0
5
10
15
20
time in minutes

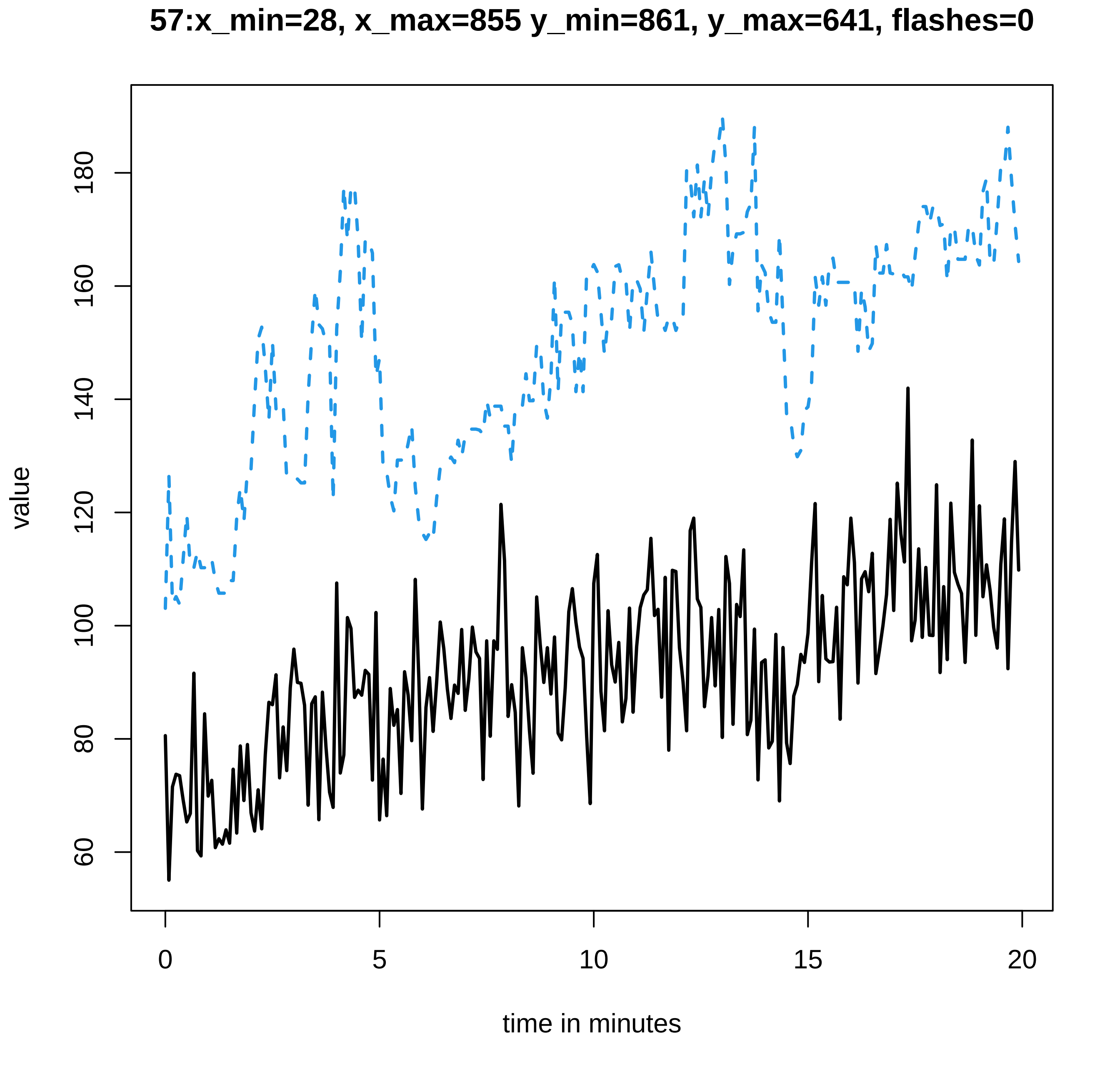
57:x_min=28, x_max=855 y_min=861, y_max=641, flashes=0
value
60
80
100
120
140
160
180
0
5
10
15
20
time in minutes

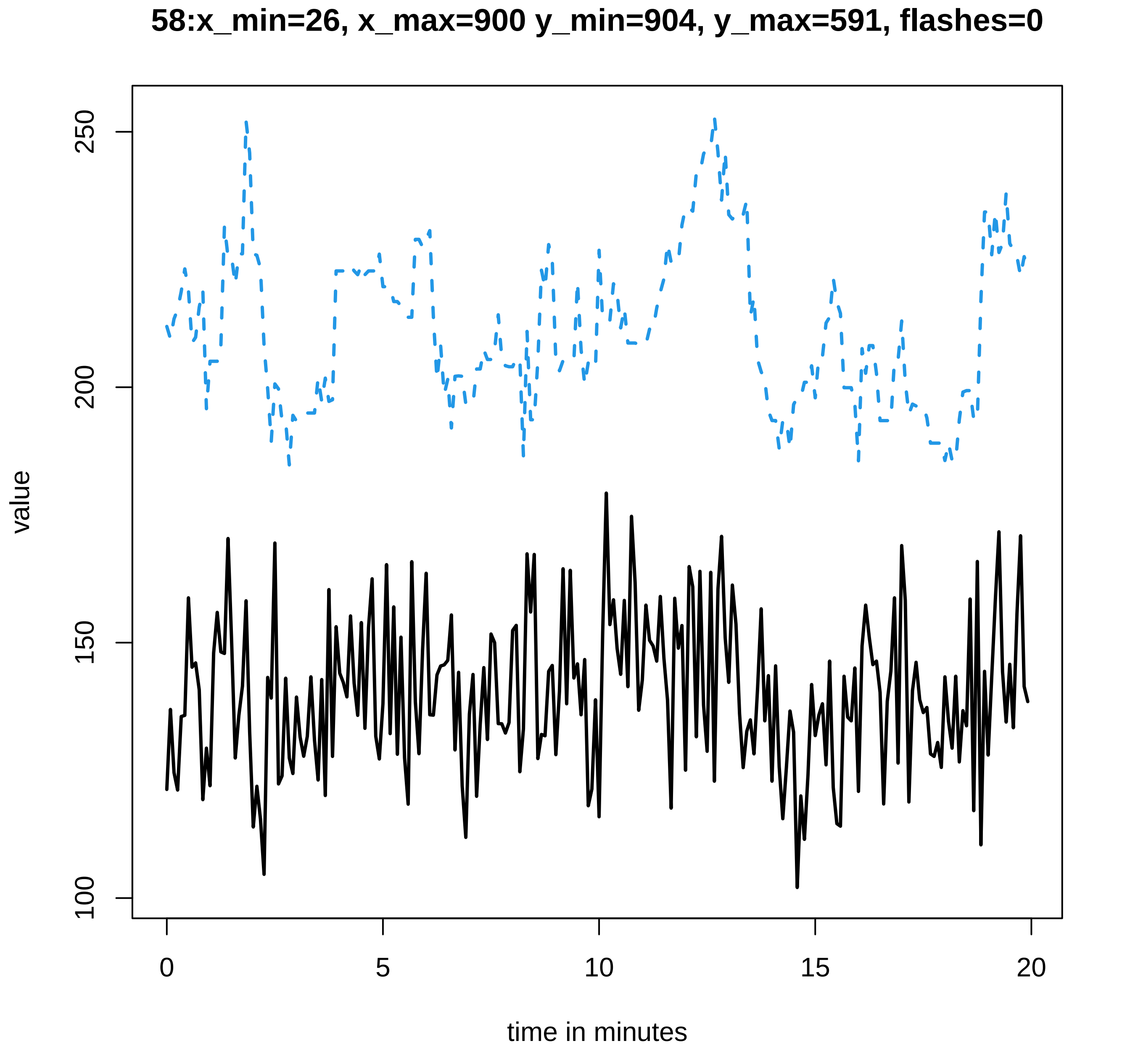
58:x_min=26, x_max=900 y_min=904, y_max=591, flashes=0
value
100
150
200
250
0
5
10
15
20
time in minutes

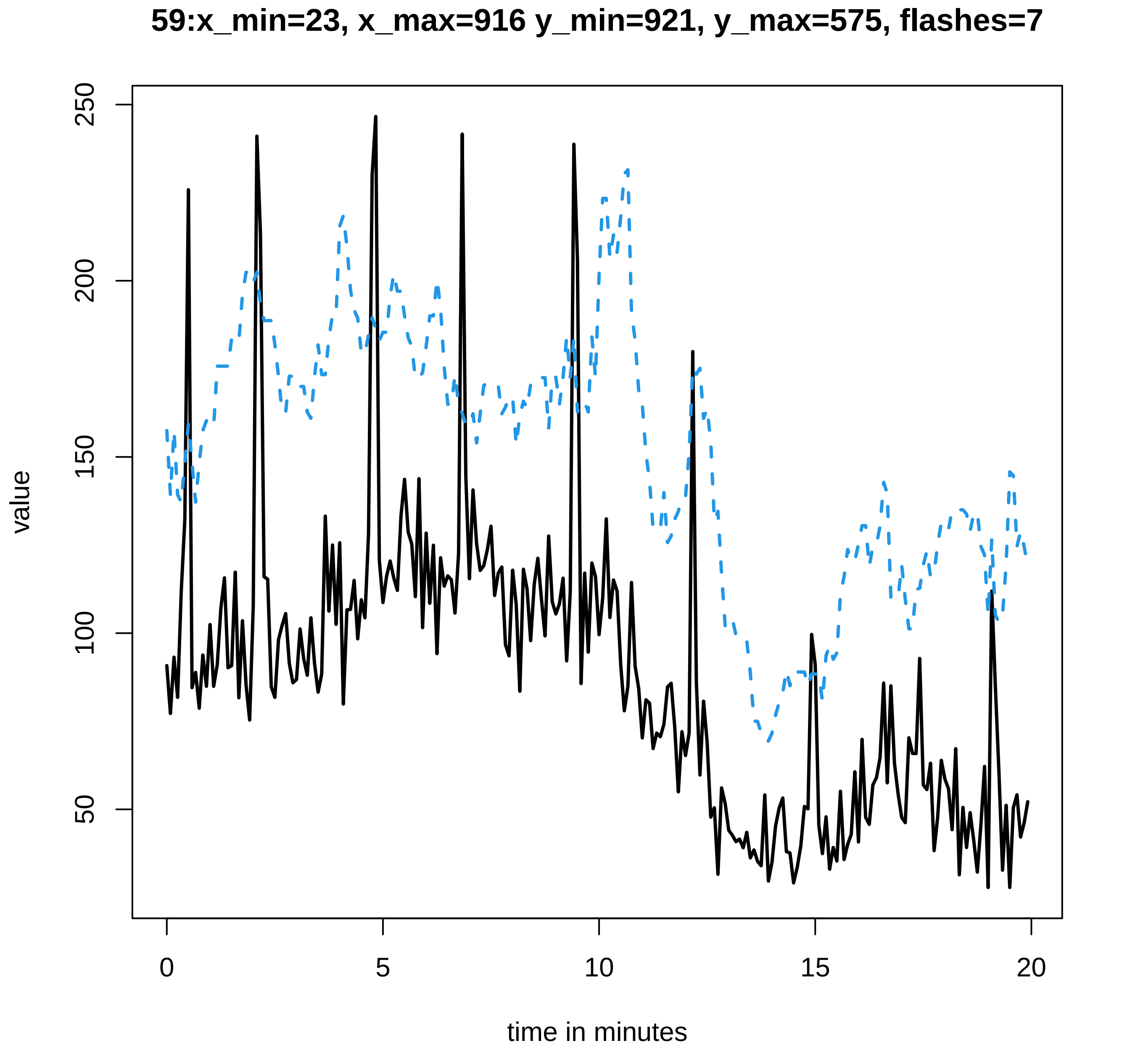
59:x_min=23, x_max=916 y_min=921, y_max=575, flashes=7
value
250
200
150
100
50
0
5
10
15
20
time in minutes

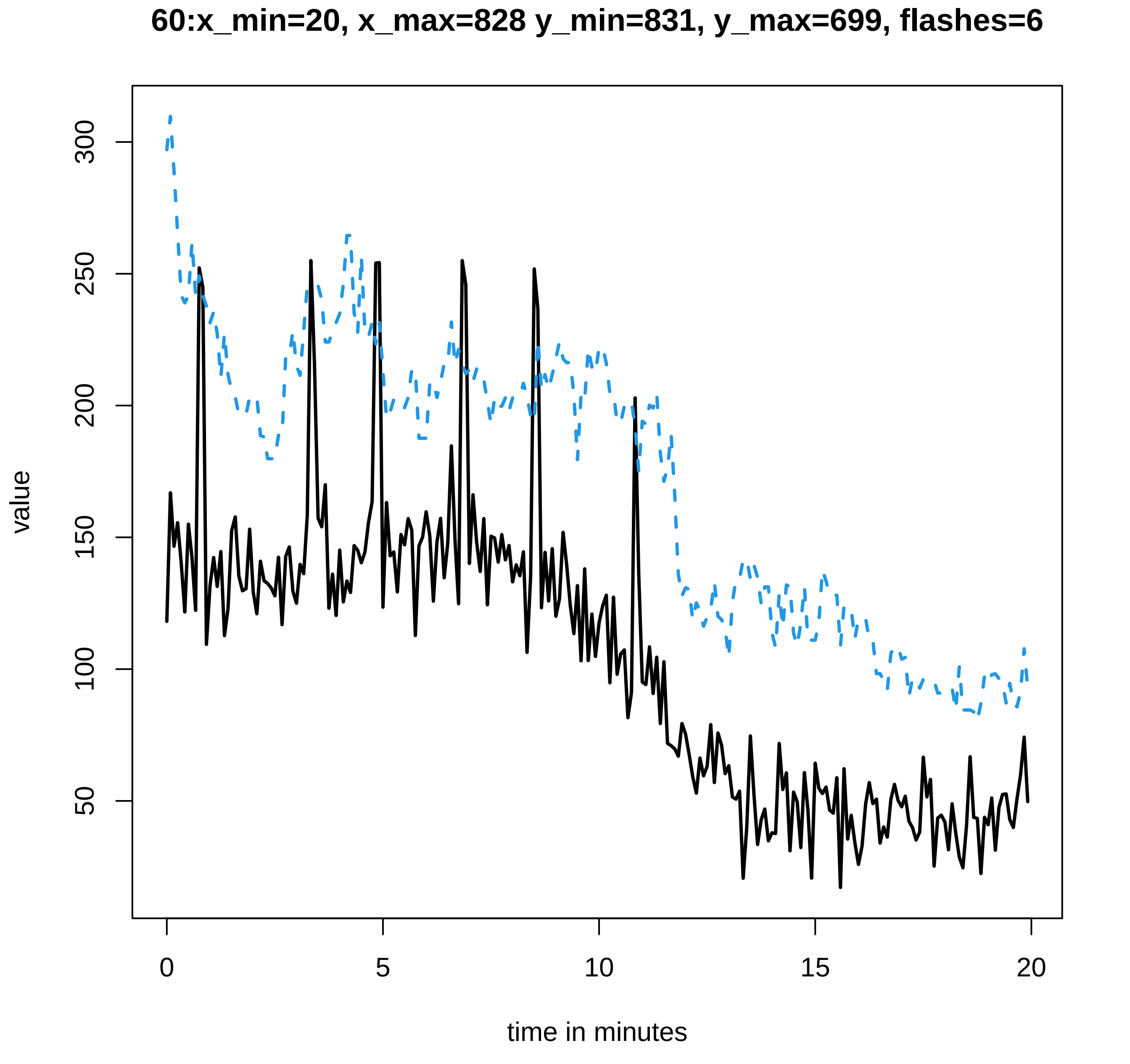
60:x_min=20, x_max=828 y_min=831, y_max=699, flashes=6
value
300
250
200
150
100
50
0
5
10
15
20
time in minutes

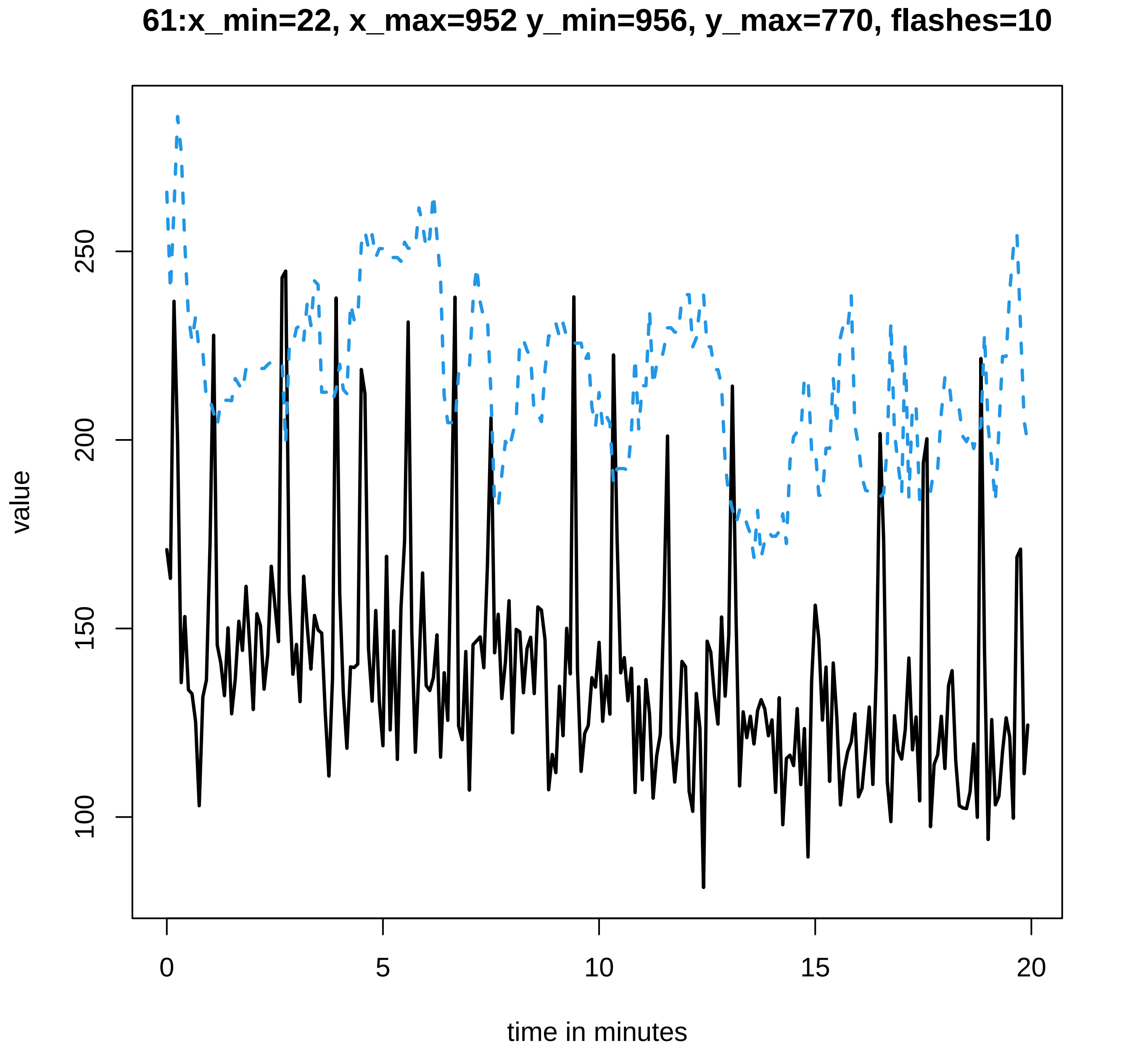
61:x_min=22, x_max=952 y_min=956, y_max=770, flashes=10
value
100
150
200
250
0
5
10
15
20
time in minutes

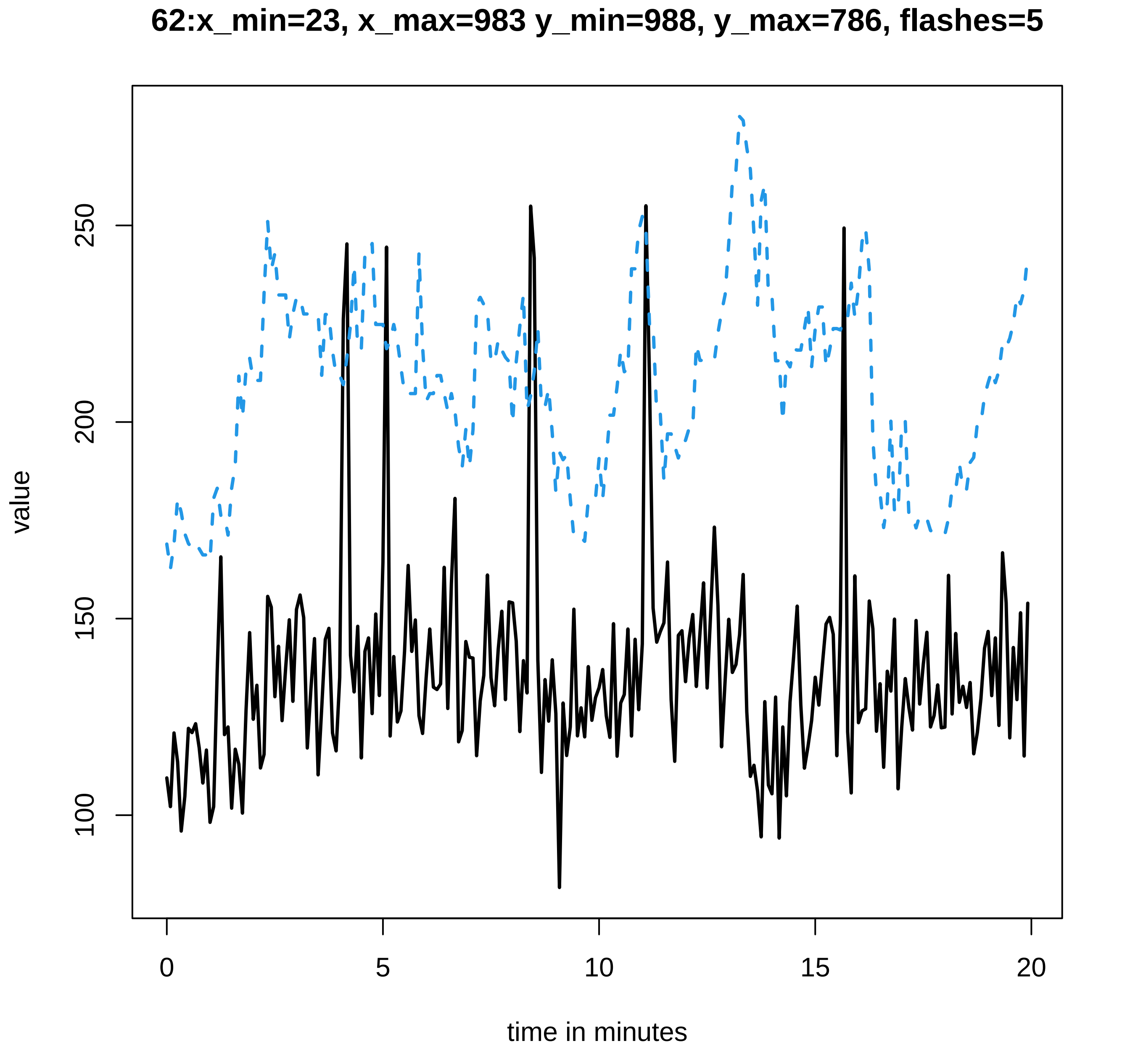
62:x_min=23, x_max=983 y_min=988, y_max=786, flashes=5
value
100
150
200
250
0
5
10
15
20
time in minutes
