## Supplementary data Calcium spiking for "LysM Receptor Proteins are Required for Ectomycorrhizal Symbiosis in Poplar": profiles-spiking-4.pdf

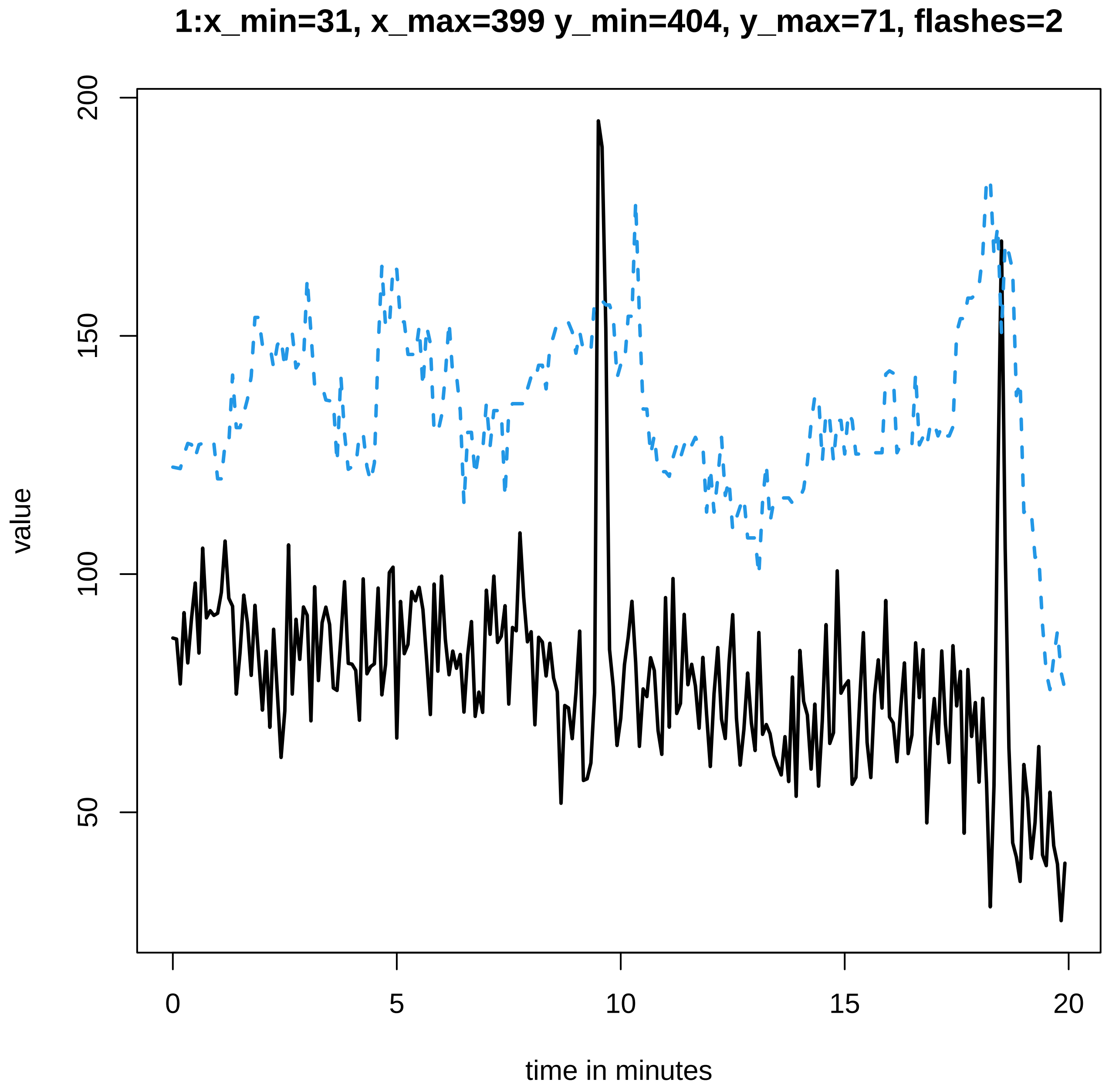

1:x_min=31, x_max=399 y_min=404, y_max=71, flashes=2
value
200
150
100
50
0
5
10
15
20
time in minutes

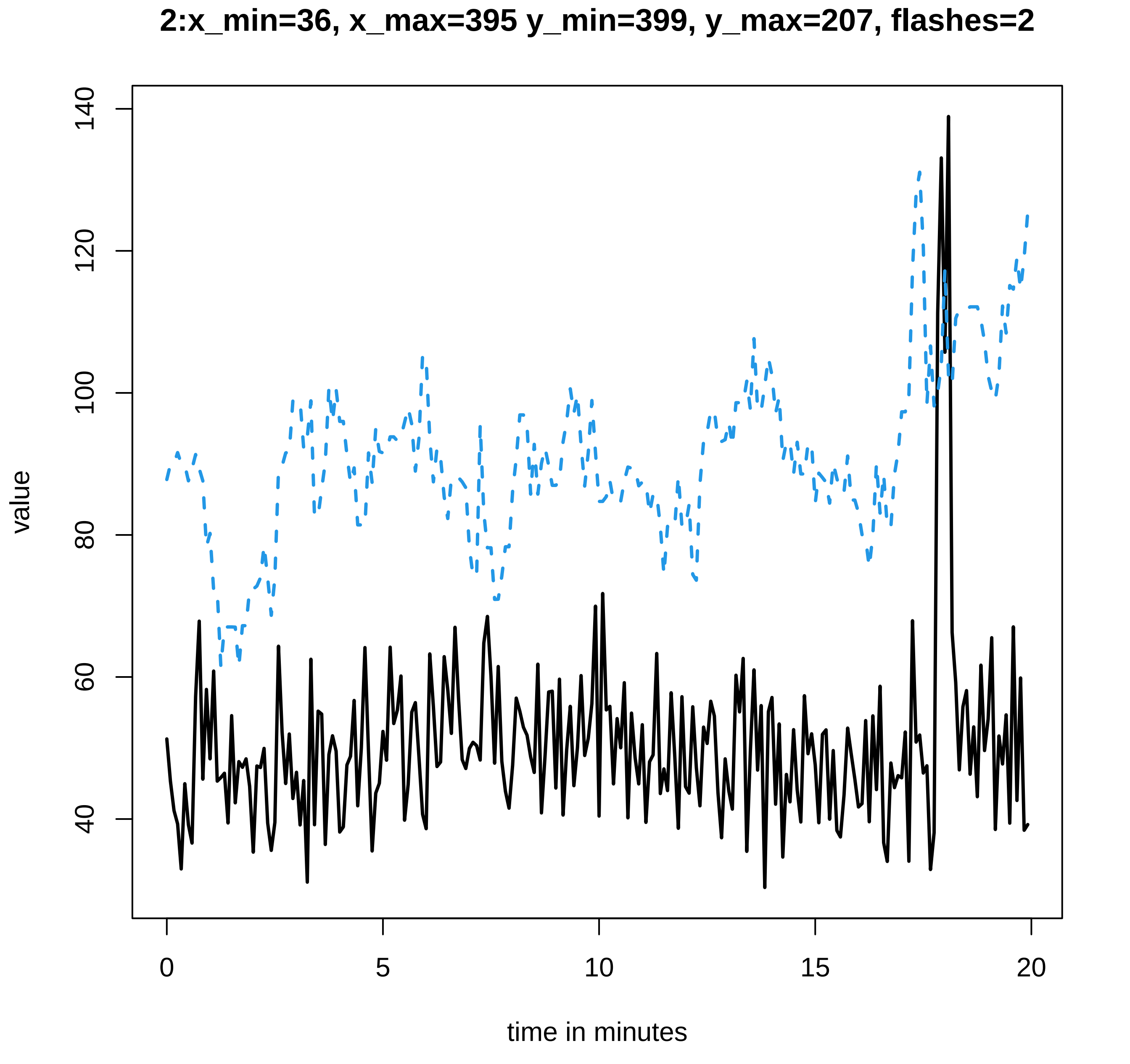
2:x_min=36, x_max=395 y_min=399, y_max=207, flashes=2
value
140
120
100
80
60
40
0
5
10
15
20
time in minutes

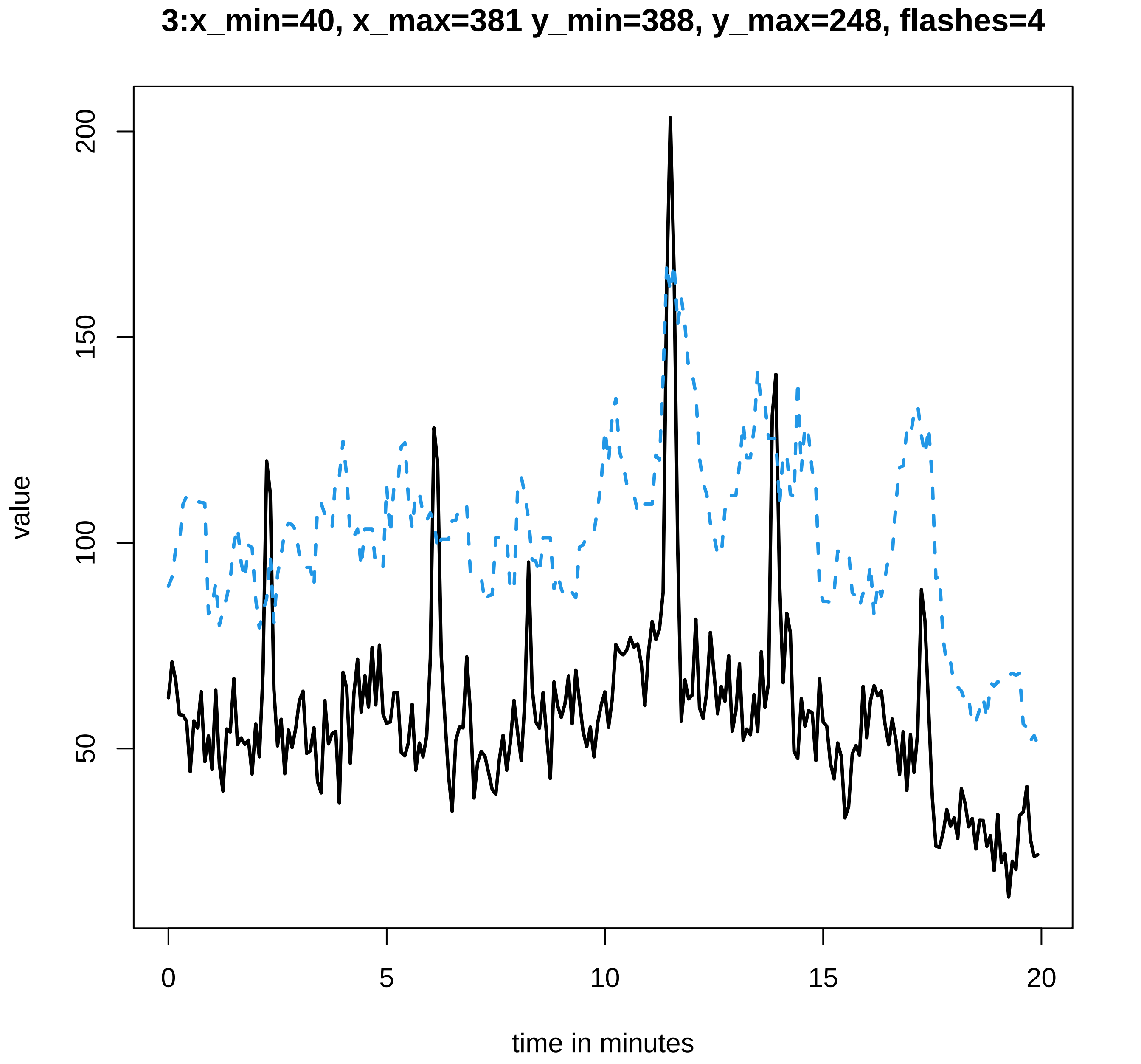
3:x_min=40, x_max=381 y_min=388, y_max=248, flashes=4
value
200
150
100
50
0
5
10
15
20
time in minutes

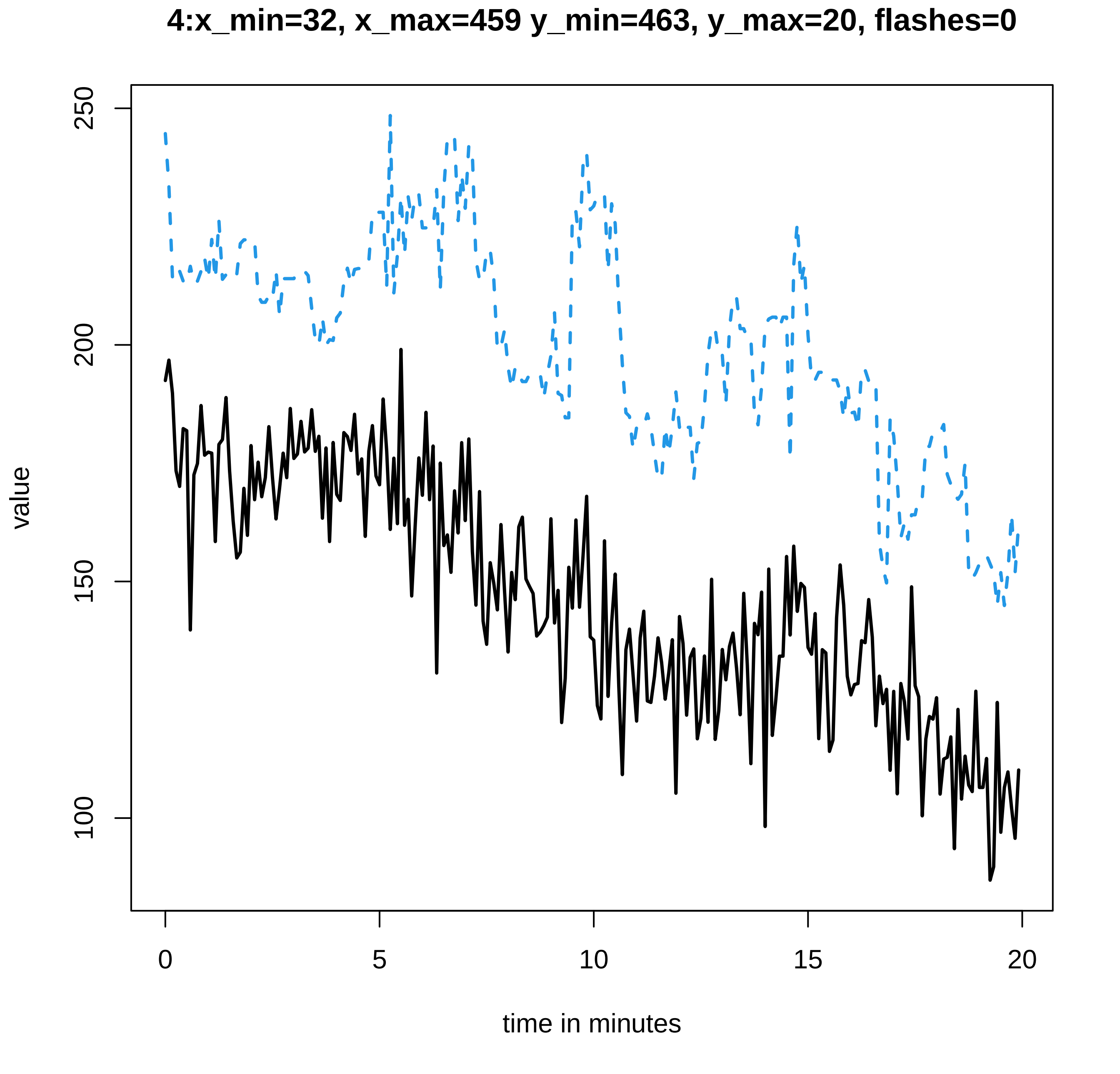
4:x_min=32, x_max=459 y_min=463, y_max=20, flashes=0
value
250
200
150
100
0
5
10
15
20
time in minutes

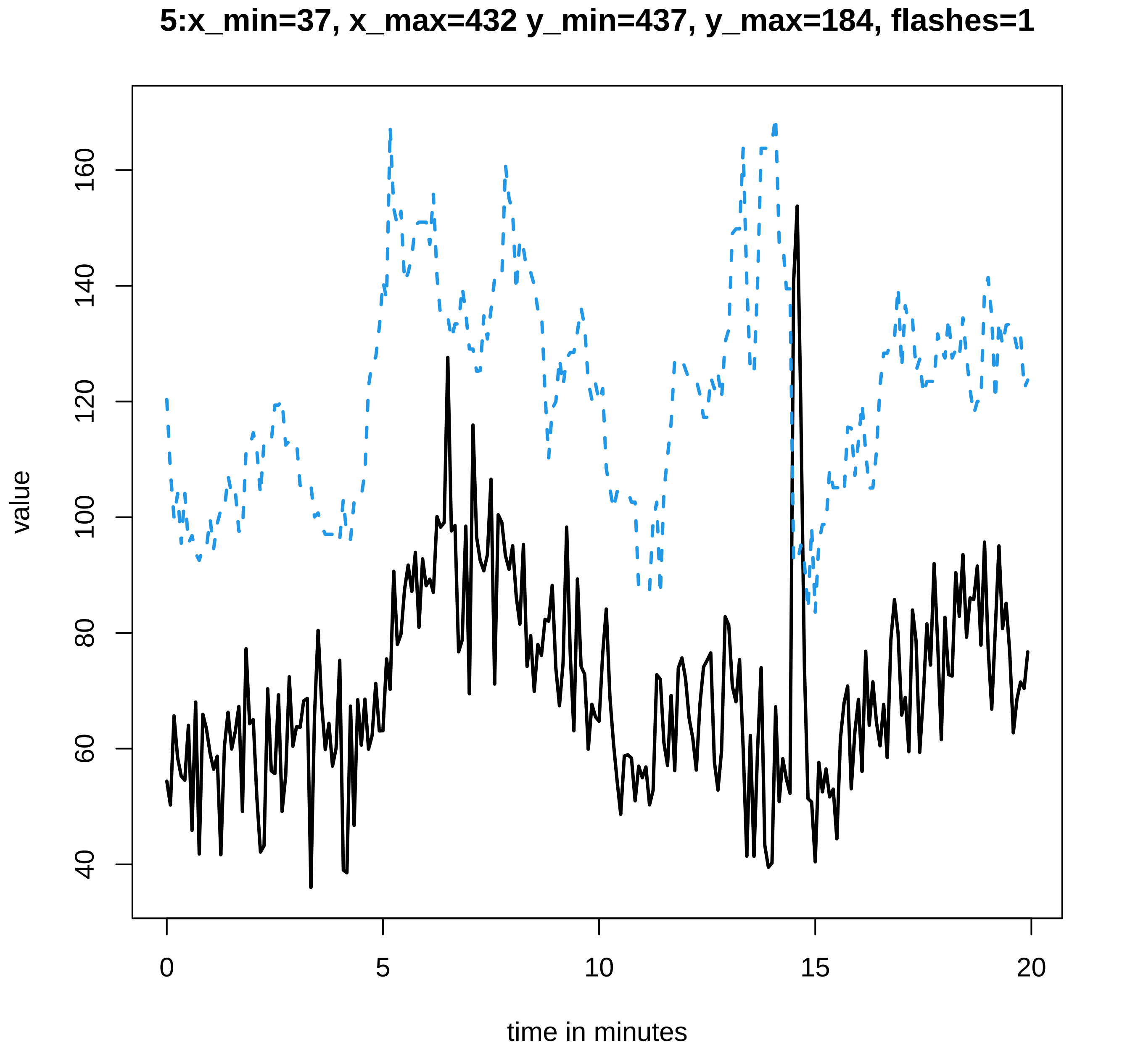

5:x_min=37, x_max=432 y_min=437, y_max=184, flashes=1
value
40
60
80
100
120
140
160
0
5
10
15
20
time in minutes

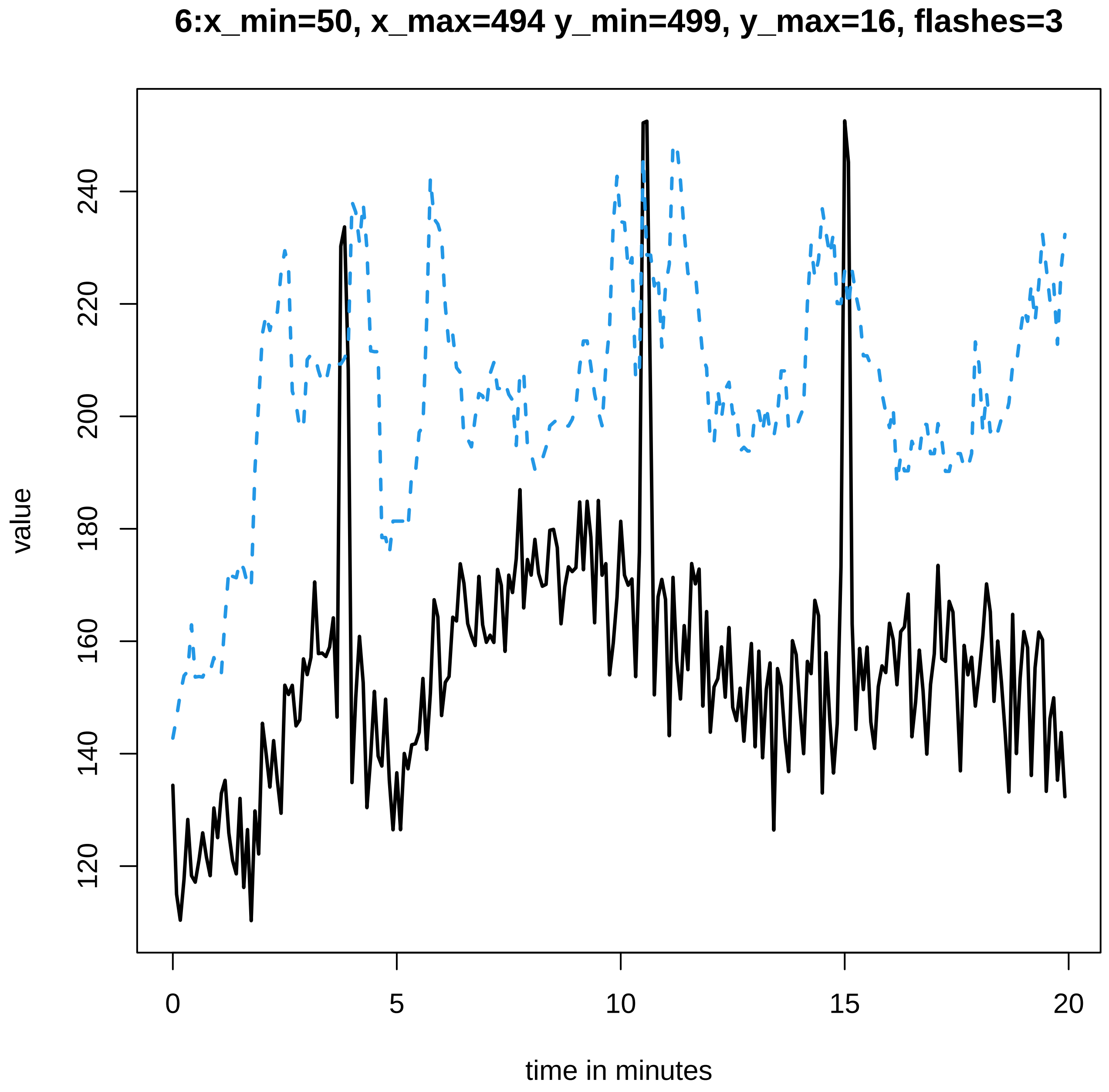

6:x_min=50, x_max=494 y_min=499, y_max=16, flashes=3
value
120
140
160
180
200
220
240
0
5
10
15
20
time in minutes

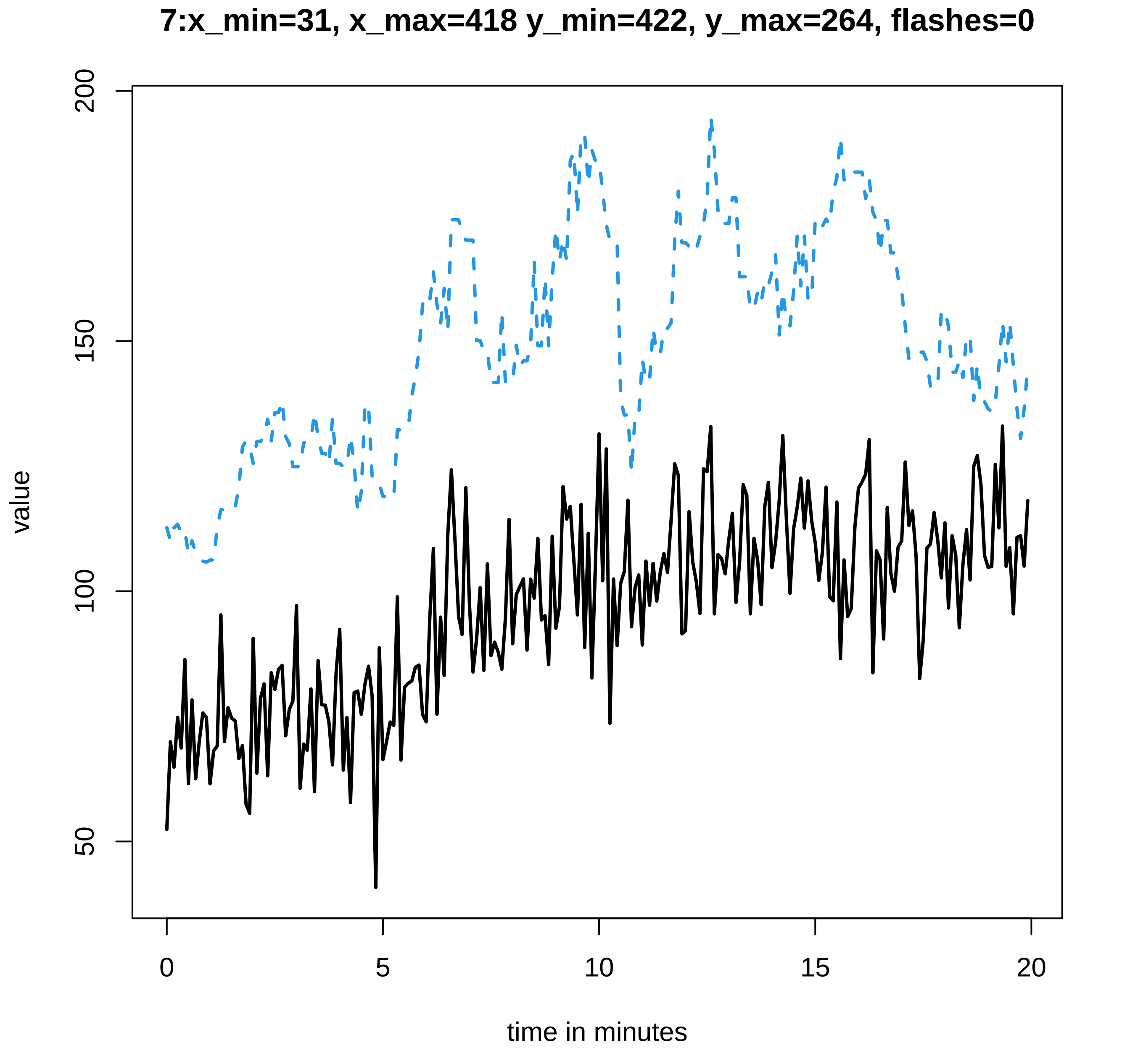

7:x_min=31, x_max=418 y_min=422, y_max=264, flashes=0
value
200
150
100
50
0
5
10
15
20
time in minutes

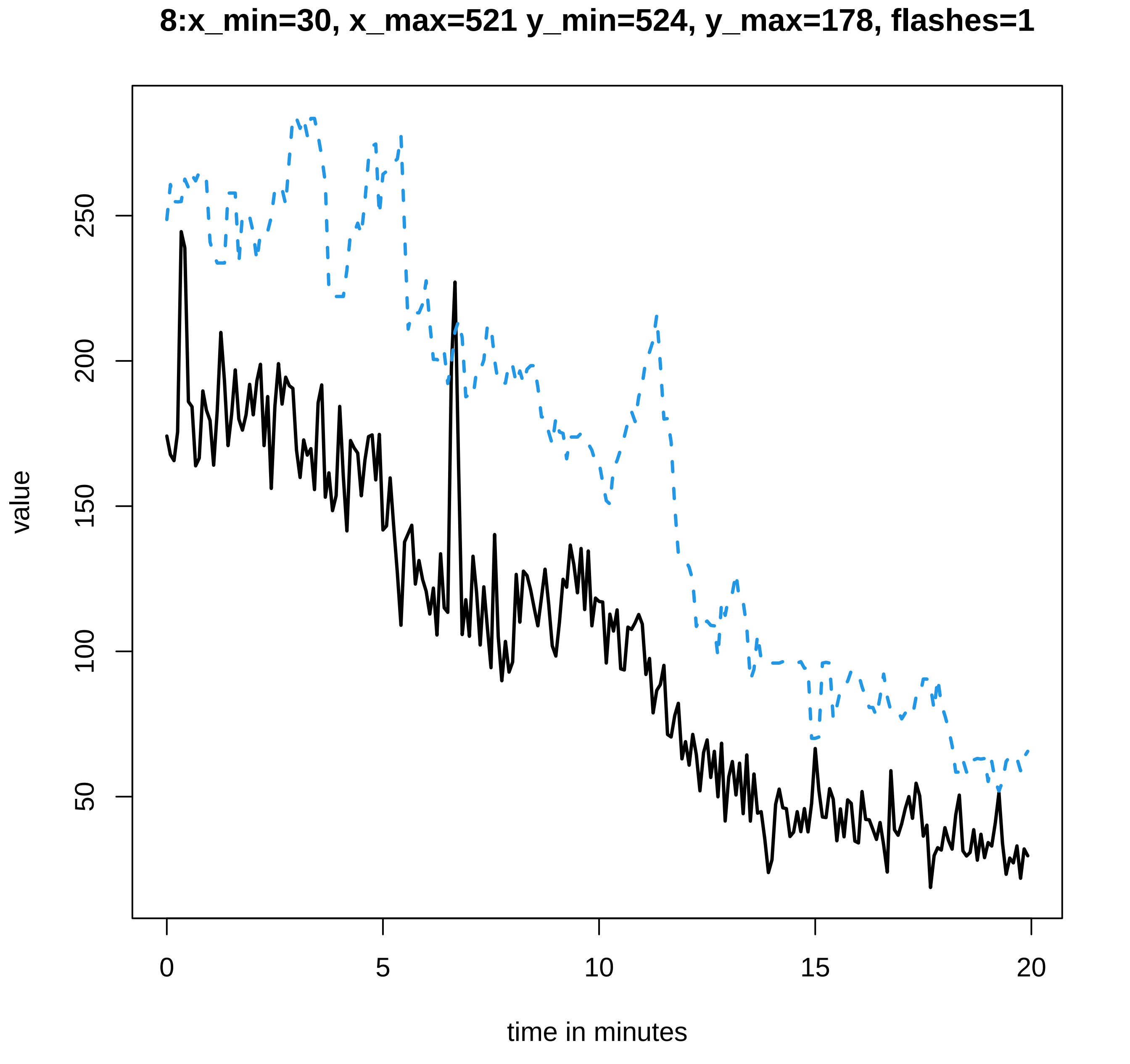

8:x_min=30, x_max=521 y_min=524, y_max=178, flashes=1
value
250
200
150
100
50
0
5
10
15
20
time in minutes

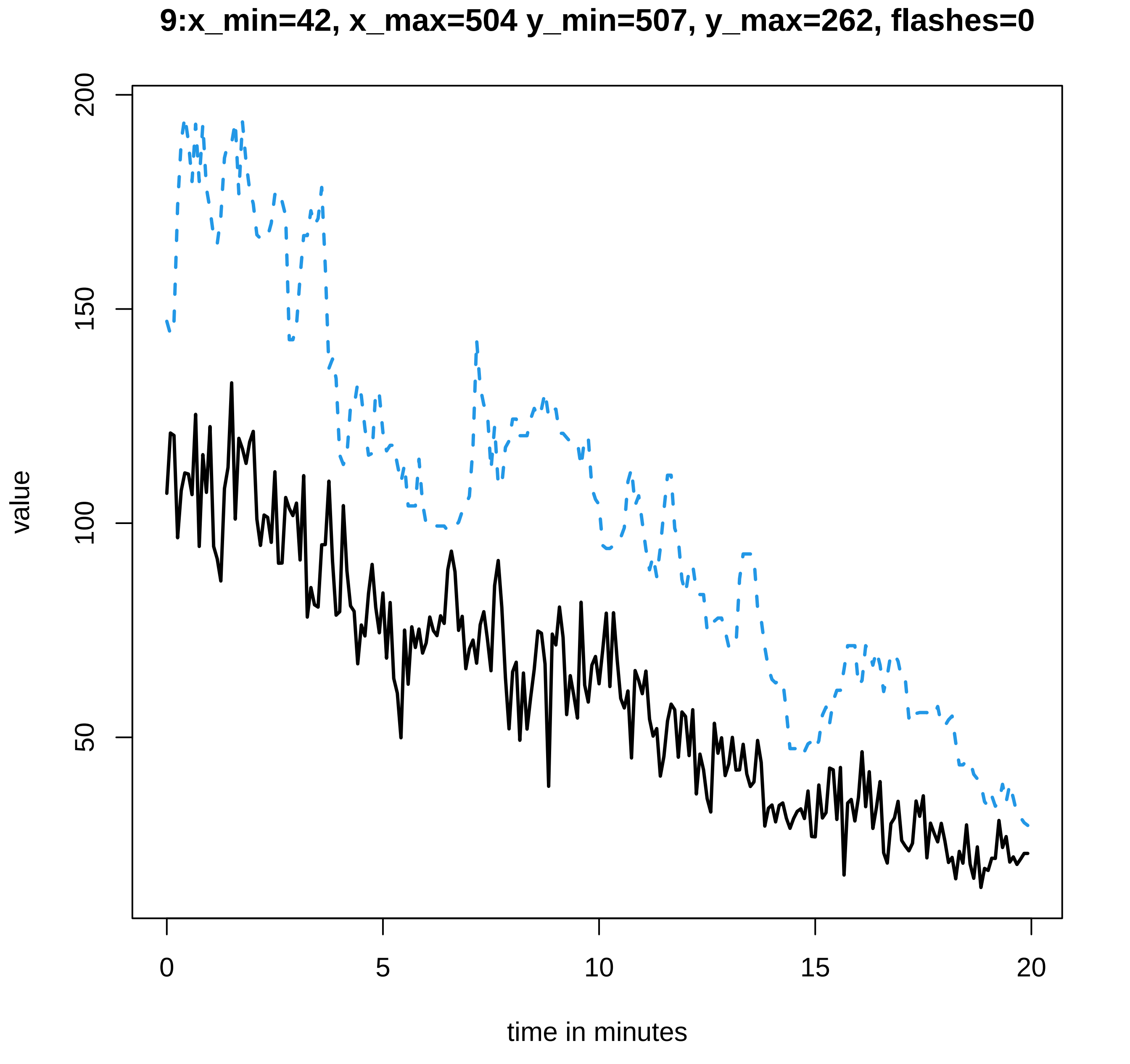

9:x_min=42, x_max=504 y_min=507, y_max=262, flashes=0
value
200
150
100
50
0
5
10
15
20
time in minutes

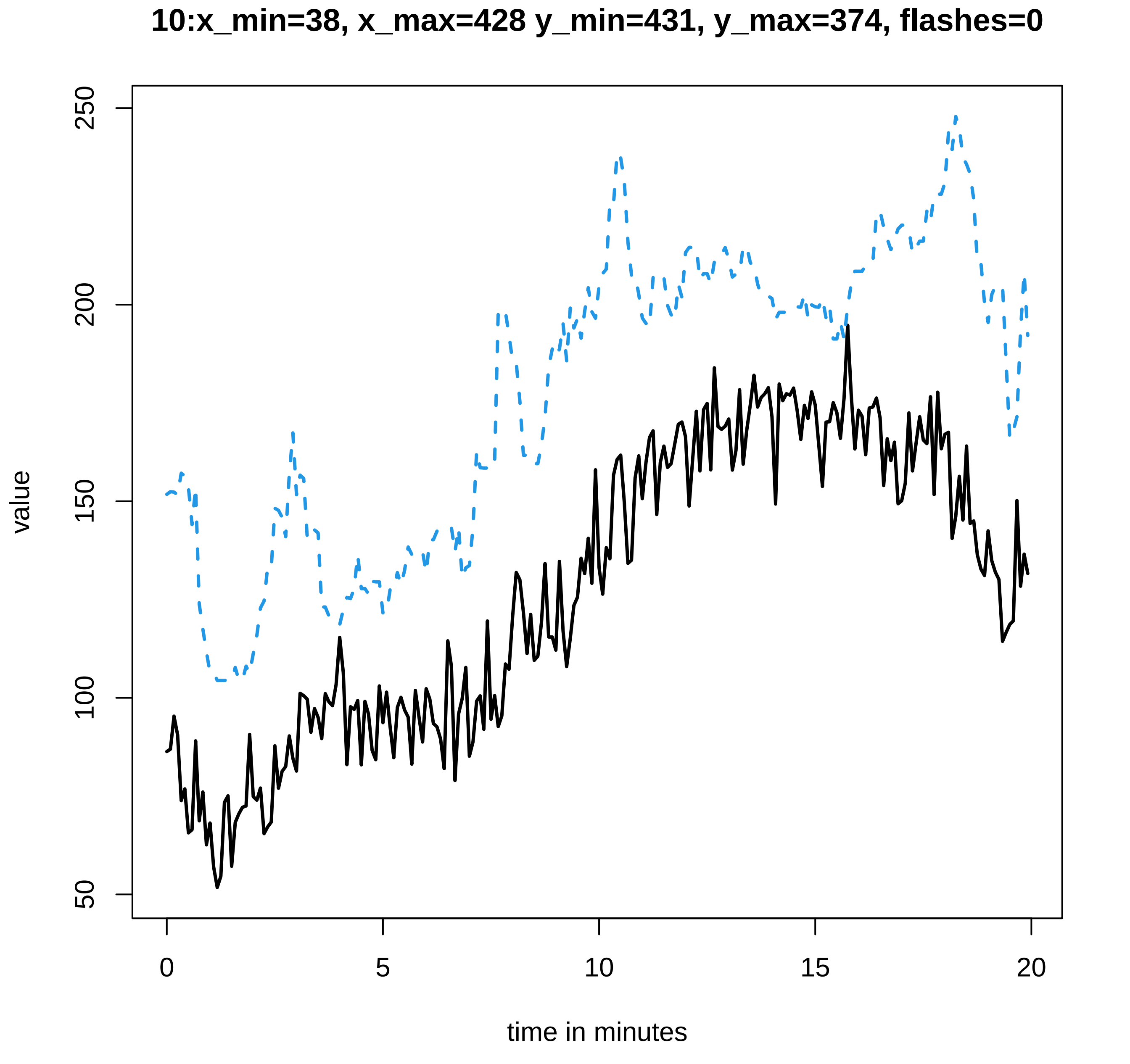
10:x_min=38, x_max=428 y_min=431, y_max=374, flashes=0
value
50
100
150
200
250
0
5
10
15
20
time in minutes

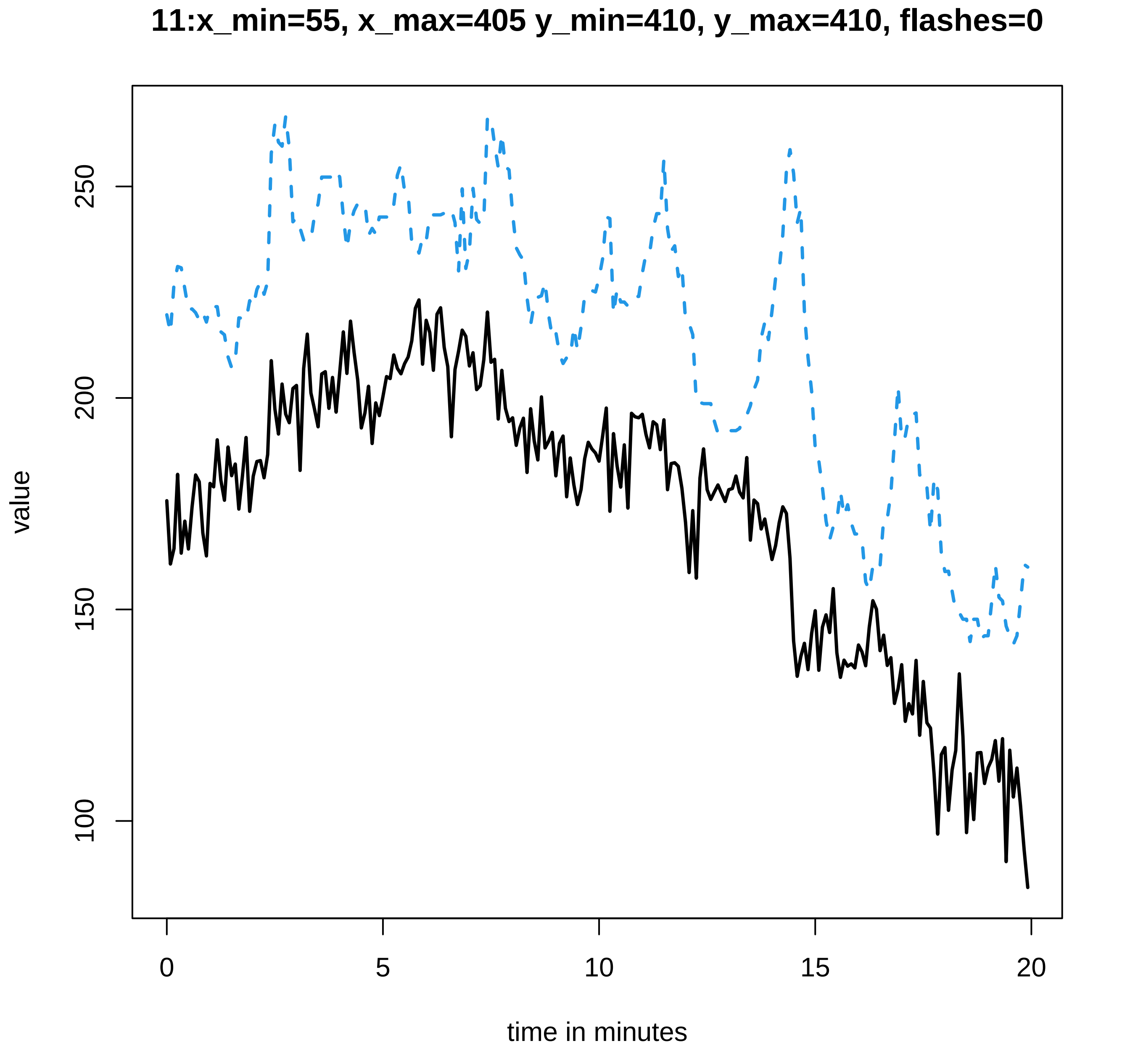
11:x_min=55, x_max=405 y_min=410, y_max=410, flashes=0
value
100
150
200
250
0
5
10
15
20
time in minutes

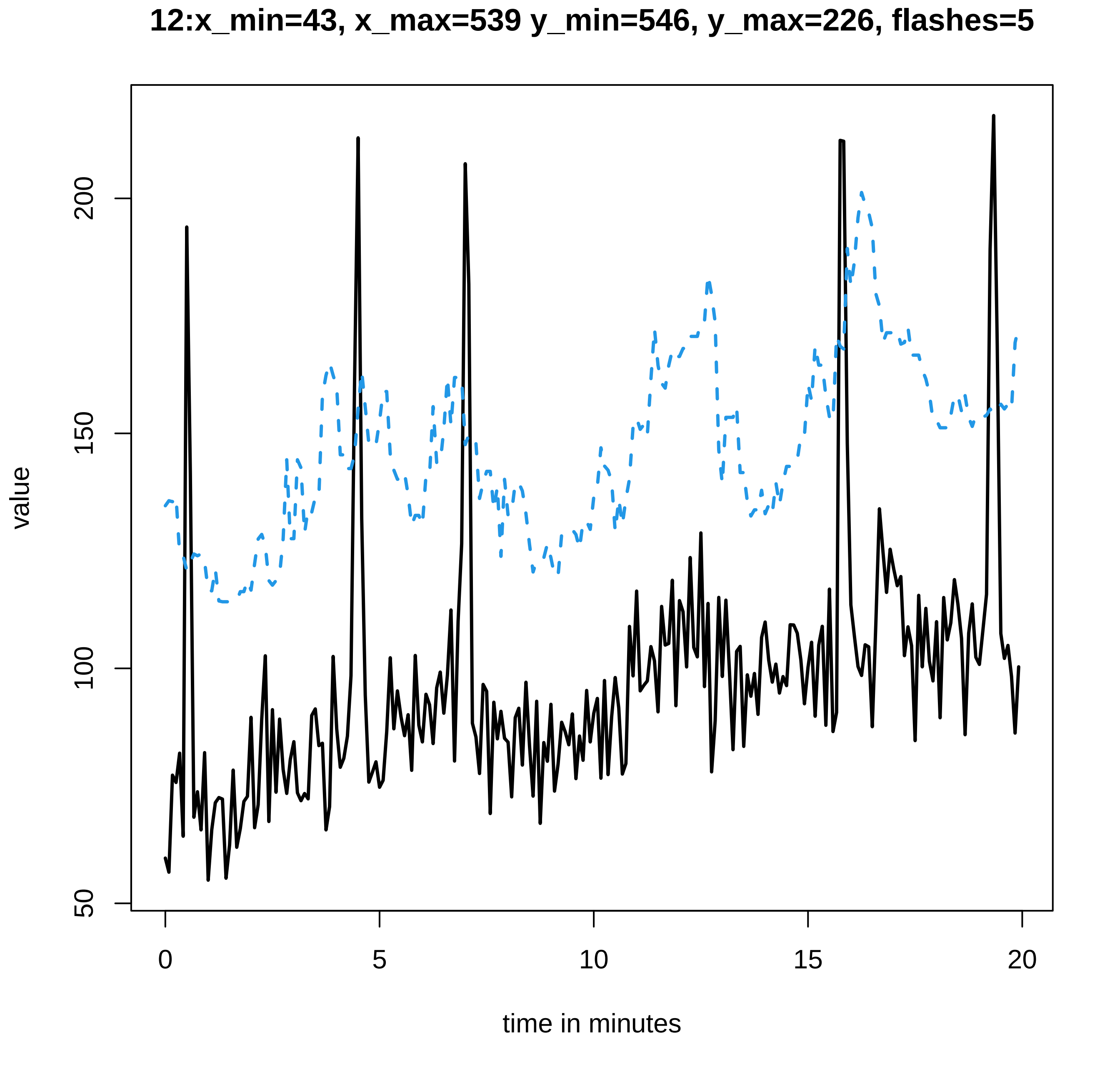
12:x_min=43, x_max=539 y_min=546, y_max=226, flashes=5
value
time in minutes
50
100
150
200
0
5
10
15
20

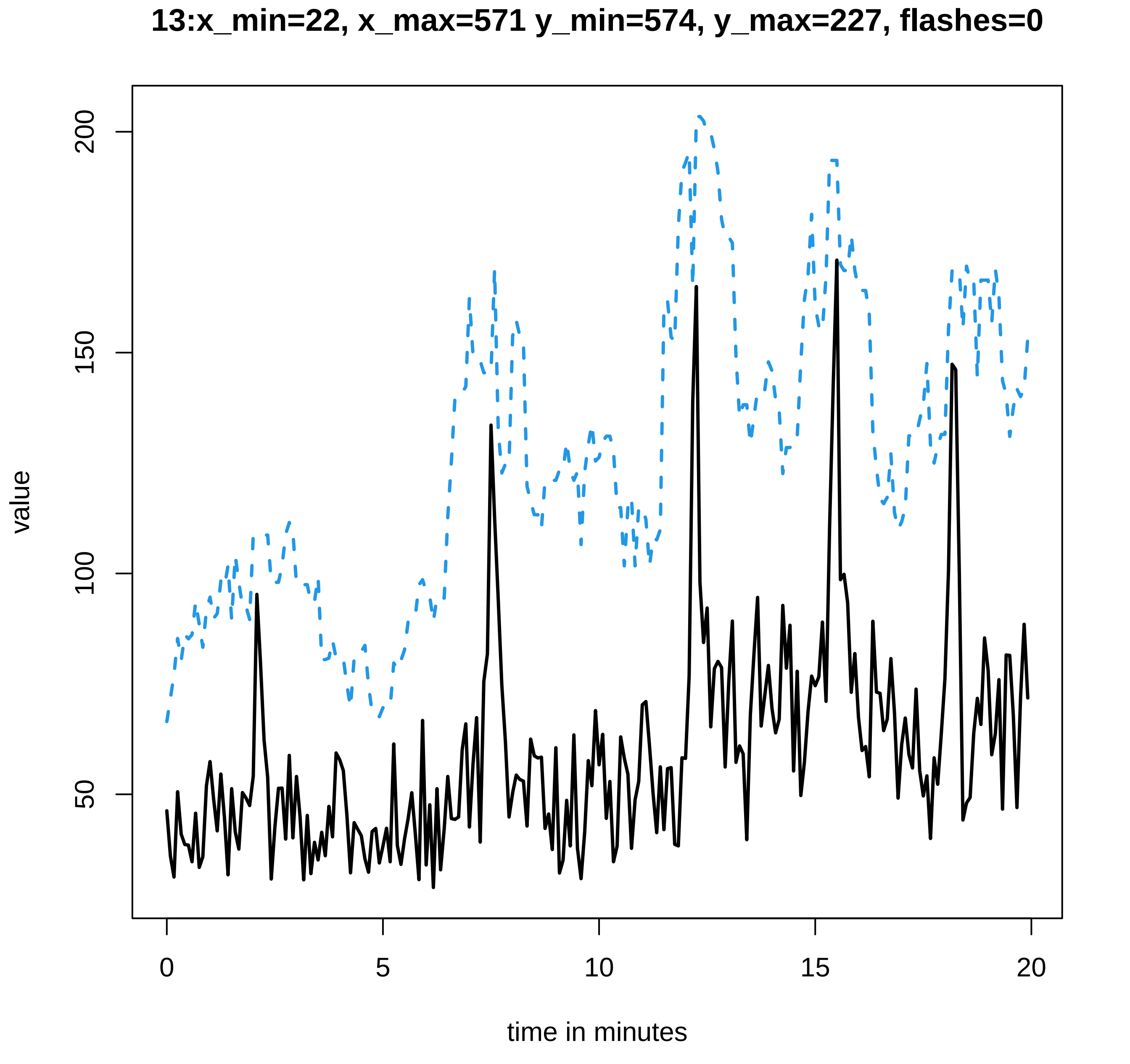

13:x_min=22, x_max=571 y_min=574, y_max=227, flashes=0
value
200
150
100
50
0
5
10
15
20
time in minutes

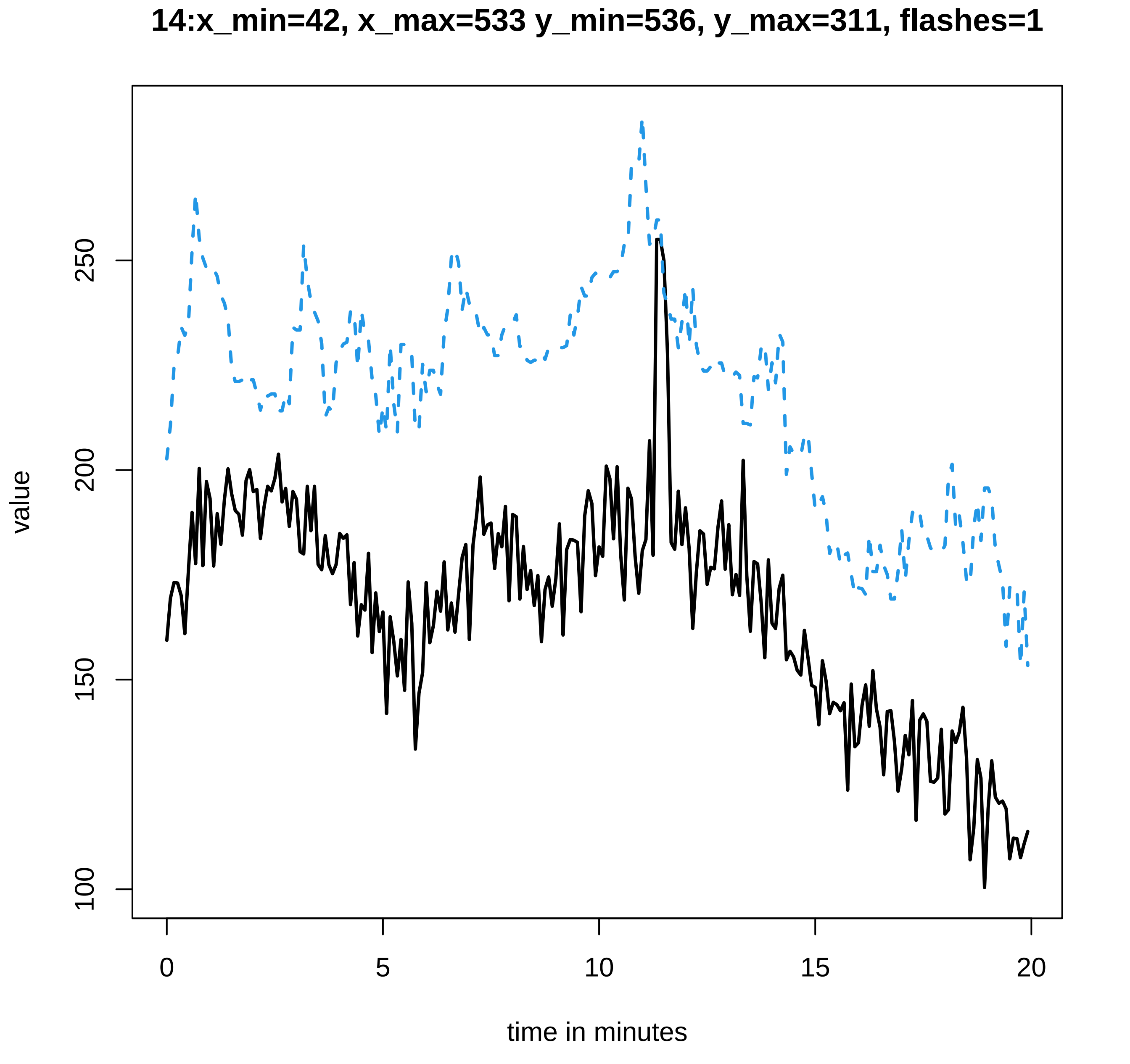

14:x_min=42, x_max=533 y_min=536, y_max=311, flashes=1
value
100
150
200
250
0
5
10
15
20
time in minutes

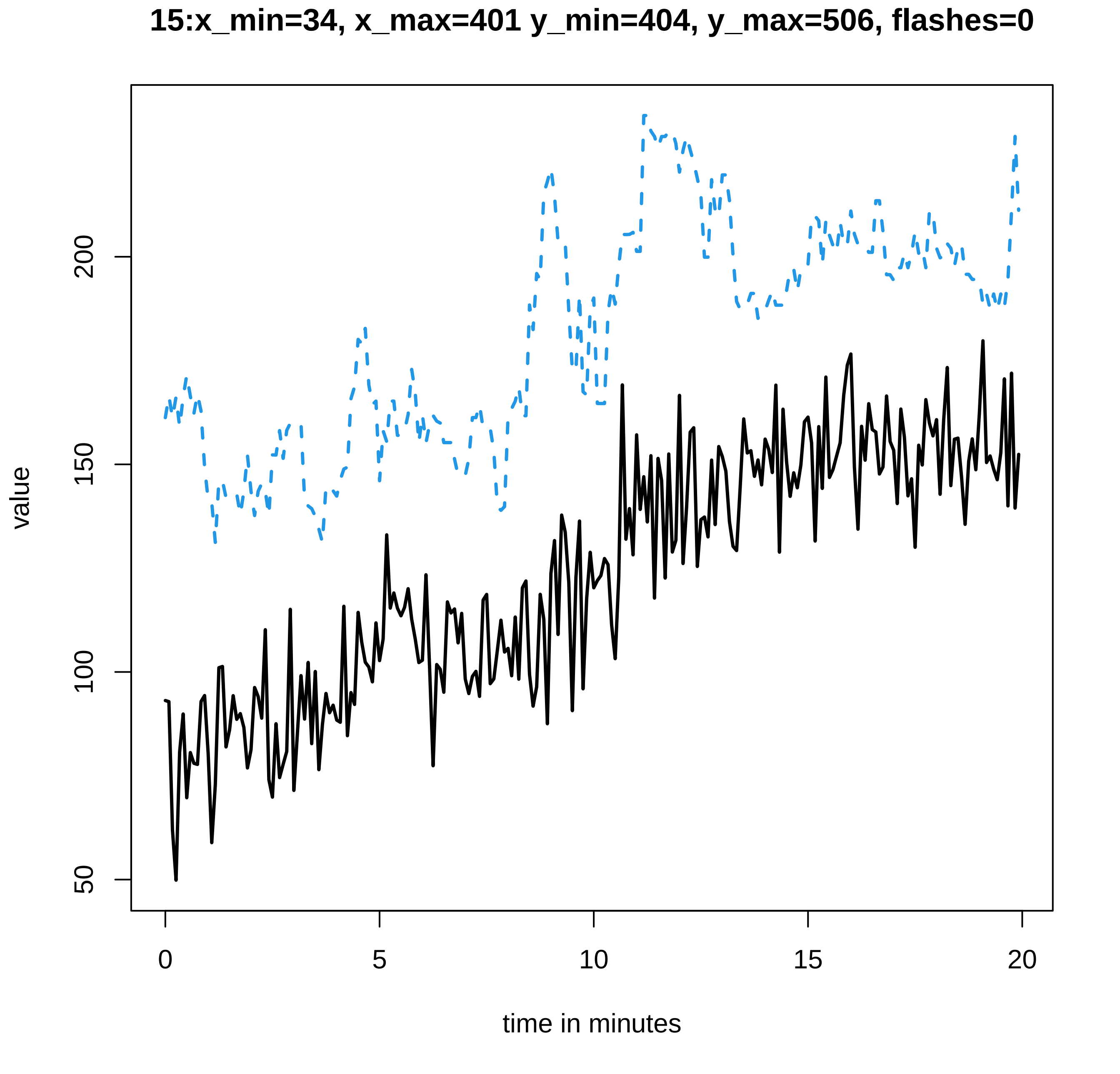

15:x_min=34, x_max=401 y_min=404, y_max=506, flashes=0
value
50
100
150
200
0
5
10
15
20
time in minutes

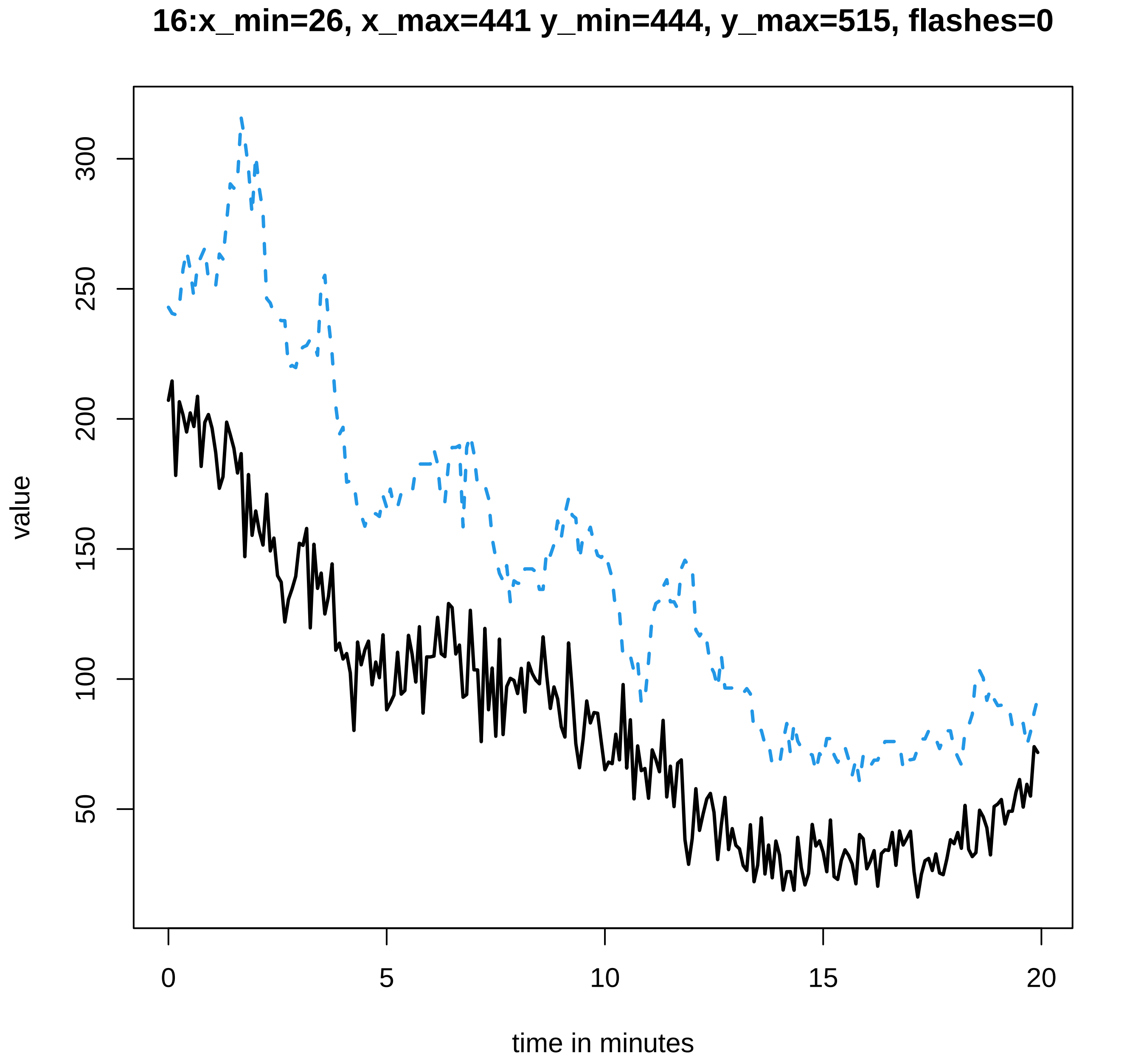
16:x_min=26, x_max=441 y_min=444, y_max=515, flashes=0
value
50
100
150
200
250
300
0
5
10
15
20
time in minutes

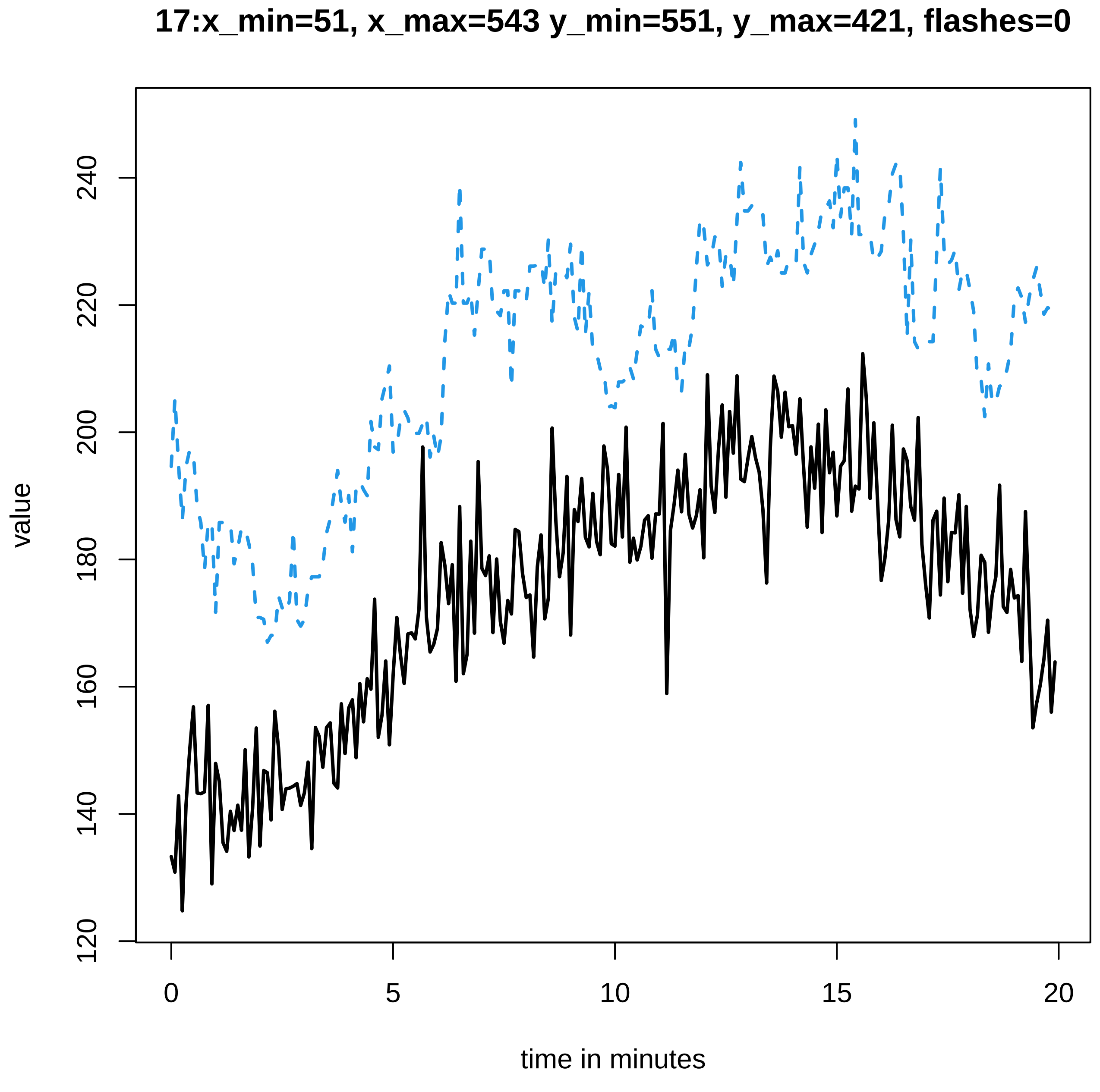

17:x_min=51, x_max=543 y_min=551, y_max=421, flashes=0
value
120
140
160
180
200
220
240
0
5
10
15
20
time in minutes

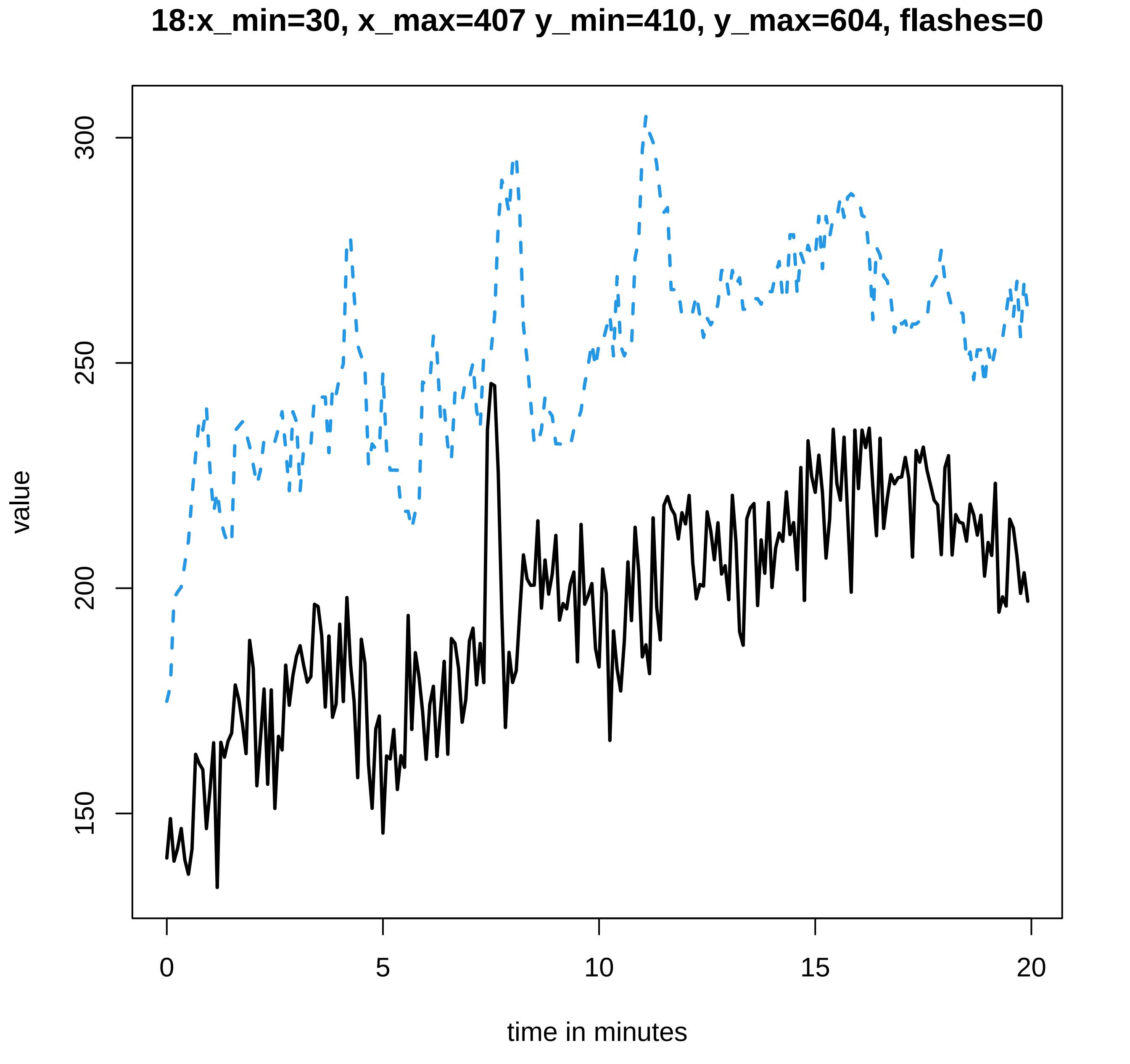
18:x_min=30, x_max=407 y_min=410, y_max=604, flashes=0
value
300
250
200
150
0
5
10
15
20
time in minutes

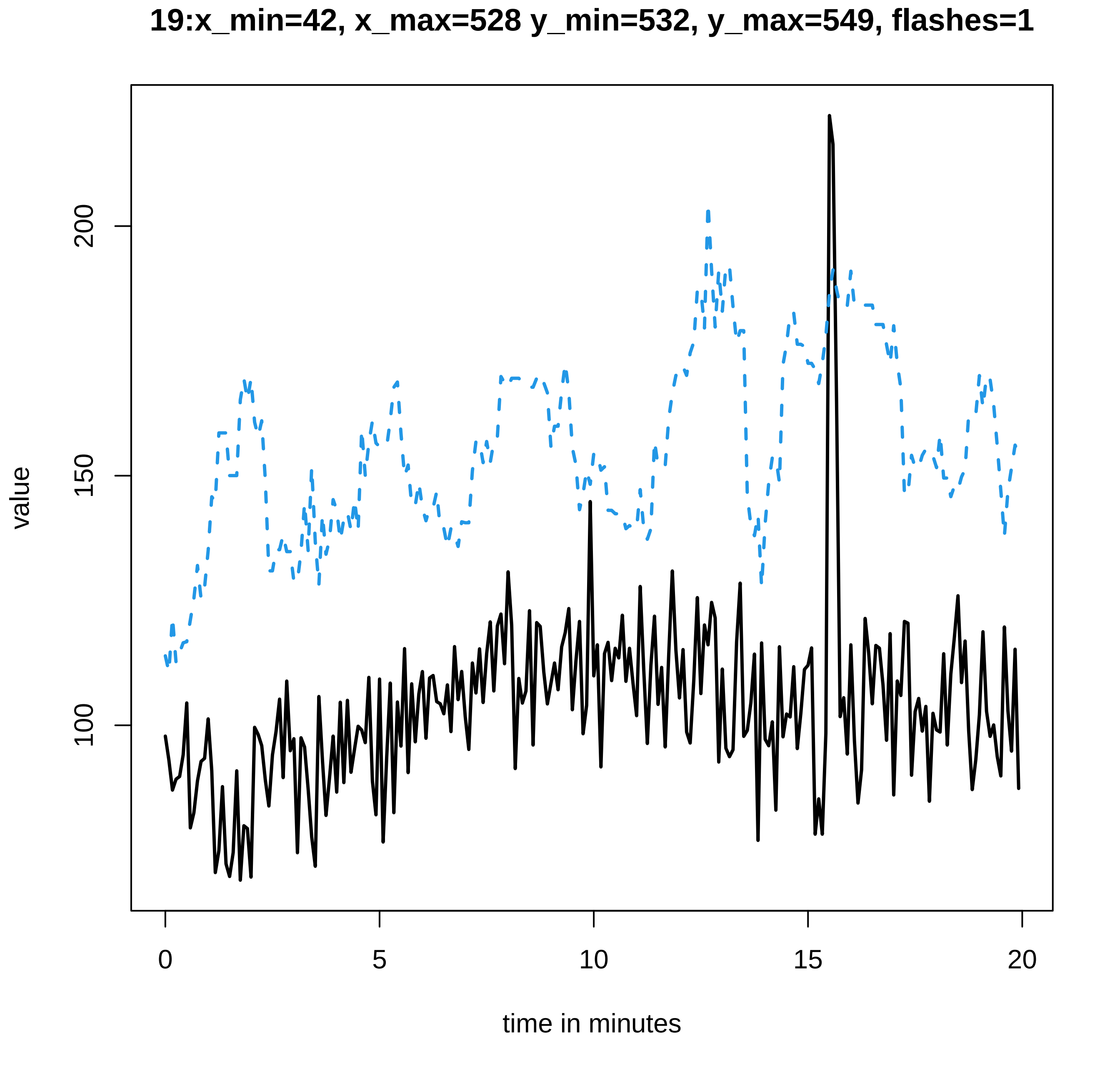
19:x_min=42, x_max=528 y_min=532, y_max=549, flashes=1
value
200
150
100
0
5
10
15
20
time in minutes

**20:x_min=60, x_max=540 y_min=546, y_max=611, flashes=1**

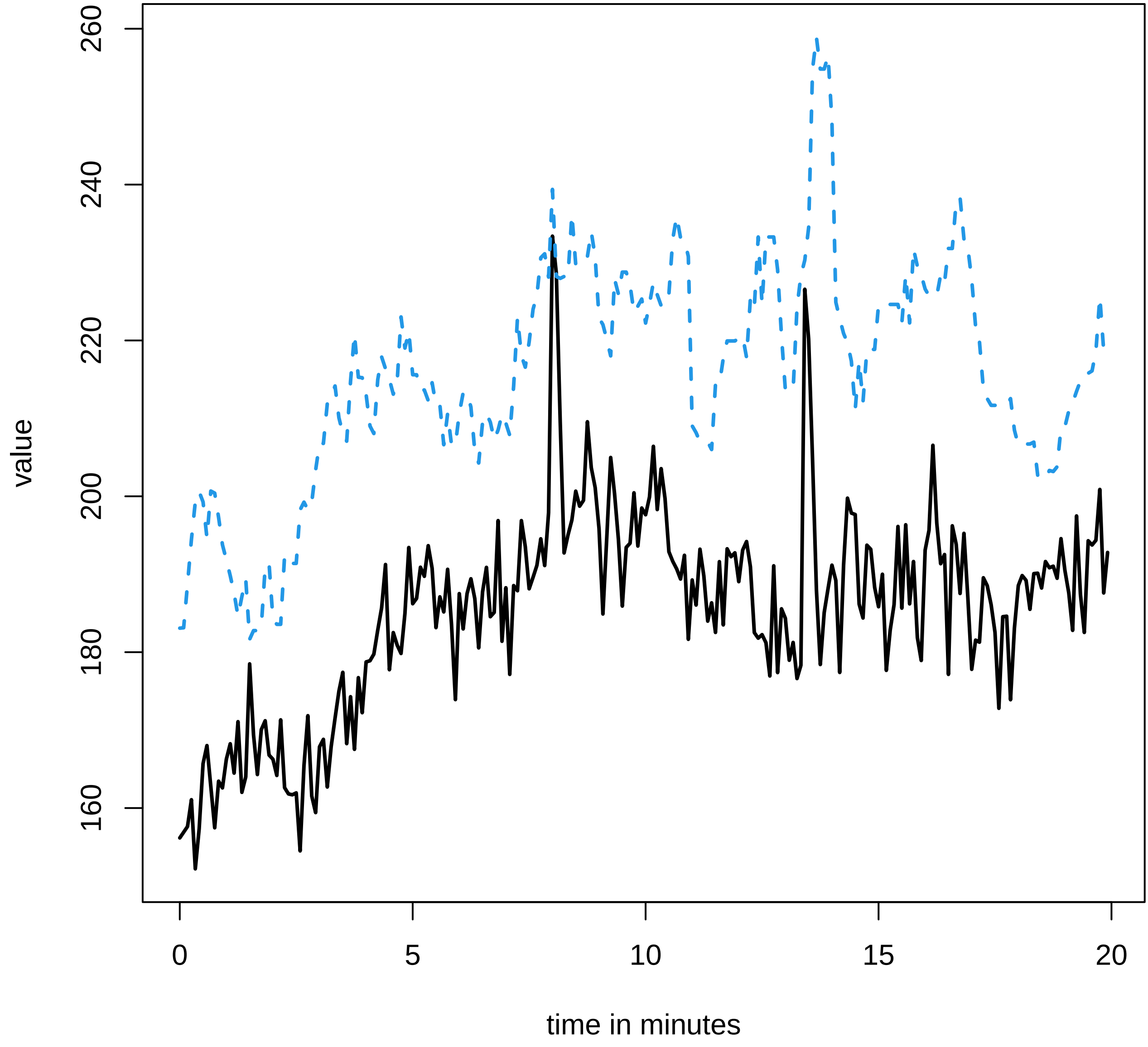

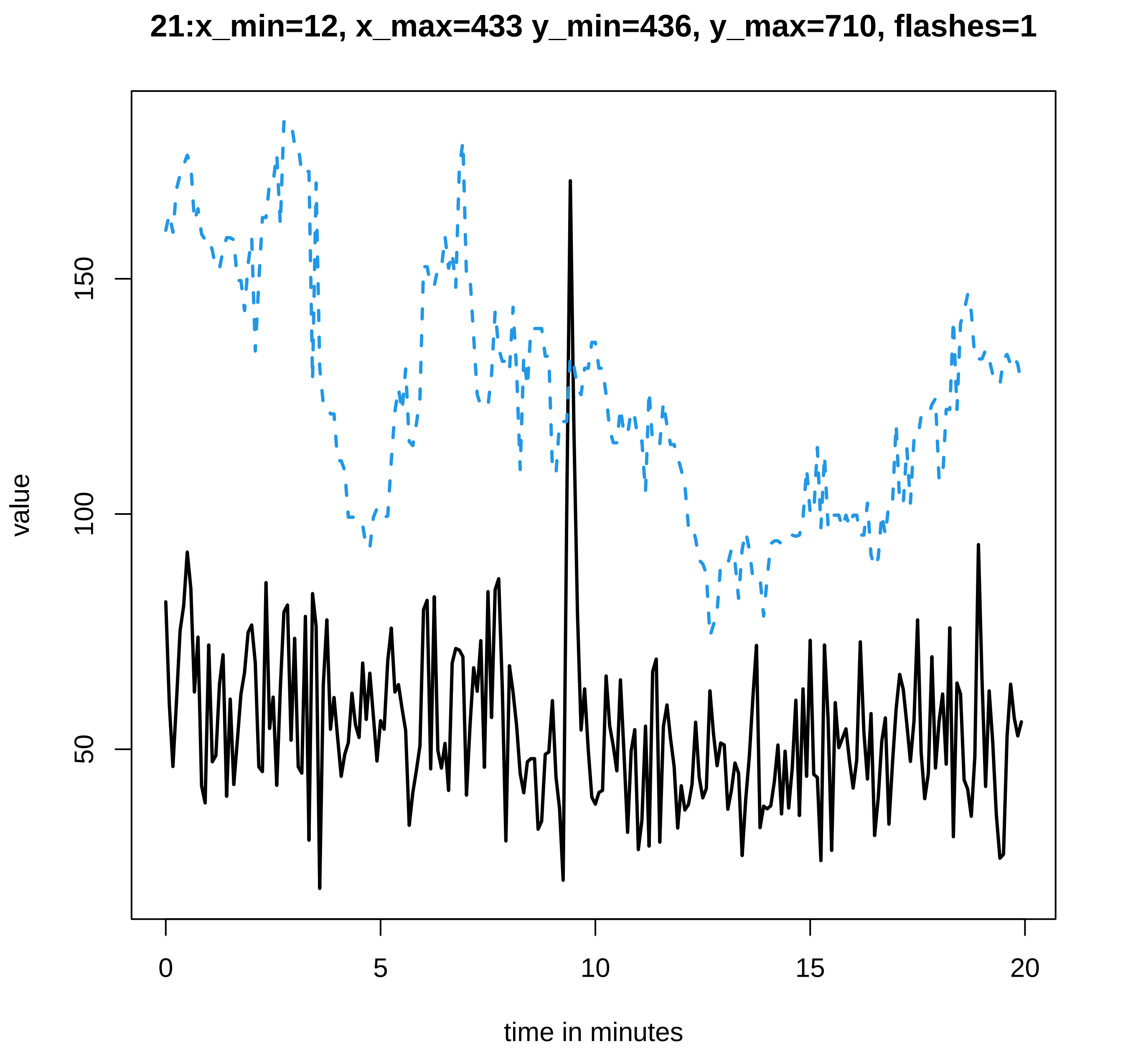
21:x_min=12, x_max=433 y_min=436, y_max=710, flashes=1
value
150
100
50
0
5
10
15
20
time in minutes

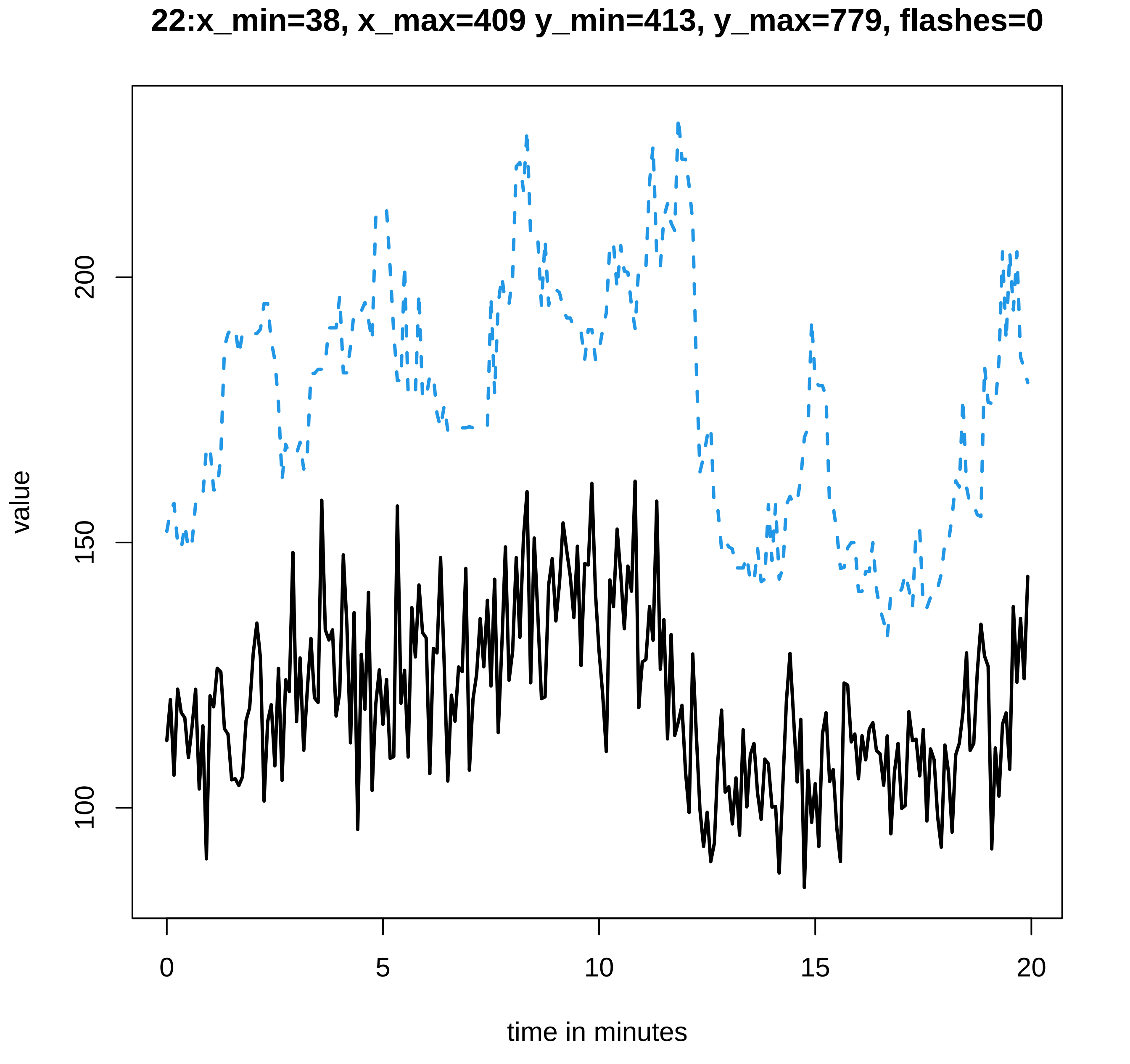
22:x_min=38, x_max=409 y_min=413, y_max=779, flashes=0
value
100
150
200
0
5
10
15
20
time in minutes

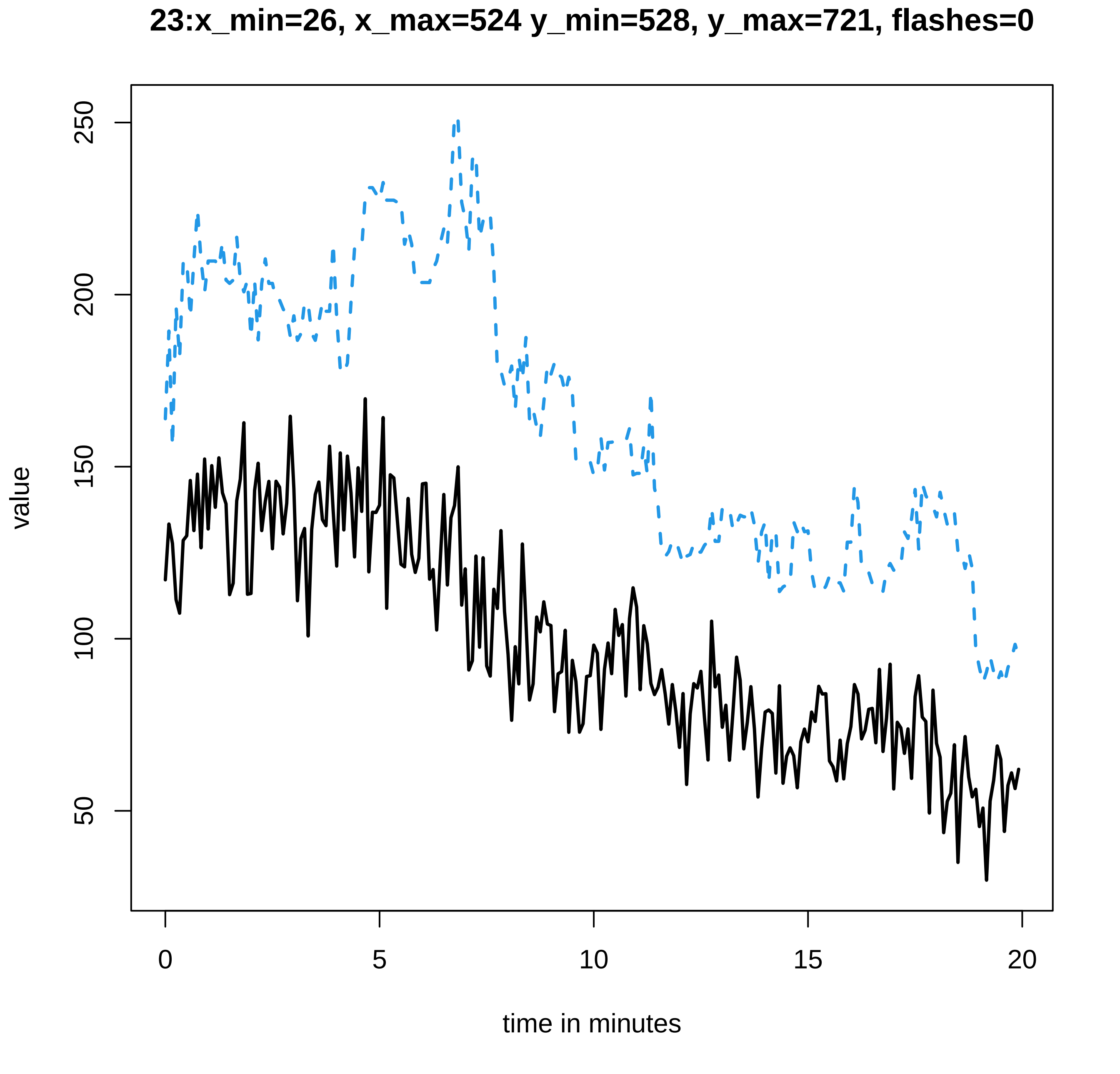

23:x_min=26, x_max=524 y_min=528, y_max=721, flashes=0
value
250
200
150
100
50
0
5
10
15
20
time in minutes

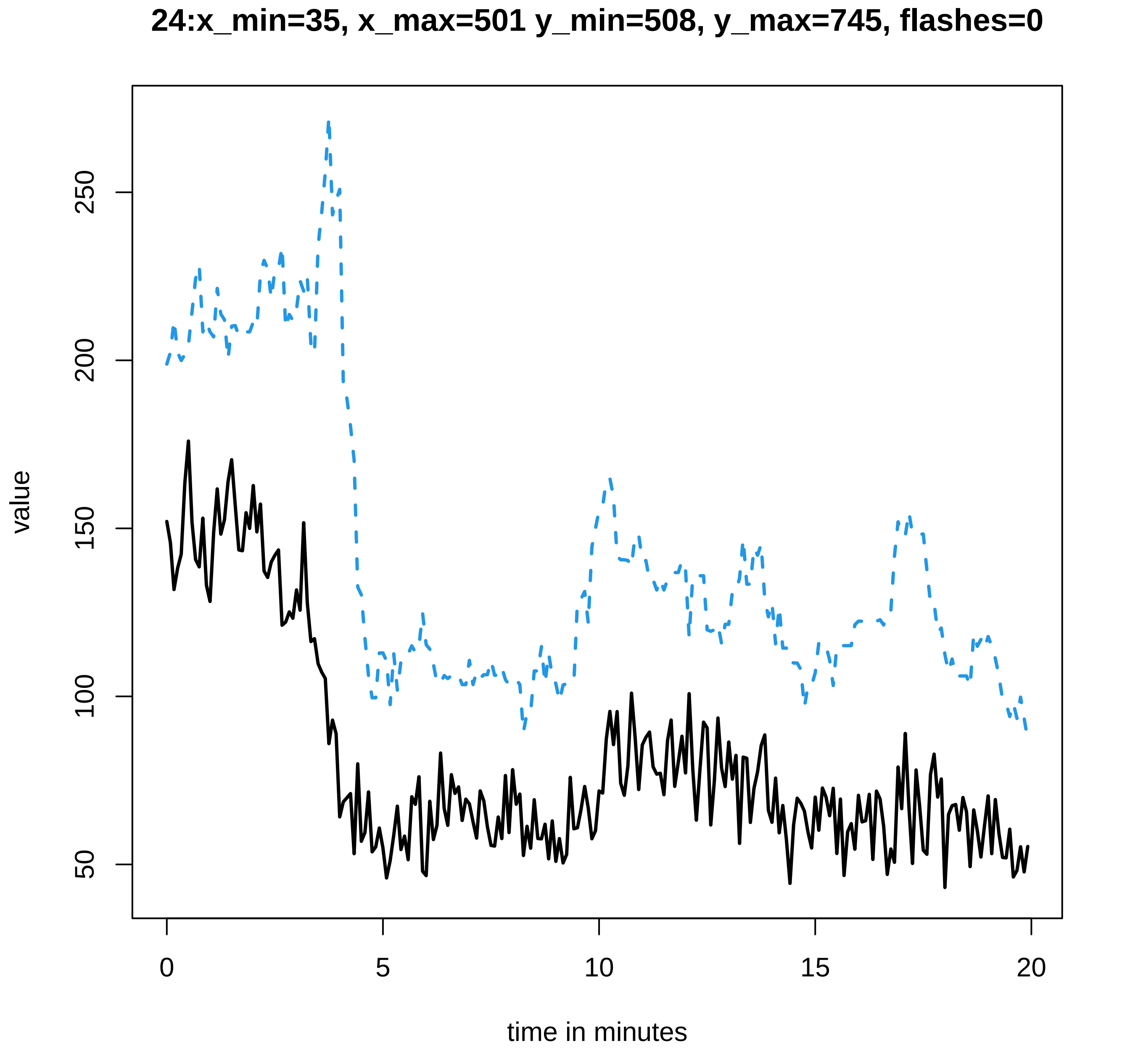

24:x_min=35, x_max=501 y_min=508, y_max=745, flashes=0
value
50
100
150
200
250
0
5
10
15
20
time in minutes

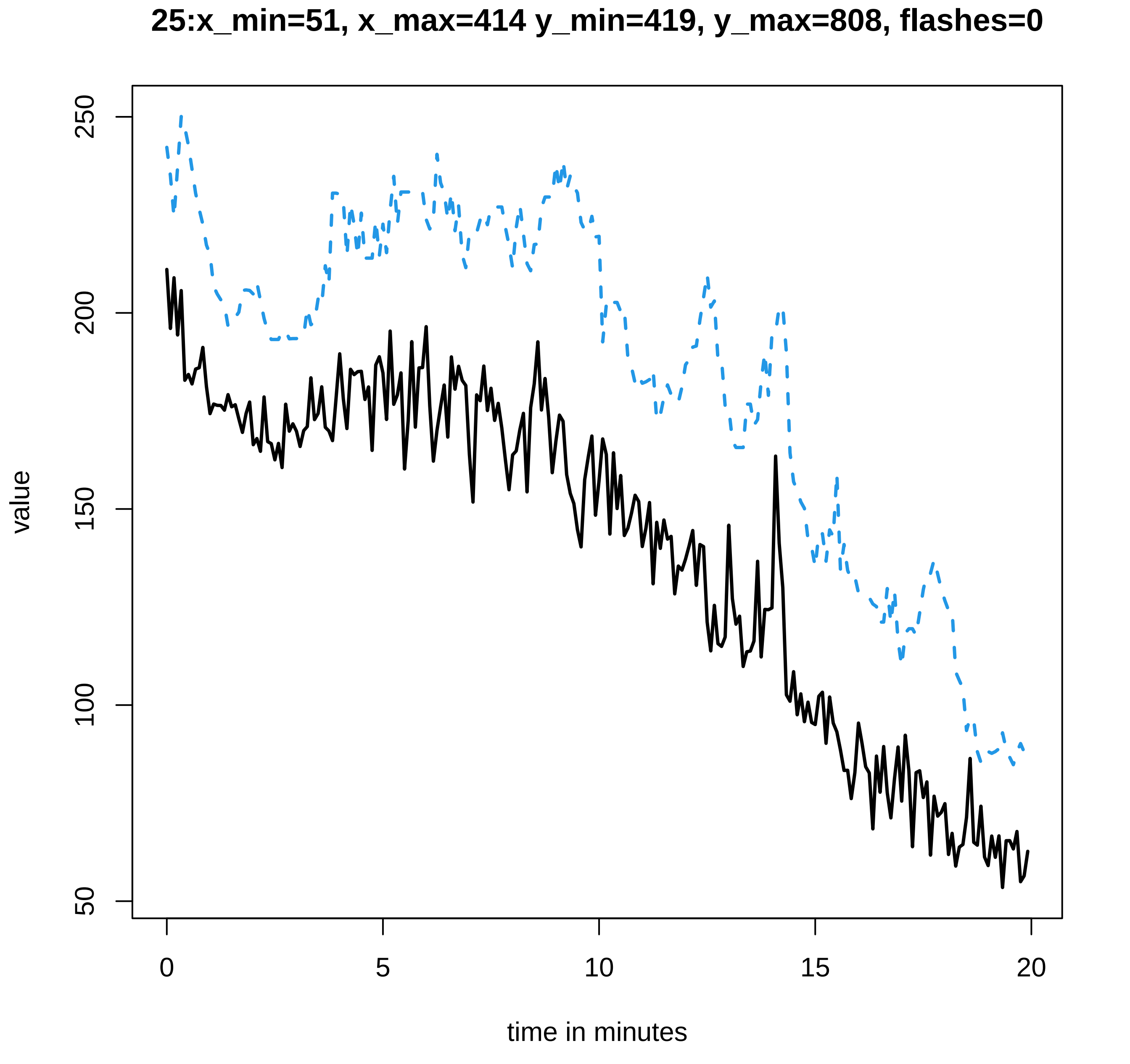
25:x_min=51, x_max=414 y_min=419, y_max=808, flashes=0
value
250
200
150
100
50
0
5
10
15
20
time in minutes

26:x_min=42, x_max=429 y_min=433, y_max=826, flashes=0

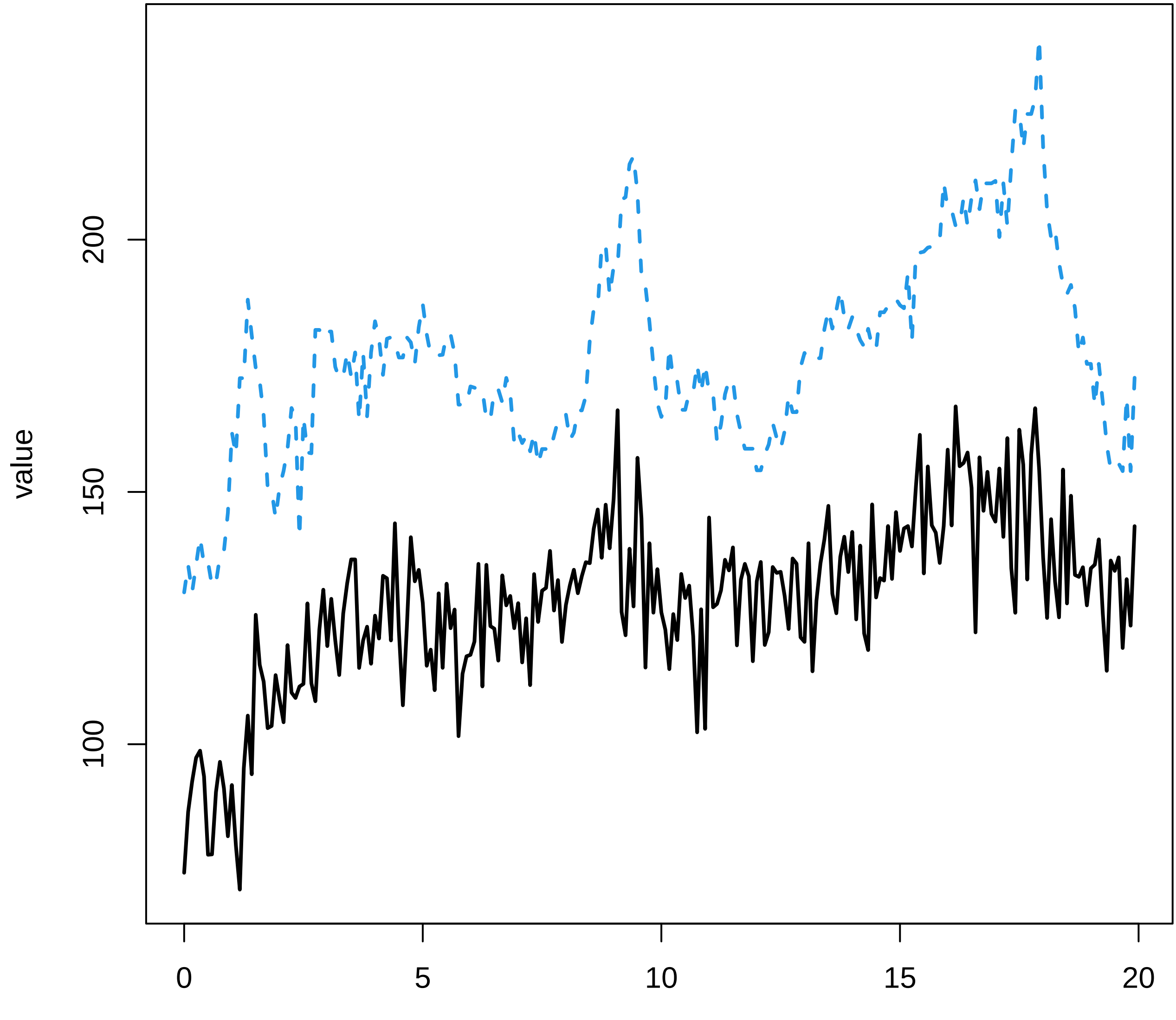

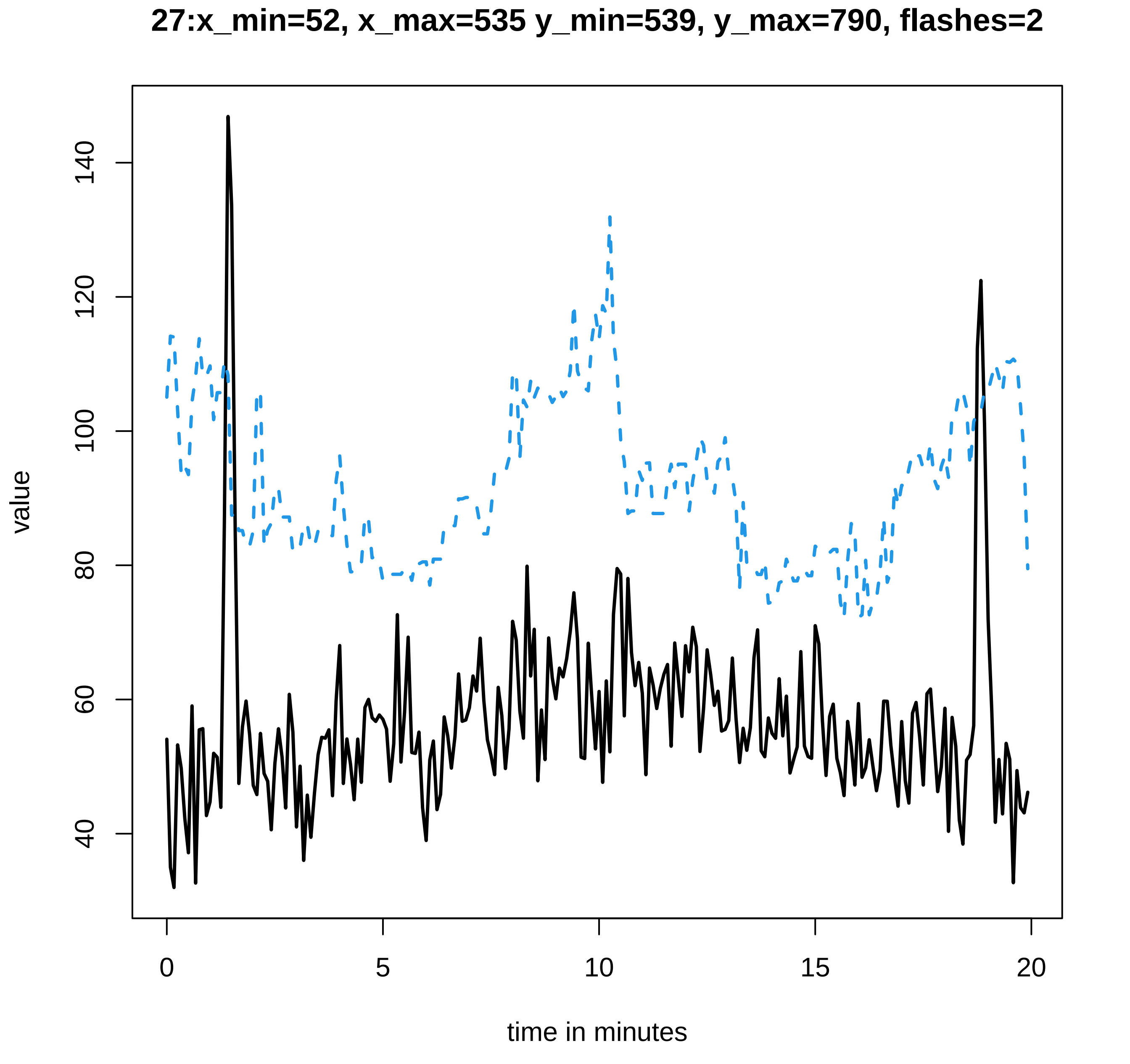

27:x_min=52, x_max=535 y_min=539, y_max=790, flashes=2
value
40
60
80
100
120
140
0
5
10
15
20
time in minutes

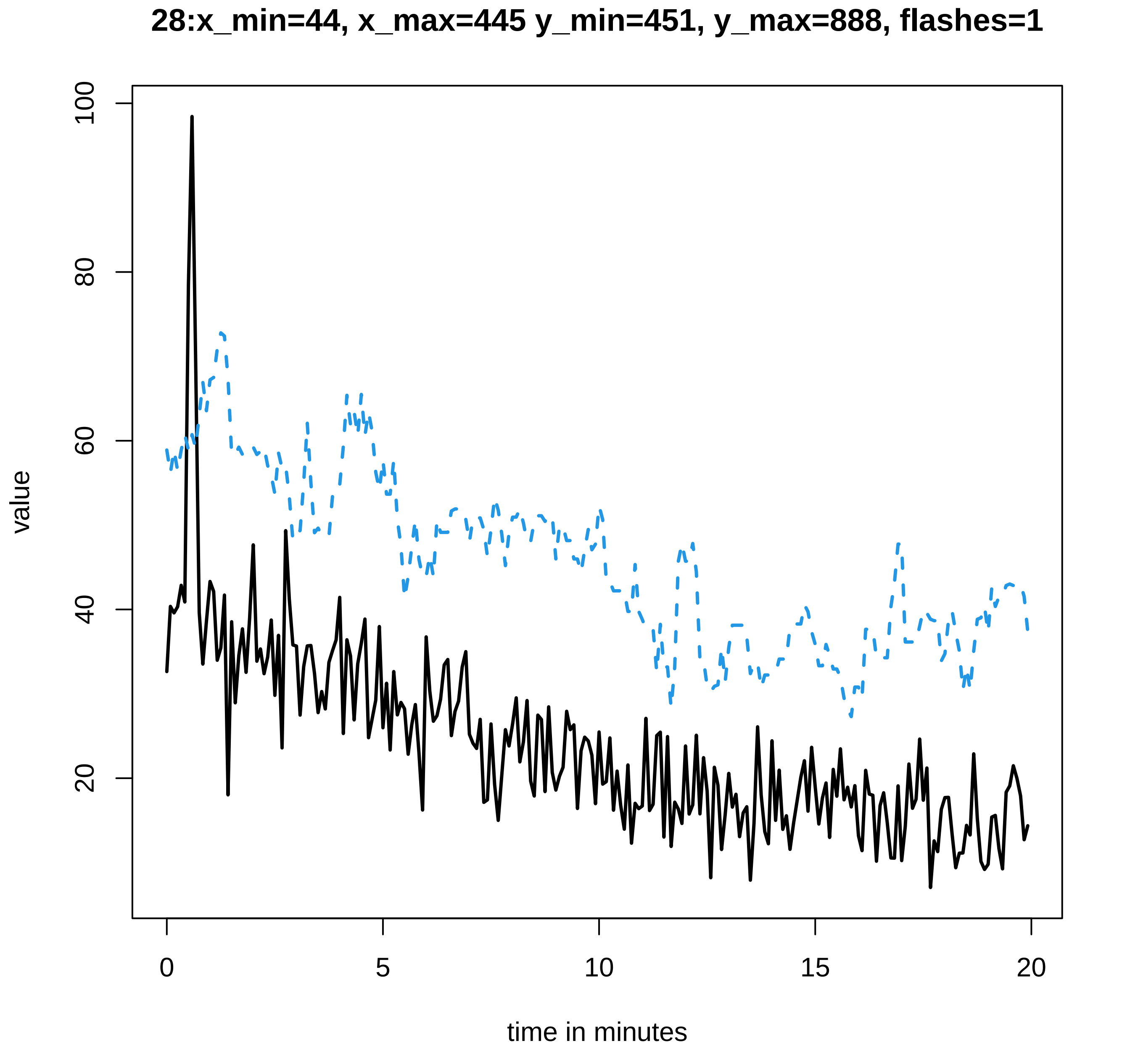
28:x_min=44, x_max=445 y_min=451, y_max=888, flashes=1
value
100
80
60
40
20
0
5
10
15
20
time in minutes

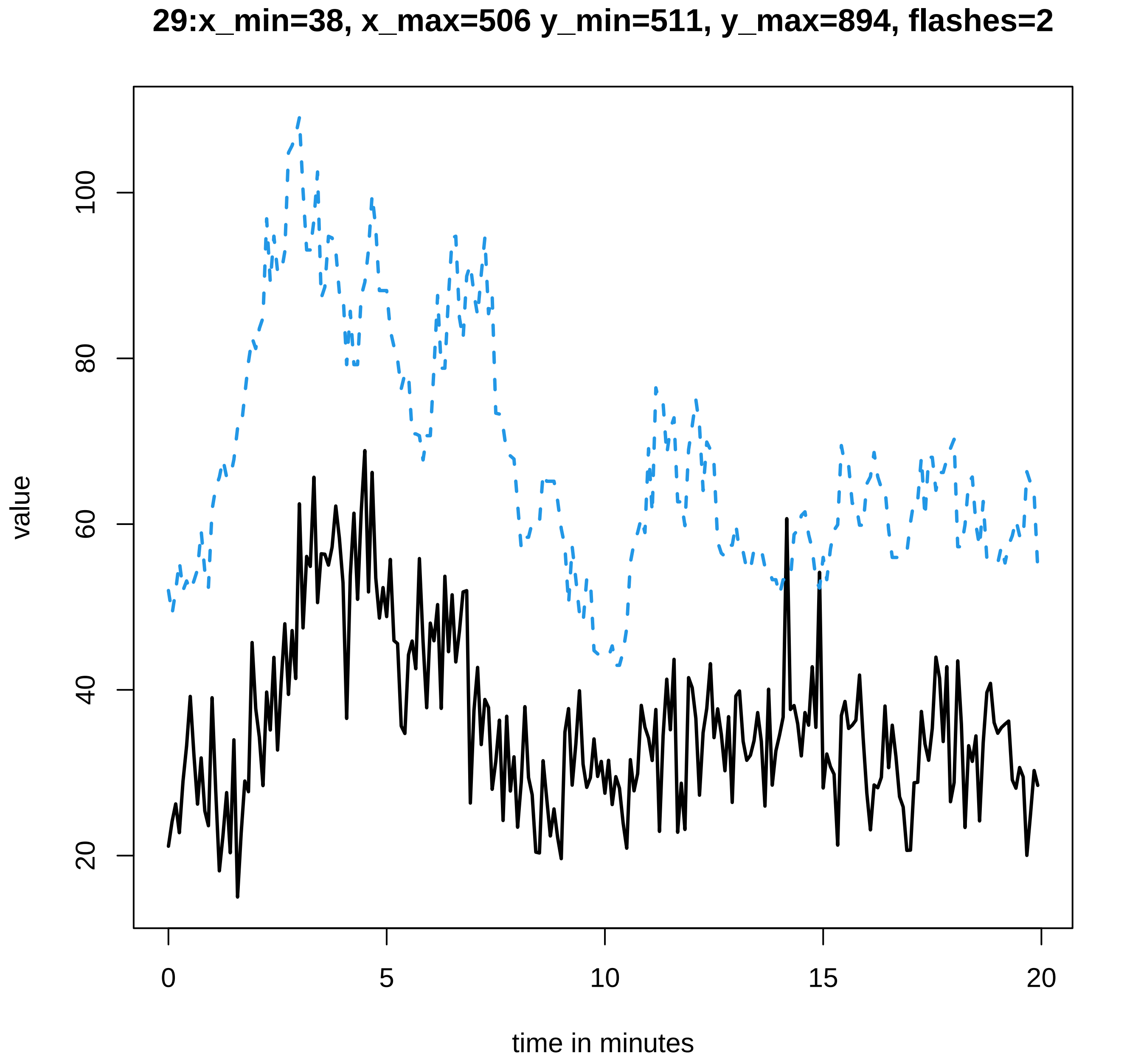
29:x_min=38, x_max=506 y_min=511, y_max=894, flashes=2
value
20
40
60
80
100
0
5
10
15
20
time in minutes

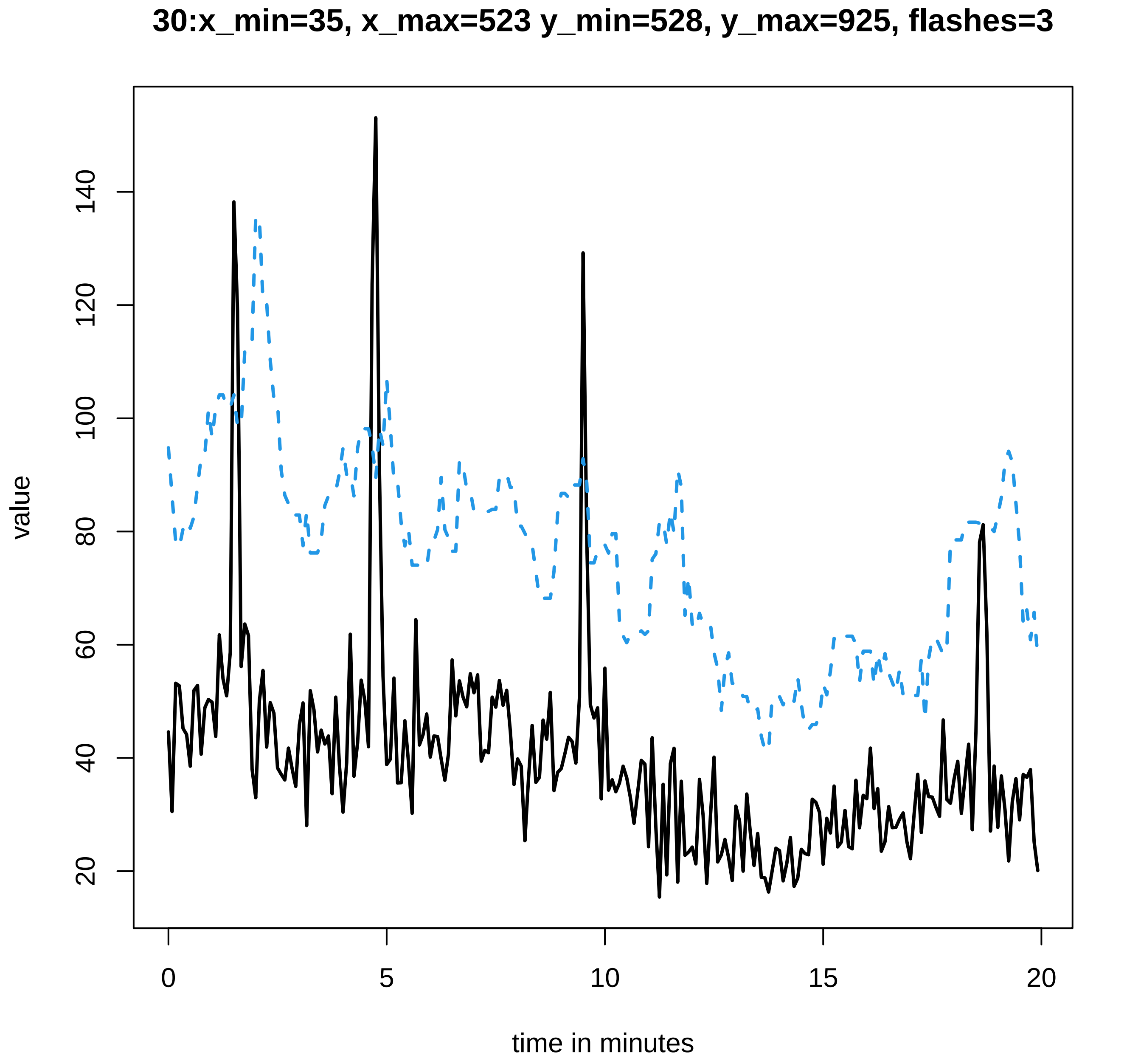
30:x_min=35, x_max=523 y_min=528, y_max=925, flashes=3
value
20
40
60
80
100
120
140
0
5
10
15
20
time in minutes
