## Supplementary data Calcium spiking for "LysM Receptor Proteins are Required for Ectomycorrhizal Symbiosis in Poplar": profiles-spiking-5.pdf

### 1:x_min=46, x_max=68 y_min=78, y_max=476, flashes=0

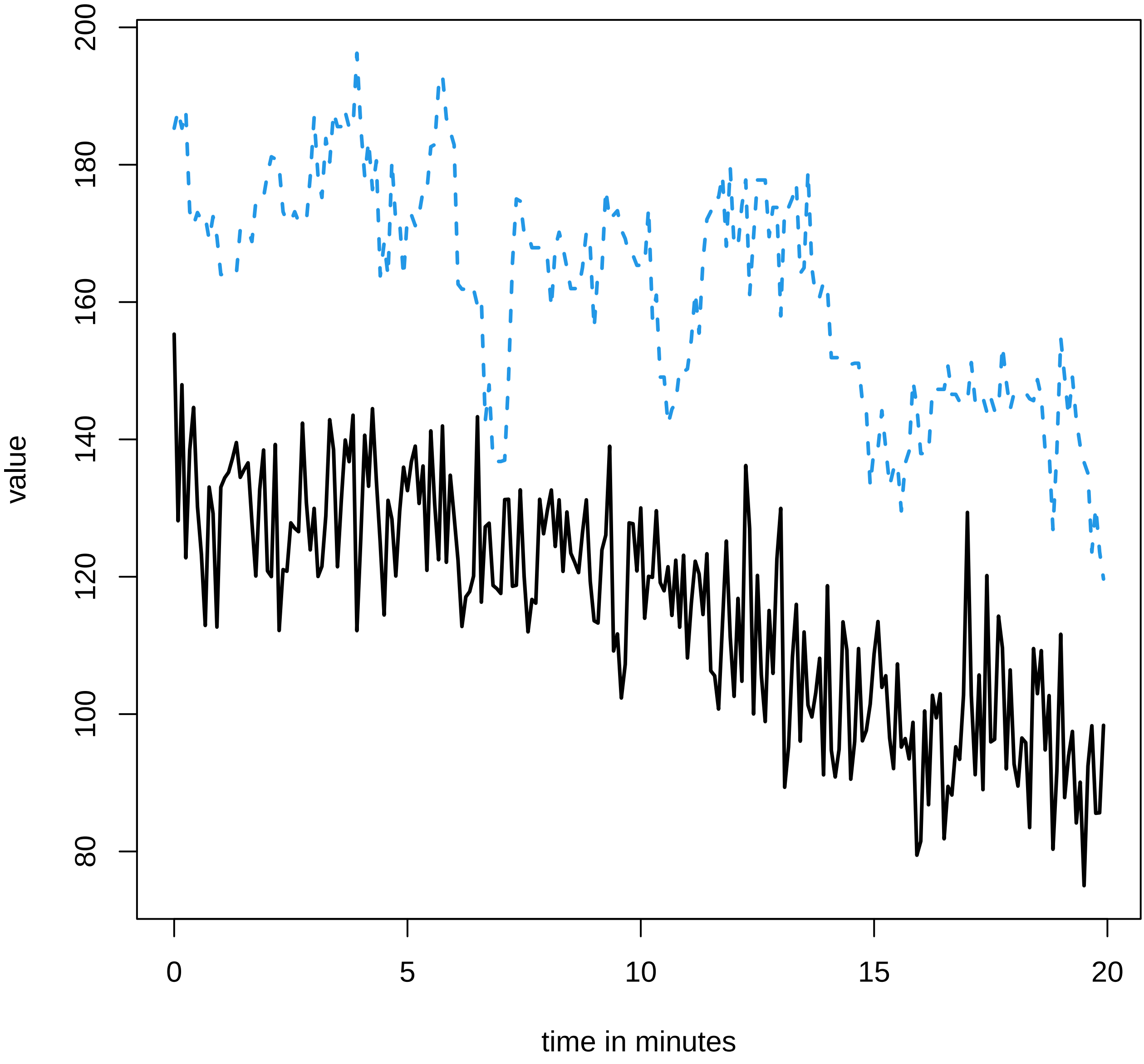

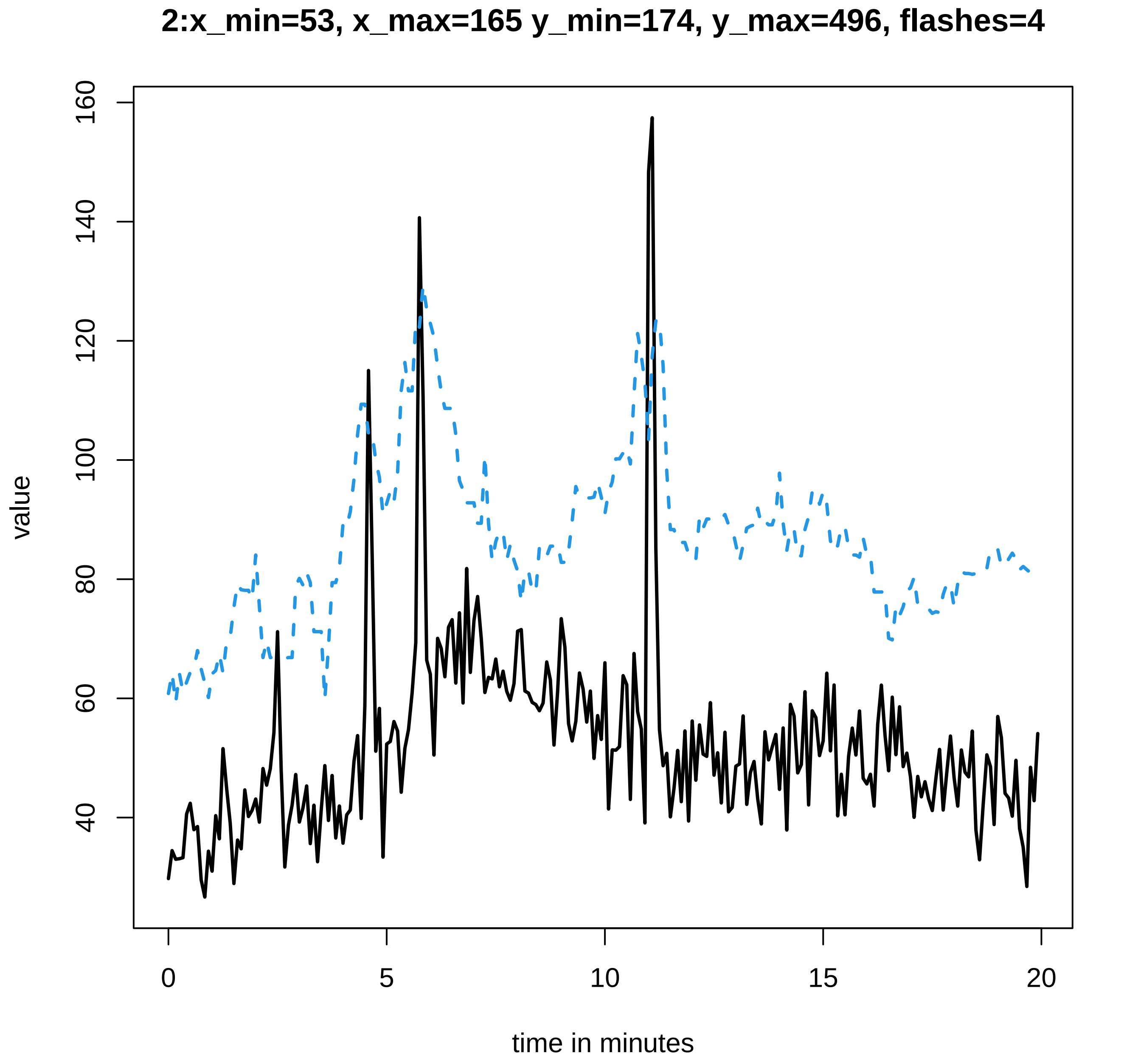
2:x_min=53, x_max=165 y_min=174, y_max=496, flashes=4
value
time in minutes
0
5
10
15
20
40
60
80
100
120
140
160

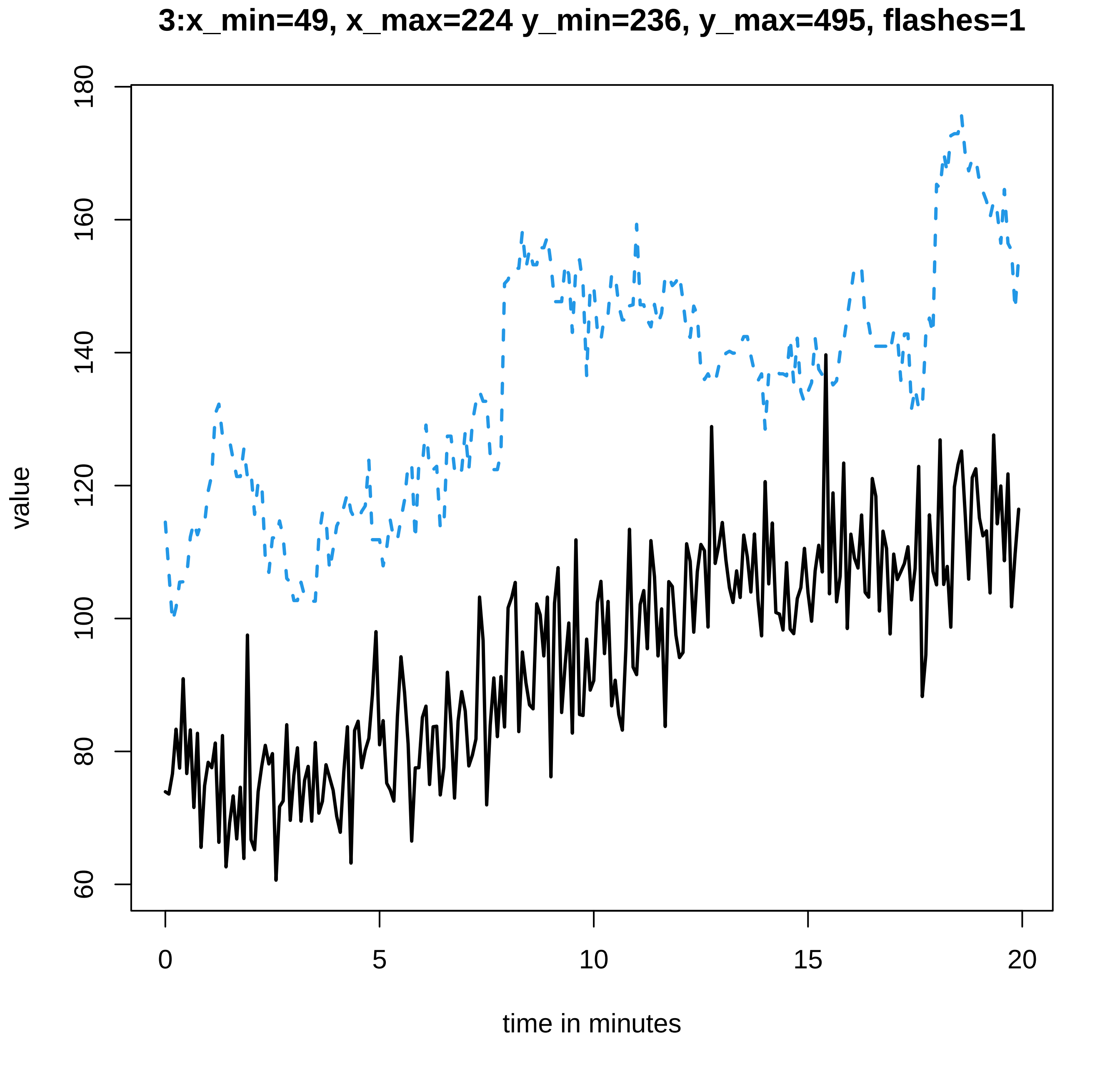
3:x_min=49, x_max=224 y_min=236, y_max=495, flashes=1
180
160
140
120
100
80
60
value
0
5
10
15
20
time in minutes

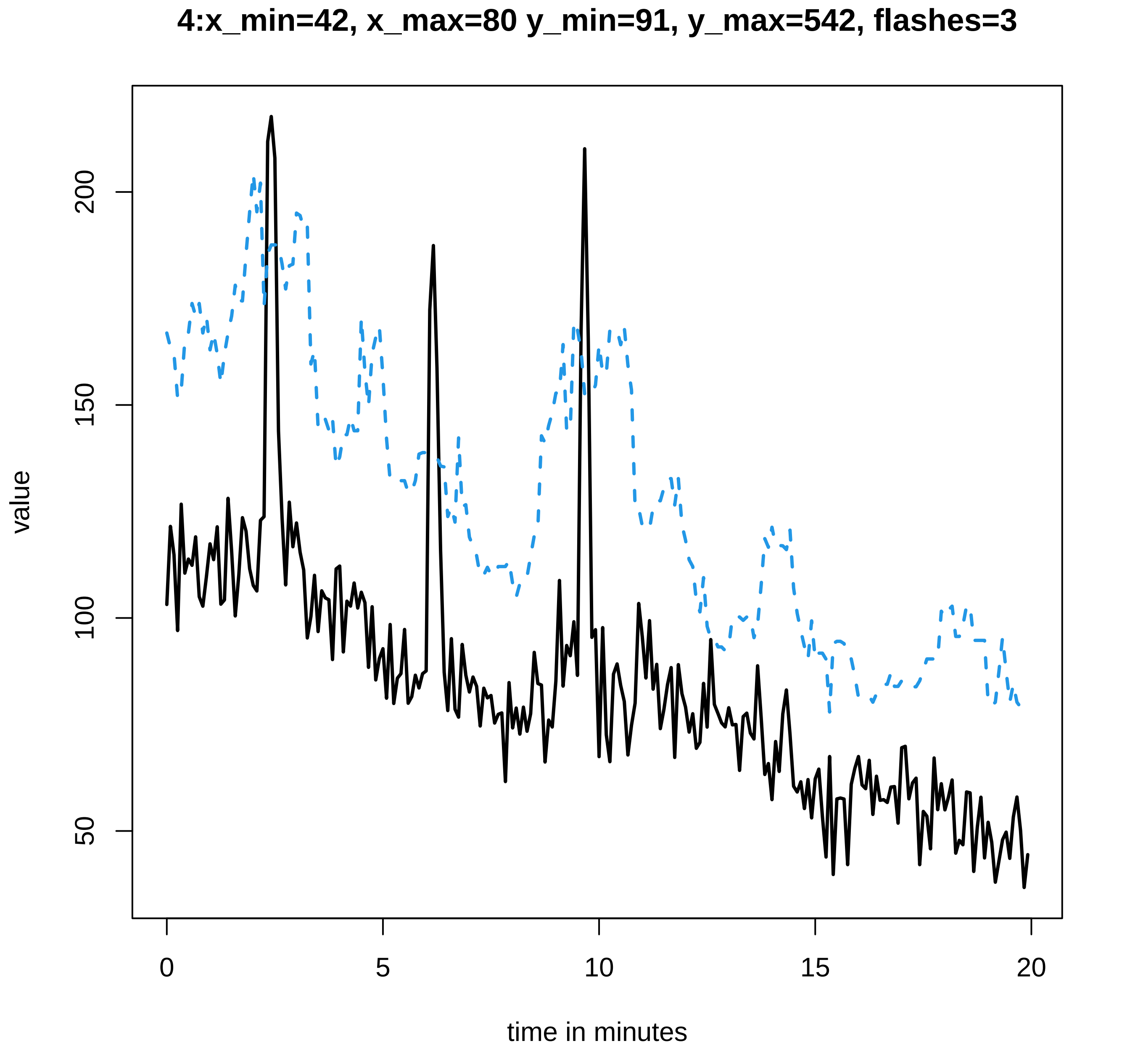

4:x_min=42, x_max=80 y_min=91, y_max=542, flashes=3
value
time in minutes
0
5
10
15
20
50
100
150
200

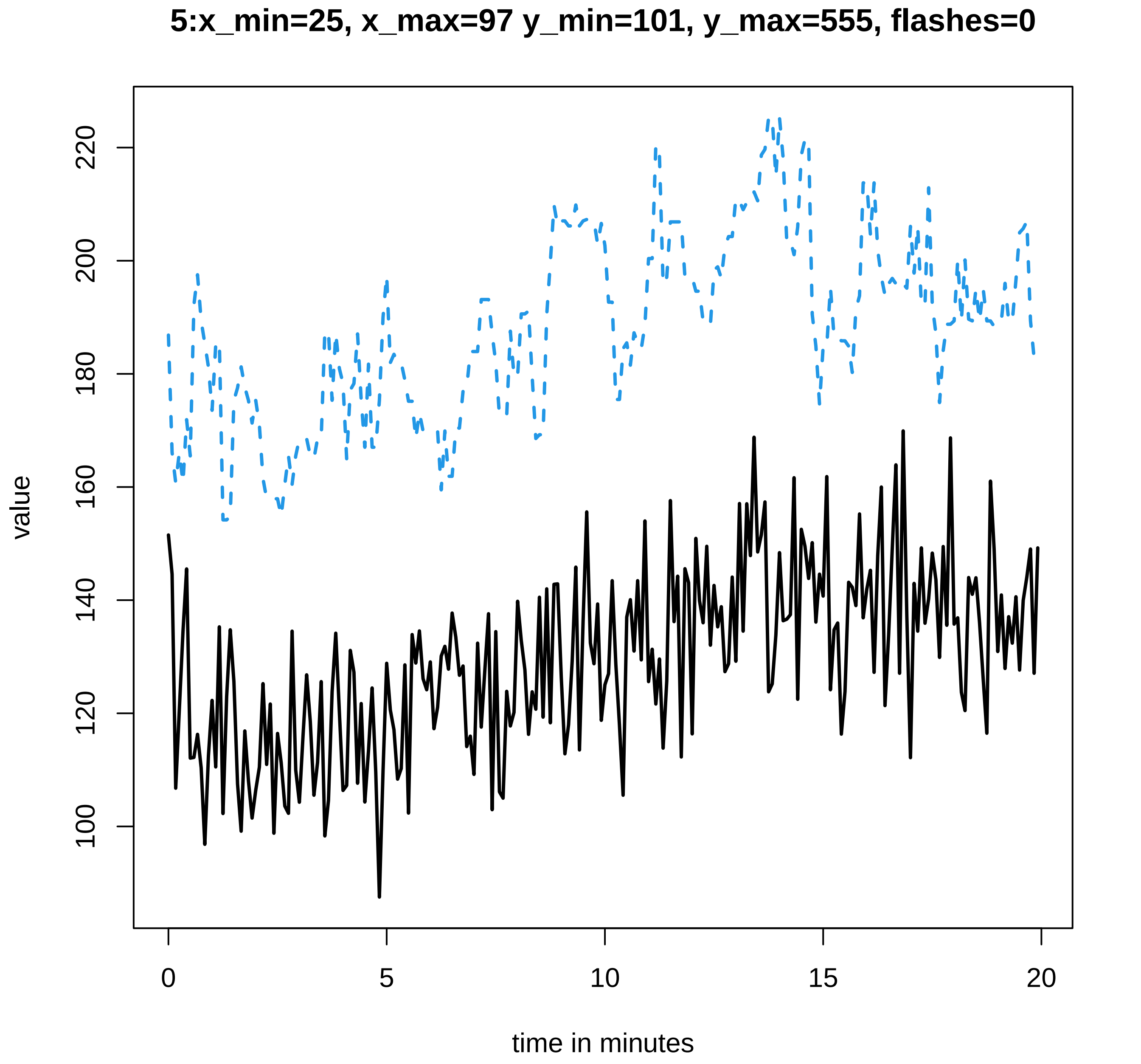
5:x_min=25, x_max=97 y_min=101, y_max=555, flashes=0
value
100
120
140
160
180
200
220
0
5
10
15
20
time in minutes

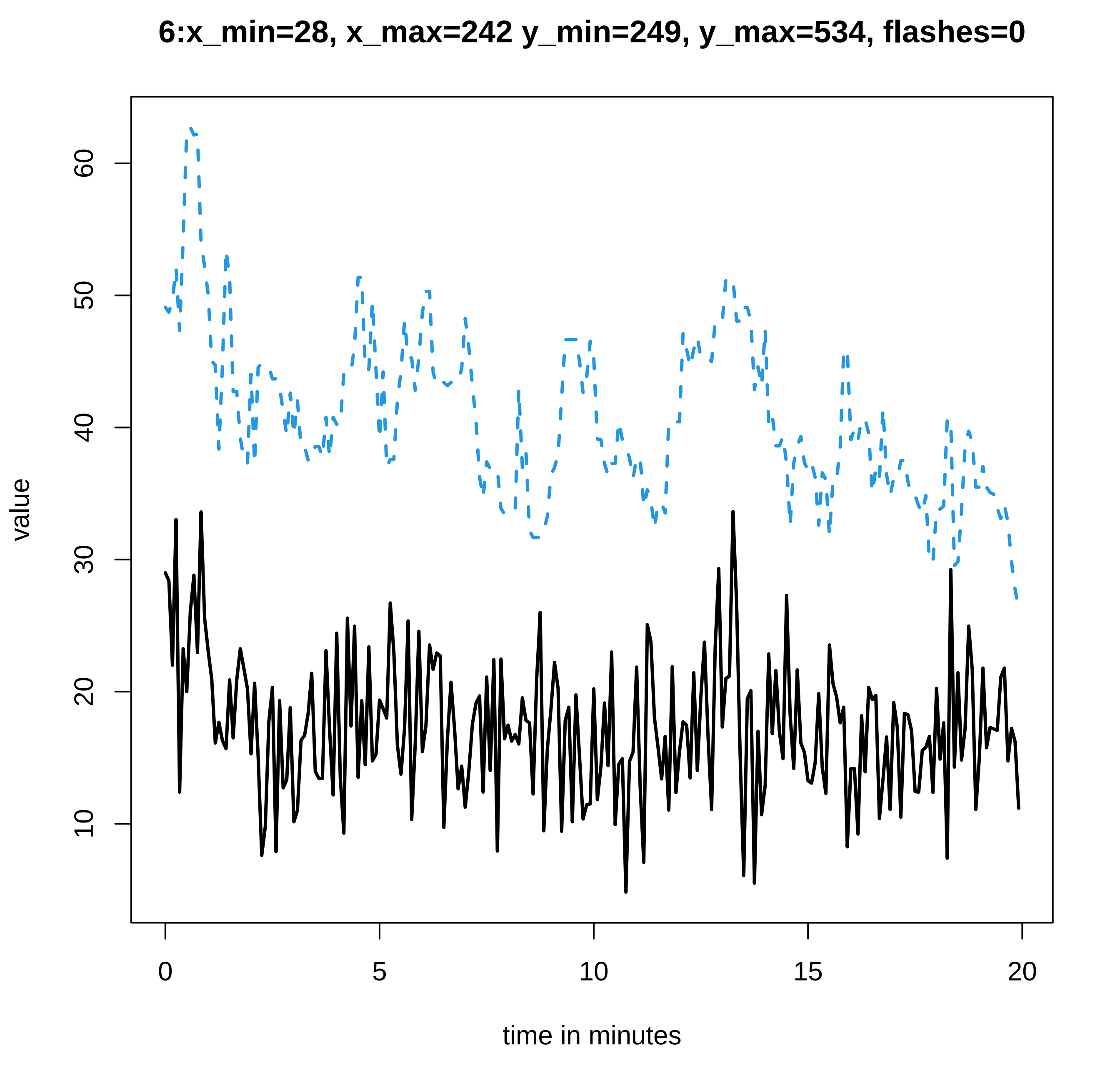
6:x_min=28, x_max=242 y_min=249, y_max=534, flashes=0
value
time in minutes
10
20
30
40
50
60
0
5
10
15
20

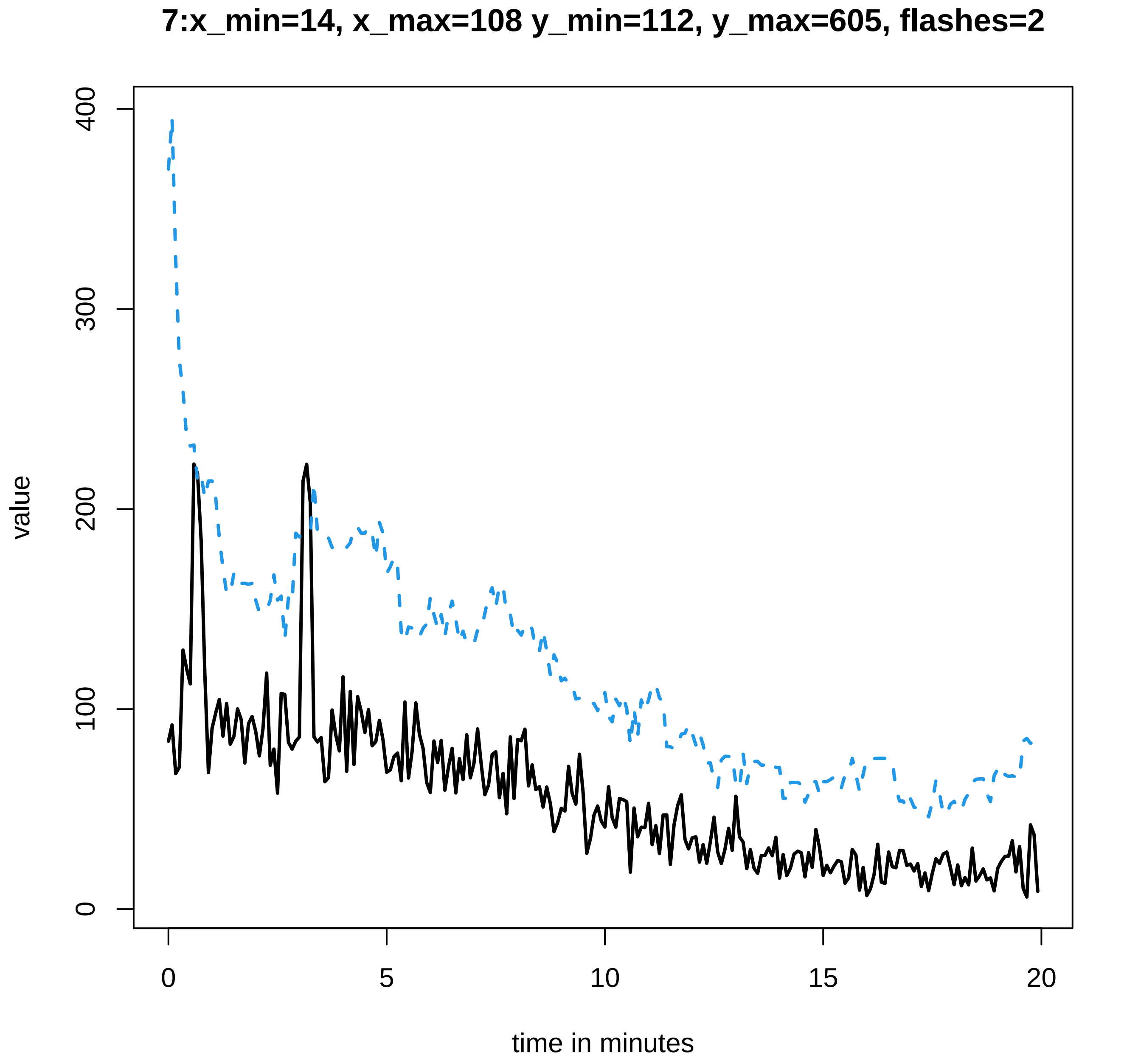
7:x_min=14, x_max=108 y_min=112, y_max=605, flashes=2
value
0
100
200
300
400
0
5
10
15
20
time in minutes

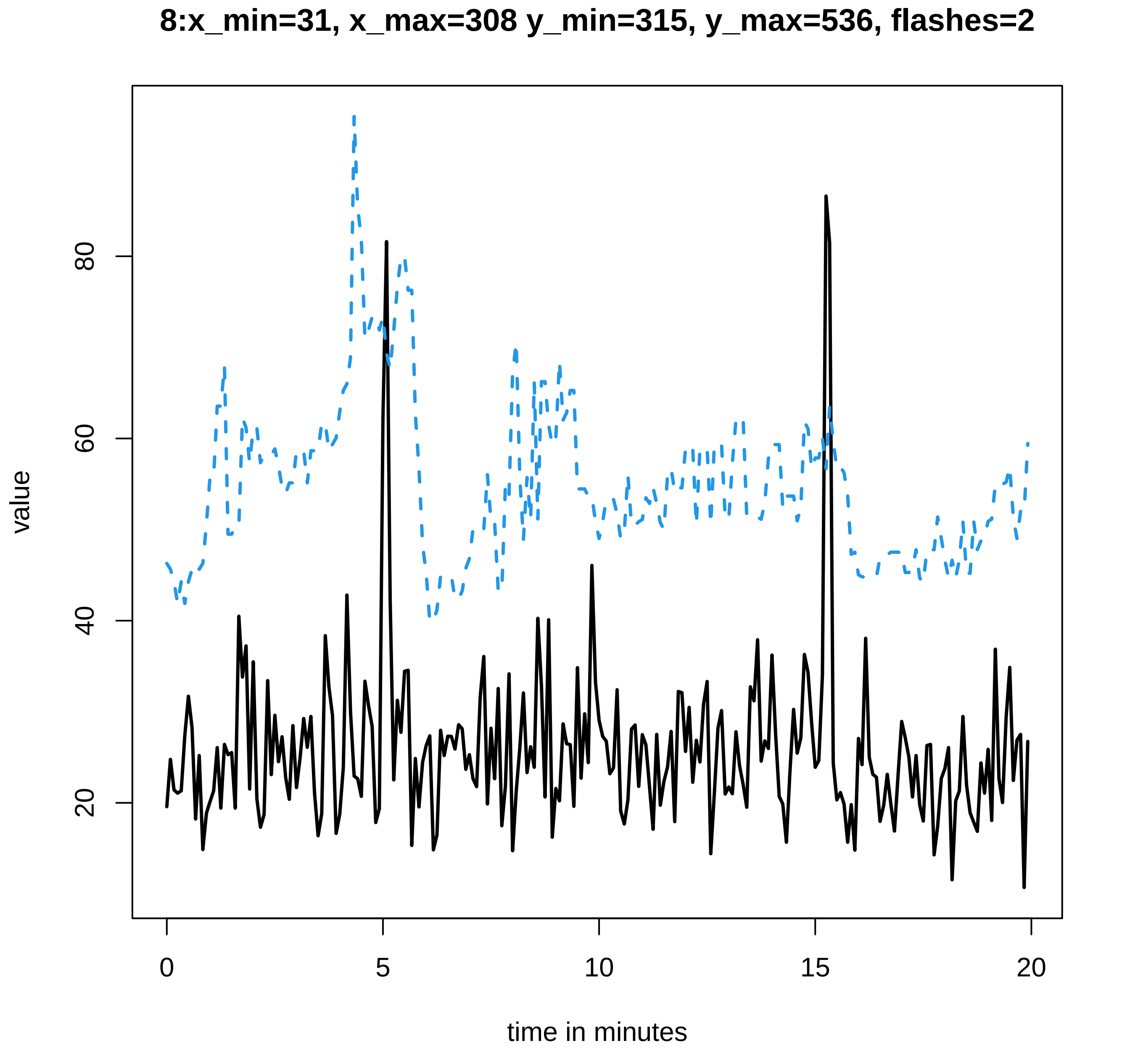
8:x_min=31, x_max=308 y_min=315, y_max=536, flashes=2
value
20
40
60
80
0
5
10
15
20
time in minutes

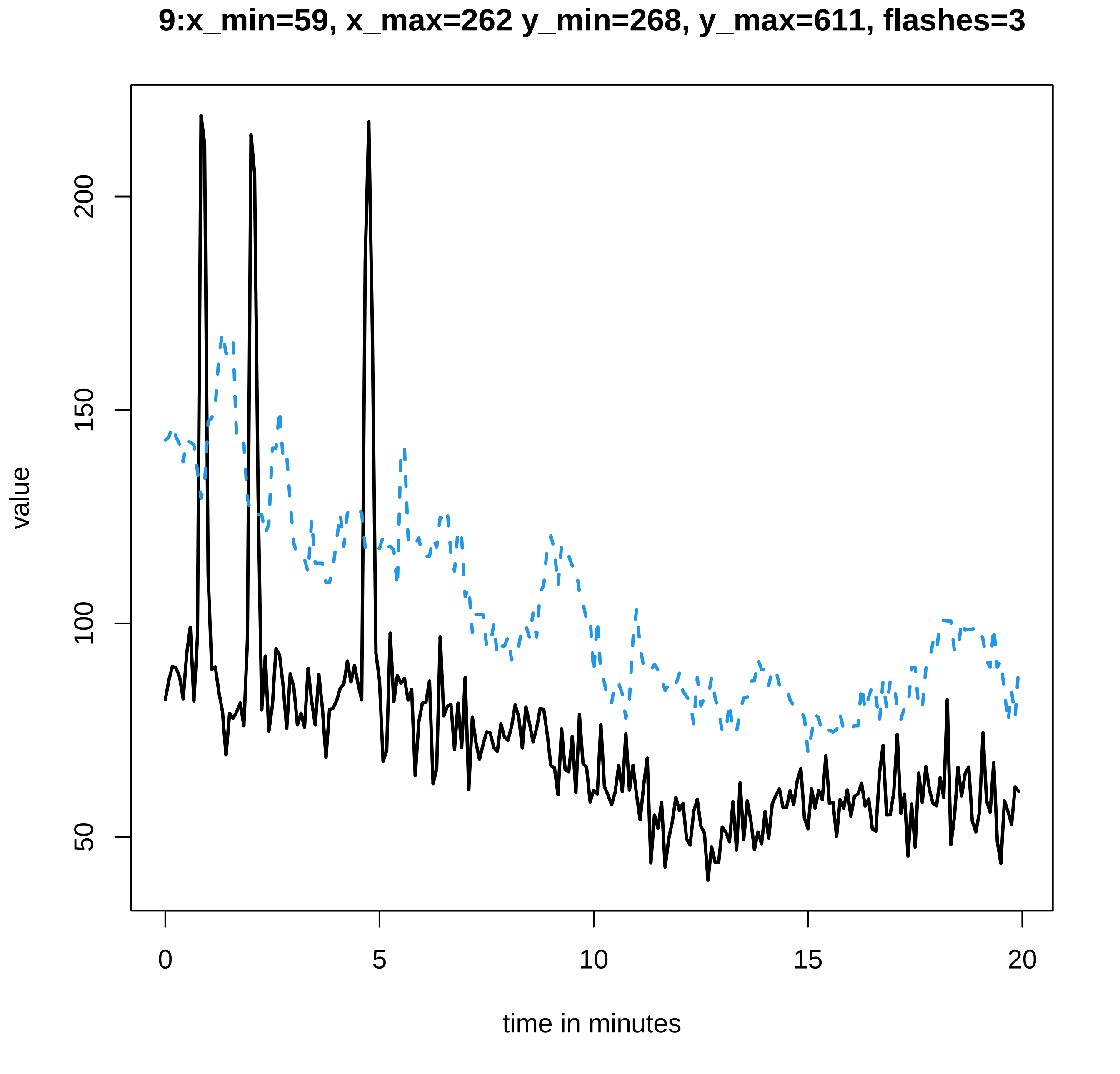

9:x_min=59, x_max=262 y_min=268, y_max=611, flashes=3
value
200
150
100
50
0
5
10
15
20
time in minutes

### 10:x_min=41, x_max=235 y_min=245, y_max=631, flashes=4

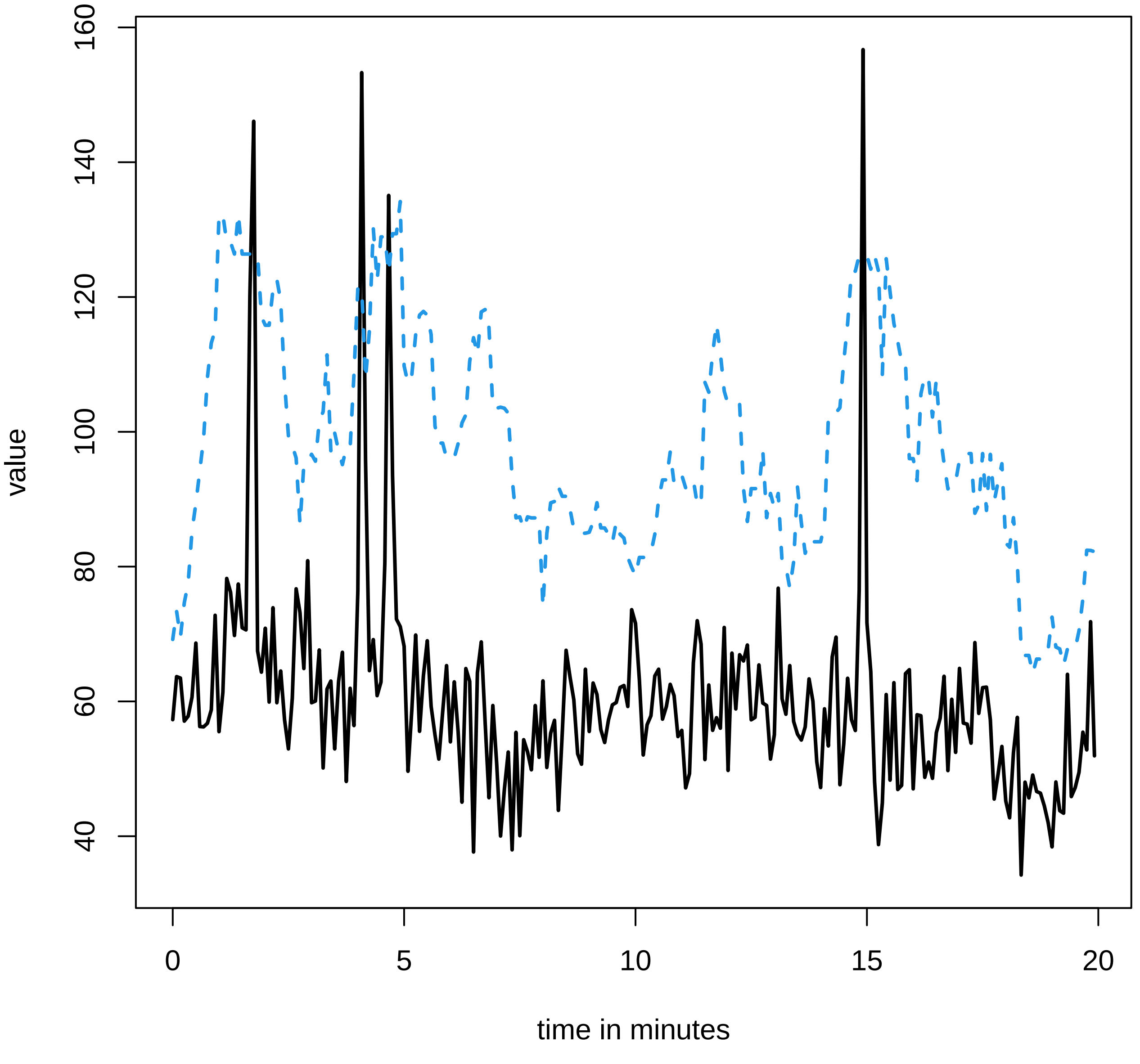

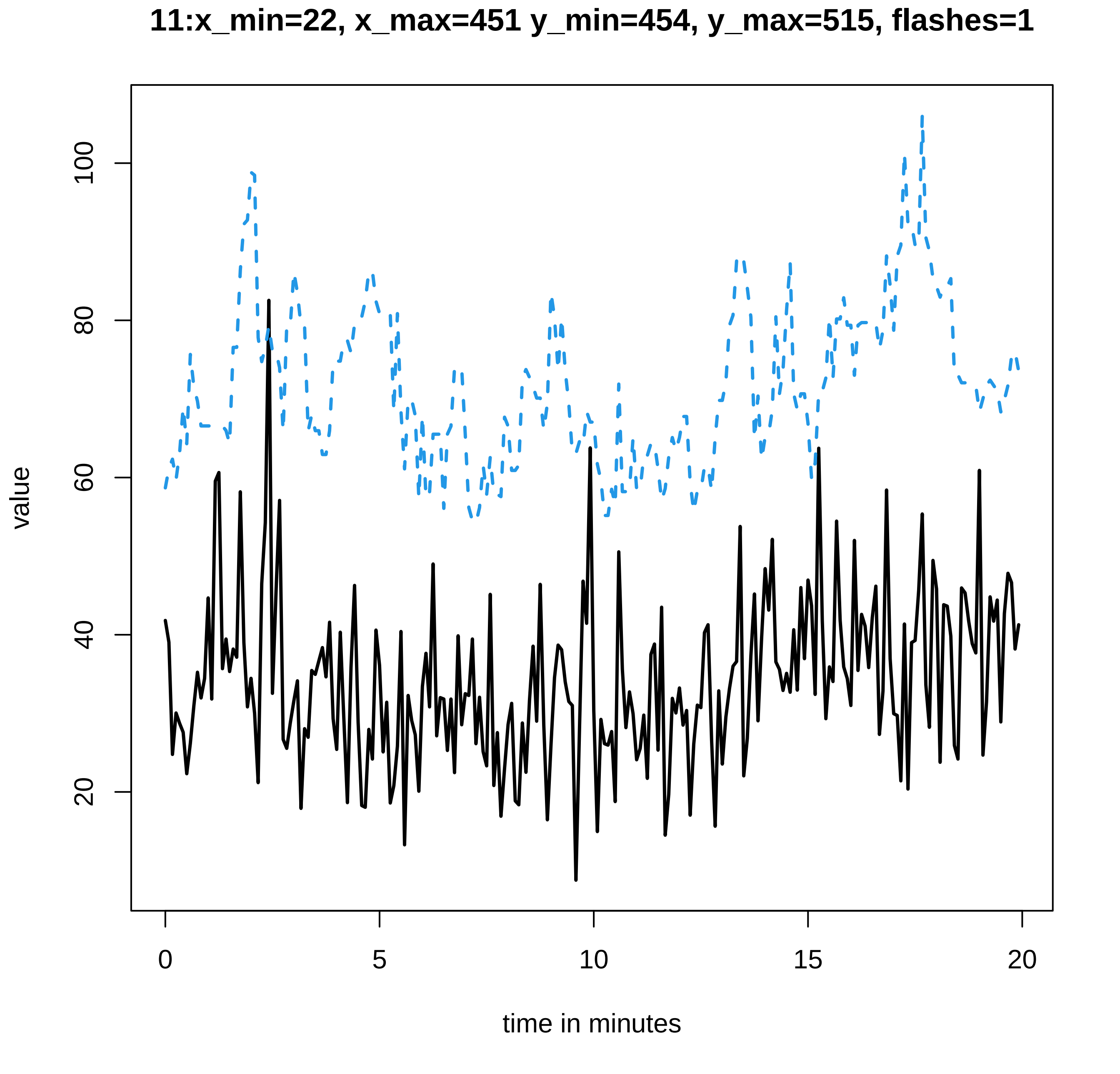
11:x_min=22, x_max=451 y_min=454, y_max=515, flashes=1
value
20
40
60
80
100
0
5
10
15
20
time in minutes

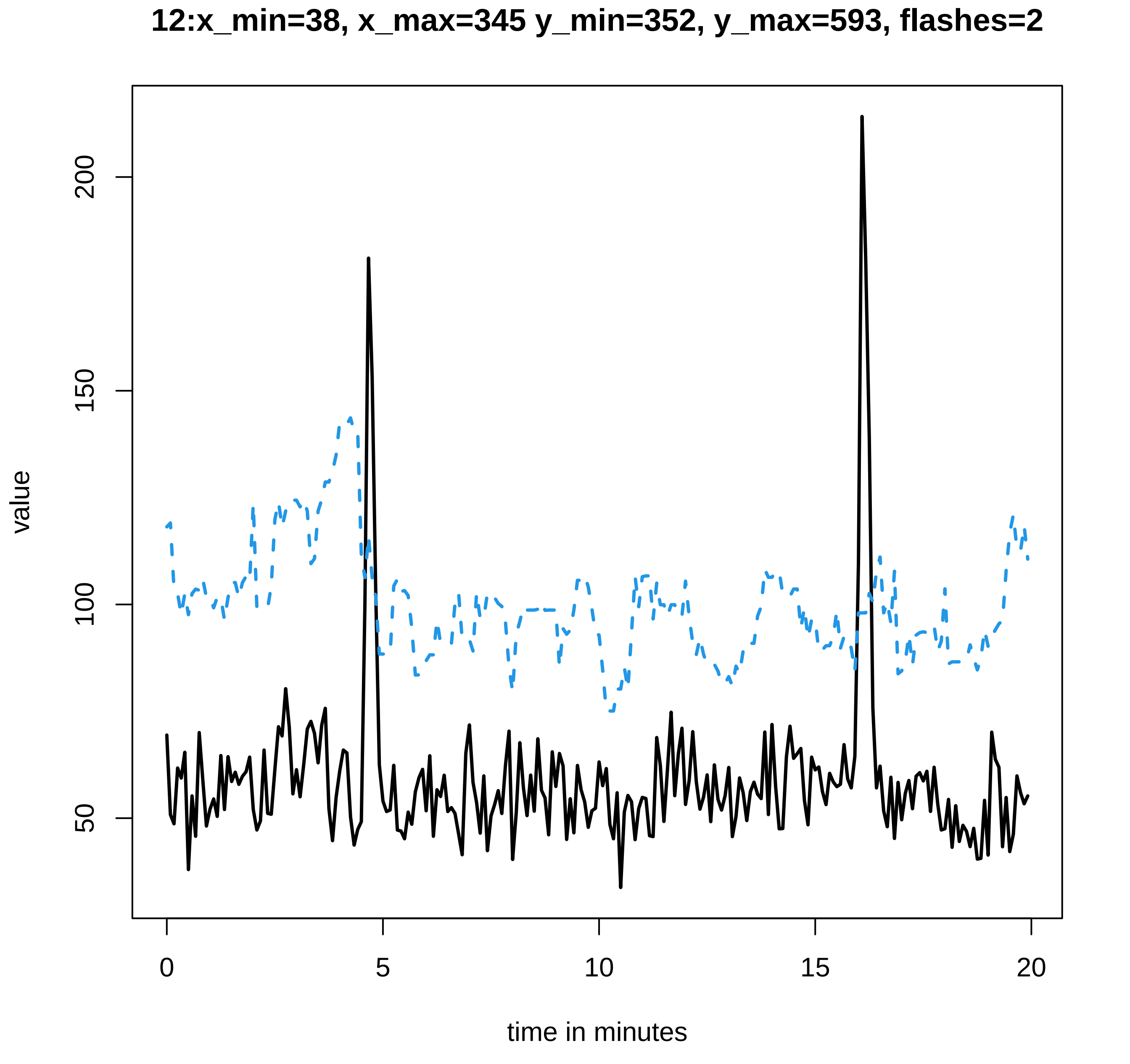
12:x_min=38, x_max=345 y_min=352, y_max=593, flashes=2
value
200
150
100
50
0
5
10
15
20
time in minutes

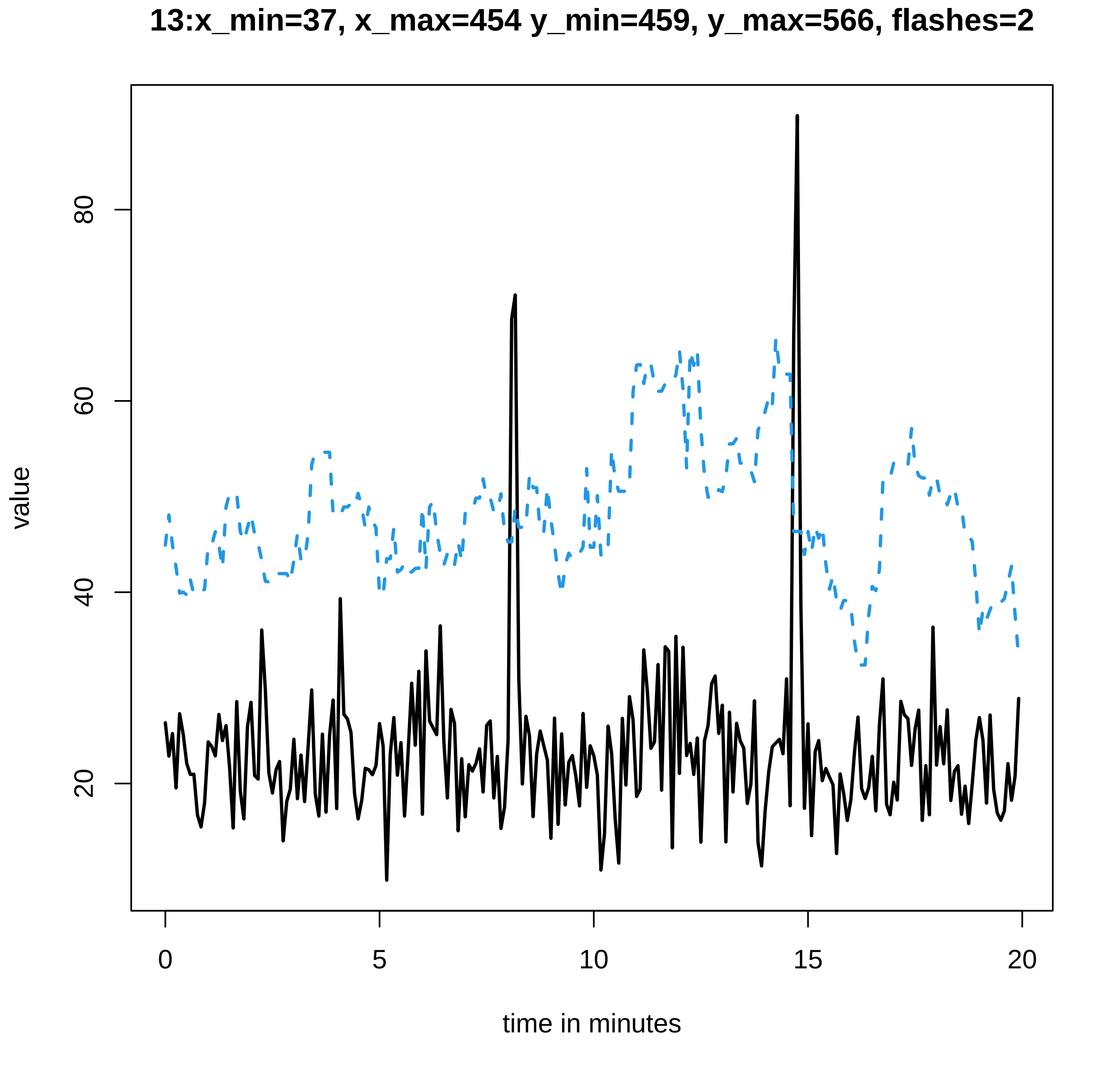

13:x_min=37, x_max=454 y_min=459, y_max=566, flashes=2
value
20
40
60
80
0
5
10
15
20
time in minutes

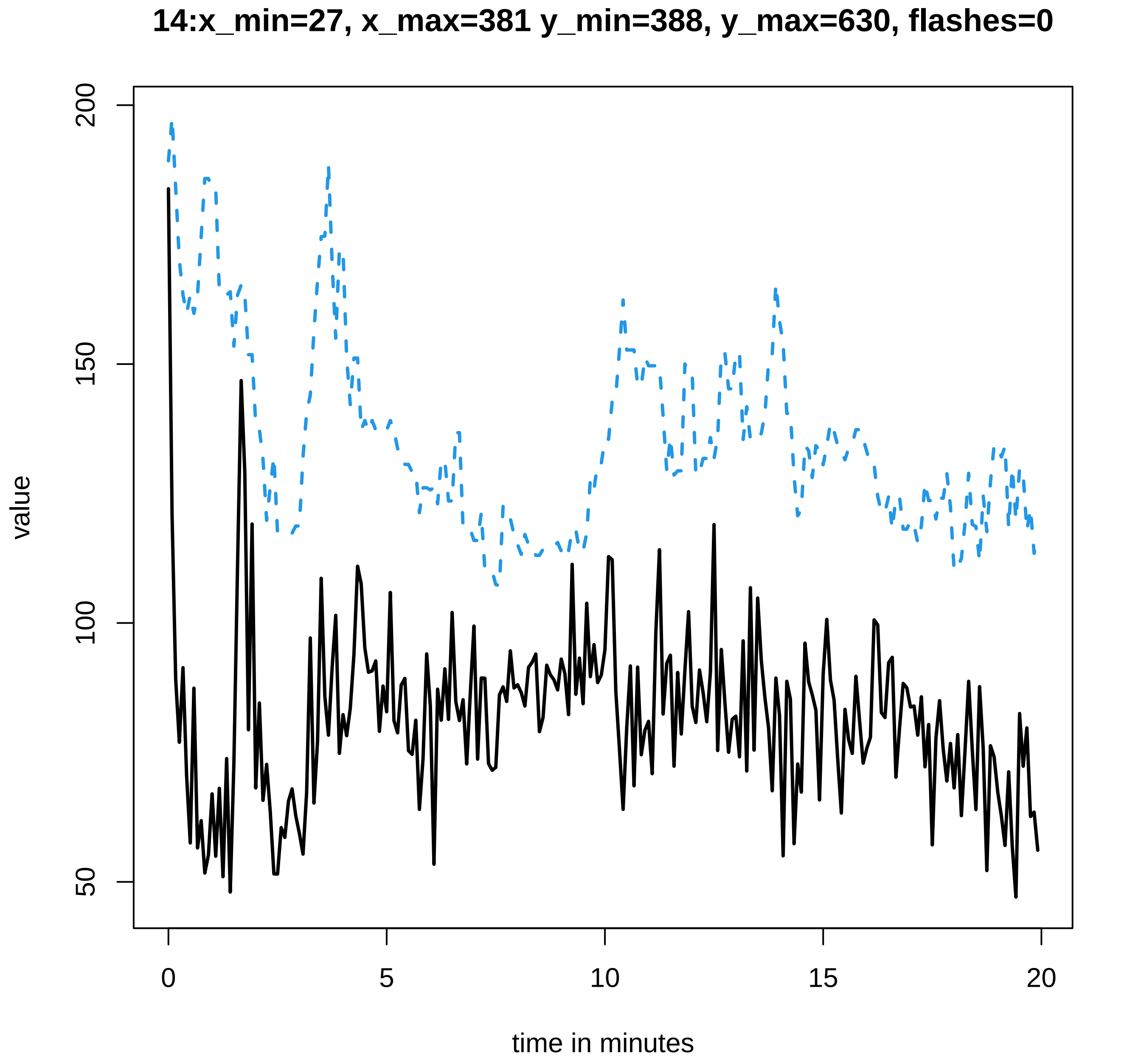
14:x_min=27, x_max=381 y_min=388, y_max=630, flashes=0
value
200
150
100
50
0
5
10
15
20
time in minutes

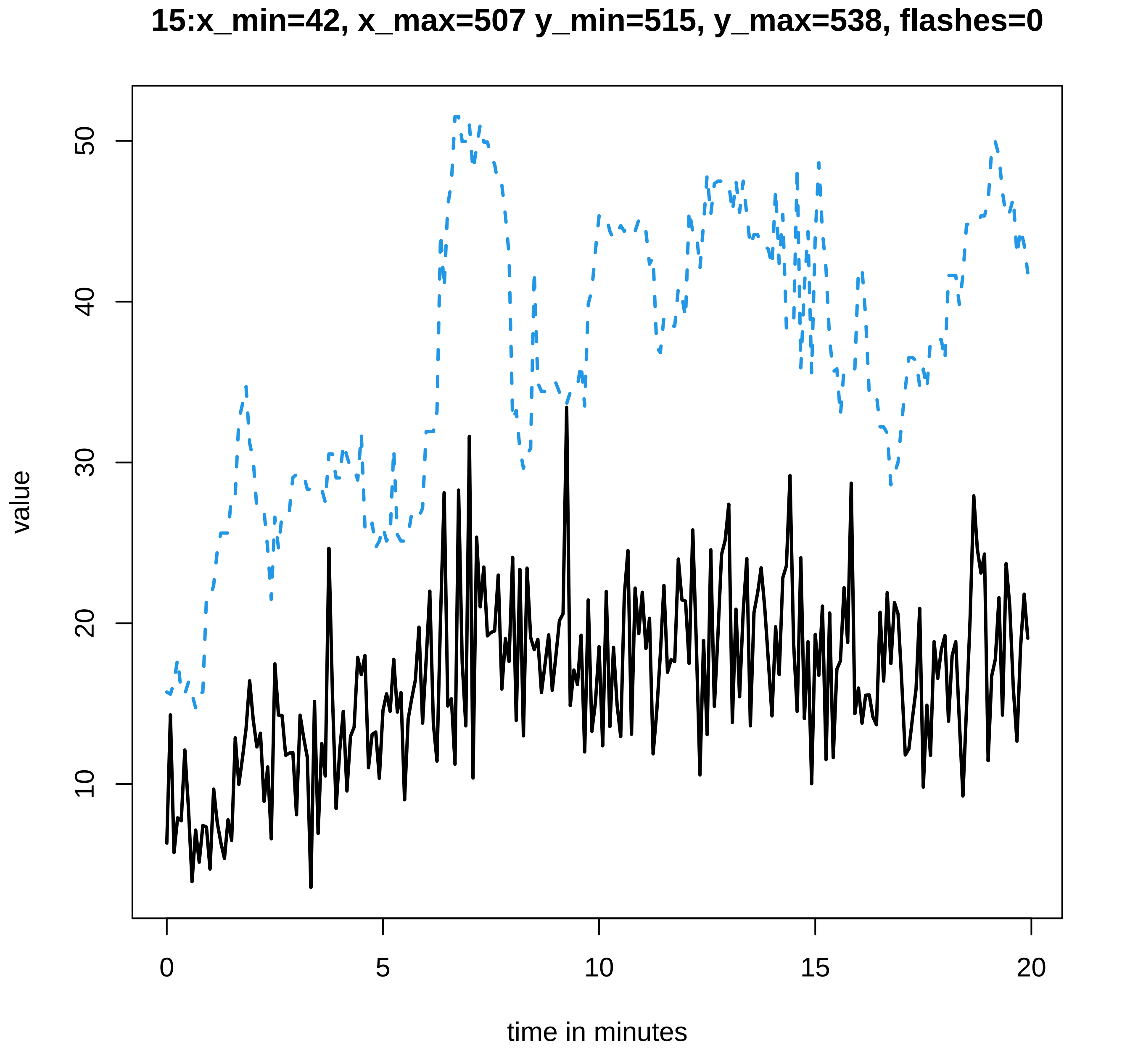
15:x_min=42, x_max=507 y_min=515, y_max=538, flashes=0
value
10
20
30
40
50
0
5
10
15
20
time in minutes

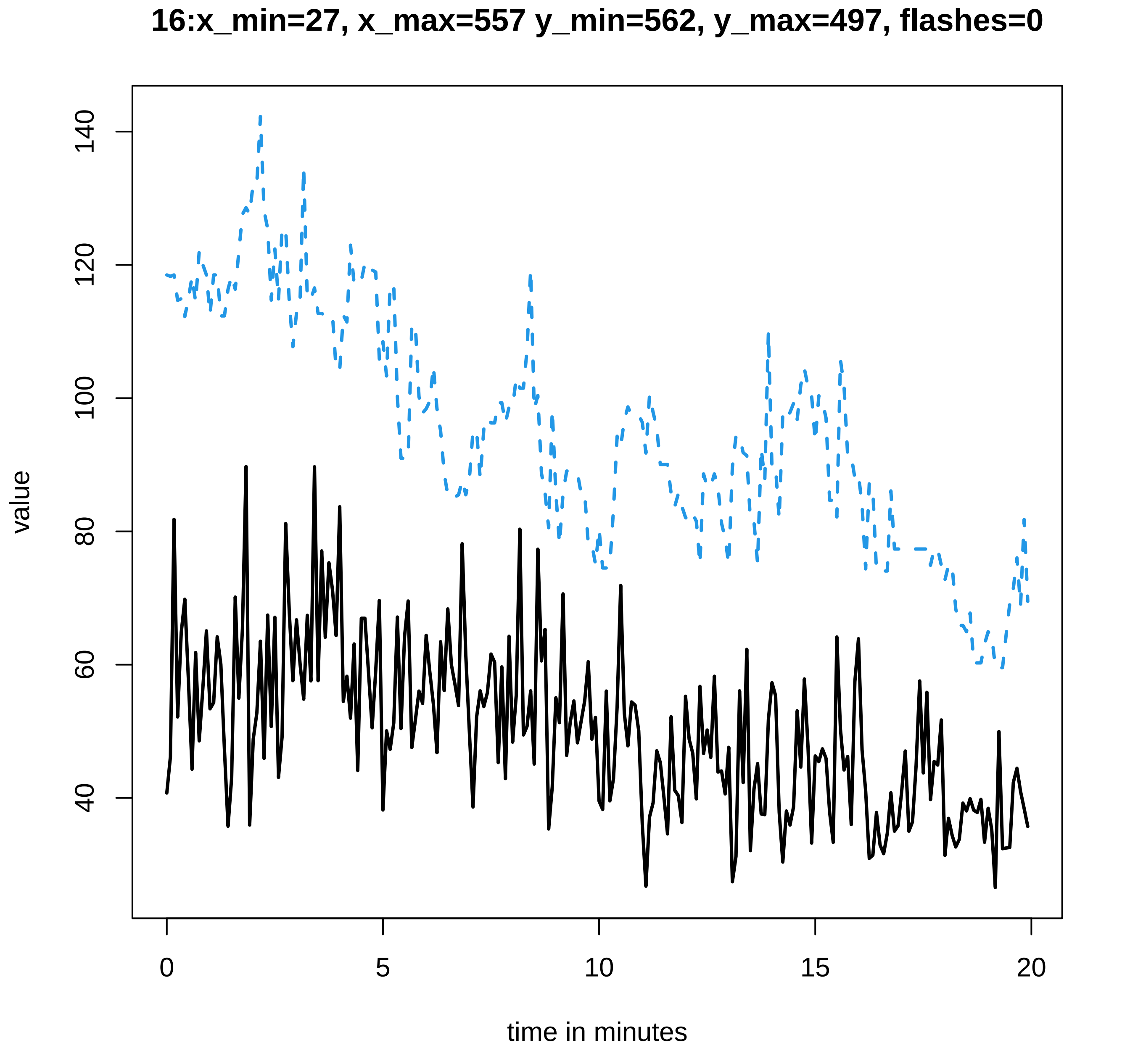
16:x_min=27, x_max=557 y_min=562, y_max=497, flashes=0
value
140
120
100
80
60
40
0
5
10
15
20
time in minutes

### 17:x_min=39, x_max=368 y_min=377, y_max=656, flashes=3

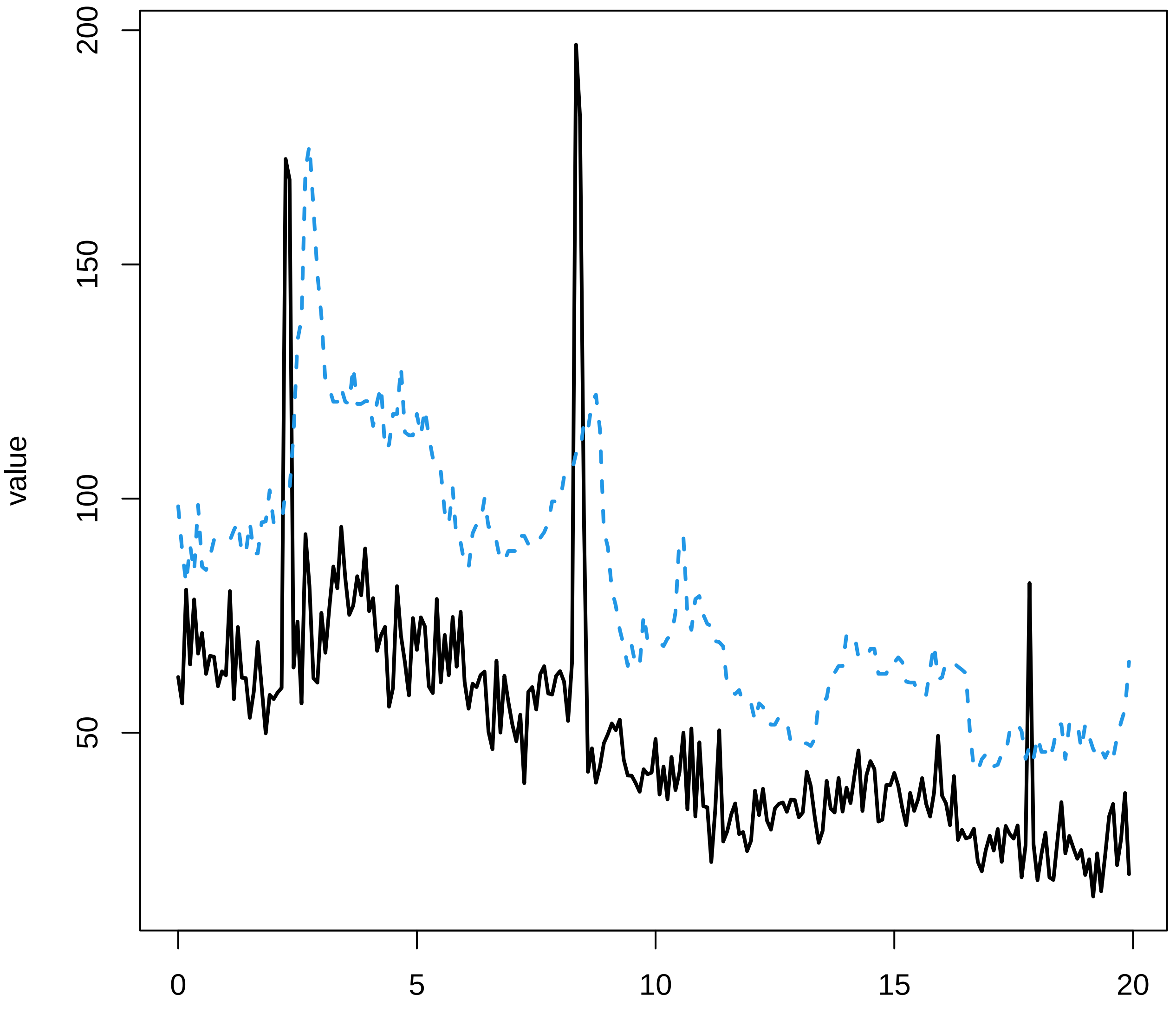

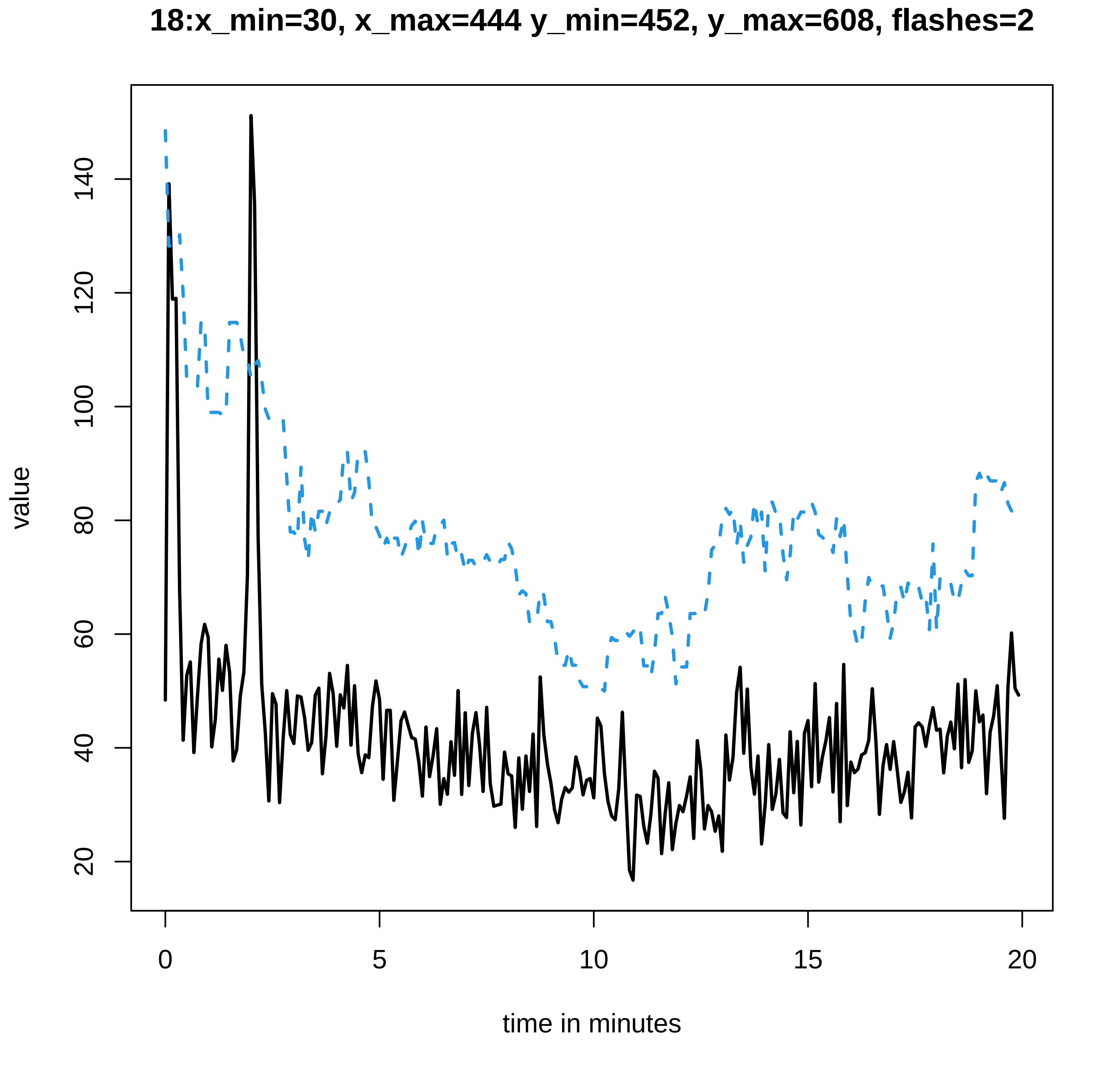

18:x_min=30, x_max=444 y_min=452, y_max=608, flashes=2
value
20
40
60
80
100
120
140
0
5
10
15
20
time in minutes

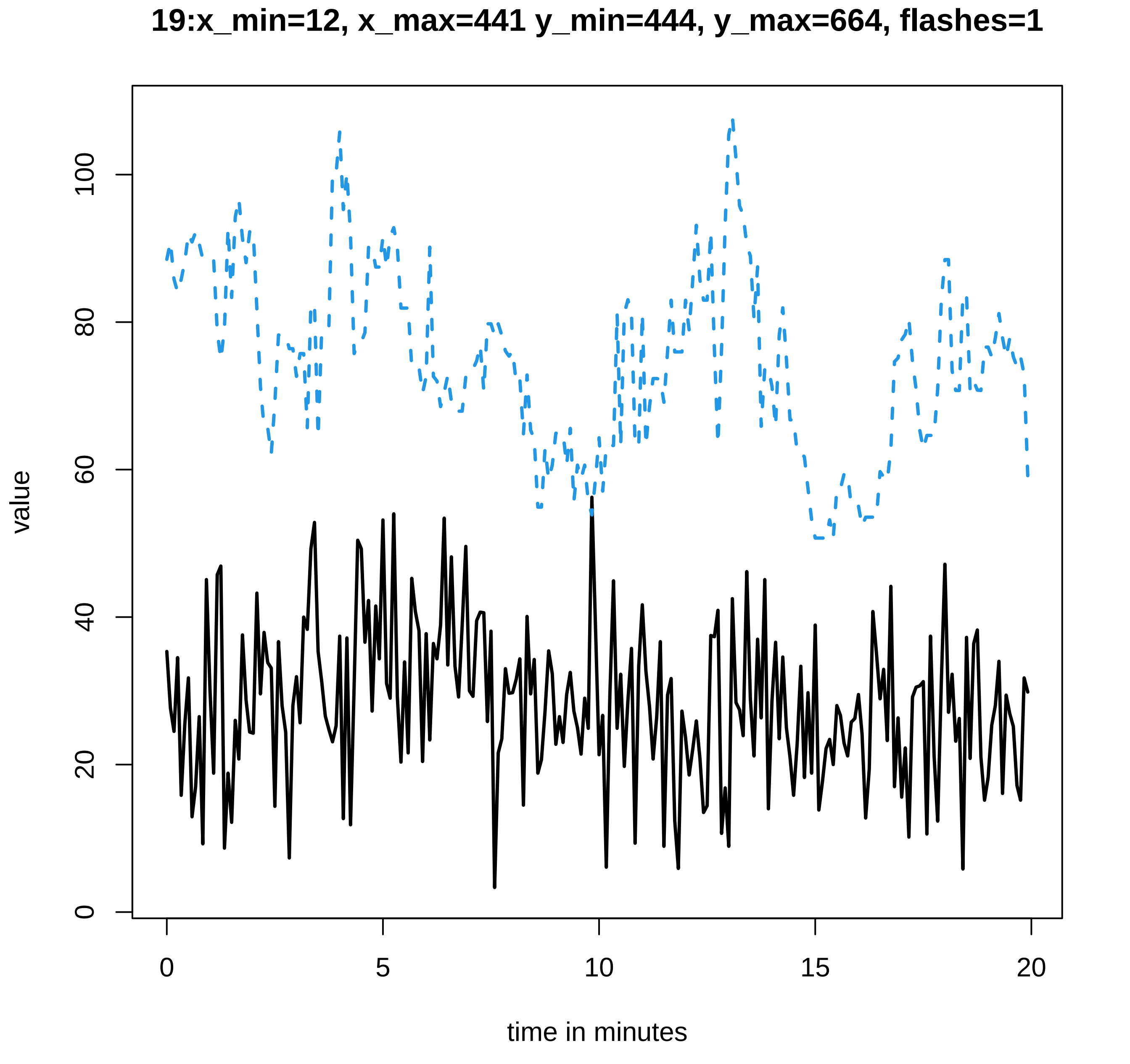

19:x_min=12, x_max=441 y_min=444, y_max=664, flashes=1
value
0
20
40
60
80
100
0
5
10
15
20
time in minutes

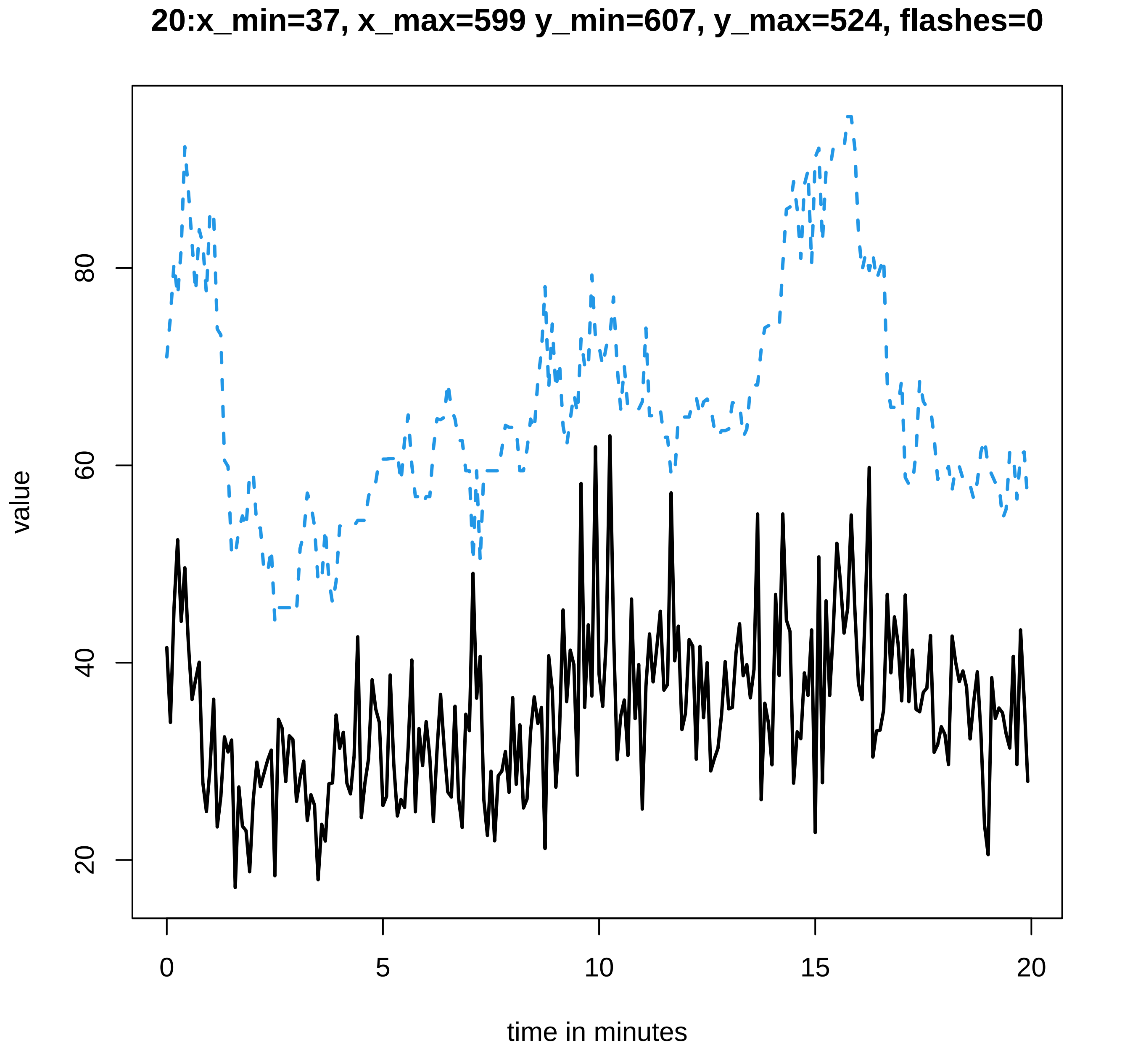
20:x_min=37, x_max=599 y_min=607, y_max=524, flashes=0
value
20
40
60
80
time in minutes
0
5
10
15
20

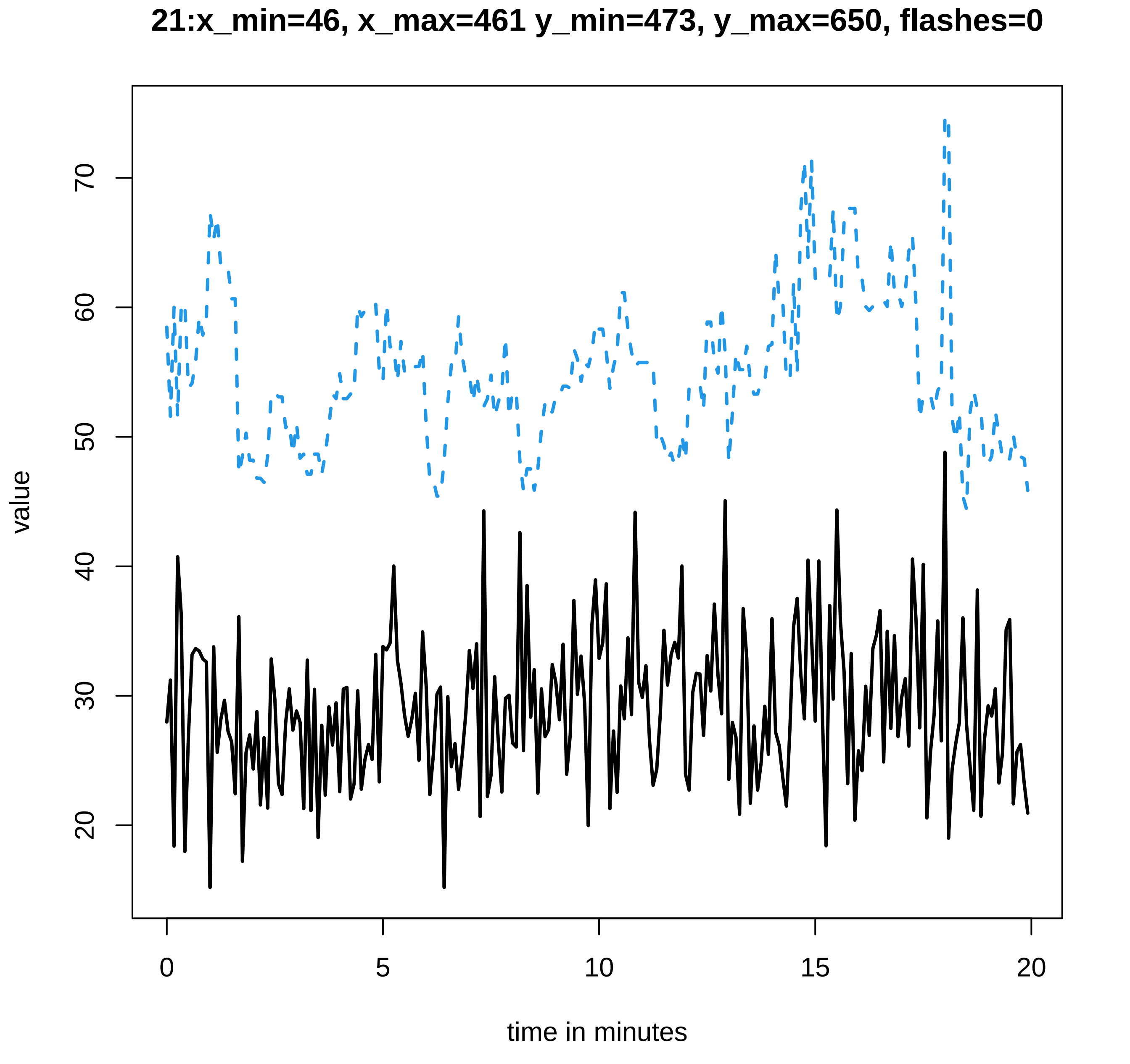
21:x_min=46, x_max=461 y_min=473, y_max=650, flashes=0
value
time in minutes
0
5
10
15
20
20
30
40
50
60
70

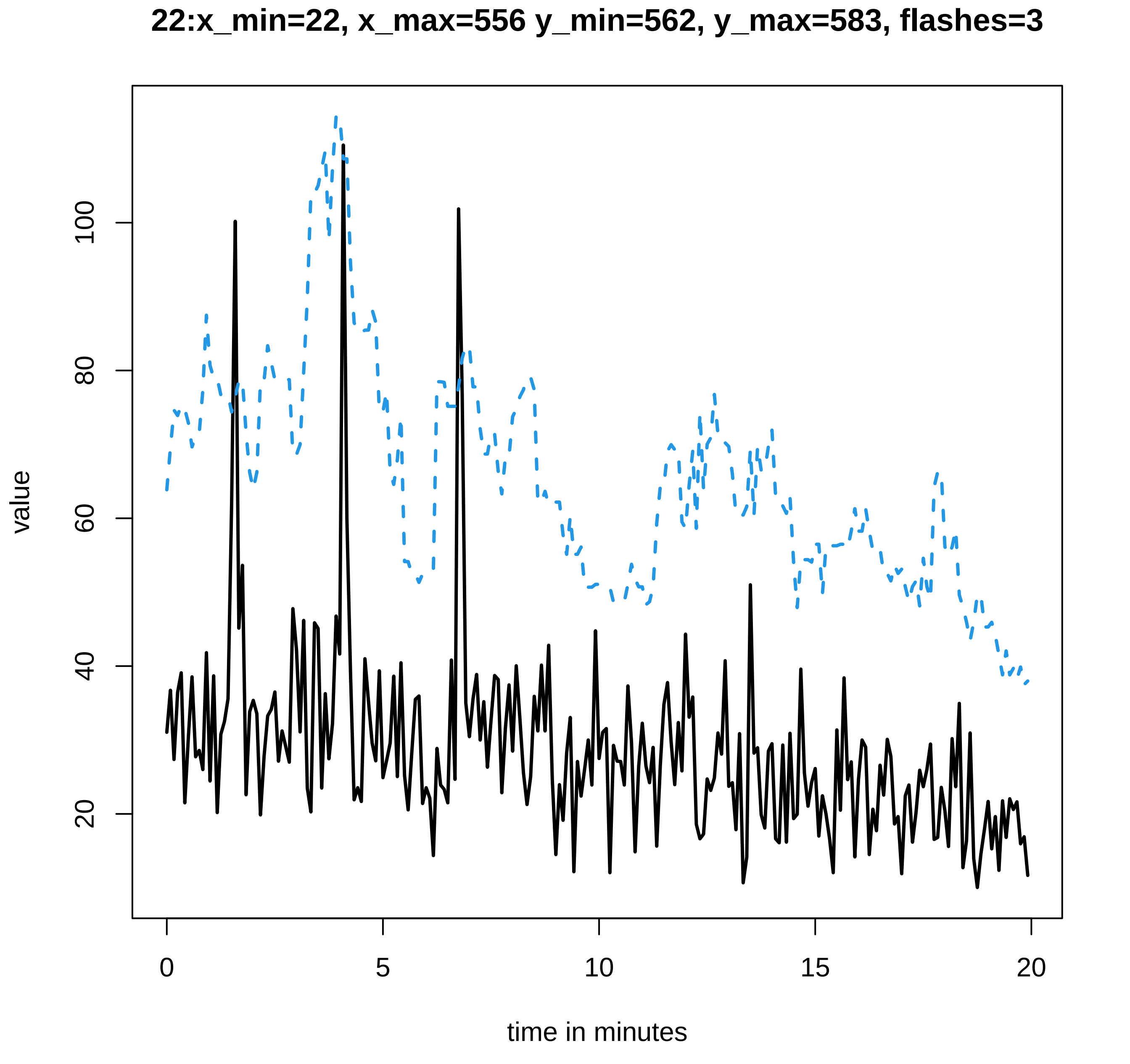
22:x_min=22, x_max=556 y_min=562, y_max=583, flashes=3
value
20
40
60
80
100
0
5
10
15
20
time in minutes

**23:x_min=36, x_max=496 y_min=504, y_max=641, flashes=4**

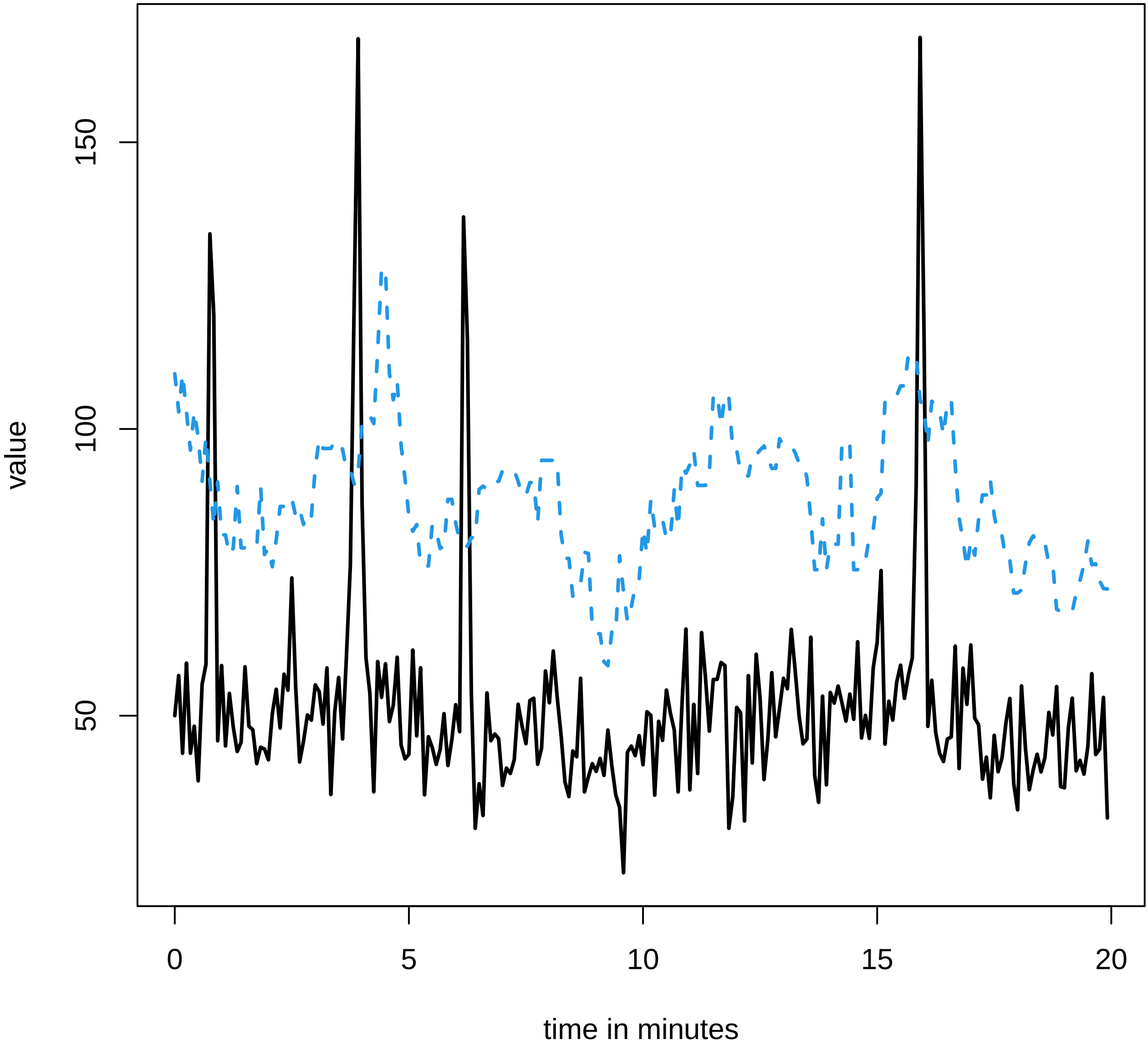

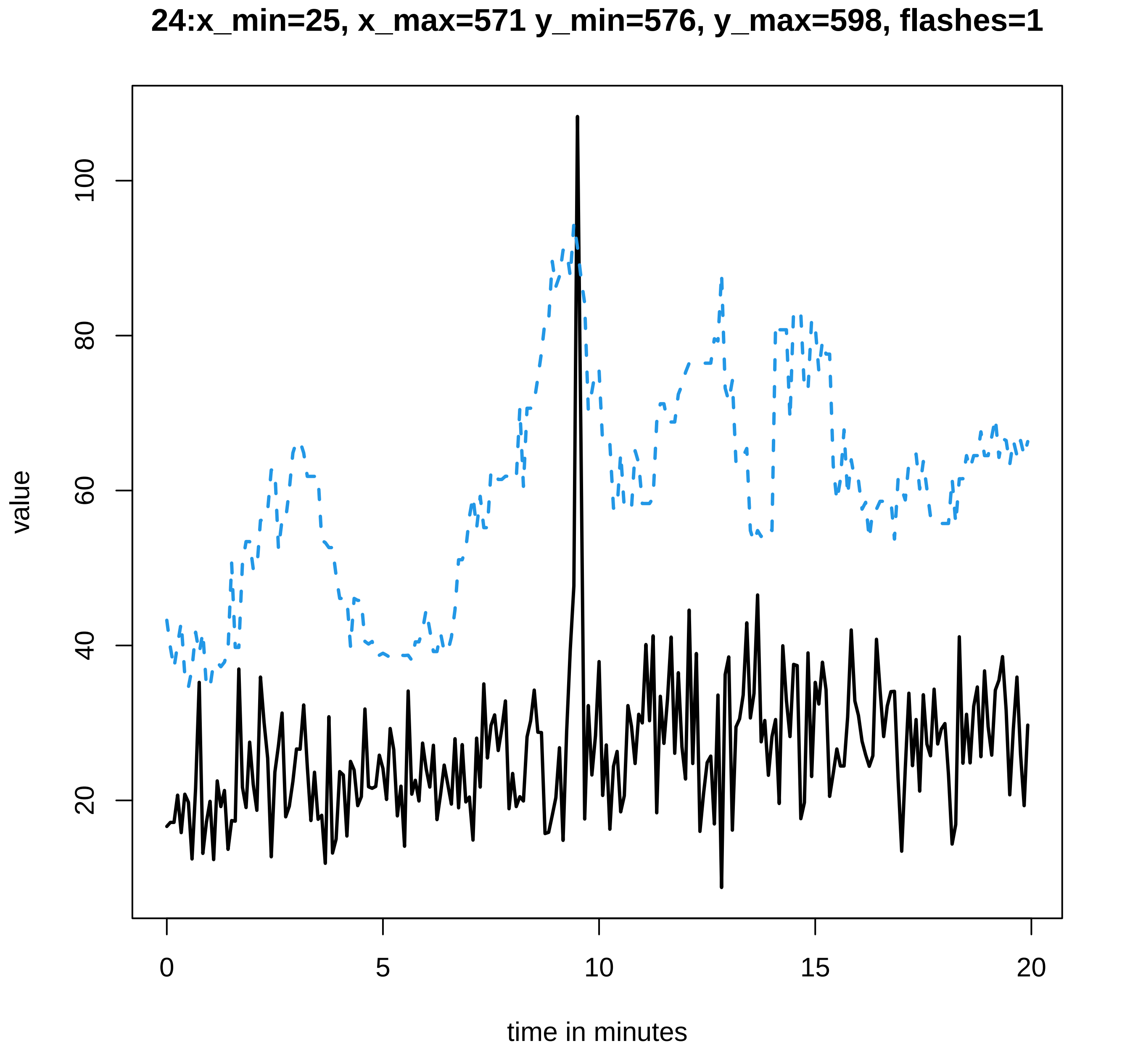
24:x_min=25, x_max=571 y_min=576, y_max=598, flashes=1
value
100
80
60
40
20
0
5
10
15
20
time in minutes

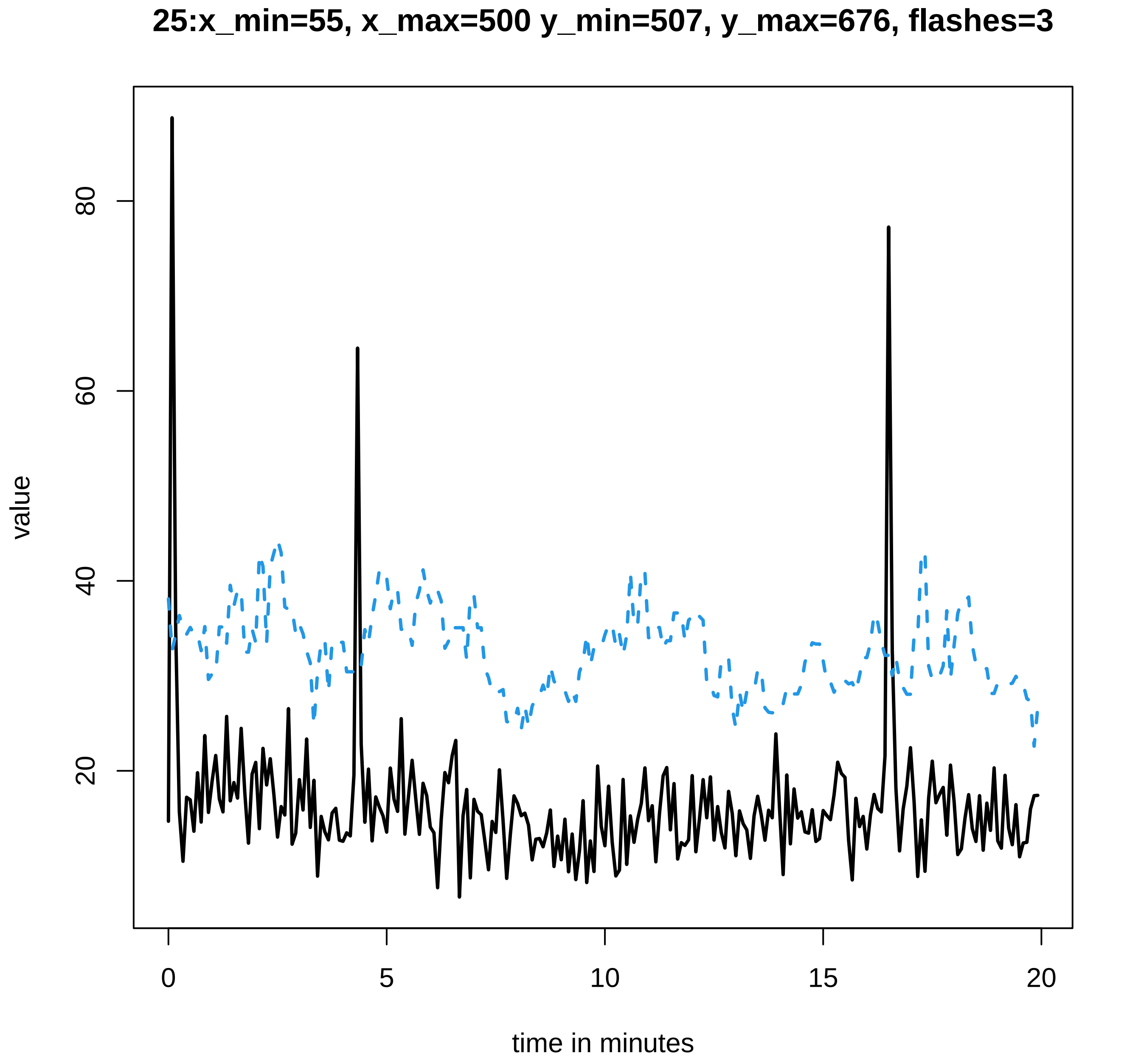
25:x_min=55, x_max=500 y_min=507, y_max=676, flashes=3
value
20
40
60
80
0
5
10
15
20
time in minutes

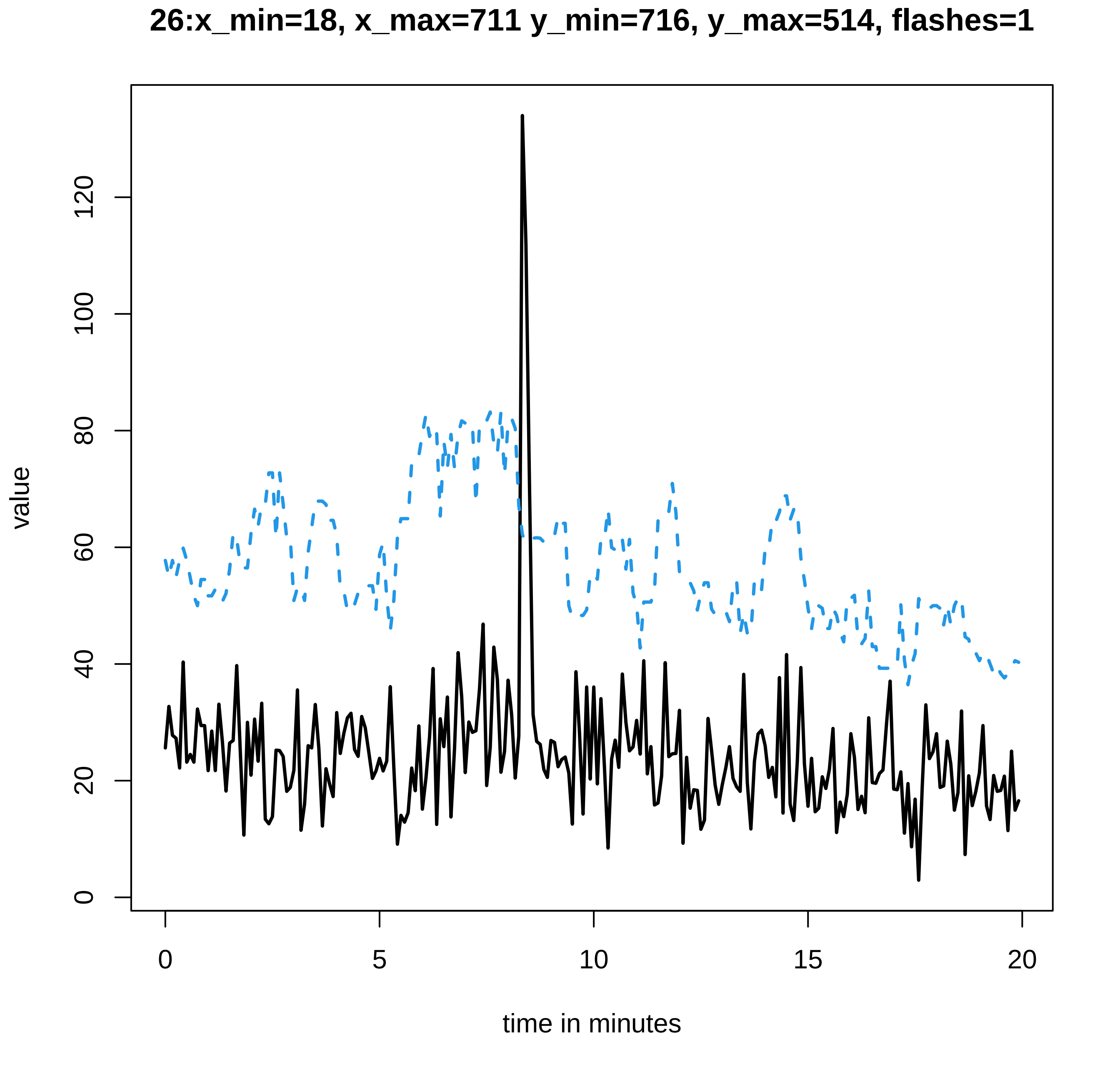

26:x_min=18, x_max=711 y_min=716, y_max=514, flashes=1
value
0
20
40
60
80
100
120
0
5
10
15
20
time in minutes

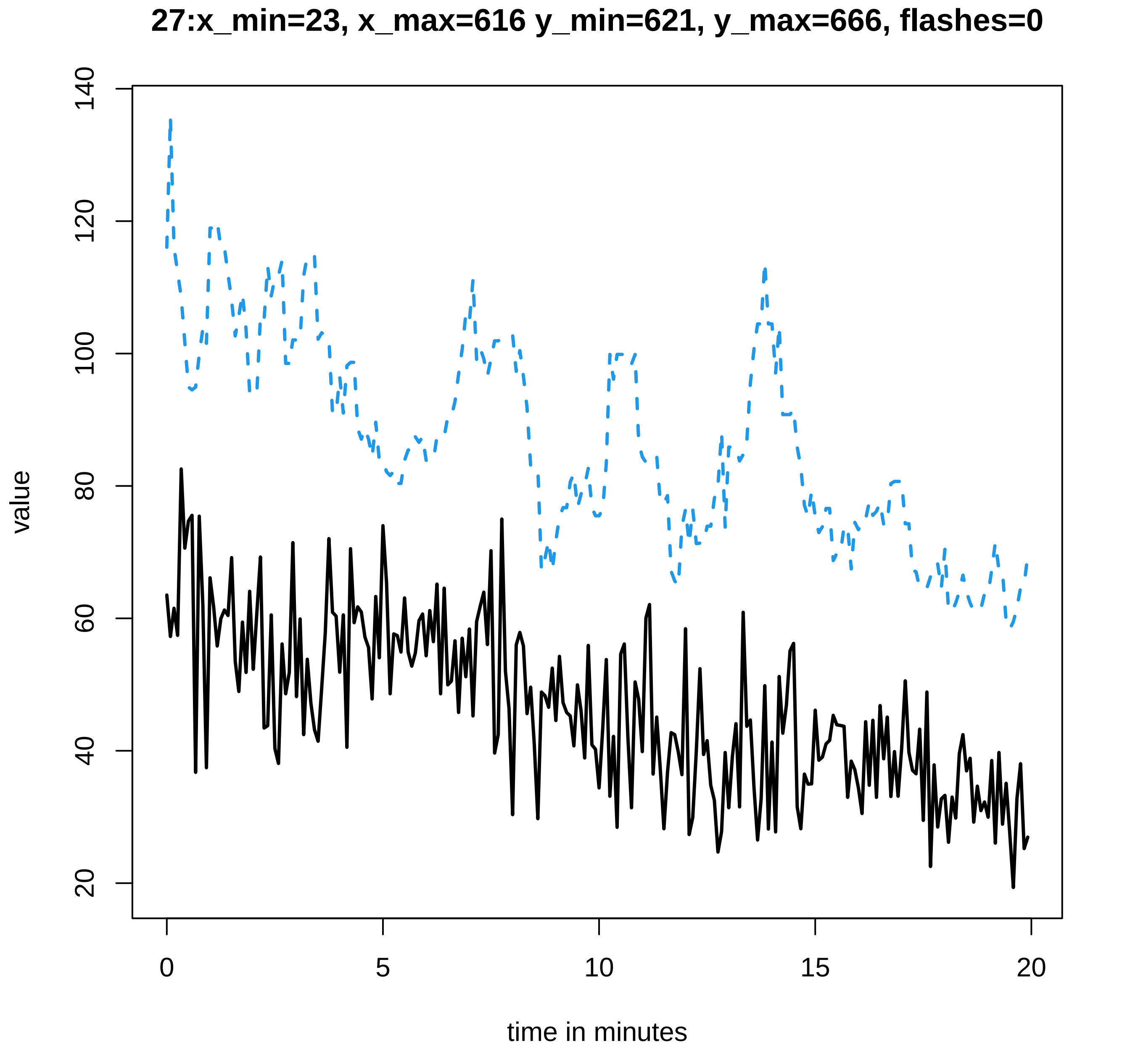
27:x_min=23, x_max=616 y_min=621, y_max=666, flashes=0
value
140
120
100
80
60
40
20
0
5
10
15
20
time in minutes

28:x_min=30, x_max=752 y_min=755, y_max=539, flashes=1

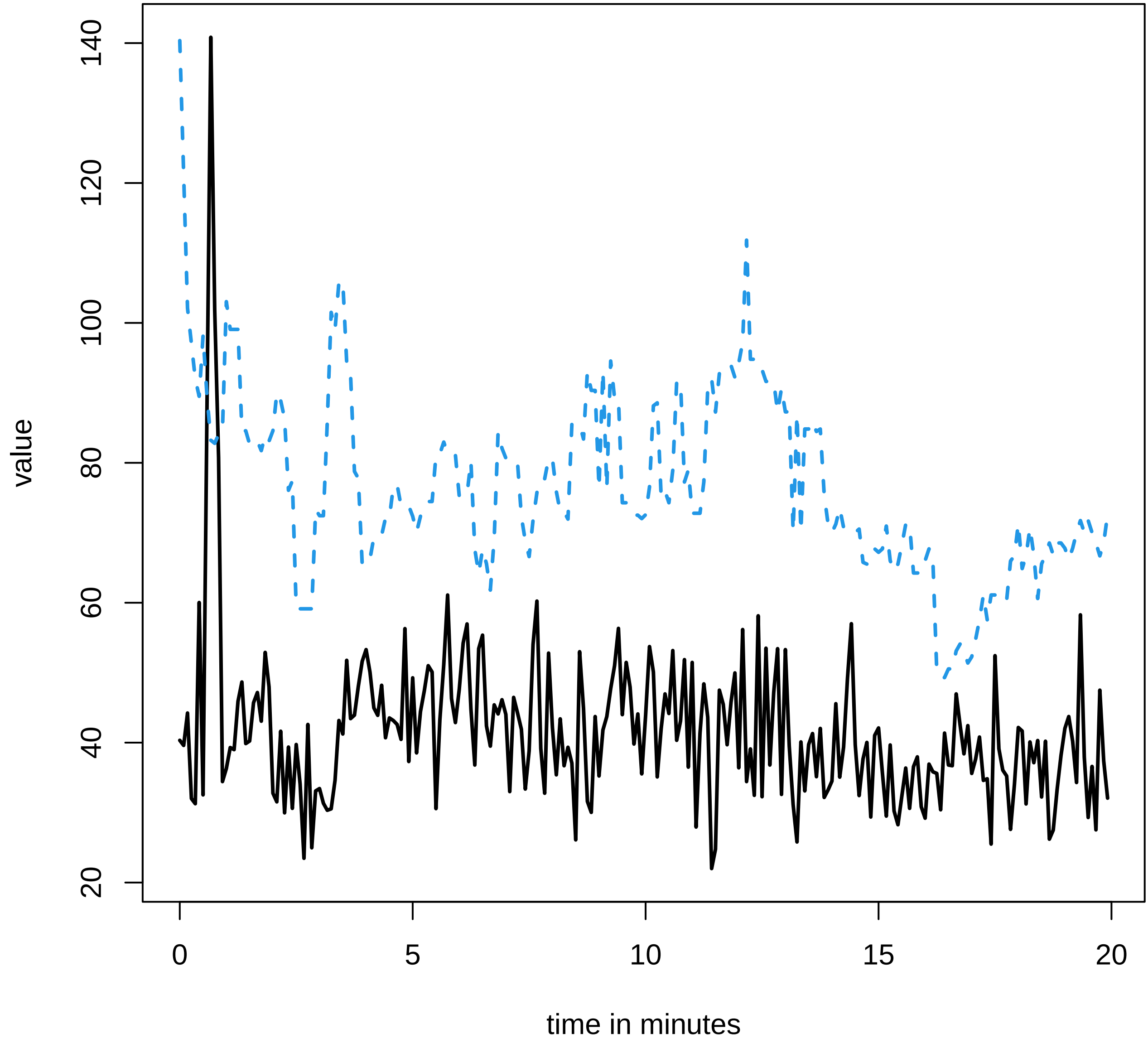

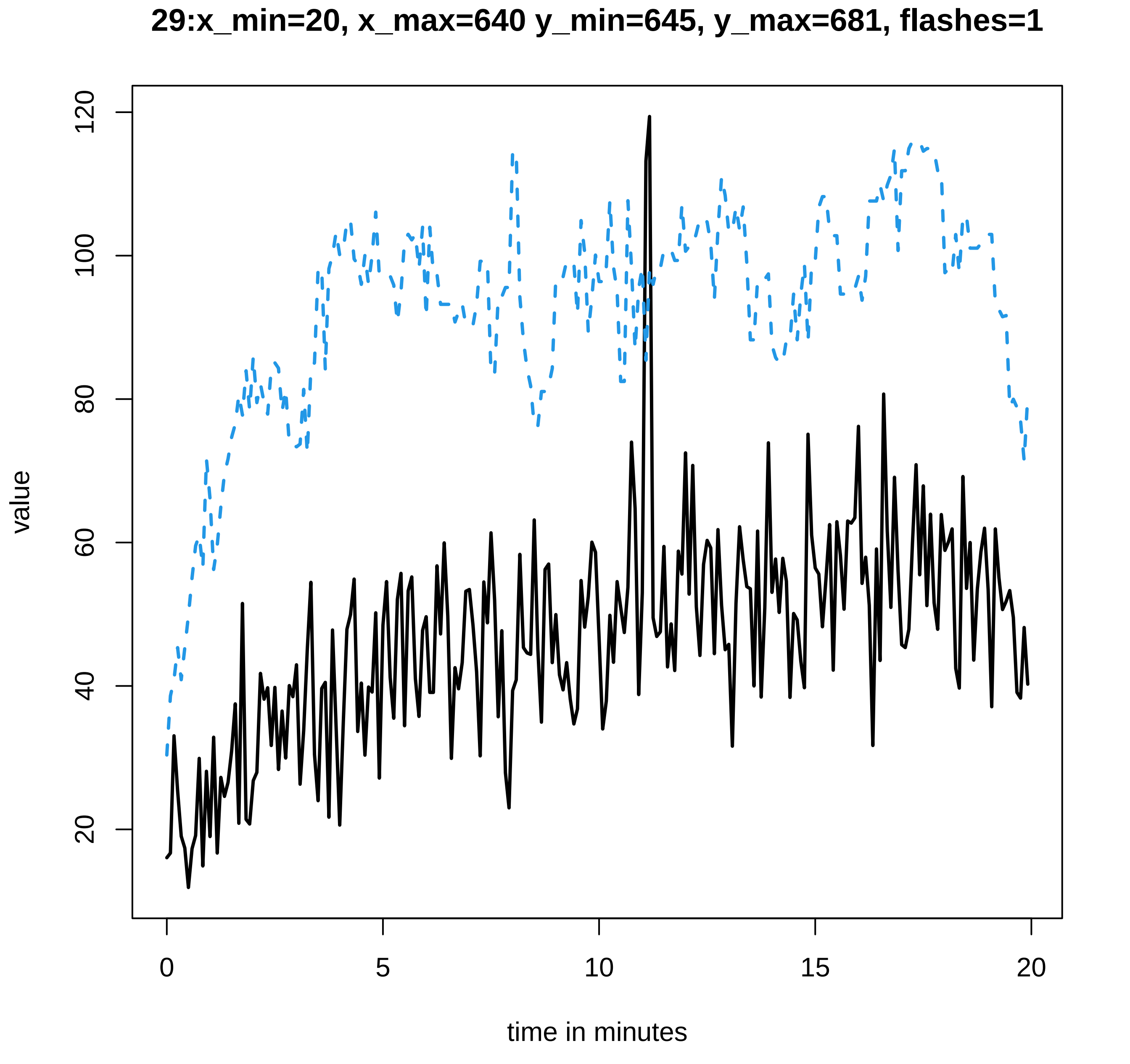
29:x_min=20, x_max=640 y_min=645, y_max=681, flashes=1
value
120
100
80
60
40
20
0
5
10
15
20
time in minutes

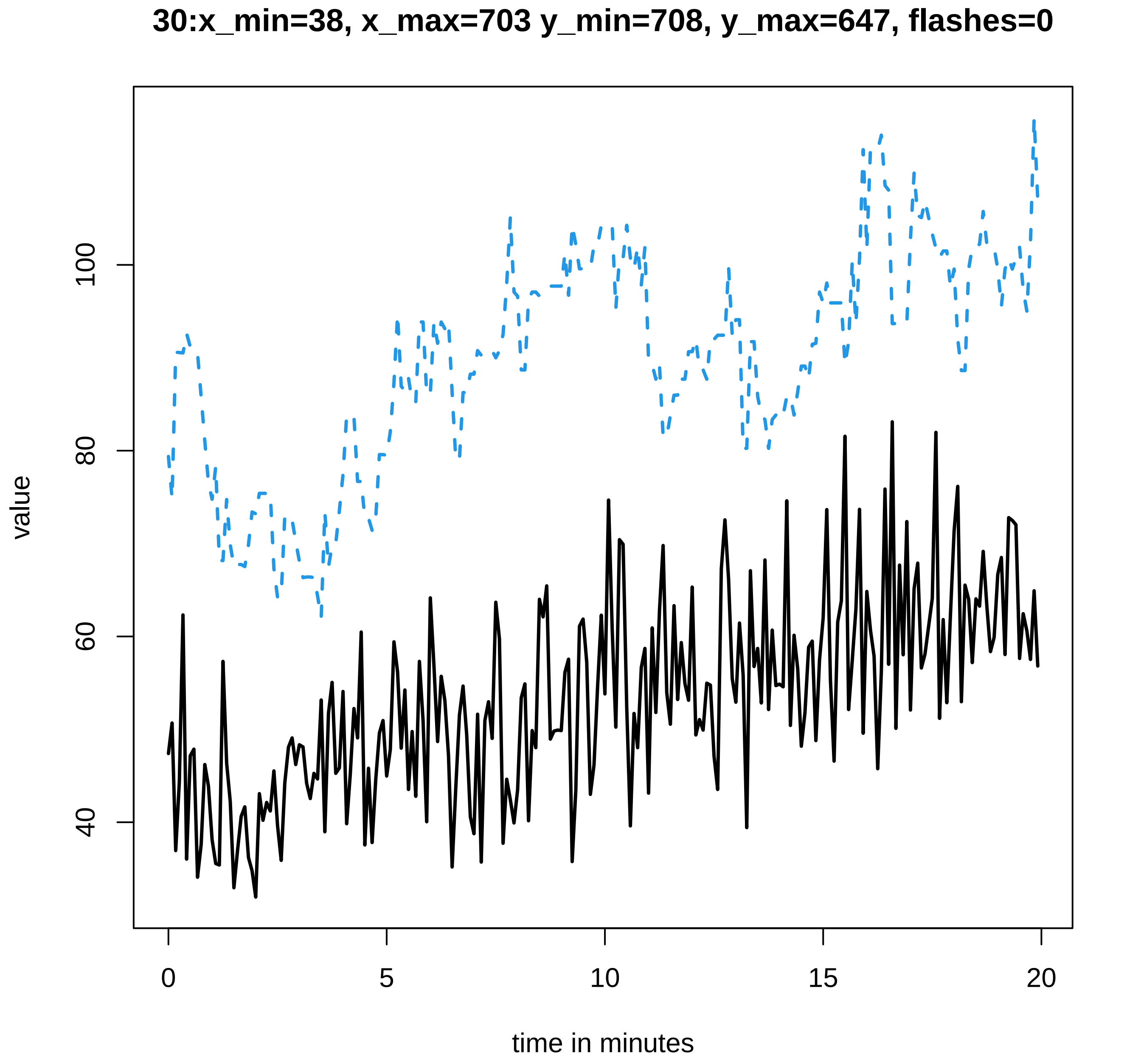
30:x_min=38, x_max=703 y_min=708, y_max=647, flashes=0
value
40
60
80
100
0
5
10
15
20
time in minutes

31:x_min=24, x_max=881 y_min=885, y_max=554, flashes=2

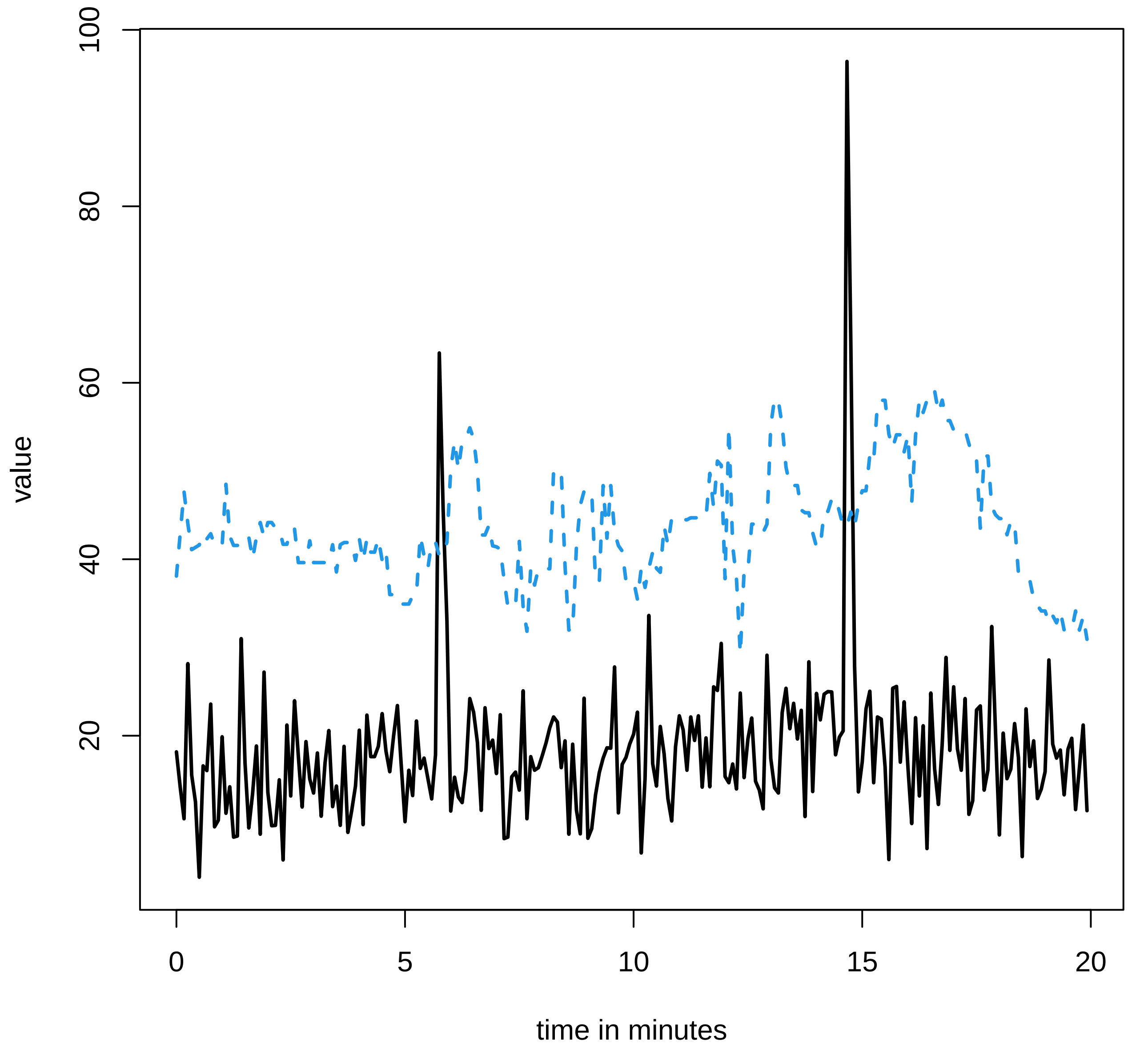

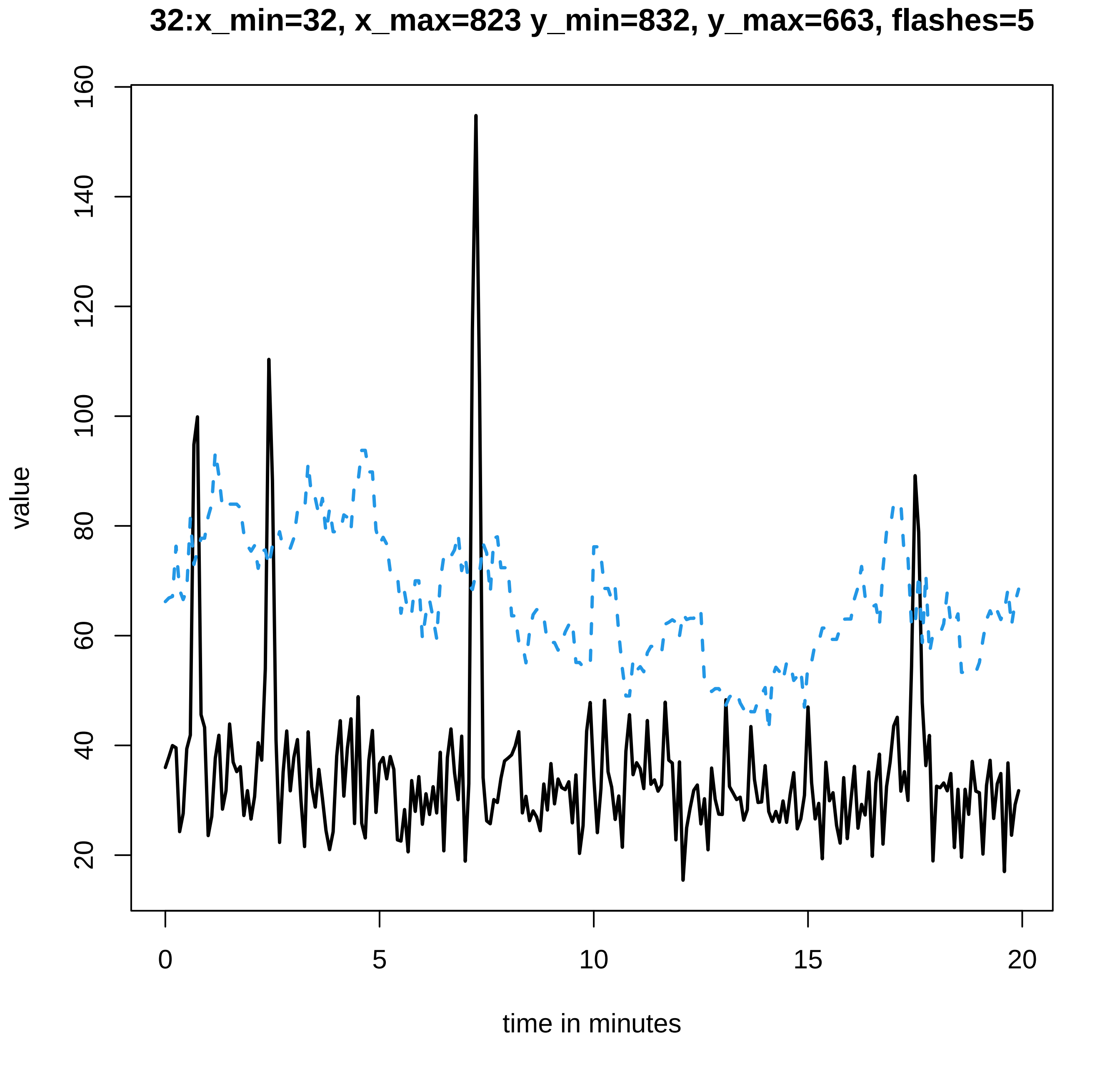

32:x_min=32, x_max=823 y_min=832, y_max=663, flashes=5
value
160
140
120
100
80
60
40
20
0
5
10
15
20
time in minutes

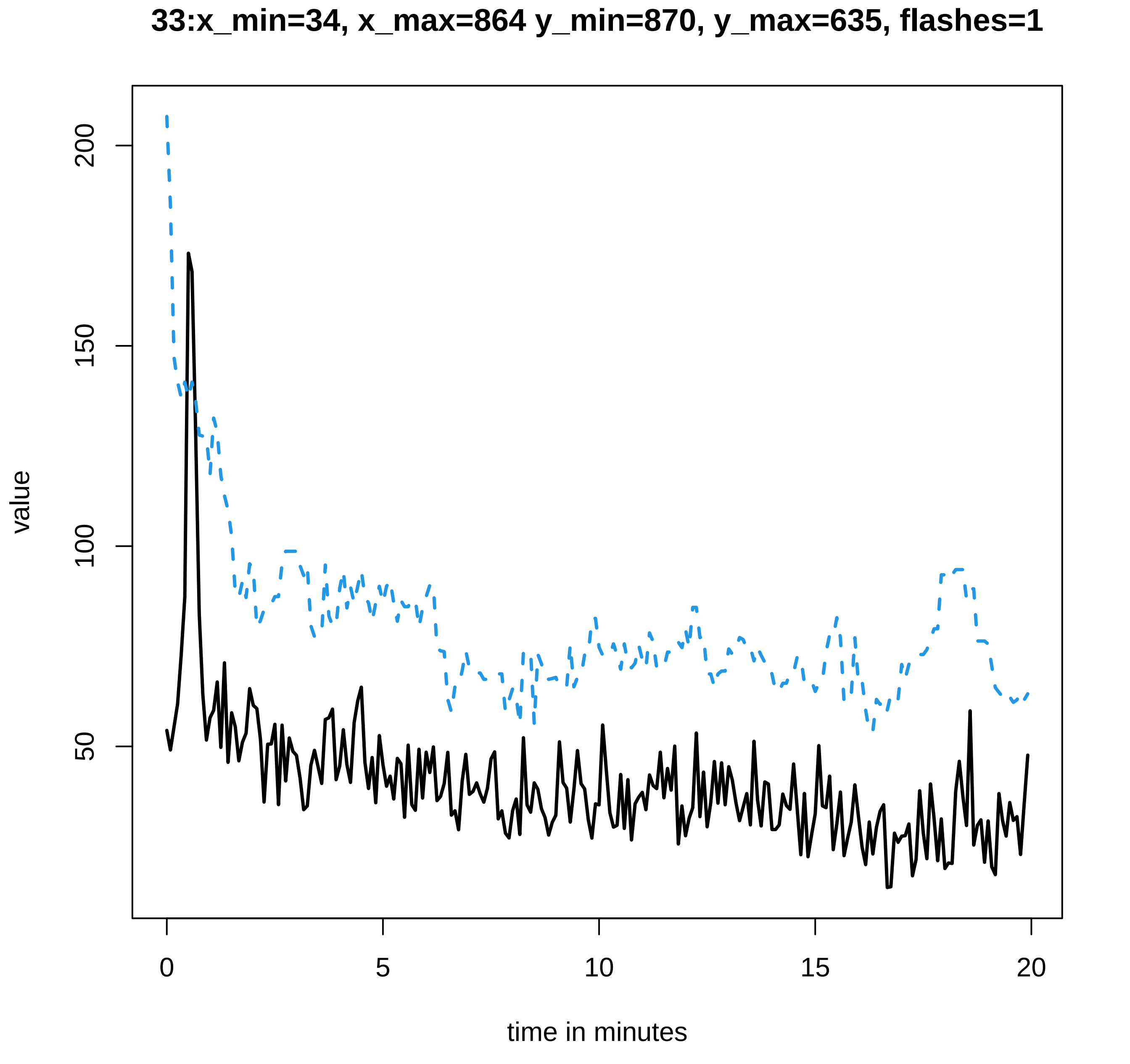

33:x_min=34, x_max=864 y_min=870, y_max=635, flashes=1
value
200
150
100
50
0
5
10
15
20
time in minutes

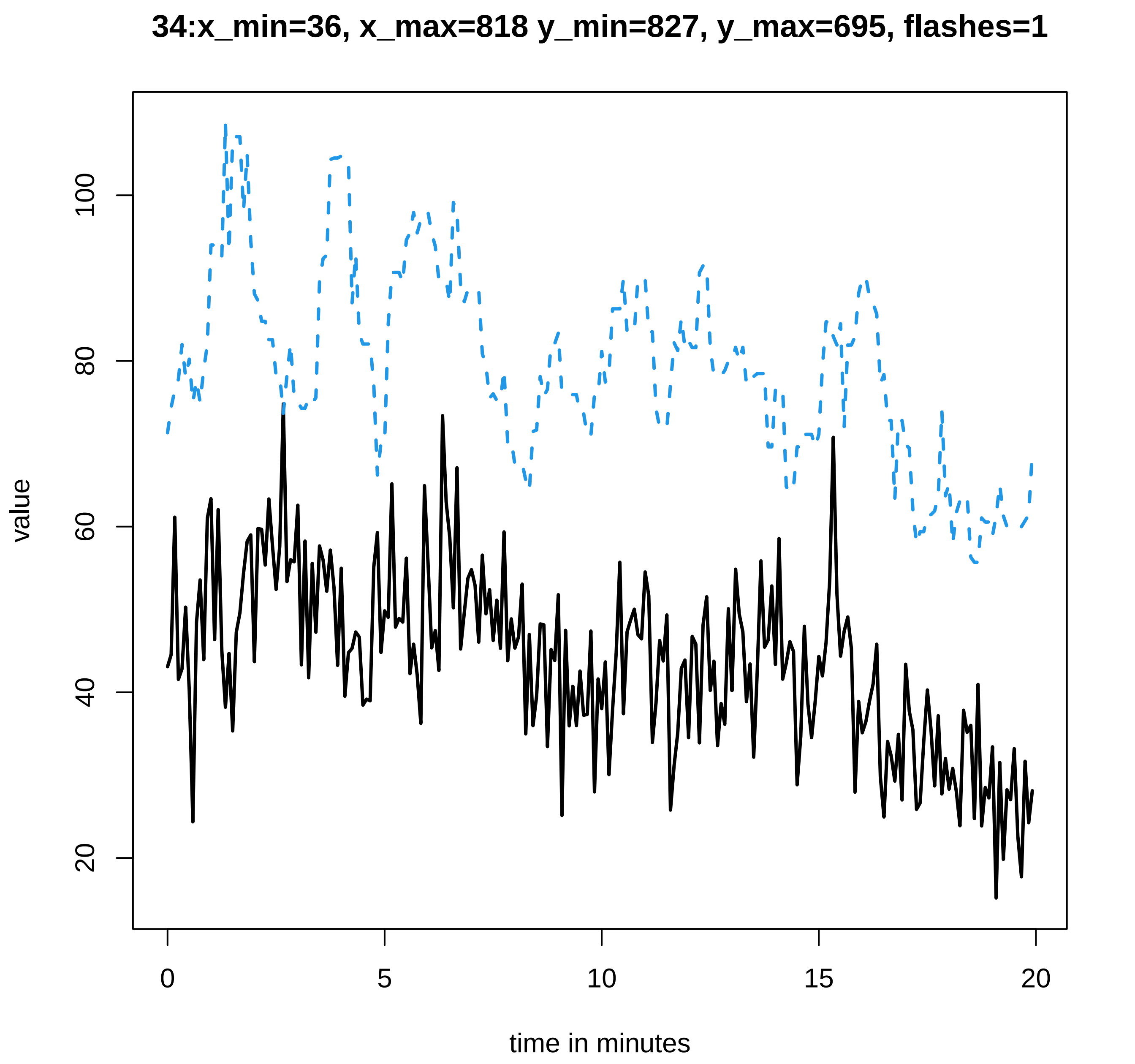

34:x_min=36, x_max=818 y_min=827, y_max=695, flashes=1
value
20
40
60
80
100
0
5
10
15
20
time in minutes

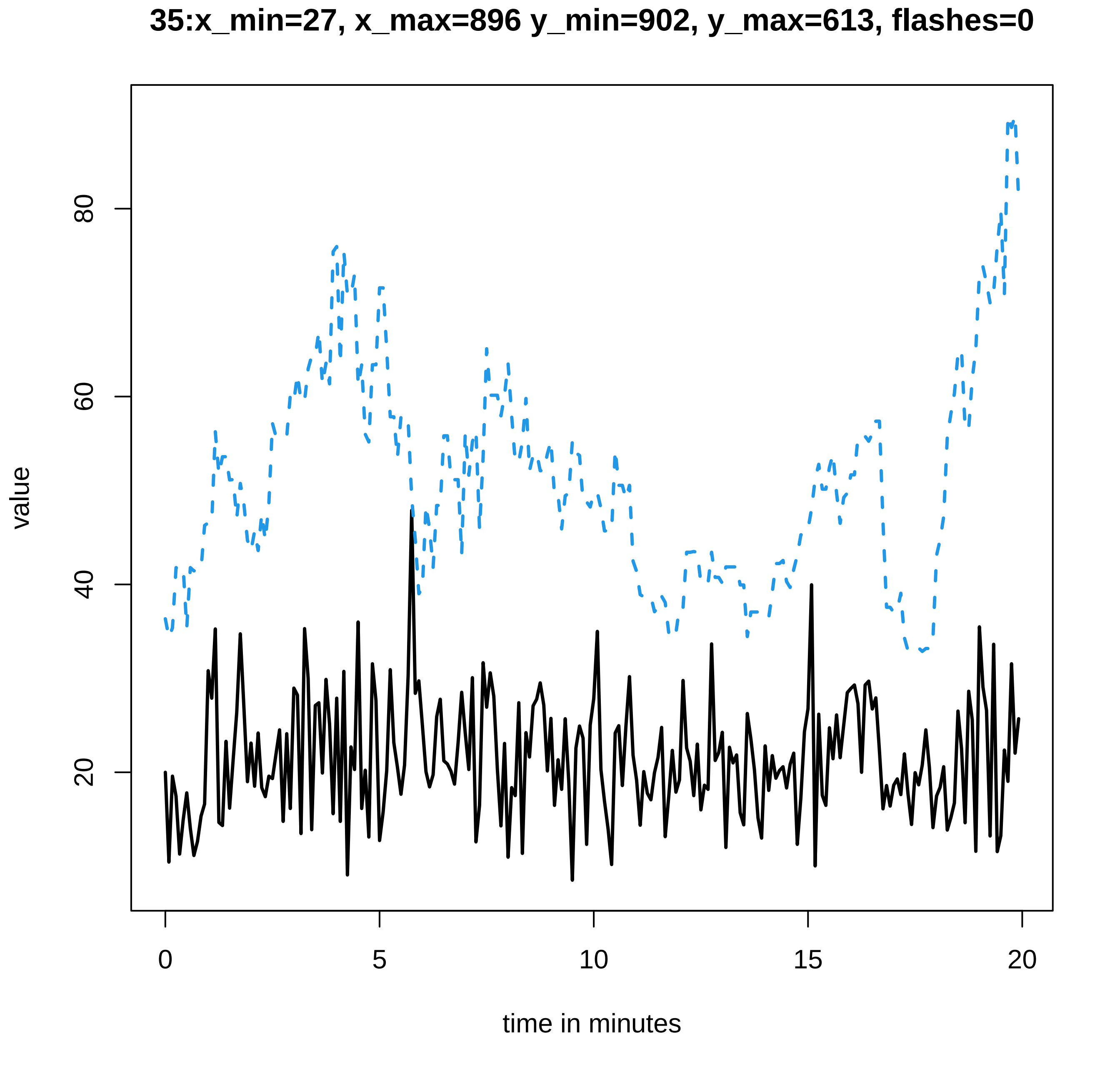
35:x_min=27, x_max=896 y_min=902, y_max=613, flashes=0
value
20
40
60
80
0
5
10
15
20
time in minutes

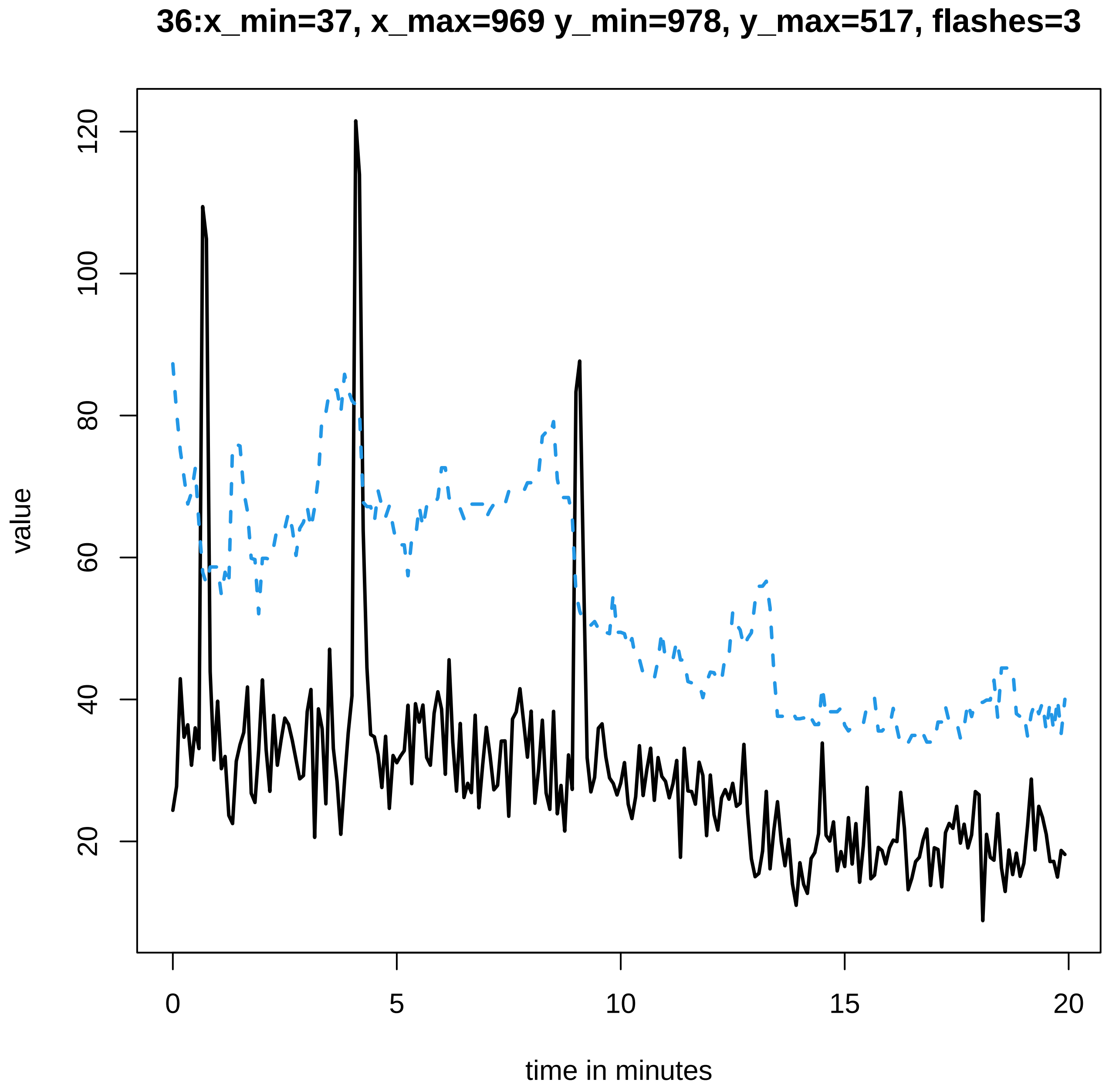

36:x_min=37, x_max=969 y_min=978, y_max=517, flashes=3
value
time in minutes
0
5
10
15
20
20
40
60
80
100
120

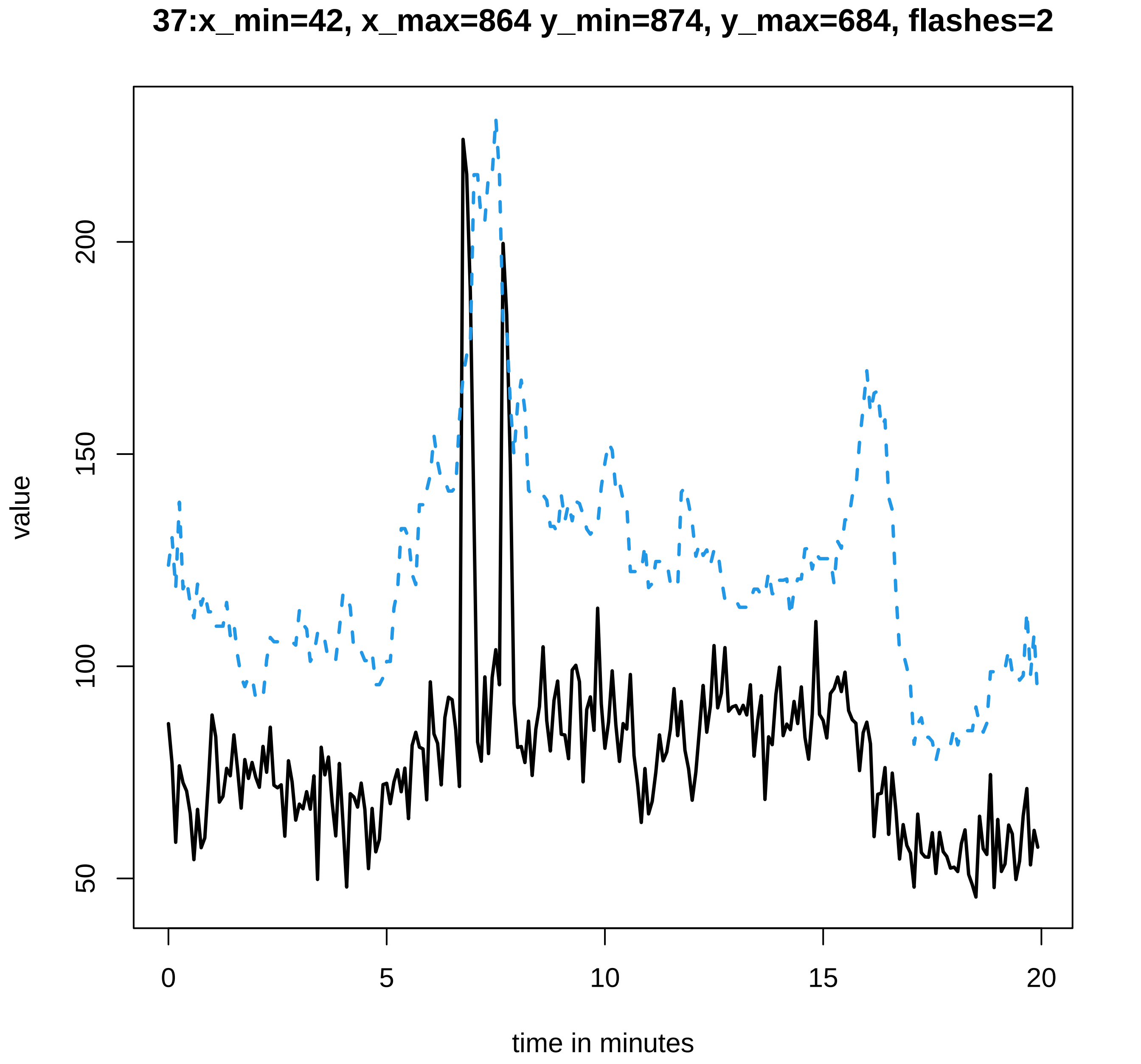
37:x_min=42, x_max=864 y_min=874, y_max=684, flashes=2
value
50
100
150
200
0
5
10
15
20
time in minutes

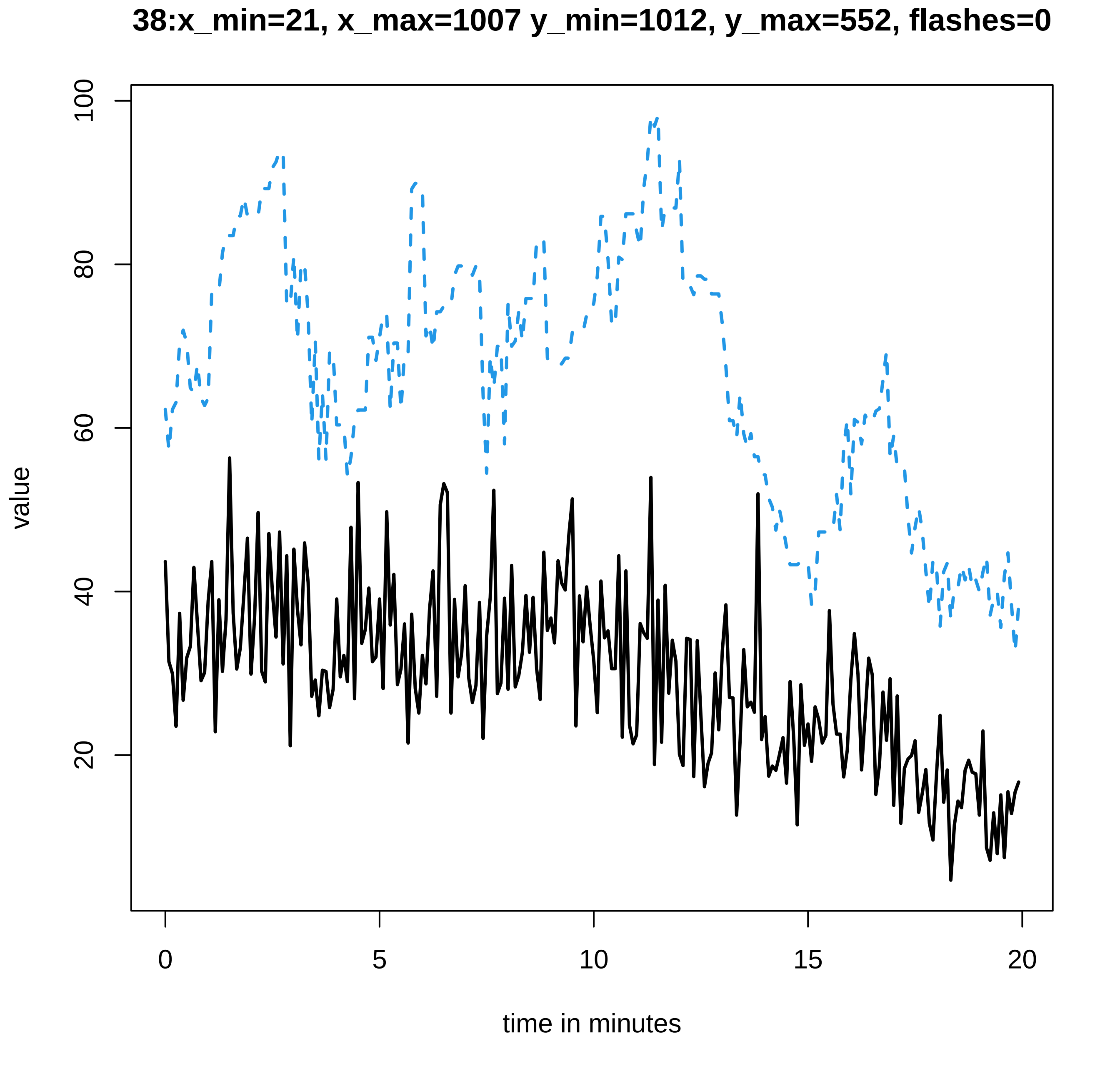
38:x_min=21, x_max=1007 y_min=1012, y_max=552, flashes=0
value
100
80
60
40
20
0
5
10
15
20
time in minutes
