## Supplementary data Calcium spiking for "LysM Receptor Proteins are Required for Ectomycorrhizal Symbiosis in Poplar": profiles-spiking-6.pdf

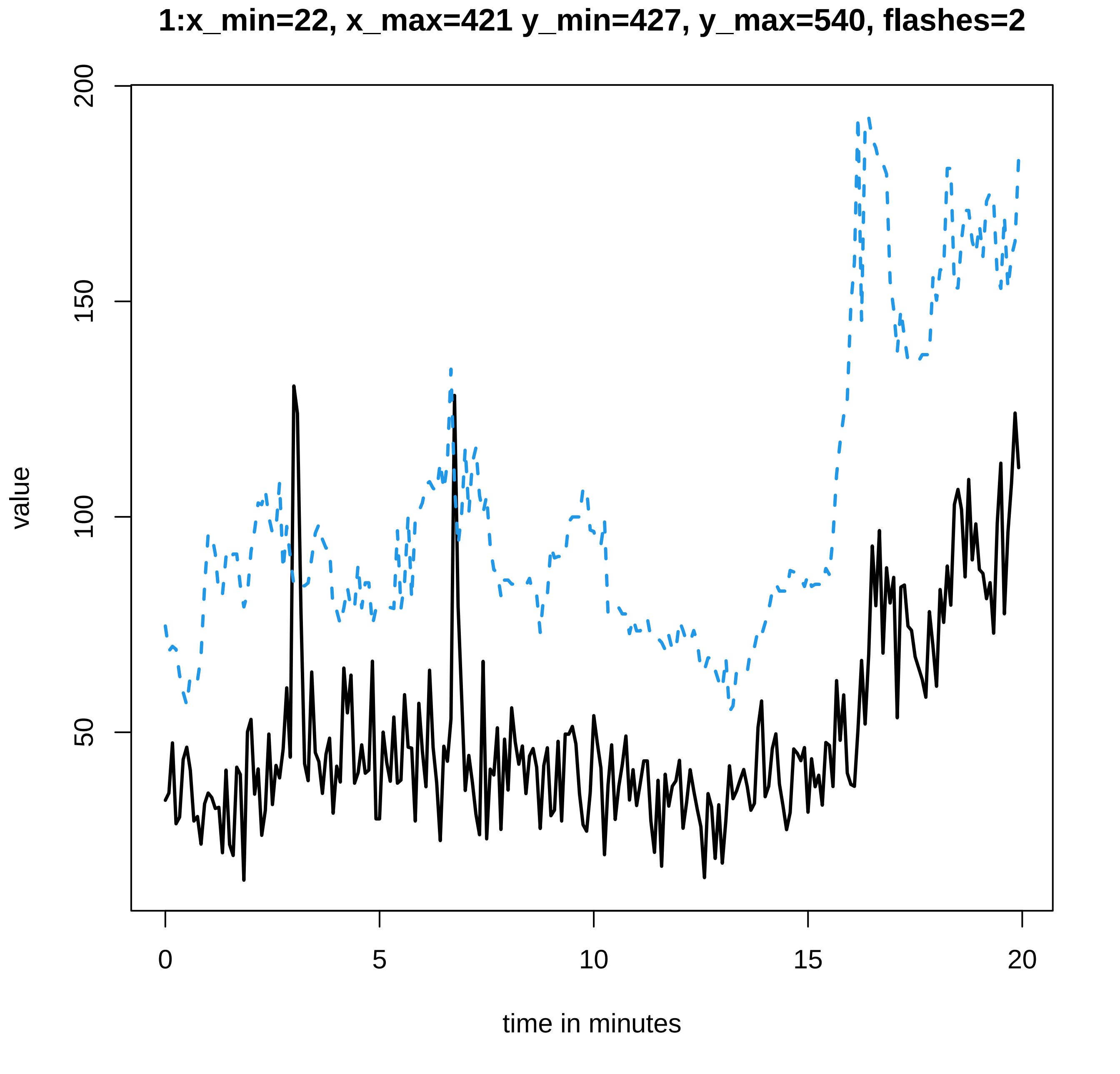

1:x_min=22, x_max=421 y_min=427, y_max=540, flashes=2
value
200
150
100
50
0
5
10
15
20
time in minutes

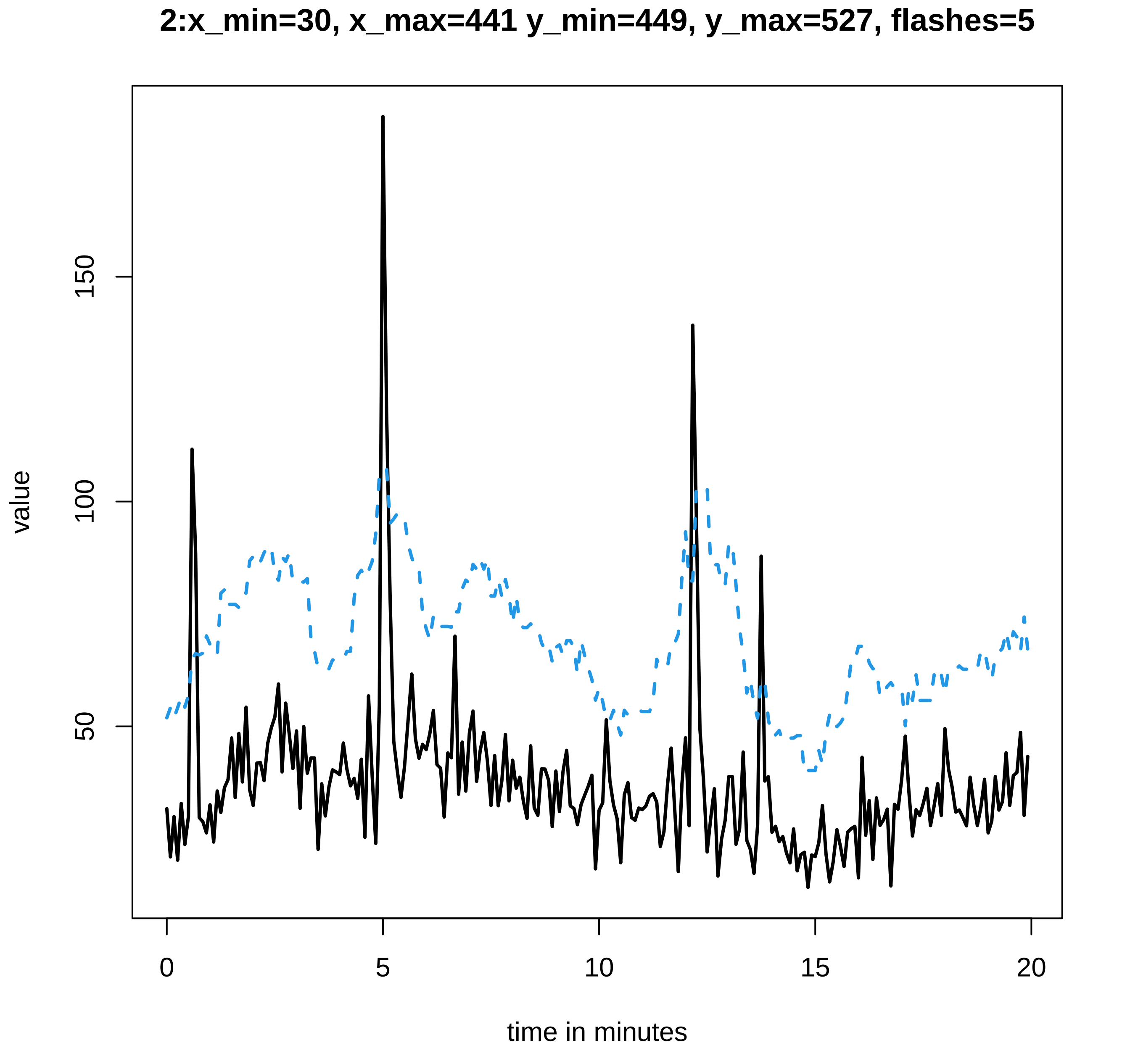
2:x_min=30, x_max=441 y_min=449, y_max=527, flashes=5
value
50
100
150
0
5
10
15
20
time in minutes

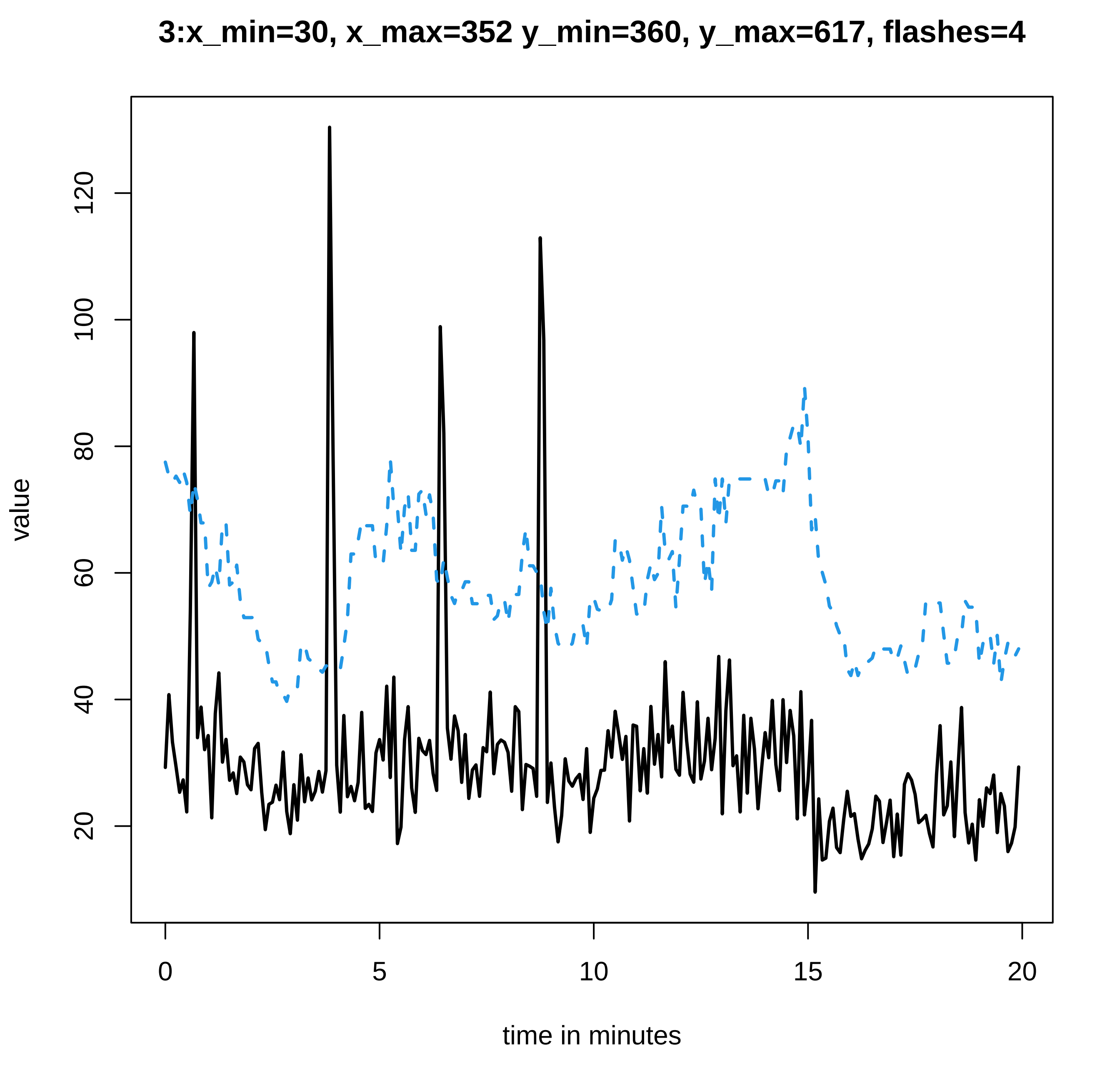
3:x_min=30, x_max=352 y_min=360, y_max=617, flashes=4
value
20
40
60
80
100
120
0
5
10
15
20
time in minutes

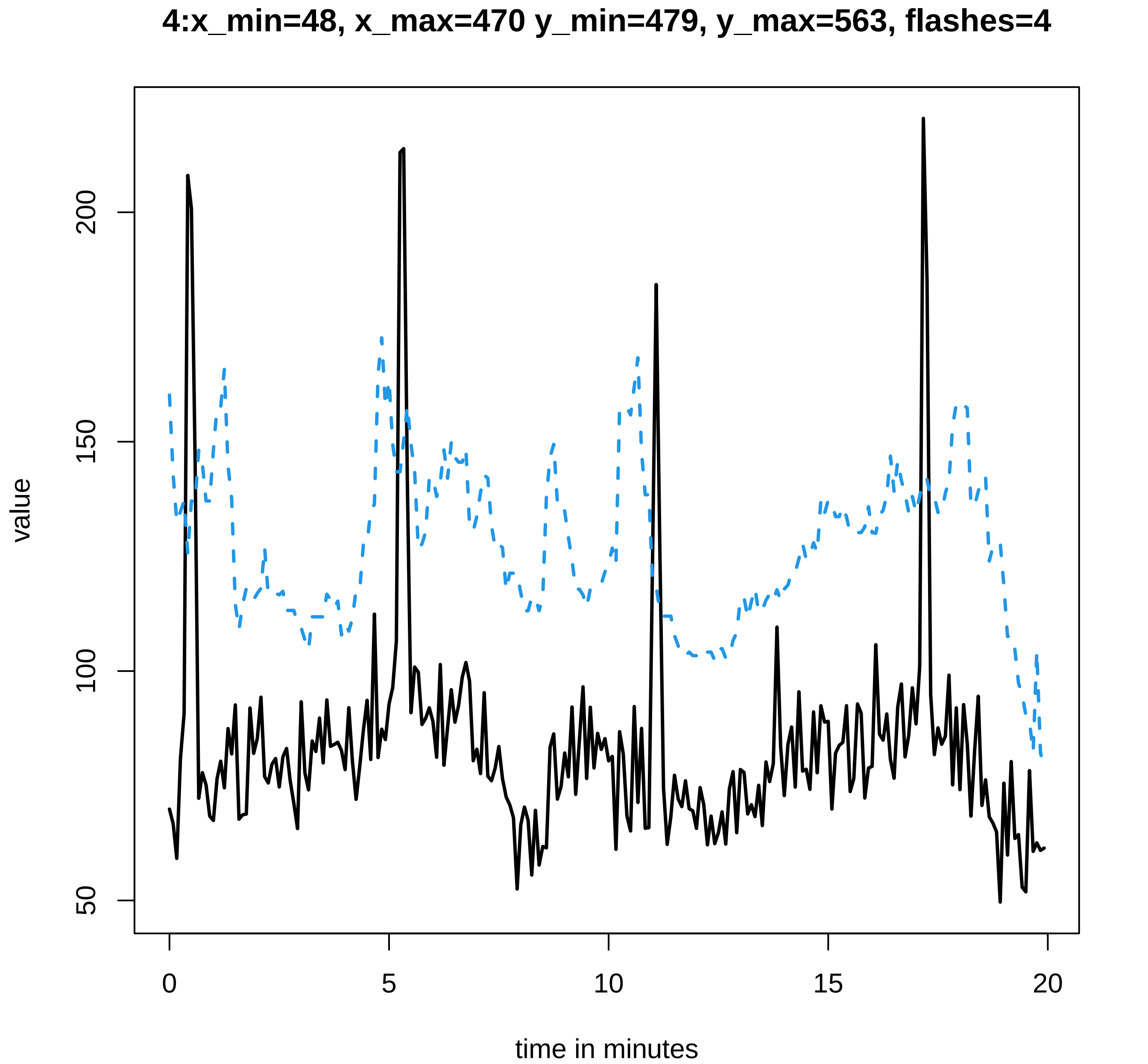
4:x_min=48, x_max=470 y_min=479, y_max=563, flashes=4
value
50
100
150
200
0
5
10
15
20
time in minutes

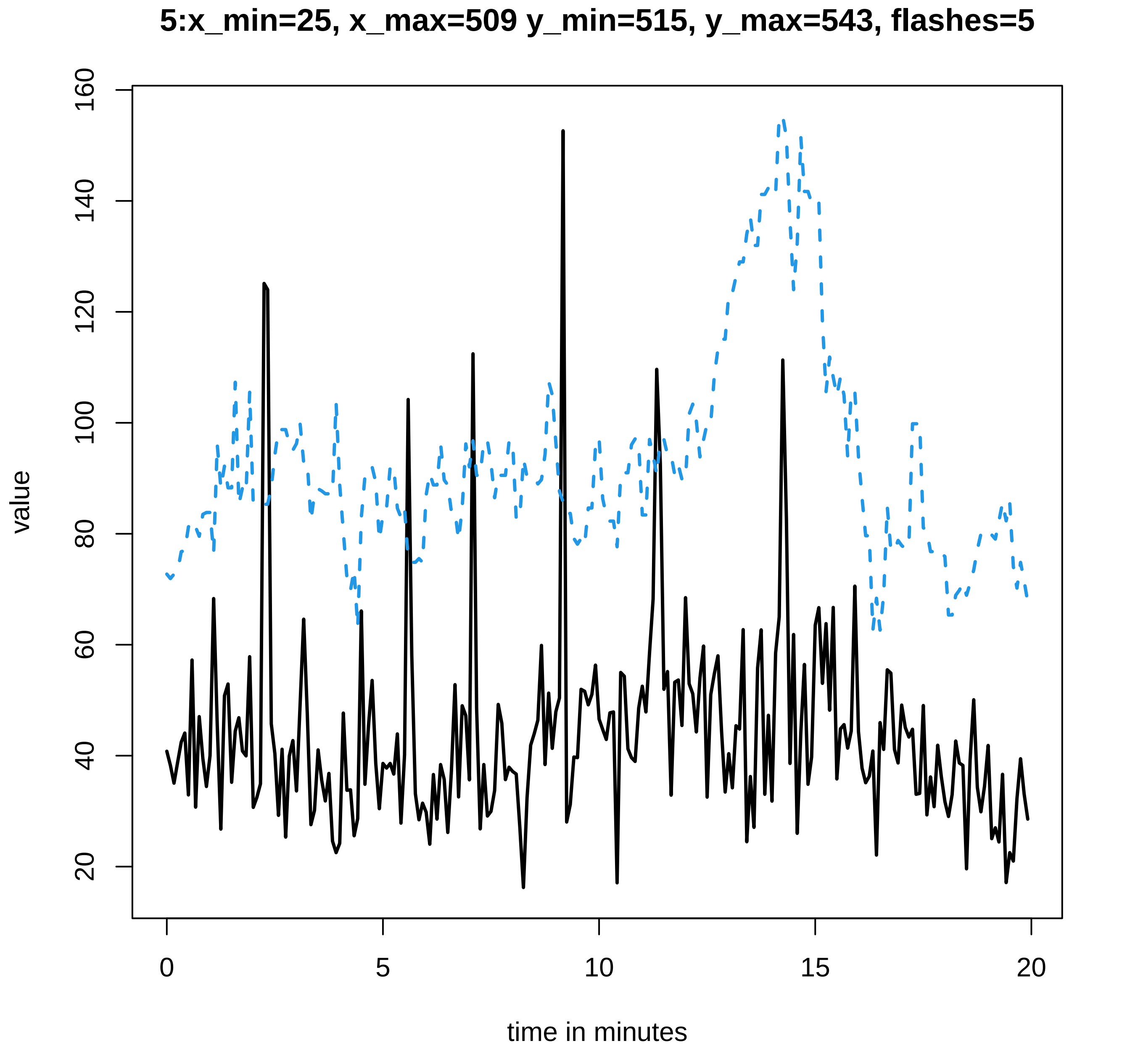
5:x_min=25, x_max=509 y_min=515, y_max=543, flashes=5
value
20
40
60
80
100
120
140
160
0
5
10
15
20
time in minutes

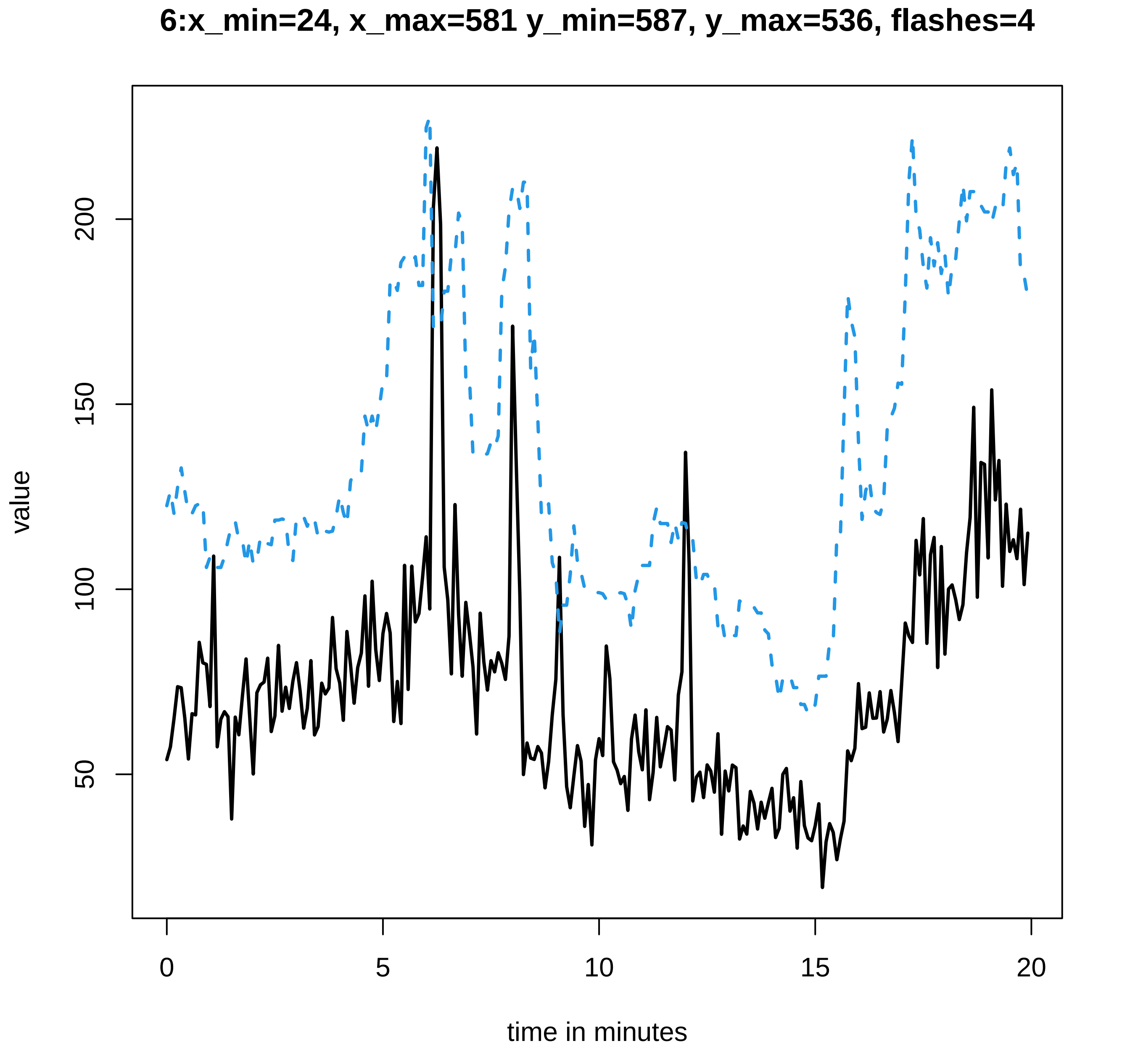

6:x_min=24, x_max=581 y_min=587, y_max=536, flashes=4
value
200
150
100
50
0
5
10
15
20
time in minutes

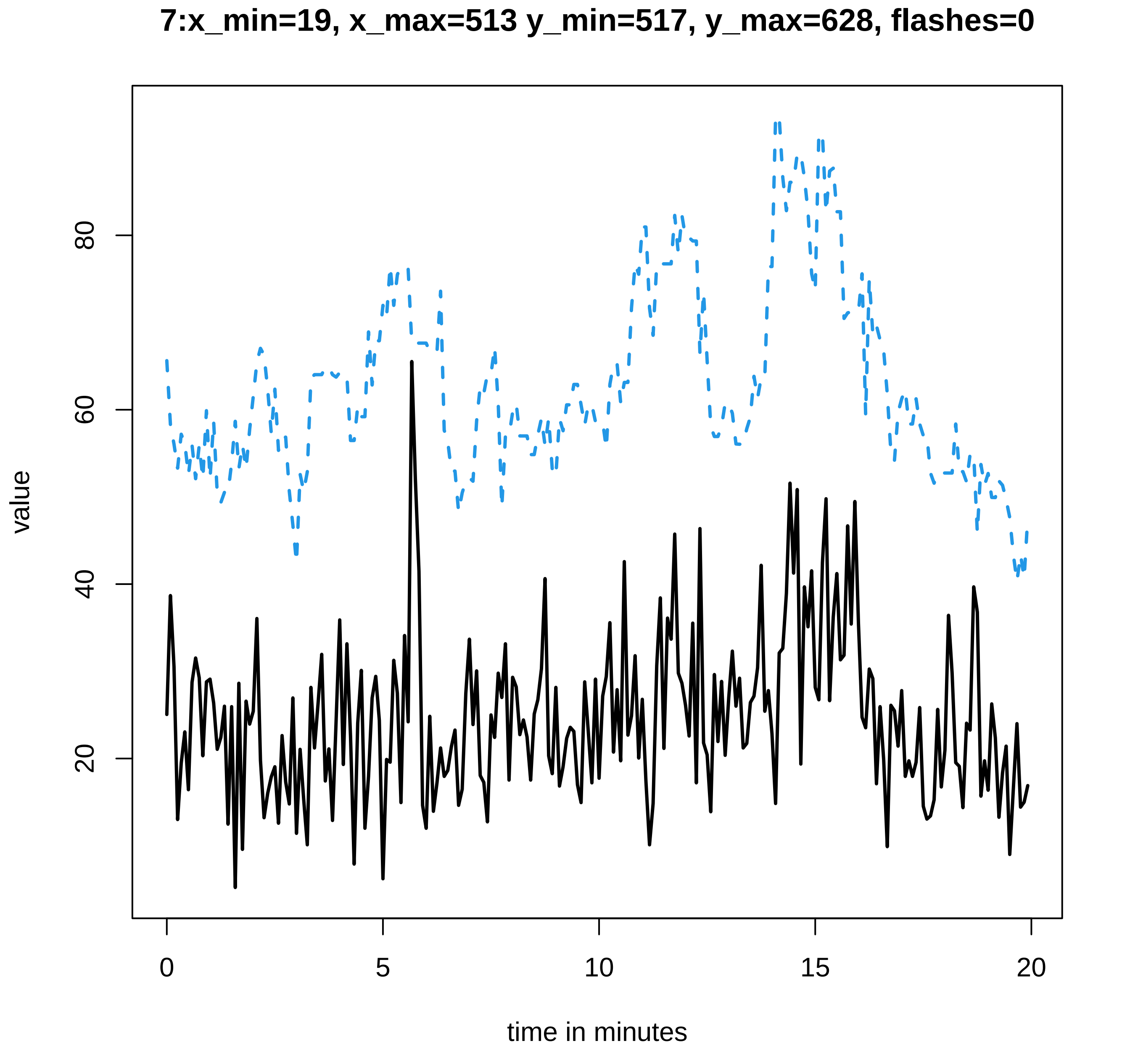
7:x_min=19, x_max=513 y_min=517, y_max=628, flashes=0
value
80
60
40
20
0
5
10
15
20
time in minutes

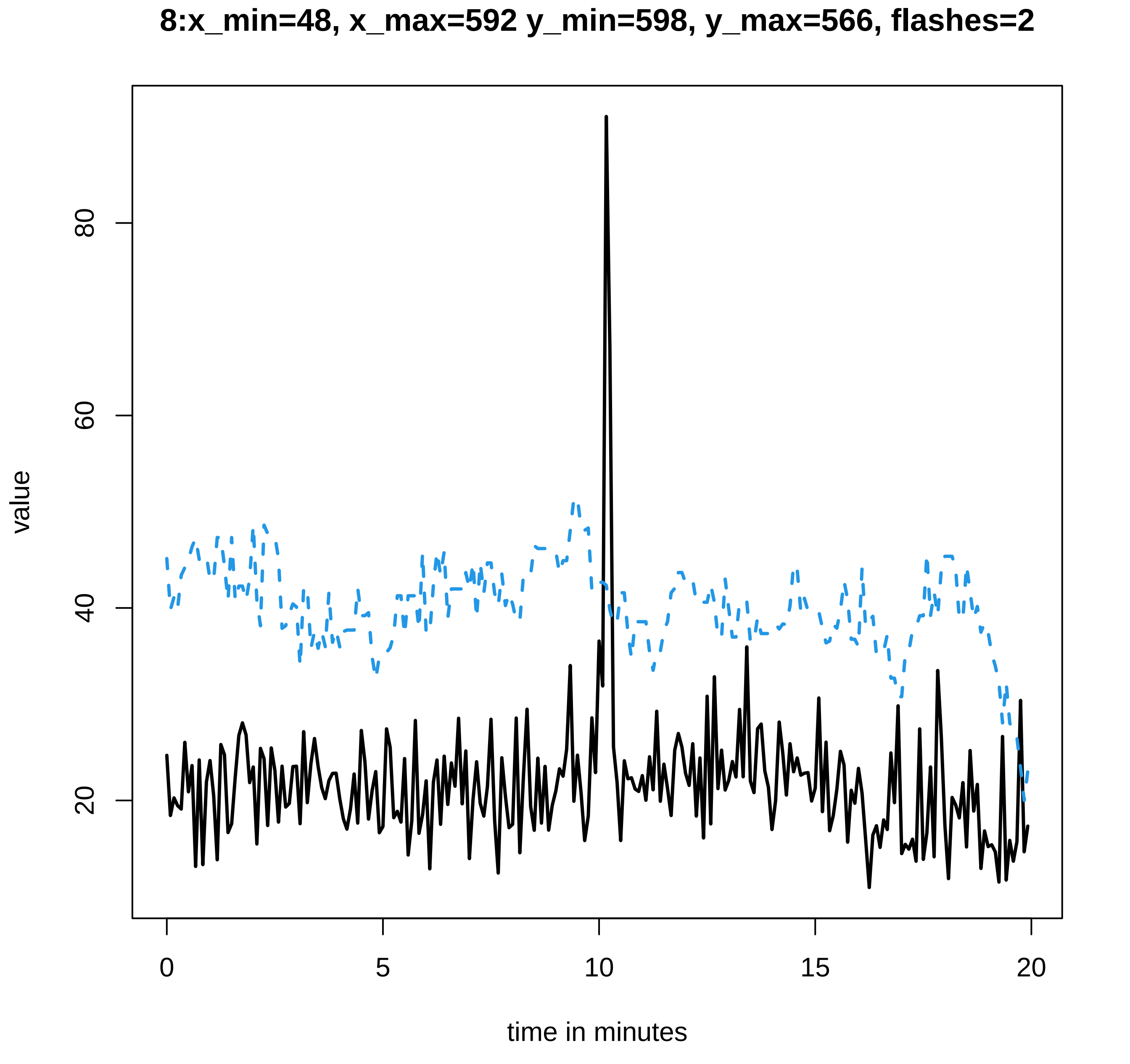
8:x_min=48, x_max=592 y_min=598, y_max=566, flashes=2
value
20
40
60
80
0
5
10
15
20
time in minutes

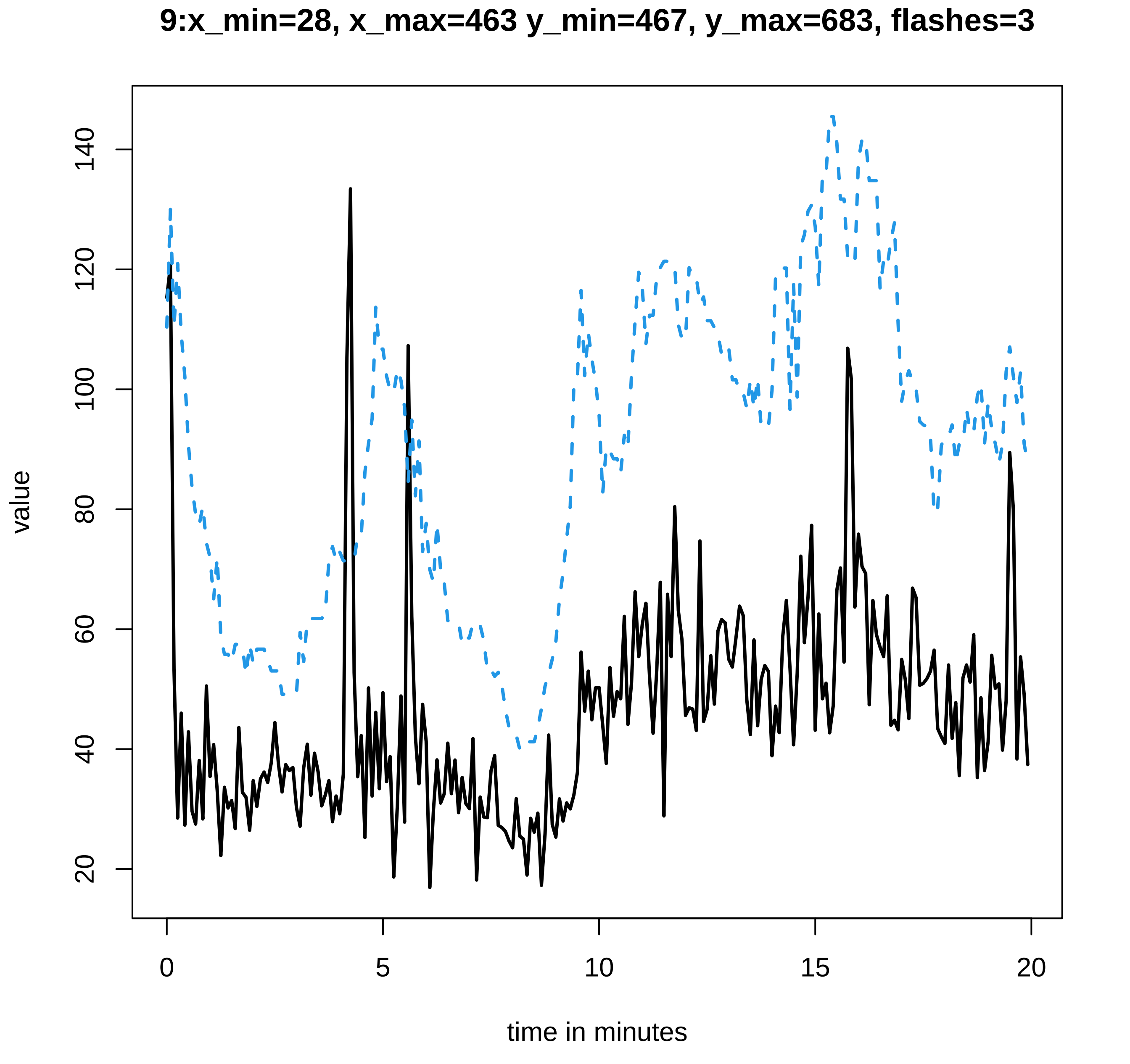

9:x_min=28, x_max=463 y_min=467, y_max=683, flashes=3
value
20
40
60
80
100
120
140
0
5
10
15
20
time in minutes

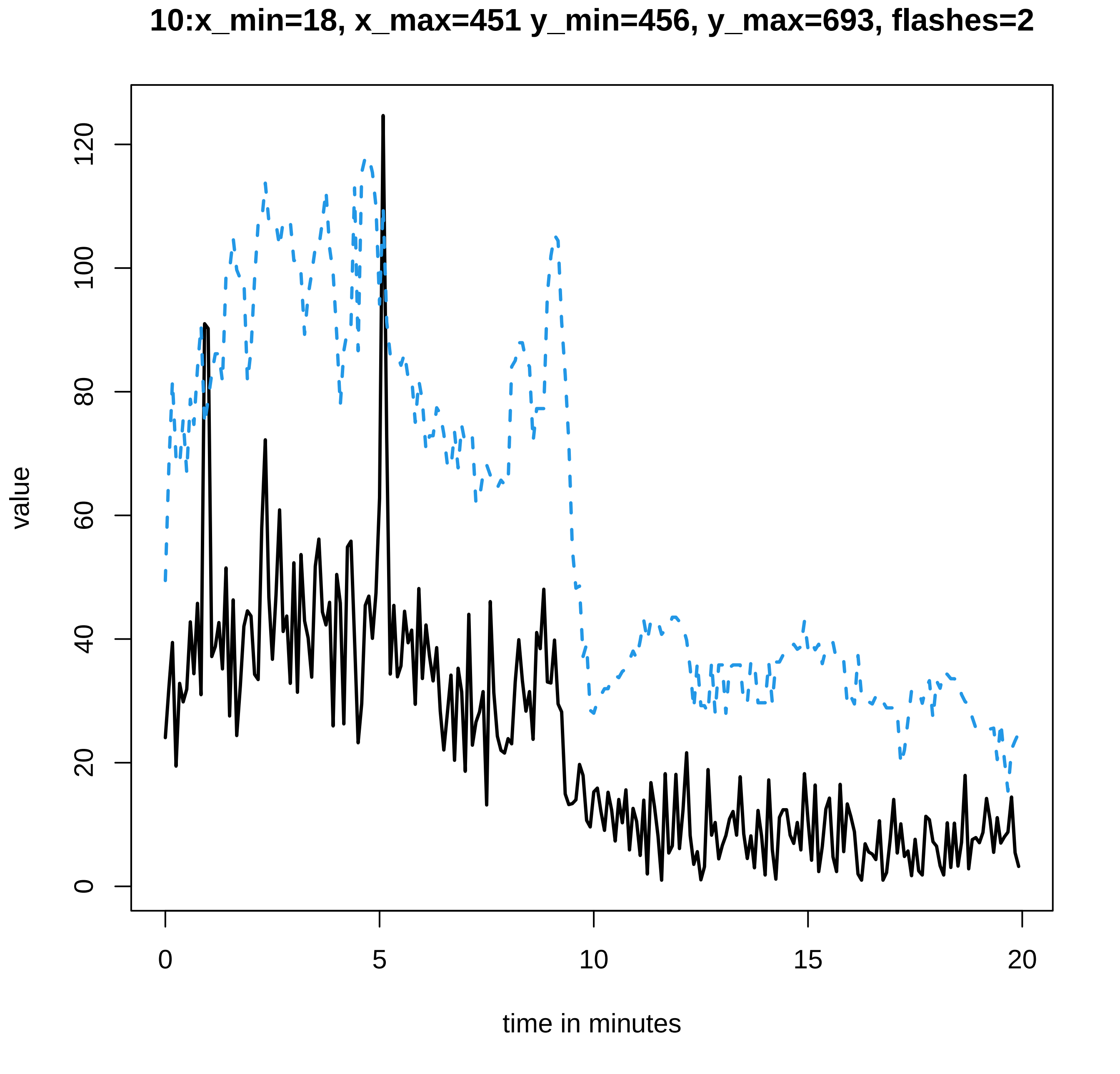

10:x_min=18, x_max=451 y_min=456, y_max=693, flashes=2
value
0
20
40
60
80
100
120
time in minutes
0
5
10
15
20

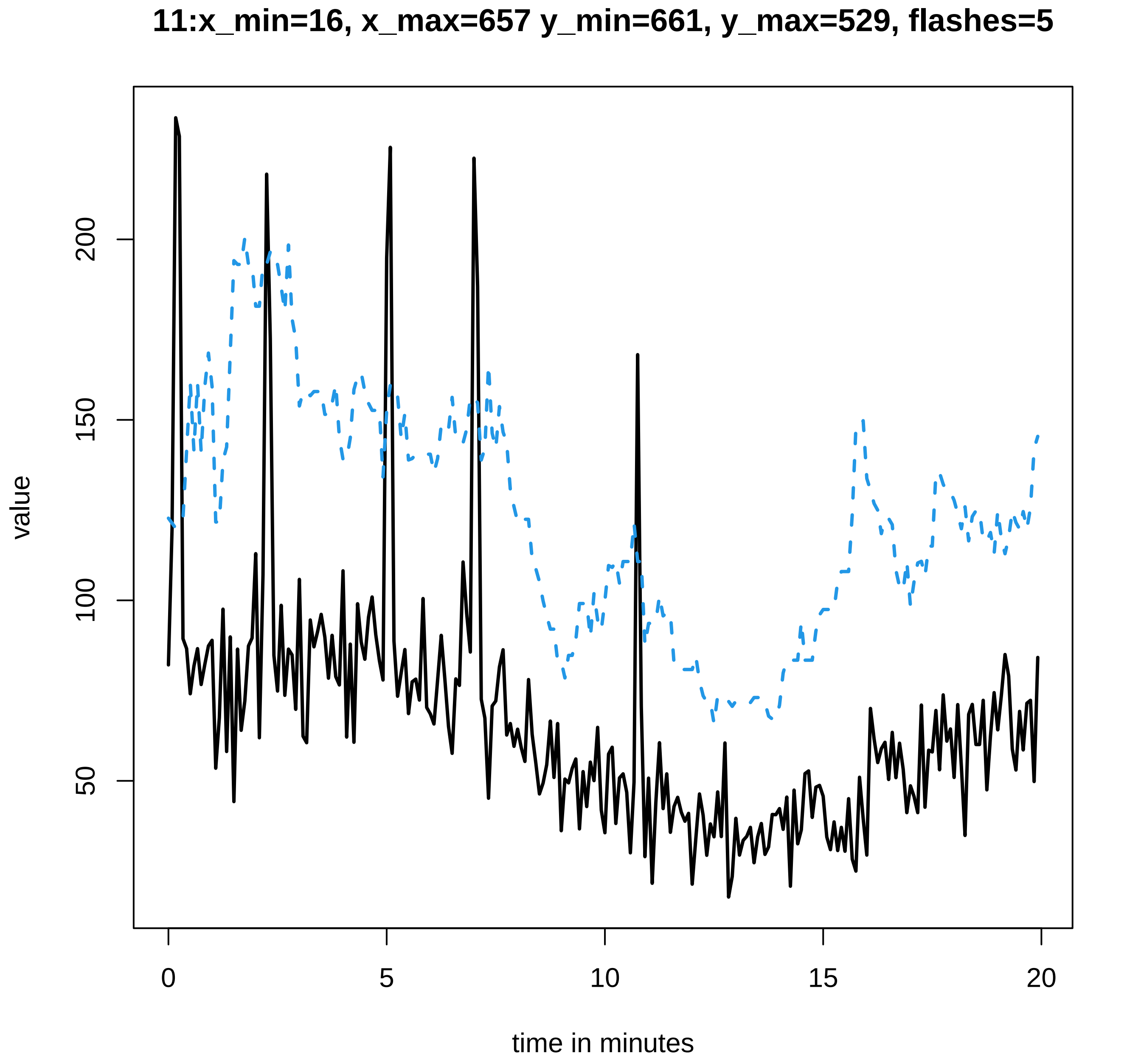
11:x_min=16, x_max=657 y_min=661, y_max=529, flashes=5
value
200
150
100
50
0
5
10
15
20
time in minutes

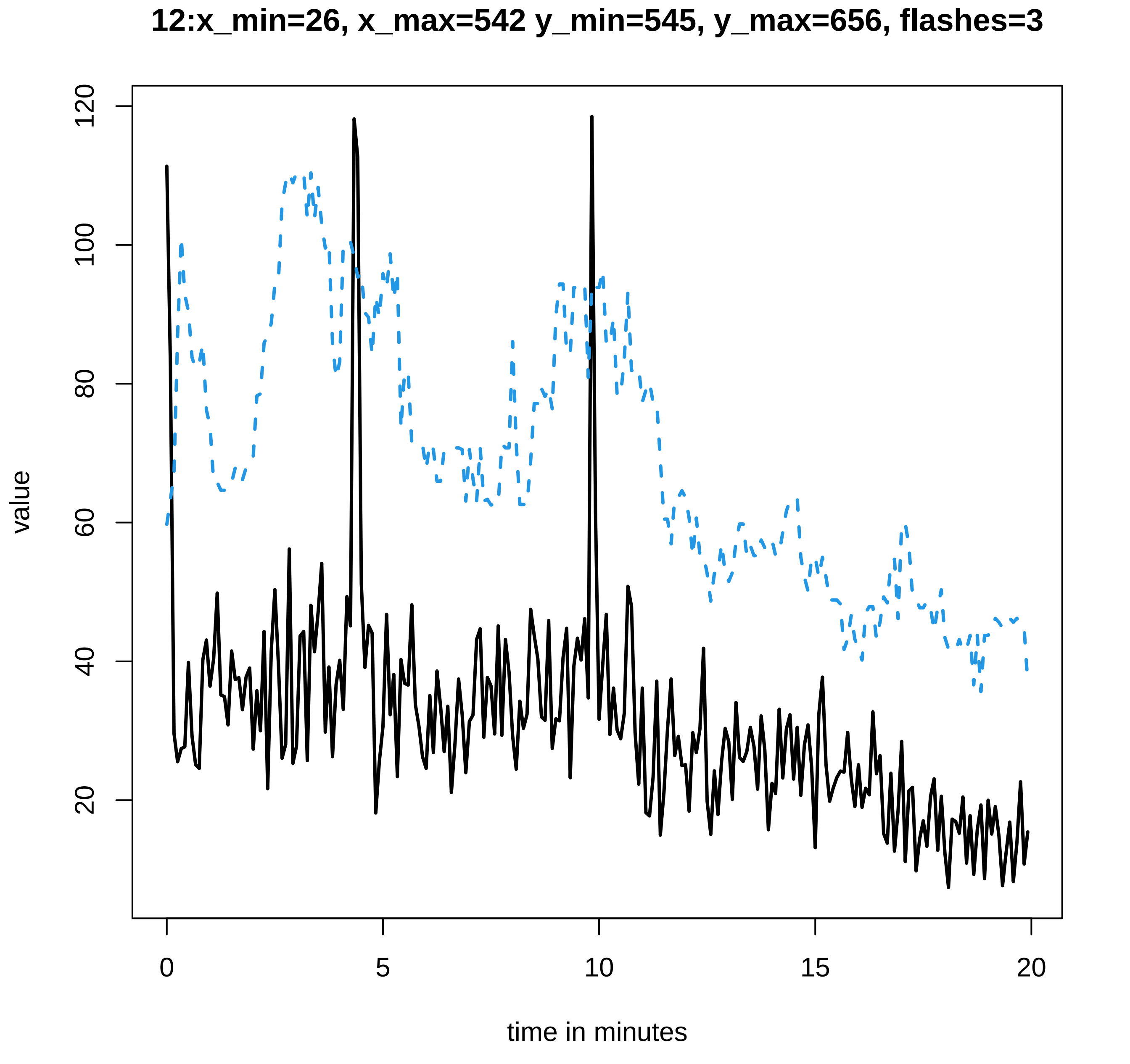
12:x_min=26, x_max=542 y_min=545, y_max=656, flashes=3
value
120
100
80
60
40
20
0
5
10
15
20
time in minutes

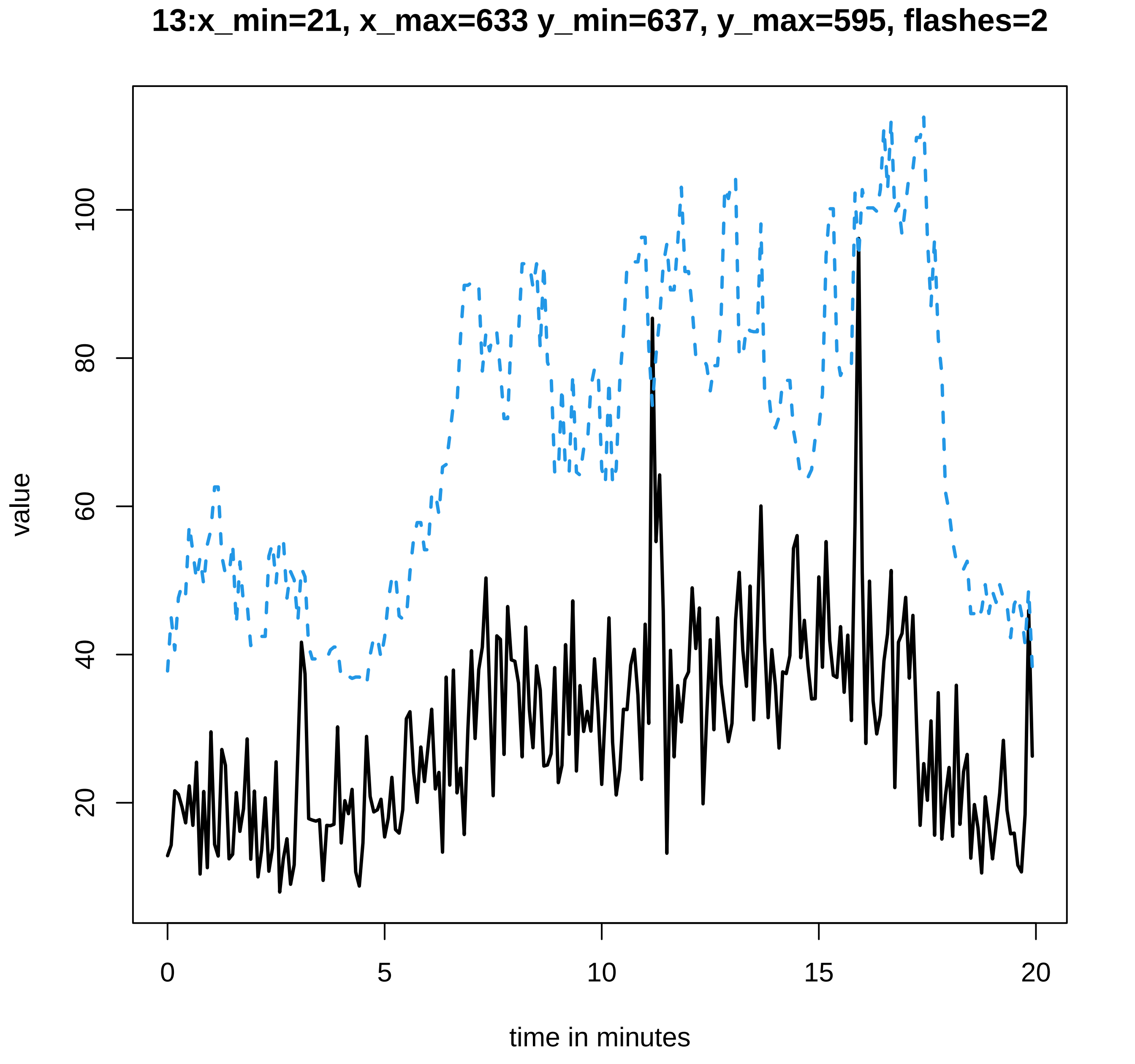
13:x_min=21, x_max=633 y_min=637, y_max=595, flashes=2
value
20
40
60
80
100
0
5
10
15
20
time in minutes

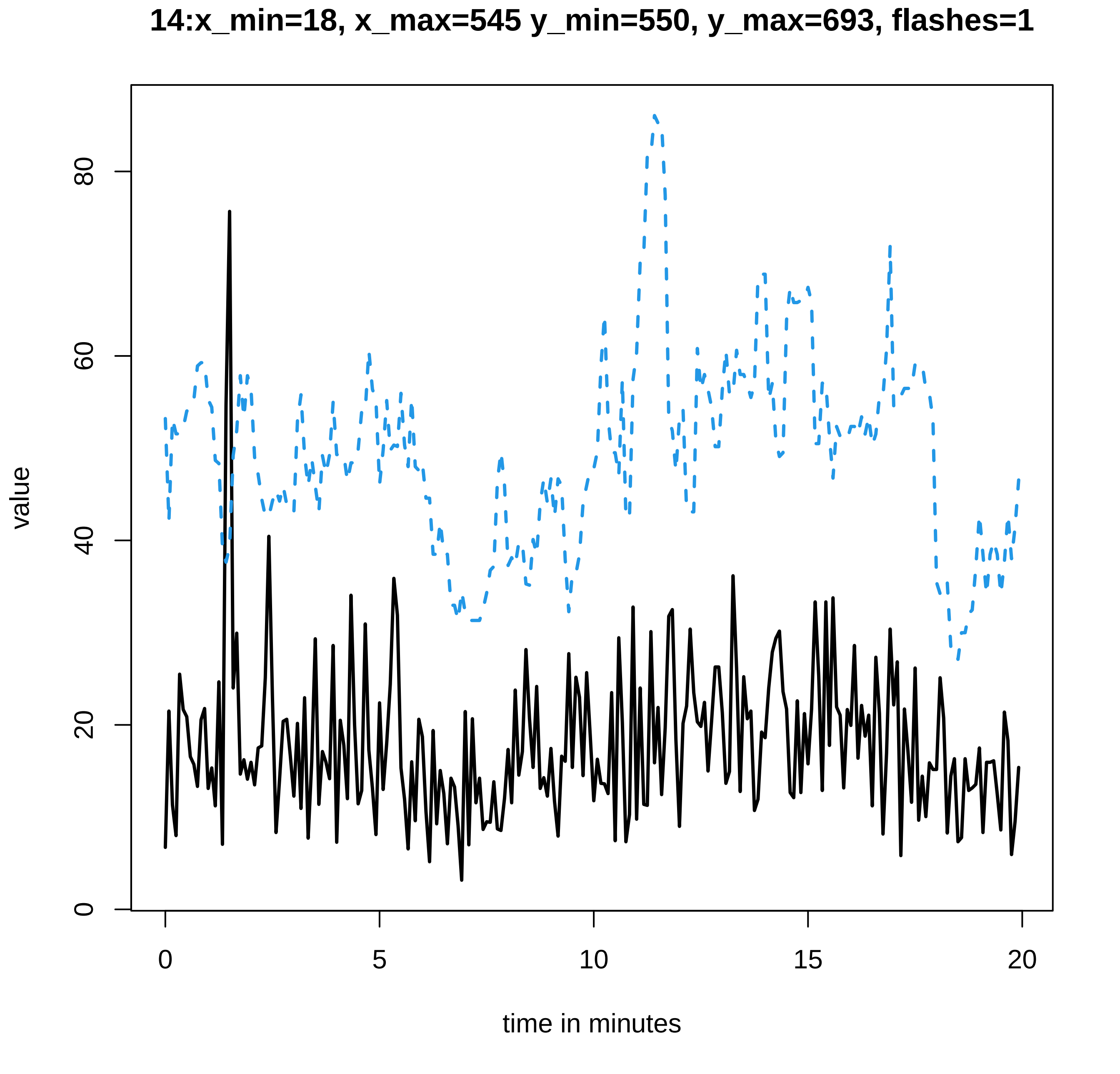

14:x_min=18, x_max=545 y_min=550, y_max=693, flashes=1
value
0
20
40
60
80
0
5
10
15
20
time in minutes

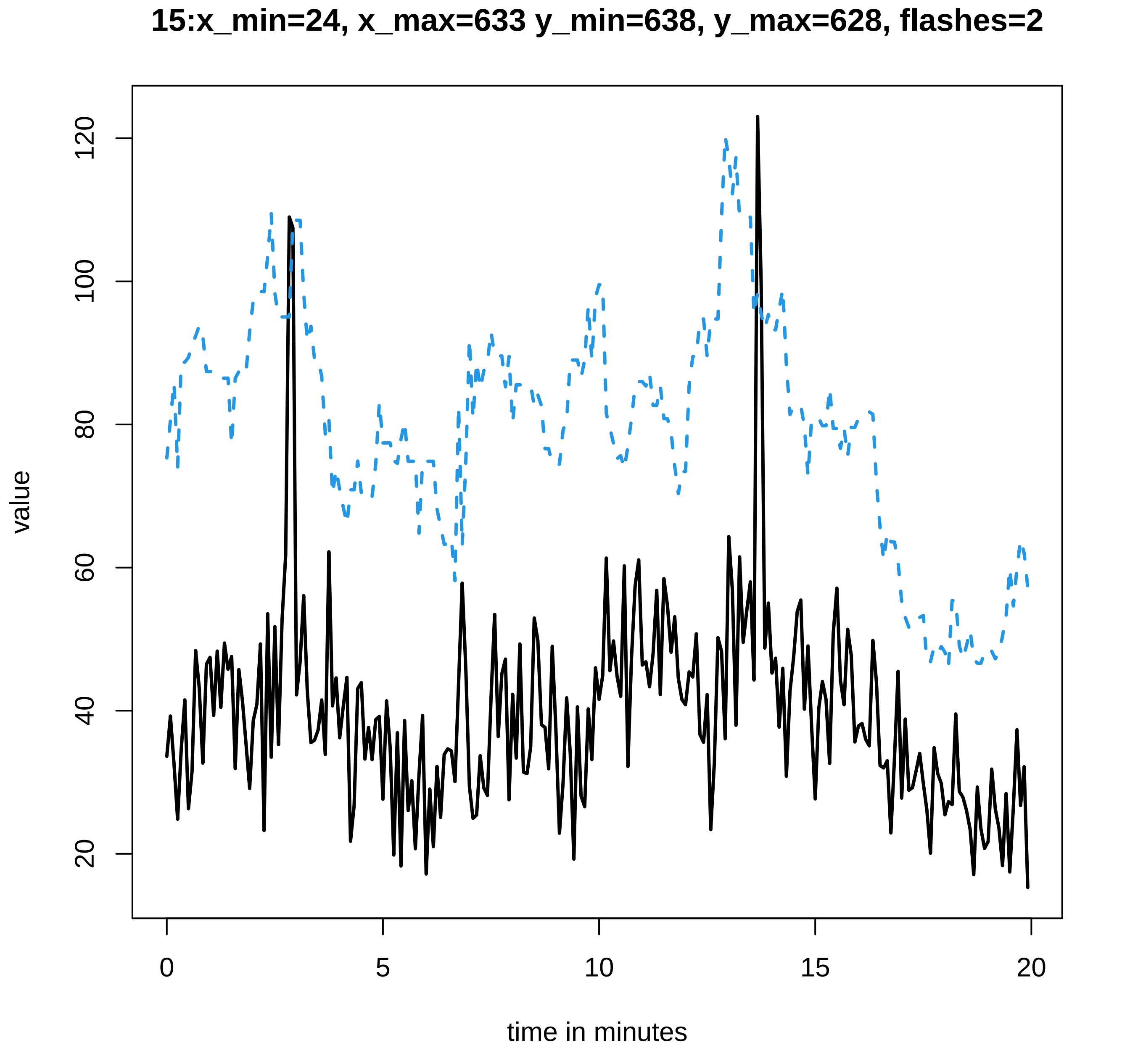
15:x_min=24, x_max=633 y_min=638, y_max=628, flashes=2
value
20
40
60
80
100
120
0
5
10
15
20
time in minutes

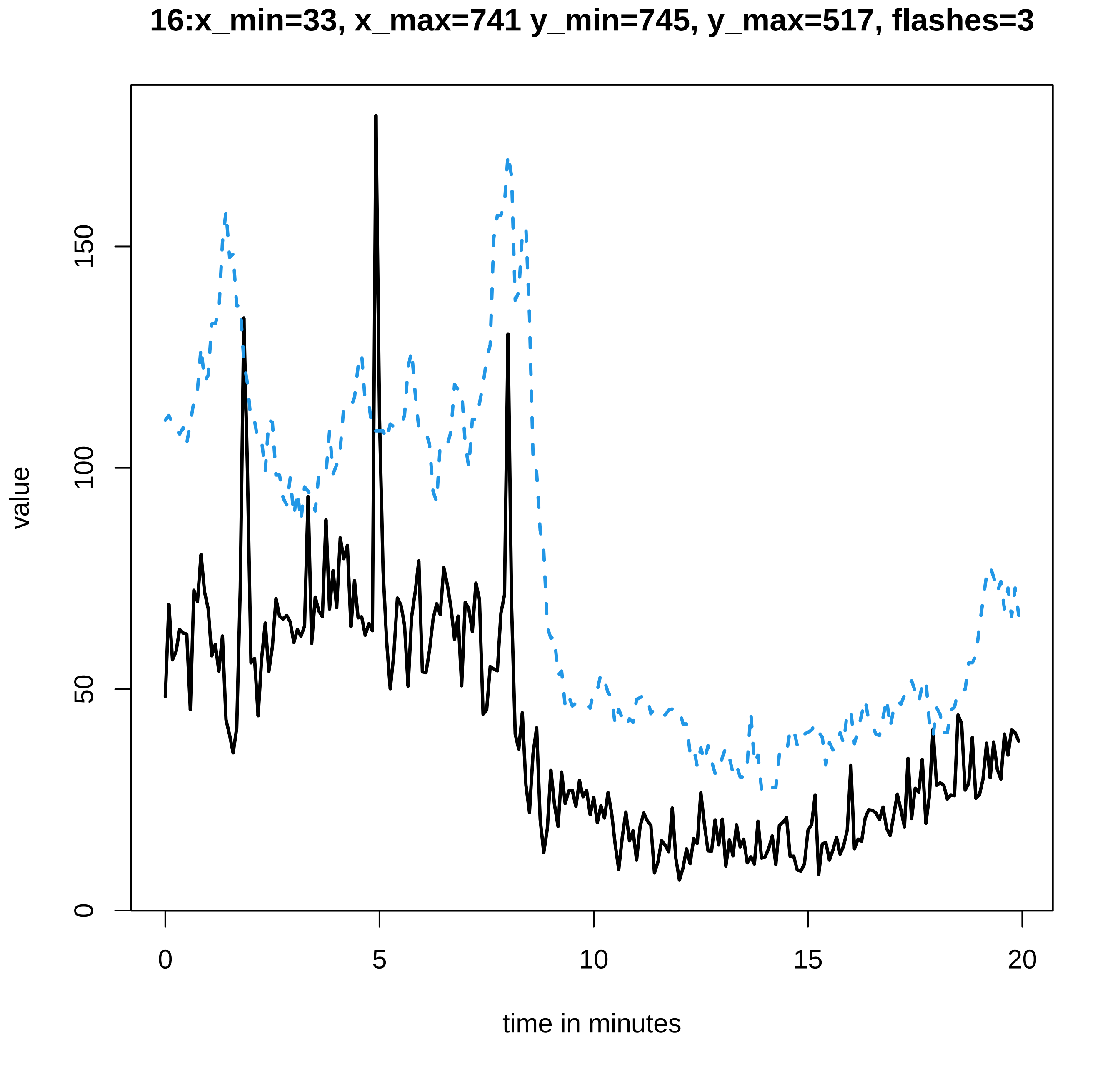
16:x_min=33, x_max=741 y_min=745, y_max=517, flashes=3
value
0
50
100
150
0
5
10
15
20
time in minutes

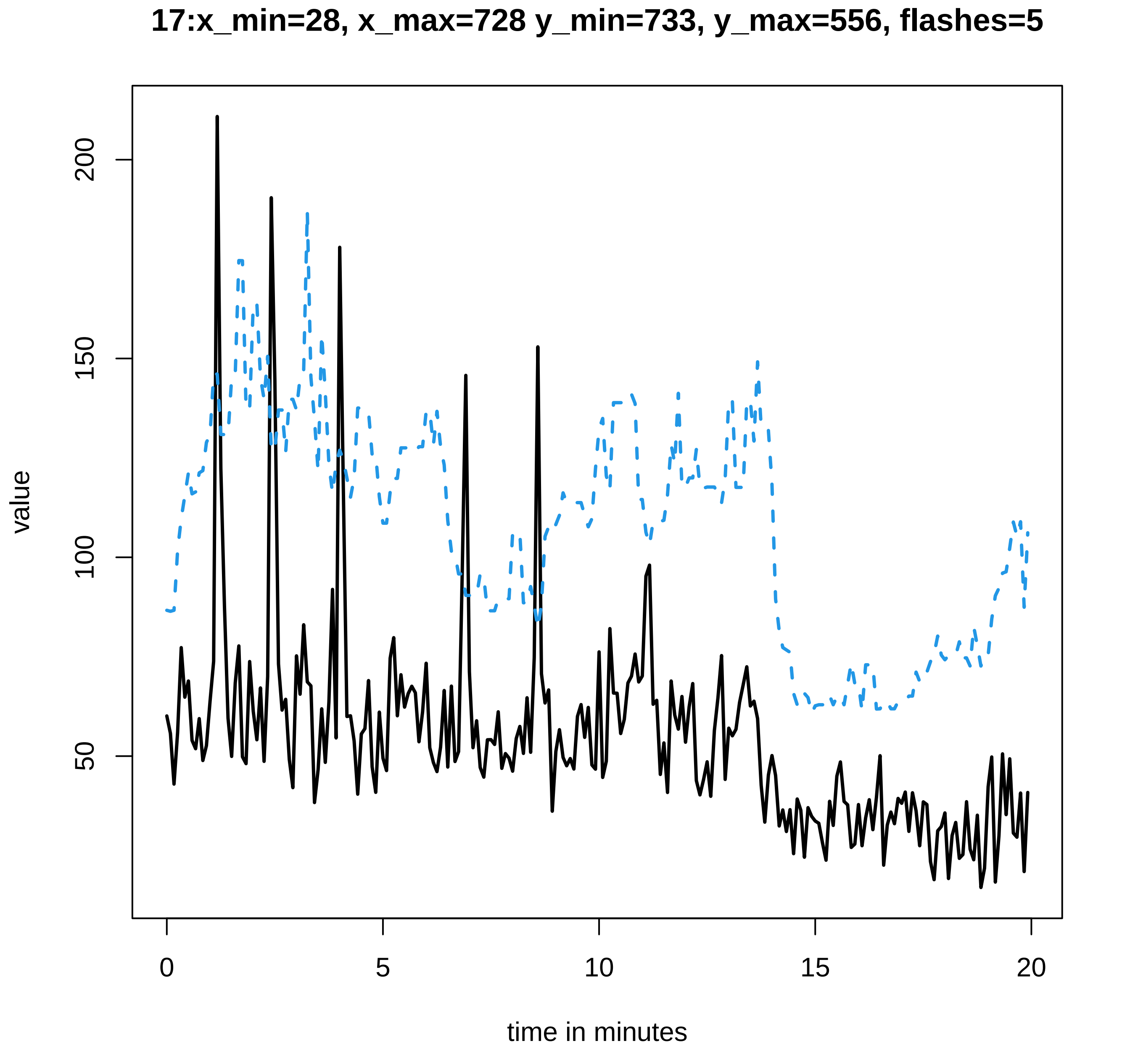

17:x_min=28, x_max=728 y_min=733, y_max=556, flashes=5
value
200
150
100
50
0
5
10
15
20
time in minutes

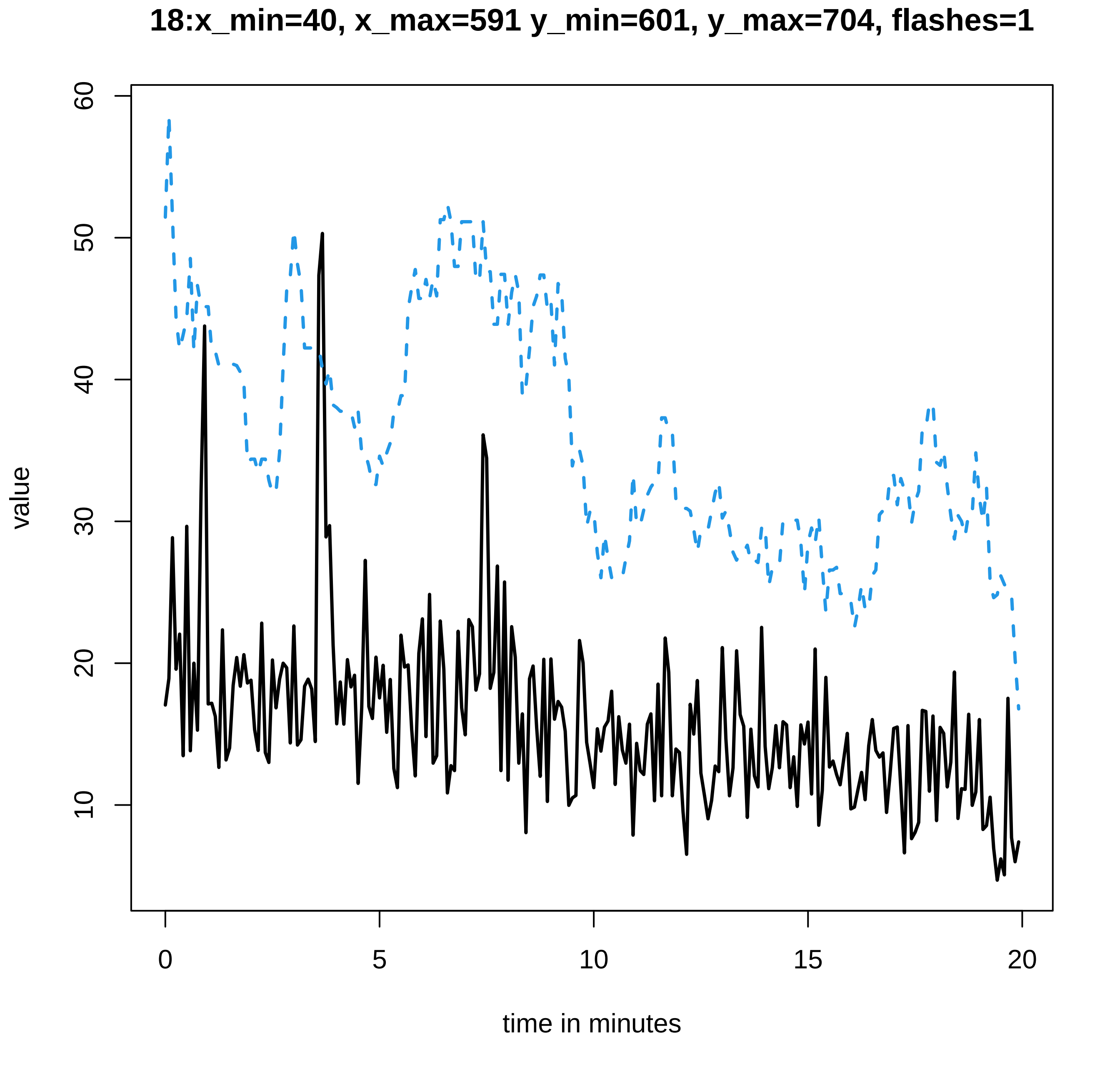

18:x_min=40, x_max=591 y_min=601, y_max=704, flashes=1
value
60
50
40
30
20
10
0
5
10
15
20
time in minutes

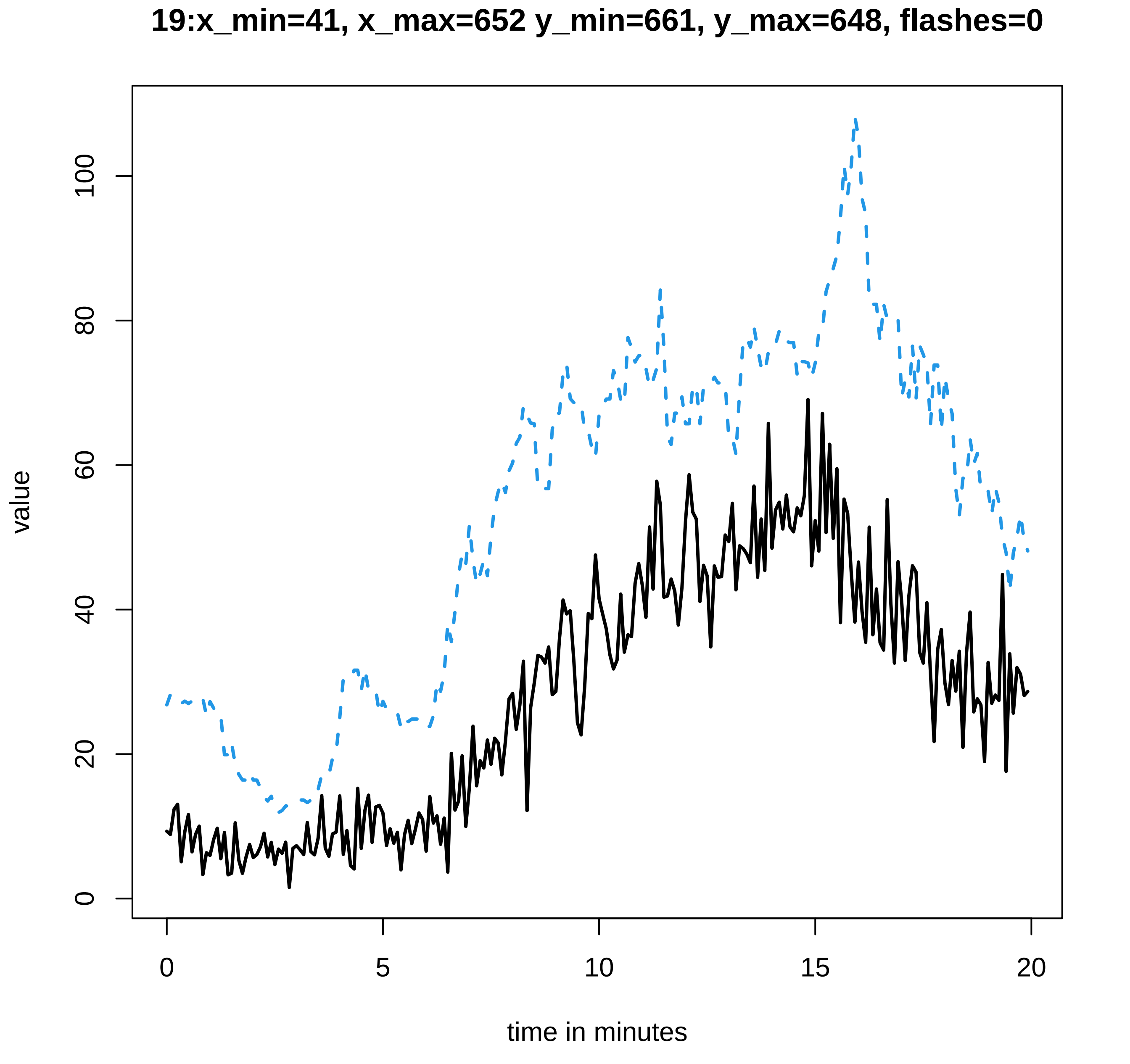
19:x_min=41, x_max=652 y_min=661, y_max=648, flashes=0
value
0
20
40
60
80
100
0
5
10
15
20
time in minutes

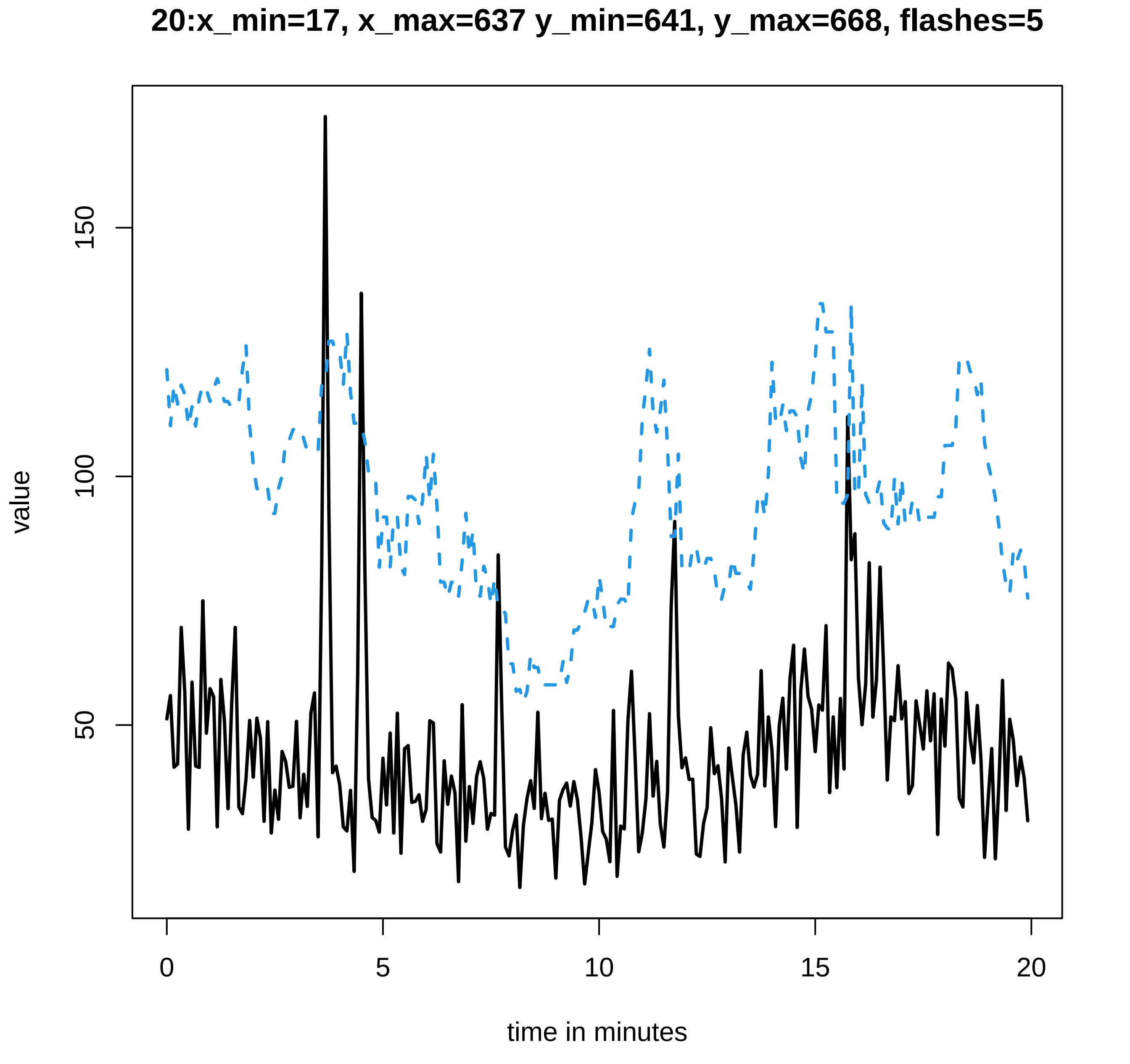
20:x_min=17, x_max=637 y_min=641, y_max=668, flashes=5
value
150
100
50
0
5
10
15
20
time in minutes

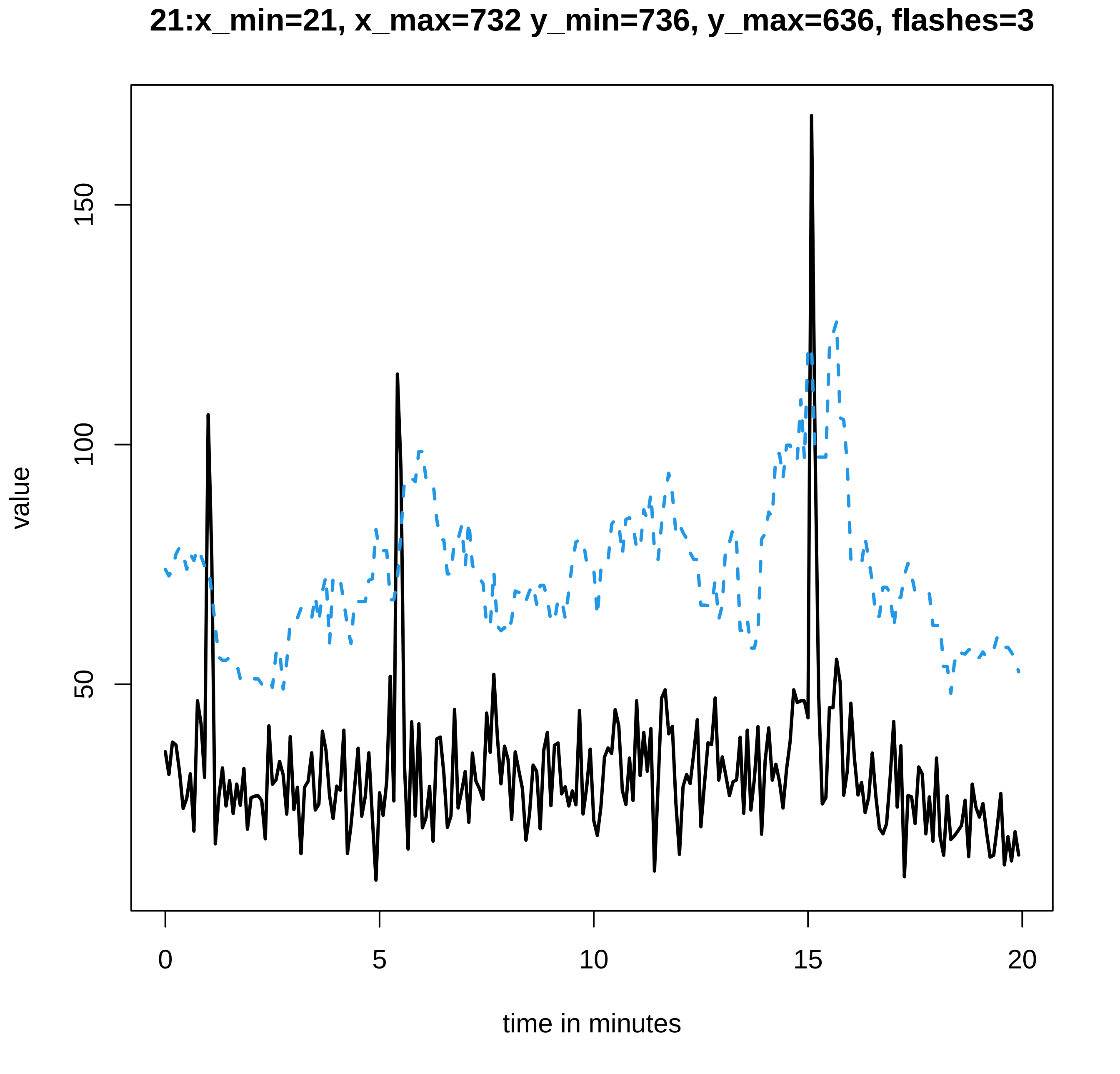

21:x_min=21, x_max=732 y_min=736, y_max=636, flashes=3
value
150
100
50
0
5
10
15
20
time in minutes

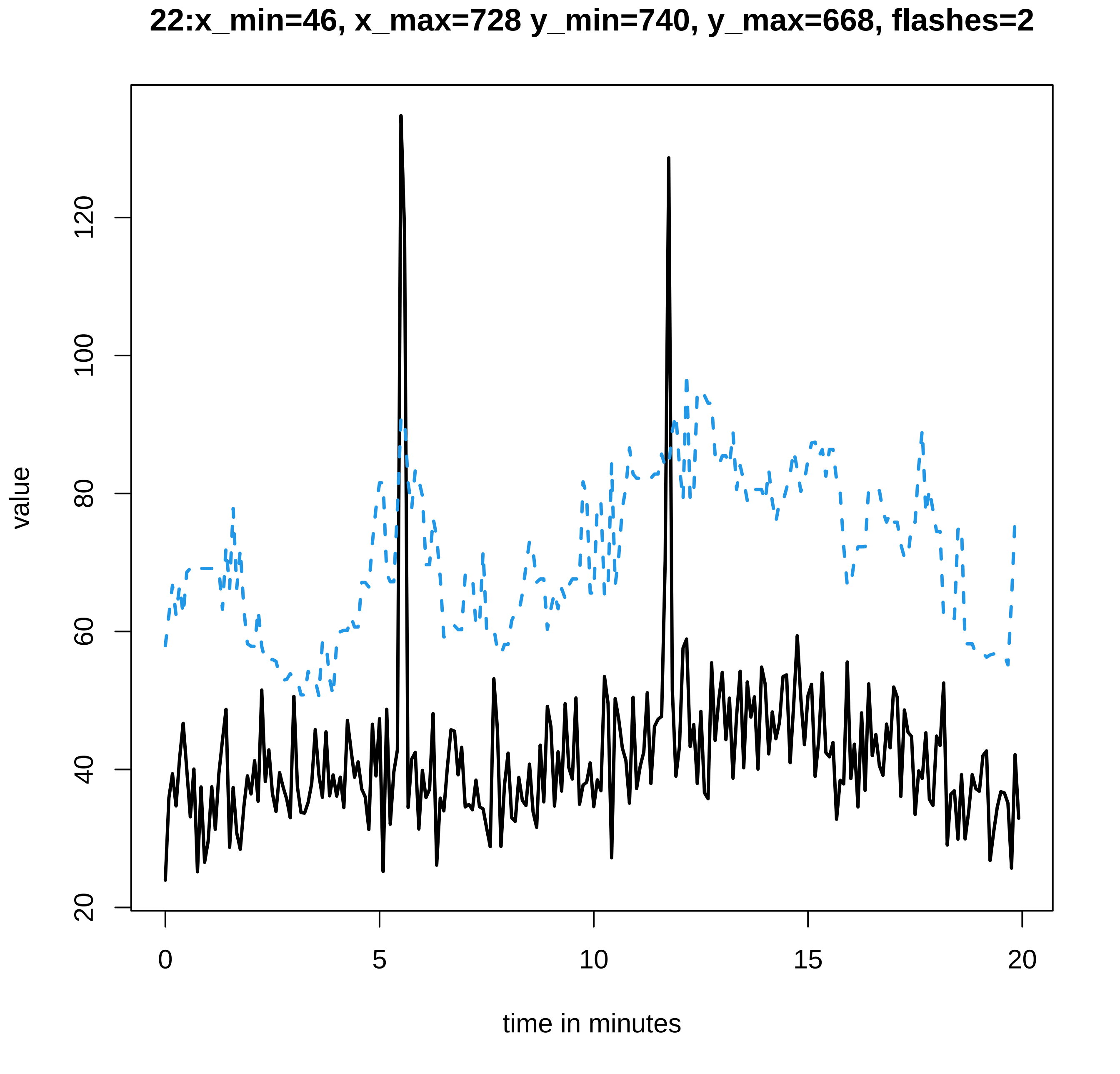
22:x_min=46, x_max=728 y_min=740, y_max=668, flashes=2
value
20
40
60
80
100
120
0
5
10
15
20
time in minutes

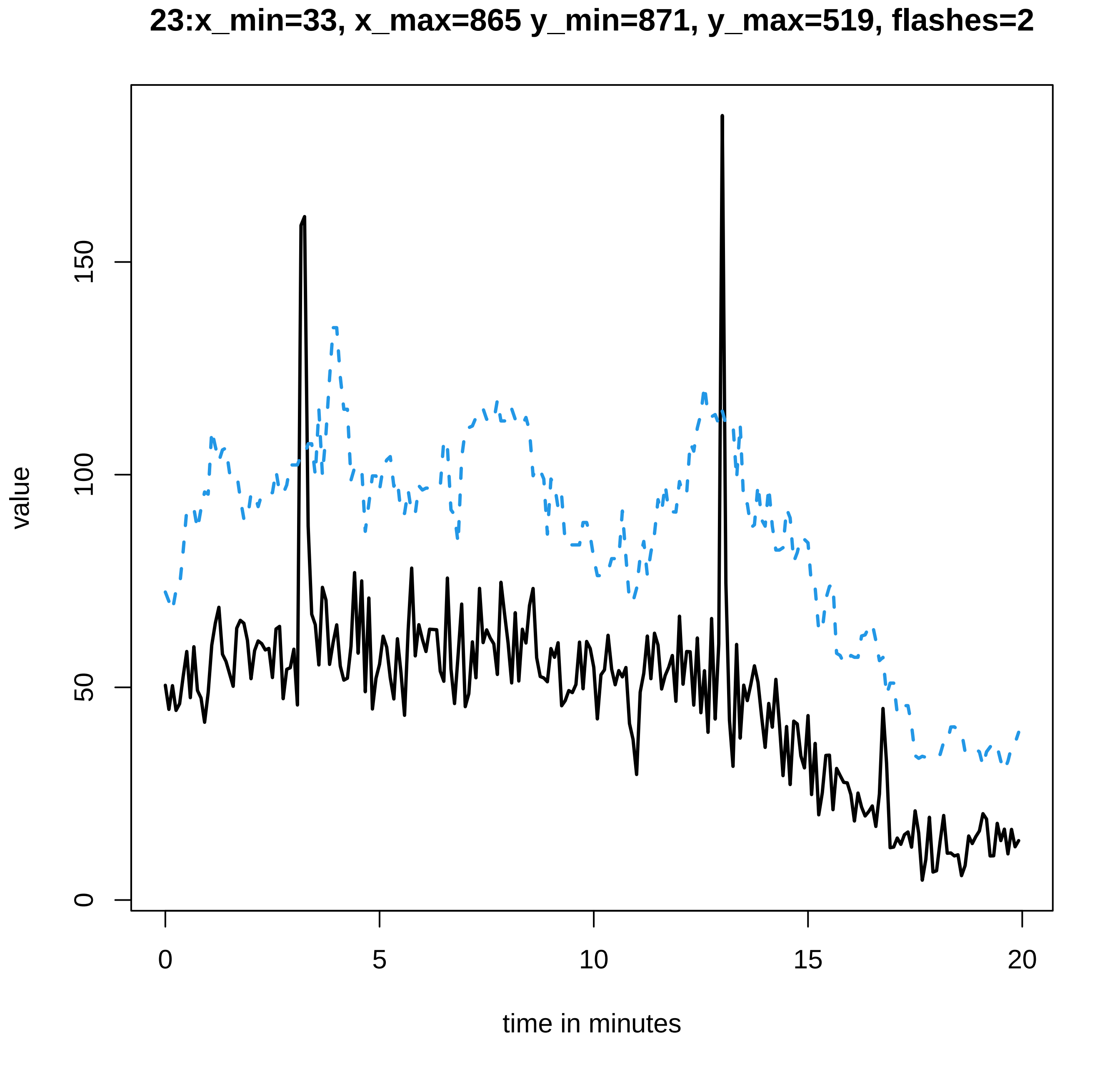

23:x_min=33, x_max=865 y_min=871, y_max=519, flashes=2
value
0
50
100
150
0
5
10
15
20
time in minutes

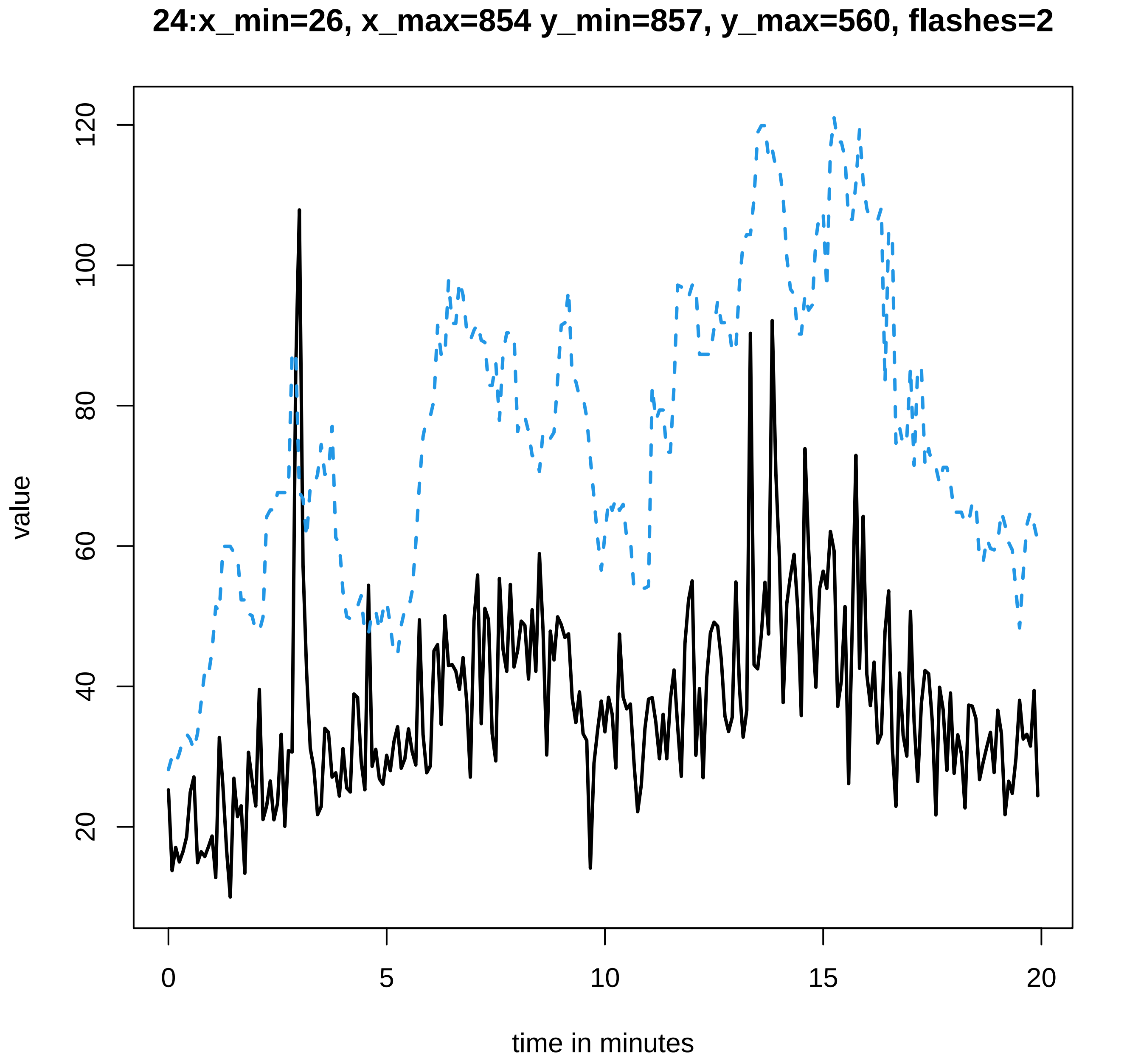
24:x_min=26, x_max=854 y_min=857, y_max=560, flashes=2
value
20
40
60
80
100
120
0
5
10
15
20
time in minutes

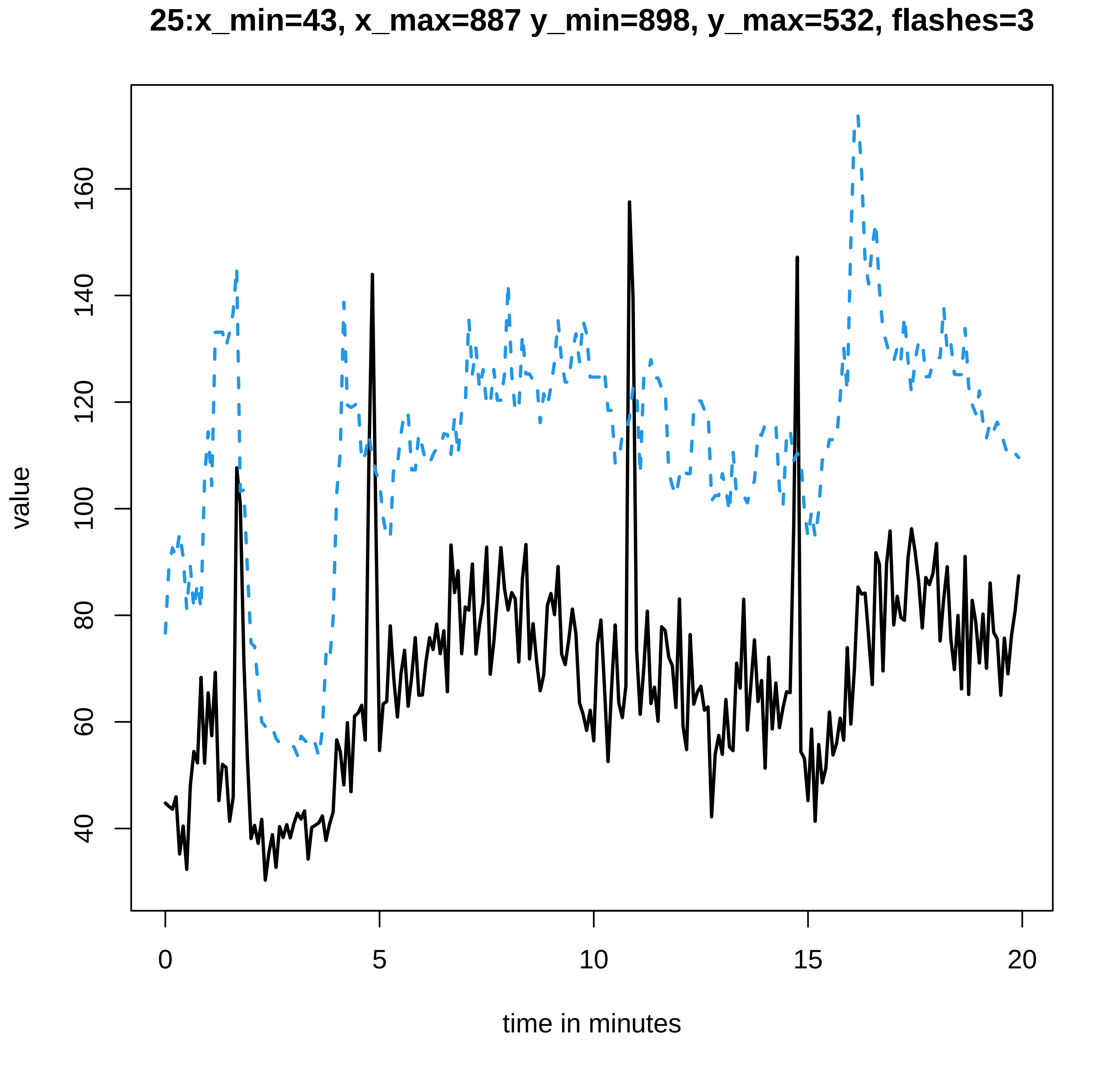

25:x_min=43, x_max=887 y_min=898, y_max=532, flashes=3
value
40
60
80
100
120
140
160
0
5
10
15
20
time in minutes

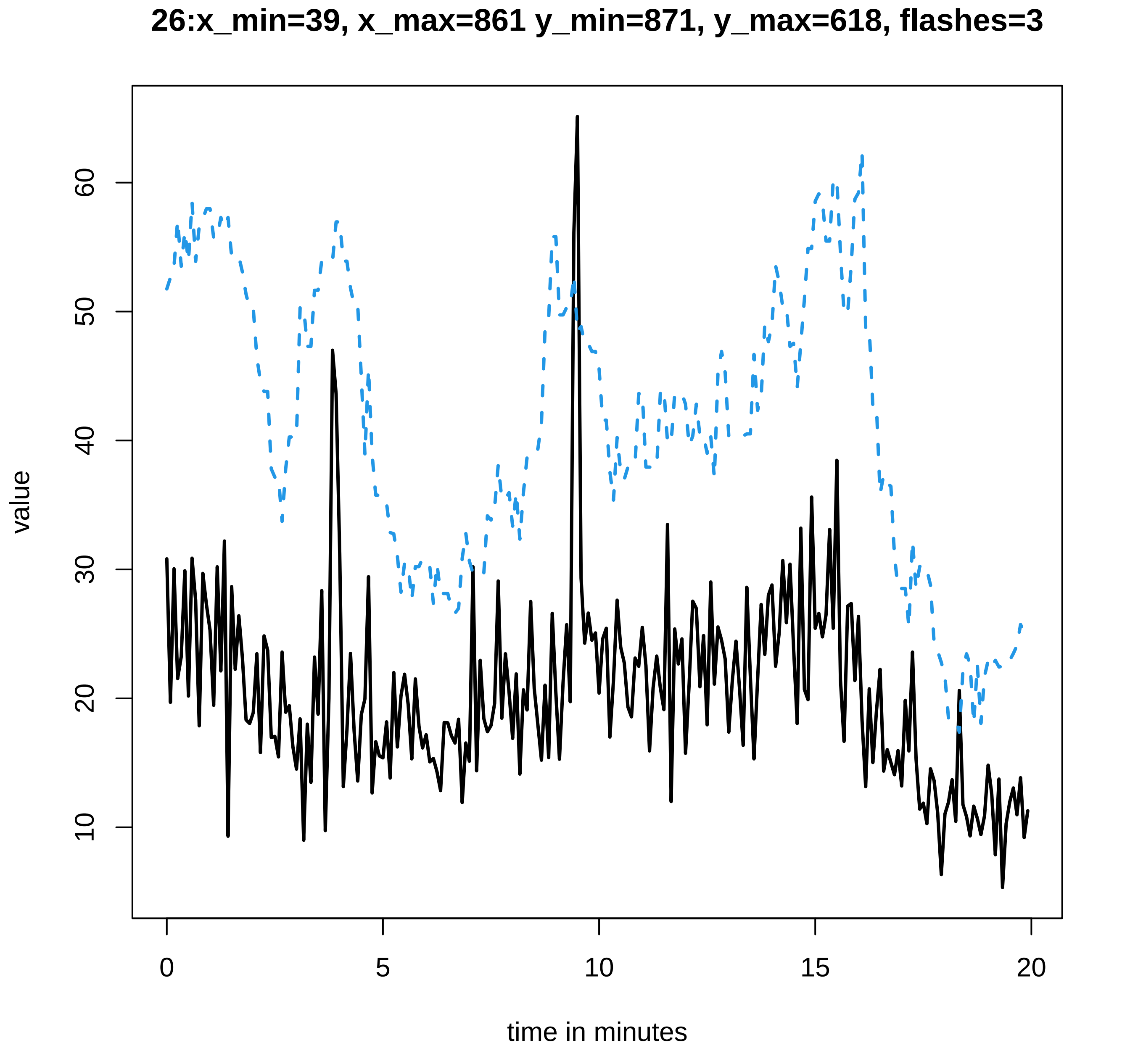
26:x_min=39, x_max=861 y_min=871, y_max=618, flashes=3
value
time in minutes
0
5
10
15
20
10
20
30
40
50
60

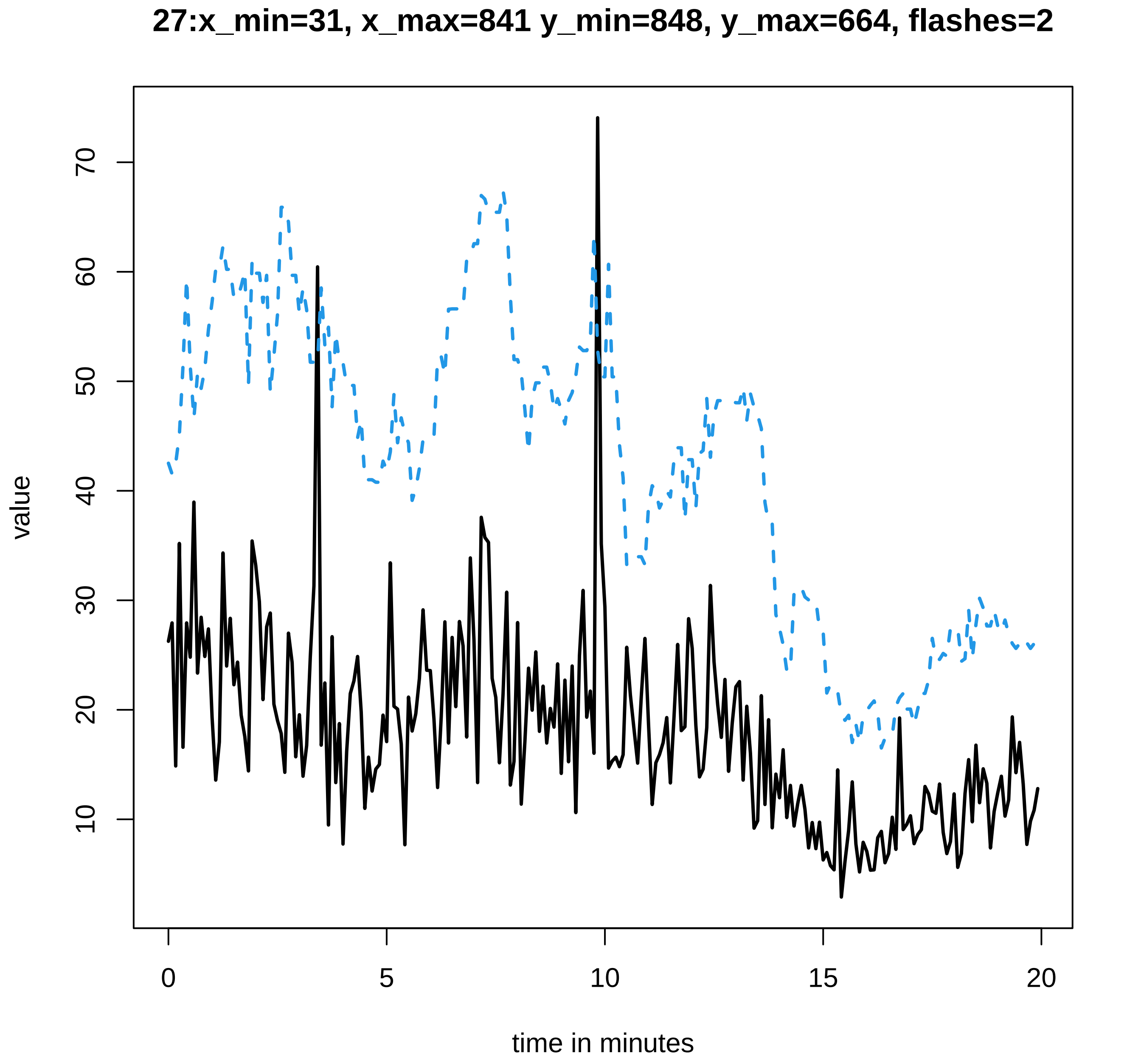
27:x_min=31, x_max=841 y_min=848, y_max=664, flashes=2
value
10
20
30
40
50
60
70
0
5
10
15
20
time in minutes

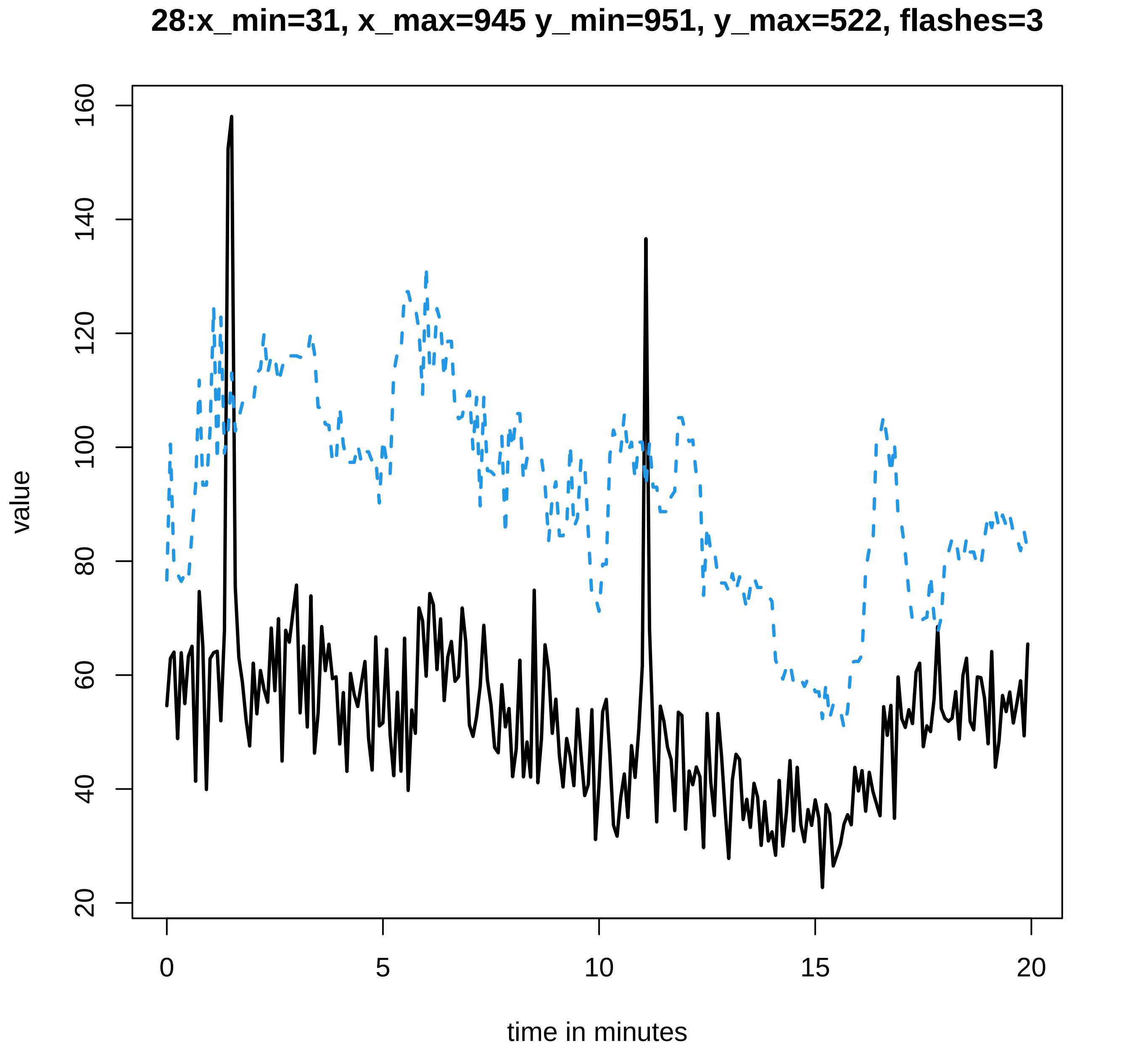
28:x_min=31, x_max=945 y_min=951, y_max=522, flashes=3
value
time in minutes
160
140
120
100
80
60
40
20
0
5
10
15
20

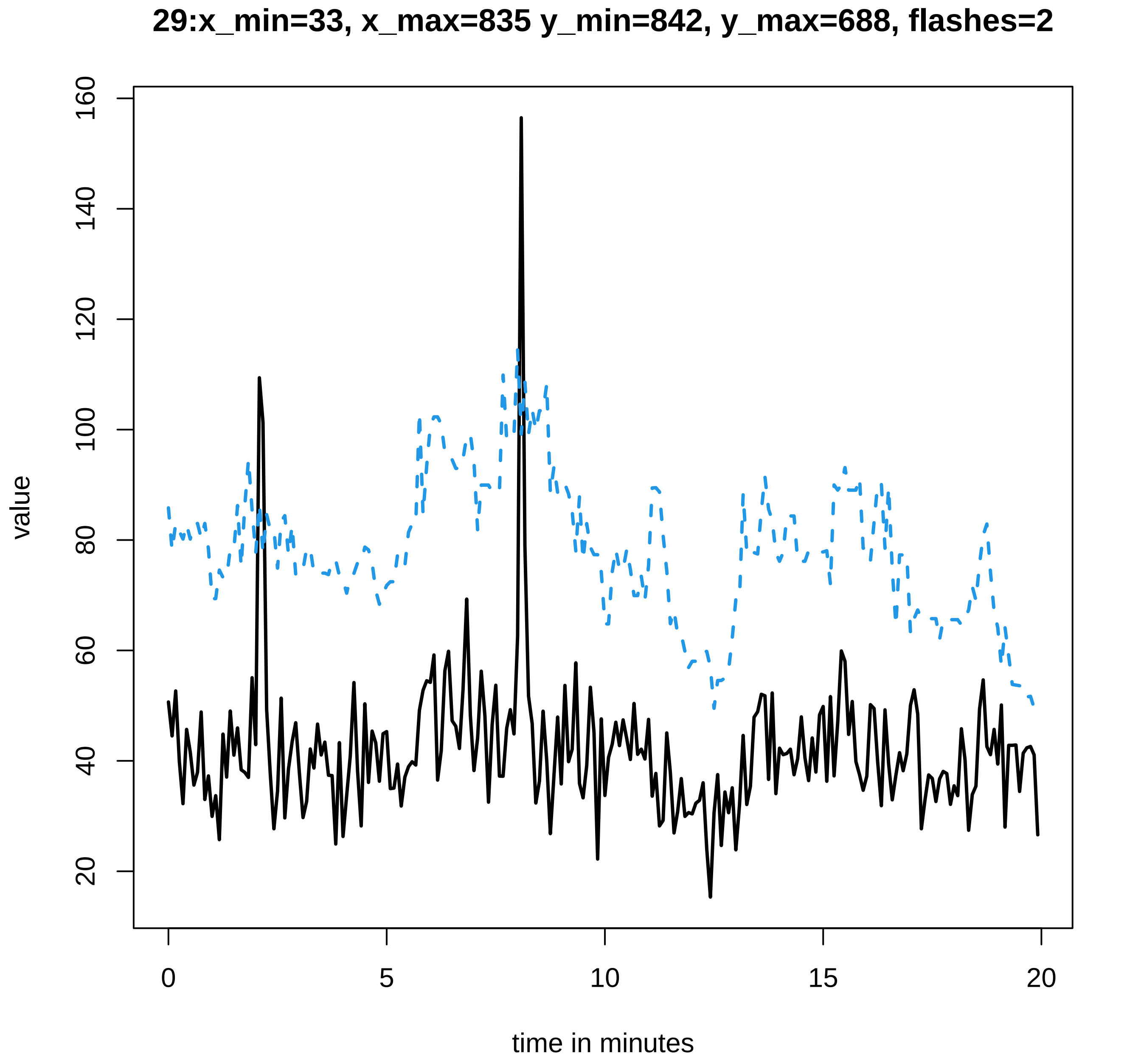
29:x_min=33, x_max=835 y_min=842, y_max=688, flashes=2
value
160
140
120
100
80
60
40
20
0
5
10
15
20
time in minutes

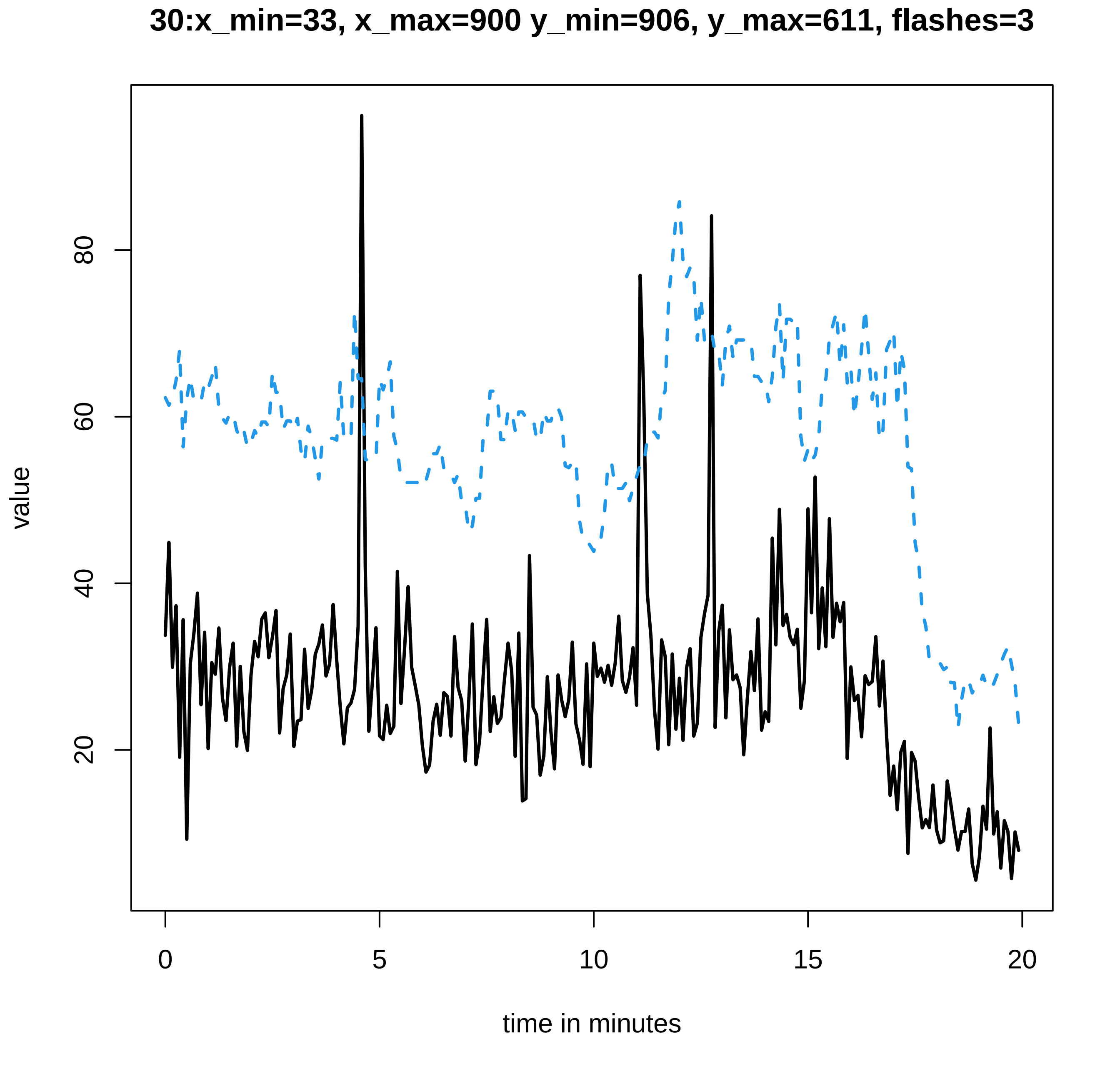
30:x_min=33, x_max=900 y_min=906, y_max=611, flashes=3
value
20
40
60
80
0
5
10
15
20
time in minutes

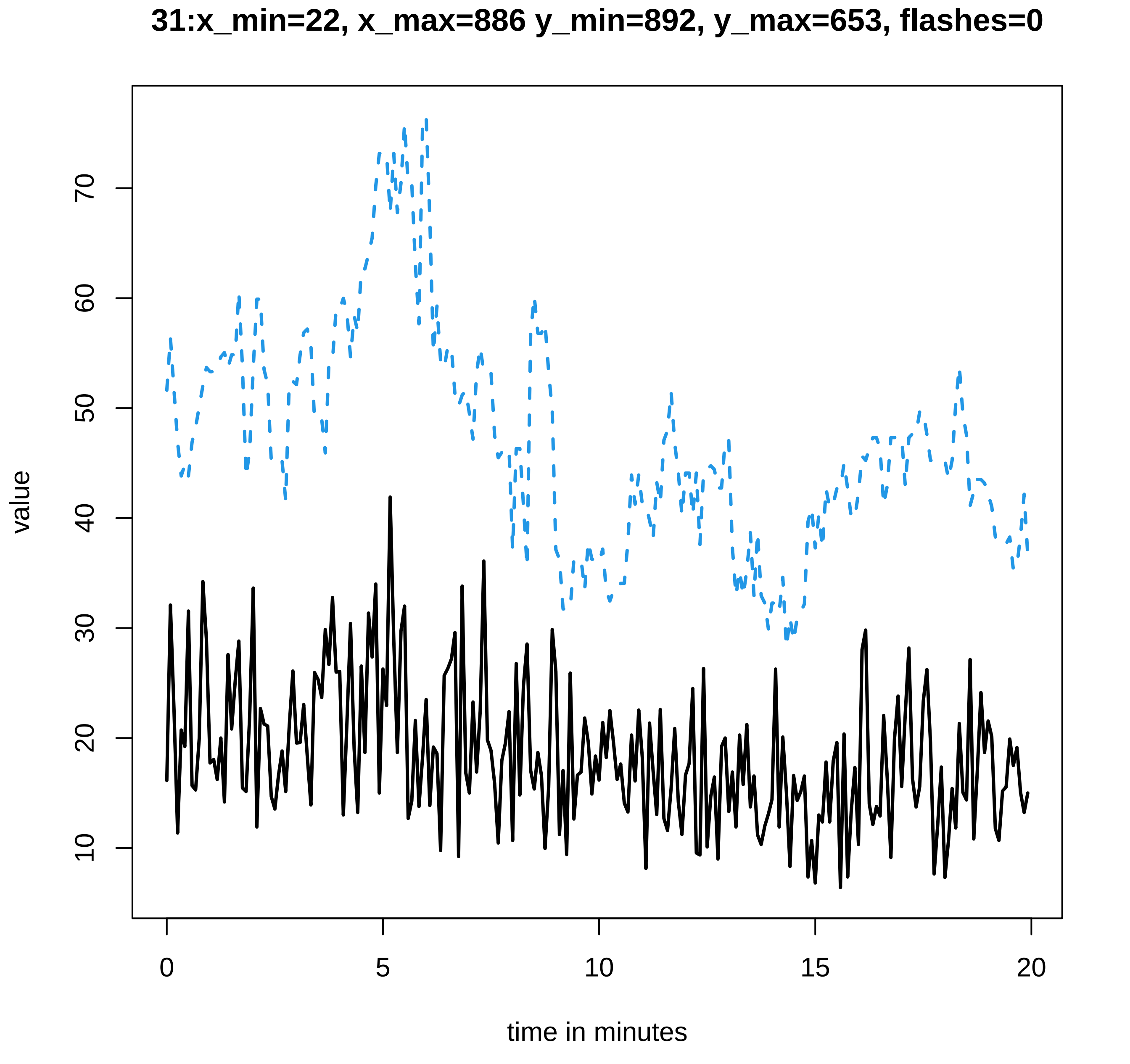
31:x_min=22, x_max=886 y_min=892, y_max=653, flashes=0
value
10
20
30
40
50
60
70
time in minutes
0
5
10
15
20

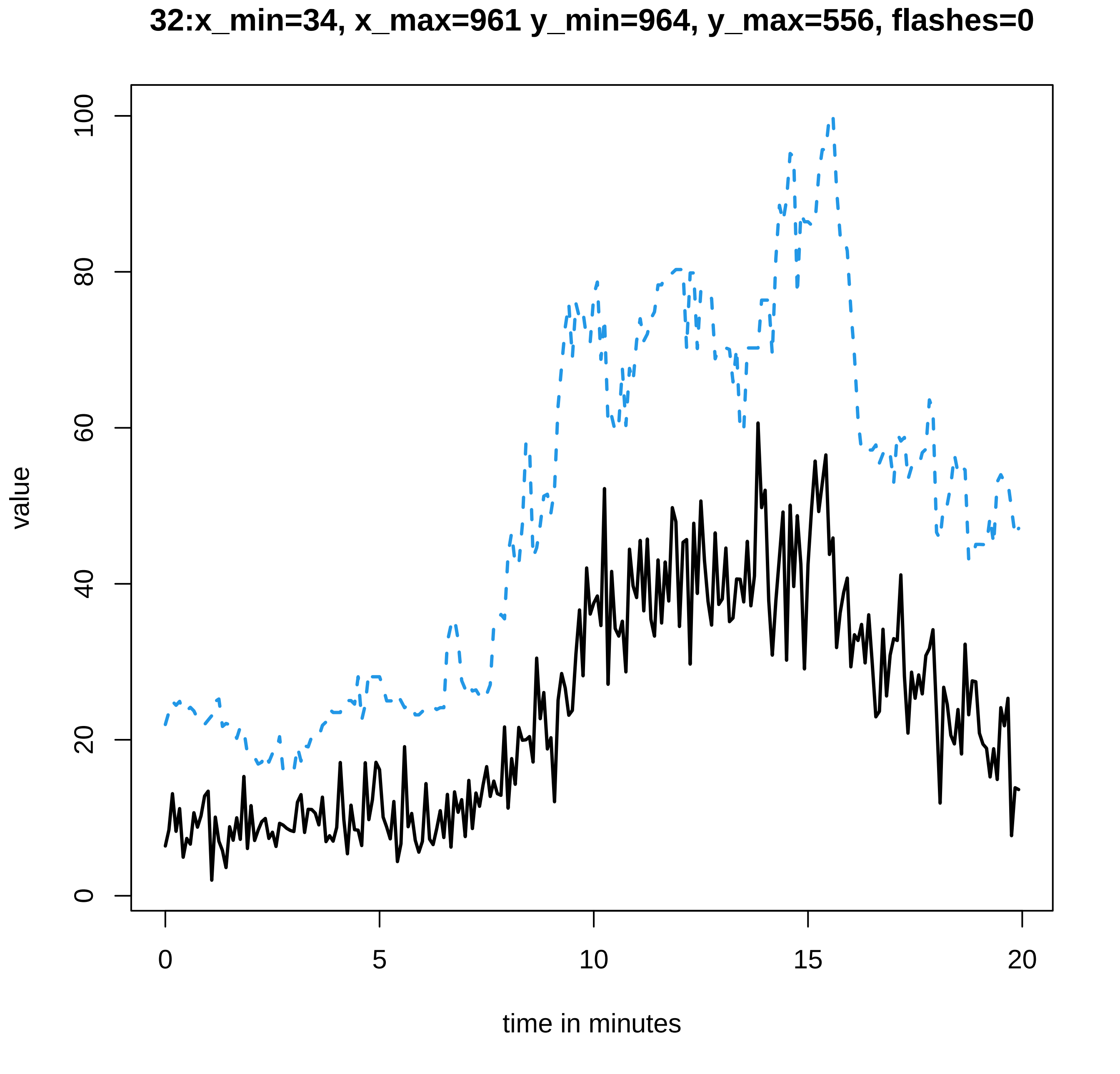

32:x_min=34, x_max=961 y_min=964, y_max=556, flashes=0
value
time in minutes
0
20
40
60
80
100
0
5
10
15
20

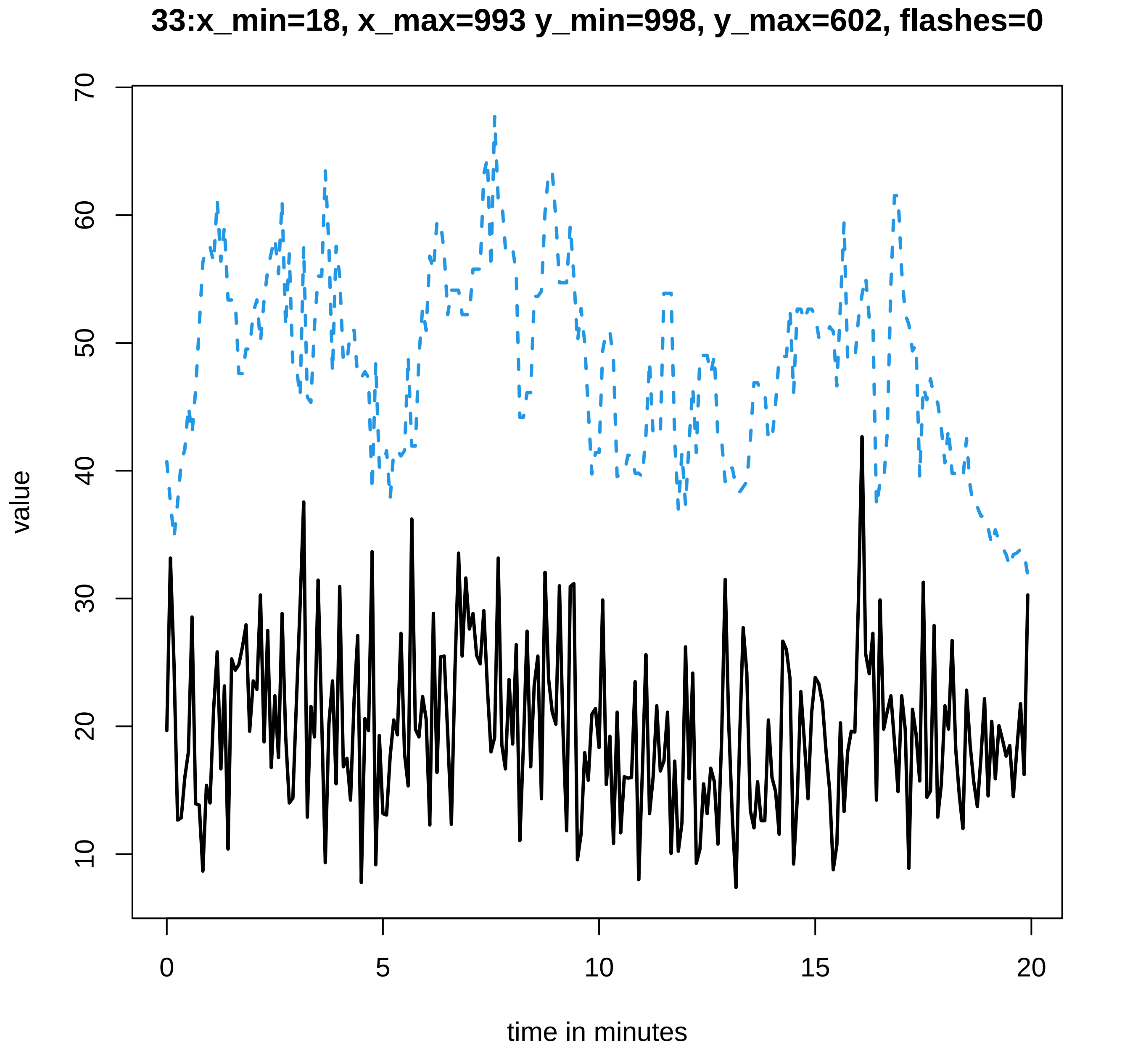
33:x_min=18, x_max=993 y_min=998, y_max=602, flashes=0
value
70
60
50
40
30
20
10
0
5
10
15
20
time in minutes

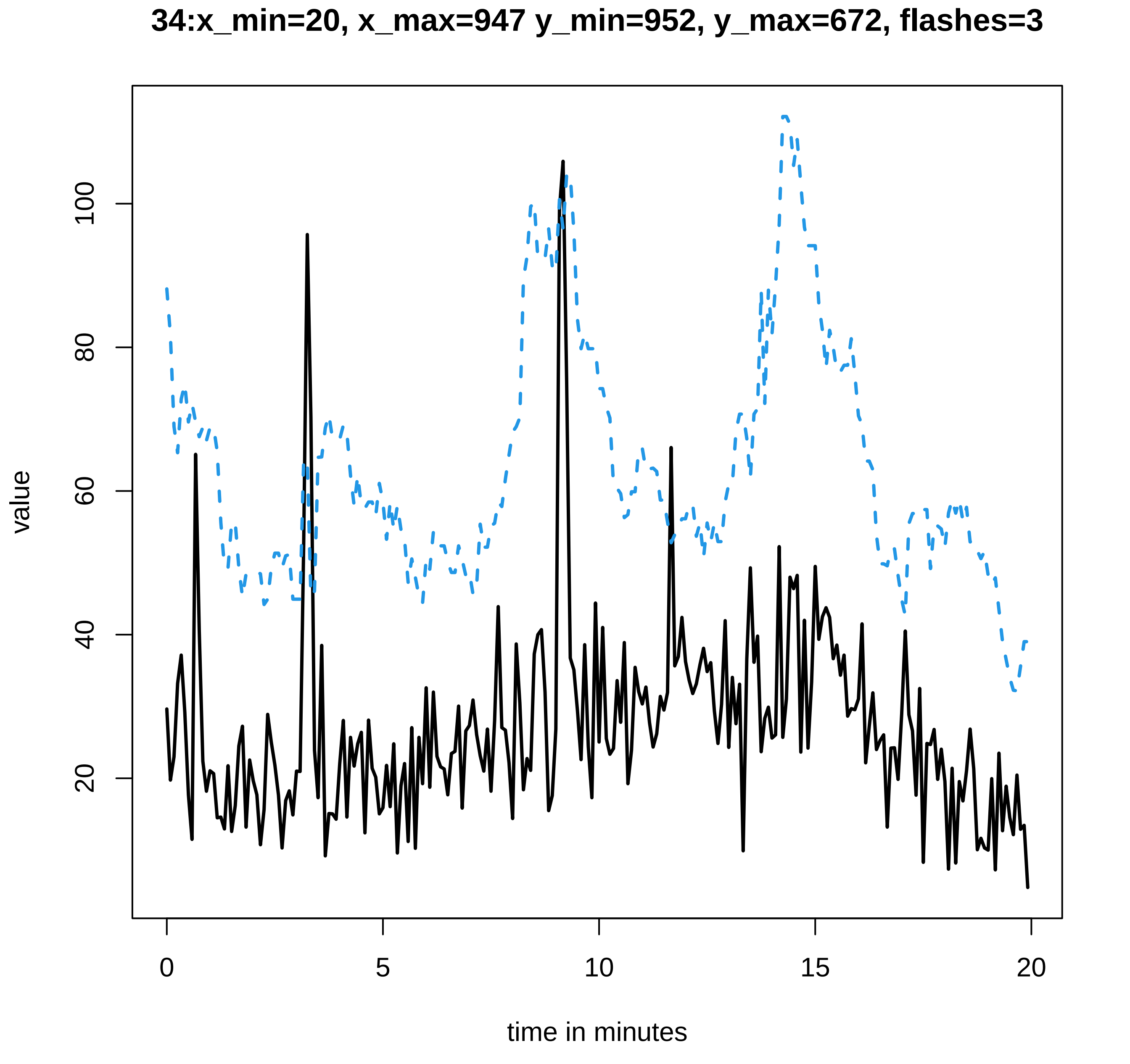
34:x_min=20, x_max=947 y_min=952, y_max=672, flashes=3
value
20
40
60
80
100
0
5
10
15
20
time in minutes

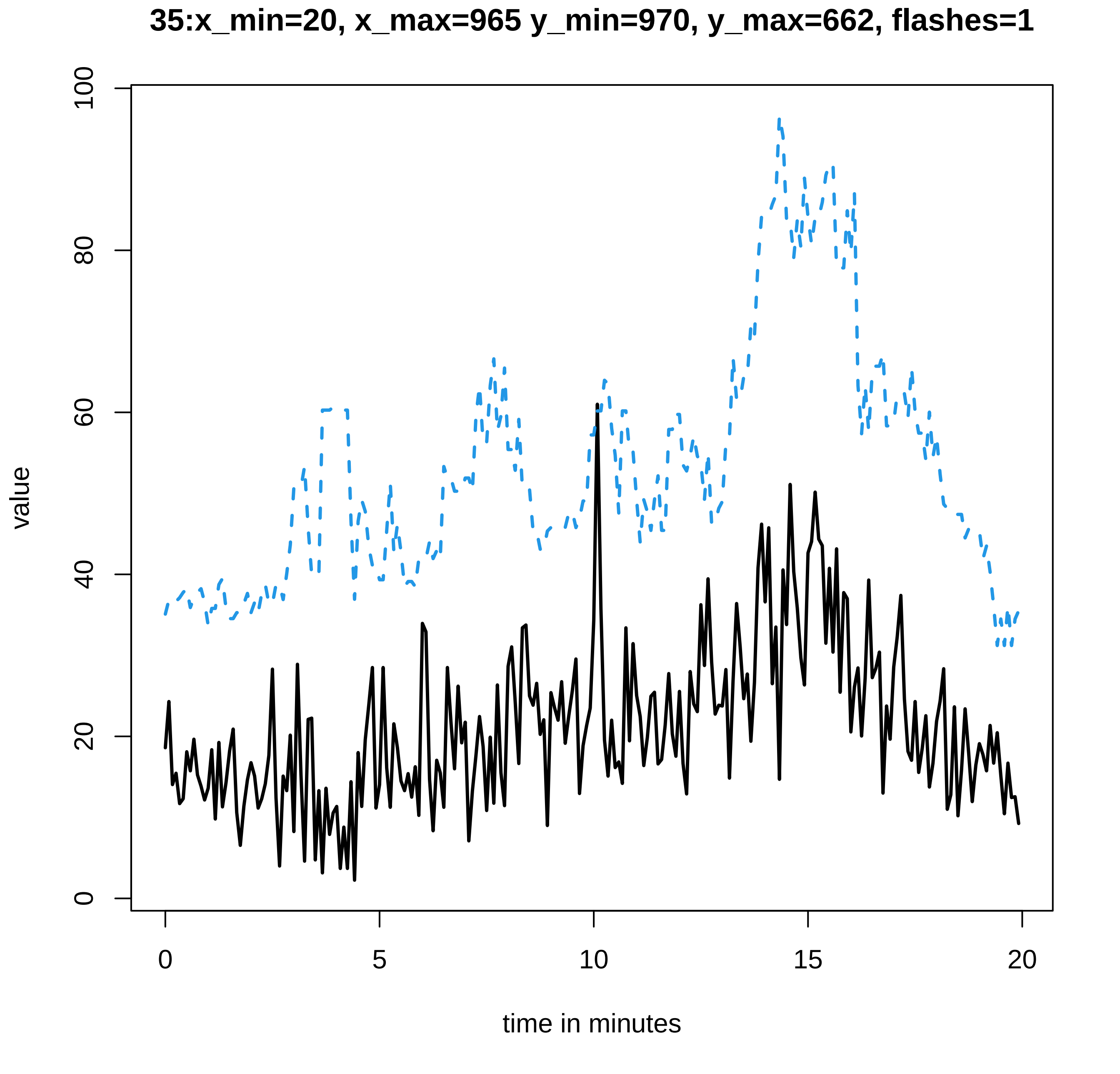

35:x_min=20, x_max=965 y_min=970, y_max=662, flashes=1
value
time in minutes
0
20
40
60
80
100
0
5
10
15
20

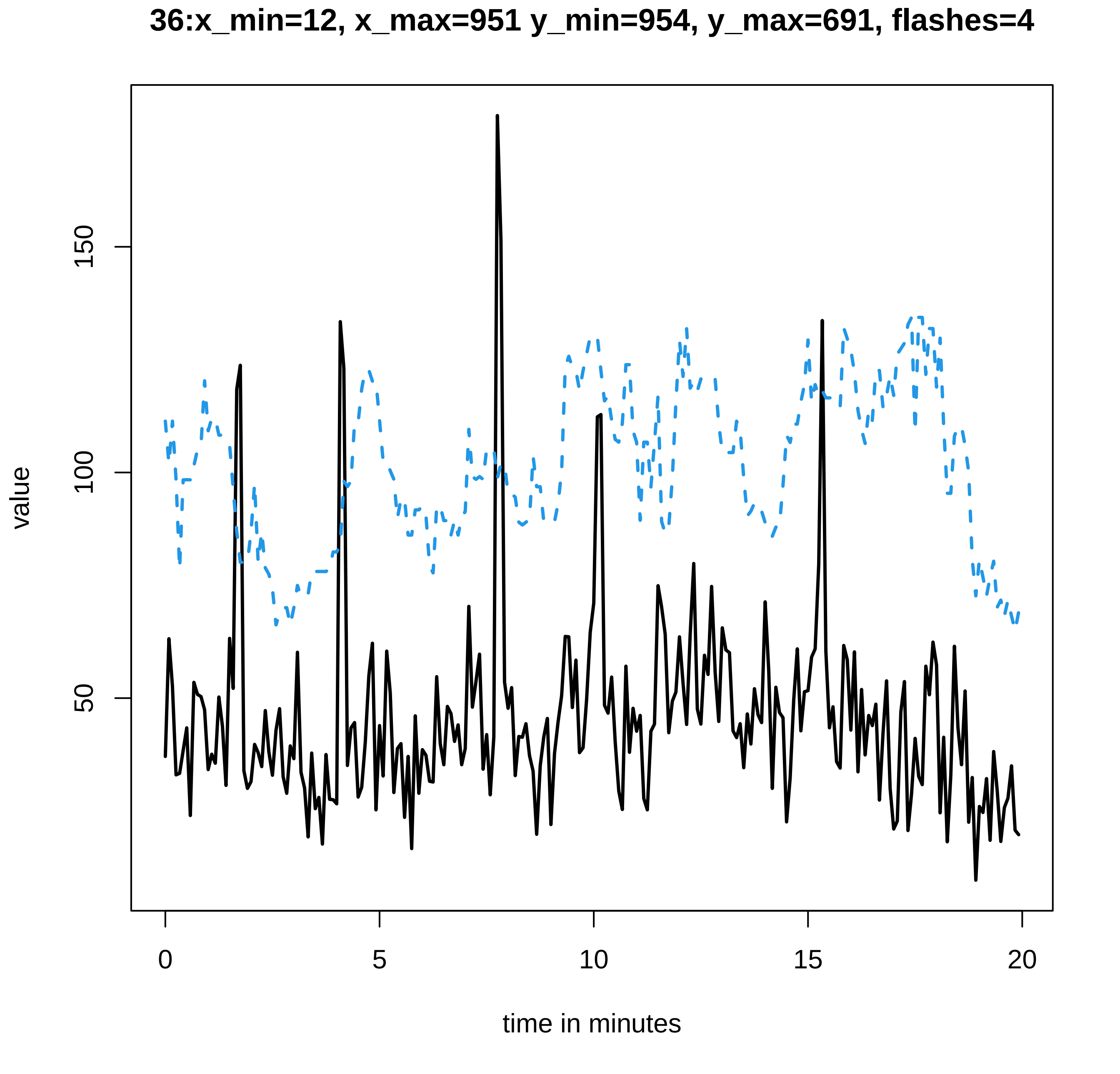
36:x_min=12, x_max=951 y_min=954, y_max=691, flashes=4
value
150
100
50
0
5
10
15
20
time in minutes

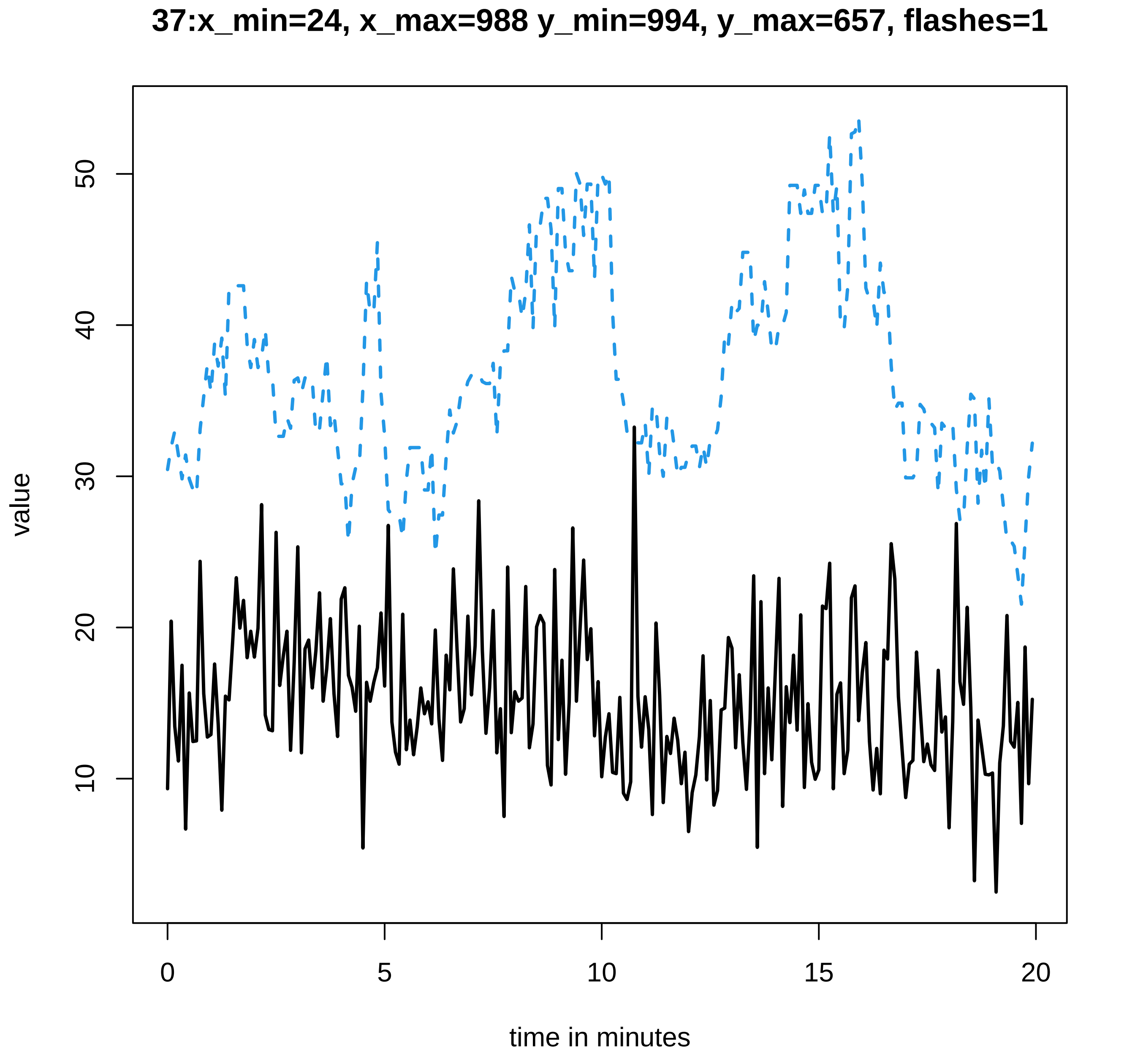
37:x_min=24, x_max=988 y_min=994, y_max=657, flashes=1
value
50
40
30
20
10
0
5
10
15
20
time in minutes

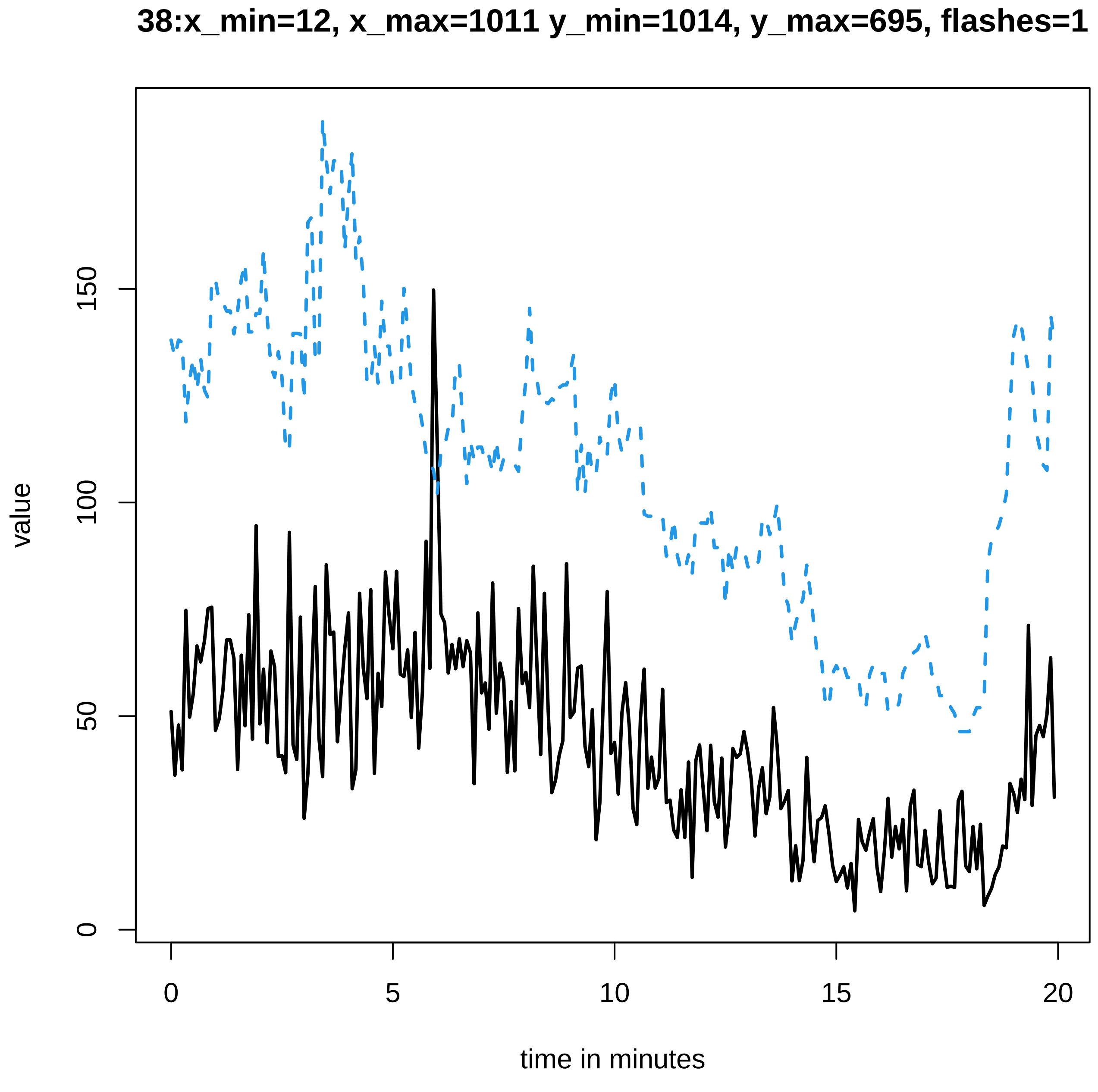
38:x_min=12, x_max=1011 y_min=1014, y_max=695, flashes=1
value
0
50
100
150
0
5
10
15
20
time in minutes
