## Supplementary data Calcium spiking for "LysM Receptor Proteins are Required for Ectomycorrhizal Symbiosis in Poplar": profiles-spiking-7.pdf

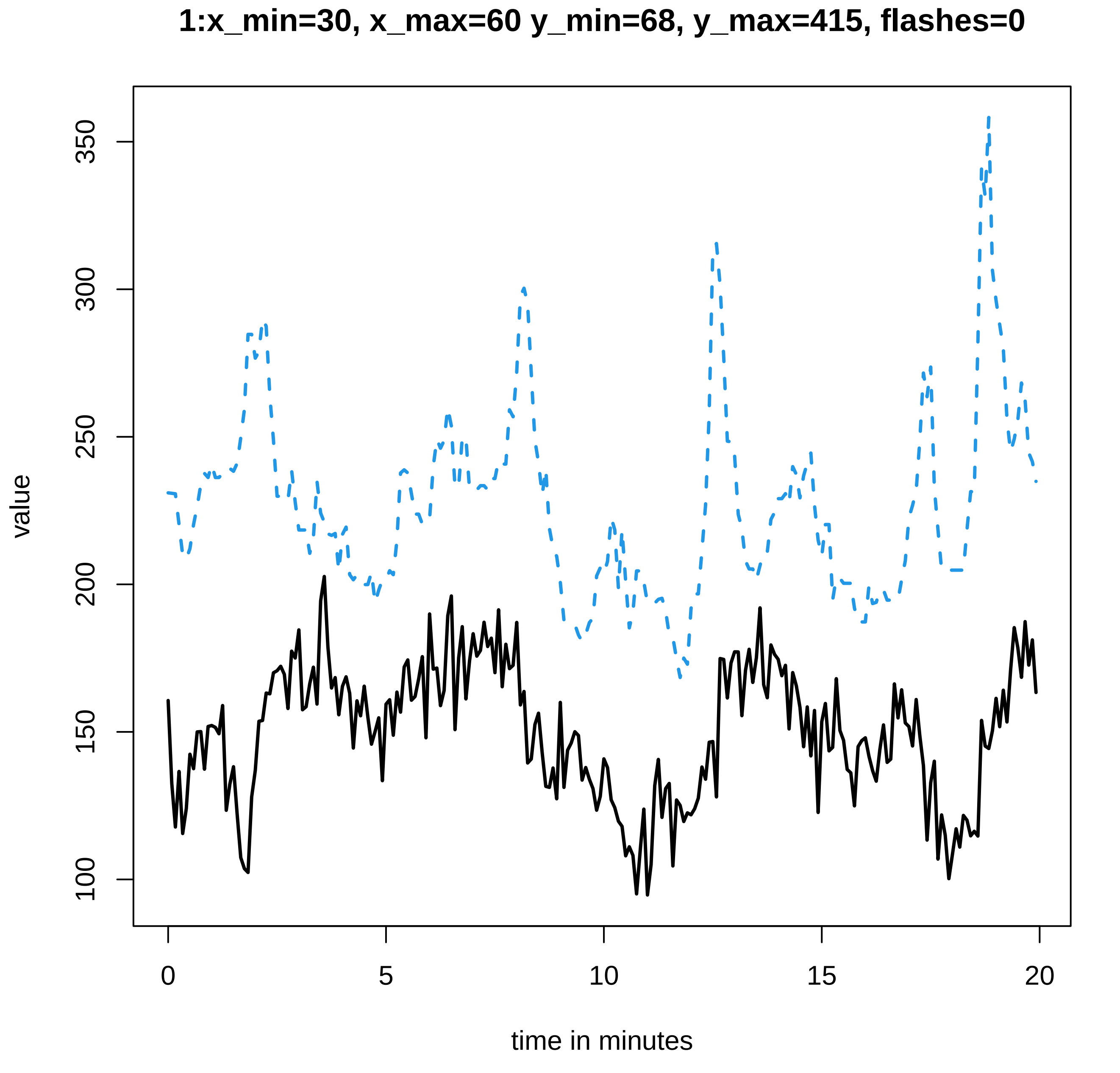
1:x_min=30, x_max=60 y_min=68, y_max=415, flashes=0
value
350
300
250
200
150
100
0
5
10
15
20
time in minutes

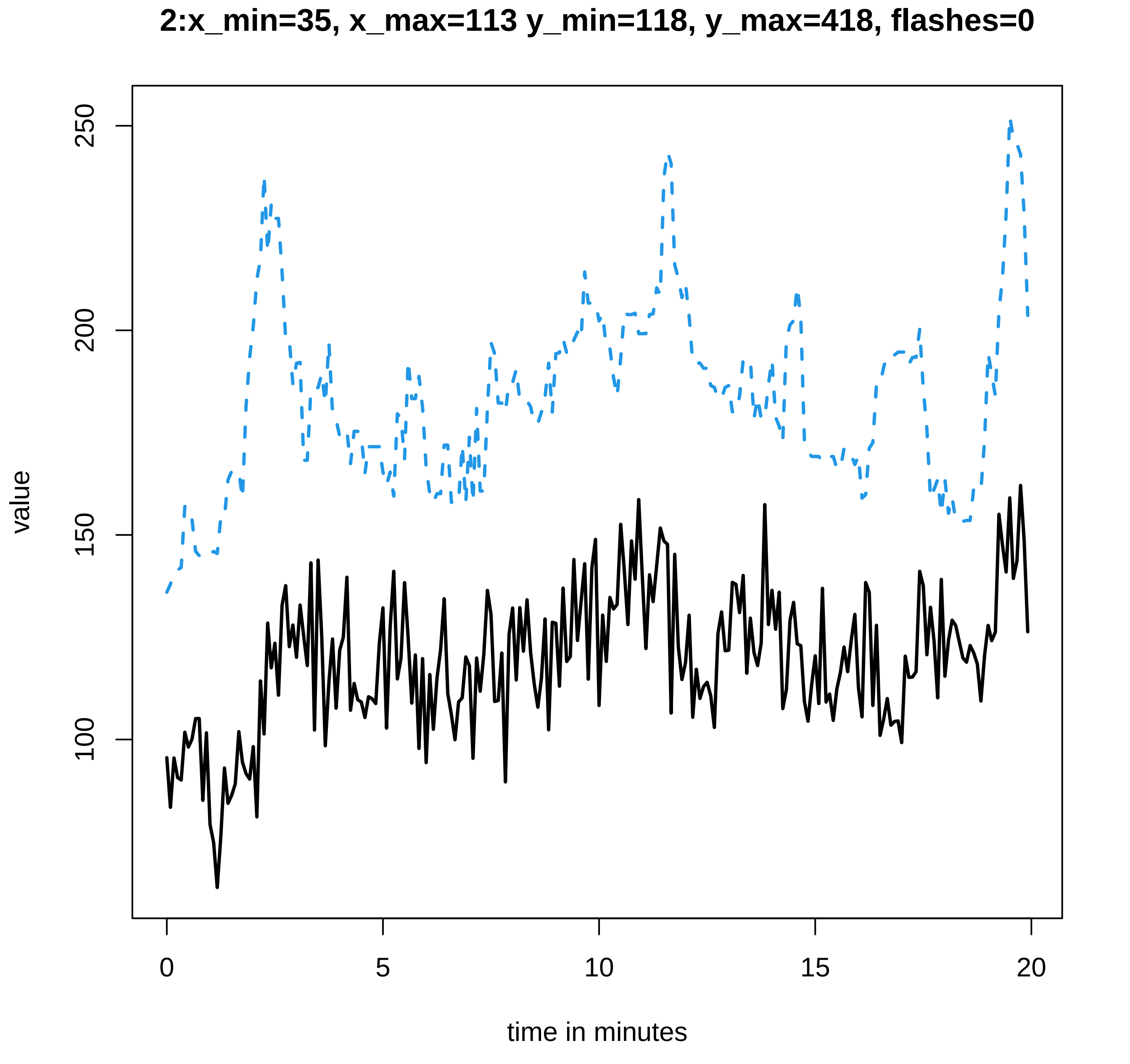

2:x_min=35, x_max=113 y_min=118, y_max=418, flashes=0
value
250
200
150
100
0
5
10
15
20
time in minutes

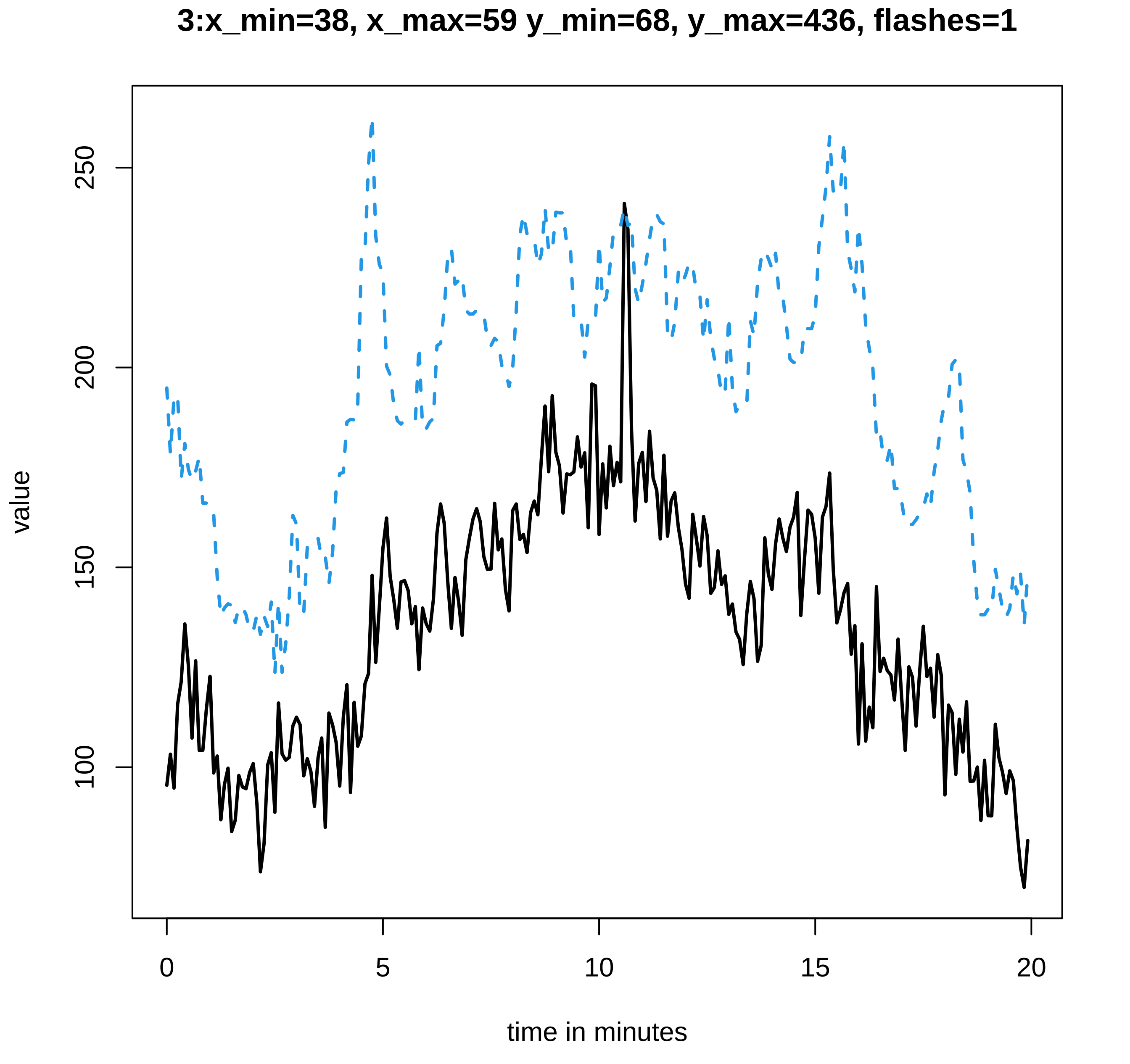
3:x_min=38, x_max=59 y_min=68, y_max=436, flashes=1
value
100
150
200
250
0
5
10
15
20
time in minutes

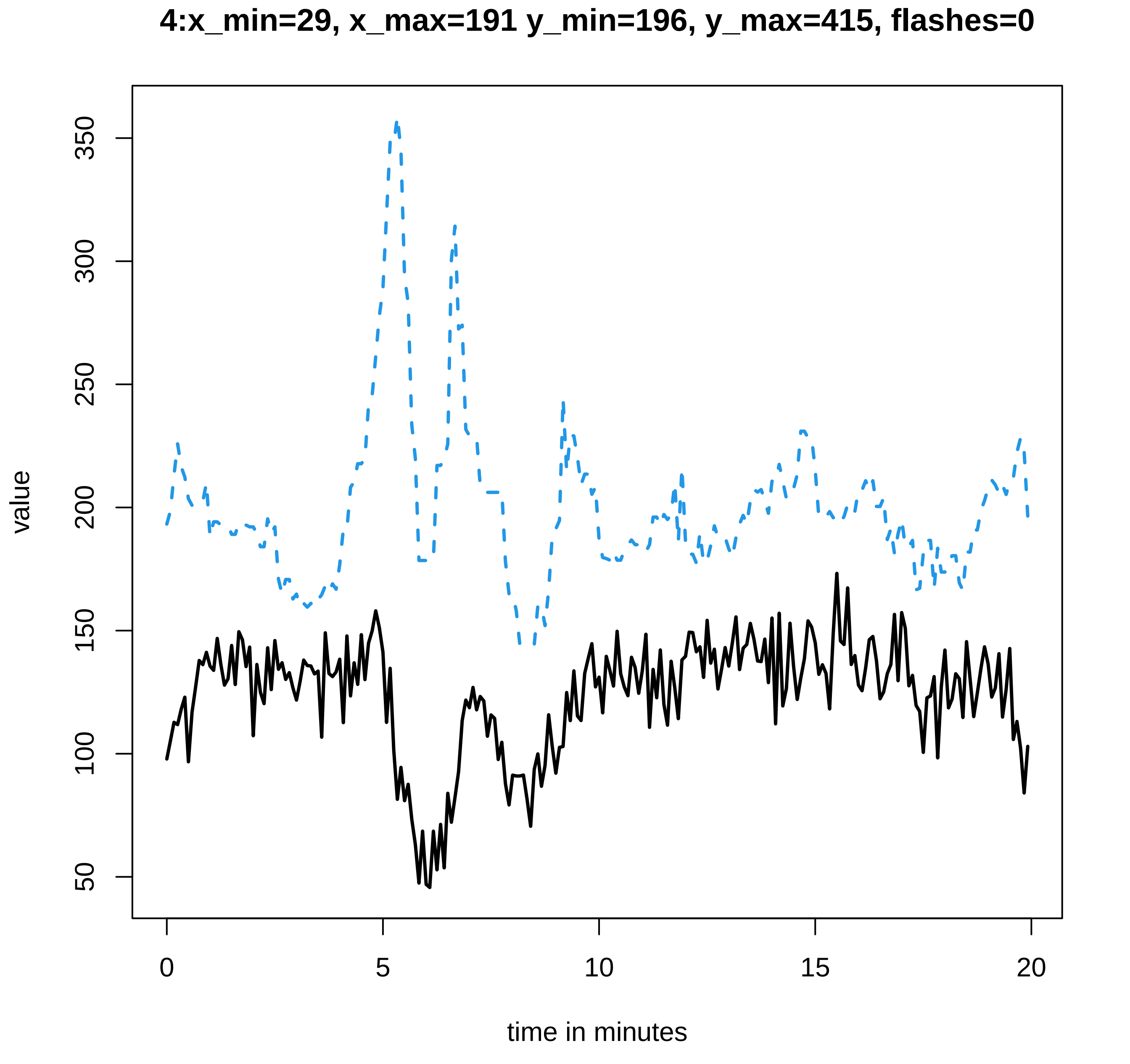

4:x_min=29, x_max=191 y_min=196, y_max=415, flashes=0
value
350
300
250
200
150
100
50
0
5
10
15
20
time in minutes

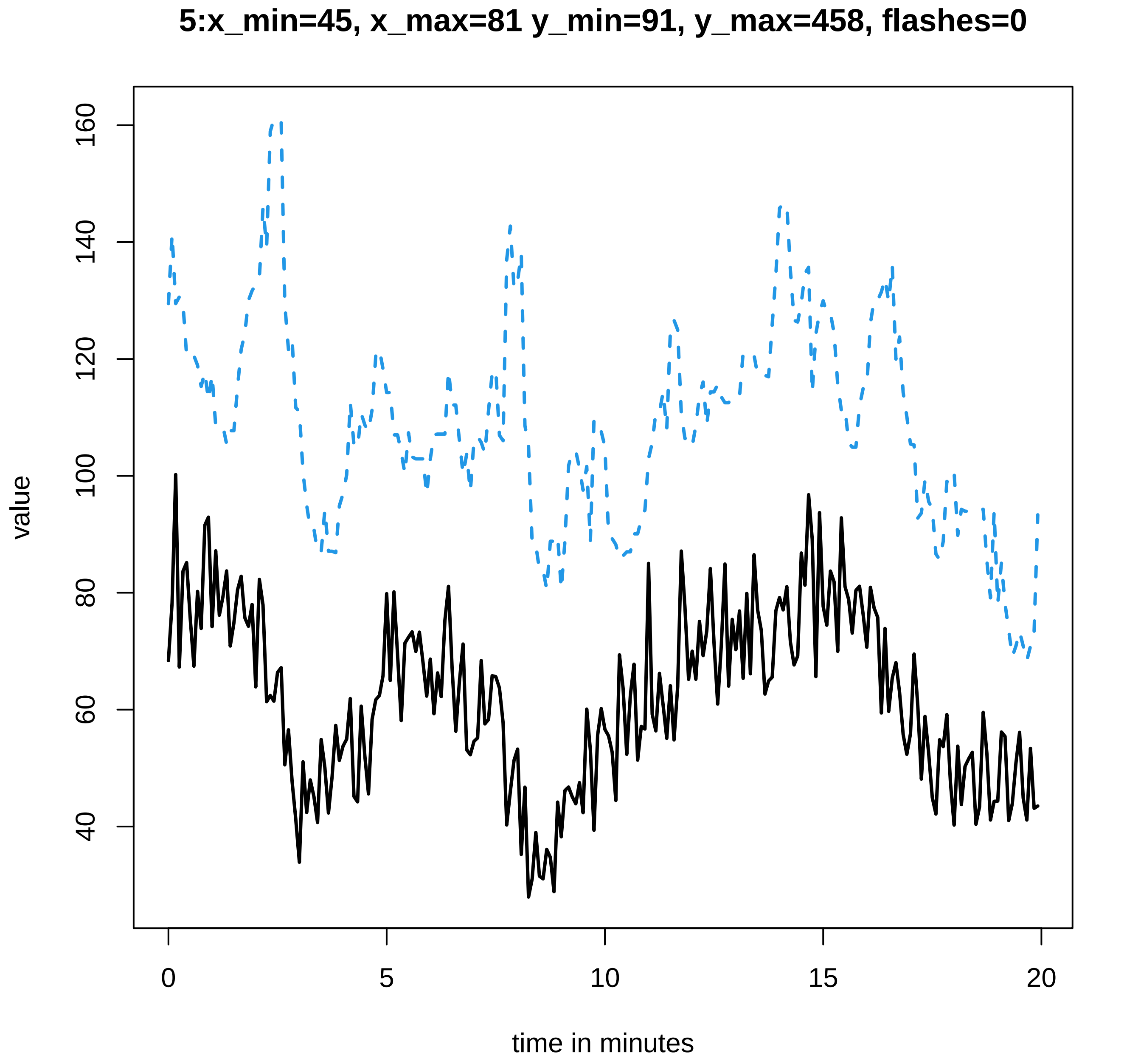
5:x_min=45, x_max=81 y_min=91, y_max=458, flashes=0
value
time in minutes
0
5
10
15
20
40
60
80
100
120
140
160

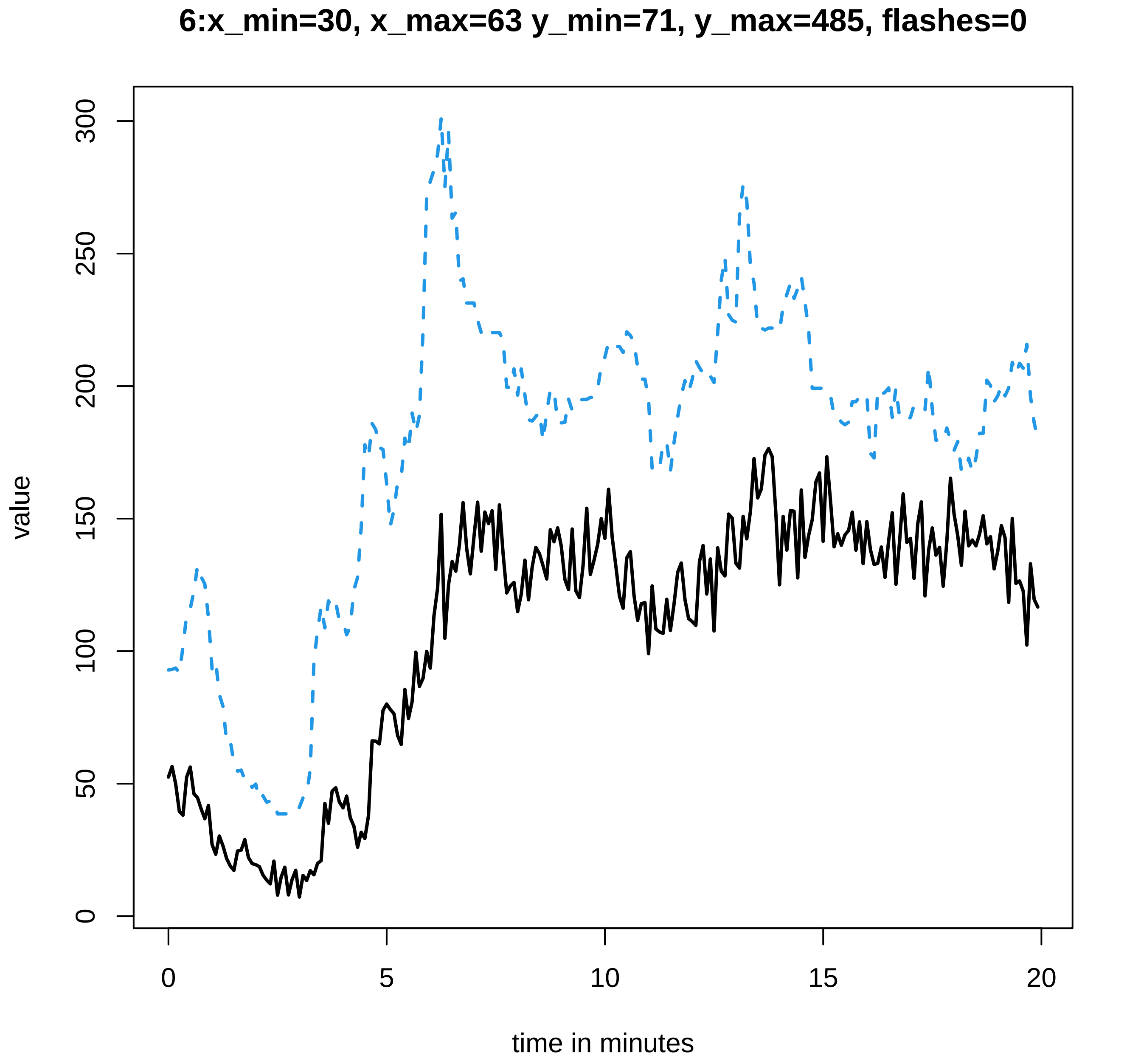
6:x_min=30, x_max=63 y_min=71, y_max=485, flashes=0
value
time in minutes
0
50
100
150
200
250
300
0
5
10
15
20

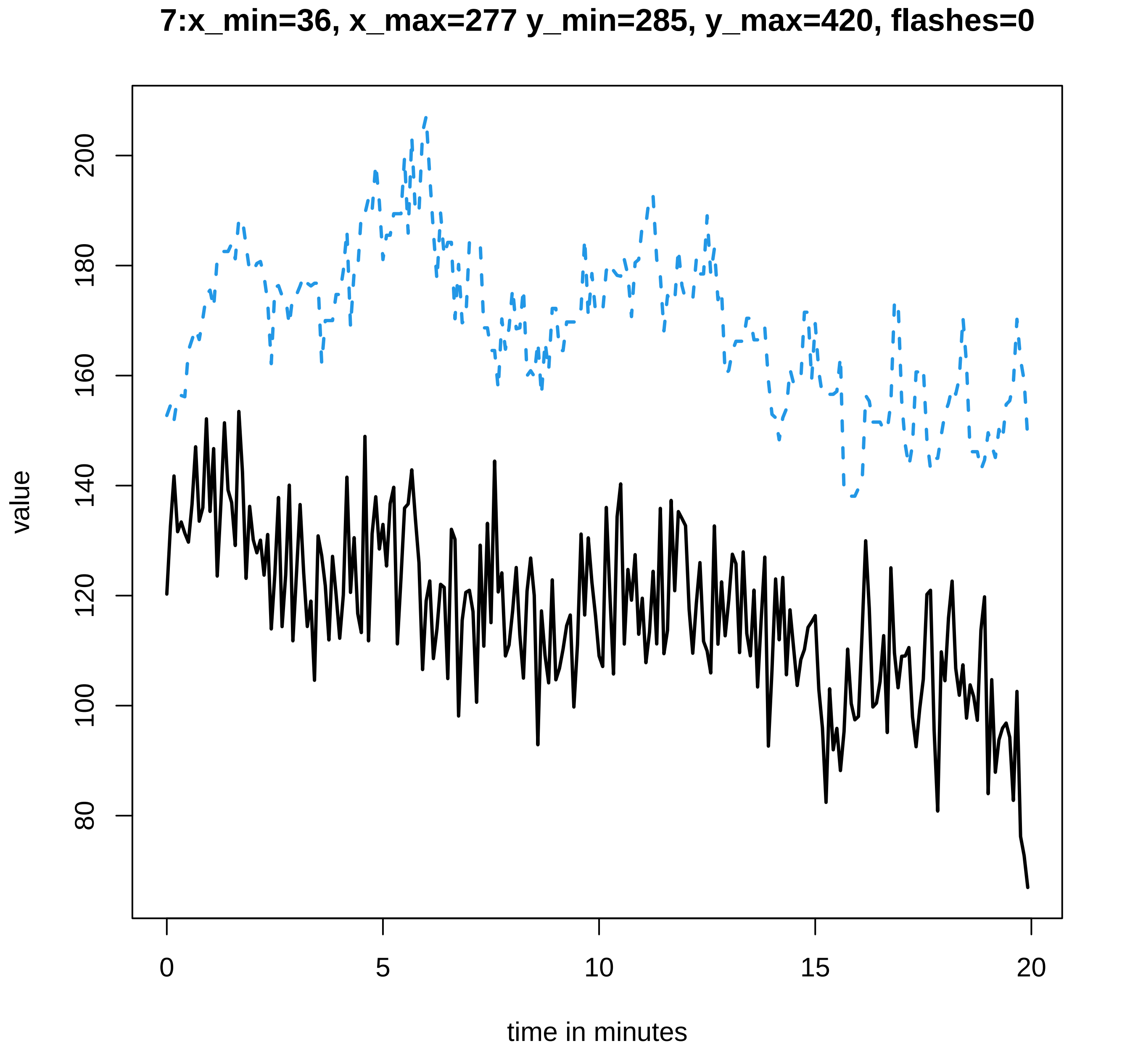

7:x_min=36, x_max=277 y_min=285, y_max=420, flashes=0
value
200
180
160
140
120
100
80
0
5
10
15
20
time in minutes

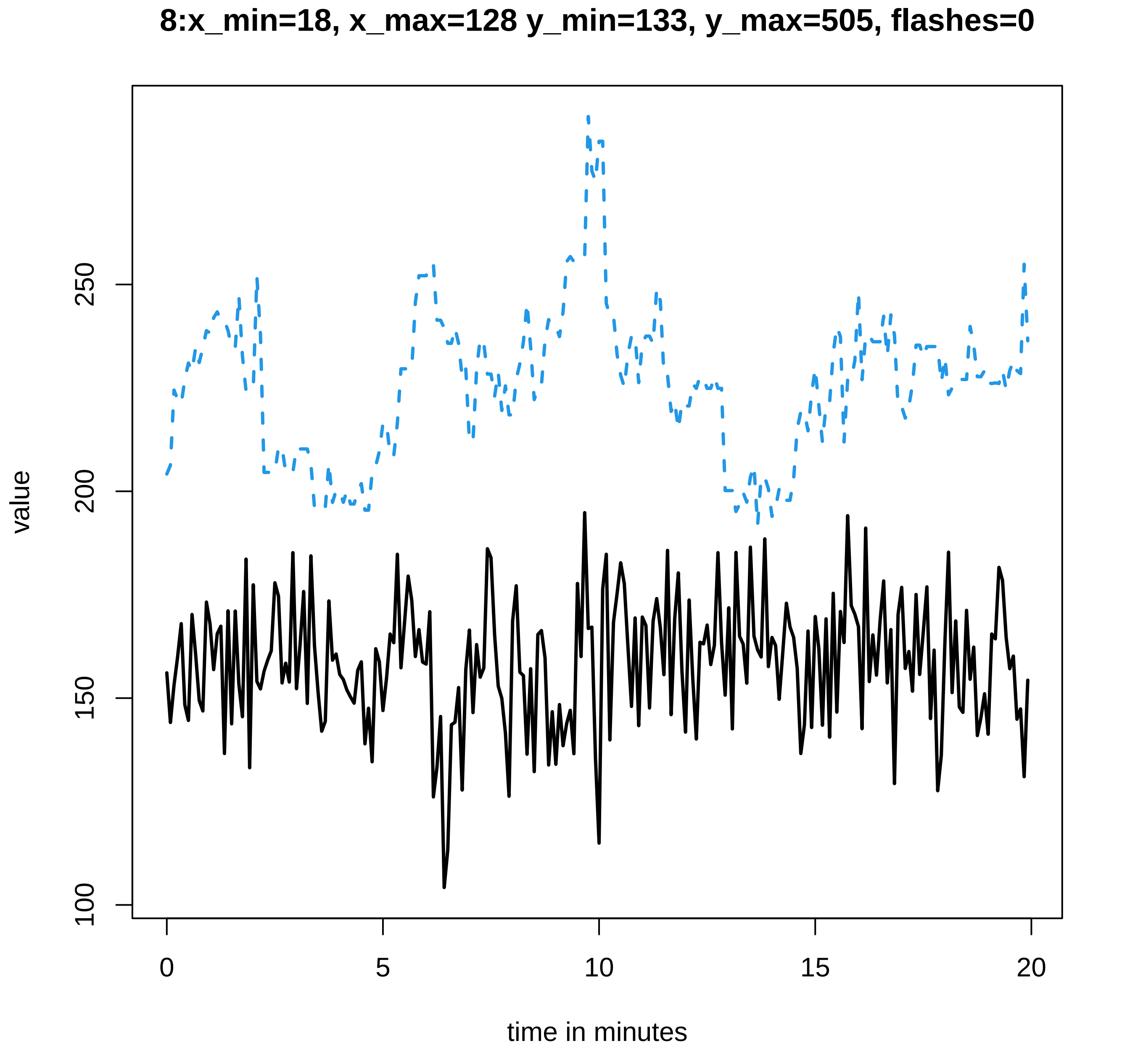
8:x_min=18, x_max=128 y_min=133, y_max=505, flashes=0
value
100
150
200
250
0
5
10
15
20
time in minutes

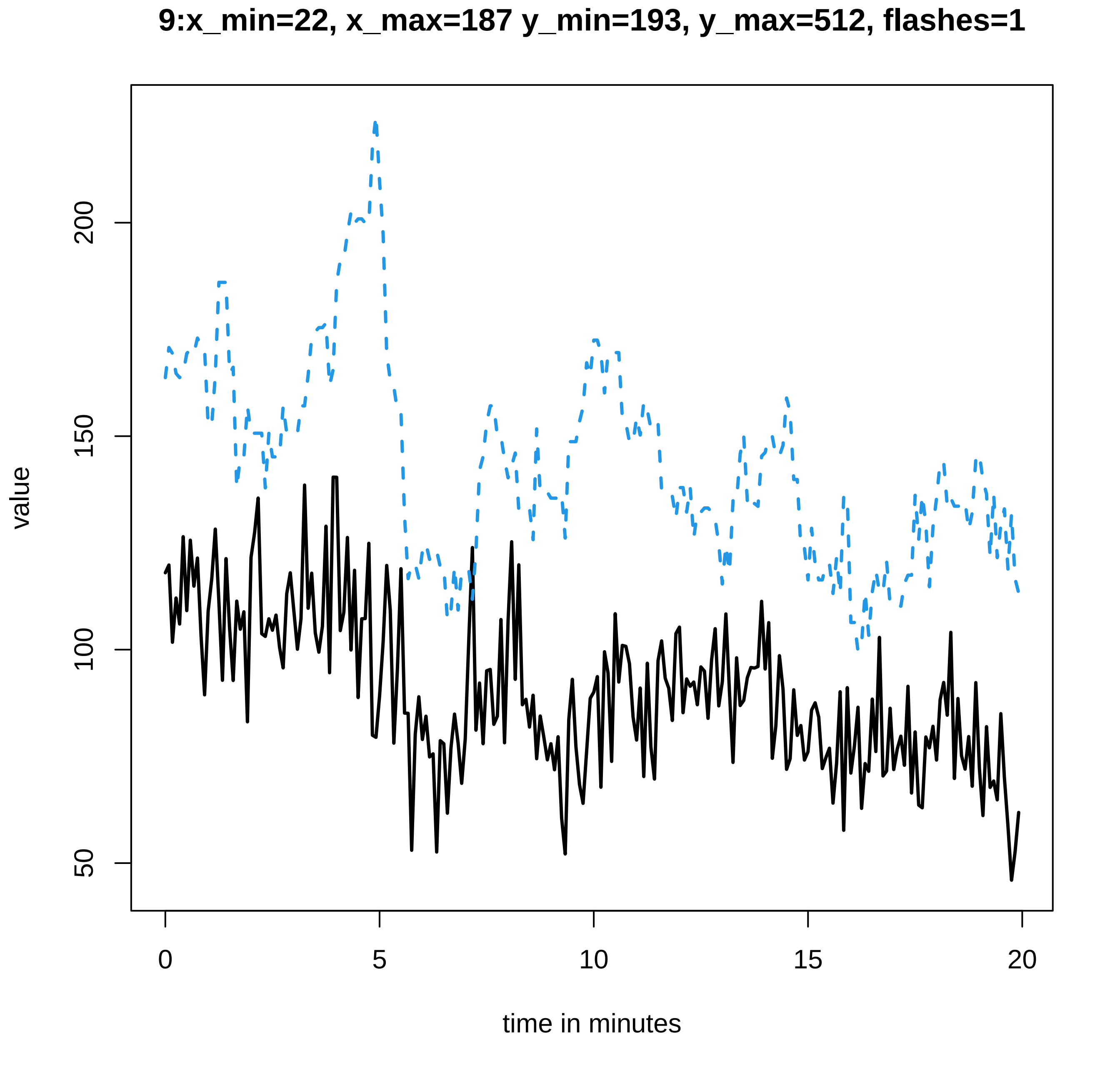

9:x_min=22, x_max=187 y_min=193, y_max=512, flashes=1
value
200
150
100
50
0
5
10
15
20
time in minutes

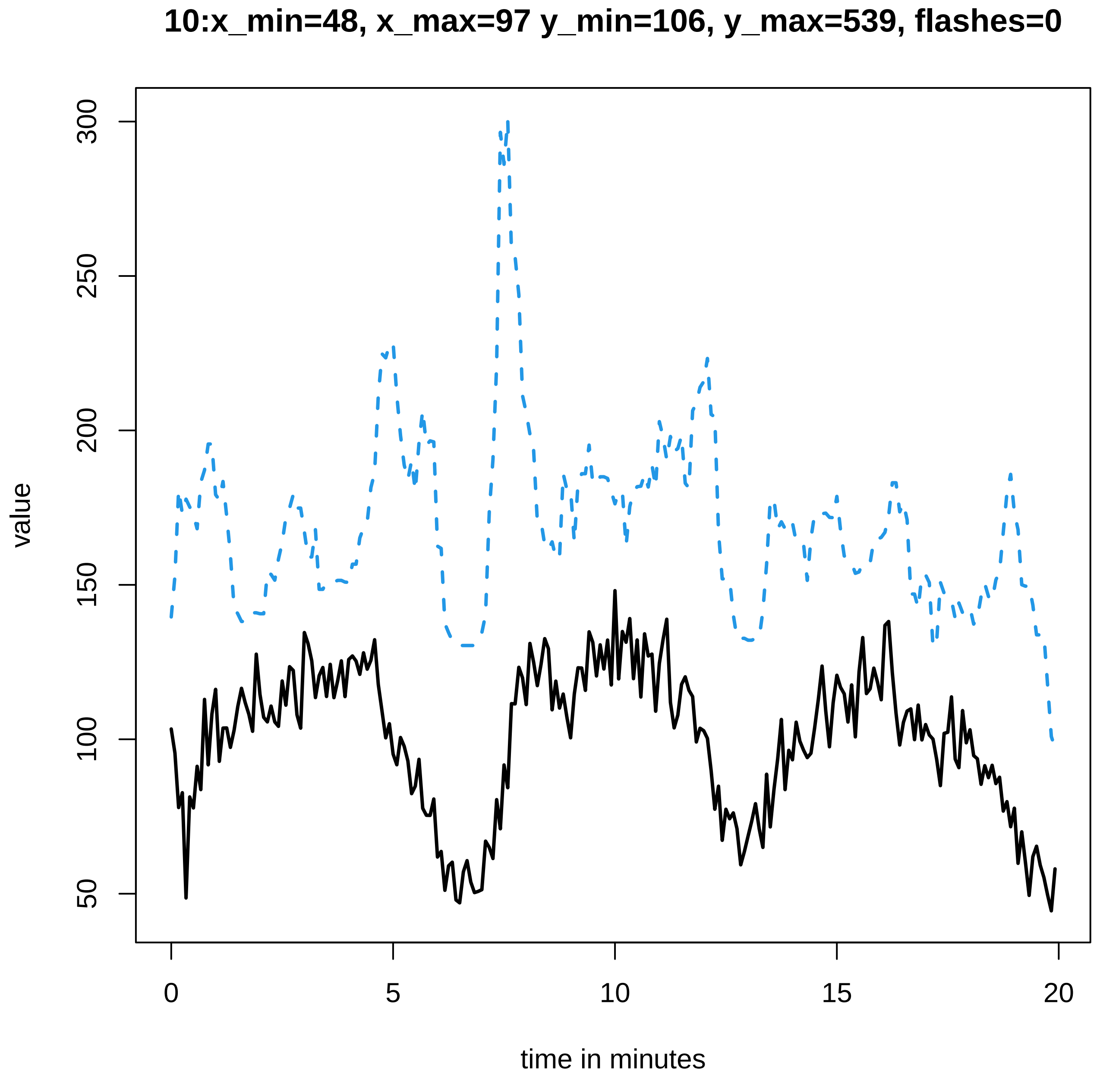
10:x_min=48, x_max=97 y_min=106, y_max=539, flashes=0
value
300
250
200
150
100
50
0
5
10
15
20
time in minutes

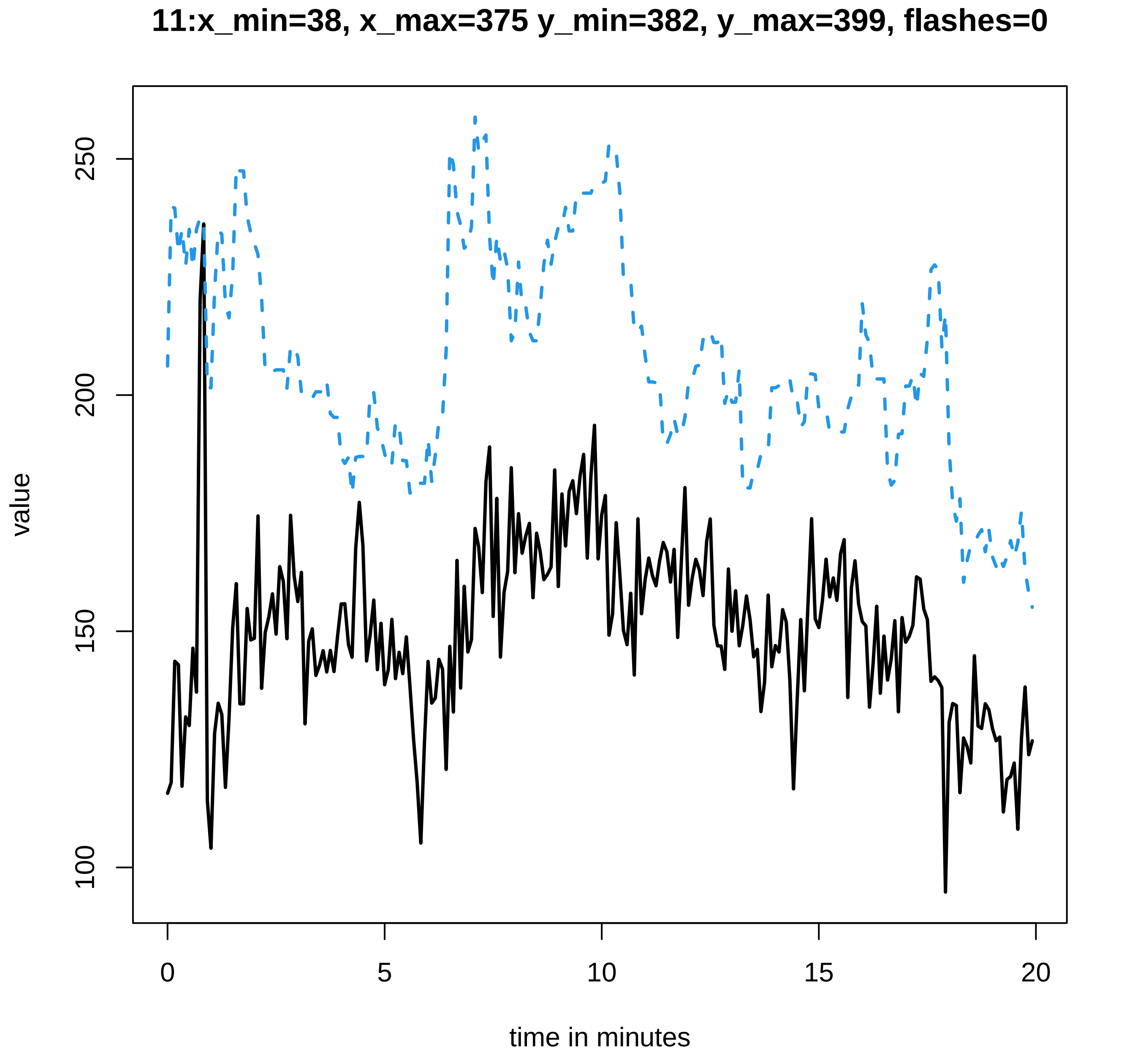

11:x_min=38, x_max=375 y_min=382, y_max=399, flashes=0
value
250
200
150
100
0
5
10
15
20
time in minutes

**12:x_min=22, x_max=296 y_min=302, y_max=467, flashes=0**

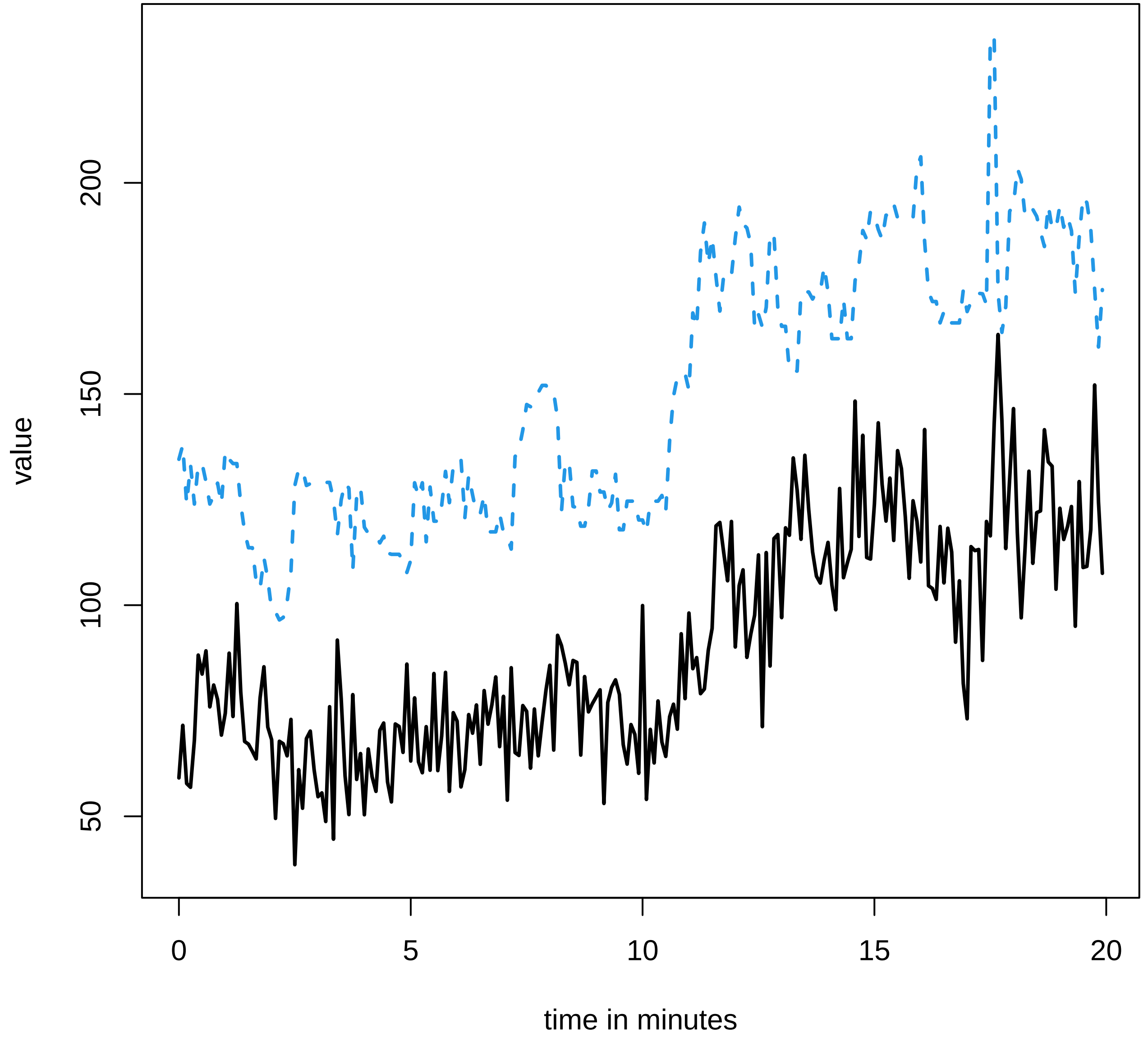

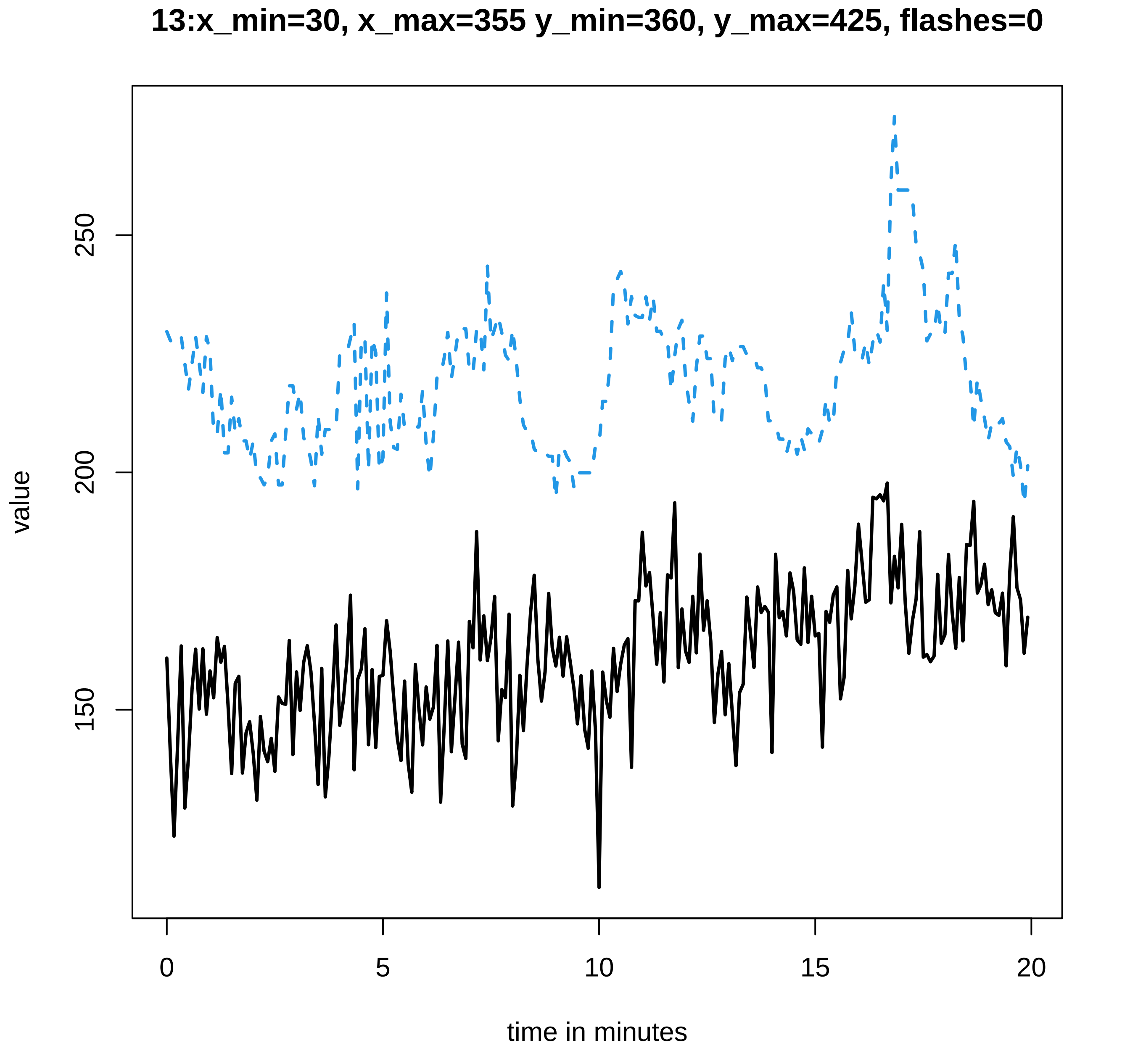

13:x_min=30, x_max=355 y_min=360, y_max=425, flashes=0
value
150
200
250
time in minutes
0
5
10
15
20

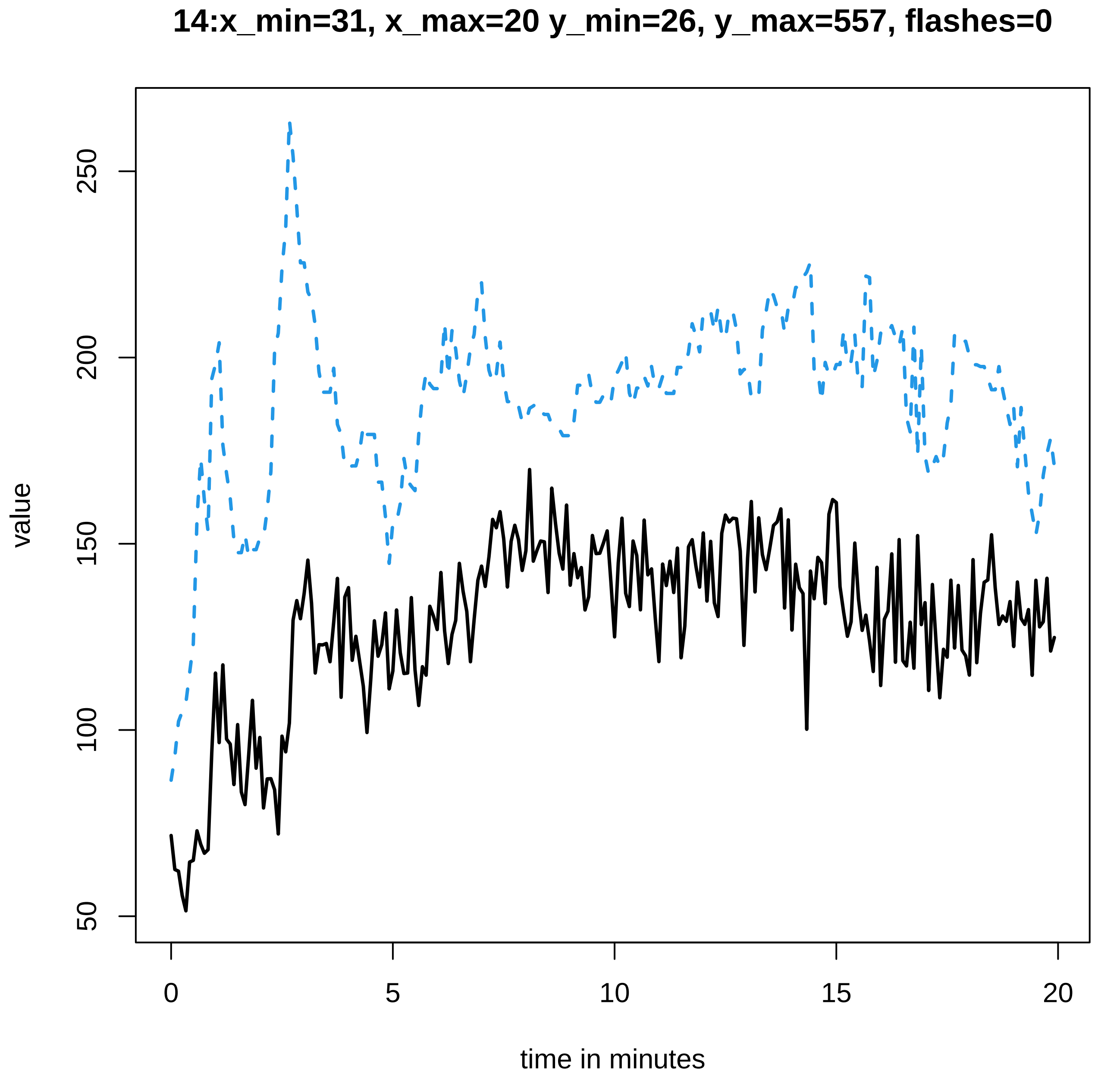
14:x_min=31, x_max=20 y_min=26, y_max=557, flashes=0
value
time in minutes
50
100
150
200
250
0
5
10
15
20

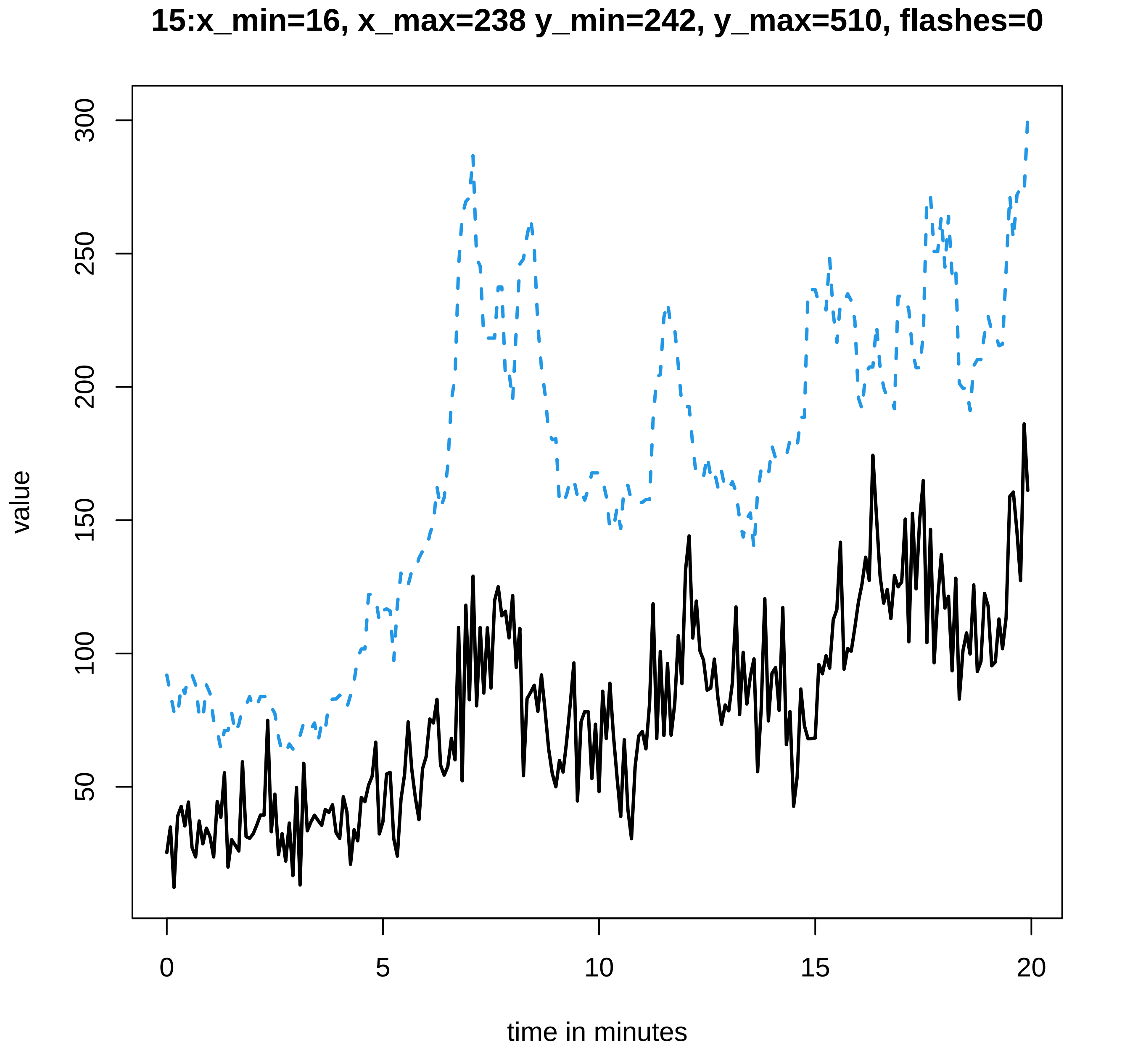
15:x_min=16, x_max=238 y_min=242, y_max=510, flashes=0
value
300
250
200
150
100
50
0
5
10
15
20
time in minutes

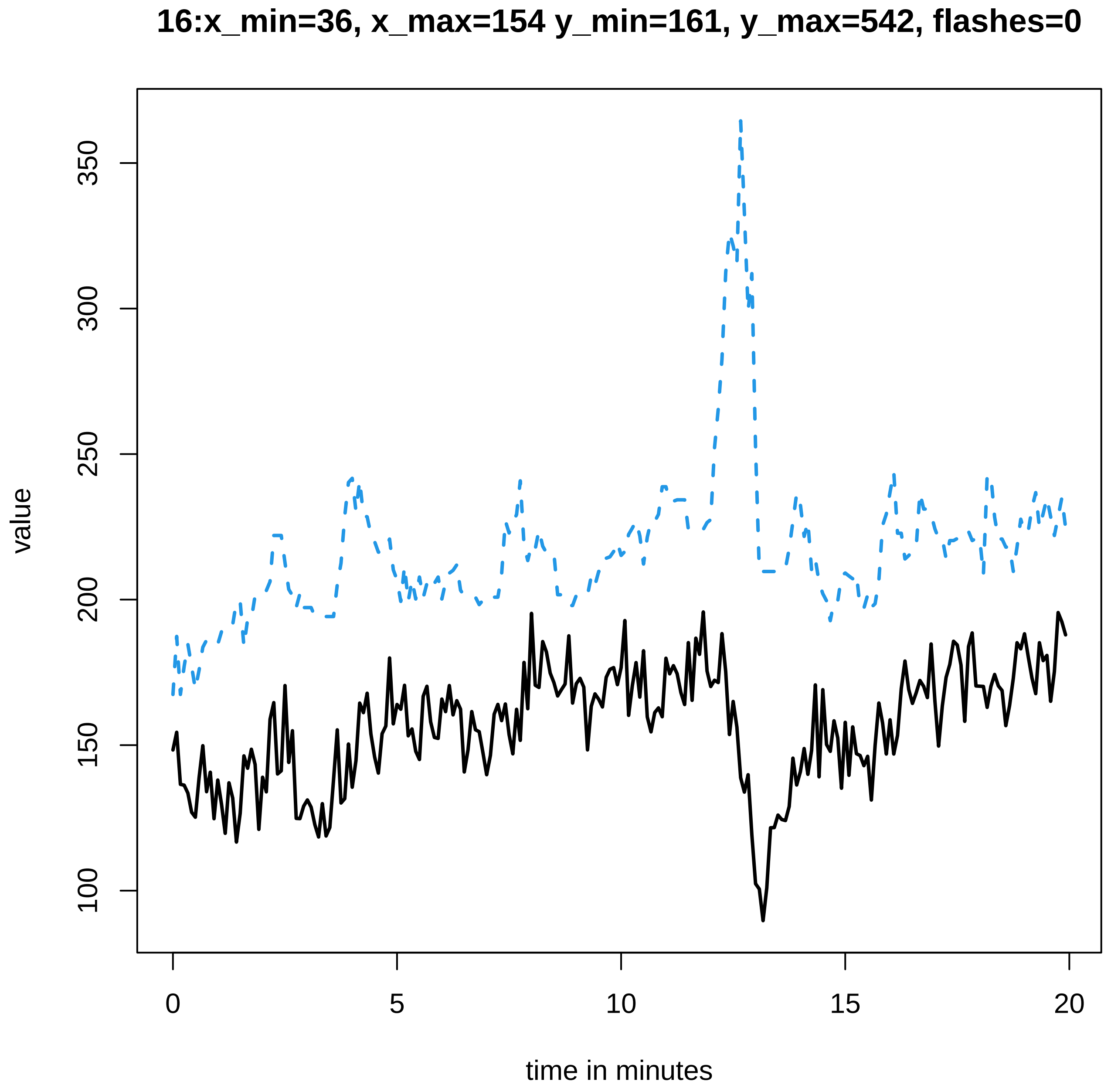
16:x_min=36, x_max=154 y_min=161, y_max=542, flashes=0
value
100
150
200
250
300
350
0
5
10
15
20
time in minutes

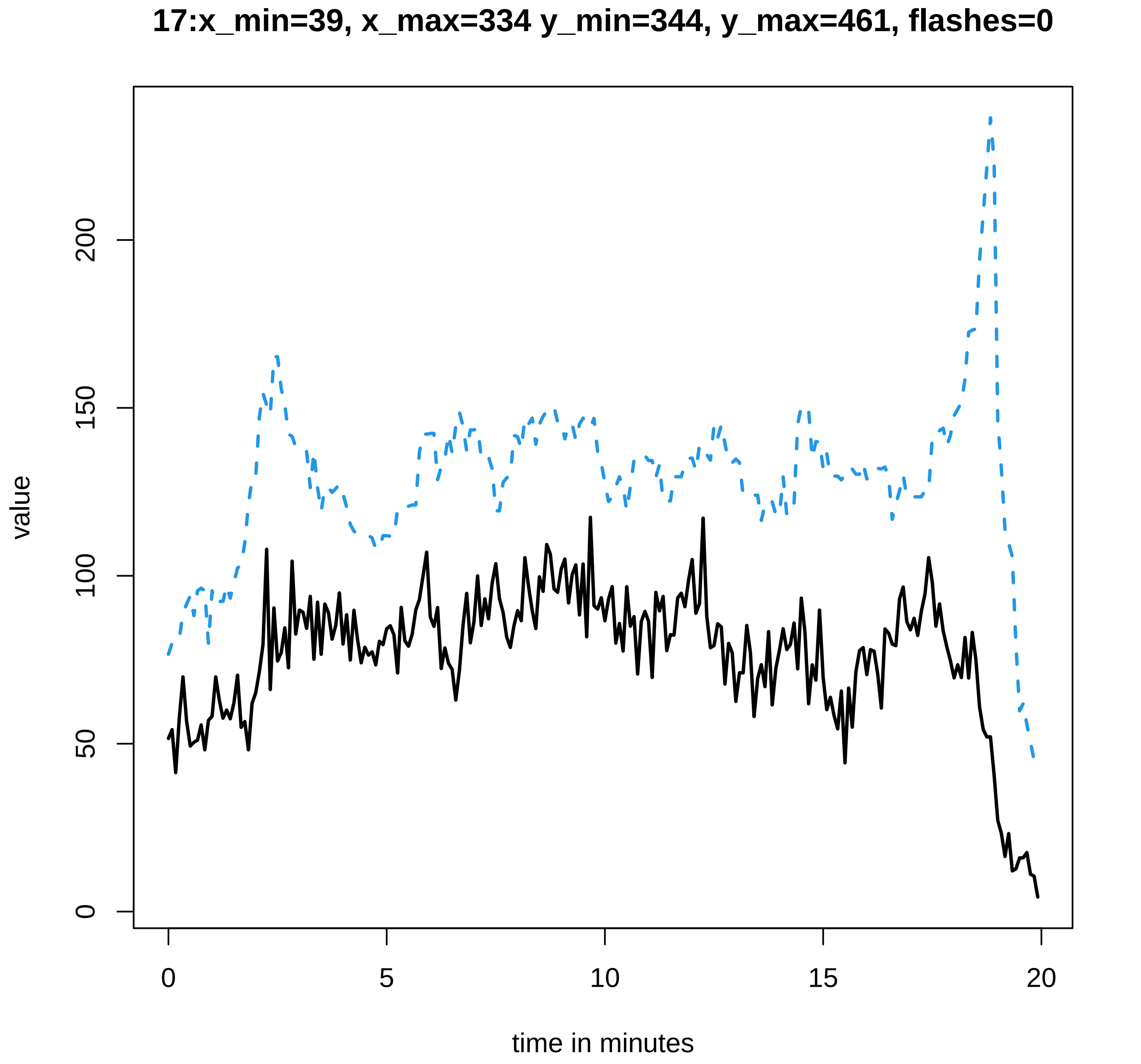

17:x_min=39, x_max=334 y_min=344, y_max=461, flashes=0
value
0
50
100
150
200
0
5
10
15
20
time in minutes

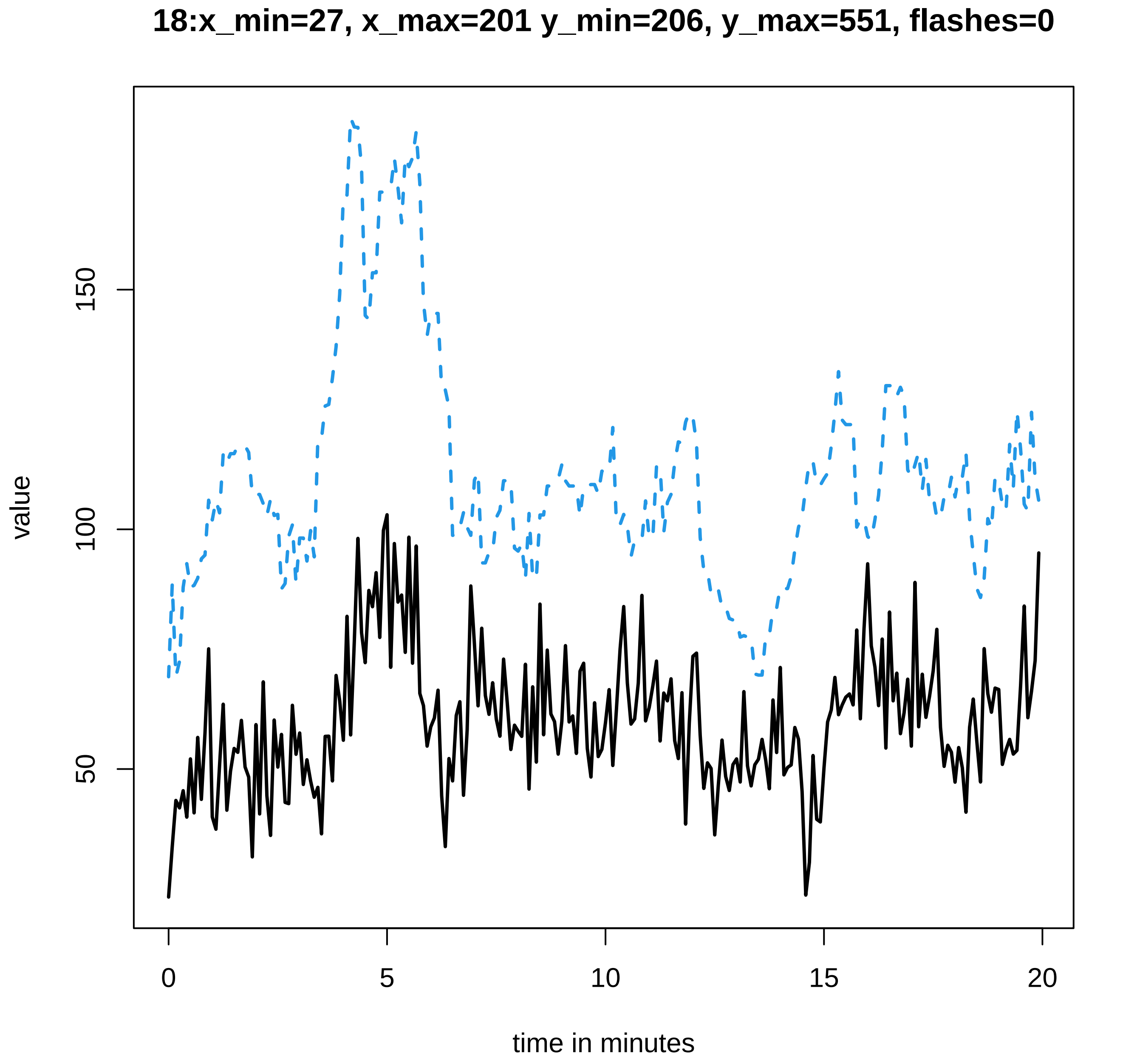
18:x_min=27, x_max=201 y_min=206, y_max=551, flashes=0
value
50
100
150
0
5
10
15
20
time in minutes

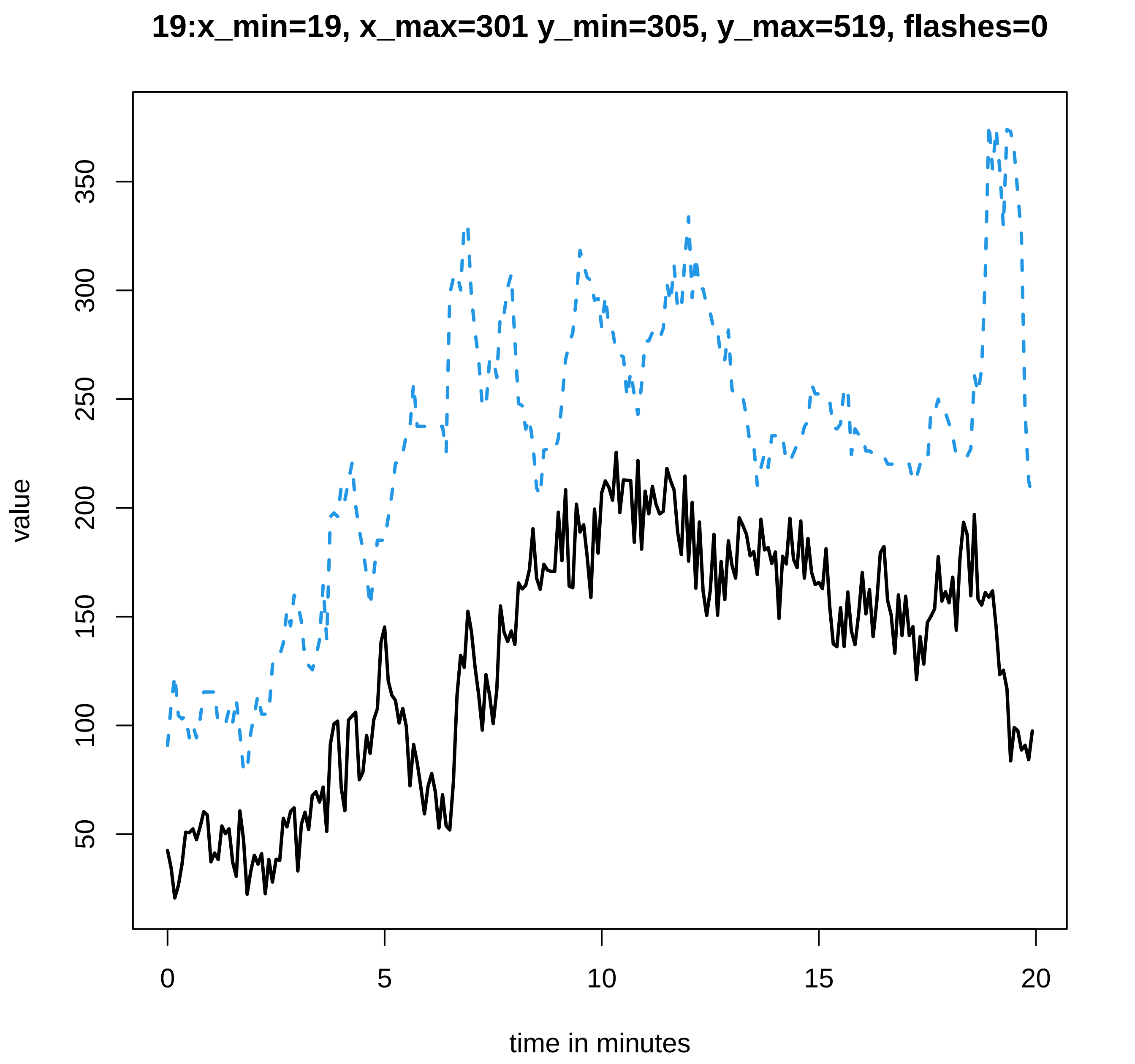

19:x_min=19, x_max=301 y_min=305, y_max=519, flashes=0
value
50
100
150
200
250
300
350
0
5
10
15
20
time in minutes

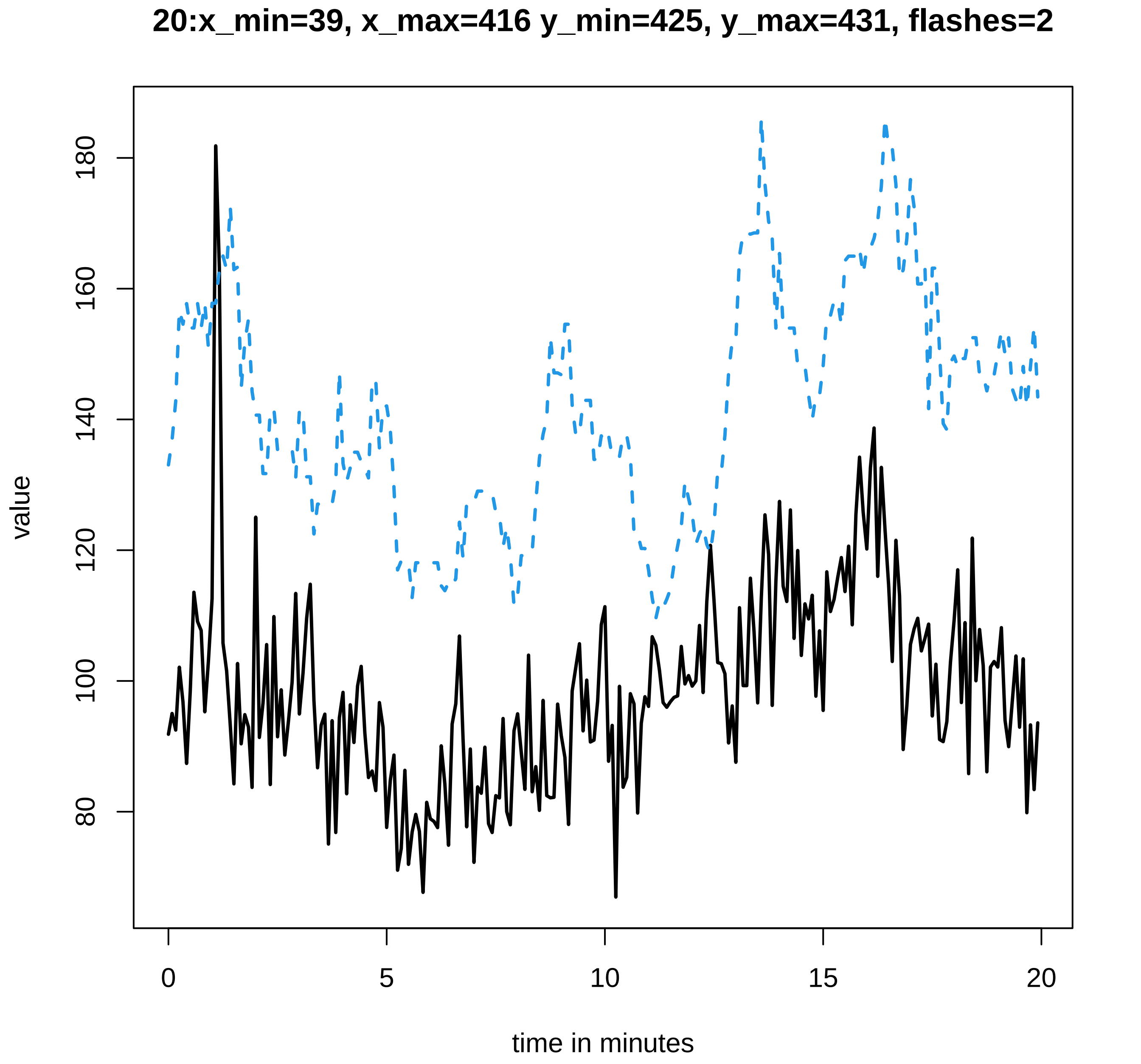

20:x_min=39, x_max=416 y_min=425, y_max=431, flashes=2
value
80
100
120
140
160
180
0
5
10
15
20
time in minutes

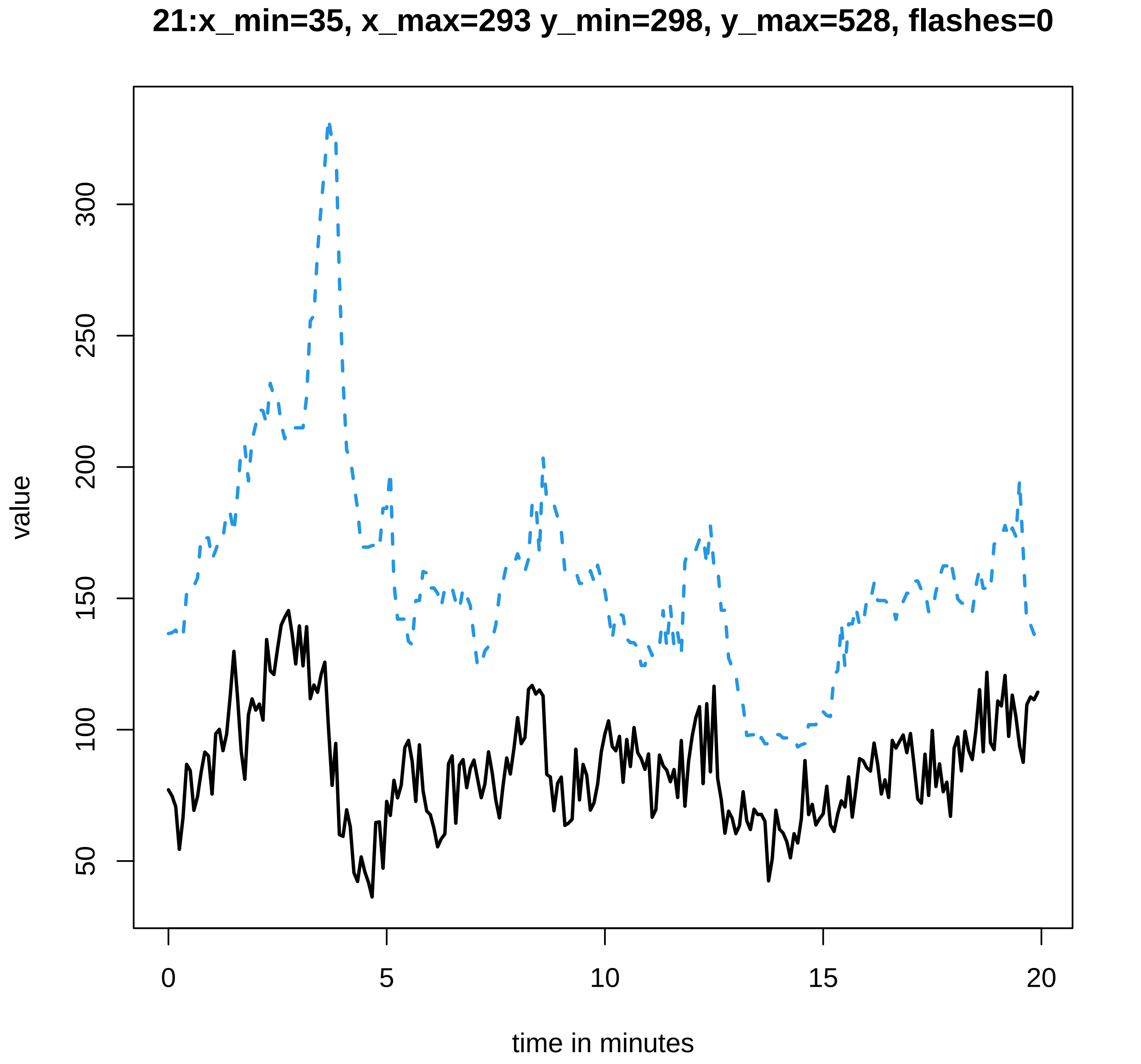
21:x_min=35, x_max=293 y_min=298, y_max=528, flashes=0
value
50
100
150
200
250
300
0
5
10
15
20
time in minutes

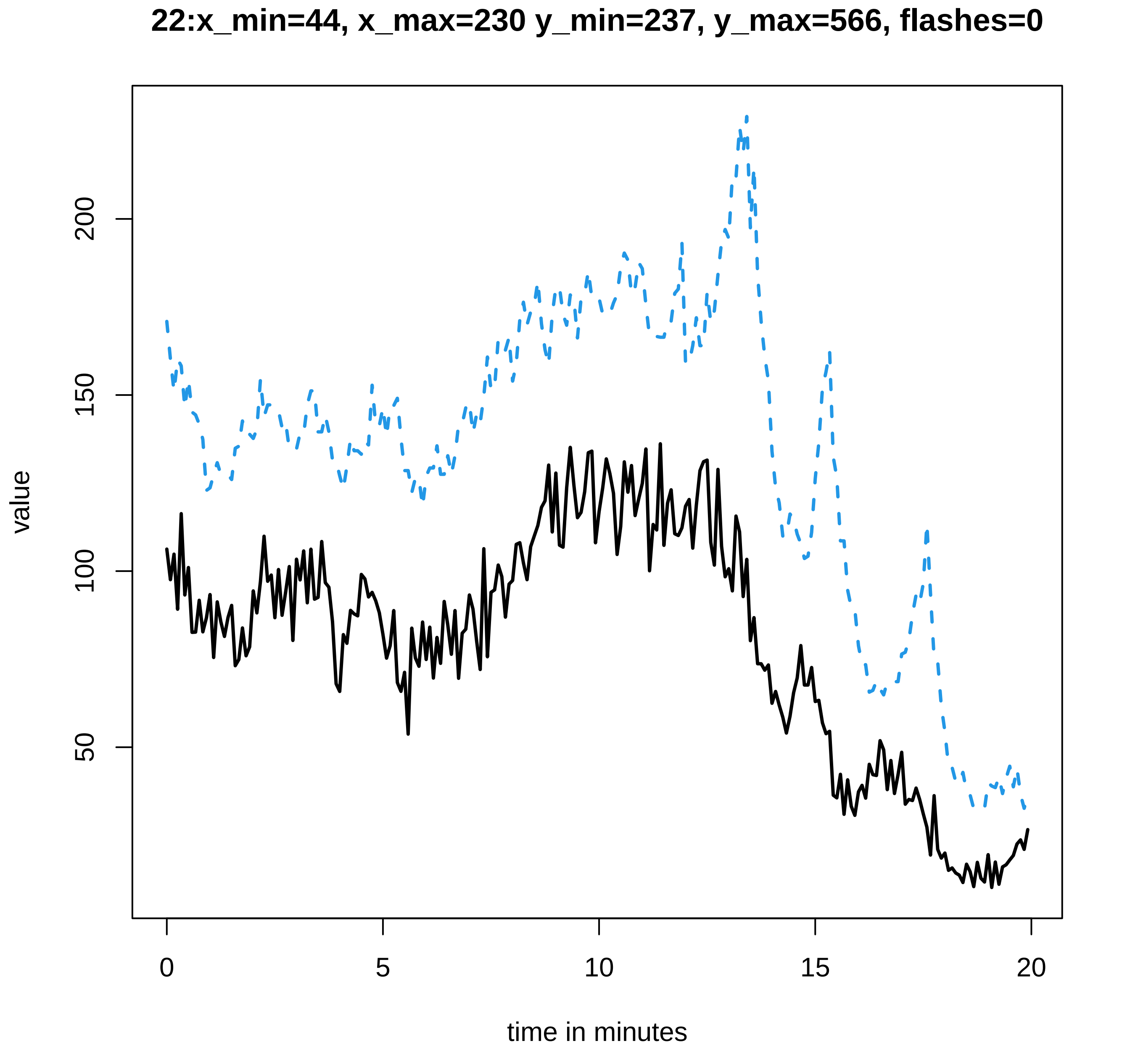

22:x_min=44, x_max=230 y_min=237, y_max=566, flashes=0
value
200
150
100
50
0
5
10
15
20
time in minutes

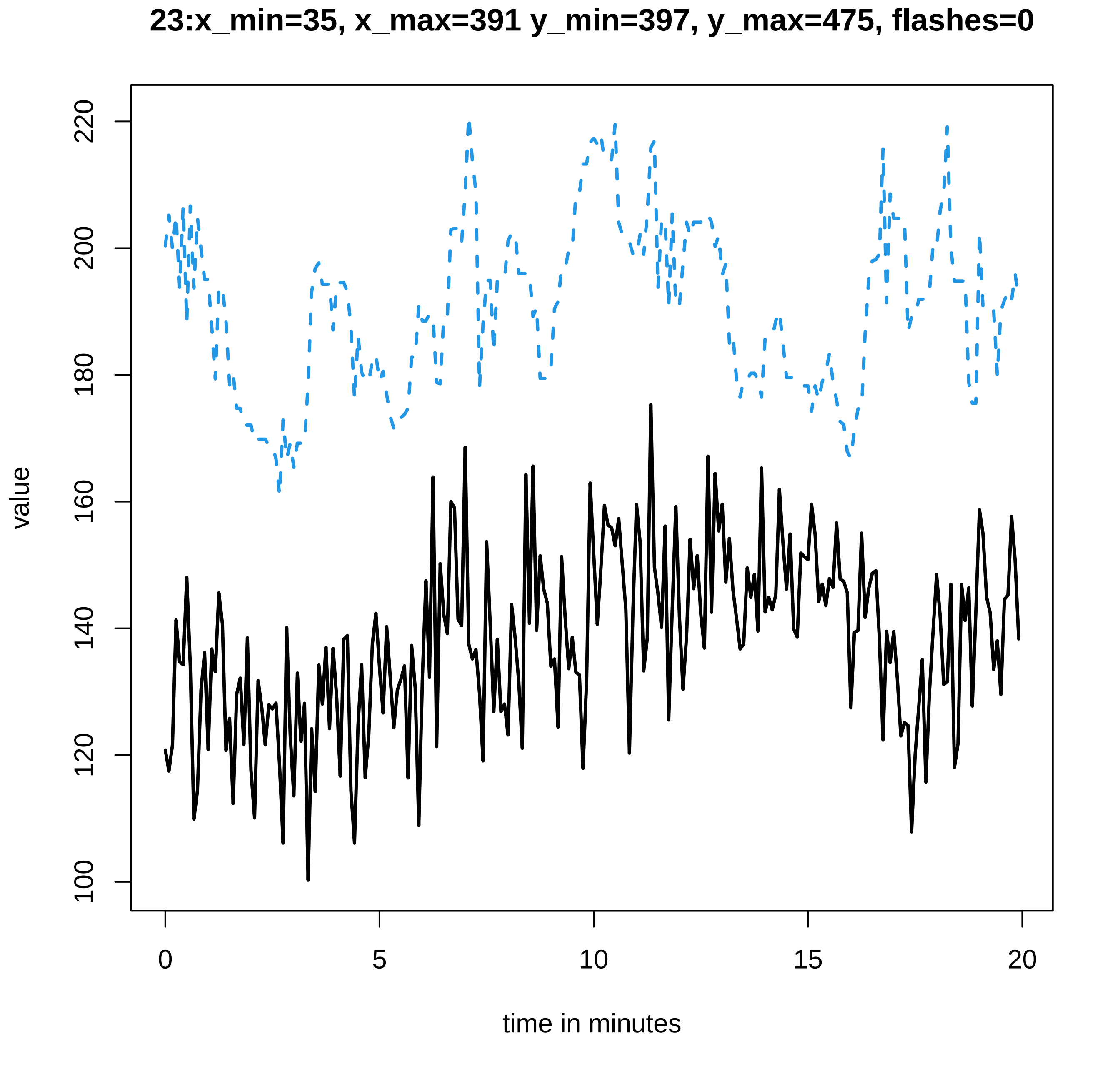
23:x_min=35, x_max=391 y_min=397, y_max=475, flashes=0
value
100
120
140
160
180
200
220
0
5
10
15
20
time in minutes

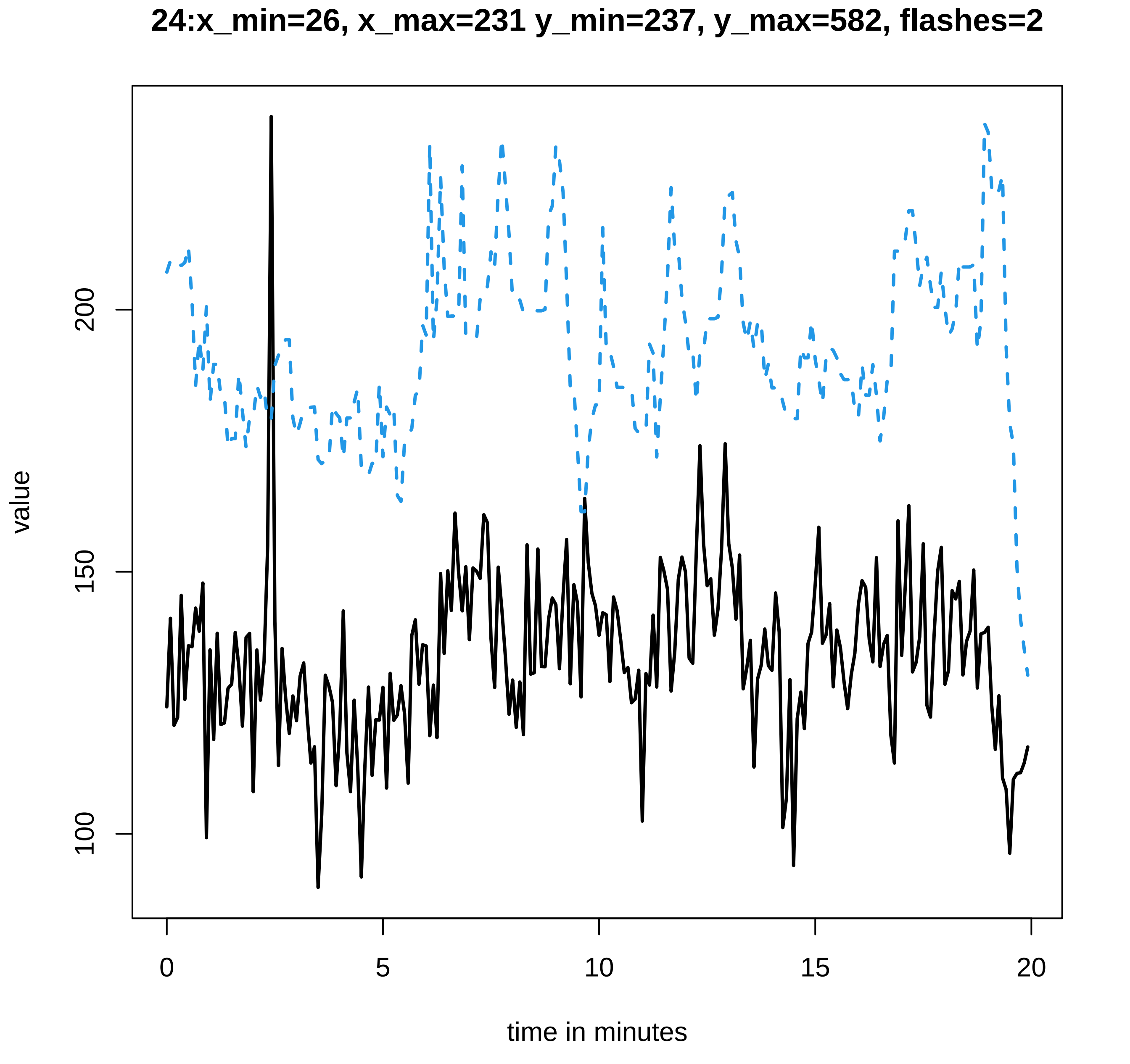

24:x_min=26, x_max=231 y_min=237, y_max=582, flashes=2
value
100
150
200
0
5
10
15
20
time in minutes

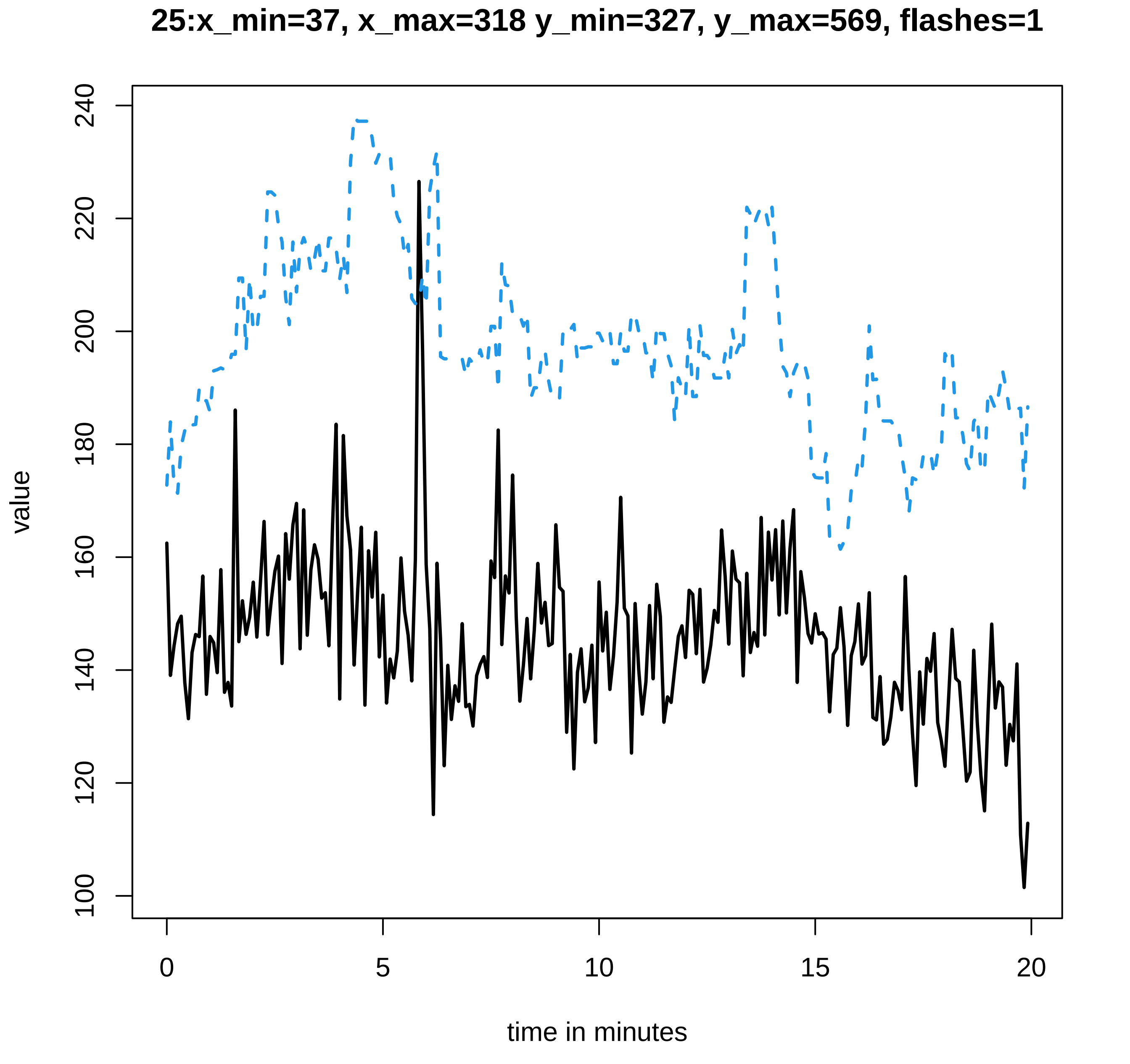

25:x_min=37, x_max=318 y_min=327, y_max=569, flashes=1
value
time in minutes
0
5
10
15
20
100
120
140
160
180
200
220
240

### 26:x_min=40, x_max=294 y_min=304, y_max=594, flashes=0

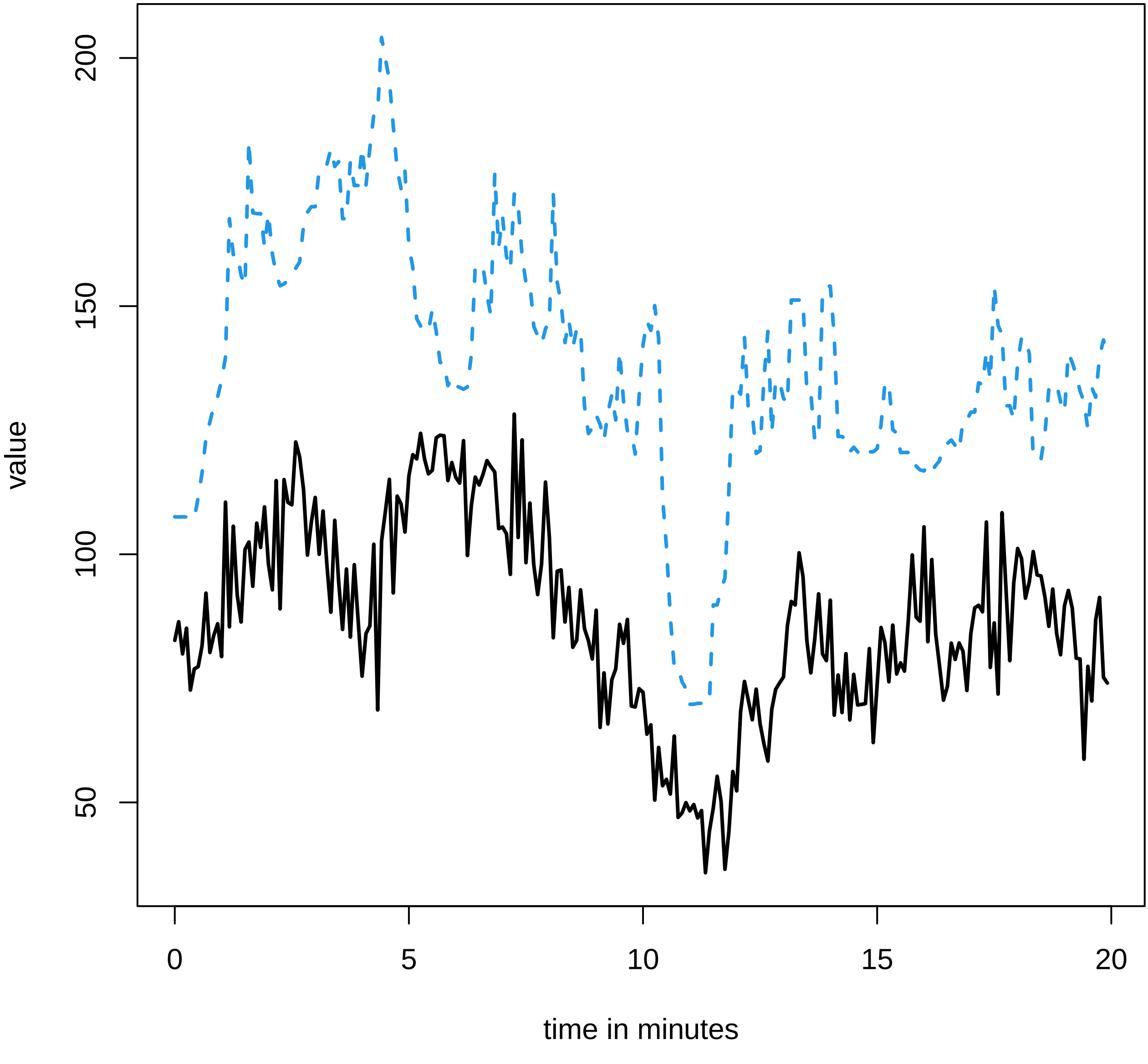

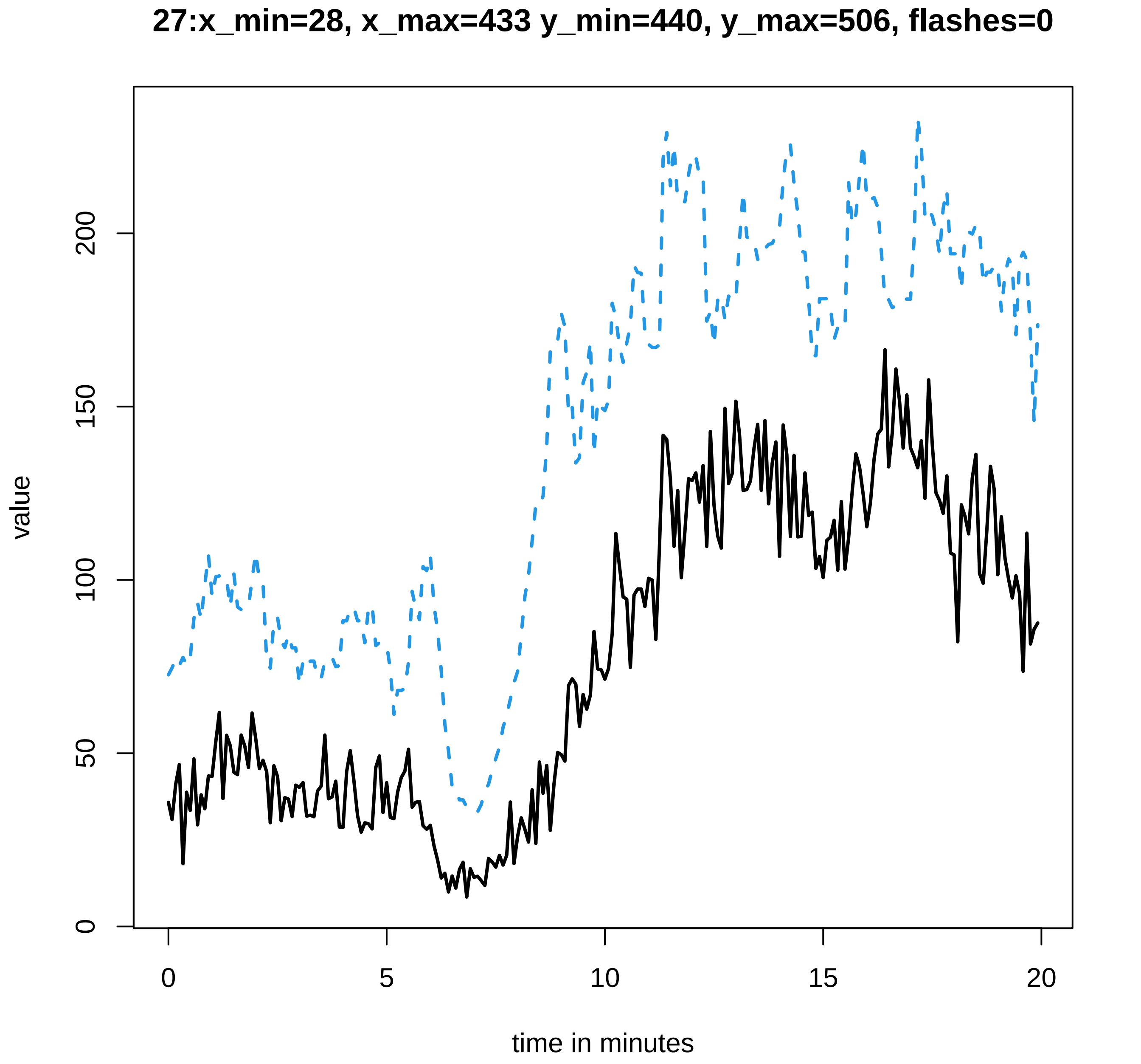

27:x_min=28, x_max=433 y_min=440, y_max=506, flashes=0
value
0
50
100
150
200
0
5
10
15
20
time in minutes

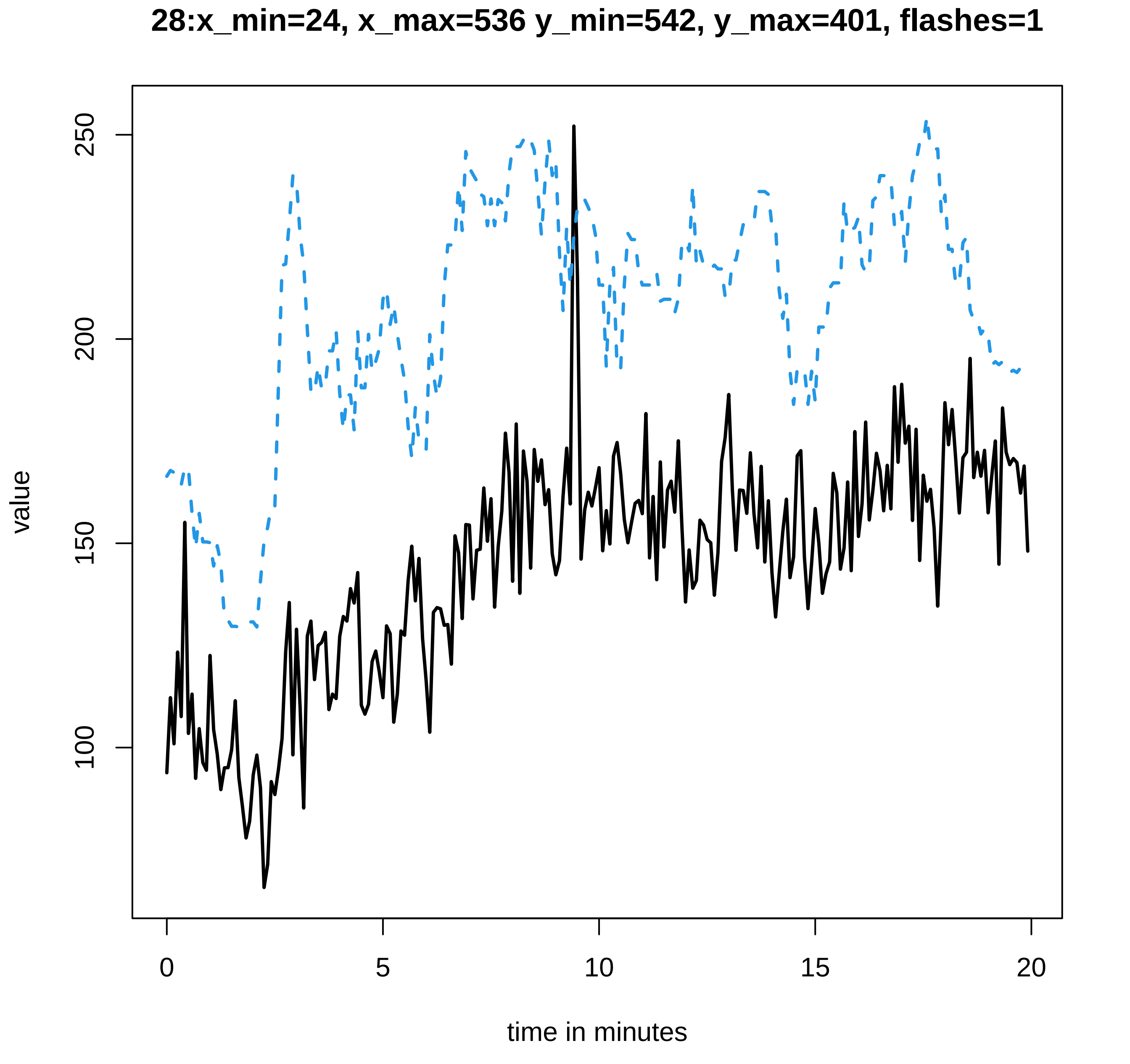
28:x_min=24, x_max=536 y_min=542, y_max=401, flashes=1
value
250
200
150
100
0
5
10
15
20
time in minutes

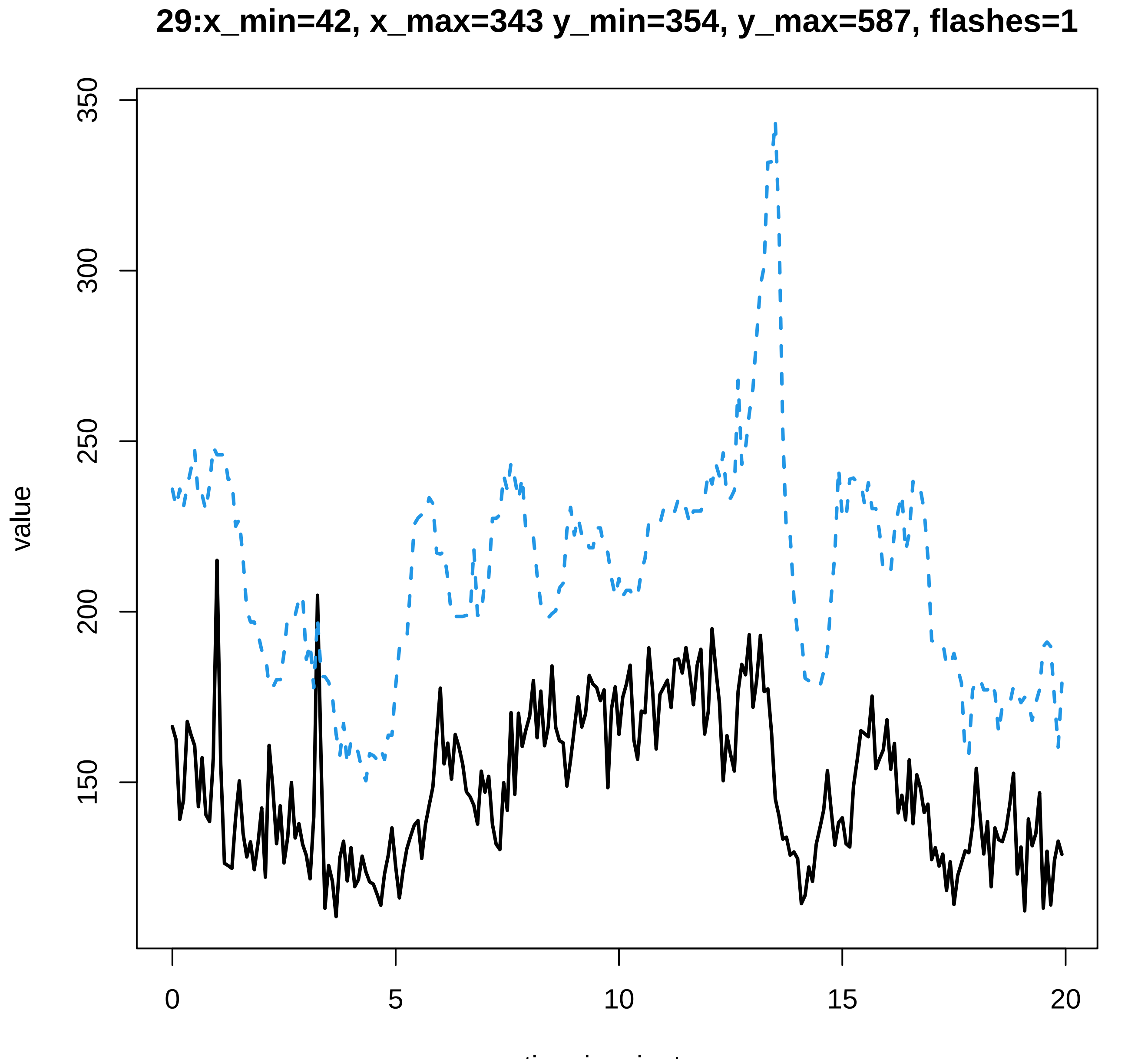
29:x_min=42, x_max=343 y_min=354, y_max=587, flashes=1
350
300
250
200
150
value
0
5
10
15
20
time in minutes

### 30:x_min=32, x_max=544 y_min=549, y_max=436, flashes=1

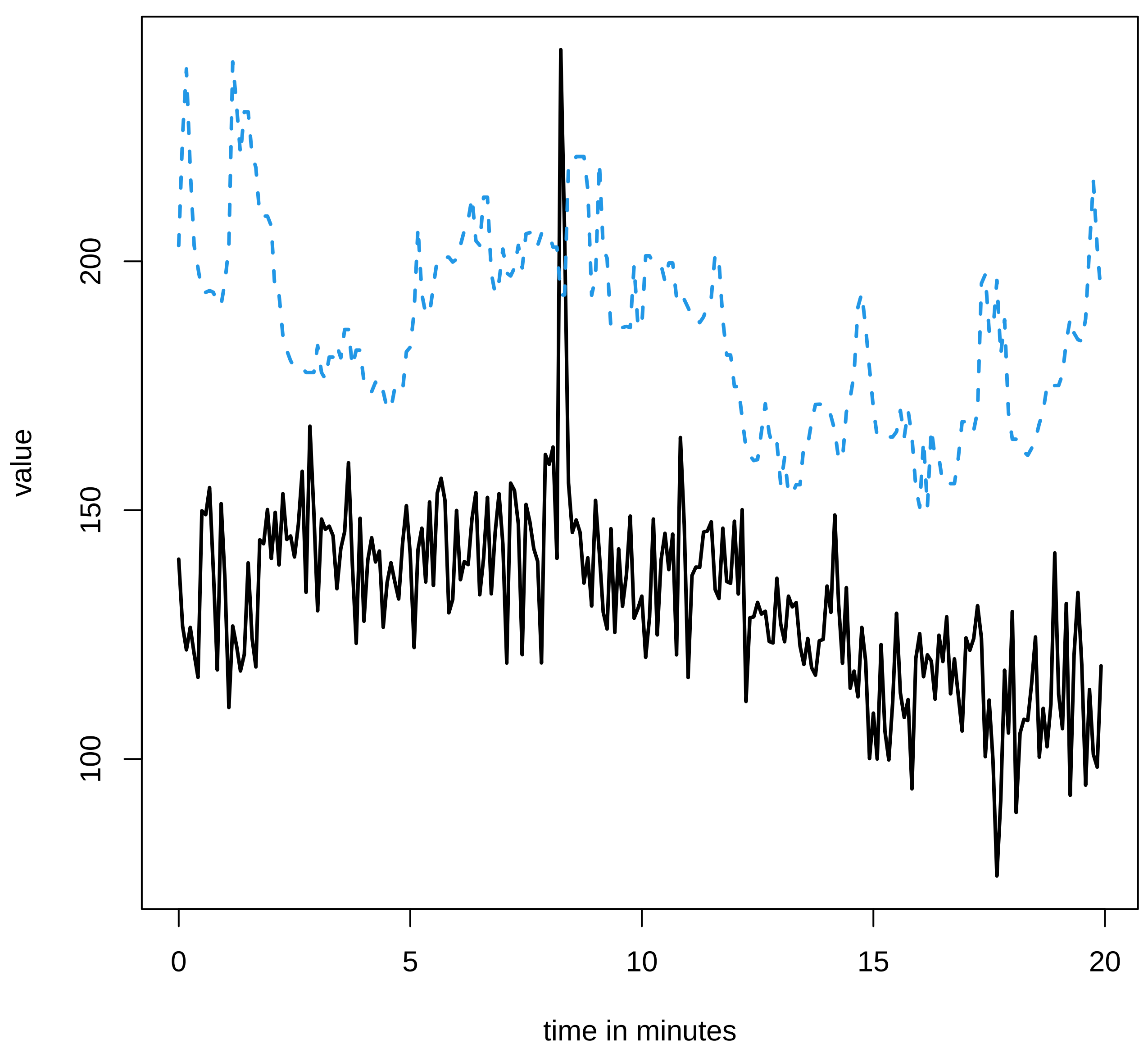

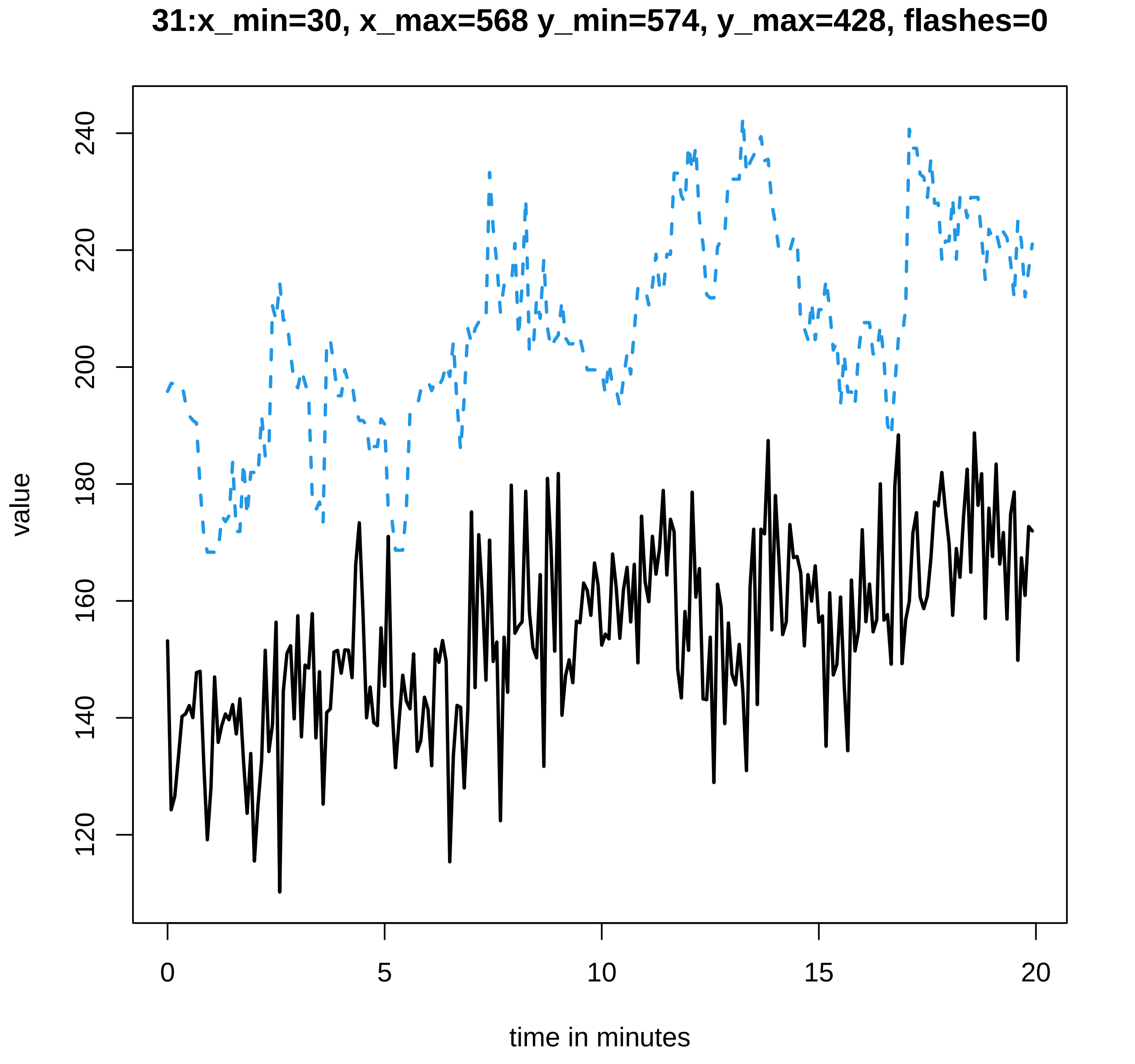
31:x_min=30, x_max=568 y_min=574, y_max=428, flashes=0
value
120
140
160
180
200
220
240
0
5
10
15
20
time in minutes

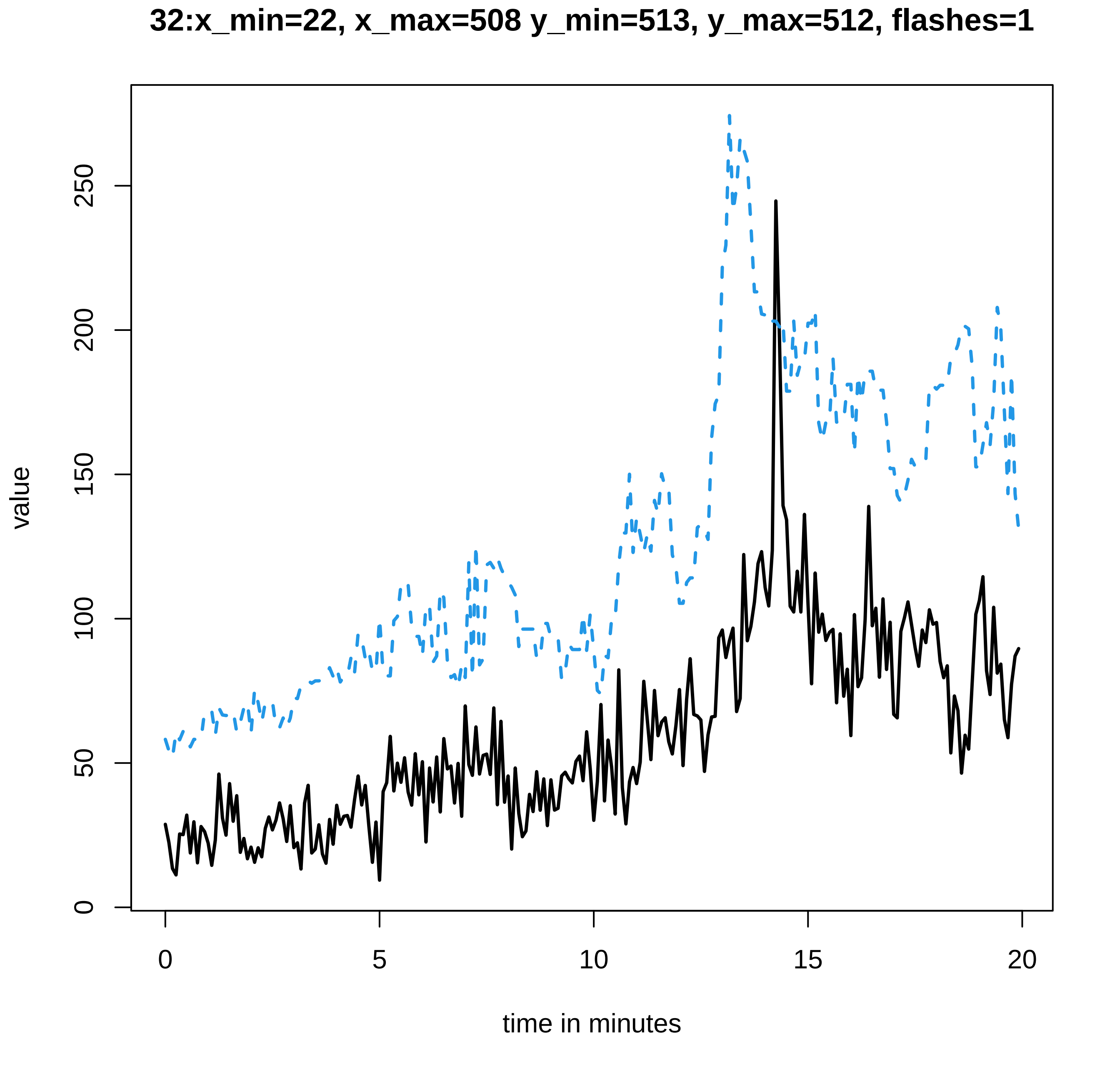
32:x_min=22, x_max=508 y_min=513, y_max=512, flashes=1
value
0
50
100
150
200
250
0
5
10
15
20
time in minutes

### 33:x_min=34, x_max=538 y_min=547, y_max=535, flashes=3

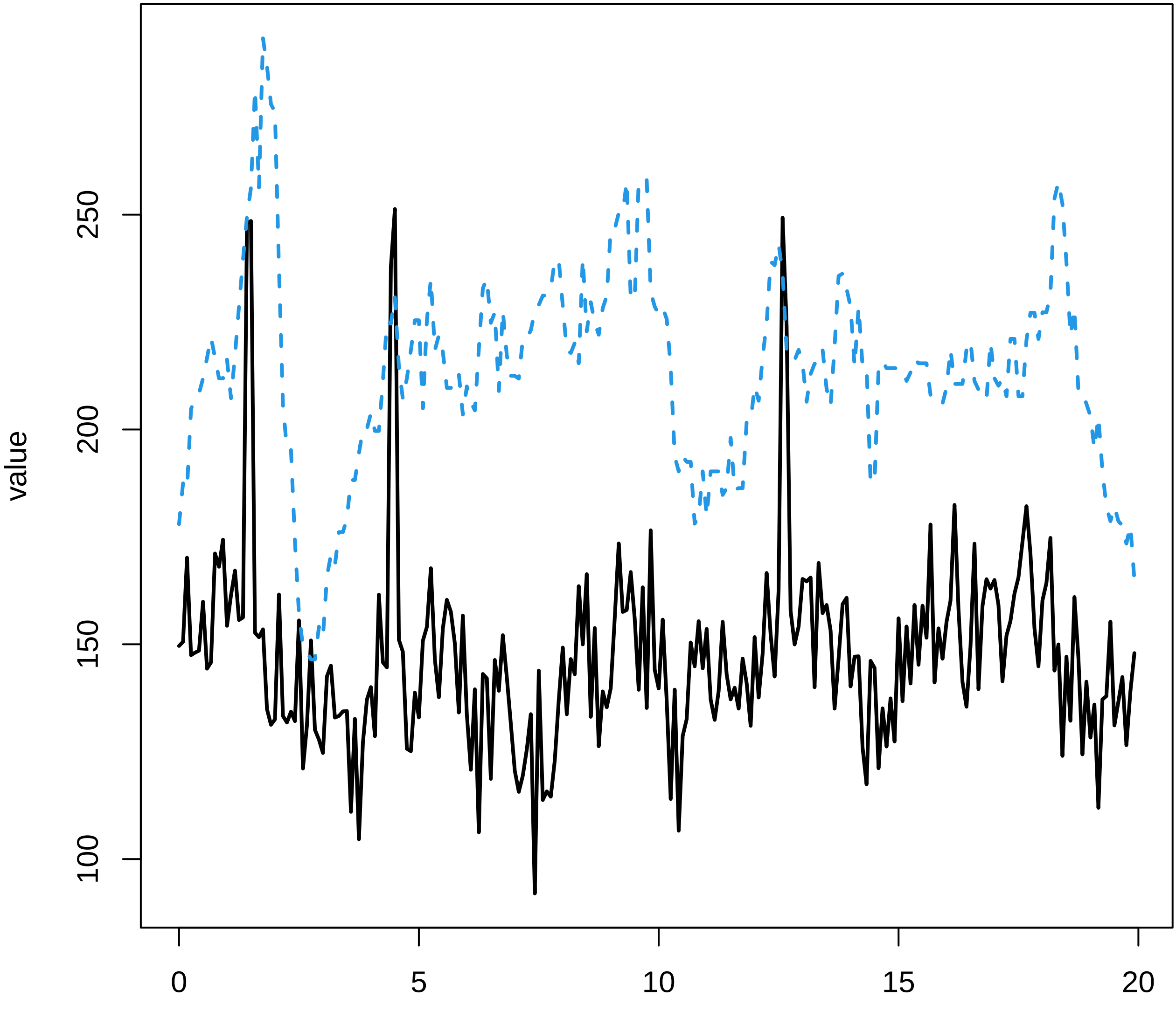

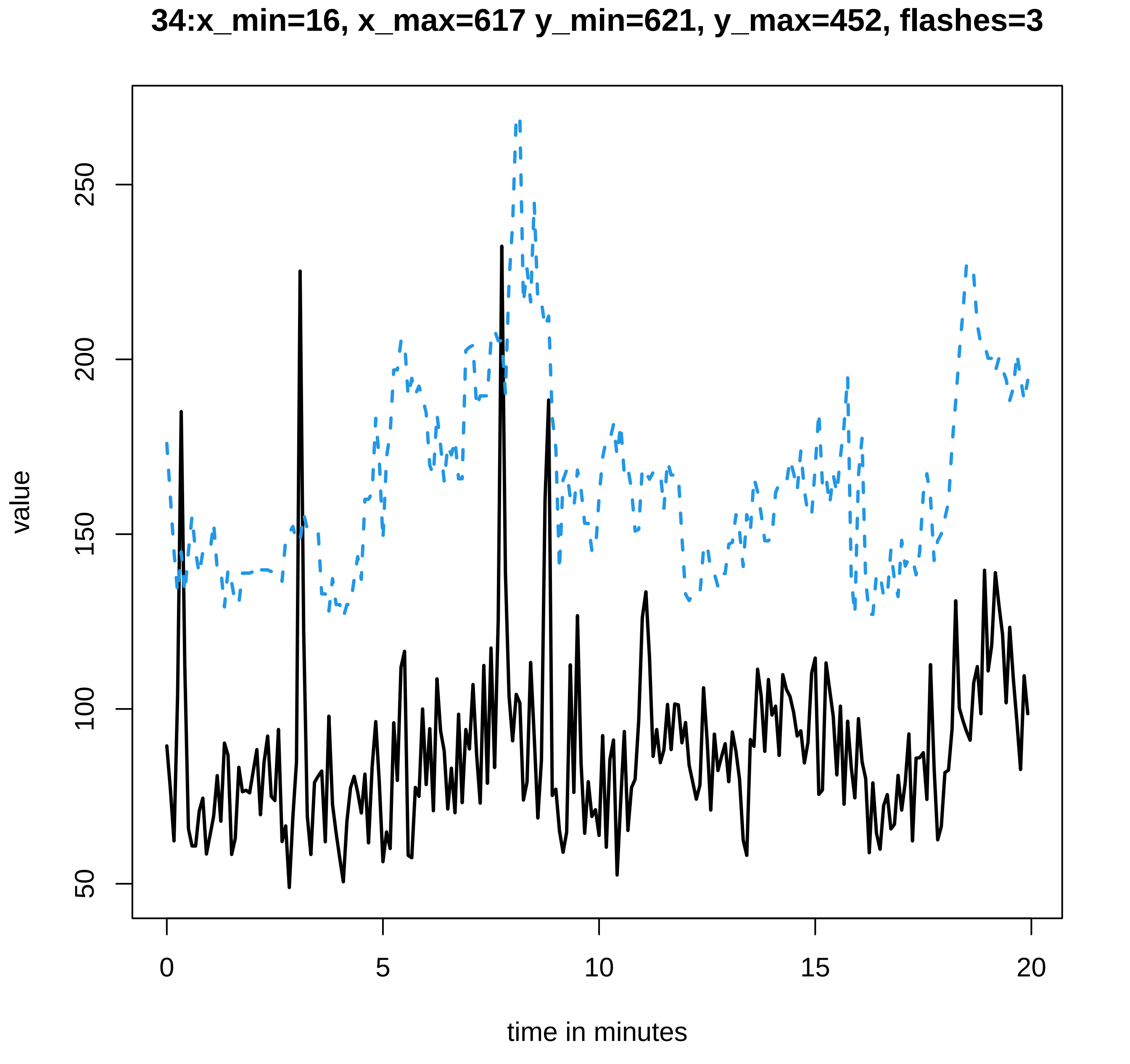
34:x_min=16, x_max=617 y_min=621, y_max=452, flashes=3
value
time in minutes
0
5
10
15
20
50
100
150
200
250

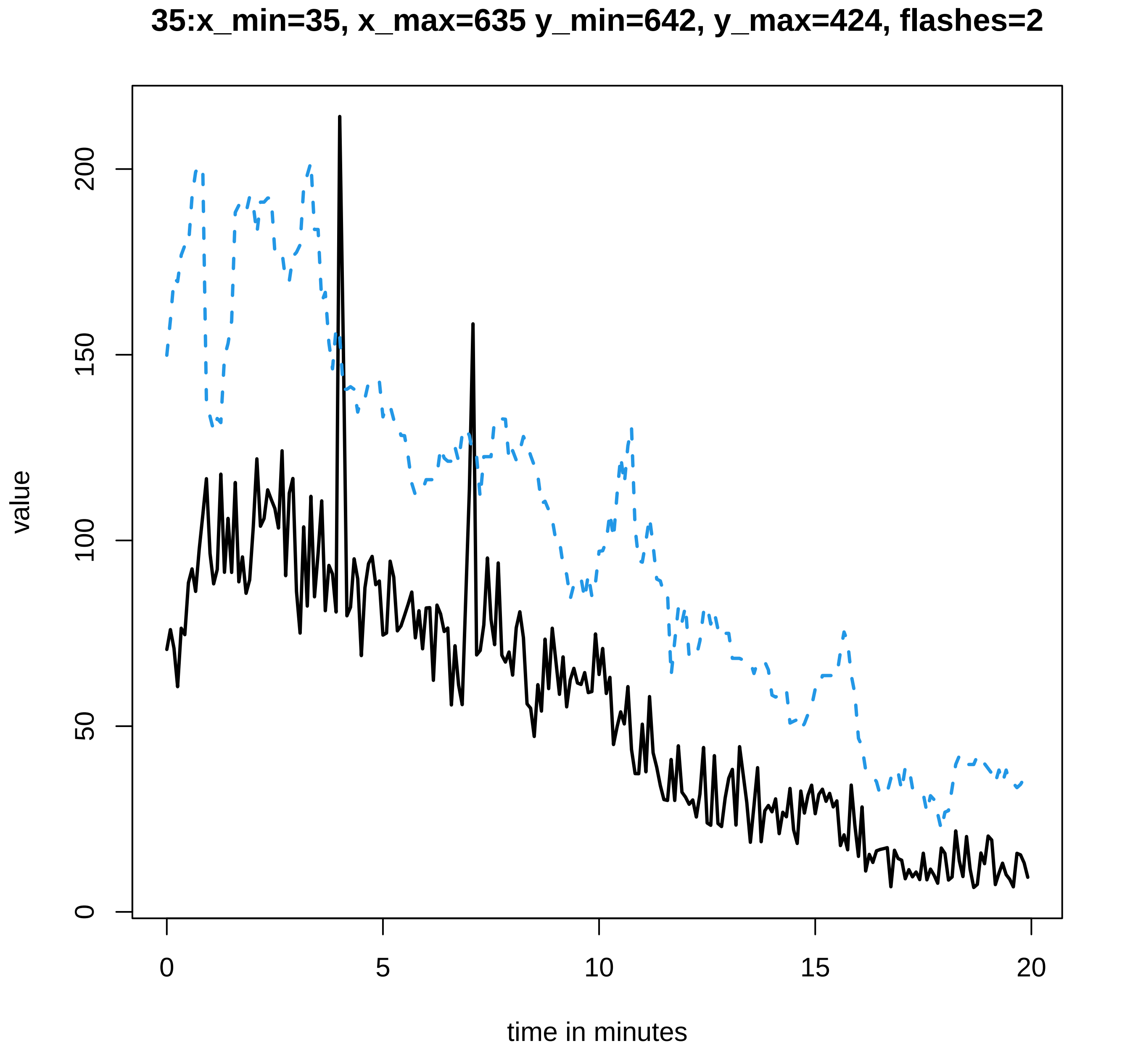
35:x_min=35, x_max=635 y_min=642, y_max=424, flashes=2
value
0
50
100
150
200
time in minutes
0
5
10
15
20

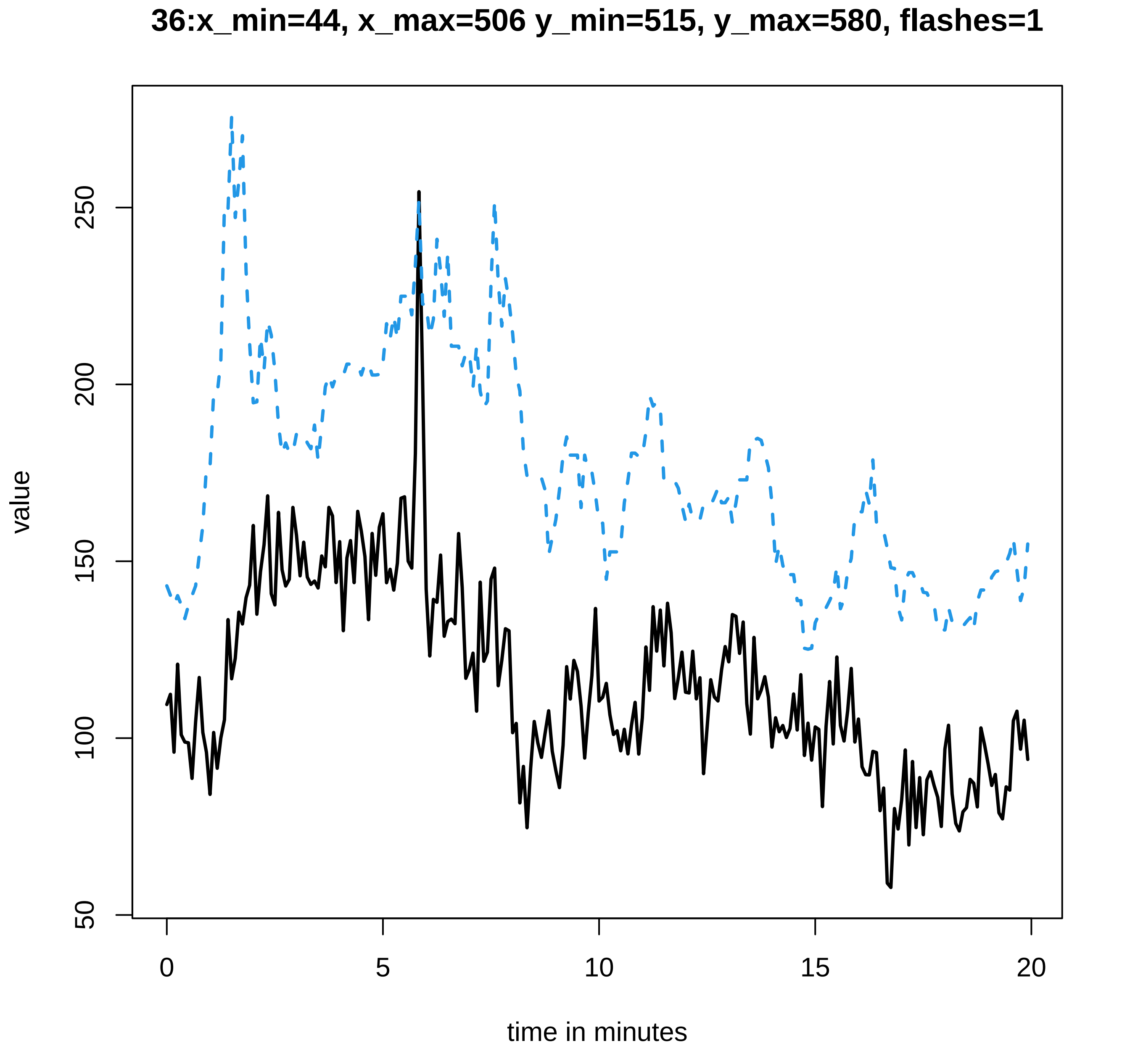

36:x_min=44, x_max=506 y_min=515, y_max=580, flashes=1
value
50
100
150
200
250
0
5
10
15
20
time in minutes

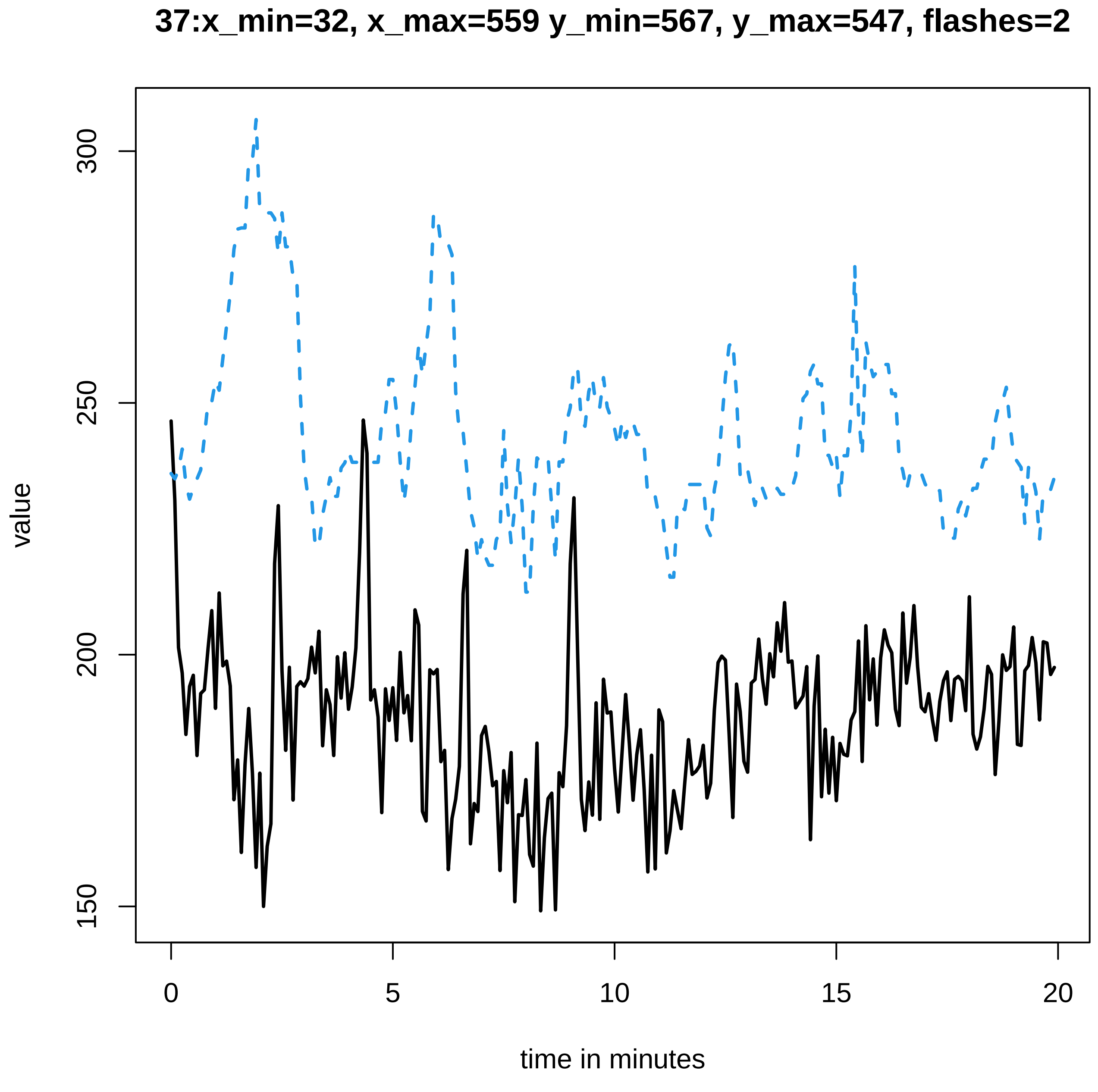
37:x_min=32, x_max=559 y_min=567, y_max=547, flashes=2
value
300
250
200
150
0
5
10
15
20
time in minutes

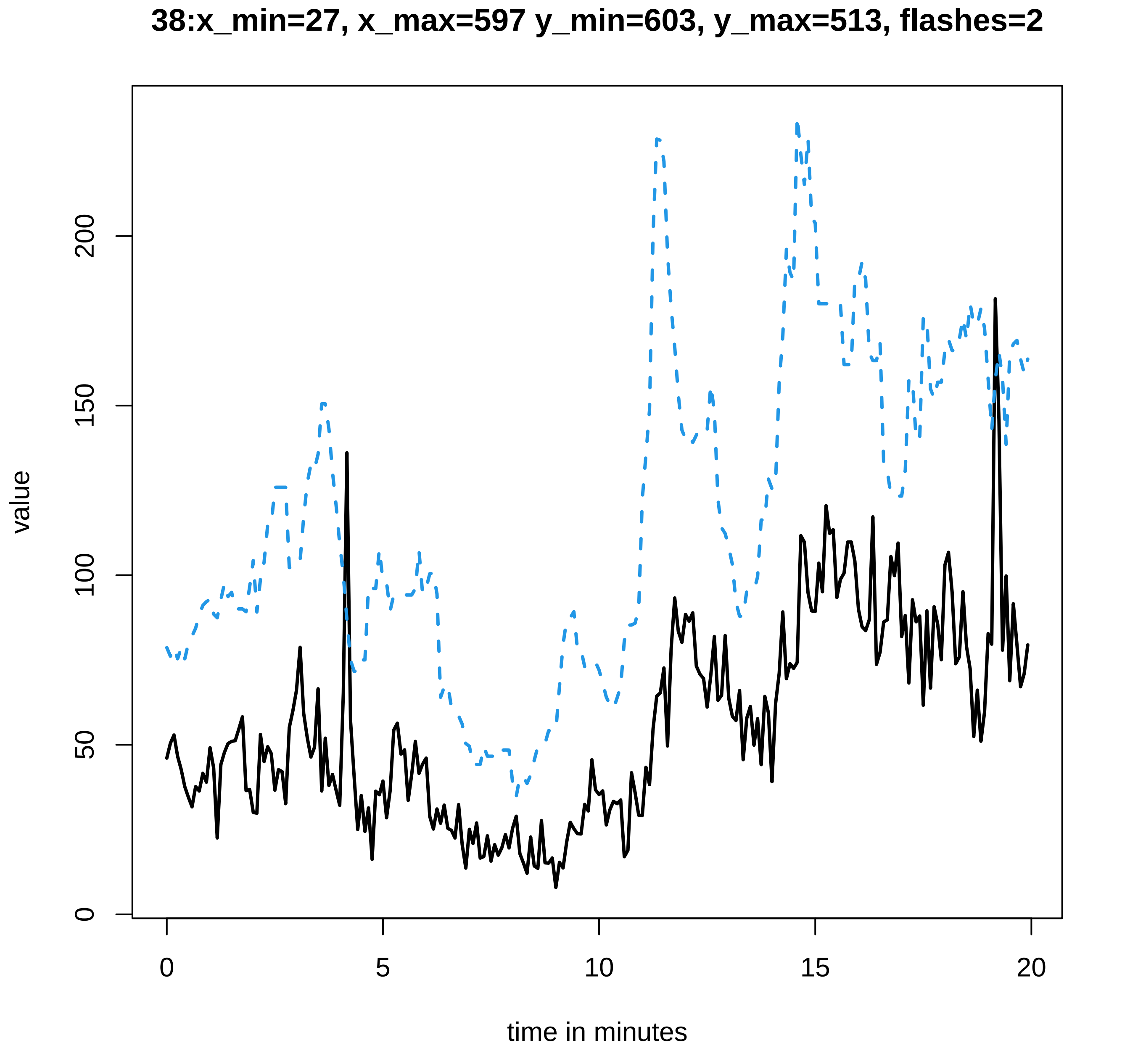
38:x_min=27, x_max=597 y_min=603, y_max=513, flashes=2
value
0
50
100
150
200
0
5
10
15
20
time in minutes

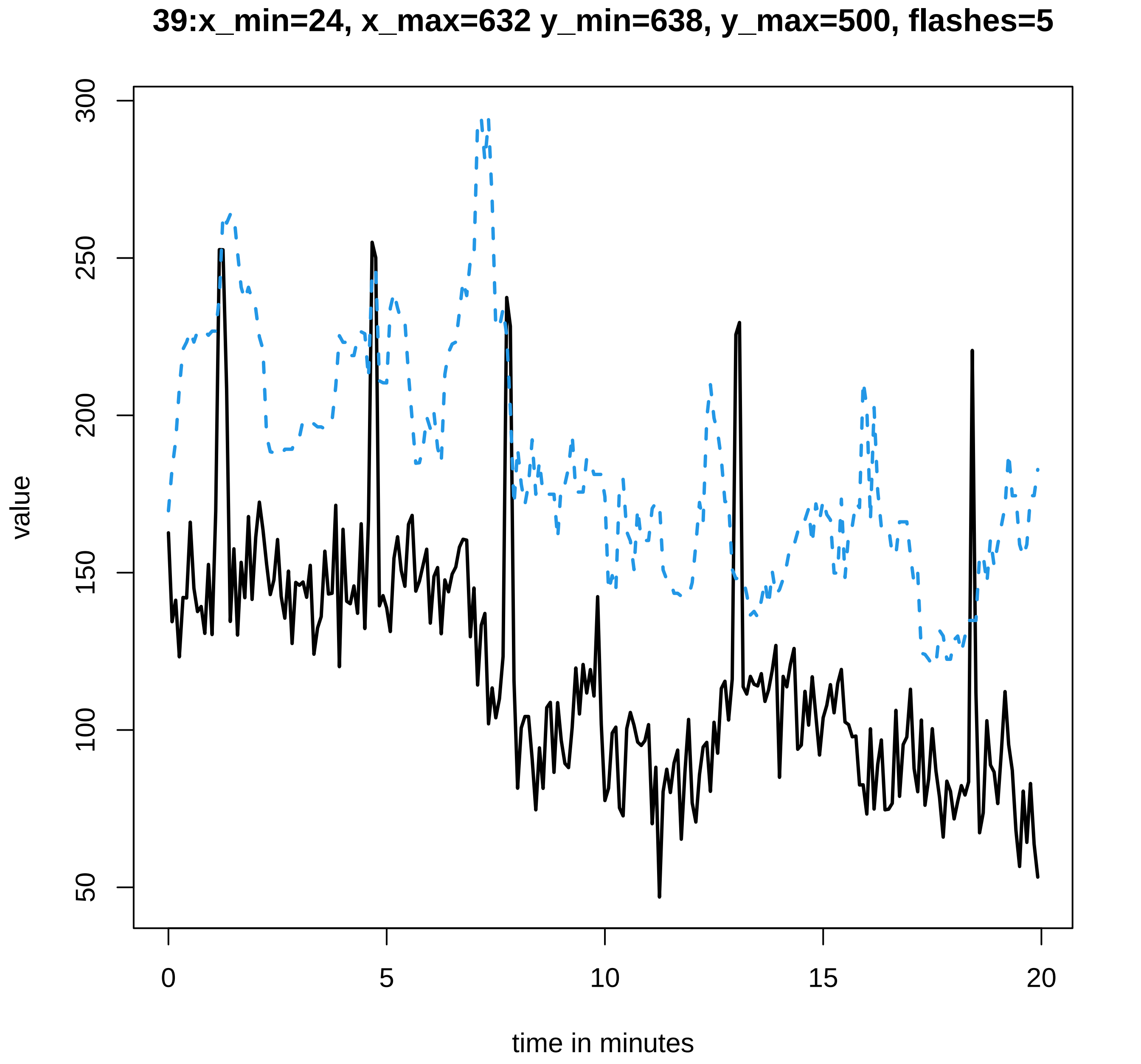

39:x_min=24, x_max=632 y_min=638, y_max=500, flashes=5
300
250
200
150
100
50
value
0
5
10
15
20
time in minutes

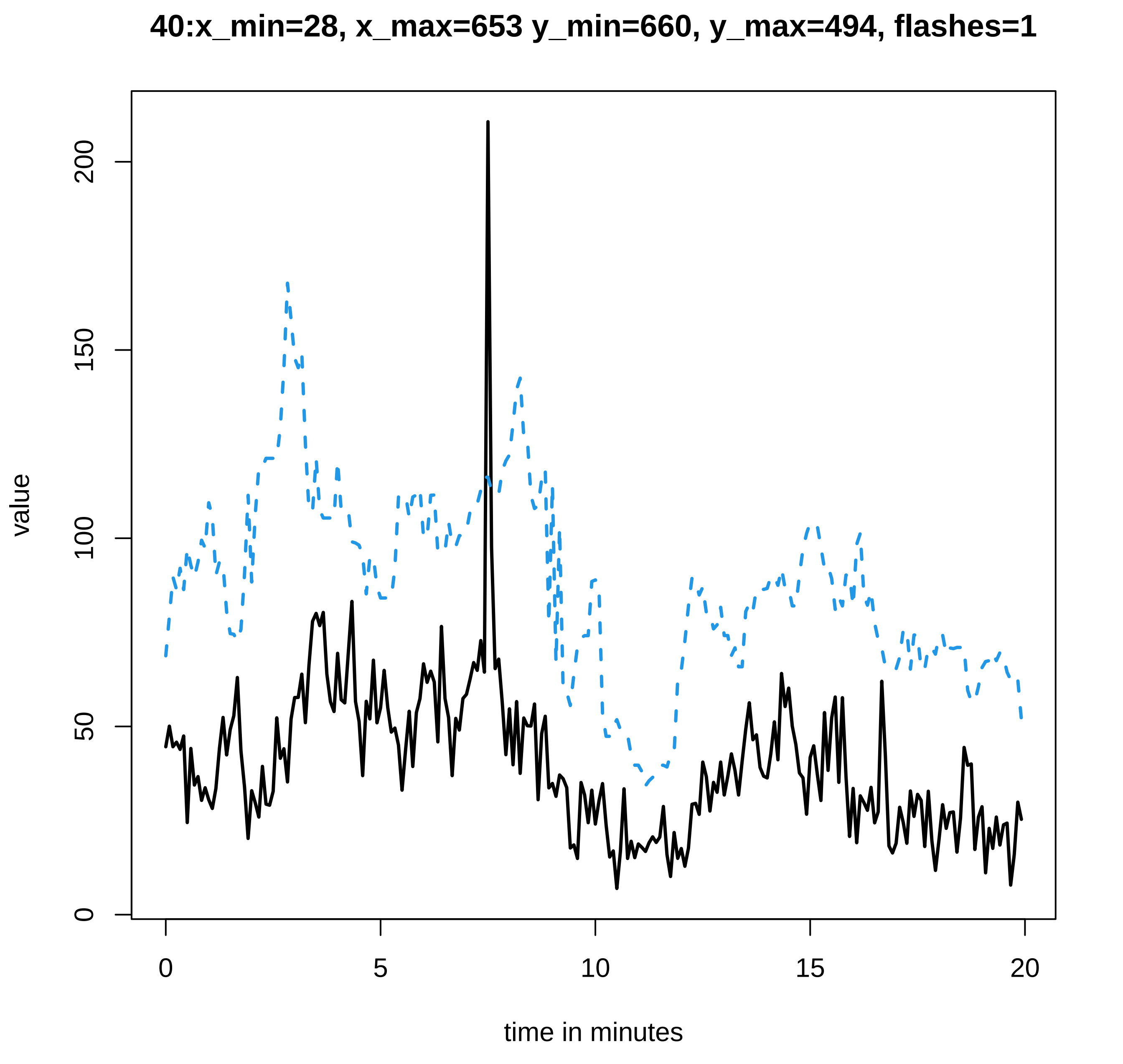
40:x_min=28, x_max=653 y_min=660, y_max=494, flashes=1
value
0
50
100
150
200
0
5
10
15
20
time in minutes

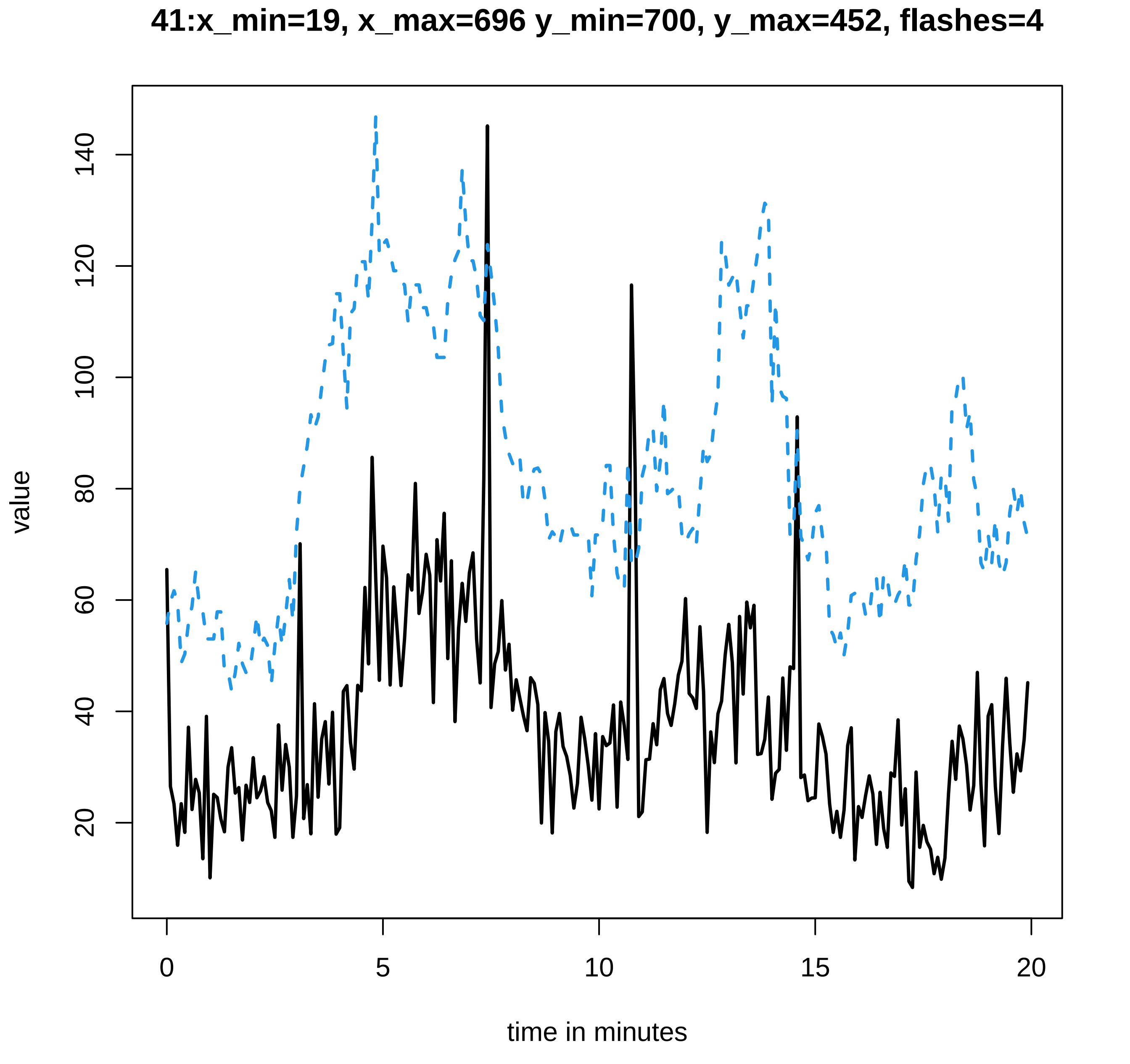
41:x_min=19, x_max=696 y_min=700, y_max=452, flashes=4
value
time in minutes
0
5
10
15
20
20
40
60
80
100
120
140

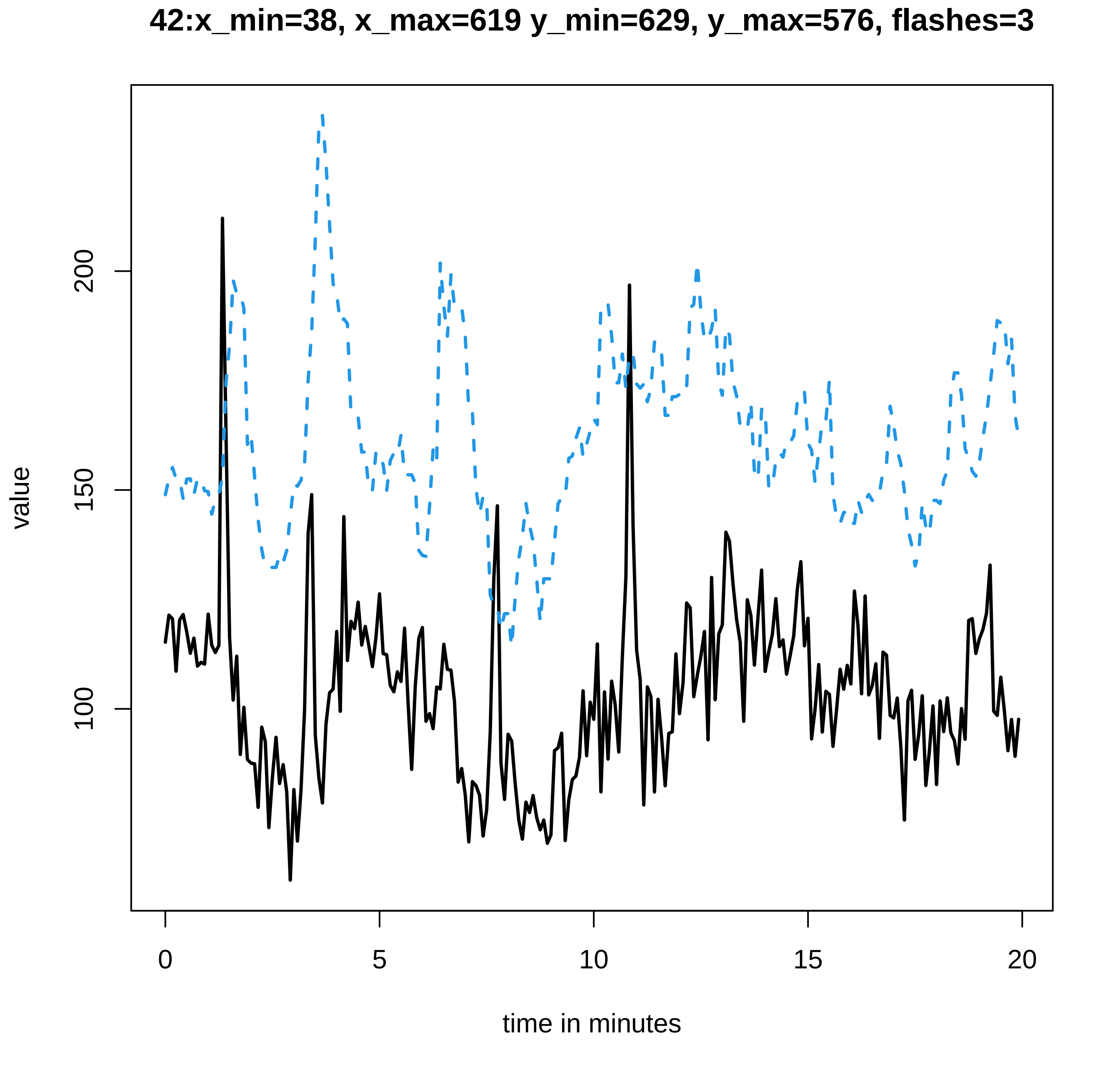
42:x_min=38, x_max=619 y_min=629, y_max=576, flashes=3
value
200
150
100
0
5
10
15
20
time in minutes

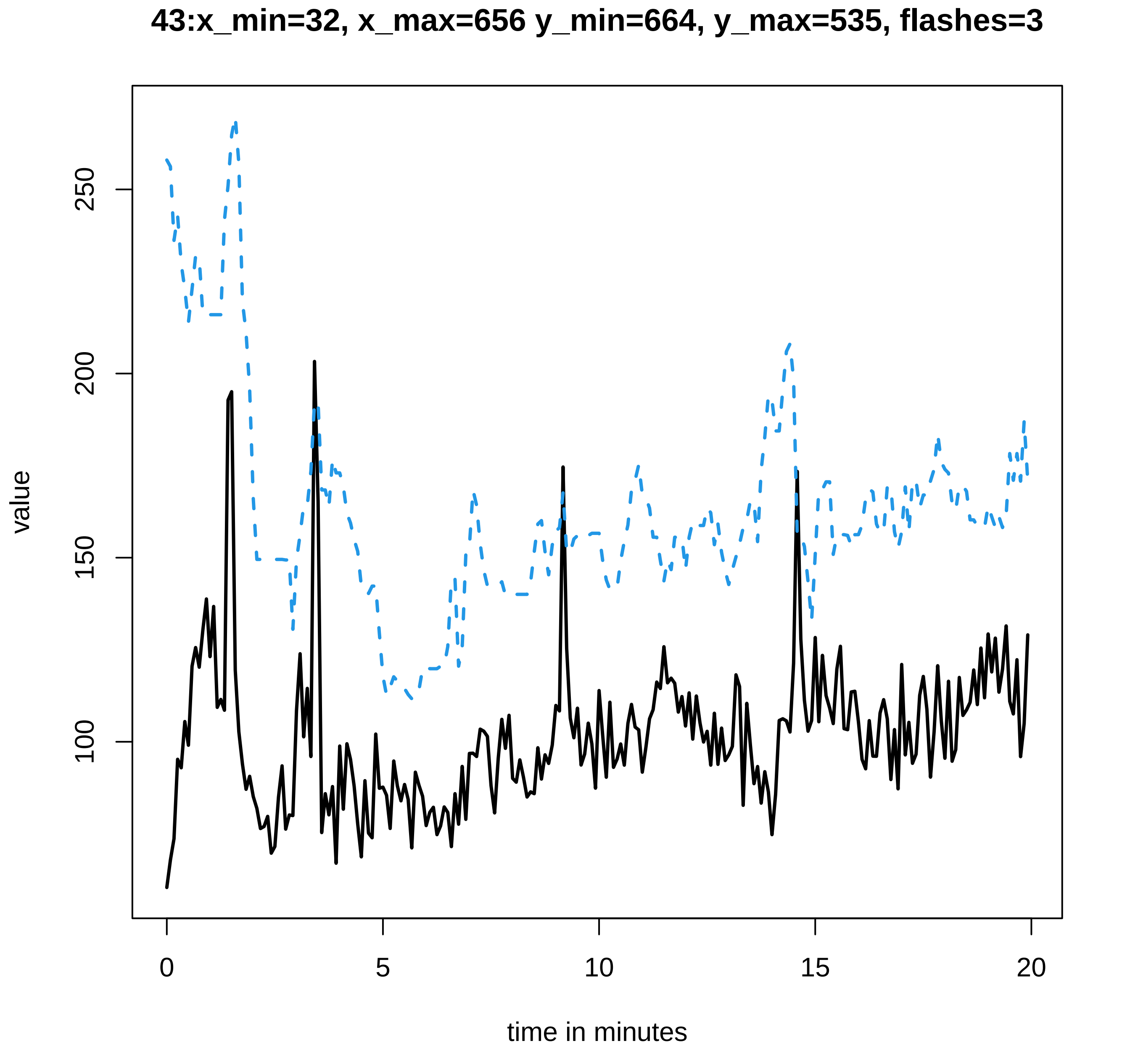
43:x_min=32, x_max=656 y_min=664, y_max=535, flashes=3
value
250
200
150
100
0
5
10
15
20
time in minutes

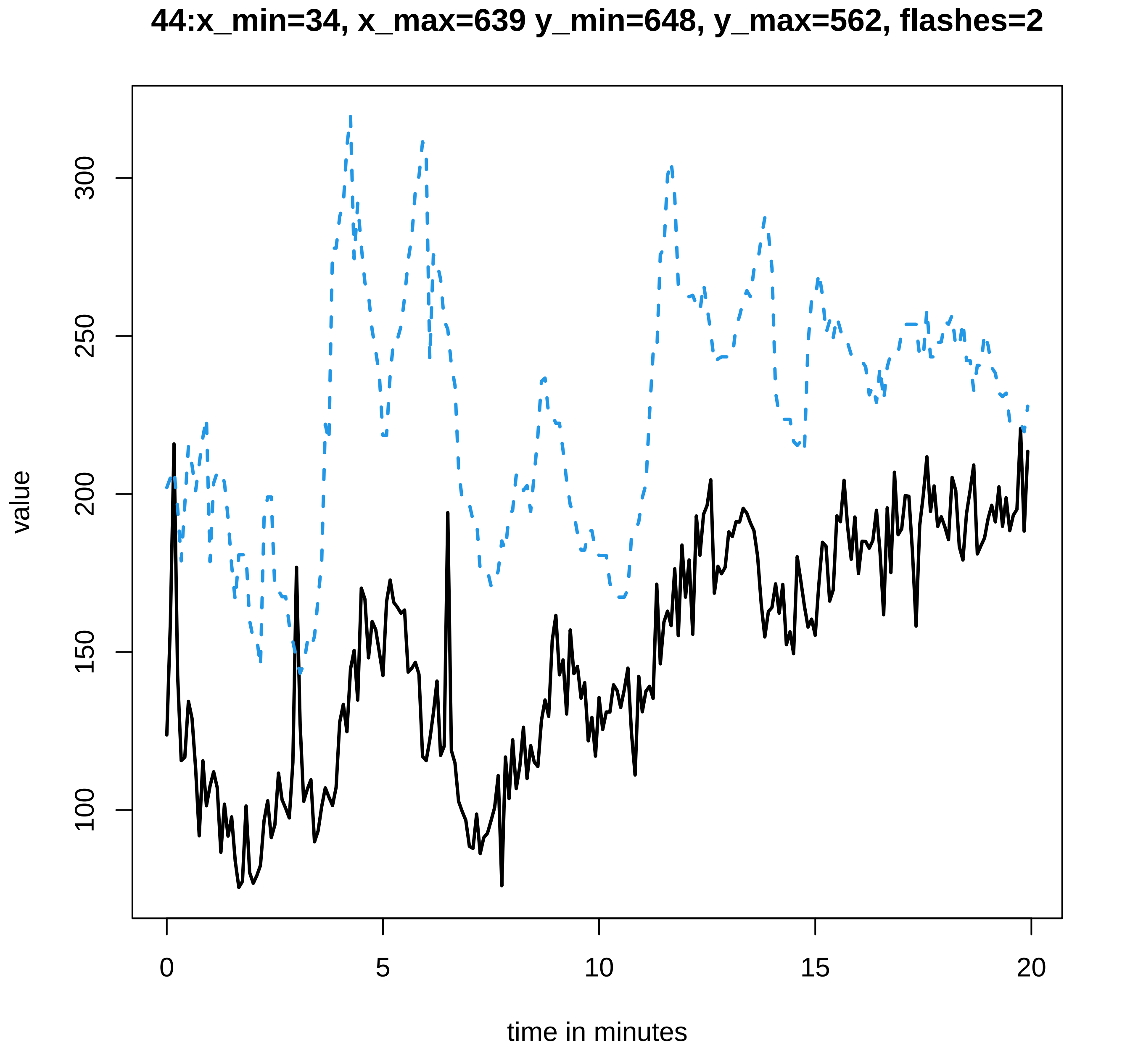
44:x_min=34, x_max=639 y_min=648, y_max=562, flashes=2
value
100
150
200
250
300
0
5
10
15
20
time in minutes

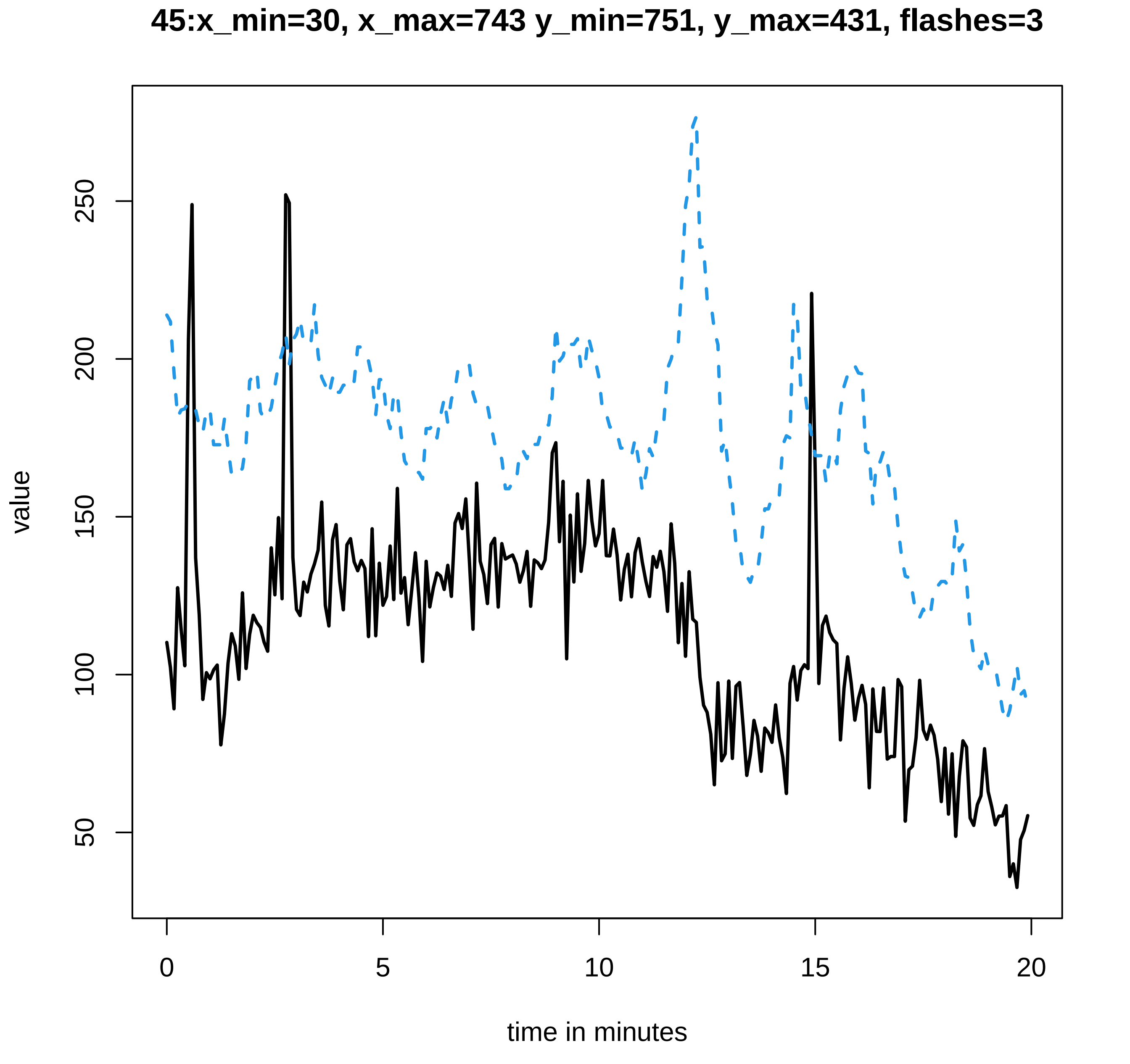
45:x_min=30, x_max=743 y_min=751, y_max=431, flashes=3
value
50
100
150
200
250
0
5
10
15
20
time in minutes

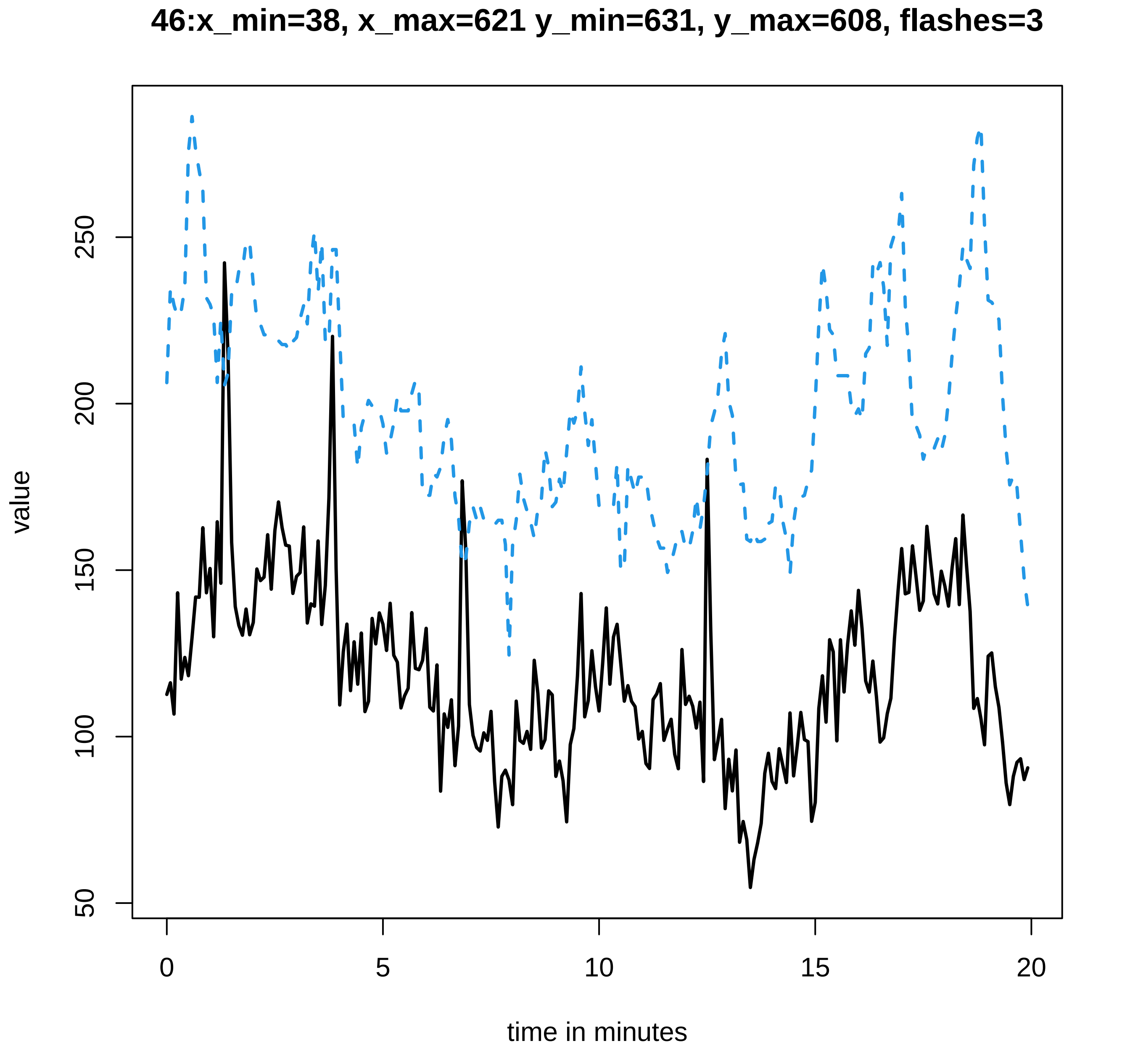

46:x_min=38, x_max=621 y_min=631, y_max=608, flashes=3
value
50
100
150
200
250
0
5
10
15
20
time in minutes

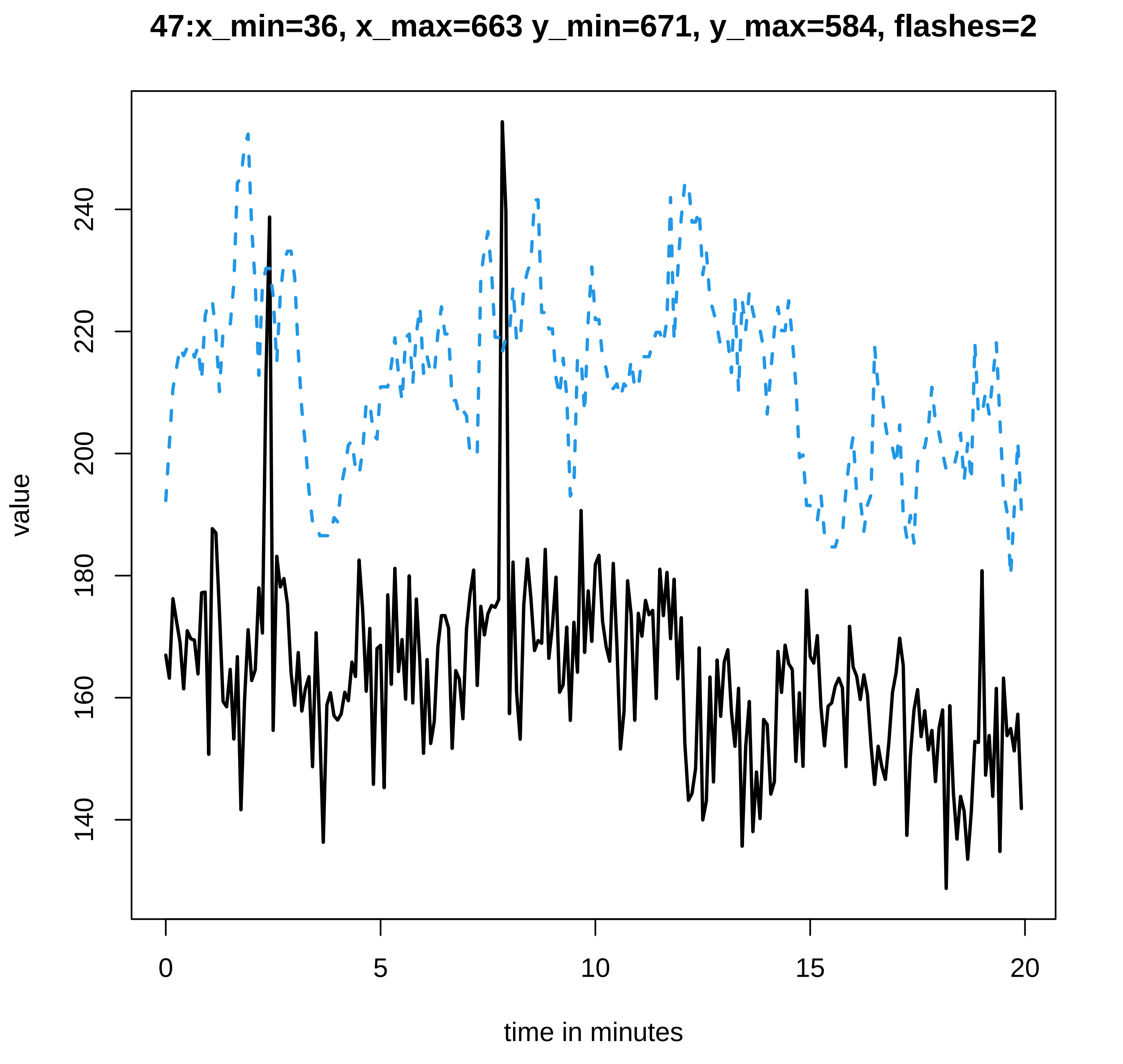

47:x_min=36, x_max=663 y_min=671, y_max=584, flashes=2
value
140
160
180
200
220
240
0
5
10
15
20
time in minutes

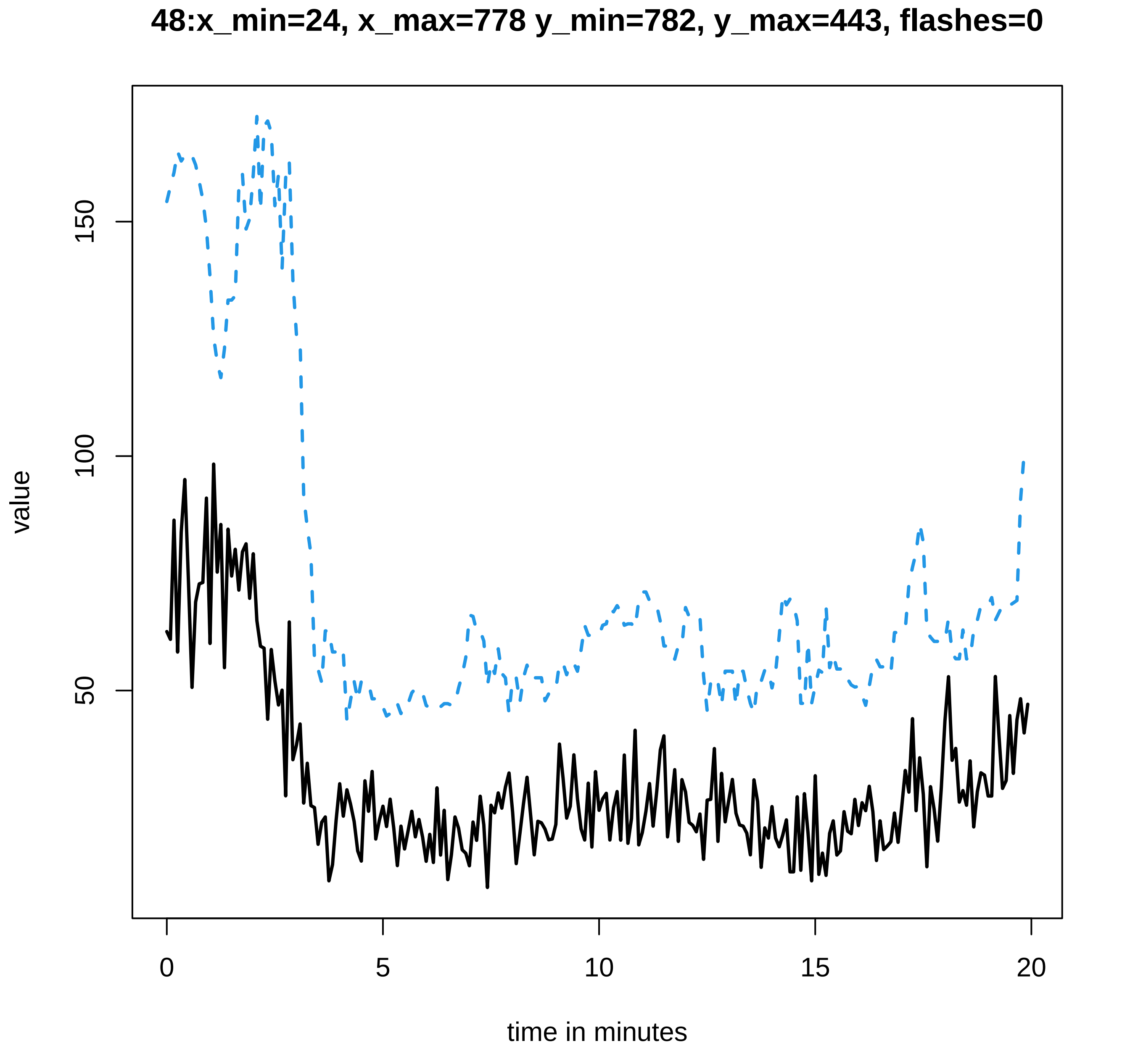

48:x_min=24, x_max=778 y_min=782, y_max=443, flashes=0
value
150
100
50
0
5
10
15
20
time in minutes

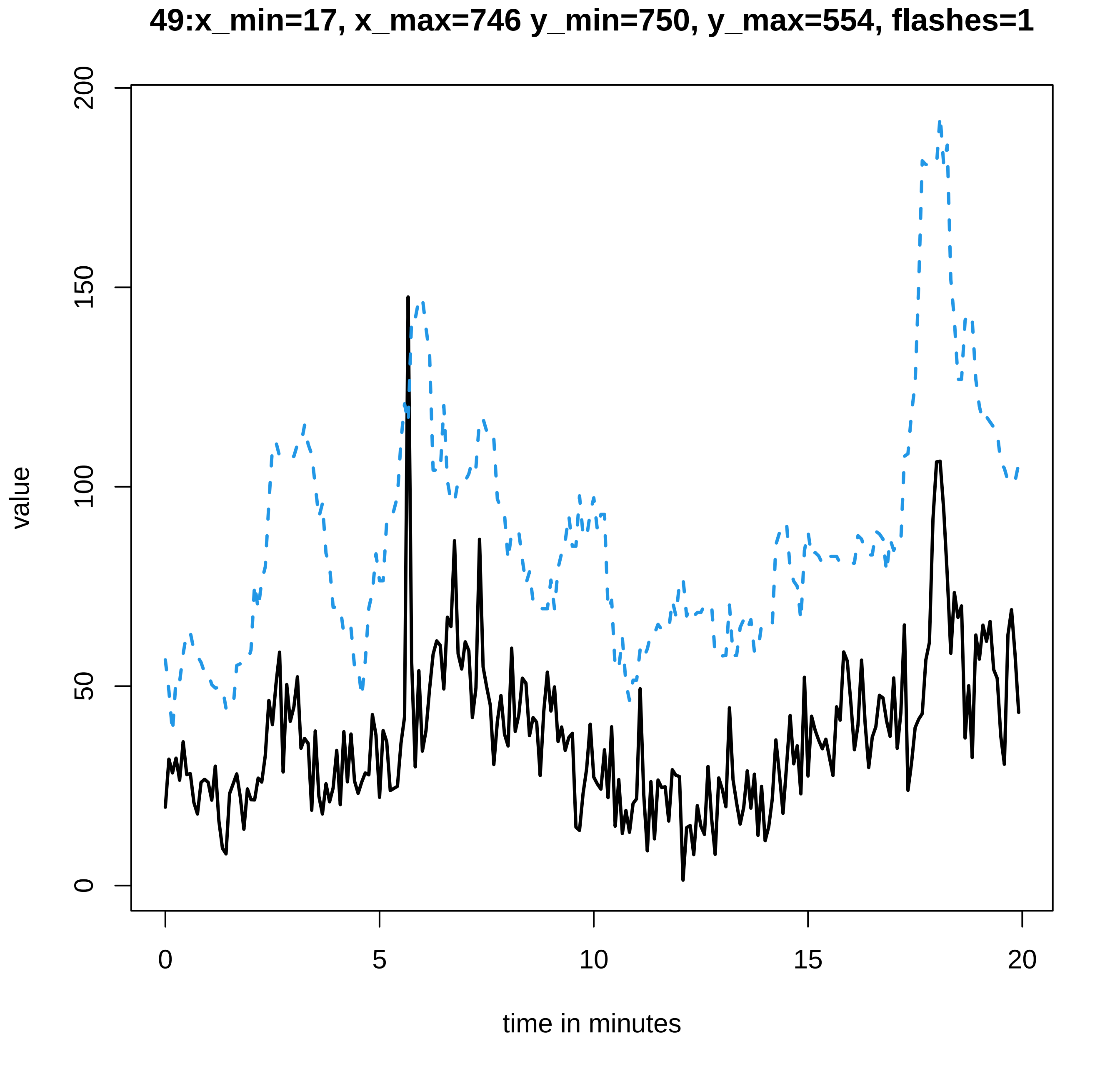
49:x_min=17, x_max=746 y_min=750, y_max=554, flashes=1
200
150
100
50
0
value
0
5
10
15
20
time in minutes

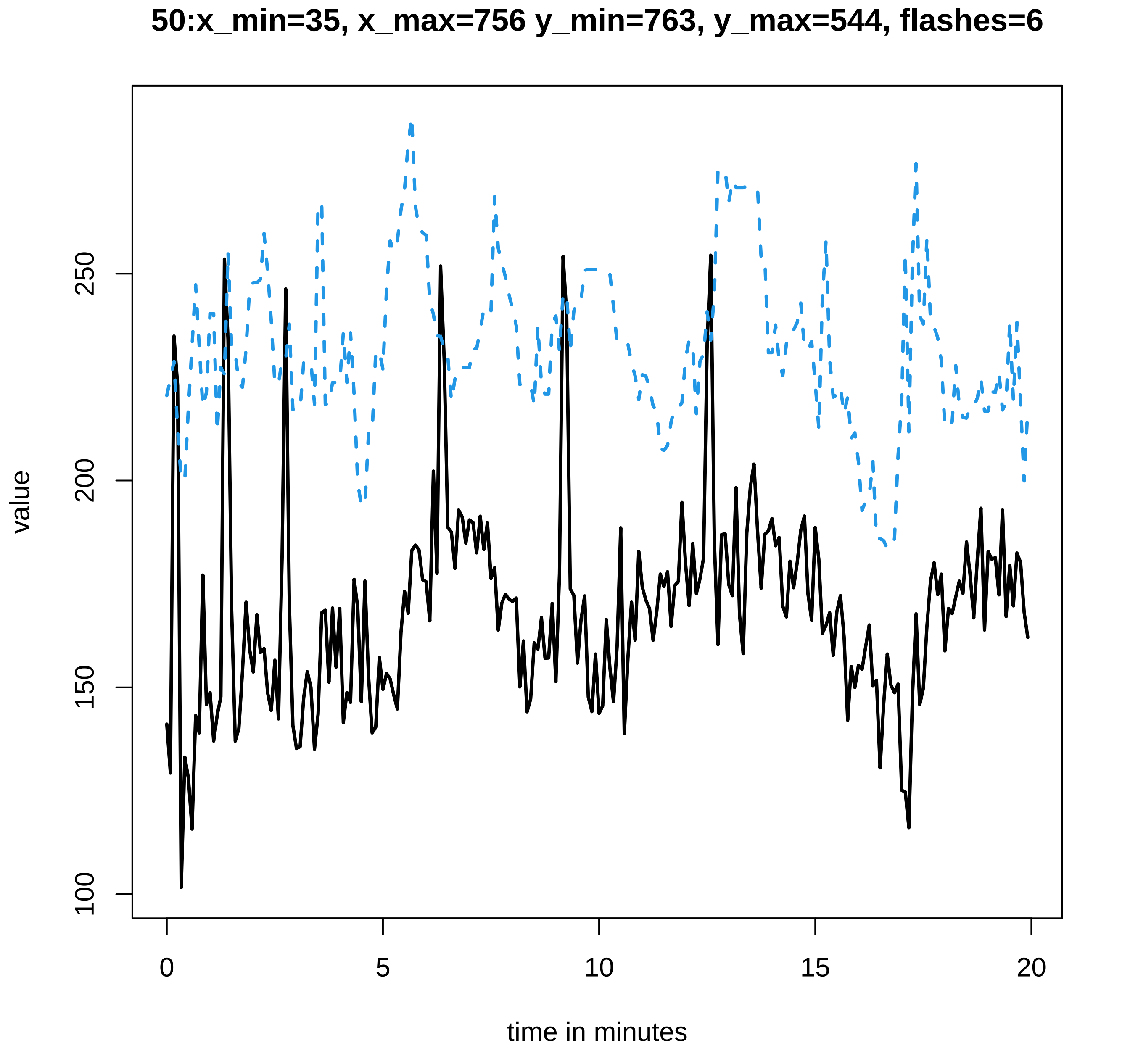
50:x_min=35, x_max=756 y_min=763, y_max=544, flashes=6
value
100
150
200
250
0
5
10
15
20
time in minutes

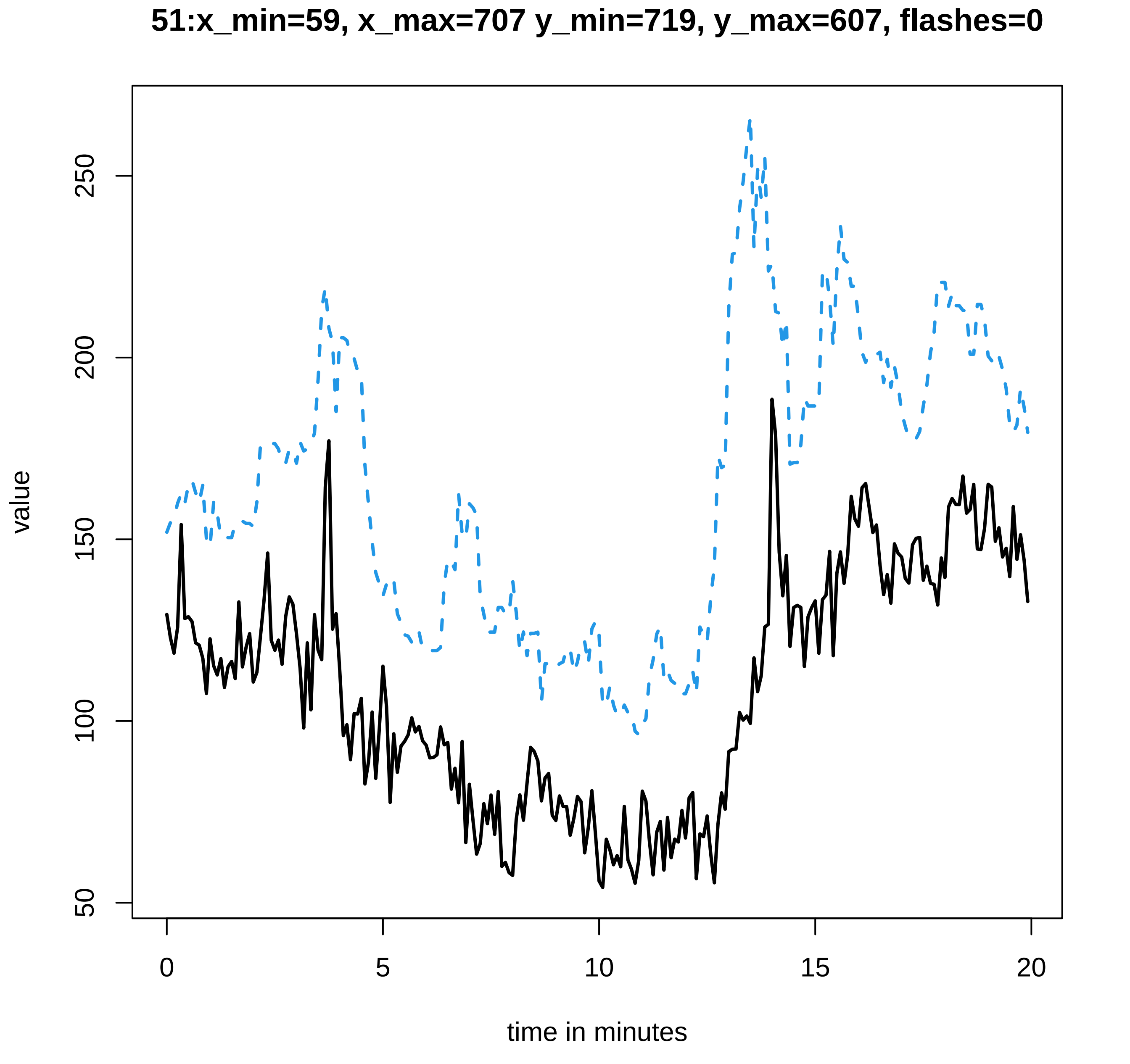
51:x_min=59, x_max=707 y_min=719, y_max=607, flashes=0
value
50
100
150
200
250
0
5
10
15
20
time in minutes

#### 52:x_min=31, x_max=831 y_min=837, y_max=473, flashes=0

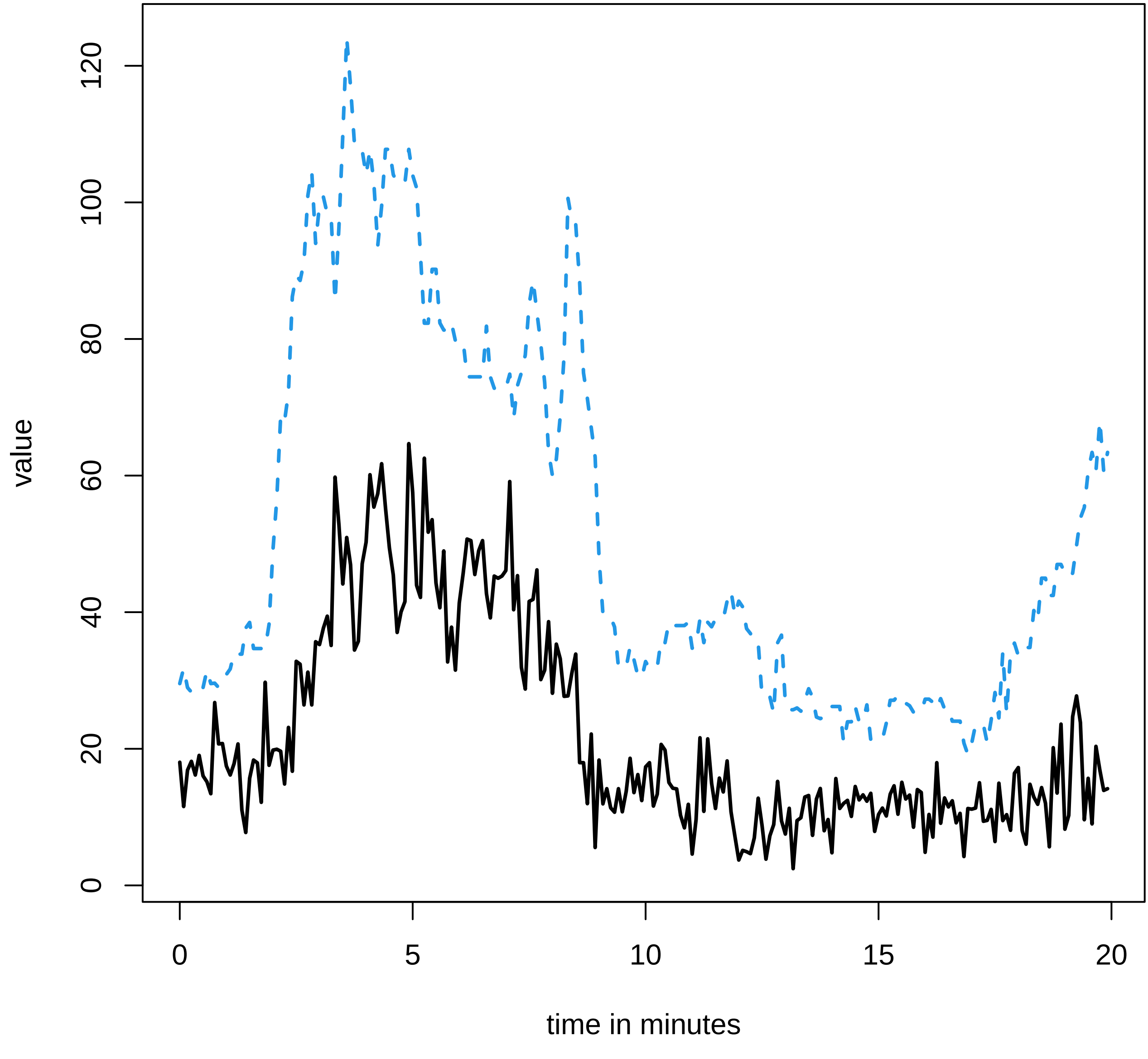

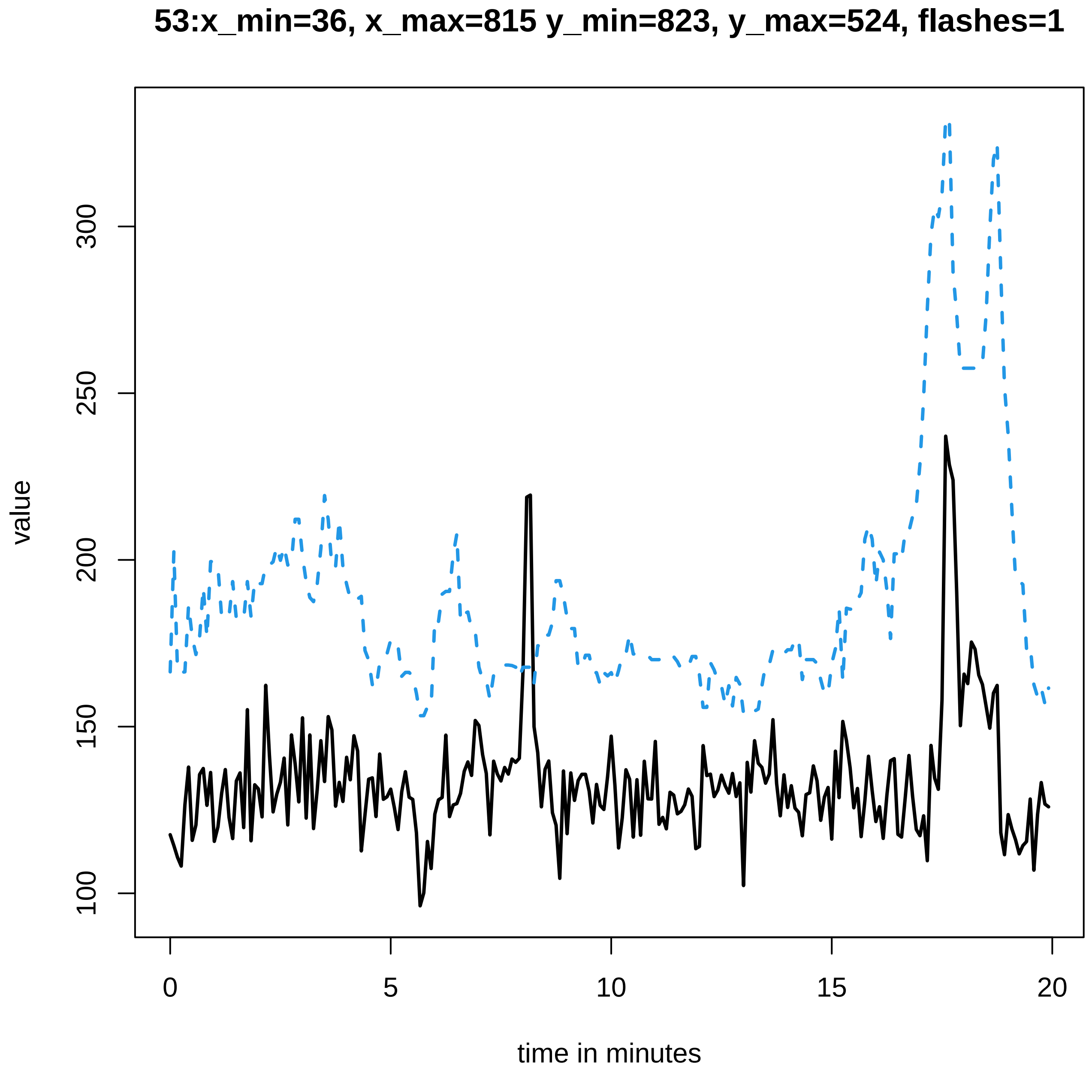
53:x_min=36, x_max=815 y_min=823, y_max=524, flashes=1
value
100
150
200
250
300
0
5
10
15
20
time in minutes

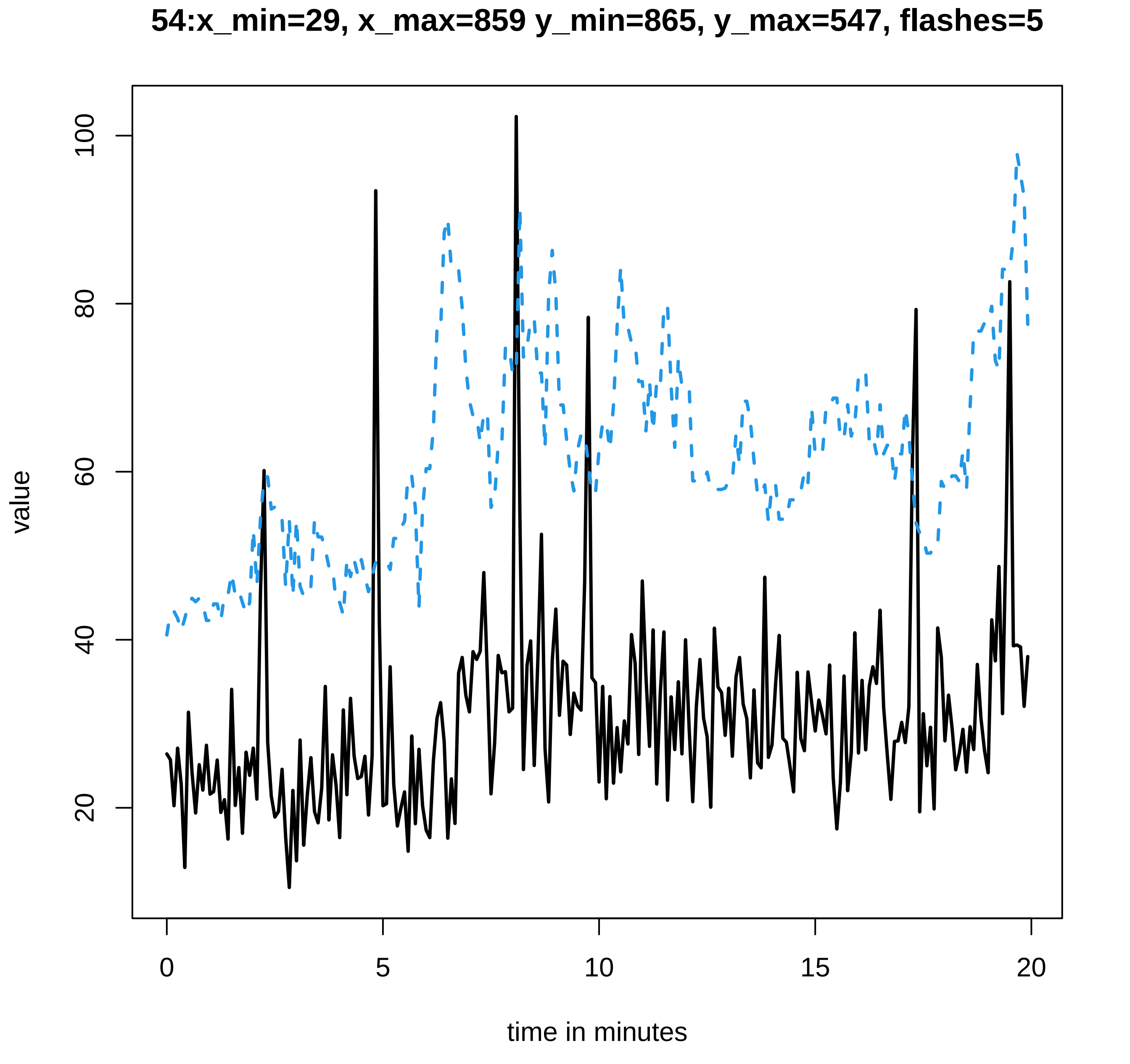
54:x_min=29, x_max=859 y_min=865, y_max=547, flashes=5
value
100
80
60
40
20
0
5
10
15
20
time in minutes

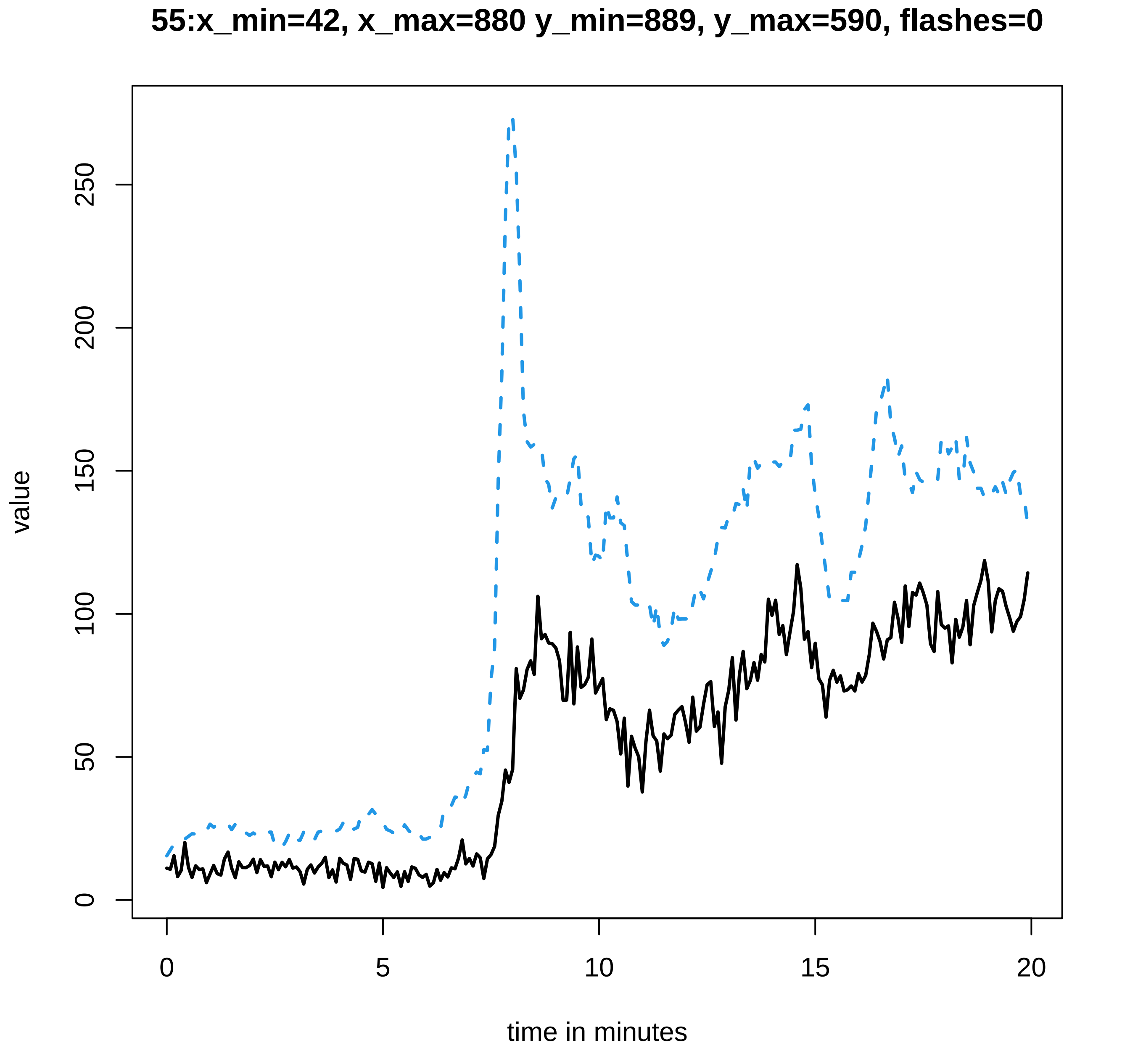
55:x_min=42, x_max=880 y_min=889, y_max=590, flashes=0
value
0
50
100
150
200
250
time in minutes
0
5
10
15
20

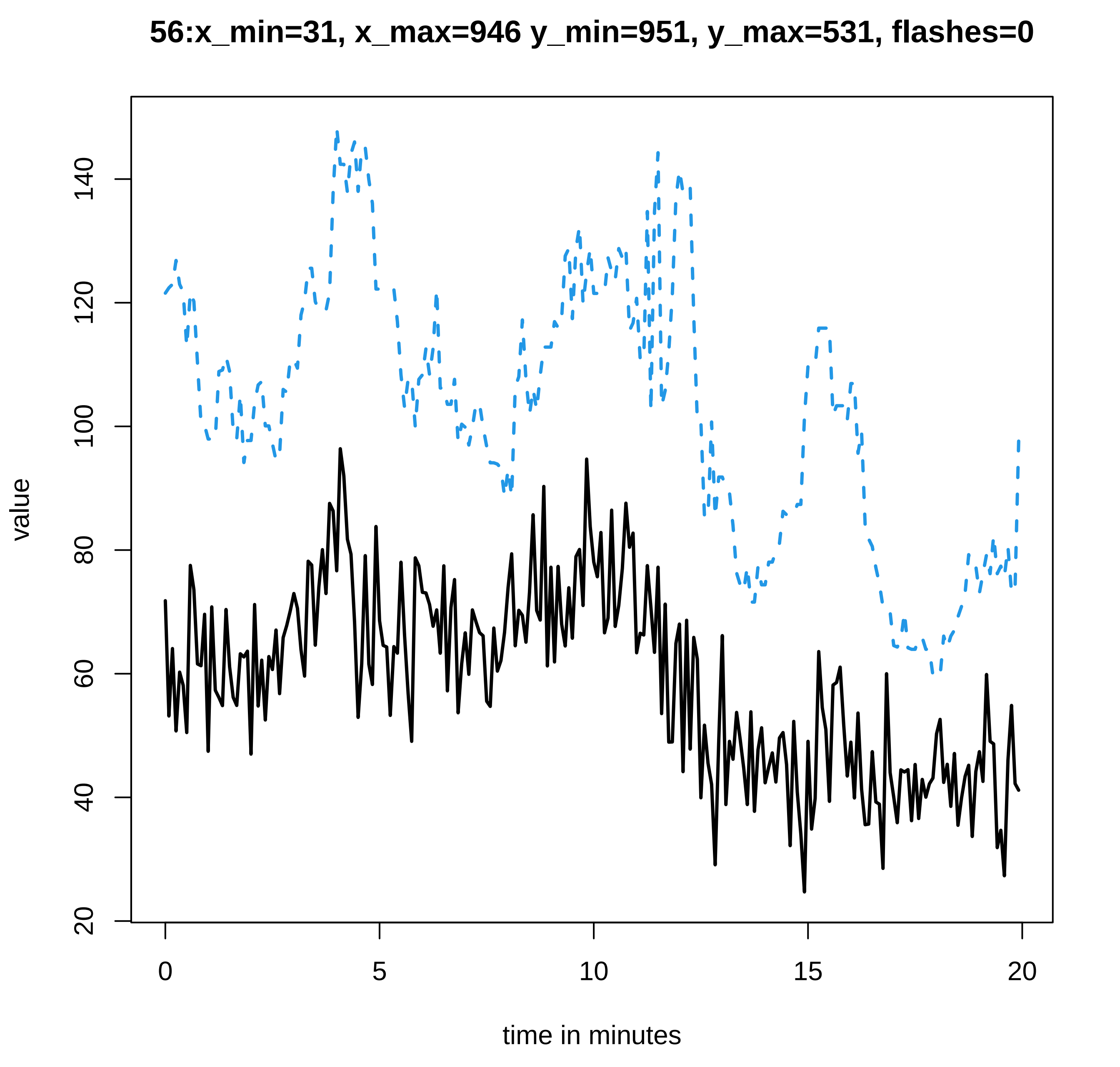
56:x_min=31, x_max=946 y_min=951, y_max=531, flashes=0
value
20
40
60
80
100
120
140
0
5
10
15
20
time in minutes

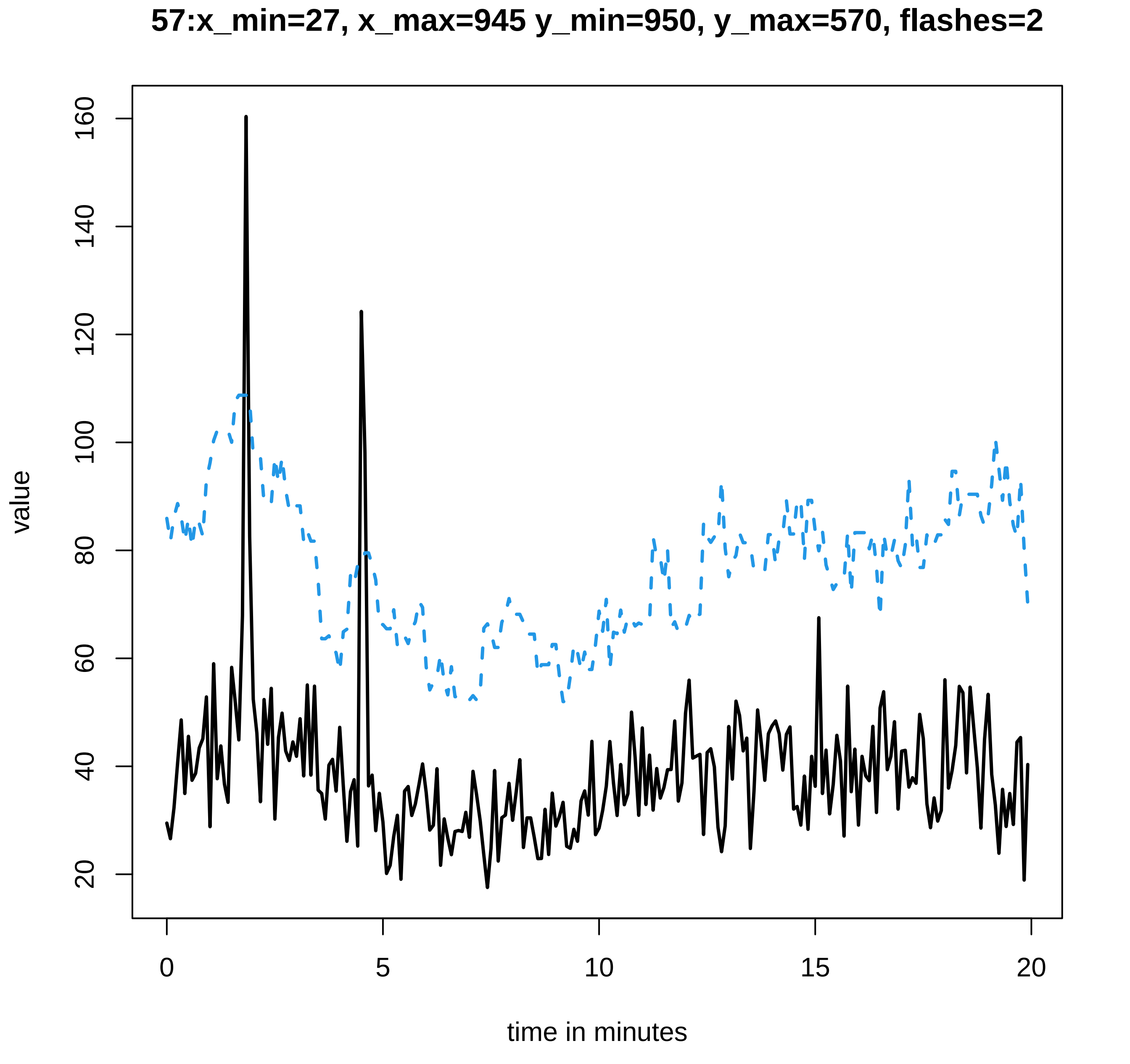

57:x_min=27, x_max=945 y_min=950, y_max=570, flashes=2
value
time in minutes
0
5
10
15
20
20
40
60
80
100
120
140
160

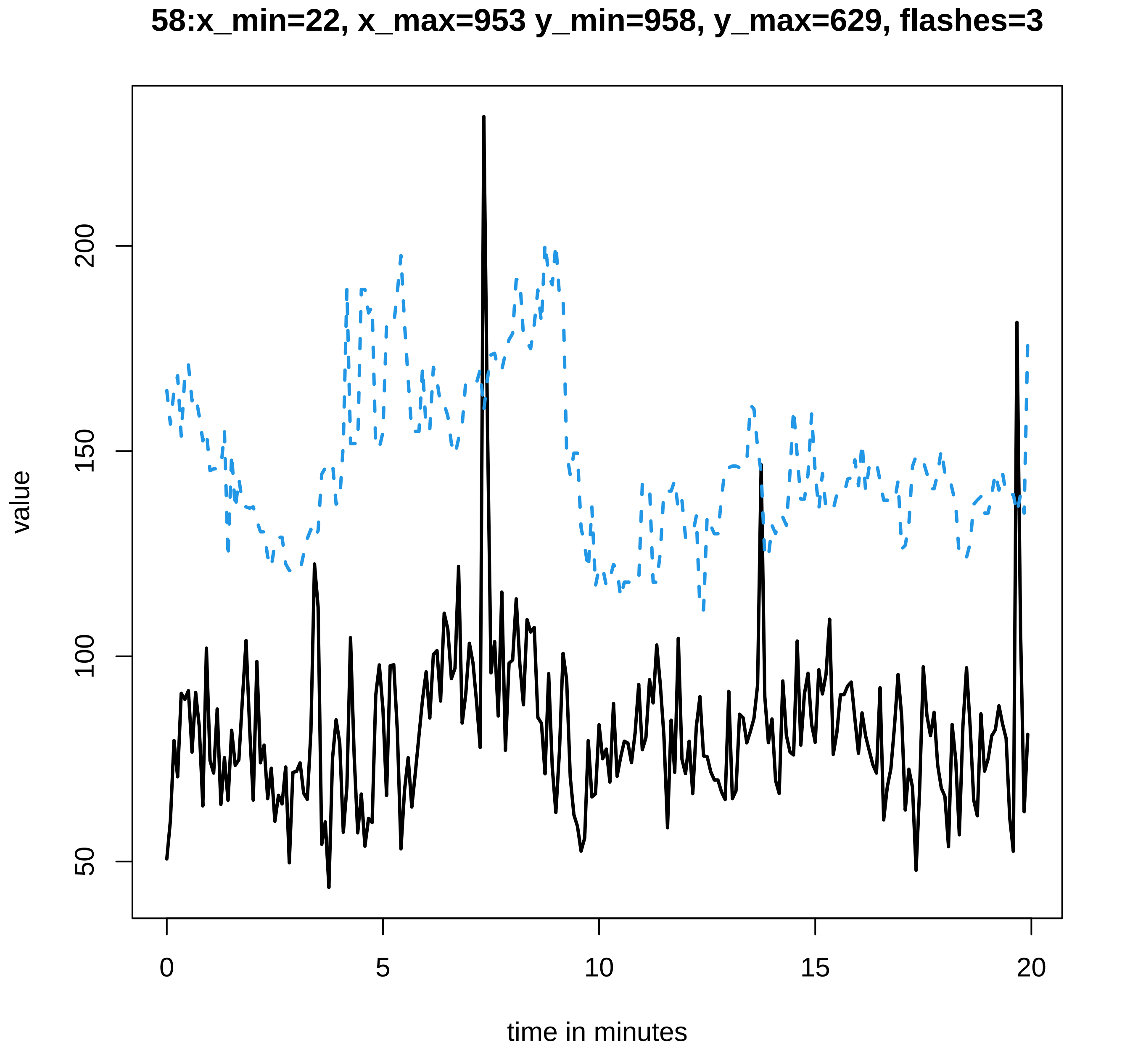
58:x_min=22, x_max=953 y_min=958, y_max=629, flashes=3
value
50
100
150
200
0
5
10
15
20
time in minutes
