## Supplementary data Calcium spiking for "LysM Receptor Proteins are Required for Ectomycorrhizal Symbiosis in Poplar": profiles-spiking-8.pdf

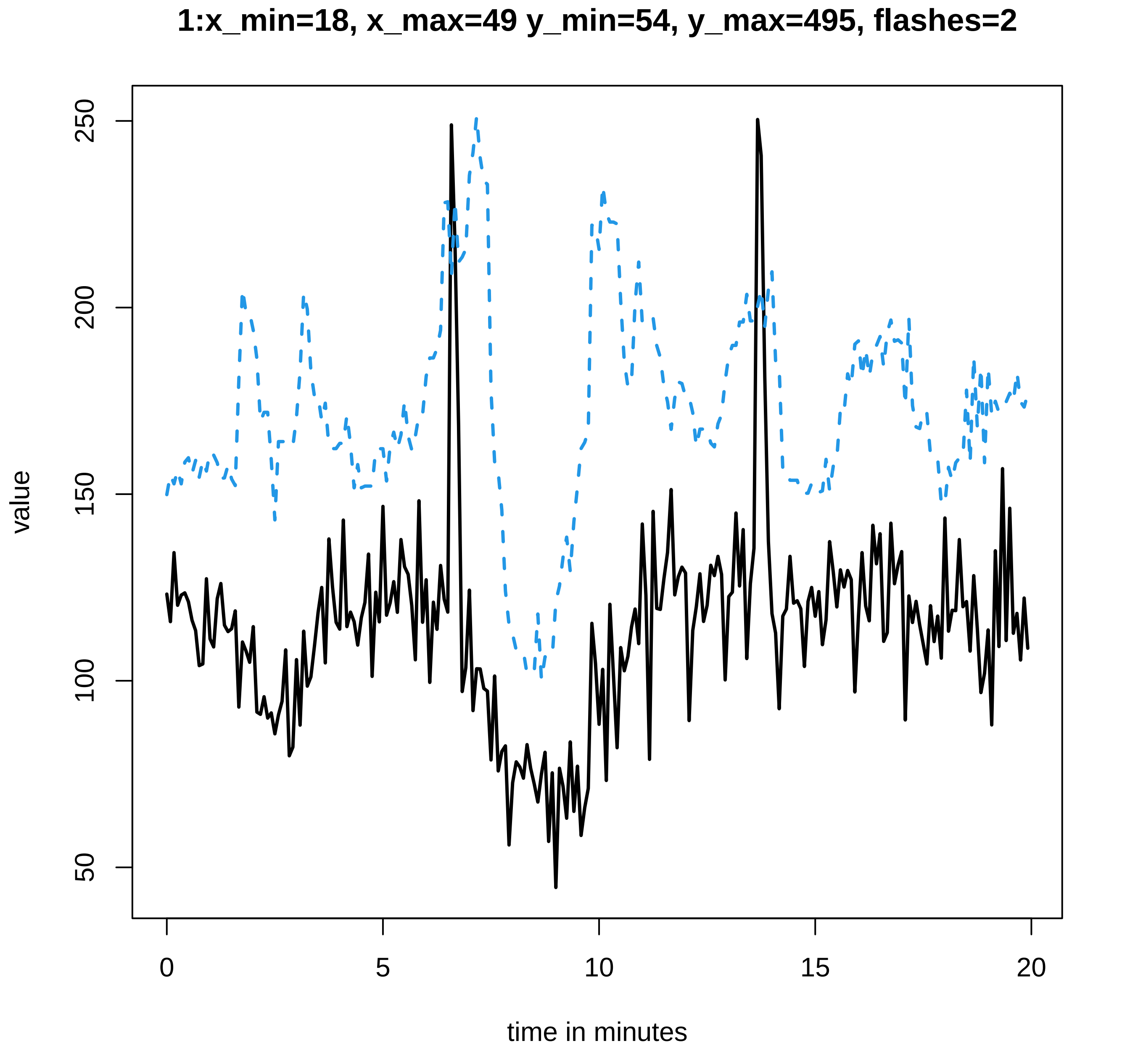

1:x_min=18, x_max=49 y_min=54, y_max=495, flashes=2
value
50
100
150
200
250
0
5
10
15
20
time in minutes

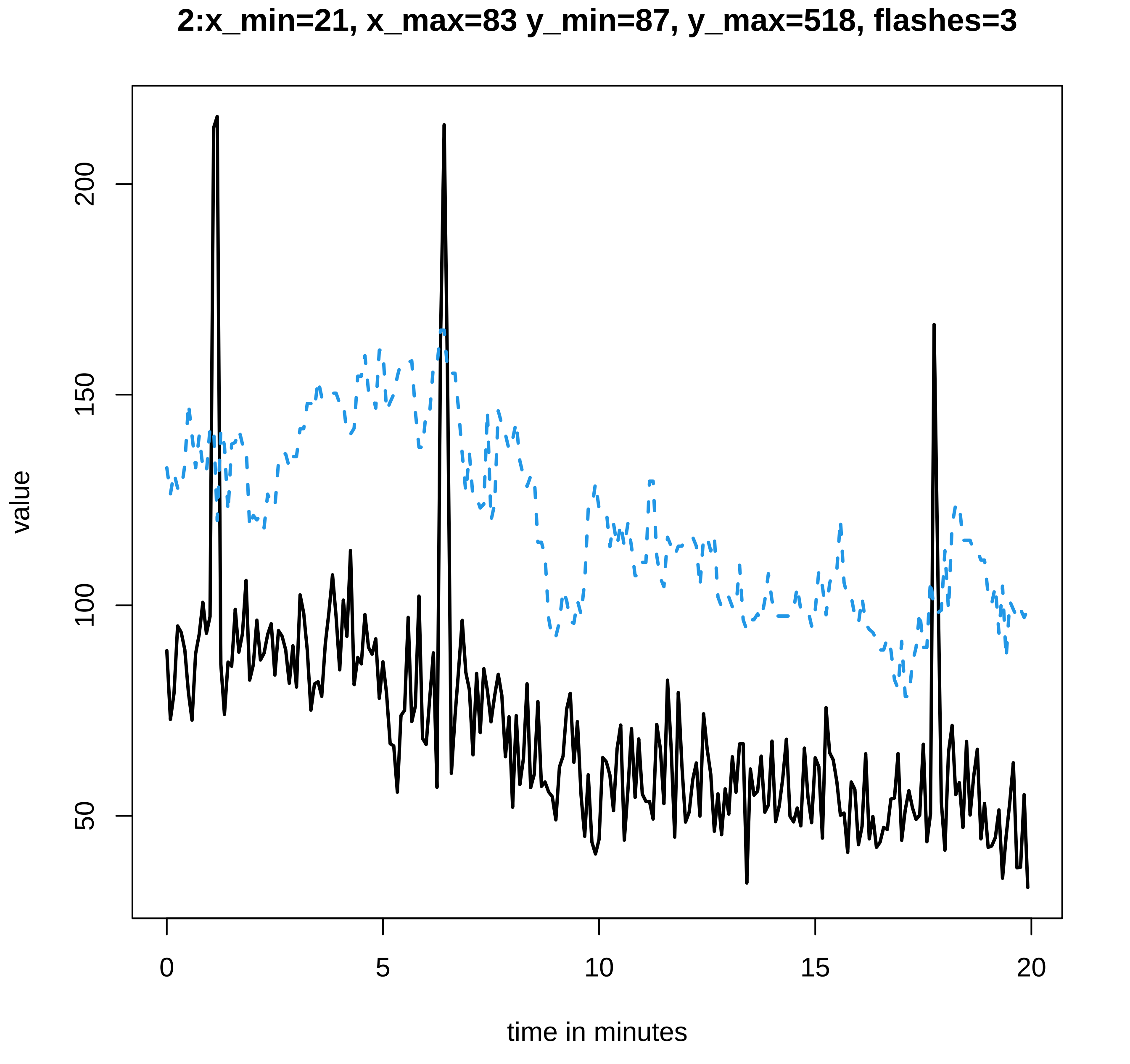

2:x_min=21, x_max=83 y_min=87, y_max=518, flashes=3
value
200
150
100
50
0
5
10
15
20
time in minutes

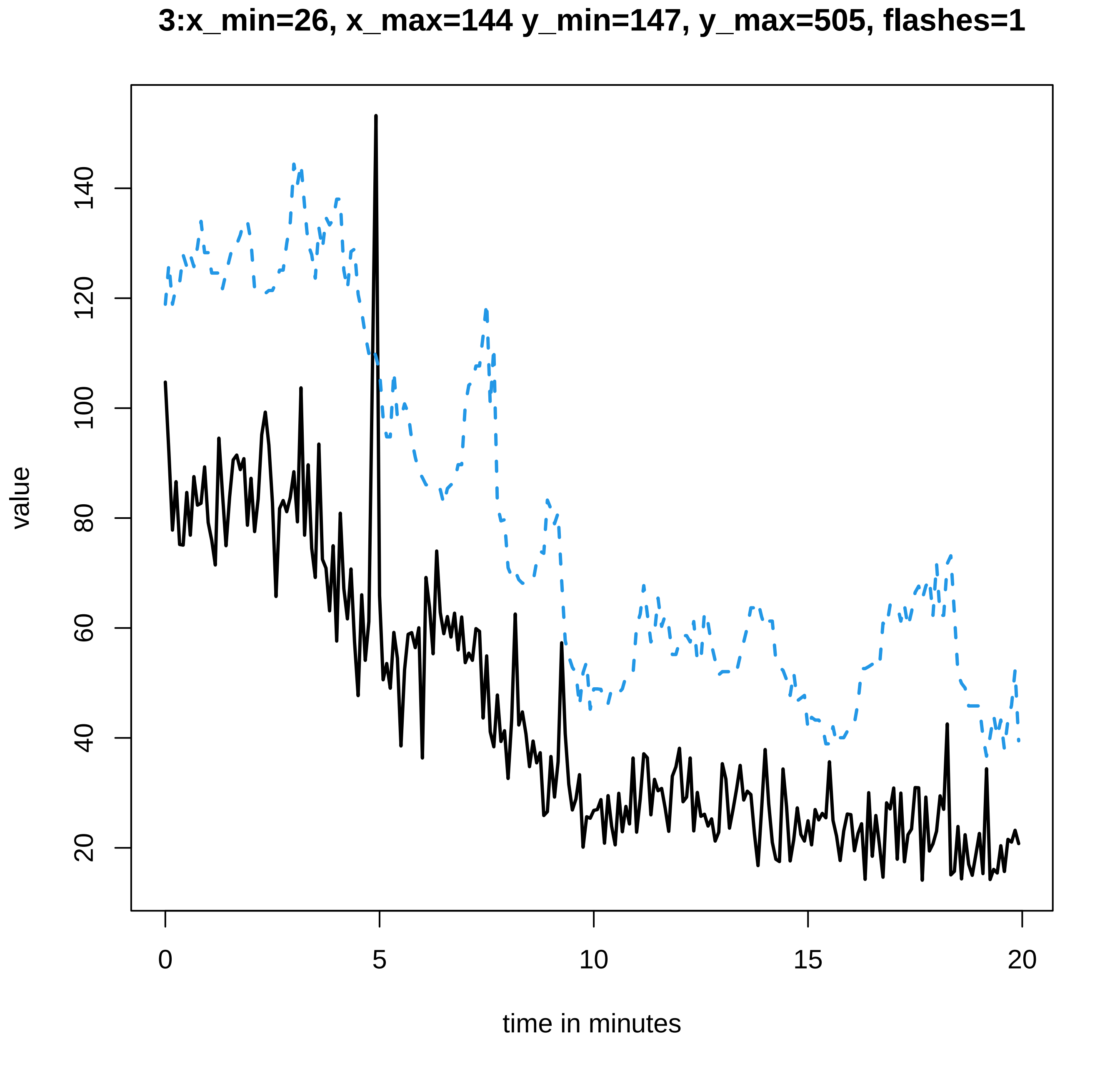

3:x_min=26, x_max=144 y_min=147, y_max=505, flashes=1
value
20
40
60
80
100
120
140
0
5
10
15
20
time in minutes

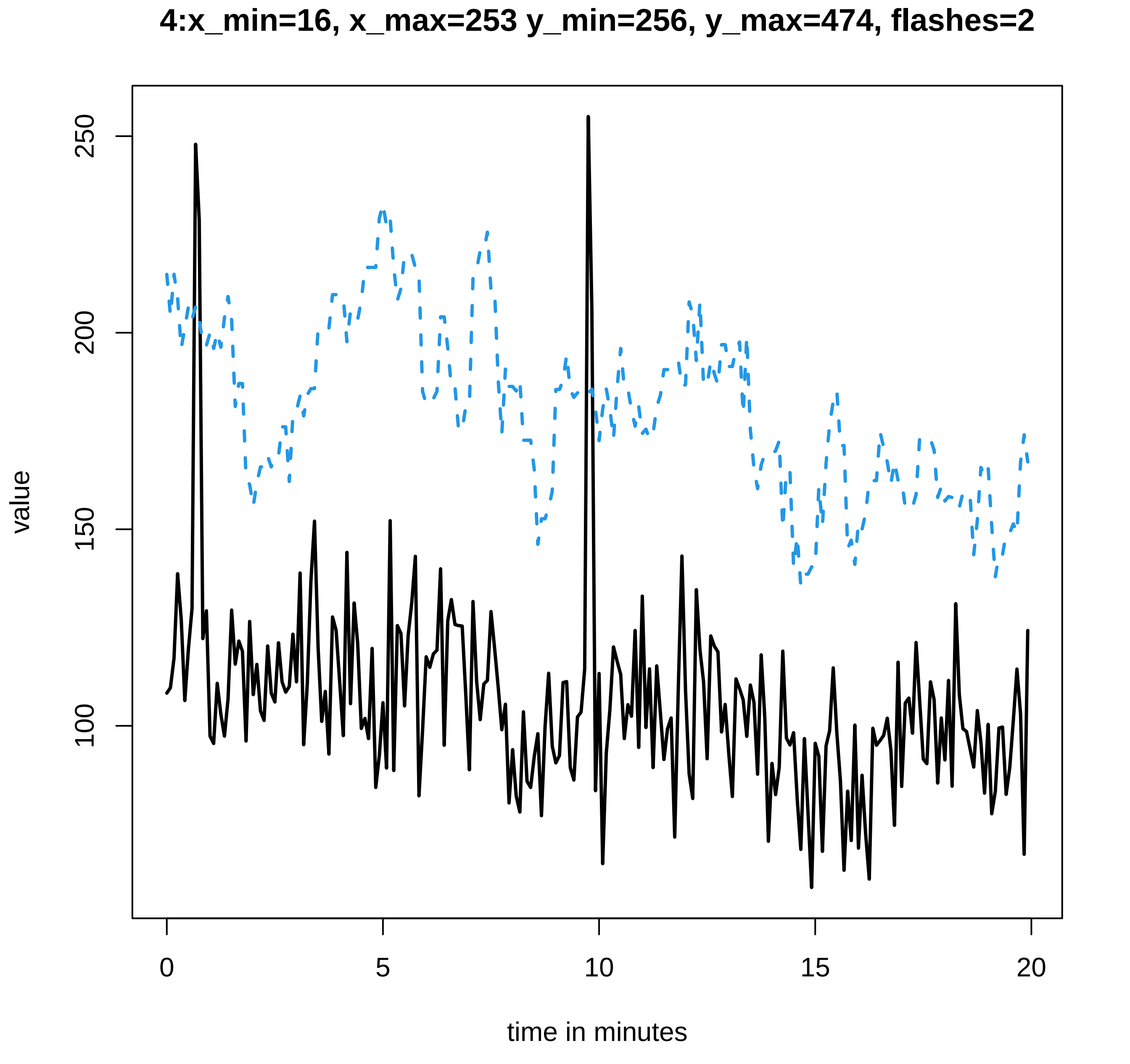

4:x_min=16, x_max=253 y_min=256, y_max=474, flashes=2
value
250
200
150
100
0
5
10
15
20
time in minutes

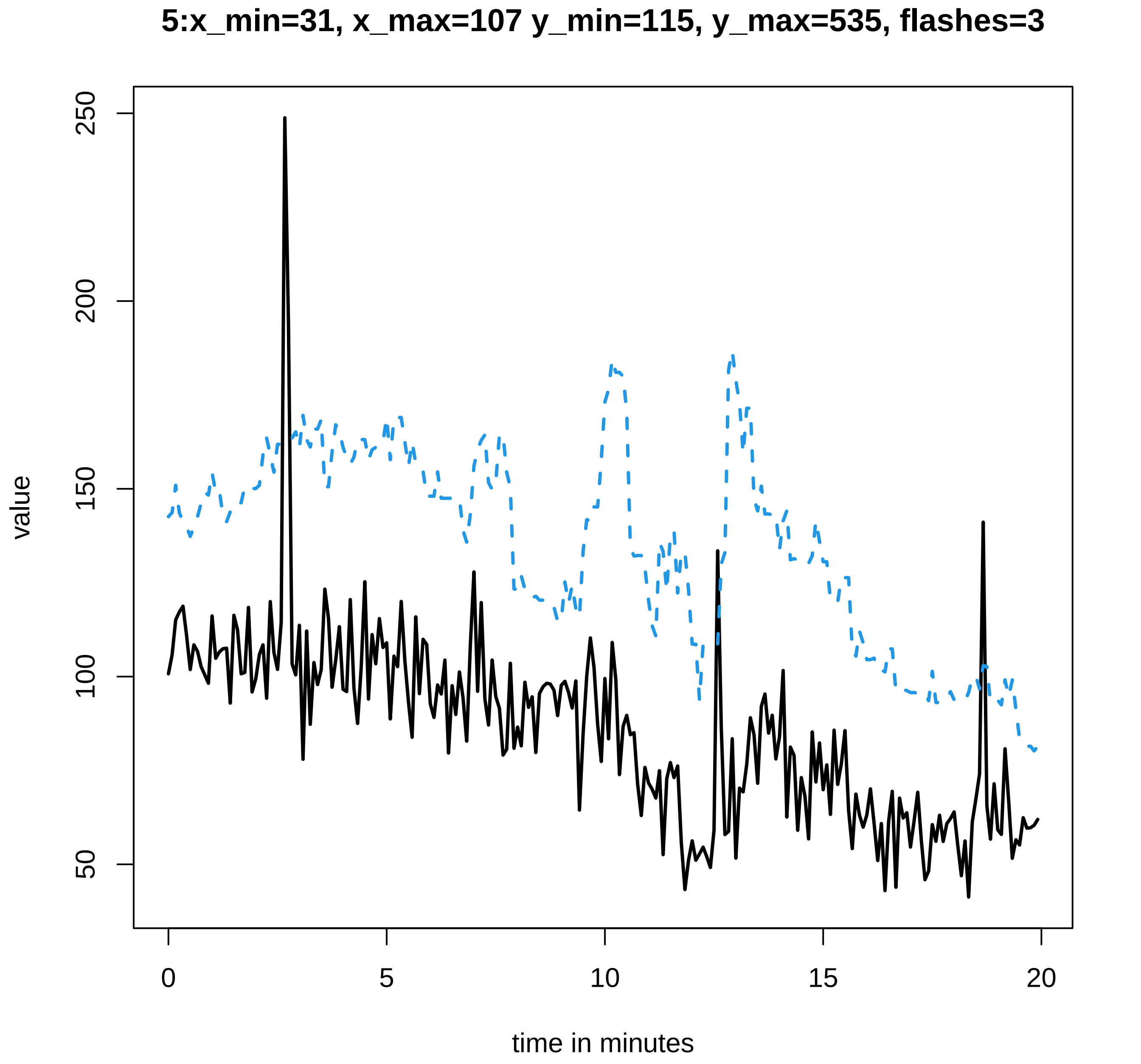

5:x_min=31, x_max=107 y_min=115, y_max=535, flashes=3
value
50
100
150
200
250
0
5
10
15
20
time in minutes

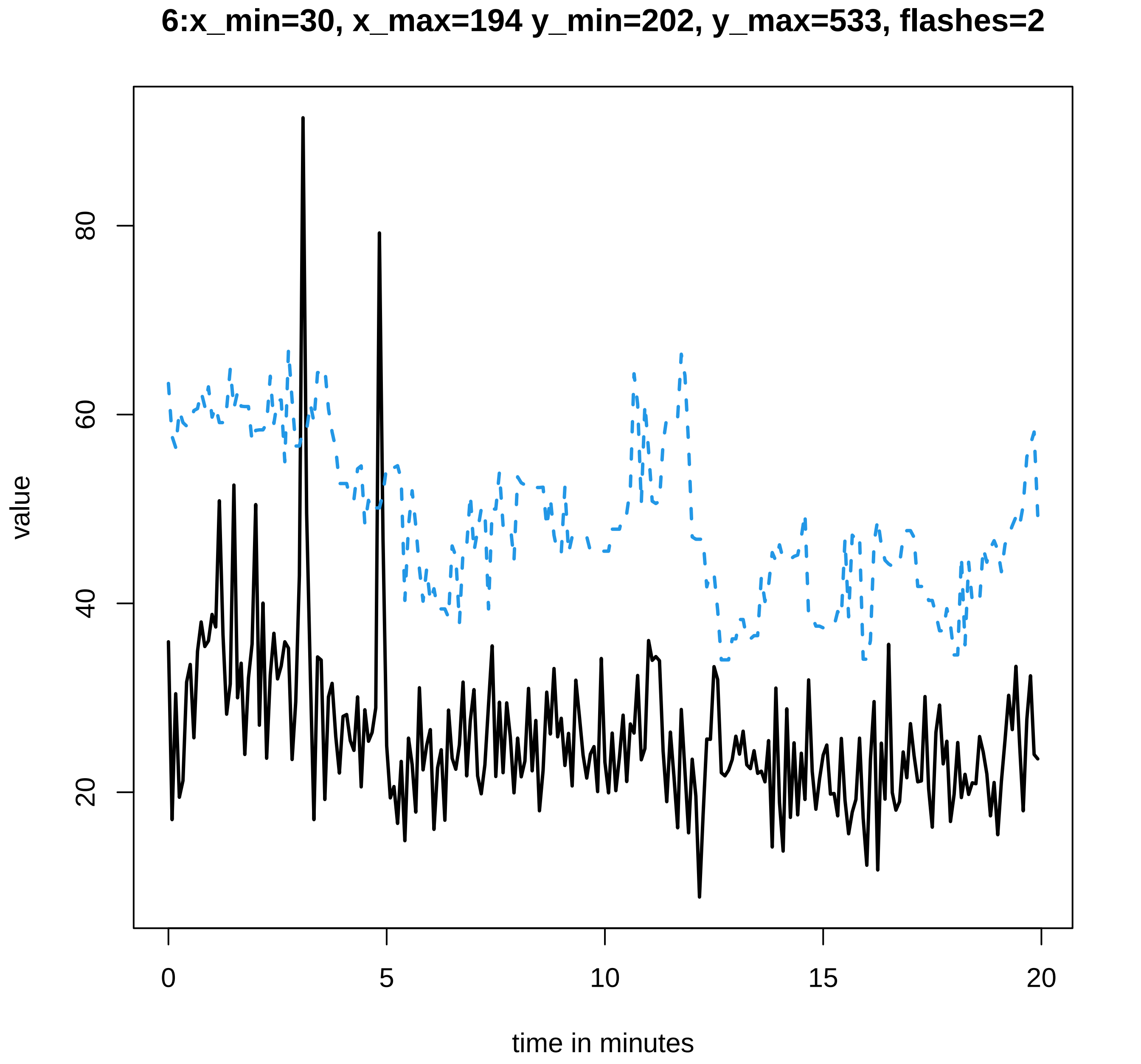

6:x_min=30, x_max=194 y_min=202, y_max=533, flashes=2
value
20
40
60
80
0
5
10
15
20
time in minutes

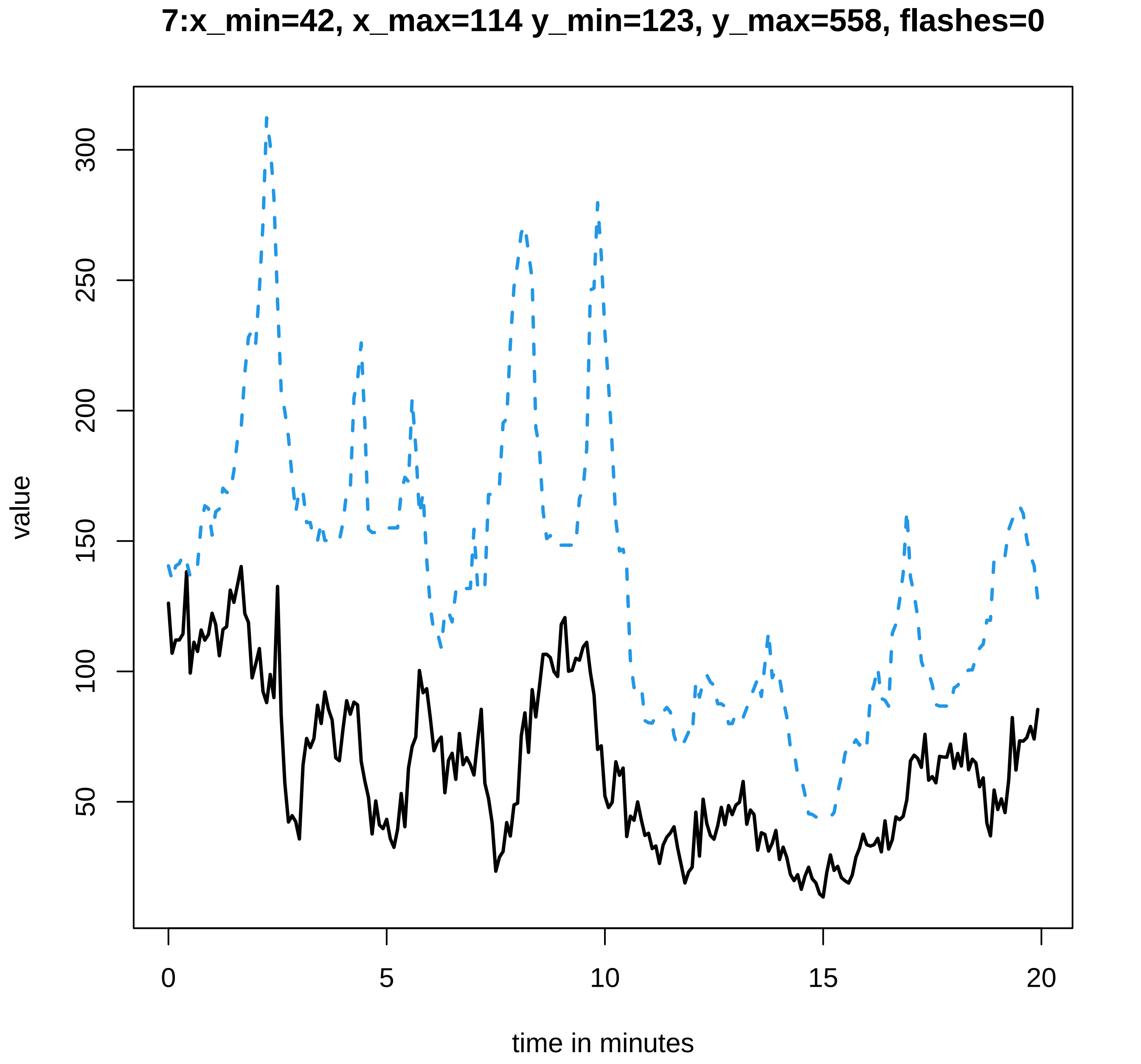

7:x_min=42, x_max=114 y_min=123, y_max=558, flashes=0
value
300
250
200
150
100
50
0
5
10
15
20
time in minutes

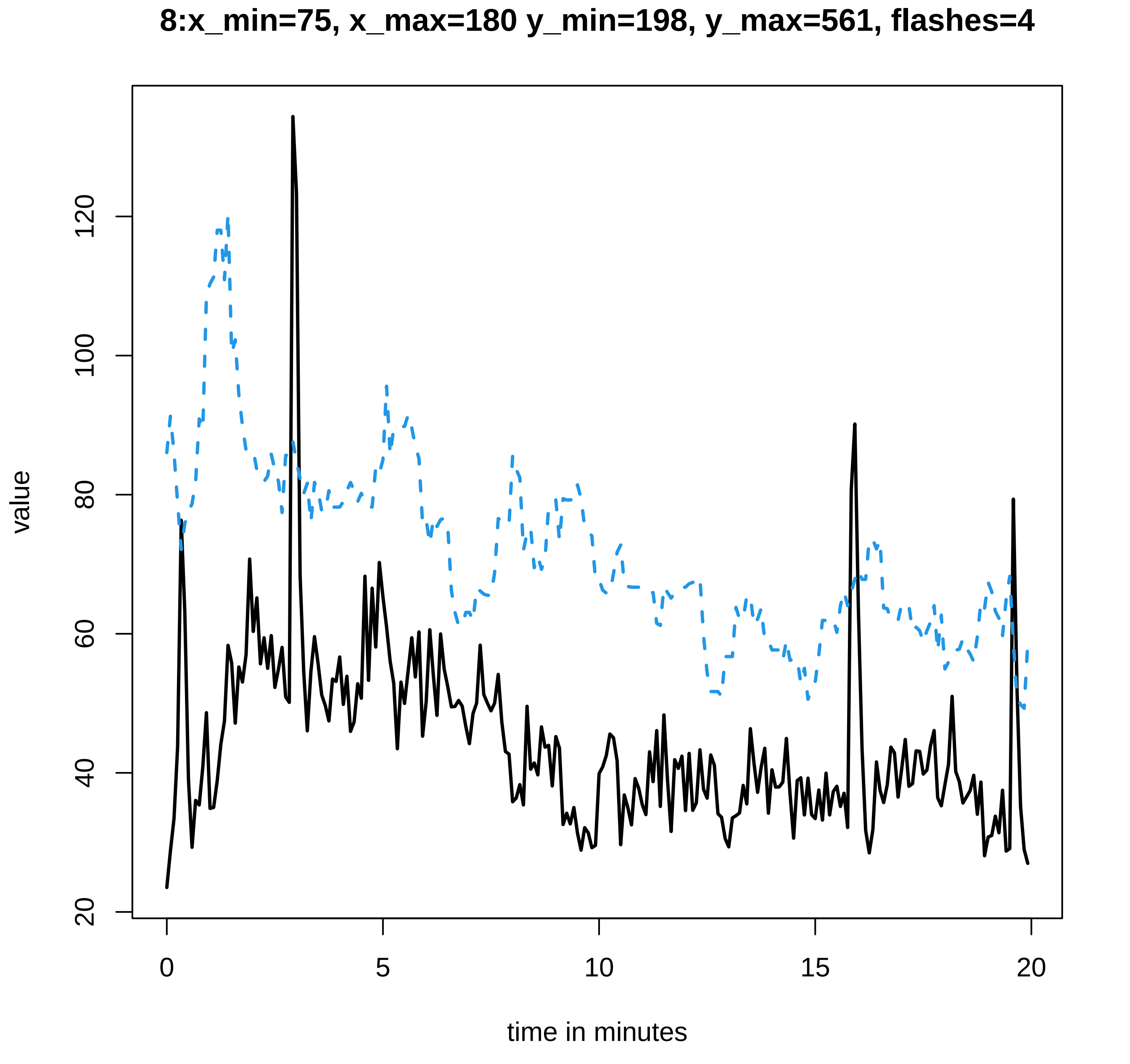

8:x_min=75, x_max=180 y_min=198, y_max=561, flashes=4
value
20
40
60
80
100
120
0
5
10
15
20
time in minutes

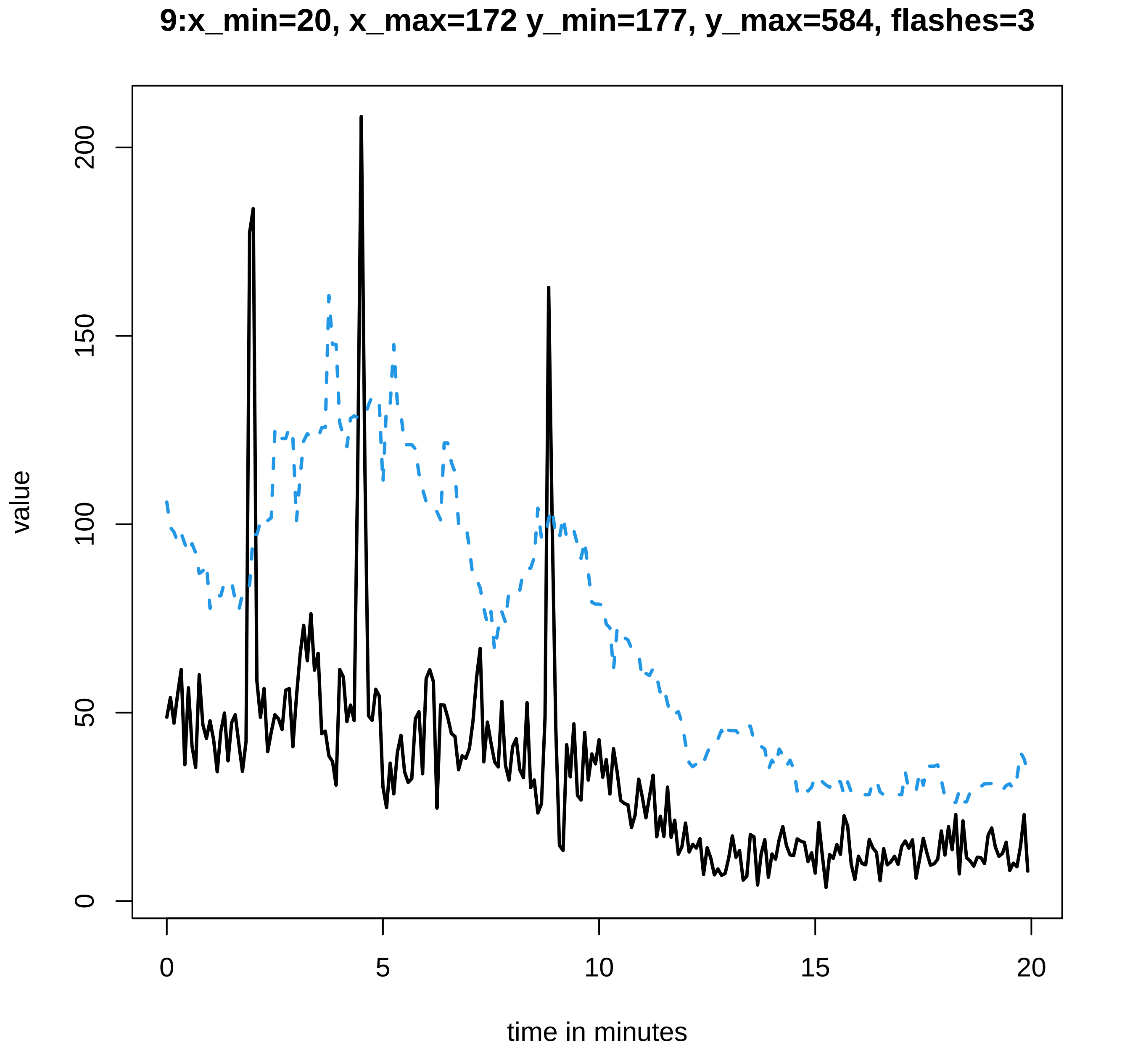

9:x_min=20, x_max=172 y_min=177, y_max=584, flashes=3
value
0
50
100
150
200
time in minutes
0
5
10
15
20

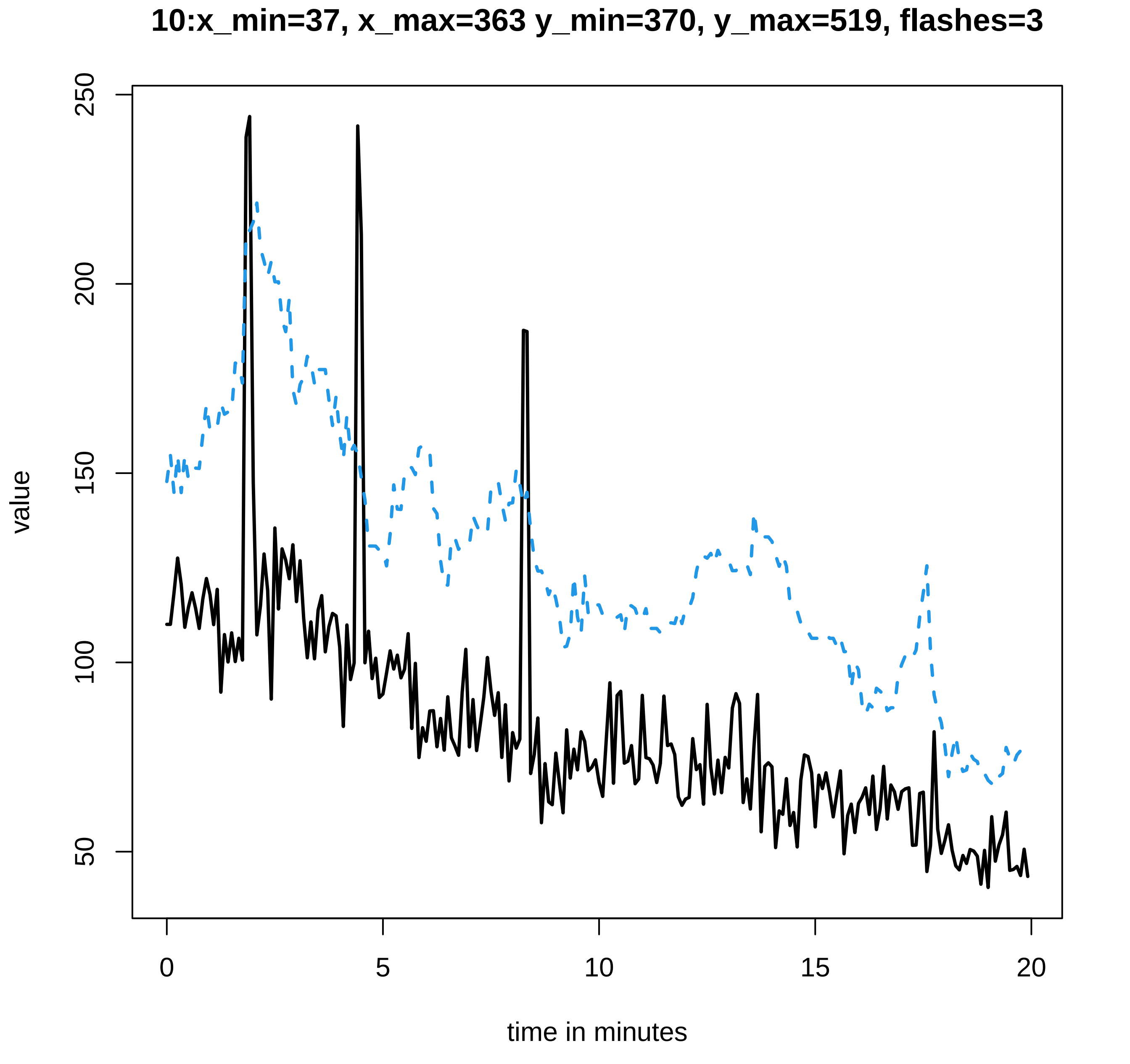
10:x_min=37, x_max=363 y_min=370, y_max=519, flashes=3
value
250
200
150
100
50
0
5
10
15
20
time in minutes

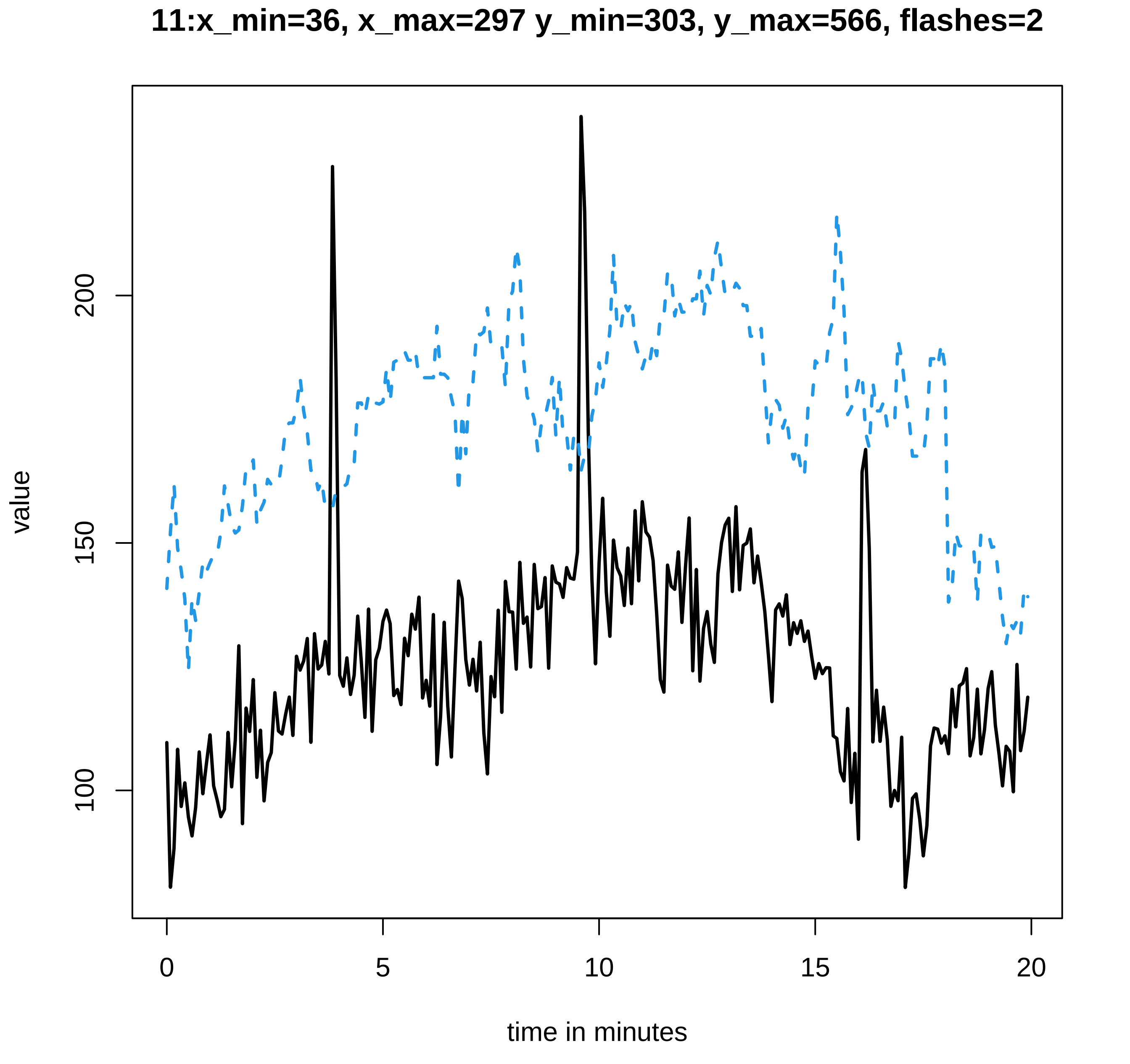

11:x_min=36, x_max=297 y_min=303, y_max=566, flashes=2
value
100
150
200
0
5
10
15
20
time in minutes

### 12:x_min=34, x_max=279 y_min=286, y_max=588, flashes=6

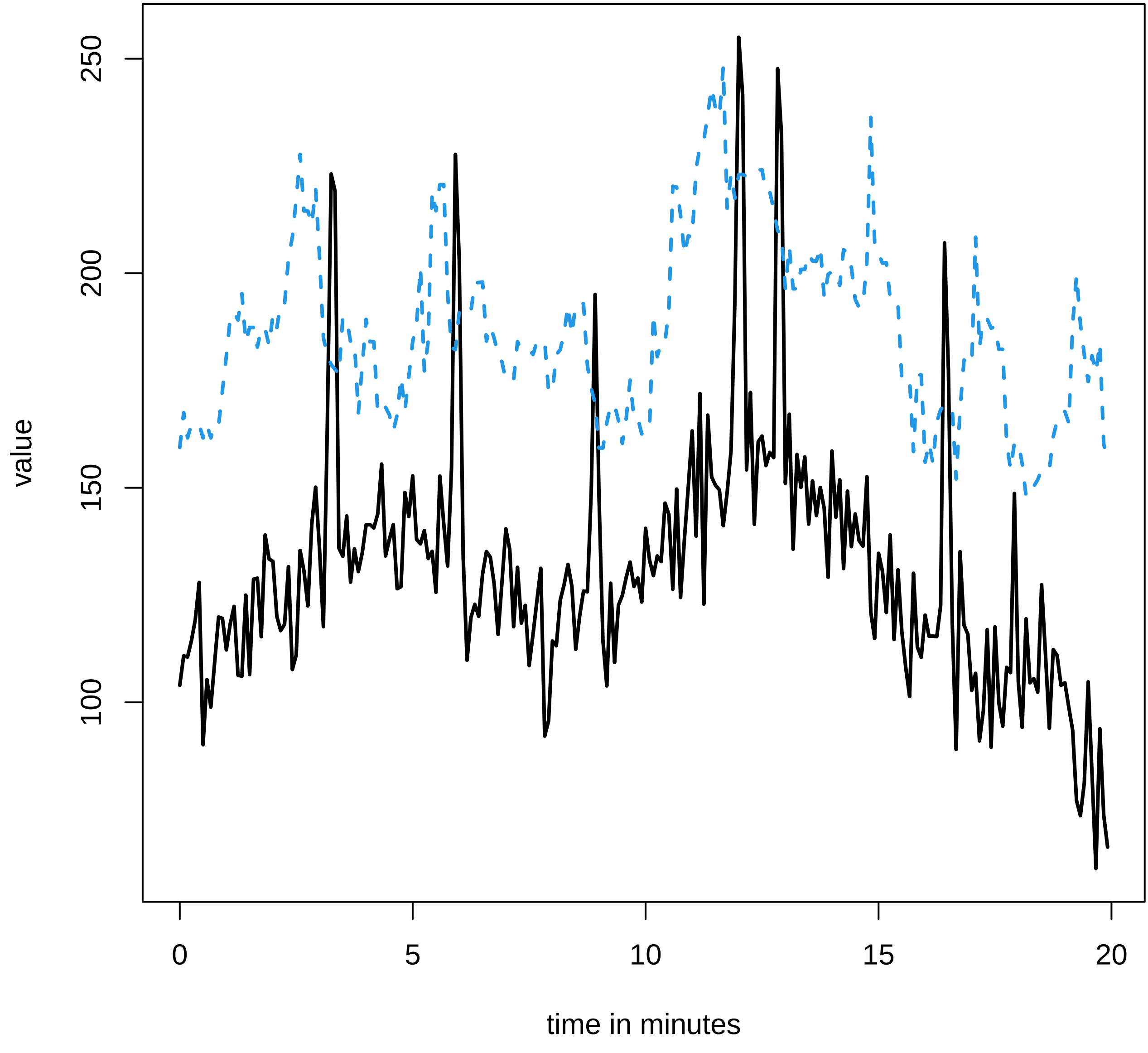

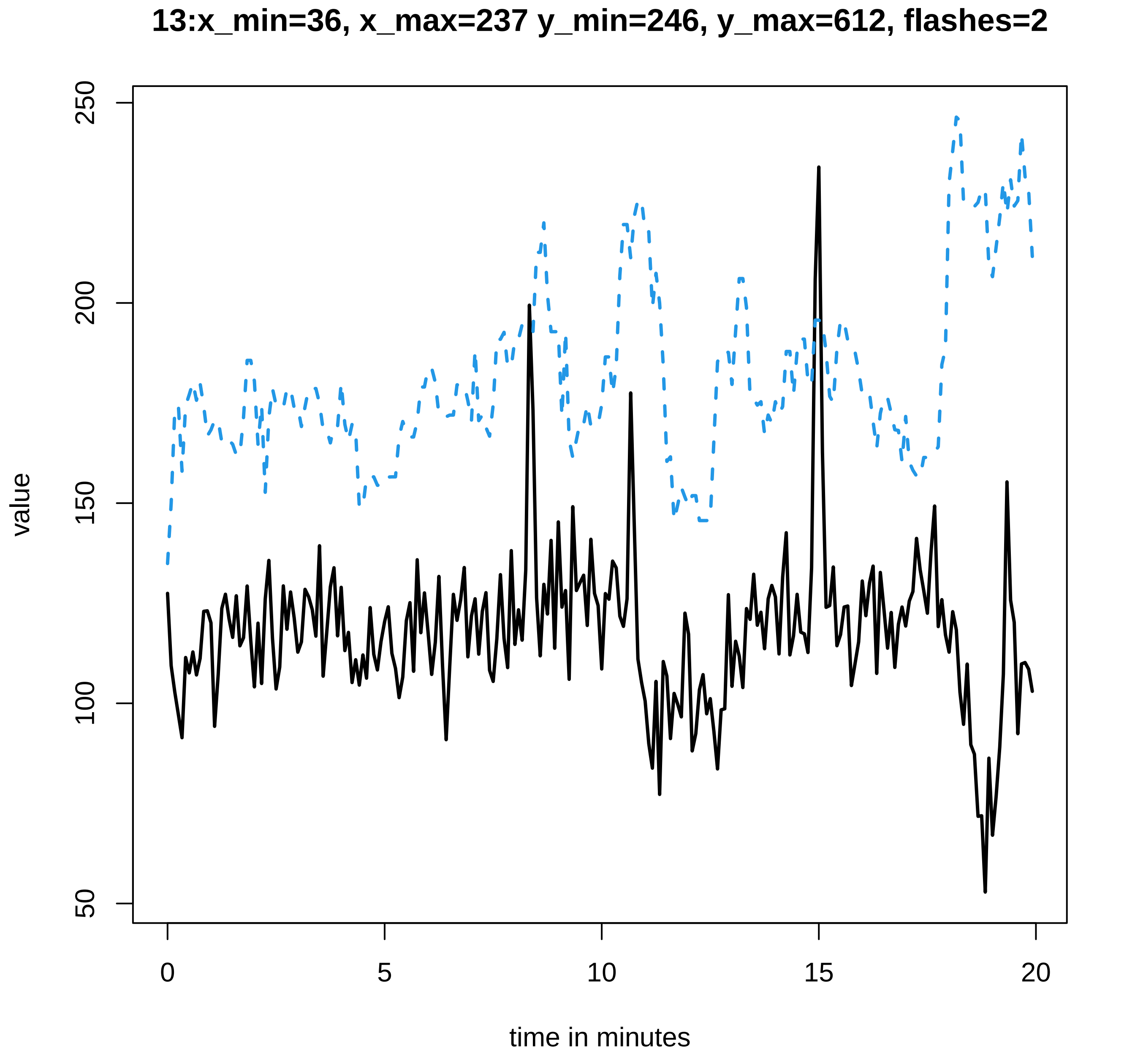
13:x_min=36, x_max=237 y_min=246, y_max=612, flashes=2
value
250
200
150
100
50
0
5
10
15
20
time in minutes

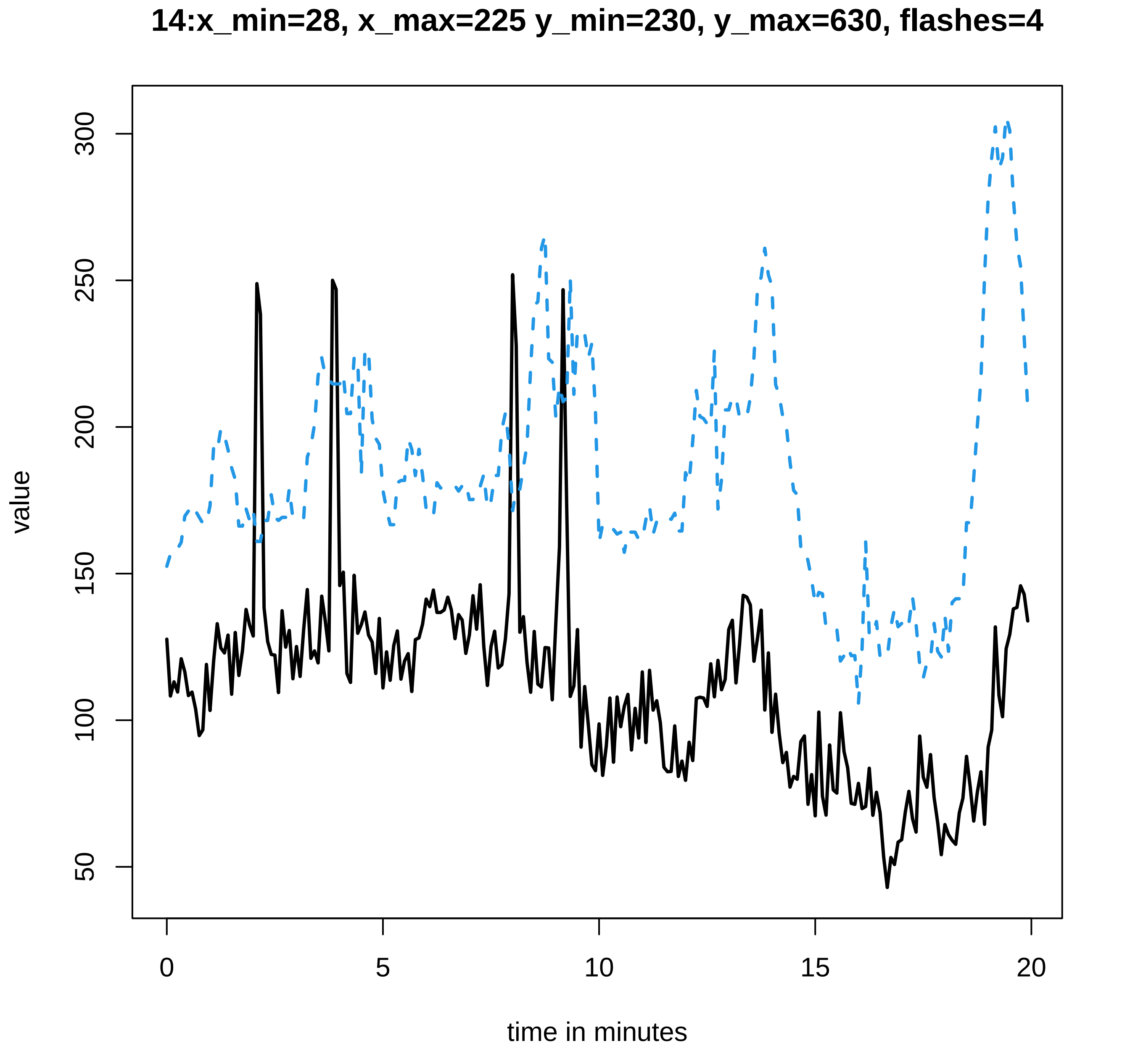
14:x_min=28, x_max=225 y_min=230, y_max=630, flashes=4
value
300
250
200
150
100
50
0
5
10
15
20
time in minutes

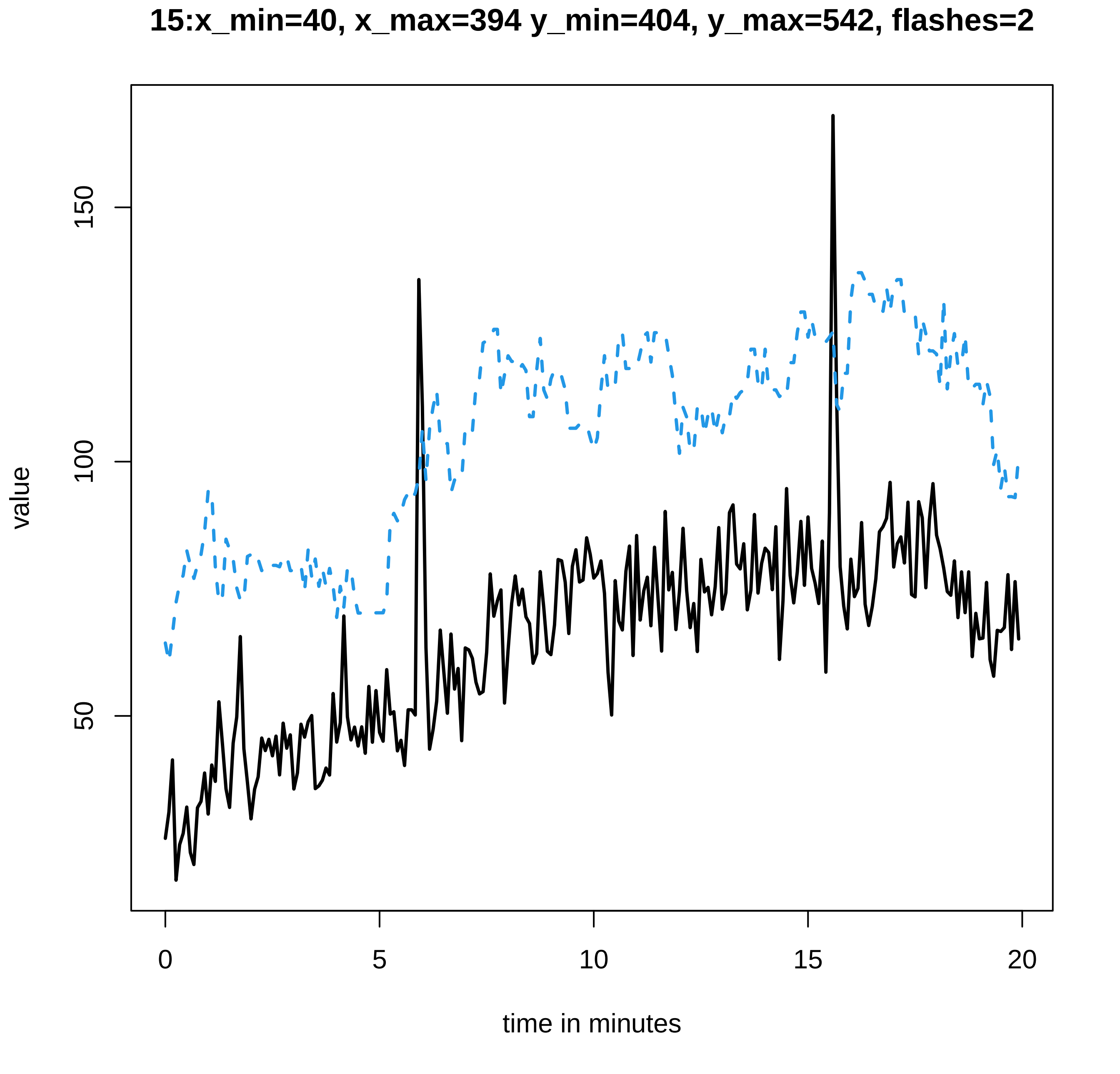

15:x_min=40, x_max=394 y_min=404, y_max=542, flashes=2
value
150
100
50
0
5
10
15
20
time in minutes

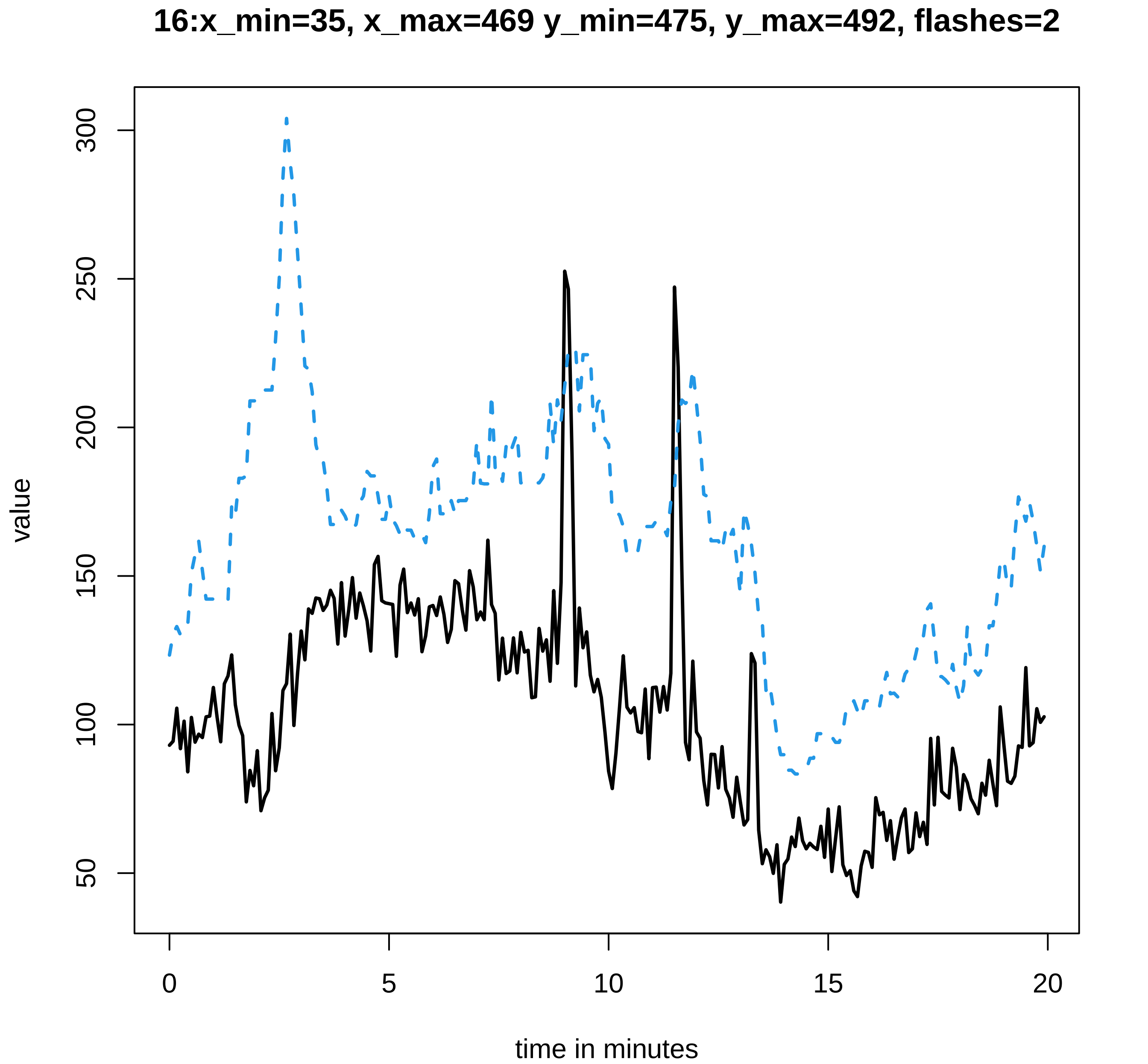
16:x_min=35, x_max=469 y_min=475, y_max=492, flashes=2
value
50
100
150
200
250
300
0
5
10
15
20
time in minutes

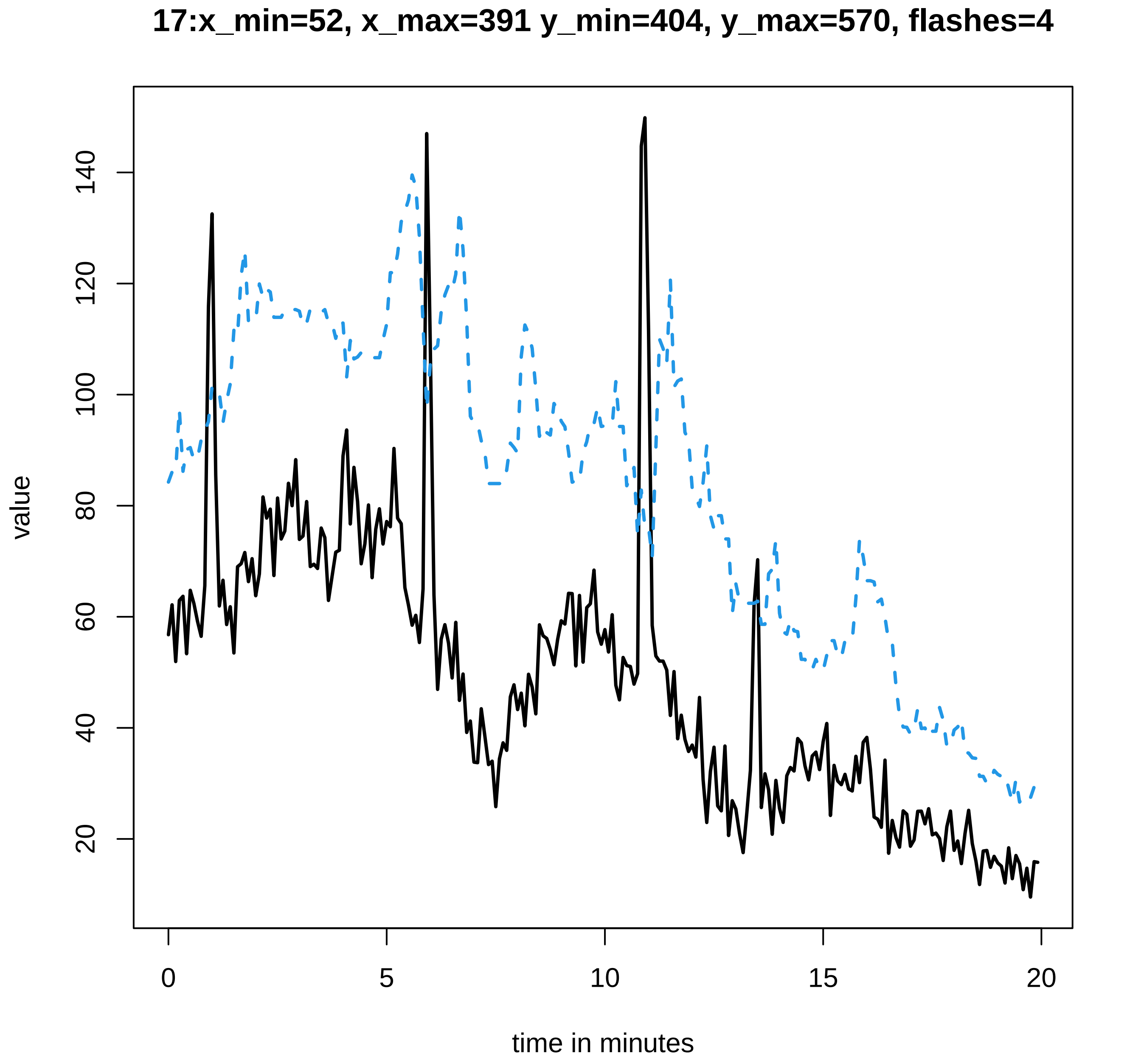

17:x_min=52, x_max=391 y_min=404, y_max=570, flashes=4
value
20
40
60
80
100
120
140
0
5
10
15
20
time in minutes

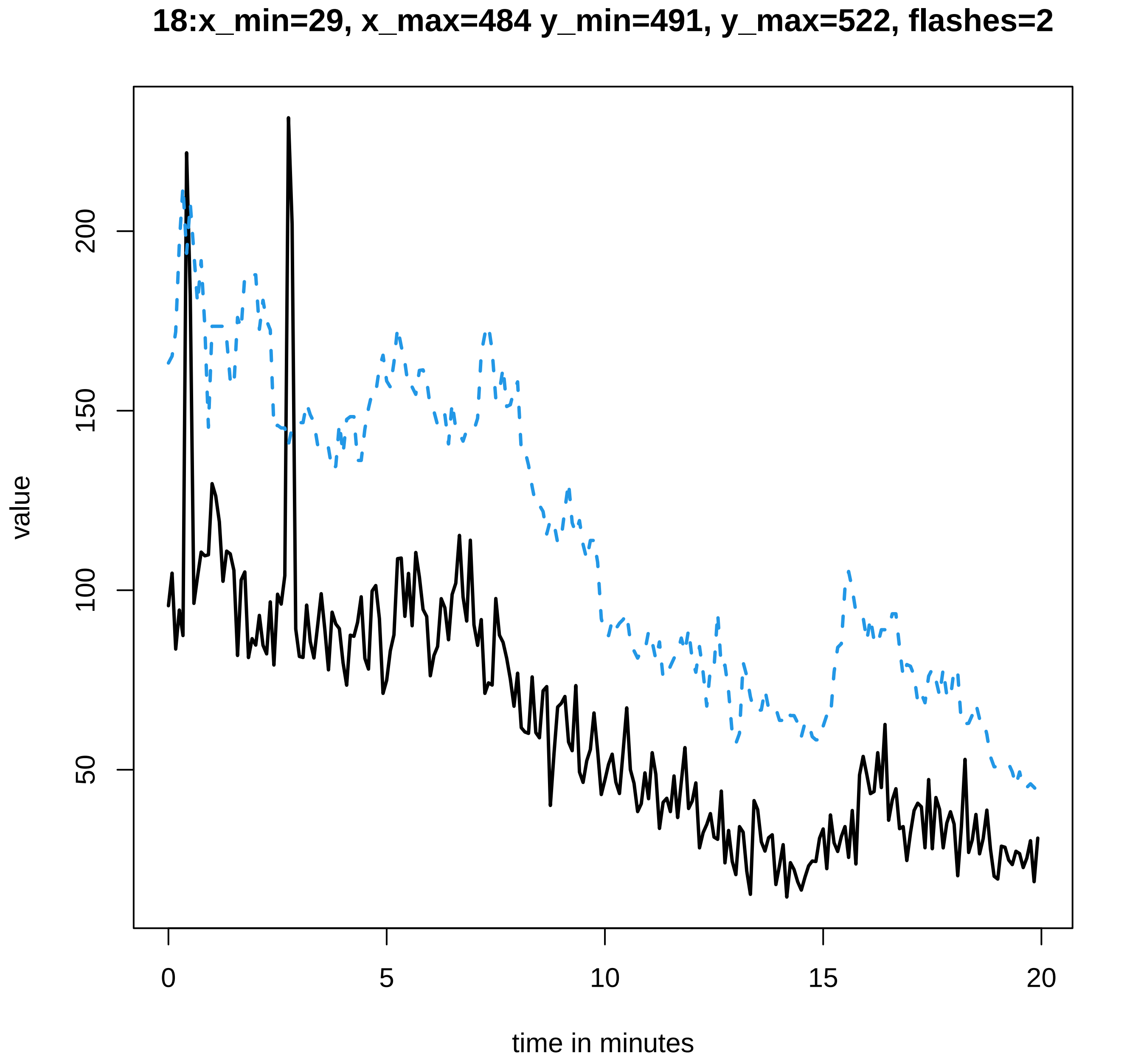

18:x_min=29, x_max=484 y_min=491, y_max=522, flashes=2
value
200
150
100
50
0
5
10
15
20
time in minutes

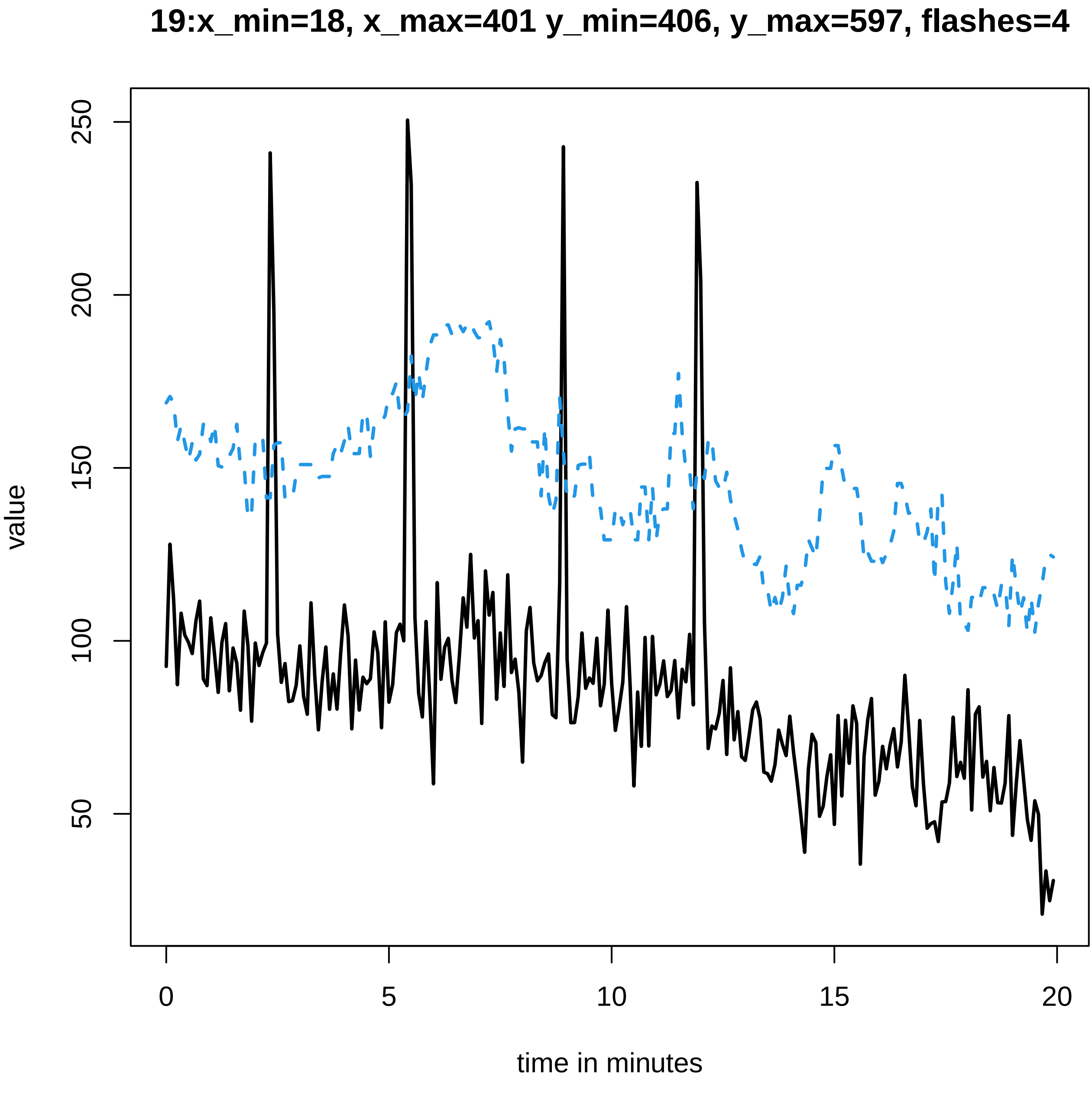
19:x_min=18, x_max=401 y_min=406, y_max=597, flashes=4
value
250
200
150
100
50
0
5
10
15
20
time in minutes

### 20:x_min=24, x_max=528 y_min=533, y_max=499, flashes=2

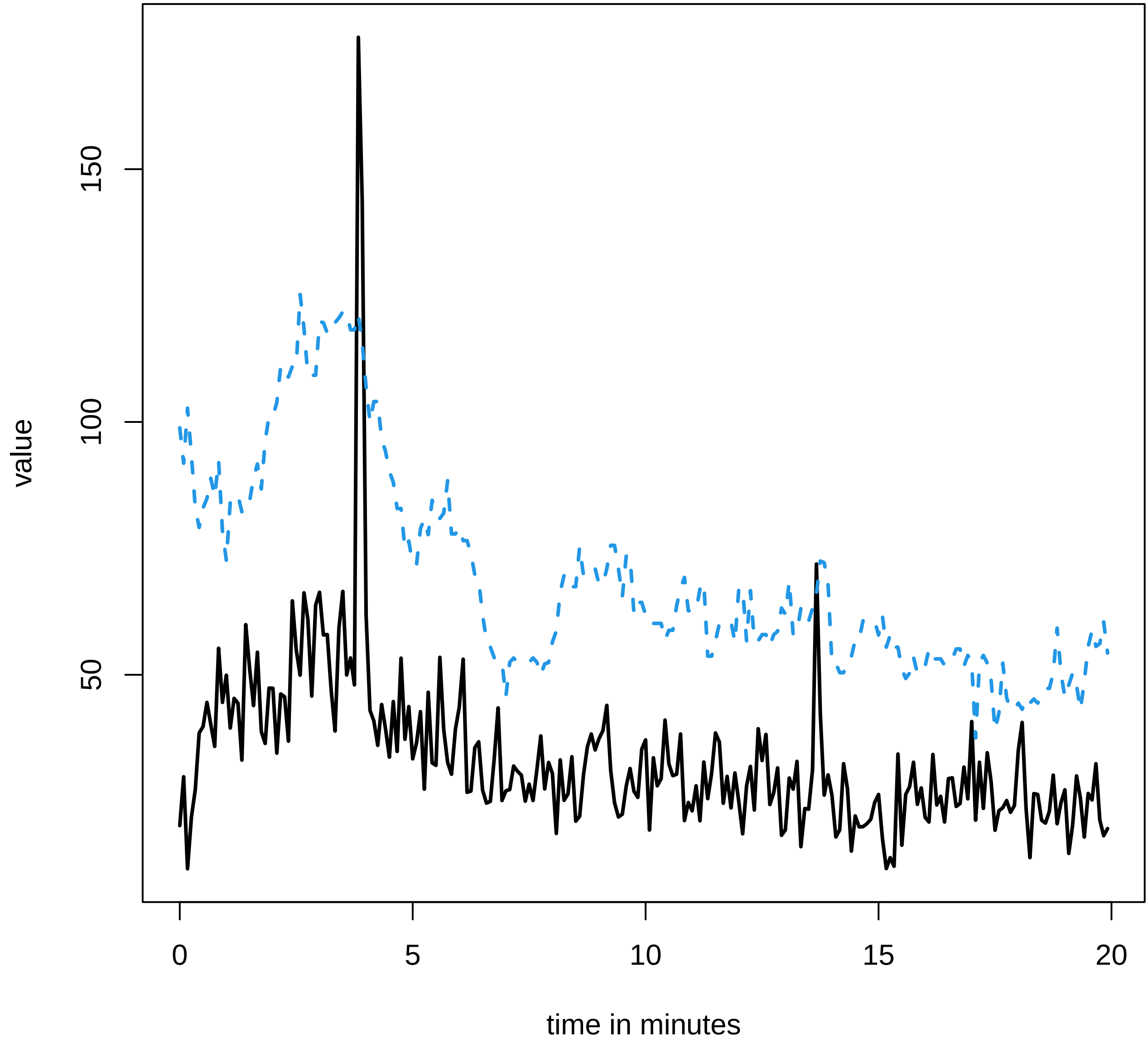

21:x_min=42, x_max=385 y_min=393, y_max=614, flashes=3

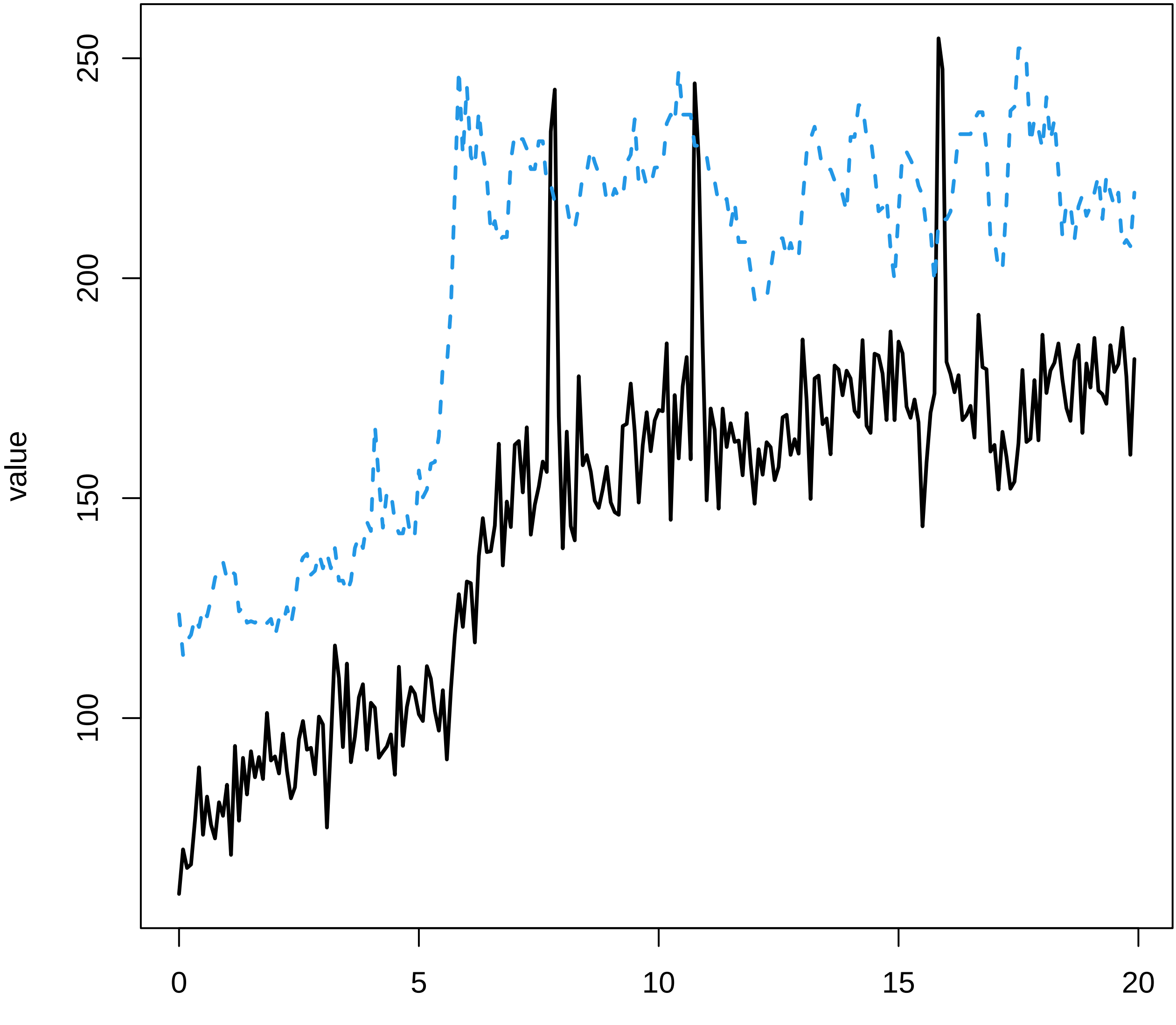

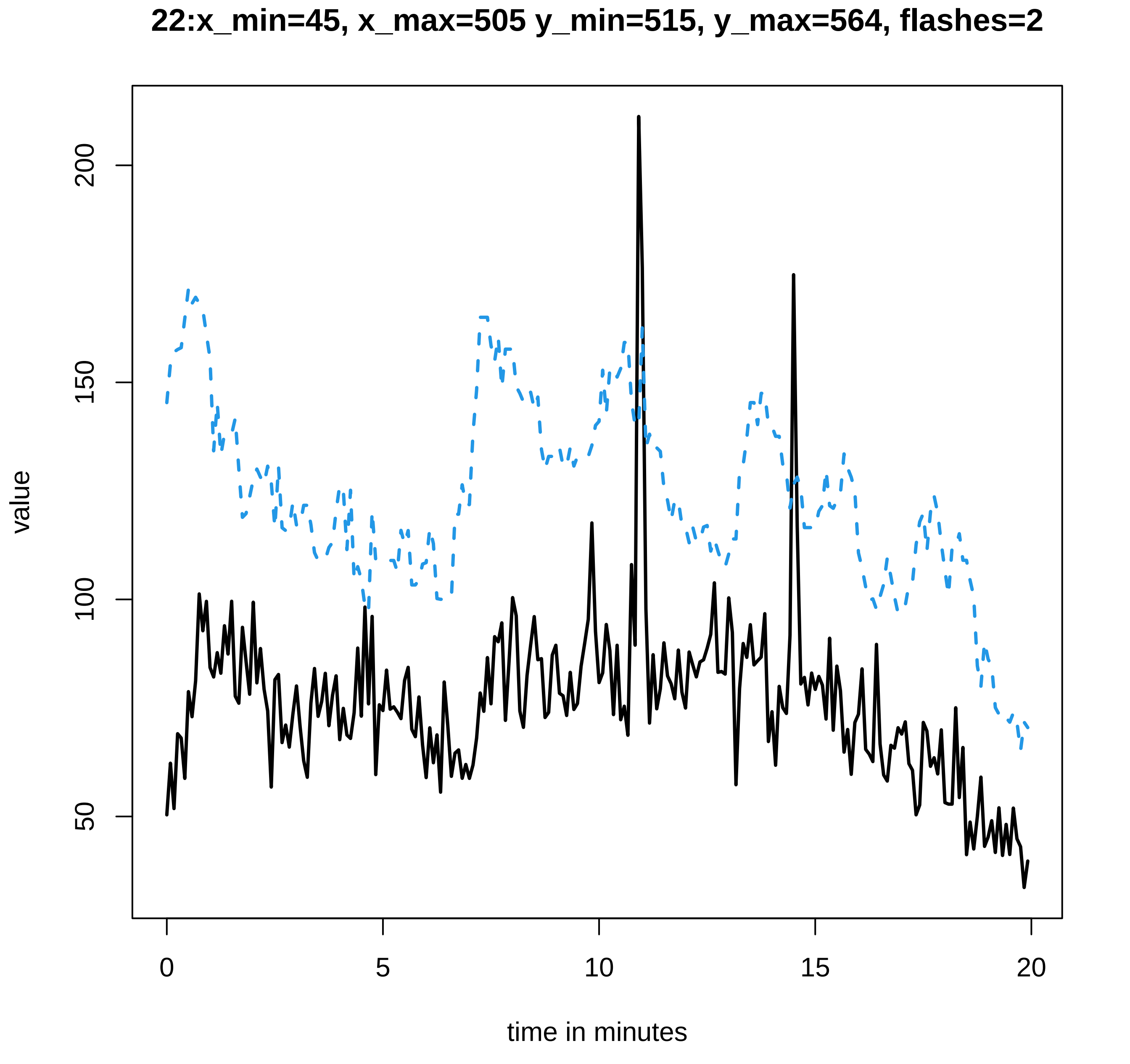
22:x_min=45, x_max=505 y_min=515, y_max=564, flashes=2
value
200
150
100
50
0
5
10
15
20
time in minutes

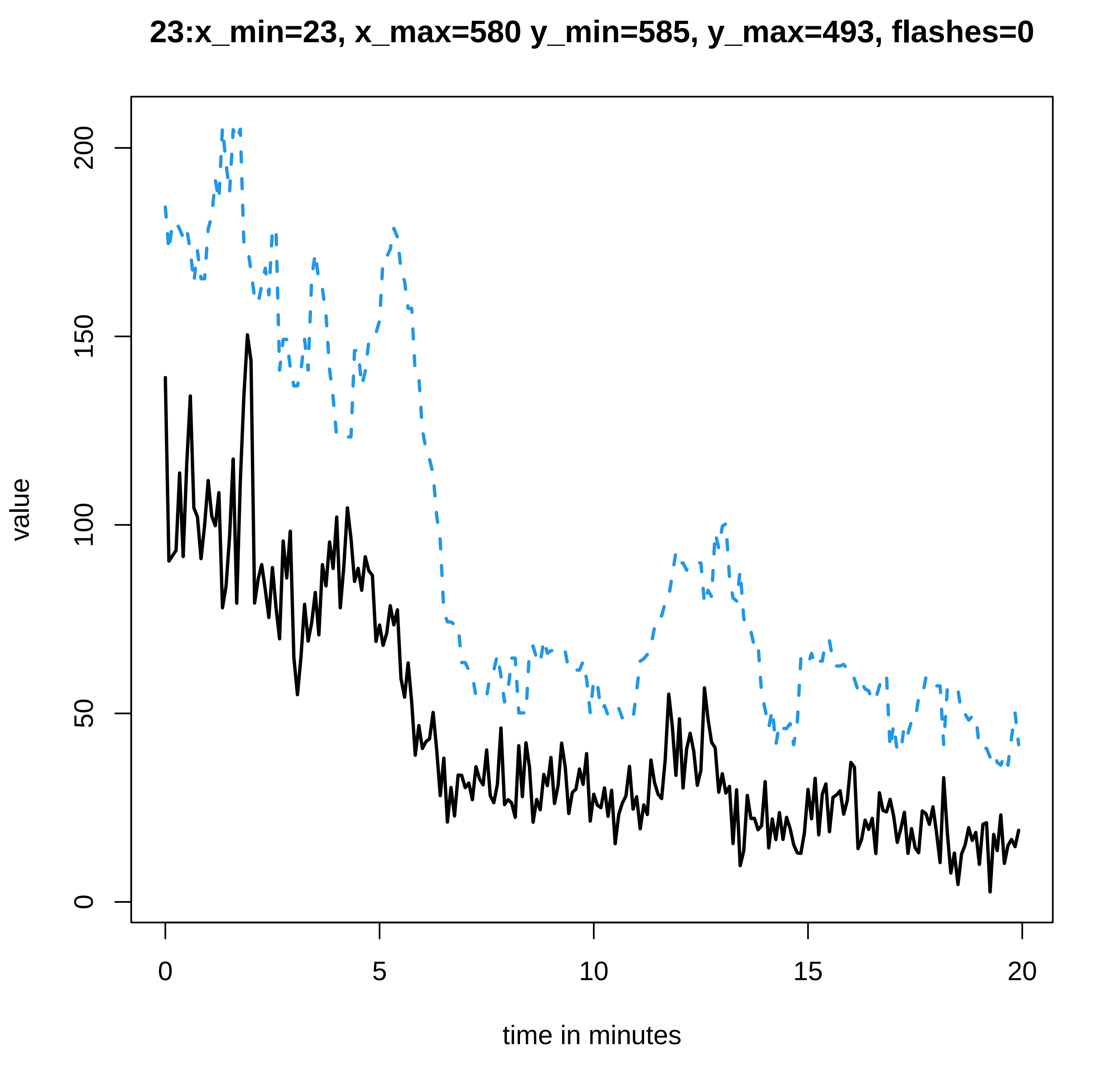
23:x_min=23, x_max=580 y_min=585, y_max=493, flashes=0
value
0
50
100
150
200
0
5
10
15
20
time in minutes

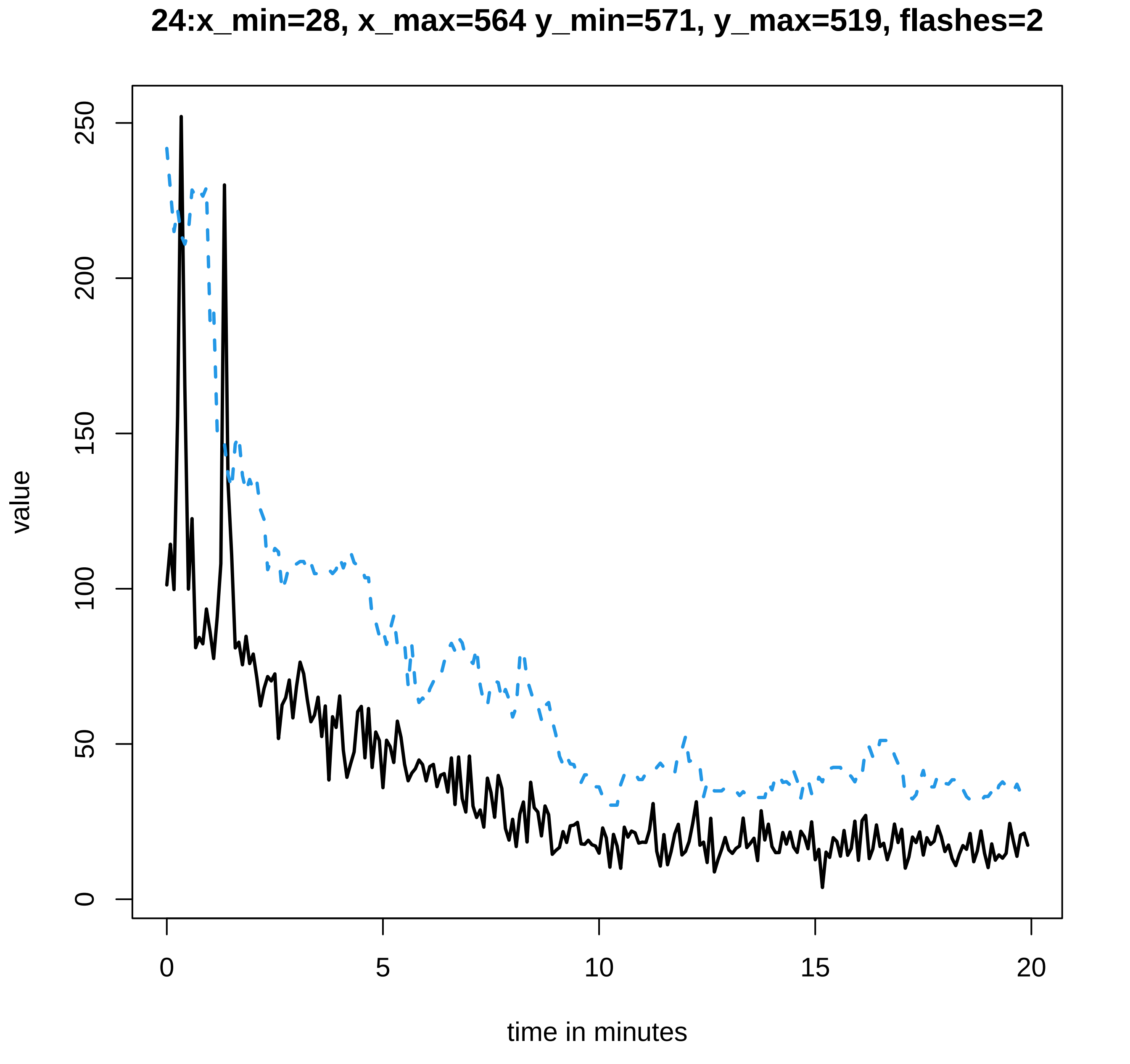
24:x_min=28, x_max=564 y_min=571, y_max=519, flashes=2
value
0
50
100
150
200
250
0
5
10
15
20
time in minutes

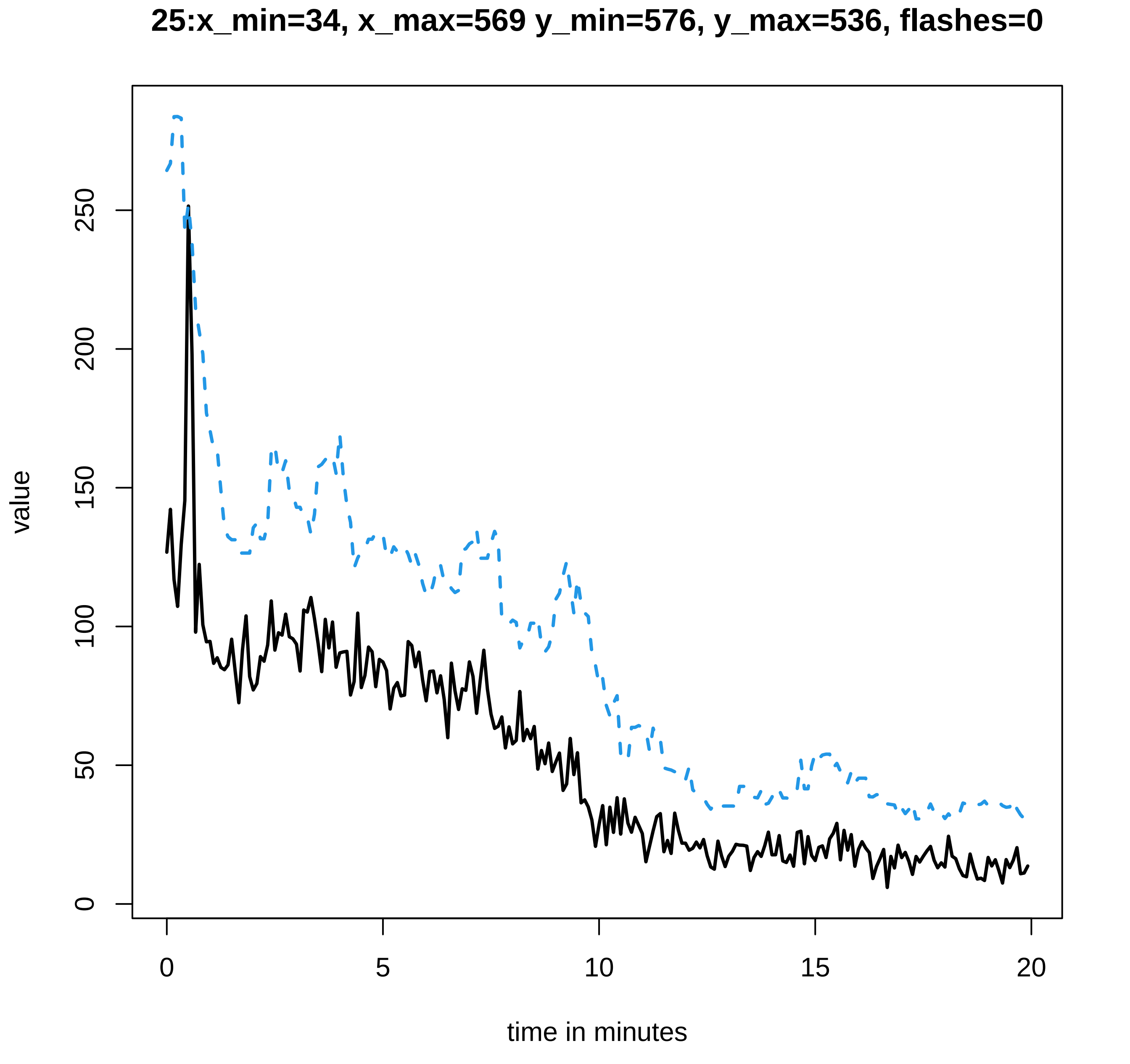

25:x_min=34, x_max=569 y_min=576, y_max=536, flashes=0
value
0
50
100
150
200
250
0
5
10
15
20
time in minutes

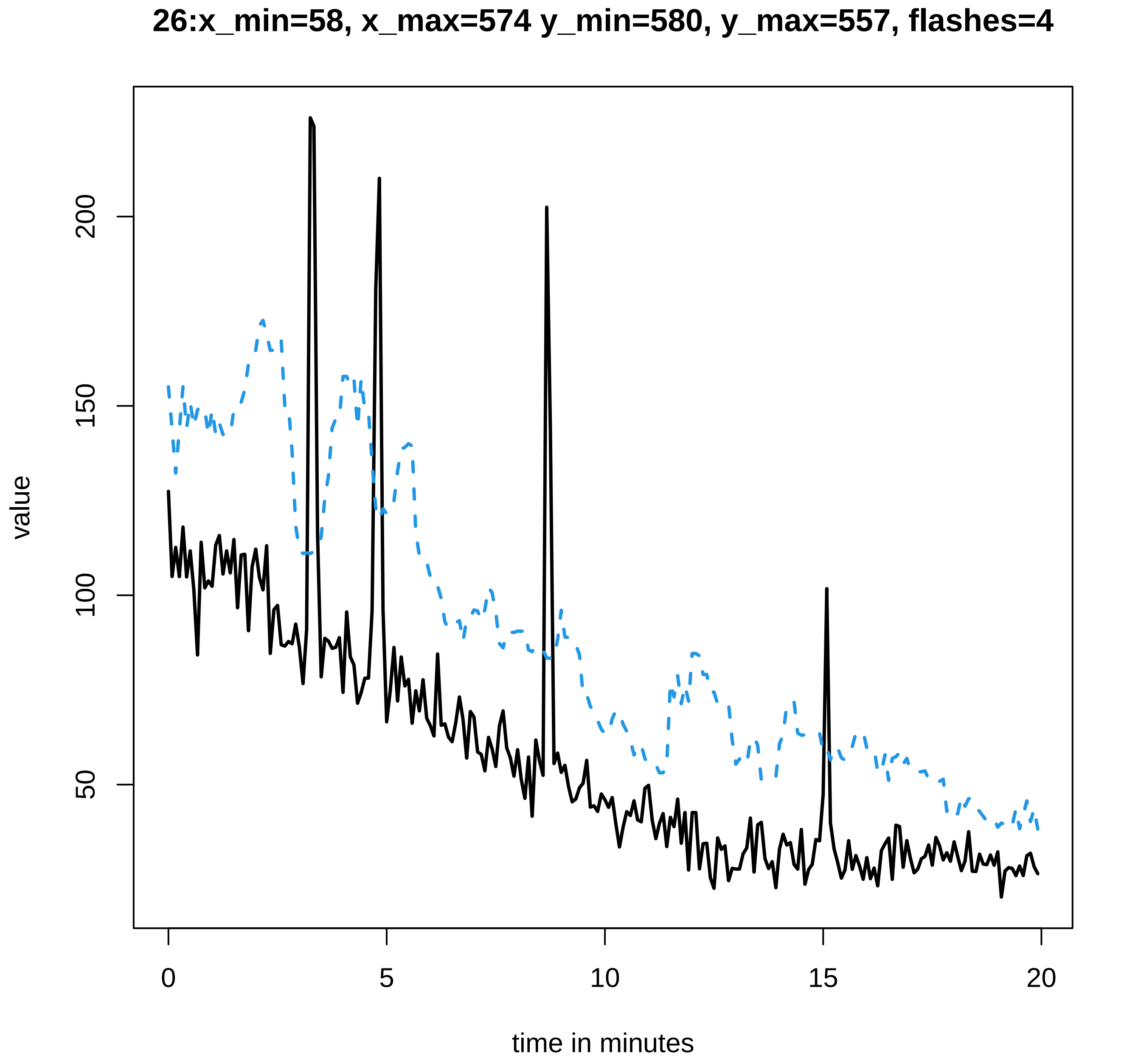
26:x_min=58, x_max=574 y_min=580, y_max=557, flashes=4
value
200
150
100
50
0
5
10
15
20
time in minutes

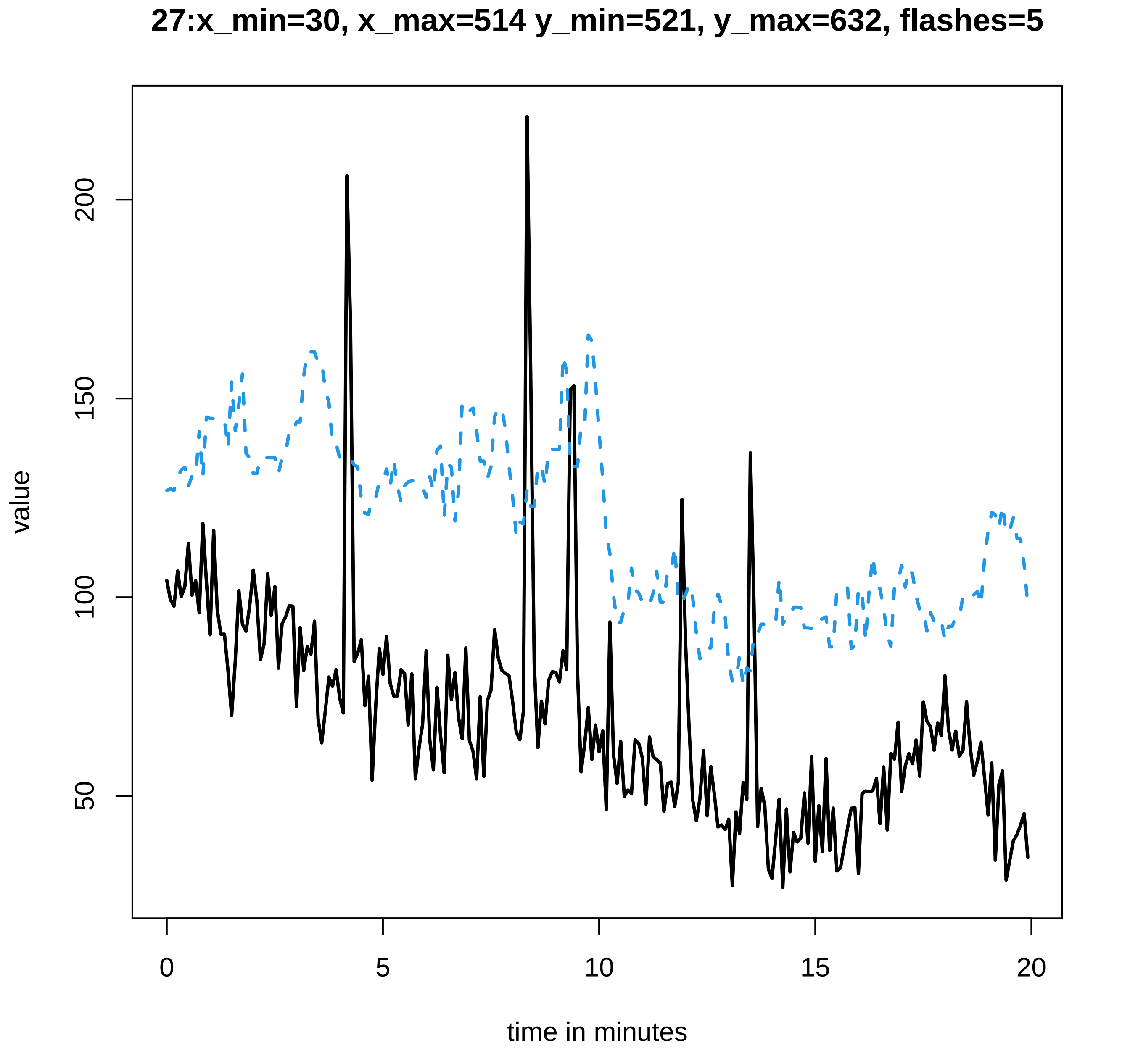
27:x_min=30, x_max=514 y_min=521, y_max=632, flashes=5
value
200
150
100
50
0
5
10
15
20
time in minutes

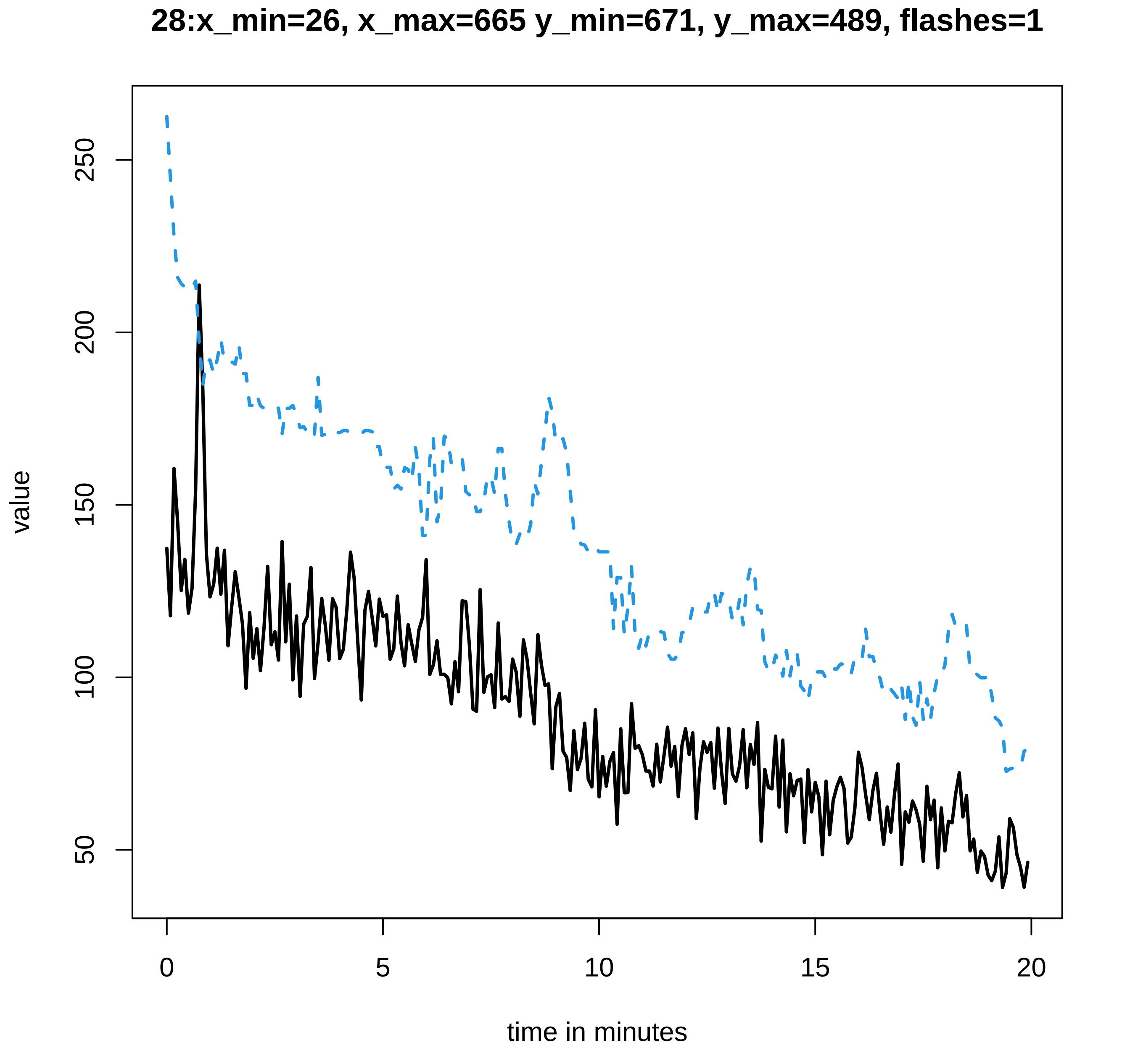
28:x_min=26, x_max=665 y_min=671, y_max=489, flashes=1
value
250
200
150
100
50
0
5
10
15
20
time in minutes

#### 29:x_min=50, x_max=564 y_min=571, y_max=602, flashes=7

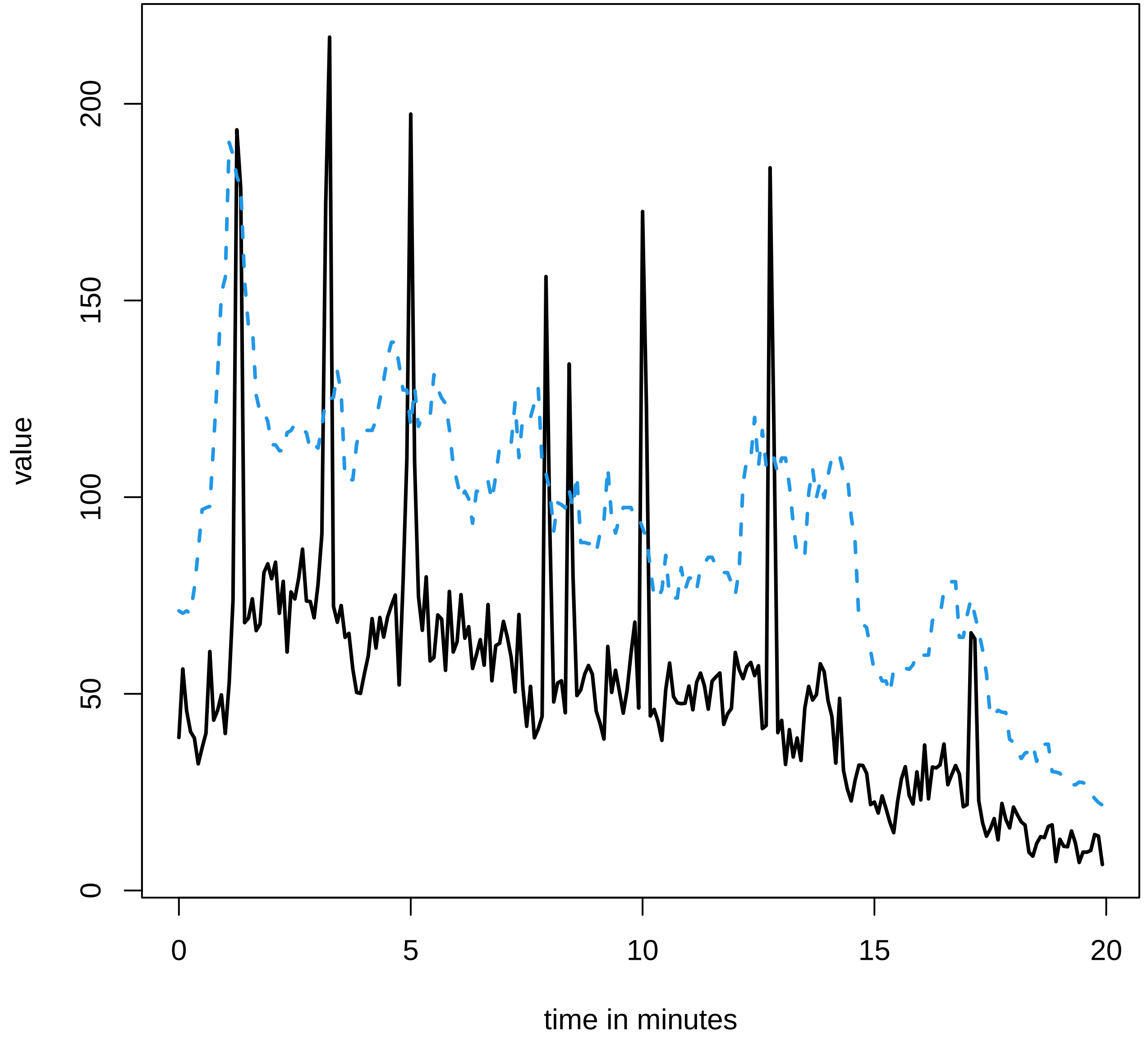

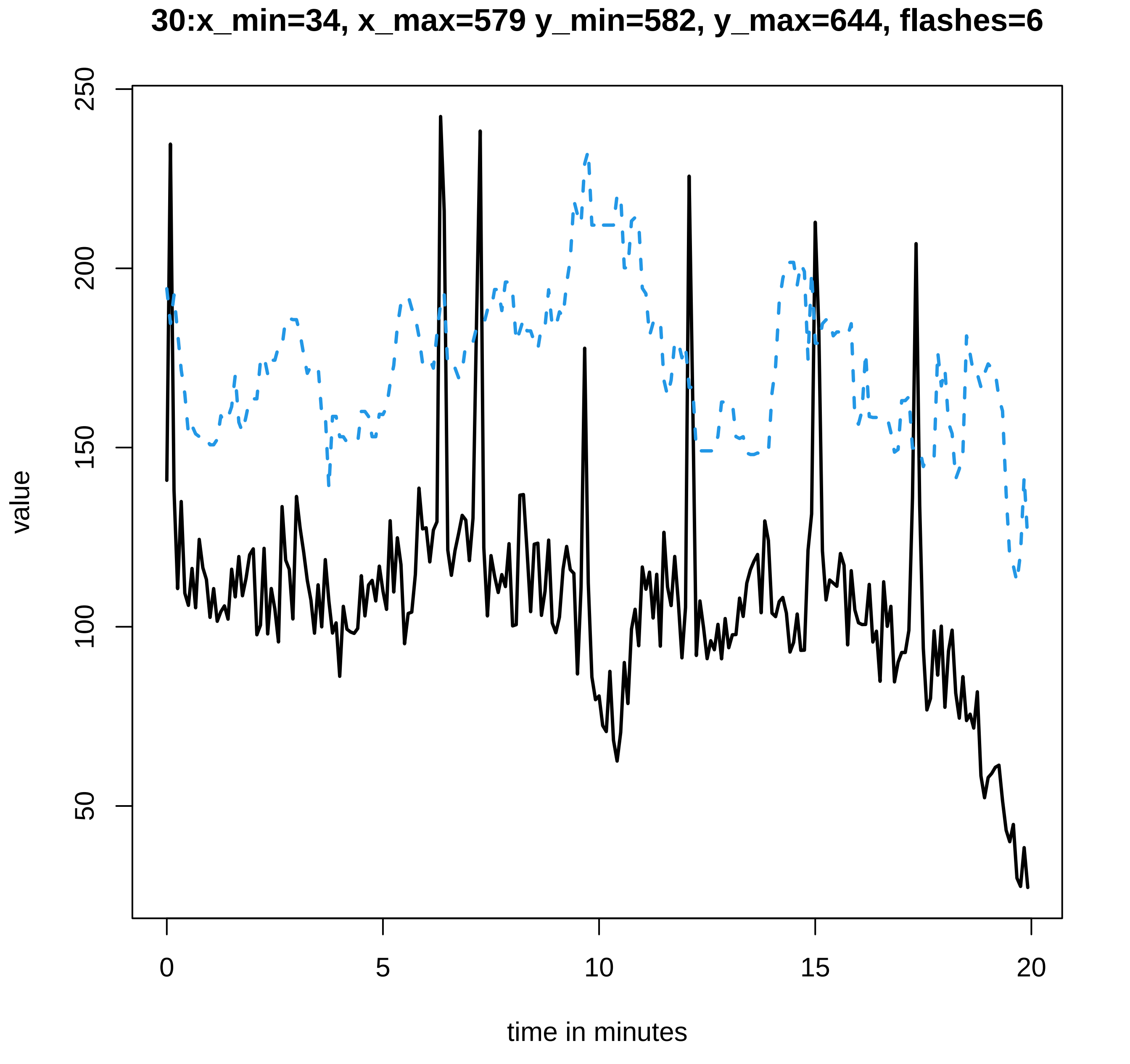

30:x_min=34, x_max=579 y_min=582, y_max=644, flashes=6
value
250
200
150
100
50
0
5
10
15
20
time in minutes

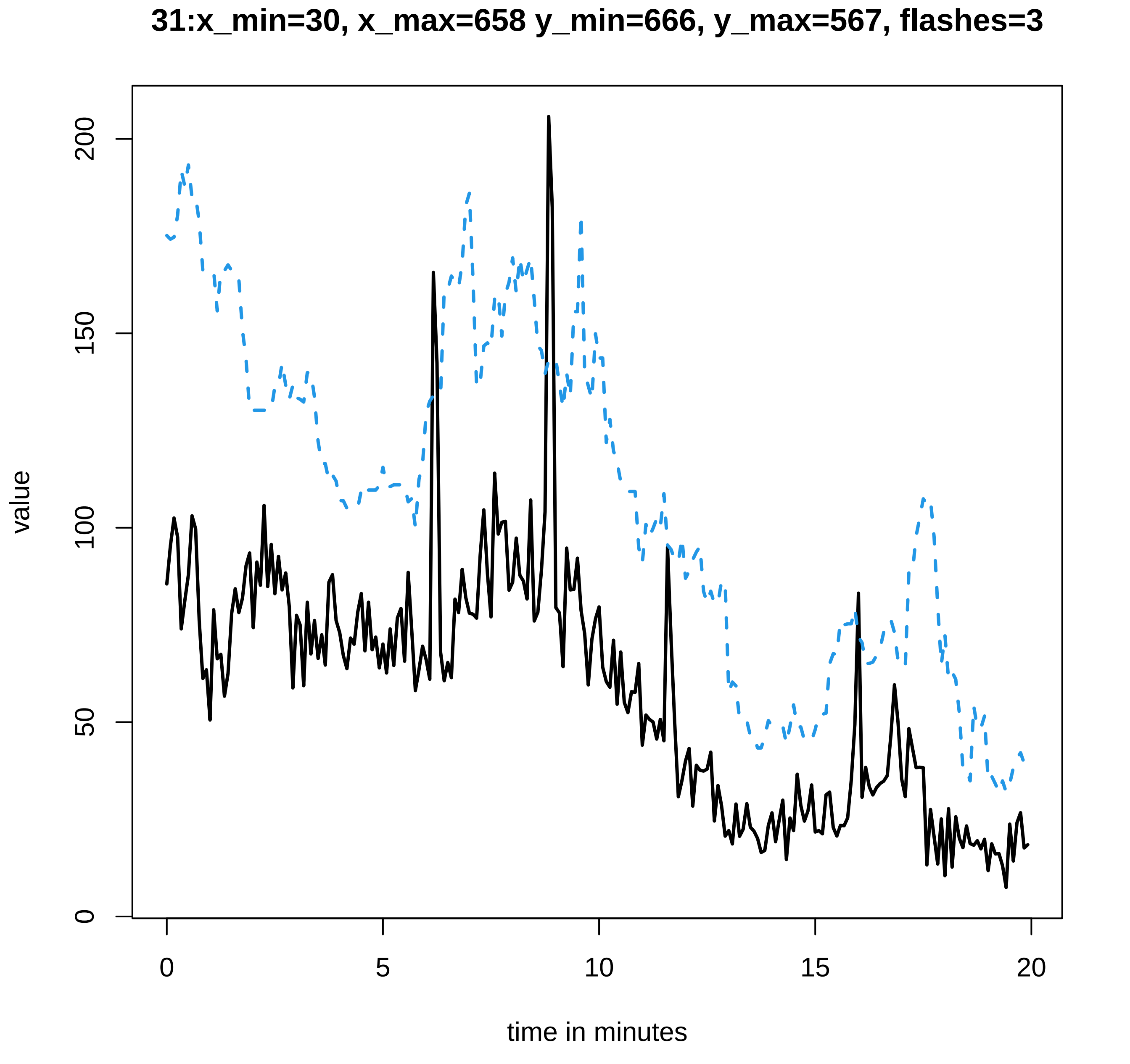

31:x_min=30, x_max=658 y_min=666, y_max=567, flashes=3
value
0
50
100
150
200
0
5
10
15
20
time in minutes

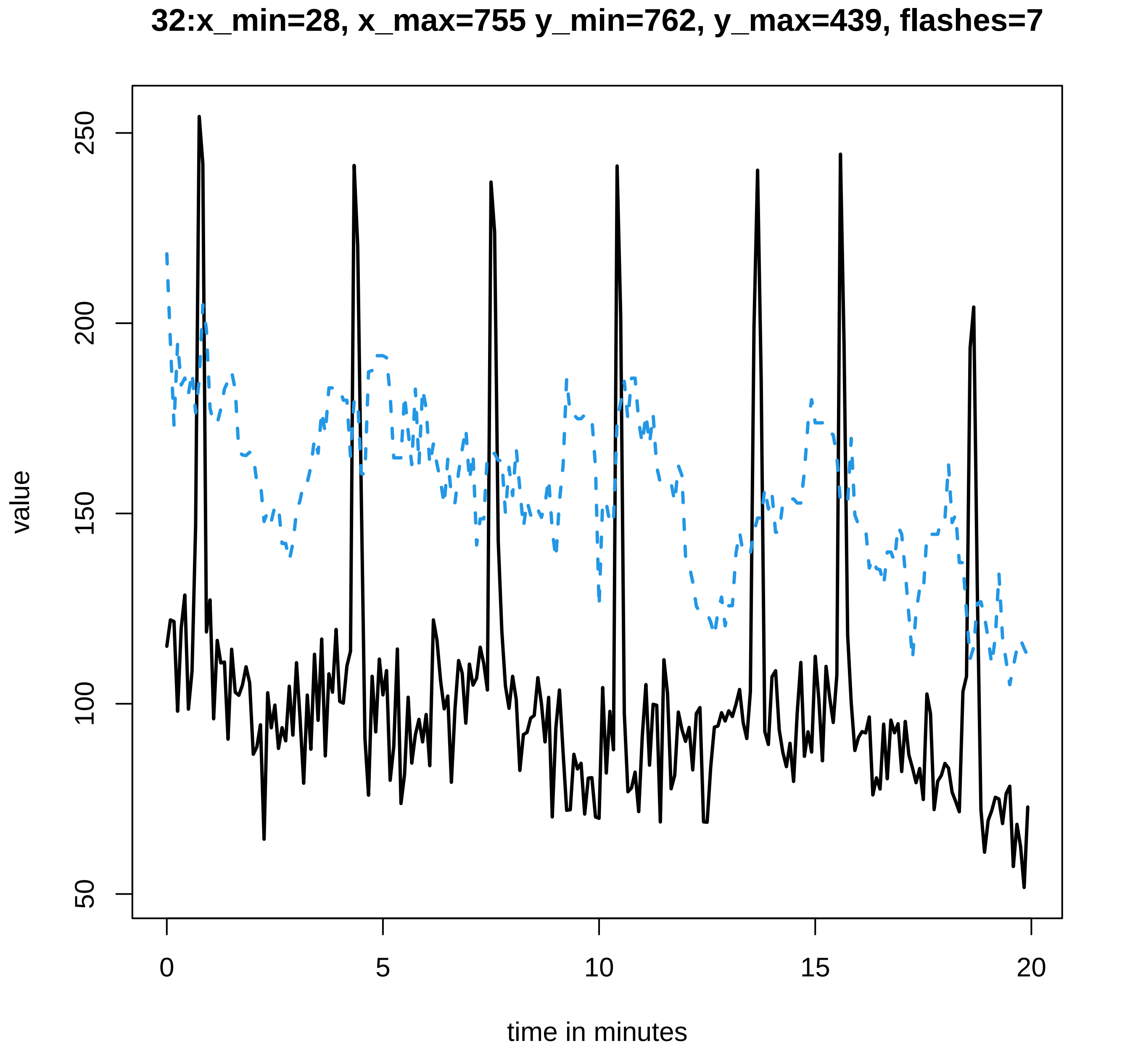

32:x_min=28, x_max=755 y_min=762, y_max=439, flashes=7
value
250
200
150
100
50
0
5
10
15
20
time in minutes

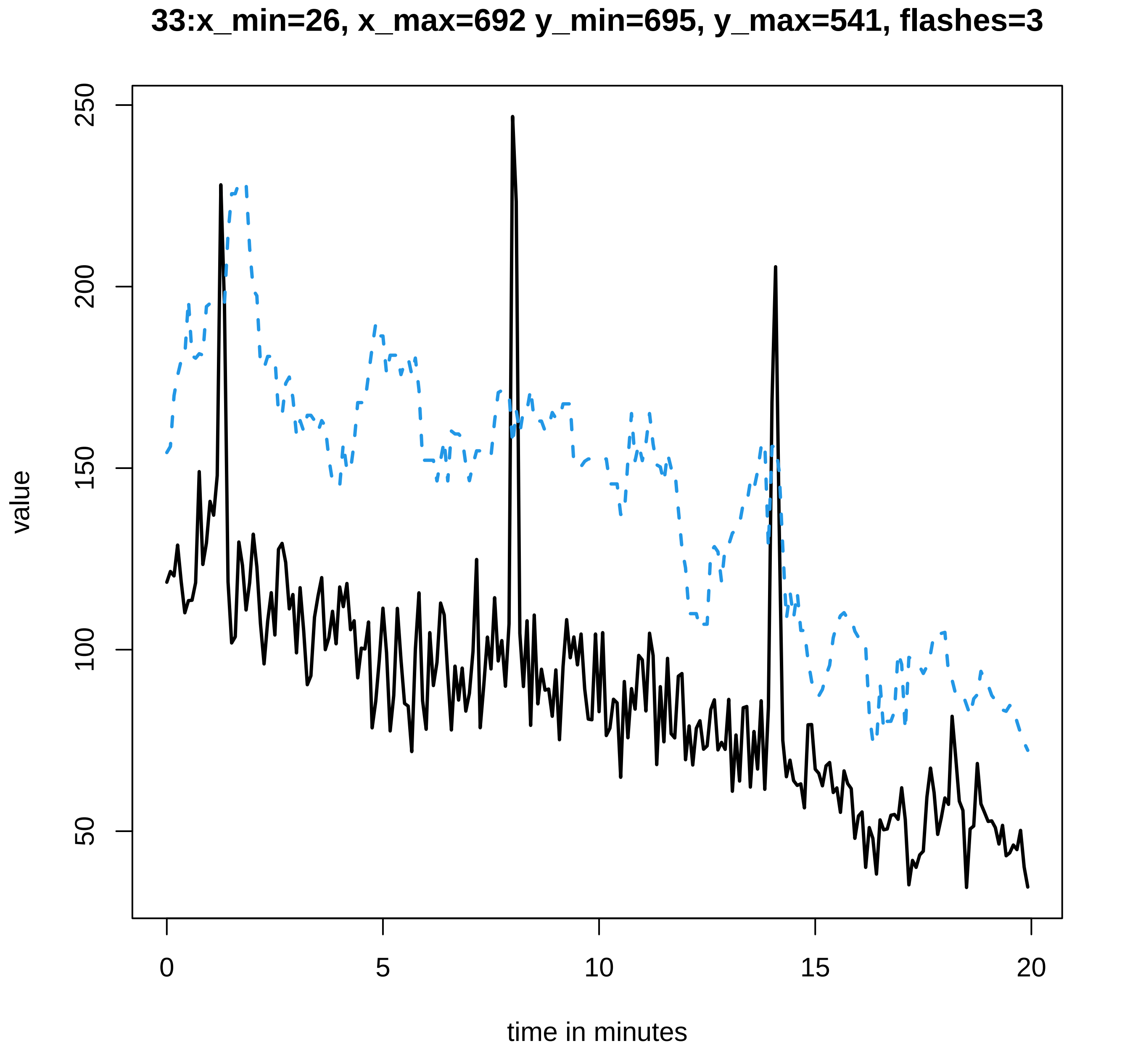
33:x_min=26, x_max=692 y_min=695, y_max=541, flashes=3
value
50
100
150
200
250
0
5
10
15
20
time in minutes

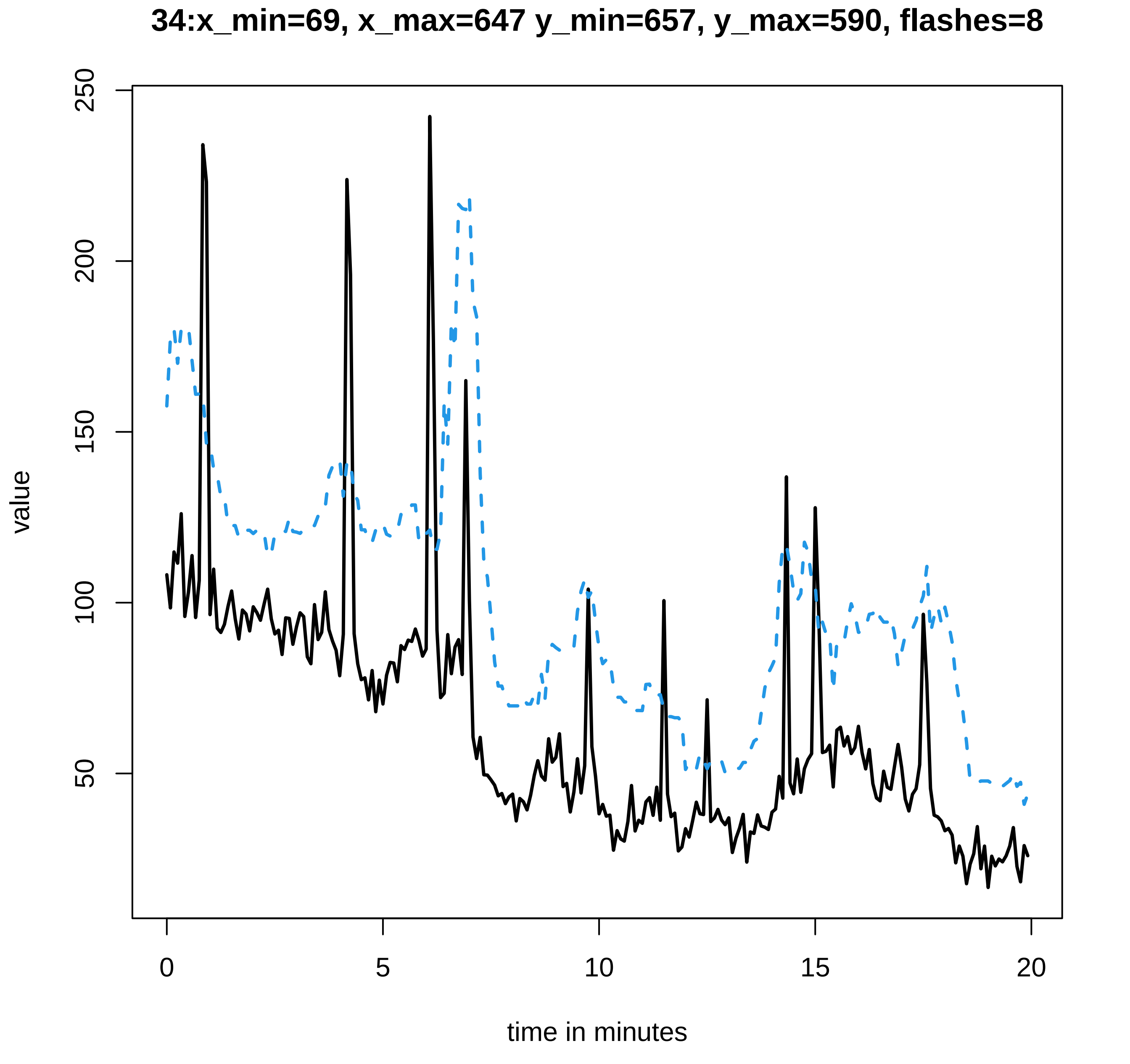
34:x_min=69, x_max=647 y_min=657, y_max=590, flashes=8
value
250
200
150
100
50
0
5
10
15
20
time in minutes

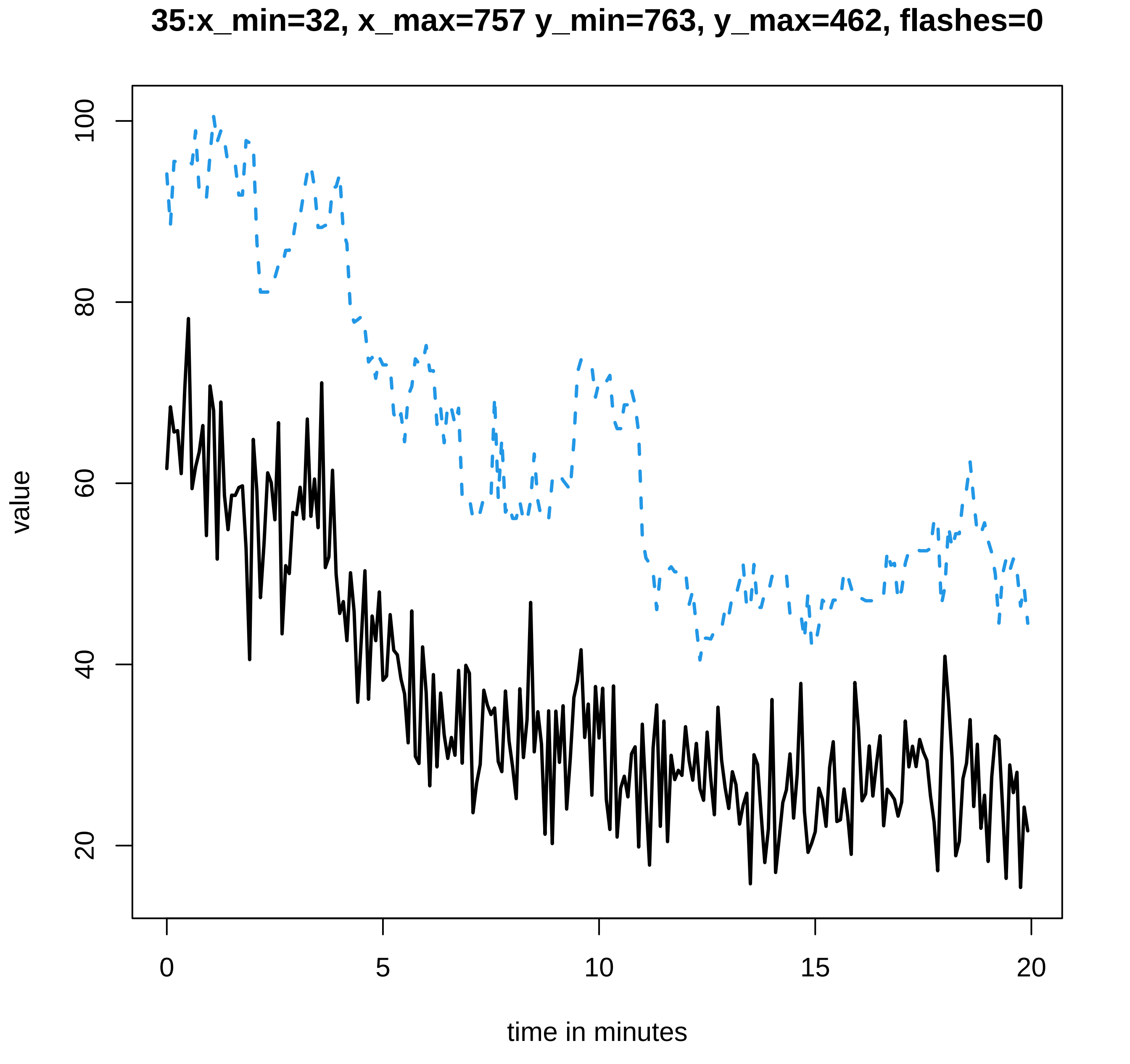
35:x_min=32, x_max=757 y_min=763, y_max=462, flashes=0
value
100
80
60
40
20
0
5
10
15
20
time in minutes

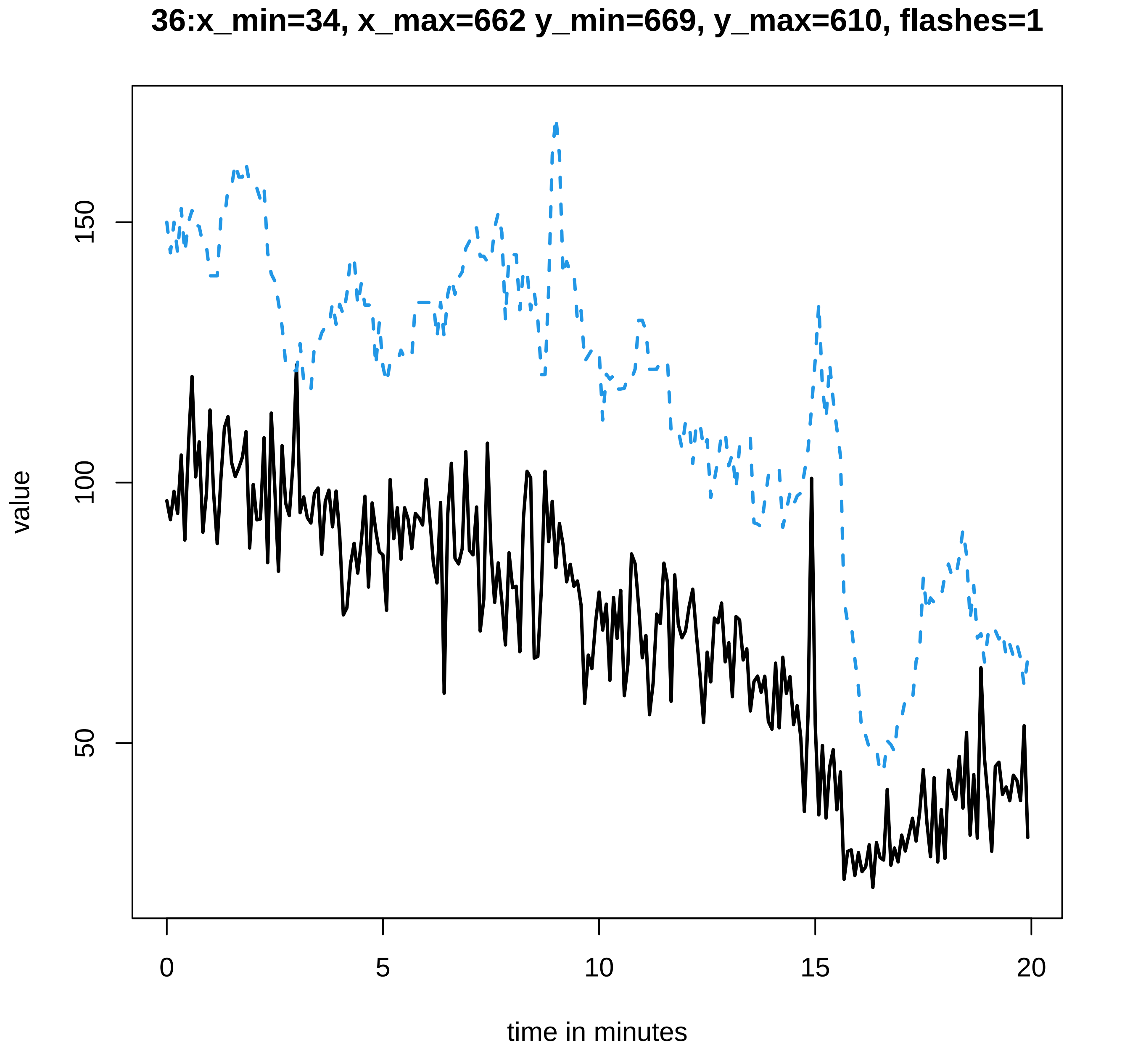
36:x_min=34, x_max=662 y_min=669, y_max=610, flashes=1
value
150
100
50
0
5
10
15
20
time in minutes

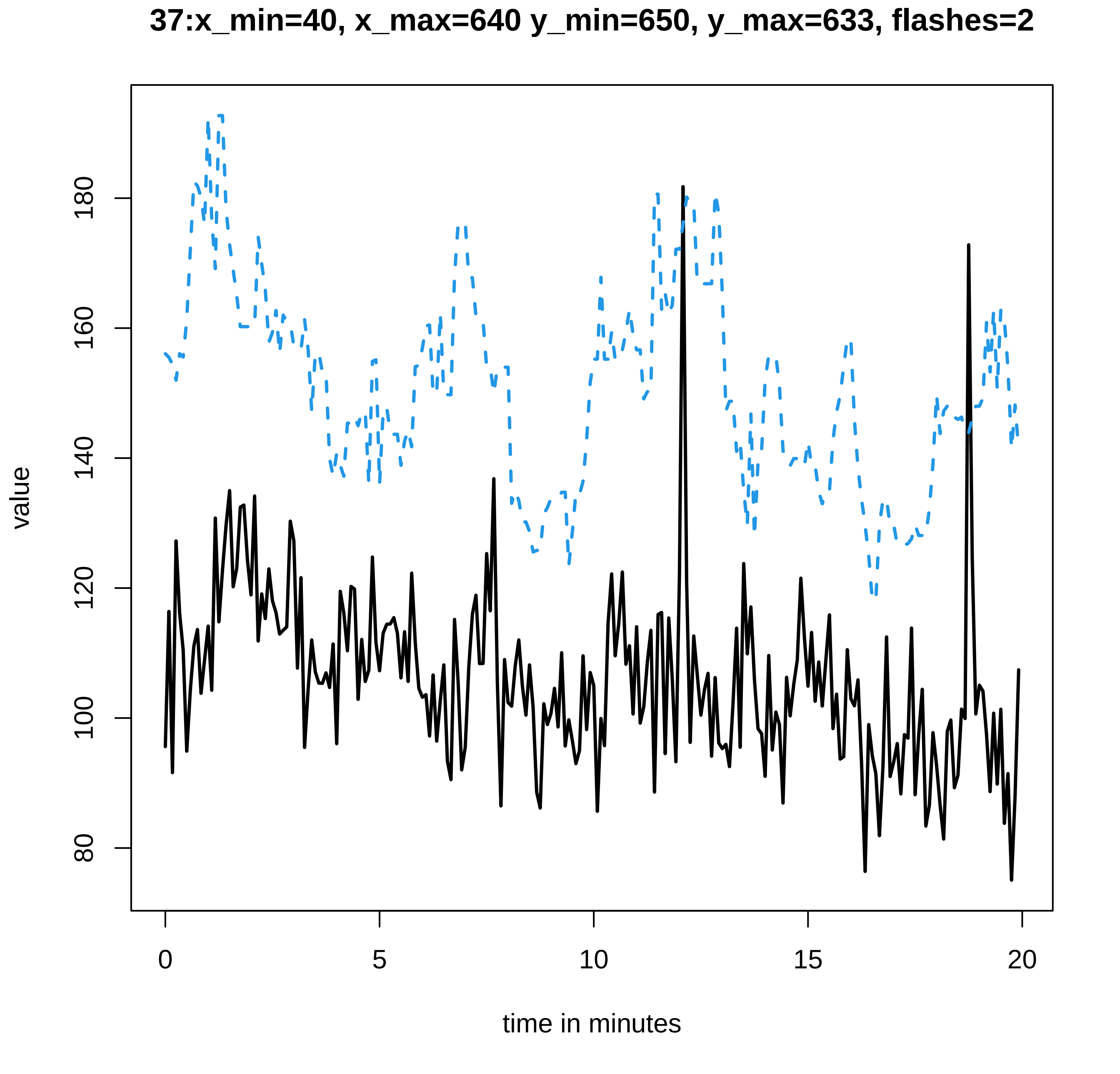

37:x_min=40, x_max=640 y_min=650, y_max=633, flashes=2
value
180
160
140
120
100
80
0
5
10
15
20
time in minutes

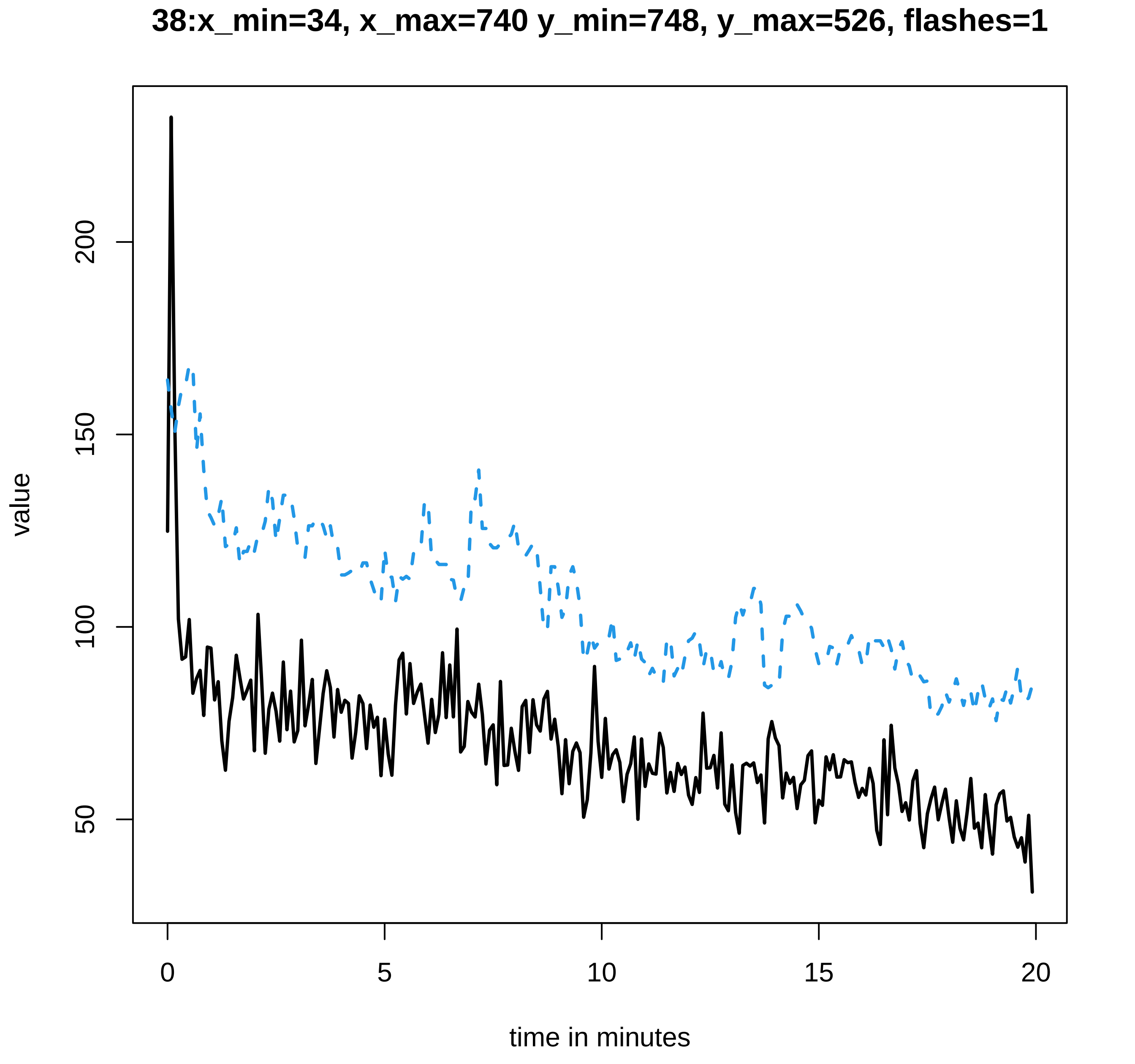

38:x_min=34, x_max=740 y_min=748, y_max=526, flashes=1
value
50
100
150
200
0
5
10
15
20
time in minutes

**39:x_min=28, x_max=700 y_min=706, y_max=587, flashes=6**

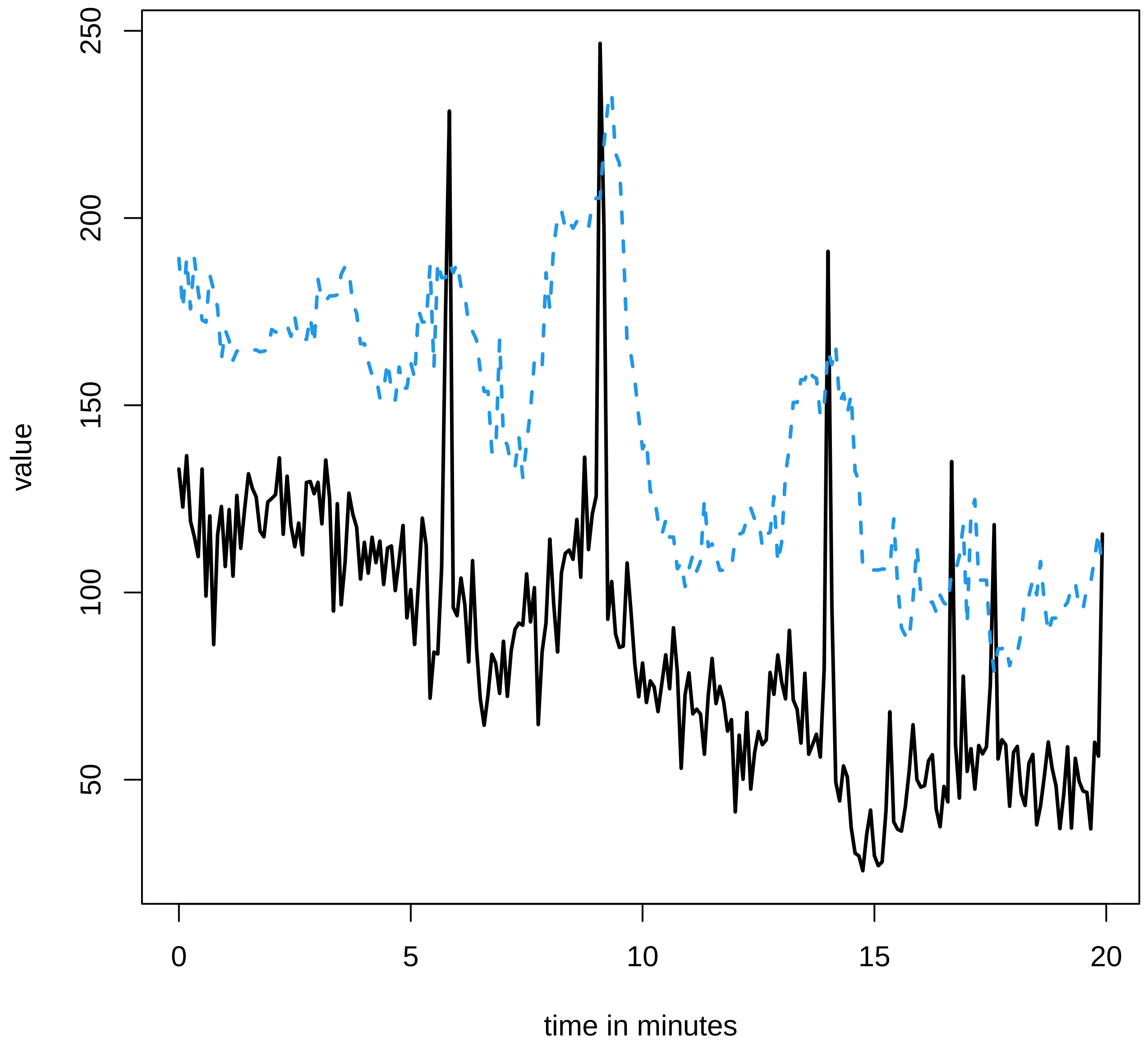

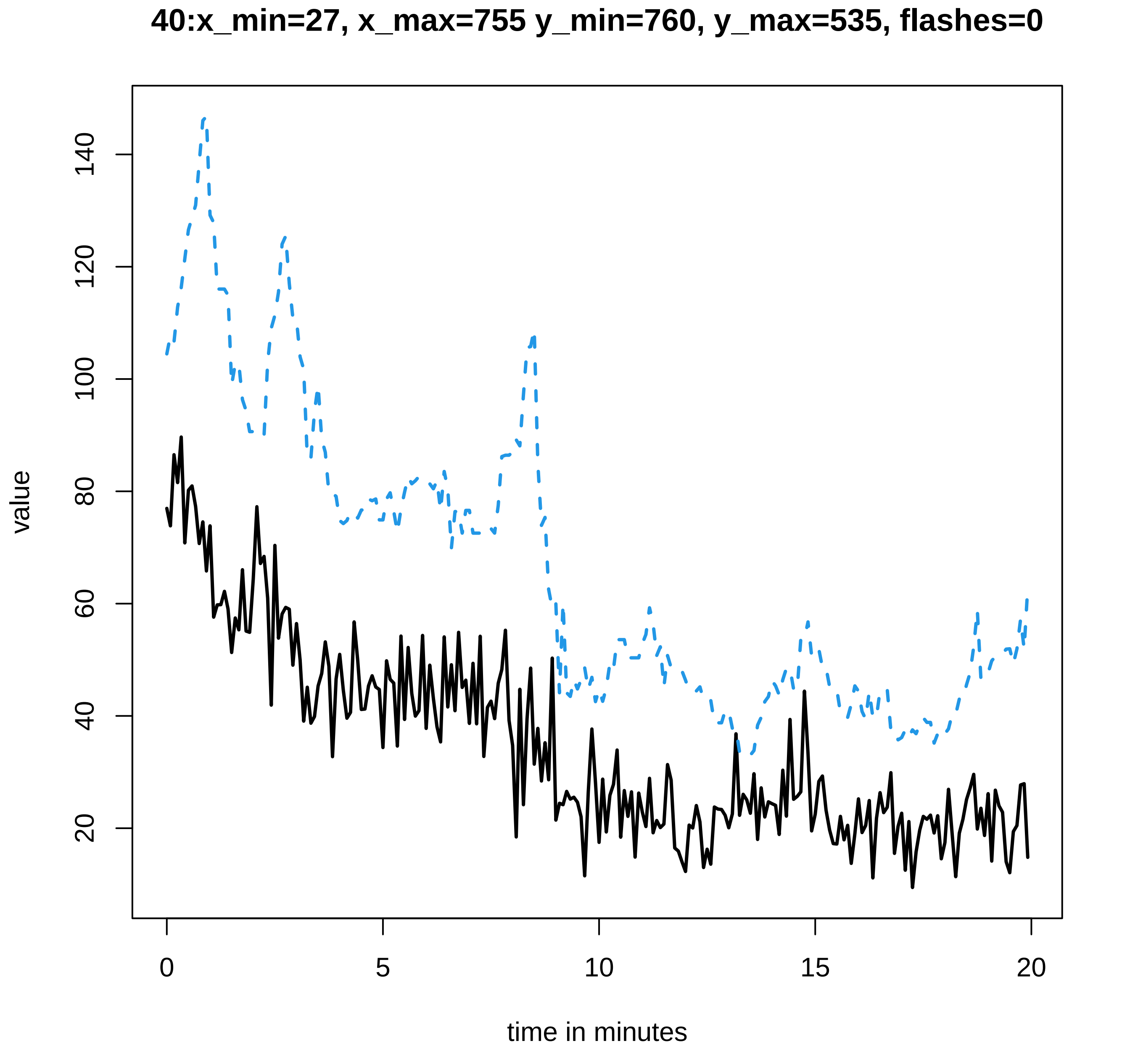
40:x_min=27, x_max=755 y_min=760, y_max=535, flashes=0
value
20
40
60
80
100
120
140
0
5
10
15
20
time in minutes

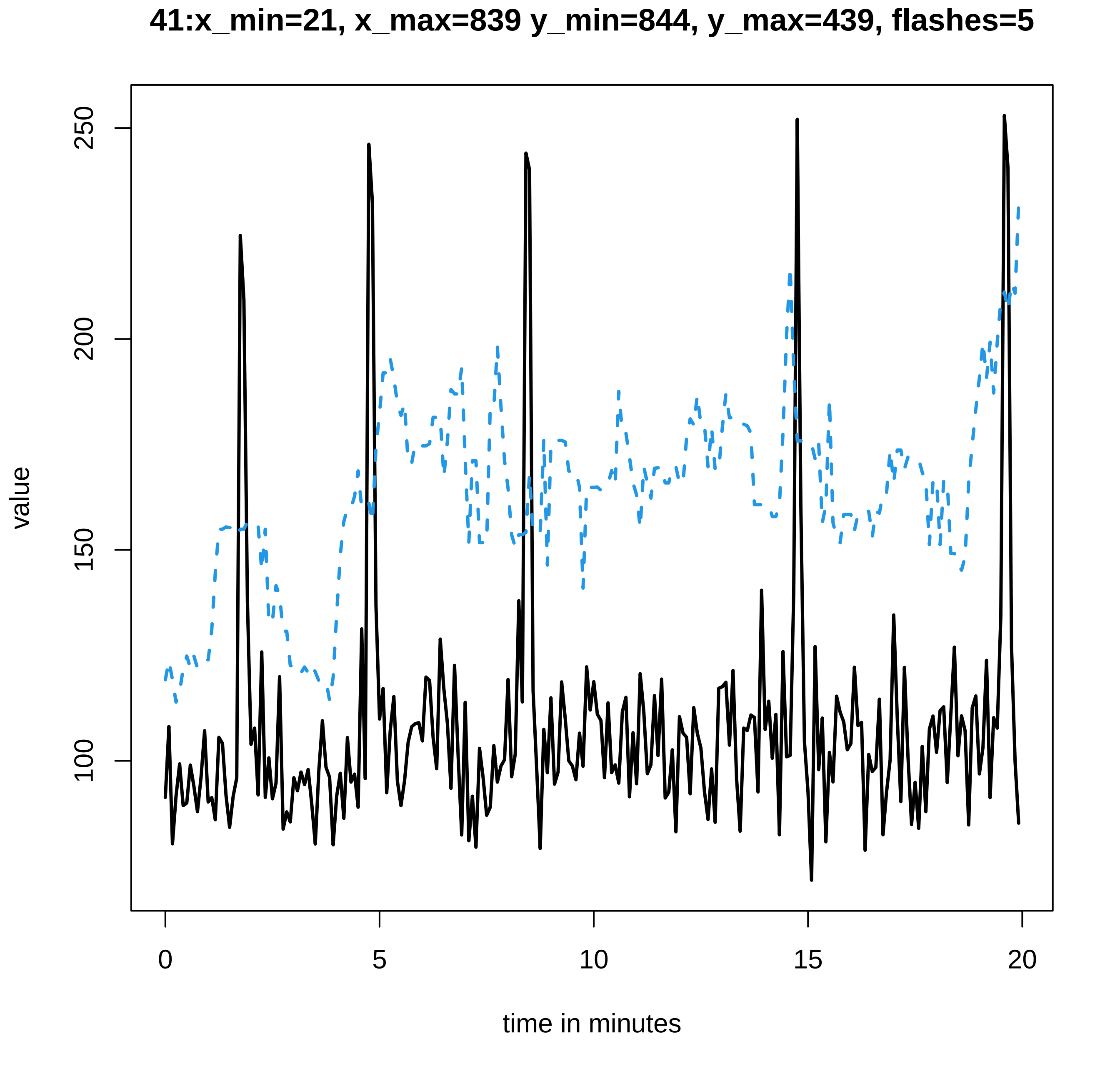

41:x_min=21, x_max=839 y_min=844, y_max=439, flashes=5
value
250
200
150
100
0
5
10
15
20
time in minutes

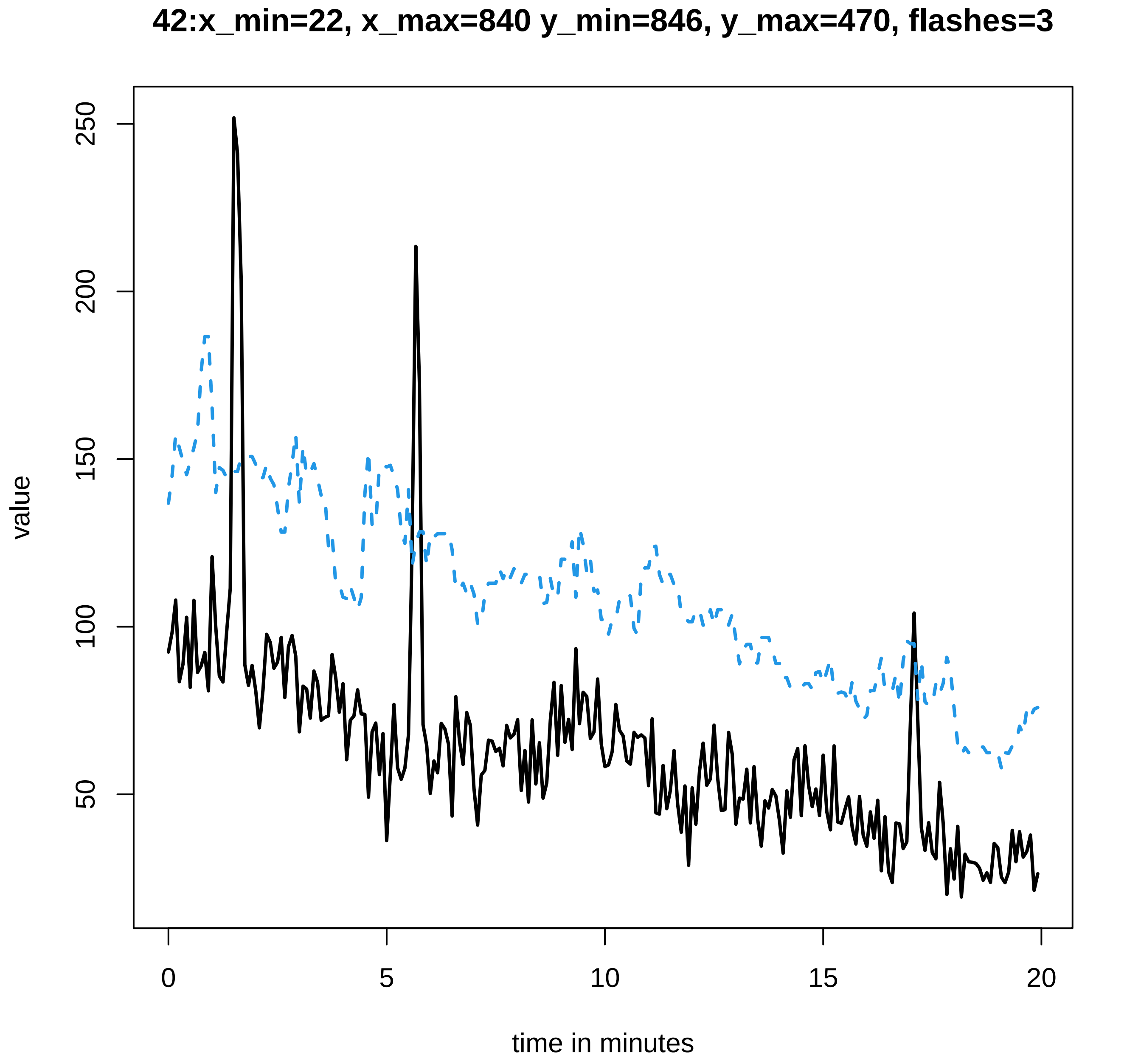
42:x_min=22, x_max=840 y_min=846, y_max=470, flashes=3
value
250
200
150
100
50
0
5
10
15
20
time in minutes

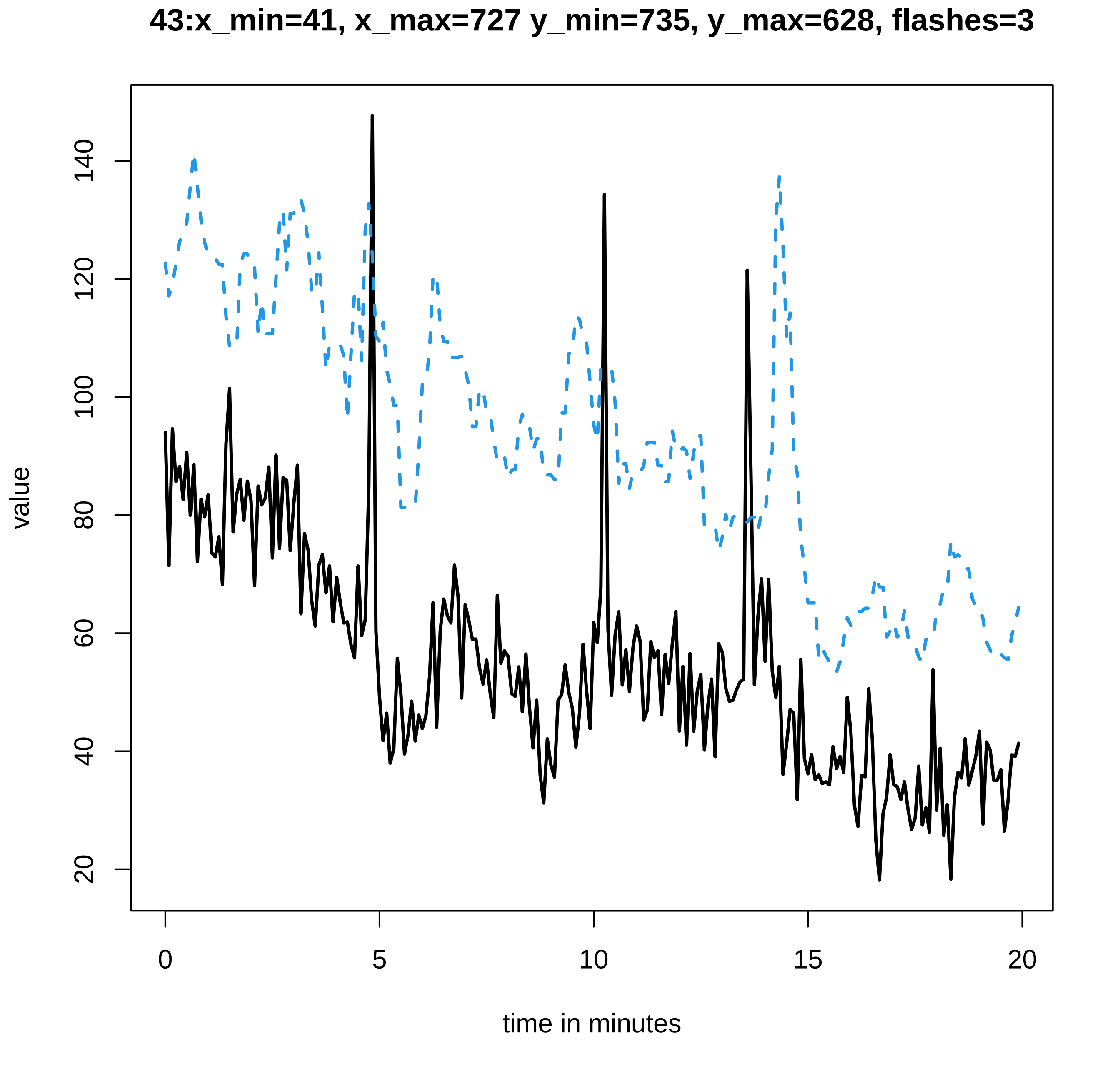
43:x_min=41, x_max=727 y_min=735, y_max=628, flashes=3
value
140
120
100
80
60
40
20
0
5
10
15
20
time in minutes

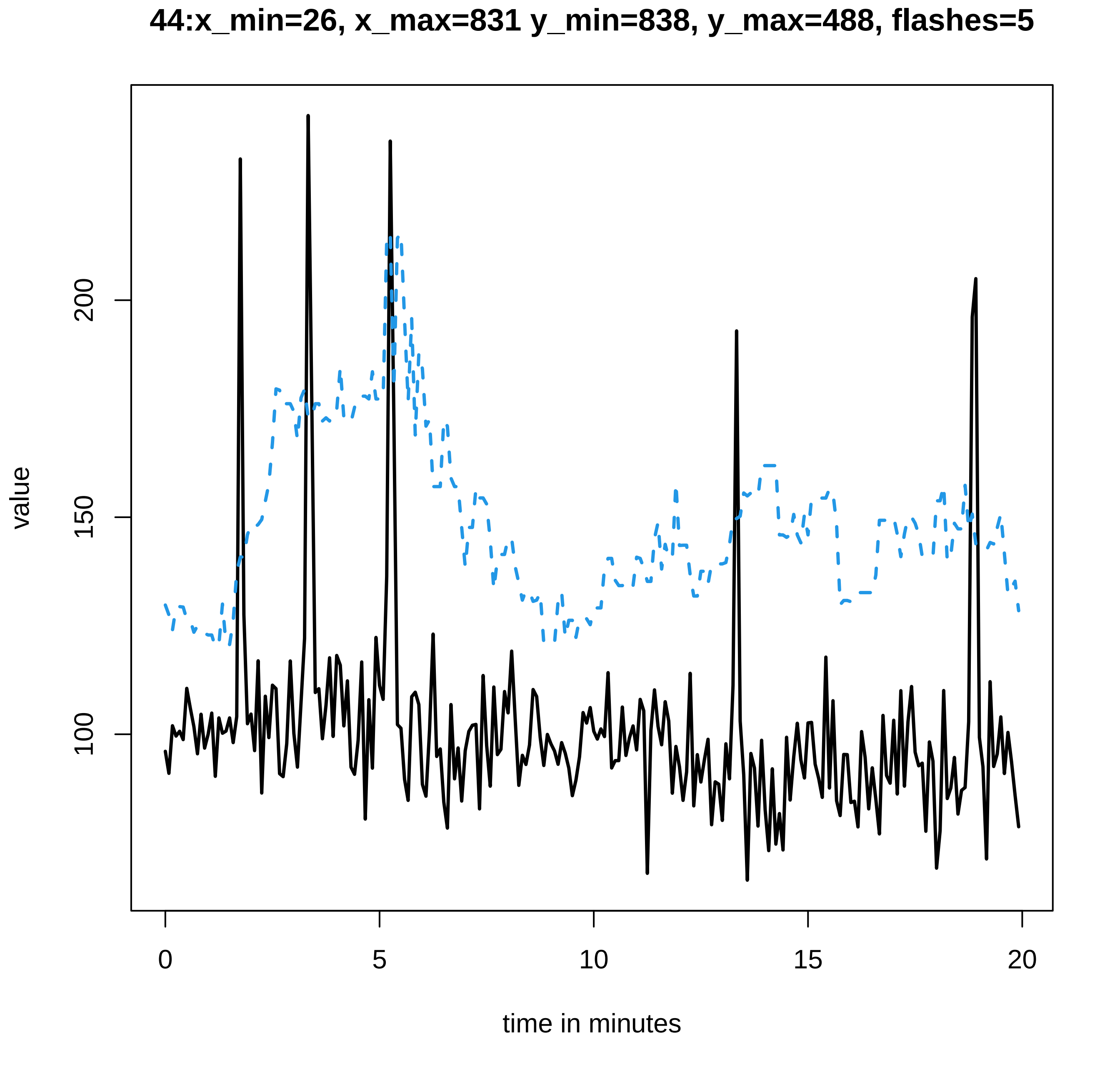

44:x_min=26, x_max=831 y_min=838, y_max=488, flashes=5
value
100
150
200
0
5
10
15
20
time in minutes

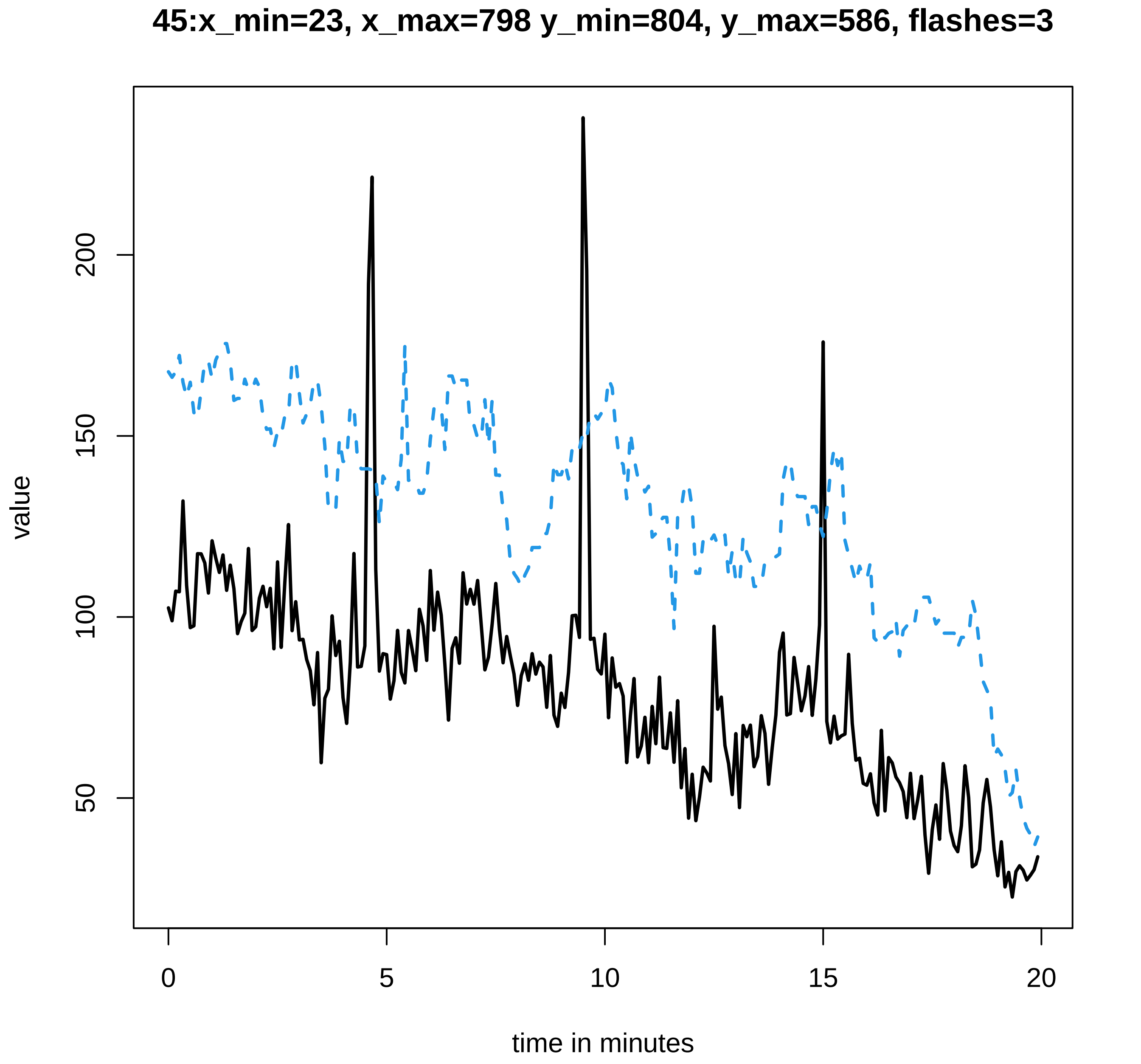
45:x_min=23, x_max=798 y_min=804, y_max=586, flashes=3
value
200
150
100
50
0
5
10
15
20
time in minutes

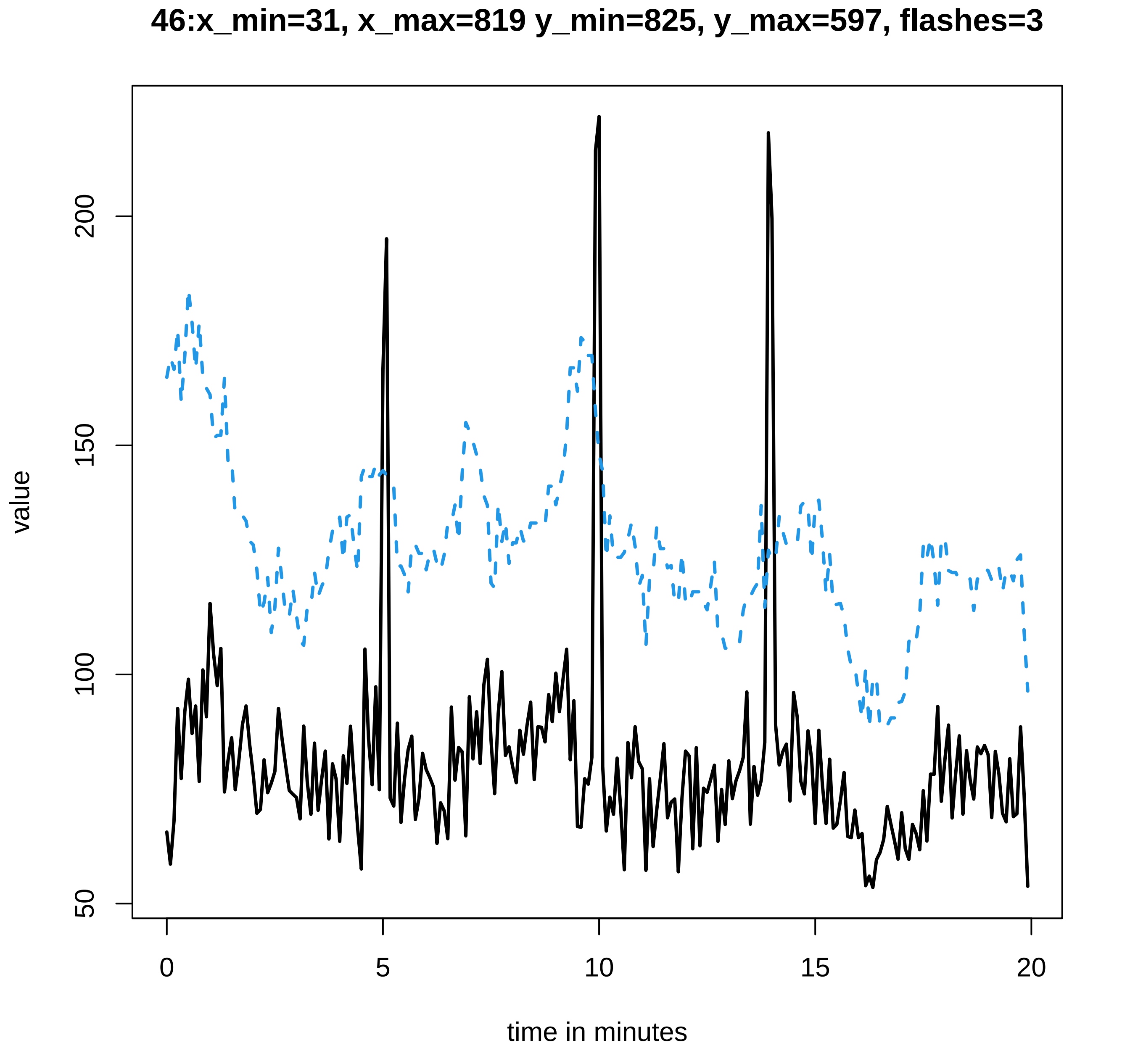

46:x_min=31, x_max=819 y_min=825, y_max=597, flashes=3
value
50
100
150
200
0
5
10
15
20
time in minutes

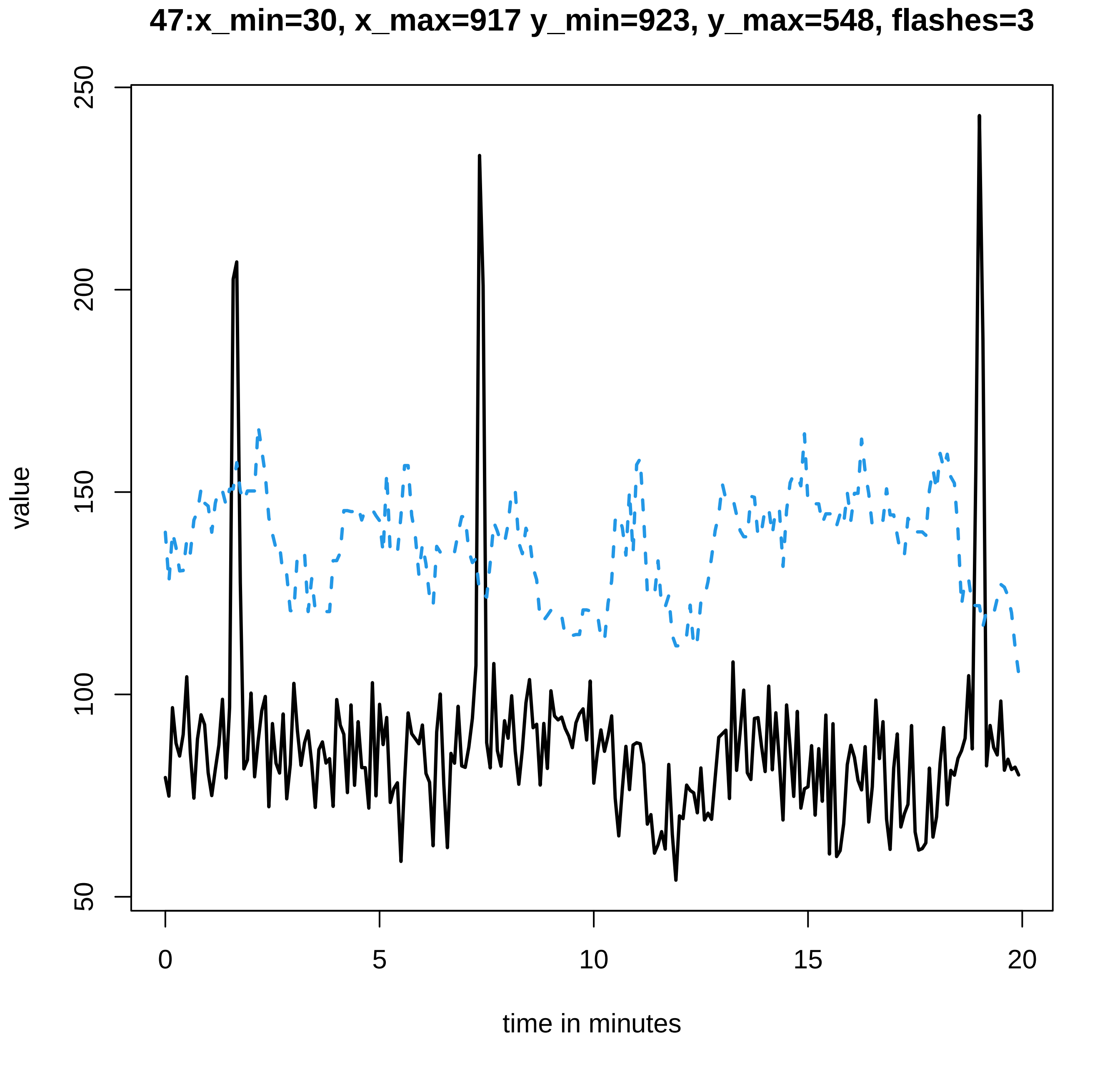
47:x_min=30, x_max=917 y_min=923, y_max=548, flashes=3
value
50
100
150
200
250
0
5
10
15
20
time in minutes
