## Supplementary data Calcium spiking for "LysM Receptor Proteins are Required for Ectomycorrhizal Symbiosis in Poplar": profiles-spiking-9.pdf

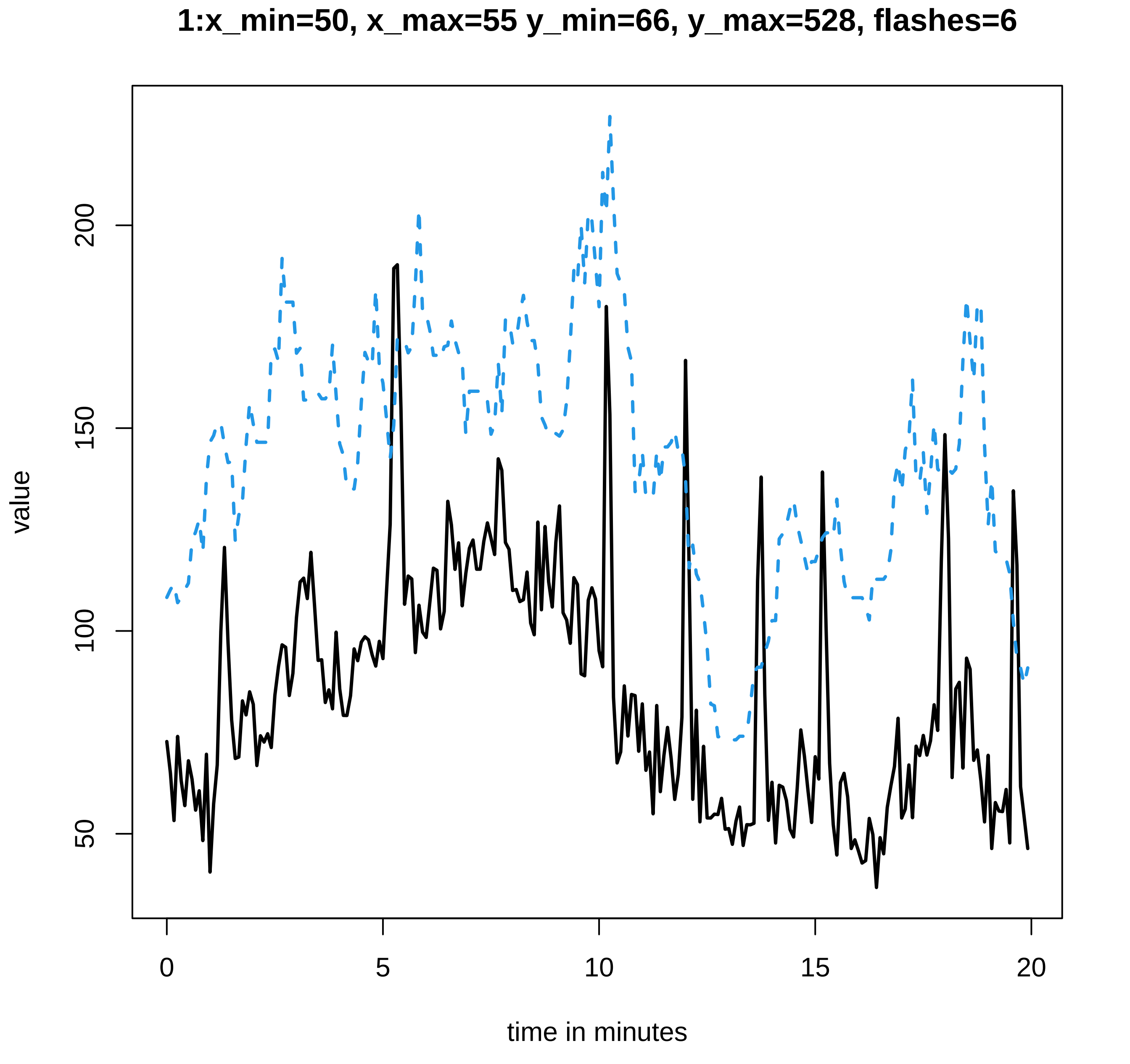
1:x_min=50, x_max=55 y_min=66, y_max=528, flashes=6
value
200
150
100
50
0
5
10
15
20
time in minutes

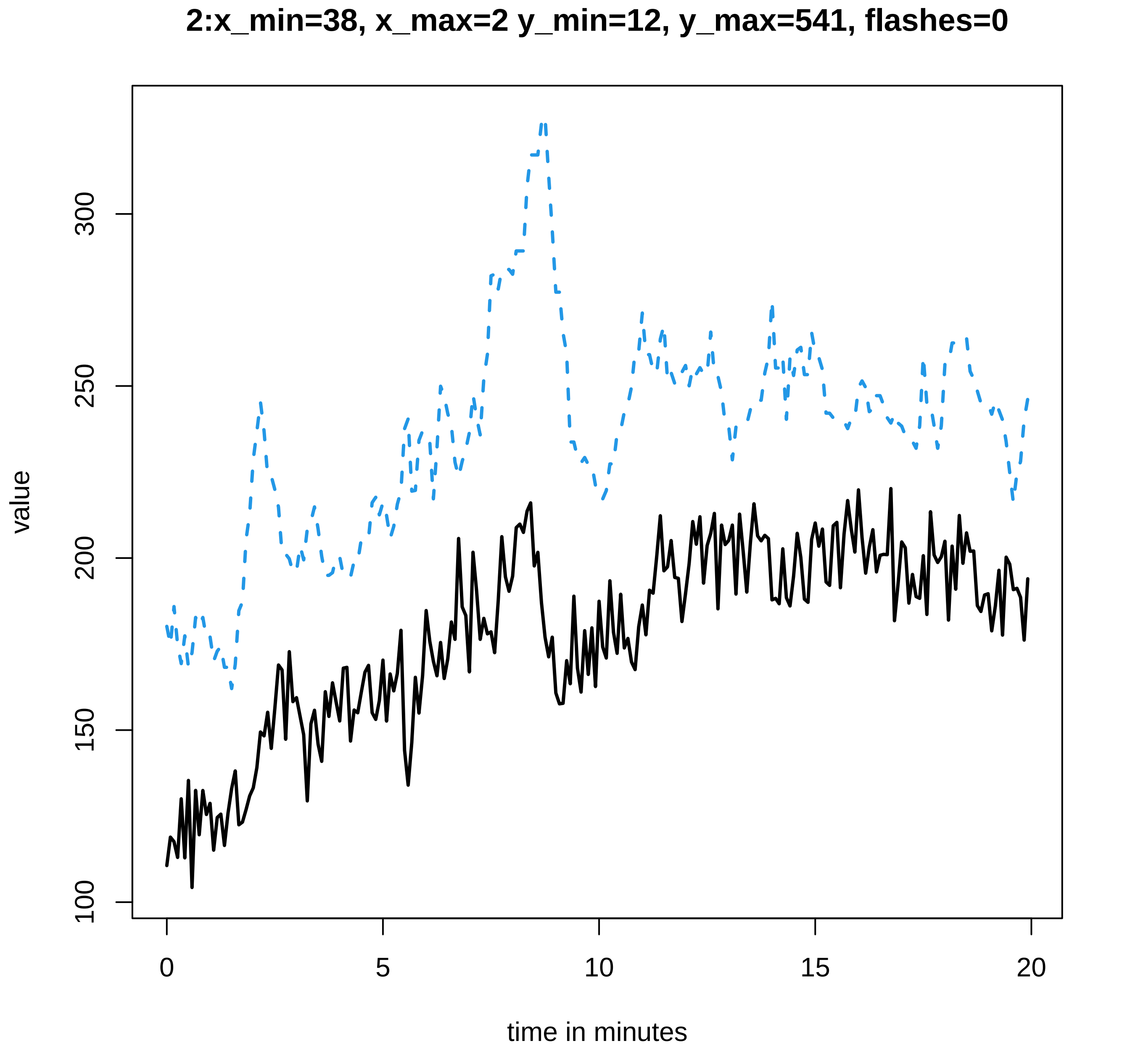

2:x_min=38, x_max=2 y_min=12, y_max=541, flashes=0
value
100
150
200
250
300
time in minutes
0
5
10
15
20

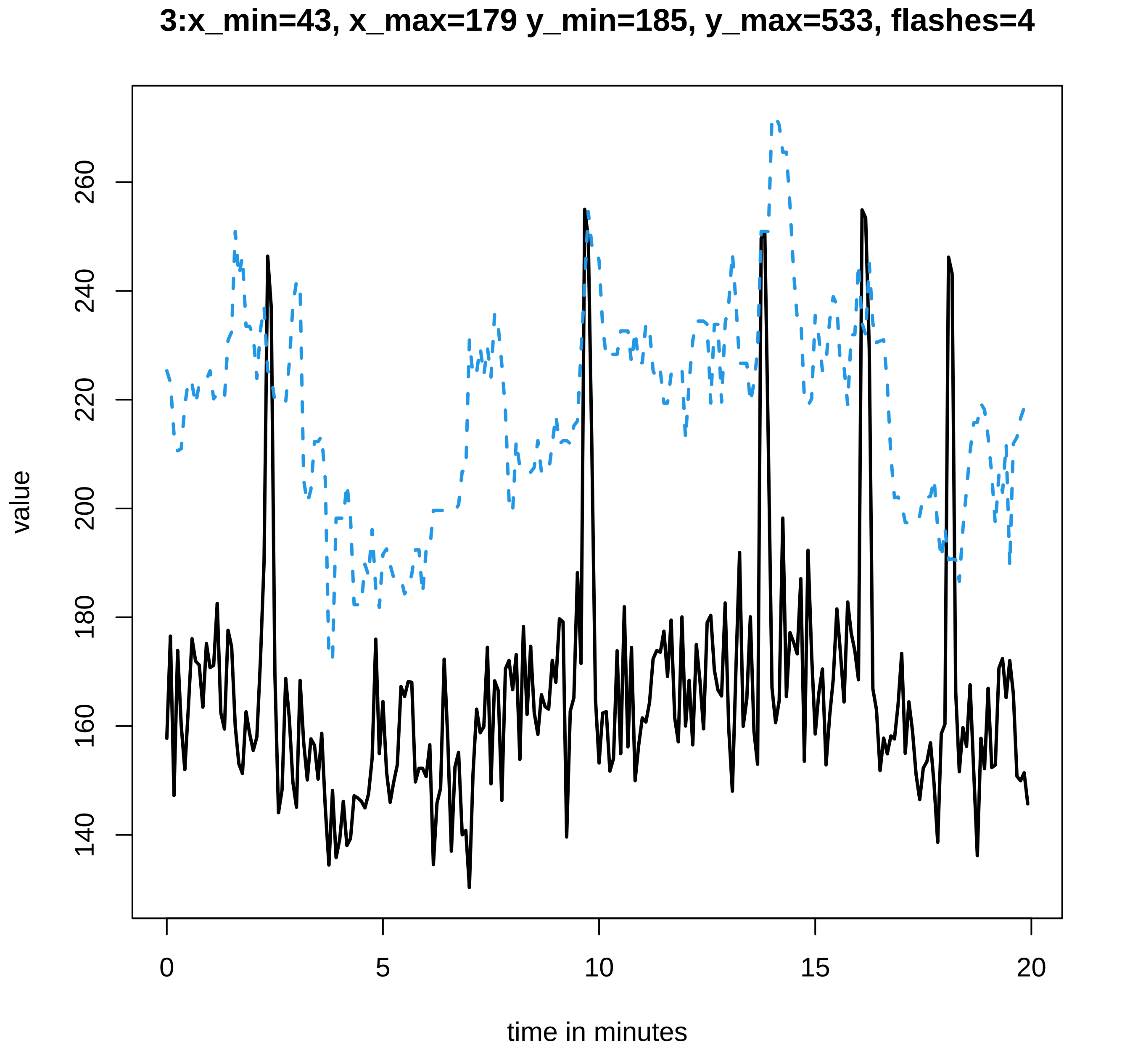

3:x_min=43, x_max=179 y_min=185, y_max=533, flashes=4
value
time in minutes
140
160
180
200
220
240
260
0
5
10
15
20

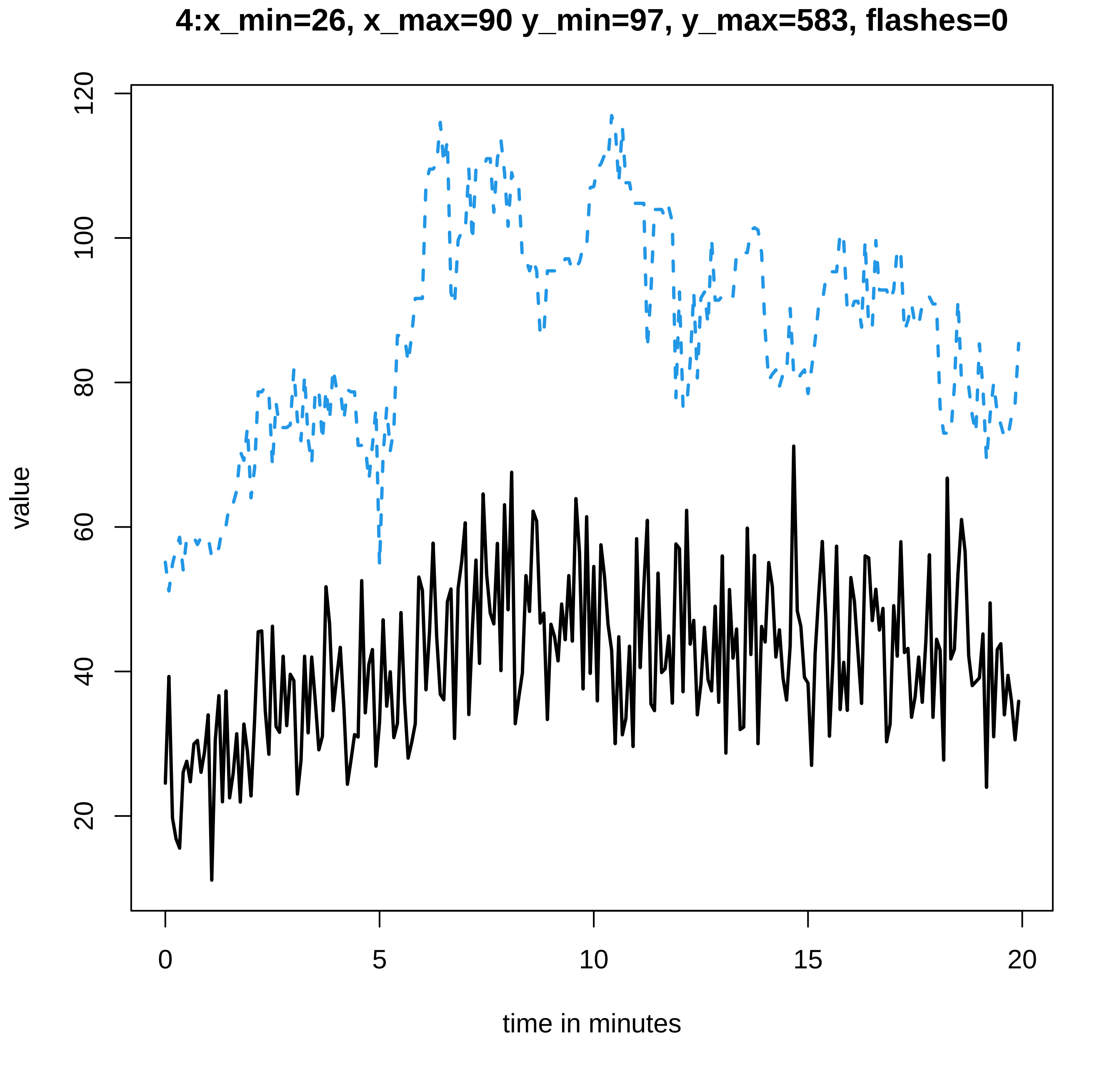

4:x_min=26, x_max=90 y_min=97, y_max=583, flashes=0
value
time in minutes
0
5
10
15
20
20
40
60
80
100
120

**5:x_min=44, x_max=194 y_min=205, y_max=606, flashes=0**

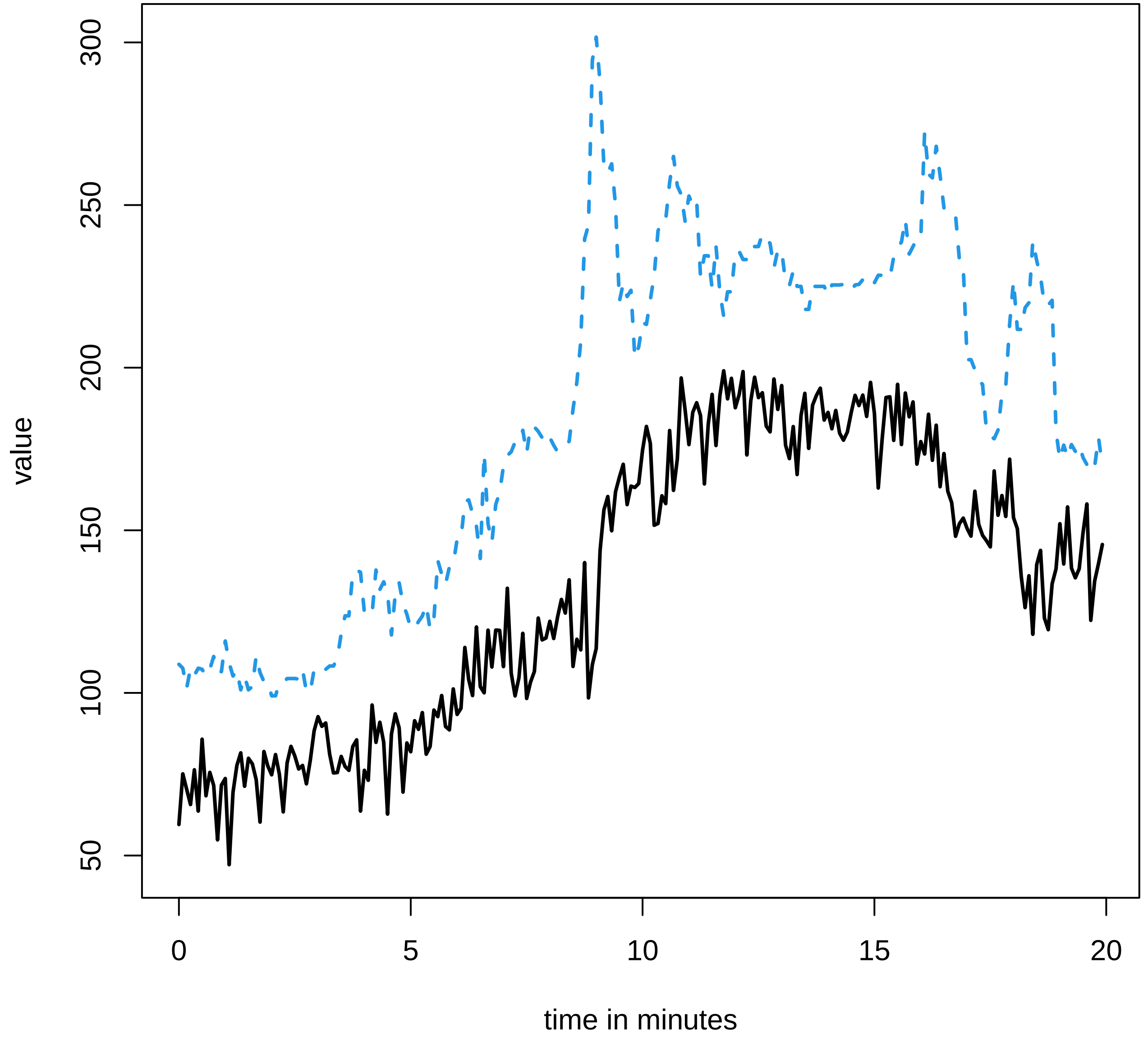

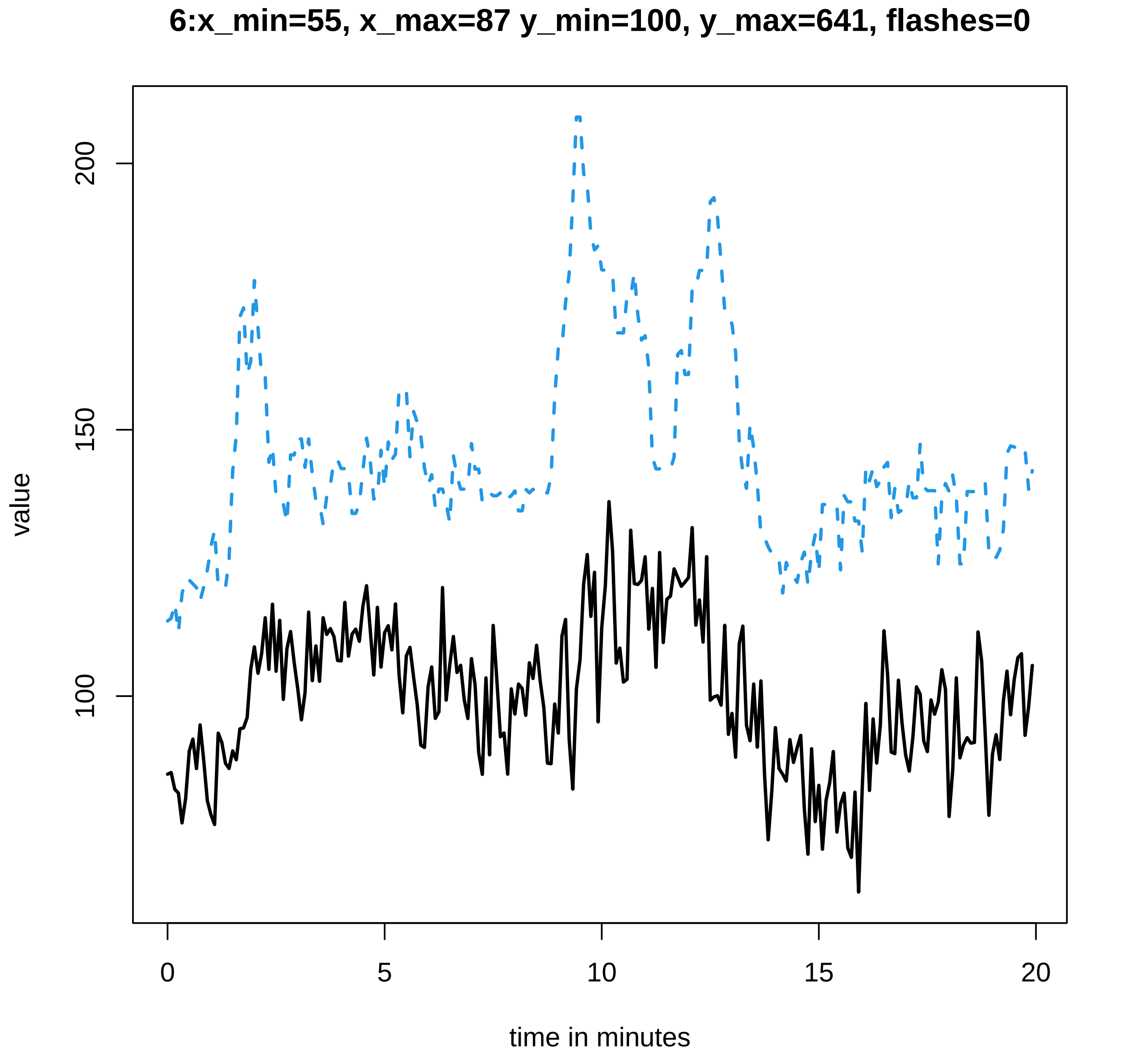

6:x_min=55, x_max=87 y_min=100, y_max=641, flashes=0
value
200
150
100
0
5
10
15
20
time in minutes

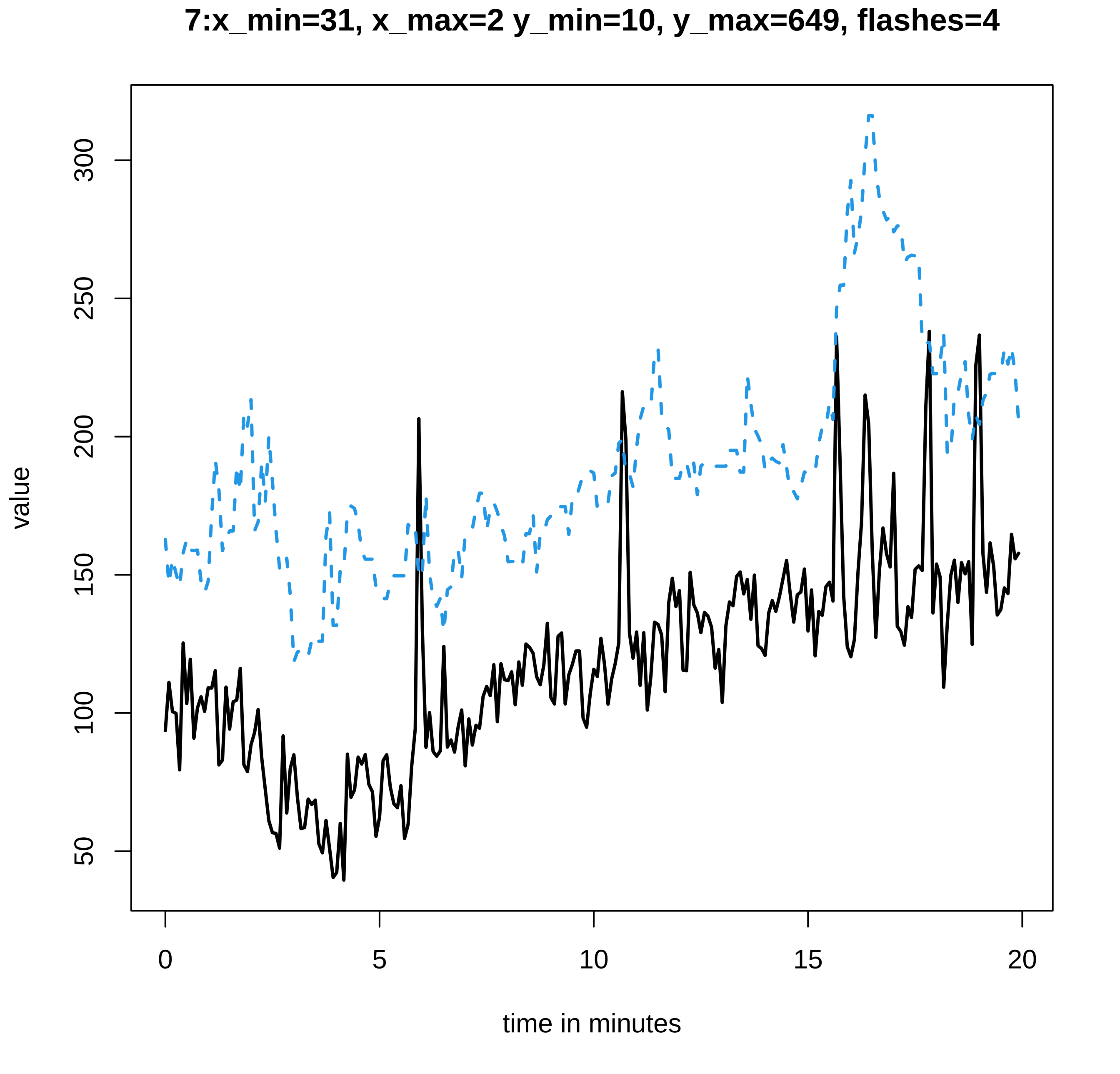

7:x_min=31, x_max=2 y_min=10, y_max=649, flashes=4
value
300
250
200
150
100
50
0
5
10
15
20
time in minutes

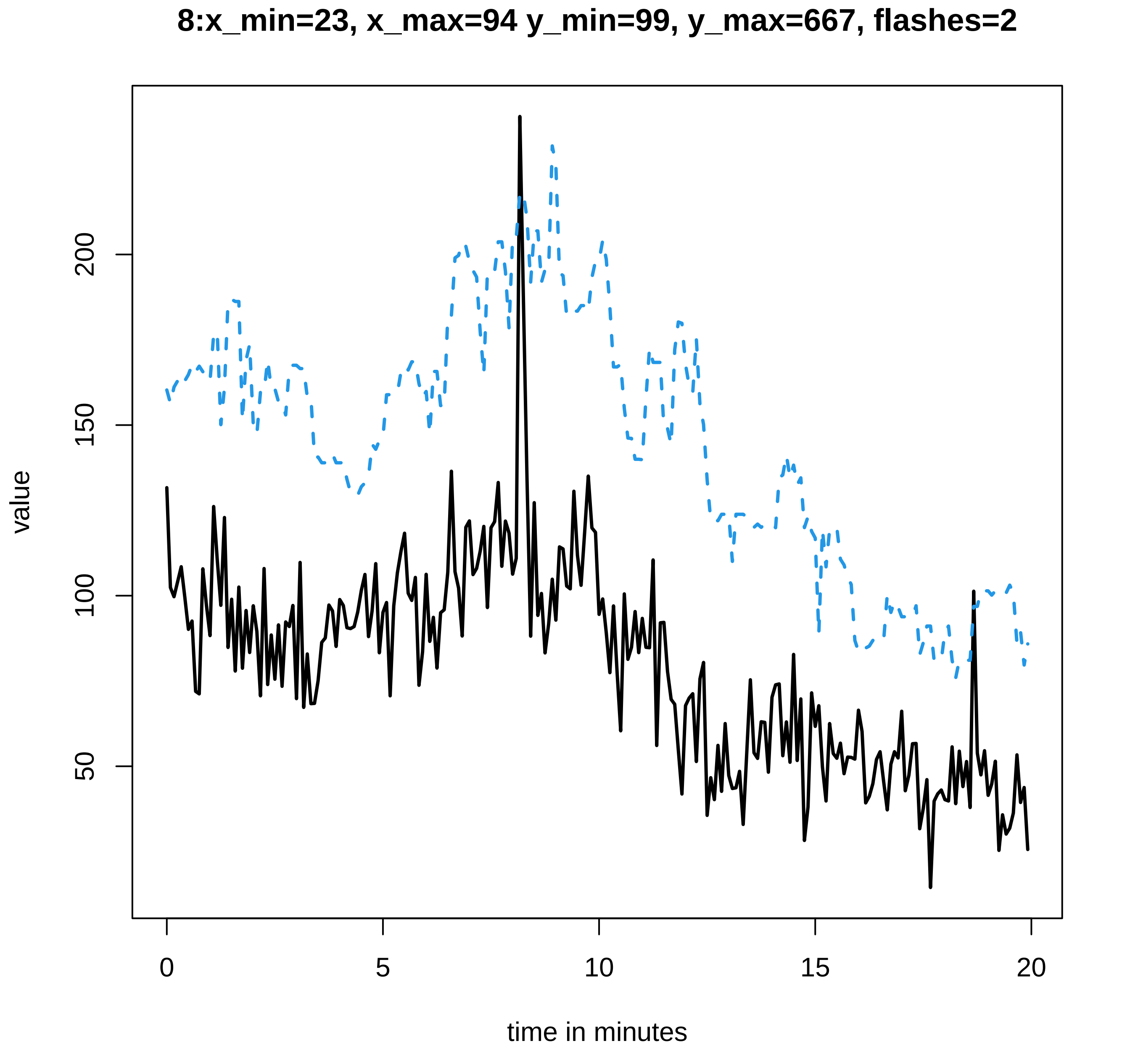

8:x_min=23, x_max=94 y_min=99, y_max=667, flashes=2
value
200
150
100
50
0
5
10
15
20
time in minutes

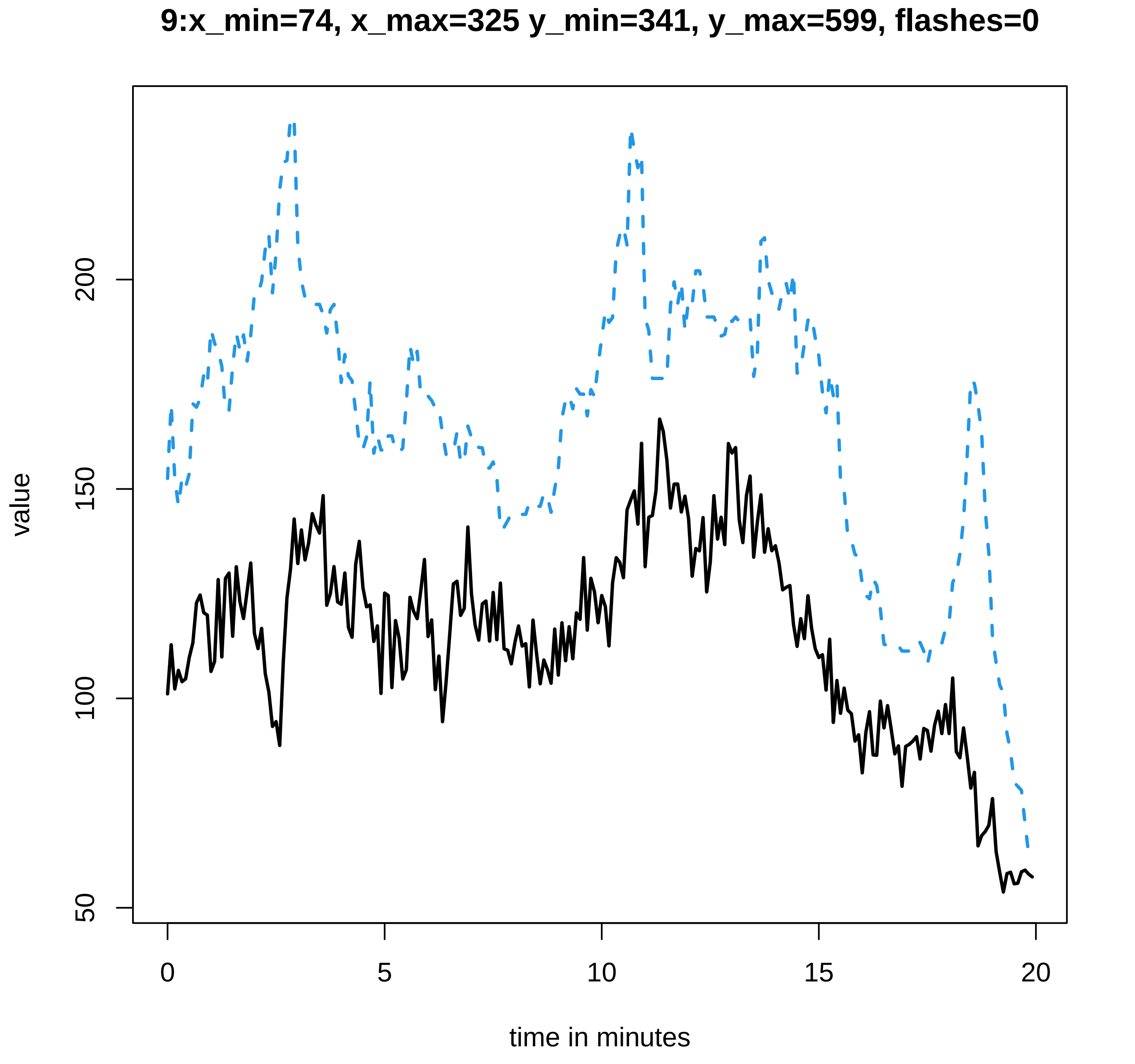
9:x_min=74, x_max=325 y_min=341, y_max=599, flashes=0
value
50
100
150
200
0
5
10
15
20
time in minutes

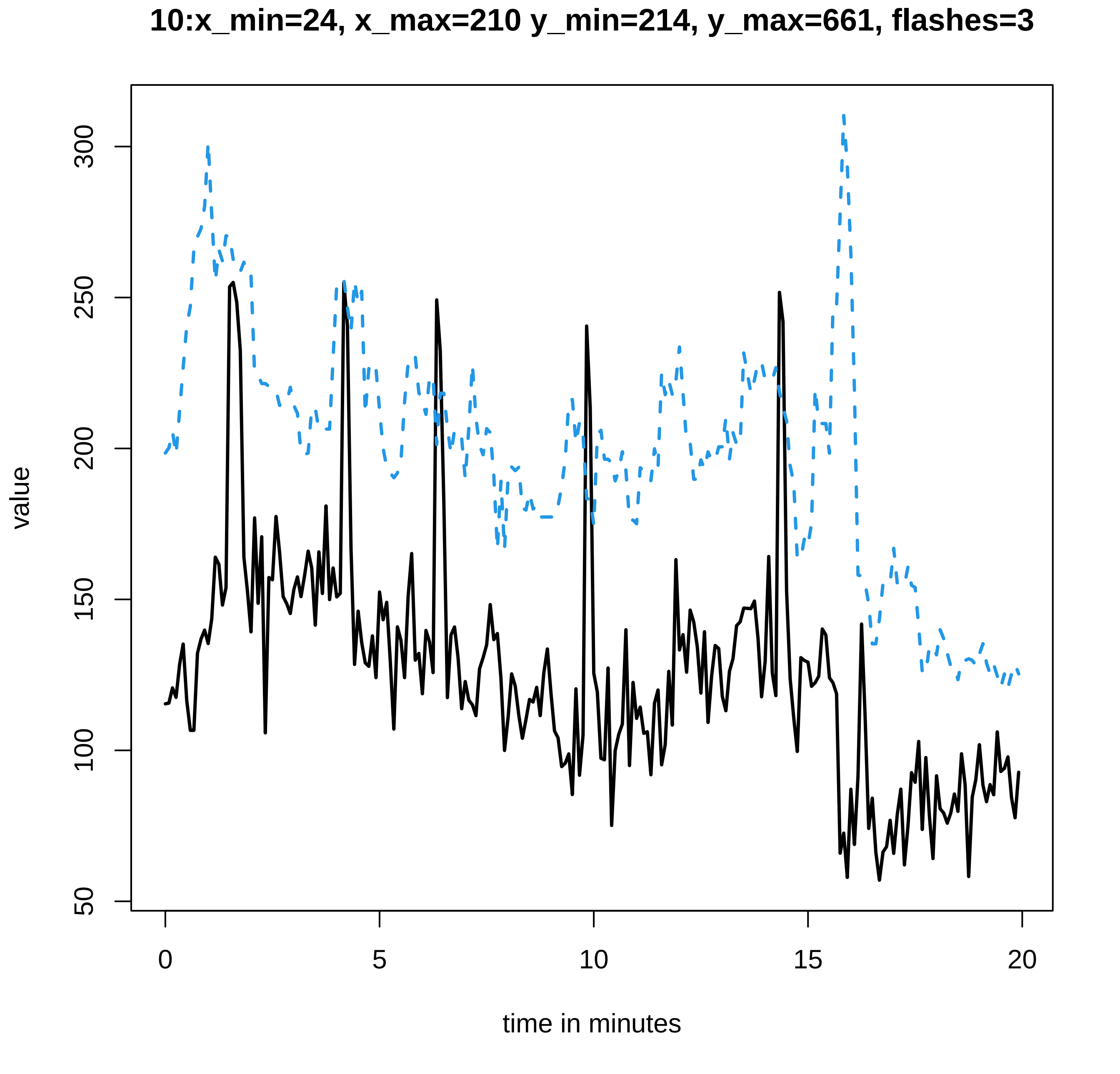

10:x_min=24, x_max=210 y_min=214, y_max=661, flashes=3
value
50
100
150
200
250
300
0
5
10
15
20
time in minutes

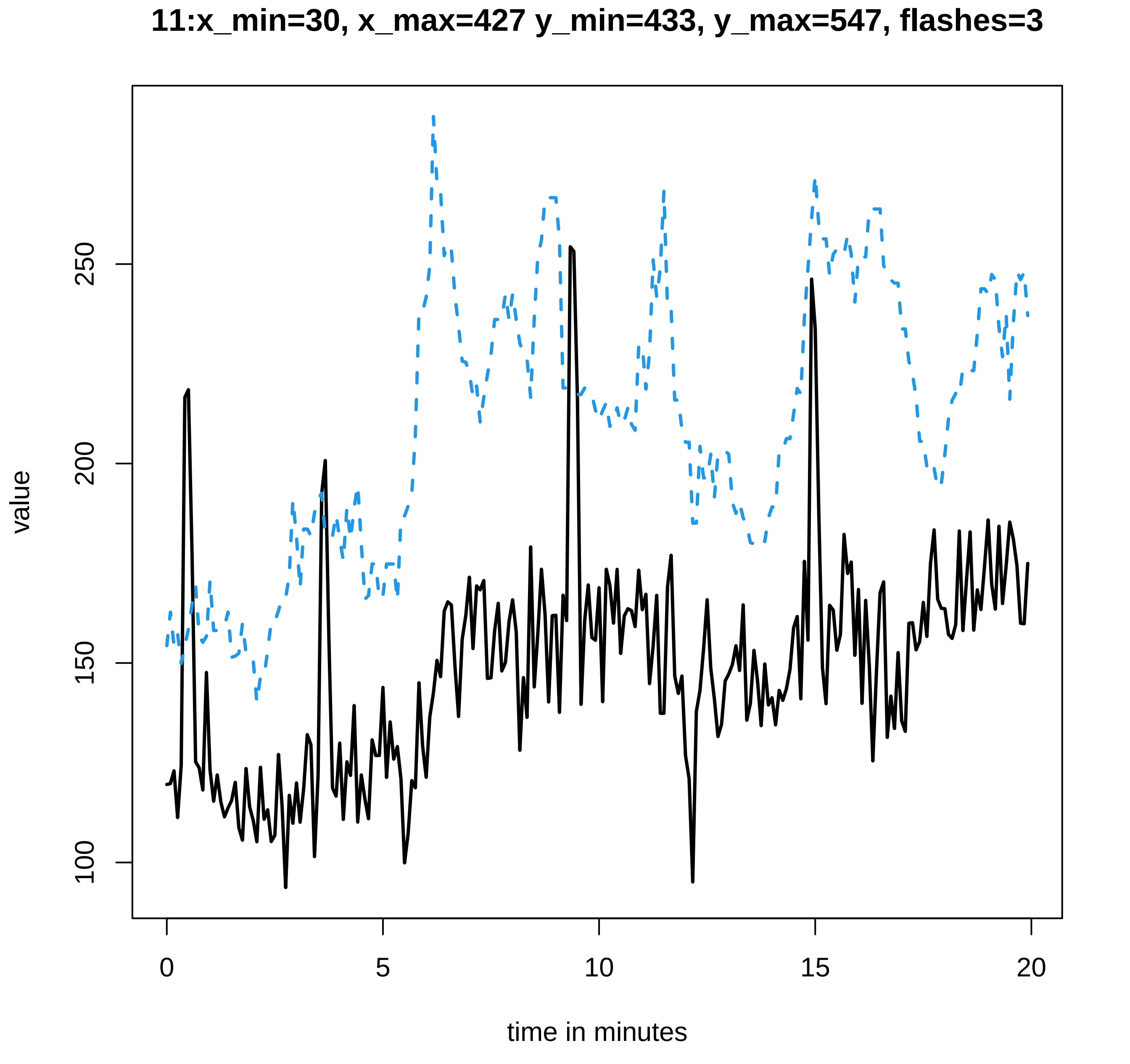
11:x_min=30, x_max=427 y_min=433, y_max=547, flashes=3
value
100
150
200
250
0
5
10
15
20
time in minutes

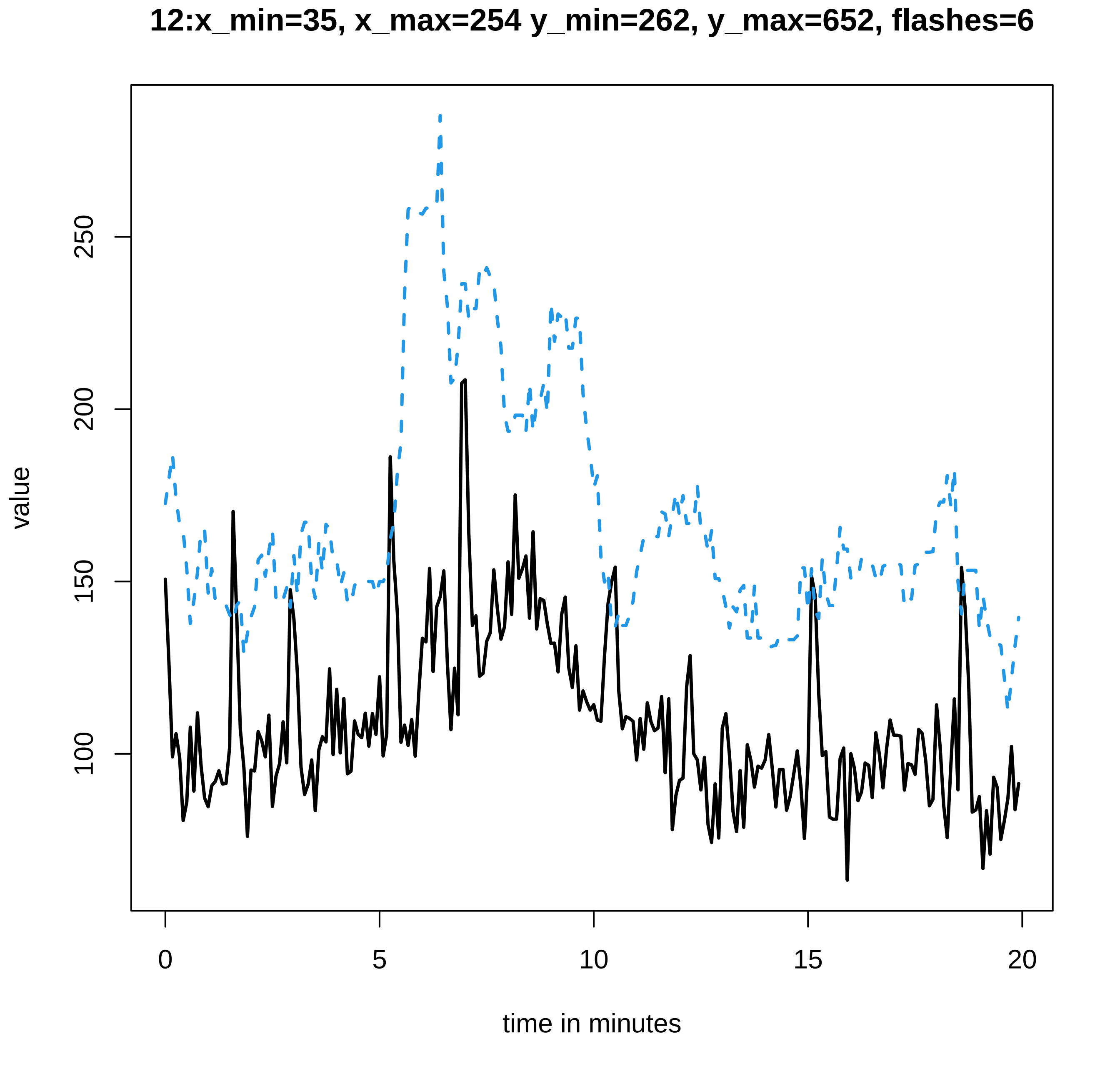
12:x_min=35, x_max=254 y_min=262, y_max=652, flashes=6
value
100
150
200
250
0
5
10
15
20
time in minutes

**13:x_min=23, x_max=484 y_min=490, y_max=530, flashes=1**

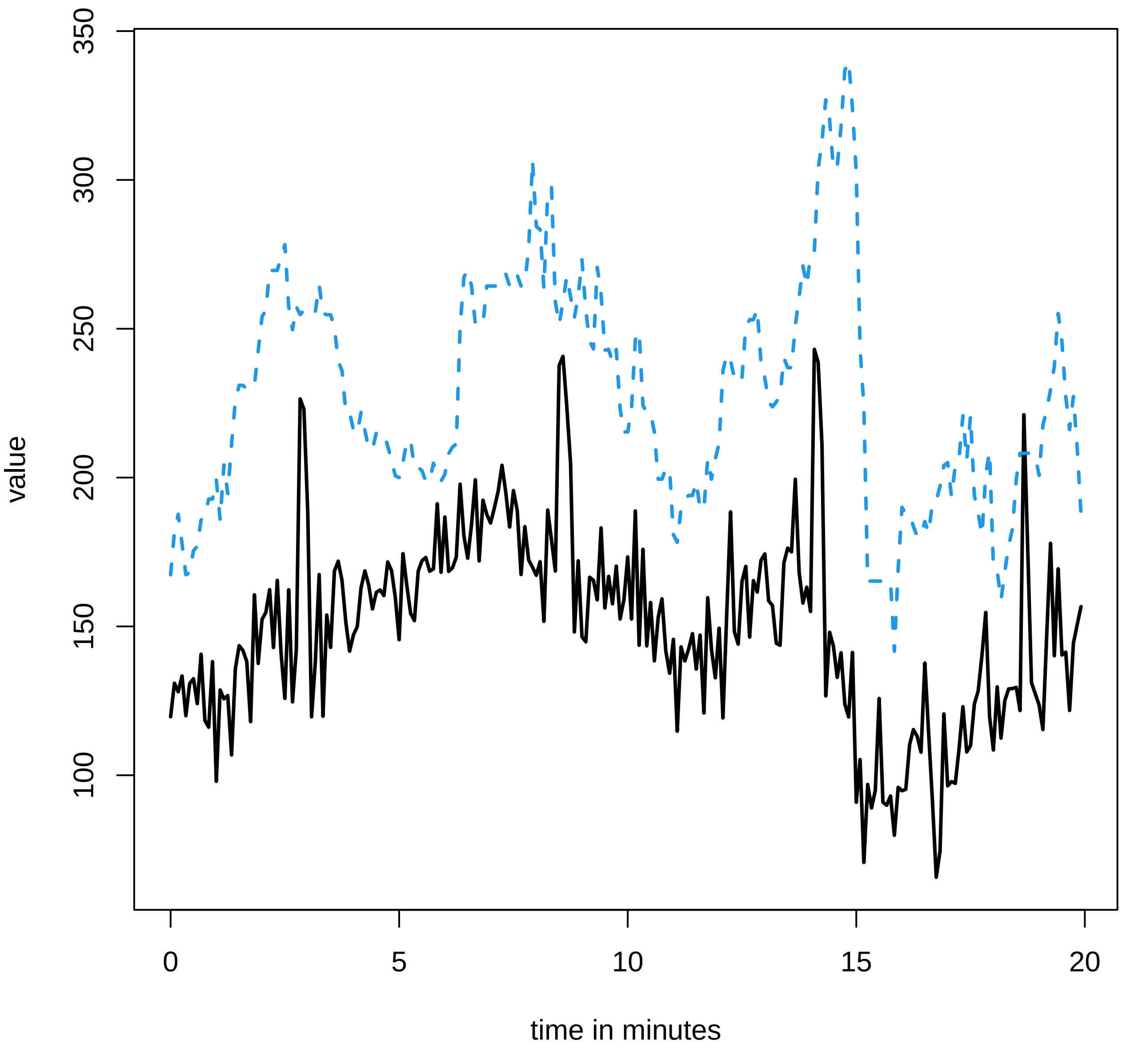

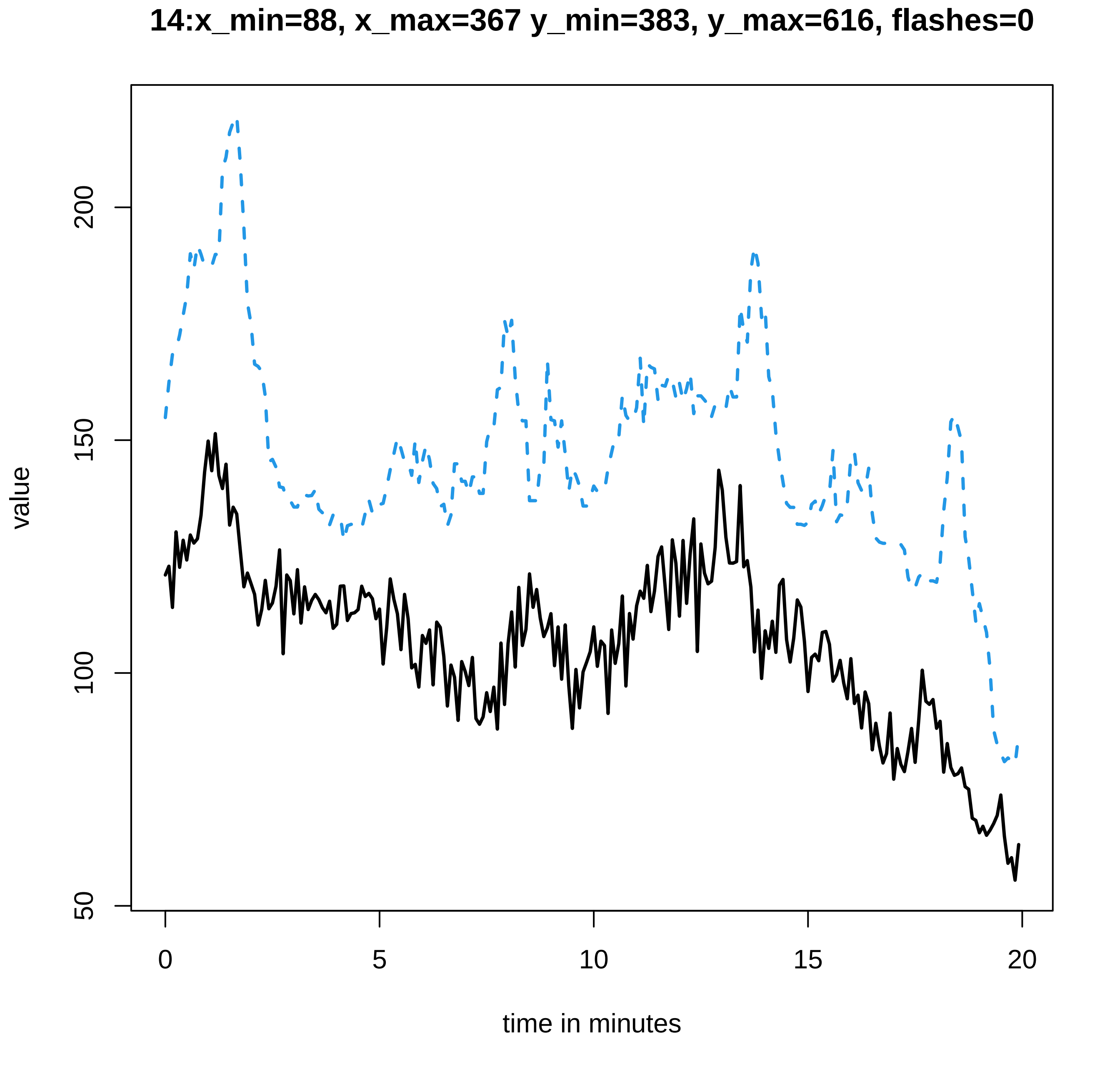
14:x_min=88, x_max=367 y_min=383, y_max=616, flashes=0
value
50
100
150
200
0
5
10
15
20
time in minutes

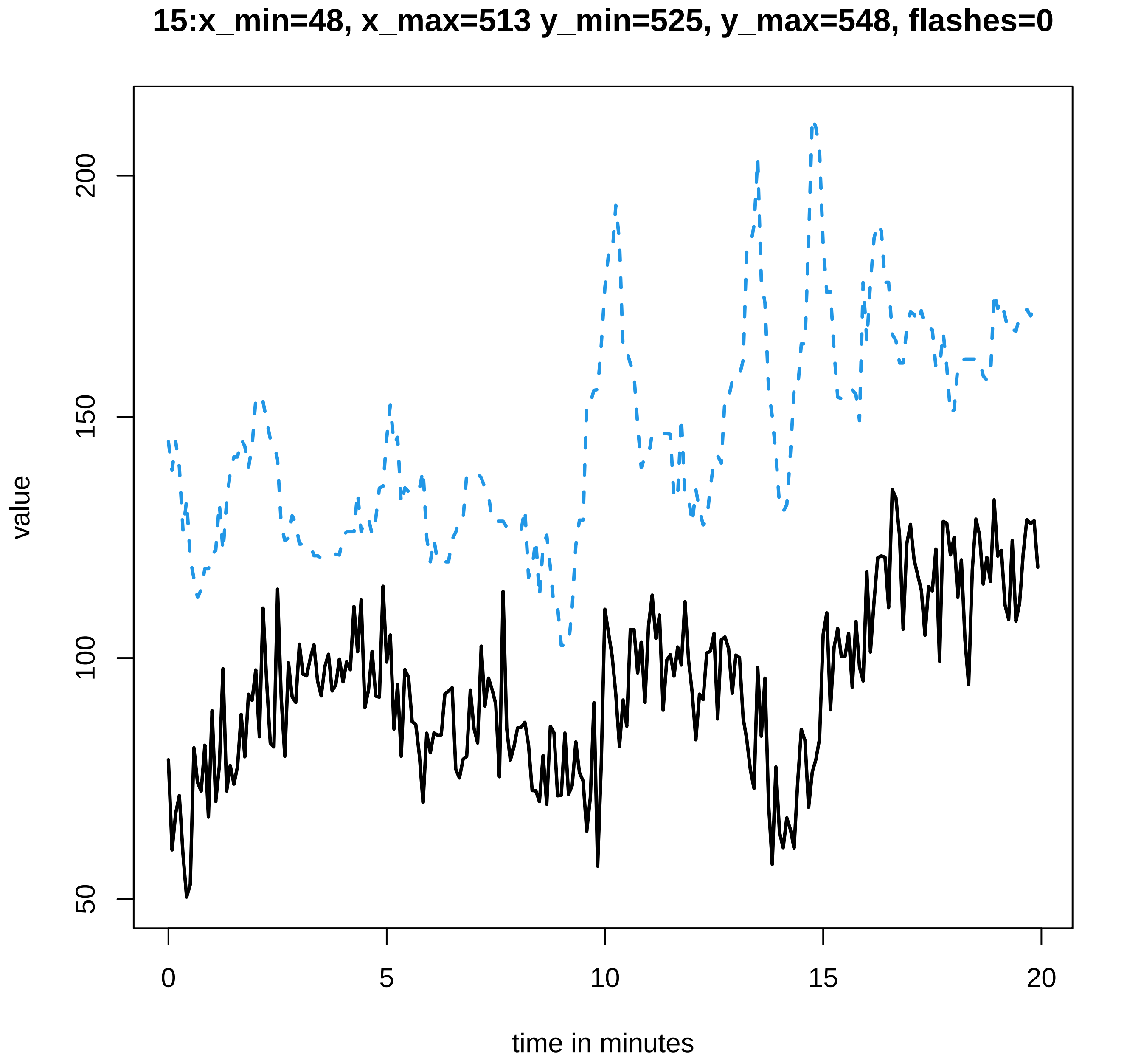

15:x_min=48, x_max=513 y_min=525, y_max=548, flashes=0
value
200
150
100
50
0
5
10
15
20
time in minutes

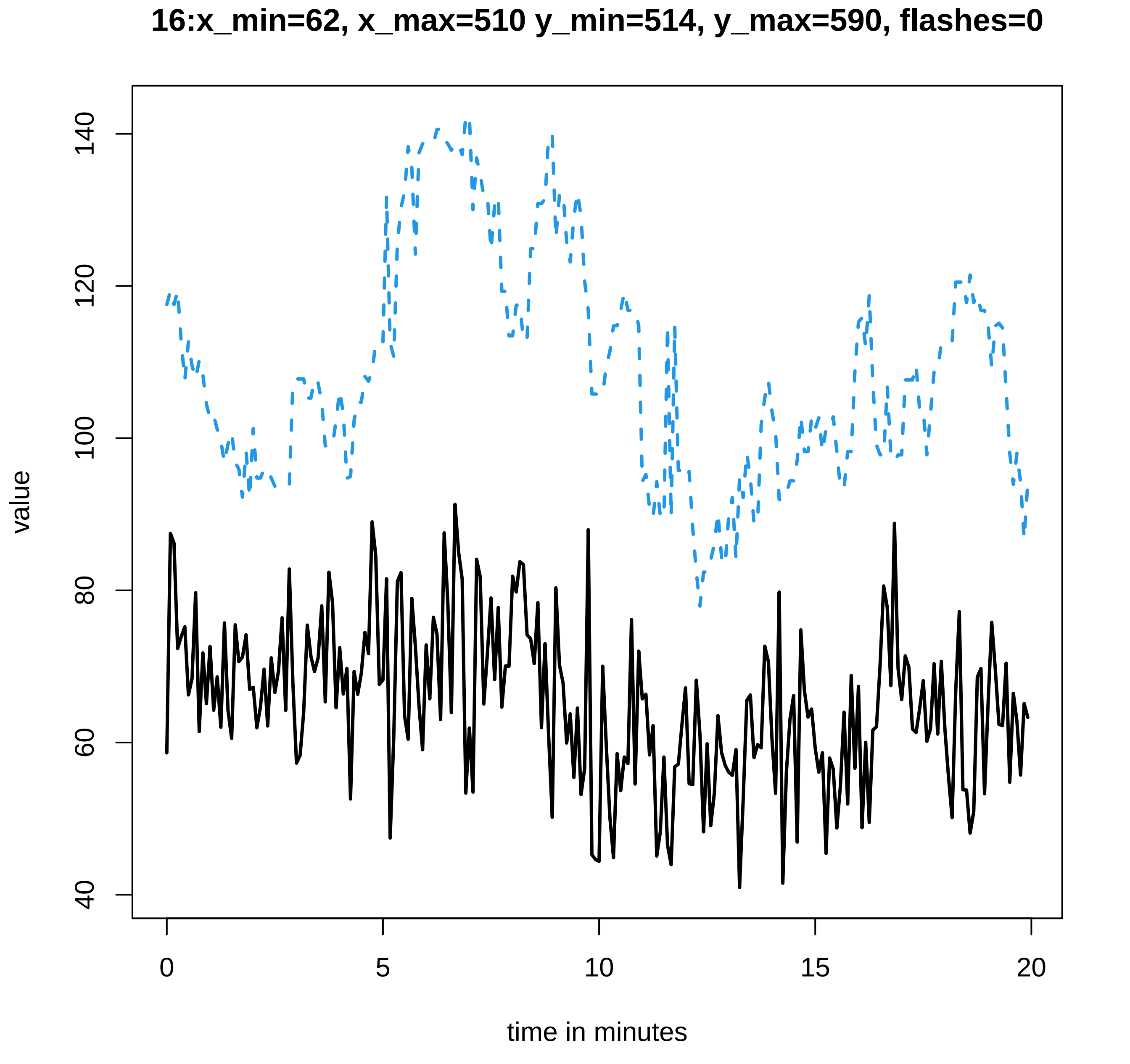

16:x_min=62, x_max=510 y_min=514, y_max=590, flashes=0
value
time in minutes
40
60
80
100
120
140
0
5
10
15
20

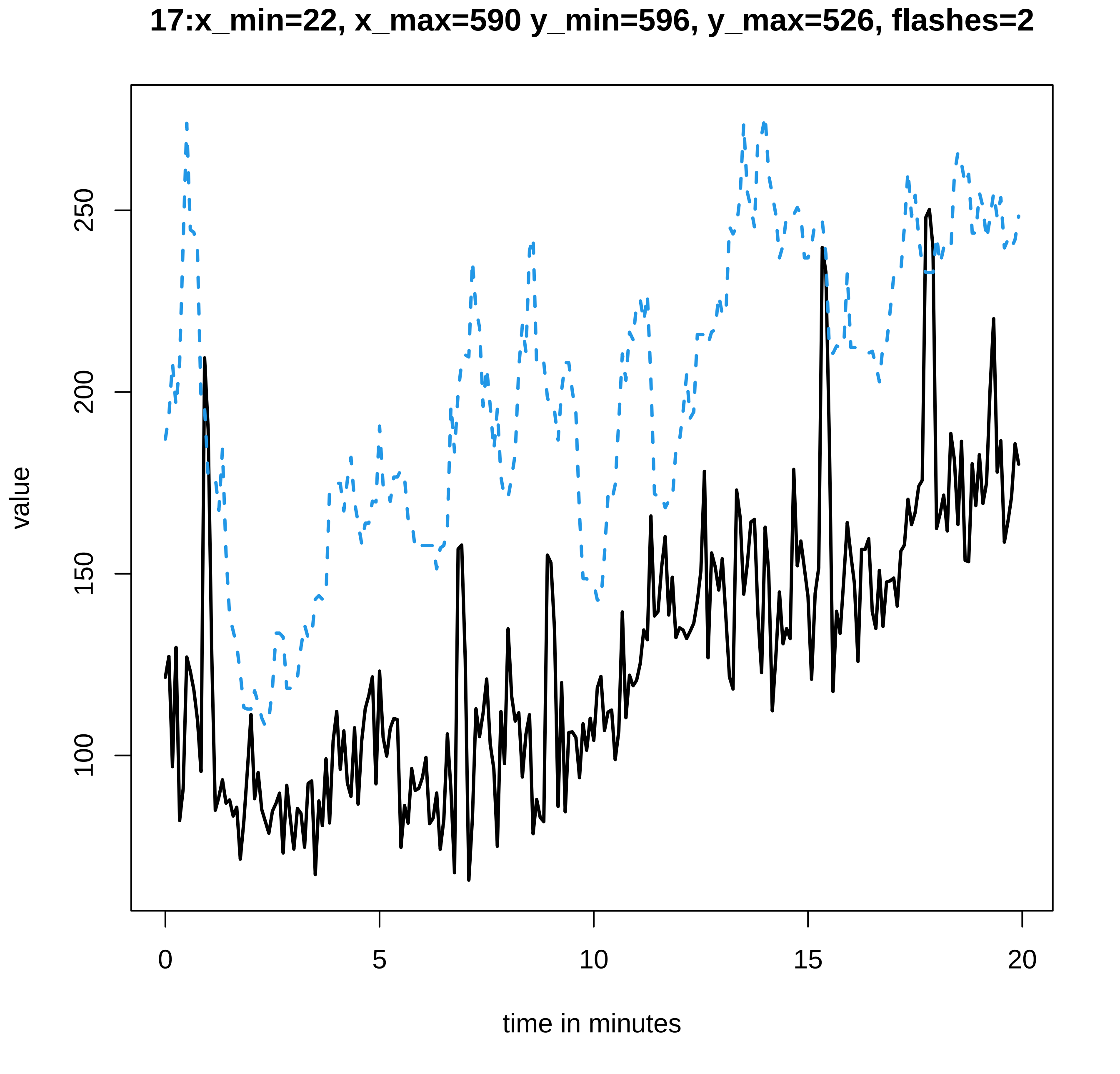

17:x_min=22, x_max=590 y_min=596, y_max=526, flashes=2
value
100
150
200
250
0
5
10
15
20
time in minutes

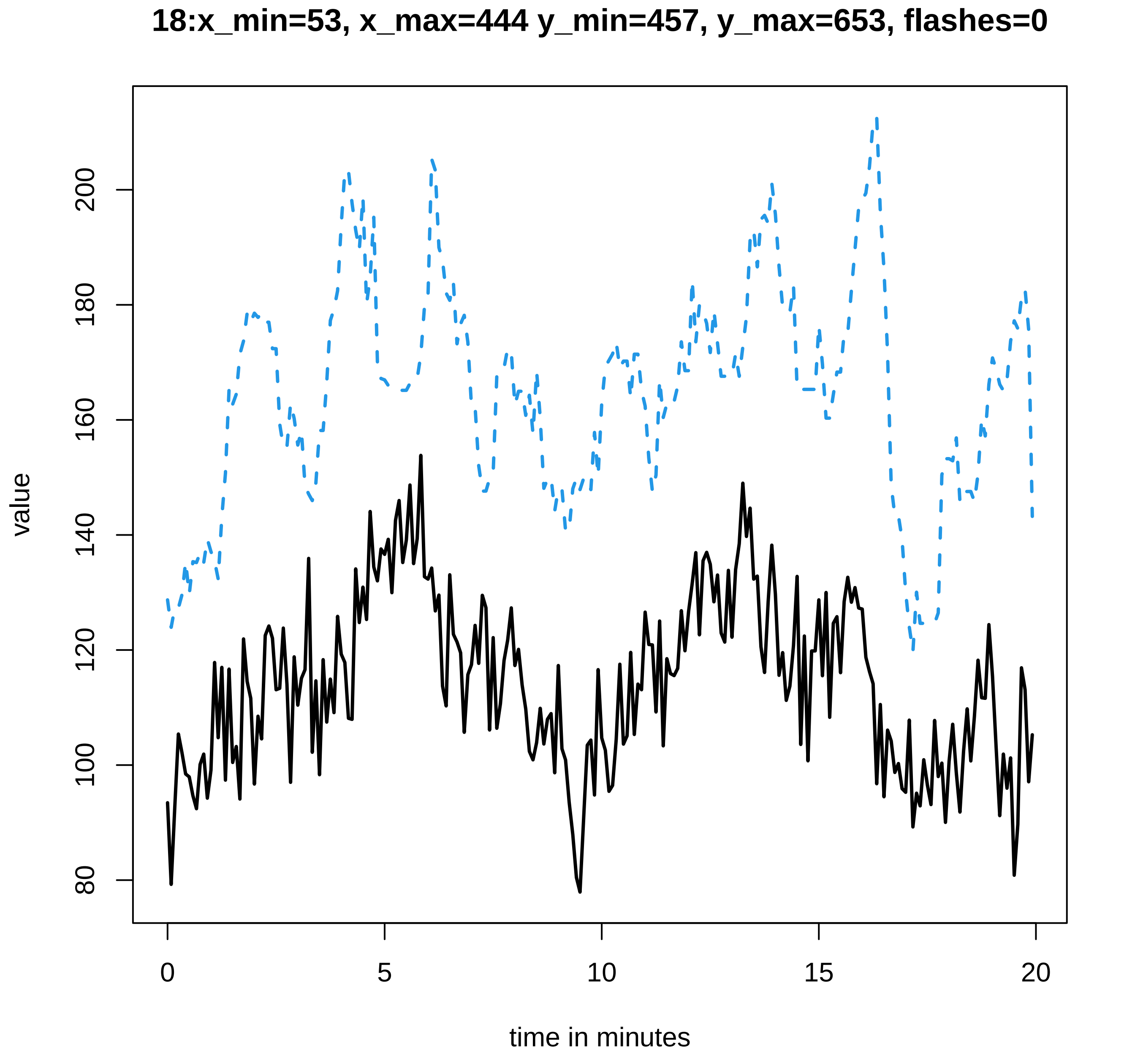

18:x_min=53, x_max=444 y_min=457, y_max=653, flashes=0
value
80
100
120
140
160
180
200
0
5
10
15
20
time in minutes

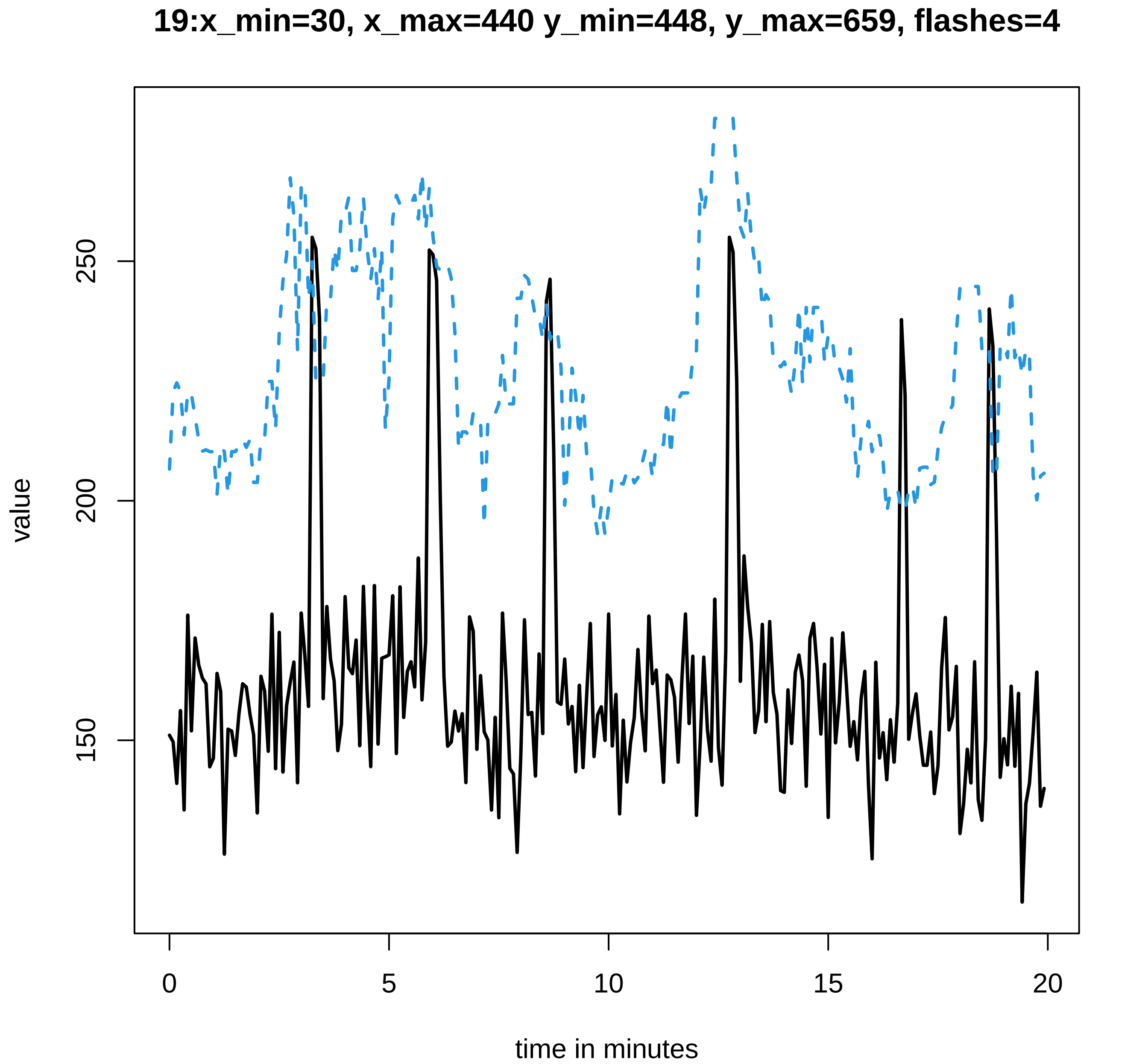
19:x_min=30, x_max=440 y_min=448, y_max=659, flashes=4
value
250
200
150
0
5
10
15
20
time in minutes

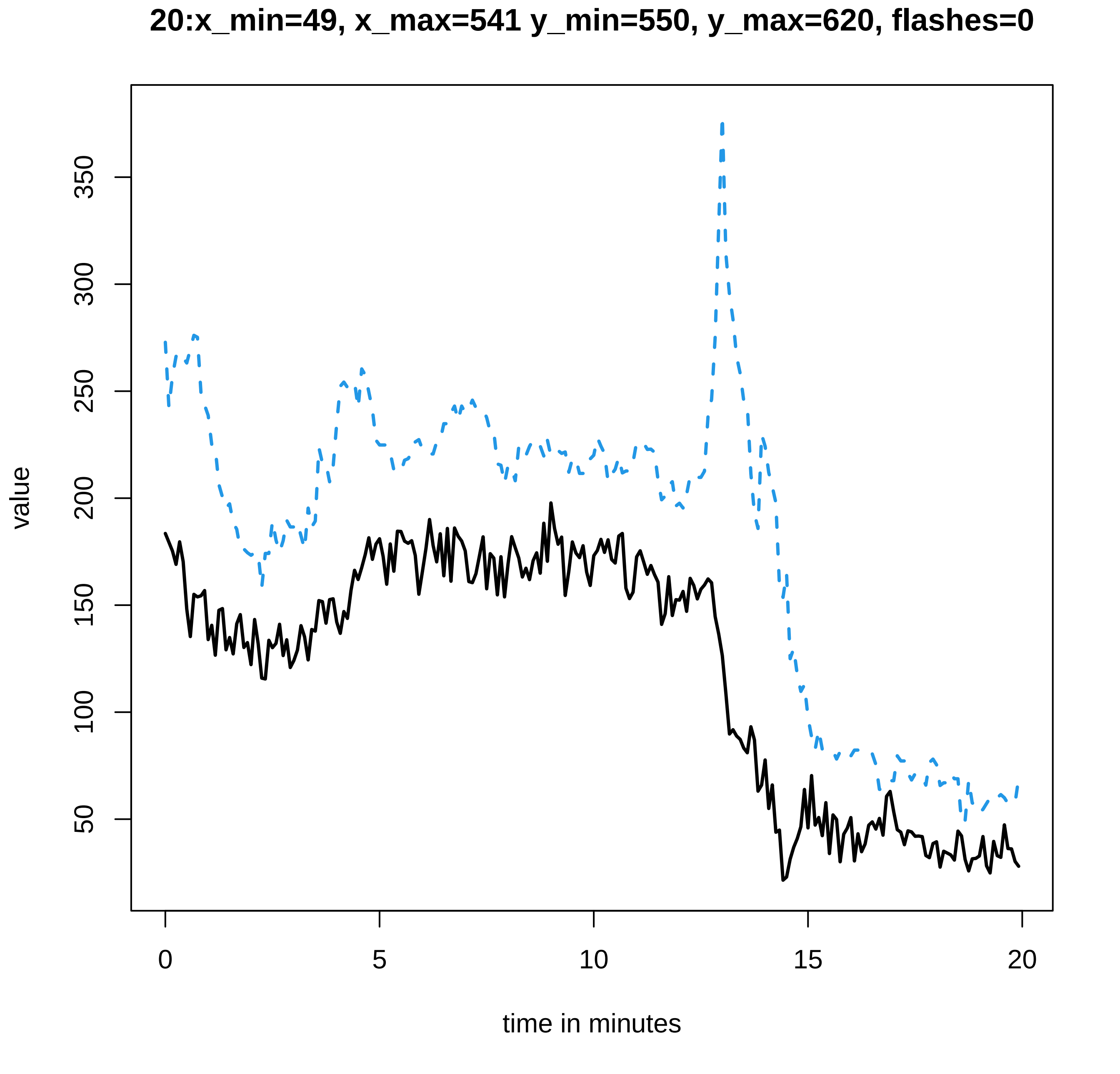

20:x_min=49, x_max=541 y_min=550, y_max=620, flashes=0
value
350
300
250
200
150
100
50
0
5
10
15
20
time in minutes

### 21:x_min=55, x_max=558 y_min=570, y_max=646, flashes=0

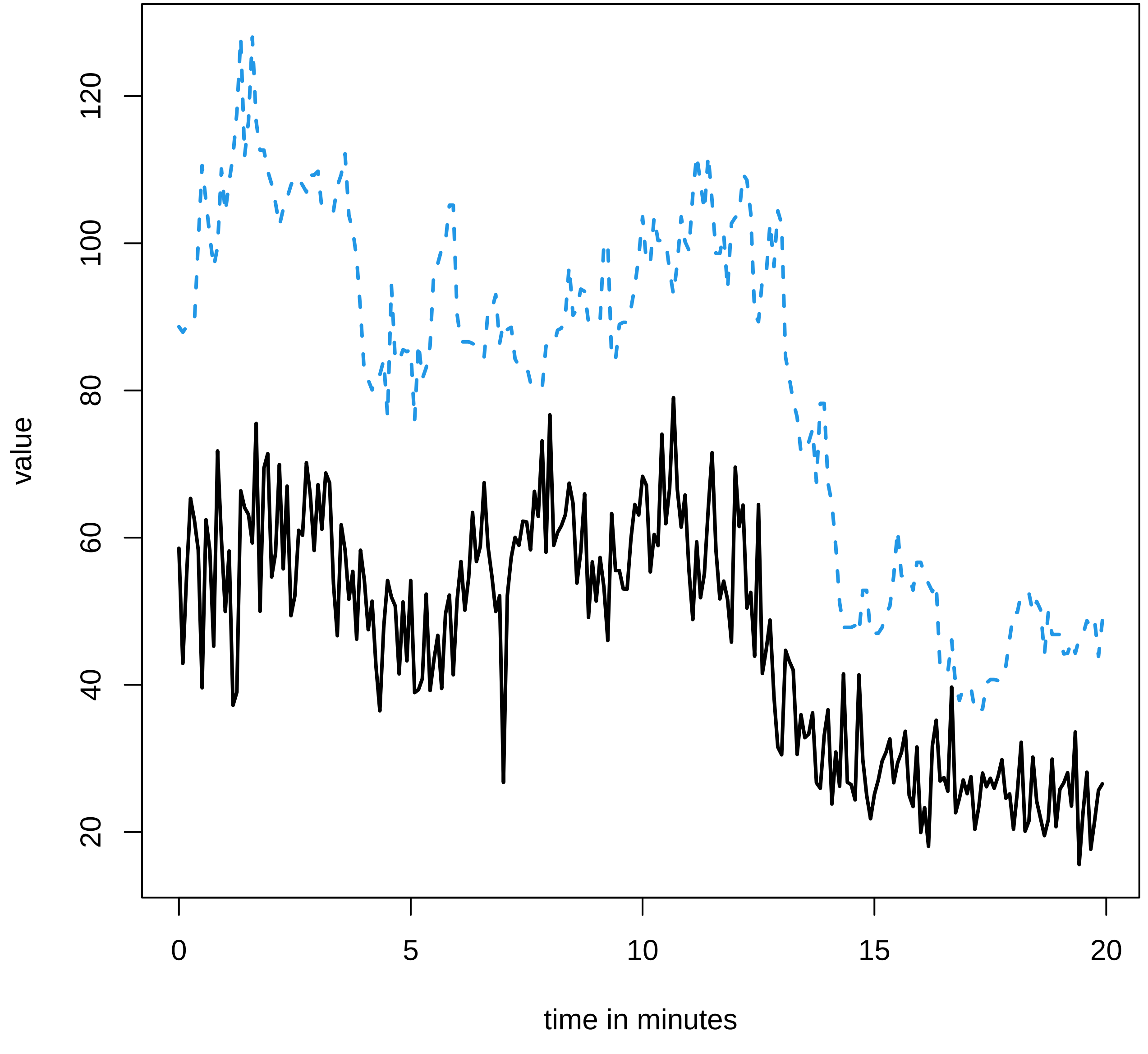

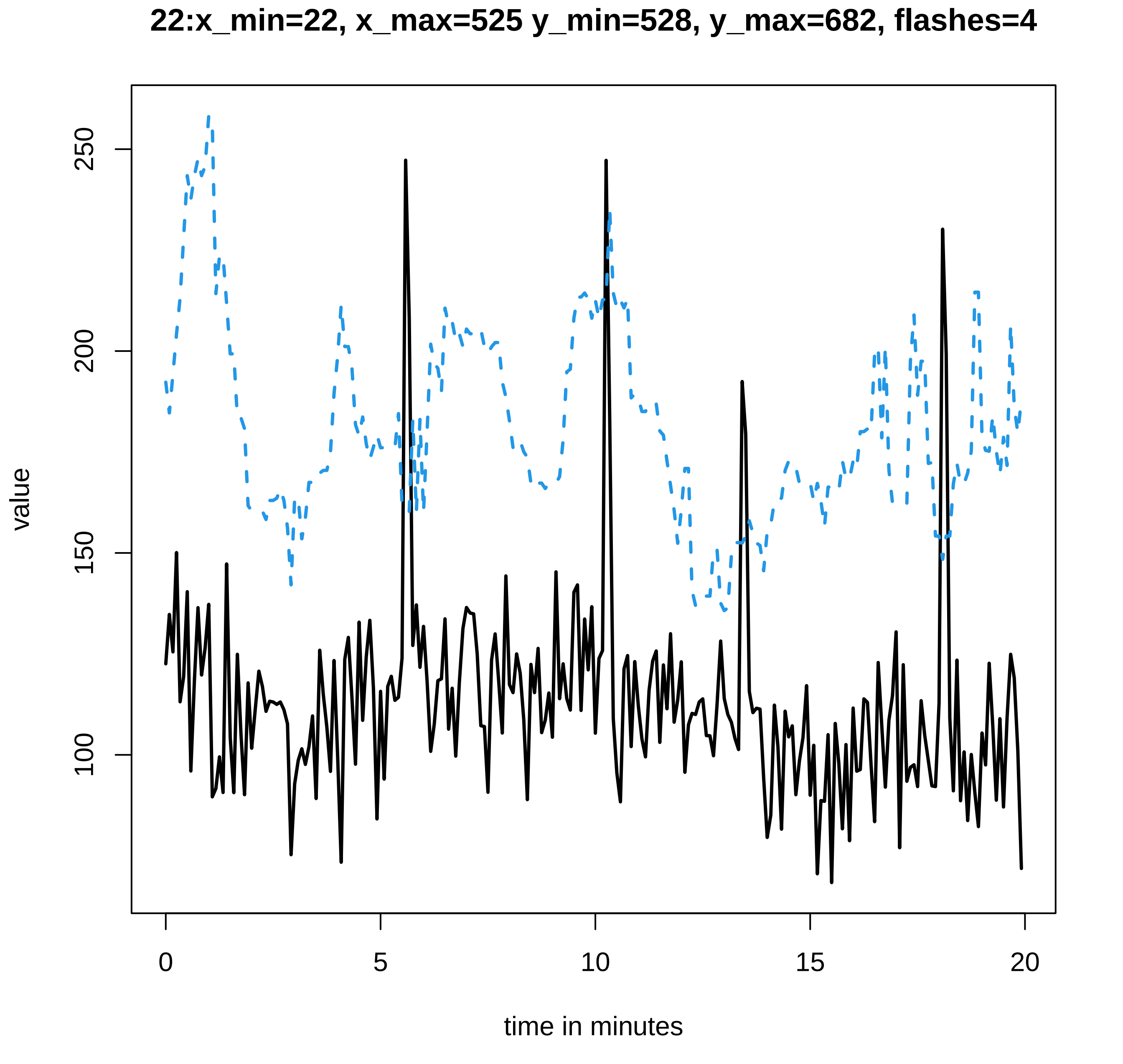
22:x_min=22, x_max=525 y_min=528, y_max=682, flashes=4
value
250
200
150
100
0
5
10
15
20
time in minutes

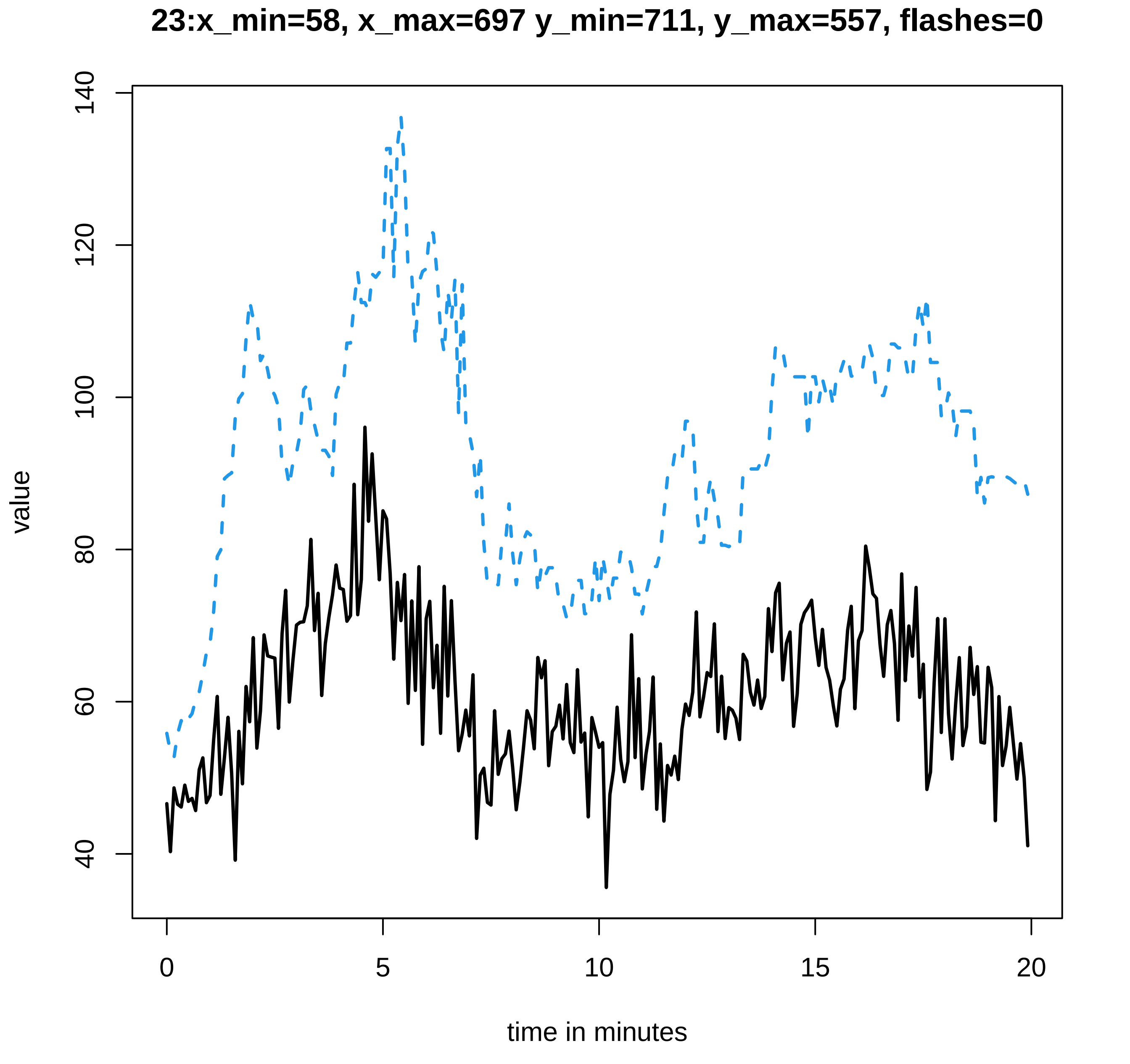
23:x_min=58, x_max=697 y_min=711, y_max=557, flashes=0
value
time in minutes
0
5
10
15
20
40
60
80
100
120
140

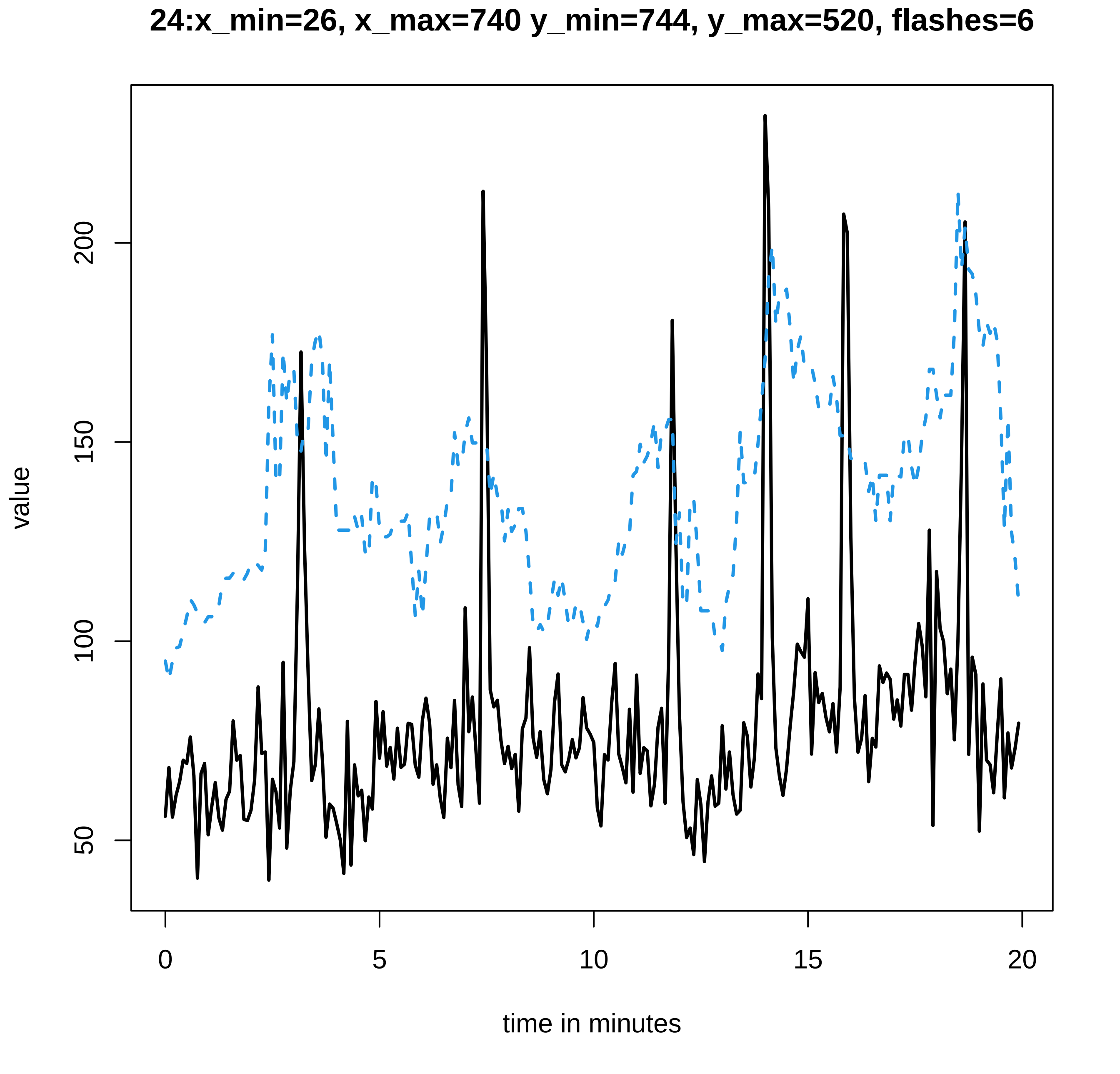

24:x_min=26, x_max=740 y_min=744, y_max=520, flashes=6
value
50
100
150
200
0
5
10
15
20
time in minutes

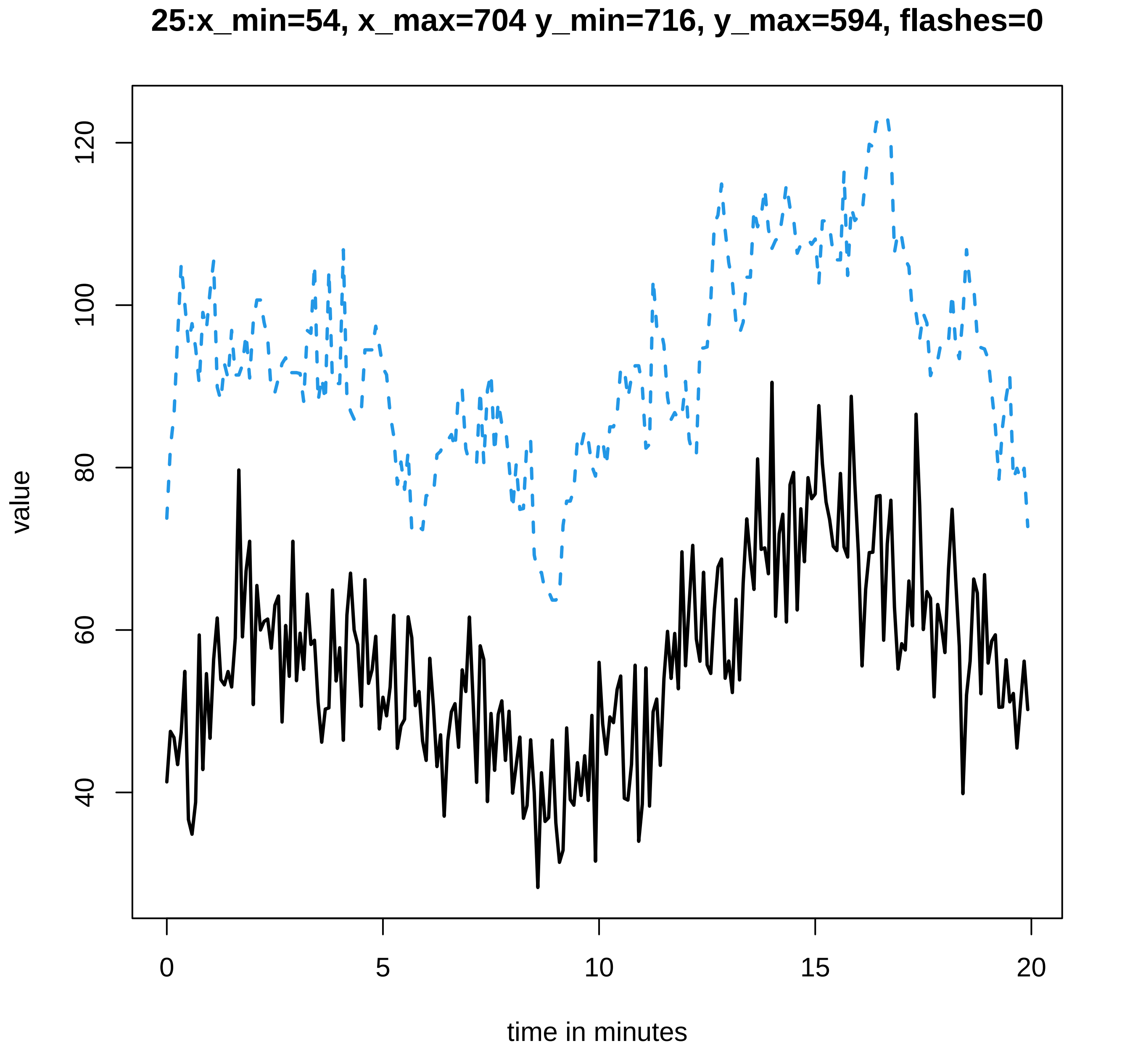
25:x_min=54, x_max=704 y_min=716, y_max=594, flashes=0
value
40
60
80
100
120
0
5
10
15
20
time in minutes

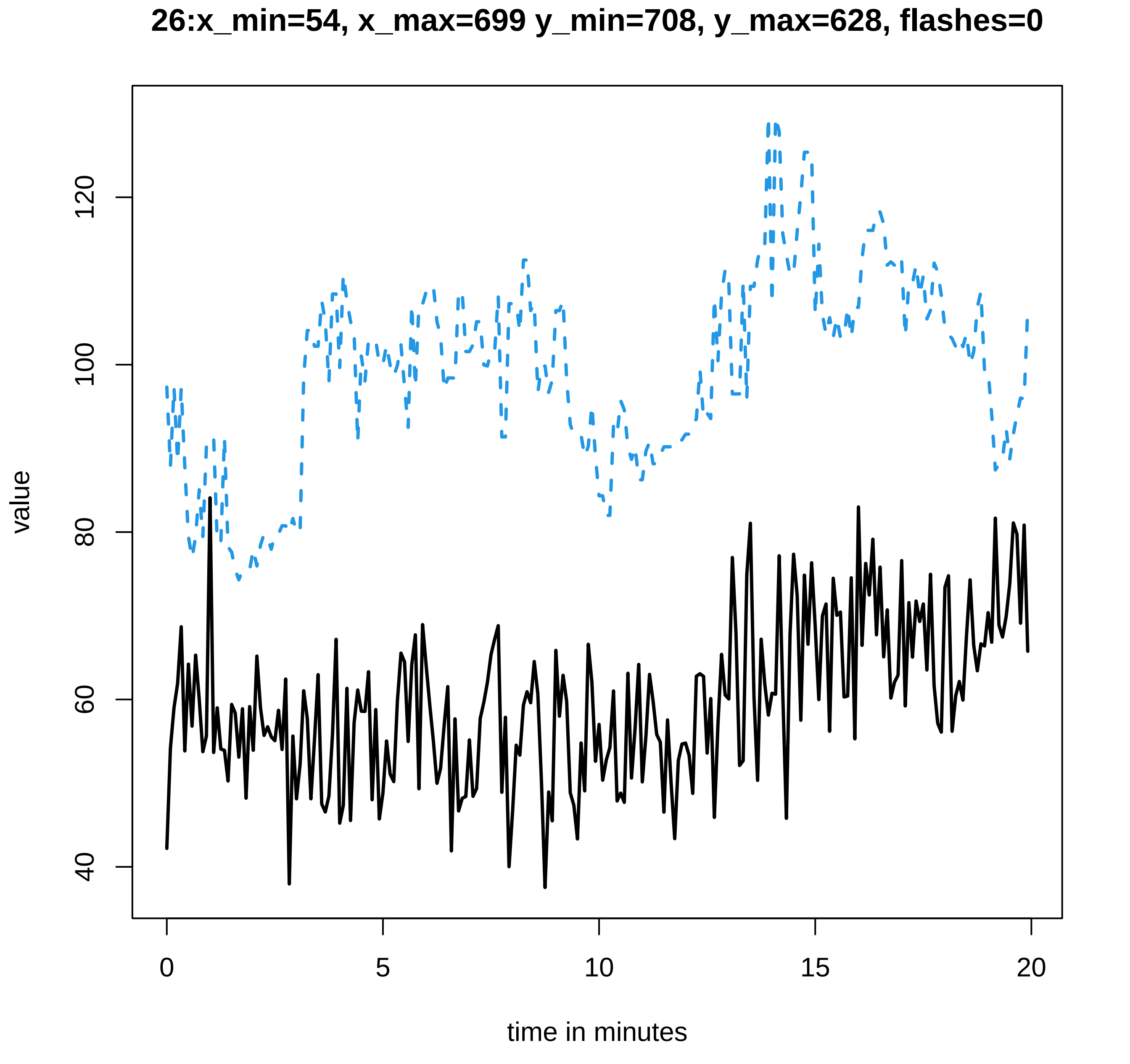
26:x_min=54, x_max=699 y_min=708, y_max=628, flashes=0
value
40
60
80
100
120
0
5
10
15
20
time in minutes

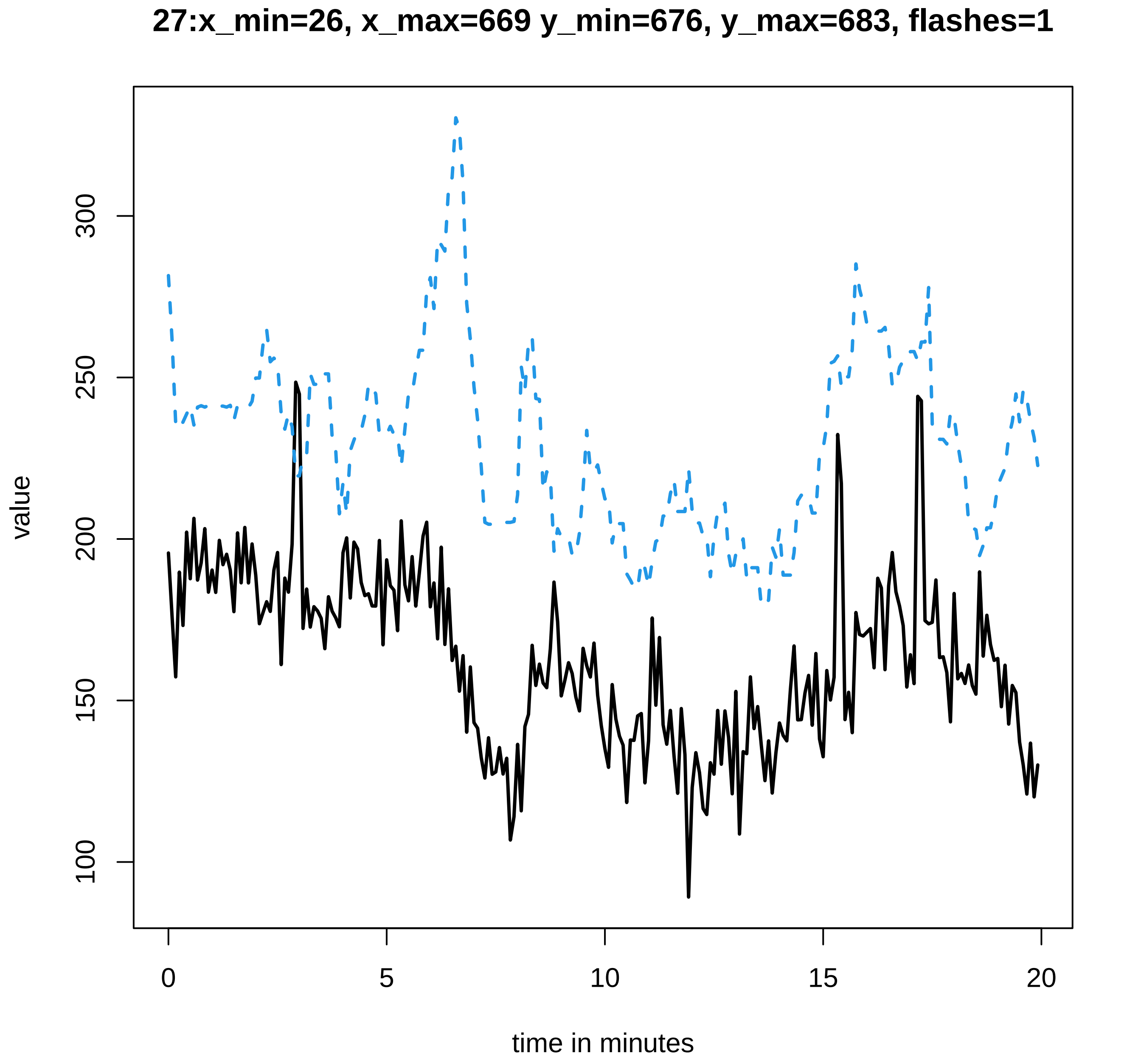
27:x_min=26, x_max=669 y_min=676, y_max=683, flashes=1
value
100
150
200
250
300
0
5
10
15
20
time in minutes

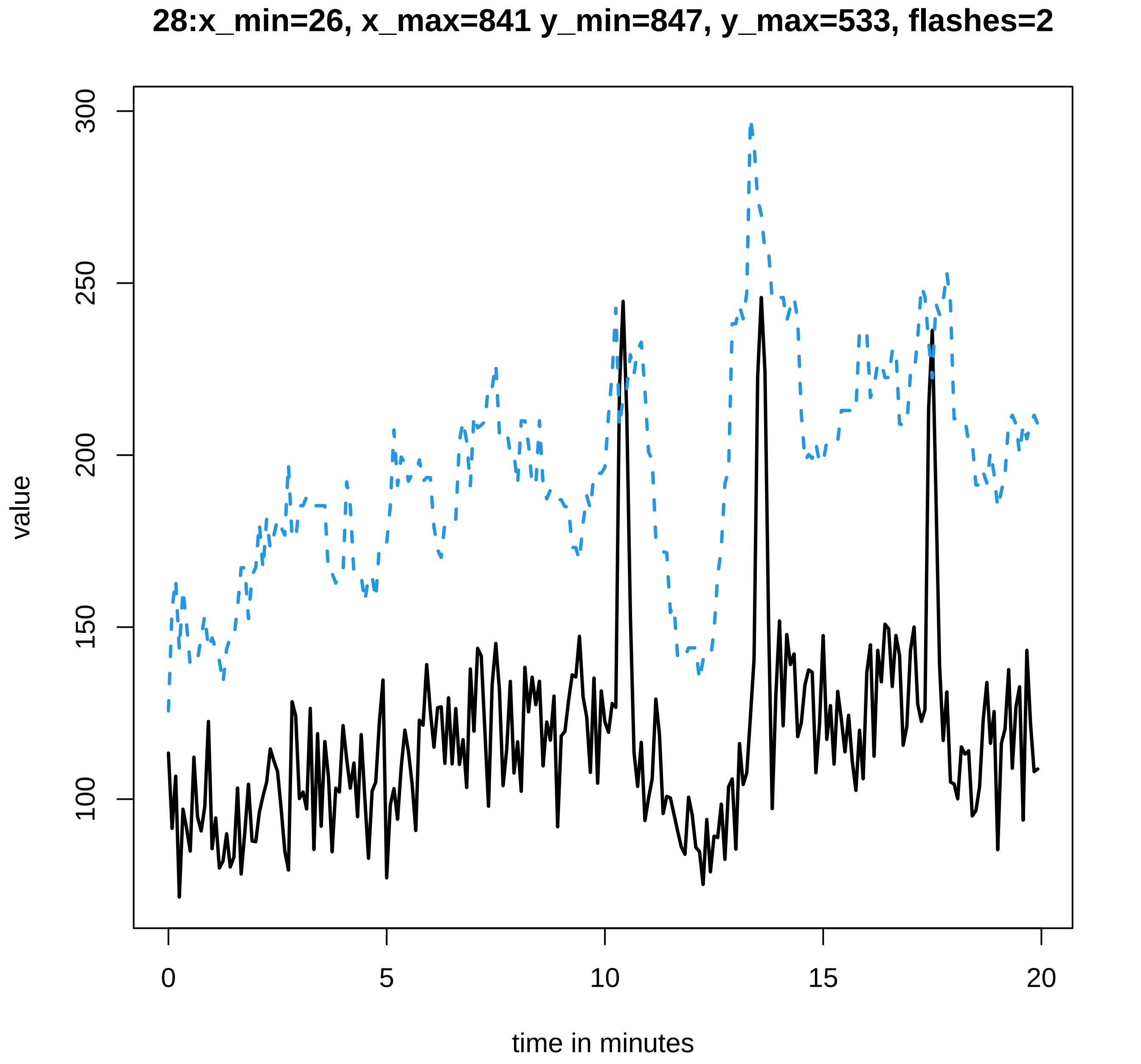
28:x_min=26, x_max=841 y_min=847, y_max=533, flashes=2
value
300
250
200
150
100
0
5
10
15
20
time in minutes

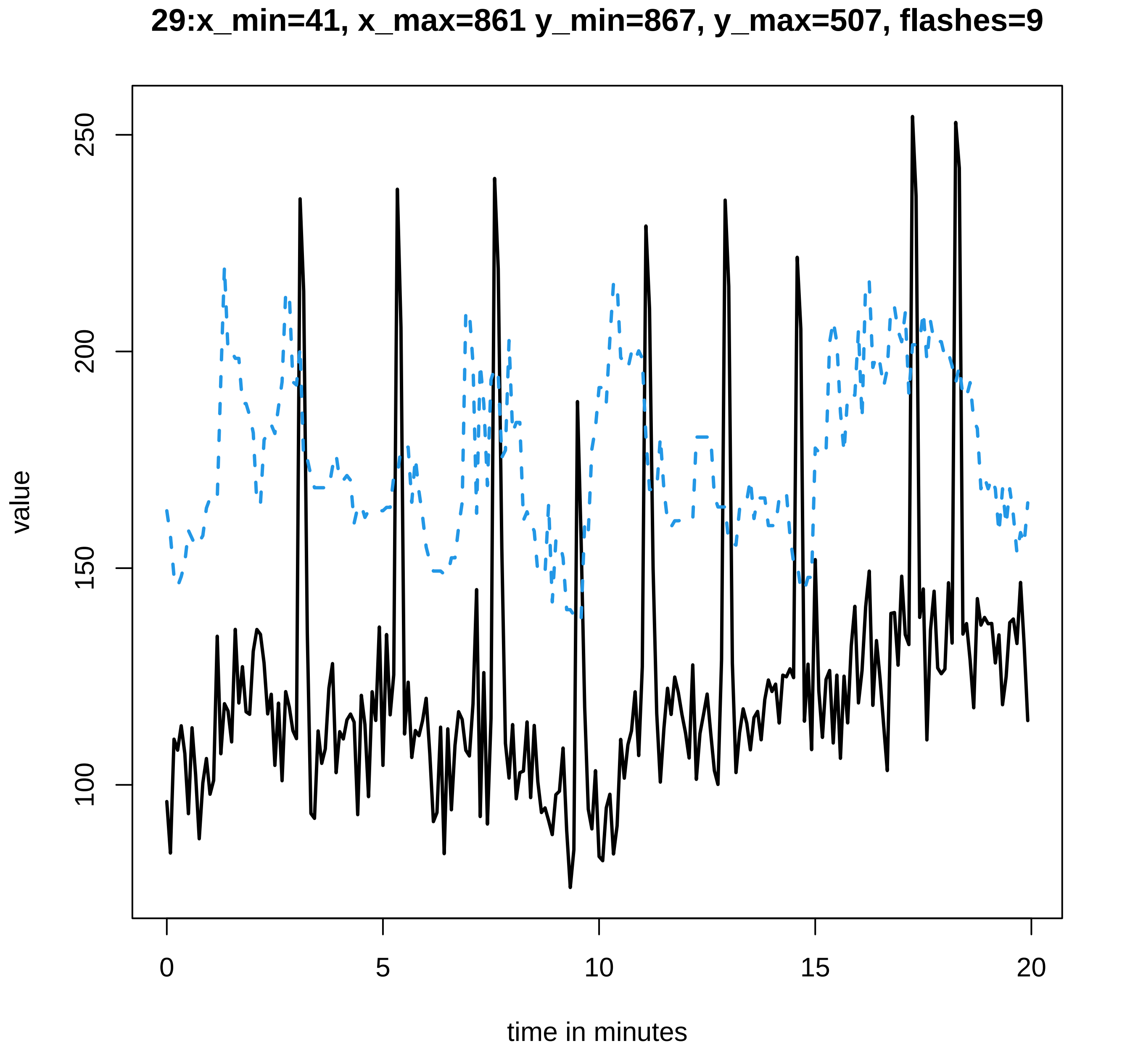
29:x_min=41, x_max=861 y_min=867, y_max=507, flashes=9
value
100
150
200
250
0
5
10
15
20
time in minutes

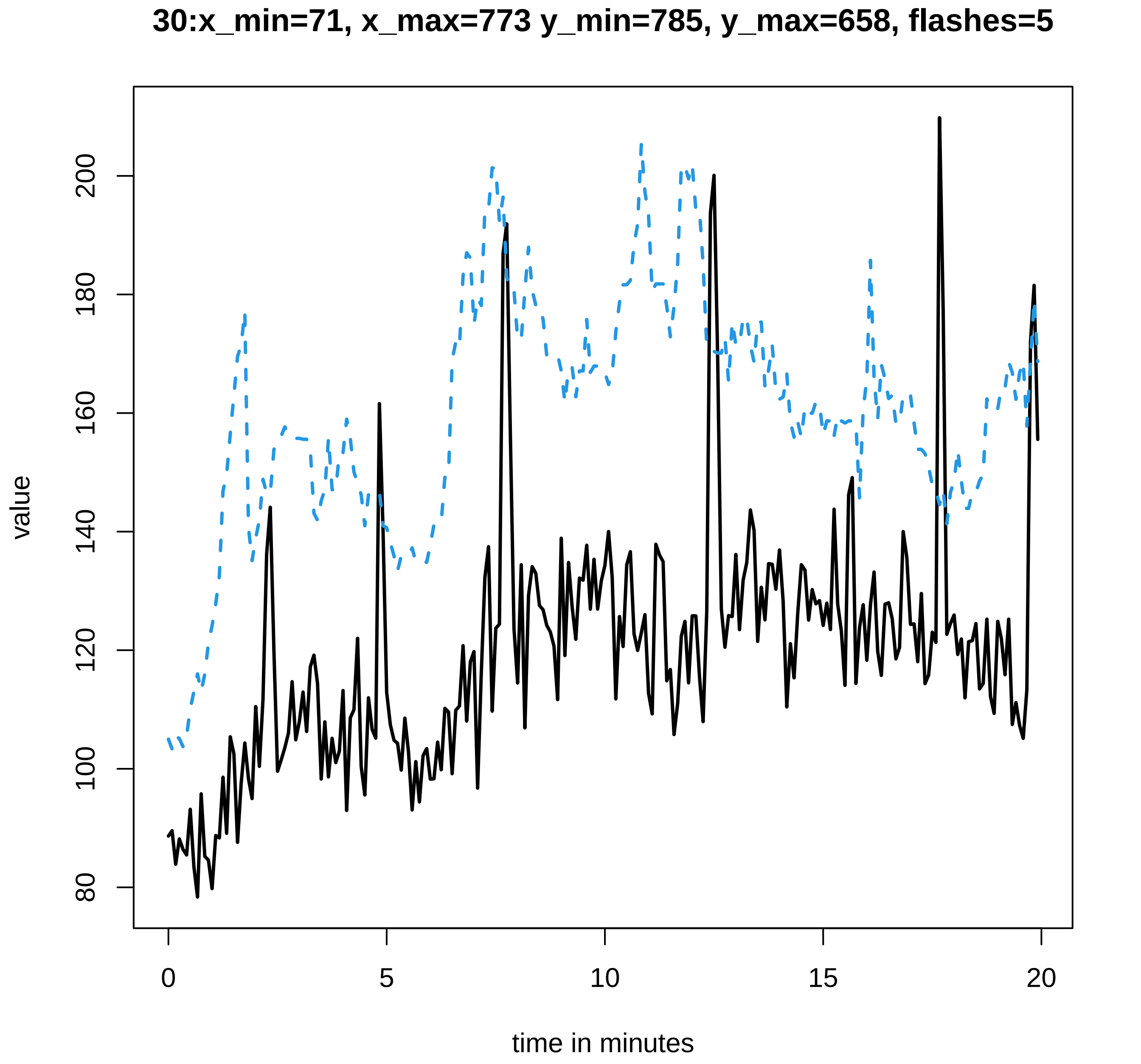

30:x_min=71, x_max=773 y_min=785, y_max=658, flashes=5
value
time in minutes
80
100
120
140
160
180
200
0
5
10
15
20

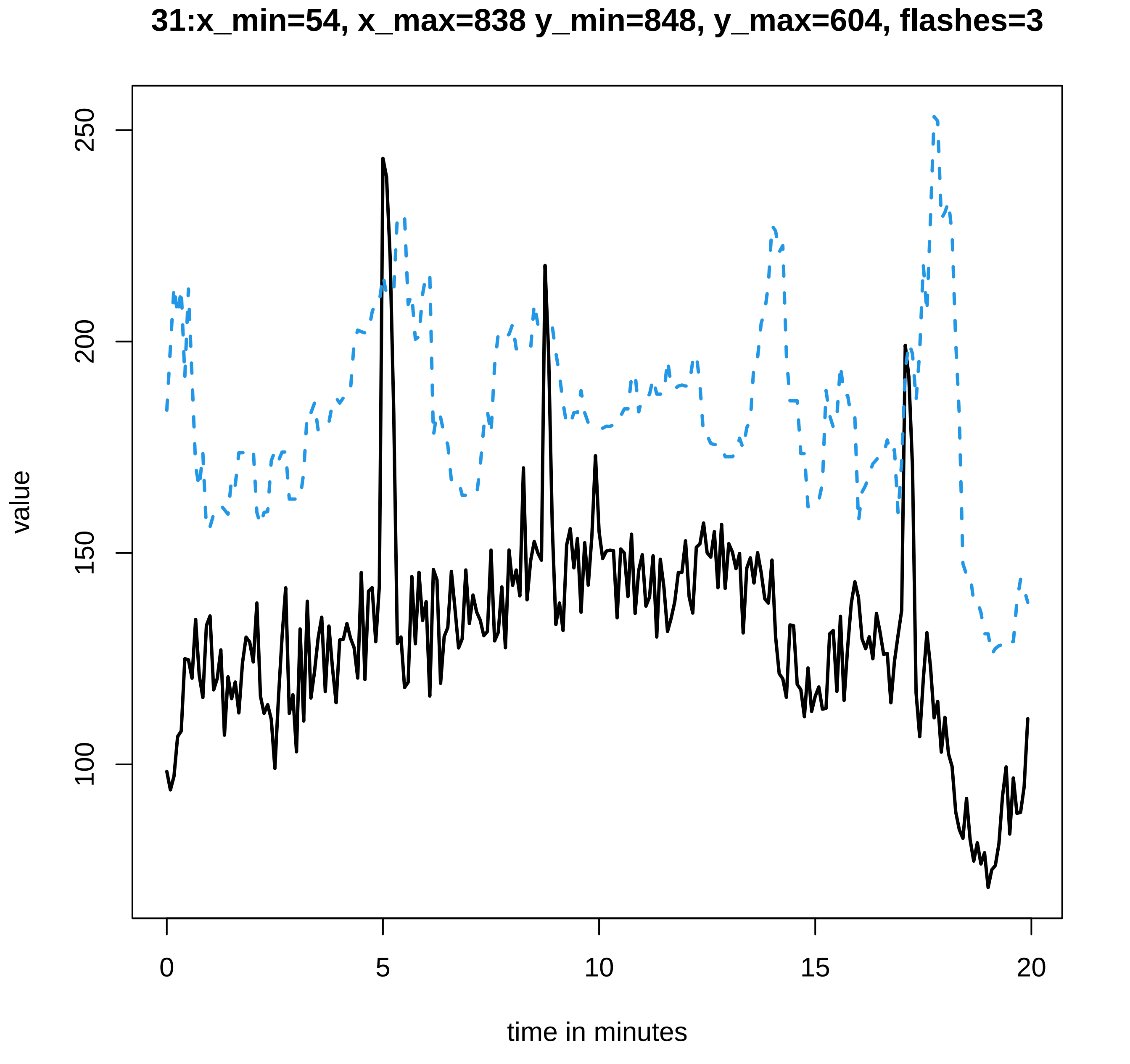
31:x_min=54, x_max=838 y_min=848, y_max=604, flashes=3
value
250
200
150
100
0
5
10
15
20
time in minutes

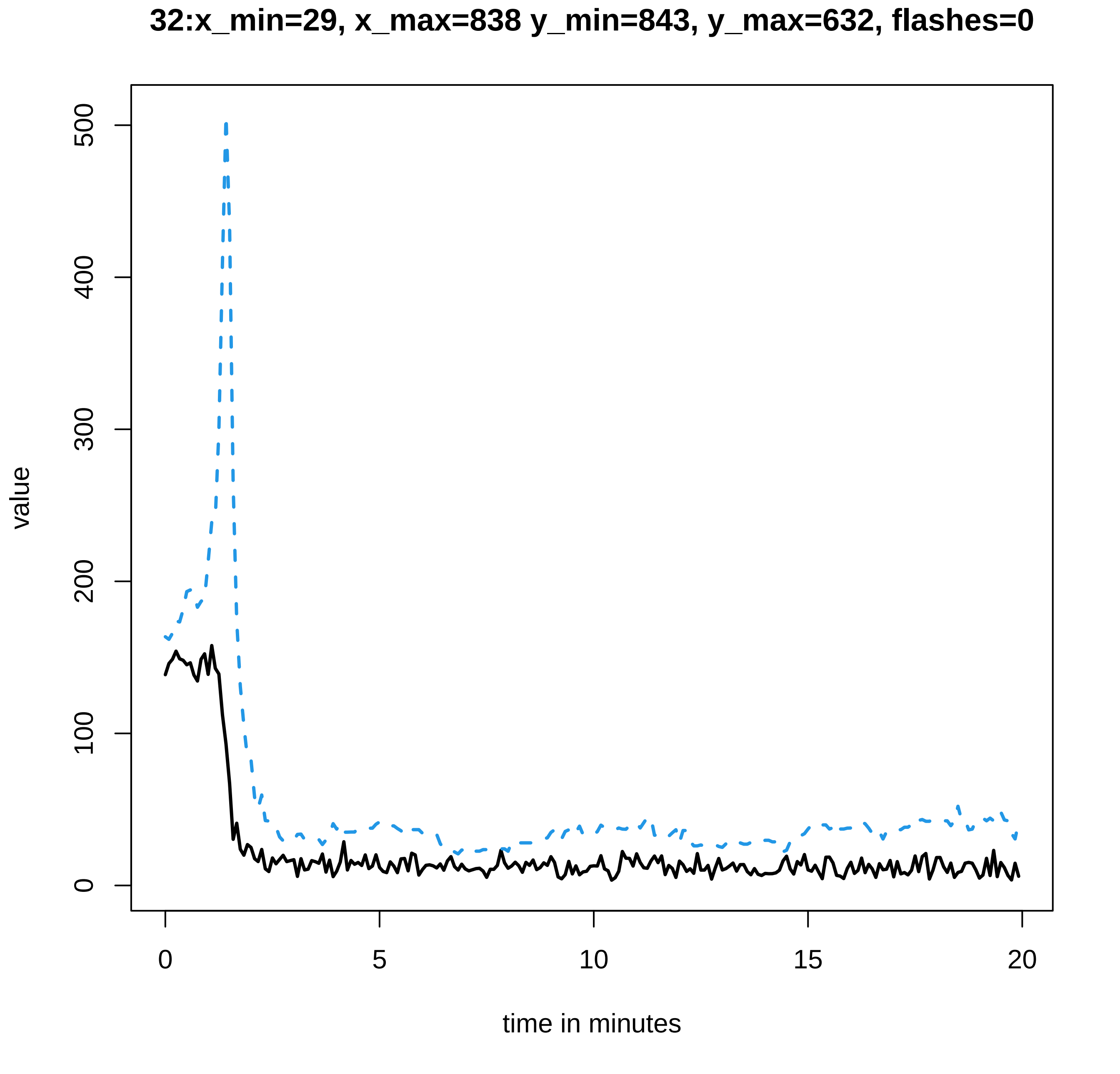

32:x_min=29, x_max=838 y_min=843, y_max=632, flashes=0
value
0
100
200
300
400
500
0
5
10
15
20
time in minutes

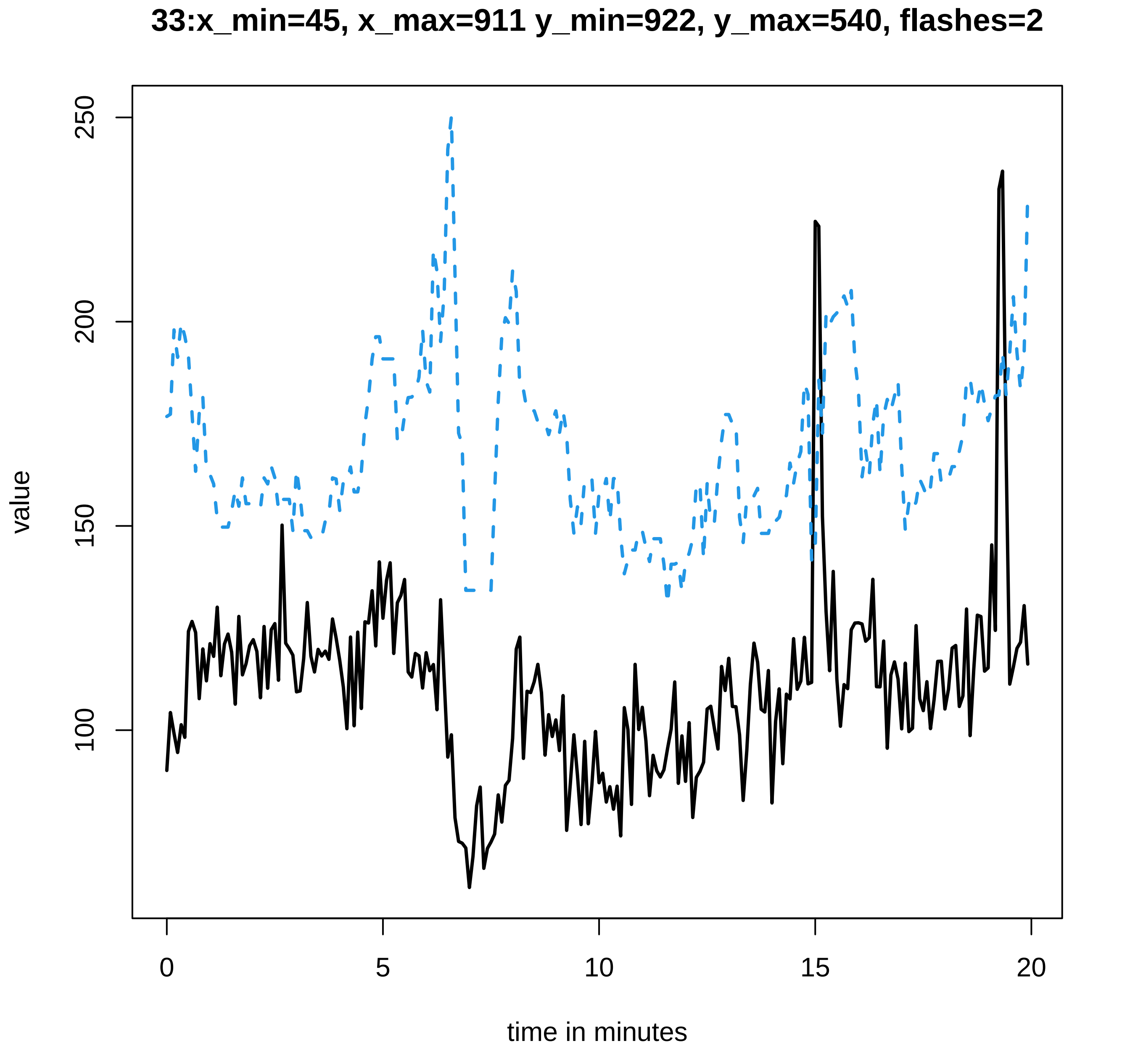
33:x_min=45, x_max=911 y_min=922, y_max=540, flashes=2
value
250
200
150
100
0
5
10
15
20
time in minutes

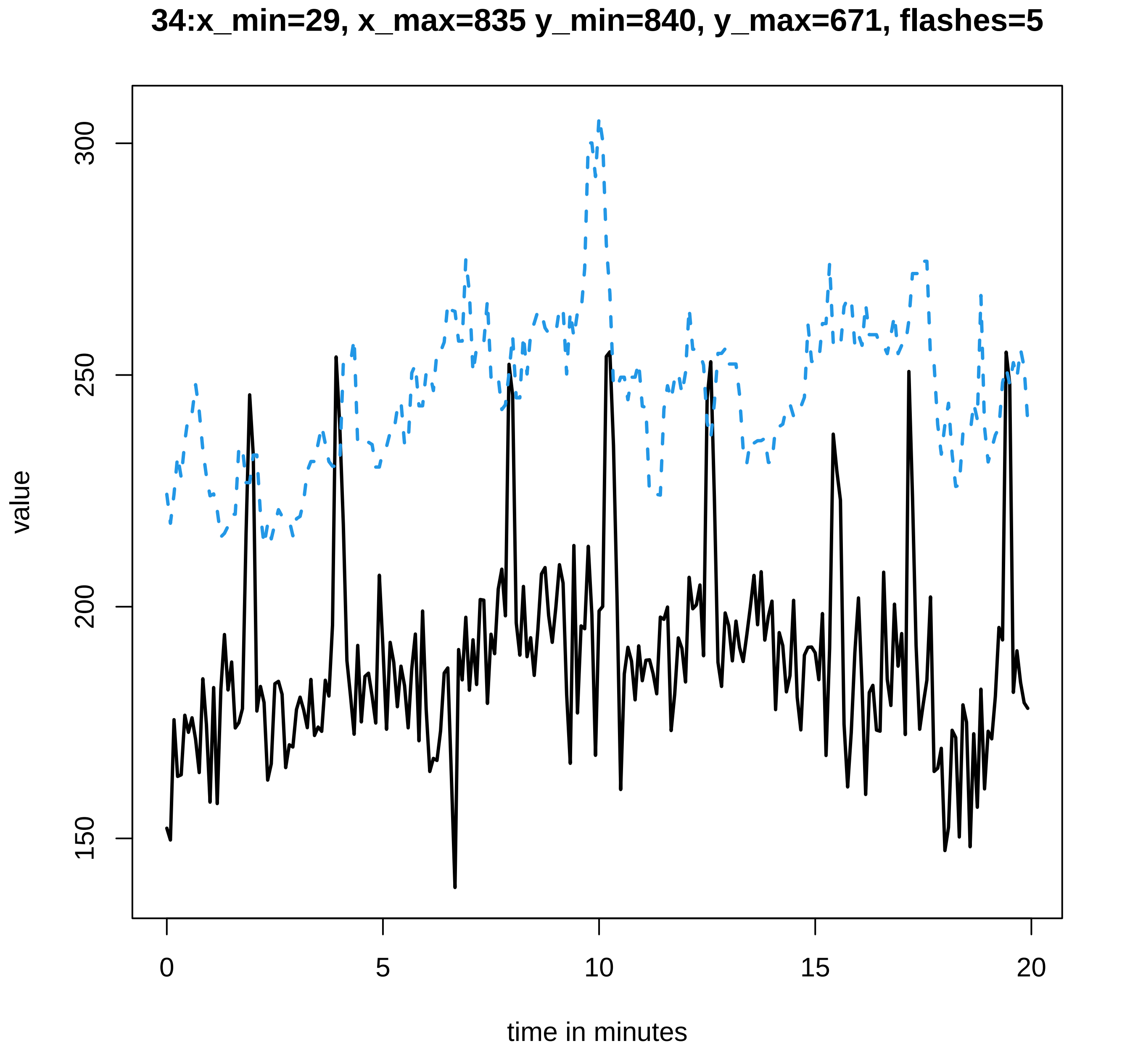

34:x_min=29, x_max=835 y_min=840, y_max=671, flashes=5
value
300
250
200
150
0
5
10
15
20
time in minutes

### 35:x_min=33, x_max=958 y_min=963, y_max=587, flashes=0

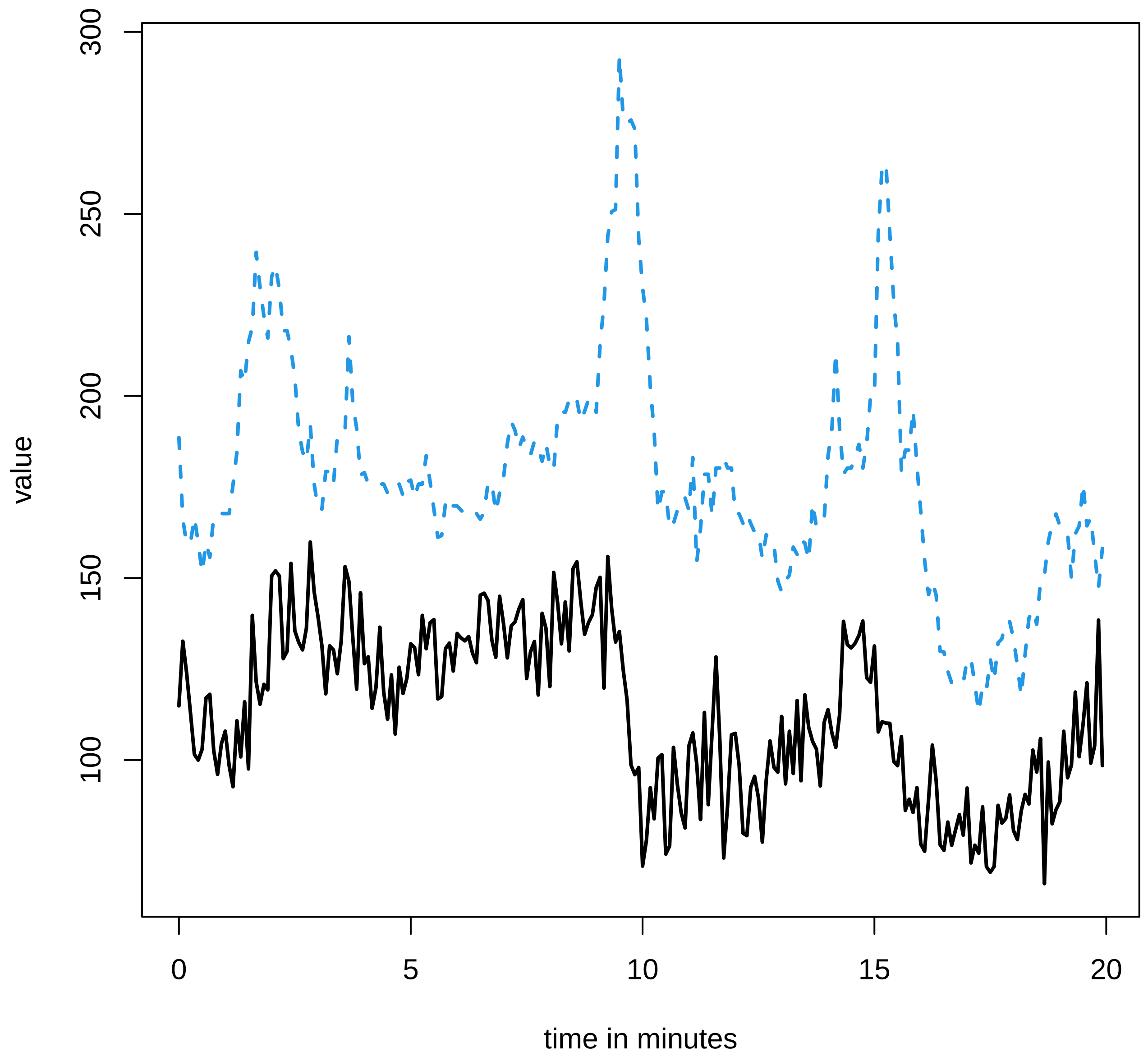

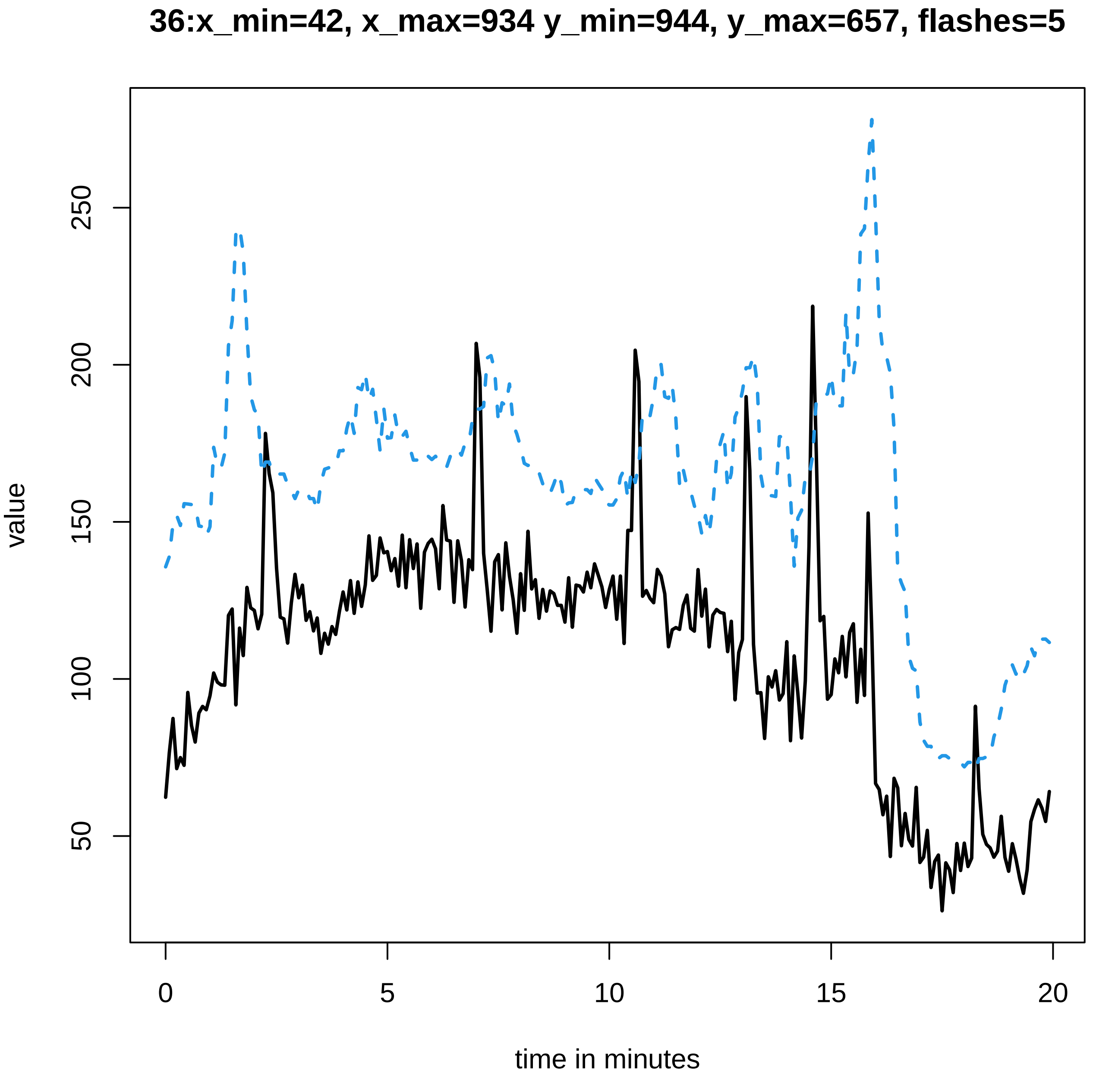
36:x_min=42, x_max=934 y_min=944, y_max=657, flashes=5
value
50
100
150
200
250
0
5
10
15
20
time in minutes

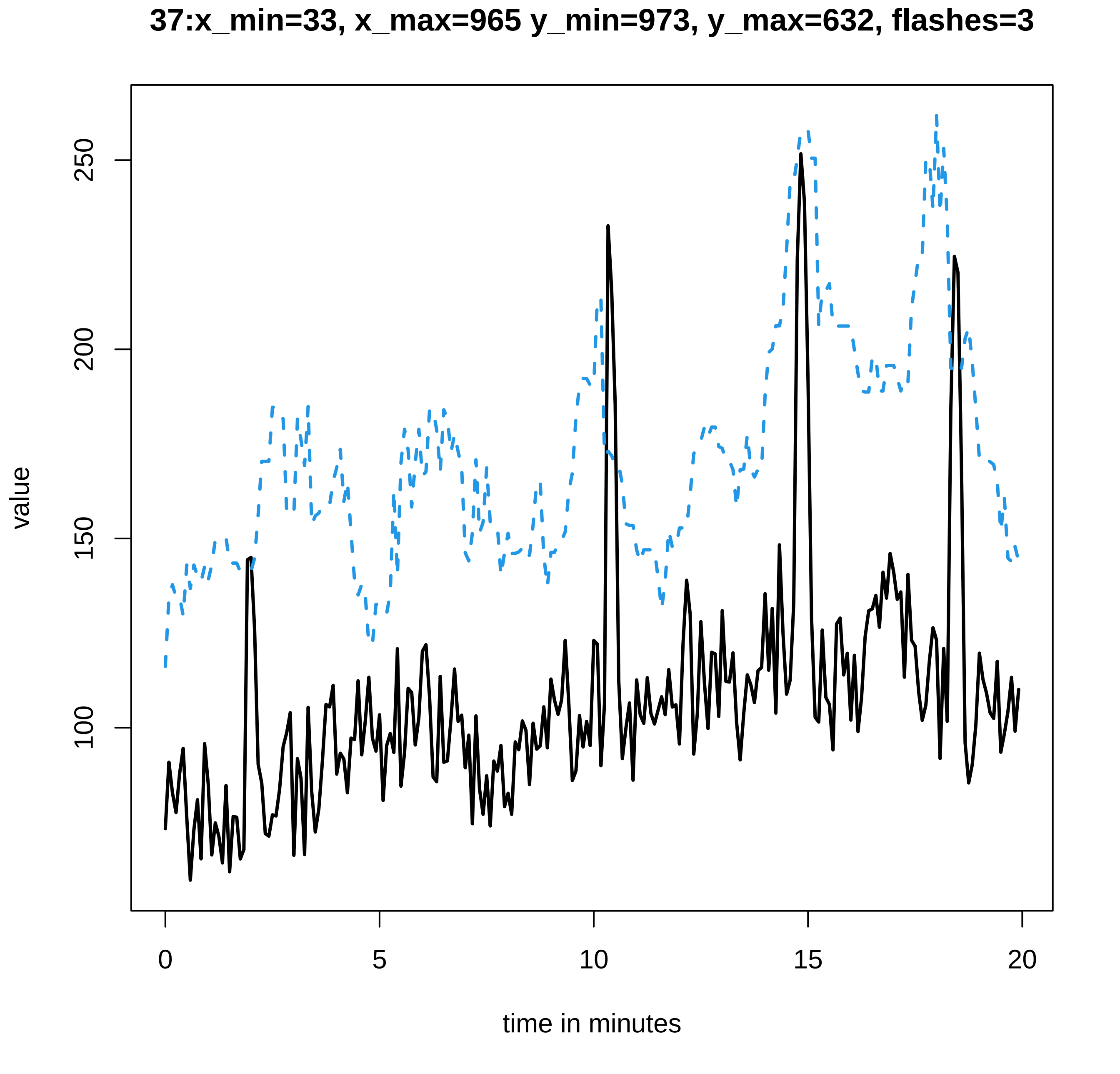

37:x_min=33, x_max=965 y_min=973, y_max=632, flashes=3
value
100
150
200
250
0
5
10
15
20
time in minutes
