## Supplementary data Calcium spiking for "LysM Receptor Proteins are Required for Ectomycorrhizal Symbiosis in Poplar": profiles-spiking-10.pdf

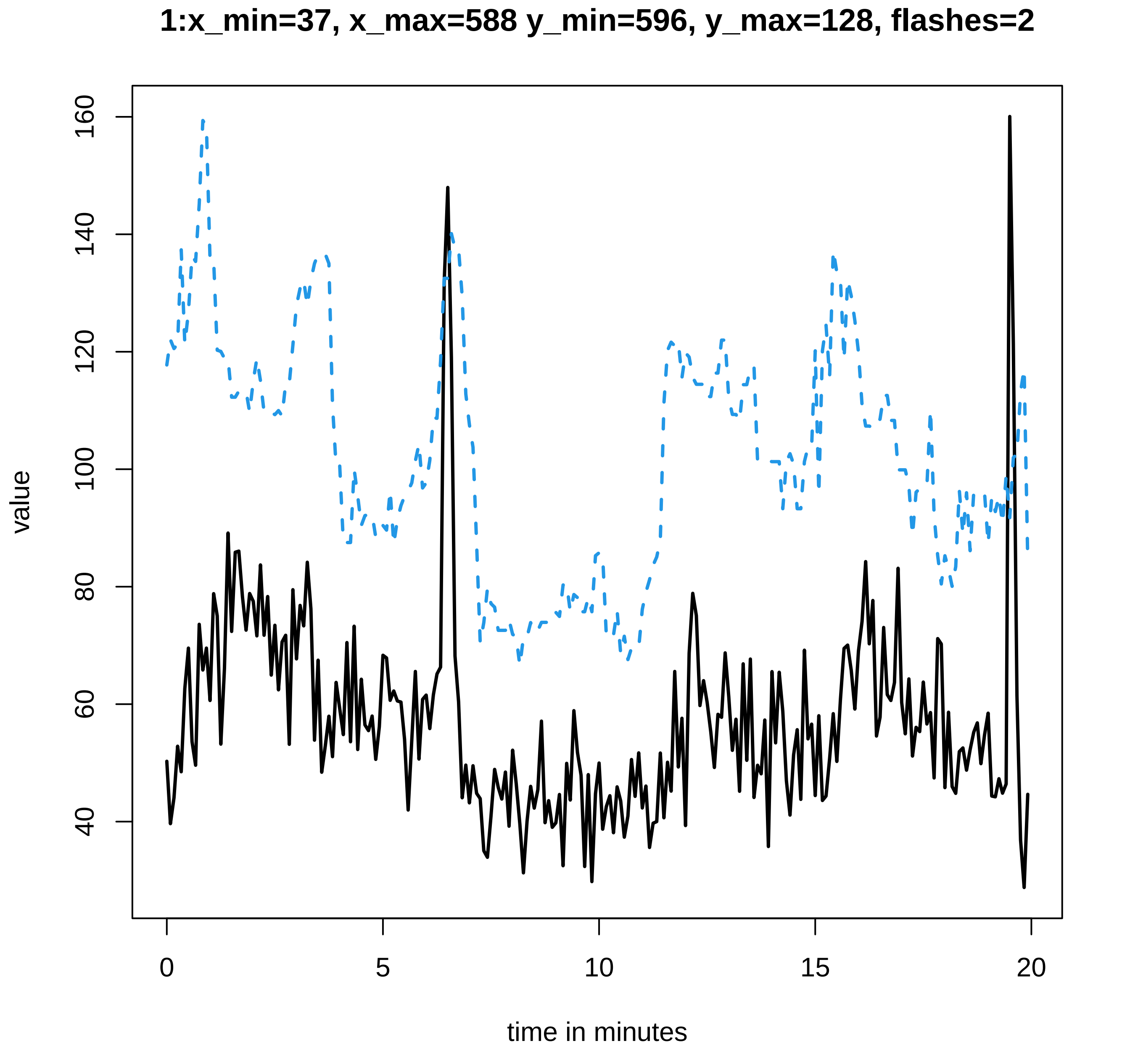
1:x_min=37, x_max=588 y_min=596, y_max=128, flashes=2
value
160
140
120
100
80
60
40
0
5
10
15
20
time in minutes

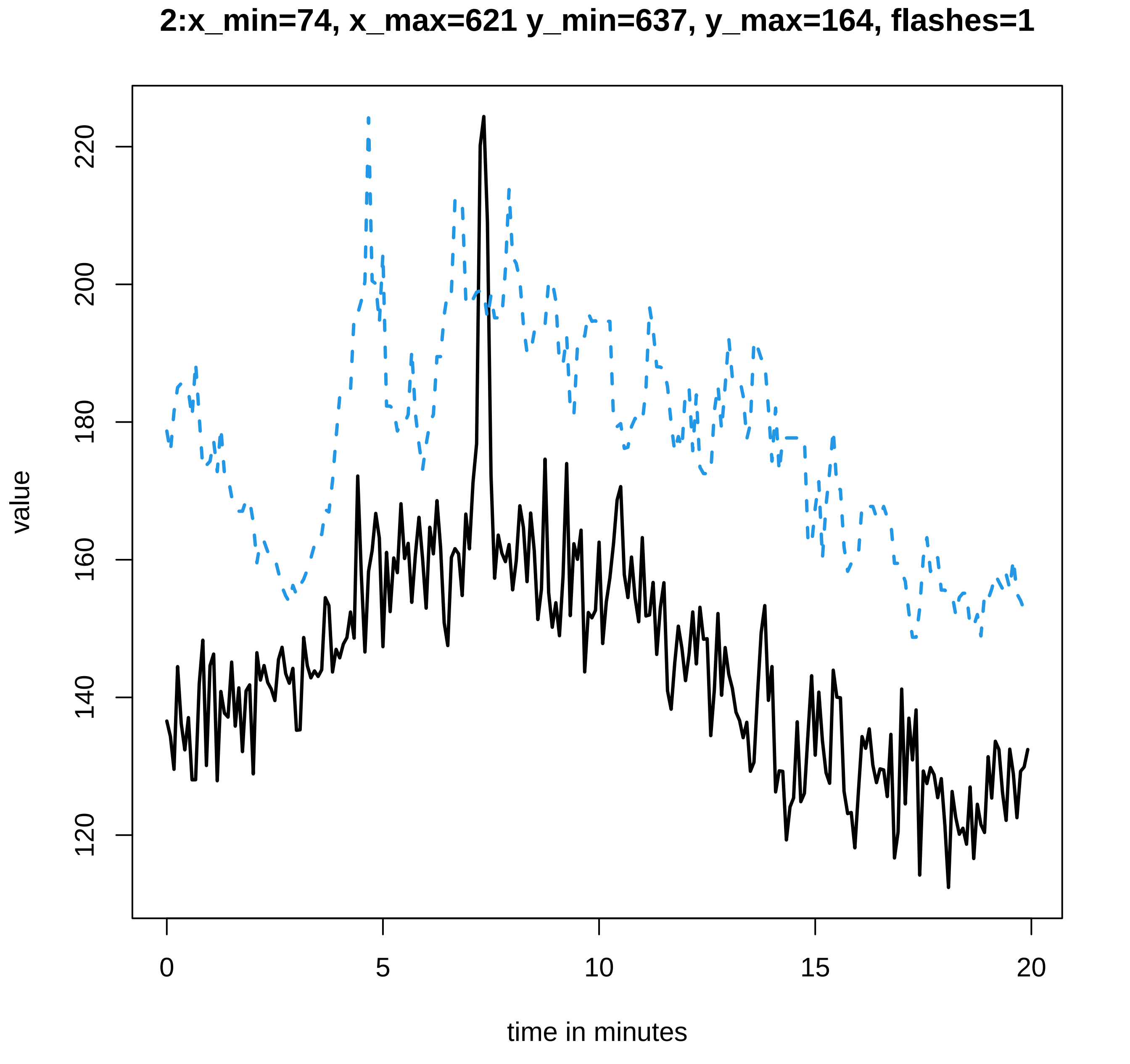
2:x_min=74, x_max=621 y_min=637, y_max=164, flashes=1
value
120
140
160
180
200
220
0
5
10
15
20
time in minutes

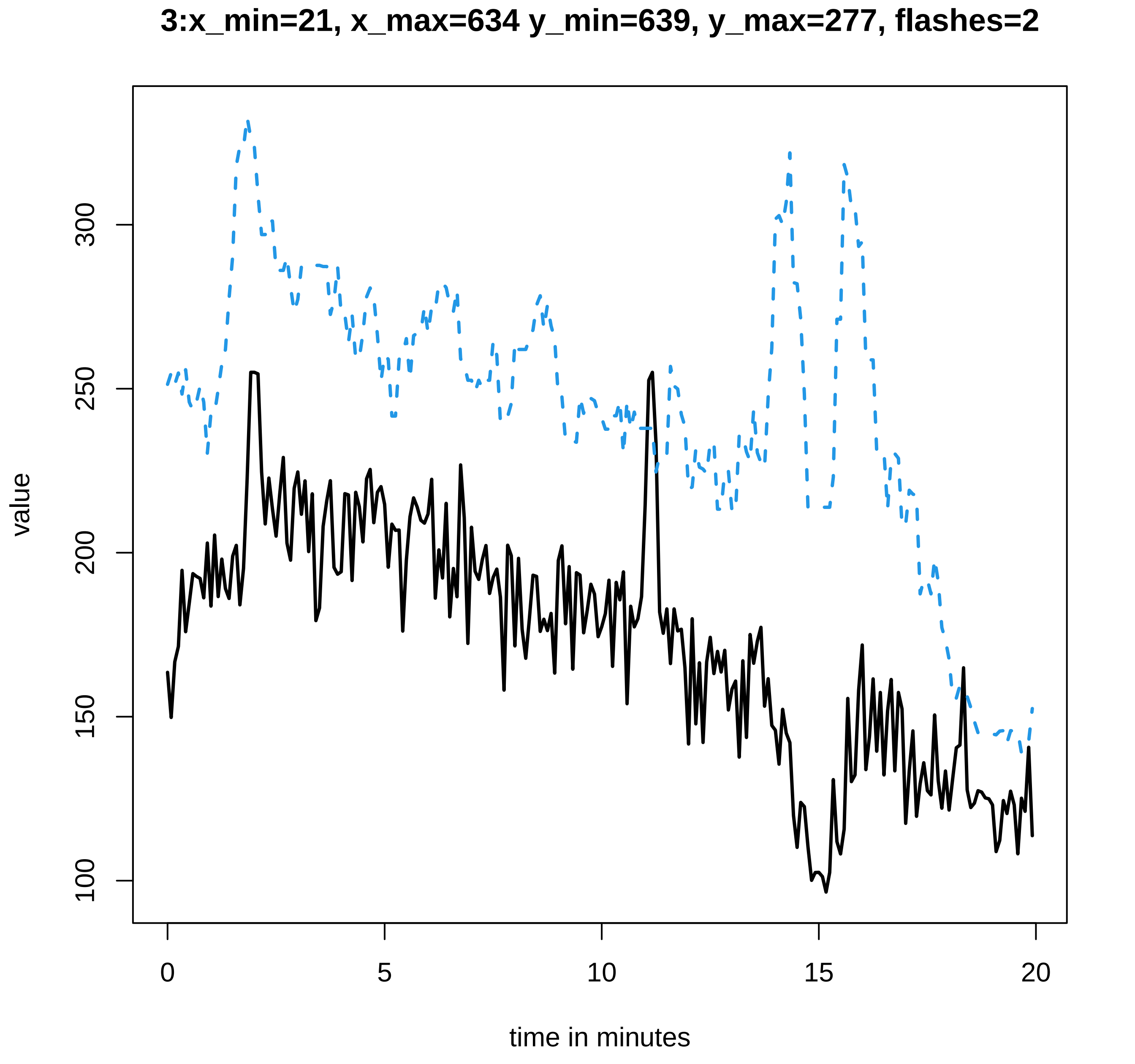

3:x_min=21, x_max=634 y_min=639, y_max=277, flashes=2
value
100
150
200
250
300
0
5
10
15
20
time in minutes

### 4:x_min=51, x_max=688 y_min=695, y_max=67, flashes=0

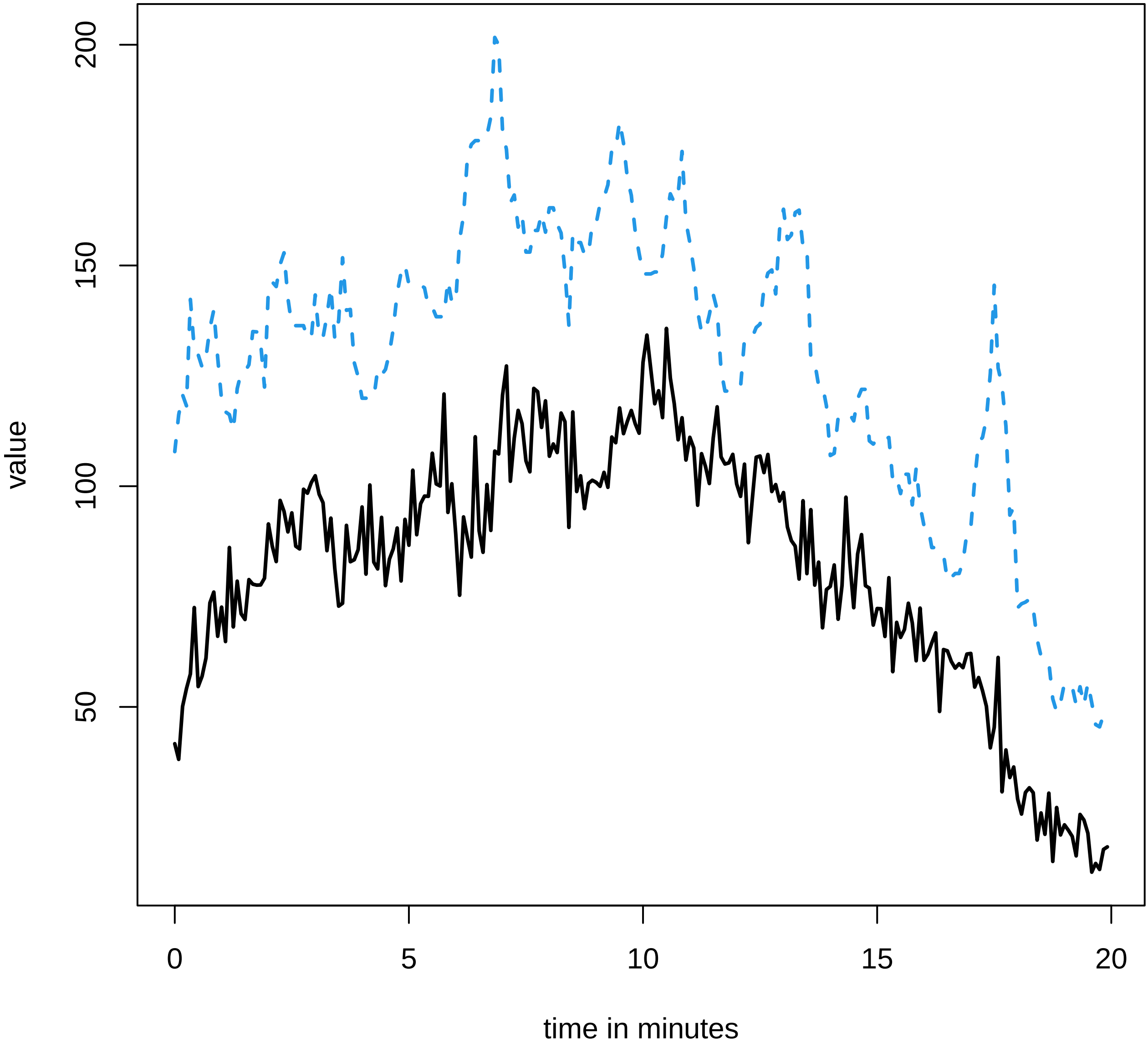

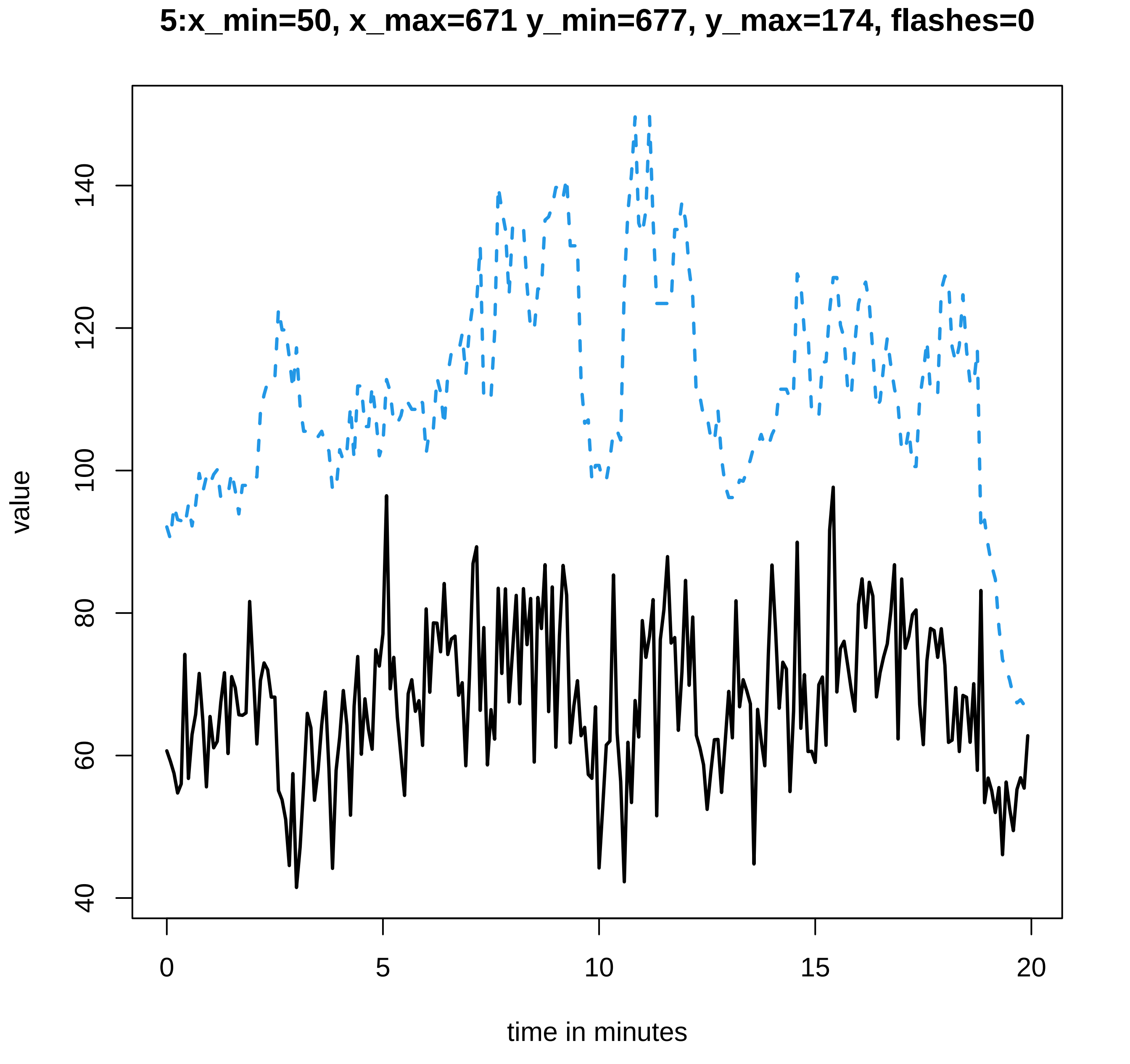
5:x_min=50, x_max=671 y_min=677, y_max=174, flashes=0
value
40
60
80
100
120
140
0
5
10
15
20
time in minutes

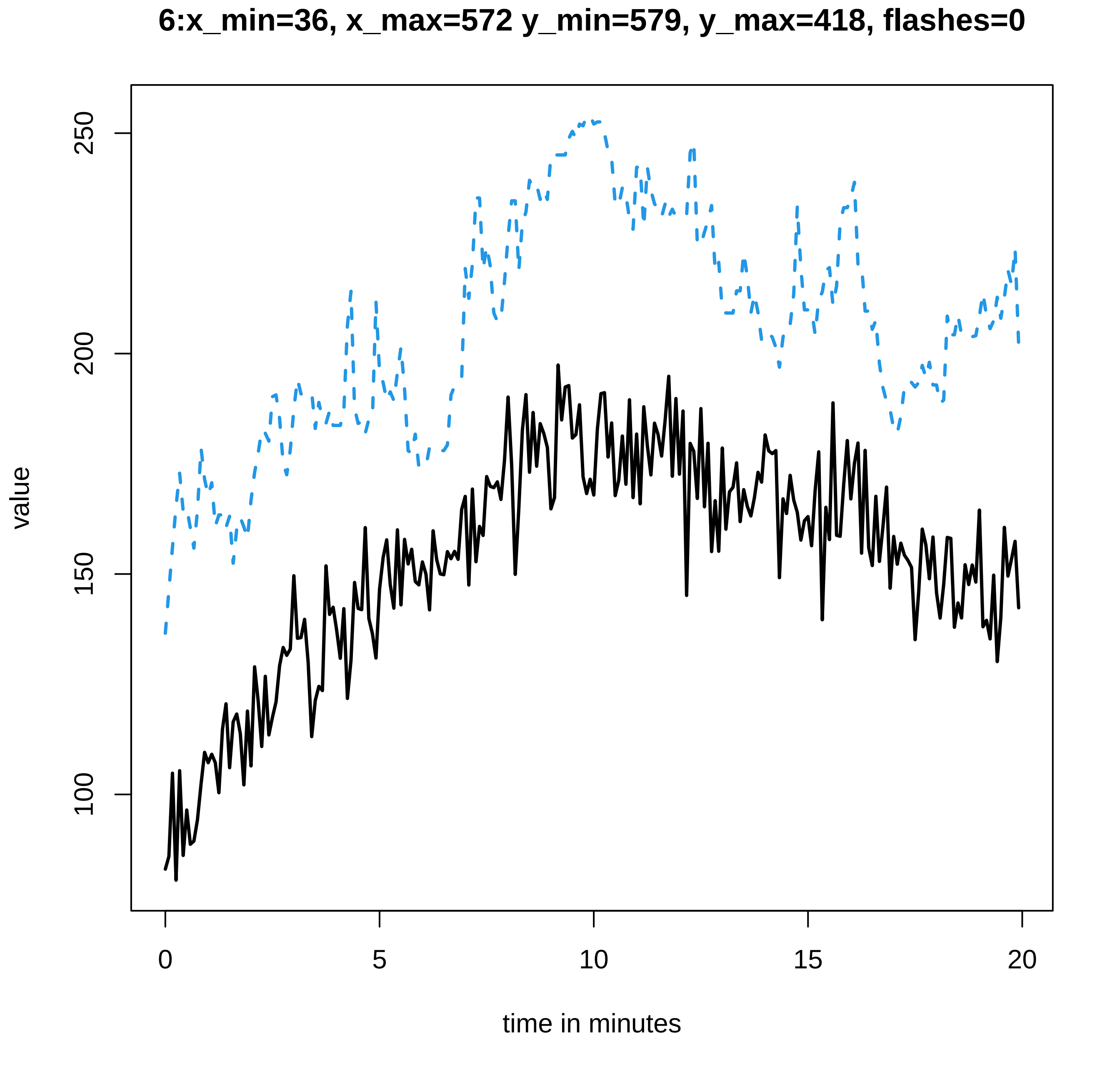
6:x_min=36, x_max=572 y_min=579, y_max=418, flashes=0
value
100
150
200
250
0
5
10
15
20
time in minutes

**7:x_min=50, x_max=625 y_min=632, y_max=348, flashes=0**

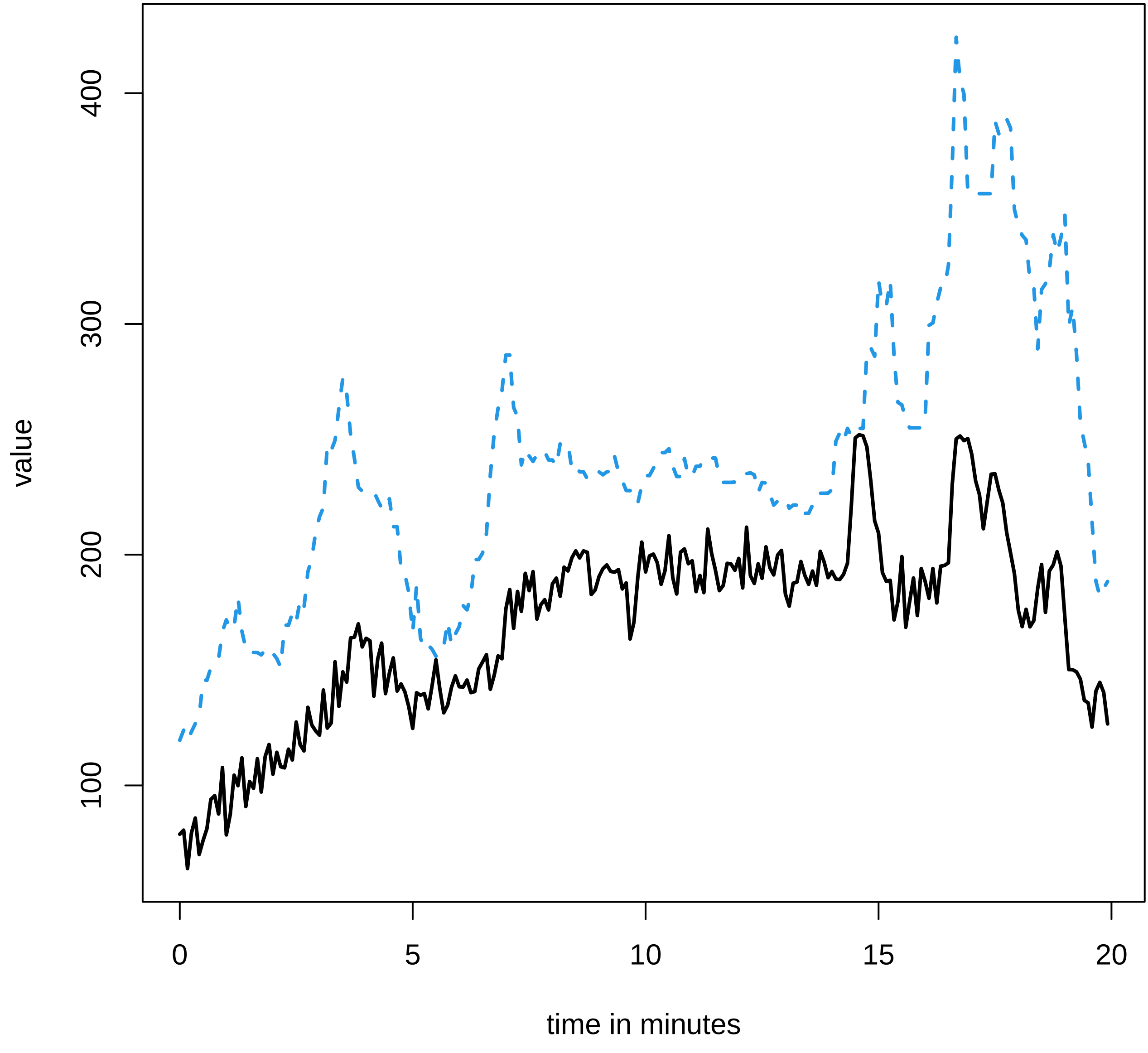

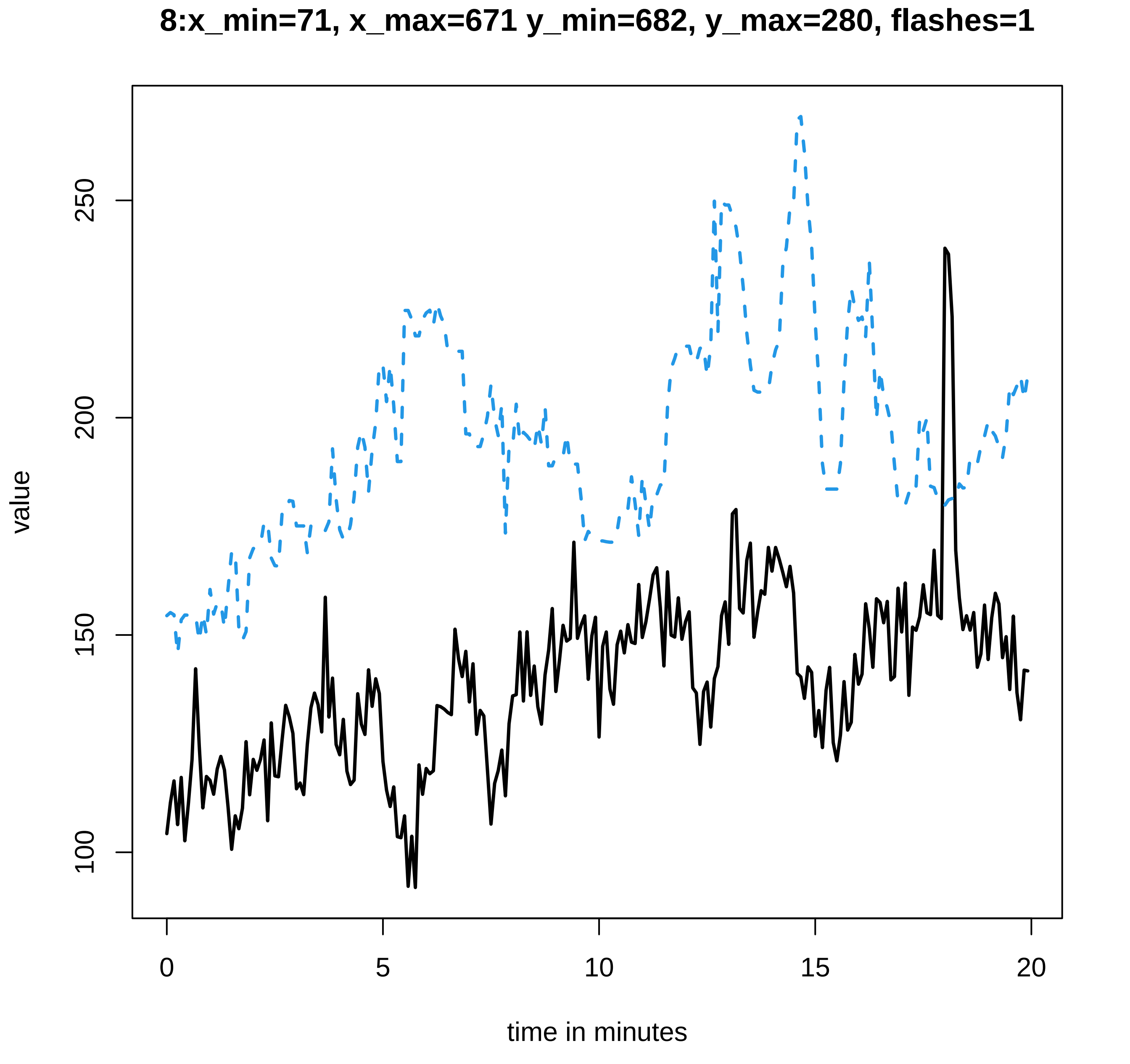
8:x_min=71, x_max=671 y_min=682, y_max=280, flashes=1
value
250
200
150
100
time in minutes
0
5
10
15
20

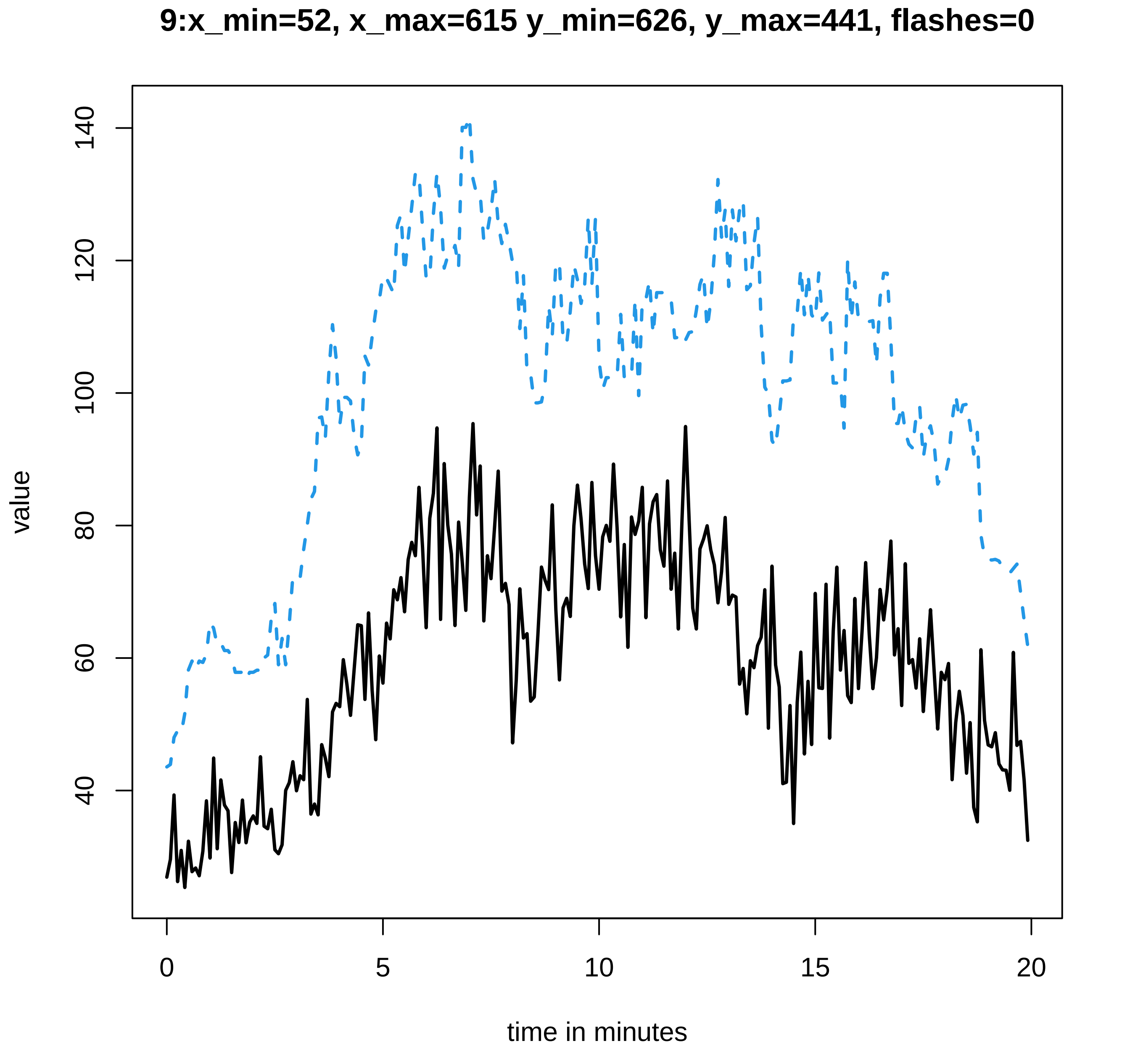
9:x_min=52, x_max=615 y_min=626, y_max=441, flashes=0
value
time in minutes
0
5
10
15
20
40
60
80
100
120
140

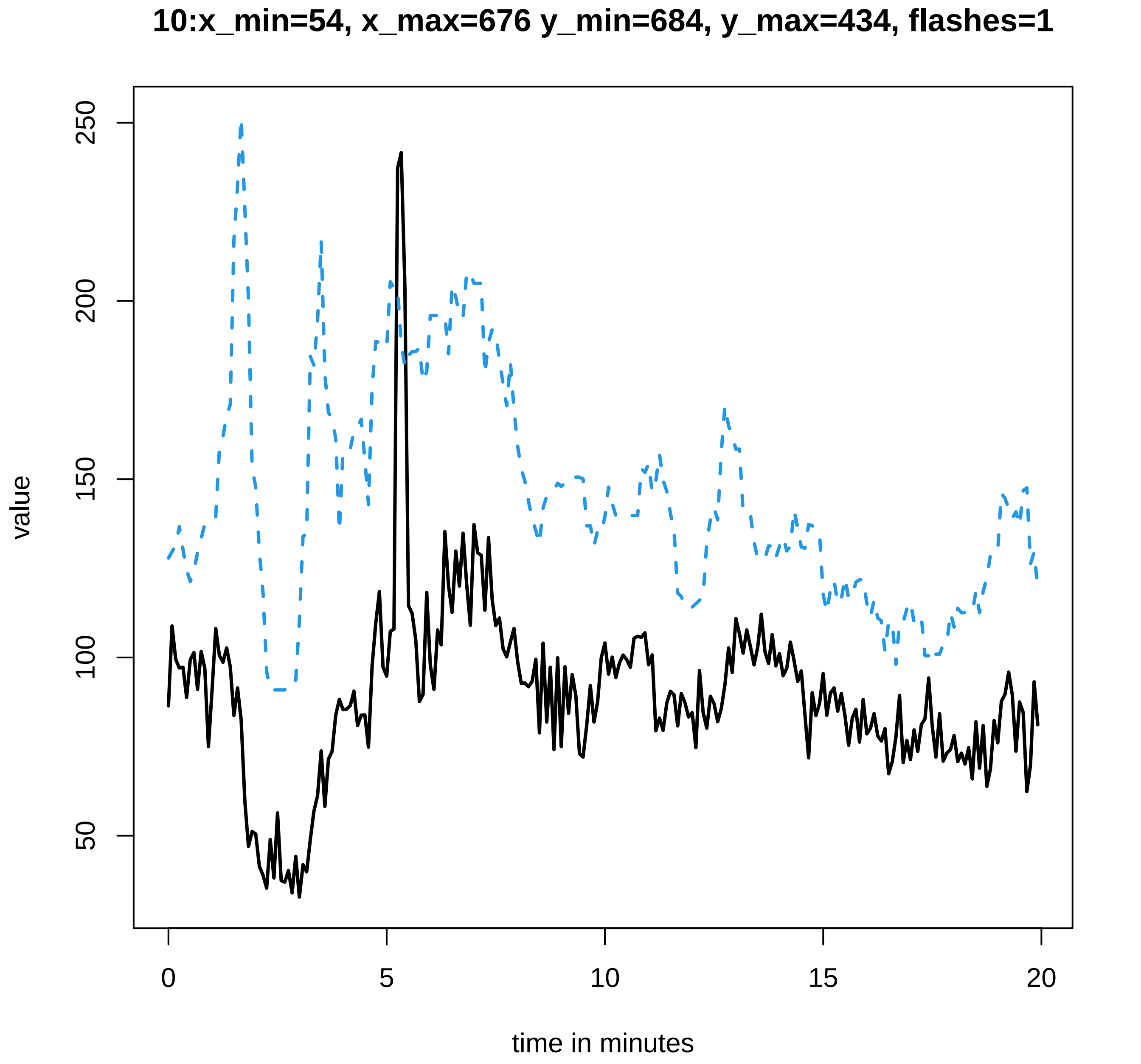
10:x_min=54, x_max=676 y_min=684, y_max=434, flashes=1
value
250
200
150
100
50
0
5
10
15
20
time in minutes

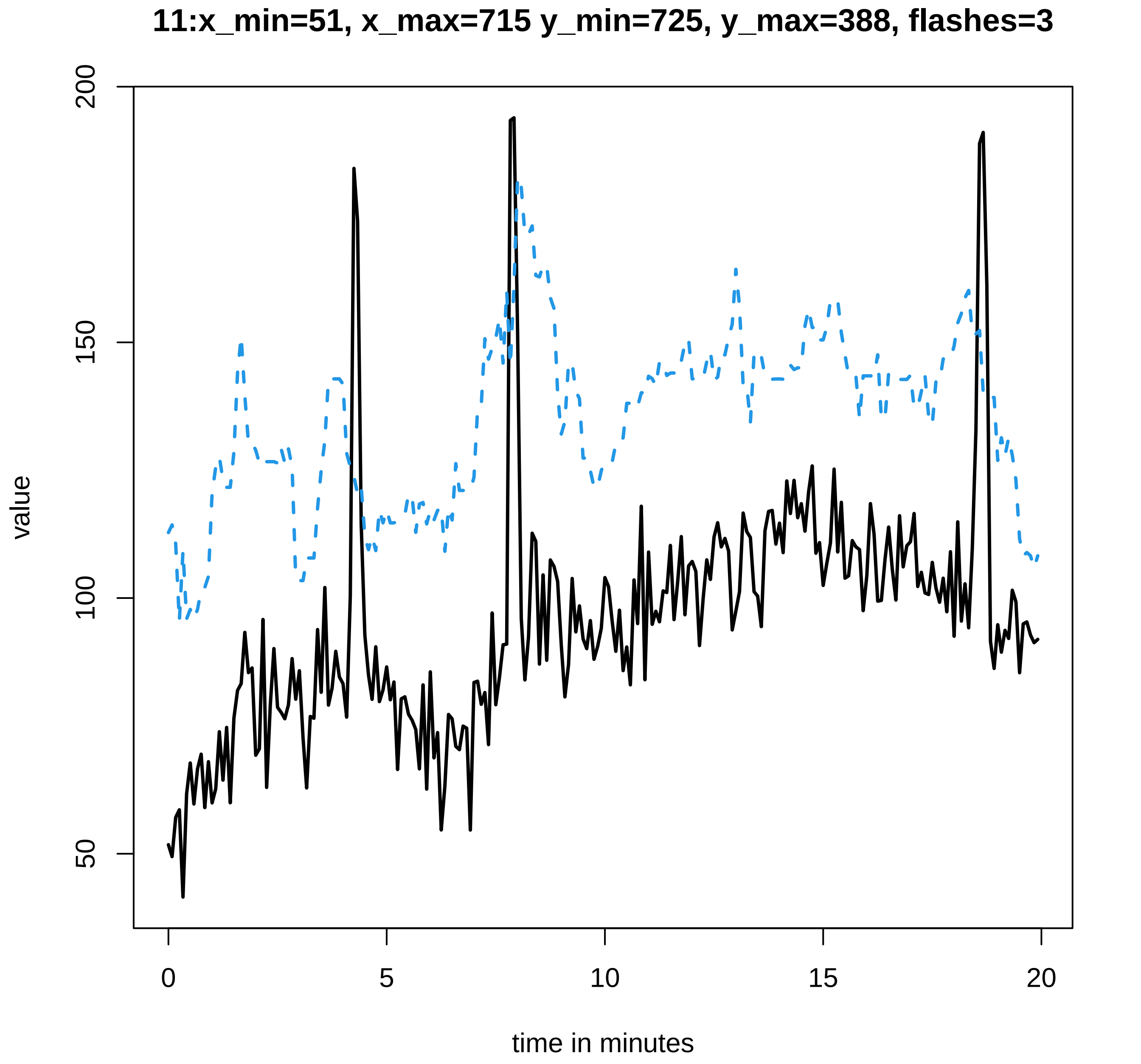
11:x_min=51, x_max=715 y_min=725, y_max=388, flashes=3
value
200
150
100
50
0
5
10
15
20
time in minutes

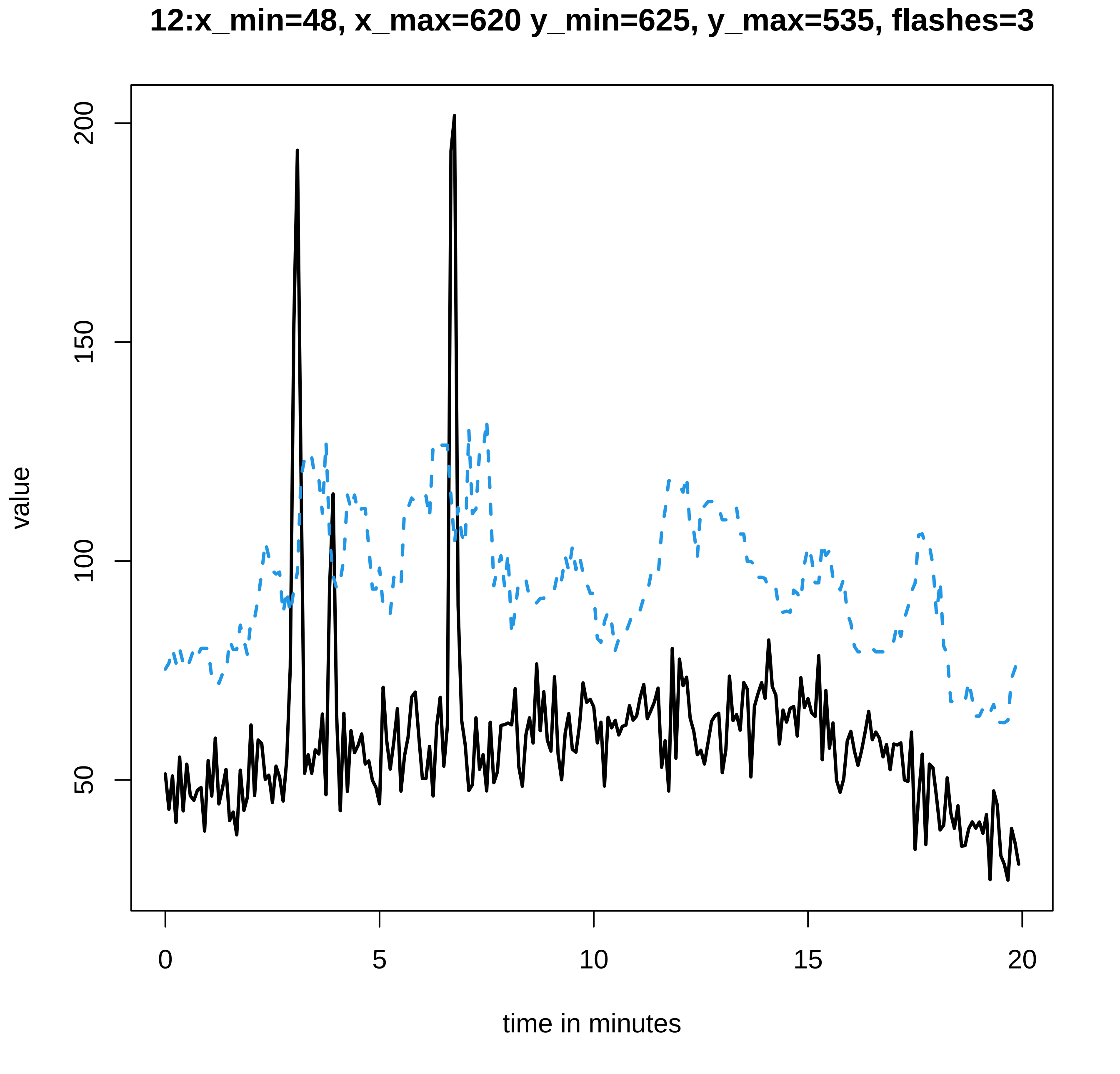
12:x_min=48, x_max=620 y_min=625, y_max=535, flashes=3
value
200
150
100
50
0
5
10
15
20
time in minutes

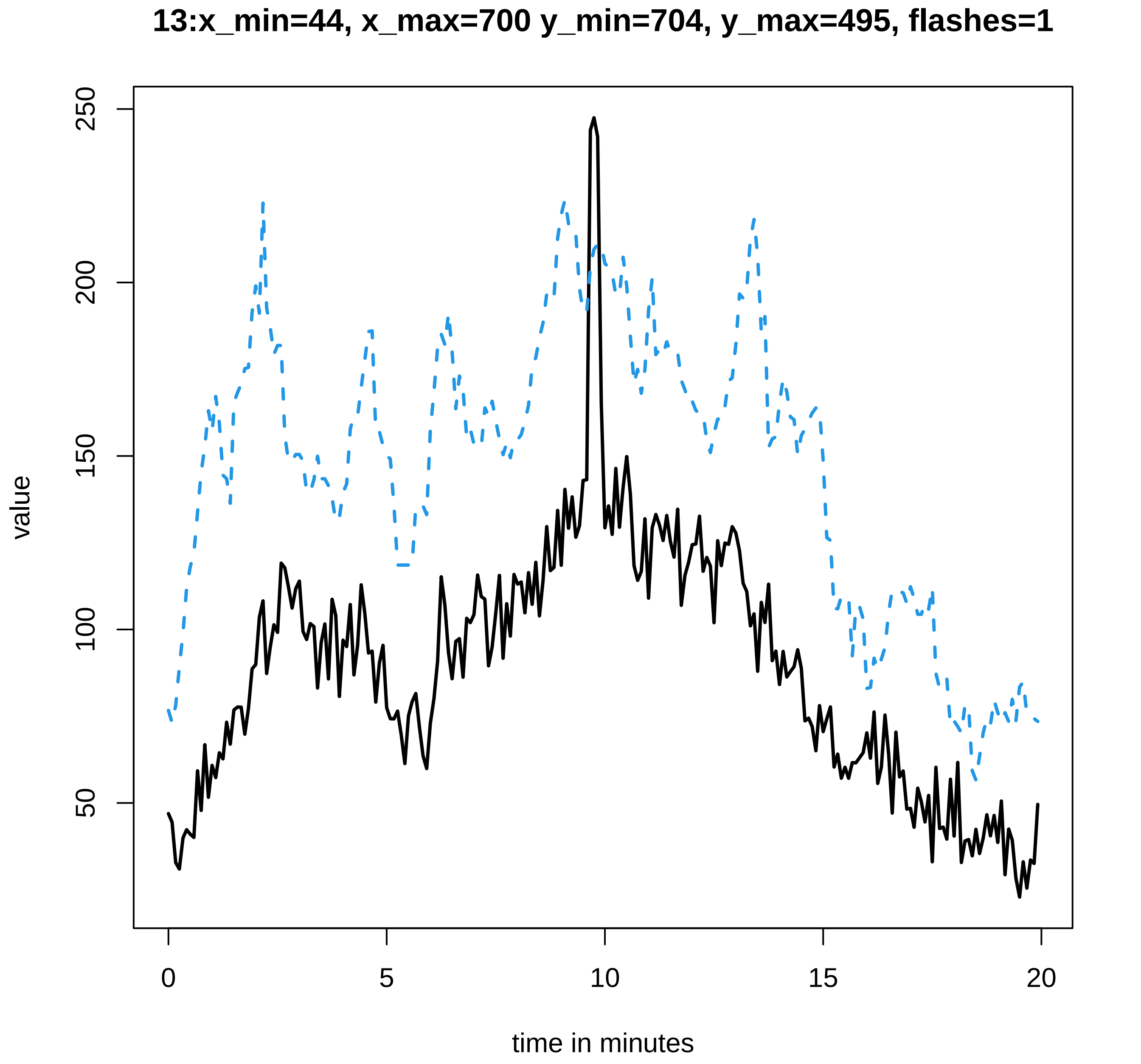

13:x_min=44, x_max=700 y_min=704, y_max=495, flashes=1
value
50
100
150
200
250
0
5
10
15
20
time in minutes

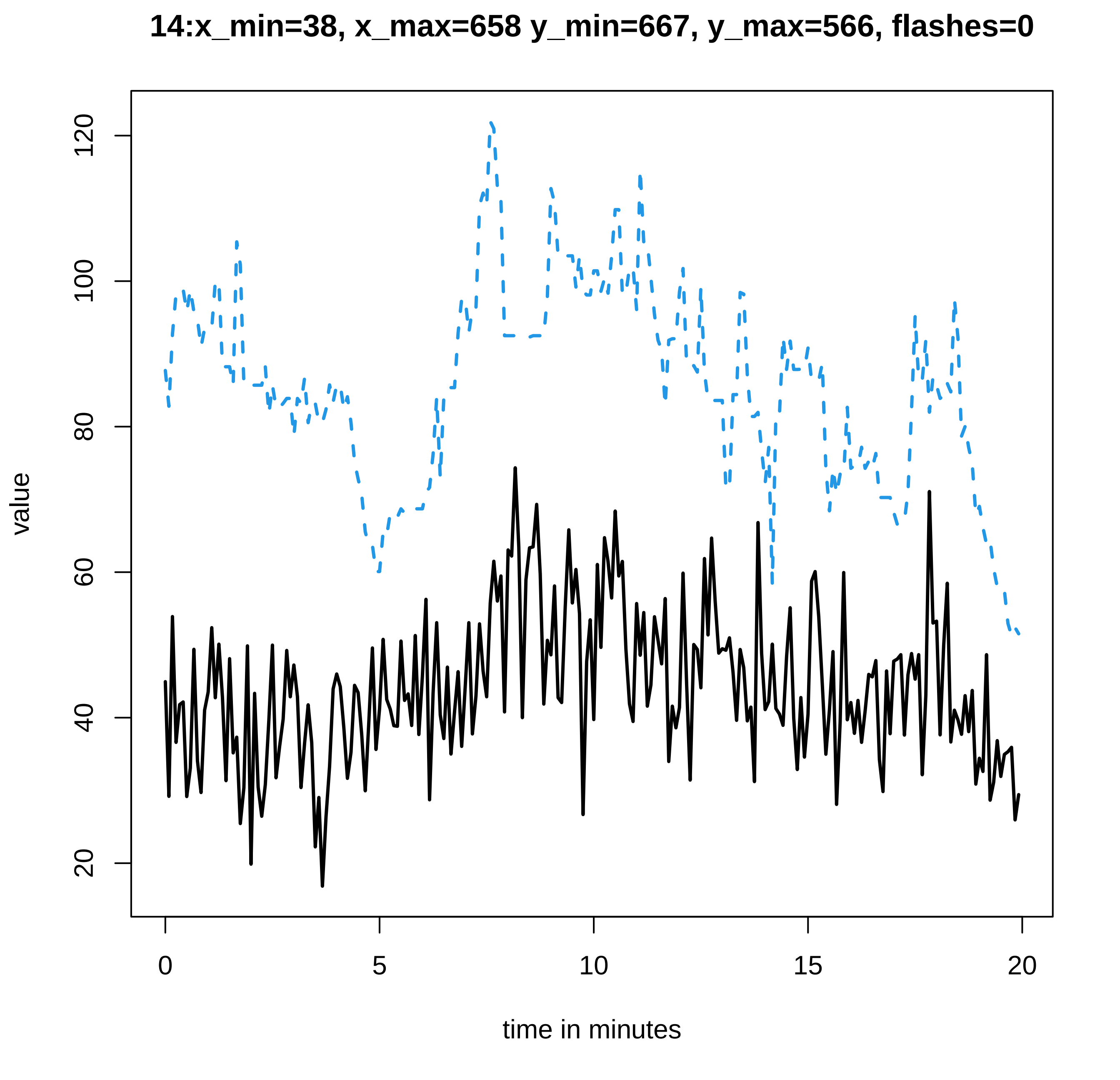
14:x_min=38, x_max=658 y_min=667, y_max=566, flashes=0
value
120
100
80
60
40
20
0
5
10
15
20
time in minutes

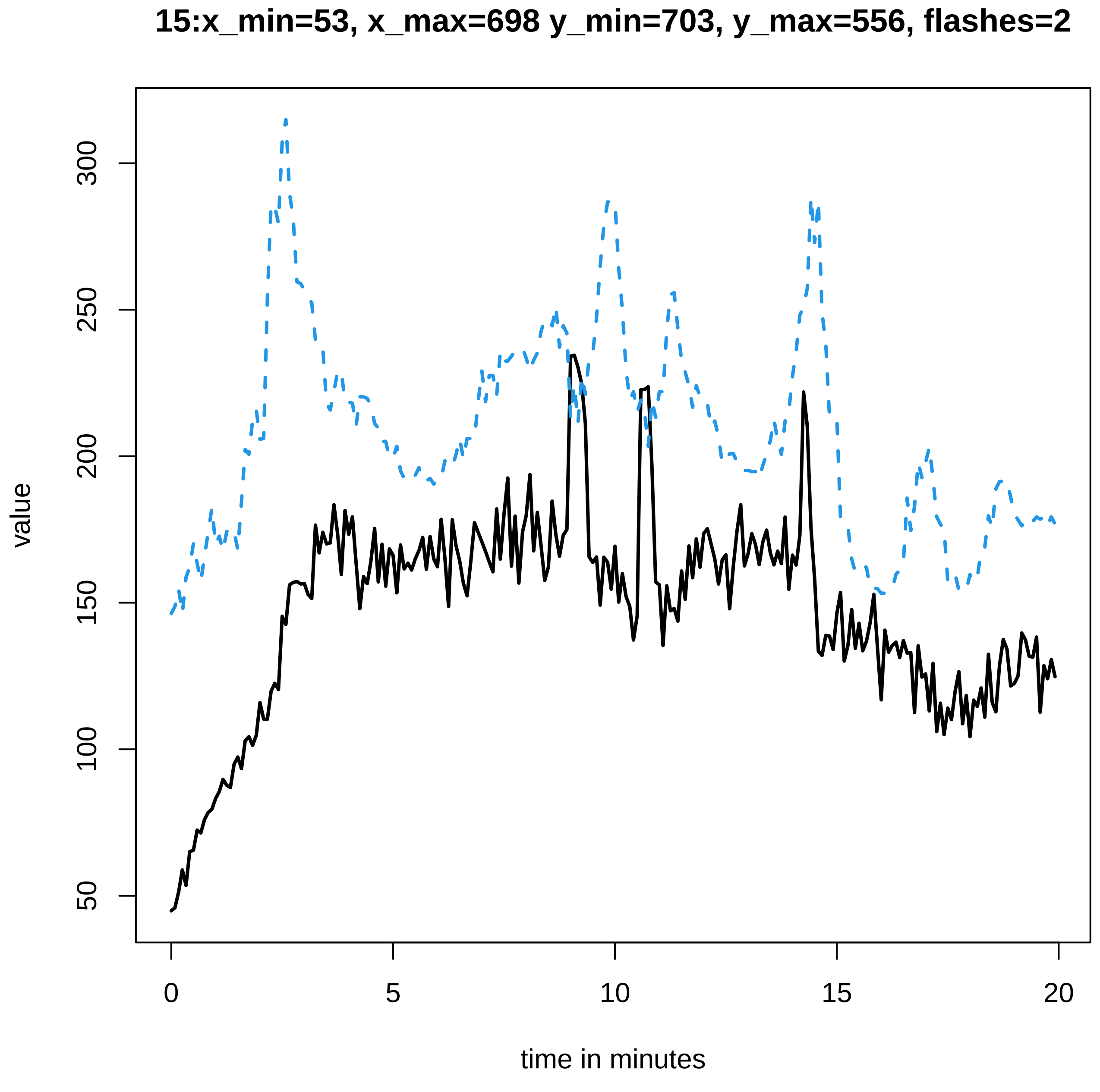
15:x_min=53, x_max=698 y_min=703, y_max=556, flashes=2
value
50
100
150
200
250
300
0
5
10
15
20
time in minutes

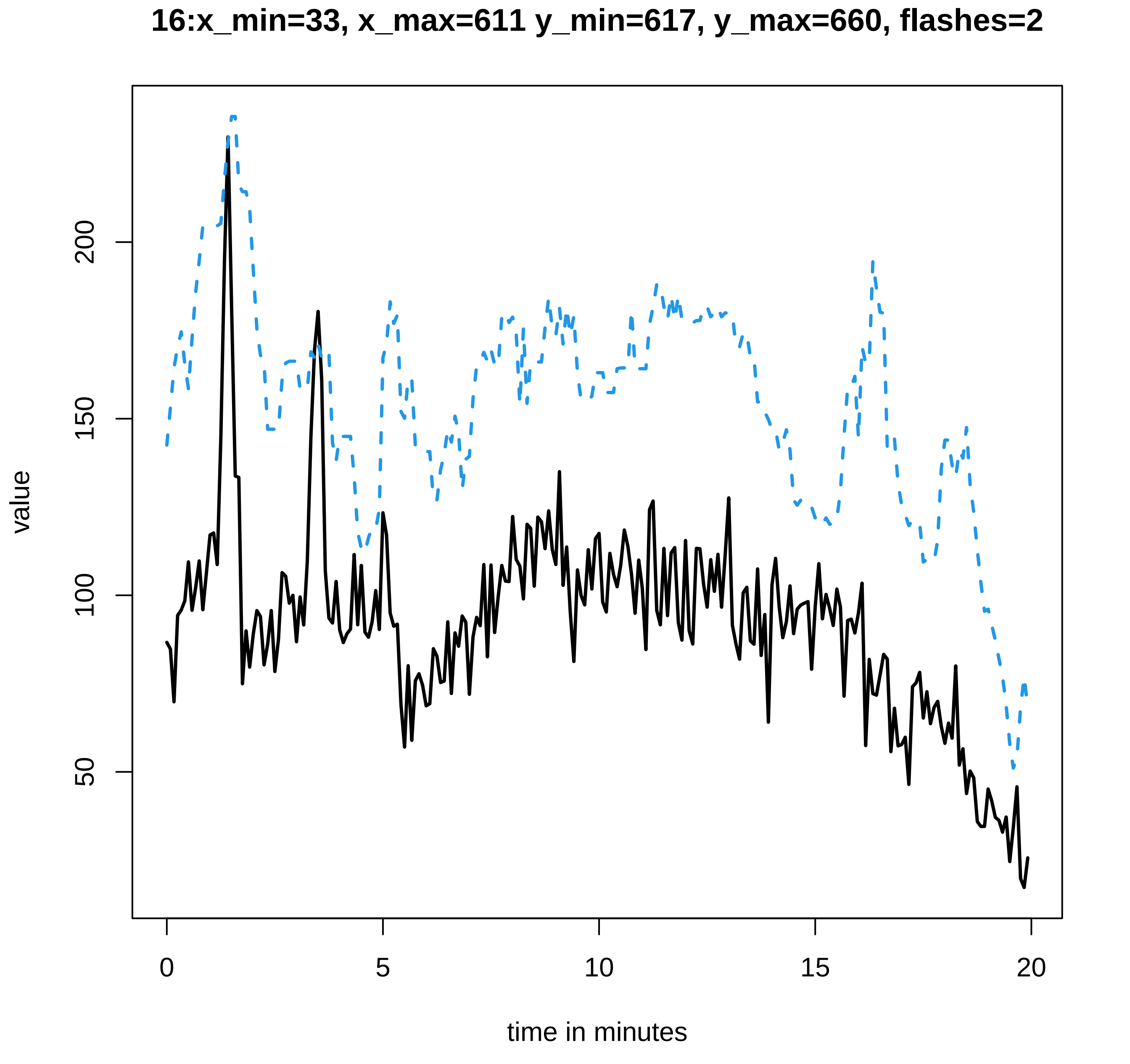

16:x_min=33, x_max=611 y_min=617, y_max=660, flashes=2
value
50
100
150
200
0
5
10
15
20
time in minutes

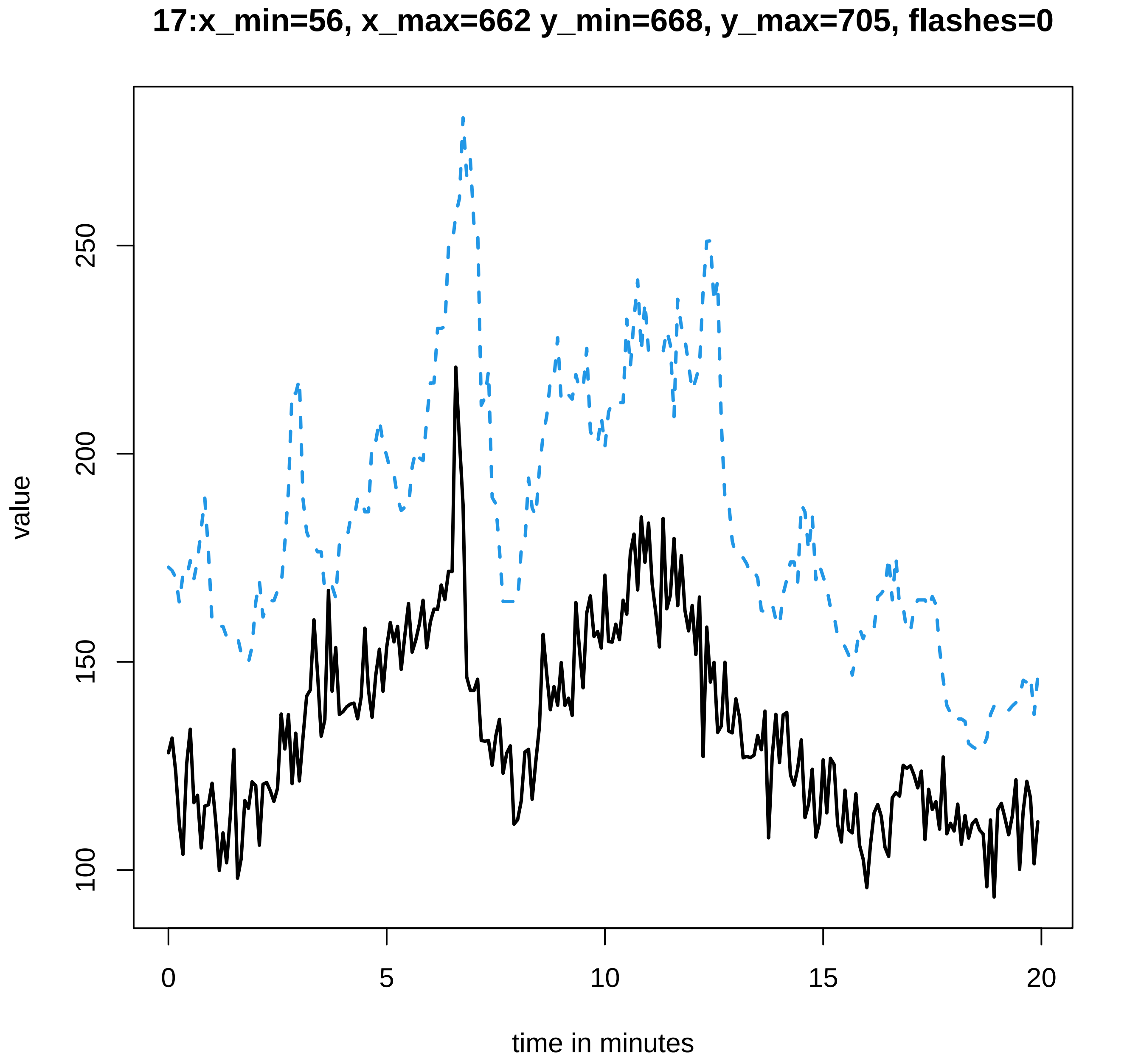

17:x_min=56, x_max=662 y_min=668, y_max=705, flashes=0
value
100
150
200
250
0
5
10
15
20
time in minutes

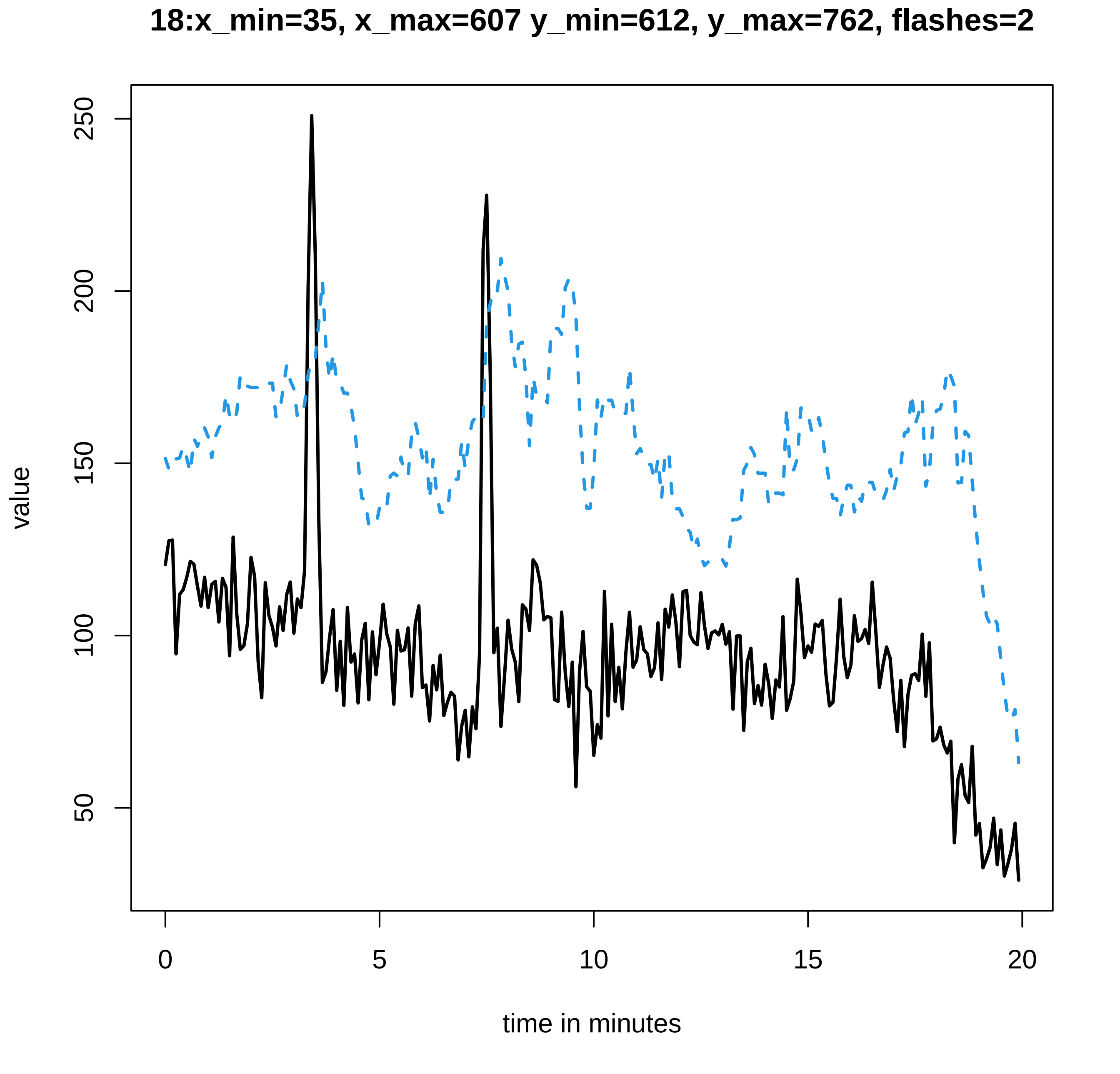
18:x_min=35, x_max=607 y_min=612, y_max=762, flashes=2
value
250
200
150
100
50
0
5
10
15
20
time in minutes

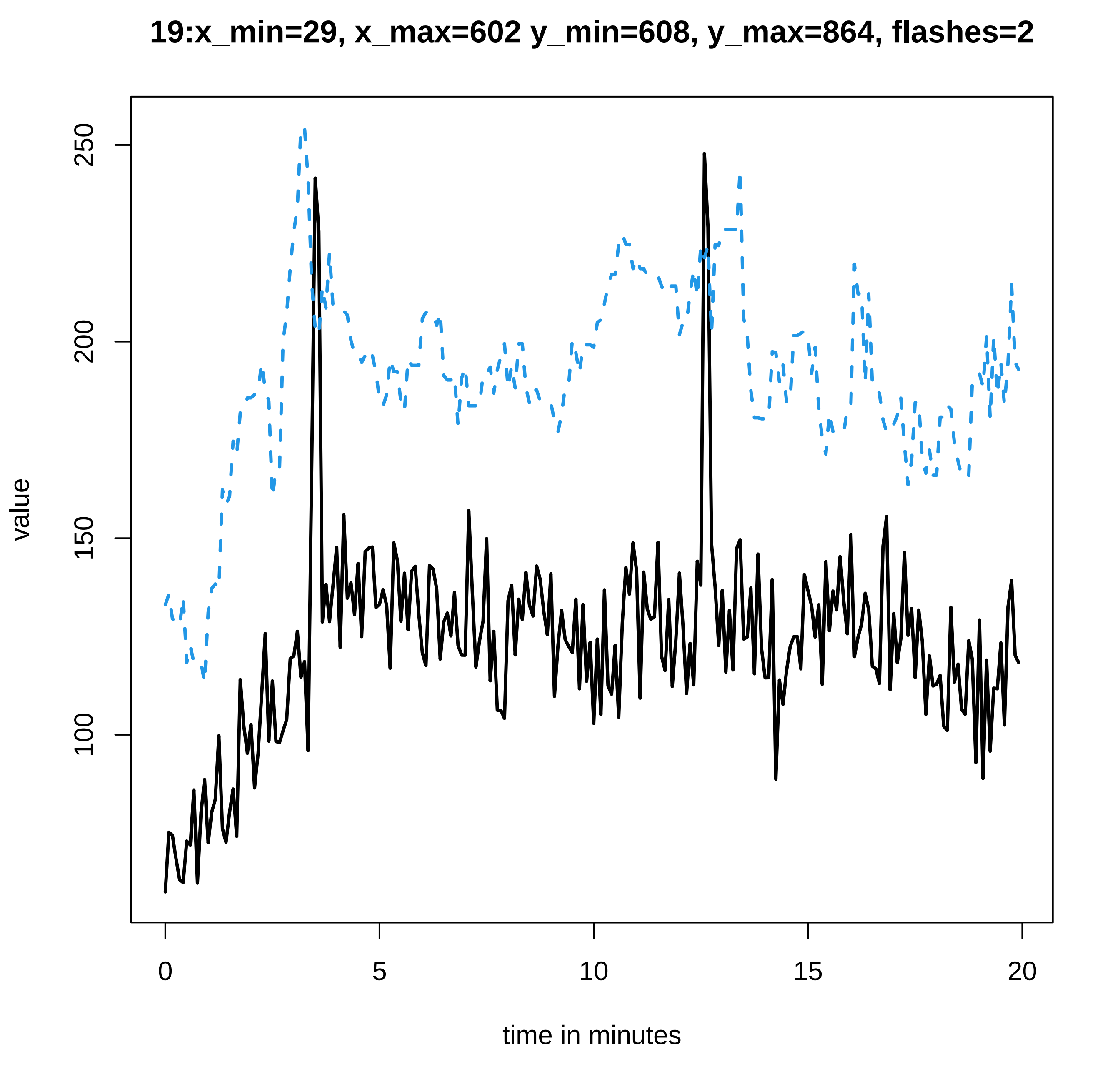
19:x_min=29, x_max=602 y_min=608, y_max=864, flashes=2
value
250
200
150
100
0
5
10
15
20
time in minutes

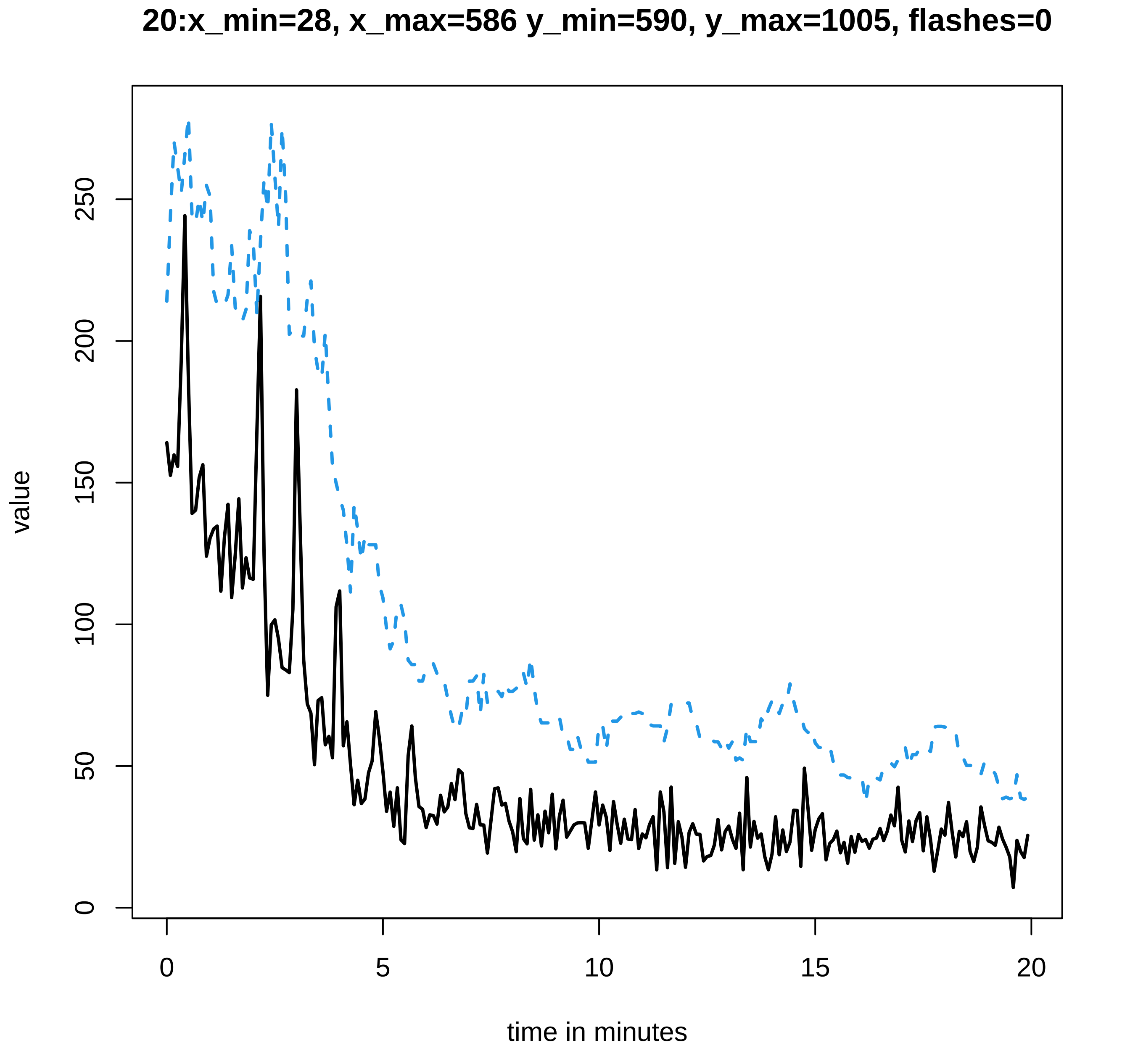
20:x_min=28, x_max=586 y_min=590, y_max=1005, flashes=0
value
0
50
100
150
200
250
0
5
10
15
20
time in minutes
