## Supplementary data Calcium spiking for "LysM Receptor Proteins are Required for Ectomycorrhizal Symbiosis in Poplar": profiles-spiking-11.pdf

#### 1:x_min=40, x_max=393 y_min=397, y_max=28, flashes=0

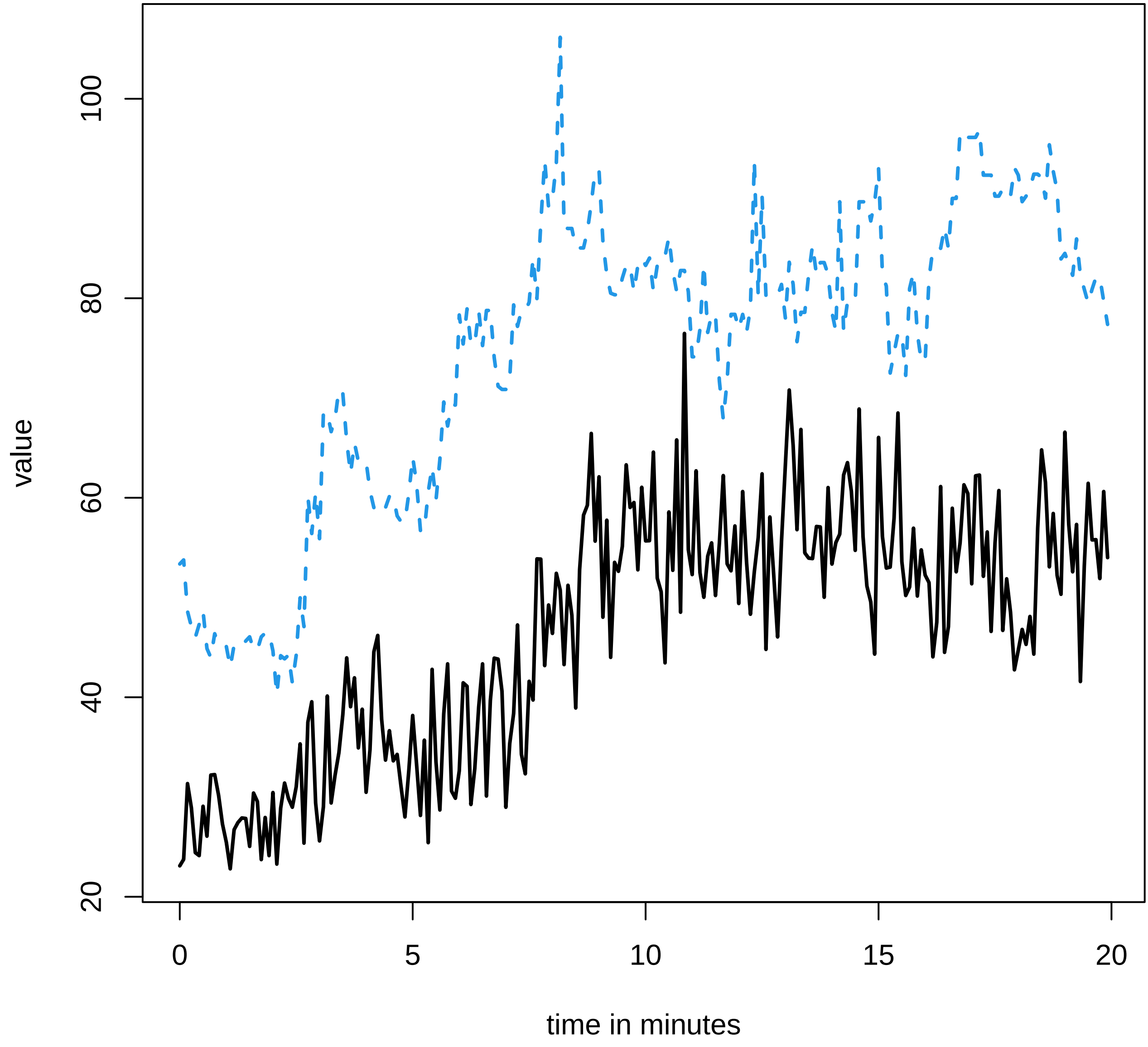

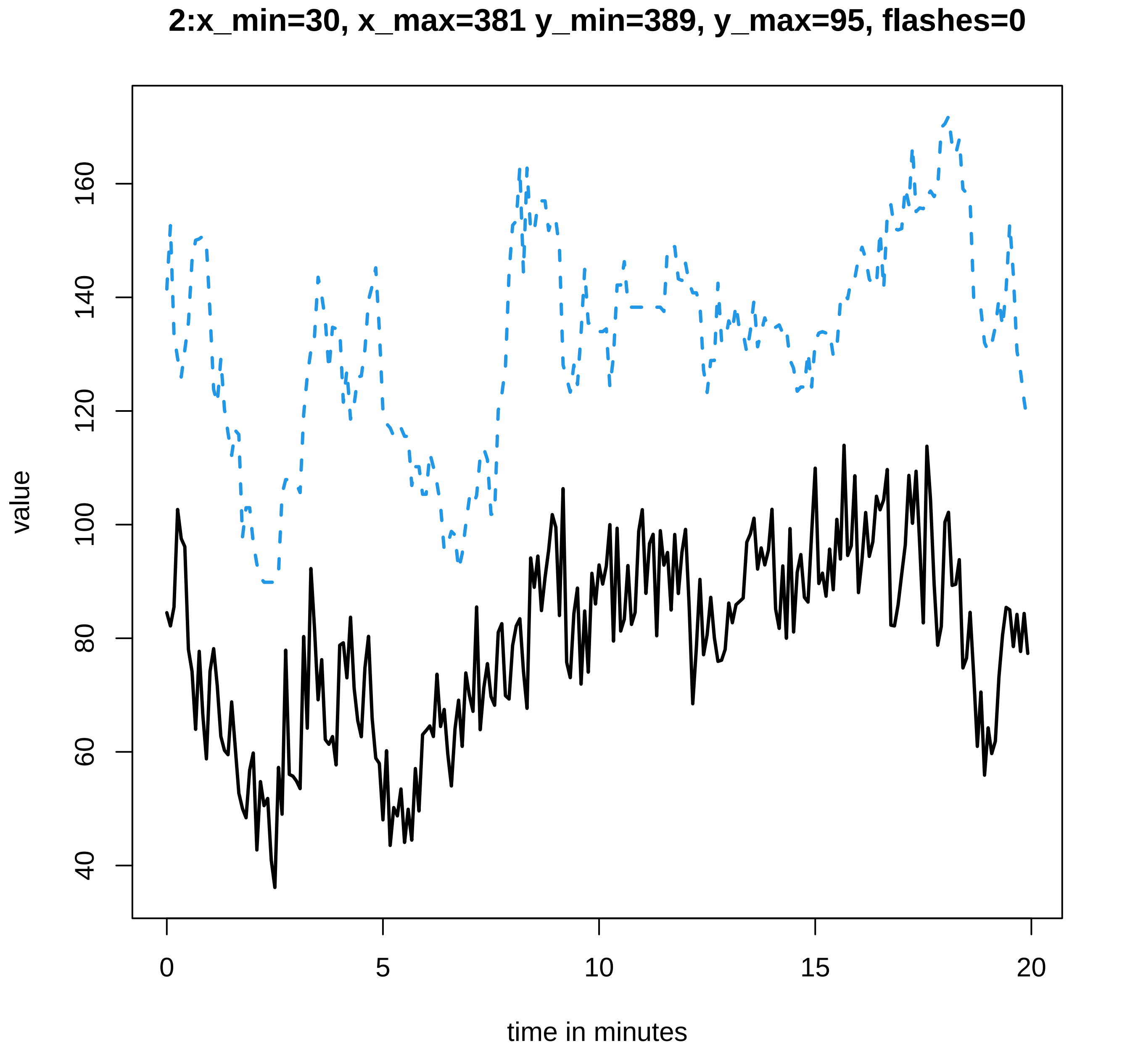
2:x_min=30, x_max=381 y_min=389, y_max=95, flashes=0
value
160
140
120
100
80
60
40
0
5
10
15
20
time in minutes

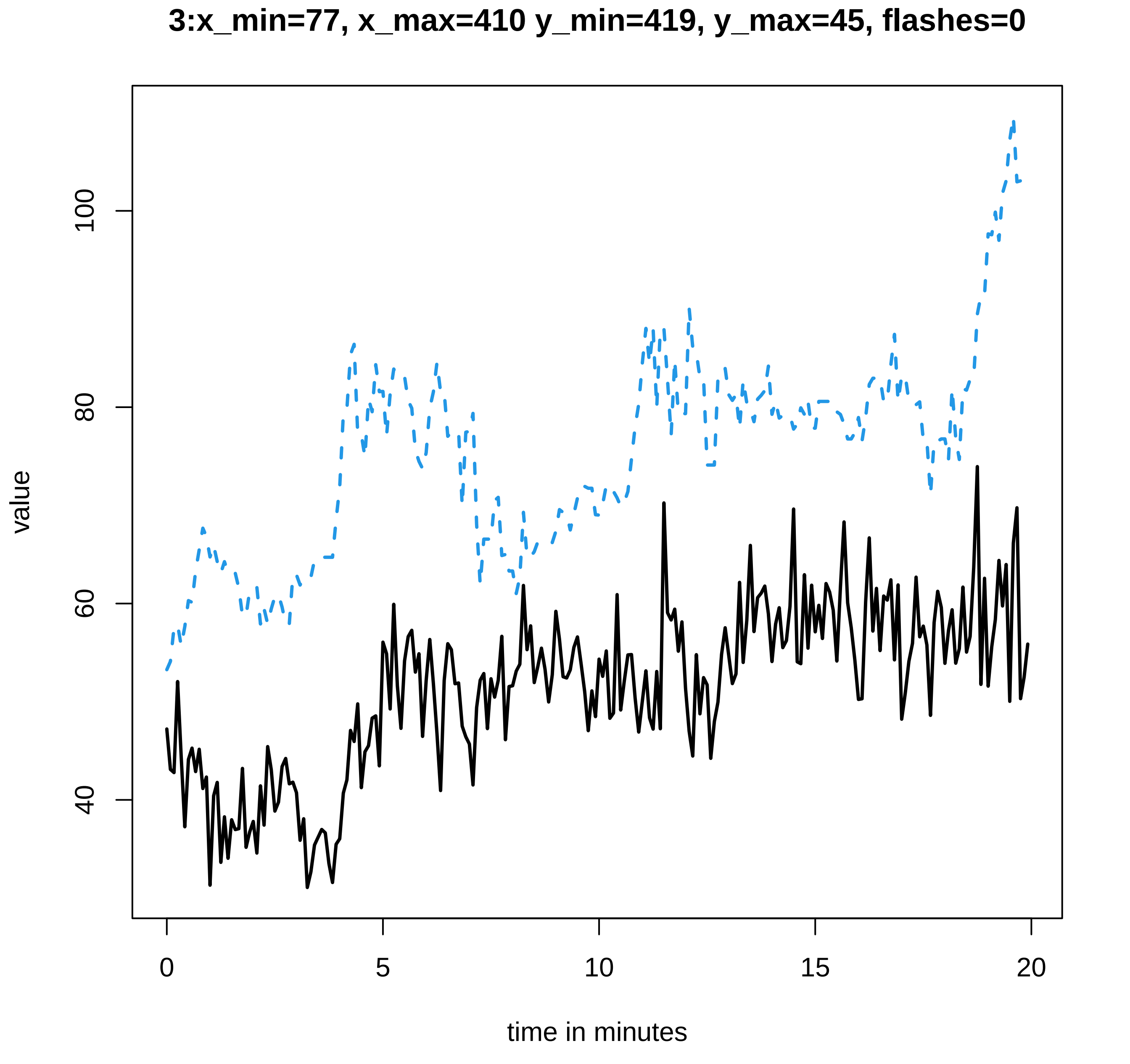

3:x_min=77, x_max=410 y_min=419, y_max=45, flashes=0
value
100
80
60
40
0
5
10
15
20
time in minutes

### 4:x_min=42, x_max=494 y_min=499, y_max=244, flashes=0

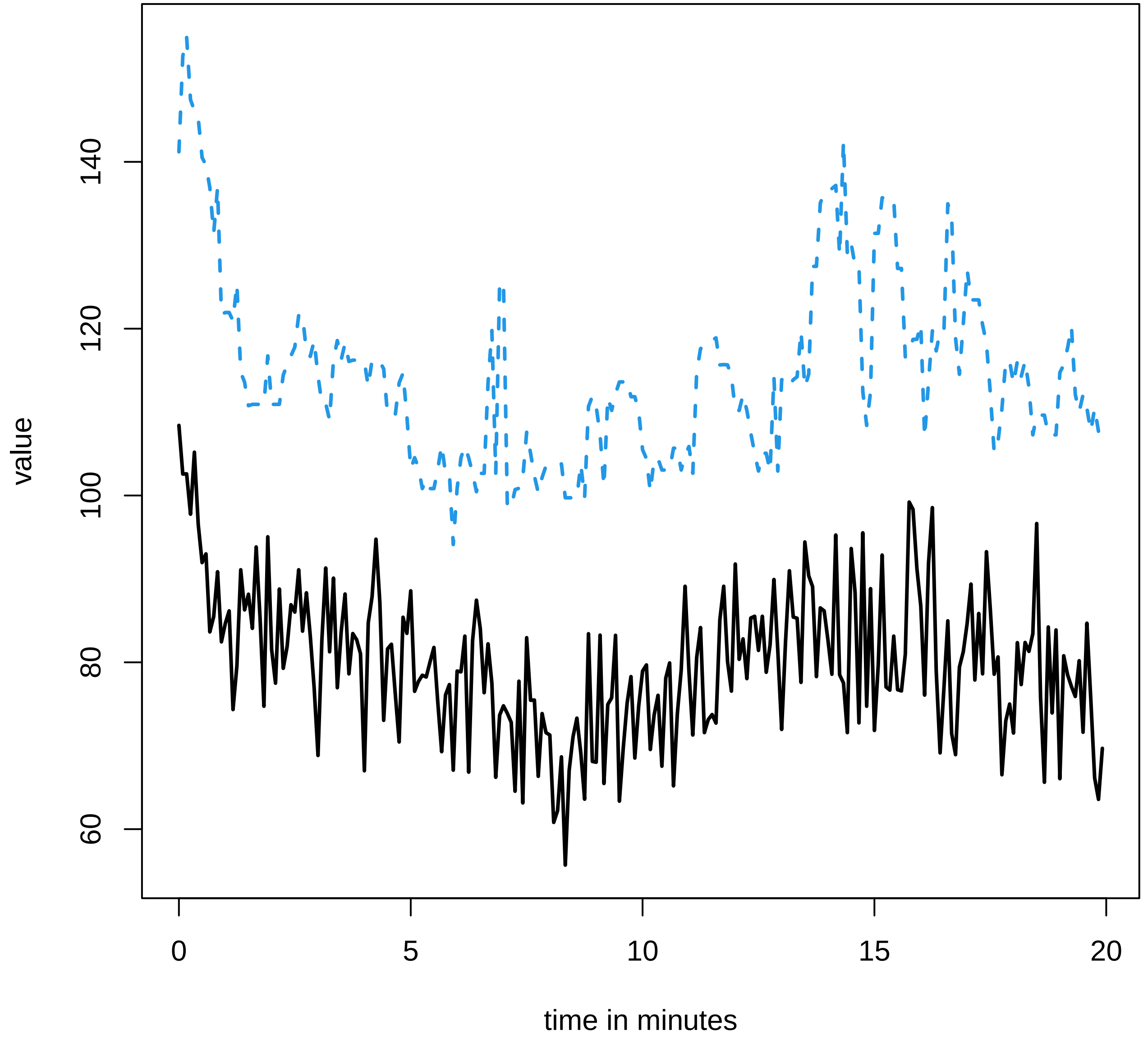

#### 5:x_min=43, x_max=513 y_min=516, y_max=219, flashes=0

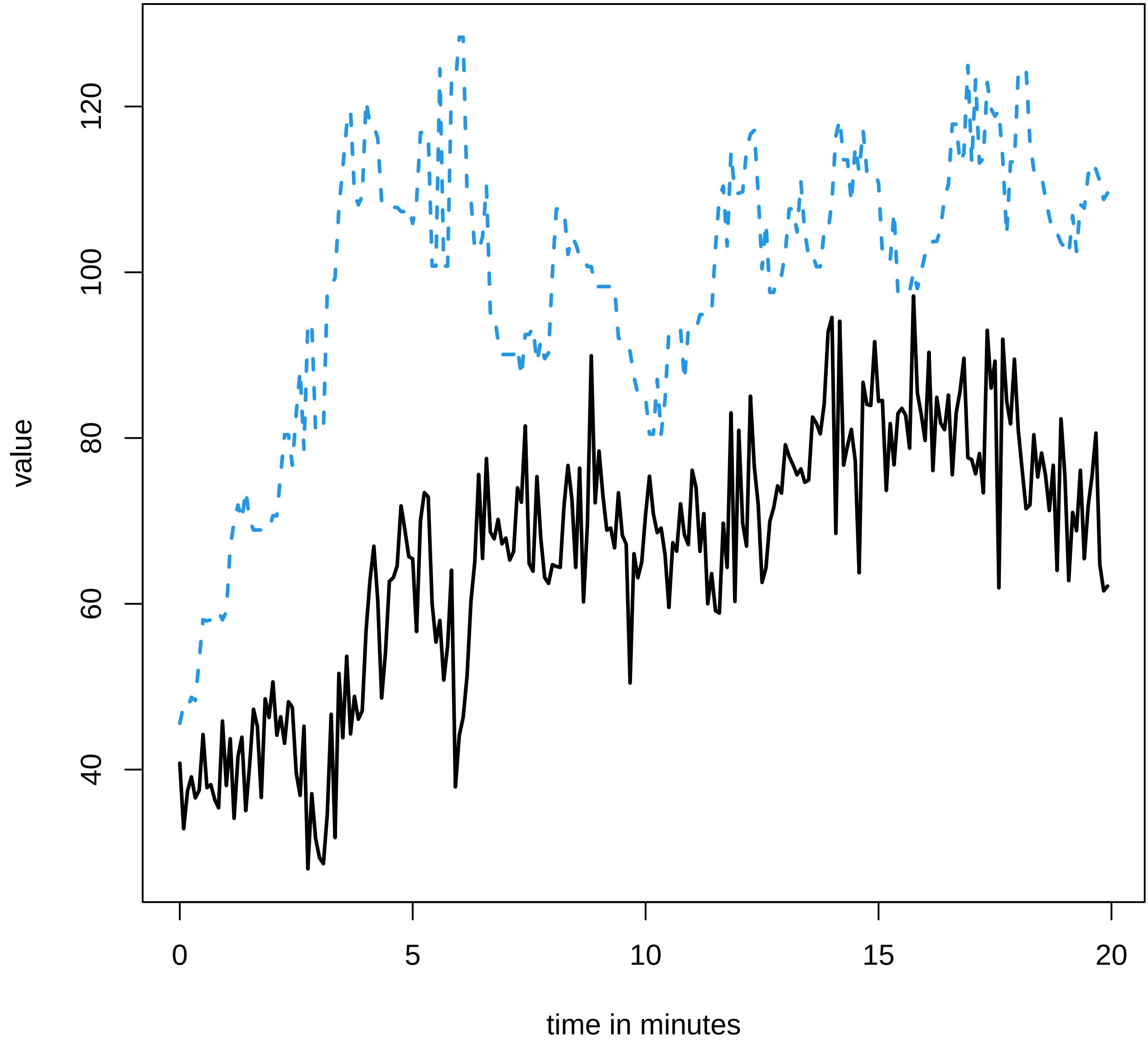

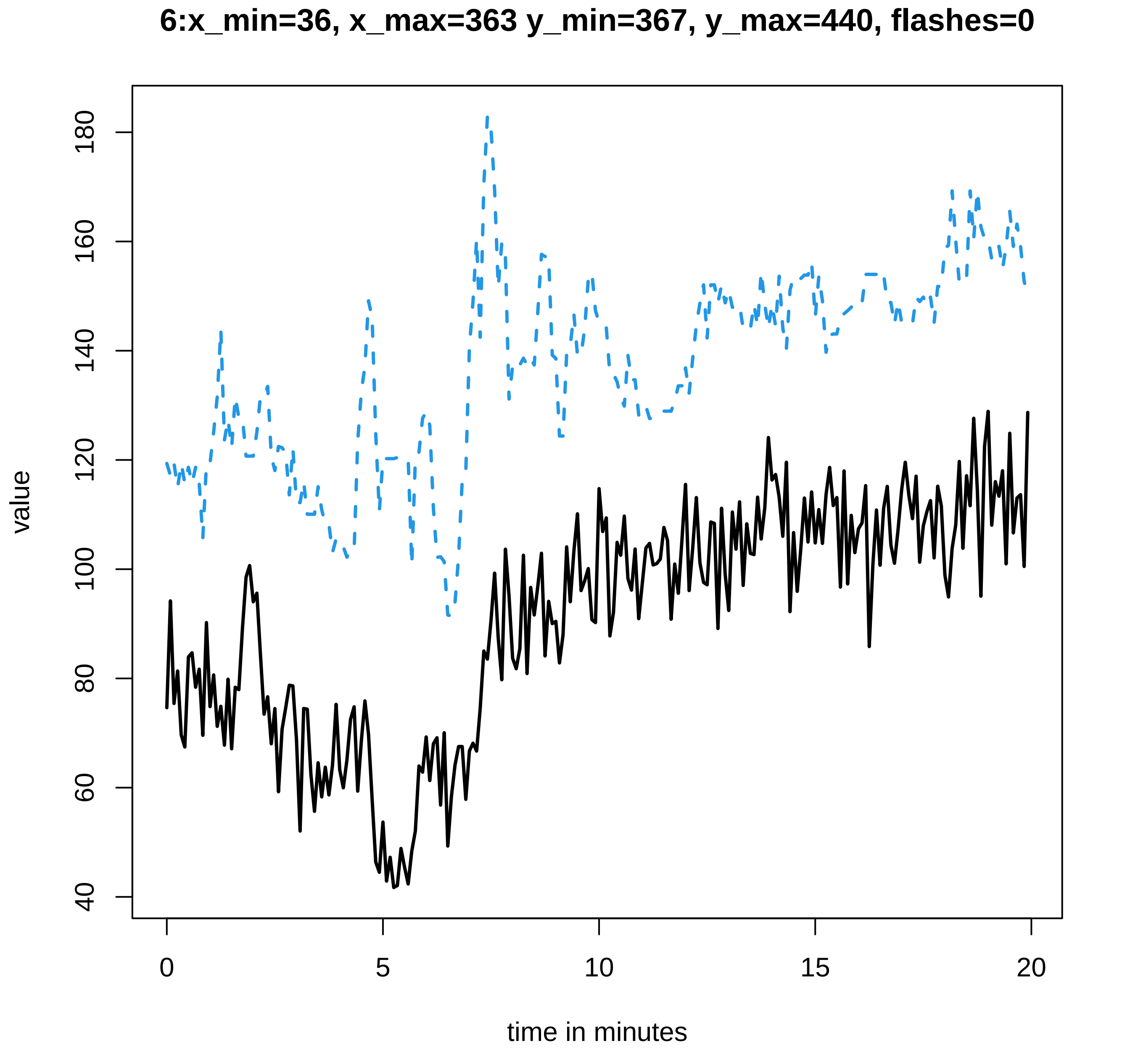

6:x_min=36, x_max=363 y_min=367, y_max=440, flashes=0
value
40
60
80
100
120
140
160
180
0
5
10
15
20
time in minutes

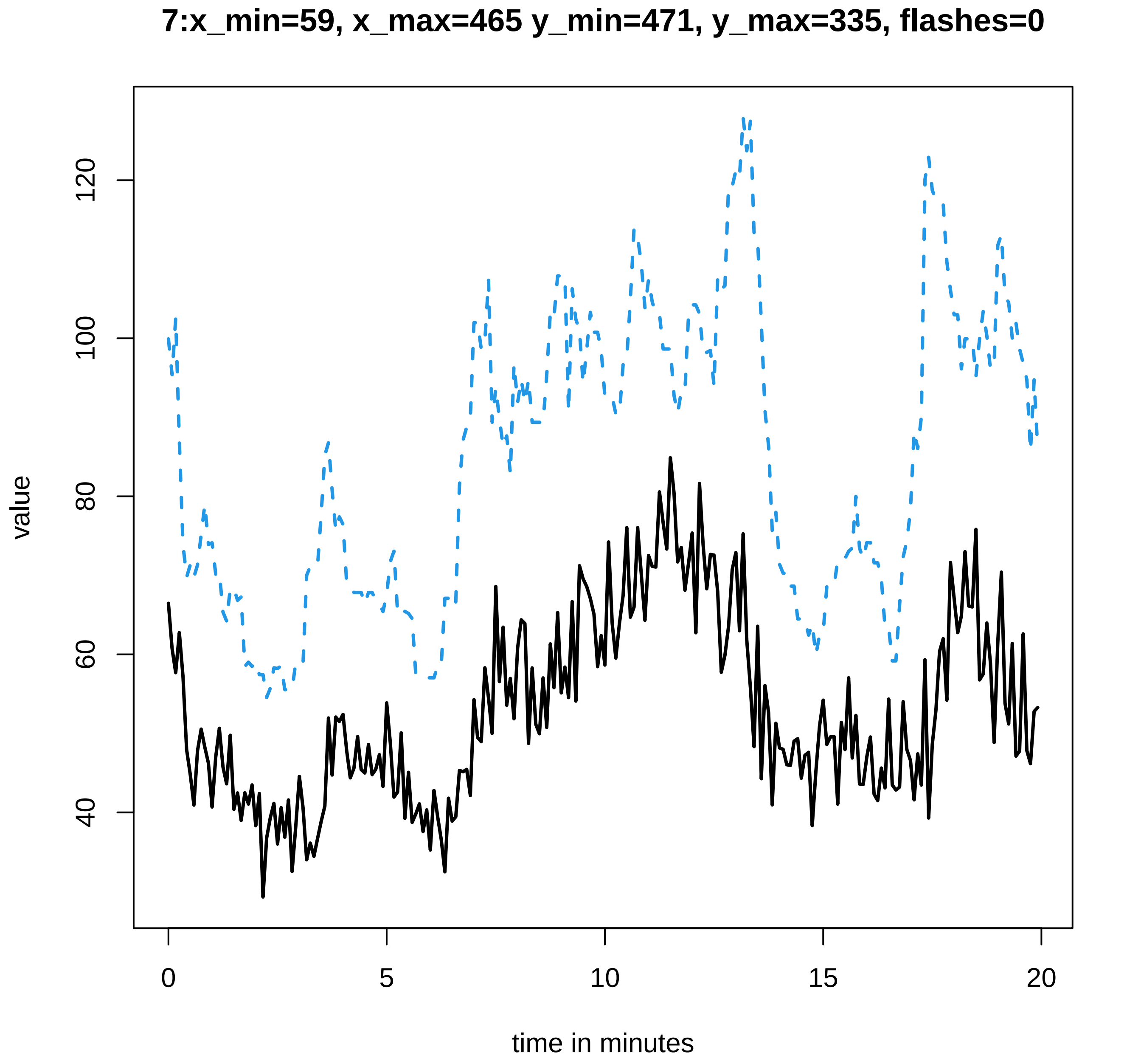

7:x_min=59, x_max=465 y_min=471, y_max=335, flashes=0
value
120
100
80
60
40
0
5
10
15
20
time in minutes

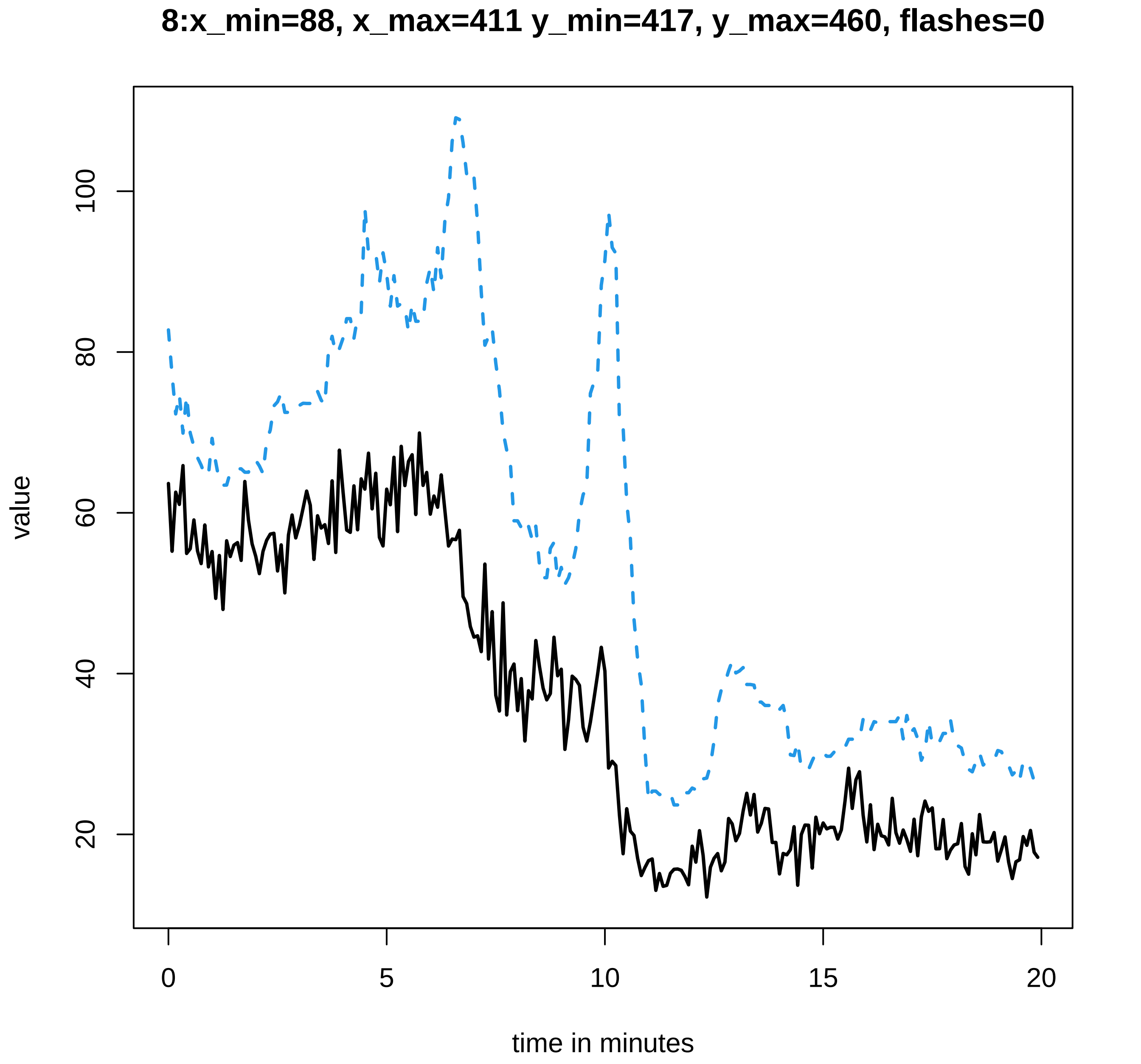
8:x_min=88, x_max=411 y_min=417, y_max=460, flashes=0
value
100
80
60
40
20
0
5
10
15
20
time in minutes

### 9:x_min=44, x_max=432 y_min=442, y_max=464, flashes=0

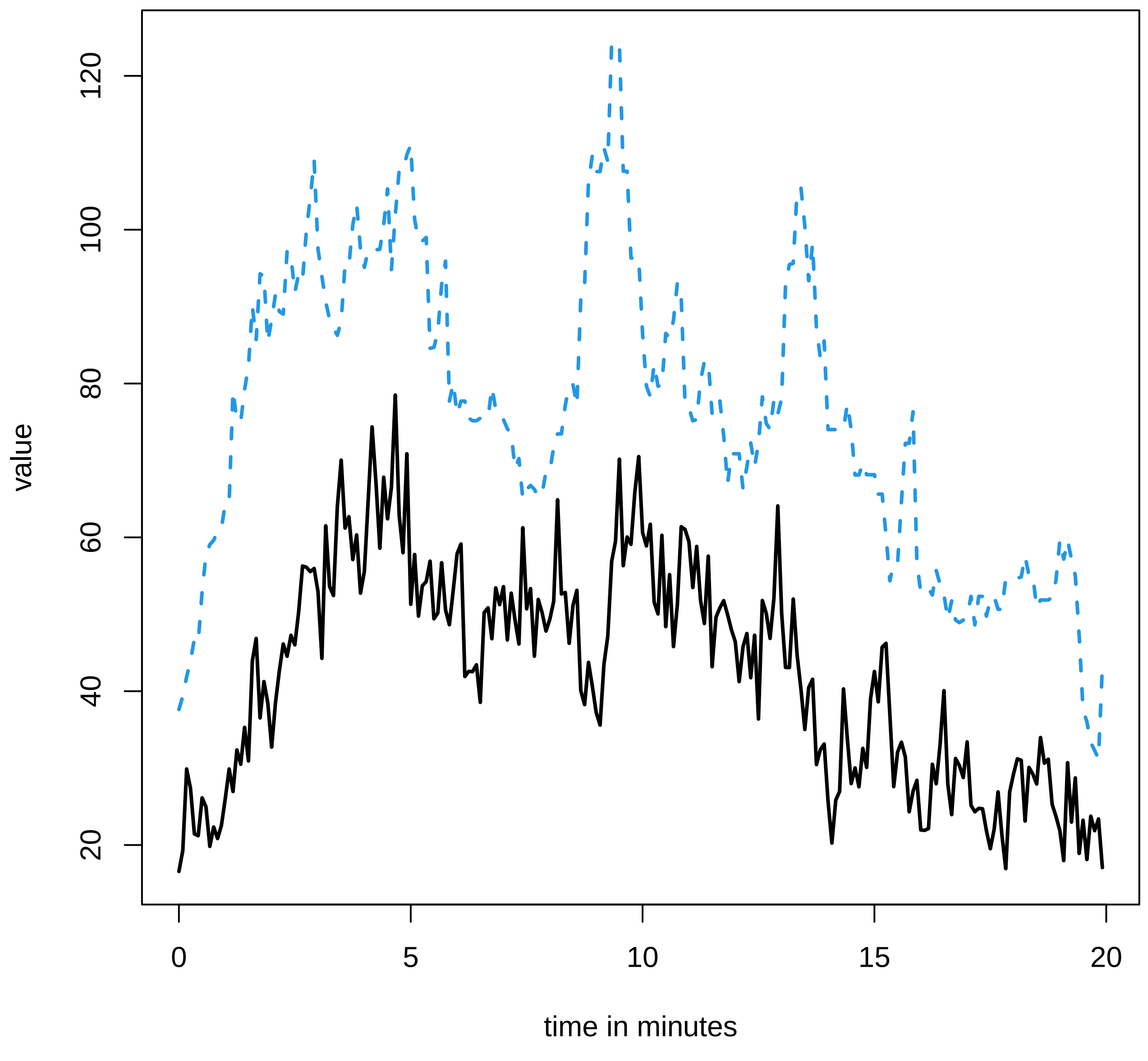

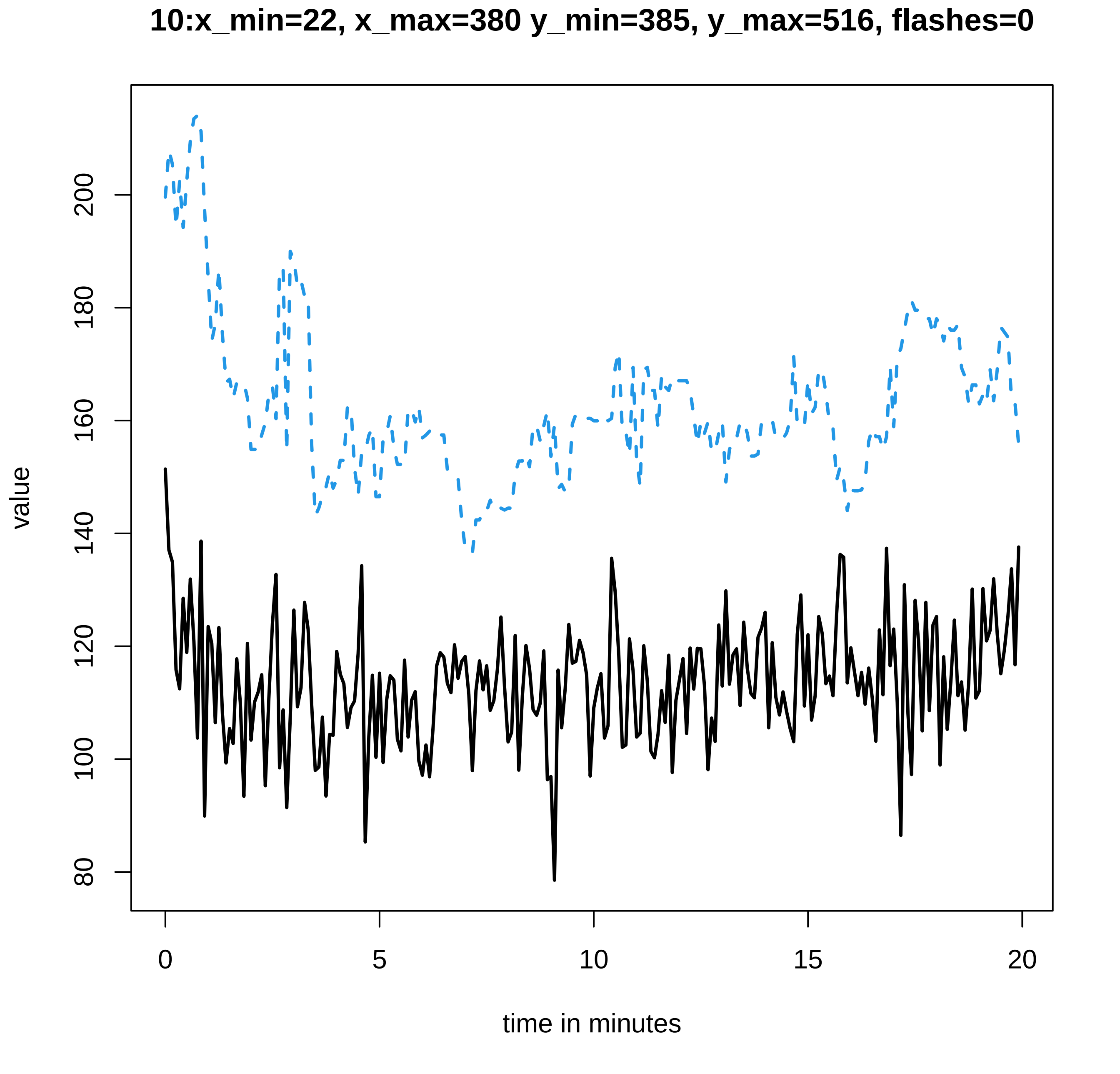
10:x_min=22, x_max=380 y_min=385, y_max=516, flashes=0
value
80
100
120
140
160
180
200
0
5
10
15
20
time in minutes

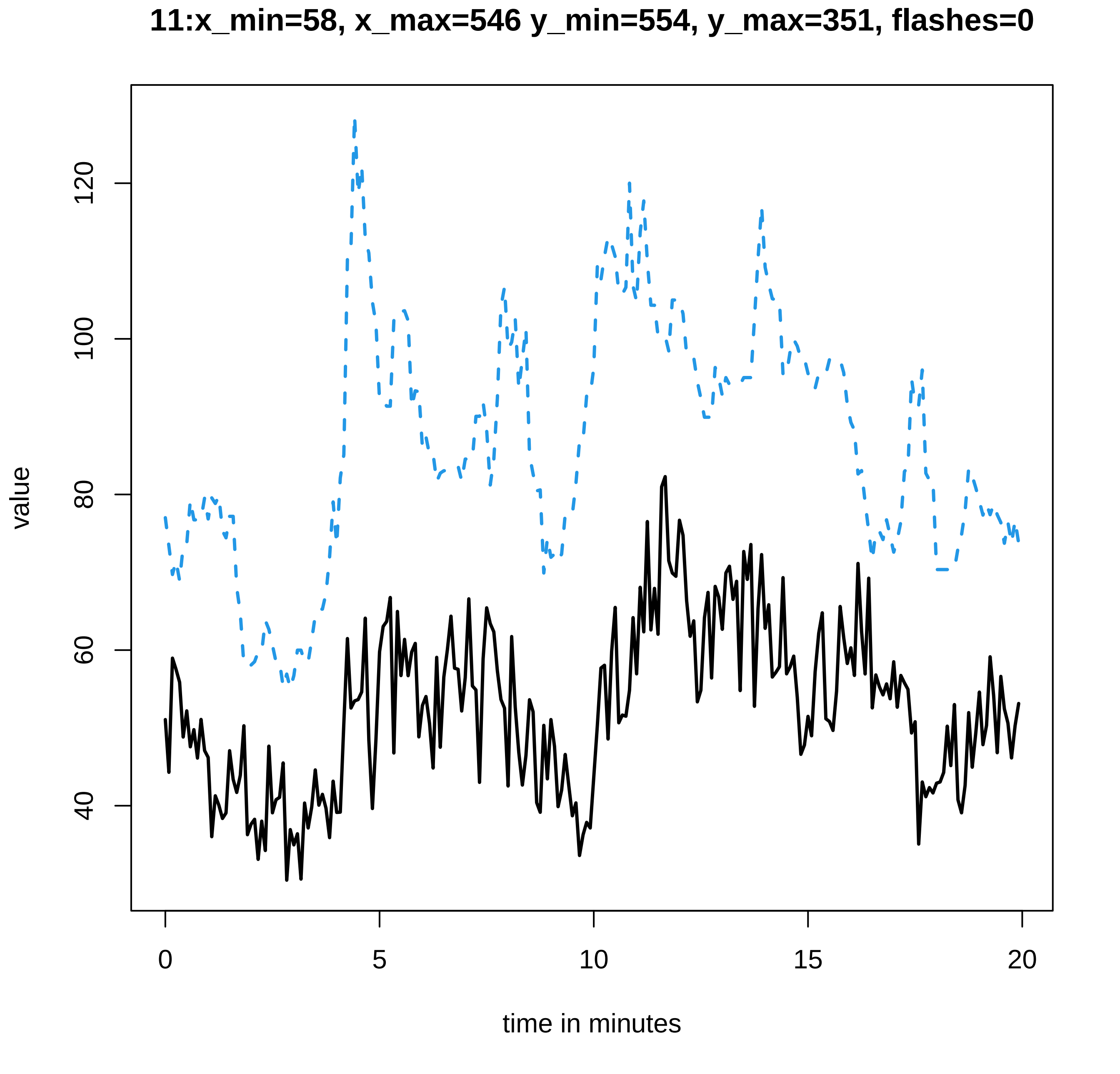

11:x_min=58, x_max=546 y_min=554, y_max=351, flashes=0
value
40
60
80
100
120
0
5
10
15
20
time in minutes

12:x_min=34, x_max=570 y_min=573, y_max=334, flashes=0

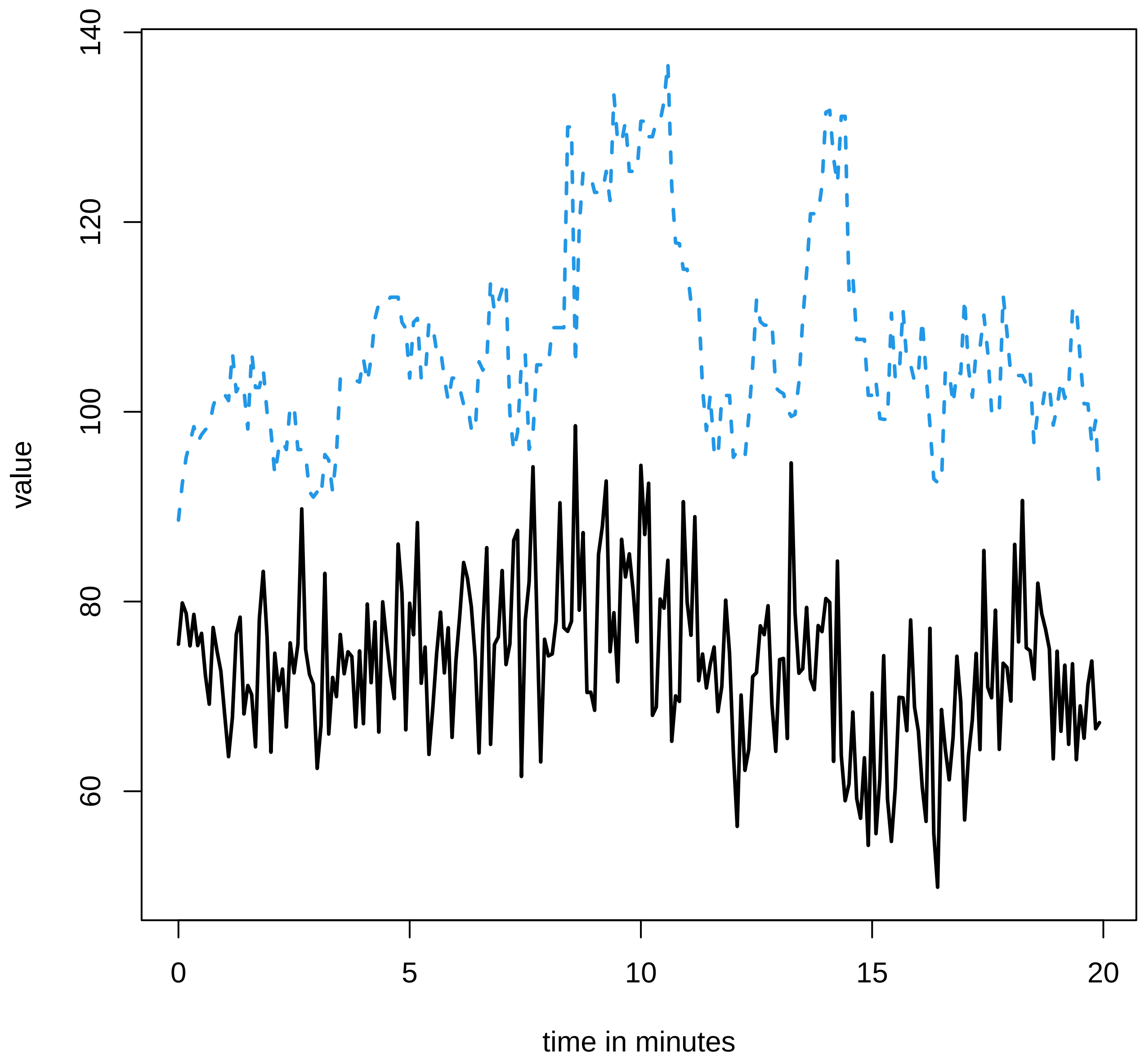

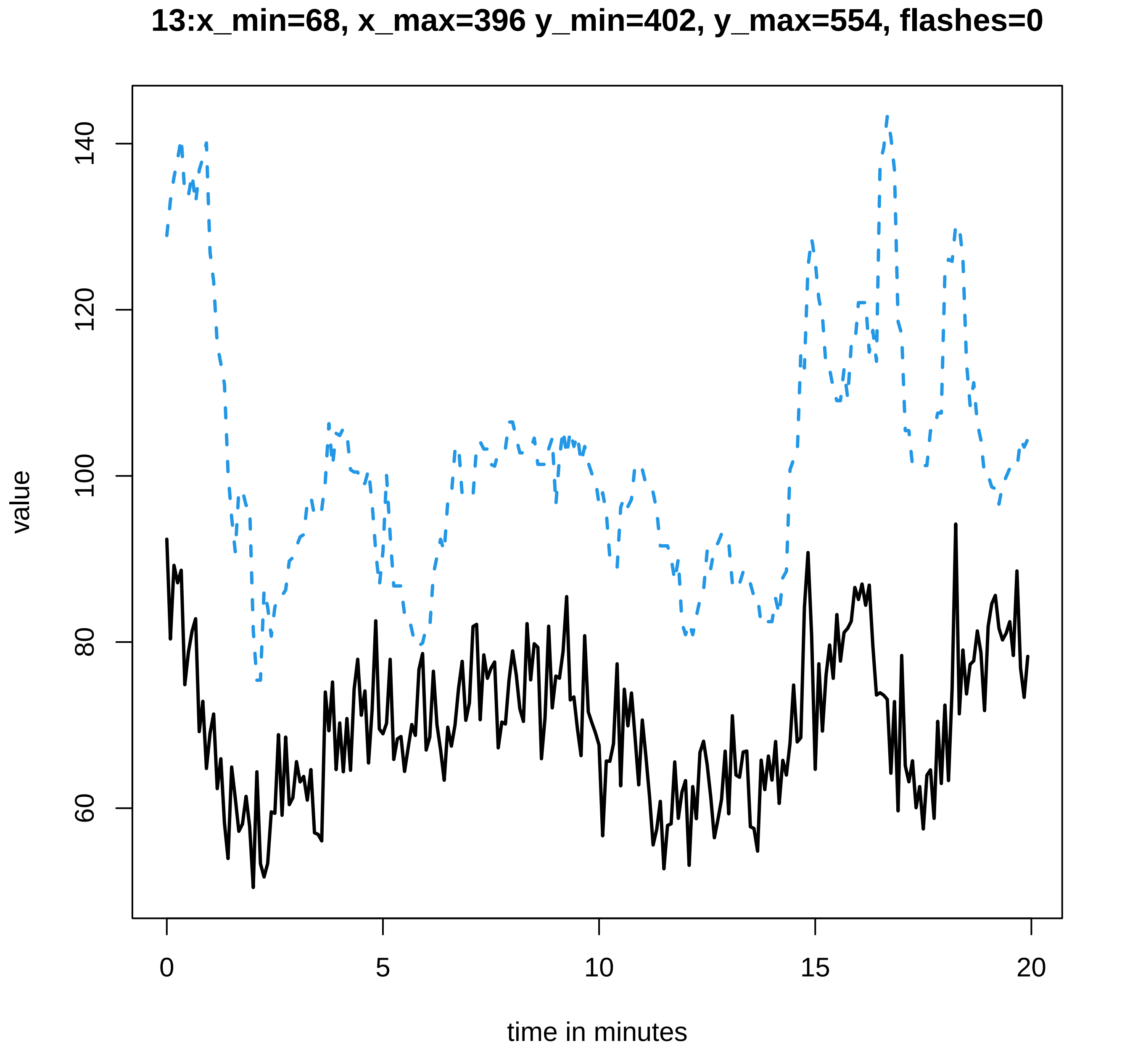

13:x_min=68, x_max=396 y_min=402, y_max=554, flashes=0
value
140
120
100
80
60
0
5
10
15
20
time in minutes

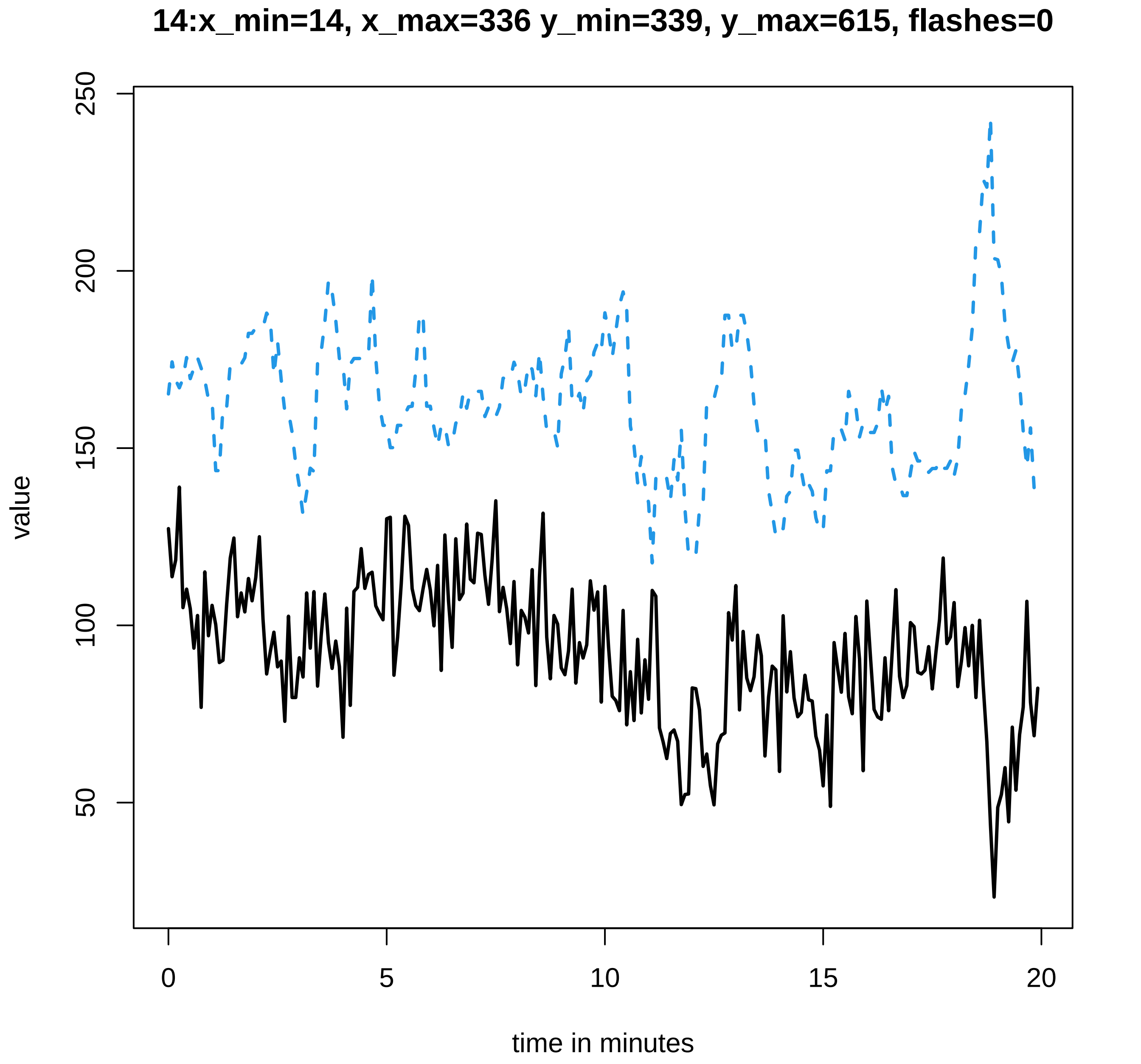
14:x_min=14, x_max=336 y_min=339, y_max=615, flashes=0
value
250
200
150
100
50
0
5
10
15
20
time in minutes

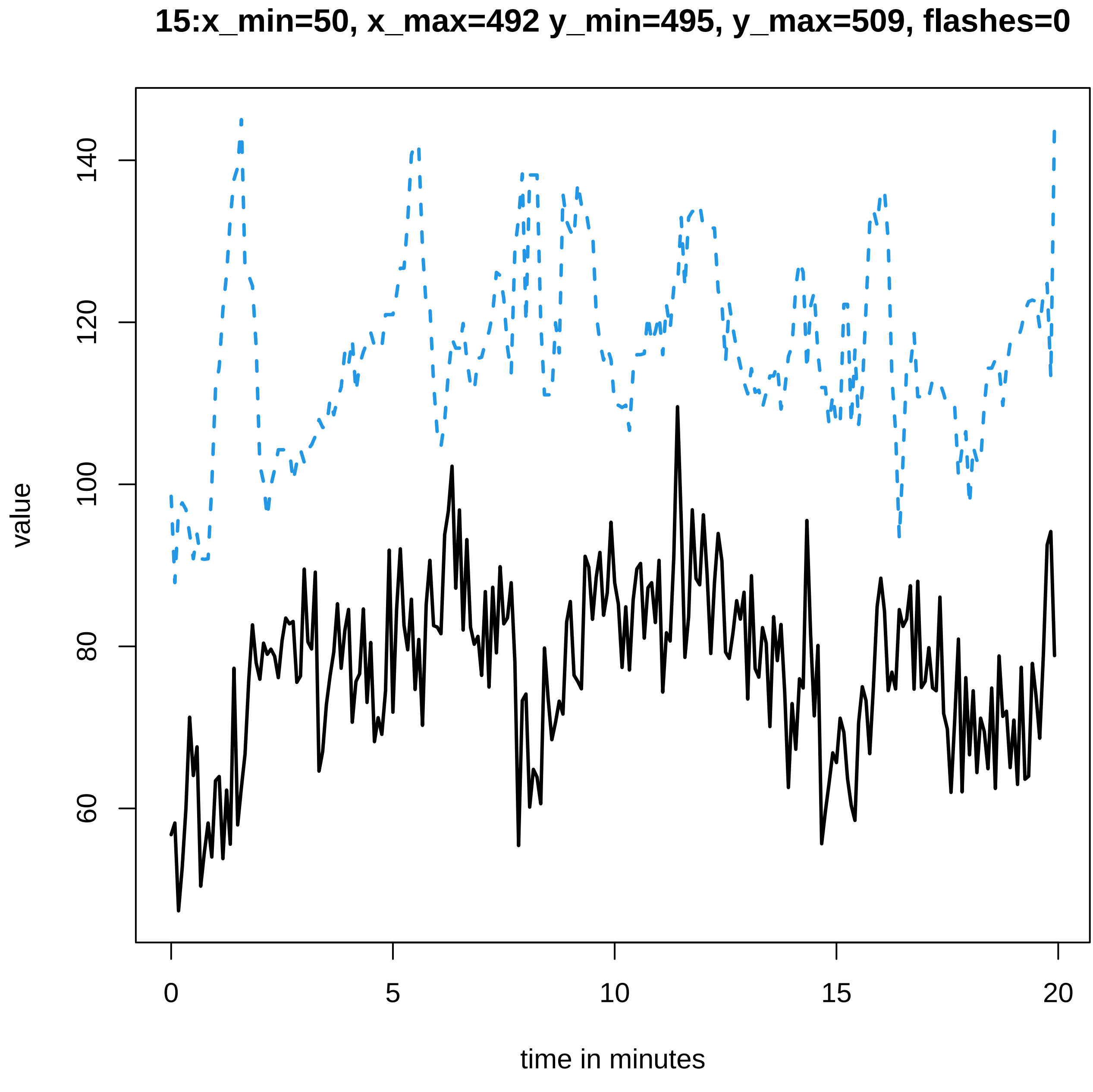

15:x_min=50, x_max=492 y_min=495, y_max=509, flashes=0
value
60
80
100
120
140
0
5
10
15
20
time in minutes

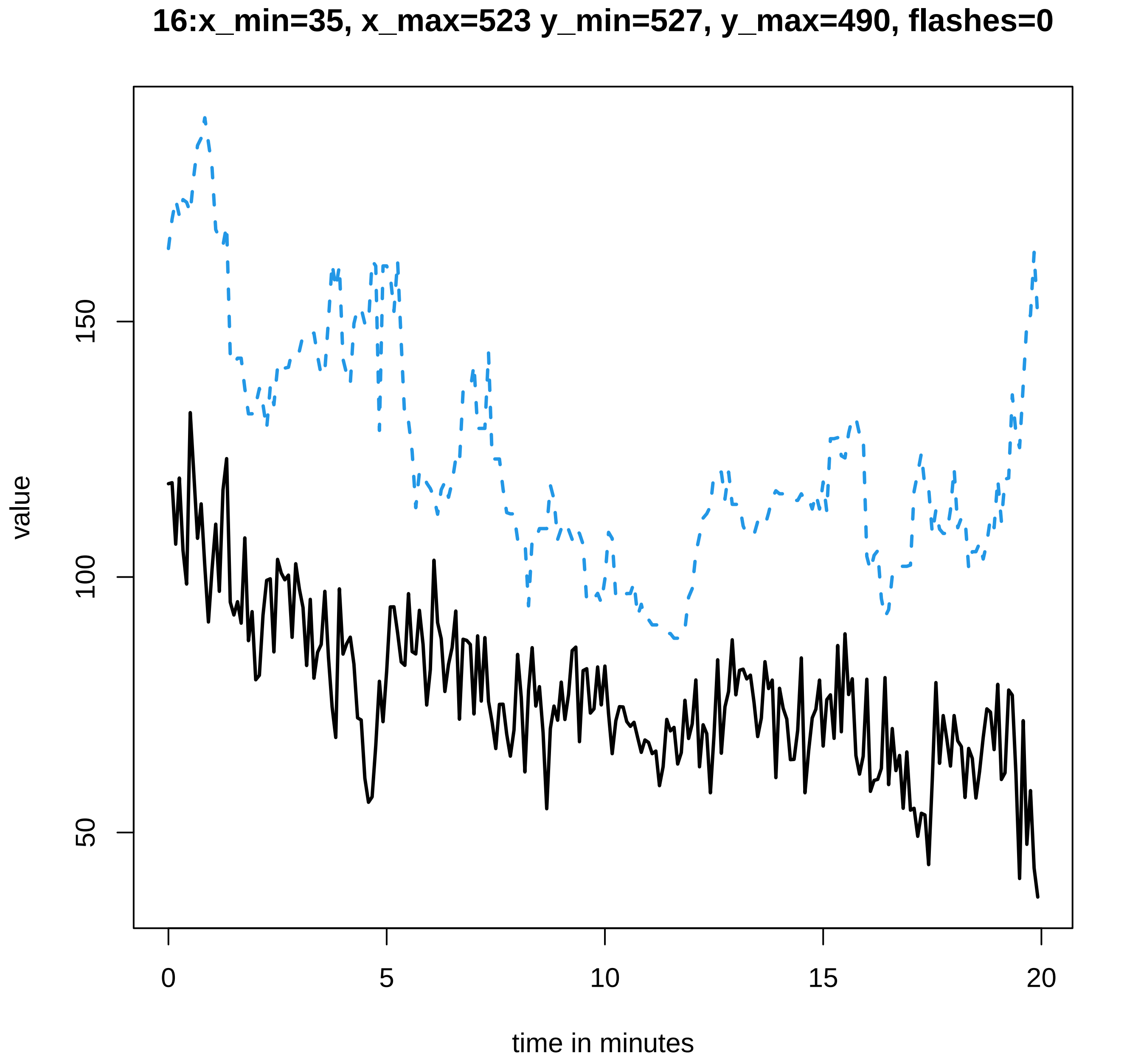
16:x_min=35, x_max=523 y_min=527, y_max=490, flashes=0
value
150
100
50
0
5
10
15
20
time in minutes

#### 17:x_min=35, x_max=547 y_min=554, y_max=509, flashes=0

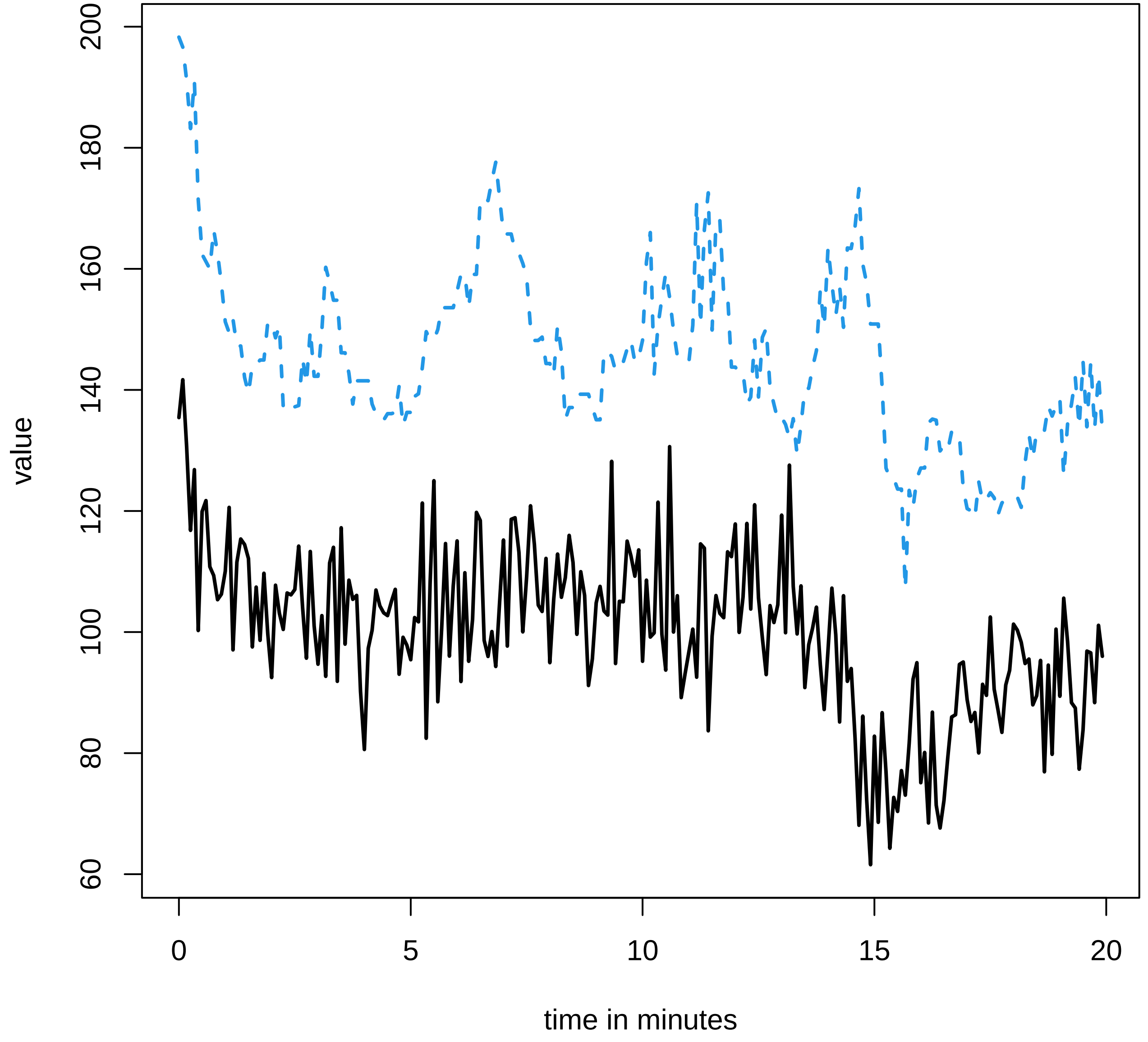

**18:x_min=34, x_max=399 y_min=402, y_max=664, flashes=0**

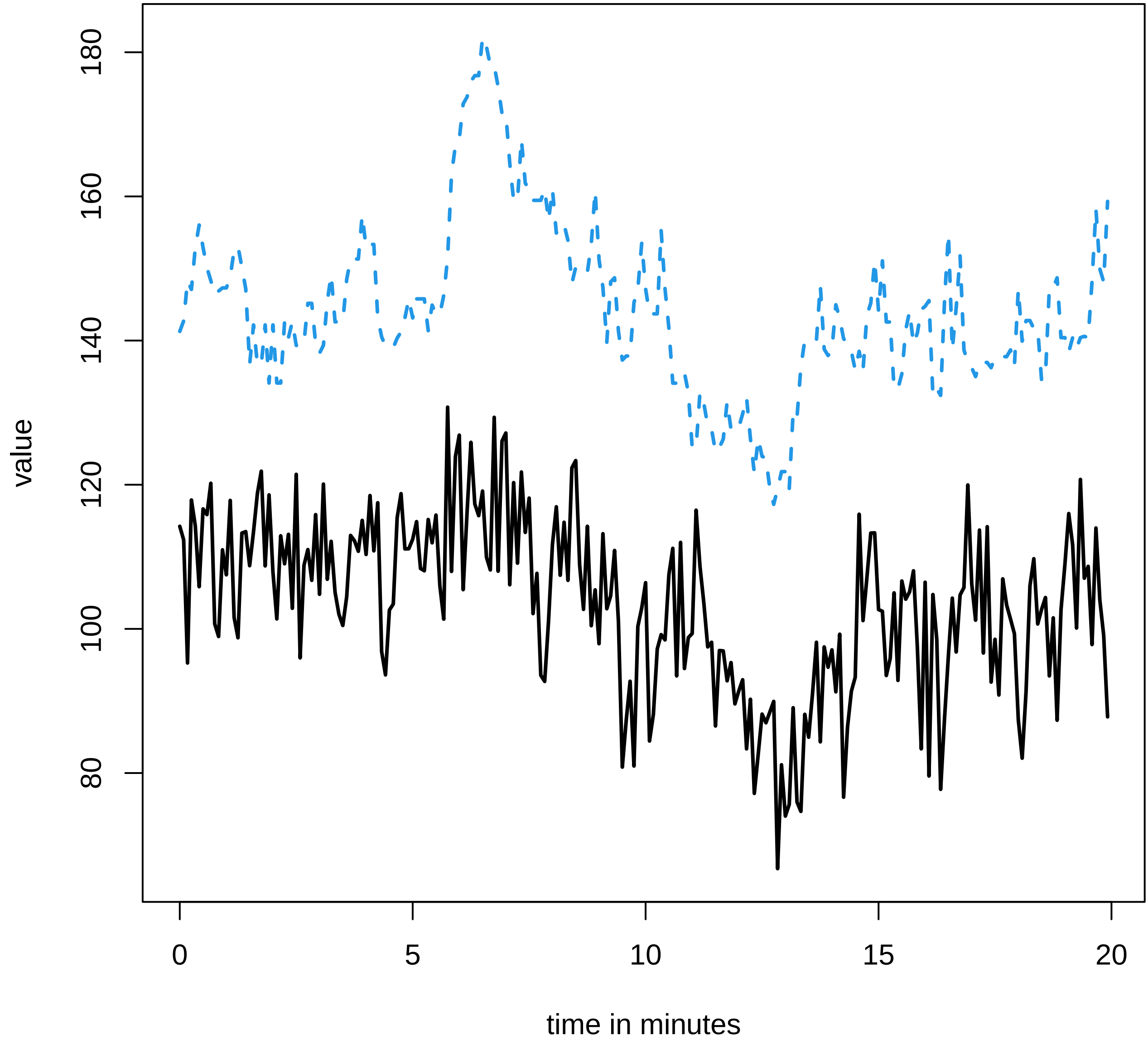

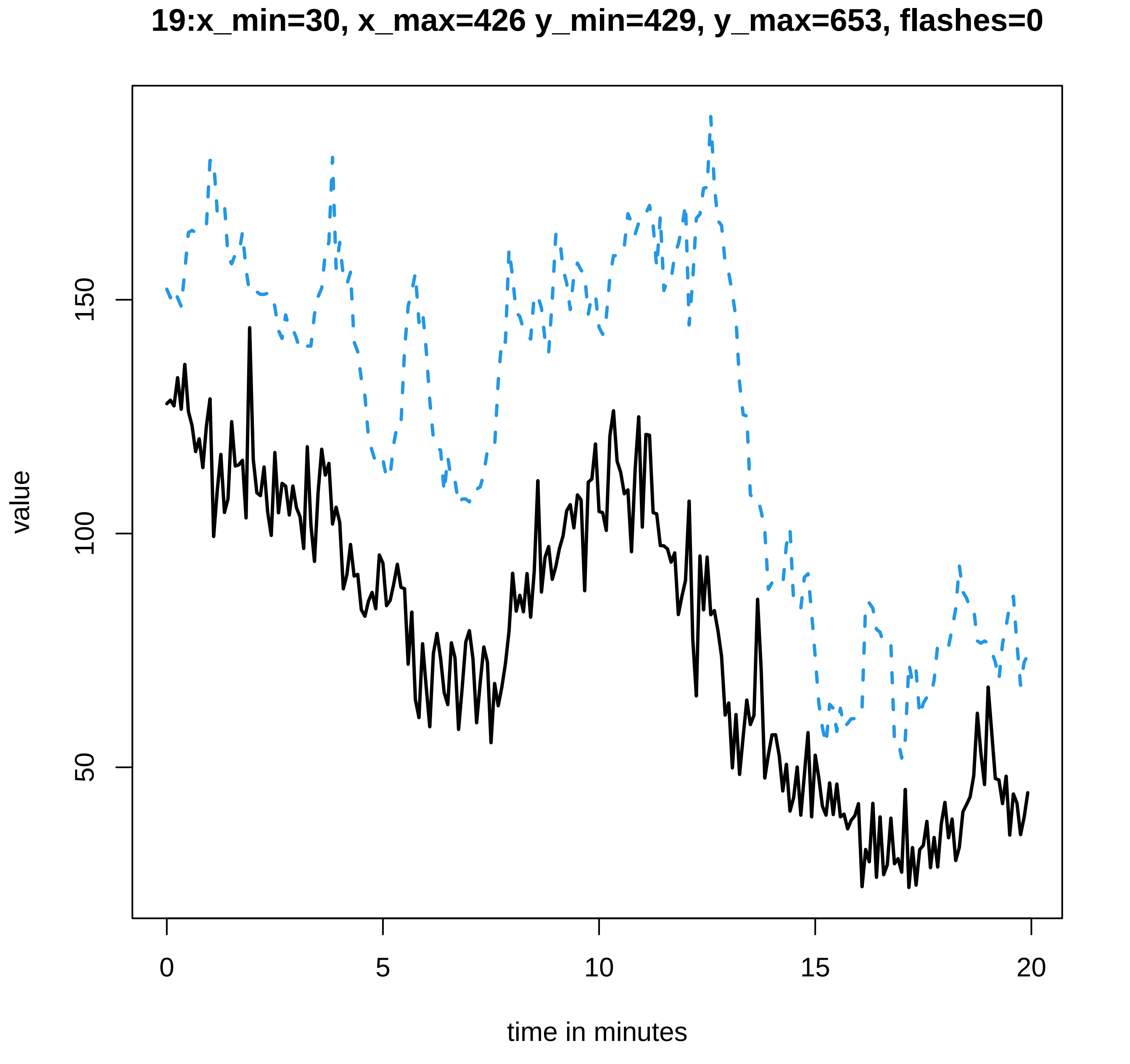

19:x_min=30, x_max=426 y_min=429, y_max=653, flashes=0
value
150
100
50
0
5
10
15
20
time in minutes

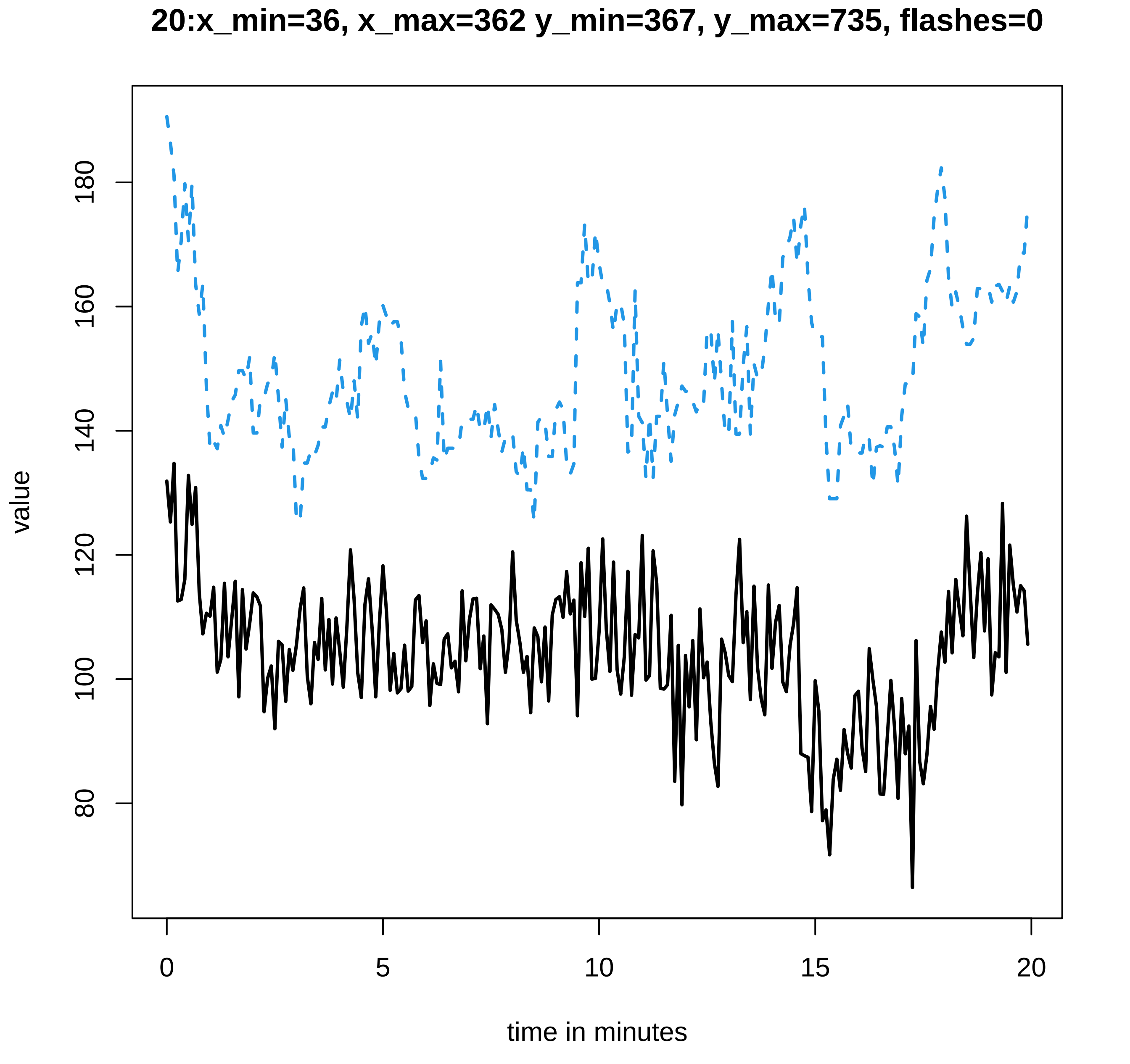

20:x_min=36, x_max=362 y_min=367, y_max=735, flashes=0
value
180
160
140
120
100
80
0
5
10
15
20
time in minutes

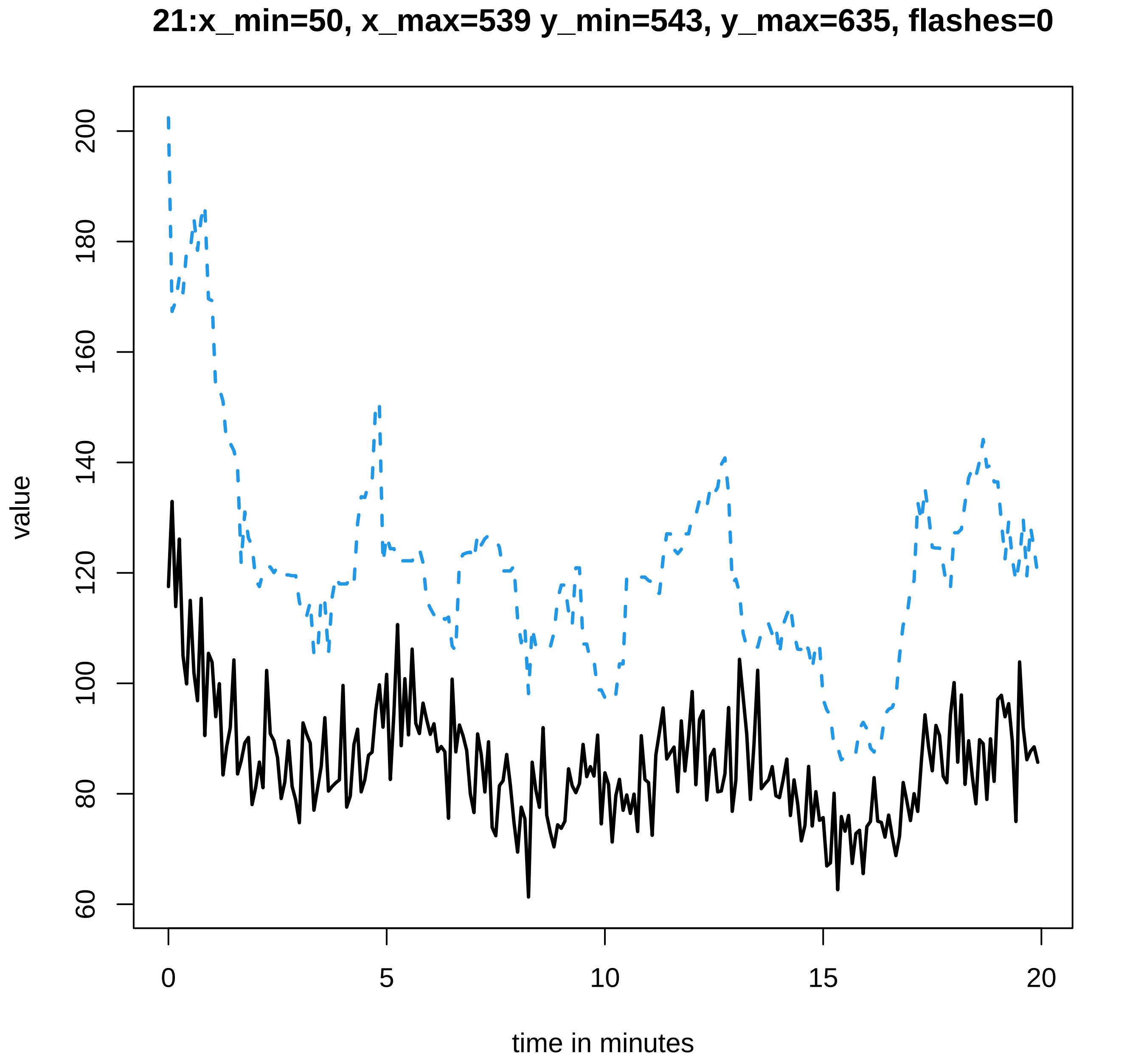
21:x_min=50, x_max=539 y_min=543, y_max=635, flashes=0
value
200
180
160
140
120
100
80
60
0
5
10
15
20
time in minutes

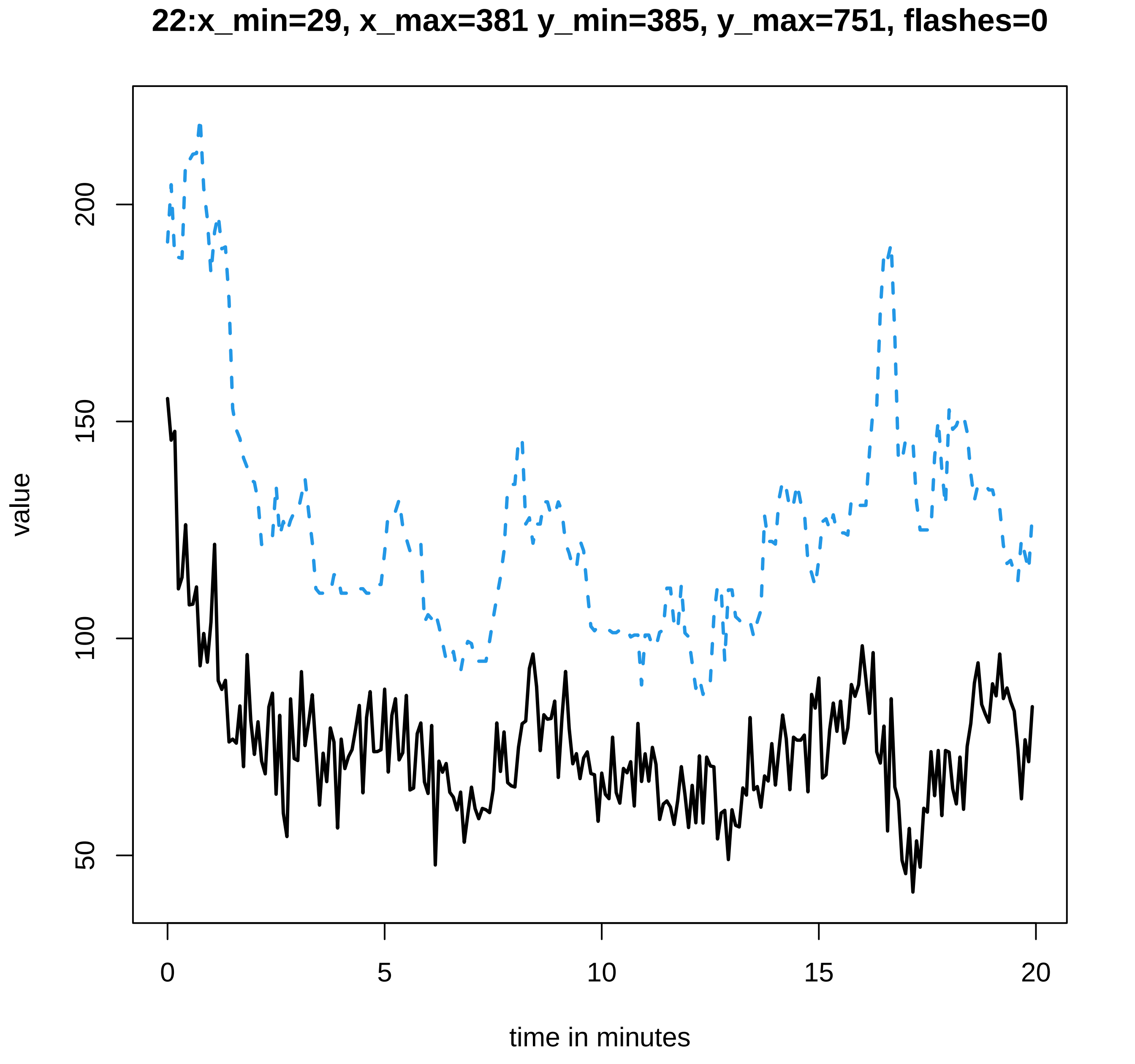
22:x_min=29, x_max=381 y_min=385, y_max=751, flashes=0
value
200
150
100
50
0
5
10
15
20
time in minutes

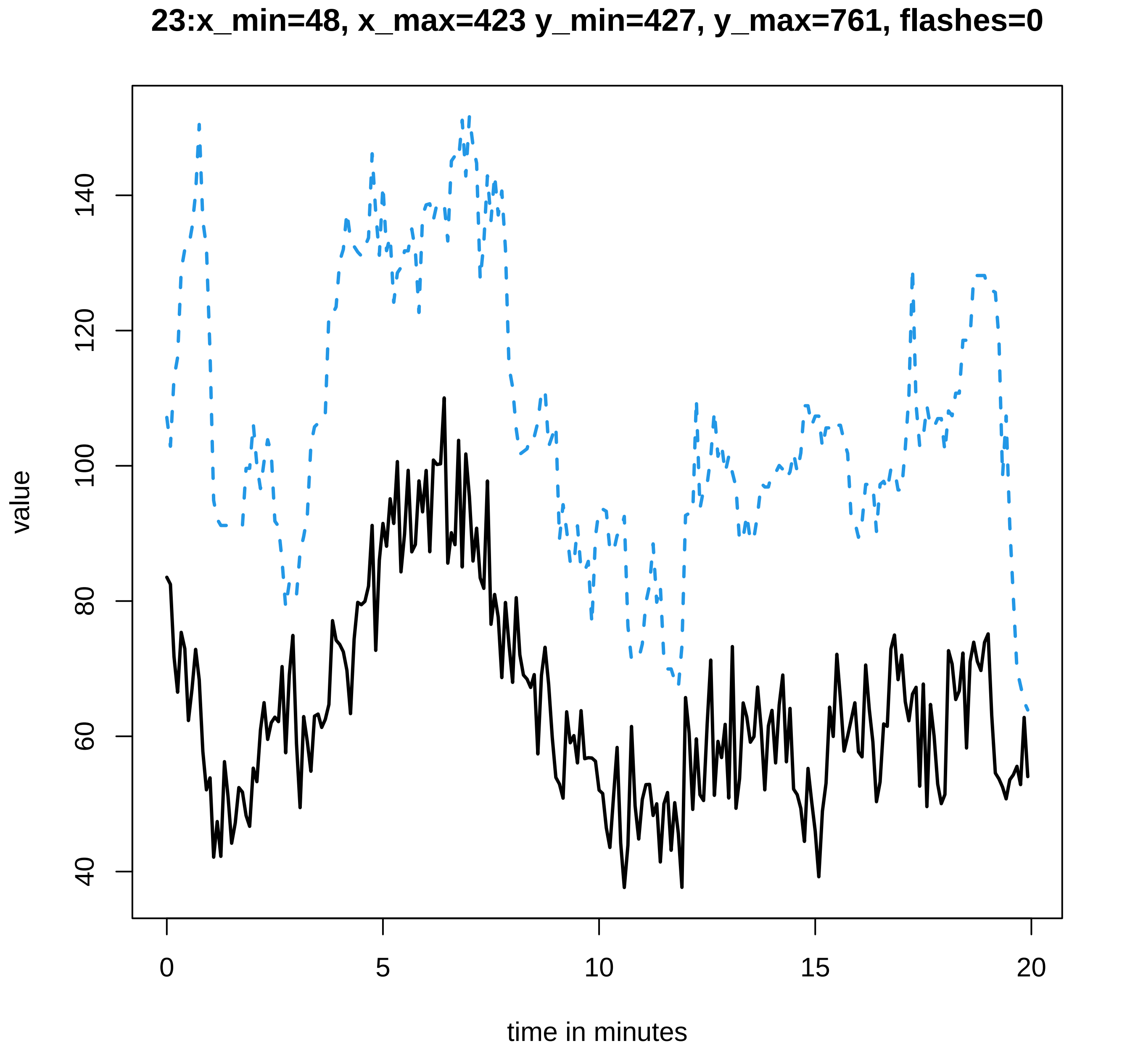

23:x_min=48, x_max=423 y_min=427, y_max=761, flashes=0
value
40
60
80
100
120
140
0
5
10
15
20
time in minutes

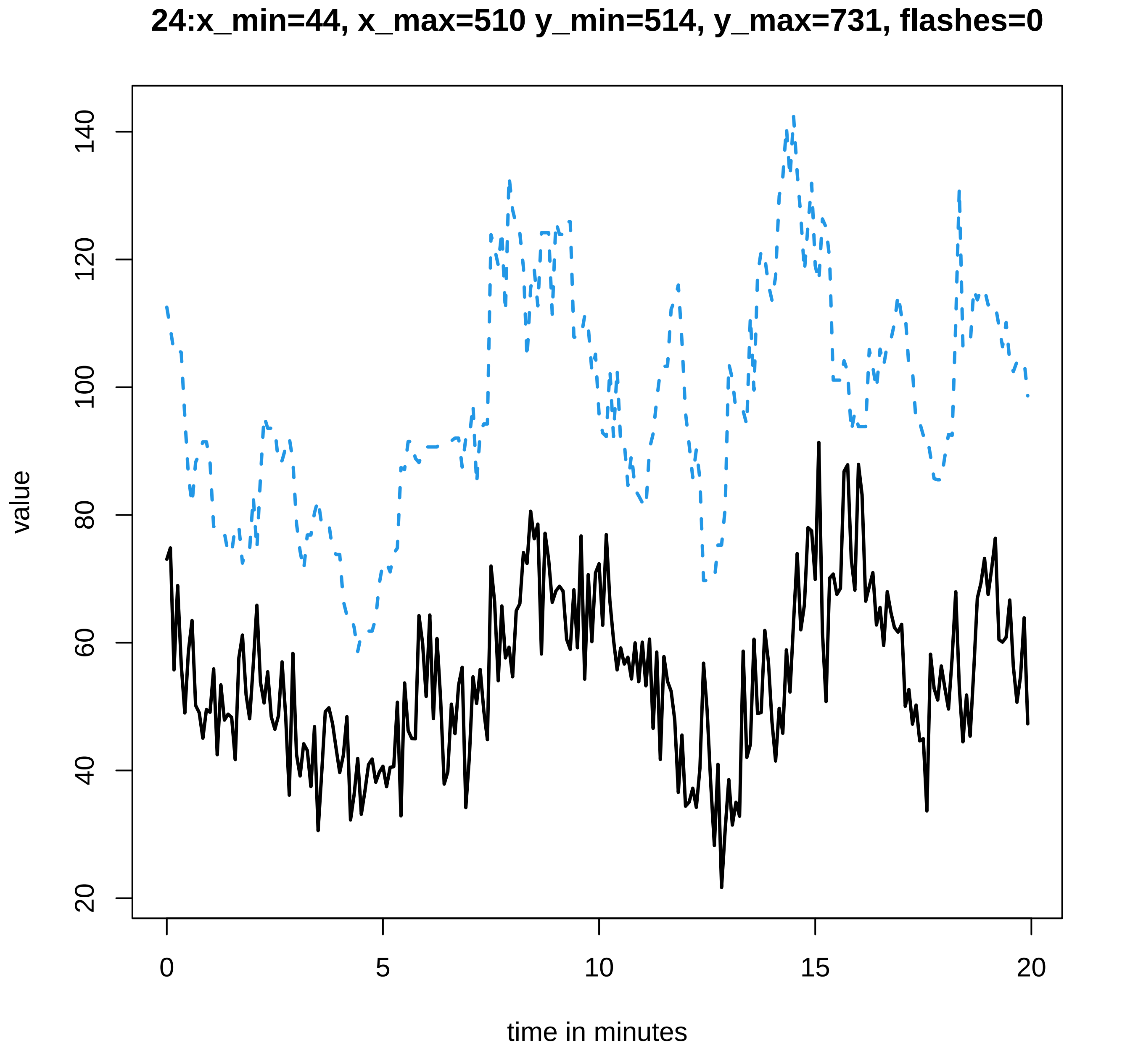

24:x_min=44, x_max=510 y_min=514, y_max=731, flashes=0
value
20
40
60
80
100
120
140
0
5
10
15
20
time in minutes

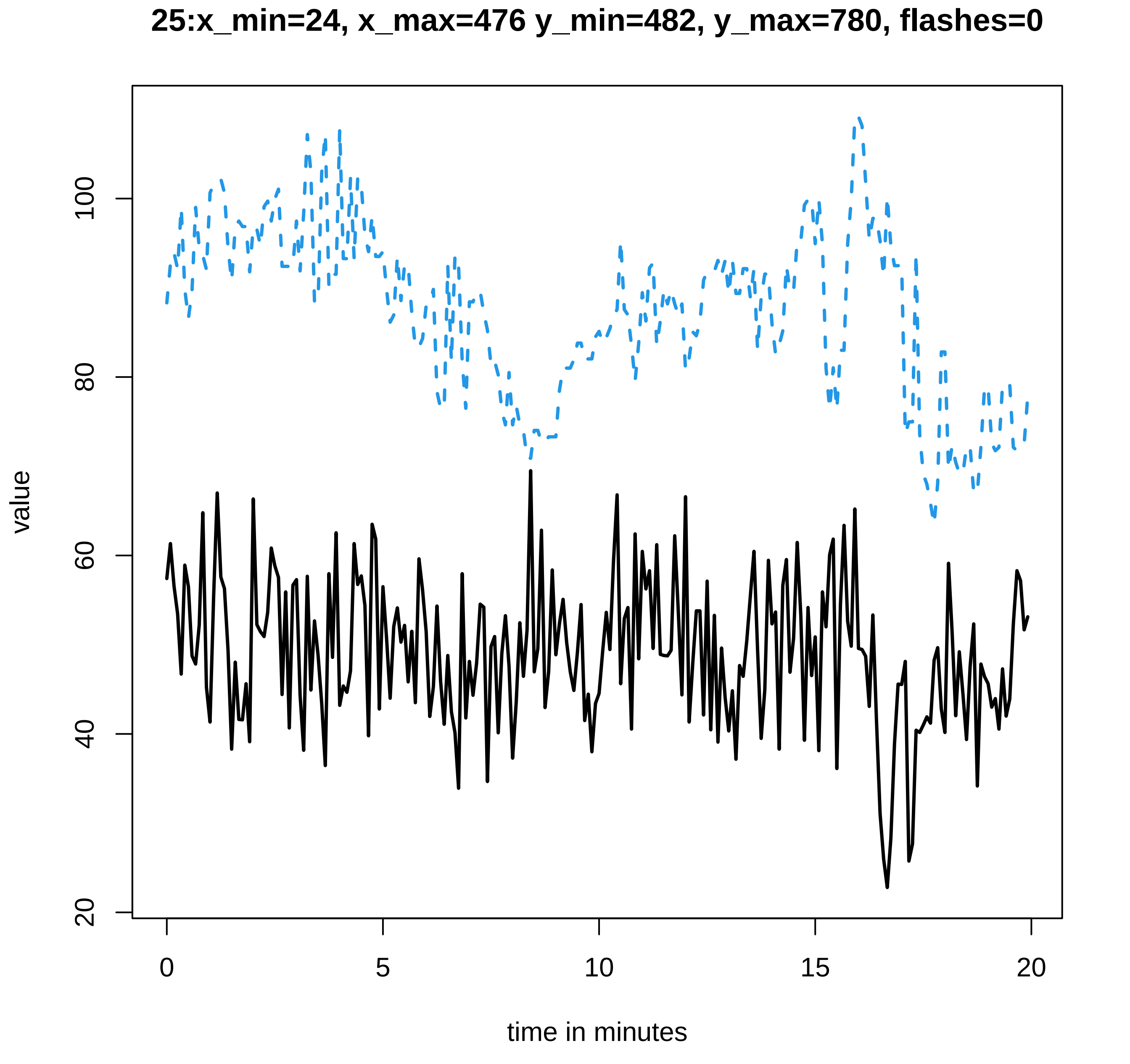

25:x_min=24, x_max=476 y_min=482, y_max=780, flashes=0
value
20
40
60
80
100
0
5
10
15
20
time in minutes

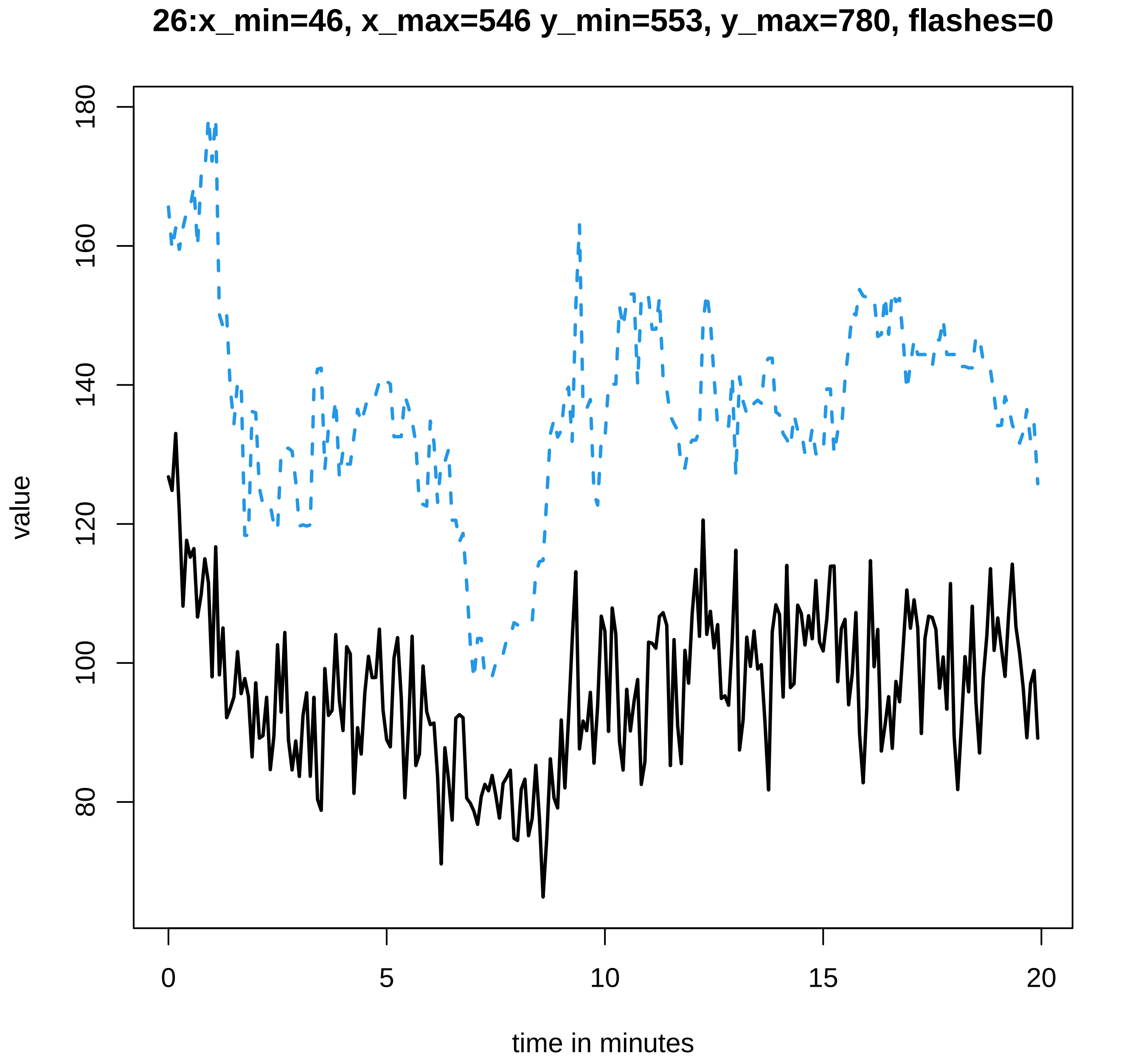
26:x_min=46, x_max=546 y_min=553, y_max=780, flashes=0
value
180
160
140
120
100
80
0
5
10
15
20
time in minutes

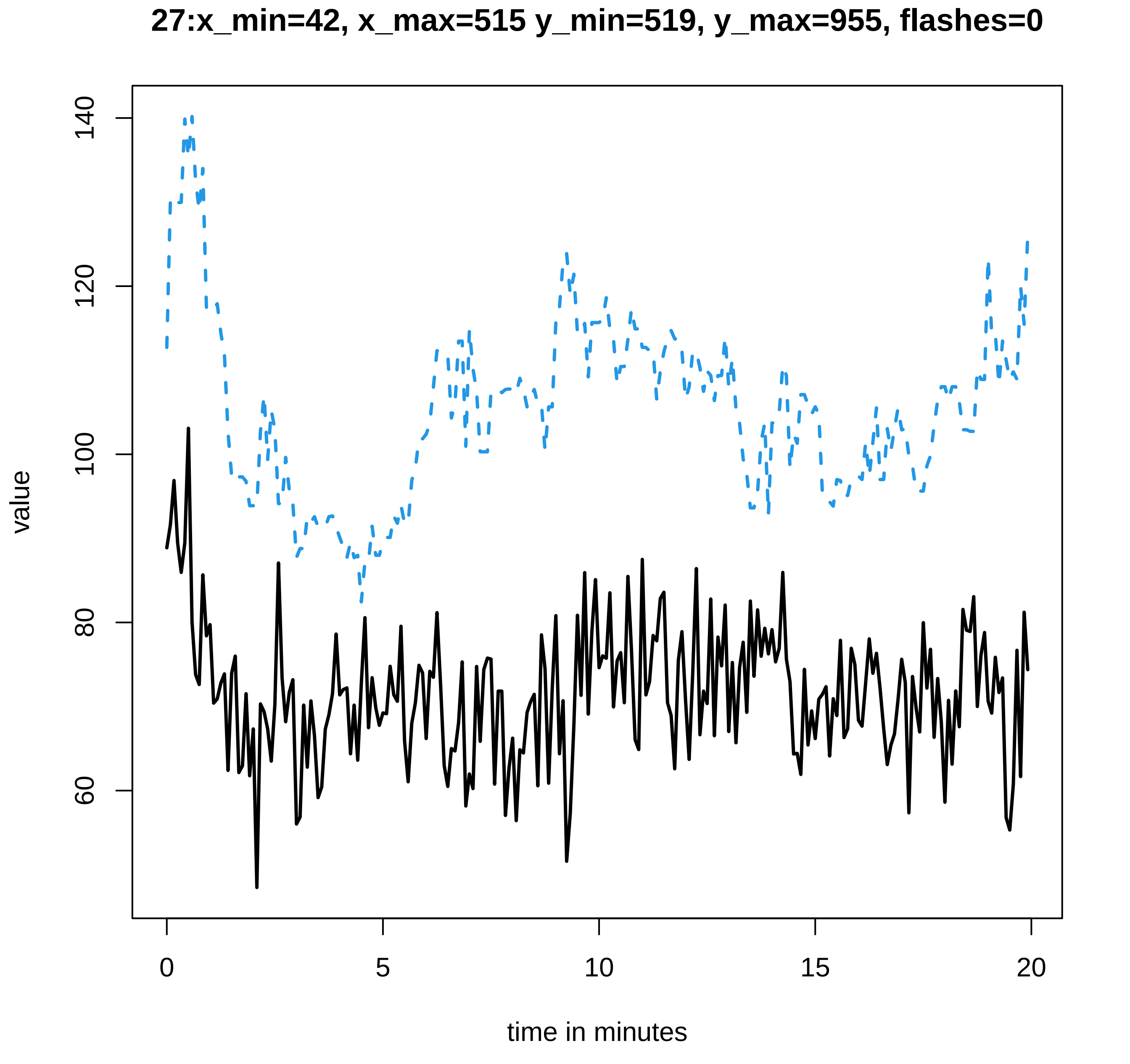
27:x_min=42, x_max=515 y_min=519, y_max=955, flashes=0
value
140
120
100
80
60
0
5
10
15
20
time in minutes
