## Supplementary data Calcium spiking for "LysM Receptor Proteins are Required for Ectomycorrhizal Symbiosis in Poplar": profiles-spiking-12.pdf

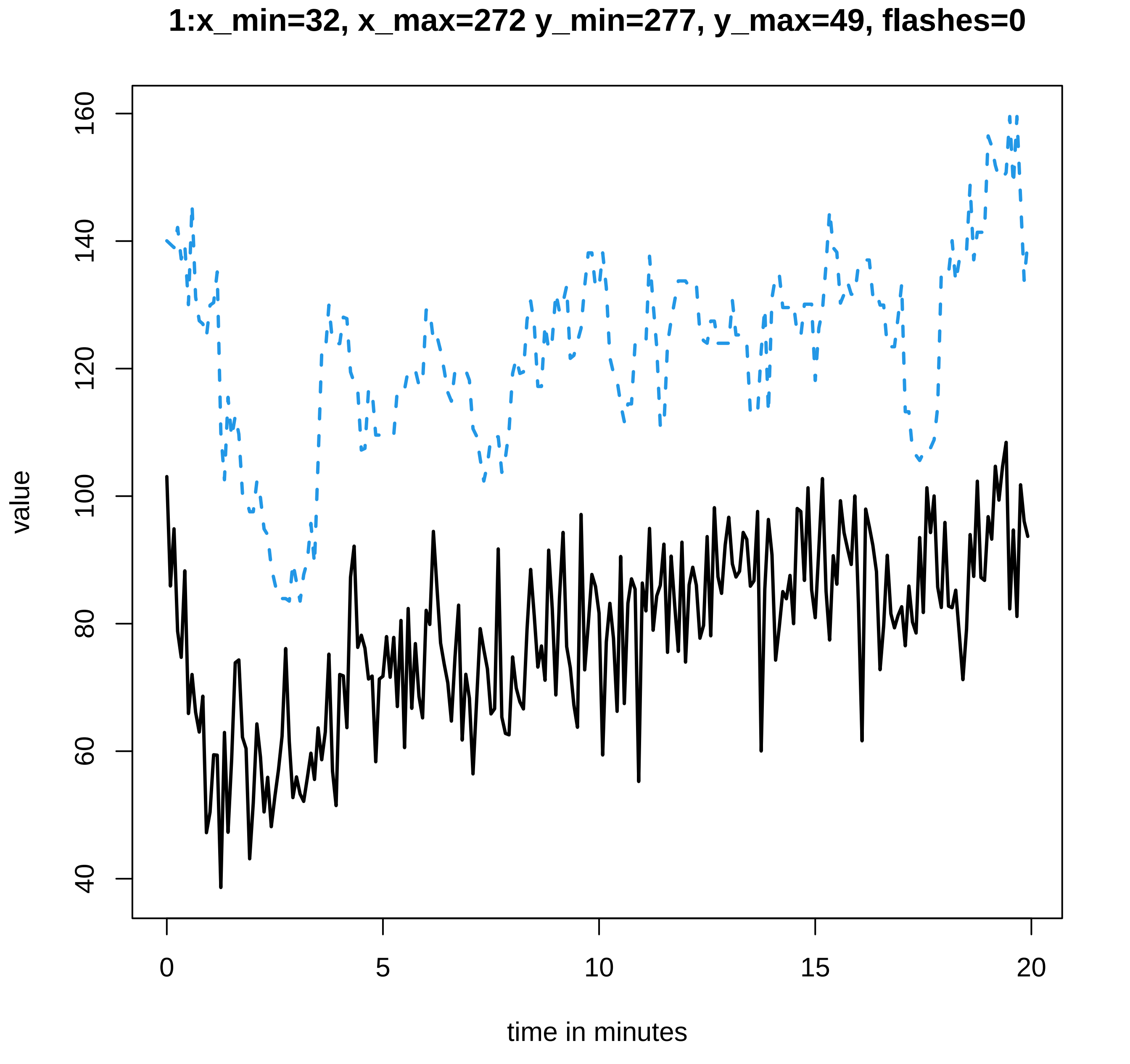

1:x_min=32, x_max=272 y_min=277, y_max=49, flashes=0
value
160
140
120
100
80
60
40
0
5
10
15
20
time in minutes

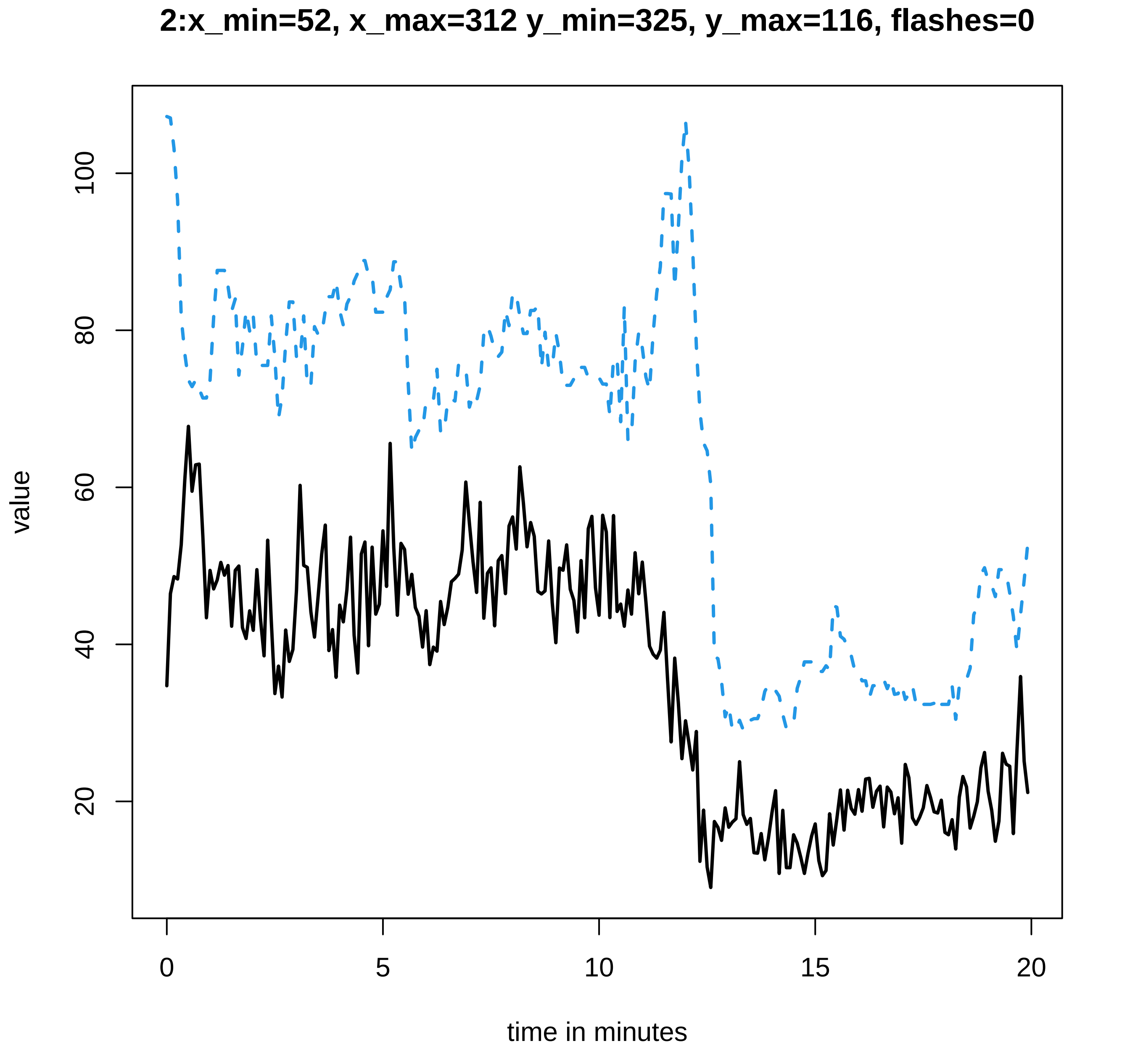

2:x_min=52, x_max=312 y_min=325, y_max=116, flashes=0
value
20
40
60
80
100
0
5
10
15
20
time in minutes

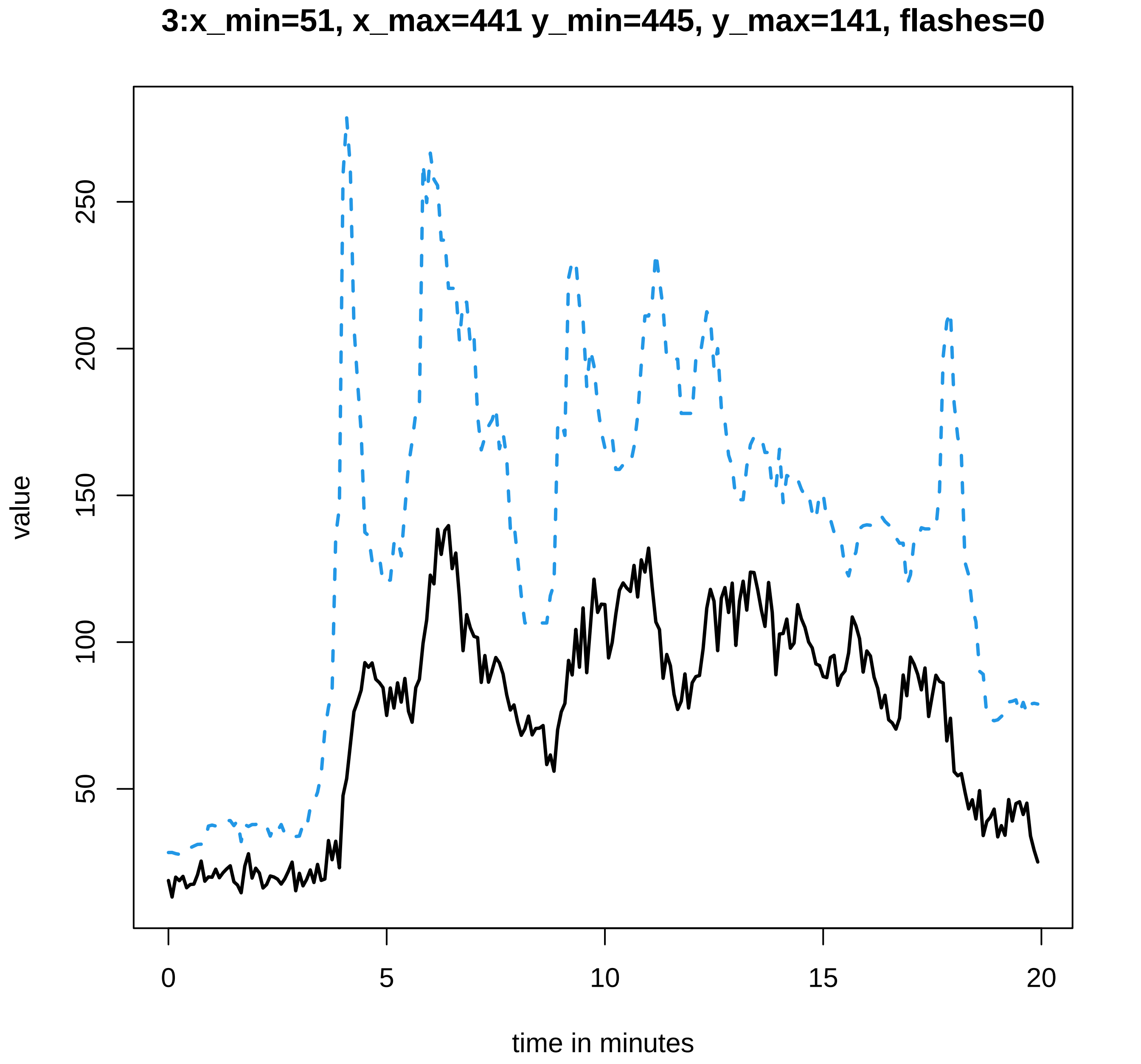
3:x_min=51, x_max=441 y_min=445, y_max=141, flashes=0
value
250
200
150
100
50
0
5
10
15
20
time in minutes

**4:x_min=48, x_max=385 y_min=389, y_max=272, flashes=0**

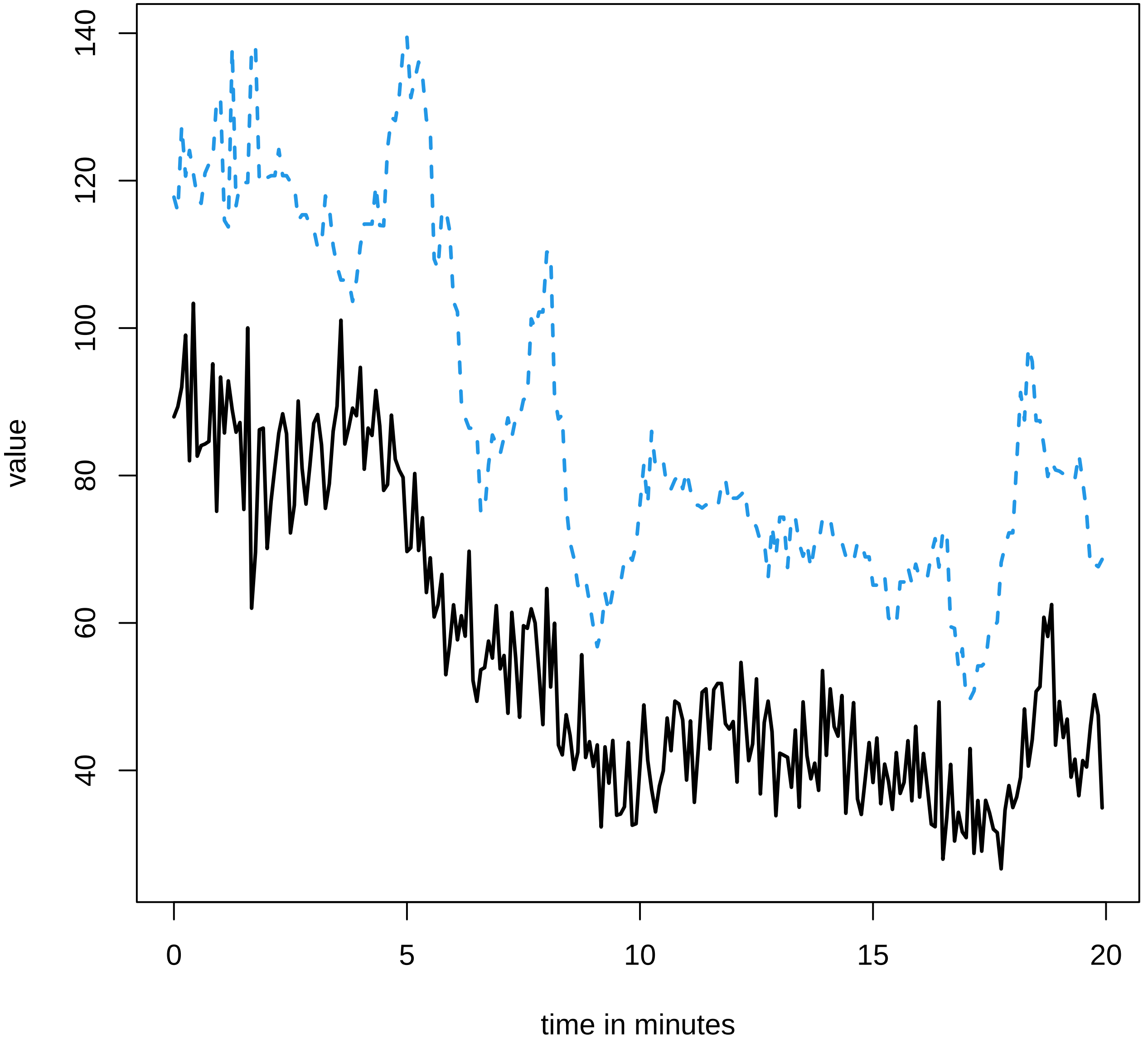

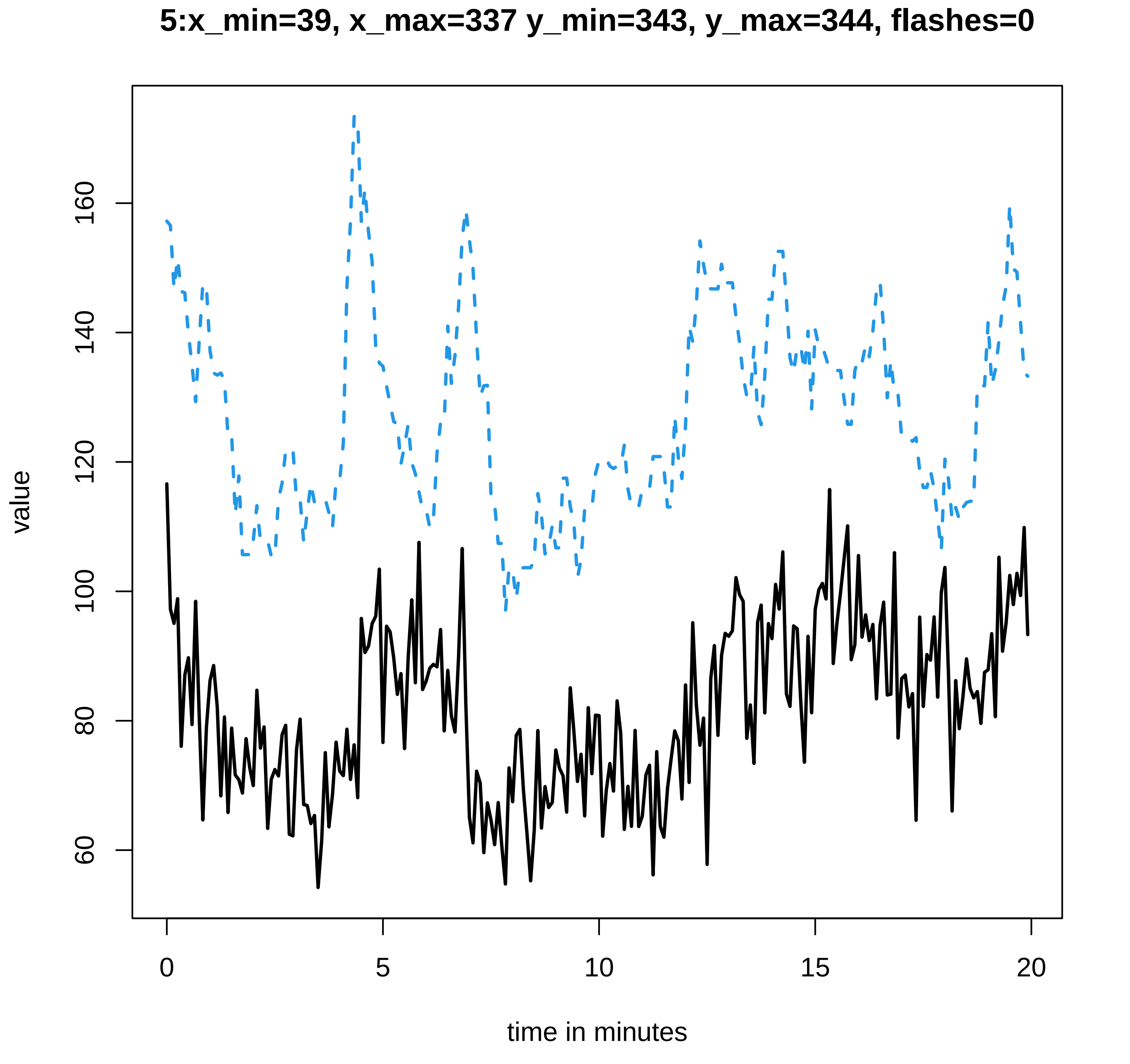

5:x_min=39, x_max=337 y_min=343, y_max=344, flashes=0
value
160
140
120
100
80
60
0
5
10
15
20
time in minutes

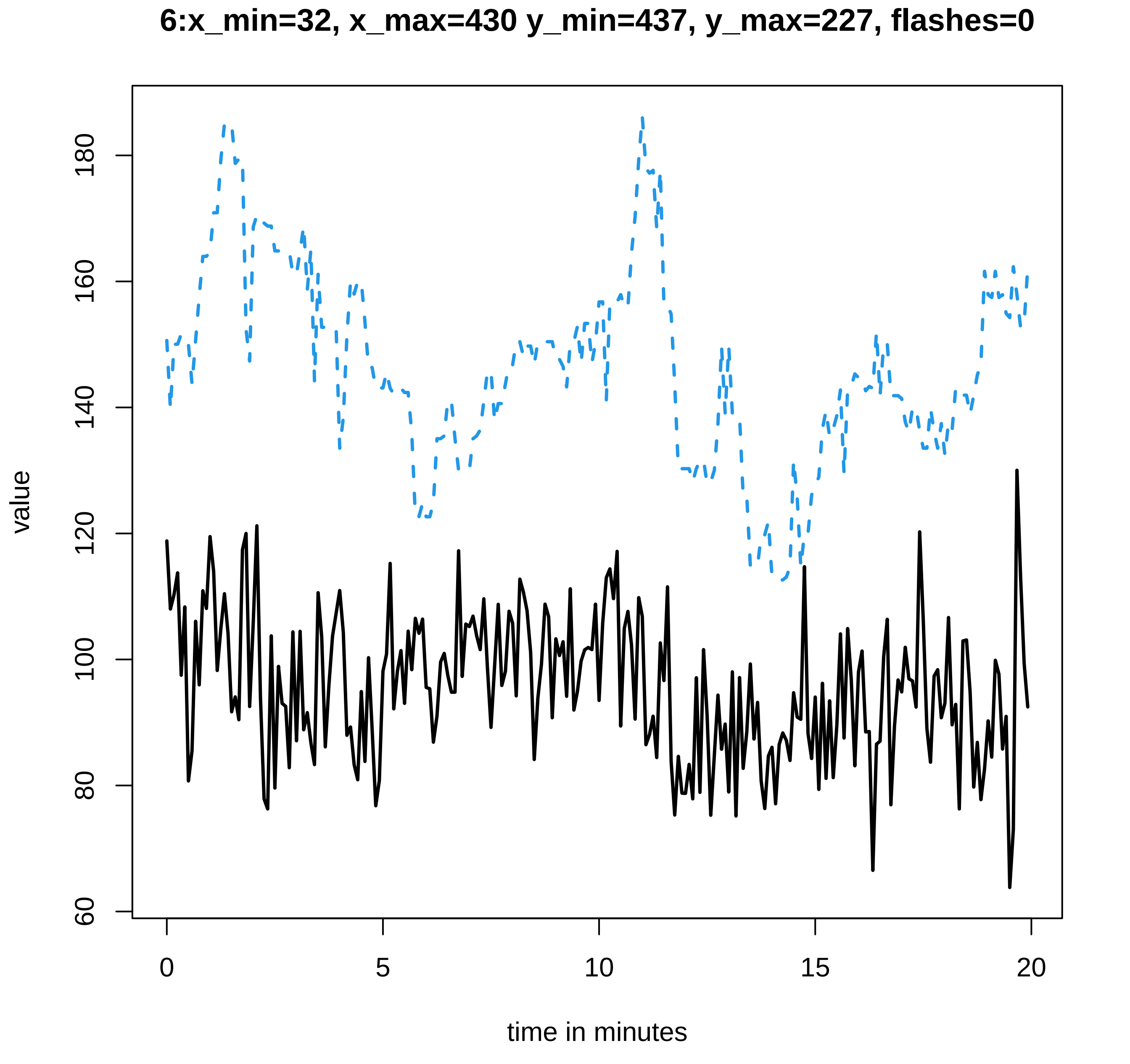

6:x_min=32, x_max=430 y_min=437, y_max=227, flashes=0
value
time in minutes
0
5
10
15
20
60
80
100
120
140
160
180

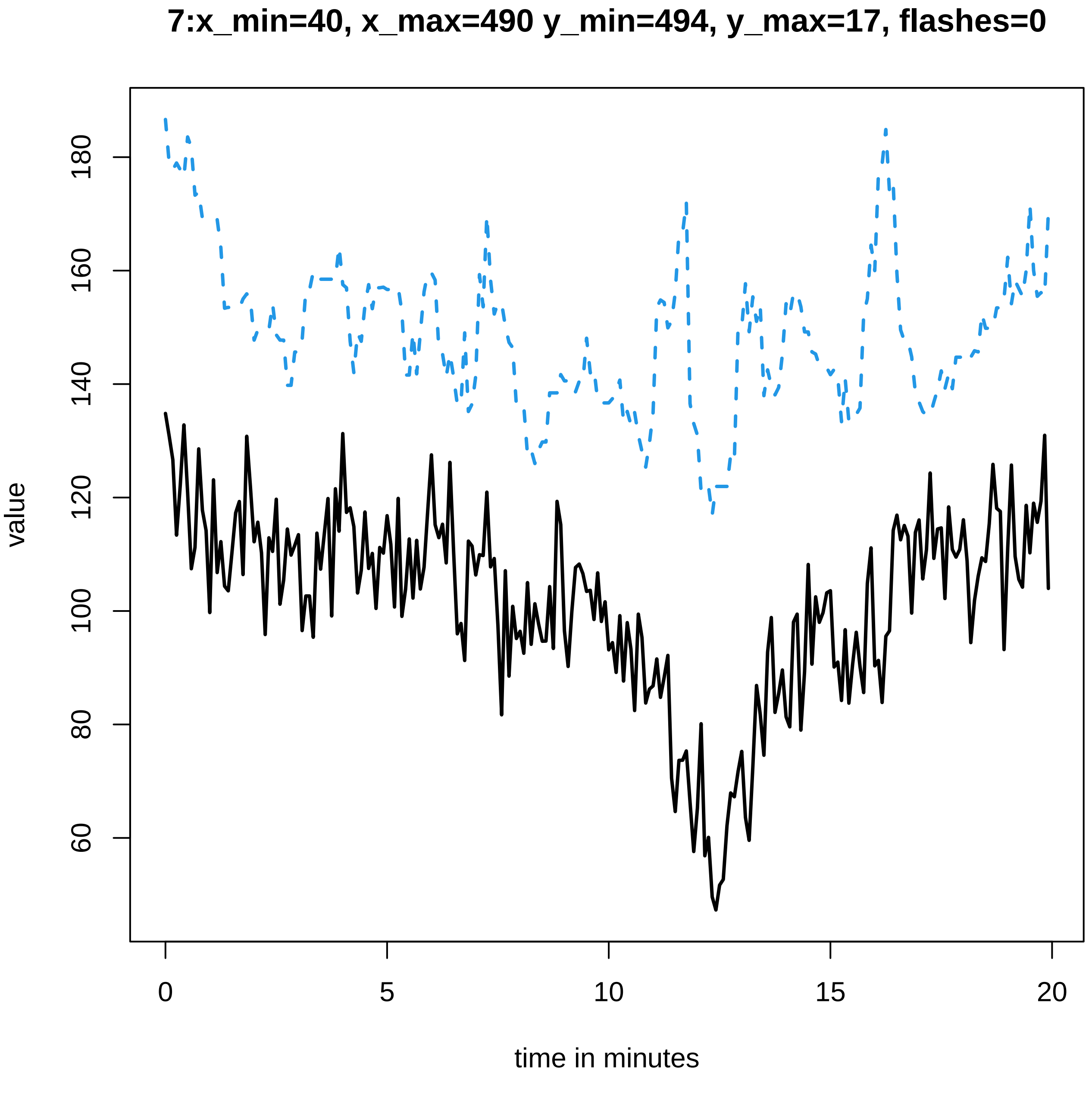
7:x_min=40, x_max=490 y_min=494, y_max=17, flashes=0
value
180
160
140
120
100
80
60
0
5
10
15
20
time in minutes

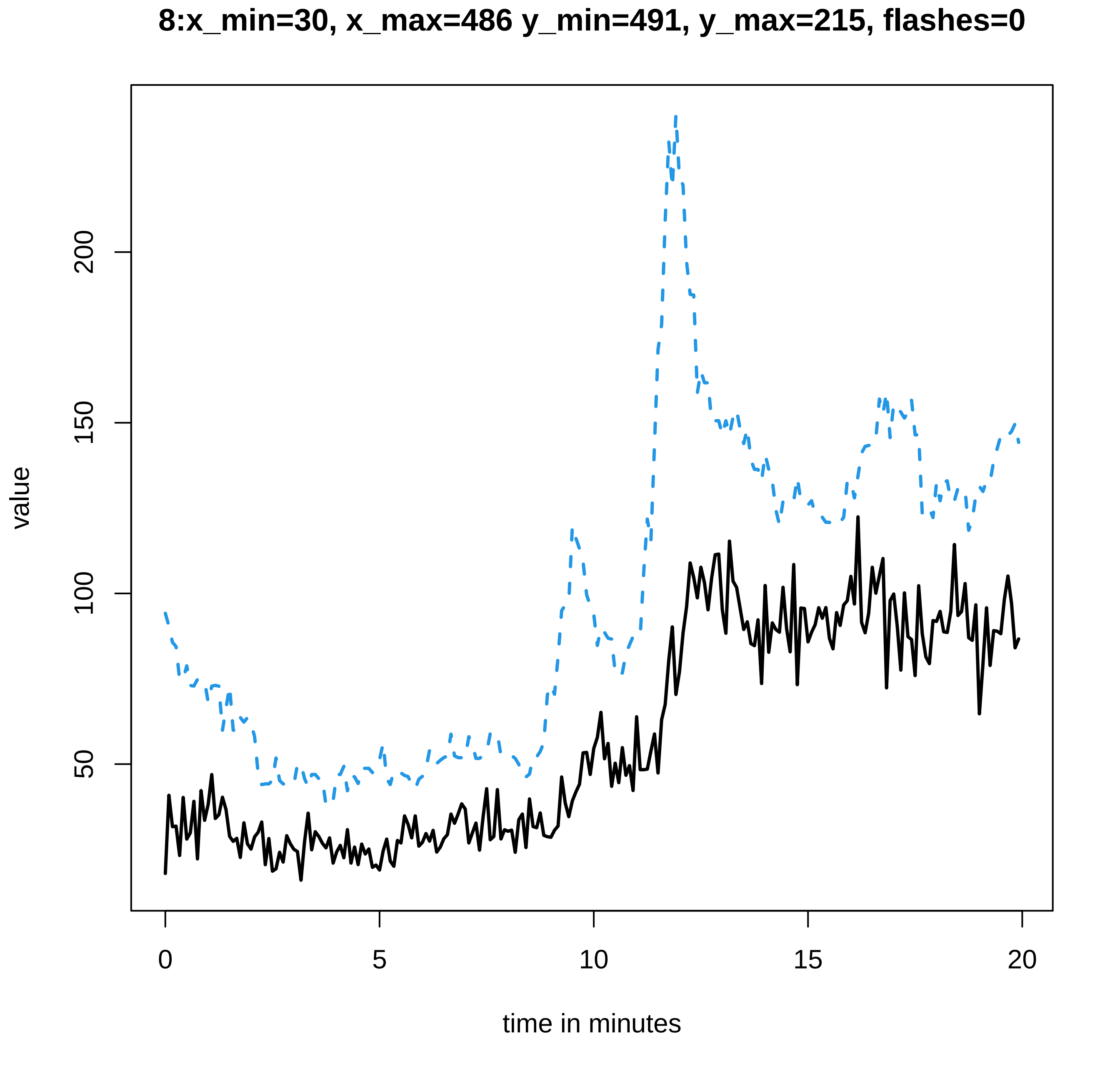

8:x_min=30, x_max=486 y_min=491, y_max=215, flashes=0
value
200
150
100
50
0
5
10
15
20
time in minutes

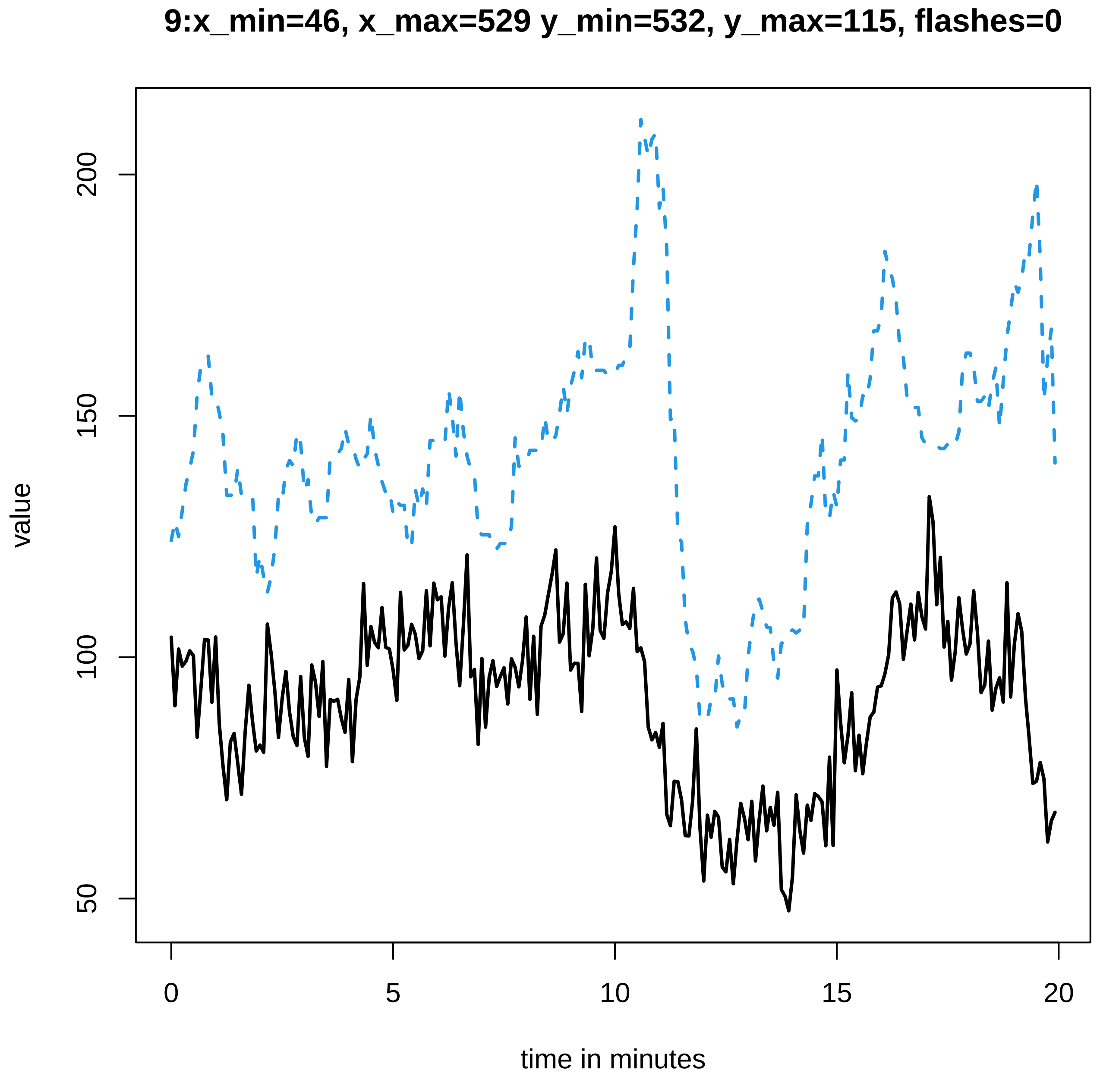

9:x_min=46, x_max=529 y_min=532, y_max=115, flashes=0
value
200
150
100
50
0
5
10
15
20
time in minutes

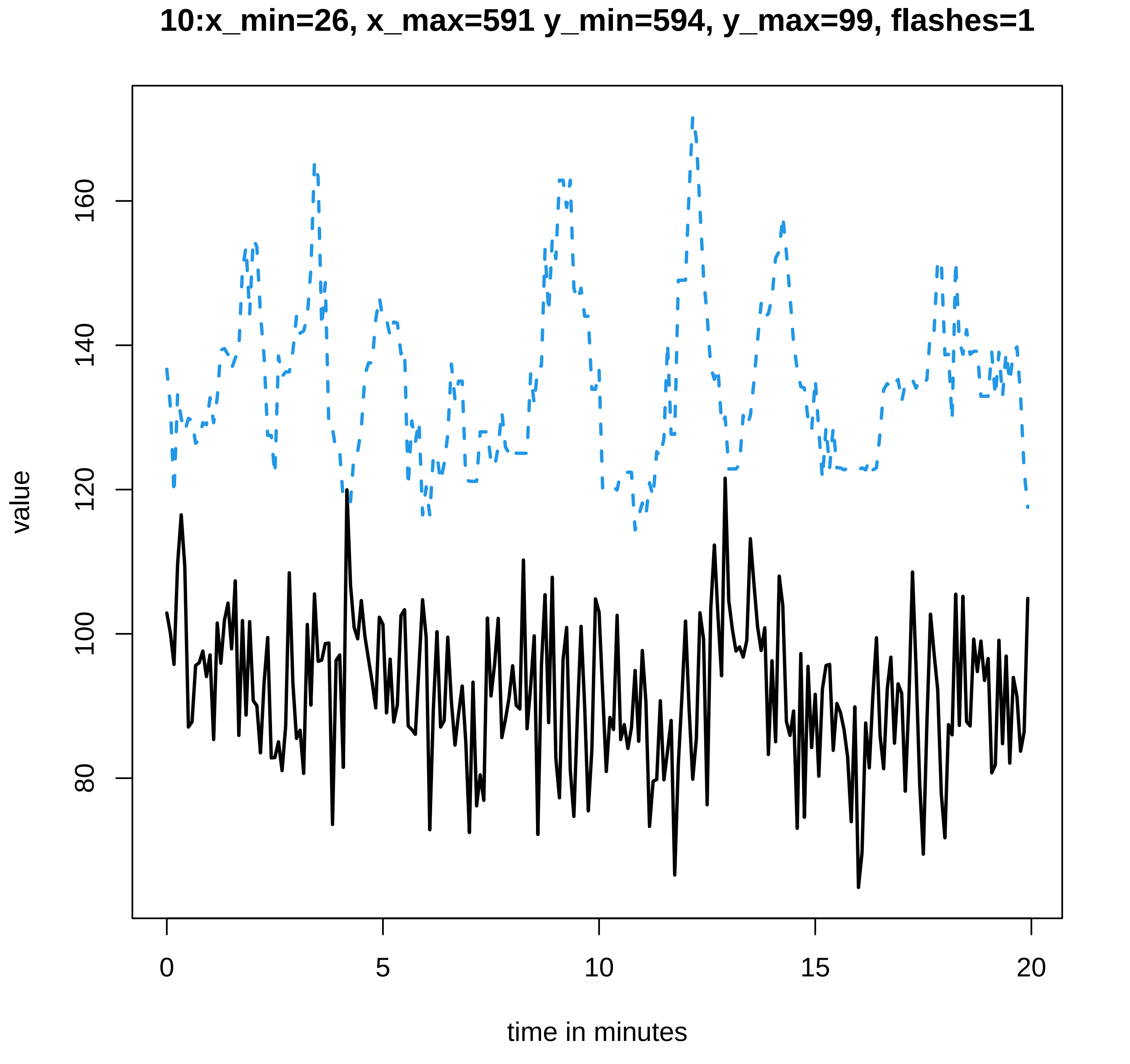
10:x_min=26, x_max=591 y_min=594, y_max=99, flashes=1
value
160
140
120
100
80
0
5
10
15
20
time in minutes

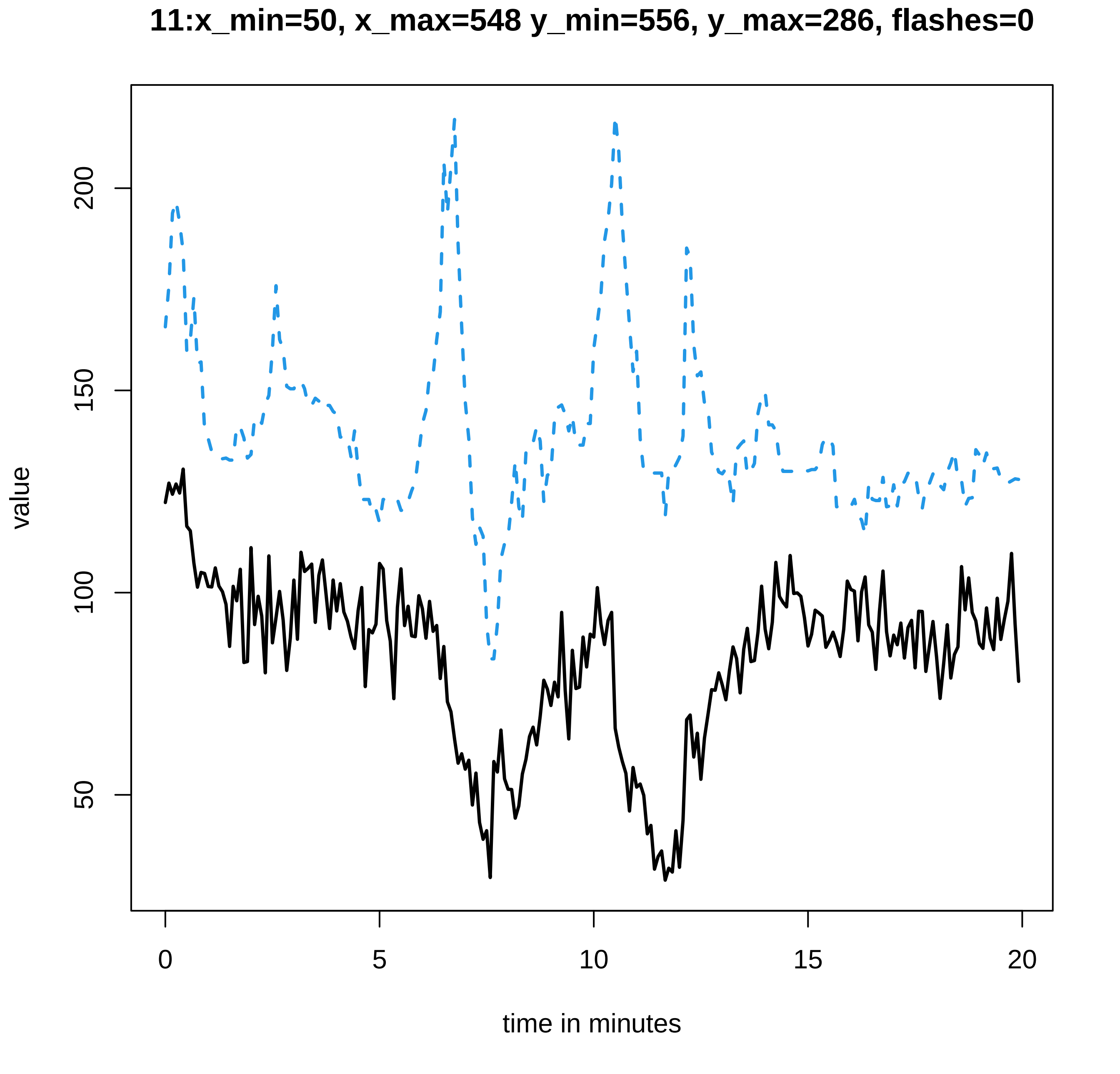

11:x_min=50, x_max=548 y_min=556, y_max=286, flashes=0
value
50
100
150
200
0
5
10
15
20
time in minutes

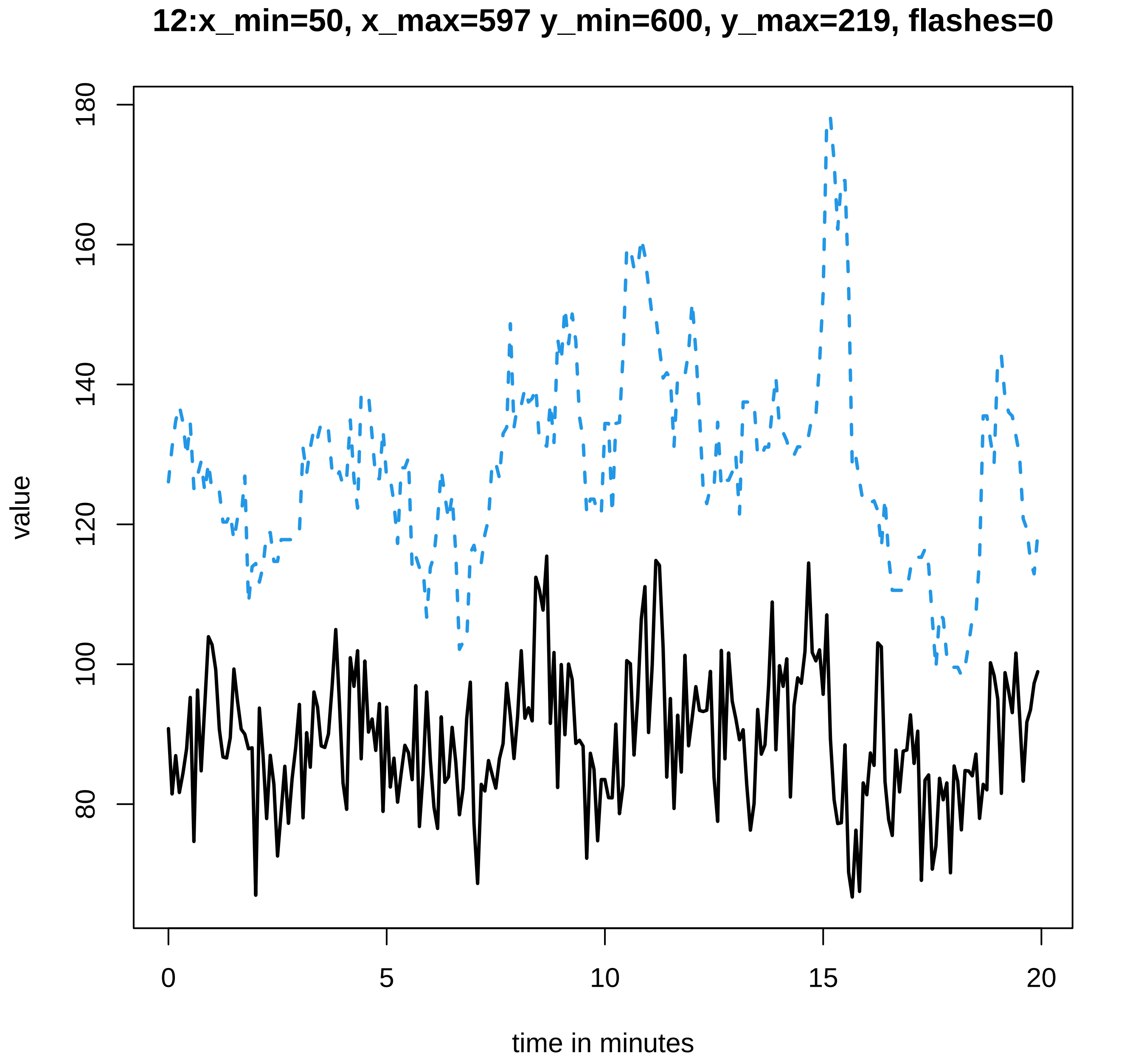
12:x_min=50, x_max=597 y_min=600, y_max=219, flashes=0
value
180
160
140
120
100
80
0
5
10
15
20
time in minutes

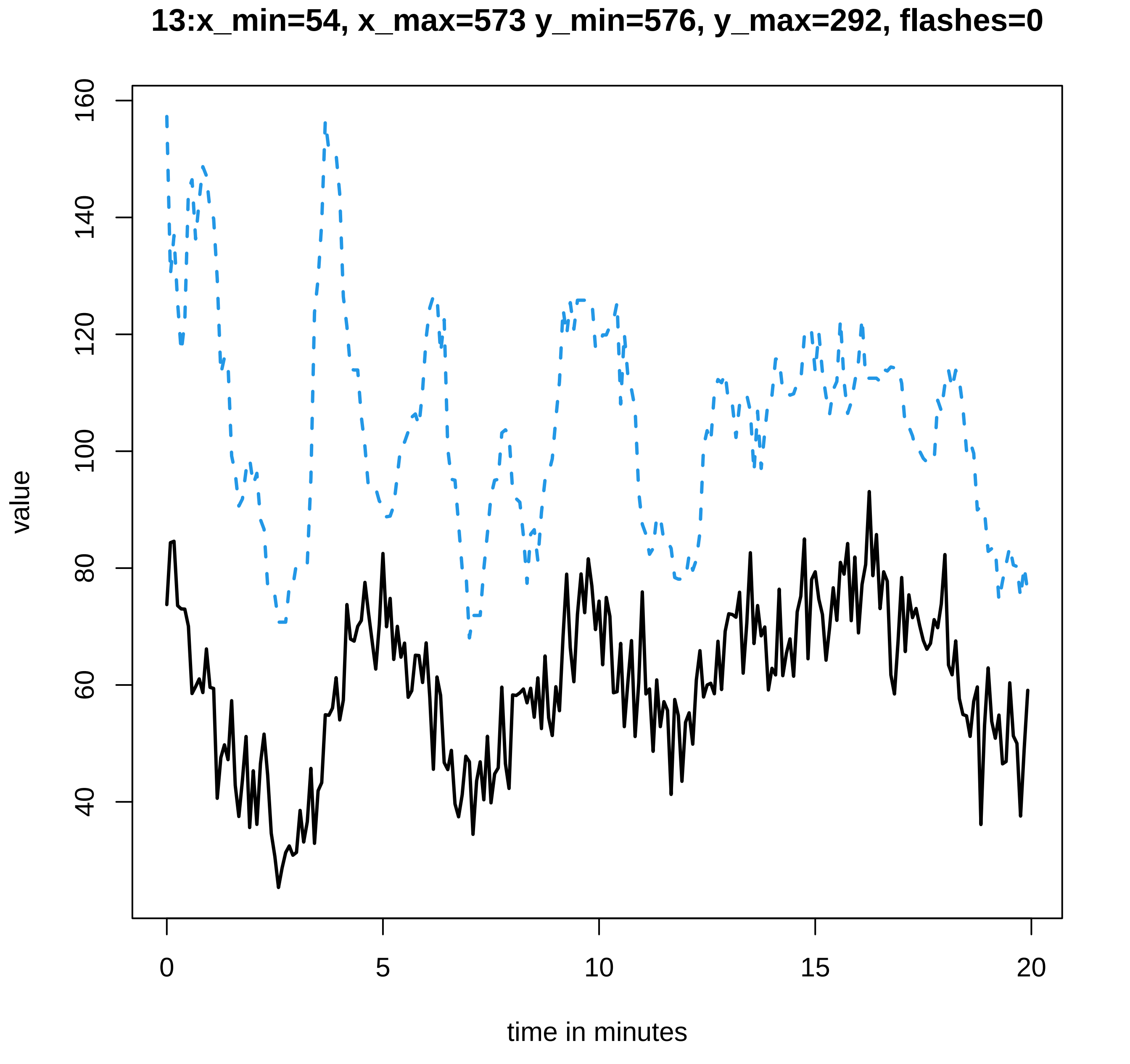
13:x_min=54, x_max=573 y_min=576, y_max=292, flashes=0
value
time in minutes
0
5
10
15
20
40
60
80
100
120
140
160

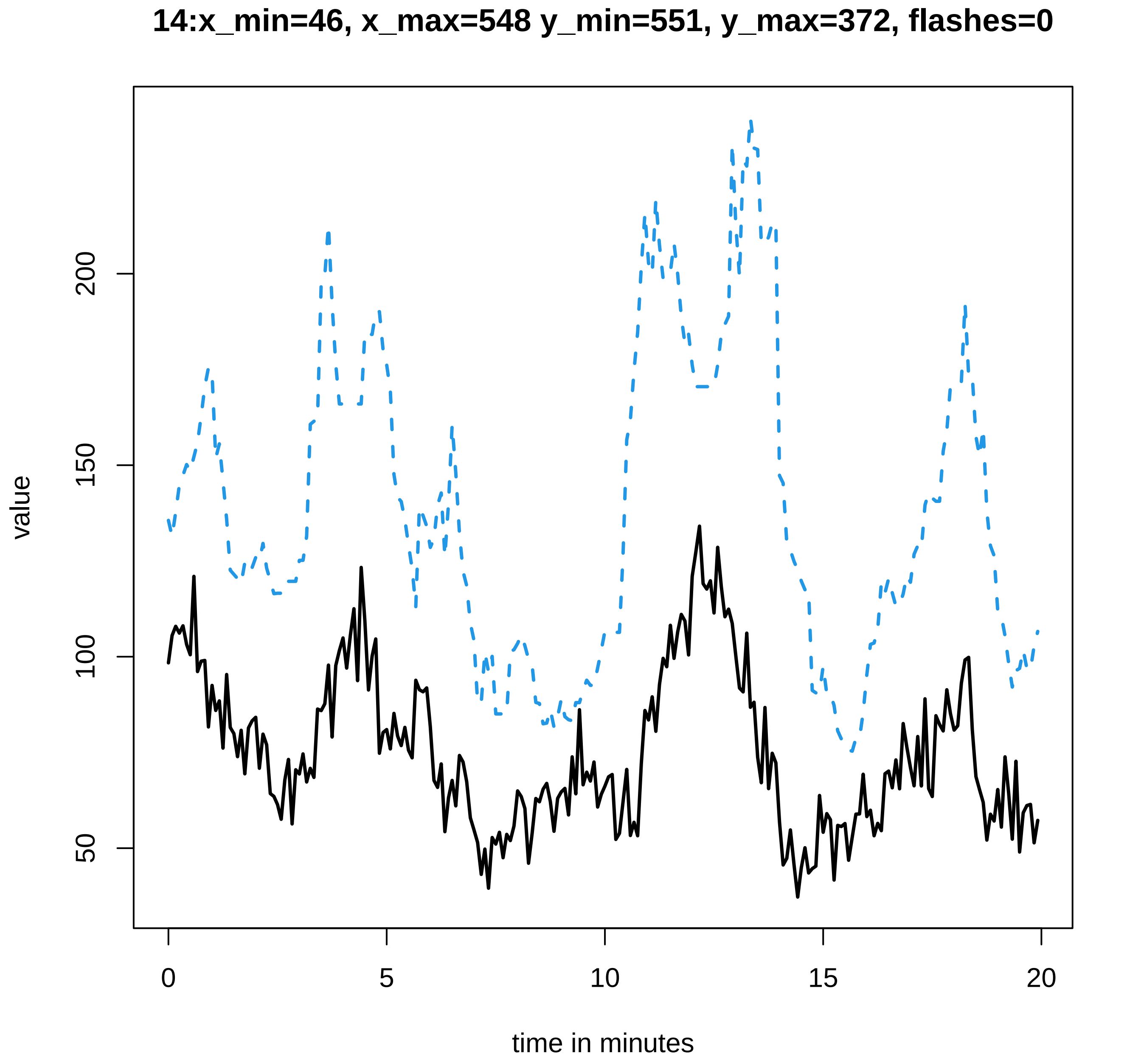
14:x_min=46, x_max=548 y_min=551, y_max=372, flashes=0
value
200
150
100
50
0
5
10
15
20
time in minutes

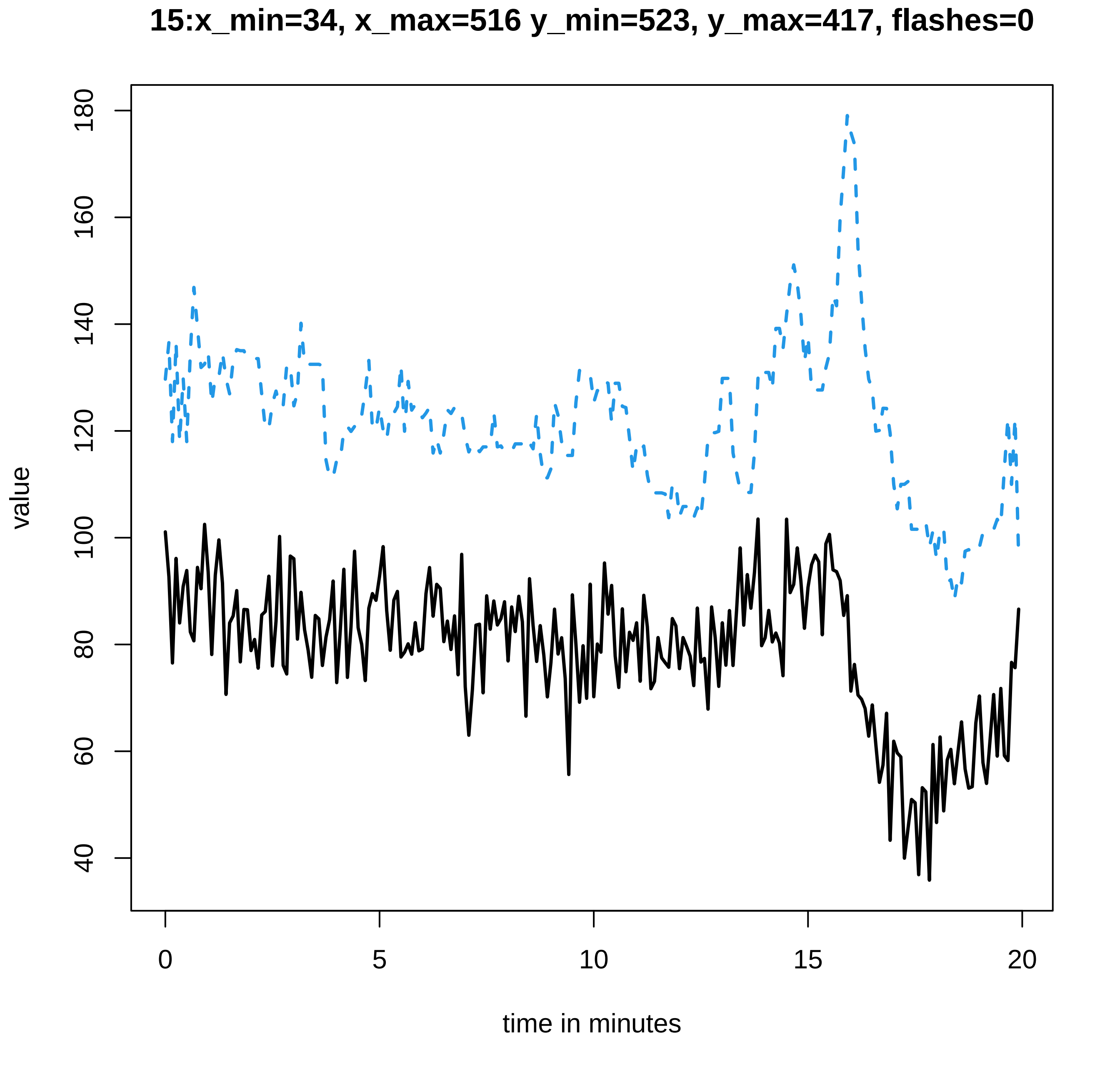
15:x_min=34, x_max=516 y_min=523, y_max=417, flashes=0
value
40
60
80
100
120
140
160
180
0
5
10
15
20
time in minutes

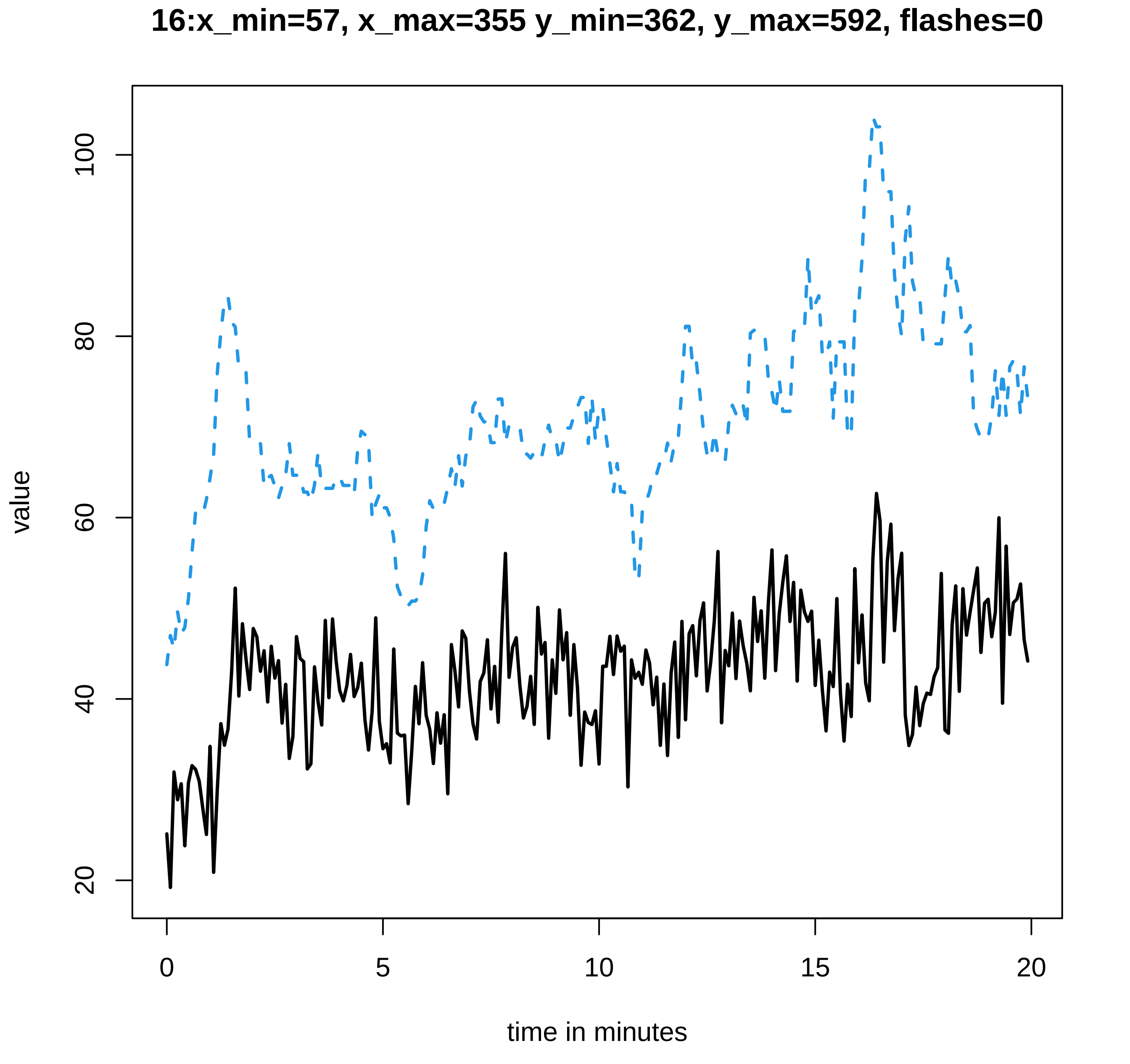
16:x_min=57, x_max=355 y_min=362, y_max=592, flashes=0
value
20
40
60
80
100
0
5
10
15
20
time in minutes

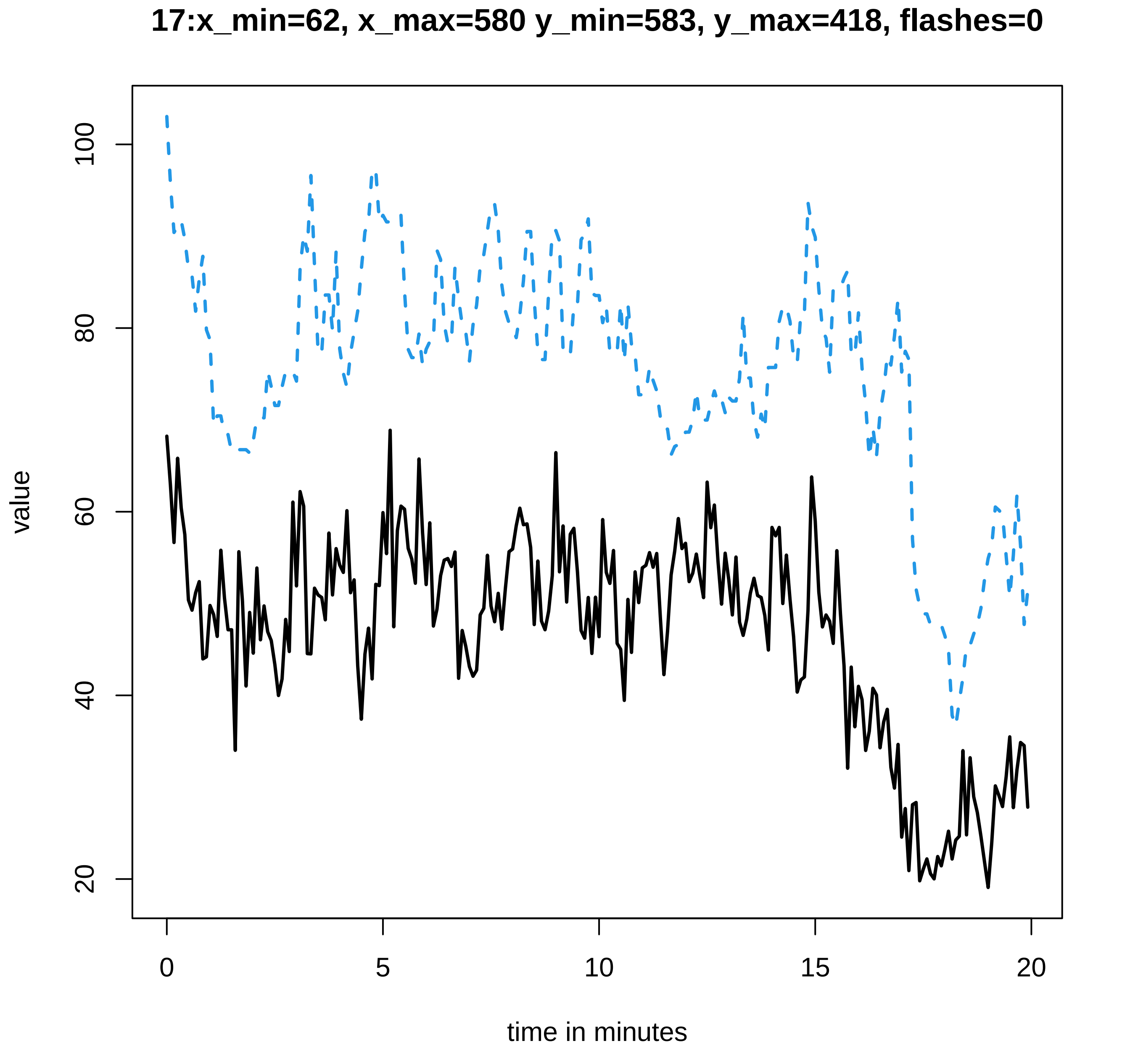
17:x_min=62, x_max=580 y_min=583, y_max=418, flashes=0
value
100
80
60
40
20
0
5
10
15
20
time in minutes

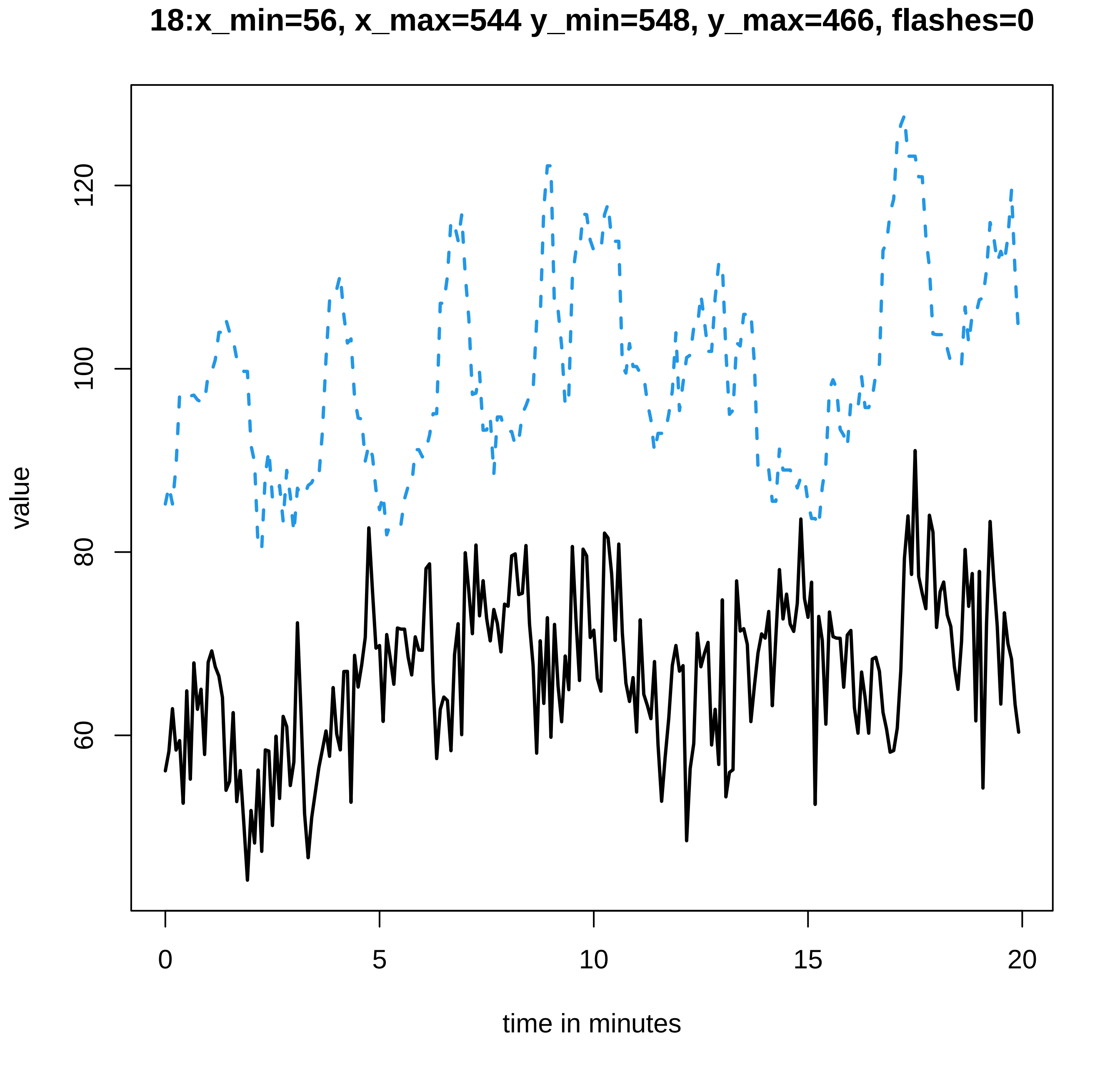
18:x_min=56, x_max=544 y_min=548, y_max=466, flashes=0
value
60
80
100
120
0
5
10
15
20
time in minutes

19:x_min=62, x_max=636 y_min=641, y_max=341, flashes=1

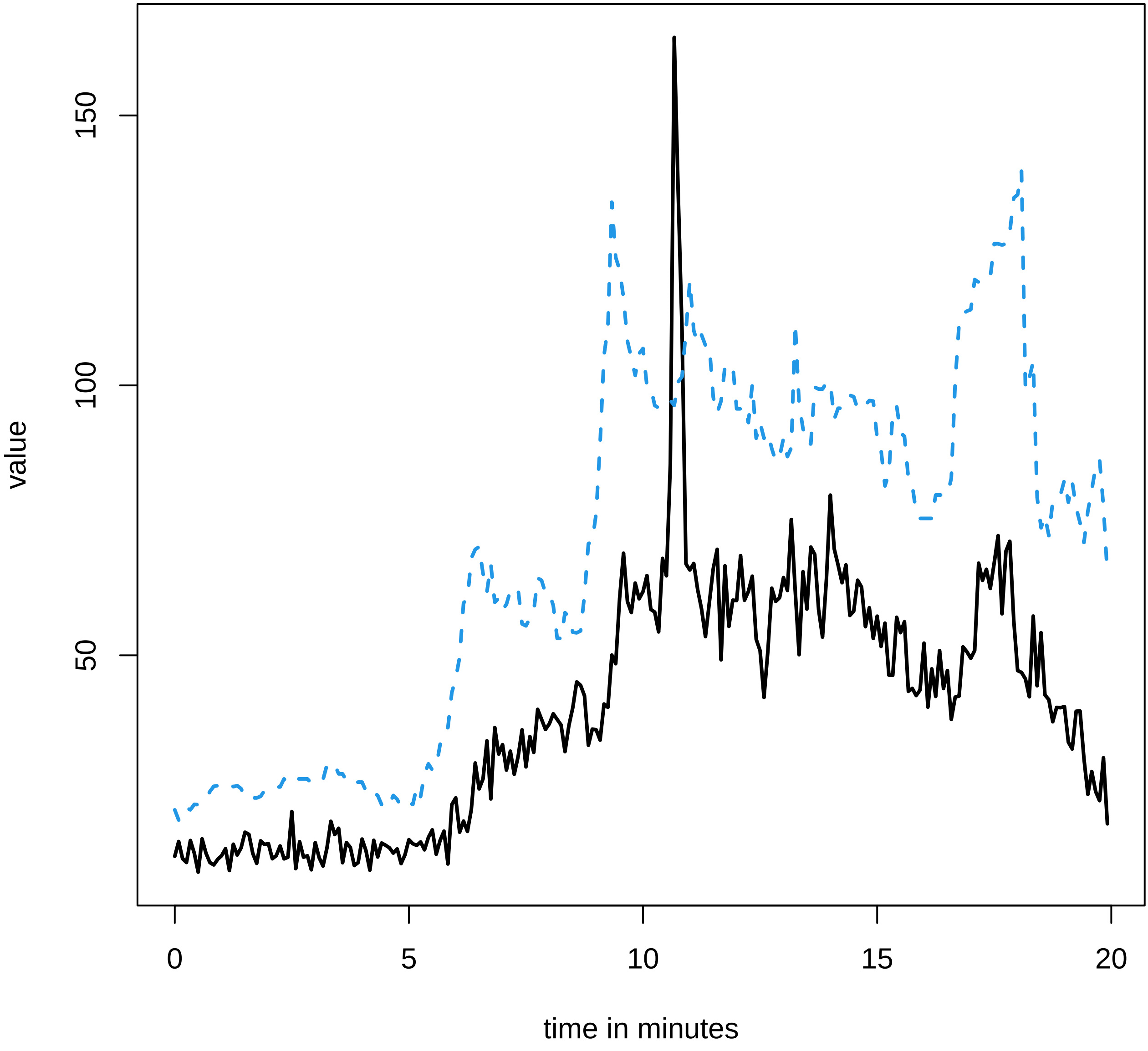

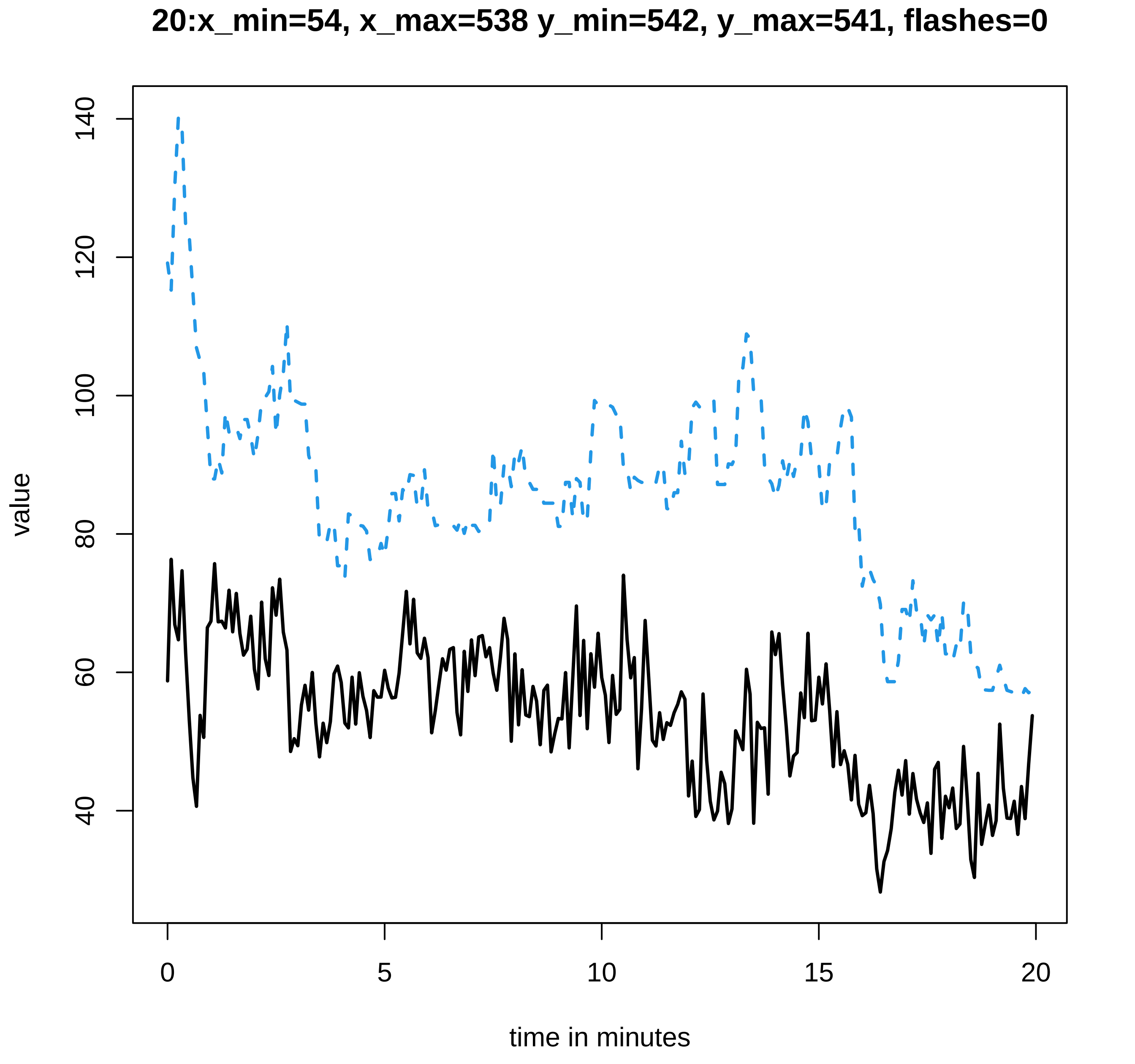
20:x_min=54, x_max=538 y_min=542, y_max=541, flashes=0
value
40
60
80
100
120
140
0
5
10
15
20
time in minutes

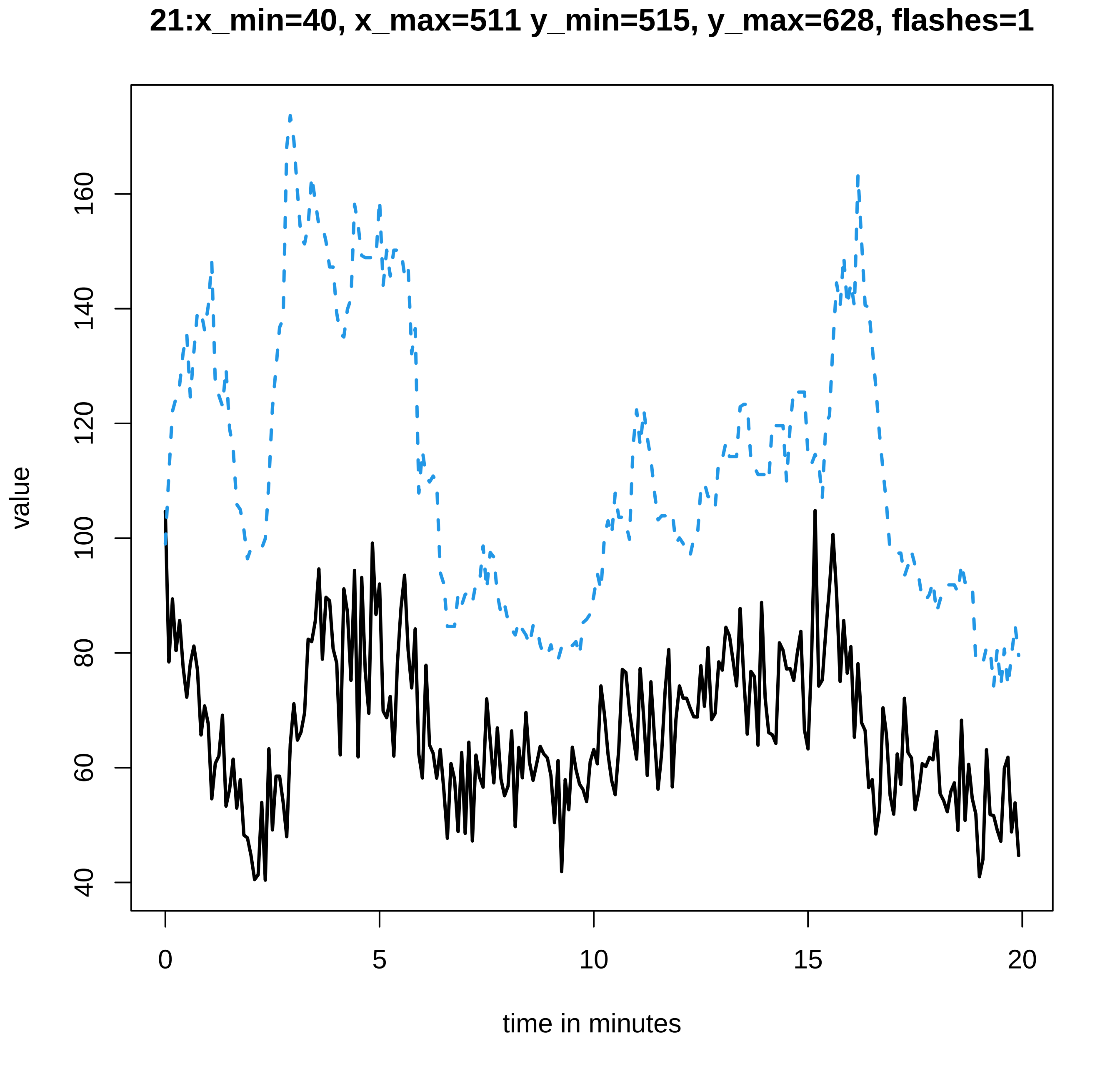
21:x_min=40, x_max=511 y_min=515, y_max=628, flashes=1
value
40
60
80
100
120
140
160
0
5
10
15
20
time in minutes

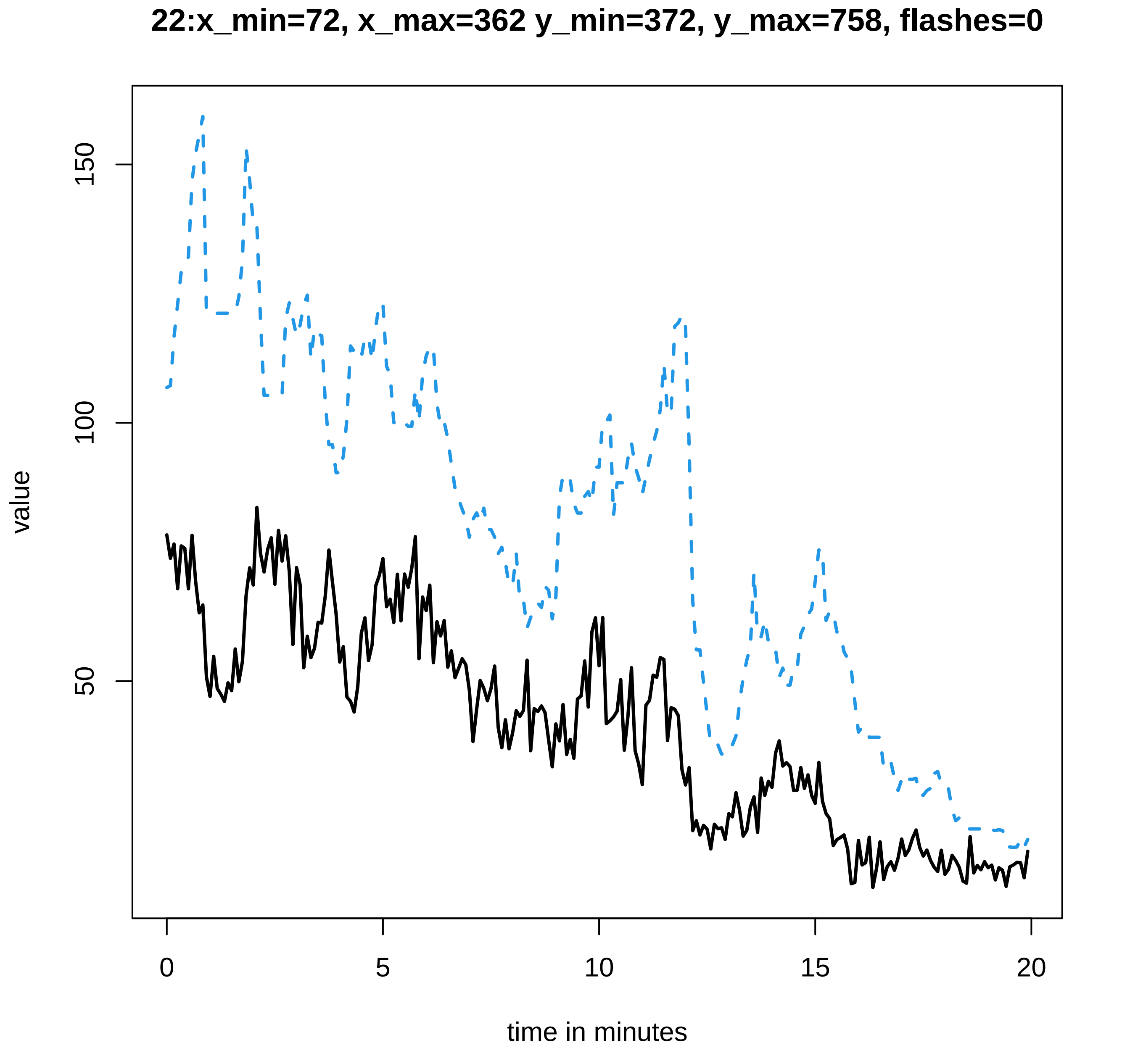

22:x_min=72, x_max=362 y_min=372, y_max=758, flashes=0
value
150
100
50
0
5
10
15
20
time in minutes

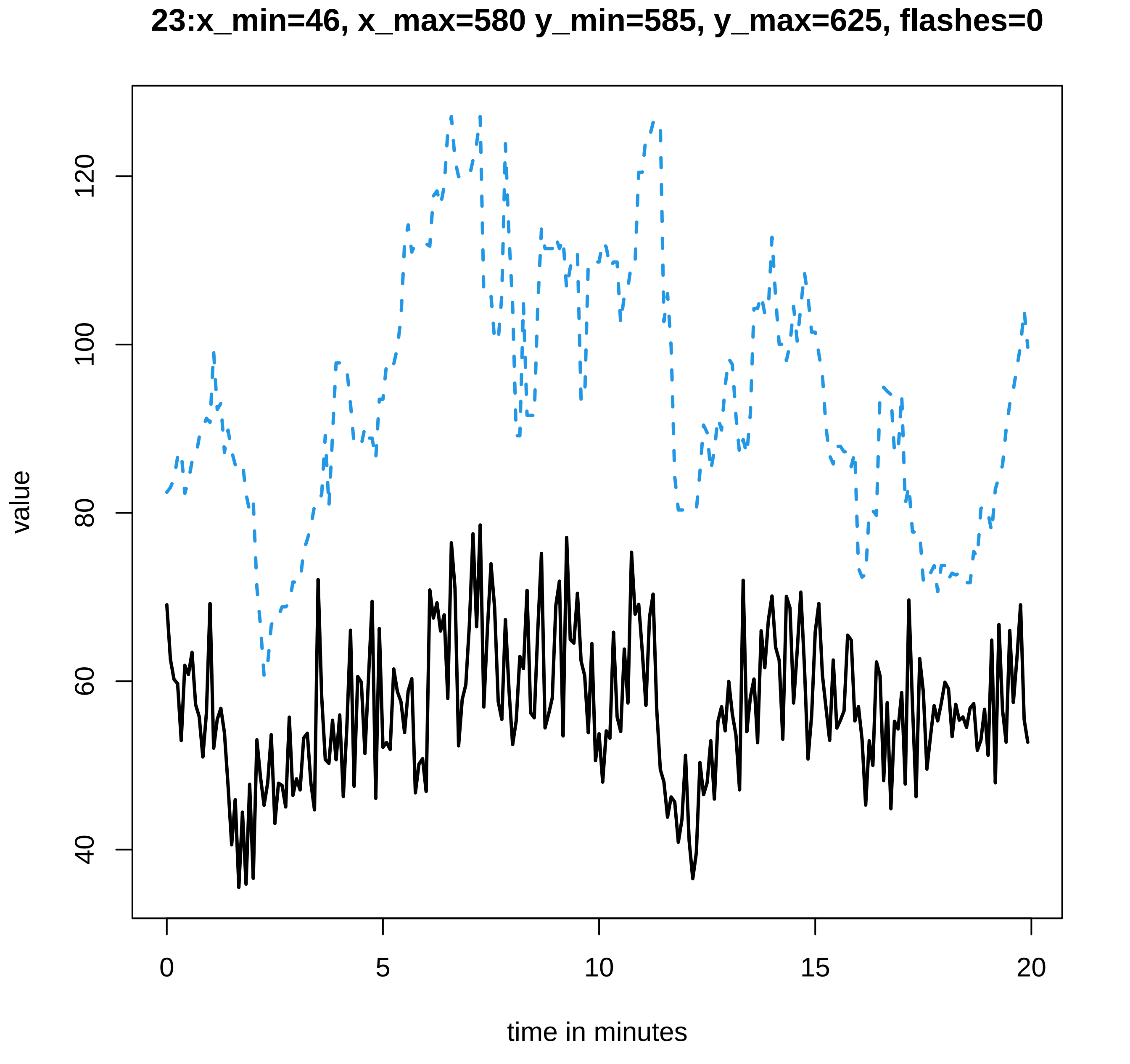
23:x_min=46, x_max=580 y_min=585, y_max=625, flashes=0
value
120
100
80
60
40
0
5
10
15
20
time in minutes

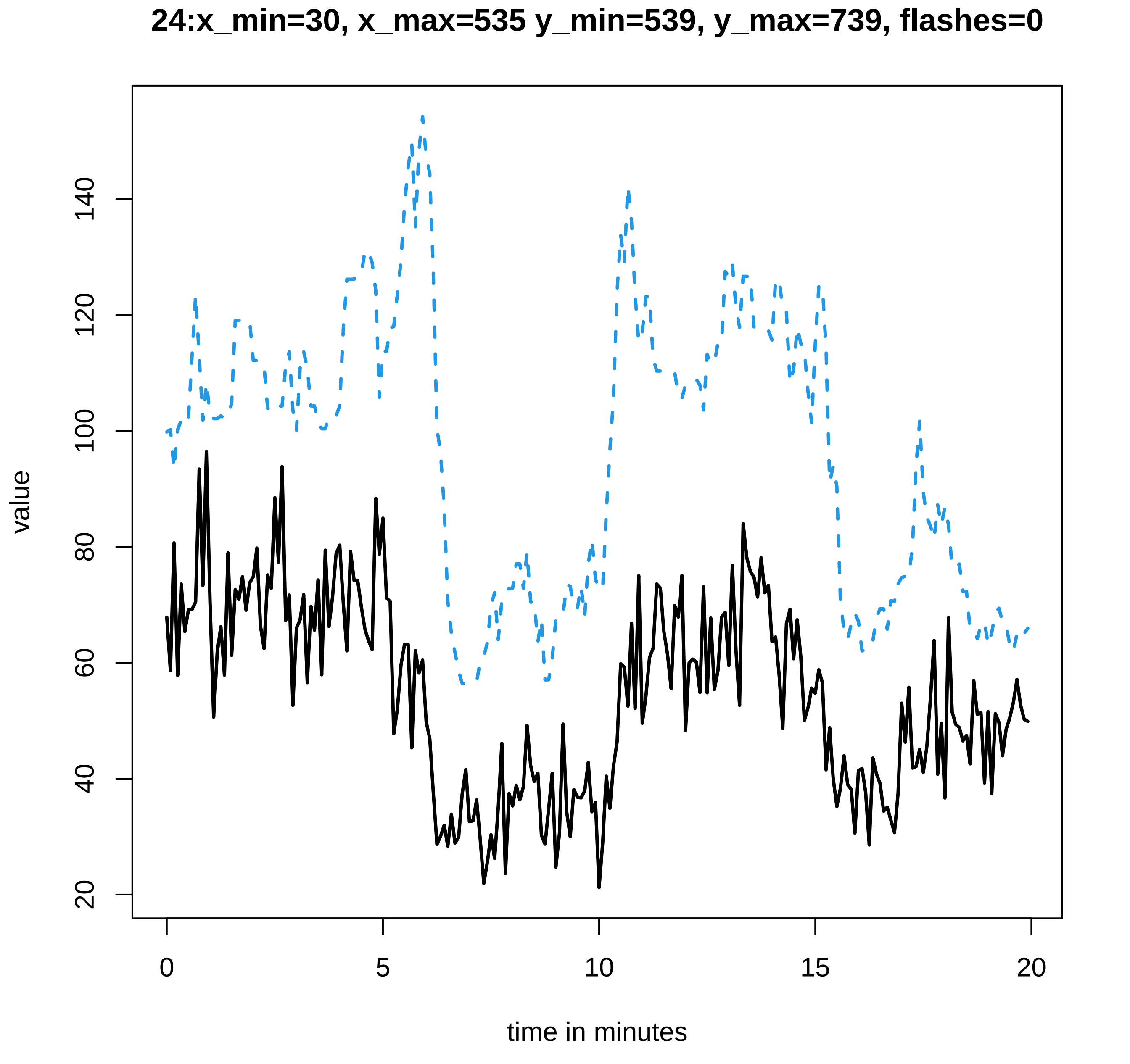
24:x_min=30, x_max=535 y_min=539, y_max=739, flashes=0
value
20
40
60
80
100
120
140
0
5
10
15
20
time in minutes

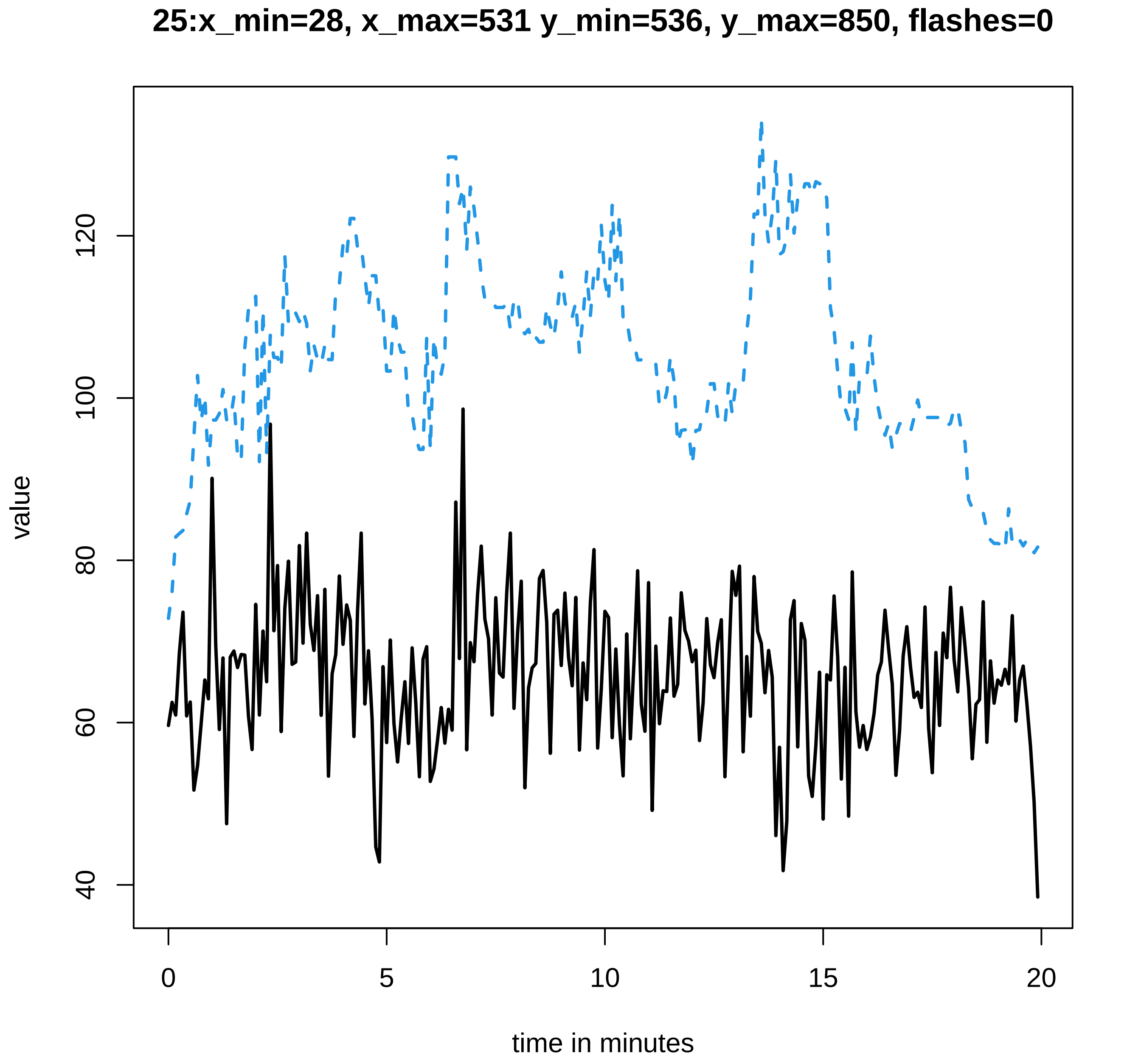
25:x_min=28, x_max=531 y_min=536, y_max=850, flashes=0
value
40
60
80
100
120
0
5
10
15
20
time in minutes

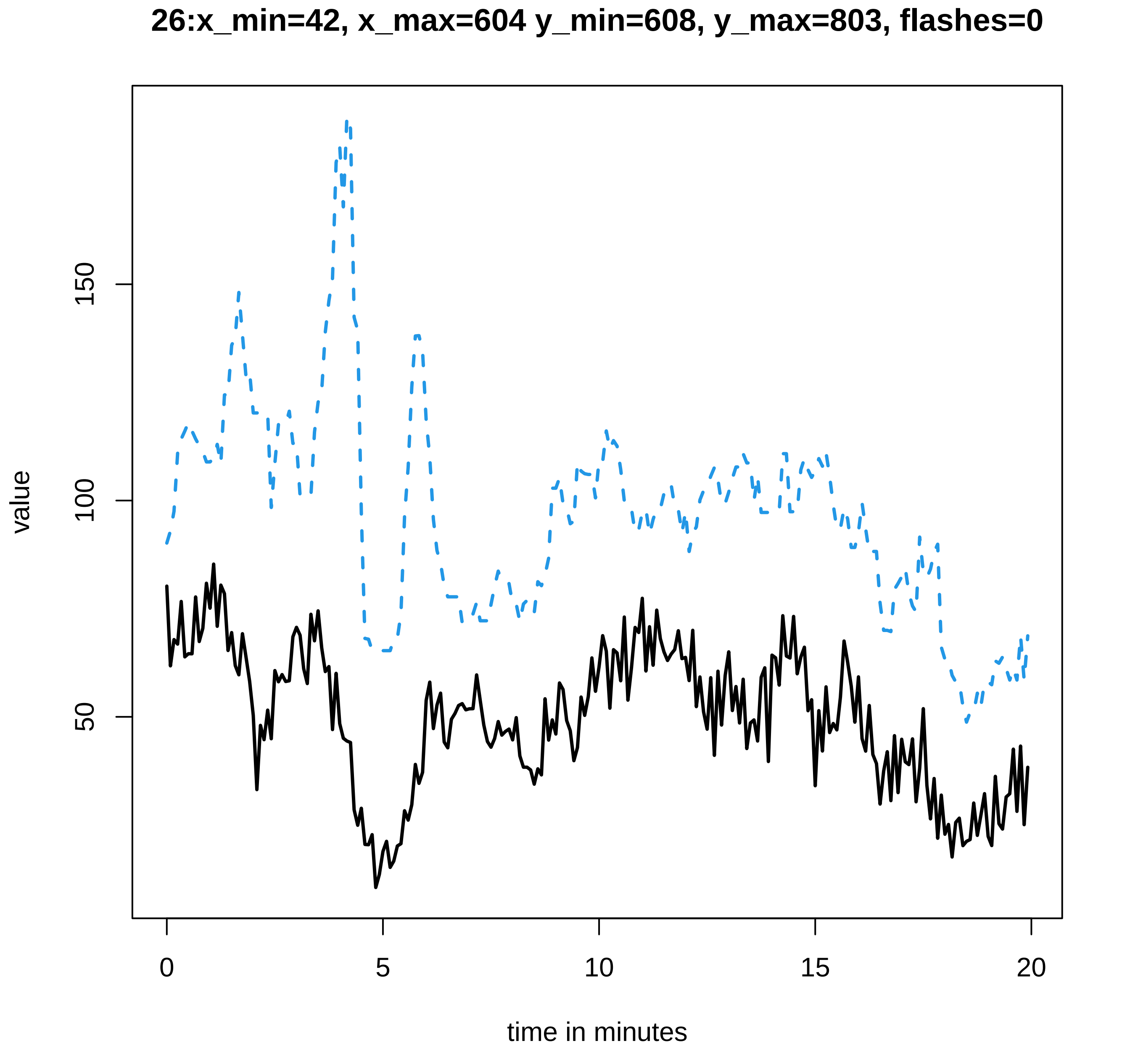
26:x_min=42, x_max=604 y_min=608, y_max=803, flashes=0
value
50
100
150
0
5
10
15
20
time in minutes

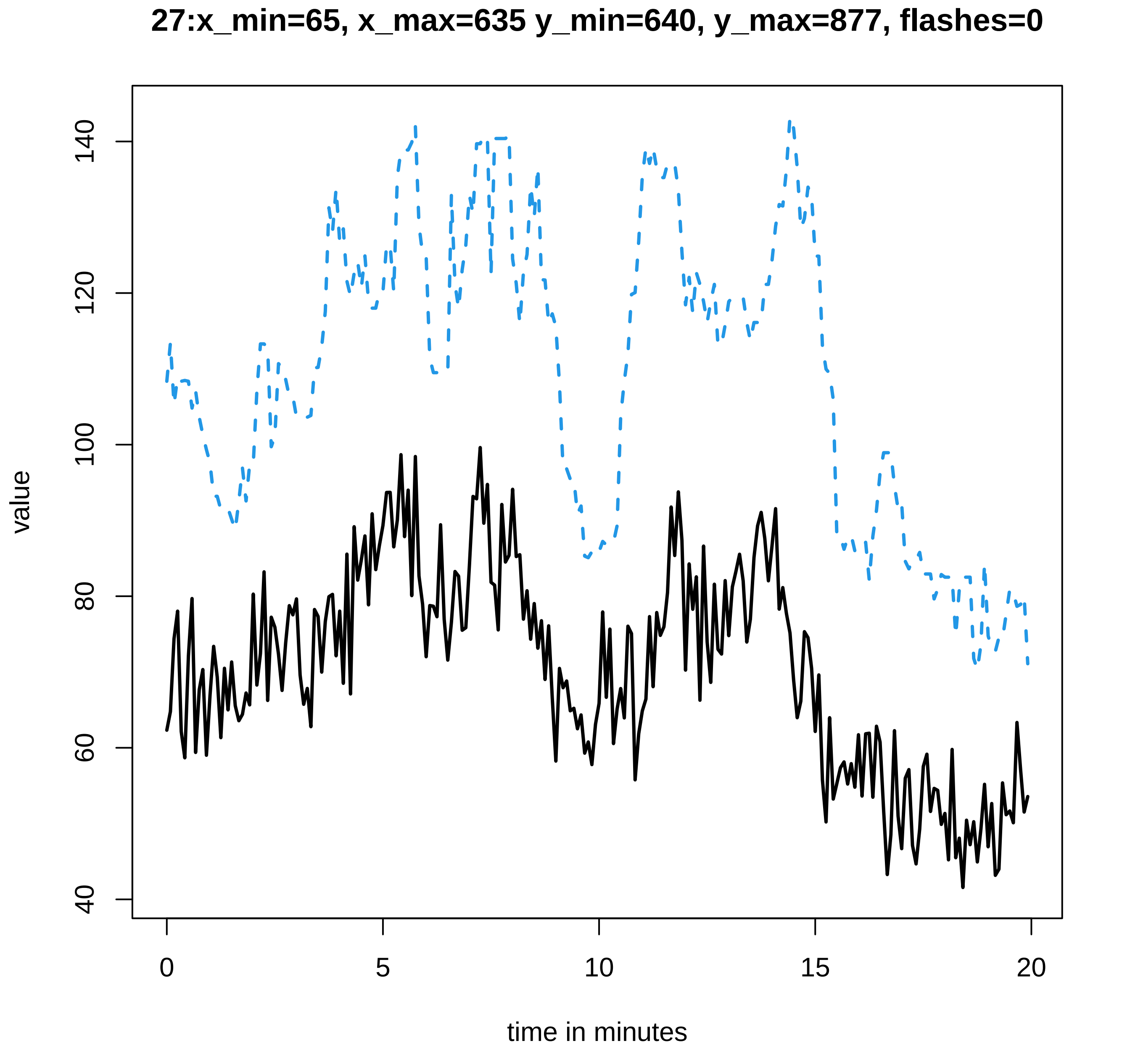

27:x_min=65, x_max=635 y_min=640, y_max=877, flashes=0
value
140
120
100
80
60
40
0
5
10
15
20
time in minutes
