## Supplementary data Calcium spiking for "LysM Receptor Proteins are Required for Ectomycorrhizal Symbiosis in Poplar": profiles-spiking-13.pdf

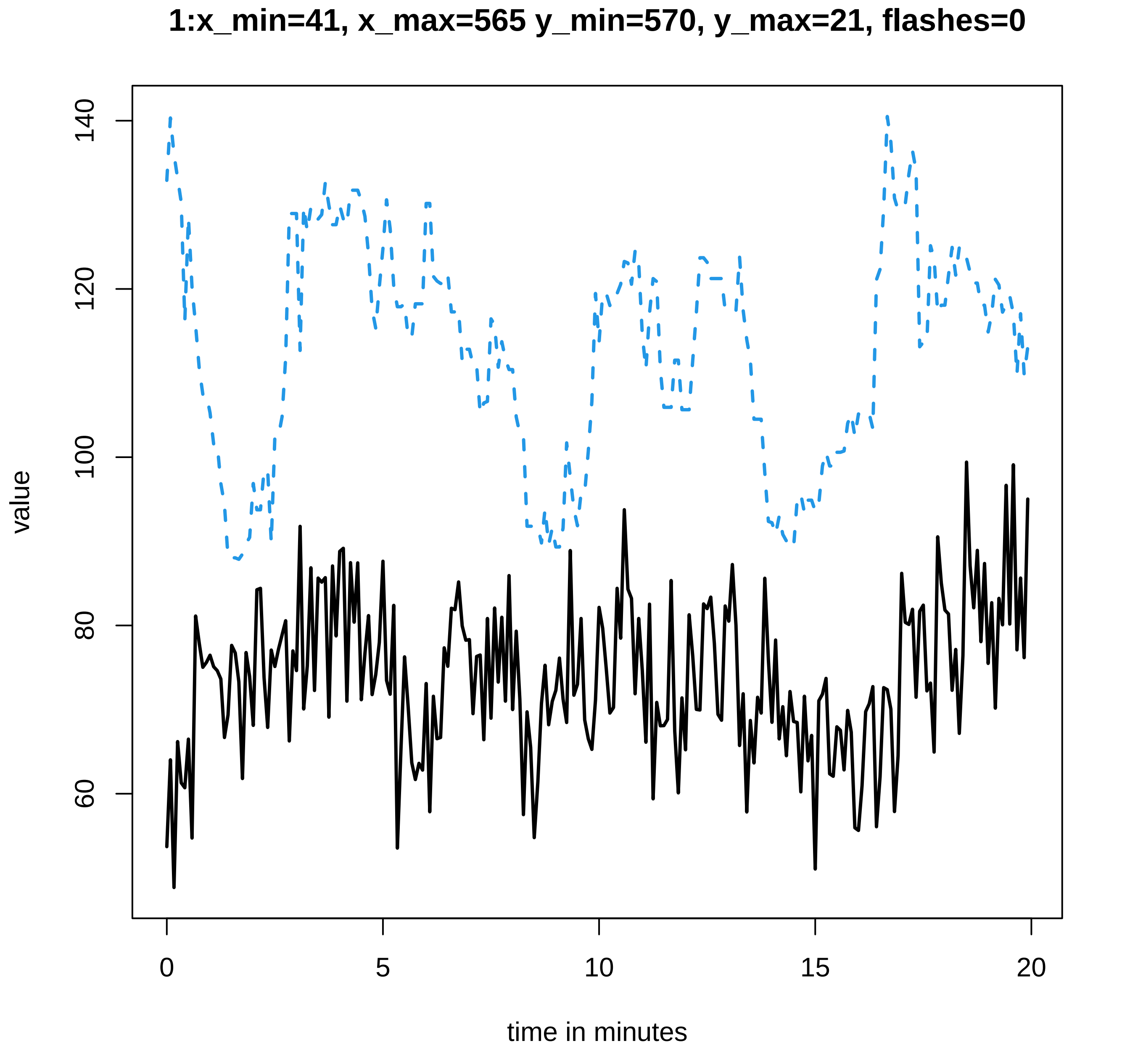
1:x_min=41, x_max=565 y_min=570, y_max=21, flashes=0
value
140
120
100
80
60
0
5
10
15
20
time in minutes

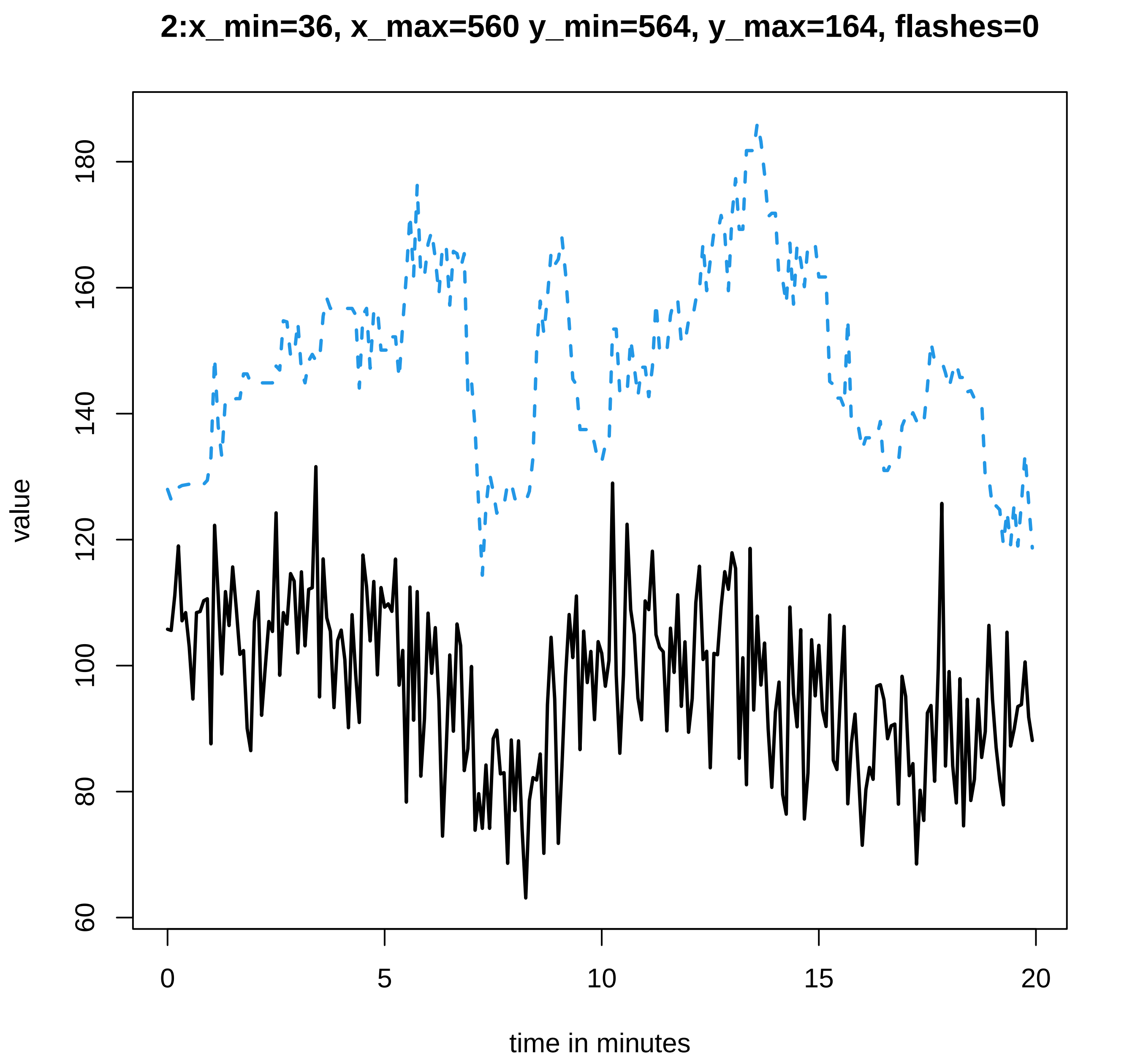
2:x_min=36, x_max=560 y_min=564, y_max=164, flashes=0
value
60
80
100
120
140
160
180
0
5
10
15
20
time in minutes

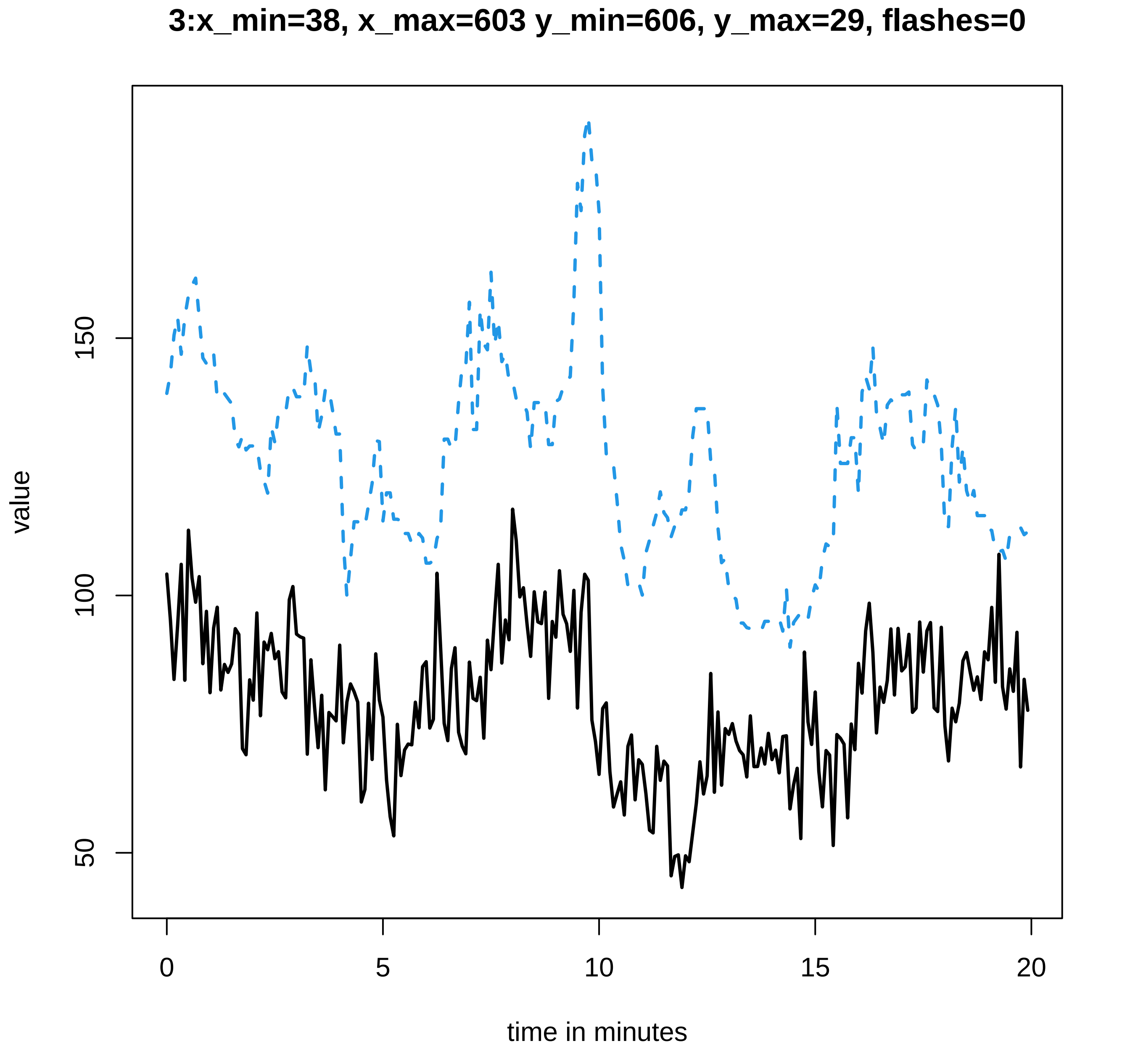

3:x_min=38, x_max=603 y_min=606, y_max=29, flashes=0
value
150
100
50
0
5
10
15
20
time in minutes

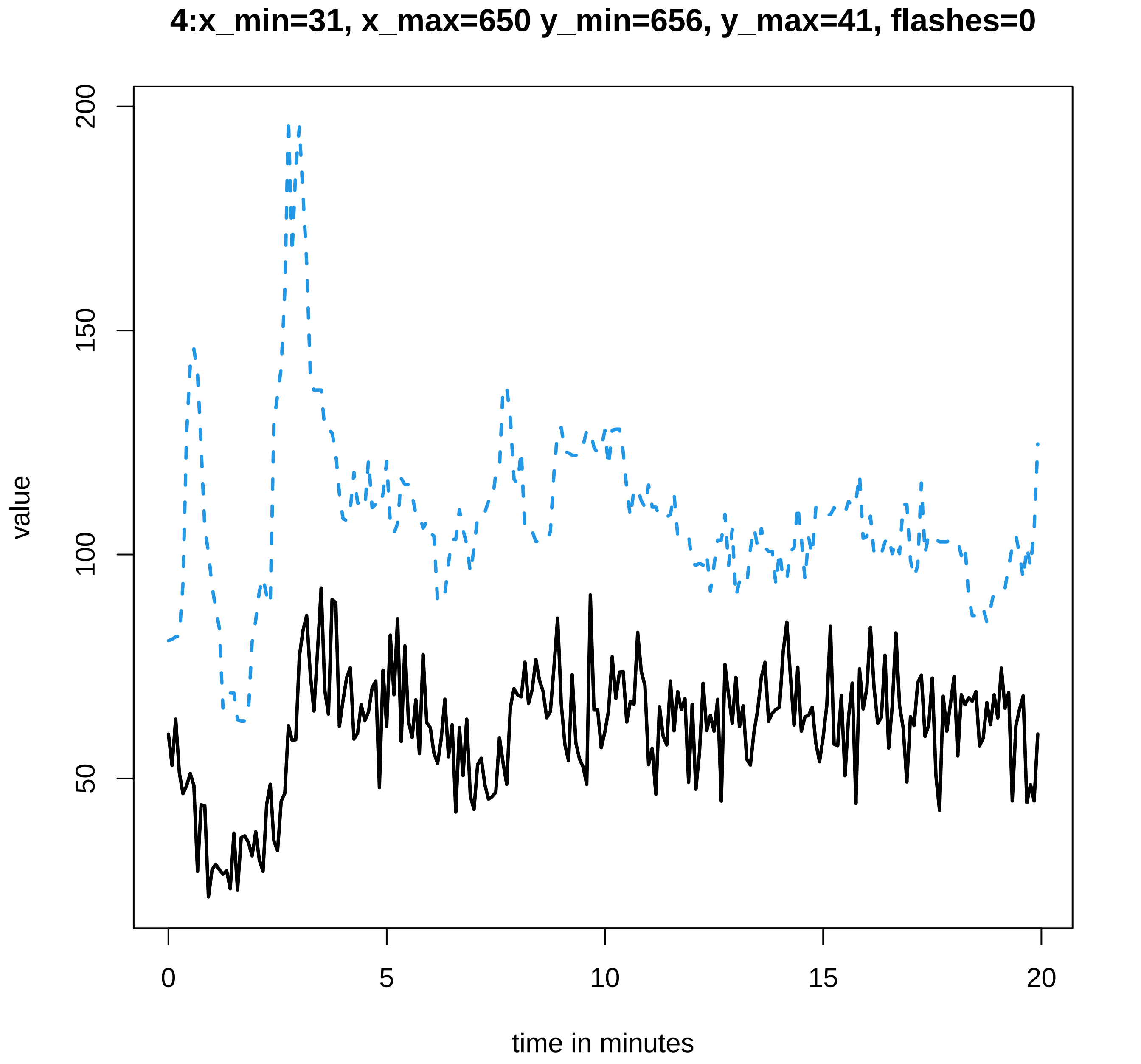
4:x_min=31, x_max=650 y_min=656, y_max=41, flashes=0
value
200
150
100
50
0
5
10
15
20
time in minutes

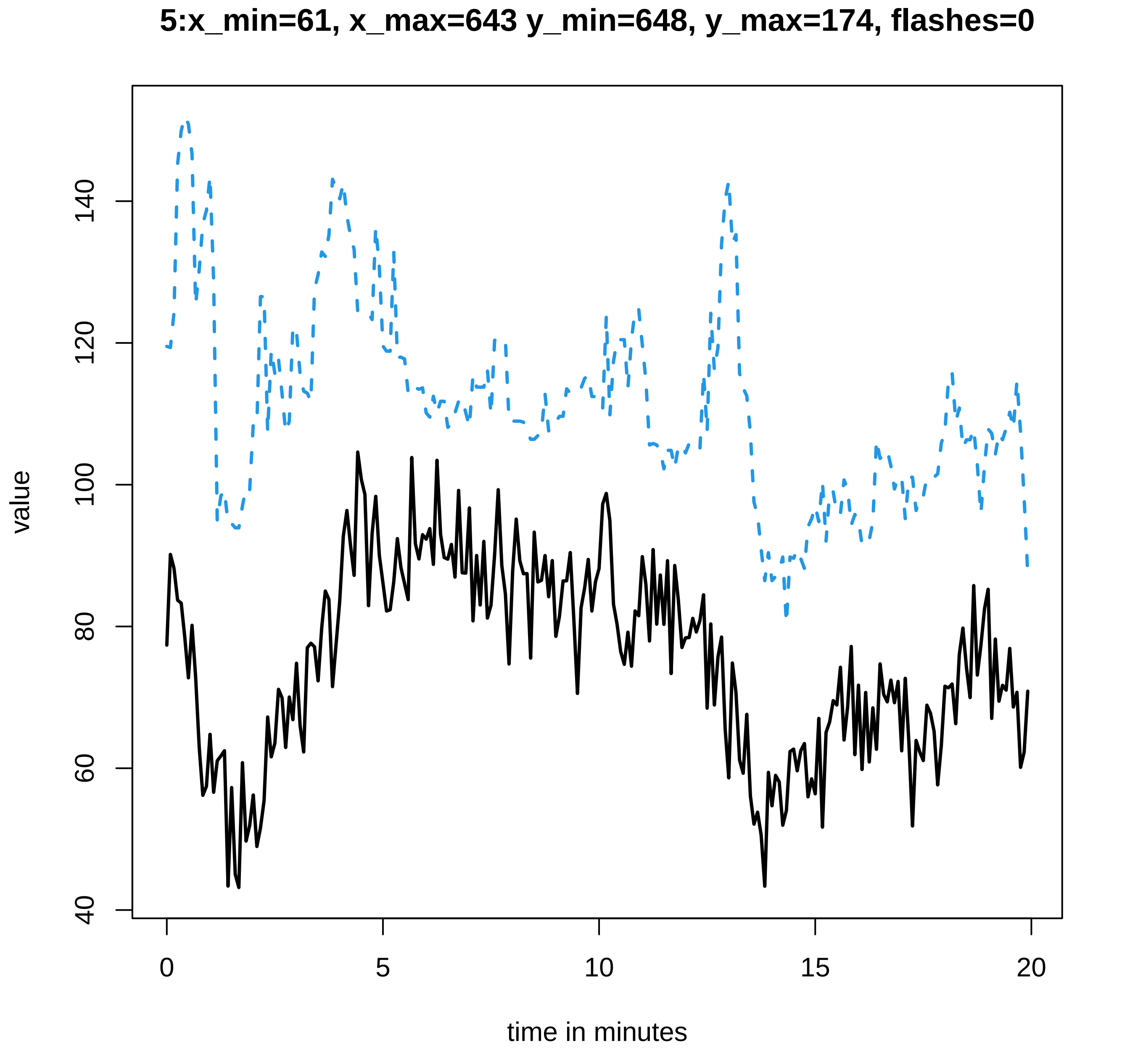
5:x_min=61, x_max=643 y_min=648, y_max=174, flashes=0
value
140
120
100
80
60
40
0
5
10
15
20
time in minutes

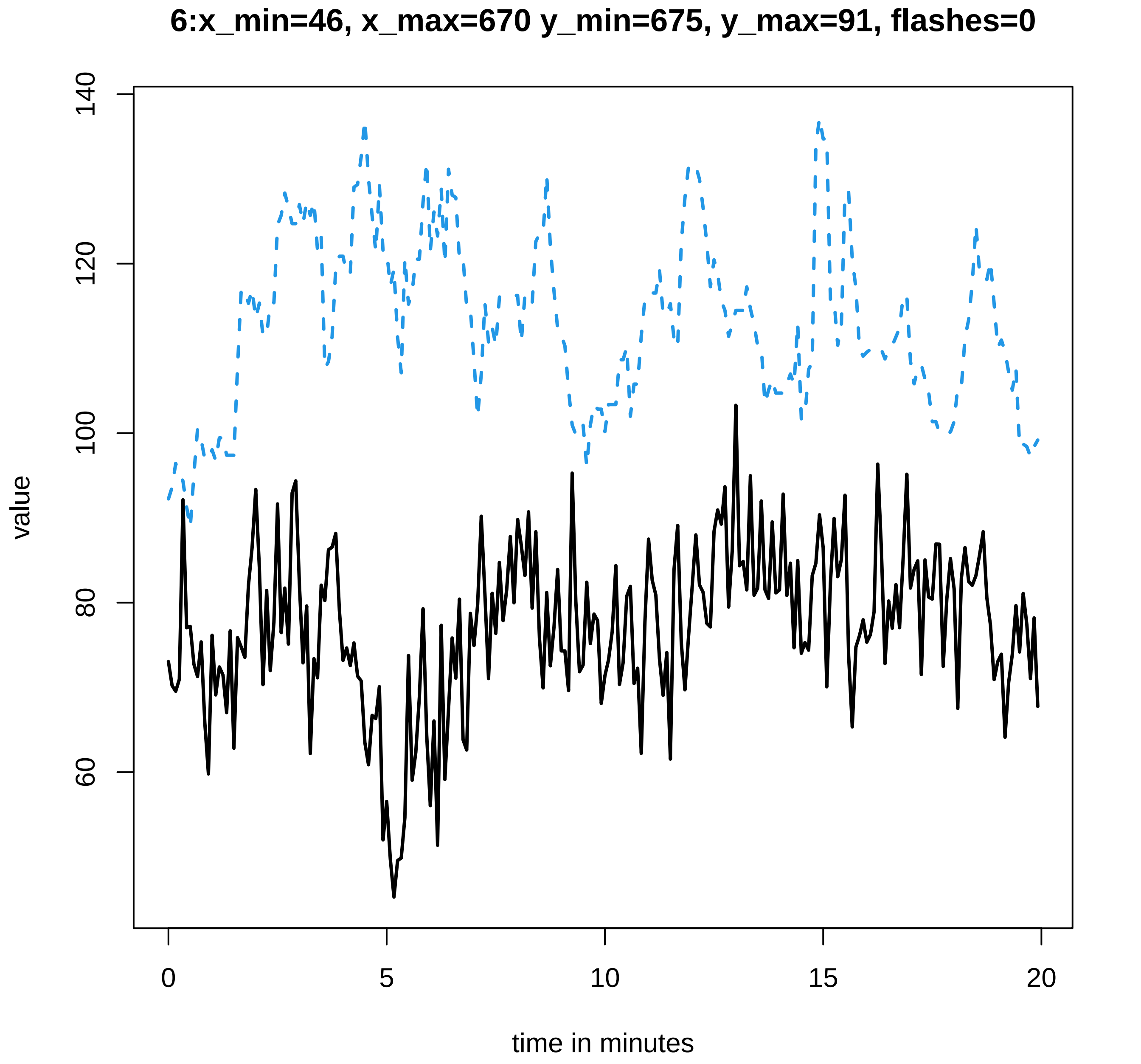
6:x_min=46, x_max=670 y_min=675, y_max=91, flashes=0
140
120
100
80
60
value
0
5
10
15
20
time in minutes

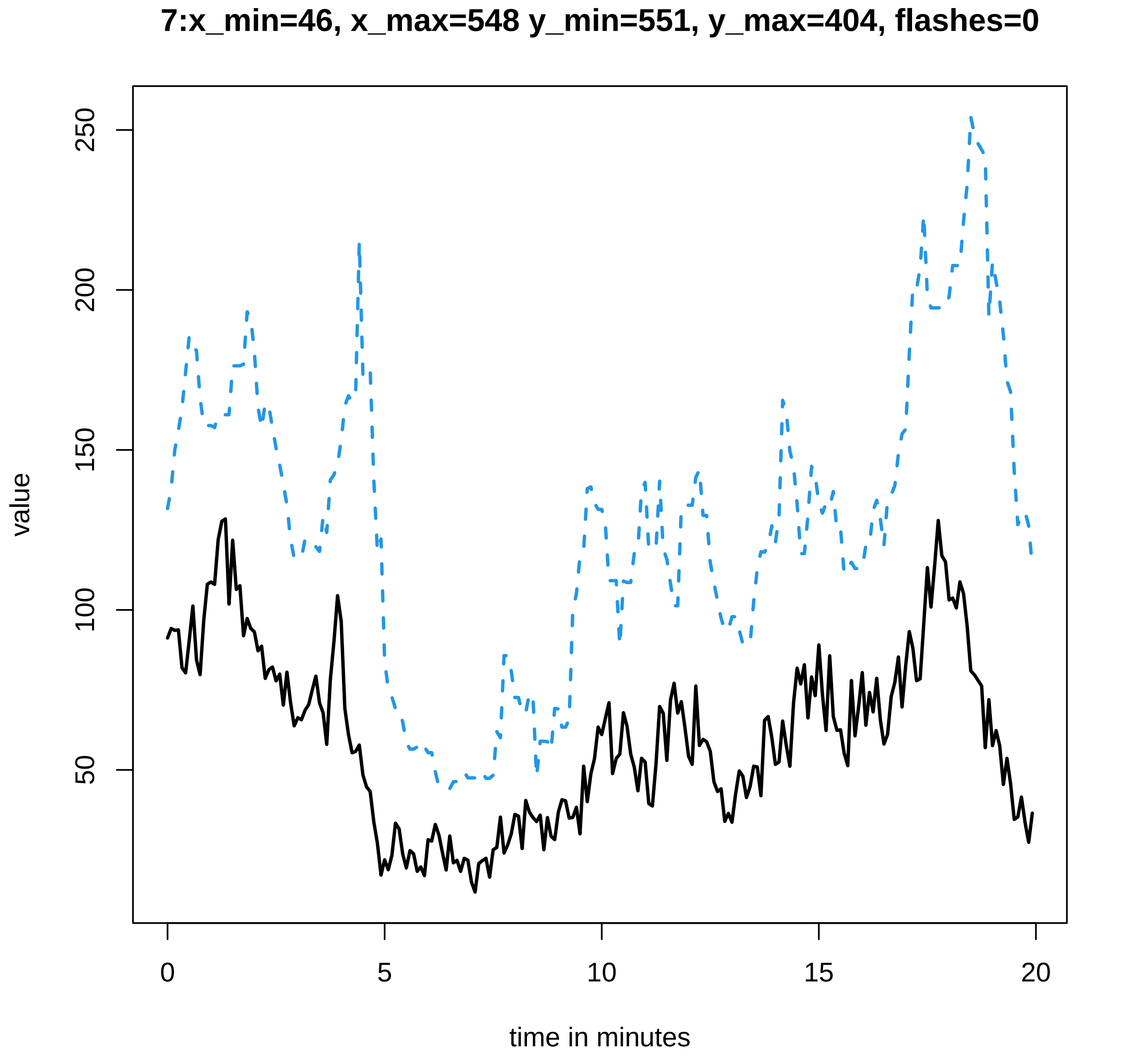
7:x_min=46, x_max=548 y_min=551, y_max=404, flashes=0
value
50
100
150
200
250
0
5
10
15
20
time in minutes

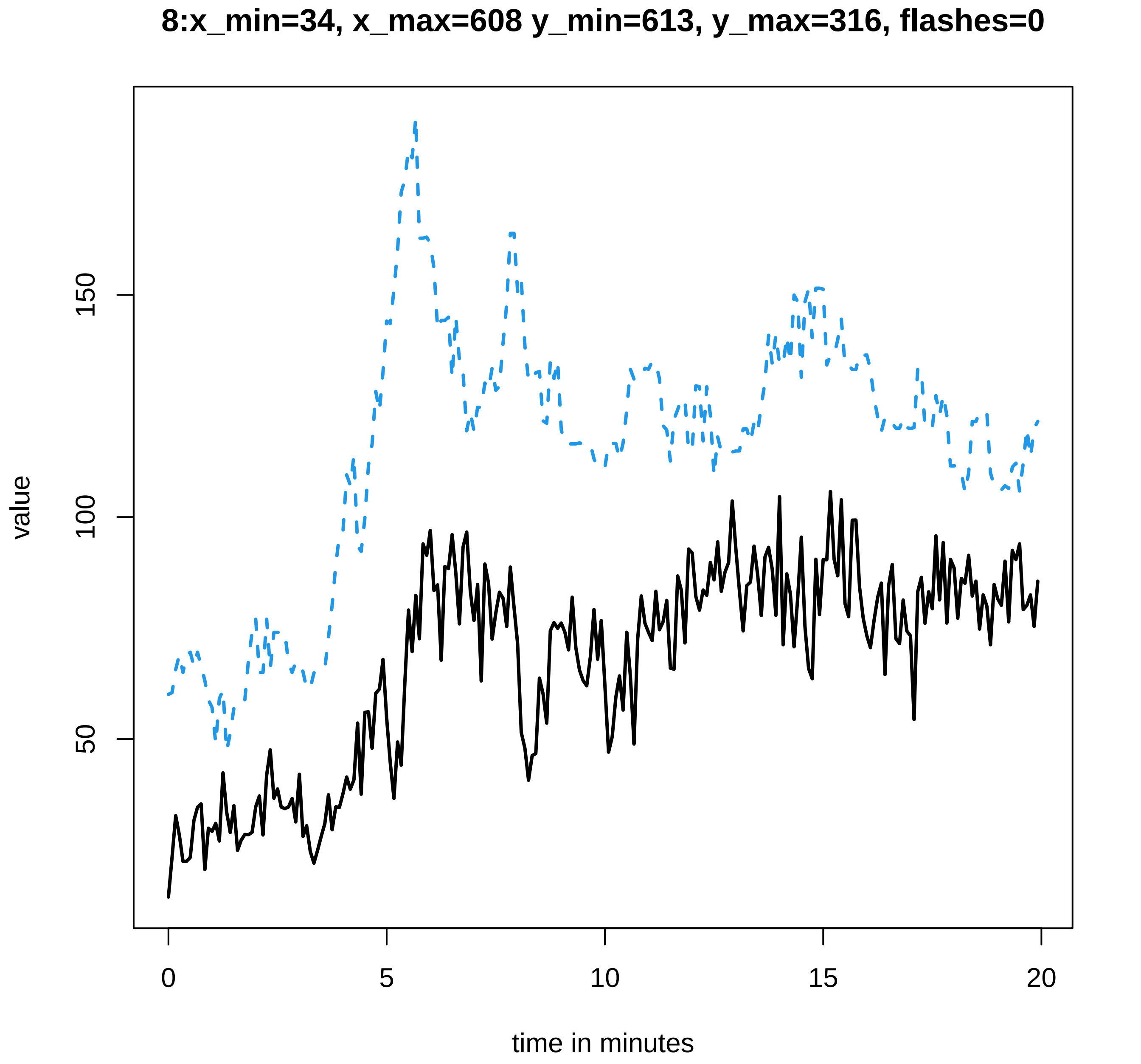
8:x_min=34, x_max=608 y_min=613, y_max=316, flashes=0
value
50
100
150
0
5
10
15
20
time in minutes

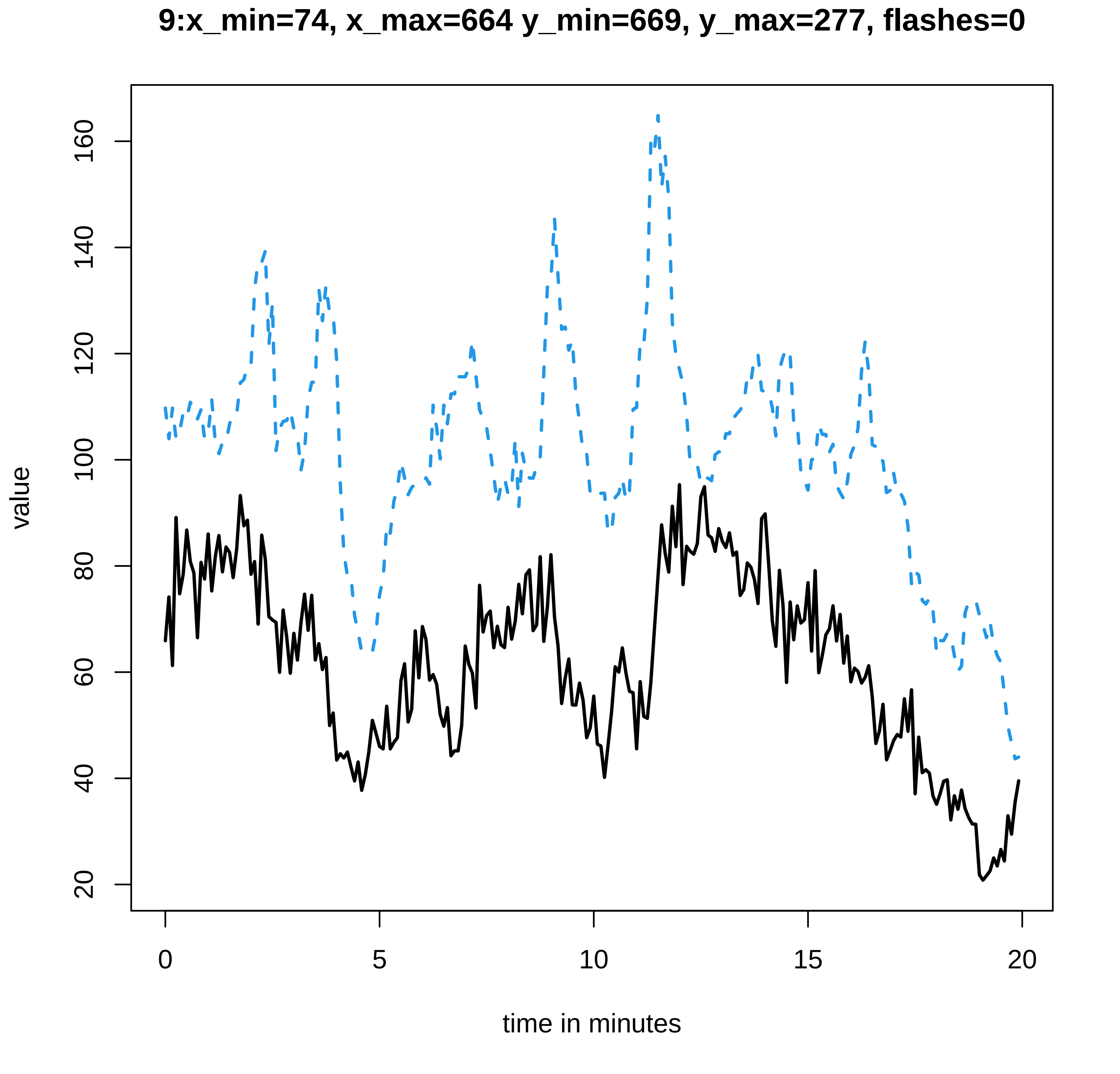
9:x_min=74, x_max=664 y_min=669, y_max=277, flashes=0
value
20
40
60
80
100
120
140
160
0
5
10
15
20
time in minutes

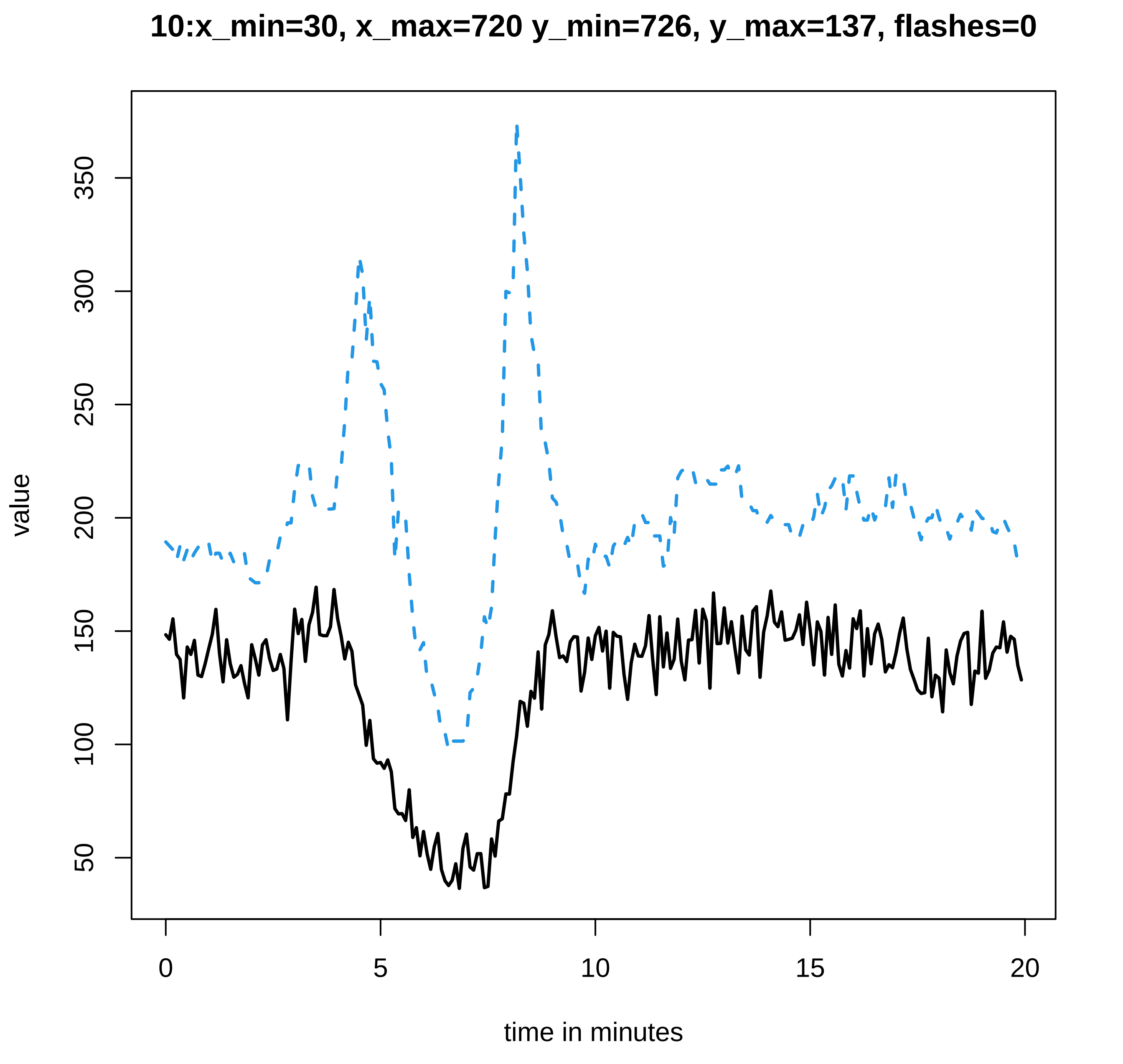

10:x_min=30, x_max=720 y_min=726, y_max=137, flashes=0
value
50
100
150
200
250
300
350
0
5
10
15
20
time in minutes

#### 11:x_min=48, x_max=524 y_min=528, y_max=531, flashes=0

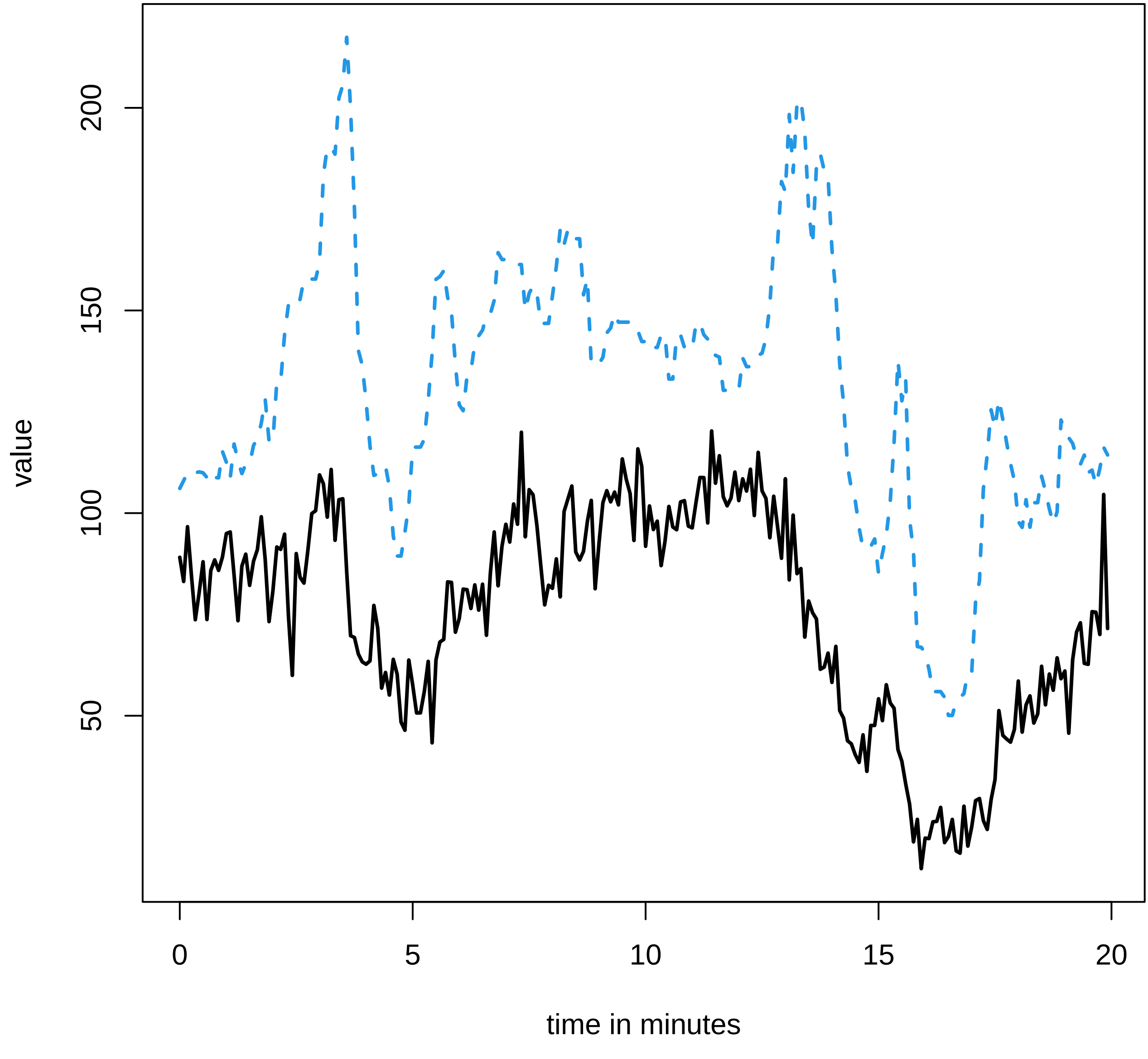

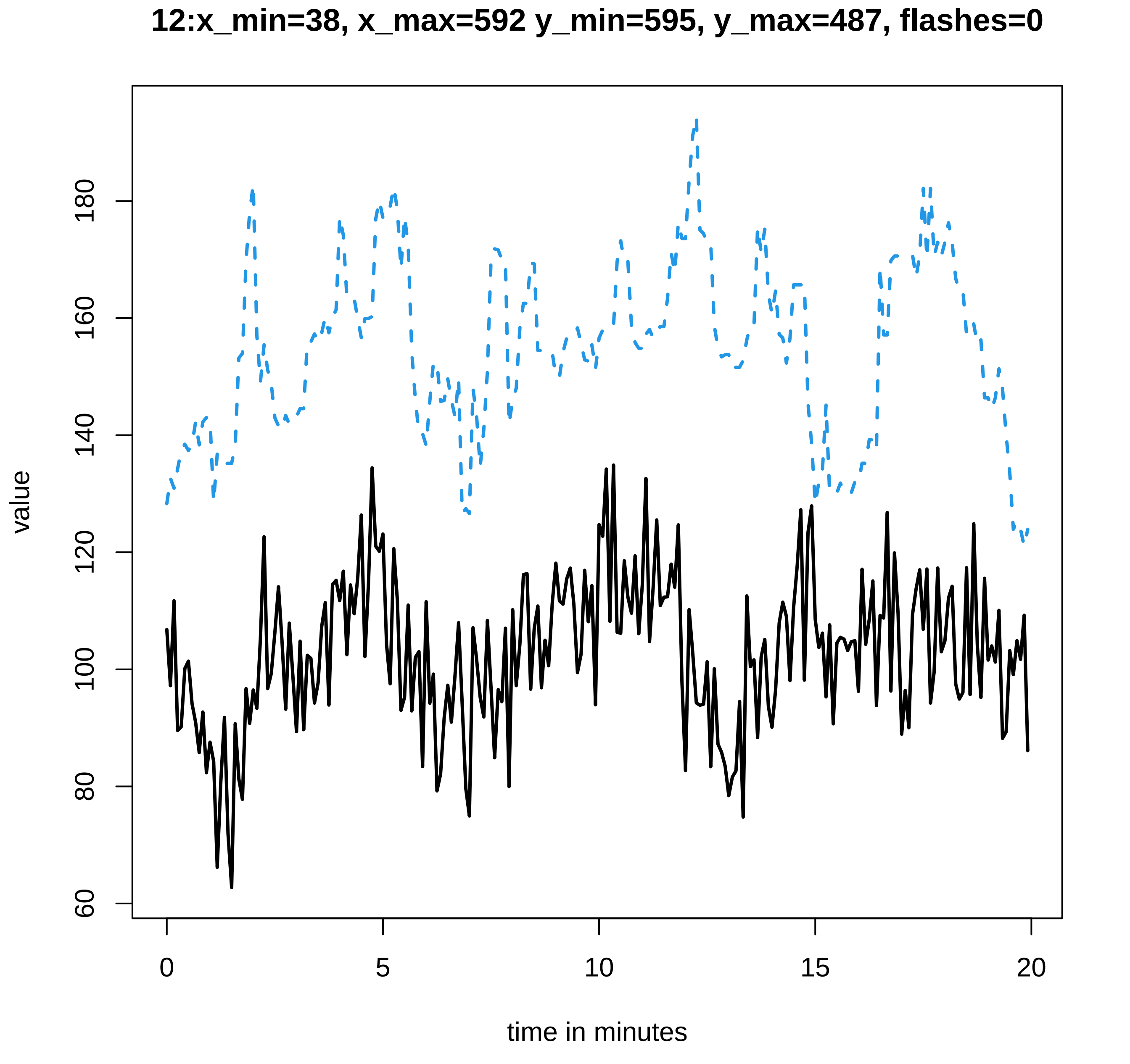

12:x_min=38, x_max=592 y_min=595, y_max=487, flashes=0
value
time in minutes
60
80
100
120
140
160
180
0
5
10
15
20

**13:x_min=42, x_max=644 y_min=650, y_max=501, flashes=0**

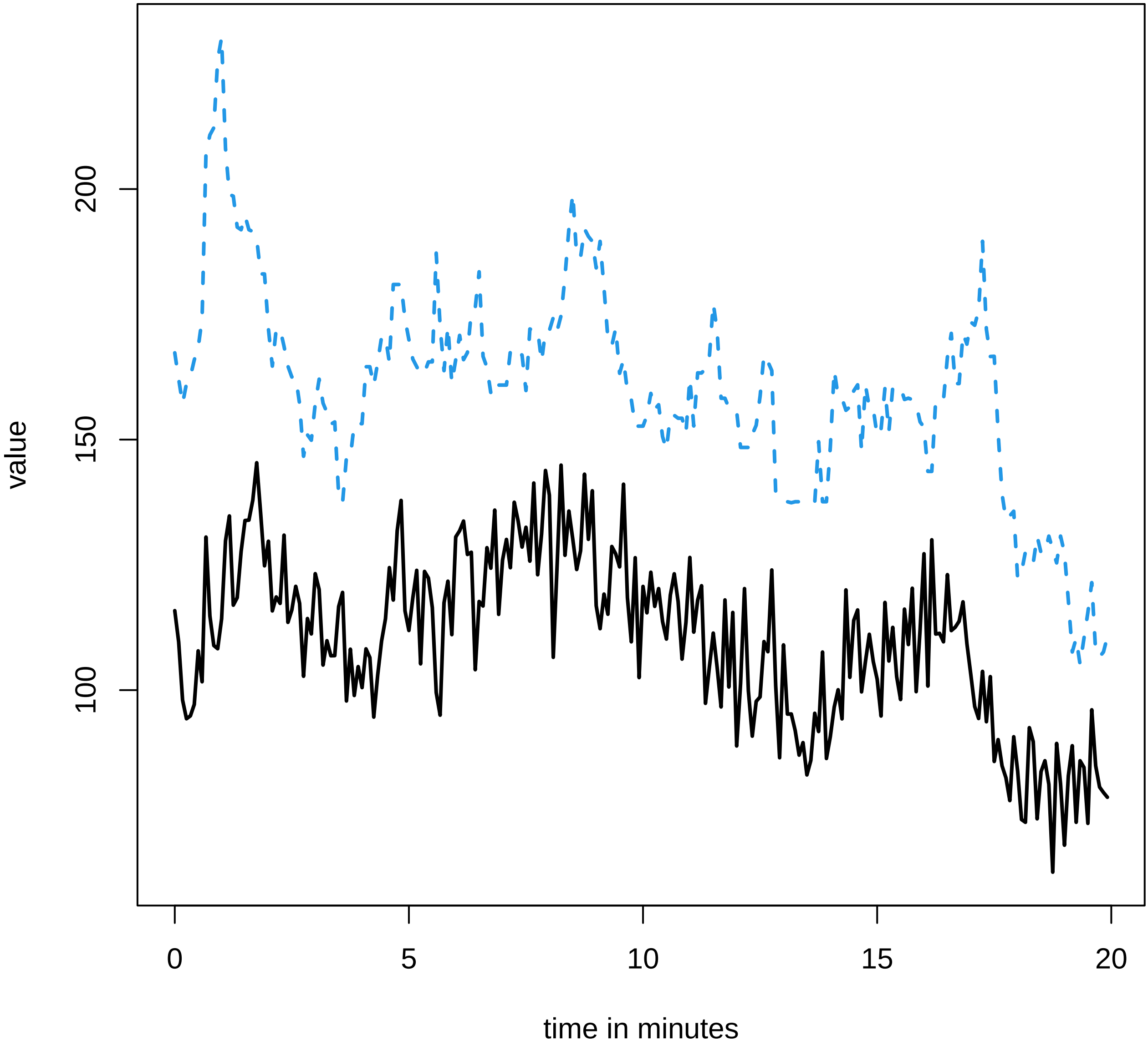

### 14:x_min=36, x_max=541 y_min=545, y_max=620, flashes=1

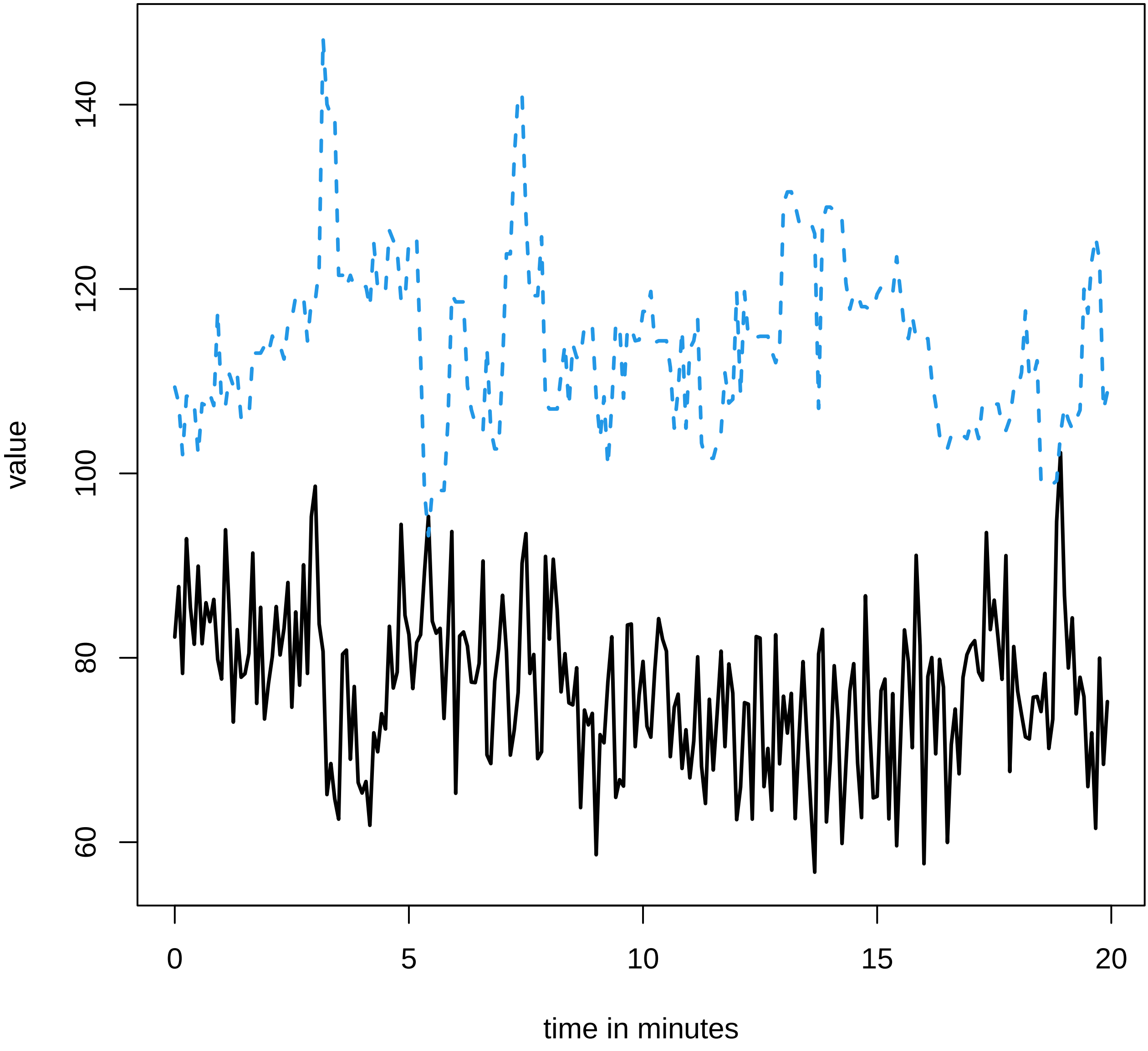

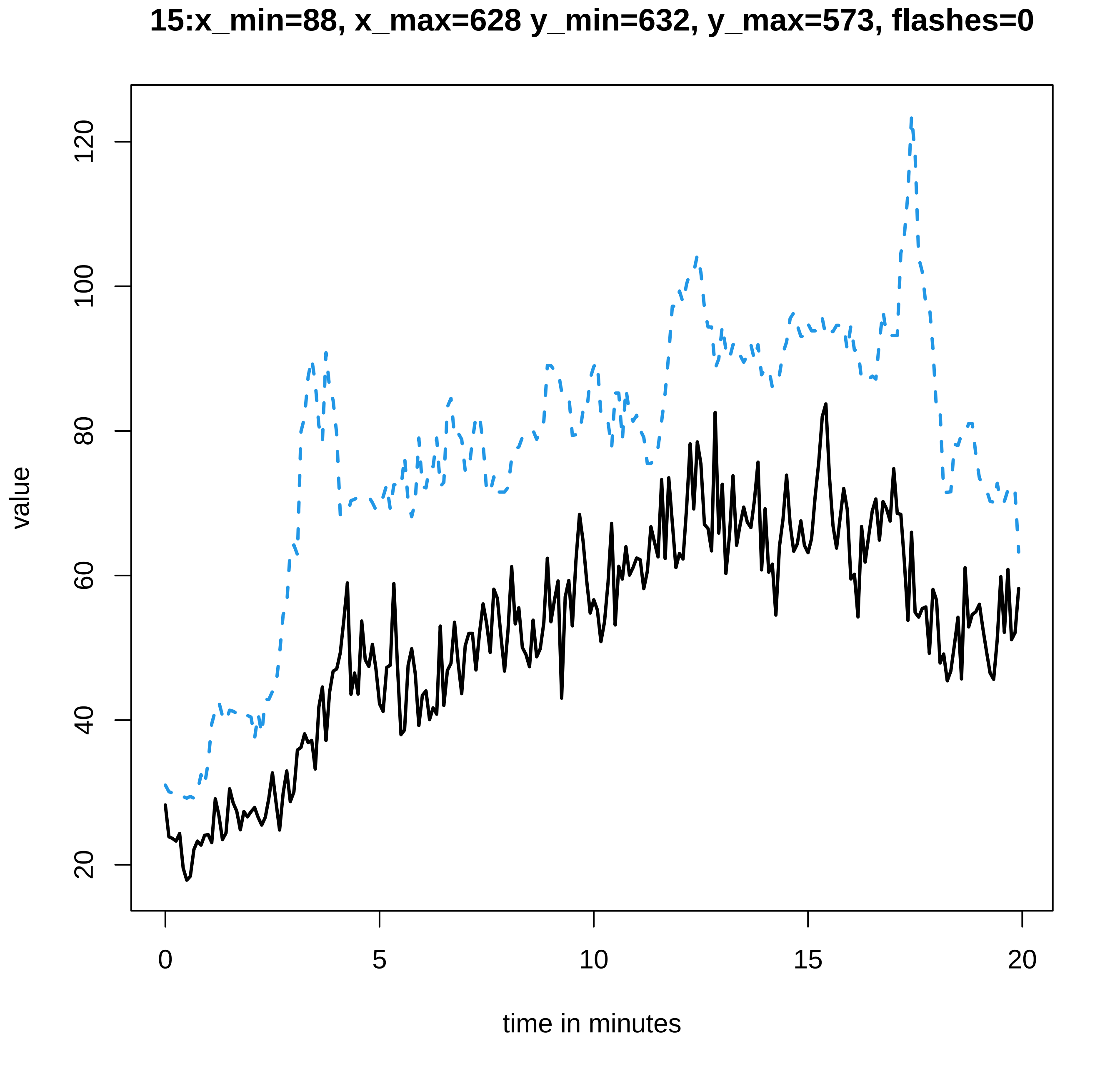
15:x_min=88, x_max=628 y_min=632, y_max=573, flashes=0
value
20
40
60
80
100
120
0
5
10
15
20
time in minutes

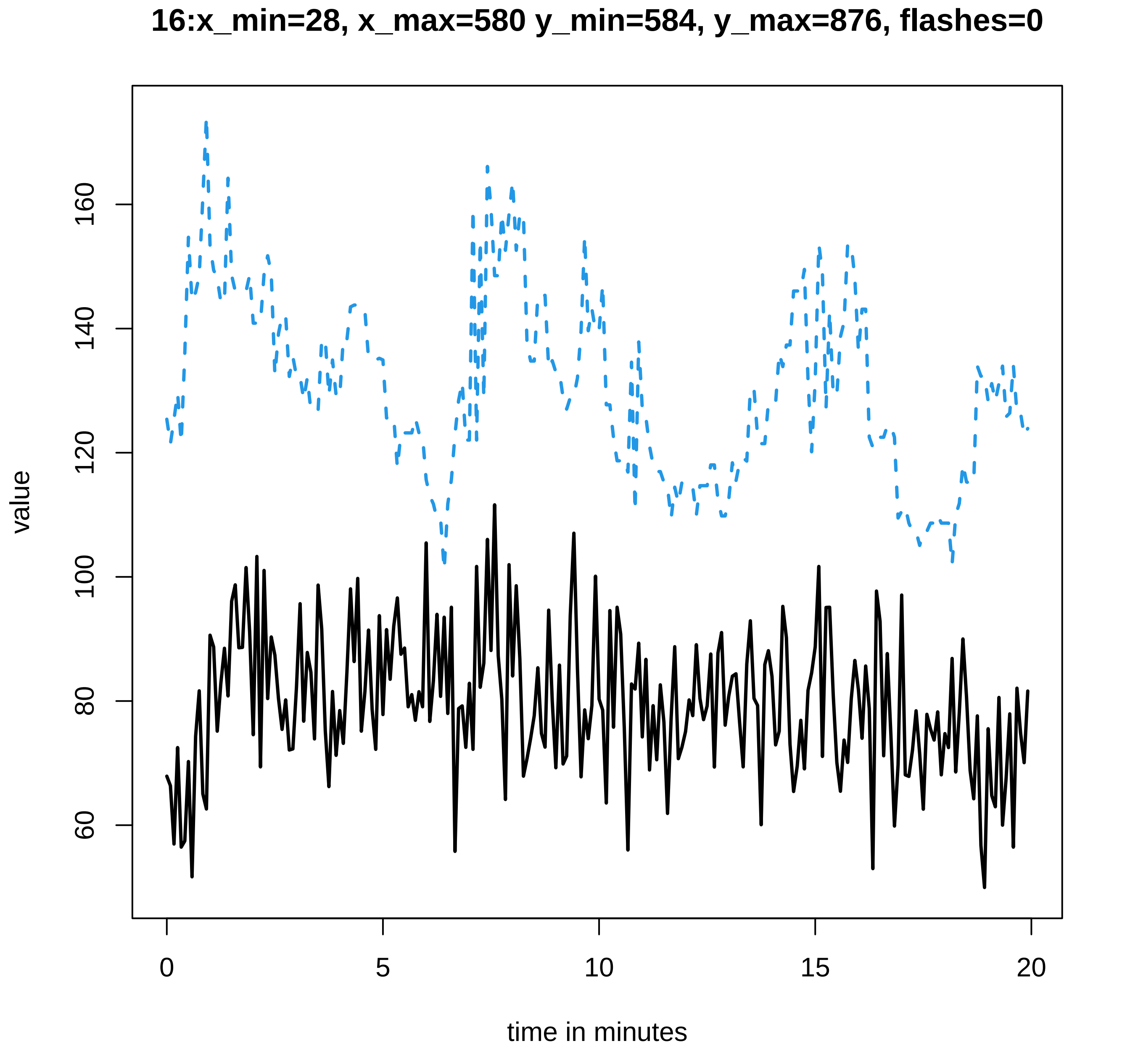
16:x_min=28, x_max=580 y_min=584, y_max=876, flashes=0
value
60
80
100
120
140
160
0
5
10
15
20
time in minutes

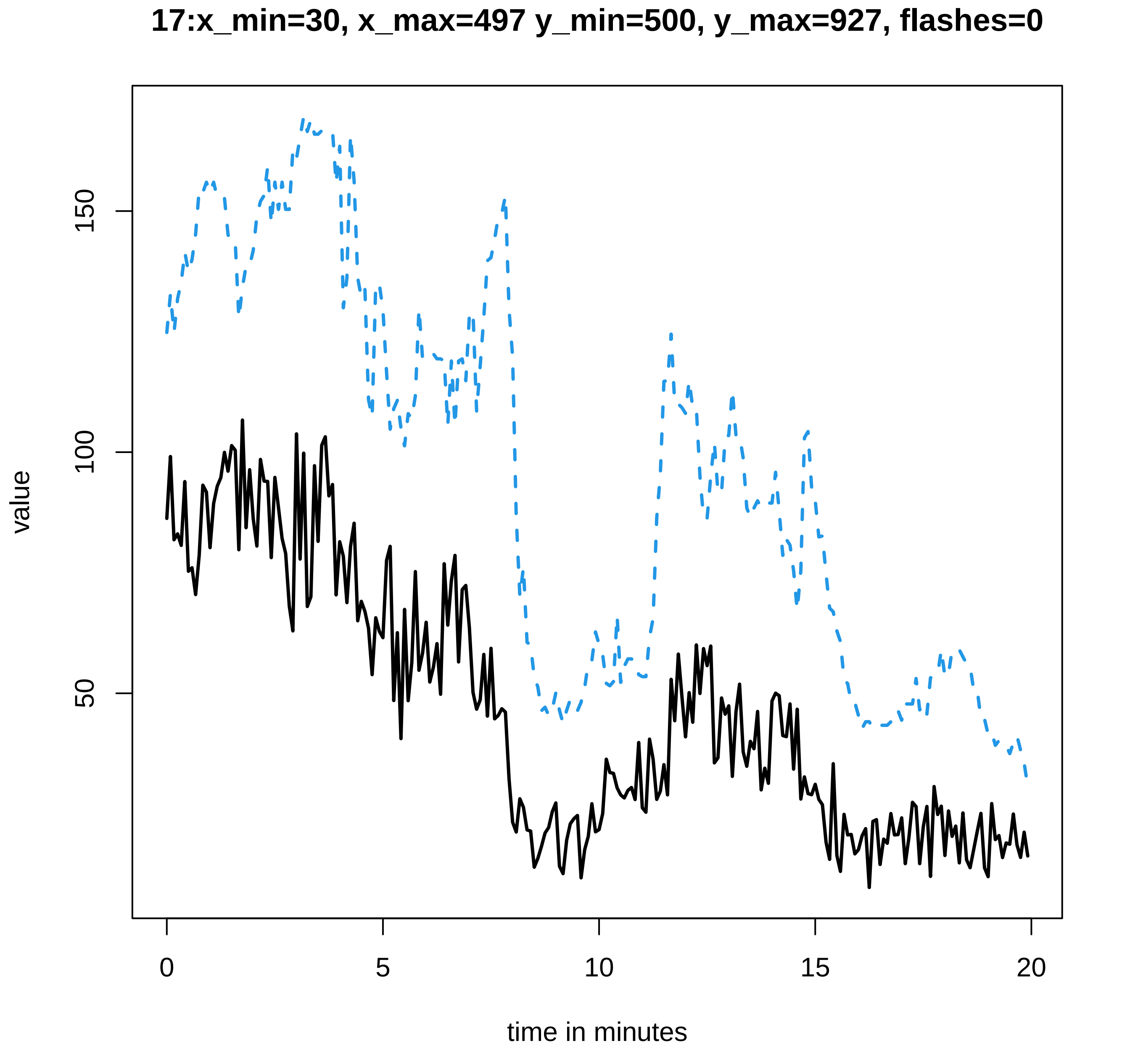
17:x_min=30, x_max=497 y_min=500, y_max=927, flashes=0
value
50
100
150
0
5
10
15
20
time in minutes

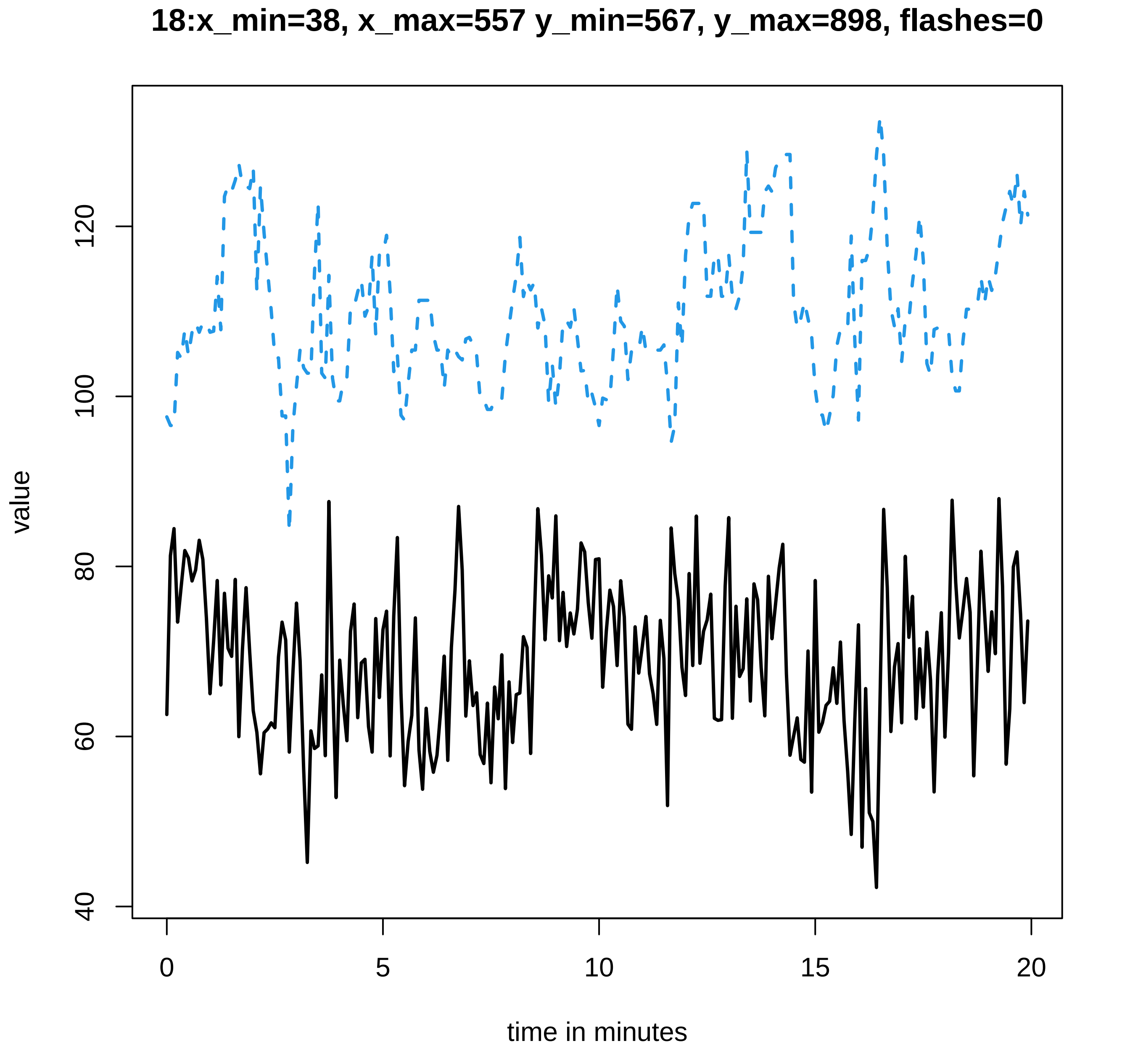

18:x_min=38, x_max=557 y_min=567, y_max=898, flashes=0
value
40
60
80
100
120
0
5
10
15
20
time in minutes

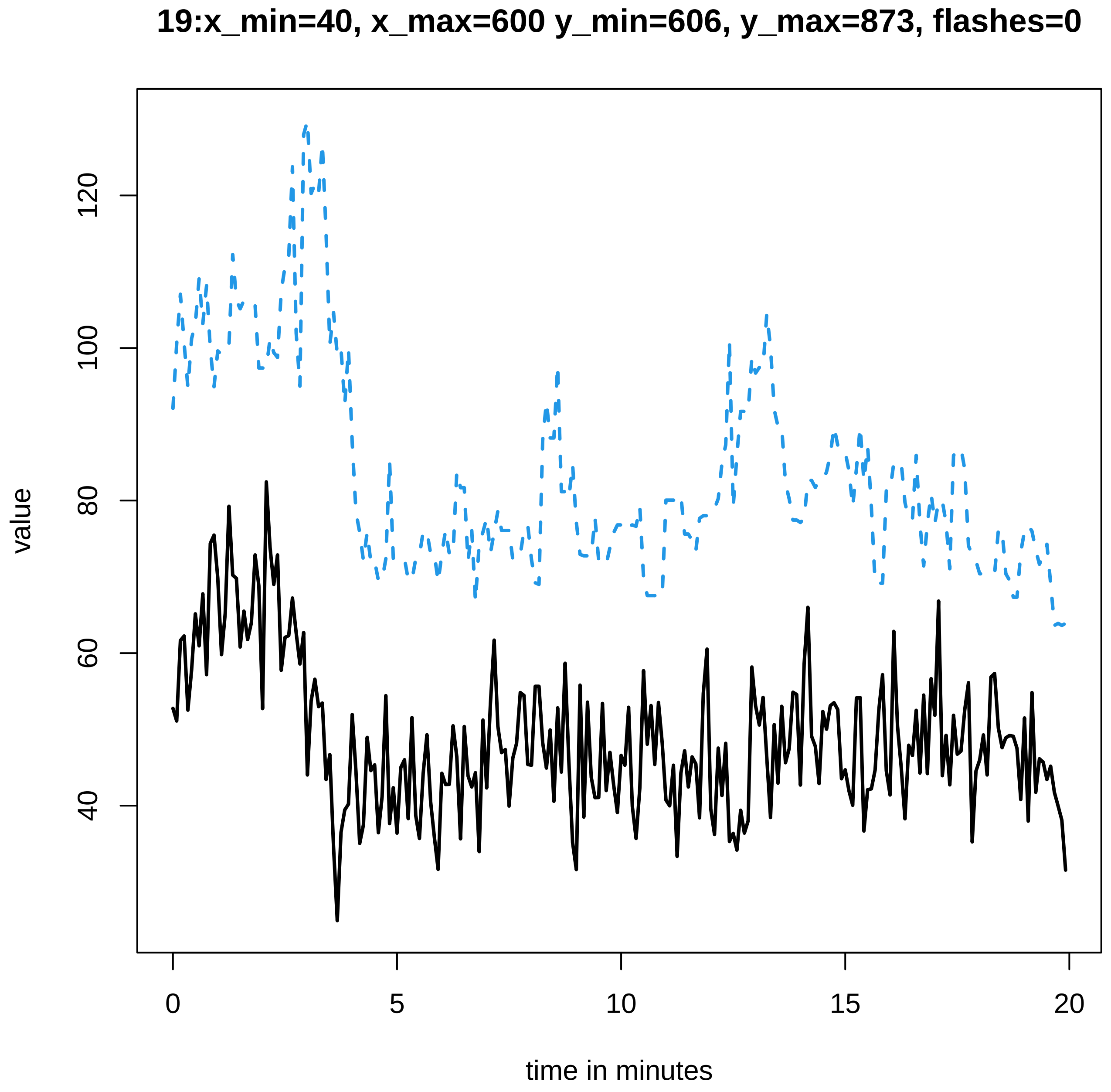
19:x_min=40, x_max=600 y_min=606, y_max=873, flashes=0
value
120
100
80
60
40
0
5
10
15
20
time in minutes

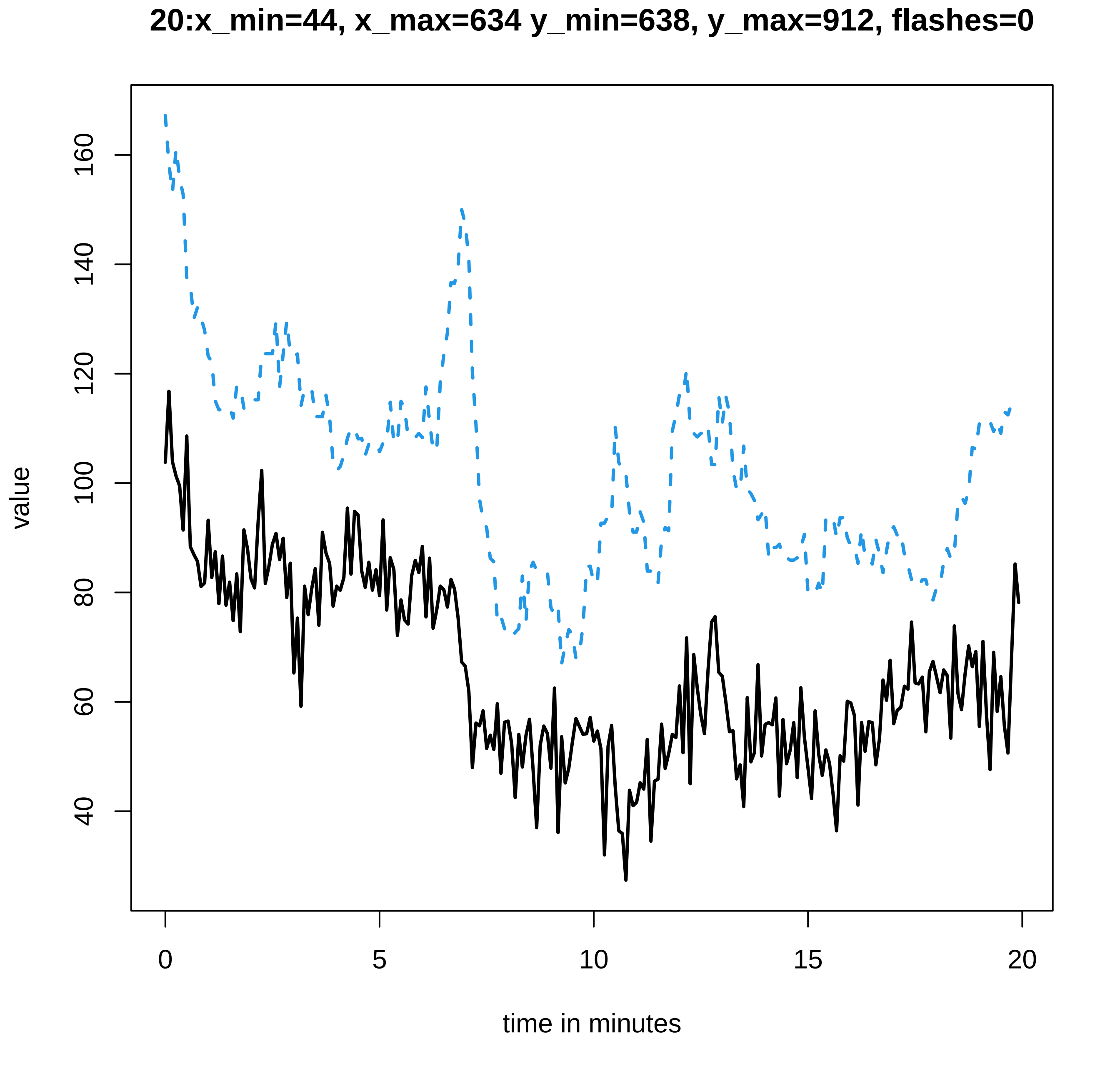

20:x_min=44, x_max=634 y_min=638, y_max=912, flashes=0
value
160
140
120
100
80
60
40
0
5
10
15
20
time in minutes

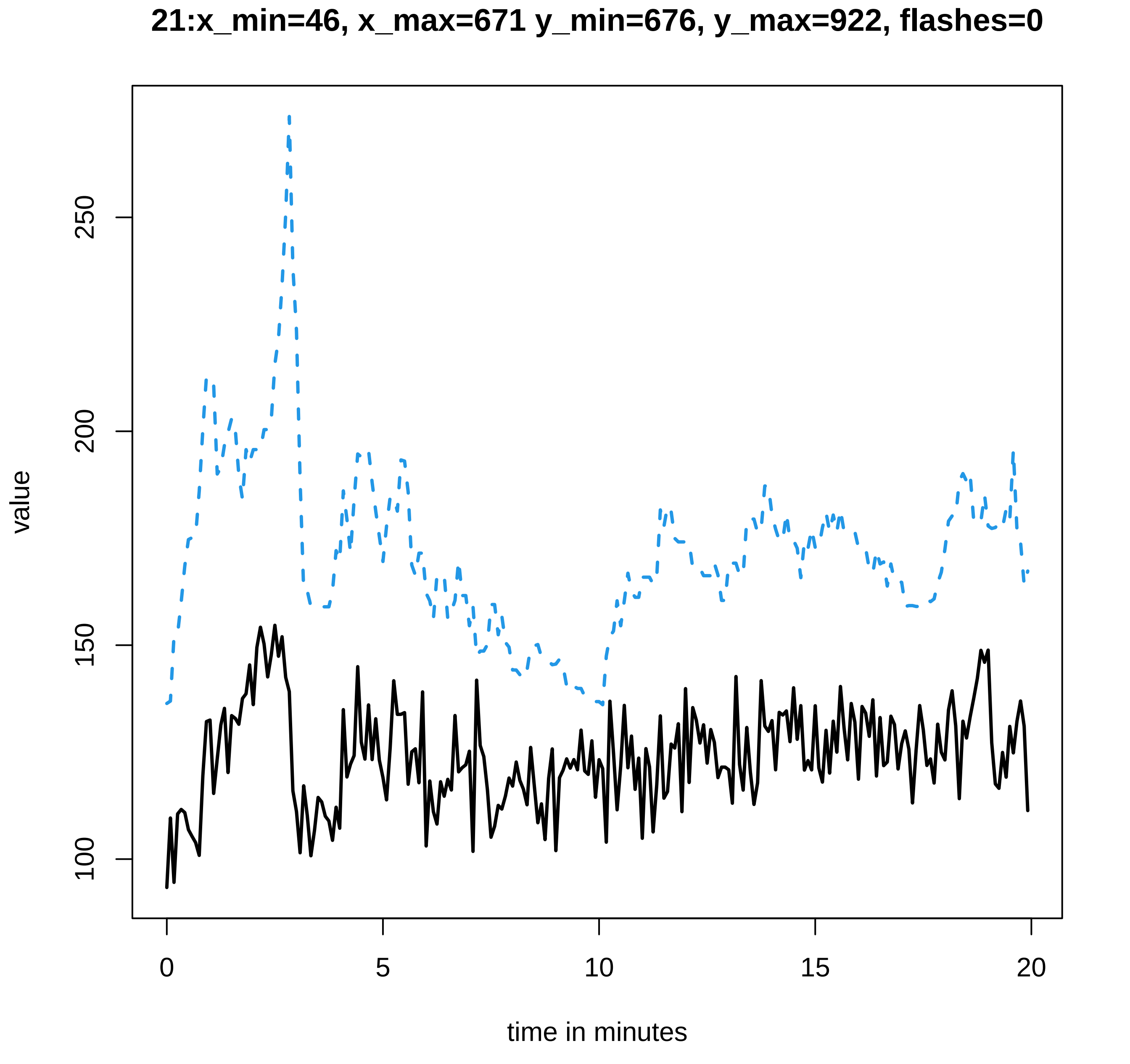
21:x_min=46, x_max=671 y_min=676, y_max=922, flashes=0
value
100
150
200
250
0
5
10
15
20
time in minutes
