## Supplementary data Calcium spiking for "LysM Receptor Proteins are Required for Ectomycorrhizal Symbiosis in Poplar": profiles-spiking-14.pdf

**1:x_min=32, x_max=410 y_min=415, y_max=91, flashes=0**

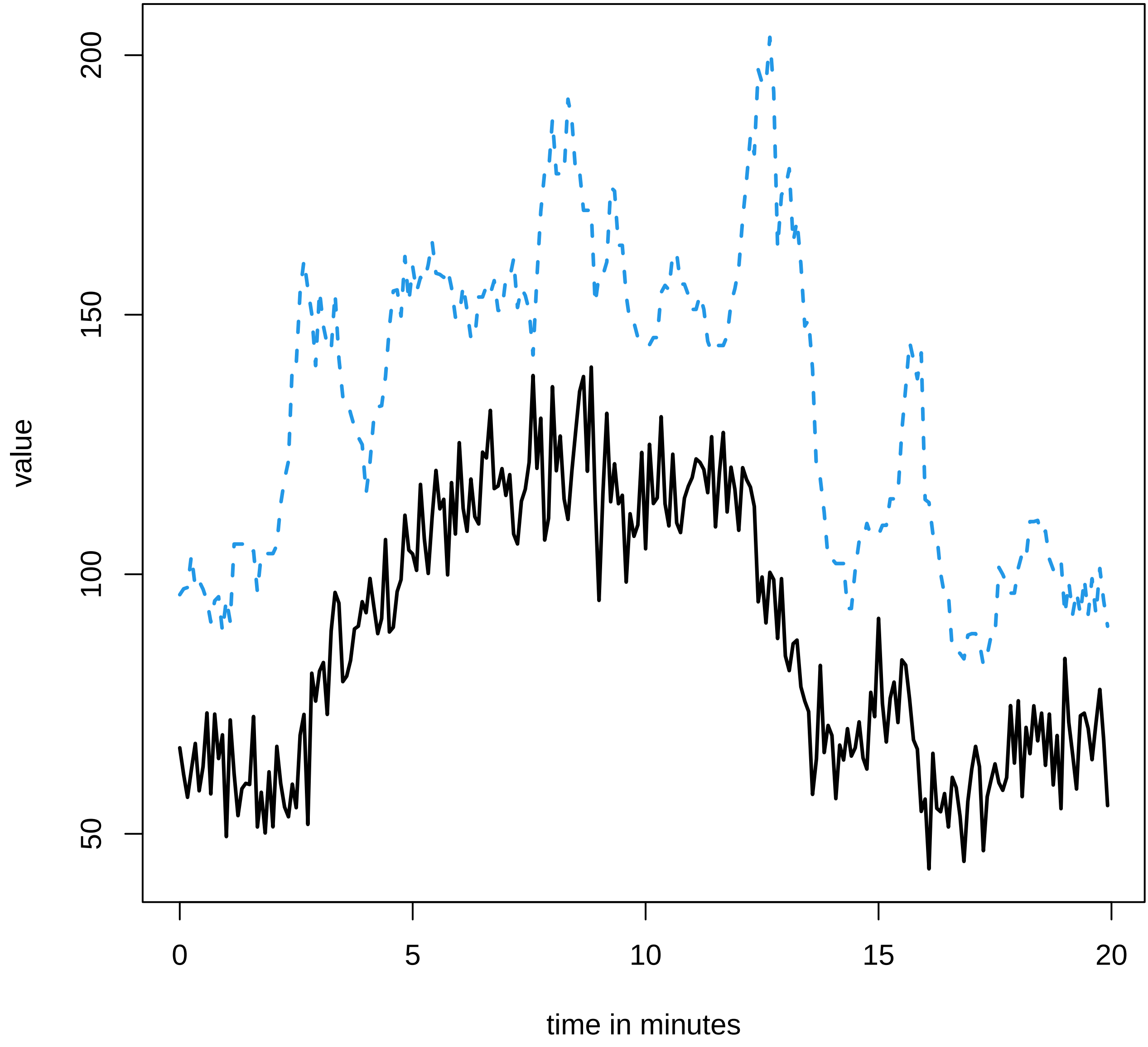

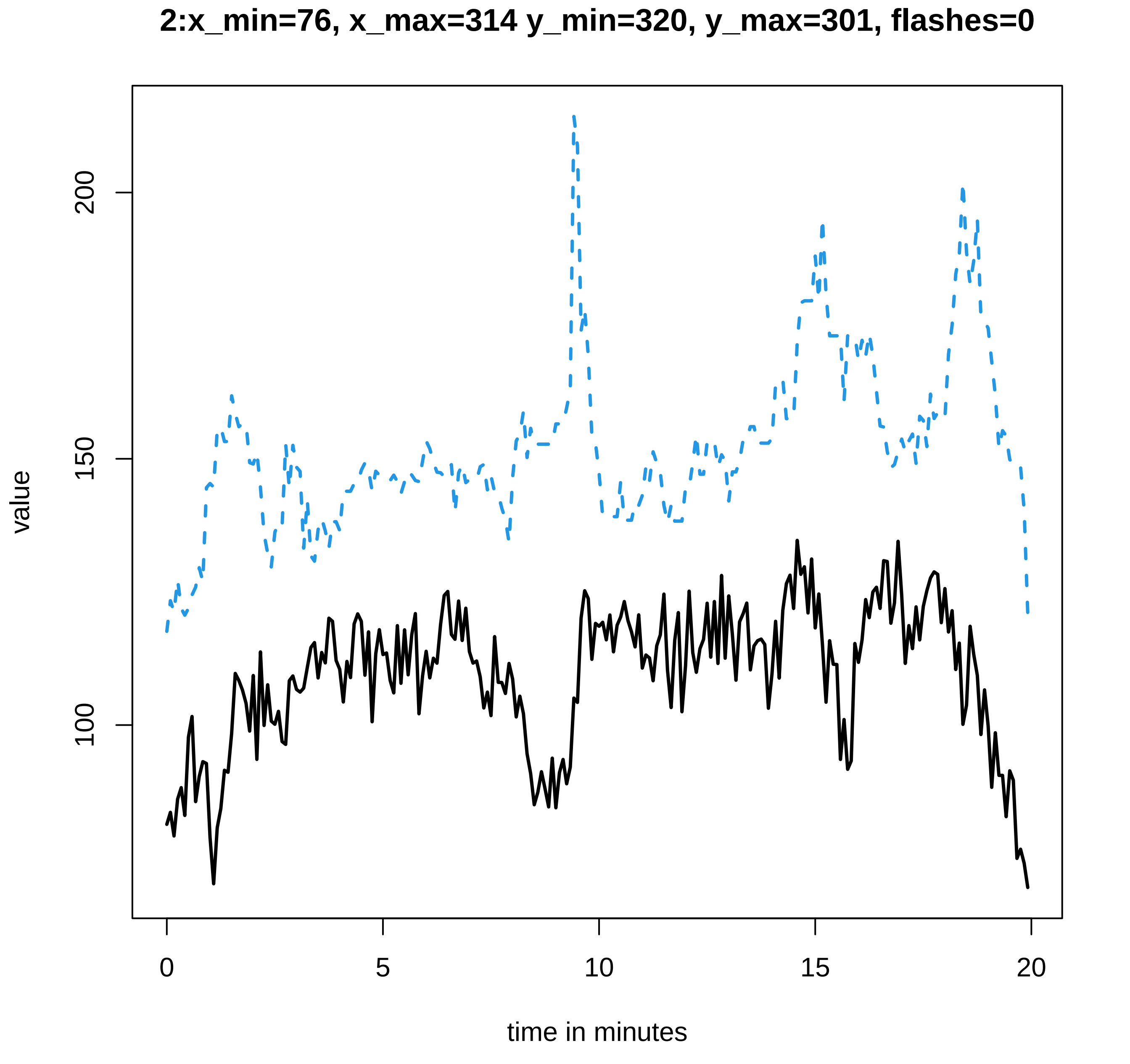

2:x_min=76, x_max=314 y_min=320, y_max=301, flashes=0
value
200
150
100
0
5
10
15
20
time in minutes

### 3:x_min=34, x_max=500 y_min=503, y_max=16, flashes=0

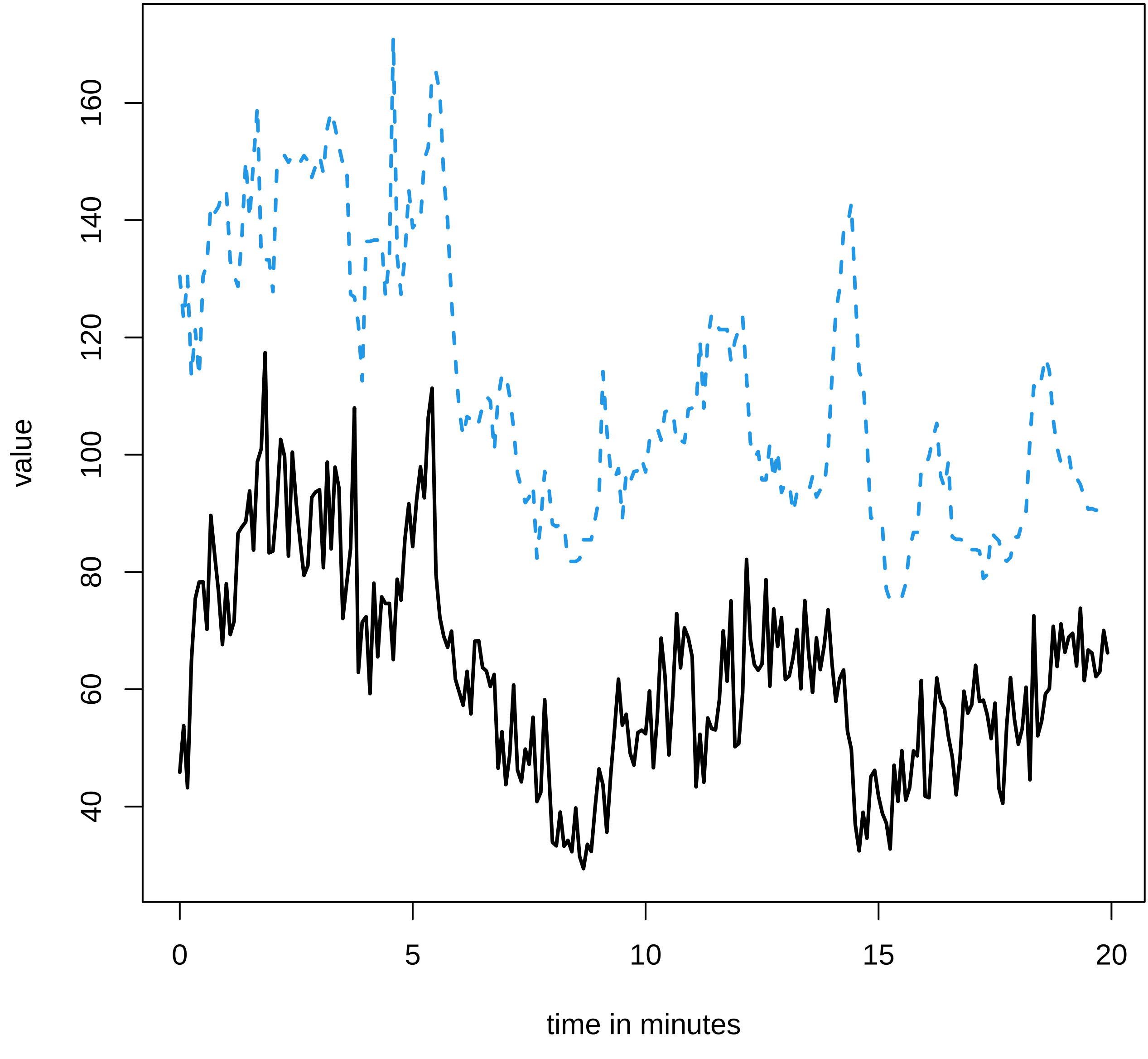

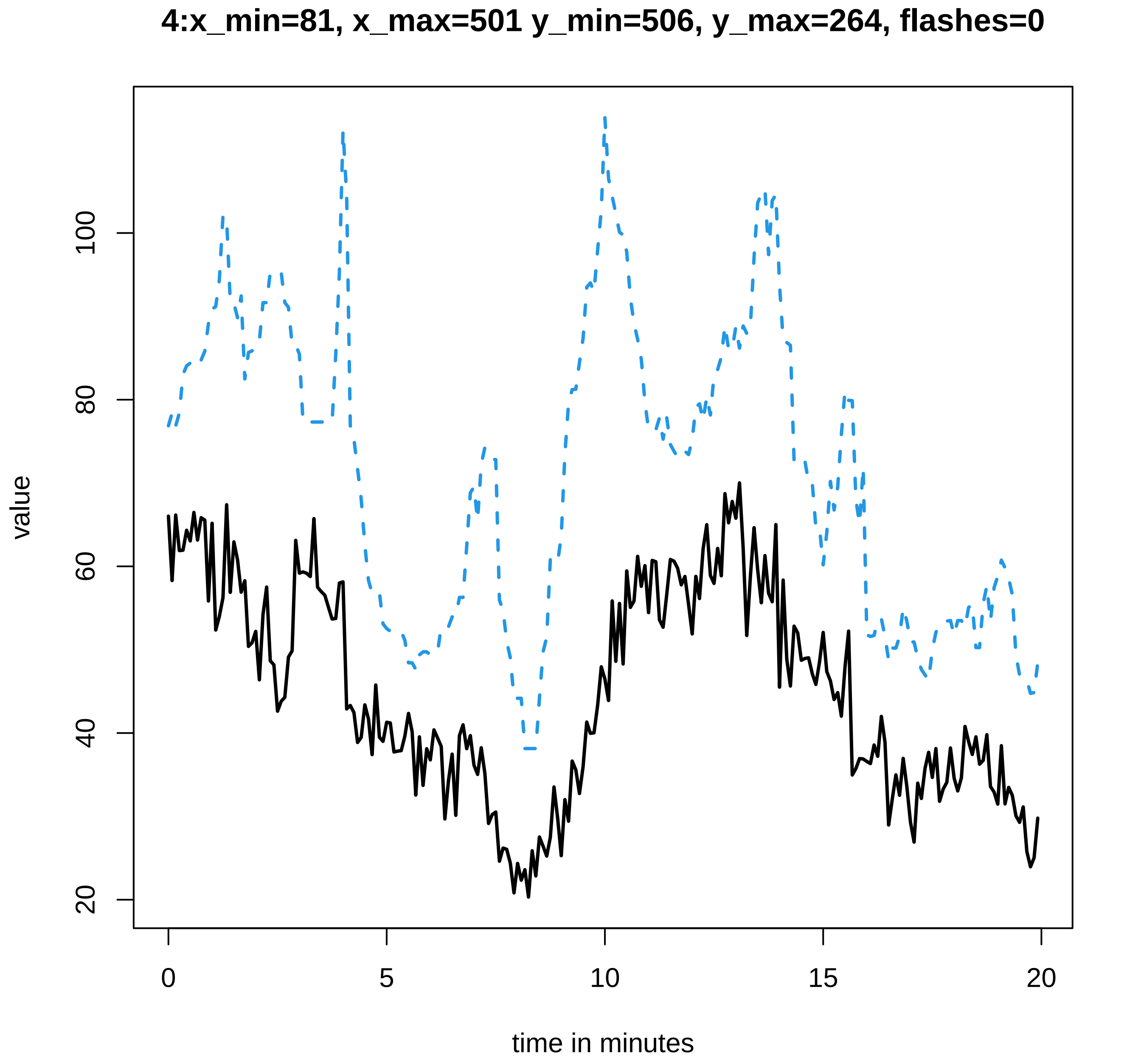
4:x_min=81, x_max=501 y_min=506, y_max=264, flashes=0
value
20
40
60
80
100
0
5
10
15
20
time in minutes

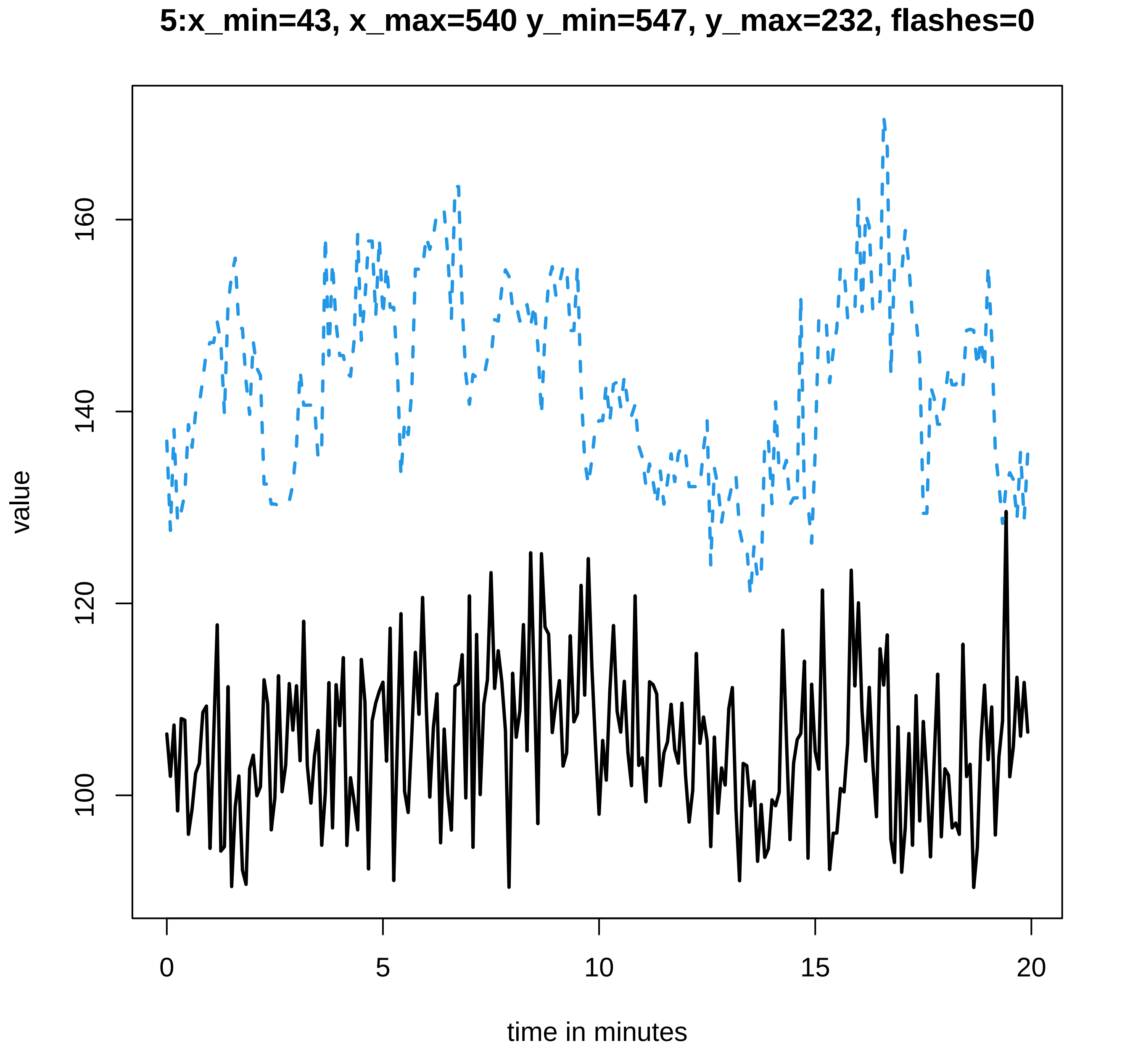

5:x_min=43, x_max=540 y_min=547, y_max=232, flashes=0
value
160
140
120
100
0
5
10
15
20
time in minutes

### 6:x_min=50, x_max=379 y_min=382, y_max=527, flashes=0

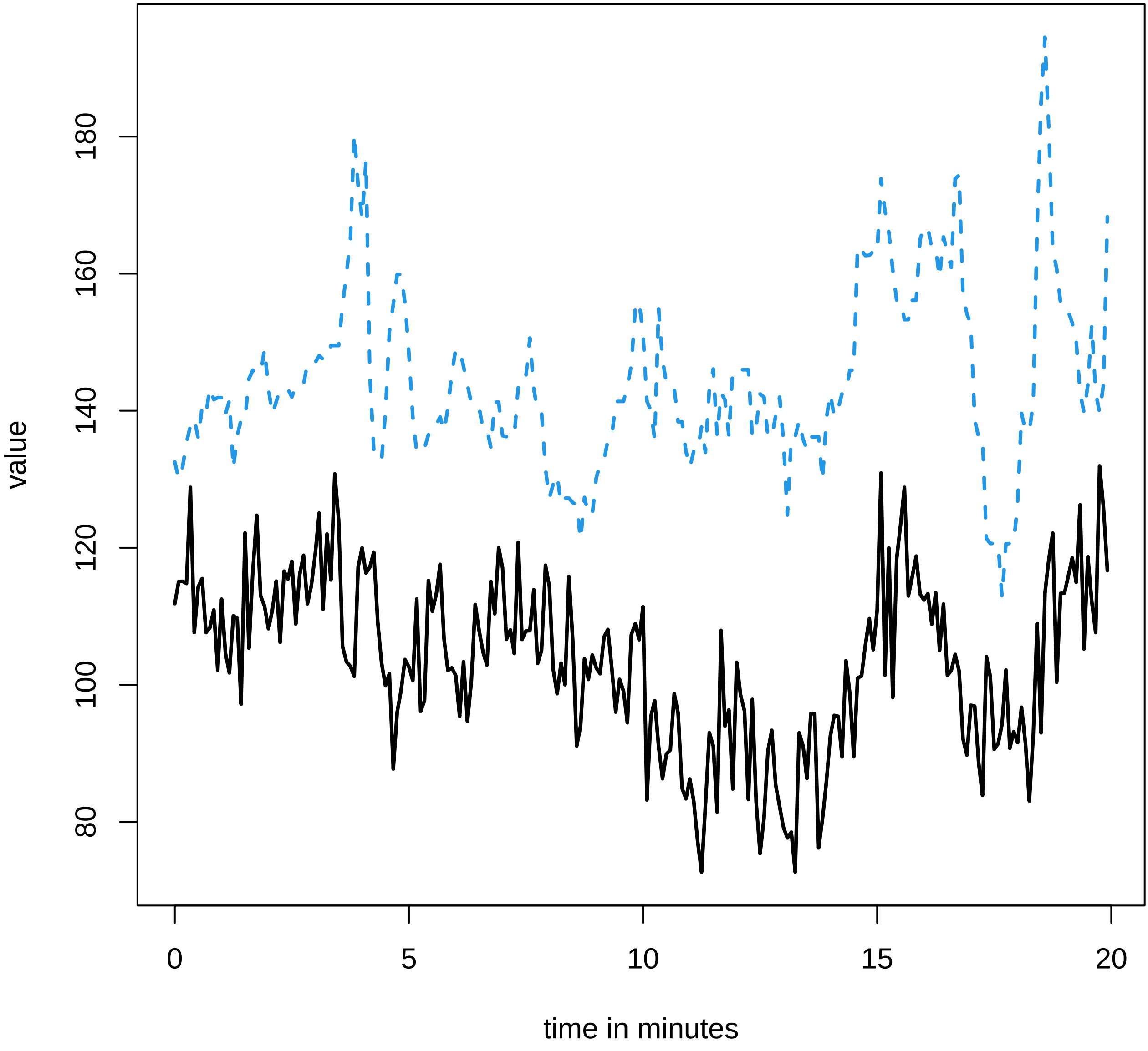

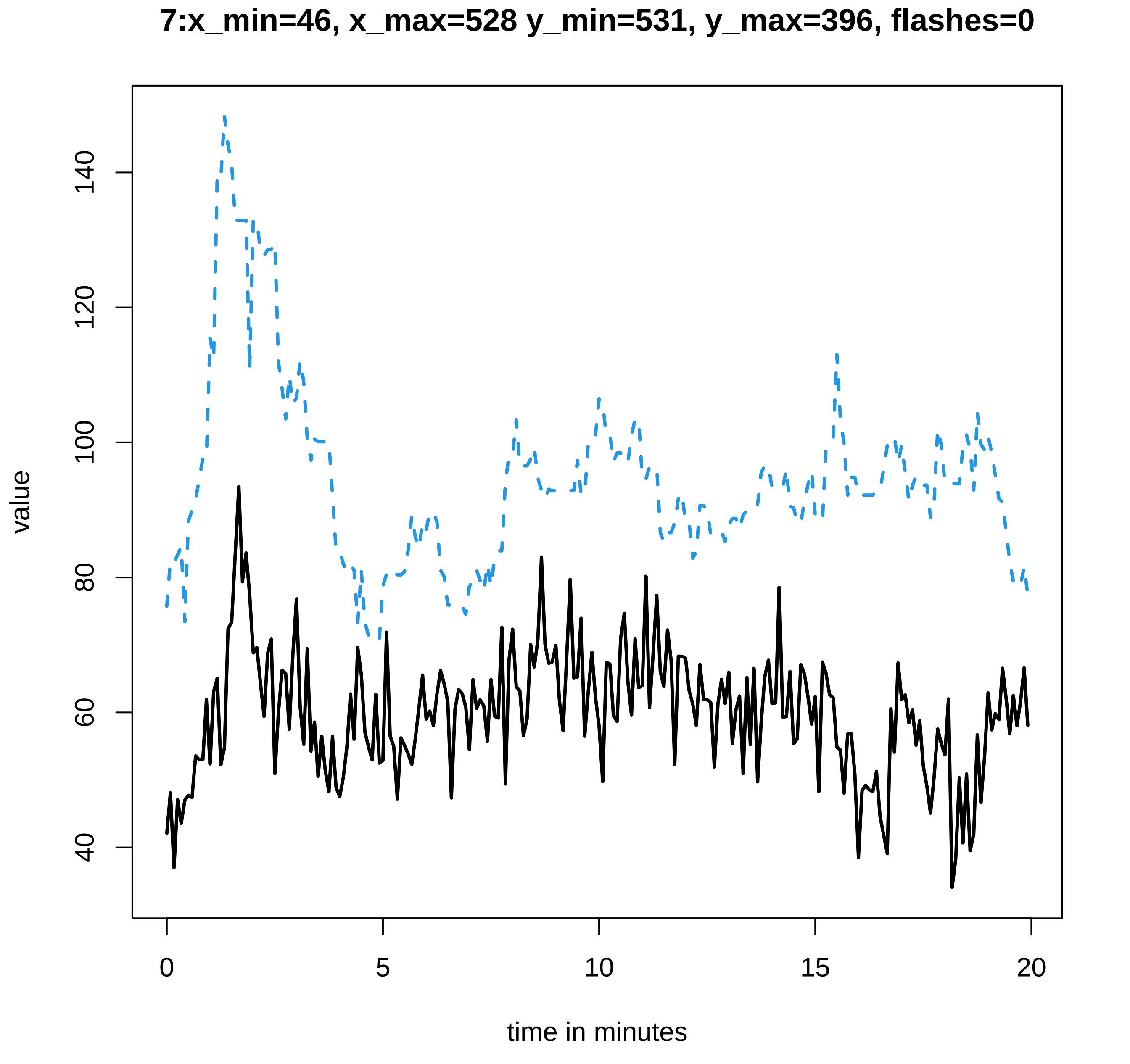
7:x_min=46, x_max=528 y_min=531, y_max=396, flashes=0
value
40
60
80
100
120
140
0
5
10
15
20
time in minutes

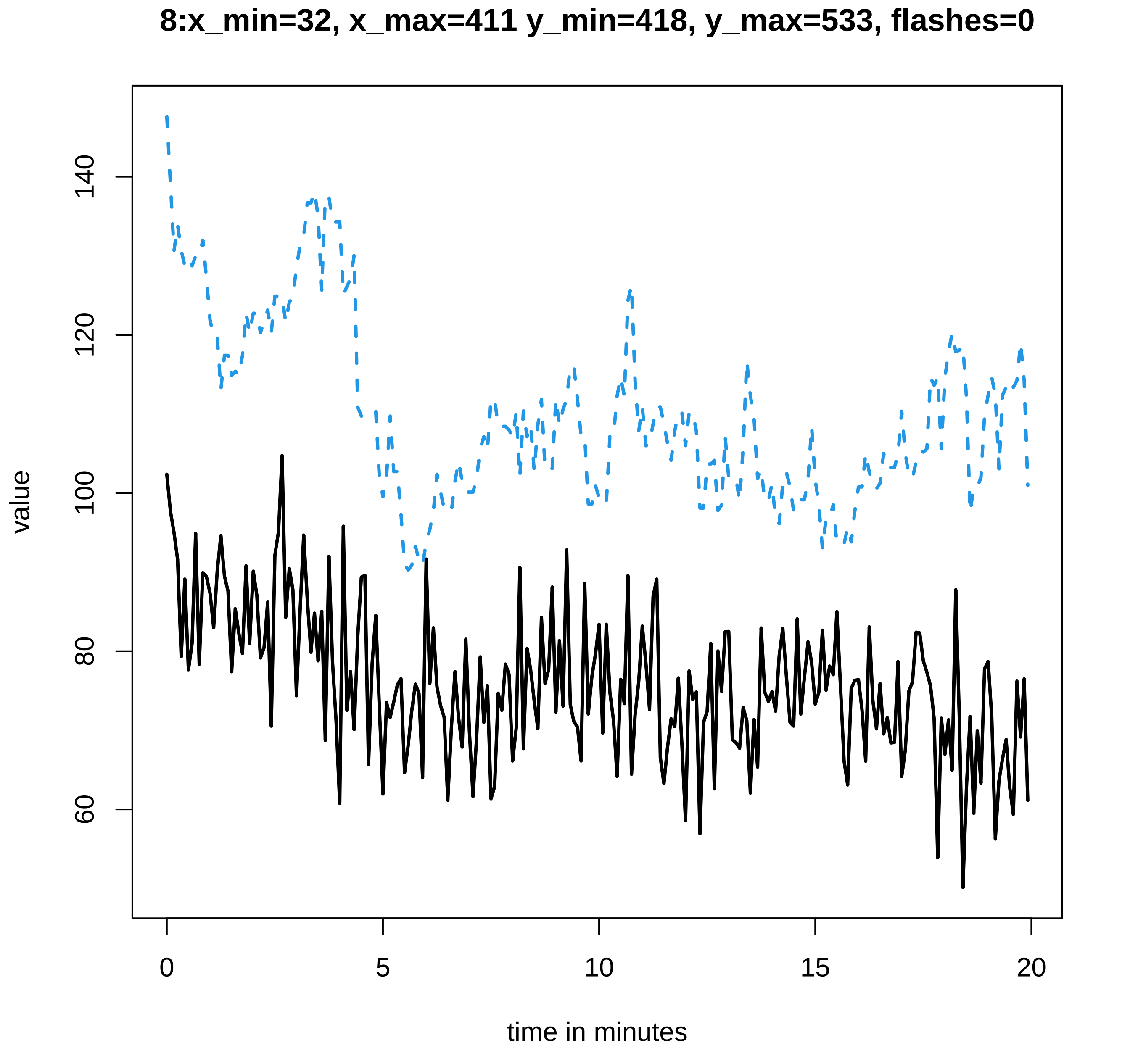
8:x_min=32, x_max=411 y_min=418, y_max=533, flashes=0
value
60
80
100
120
140
0
5
10
15
20
time in minutes

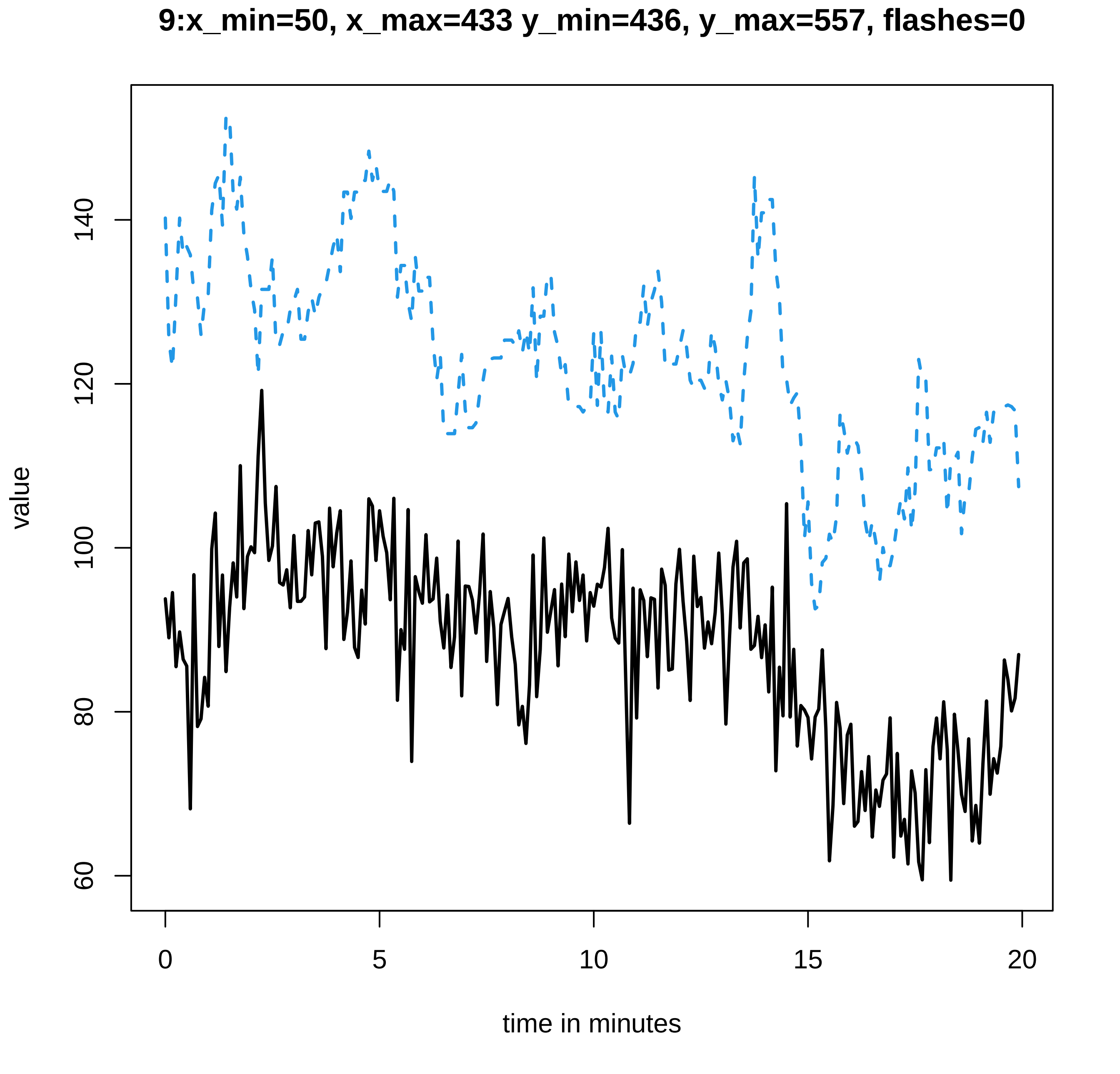
9:x_min=50, x_max=433 y_min=436, y_max=557, flashes=0
value
time in minutes
0
5
10
15
20
60
80
100
120
140

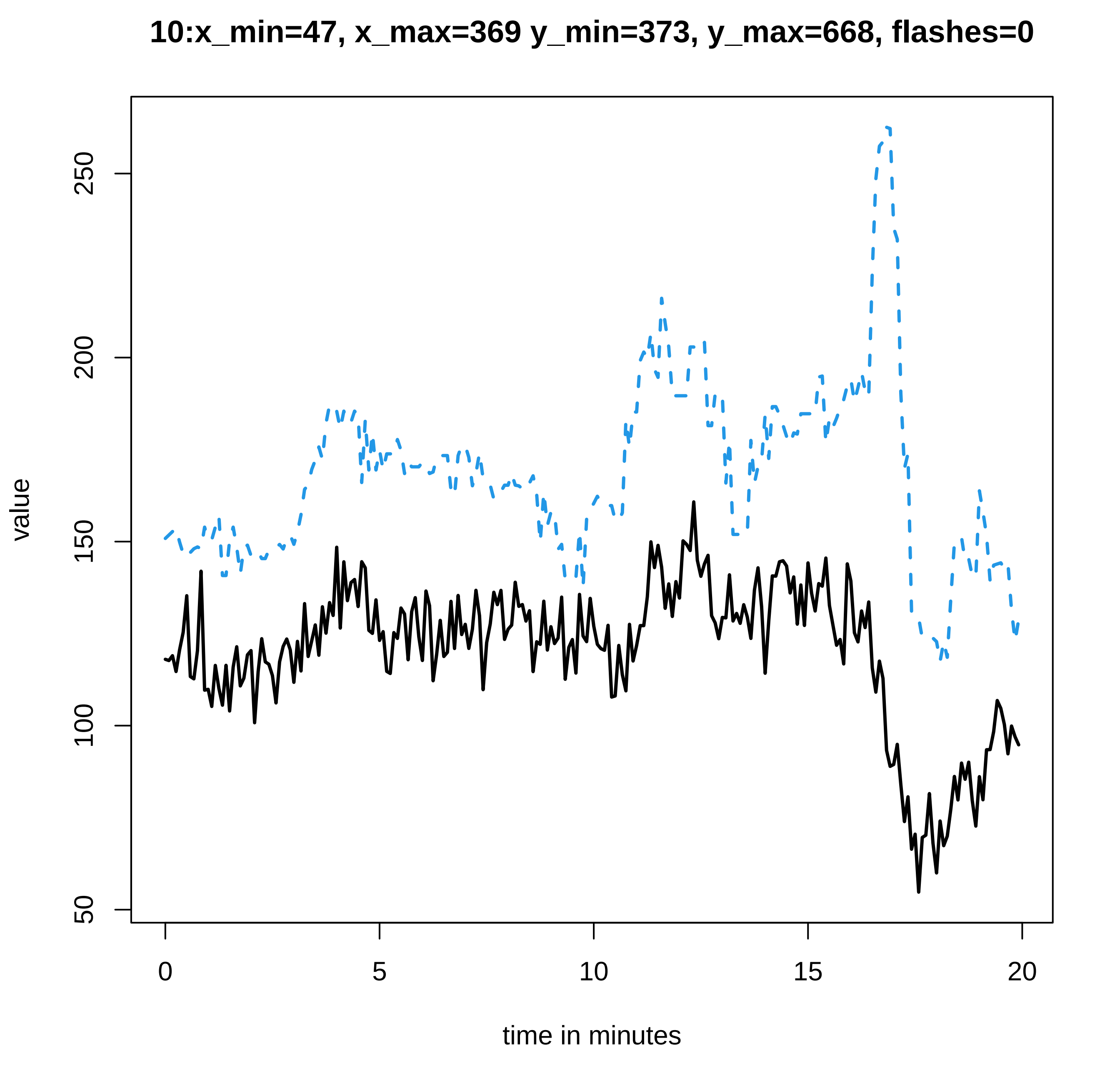

10:x_min=47, x_max=369 y_min=373, y_max=668, flashes=0
value
250
200
150
100
50
0
5
10
15
20
time in minutes

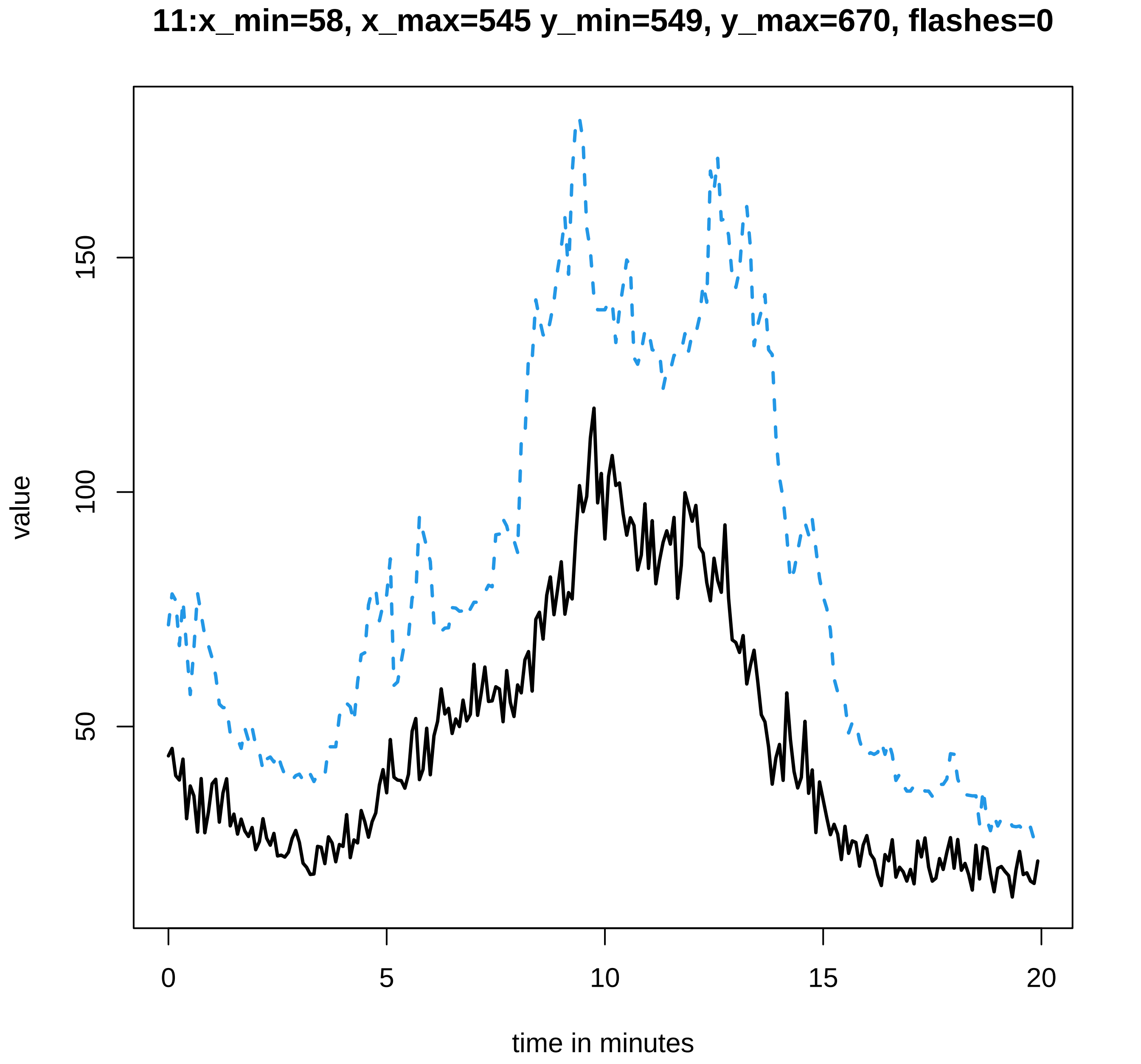
11:x_min=58, x_max=545 y_min=549, y_max=670, flashes=0
value
50
100
150
0
5
10
15
20
time in minutes

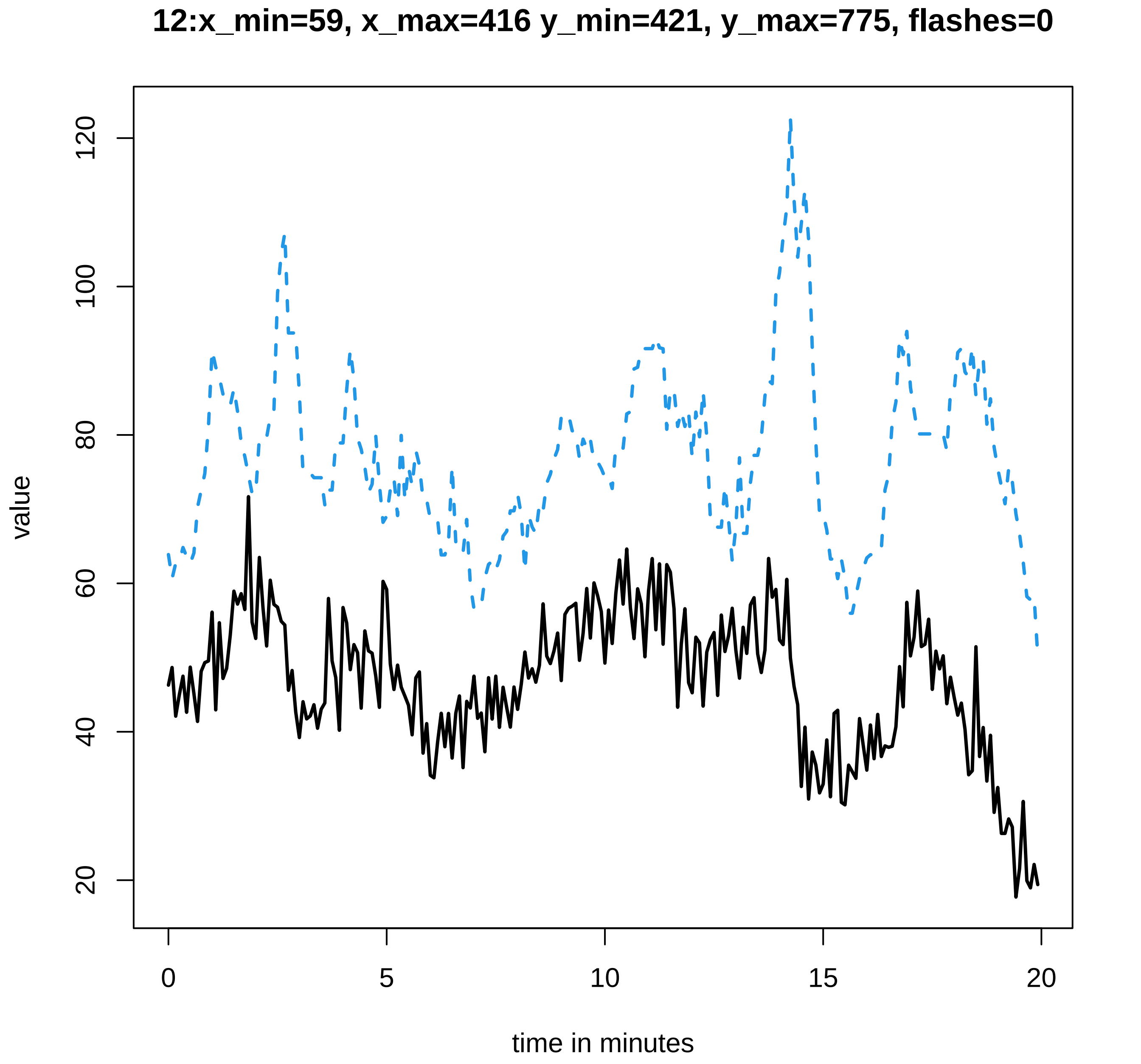
12:x_min=59, x_max=416 y_min=421, y_max=775, flashes=0
value
120
100
80
60
40
20
0
5
10
15
20
time in minutes

#### 13:x_min=88, x_max=470 y_min=474, y_max=753, flashes=0

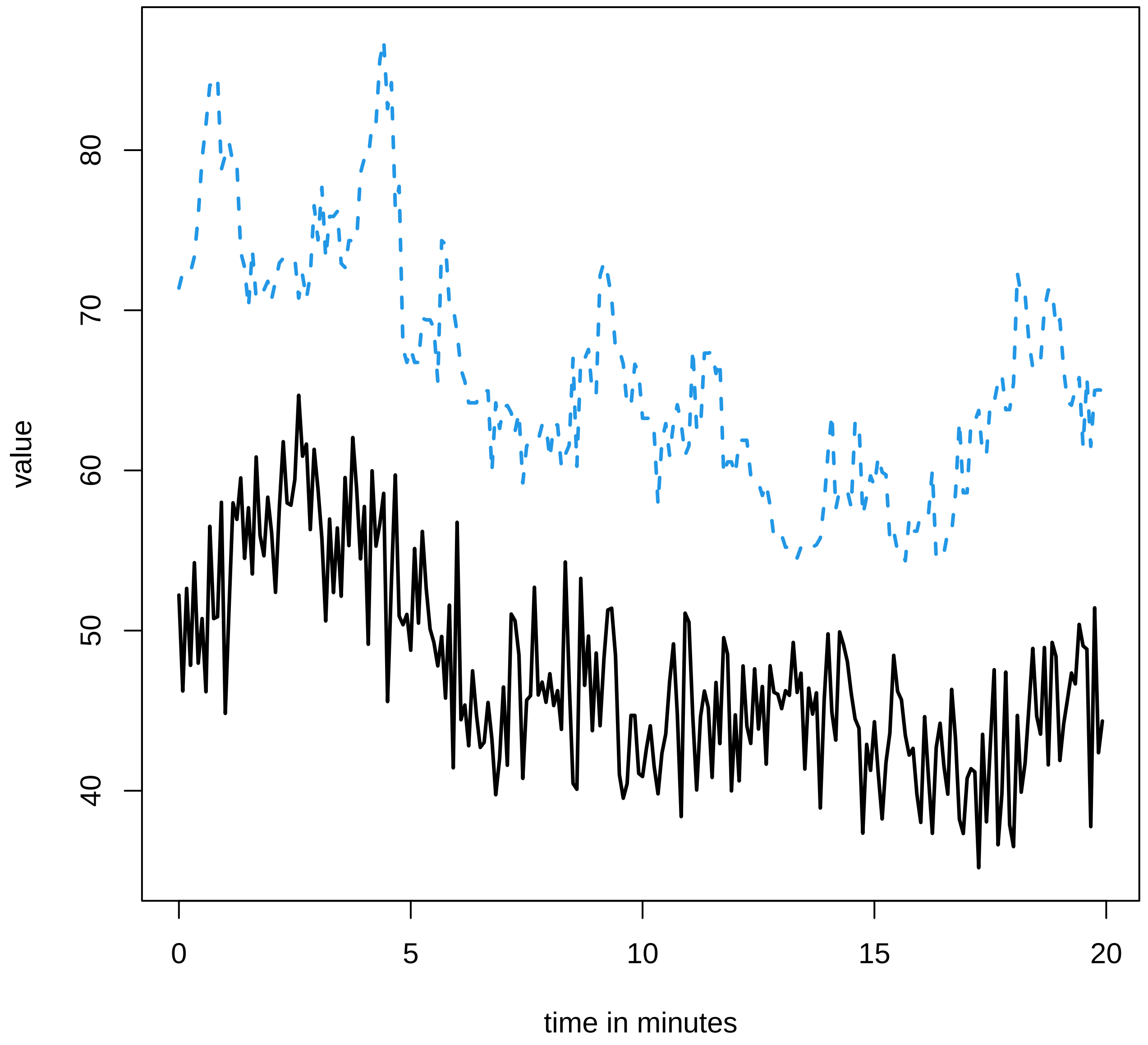

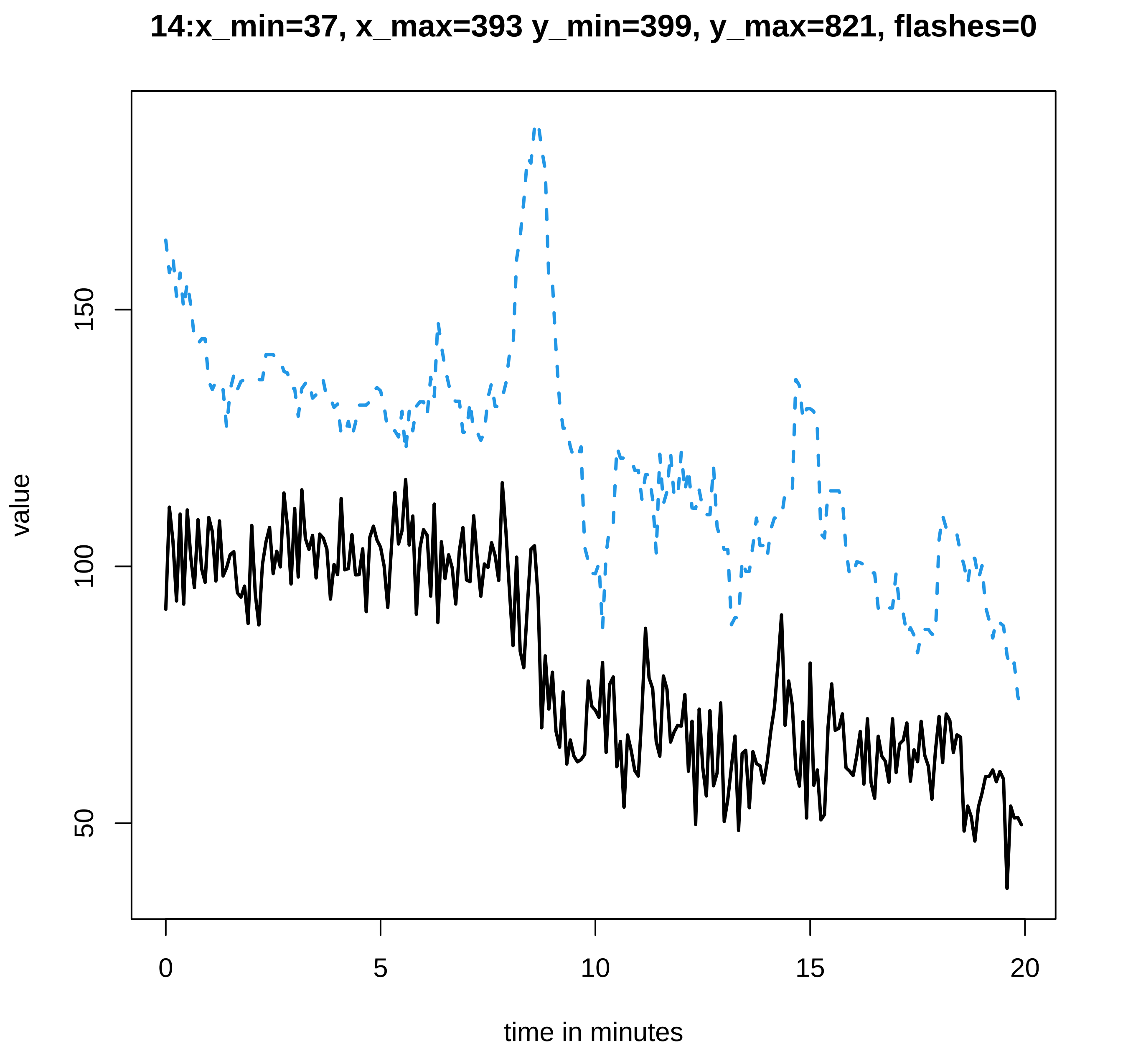

14:x_min=37, x_max=393 y_min=399, y_max=821, flashes=0
value
150
100
50
0
5
10
15
20
time in minutes

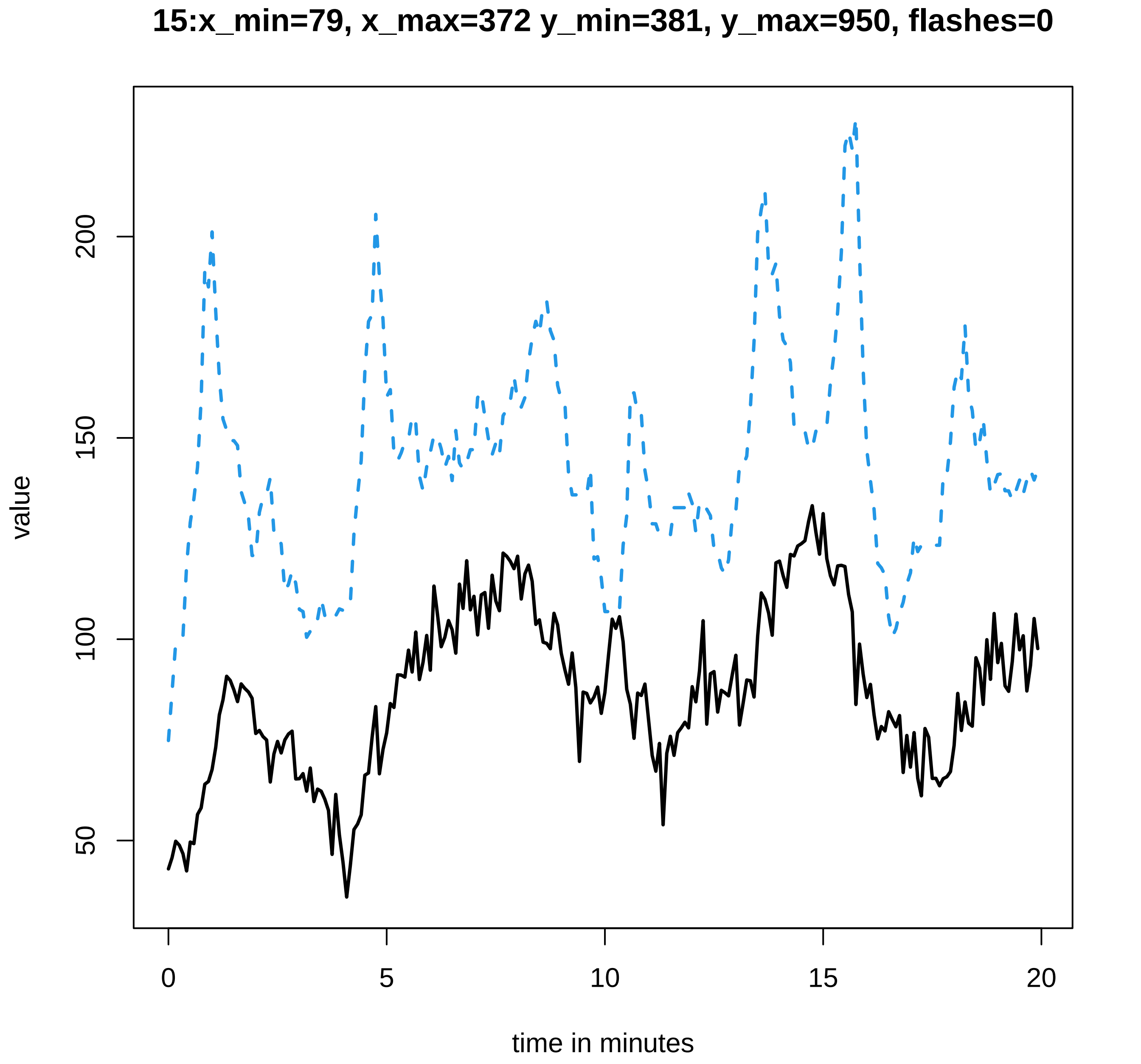

15:x_min=79, x_max=372 y_min=381, y_max=950, flashes=0
value
50
100
150
200
0
5
10
15
20
time in minutes

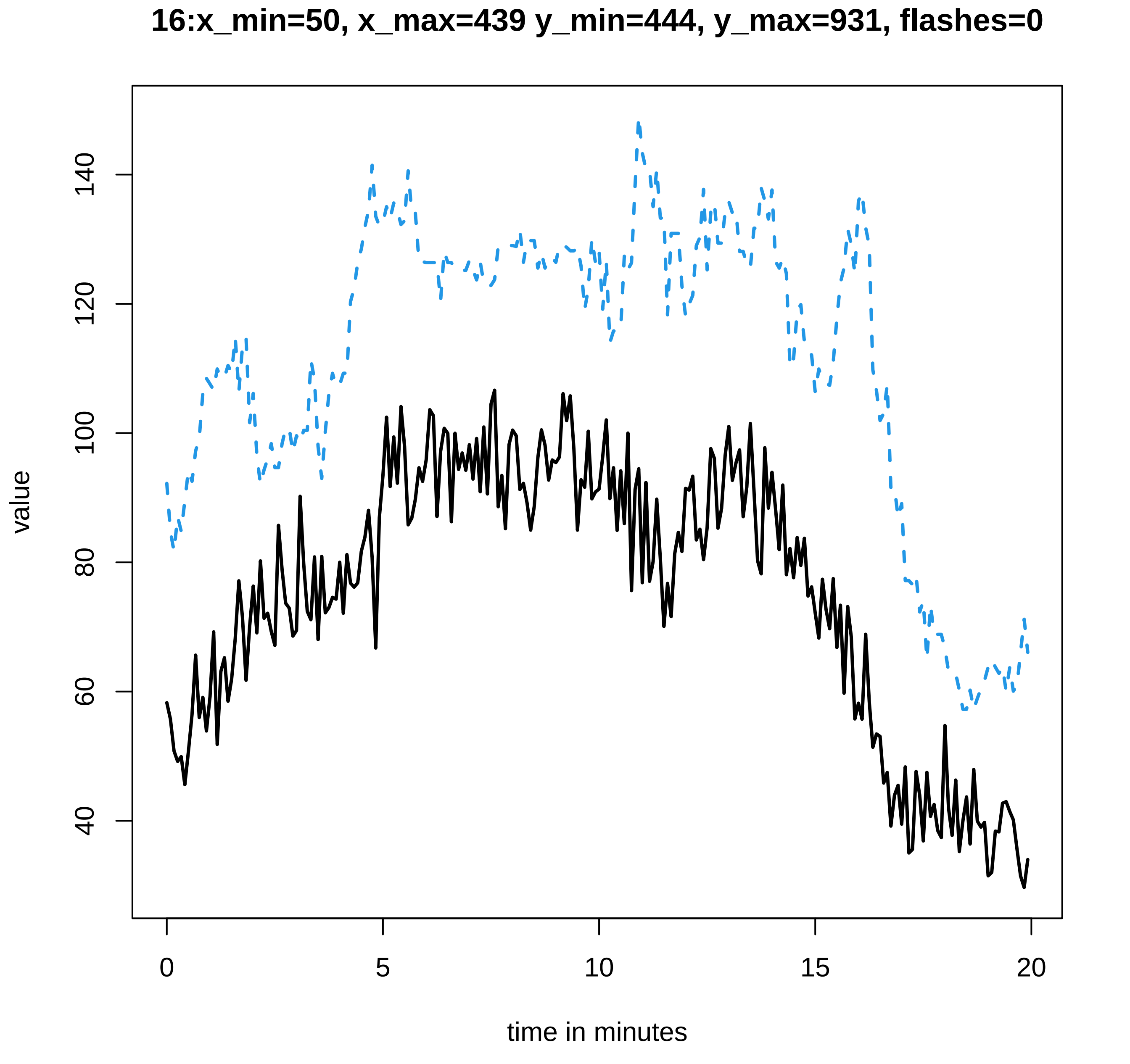
16:x_min=50, x_max=439 y_min=444, y_max=931, flashes=0
value
40
60
80
100
120
140
0
5
10
15
20
time in minutes

#### 17:x_min=44, x_max=546 y_min=550, y_max=889, flashes=0

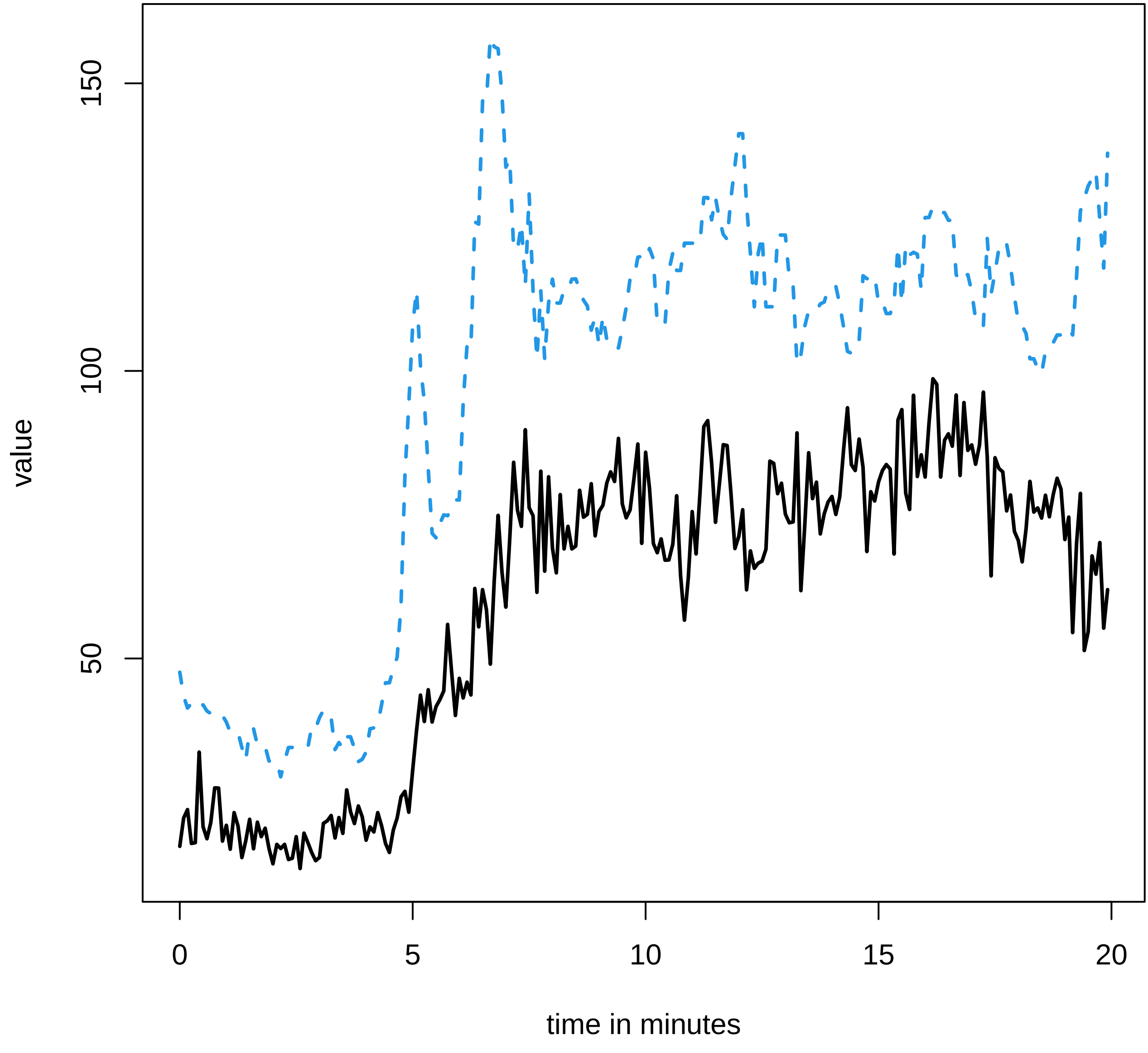

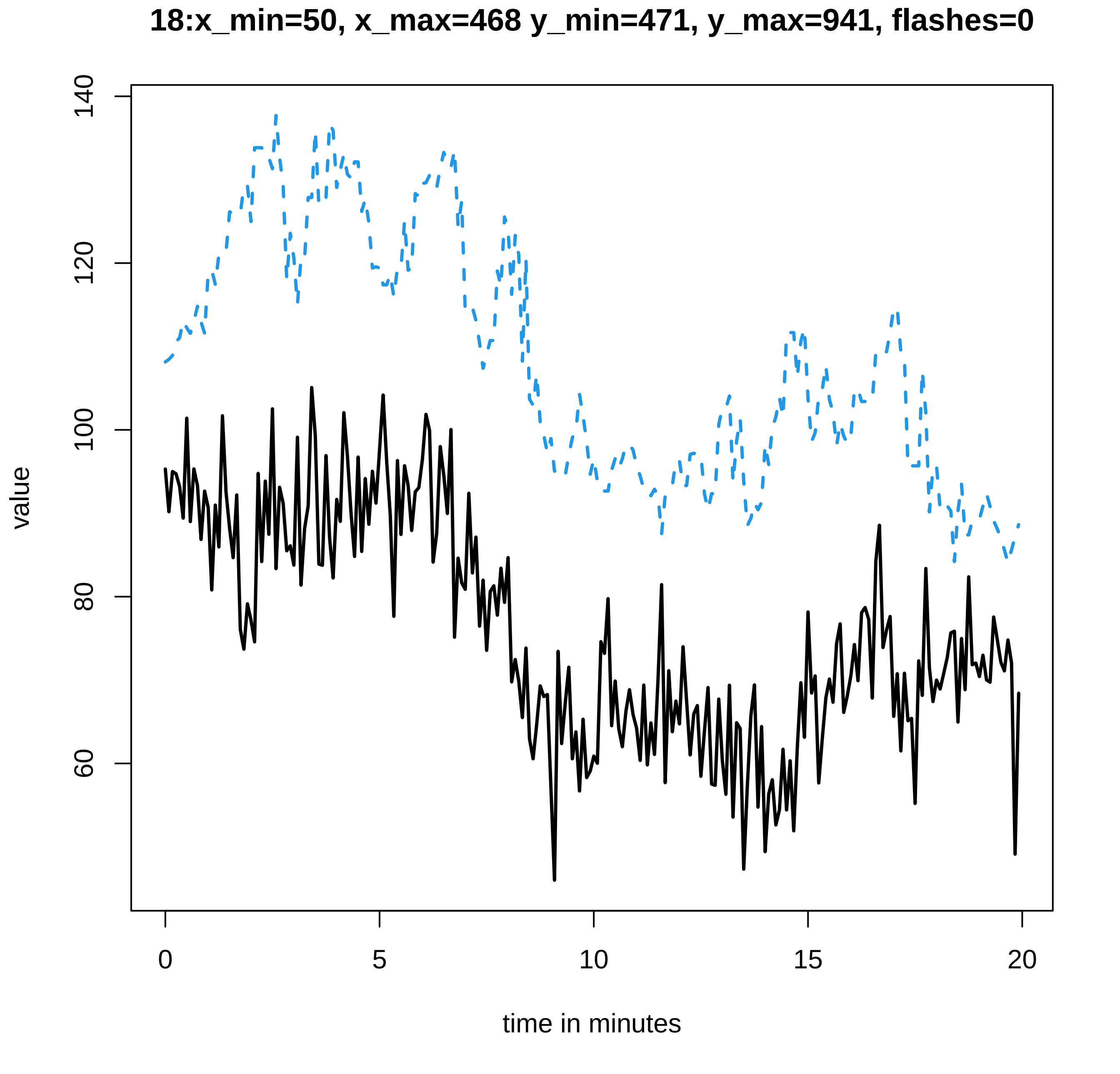

18:x_min=50, x_max=468 y_min=471, y_max=941, flashes=0
140
120
100
80
60
value
0
5
10
15
20
time in minutes

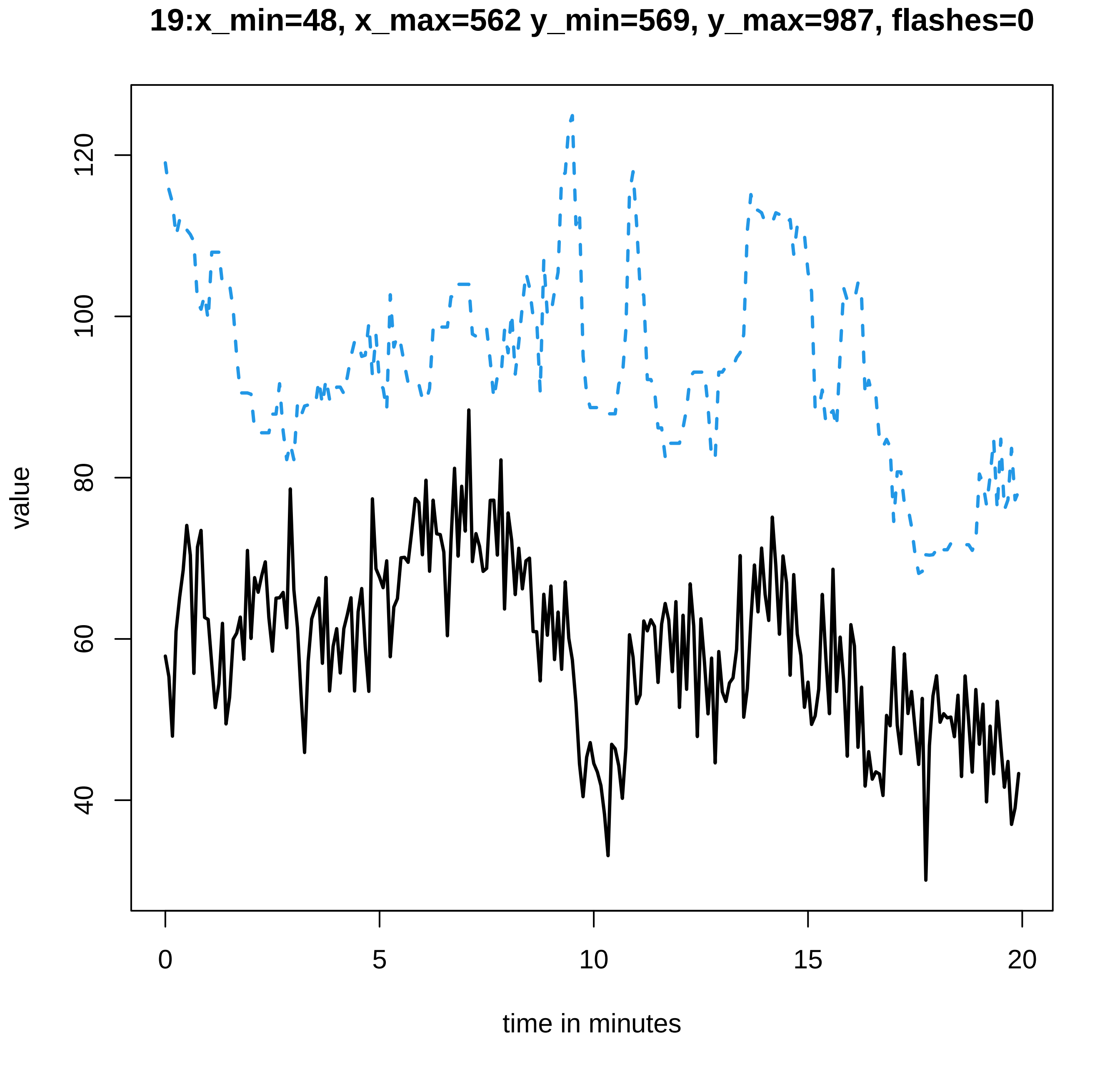
19:x_min=48, x_max=562 y_min=569, y_max=987, flashes=0
value
120
100
80
60
40
0
5
10
15
20
time in minutes
