## Supplementary data Calcium spiking for "LysM Receptor Proteins are Required for Ectomycorrhizal Symbiosis in Poplar": profiles-spiking-15.pdf

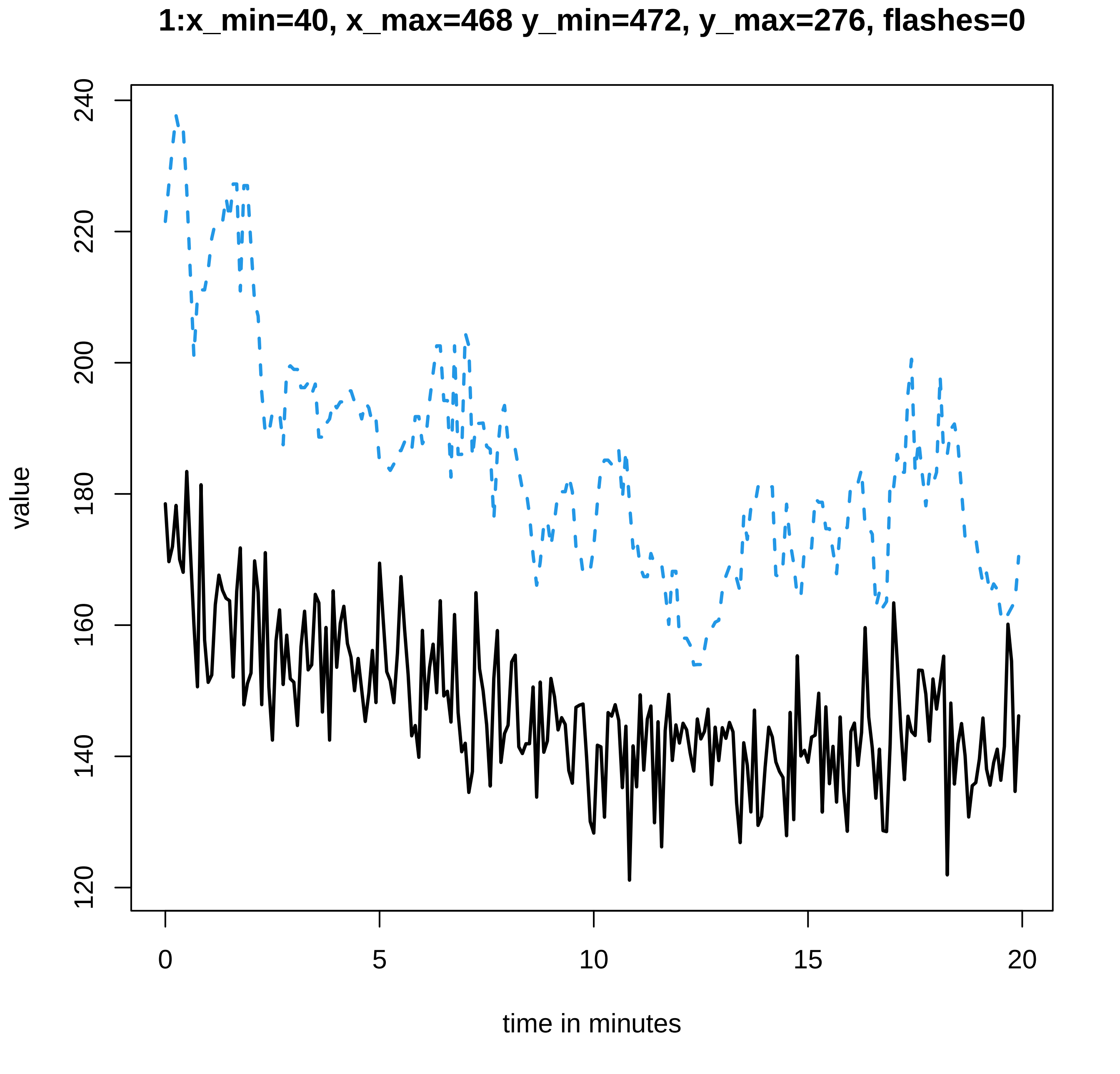

1:x_min=40, x_max=468 y_min=472, y_max=276, flashes=0
value
240
220
200
180
160
140
120
0
5
10
15
20
time in minutes

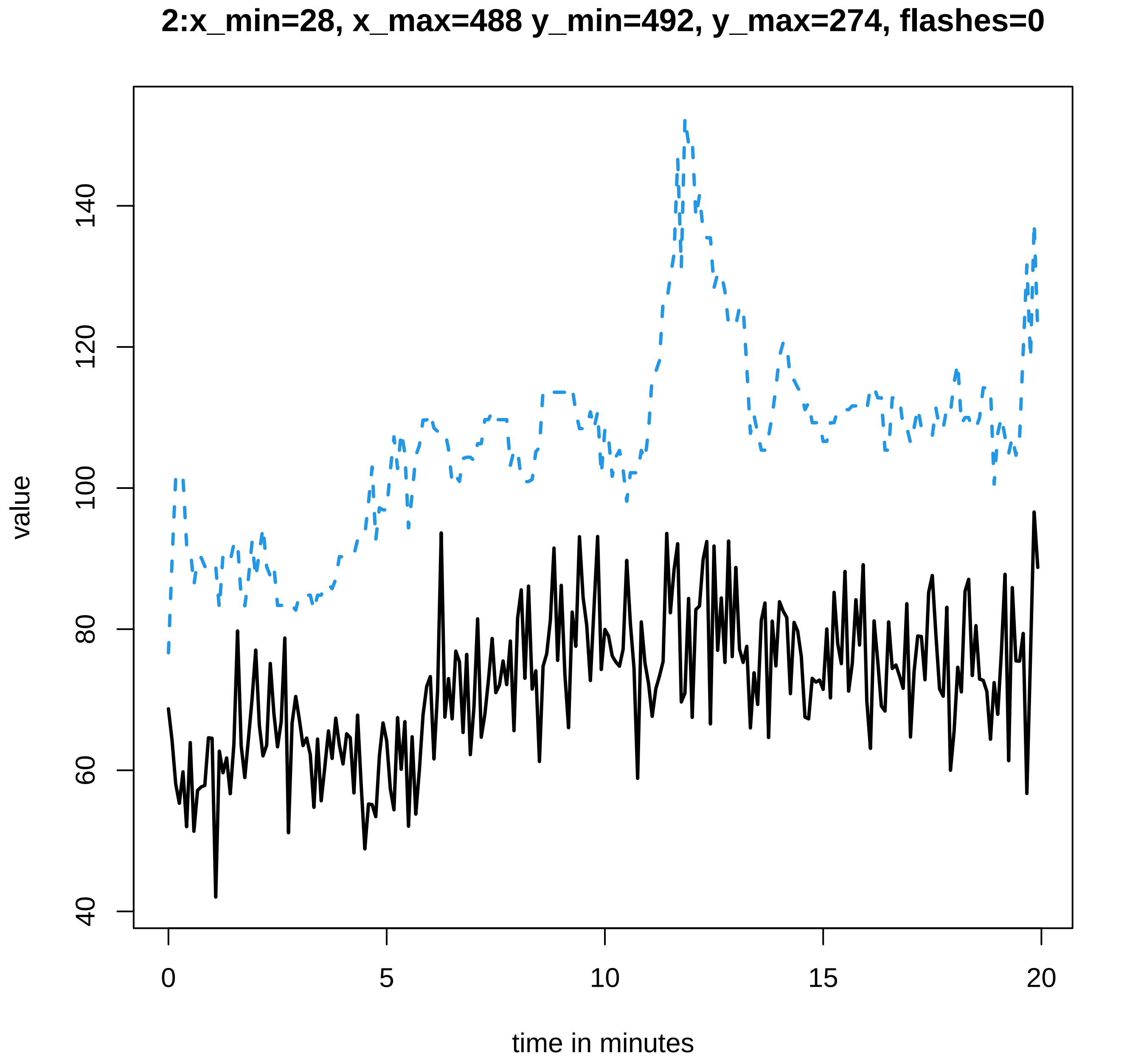

2:x_min=28, x_max=488 y_min=492, y_max=274, flashes=0
value
40
60
80
100
120
140
0
5
10
15
20
time in minutes

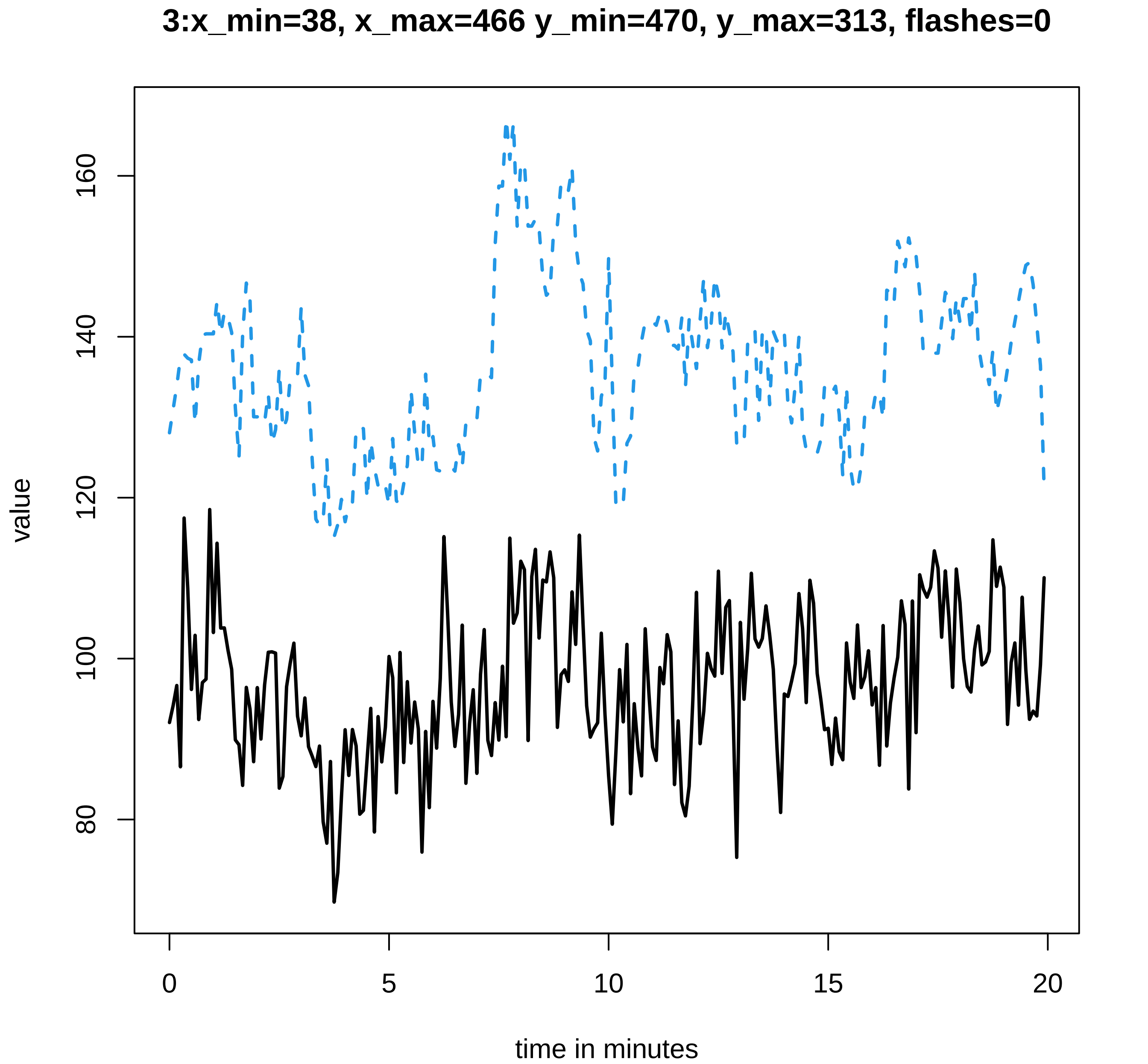

3:x_min=38, x_max=466 y_min=470, y_max=313, flashes=0
value
160
140
120
100
80
0
5
10
15
20
time in minutes

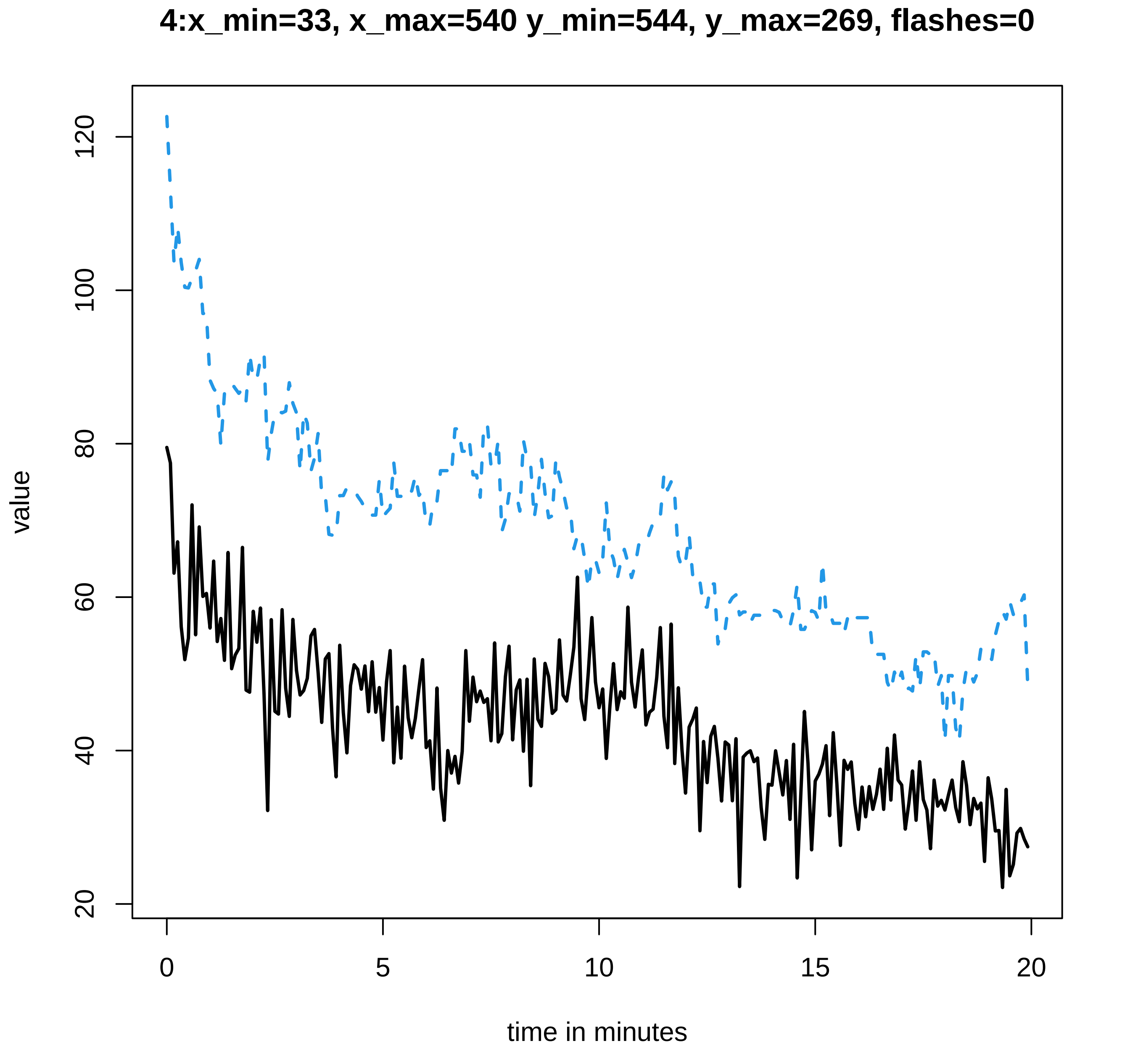
4:x_min=33, x_max=540 y_min=544, y_max=269, flashes=0
value
time in minutes
0
5
10
15
20
20
40
60
80
100
120

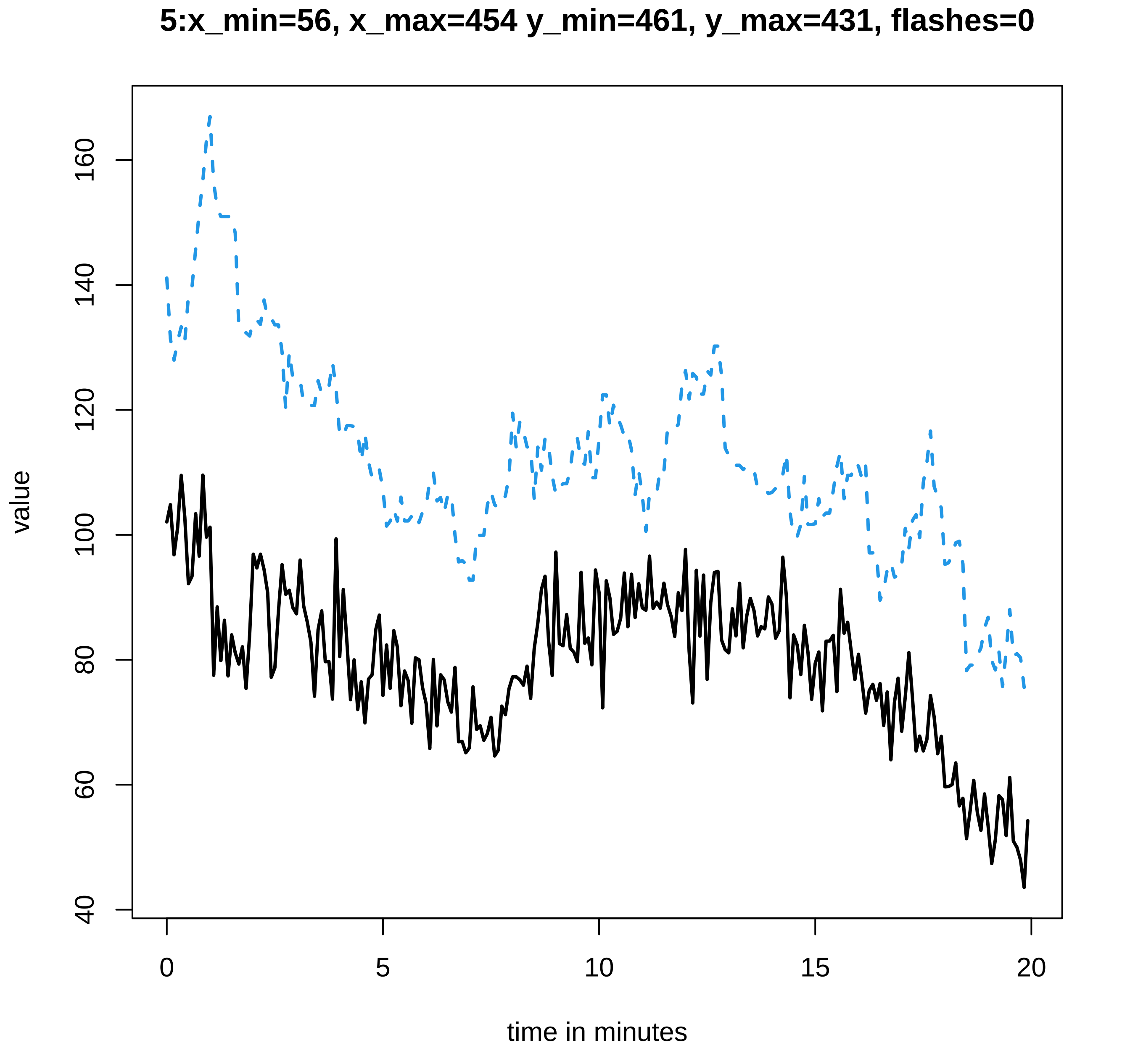

5:x_min=56, x_max=454 y_min=461, y_max=431, flashes=0
value
40
60
80
100
120
140
160
0
5
10
15
20
time in minutes

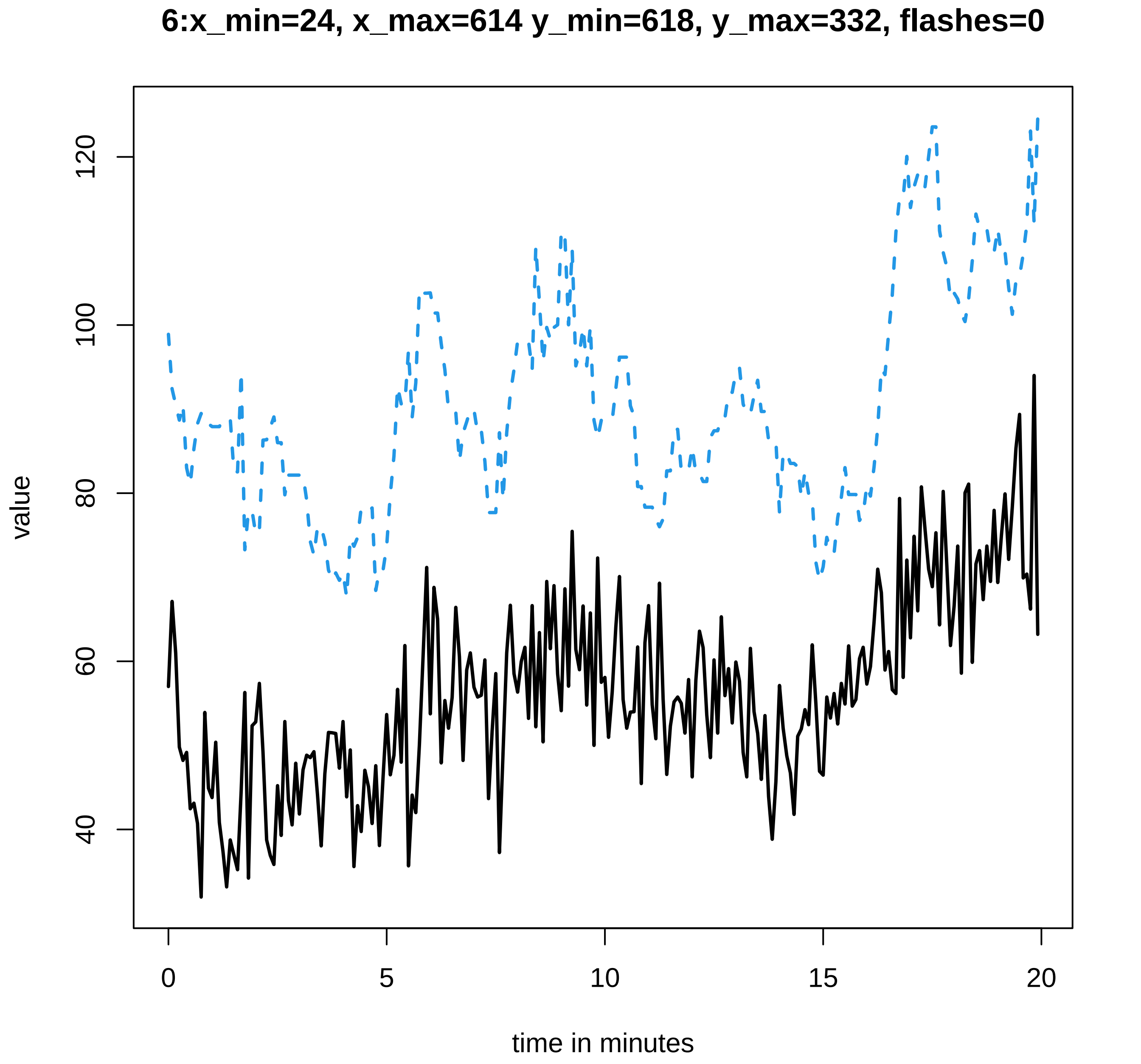
6:x_min=24, x_max=614 y_min=618, y_max=332, flashes=0
value
120
100
80
60
40
0
5
10
15
20
time in minutes

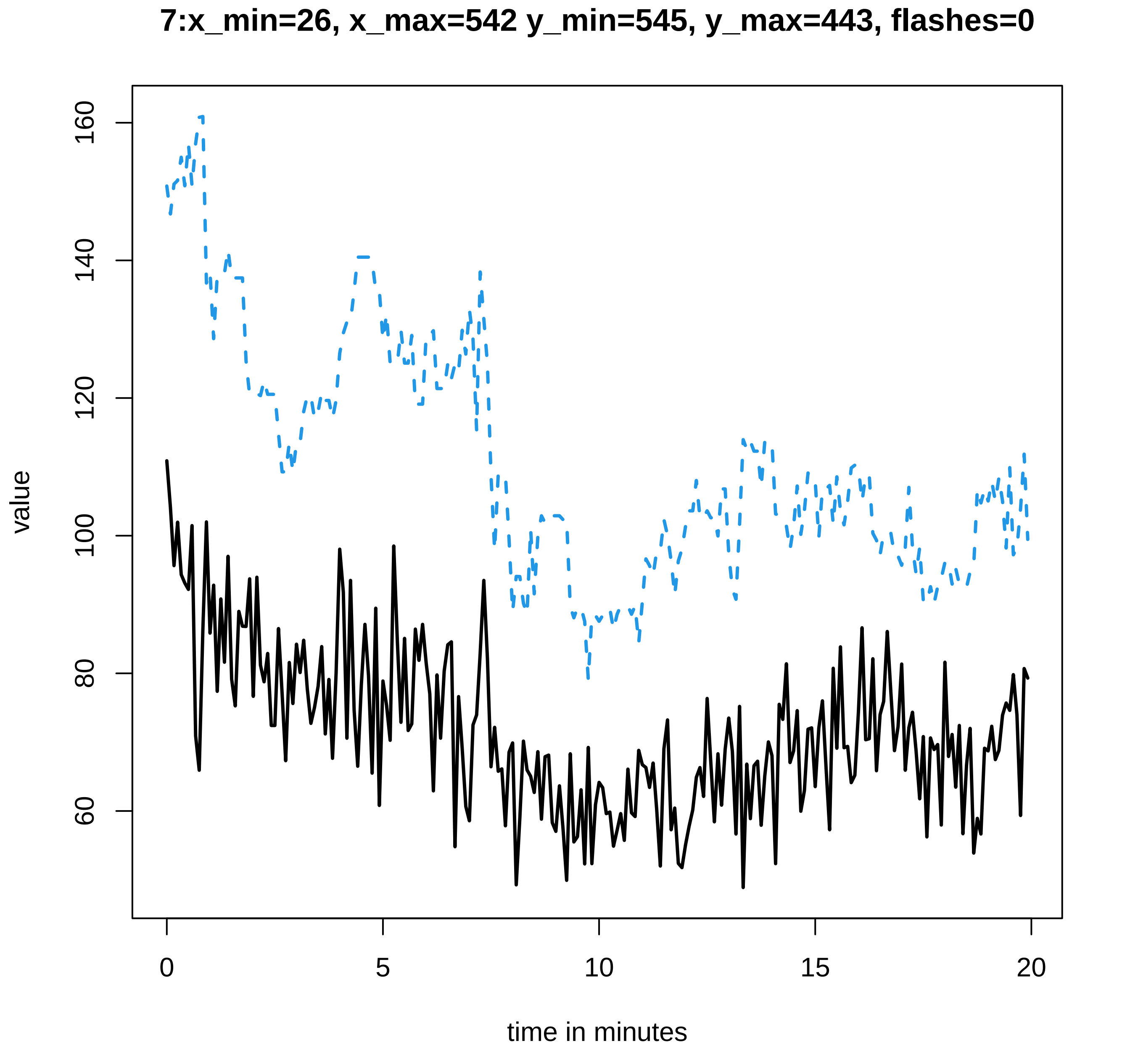
7:x_min=26, x_max=542 y_min=545, y_max=443, flashes=0
value
160
140
120
100
80
60
0
5
10
15
20
time in minutes

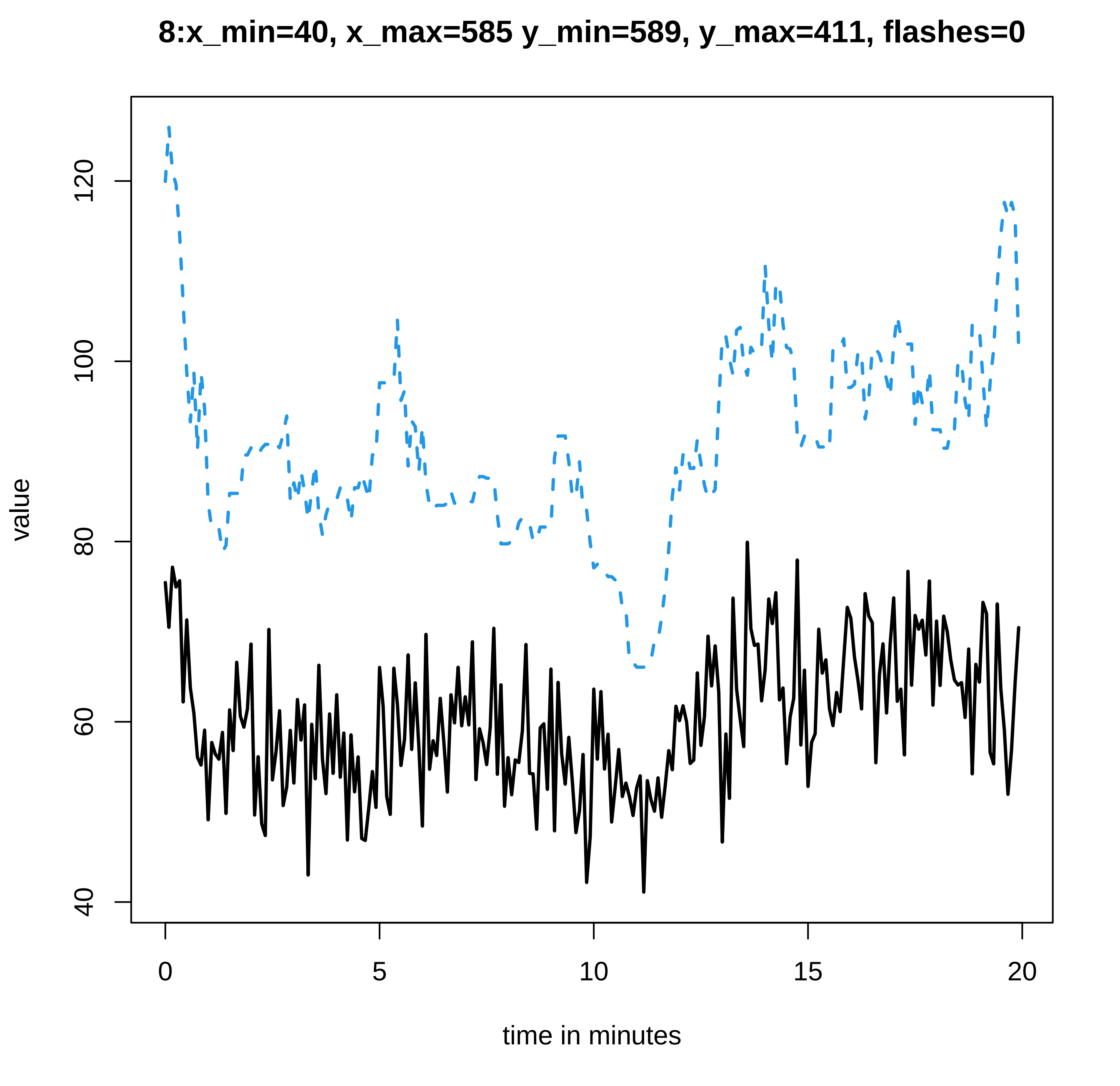
8:x_min=40, x_max=585 y_min=589, y_max=411, flashes=0
value
40
60
80
100
120
0
5
10
15
20
time in minutes

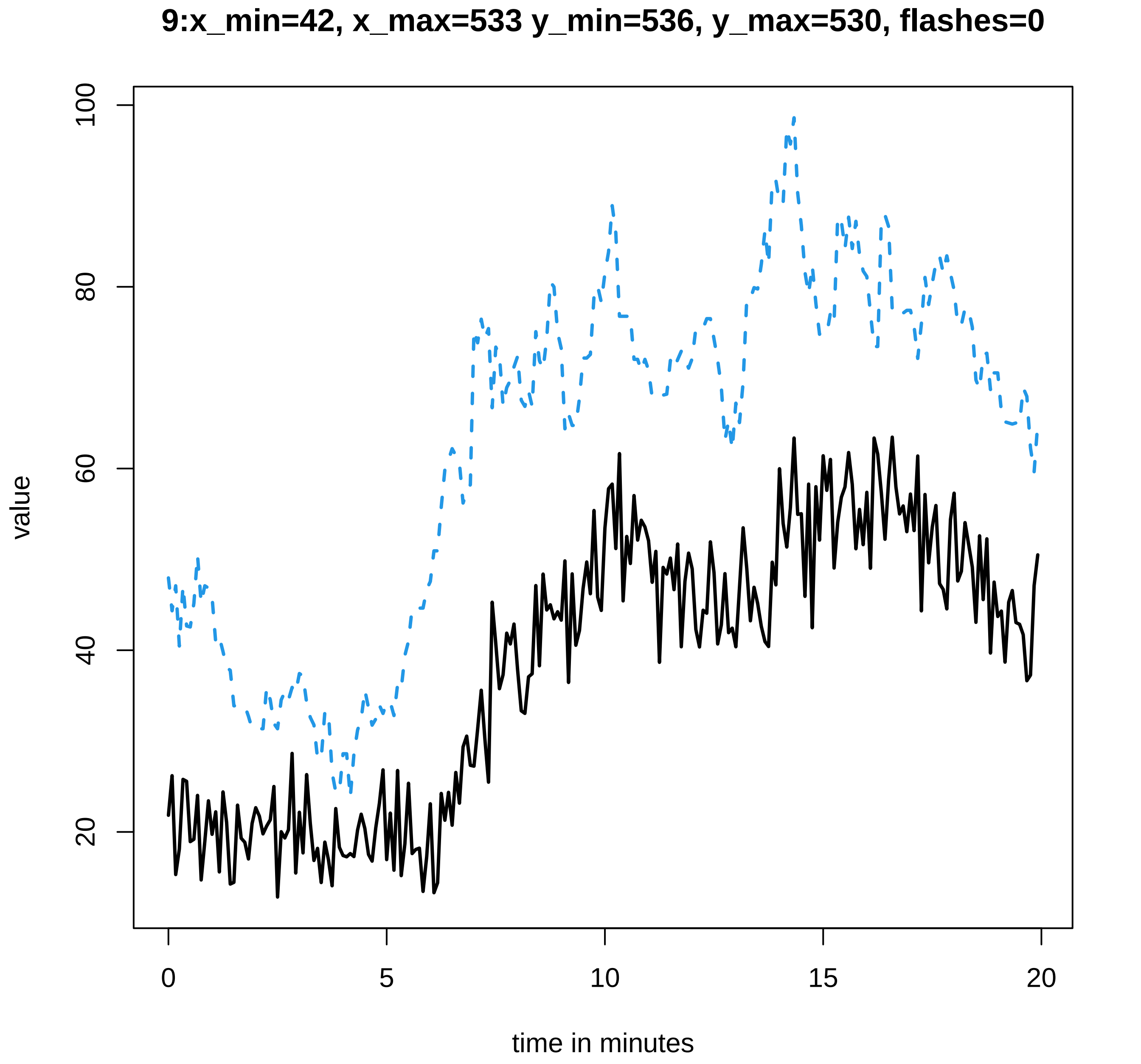
9:x_min=42, x_max=533 y_min=536, y_max=530, flashes=0
value
100
80
60
40
20
0
5
10
15
20
time in minutes

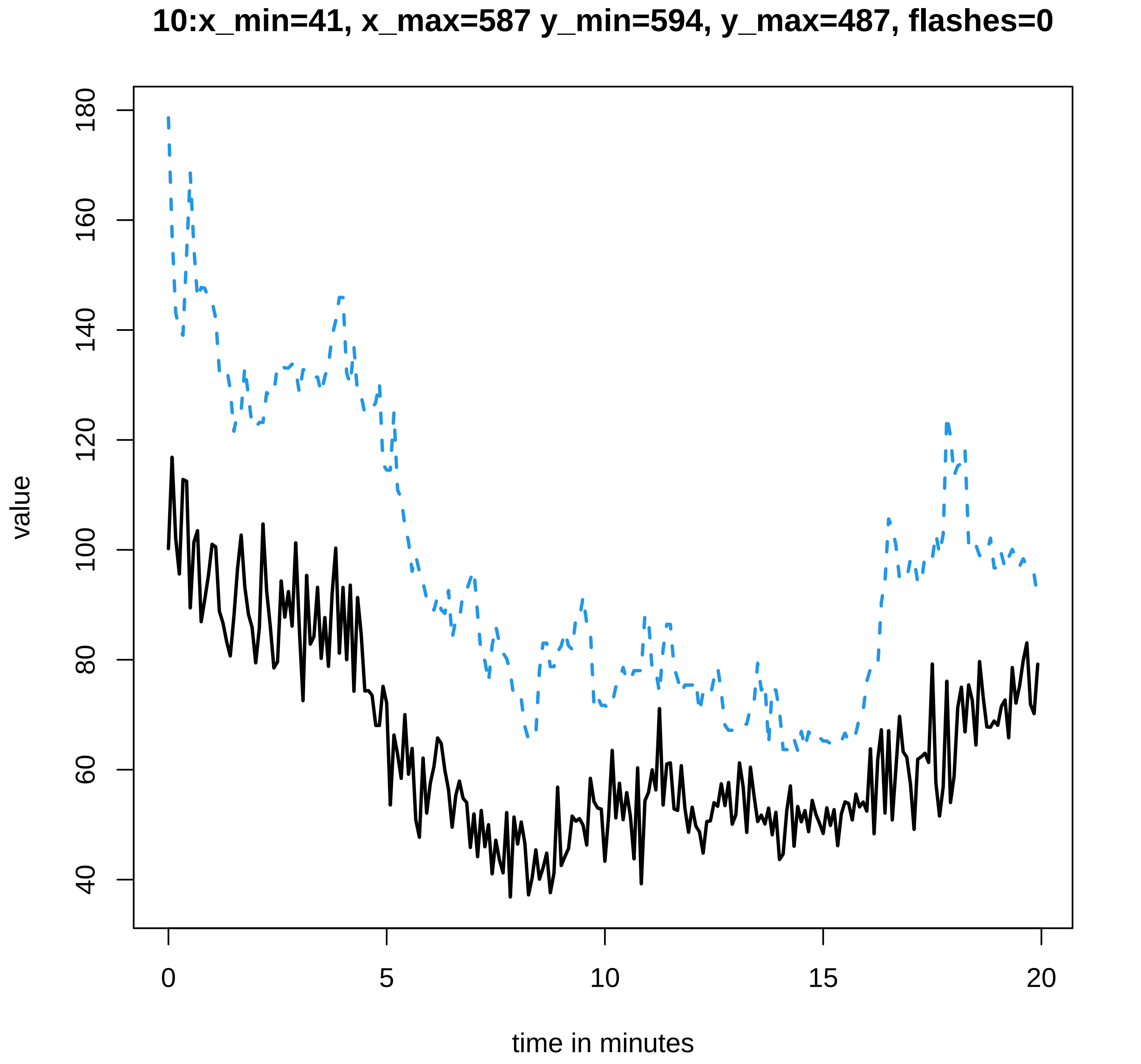
10:x_min=41, x_max=587 y_min=594, y_max=487, flashes=0
value
180
160
140
120
100
80
60
40
0
5
10
15
20
time in minutes

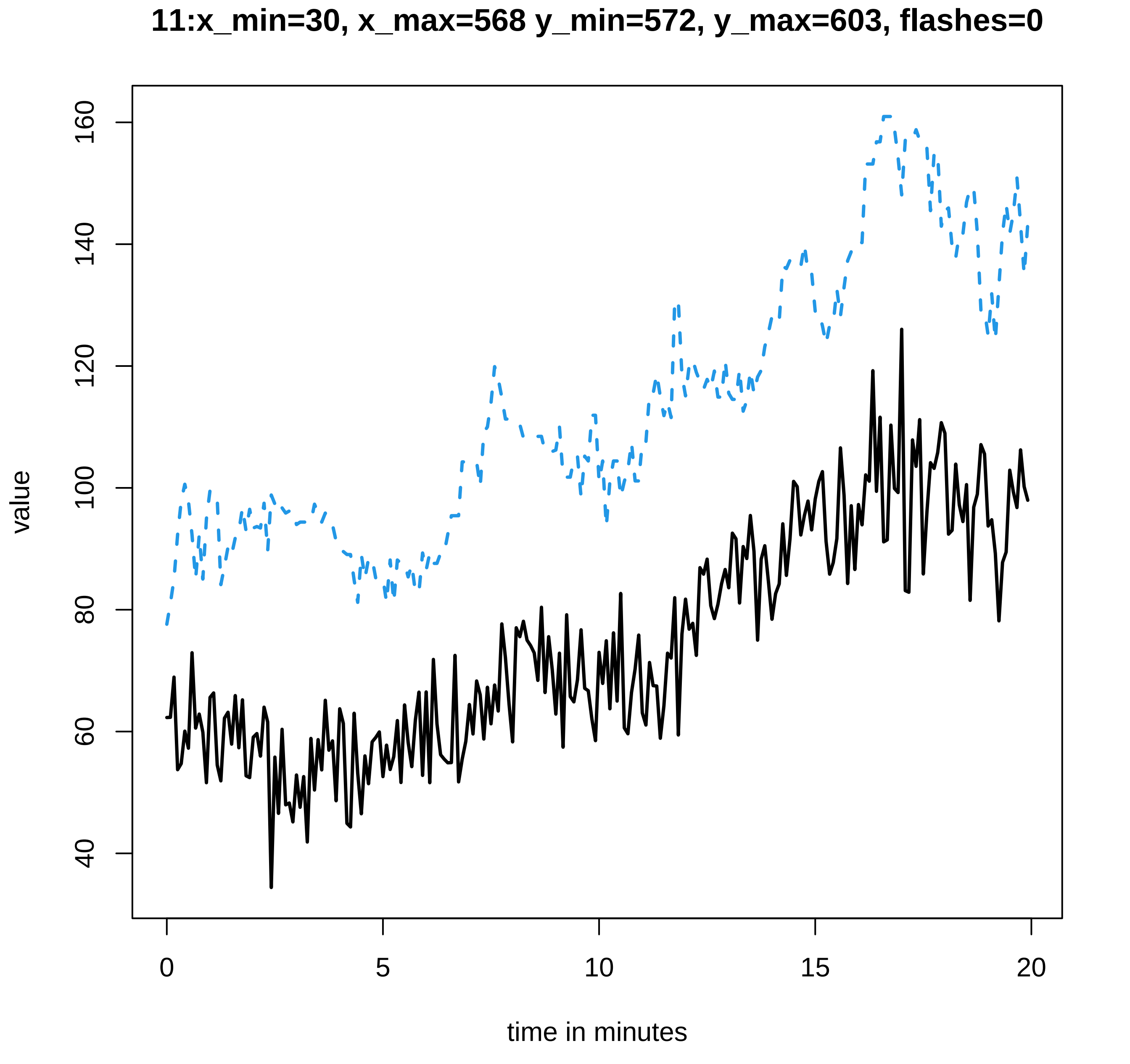

11:x_min=30, x_max=568 y_min=572, y_max=603, flashes=0
value
time in minutes
0
5
10
15
20
40
60
80
100
120
140
160

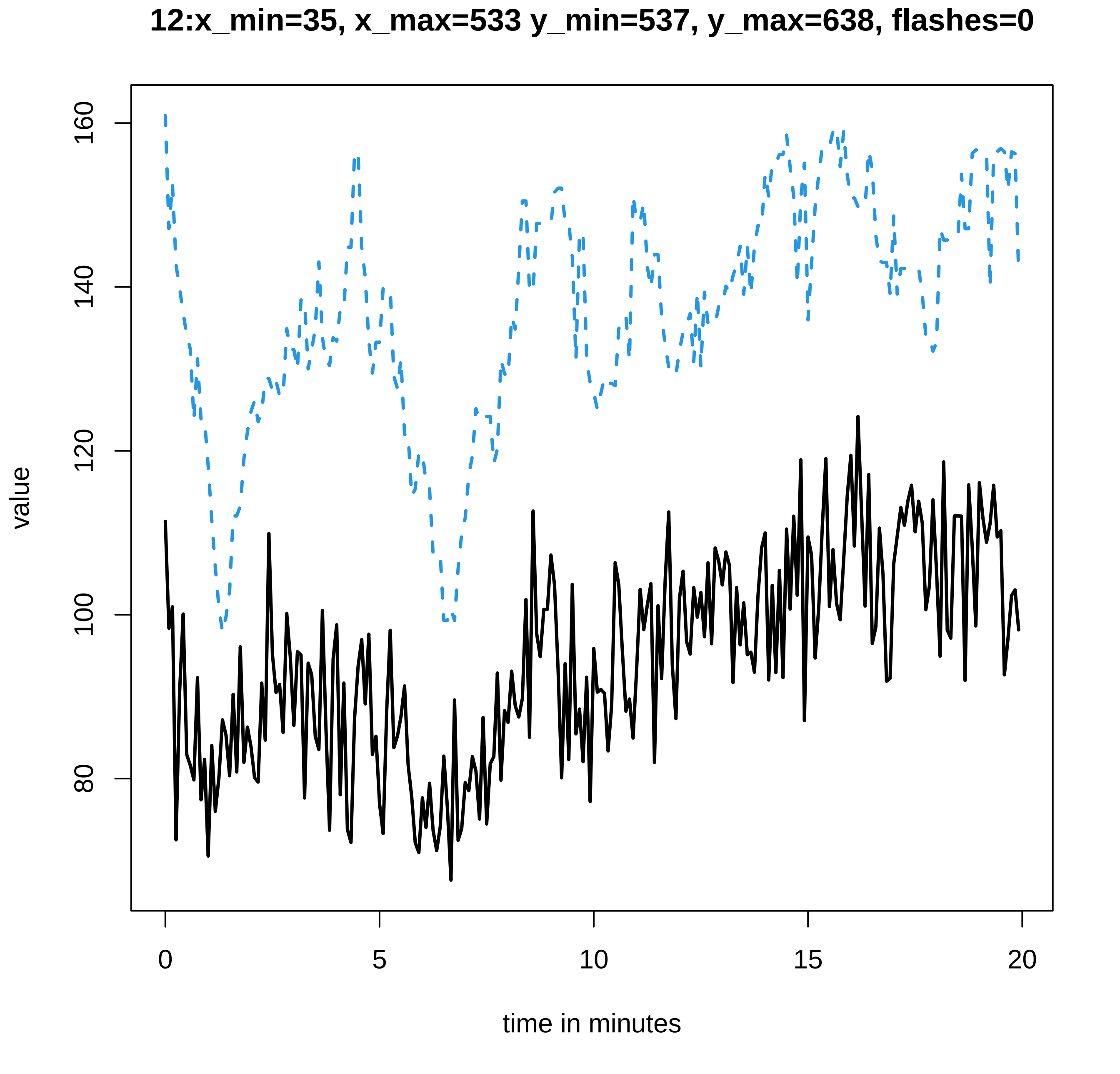
12:x_min=35, x_max=533 y_min=537, y_max=638, flashes=0
value
160
140
120
100
80
0
5
10
15
20
time in minutes

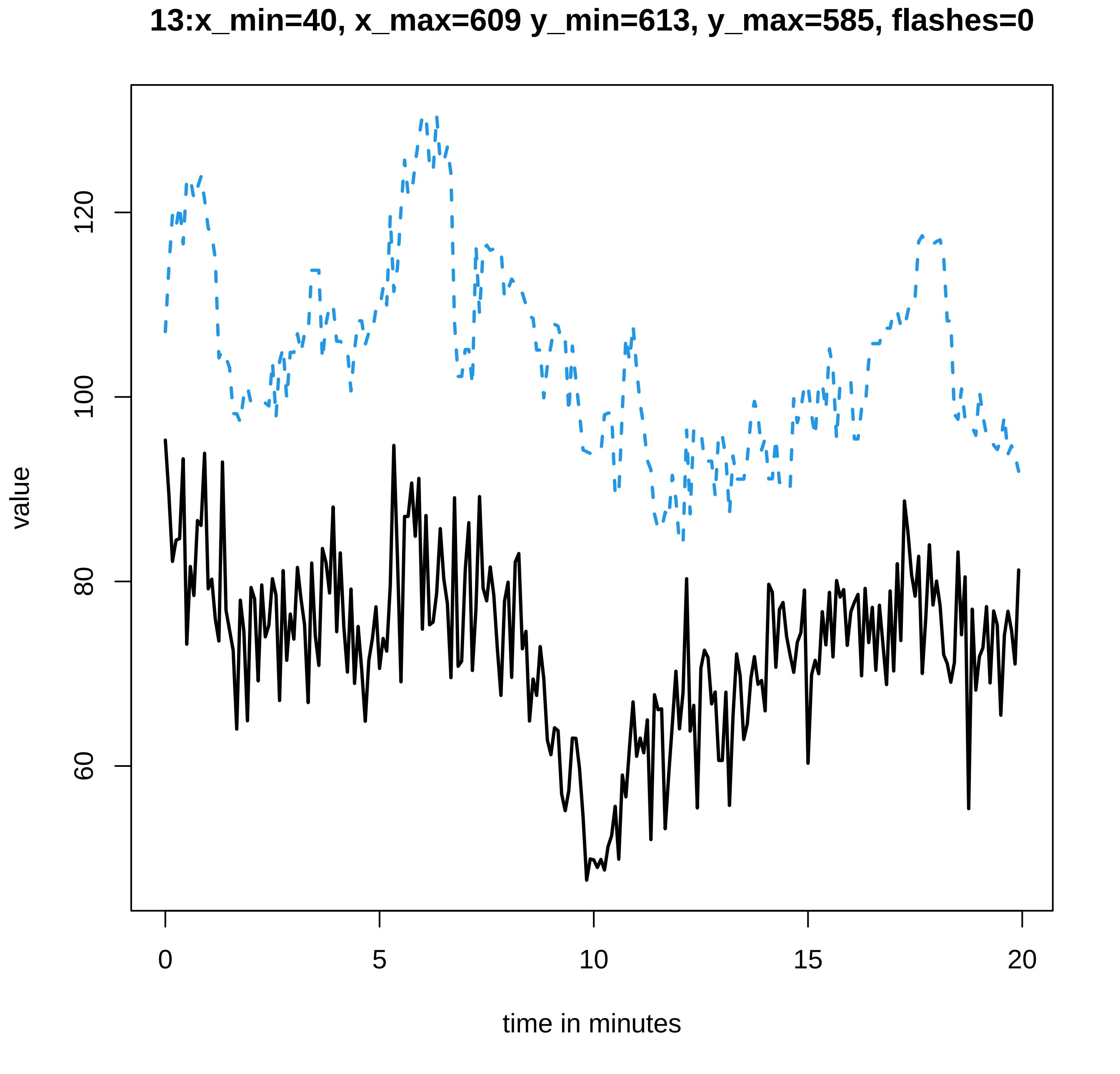
13:x_min=40, x_max=609 y_min=613, y_max=585, flashes=0
value
time in minutes
0
5
10
15
20
60
80
100
120

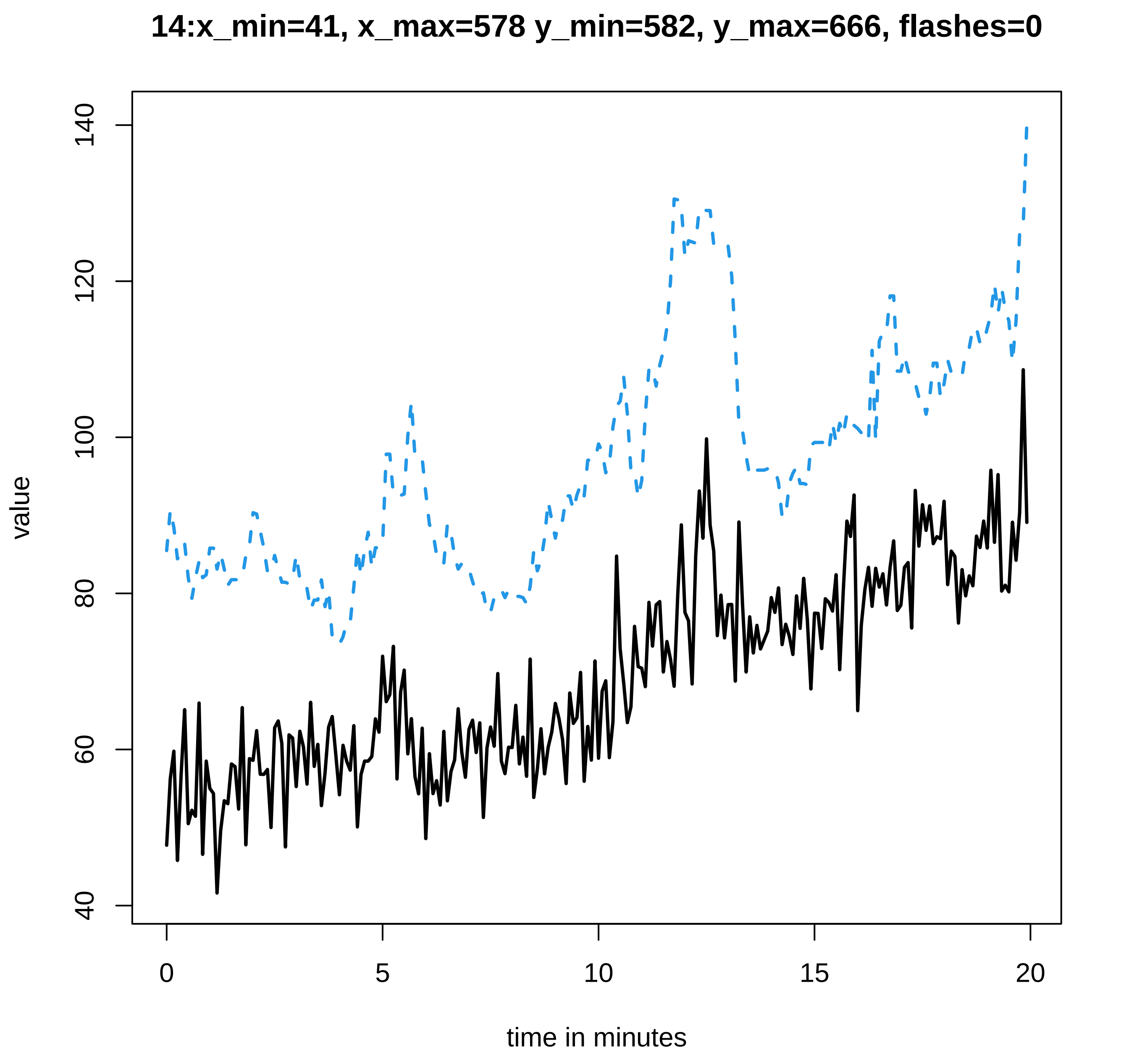
14:x_min=41, x_max=578 y_min=582, y_max=666, flashes=0
value
140
120
100
80
60
40
0
5
10
15
20
time in minutes

**15:x_min=35, x_max=485 y_min=491, y_max=894, flashes=0**

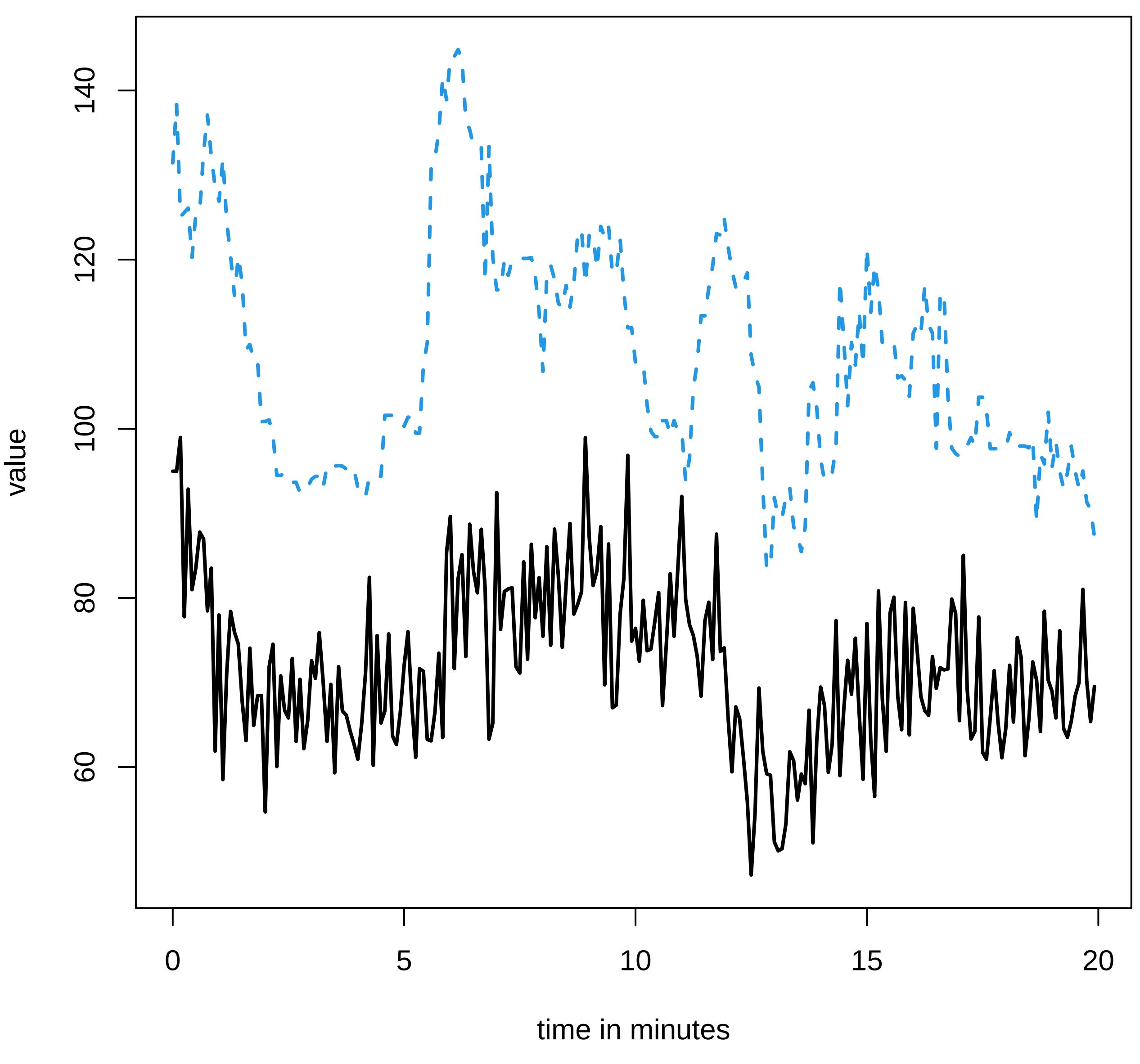

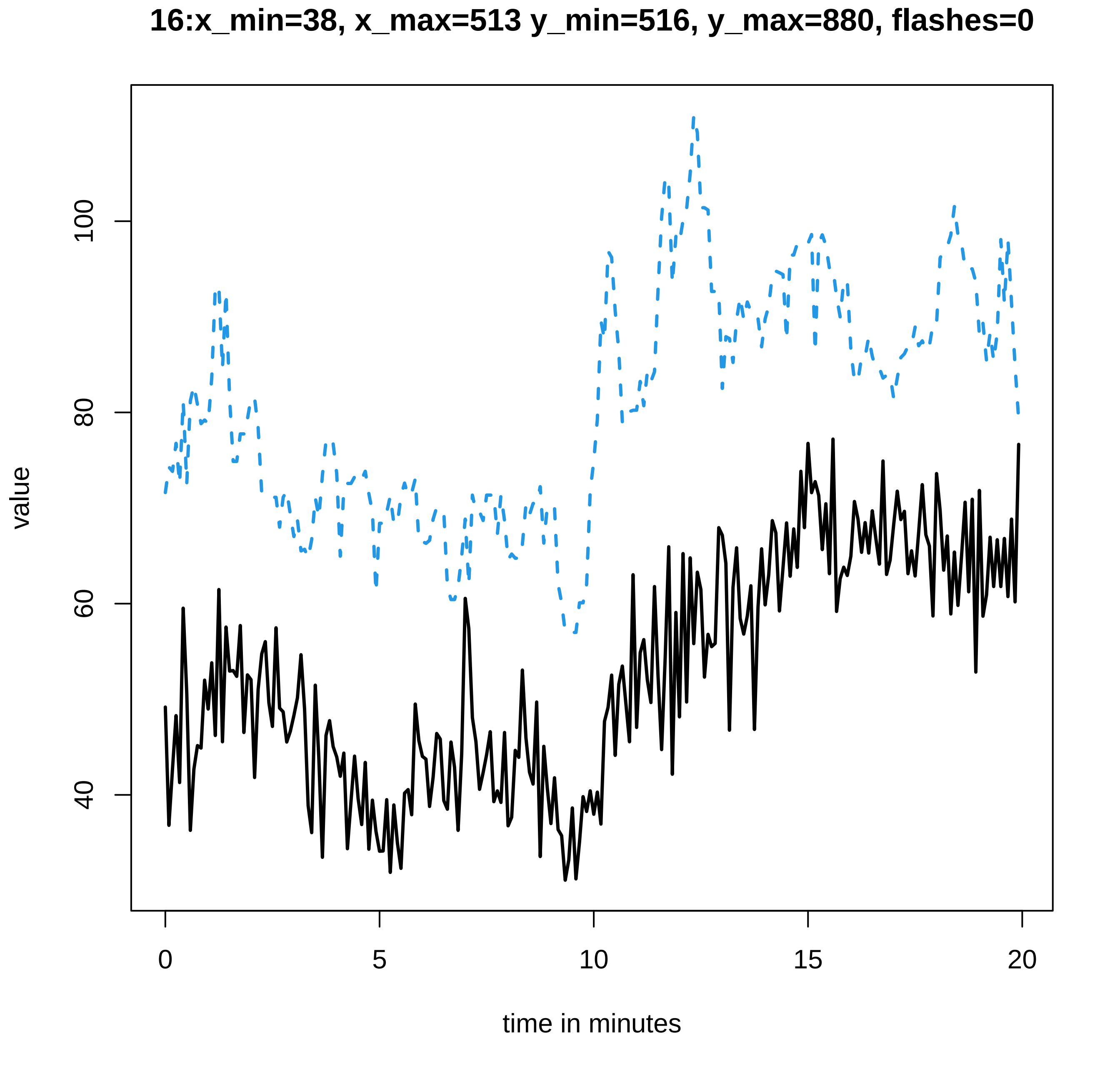
16:x_min=38, x_max=513 y_min=516, y_max=880, flashes=0
value
40
60
80
100
0
5
10
15
20
time in minutes
