## Supplementary data Calcium spiking for "LysM Receptor Proteins are Required for Ectomycorrhizal Symbiosis in Poplar": profiles-spiking-16.pdf

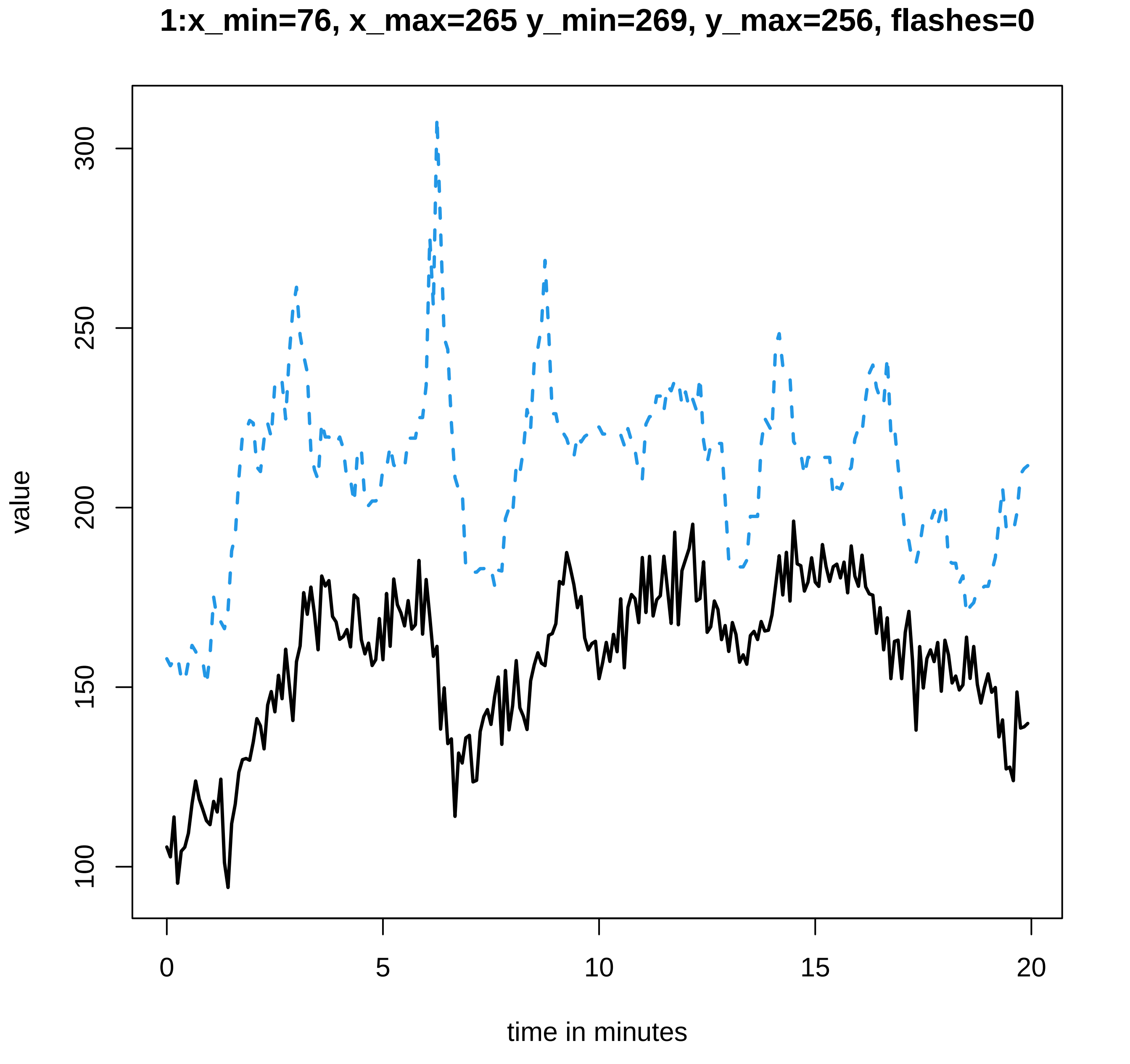

1:x_min=76, x_max=265 y_min=269, y_max=256, flashes=0
value
300
250
200
150
100
0
5
10
15
20
time in minutes

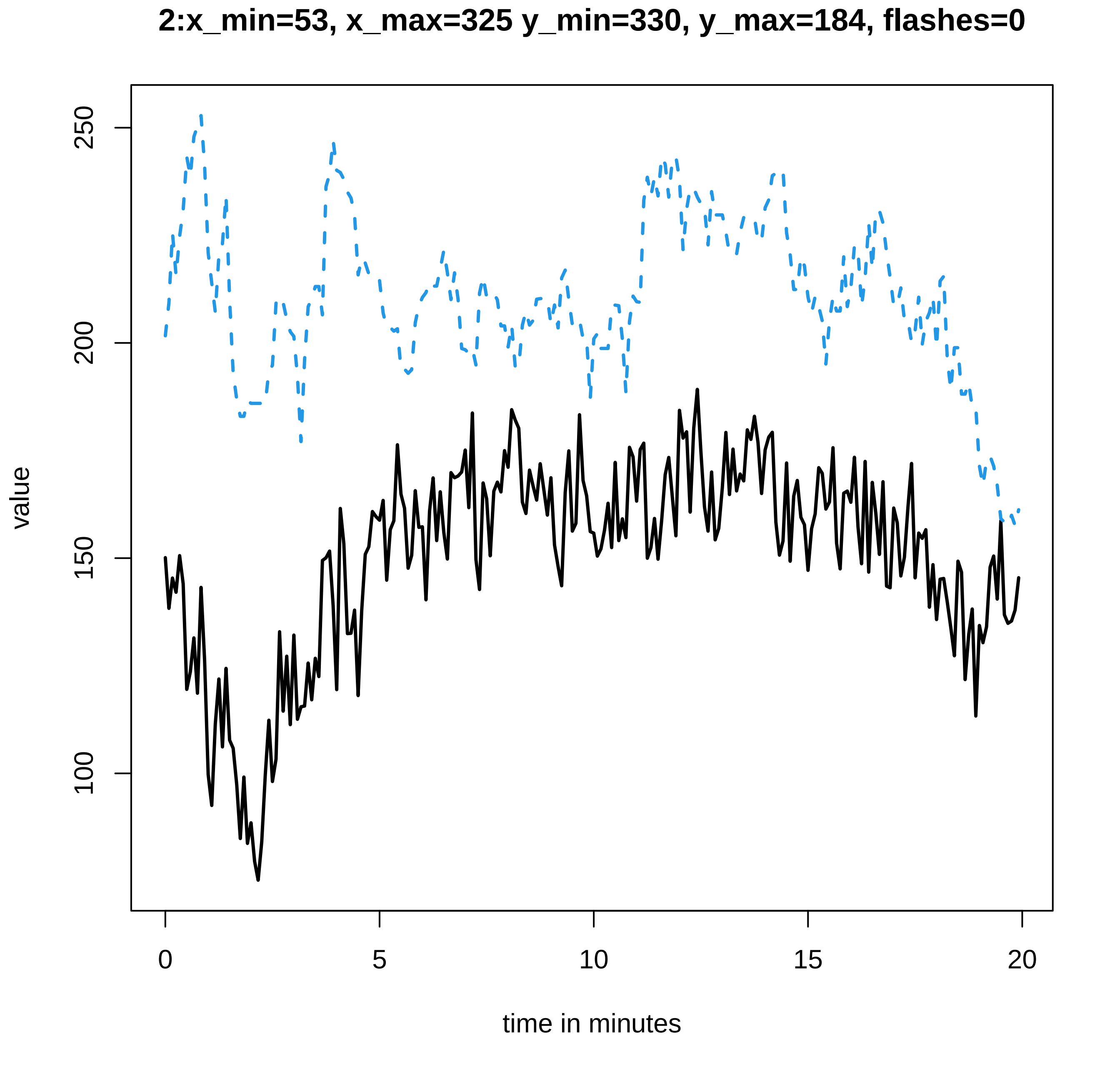
2:x_min=53, x_max=325 y_min=330, y_max=184, flashes=0
value
250
200
150
100
0
5
10
15
20
time in minutes

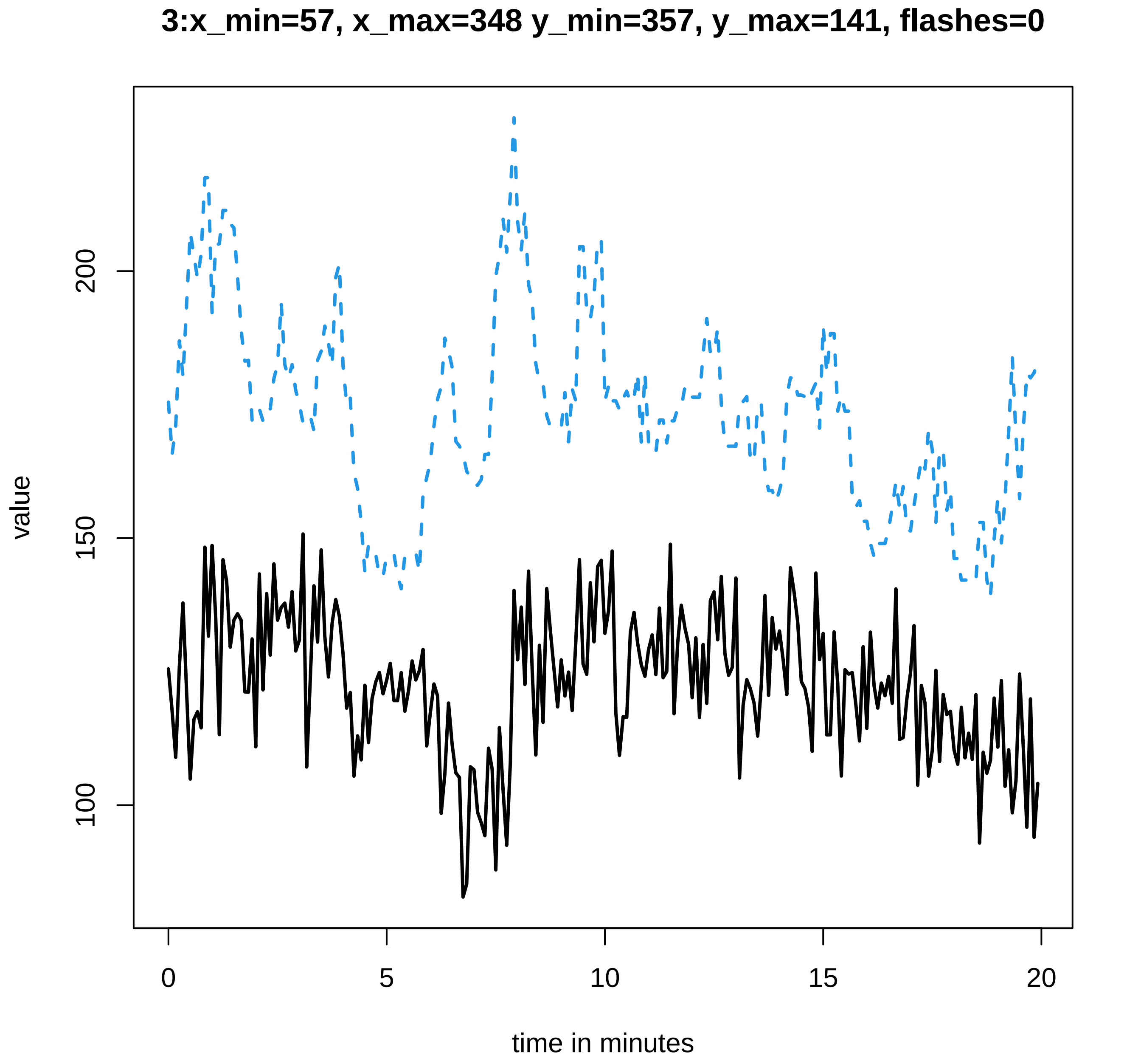
3:x_min=57, x_max=348 y_min=357, y_max=141, flashes=0
value
200
150
100
0
5
10
15
20
time in minutes

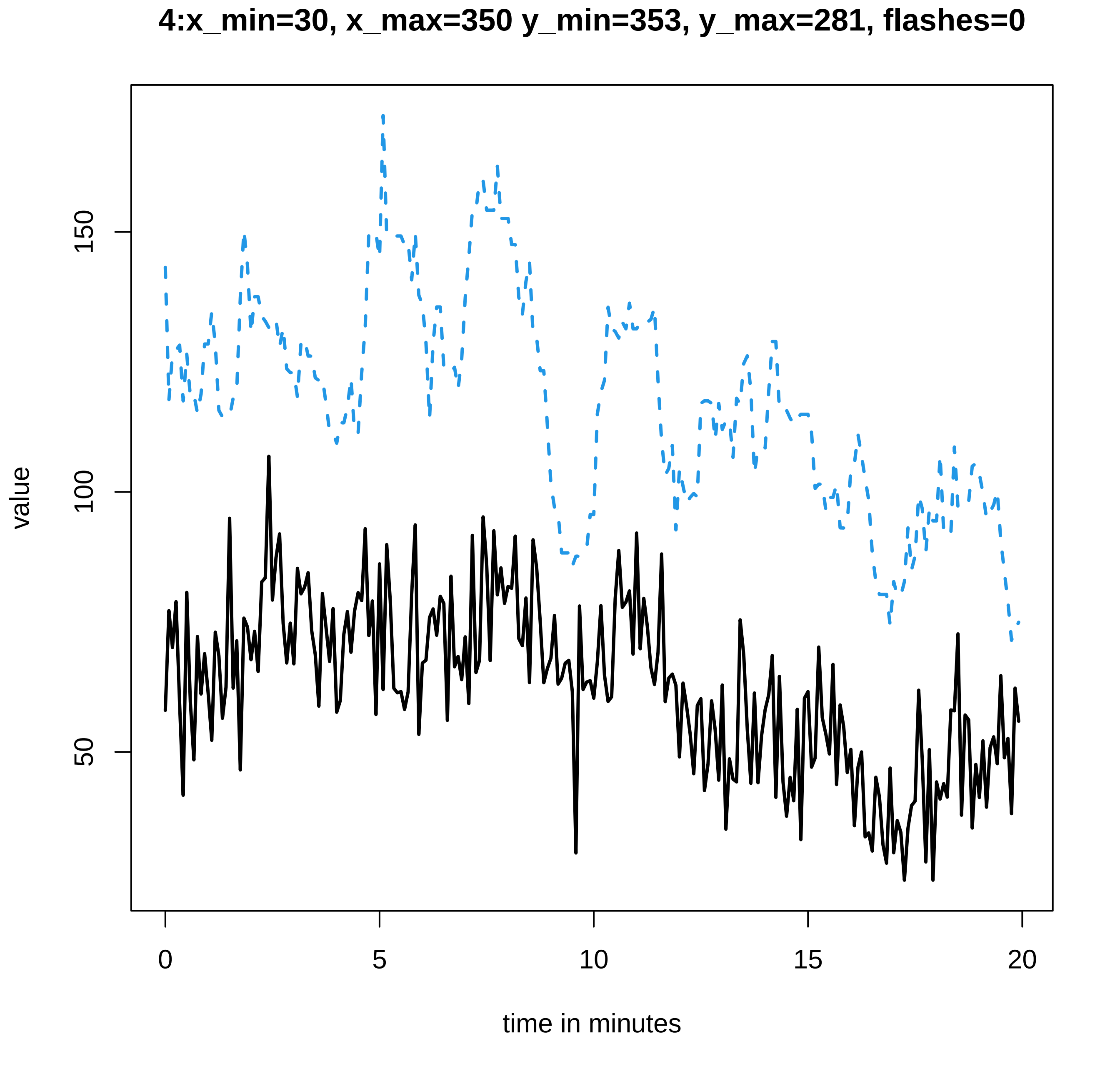
4:x_min=30, x_max=350 y_min=353, y_max=281, flashes=0
value
150
100
50
0
5
10
15
20
time in minutes

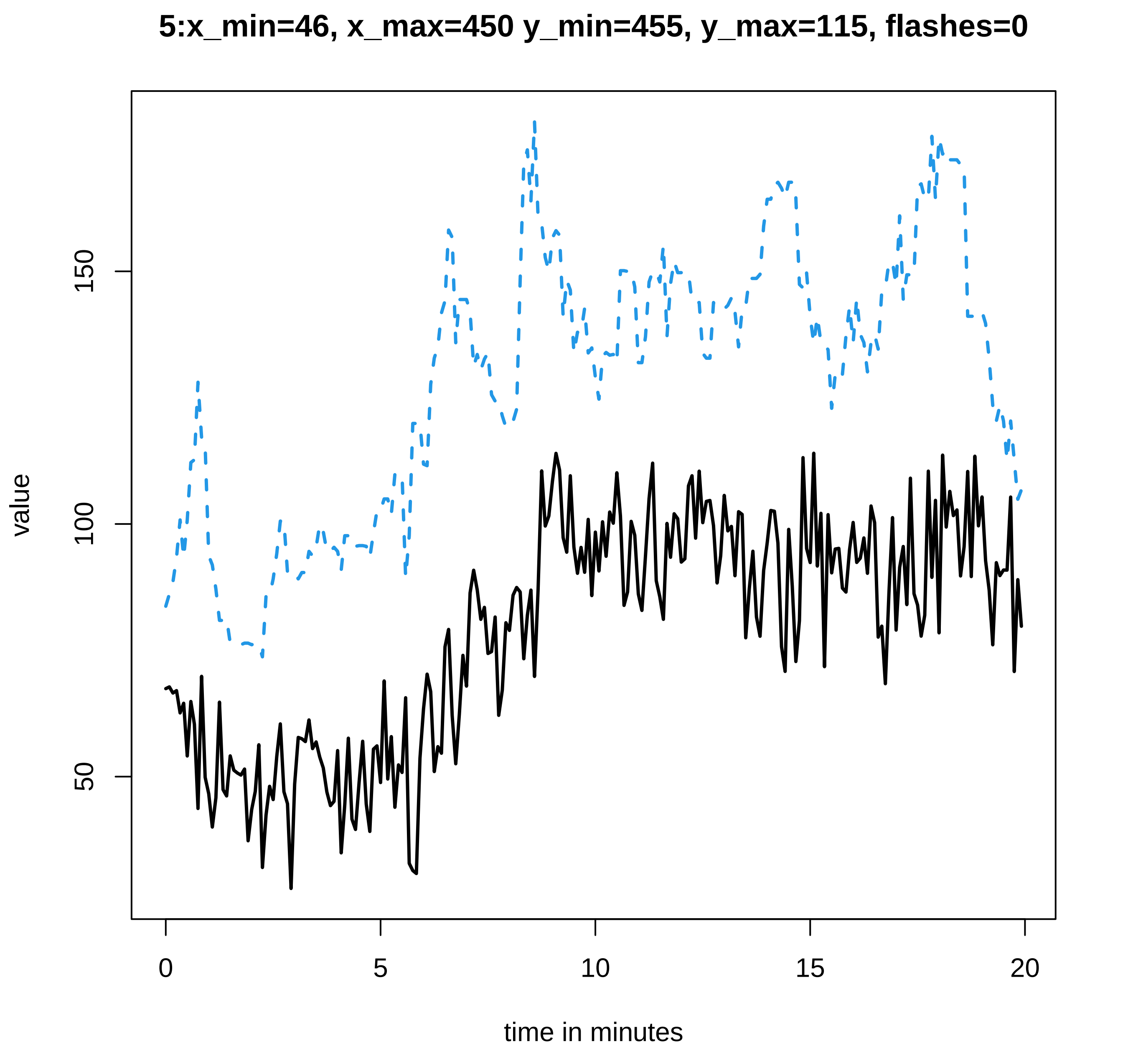
5:x_min=46, x_max=450 y_min=455, y_max=115, flashes=0
value
50
100
150
0
5
10
15
20
time in minutes

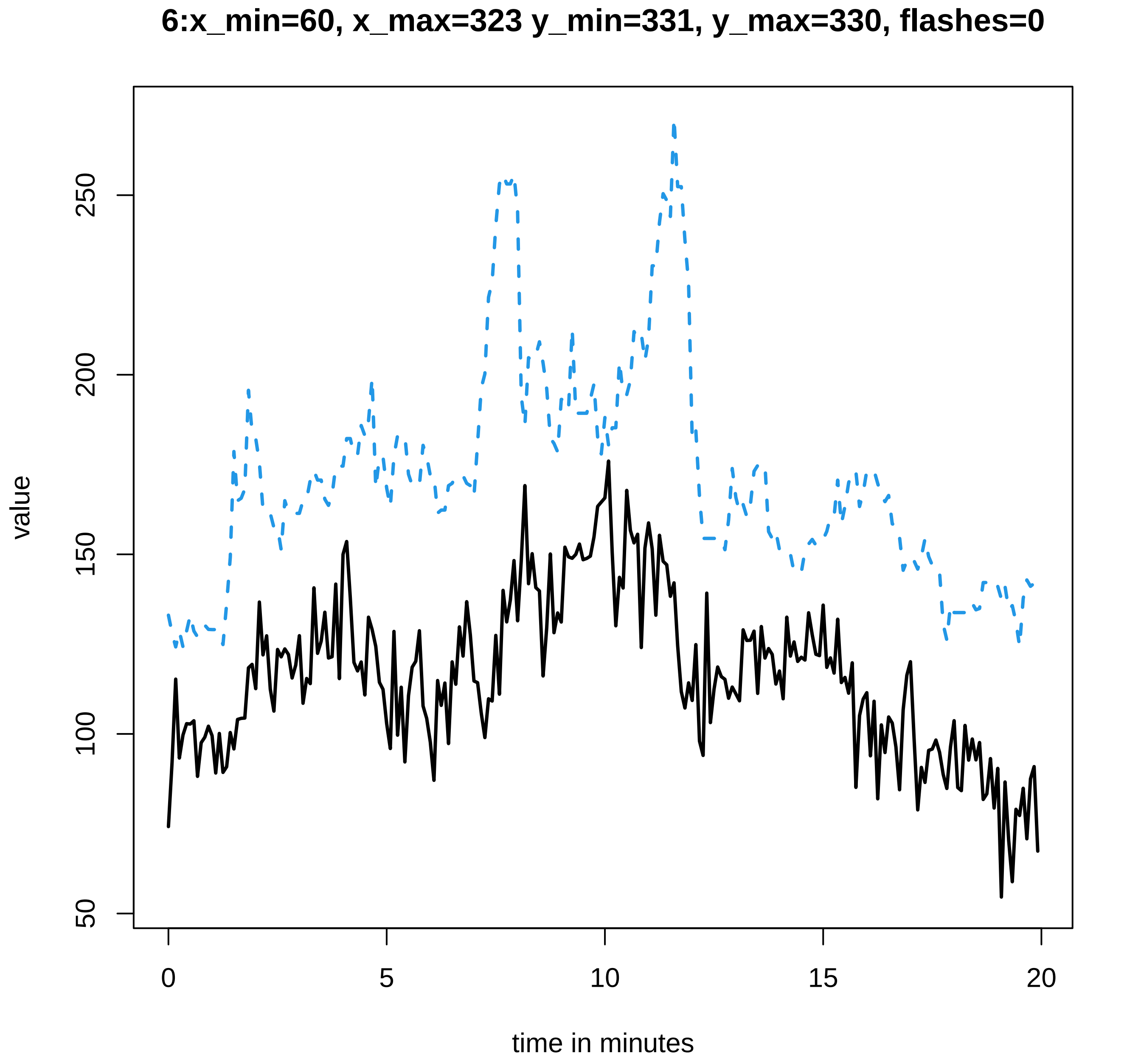
6:x_min=60, x_max=323 y_min=331, y_max=330, flashes=0
value
50
100
150
200
250
0
5
10
15
20
time in minutes

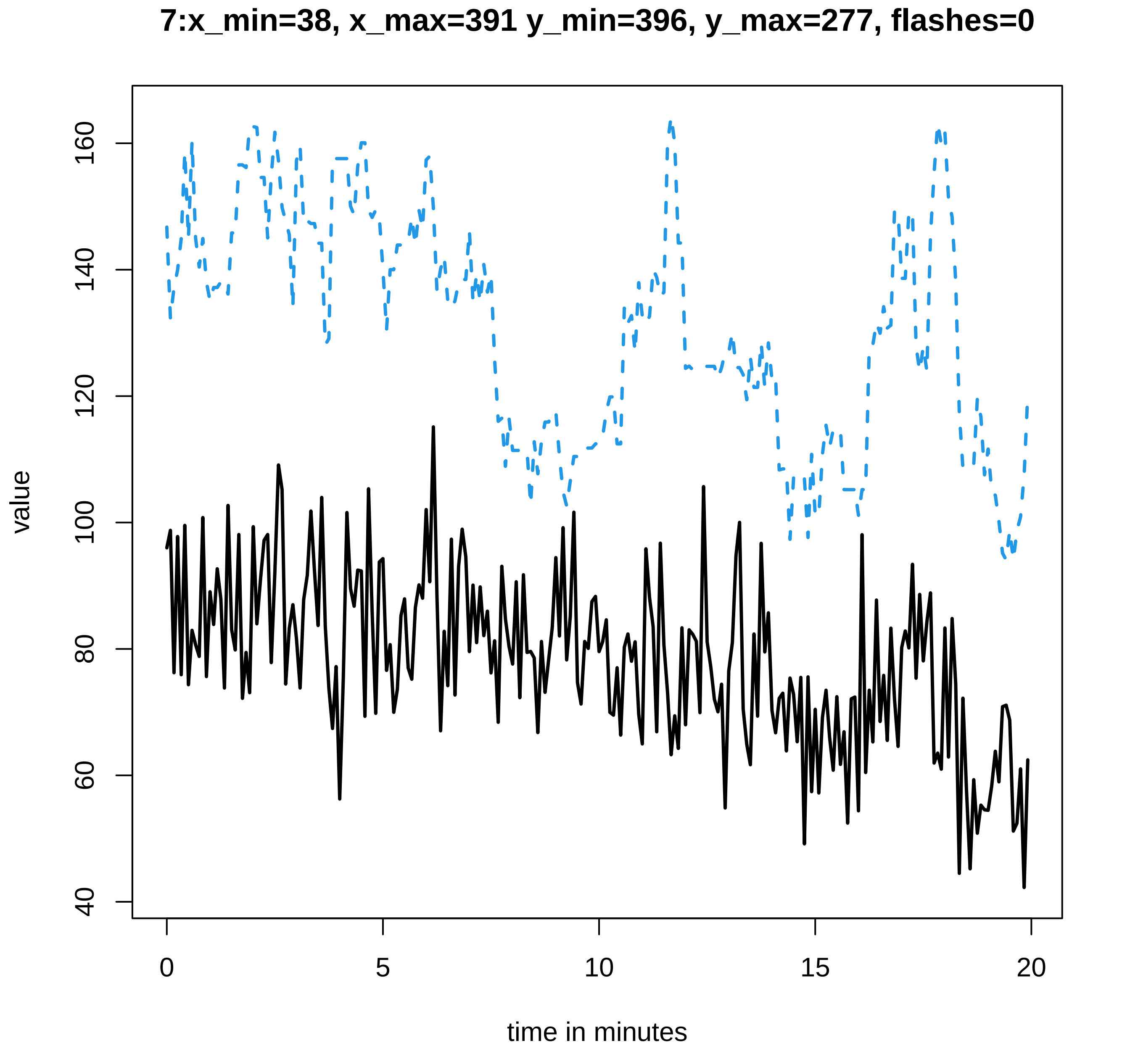
7:x_min=38, x_max=391 y_min=396, y_max=277, flashes=0
value
time in minutes
160
140
120
100
80
60
40
0
5
10
15
20

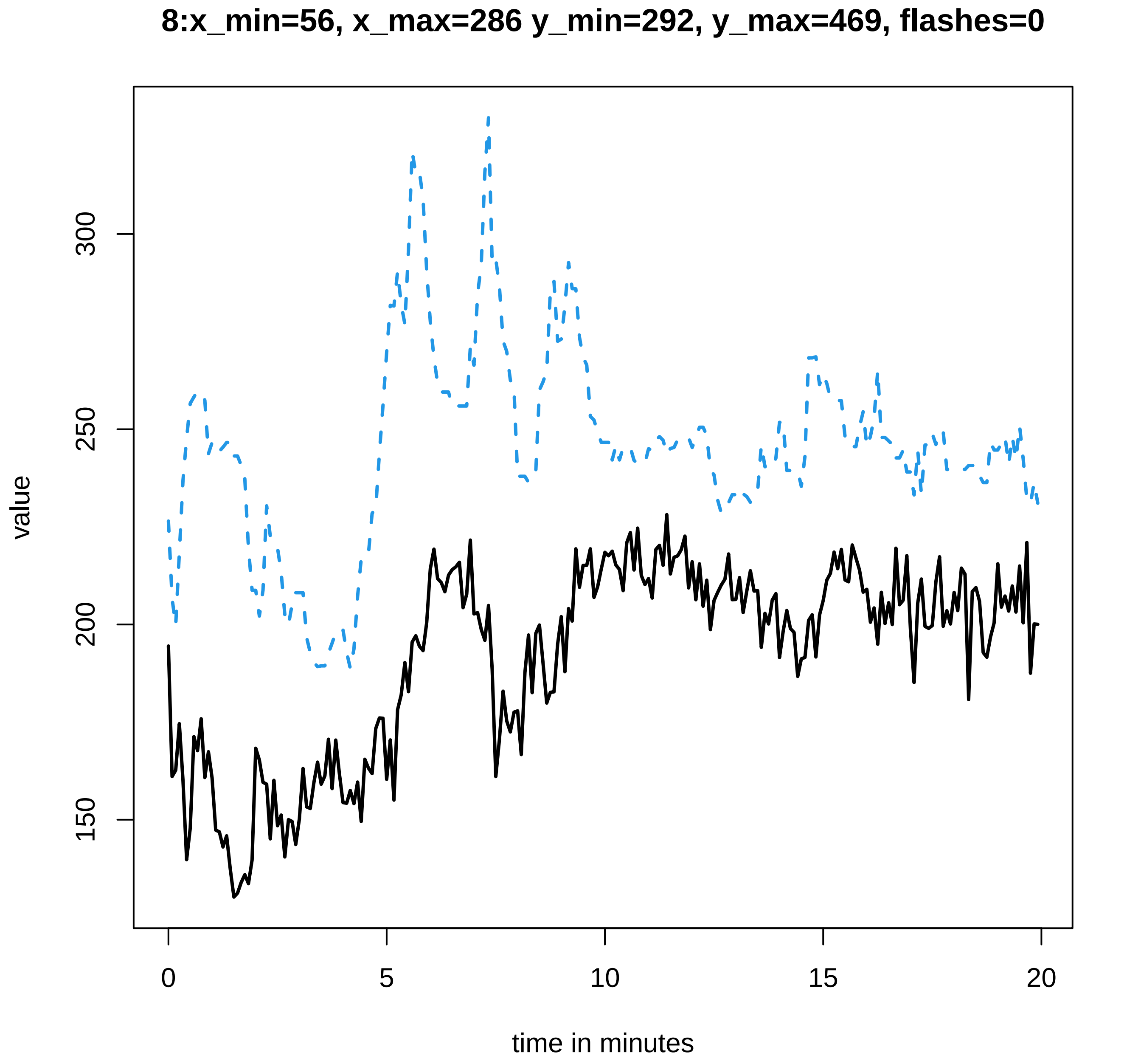

8:x_min=56, x_max=286 y_min=292, y_max=469, flashes=0
value
150
200
250
300
0
5
10
15
20
time in minutes

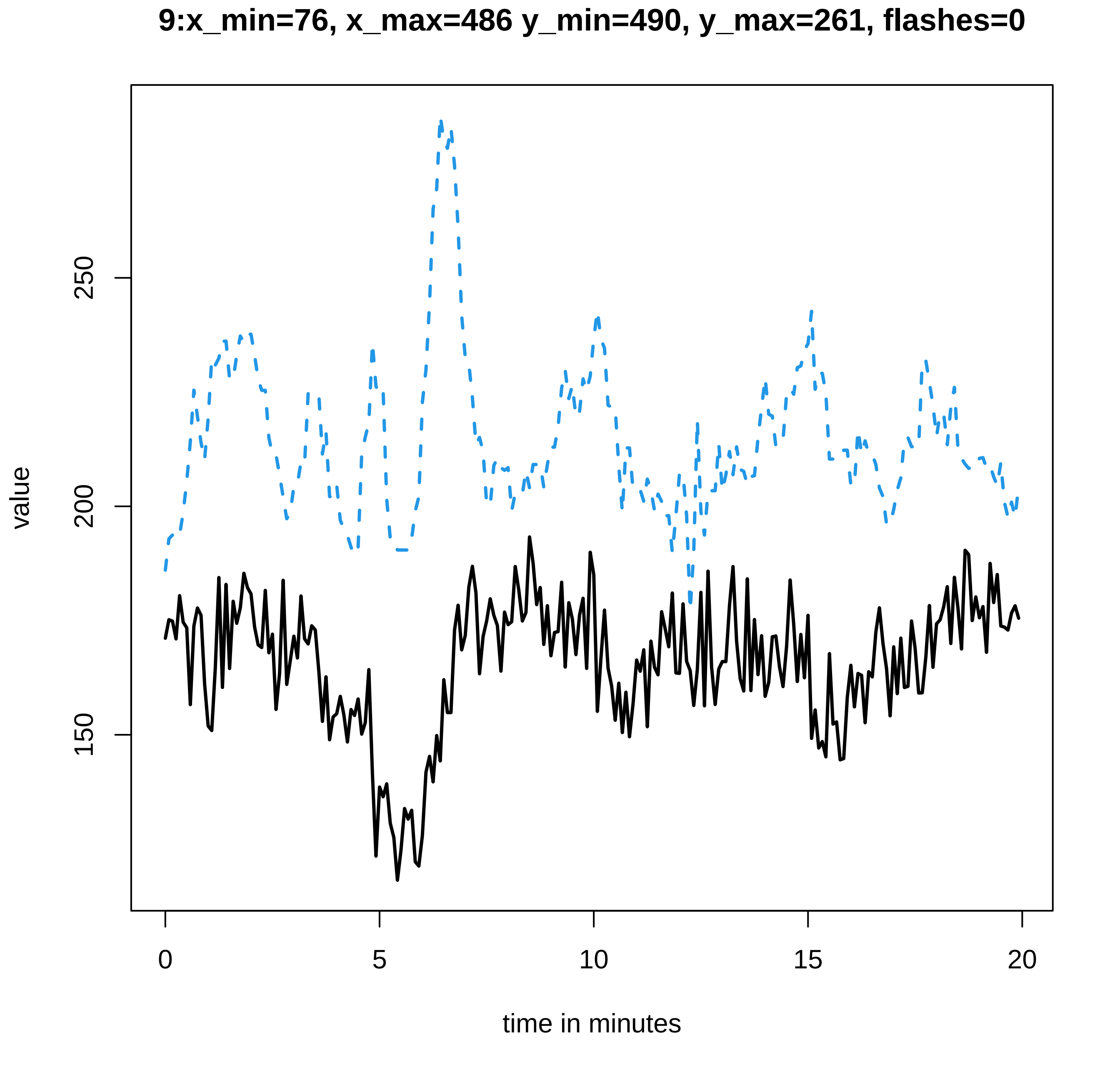
9:x_min=76, x_max=486 y_min=490, y_max=261, flashes=0
value
250
200
150
0
5
10
15
20
time in minutes

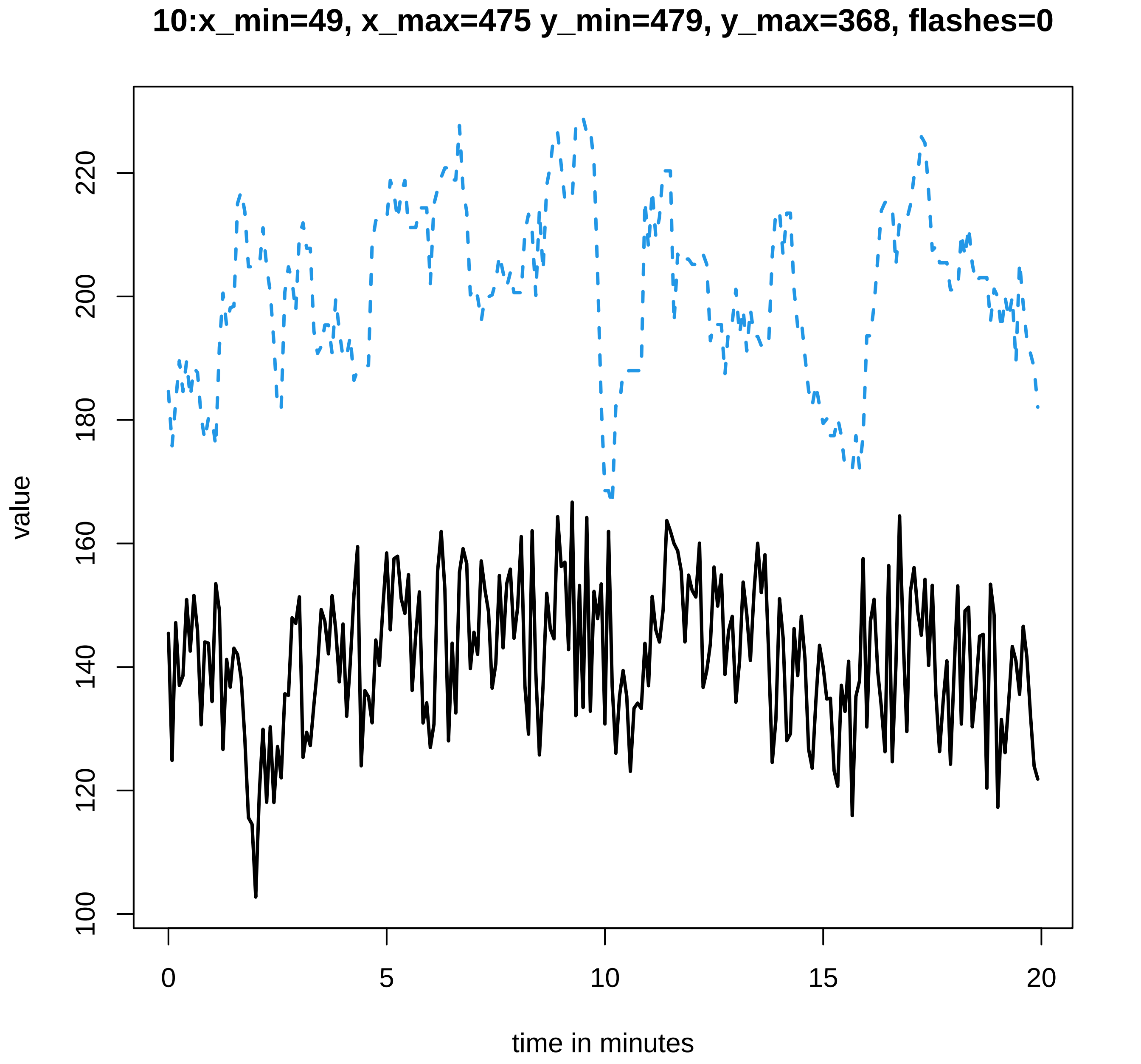
10:x_min=49, x_max=475 y_min=479, y_max=368, flashes=0
value
100
120
140
160
180
200
220
0
5
10
15
20
time in minutes

### 11:x_min=72, x_max=362 y_min=366, y_max=478, flashes=0

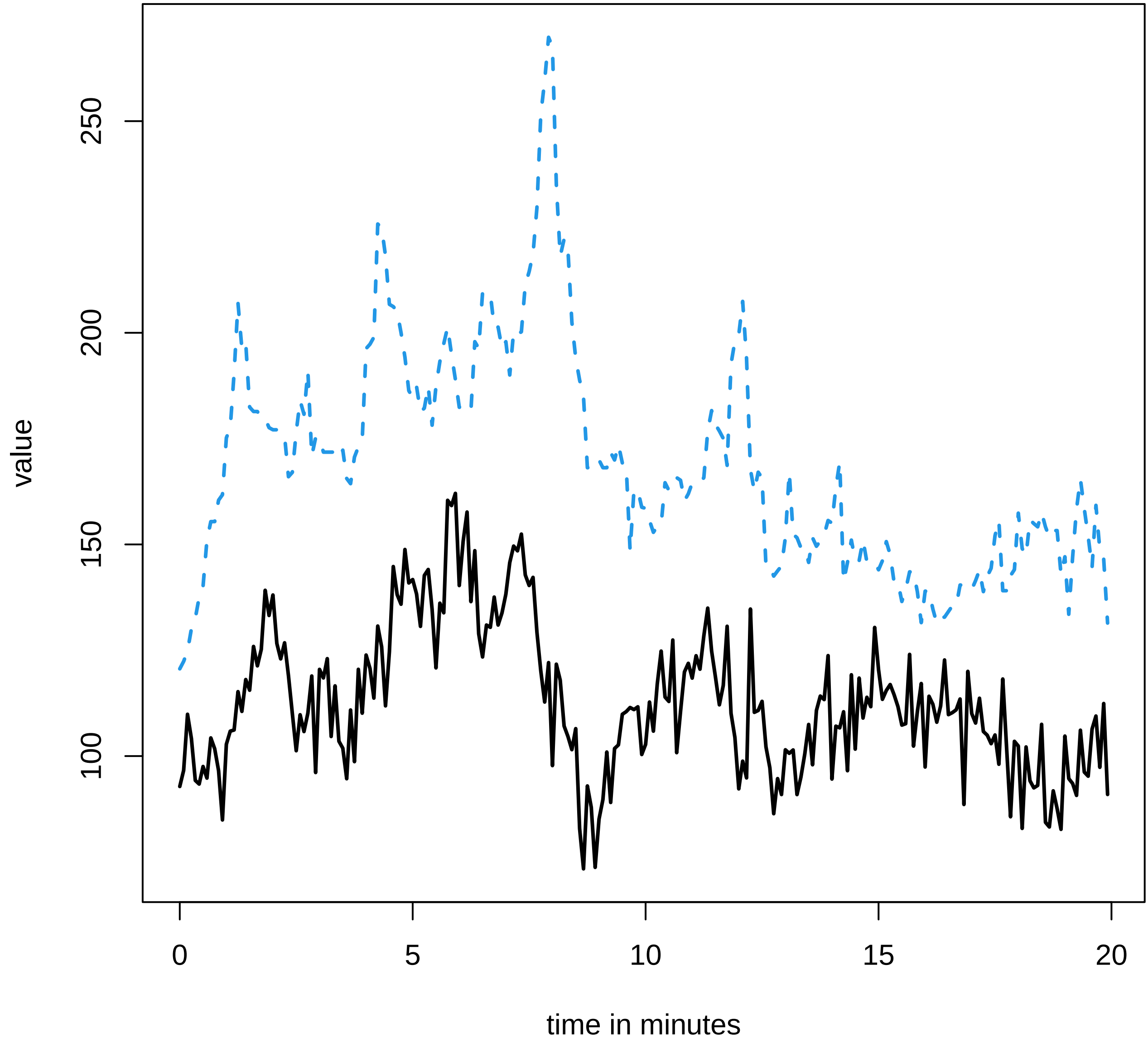

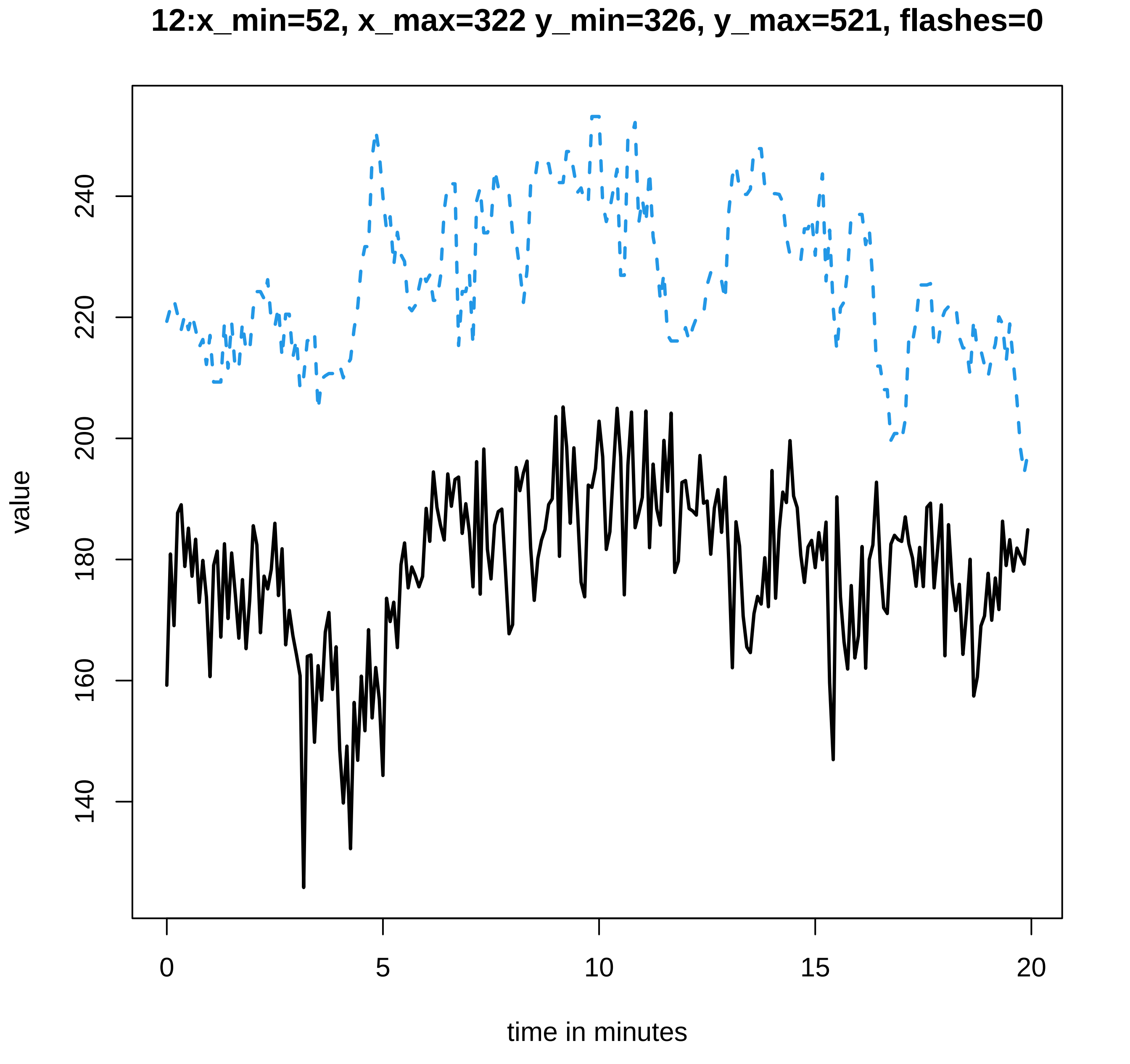

12:x_min=52, x_max=322 y_min=326, y_max=521, flashes=0
value
140
160
180
200
220
240
0
5
10
15
20
time in minutes

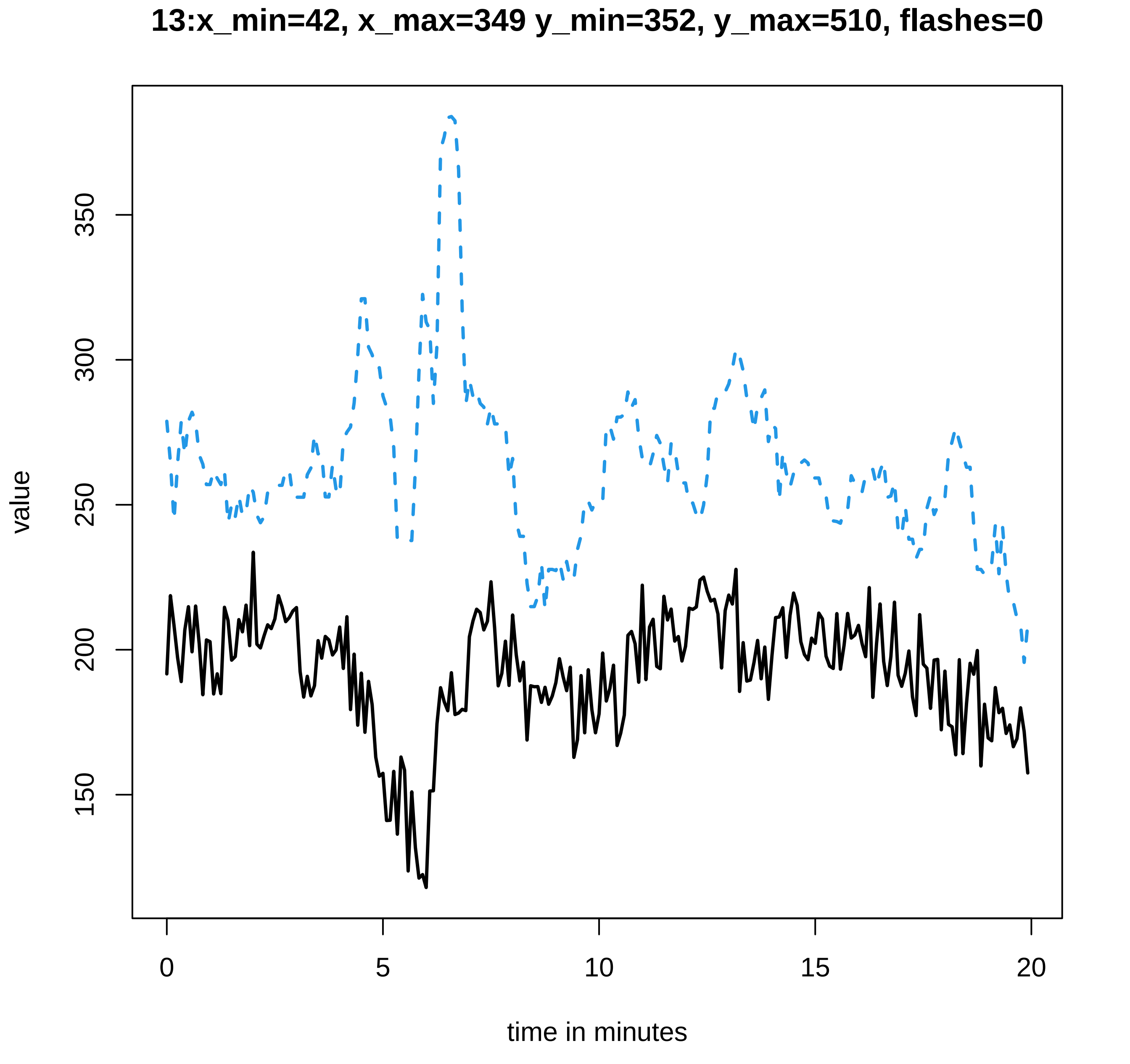
13:x_min=42, x_max=349 y_min=352, y_max=510, flashes=0
value
350
300
250
200
150
0
5
10
15
20
time in minutes

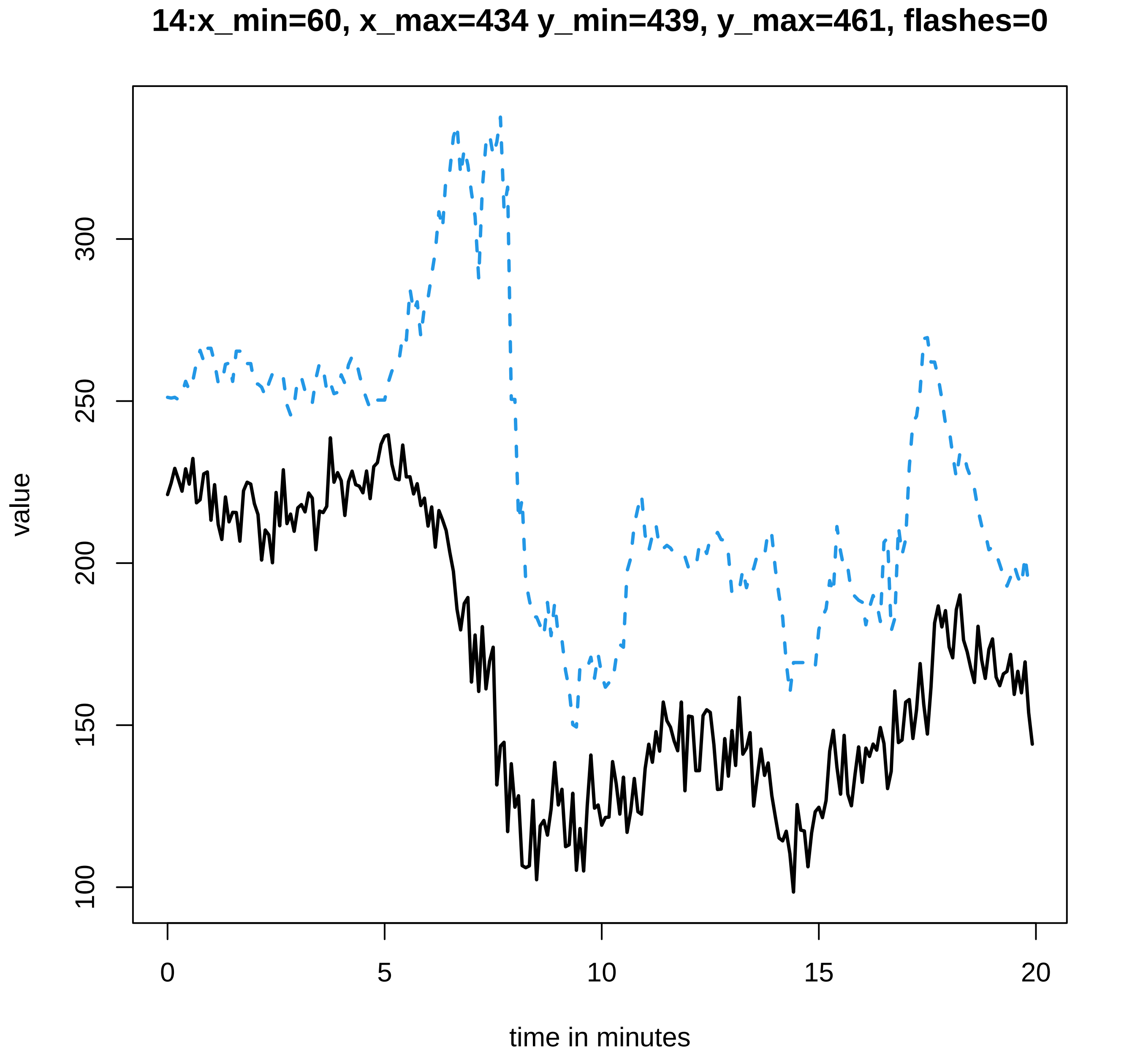

14:x_min=60, x_max=434 y_min=439, y_max=461, flashes=0
value
300
250
200
150
100
0
5
10
15
20
time in minutes

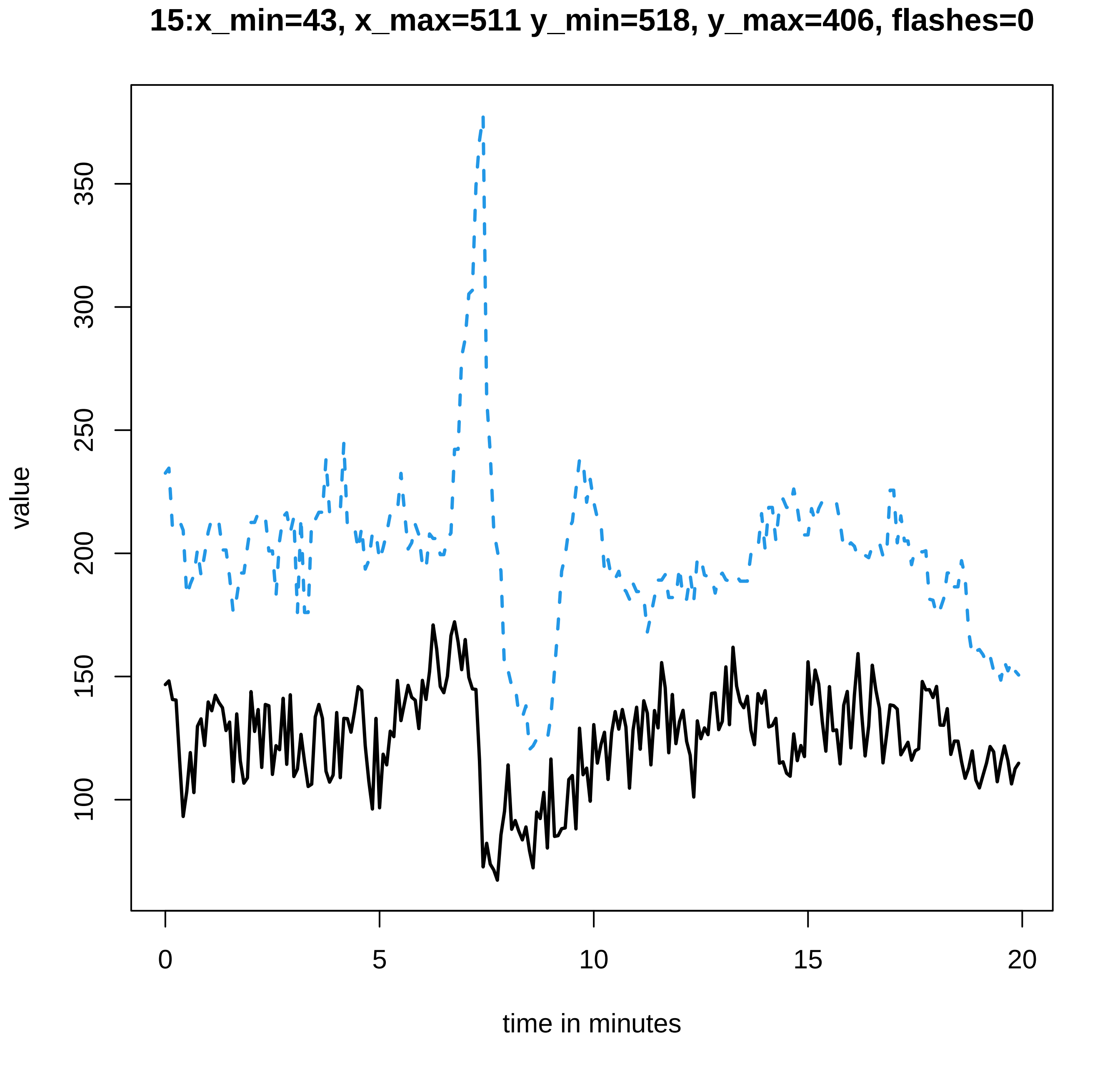

15:x_min=43, x_max=511 y_min=518, y_max=406, flashes=0
value
100
150
200
250
300
350
0
5
10
15
20
time in minutes

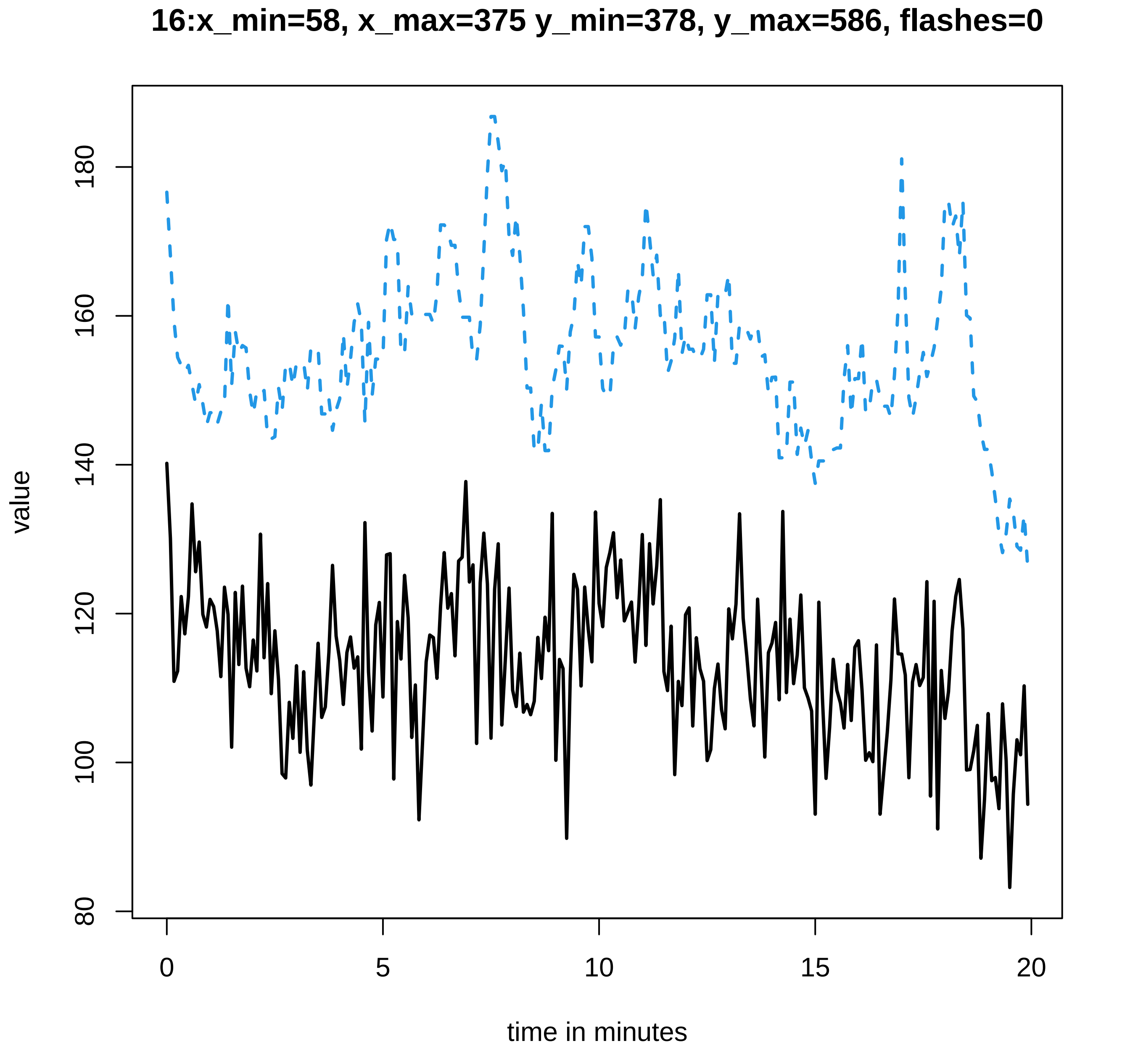
16:x_min=58, x_max=375 y_min=378, y_max=586, flashes=0
value
time in minutes
80
100
120
140
160
180
0
5
10
15
20

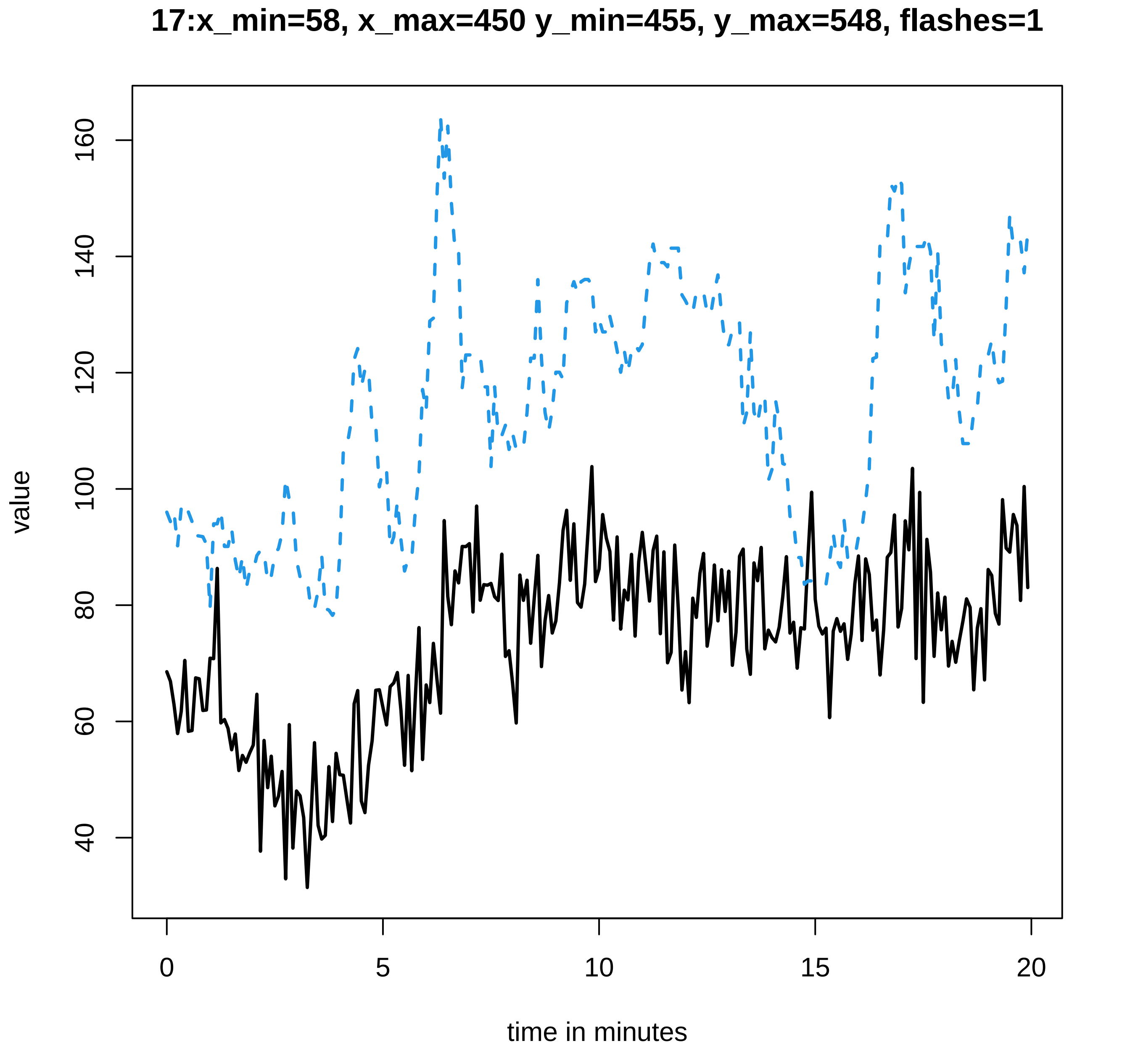

17:x_min=58, x_max=450 y_min=455, y_max=548, flashes=1
value
160
140
120
100
80
60
40
0
5
10
15
20
time in minutes

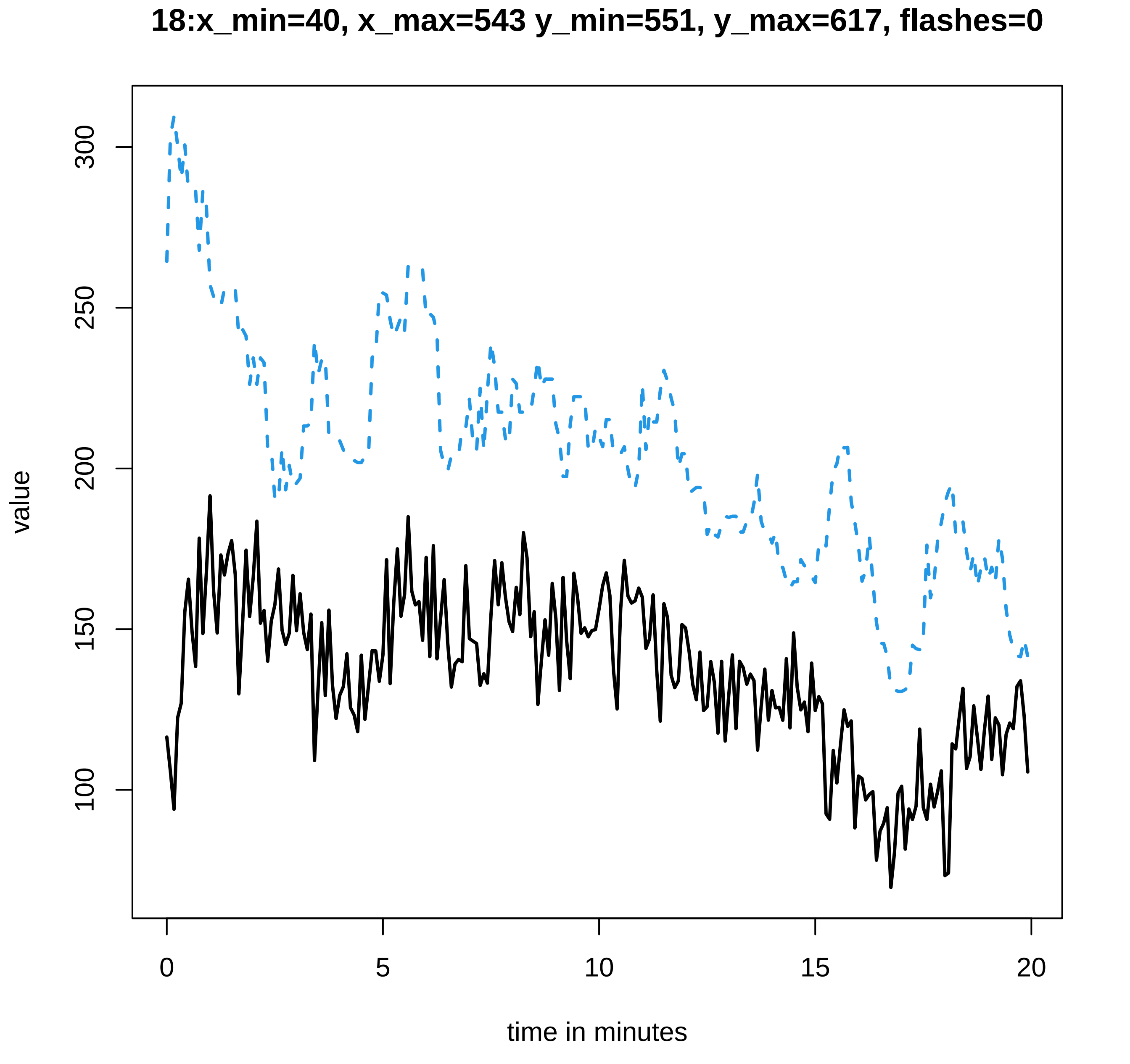
18:x_min=40, x_max=543 y_min=551, y_max=617, flashes=0
value
300
250
200
150
100
0
5
10
15
20
time in minutes

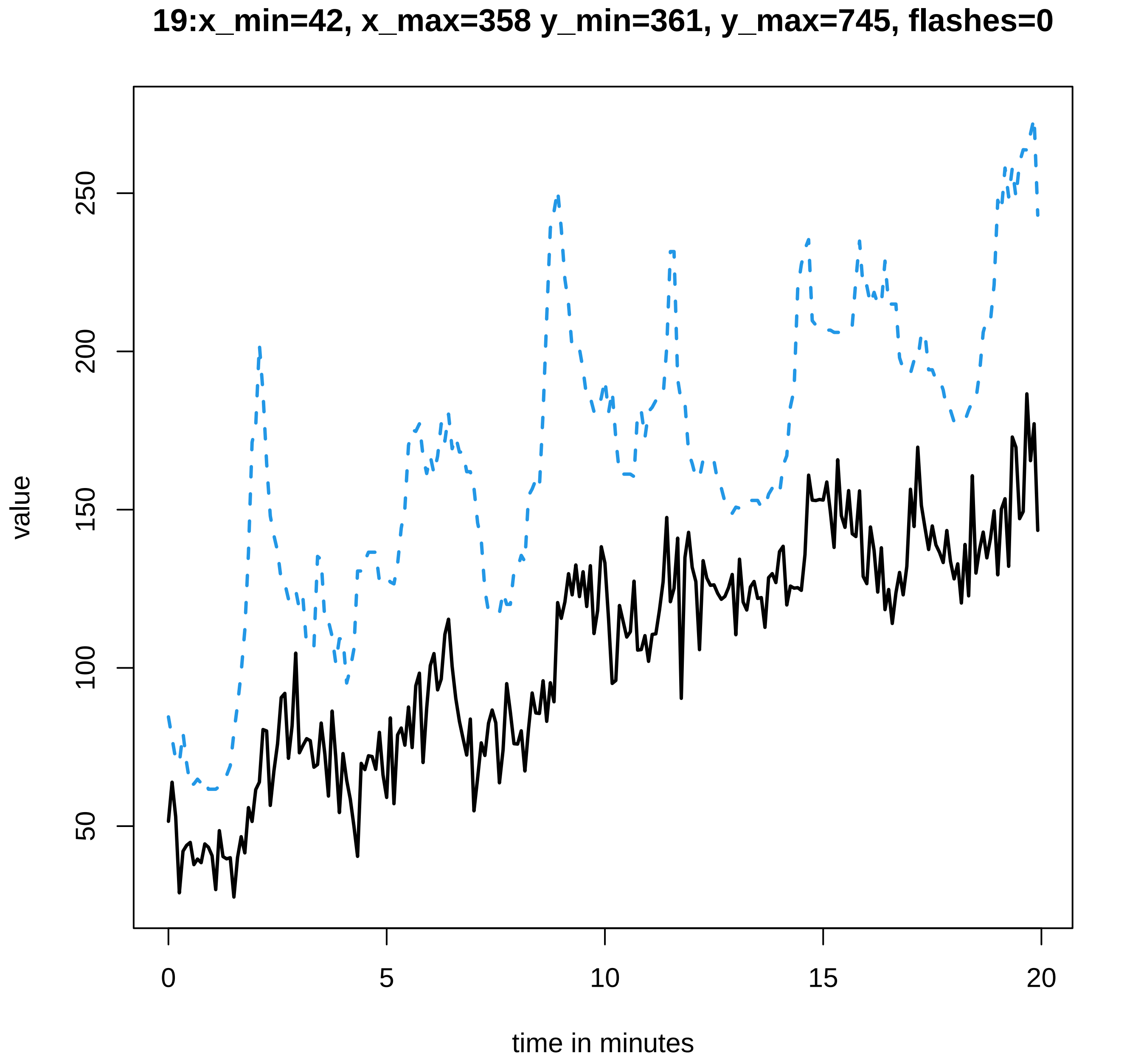
19:x_min=42, x_max=358 y_min=361, y_max=745, flashes=0
value
50
100
150
200
250
0
5
10
15
20
time in minutes

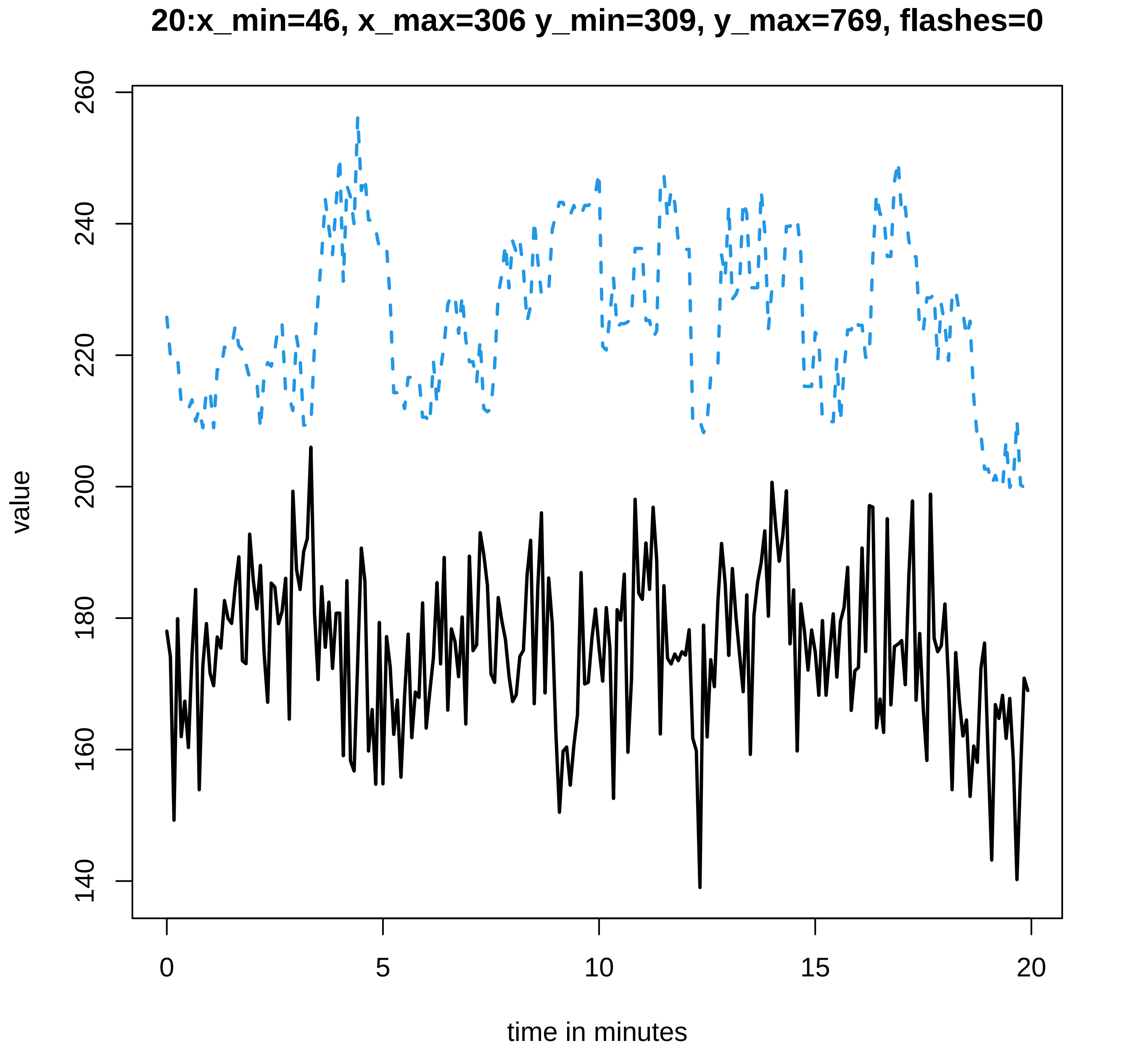
20:x_min=46, x_max=306 y_min=309, y_max=769, flashes=0
value
260
240
220
200
180
160
140
0
5
10
15
20
time in minutes

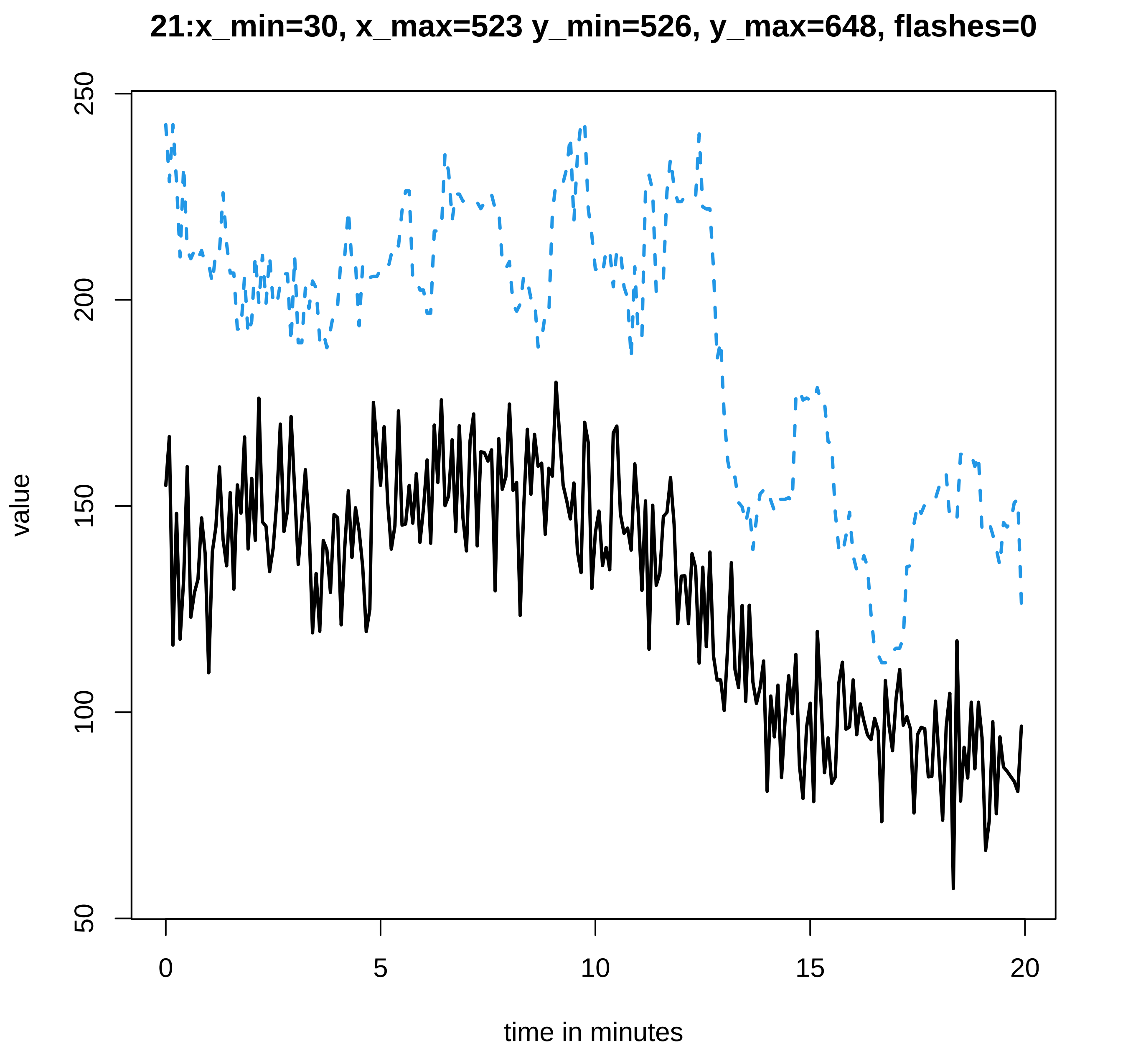

21:x_min=30, x_max=523 y_min=526, y_max=648, flashes=0
value
250
200
150
100
50
0
5
10
15
20
time in minutes

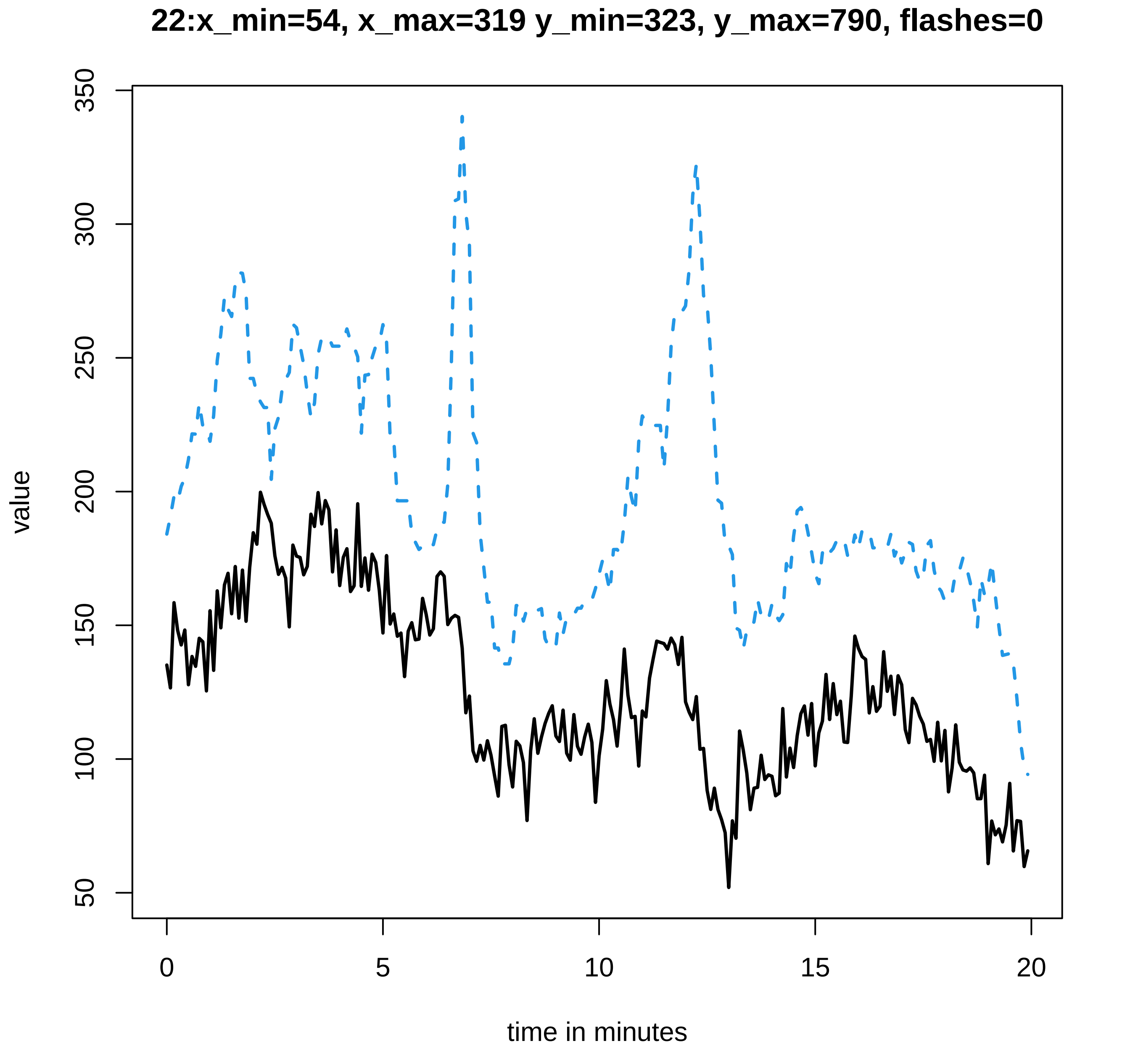

22:x_min=54, x_max=319 y_min=323, y_max=790, flashes=0
value
350
300
250
200
150
100
50
0
5
10
15
20
time in minutes

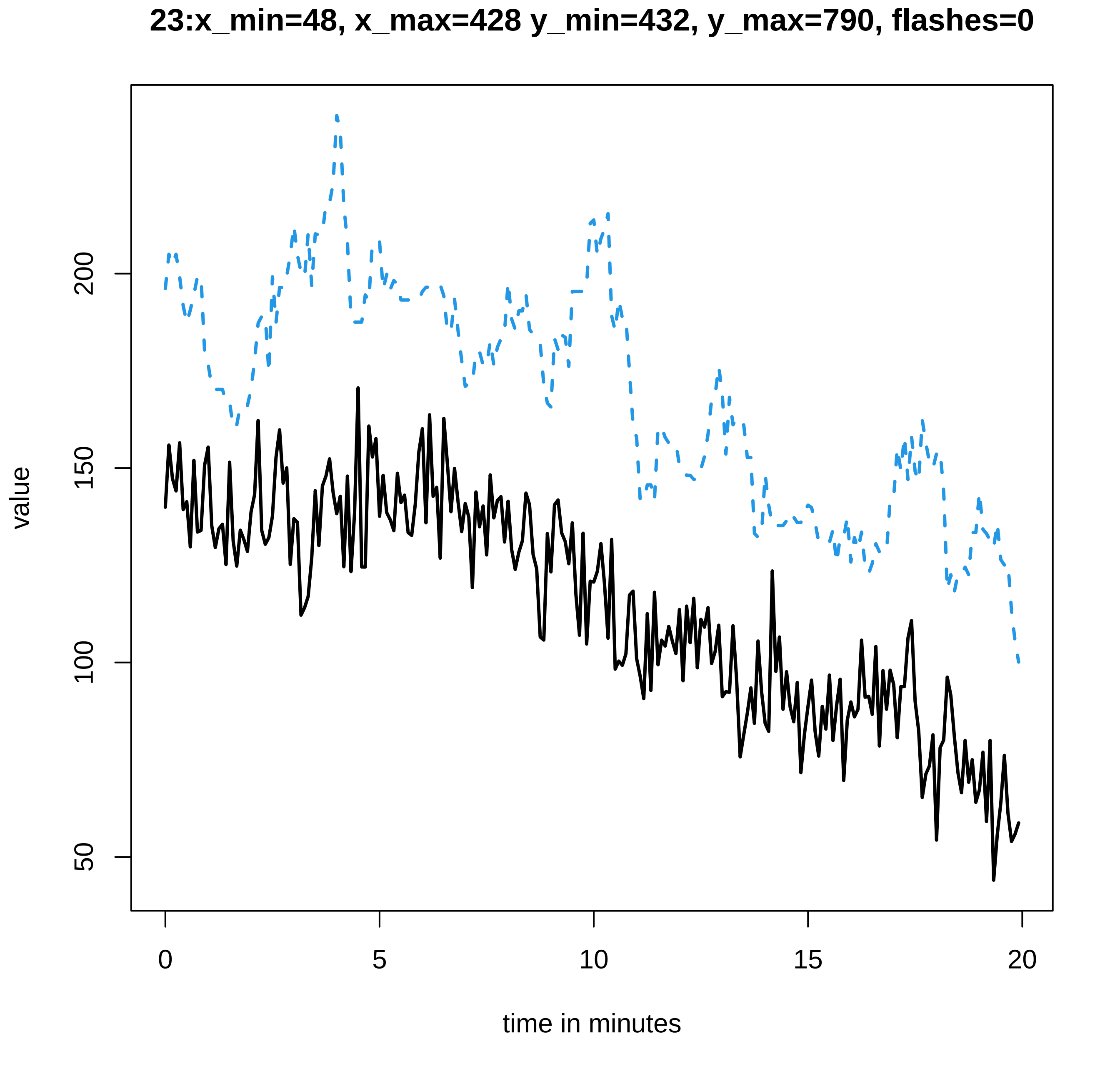
23:x_min=48, x_max=428 y_min=432, y_max=790, flashes=0
value
200
150
100
50
0
5
10
15
20
time in minutes

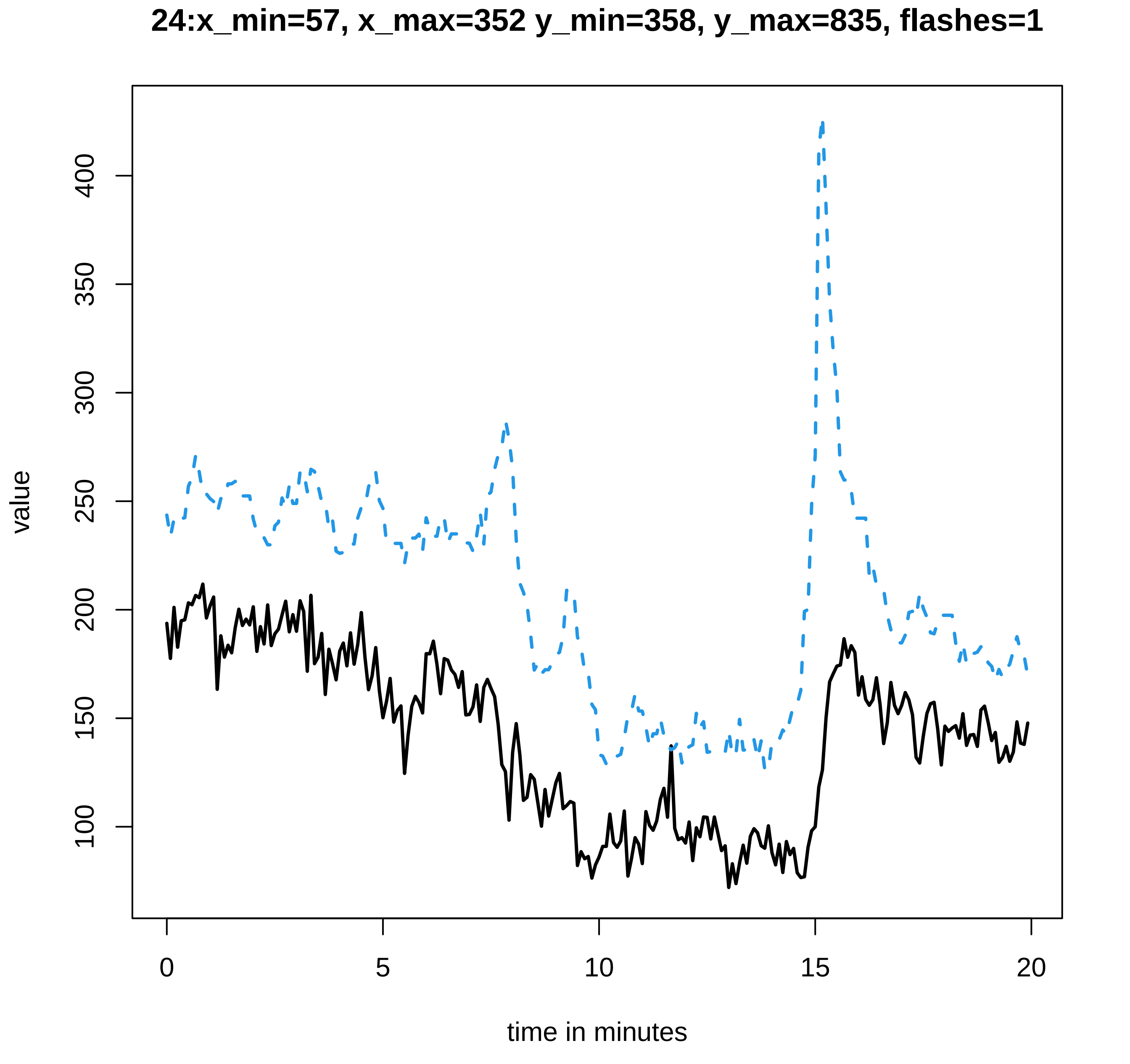

24:x_min=57, x_max=352 y_min=358, y_max=835, flashes=1
value
100
150
200
250
300
350
400
0
5
10
15
20
time in minutes

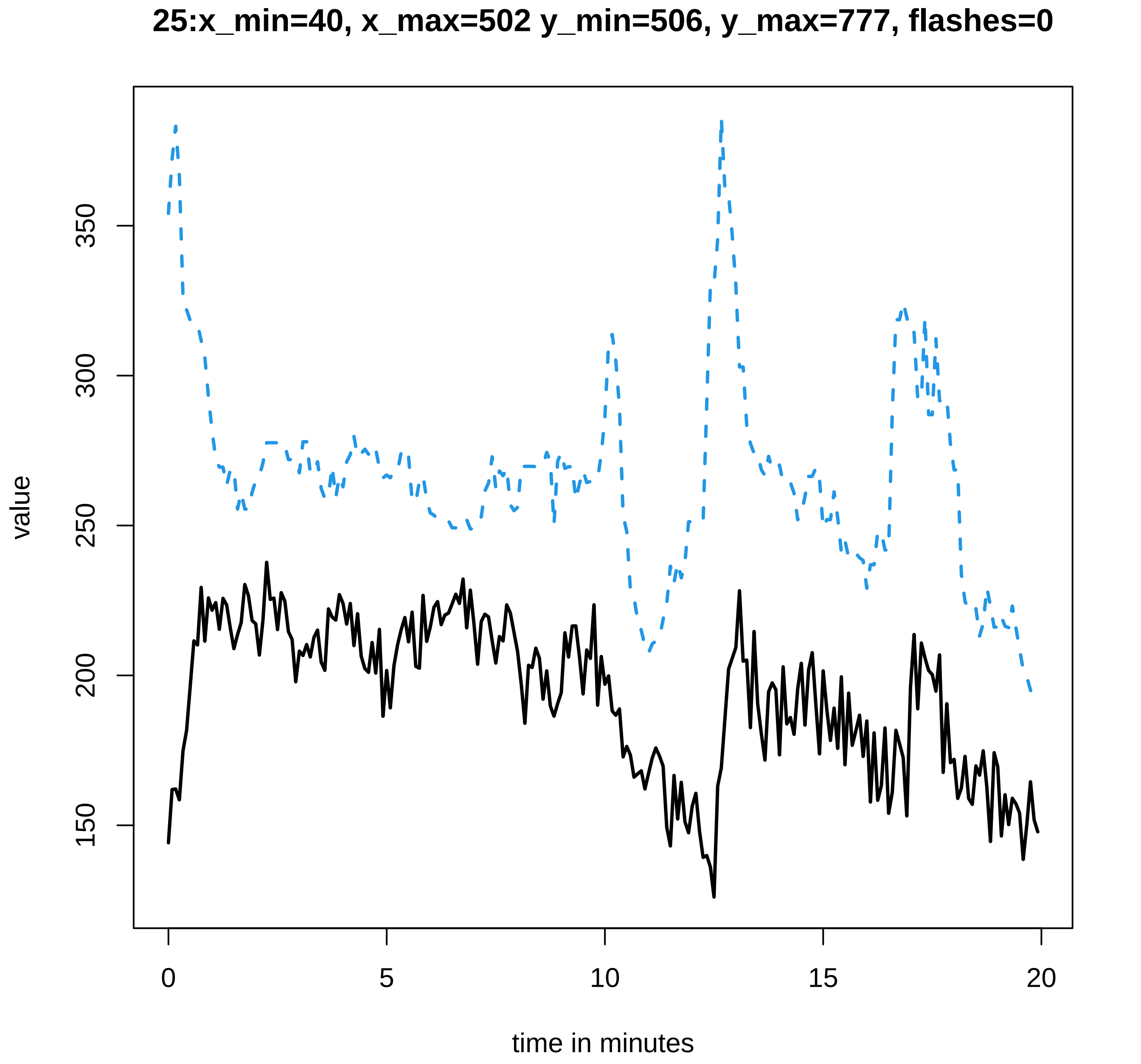

25:x_min=40, x_max=502 y_min=506, y_max=777, flashes=0
value
150
200
250
300
350
0
5
10
15
20
time in minutes

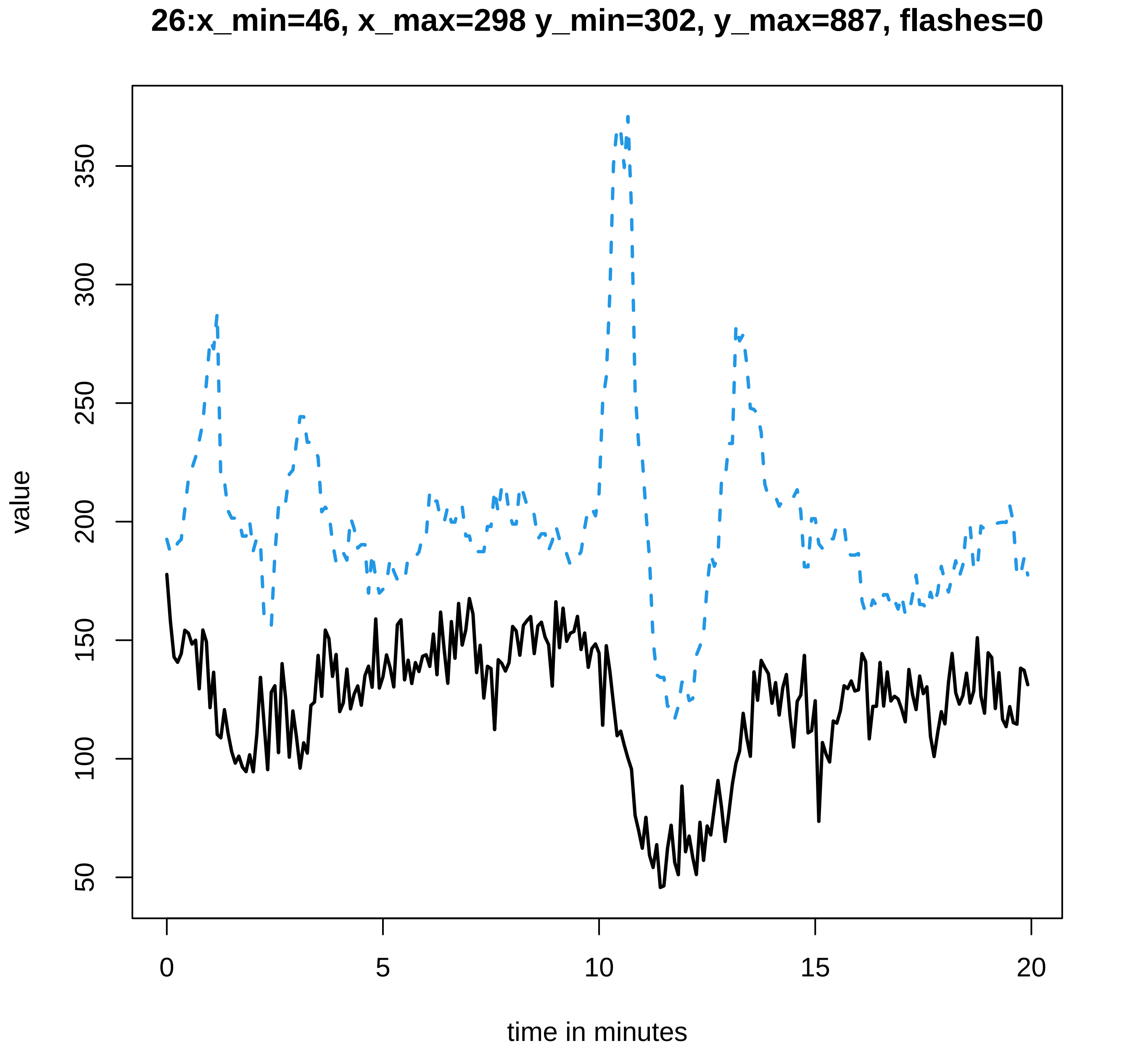
26:x_min=46, x_max=298 y_min=302, y_max=887, flashes=0
value
50
100
150
200
250
300
350
0
5
10
15
20
time in minutes

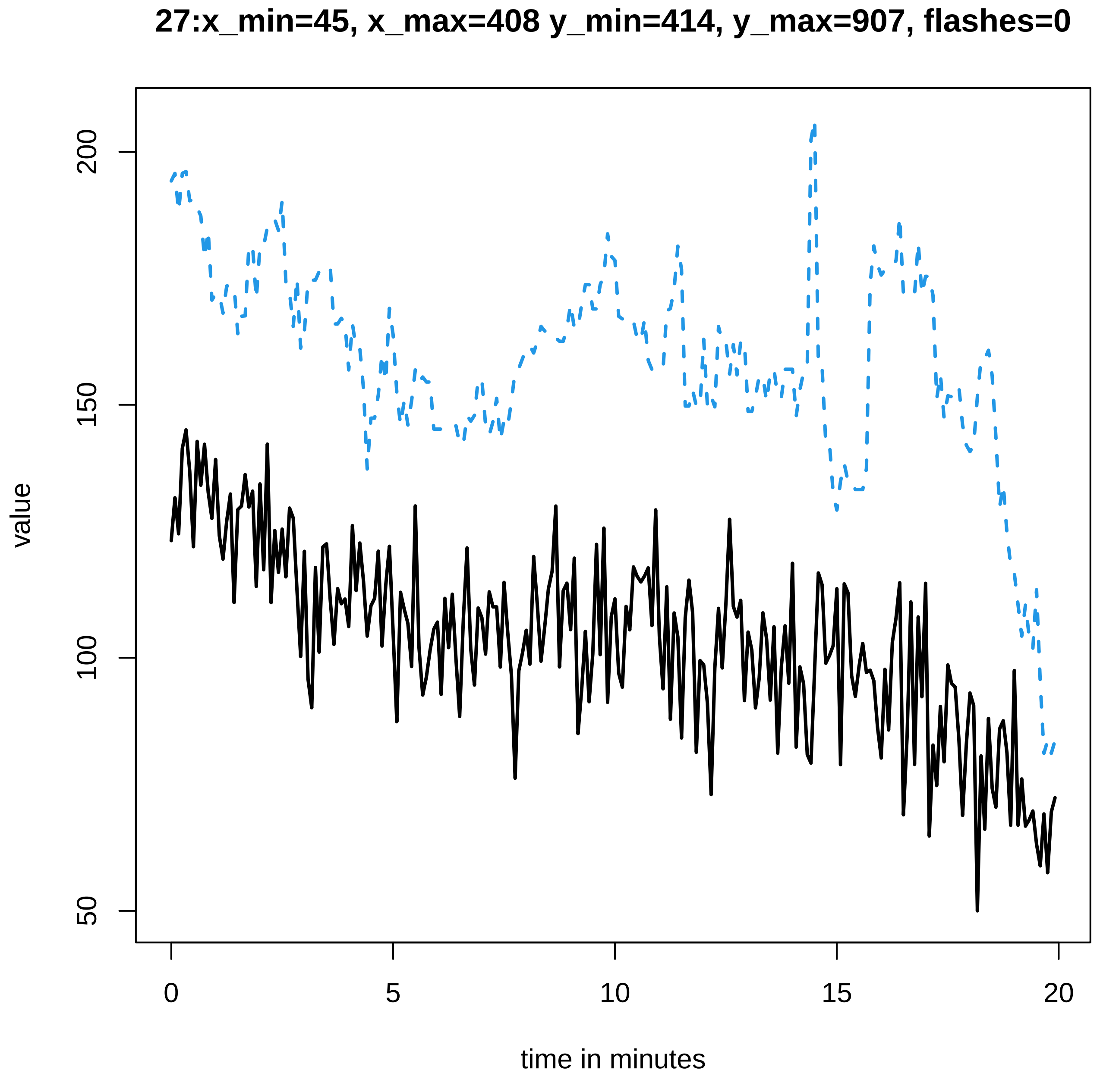
27:x_min=45, x_max=408 y_min=414, y_max=907, flashes=0
value
200
150
100
50
0
5
10
15
20
time in minutes

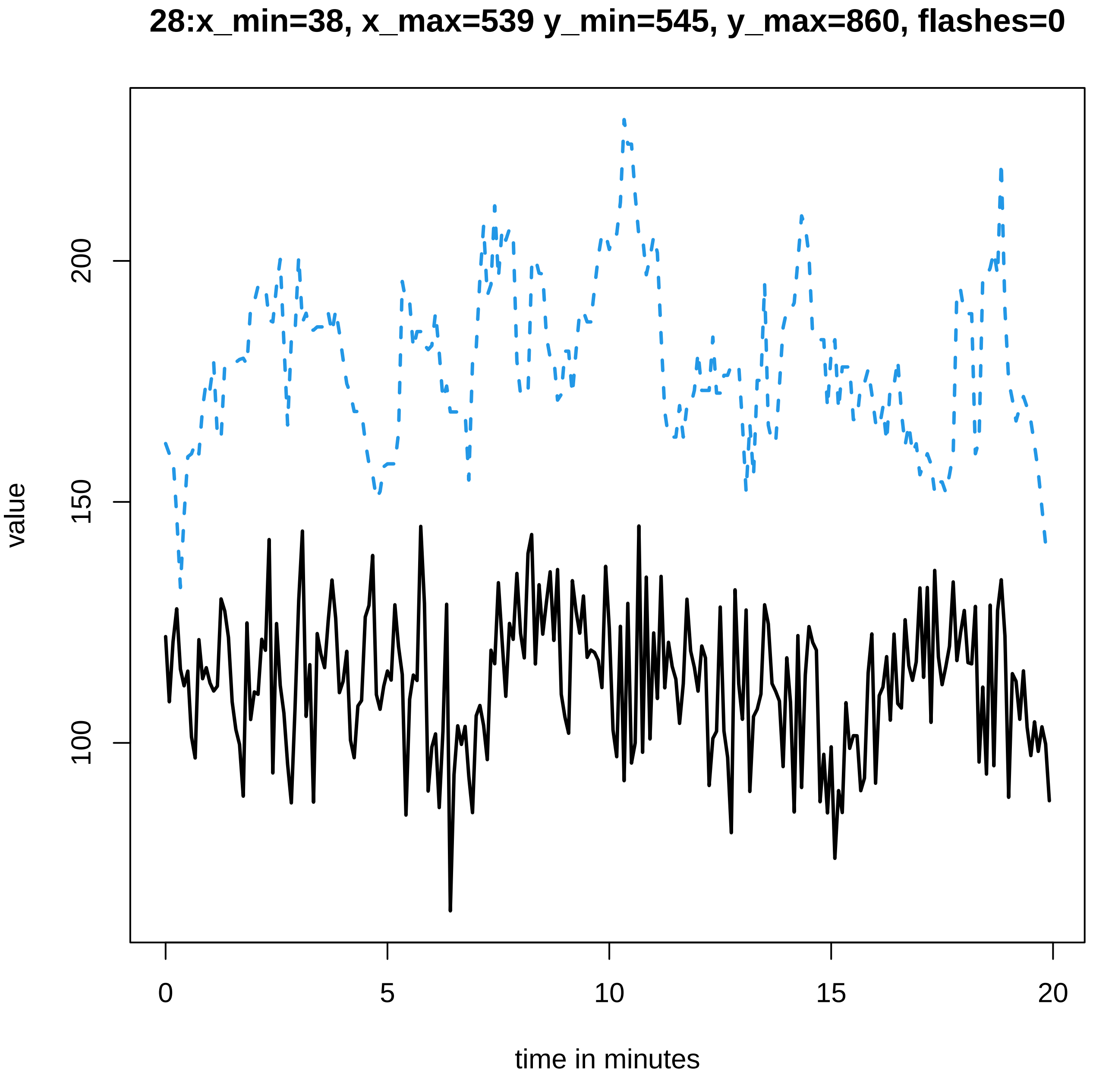

28:x_min=38, x_max=539 y_min=545, y_max=860, flashes=0
value
200
150
100
0
5
10
15
20
time in minutes

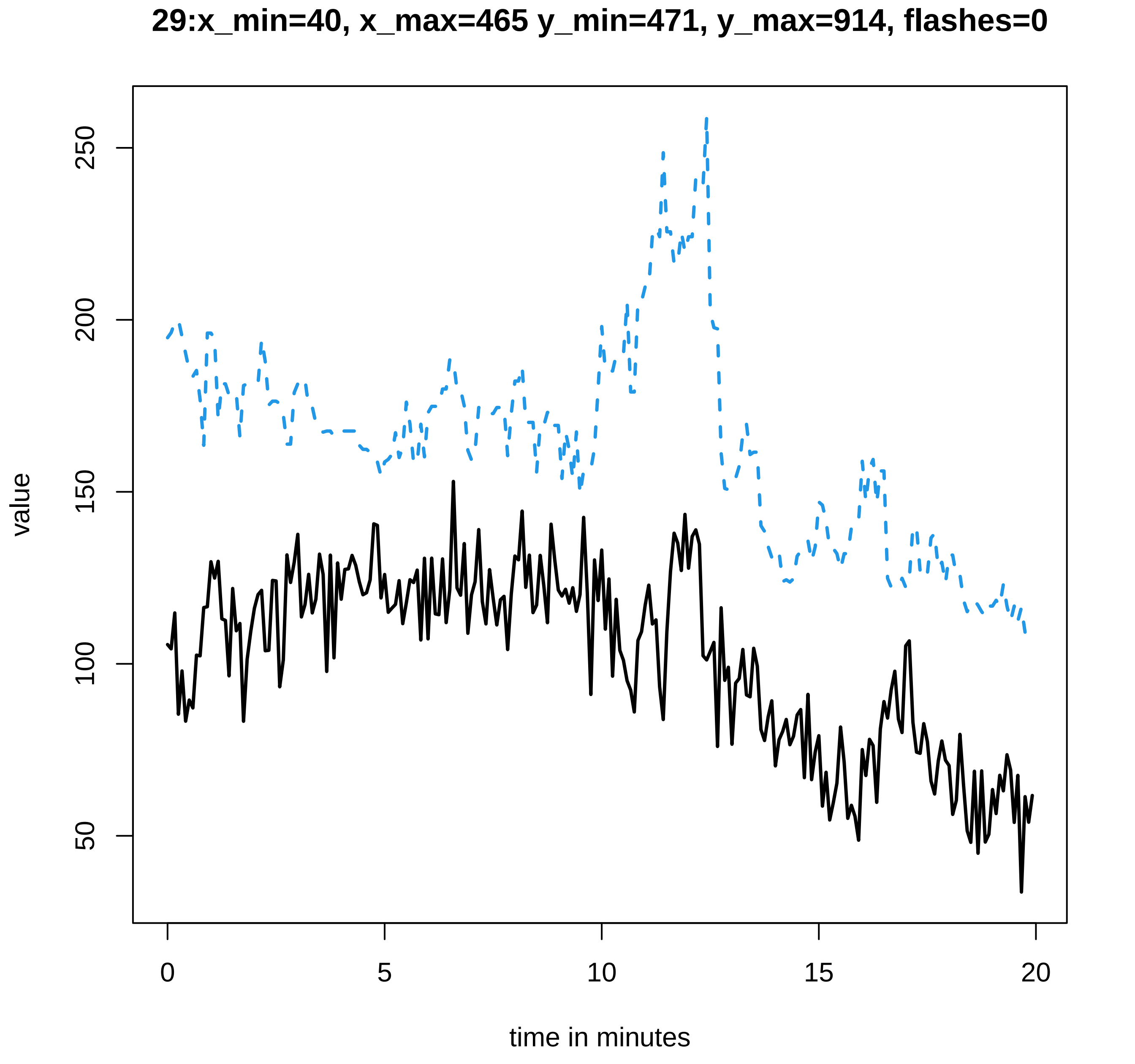
29:x_min=40, x_max=465 y_min=471, y_max=914, flashes=0
value
250
200
150
100
50
0
5
10
15
20
time in minutes

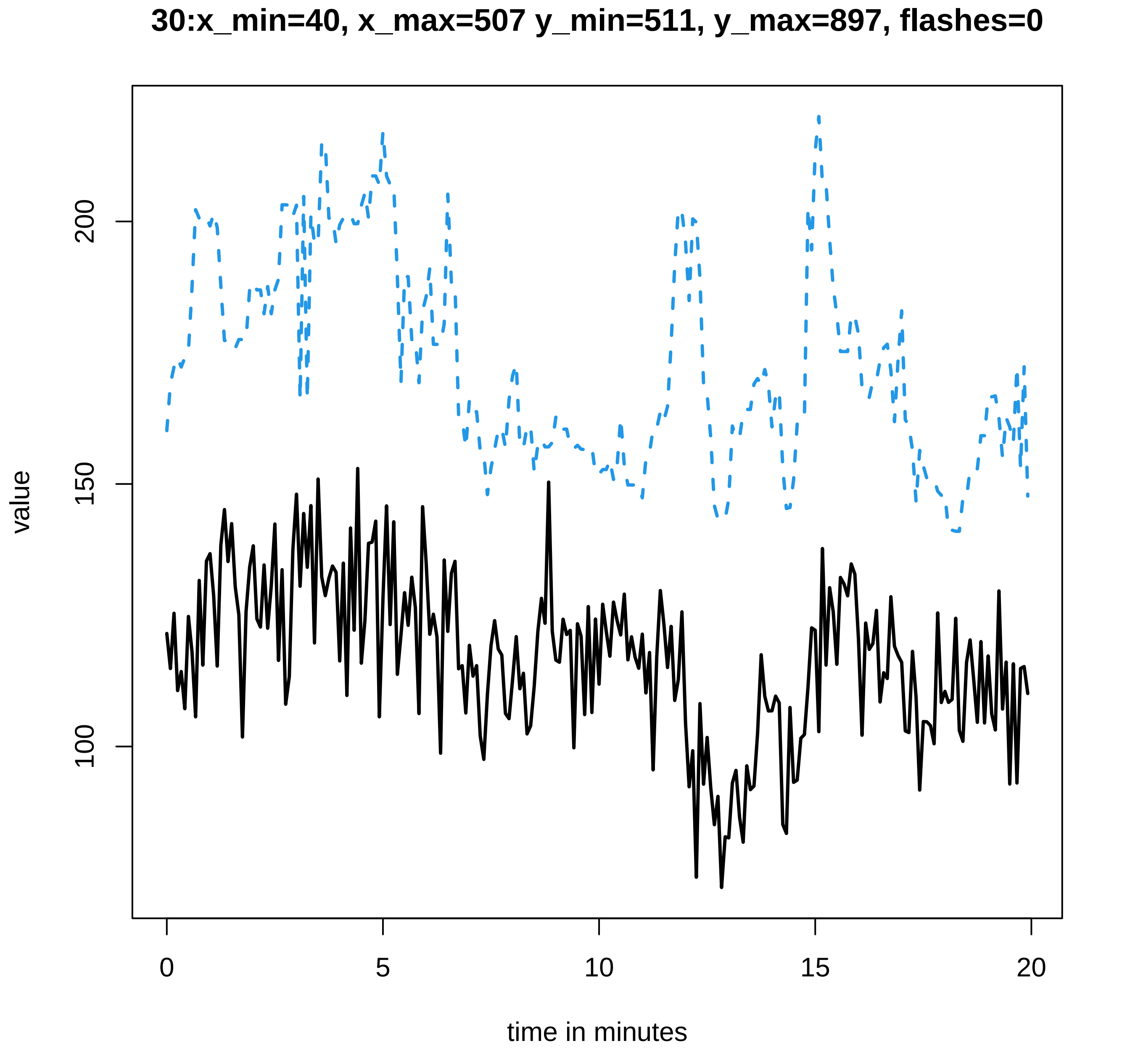
30:x_min=40, x_max=507 y_min=511, y_max=897, flashes=0
value
100
150
200
0
5
10
15
20
time in minutes

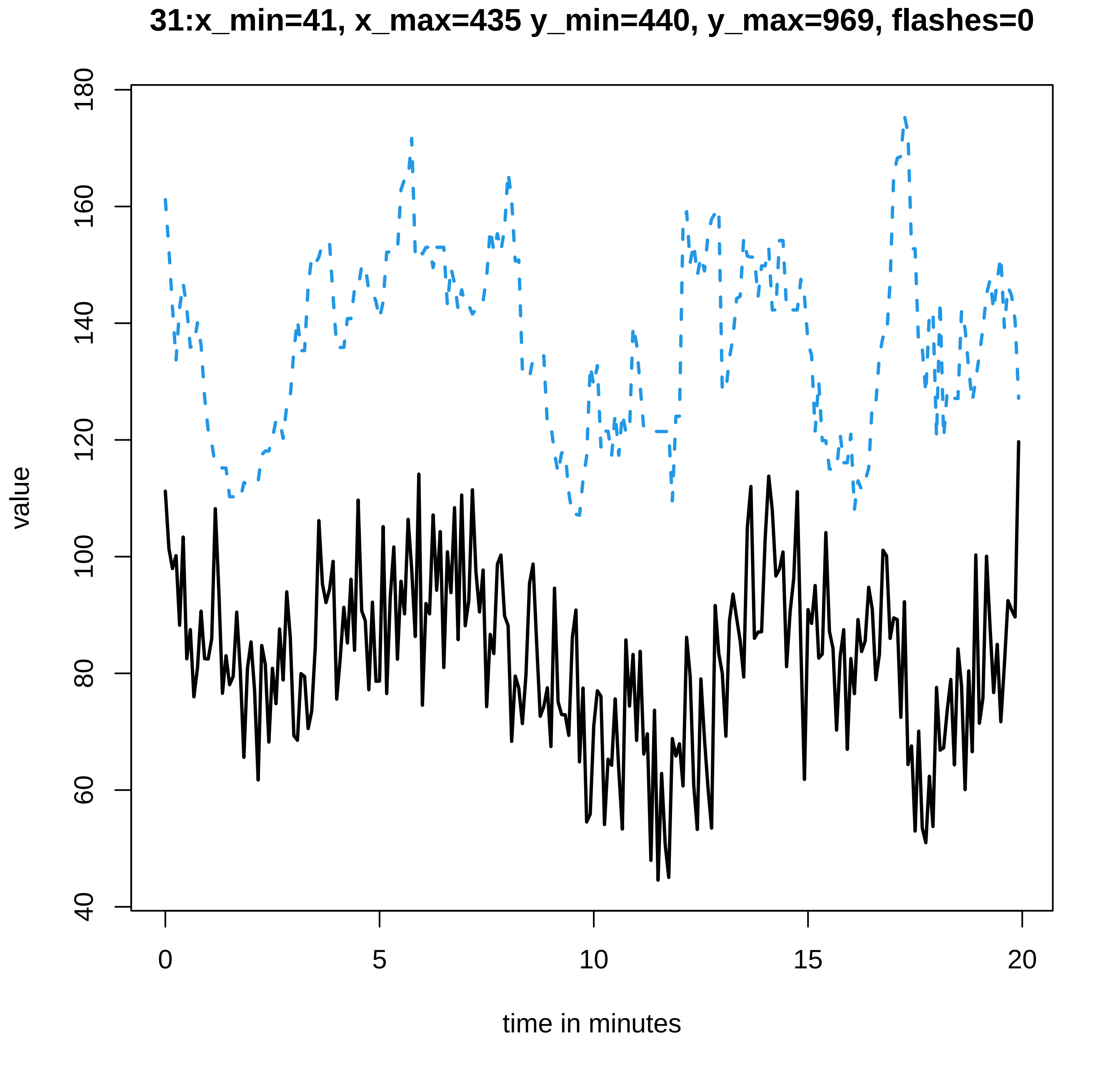
31:x_min=41, x_max=435 y_min=440, y_max=969, flashes=0
value
time in minutes
0
5
10
15
20
40
60
80
100
120
140
160
180

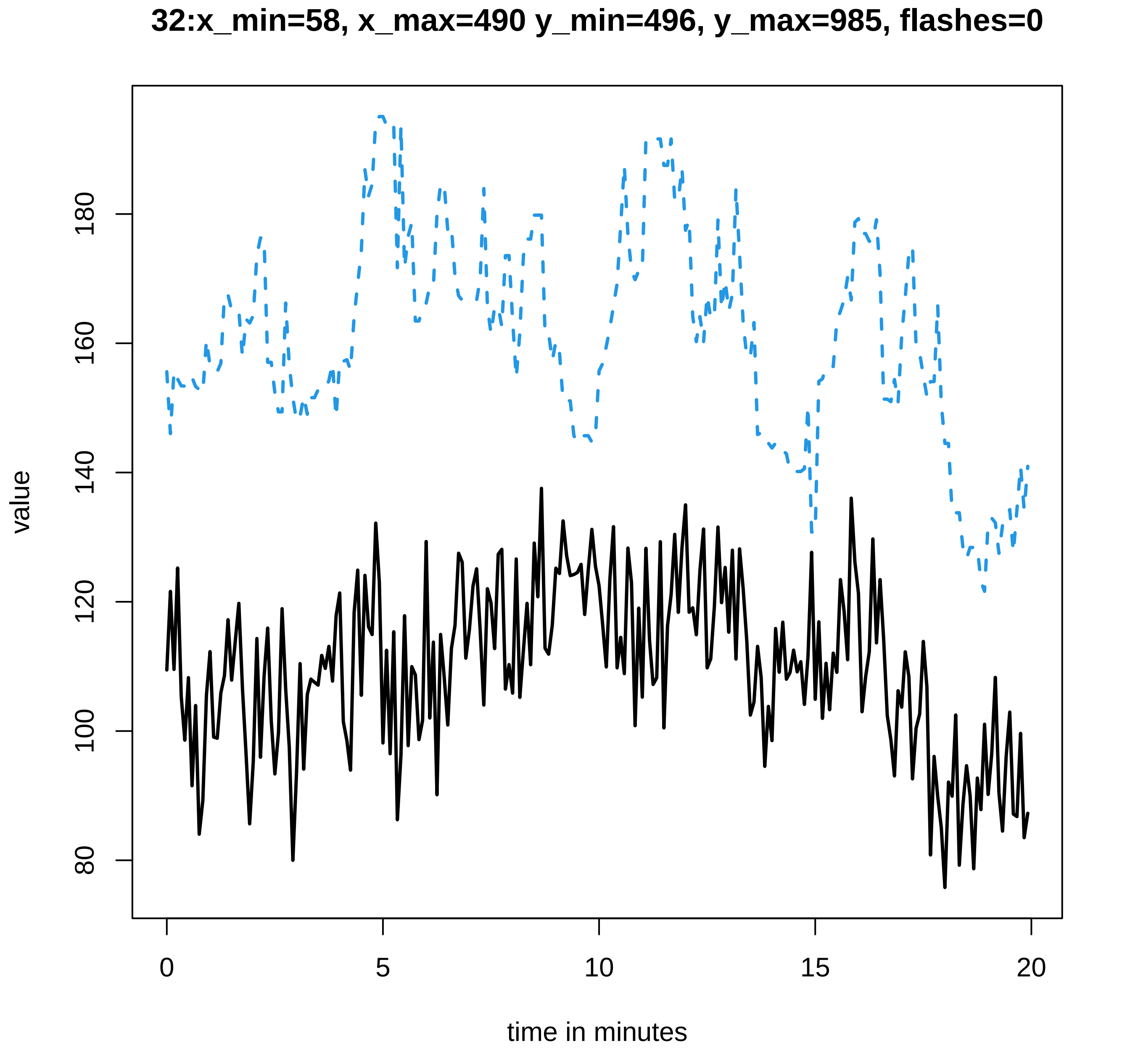

32:x_min=58, x_max=490 y_min=496, y_max=985, flashes=0
value
80
100
120
140
160
180
0
5
10
15
20
time in minutes

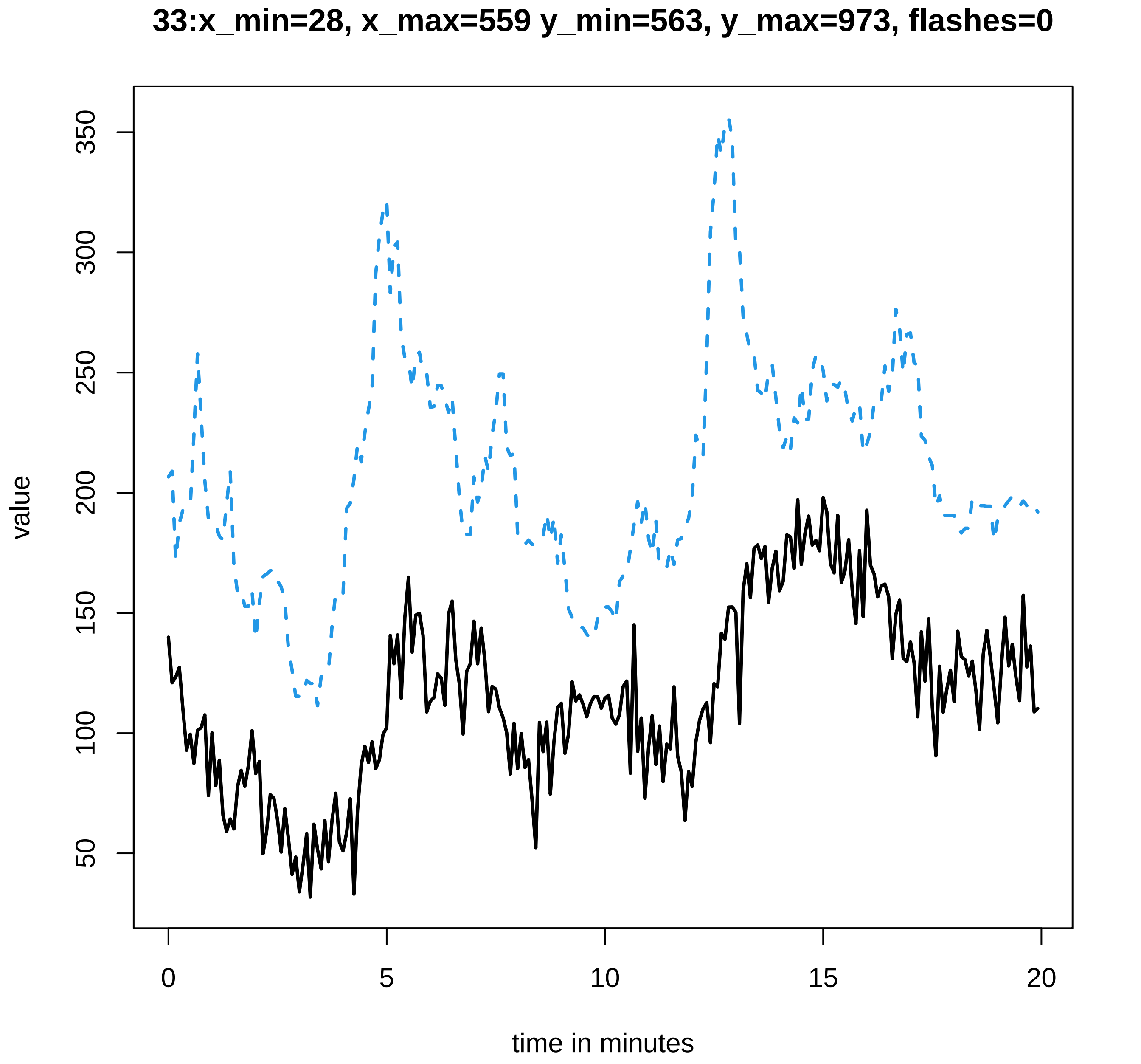
33:x_min=28, x_max=559 y_min=563, y_max=973, flashes=0
value
50
100
150
200
250
300
350
0
5
10
15
20
time in minutes
