## Supplementary data Calcium spiking for "LysM Receptor Proteins are Required for Ectomycorrhizal Symbiosis in Poplar": profiles-spiking-17.pdf

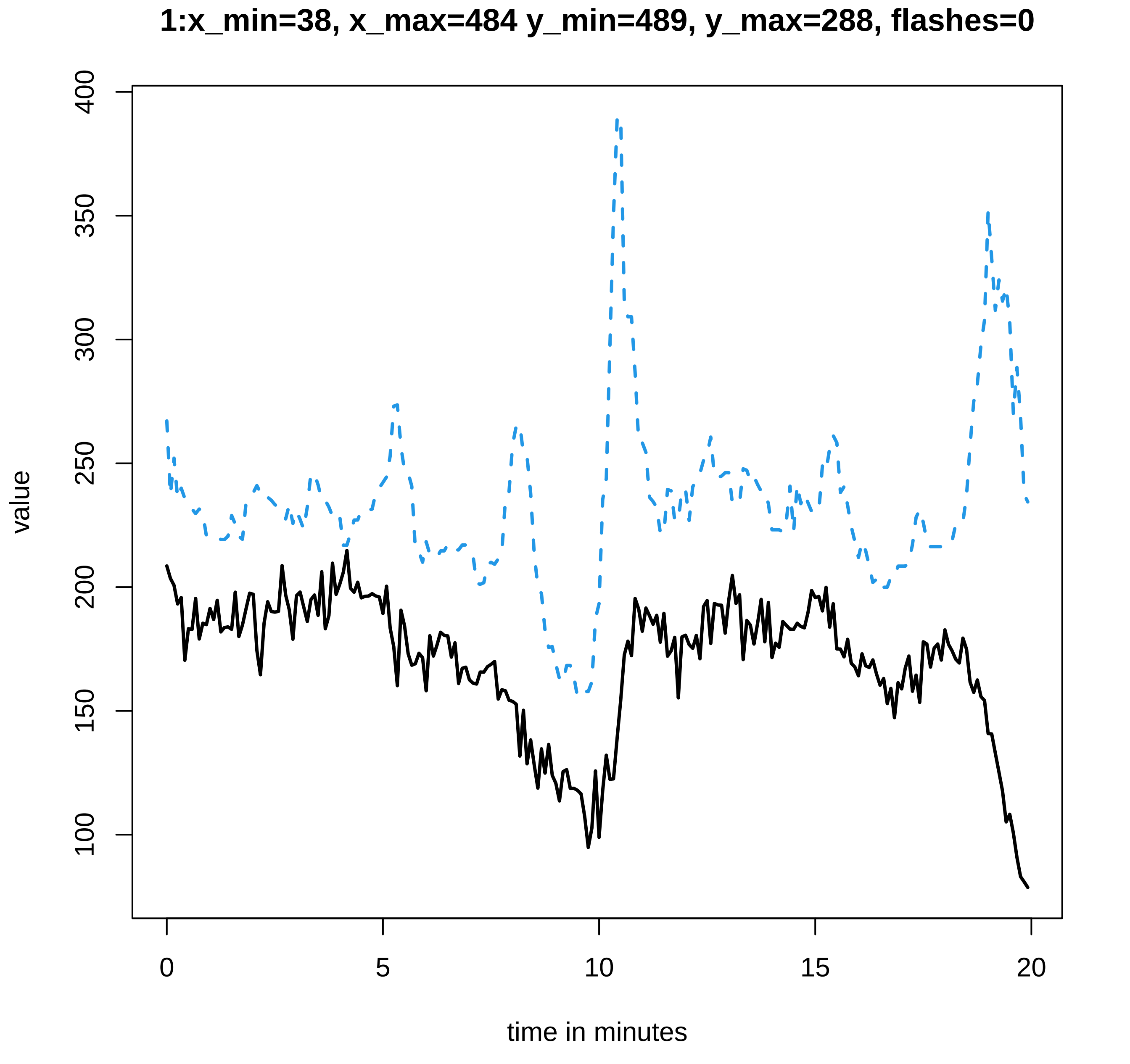
1:x_min=38, x_max=484 y_min=489, y_max=288, flashes=0
value
400
350
300
250
200
150
100
0
5
10
15
20
time in minutes

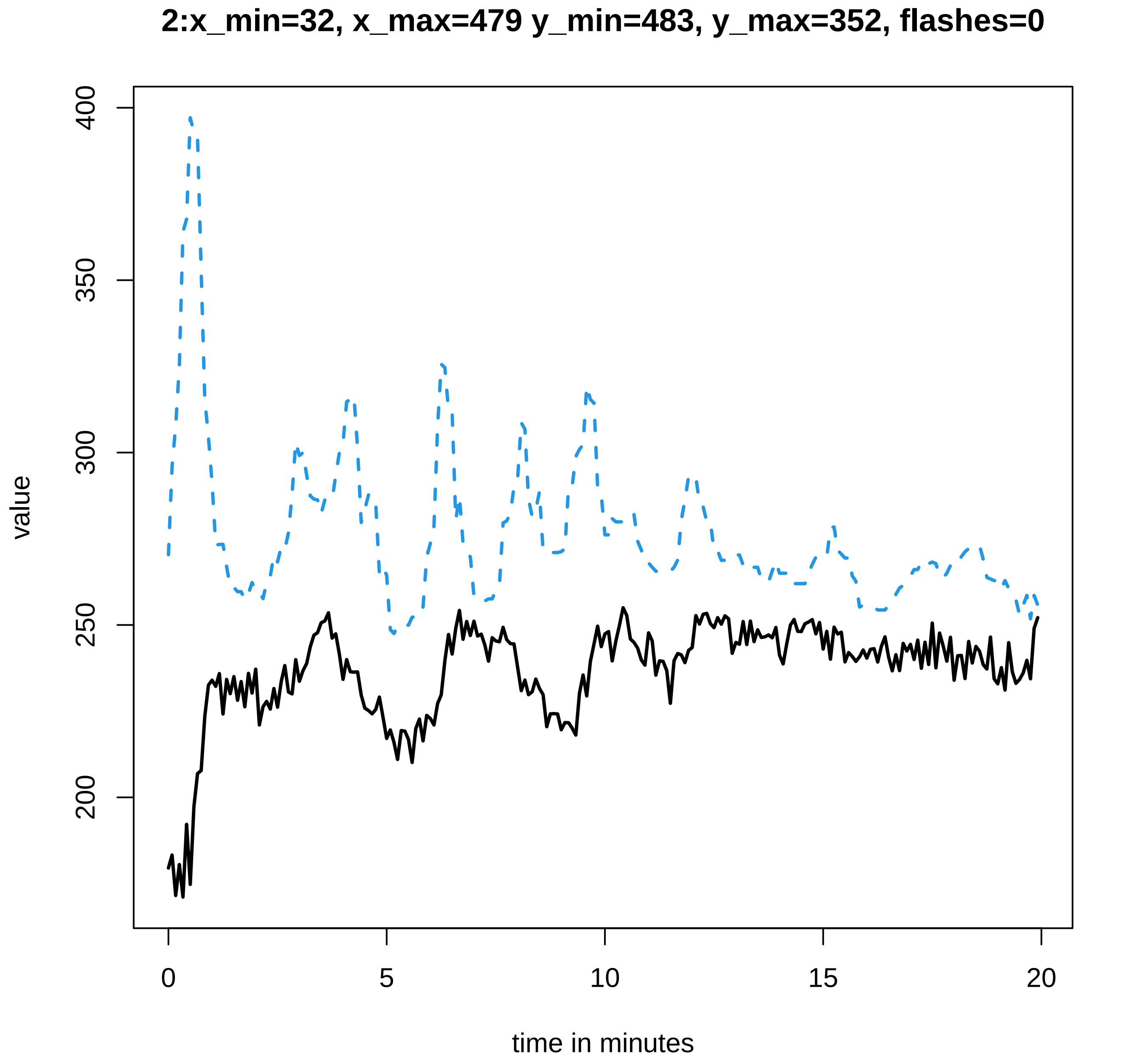
2:x_min=32, x_max=479 y_min=483, y_max=352, flashes=0
value
400
350
300
250
200
0
5
10
15
20
time in minutes

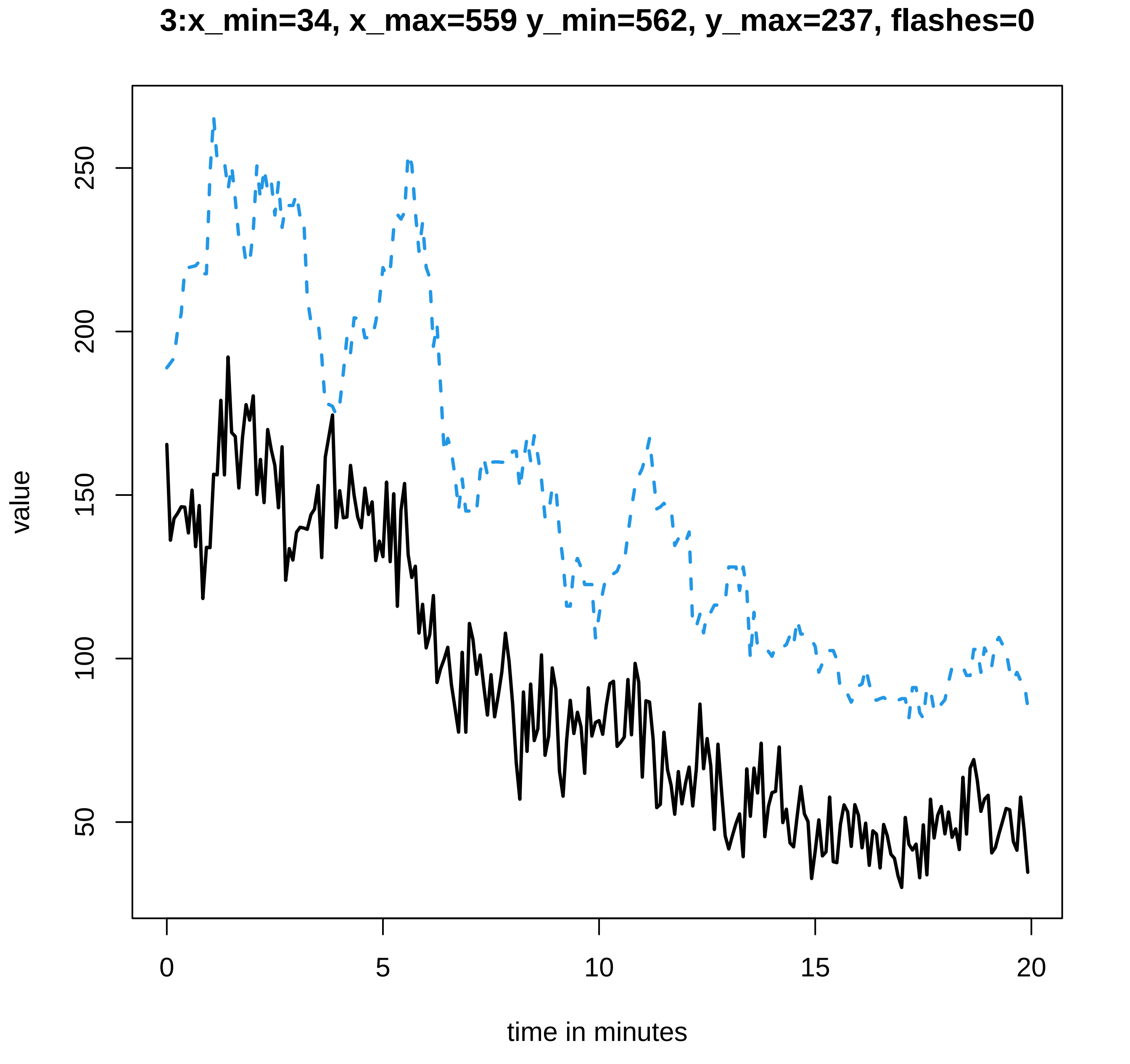
3:x_min=34, x_max=559 y_min=562, y_max=237, flashes=0
value
250
200
150
100
50
0
5
10
15
20
time in minutes

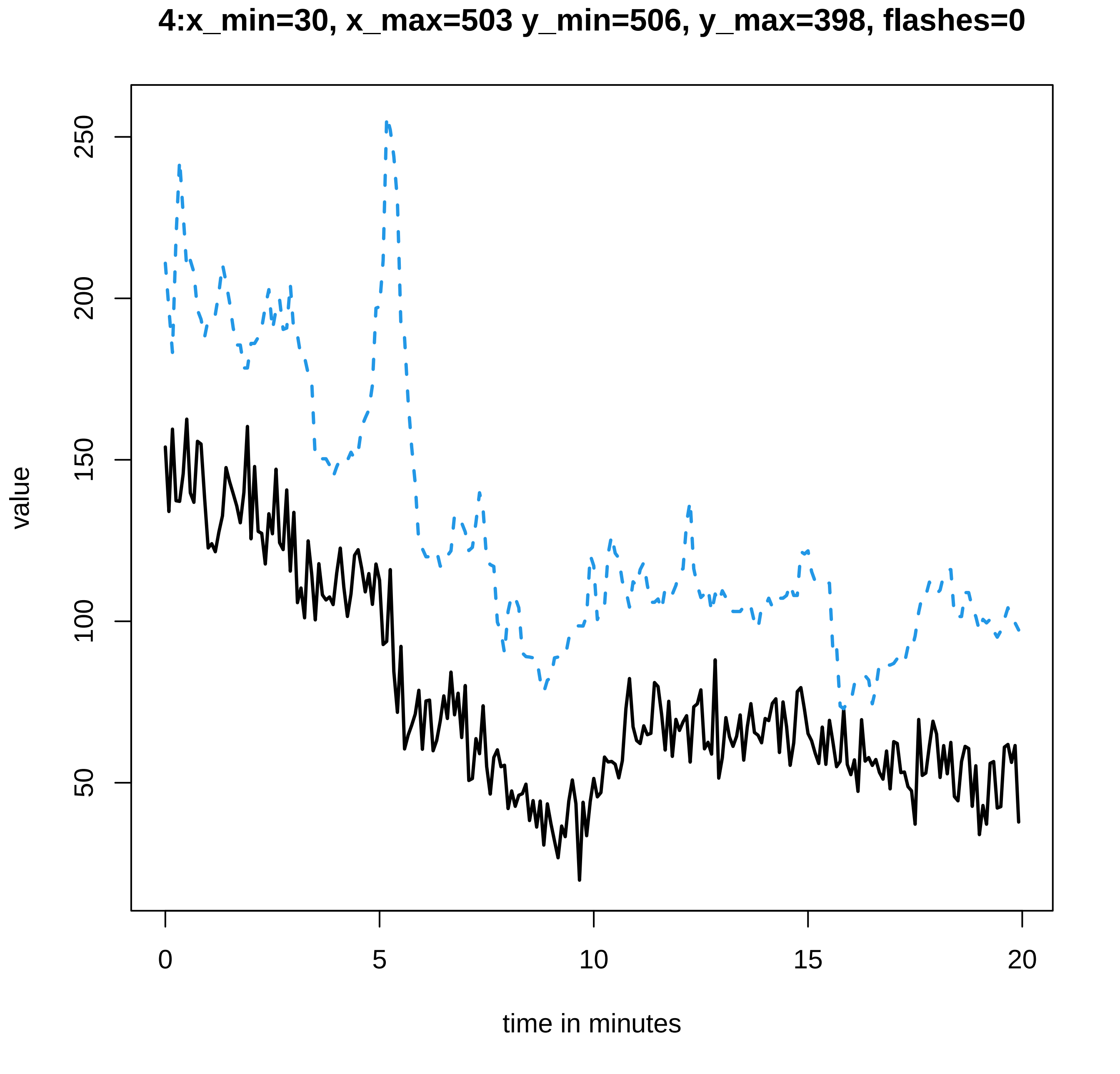
4:x_min=30, x_max=503 y_min=506, y_max=398, flashes=0
value
250
200
150
100
50
0
5
10
15
20
time in minutes

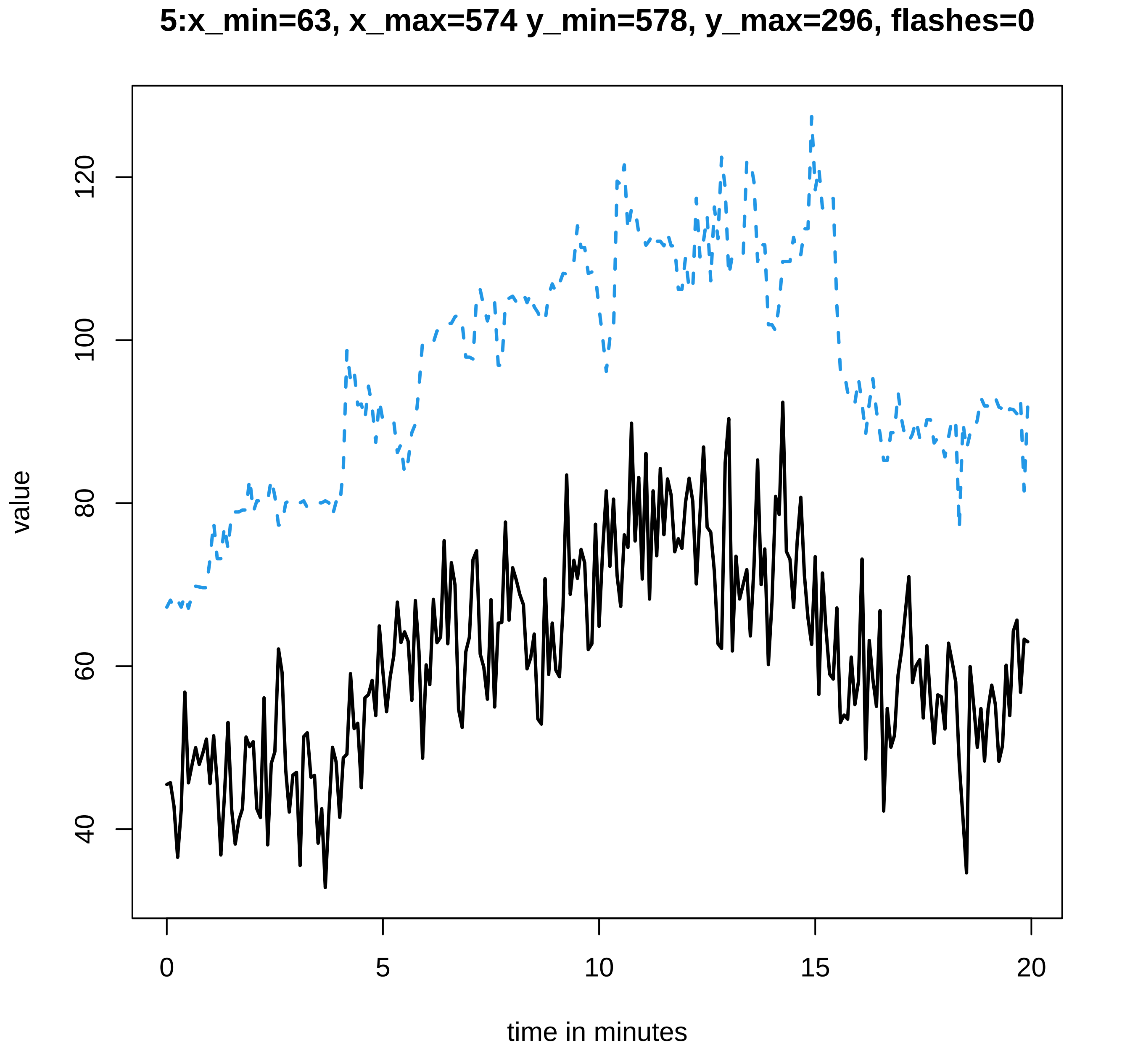
5:x_min=63, x_max=574 y_min=578, y_max=296, flashes=0
value
120
100
80
60
40
0
5
10
15
20
time in minutes

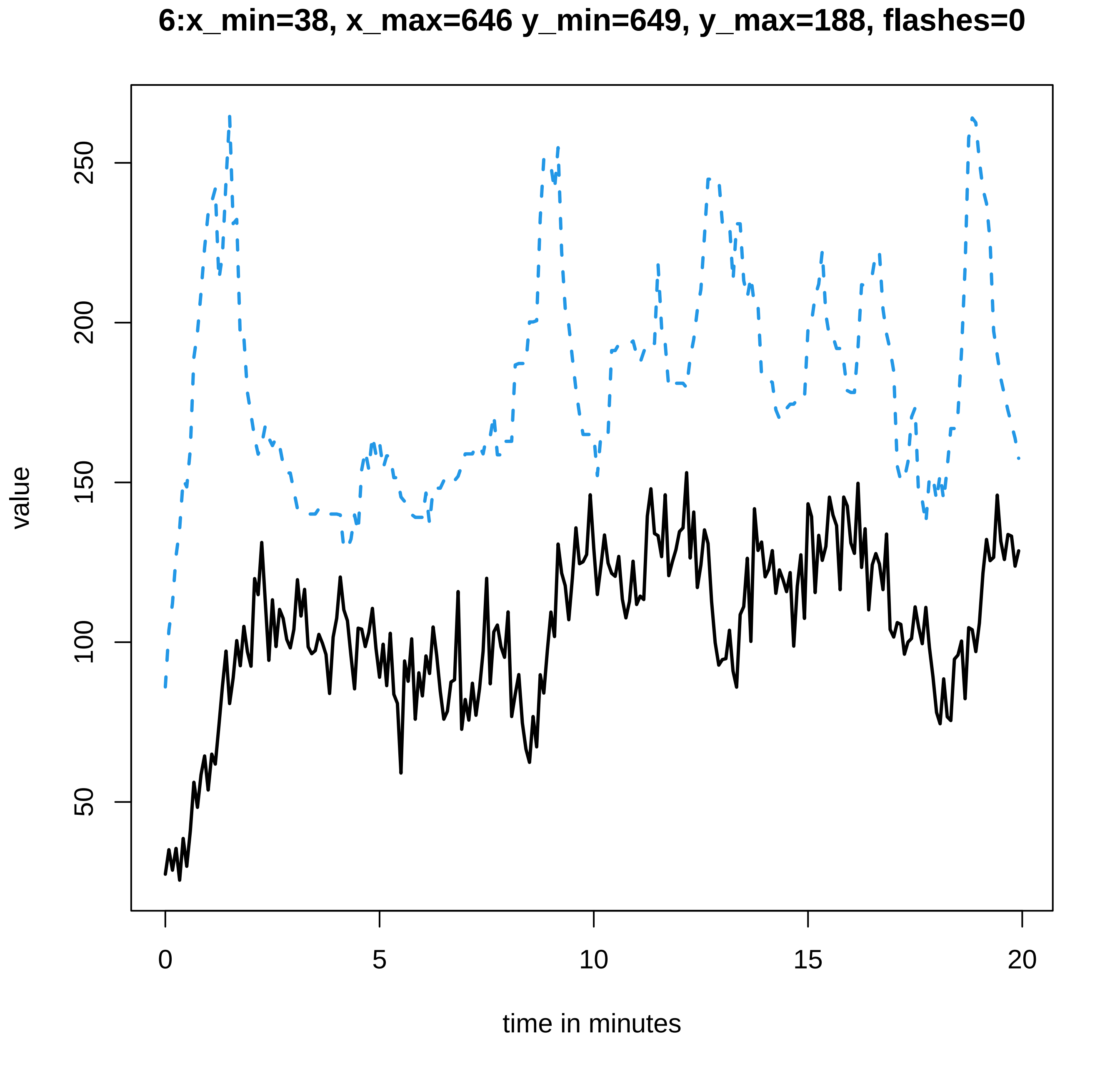

6:x_min=38, x_max=646 y_min=649, y_max=188, flashes=0
value
50
100
150
200
250
0
5
10
15
20
time in minutes

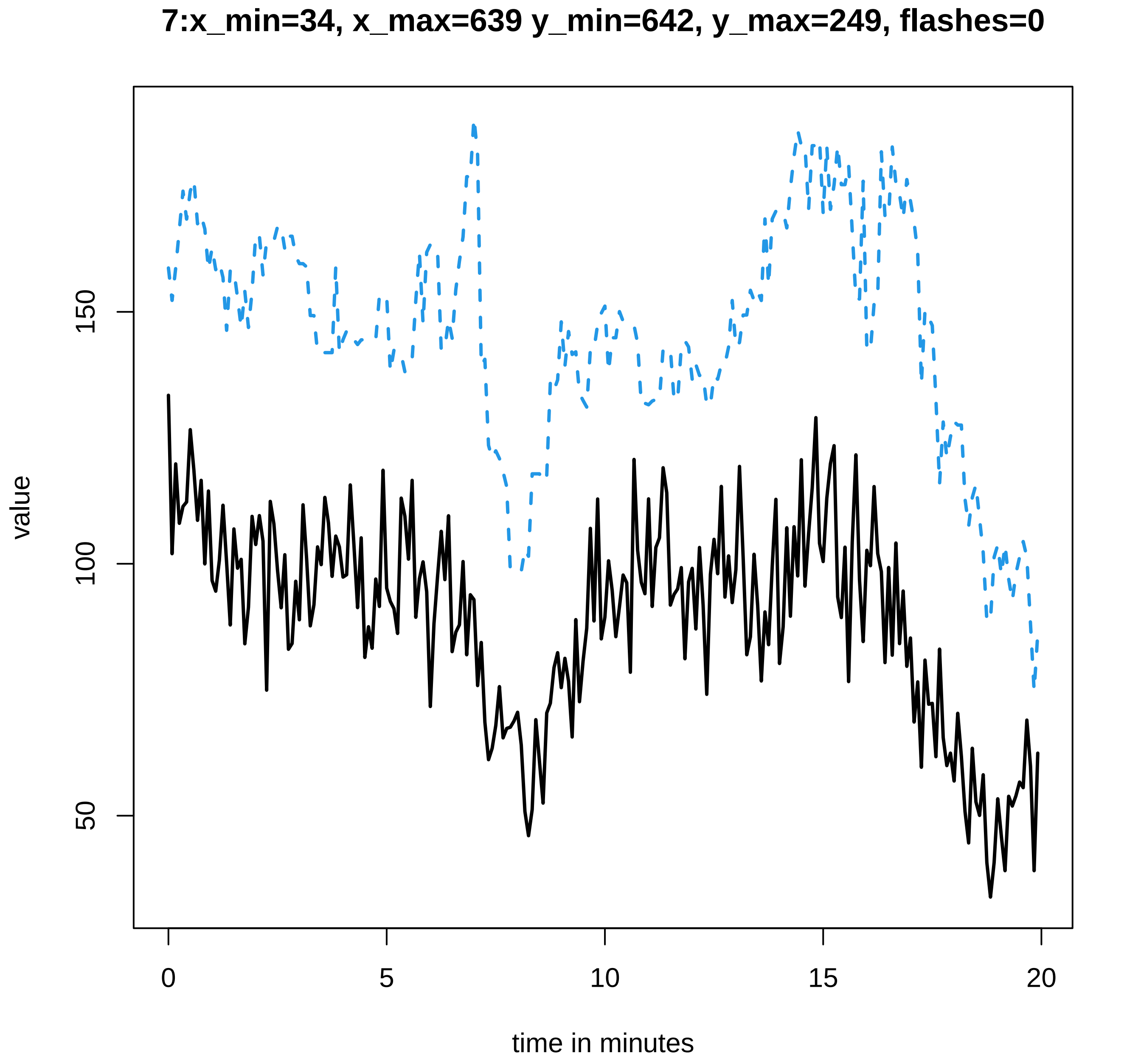

7:x_min=34, x_max=639 y_min=642, y_max=249, flashes=0
value
50
100
150
0
5
10
15
20
time in minutes

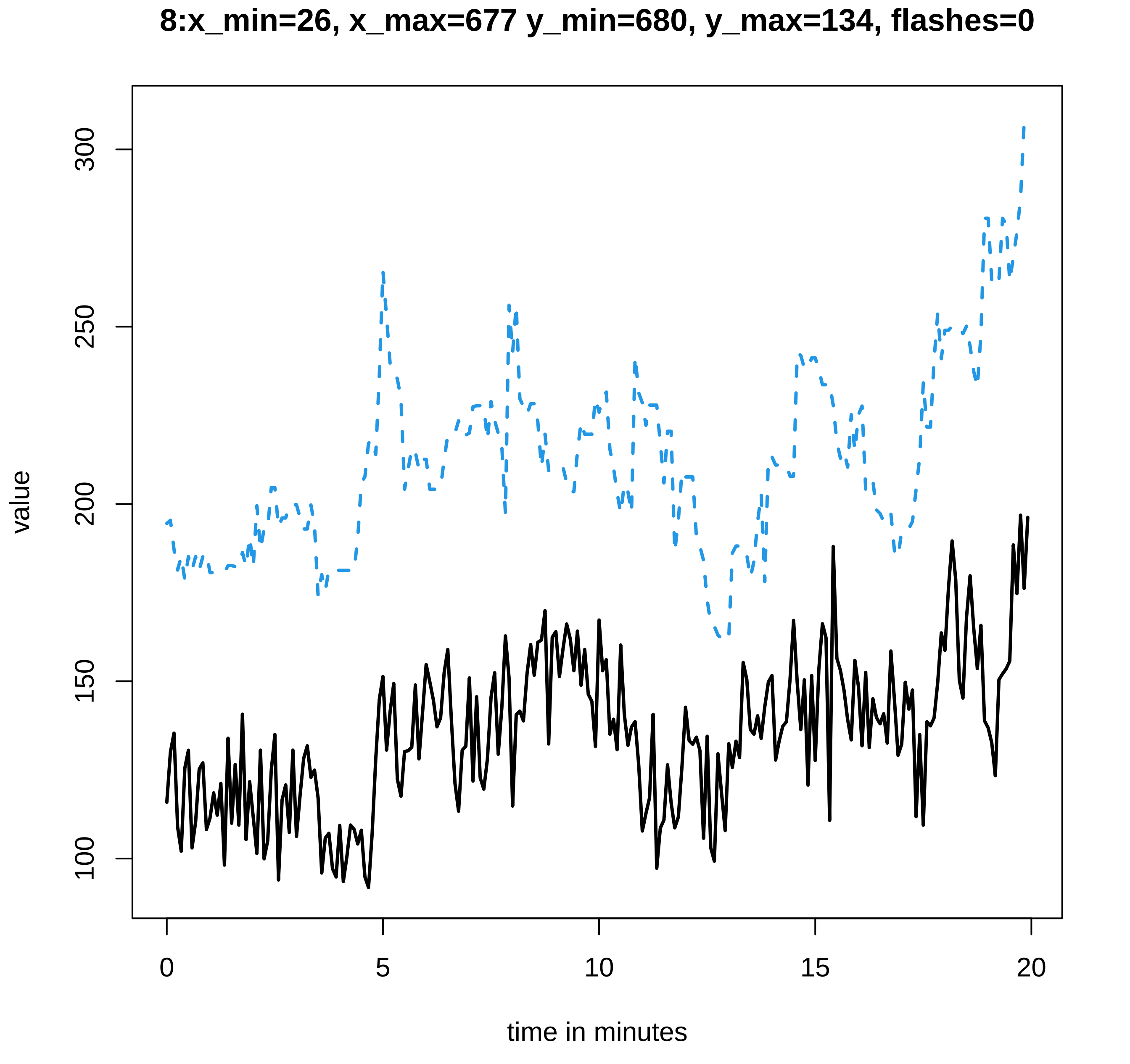
8:x_min=26, x_max=677 y_min=680, y_max=134, flashes=0
value
300
250
200
150
100
0
5
10
15
20
time in minutes

### 9:x_min=32, x_max=673 y_min=677, y_max=216, flashes=0

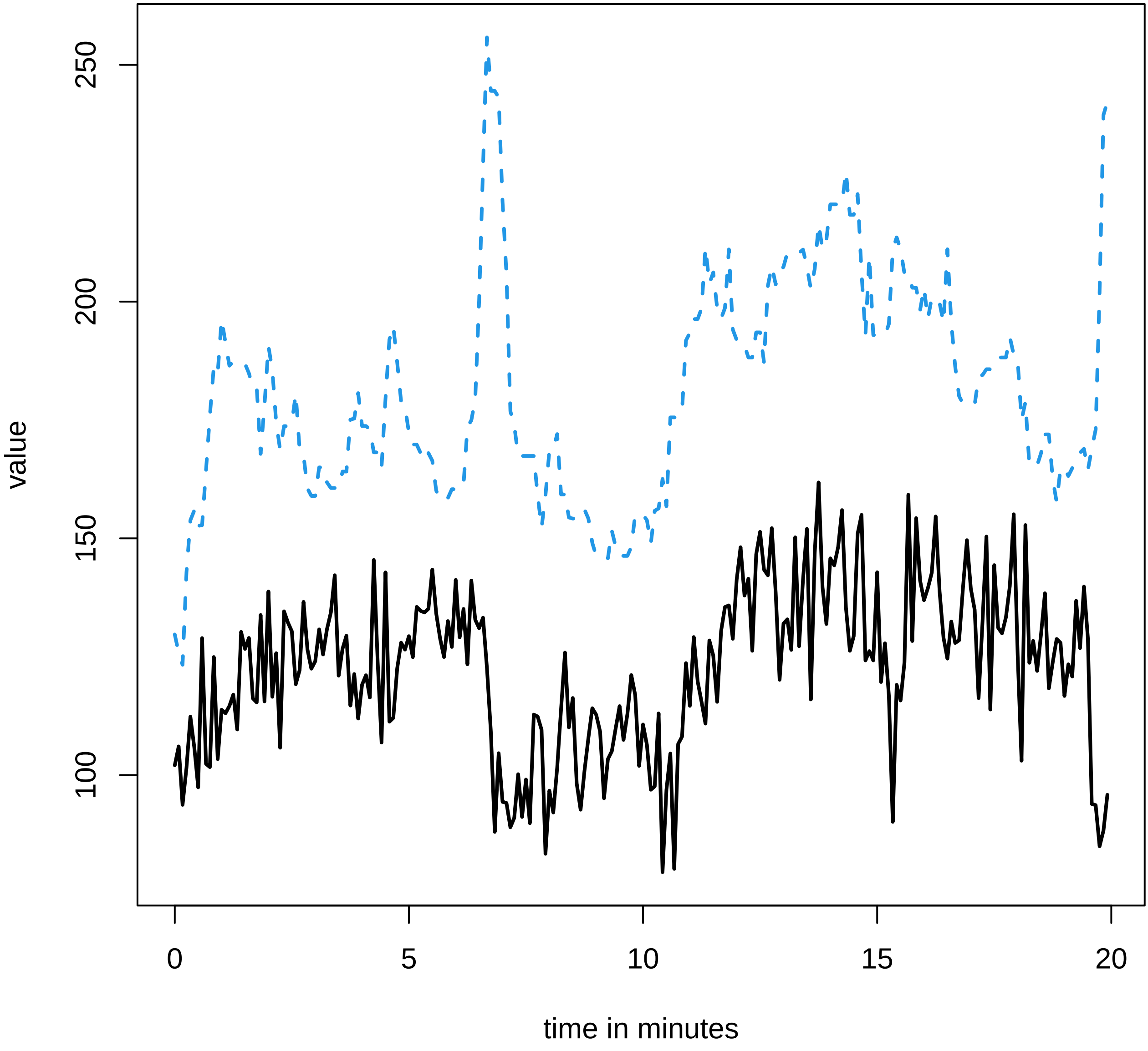

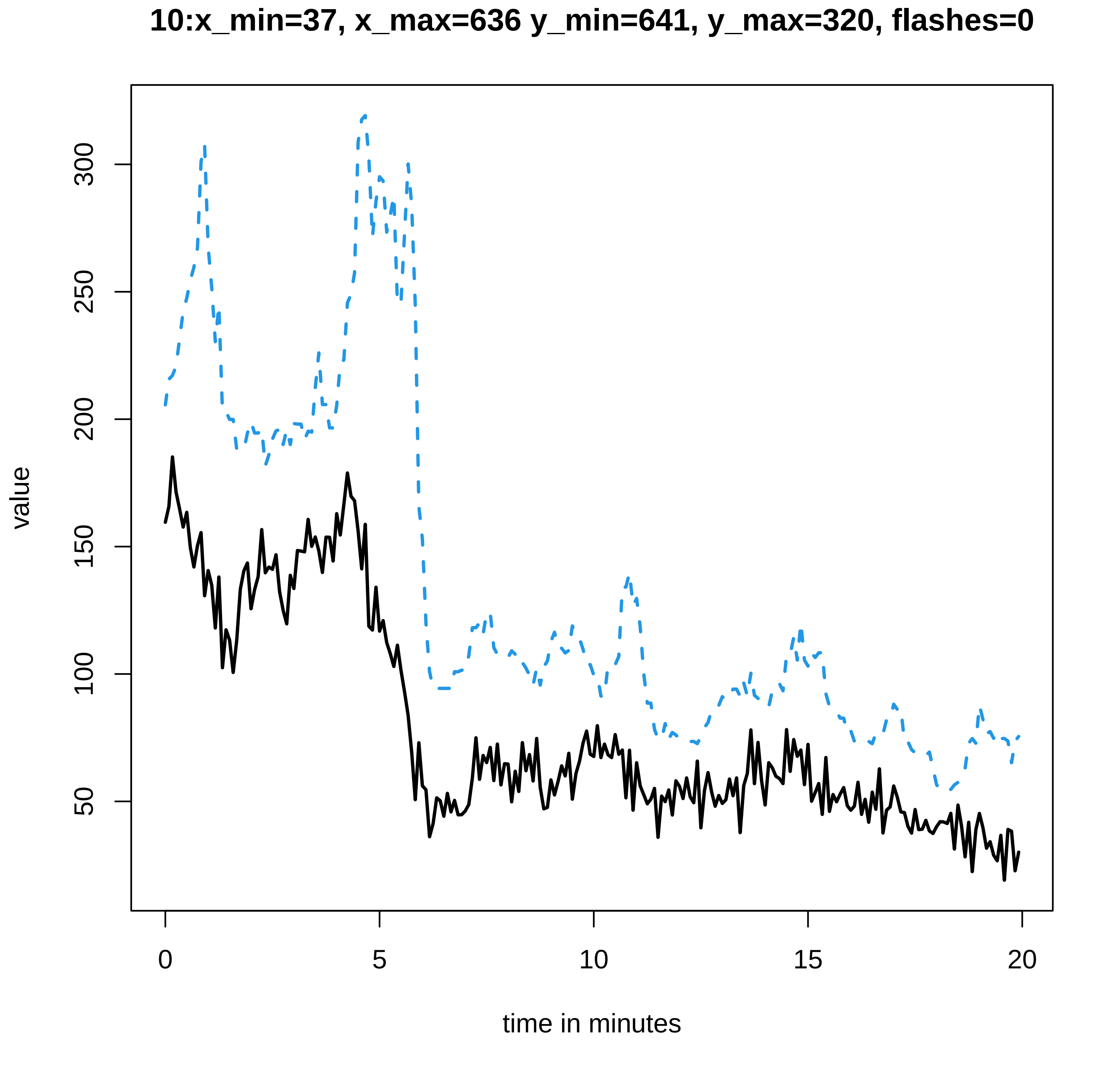
10:x_min=37, x_max=636 y_min=641, y_max=320, flashes=0
value
300
250
200
150
100
50
0
5
10
15
20
time in minutes

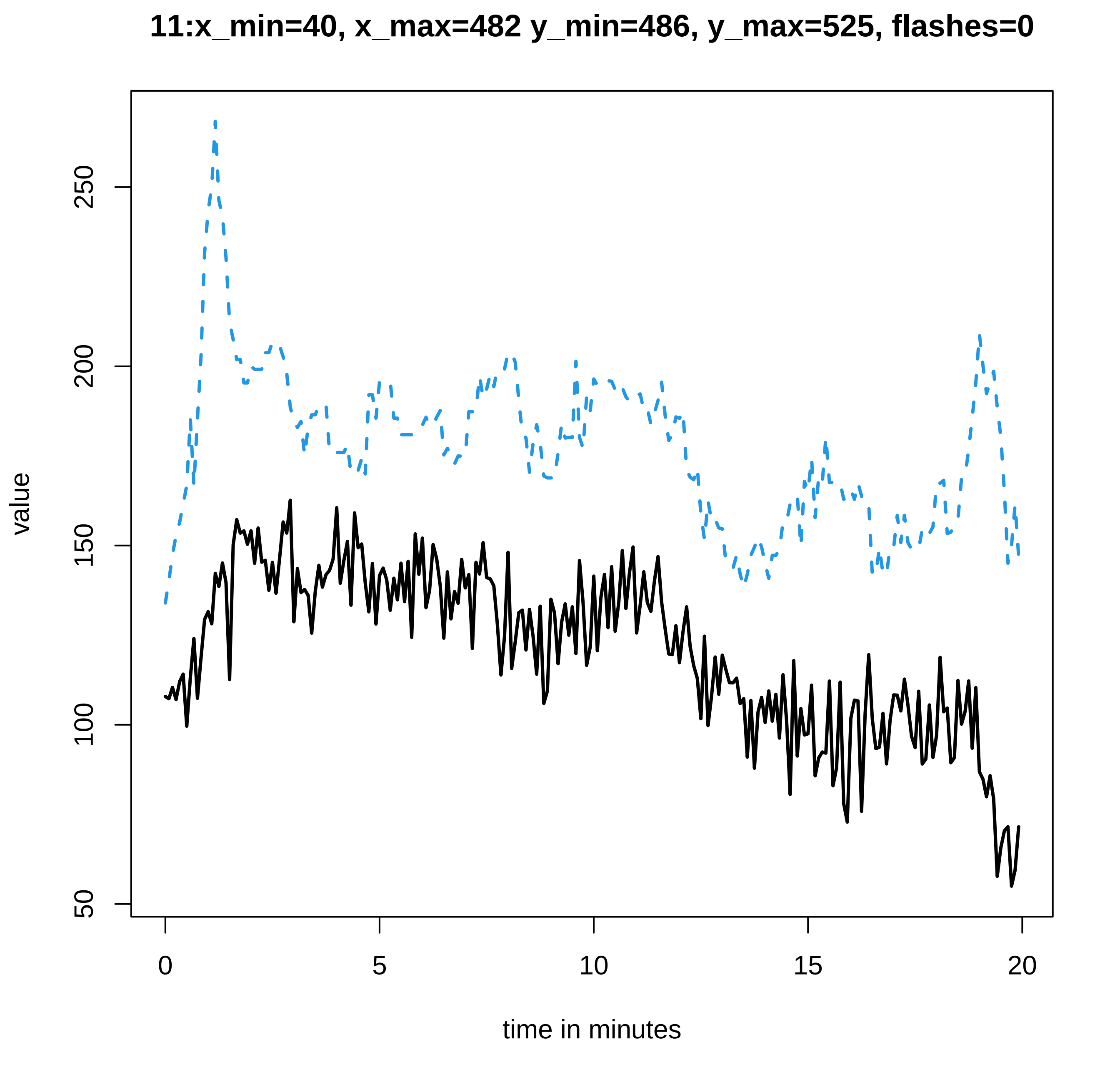
11:x_min=40, x_max=482 y_min=486, y_max=525, flashes=0
value
50
100
150
200
250
0
5
10
15
20
time in minutes

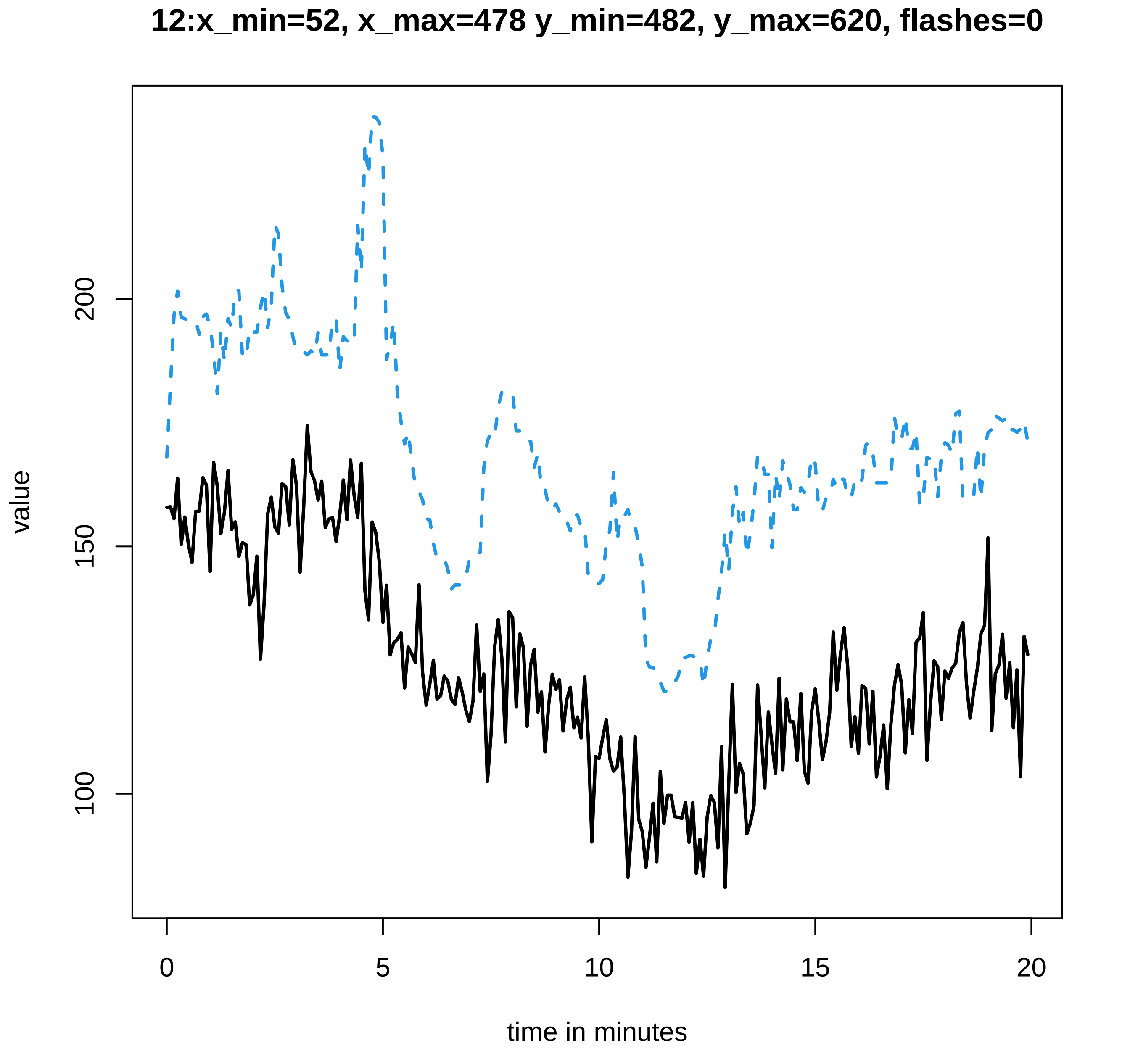

12:x_min=52, x_max=478 y_min=482, y_max=620, flashes=0
value
100
150
200
0
5
10
15
20
time in minutes

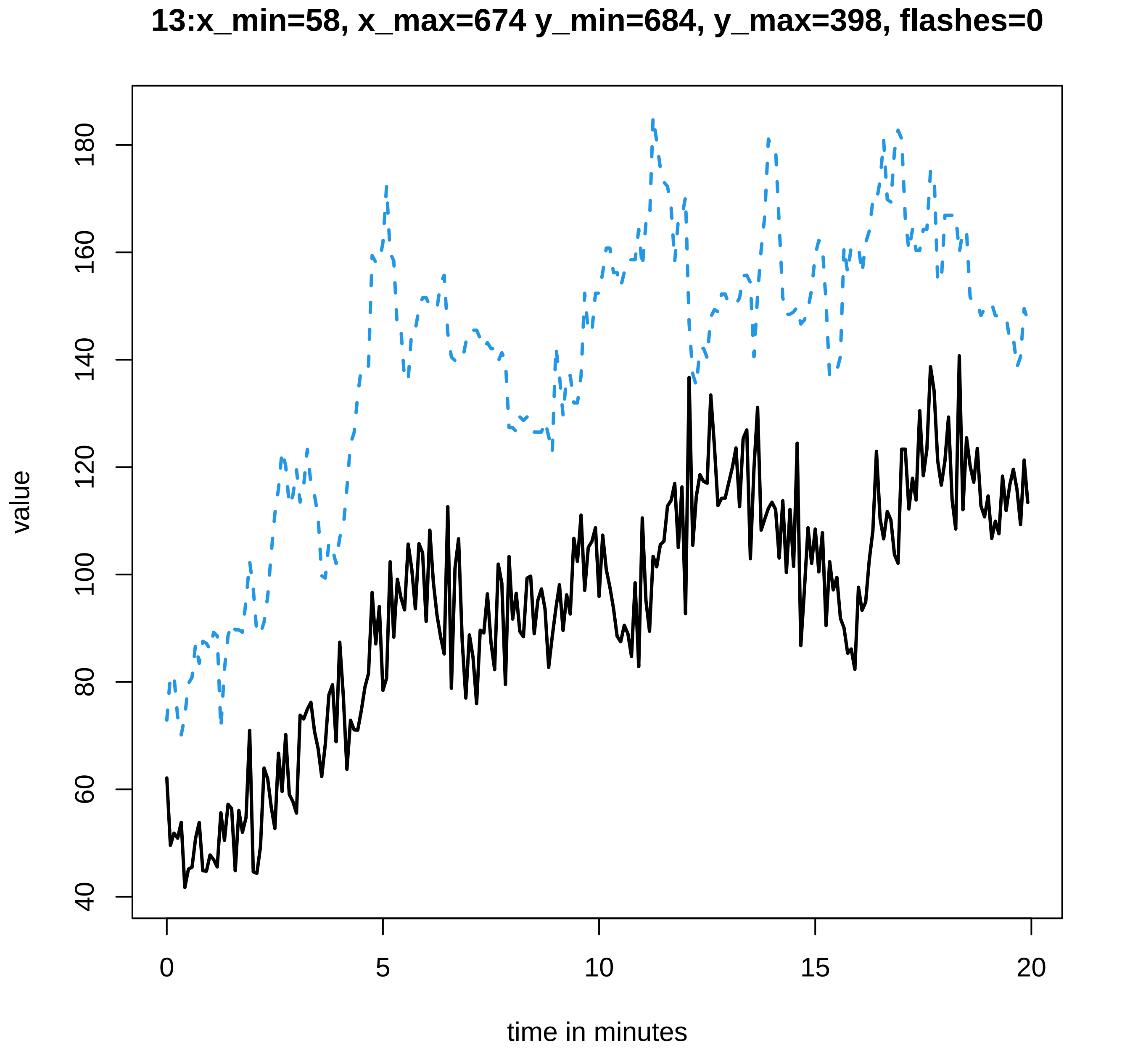
13:x_min=58, x_max=674 y_min=684, y_max=398, flashes=0
value
40
60
80
100
120
140
160
180
0
5
10
15
20
time in minutes

#### 14:x_min=34, x_max=599 y_min=602, y_max=515, flashes=0

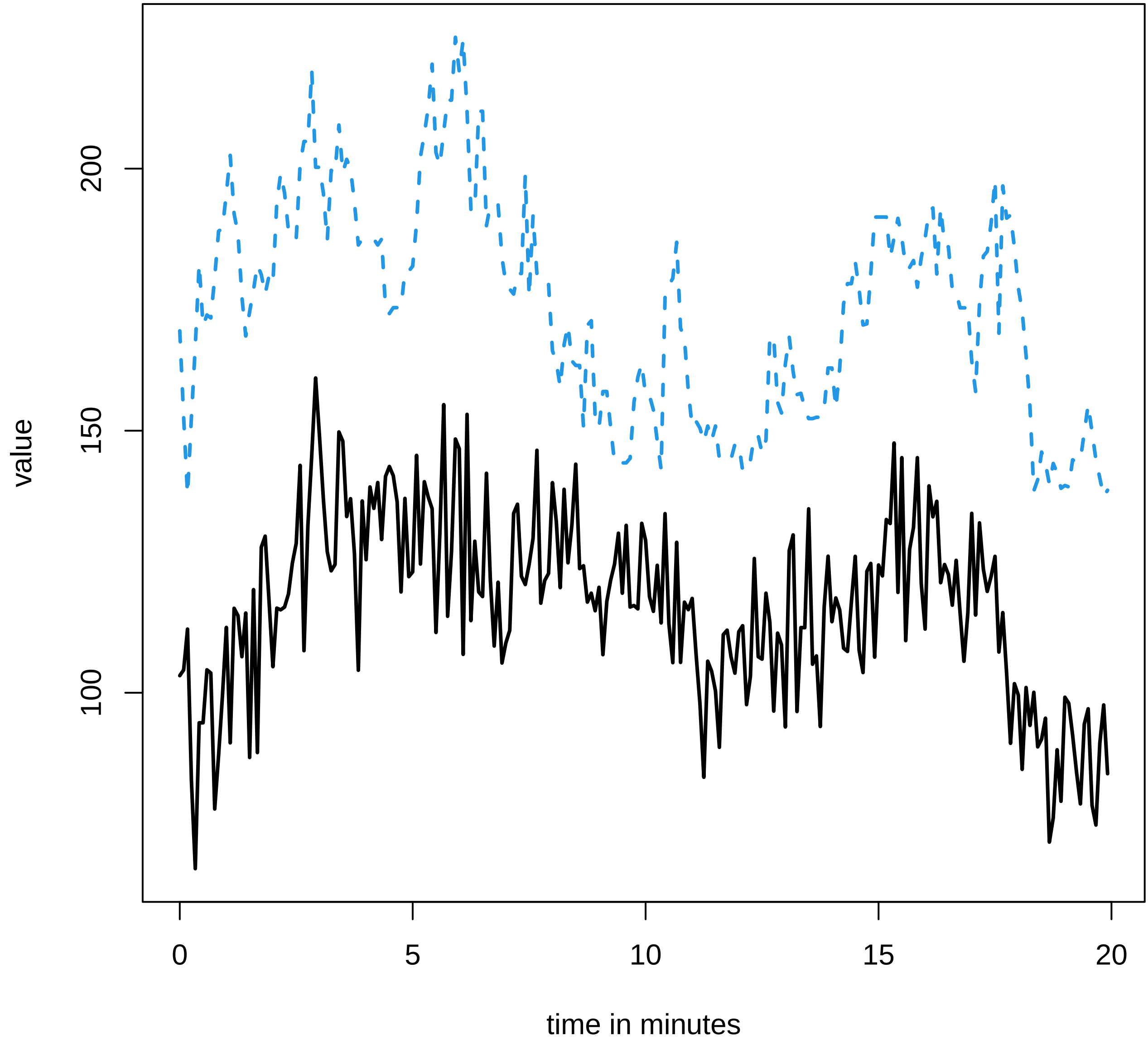

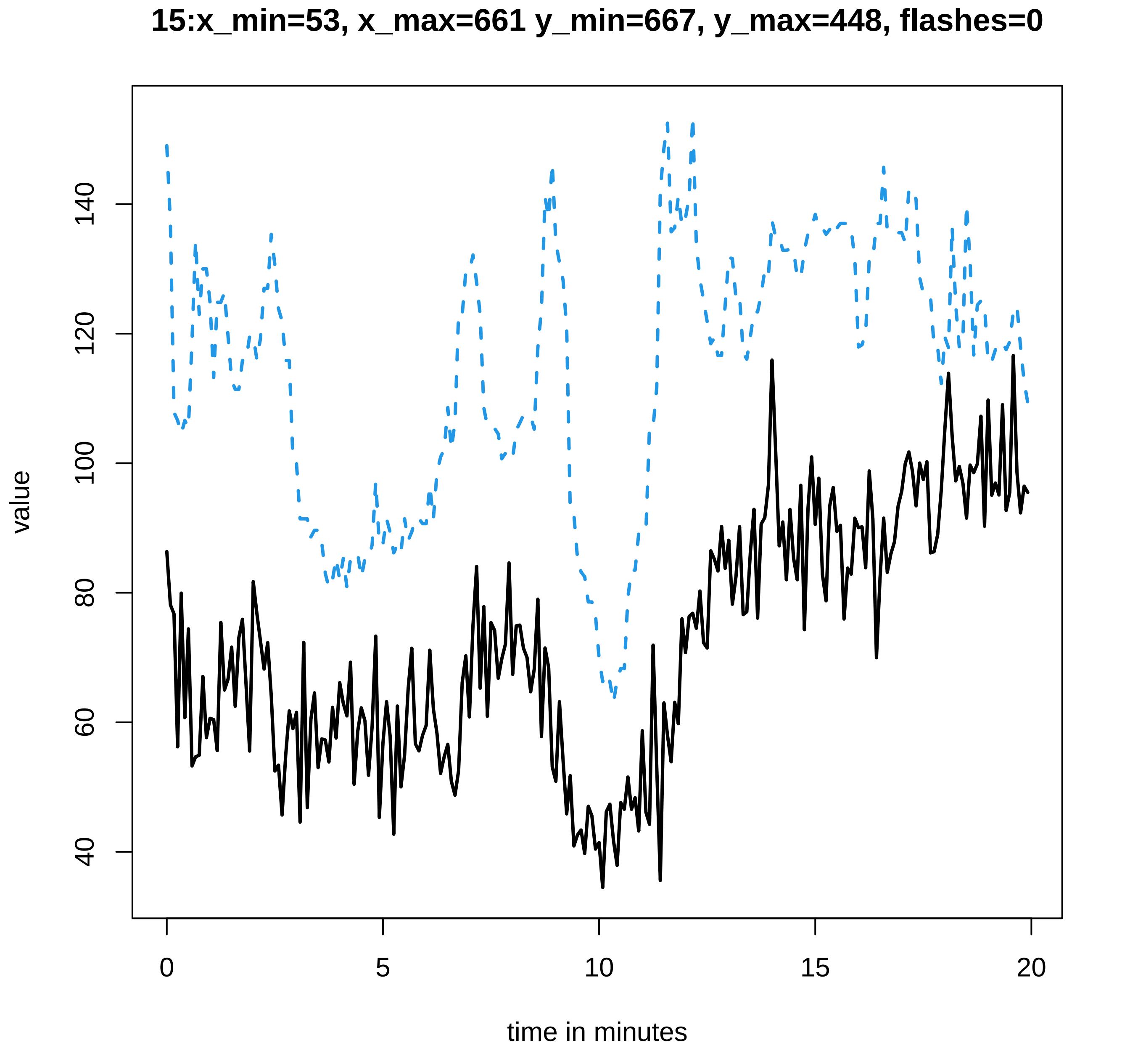

15:x_min=53, x_max=661 y_min=667, y_max=448, flashes=0
value
40
60
80
100
120
140
0
5
10
15
20
time in minutes

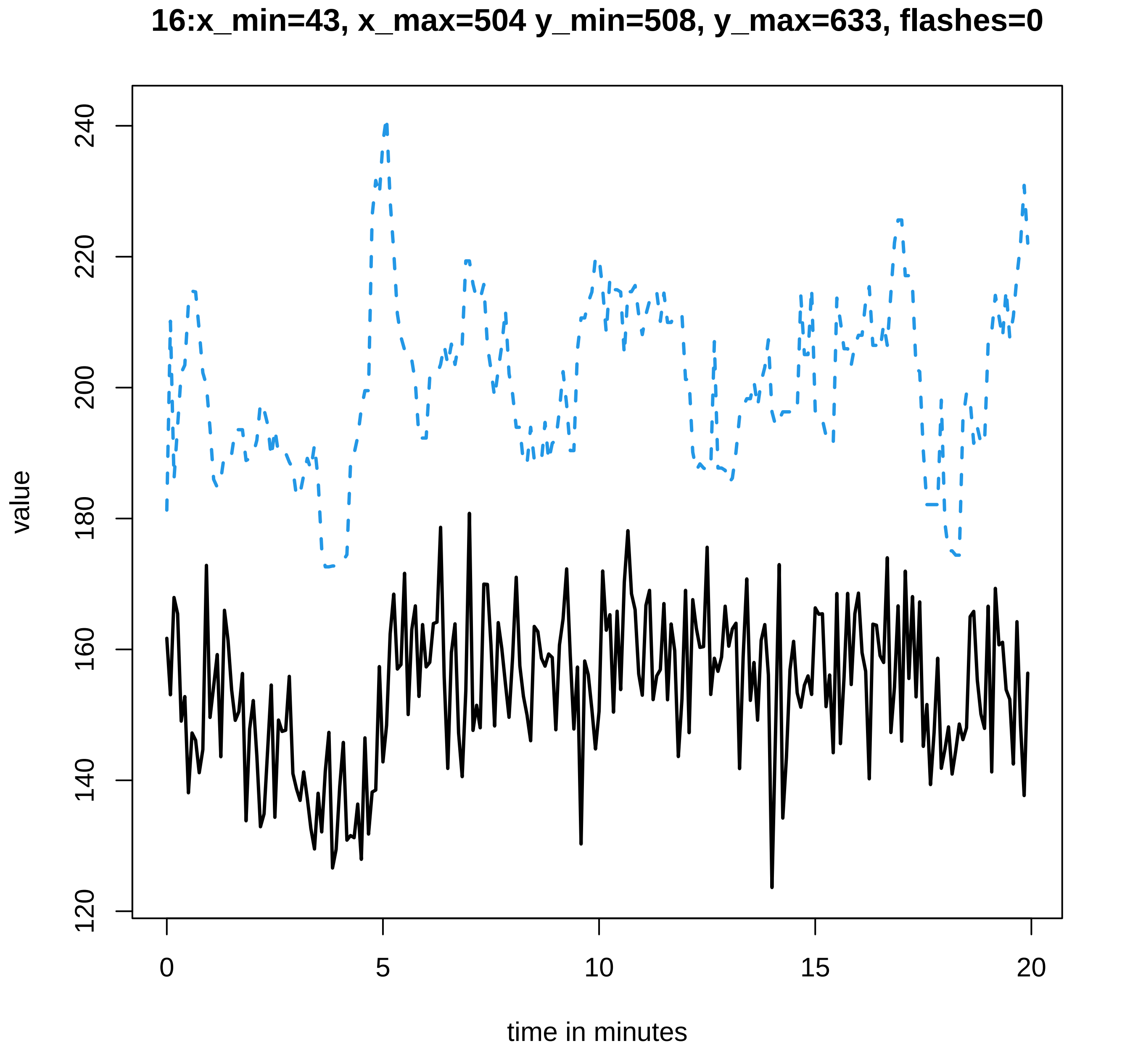
16:x_min=43, x_max=504 y_min=508, y_max=633, flashes=0
value
time in minutes
120
140
160
180
200
220
240
0
5
10
15
20

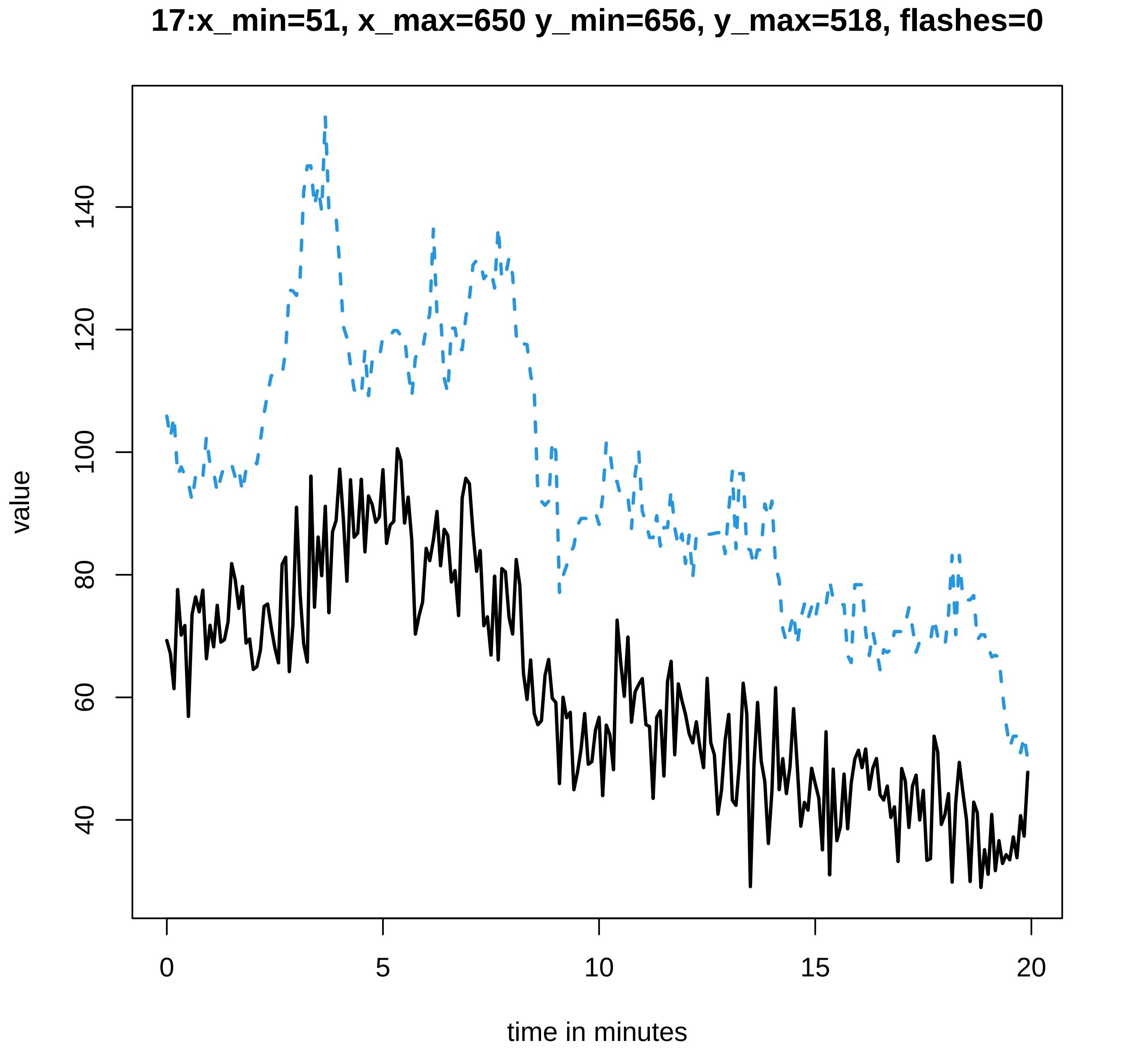
17:x_min=51, x_max=650 y_min=656, y_max=518, flashes=0
value
40
60
80
100
120
140
0
5
10
15
20
time in minutes

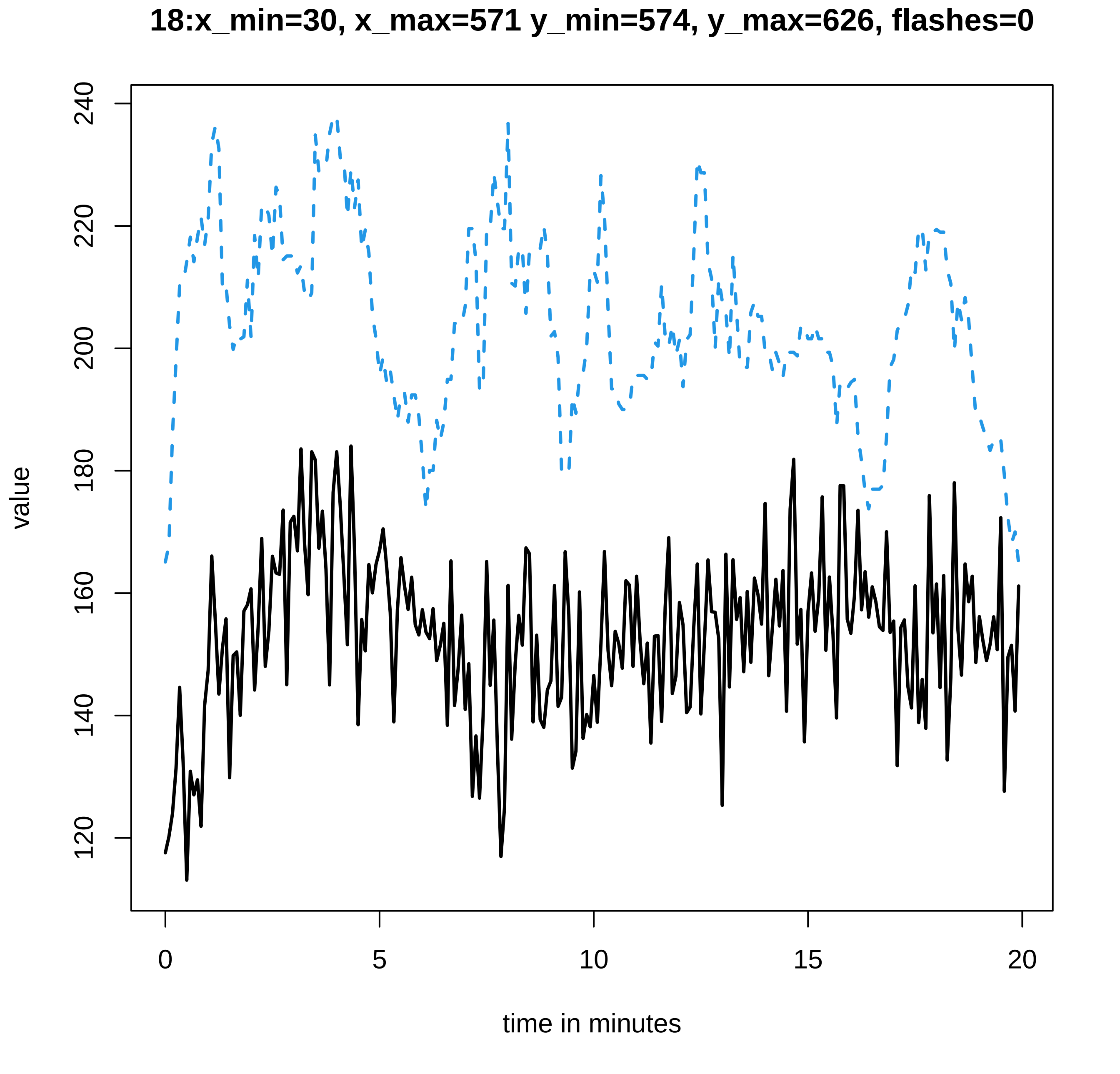
18:x_min=30, x_max=571 y_min=574, y_max=626, flashes=0
value
time in minutes
120
140
160
180
200
220
240
0
5
10
15
20

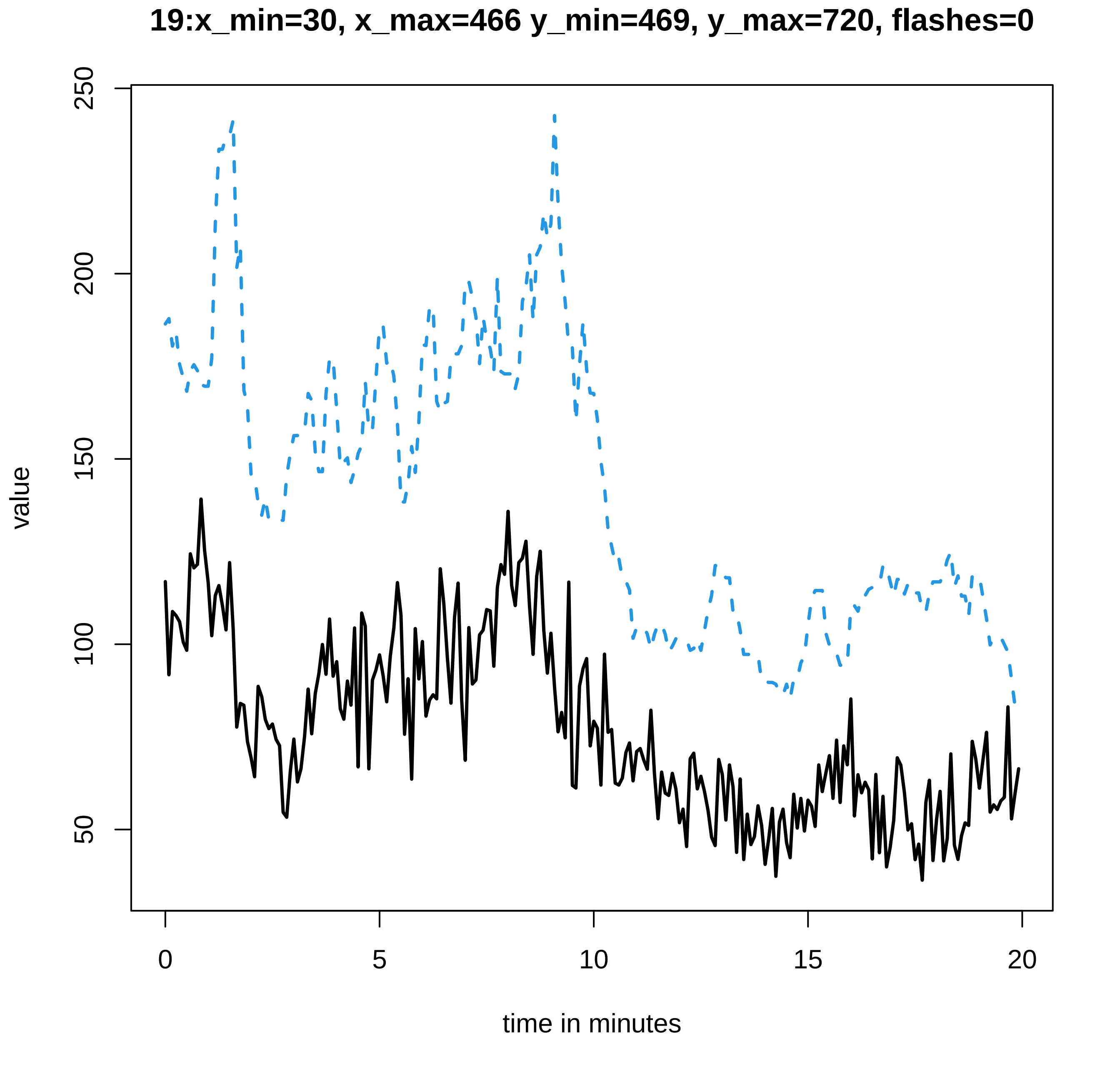
19:x_min=30, x_max=466 y_min=469, y_max=720, flashes=0
value
250
200
150
100
50
0
5
10
15
20
time in minutes

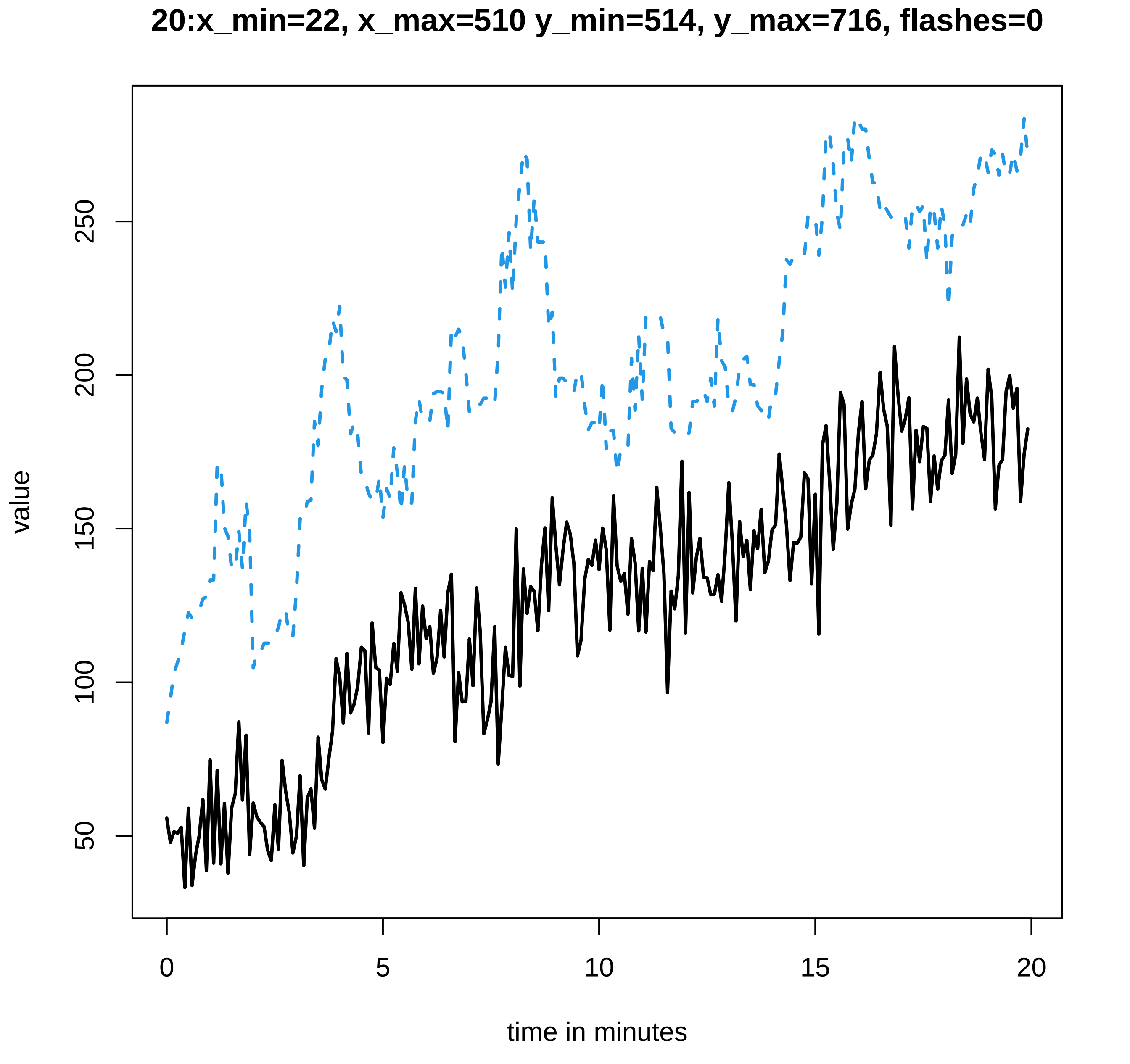
20:x_min=22, x_max=510 y_min=514, y_max=716, flashes=0
value
50
100
150
200
250
0
5
10
15
20
time in minutes

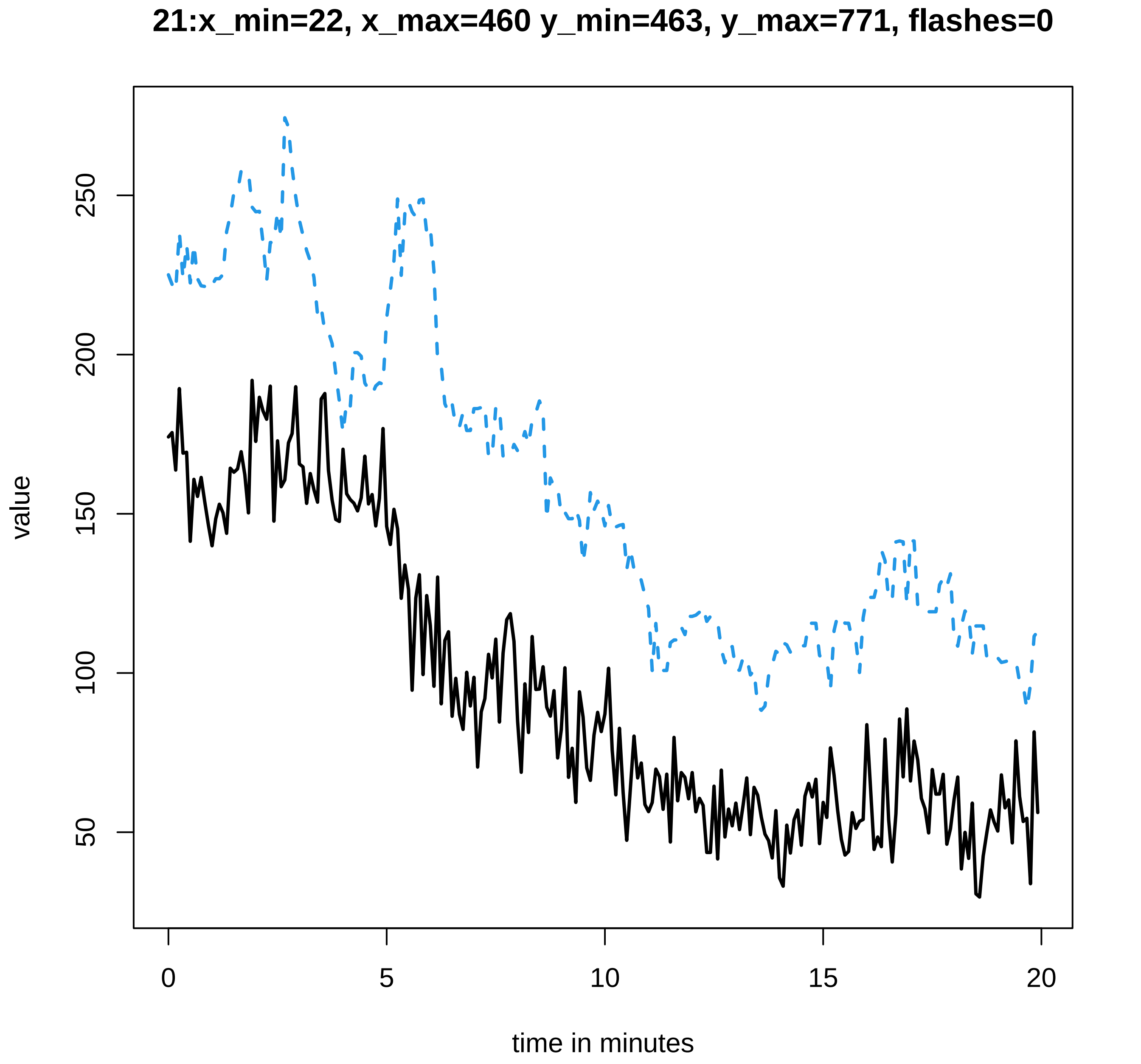
21:x_min=22, x_max=460 y_min=463, y_max=771, flashes=0
value
50
100
150
200
250
0
5
10
15
20
time in minutes

#### 22:x_min=36, x_max=659 y_min=663, y_max=665, flashes=0

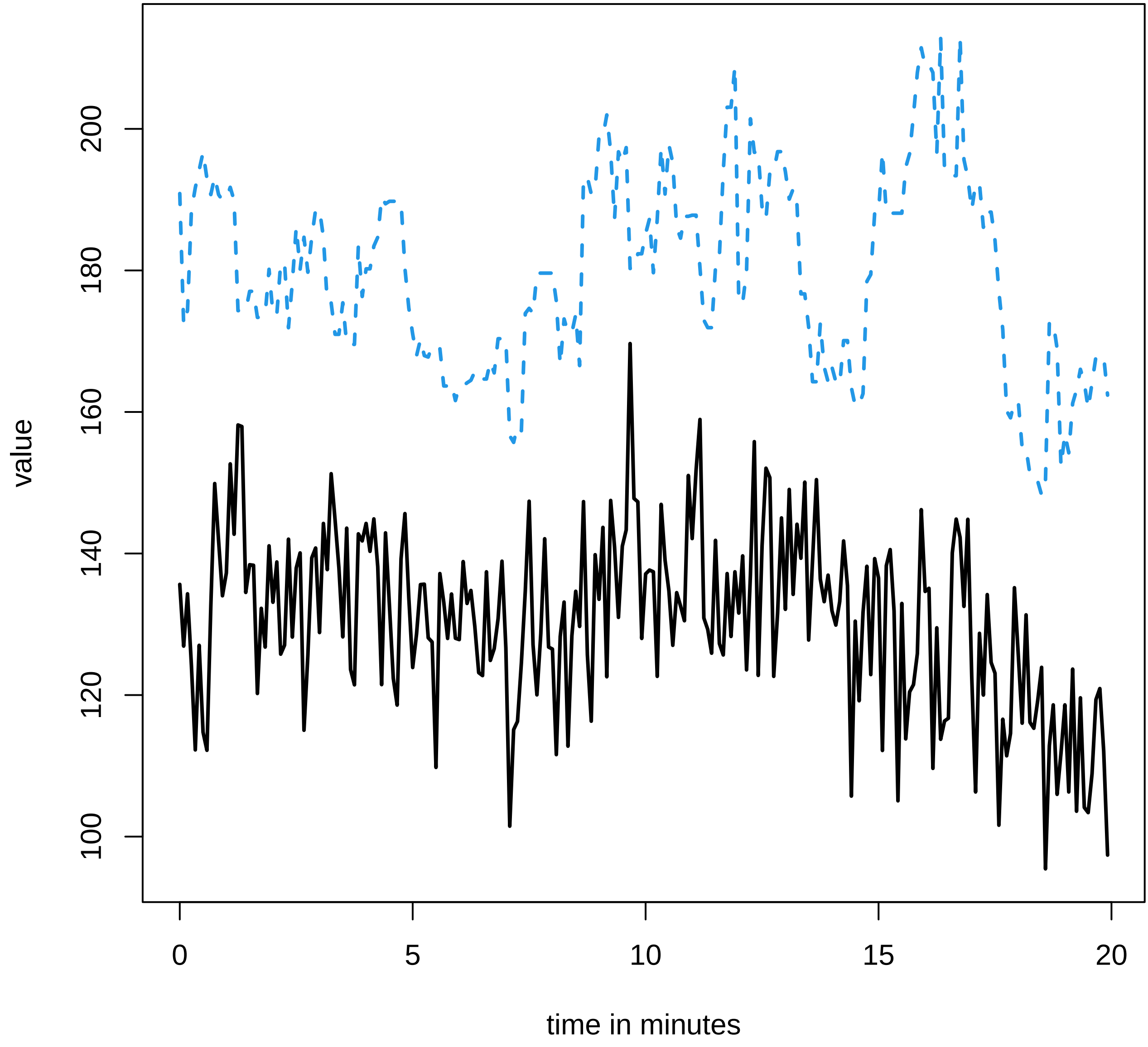

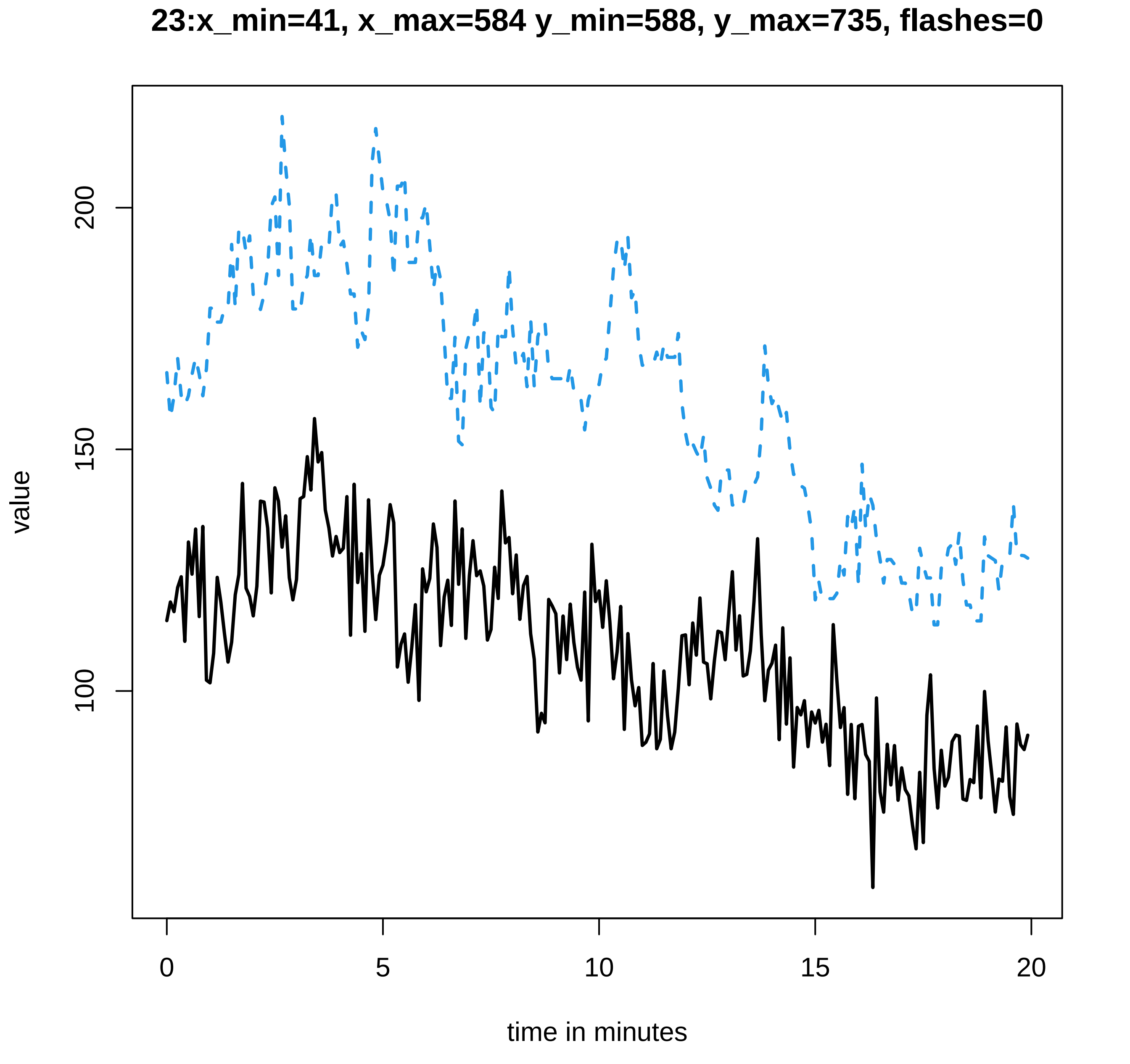
23:x_min=41, x_max=584 y_min=588, y_max=735, flashes=0
value
200
150
100
0
5
10
15
20
time in minutes

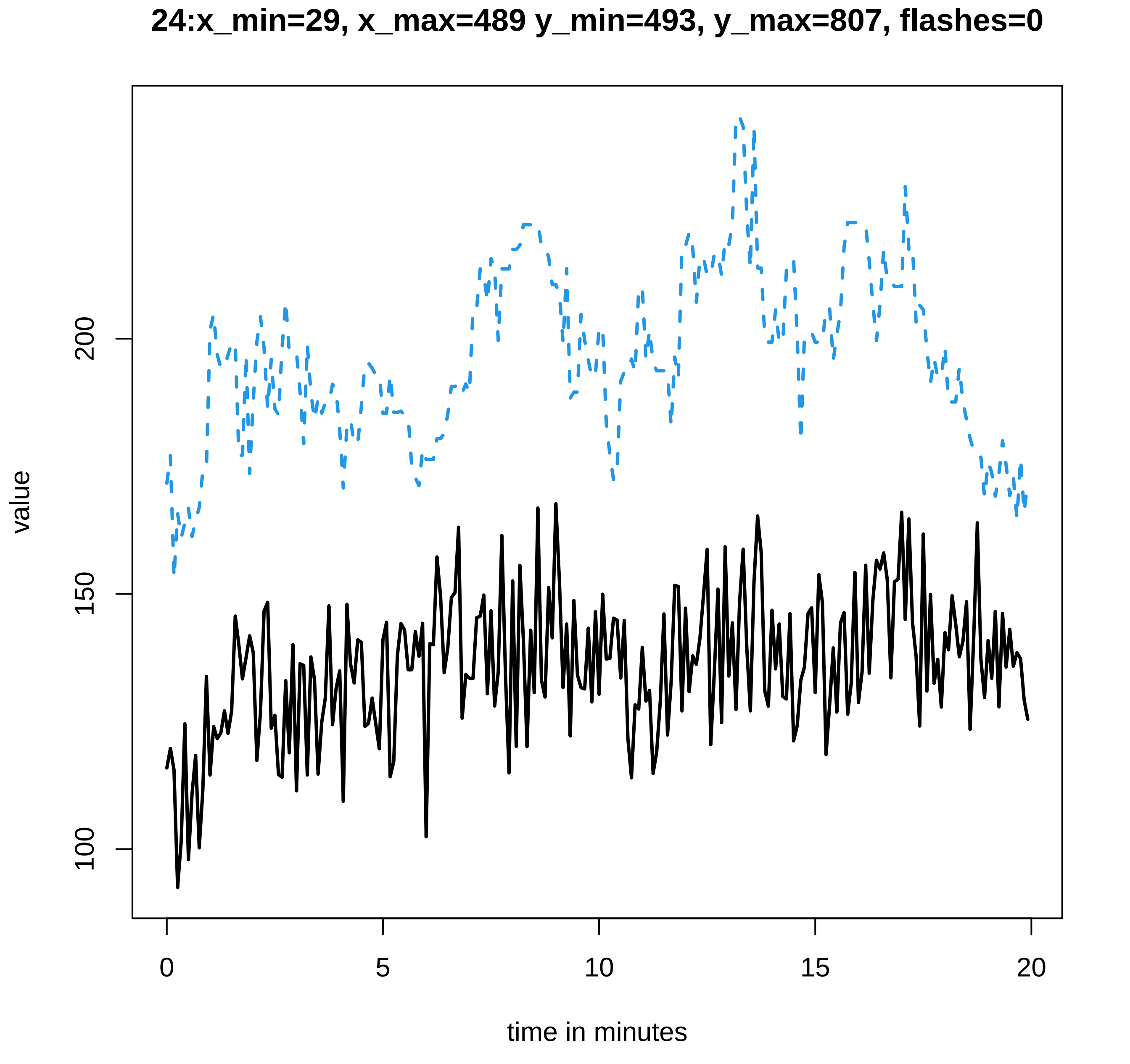

24:x_min=29, x_max=489 y_min=493, y_max=807, flashes=0
value
100
150
200
0
5
10
15
20
time in minutes

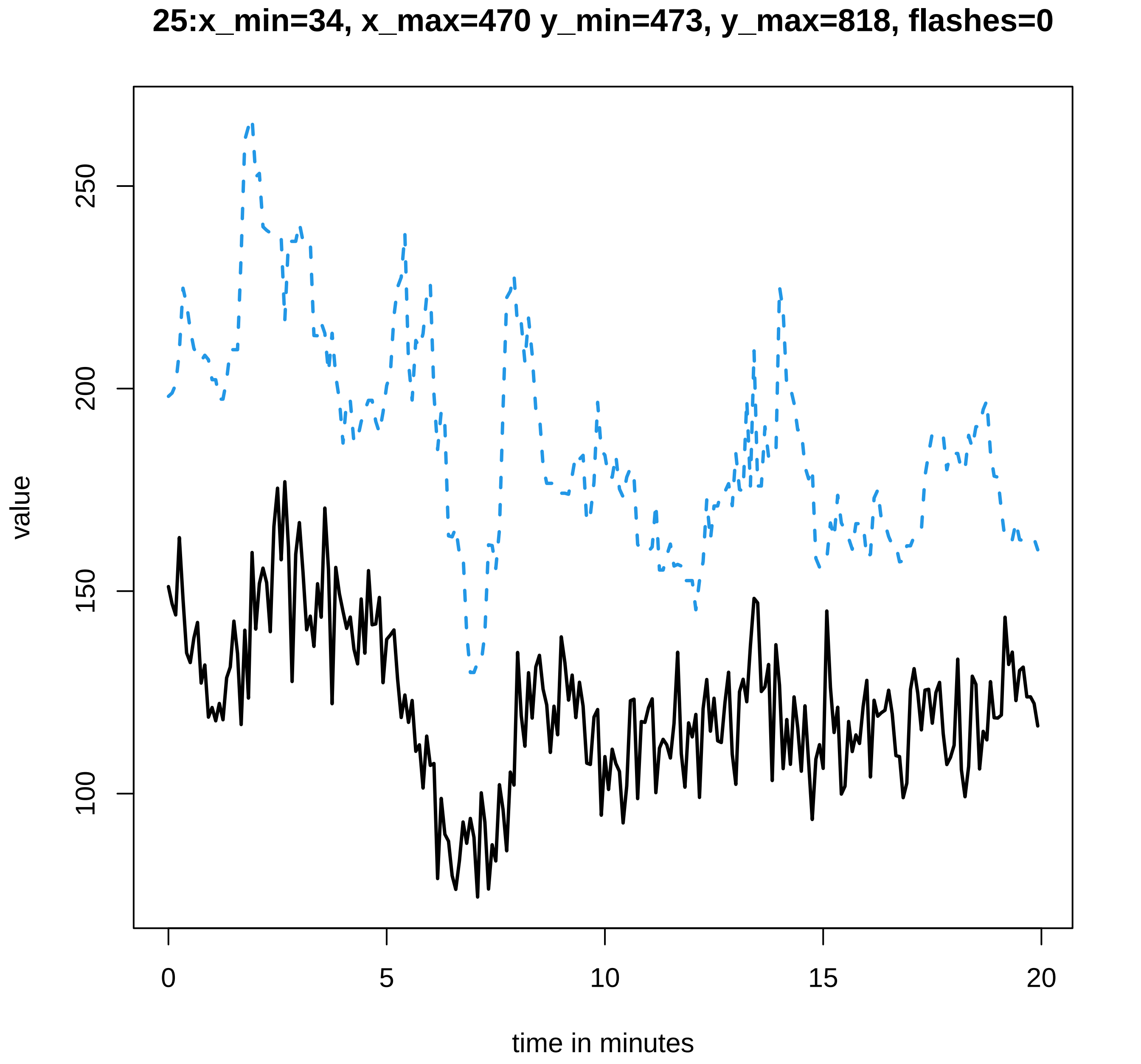

25:x_min=34, x_max=470 y_min=473, y_max=818, flashes=0
value
100
150
200
250
0
5
10
15
20
time in minutes

**26:x_min=26, x_max=647 y_min=650, y_max=750, flashes=1**

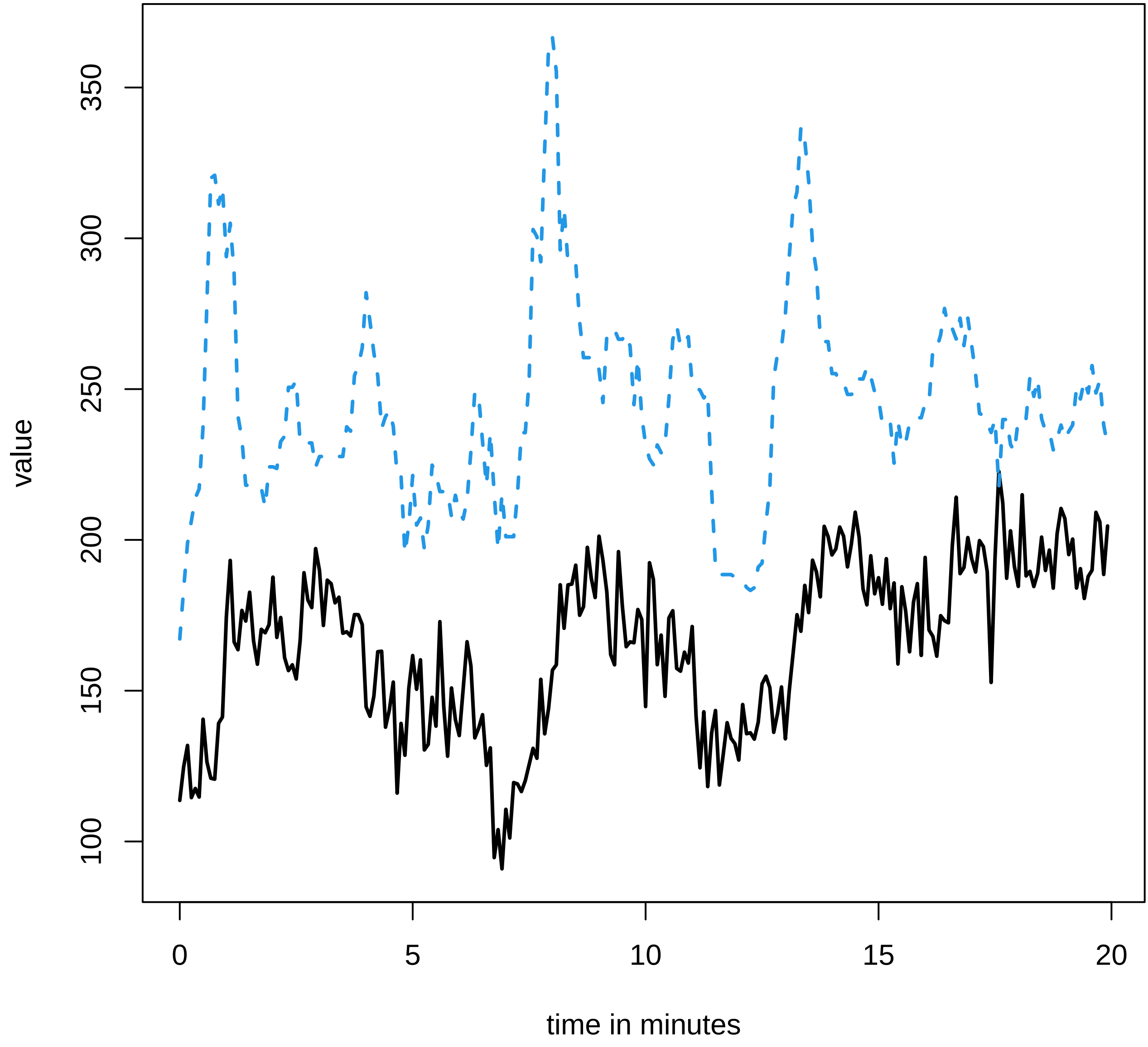

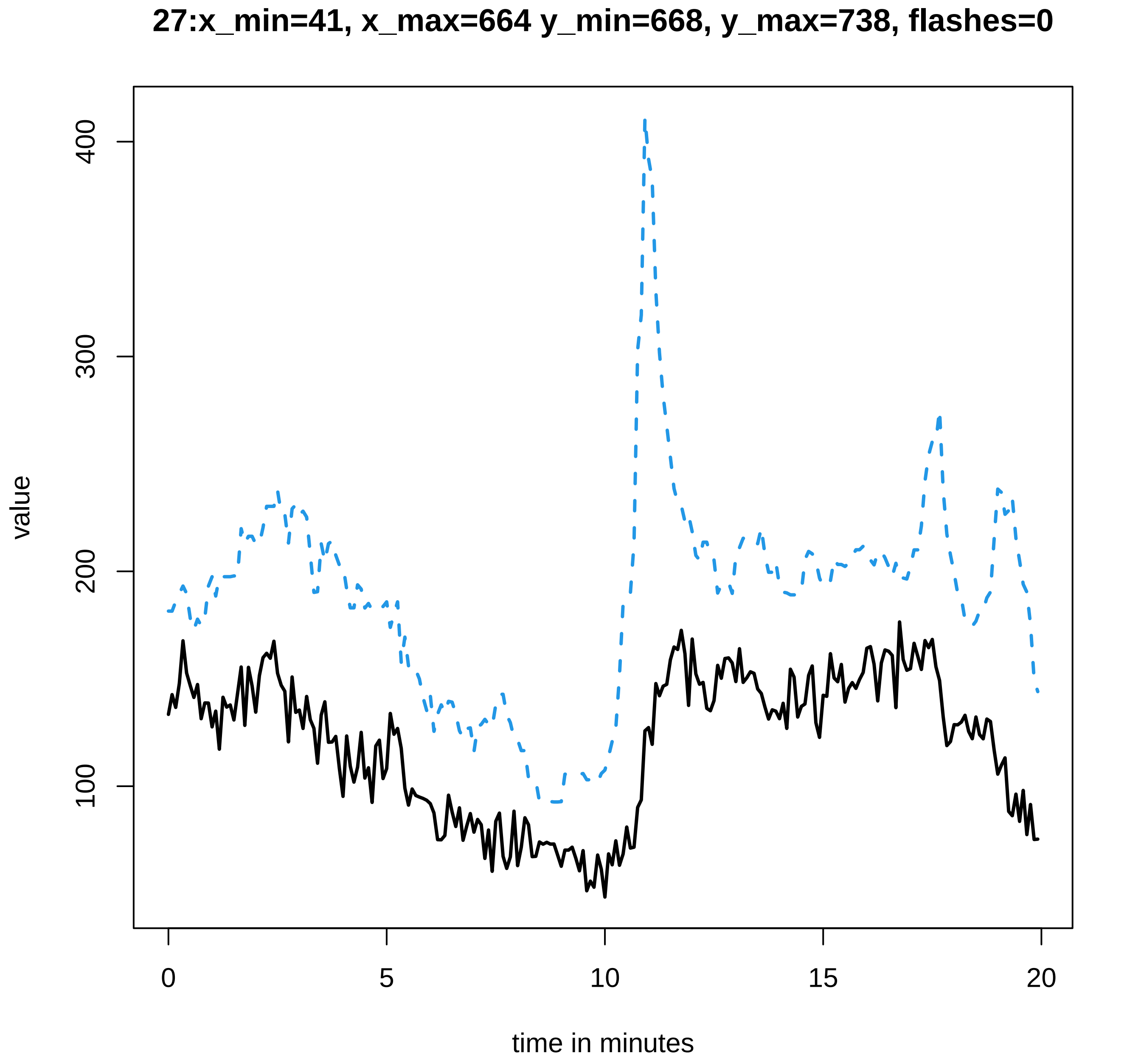
27:x_min=41, x_max=664 y_min=668, y_max=738, flashes=0
value
100
200
300
400
0
5
10
15
20
time in minutes

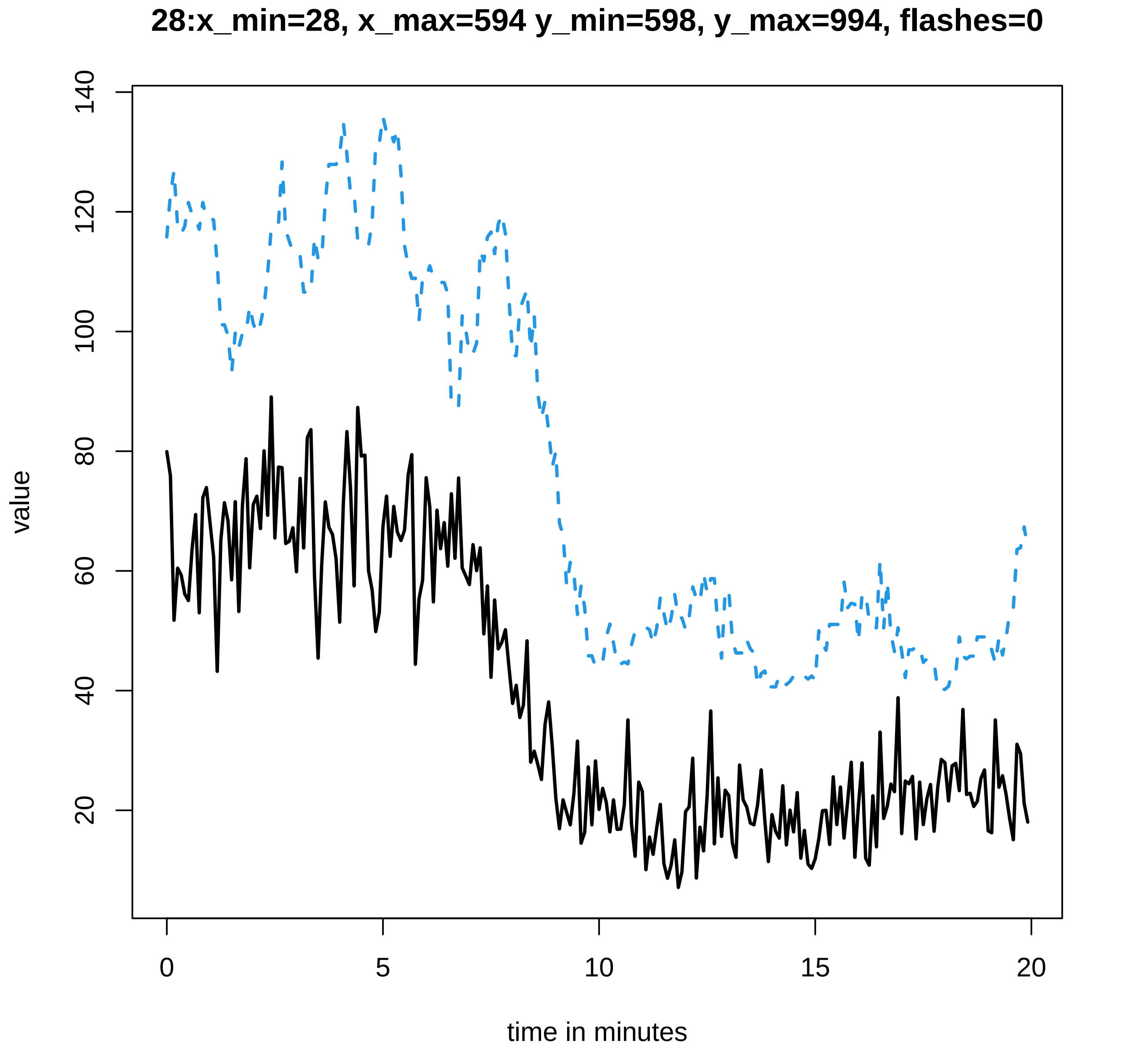
28:x_min=28, x_max=594 y_min=598, y_max=994, flashes=0
value
20
40
60
80
100
120
140
0
5
10
15
20
time in minutes
