## Supplementary data Calcium spiking for "LysM Receptor Proteins are Required for Ectomycorrhizal Symbiosis in Poplar": profiles-spiking-18.pdf

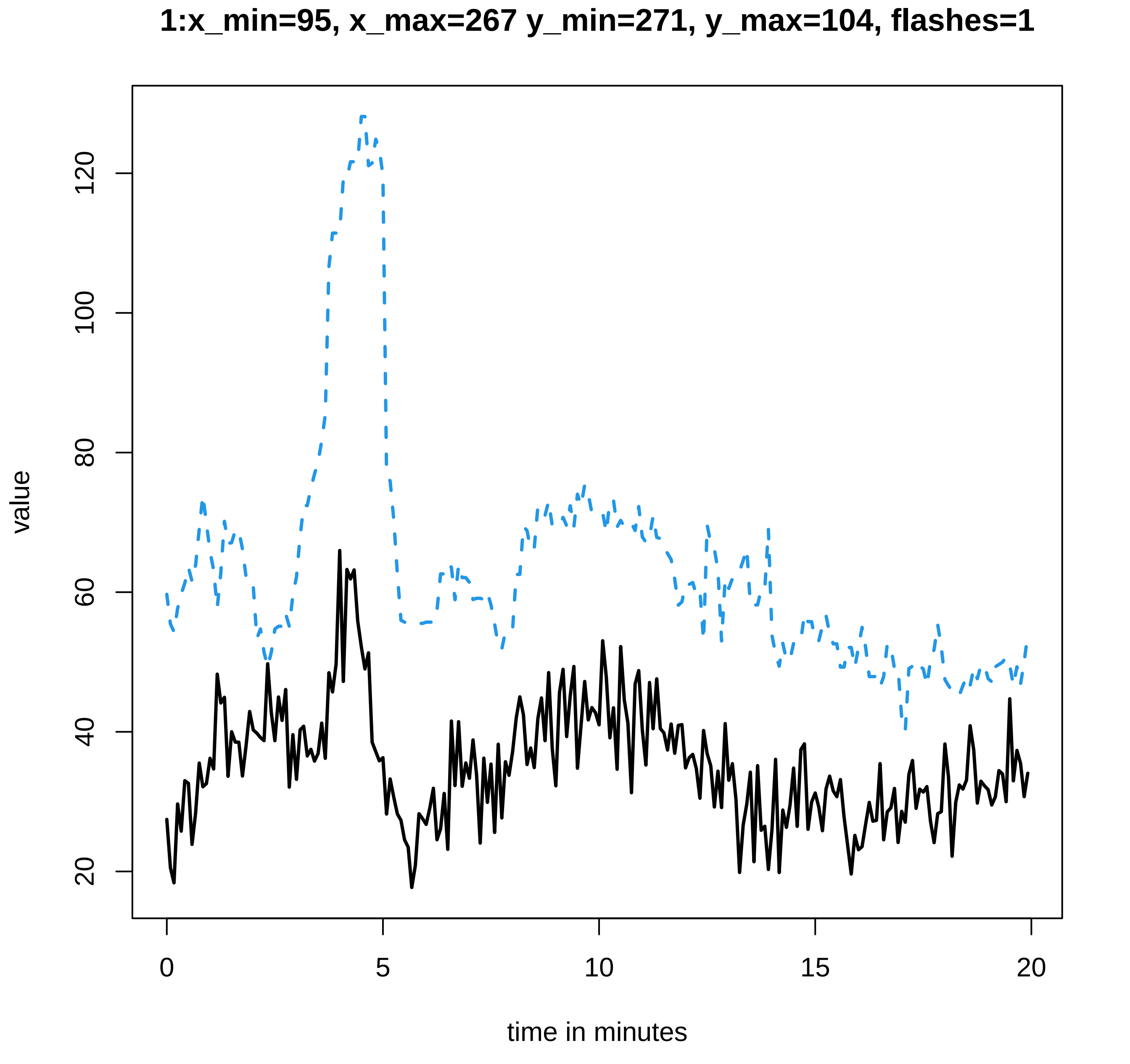
1:x_min=95, x_max=267 y_min=271, y_max=104, flashes=1
value
time in minutes
20
40
60
80
100
120
0
5
10
15
20

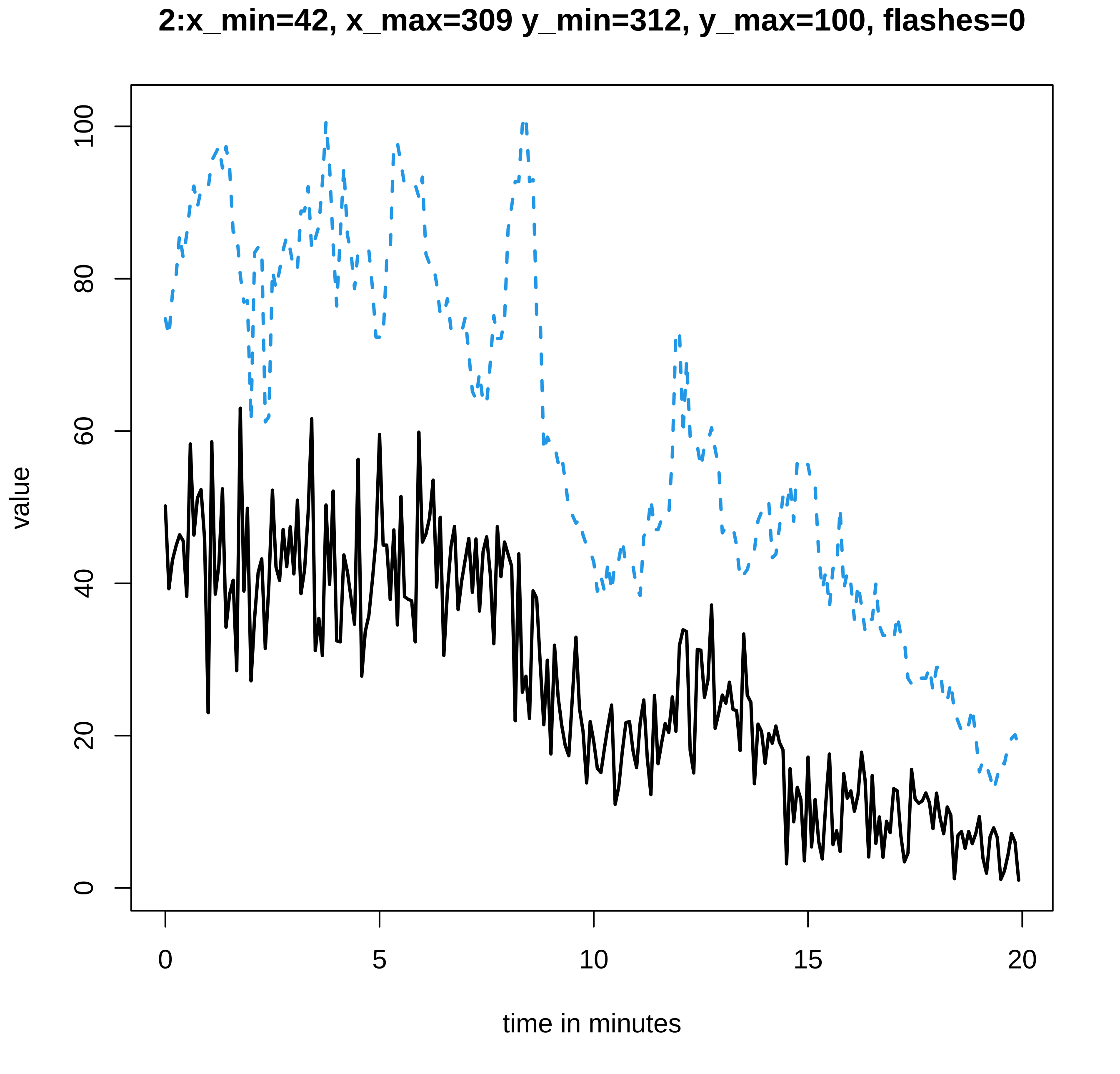

2:x_min=42, x_max=309 y_min=312, y_max=100, flashes=0
value
time in minutes
0
5
10
15
20
0
20
40
60
80
100

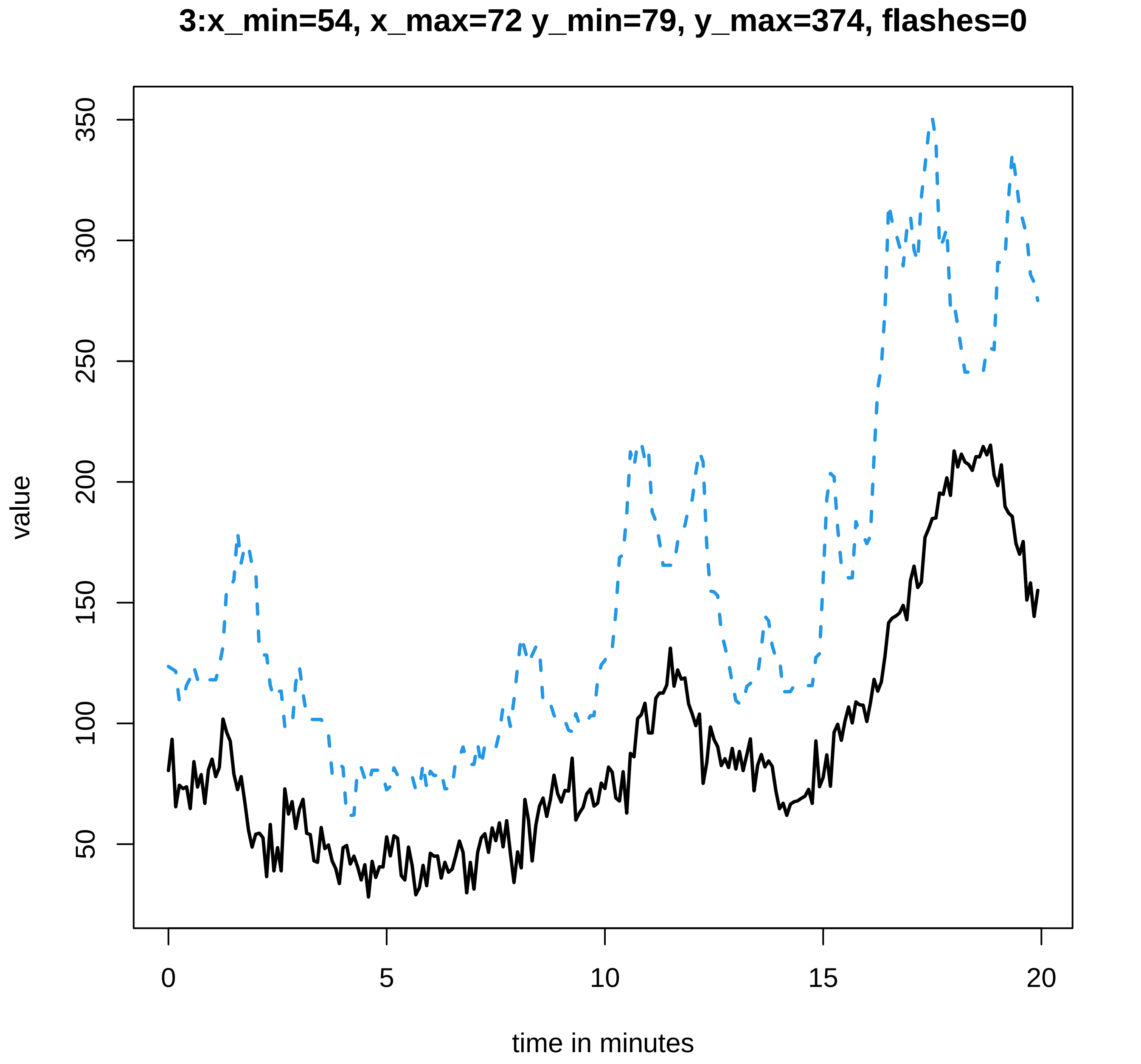
3:x_min=54, x_max=72 y_min=79, y_max=374, flashes=0
value
350
300
250
200
150
100
50
0
5
10
15
20
time in minutes

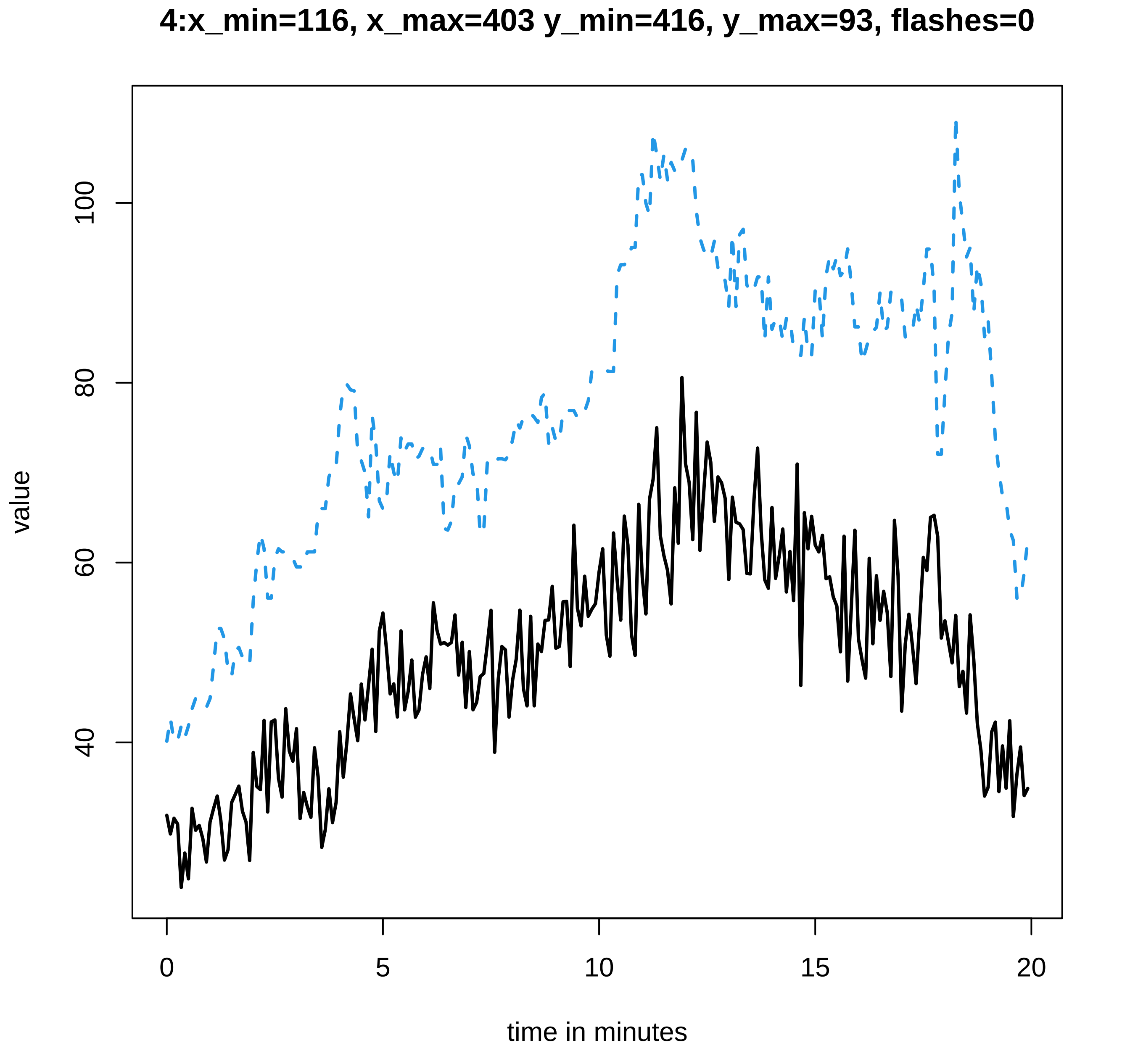
4:x_min=116, x_max=403 y_min=416, y_max=93, flashes=0
value
100
80
60
40
0
5
10
15
20
time in minutes

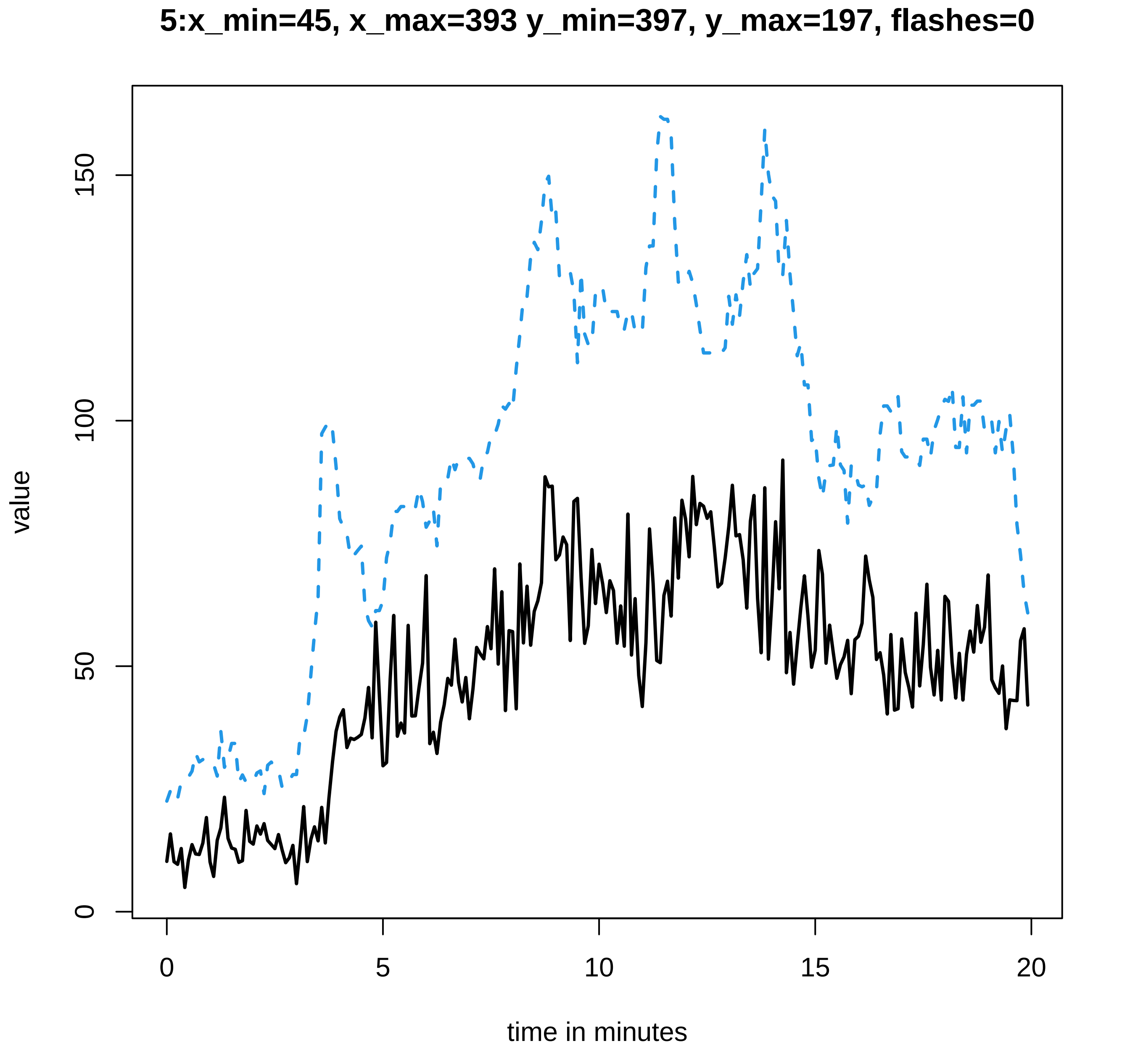
5:x_min=45, x_max=393 y_min=397, y_max=197, flashes=0
value
0
50
100
150
time in minutes
0
5
10
15
20

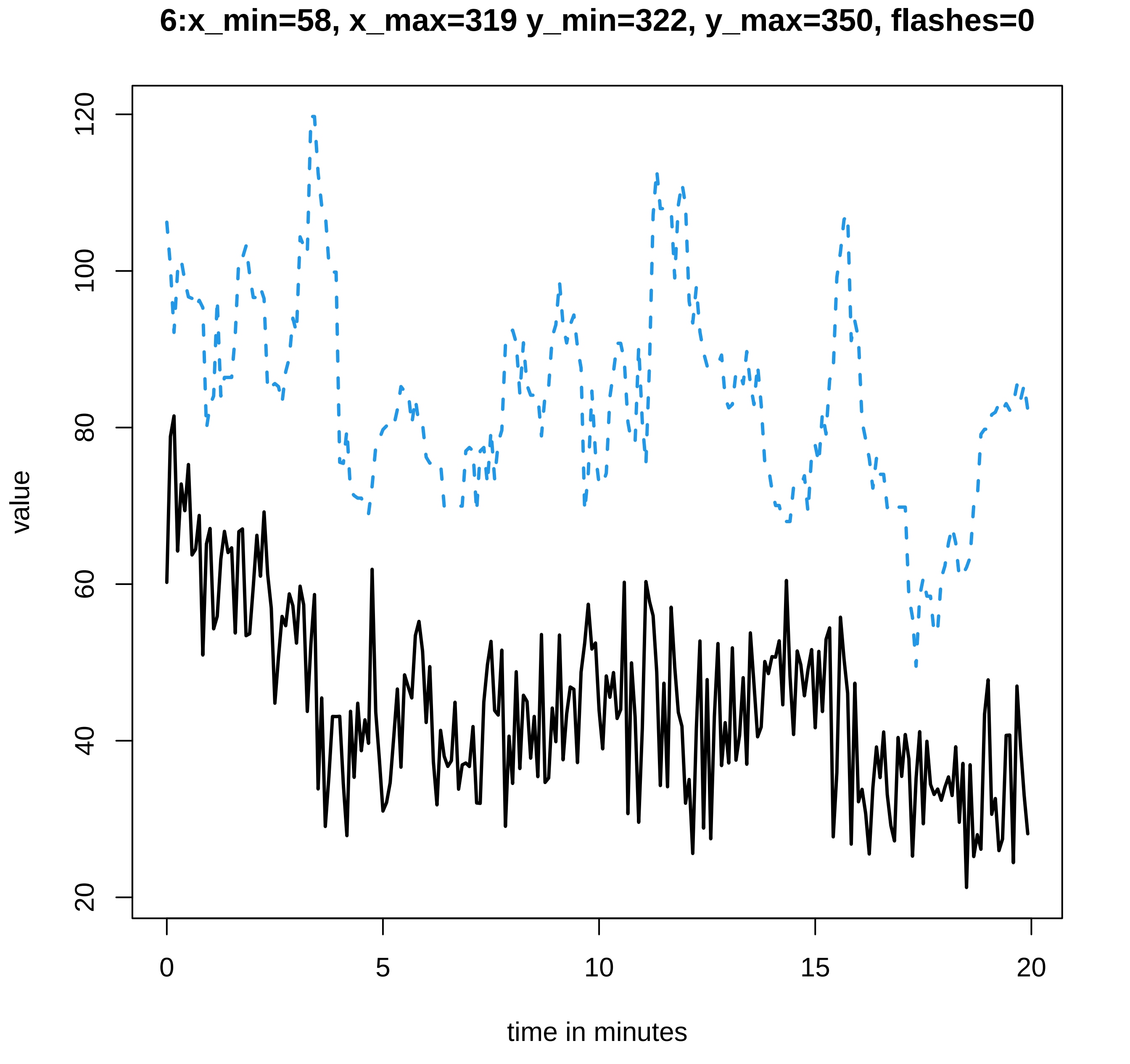
6:x_min=58, x_max=319 y_min=322, y_max=350, flashes=0
value
120
100
80
60
40
20
0
5
10
15
20
time in minutes

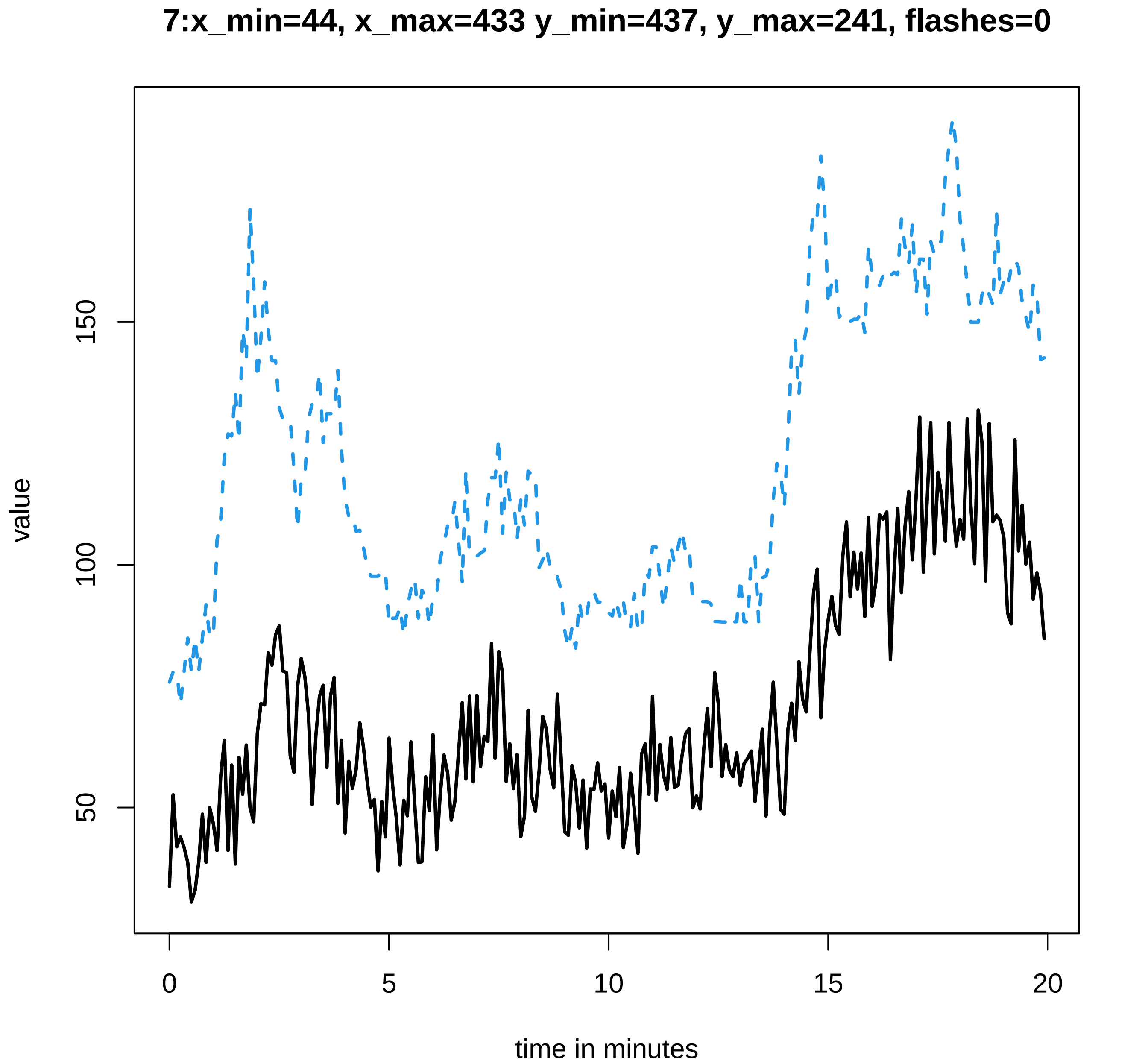

7:x_min=44, x_max=433 y_min=437, y_max=241, flashes=0
value
150
100
50
0
5
10
15
20
time in minutes

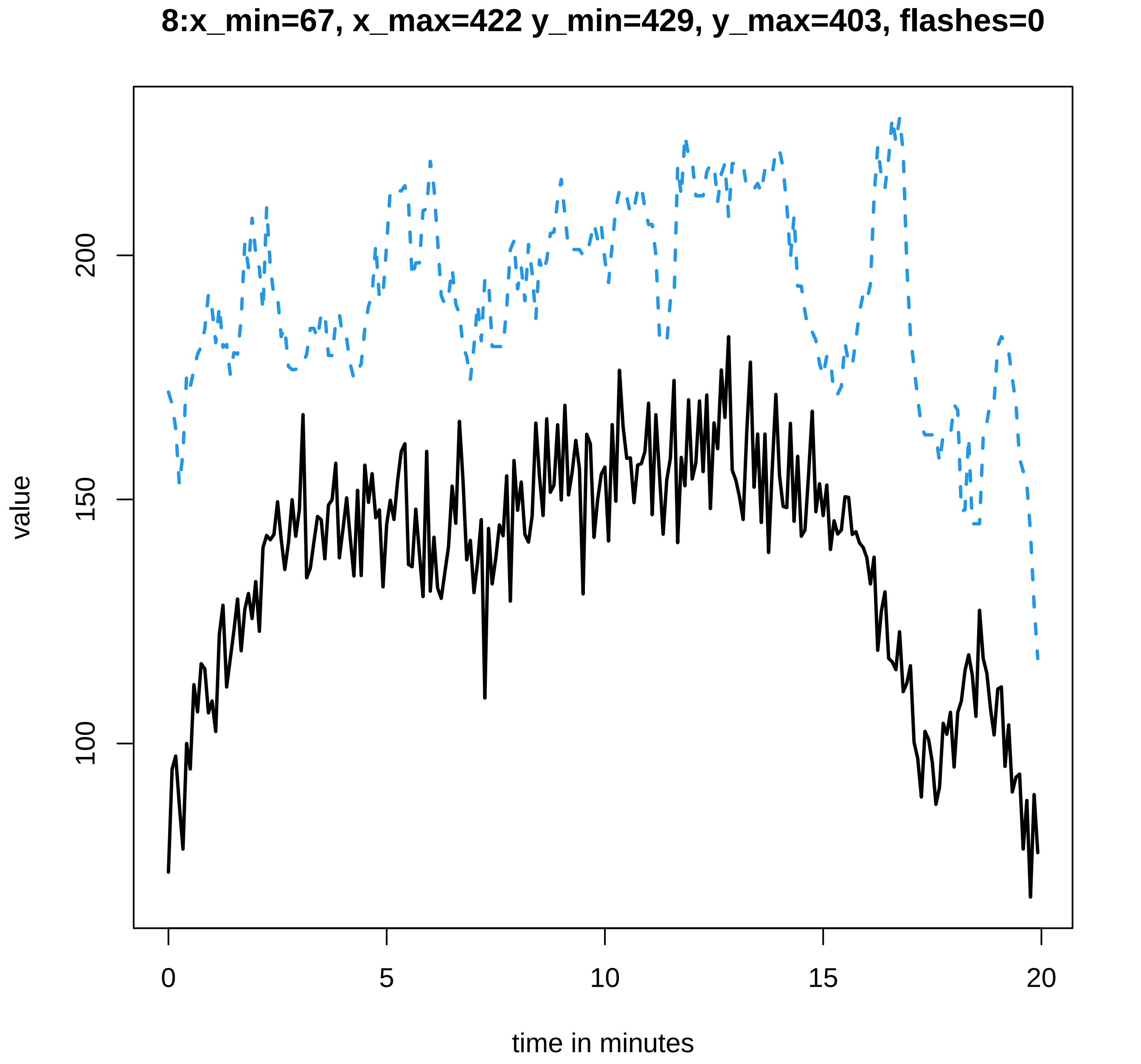

8:x_min=67, x_max=422 y_min=429, y_max=403, flashes=0
value
100
150
200
time in minutes
0
5
10
15
20

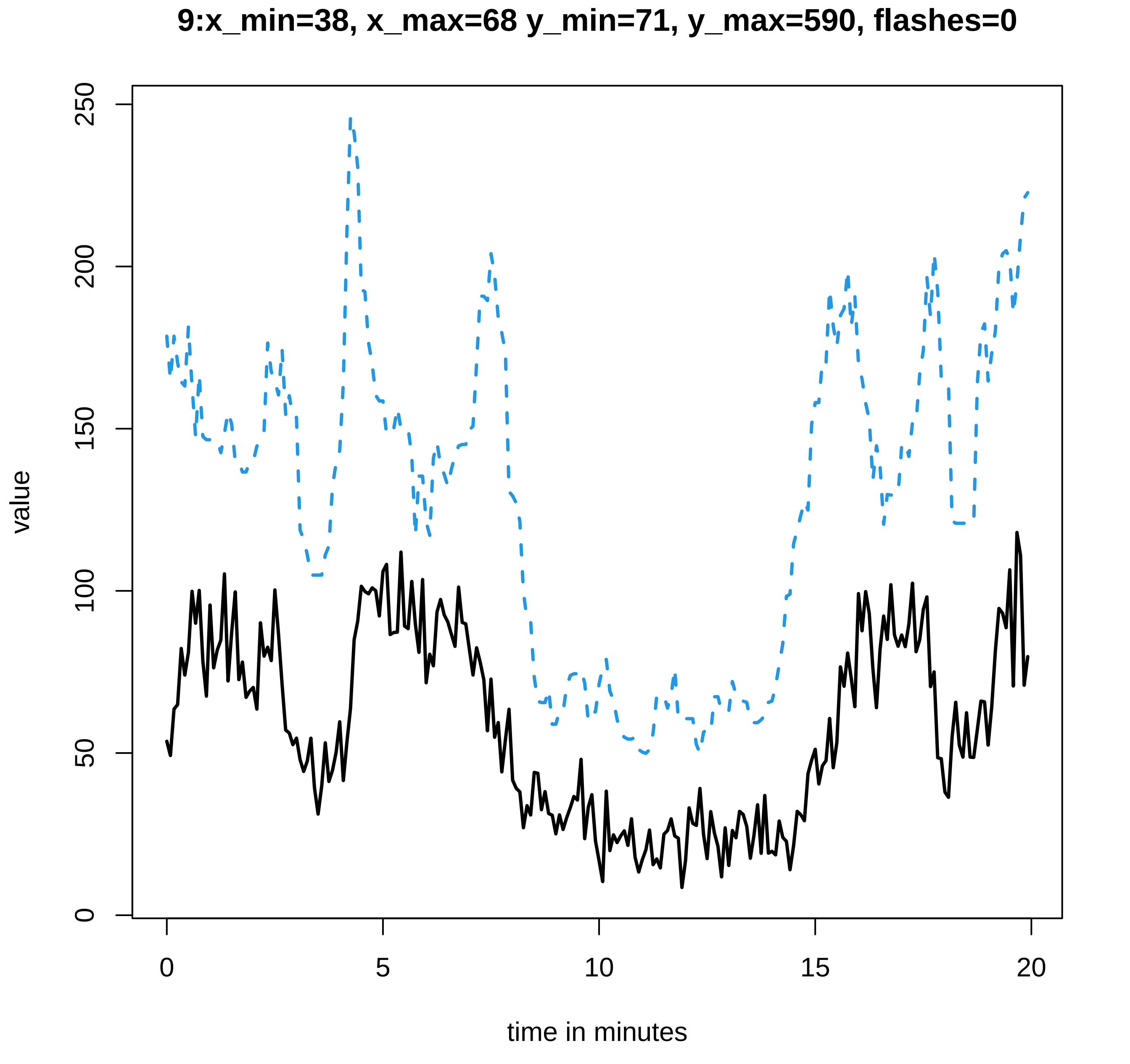

9:x_min=38, x_max=68 y_min=71, y_max=590, flashes=0
value
0
50
100
150
200
250
0
5
10
15
20
time in minutes

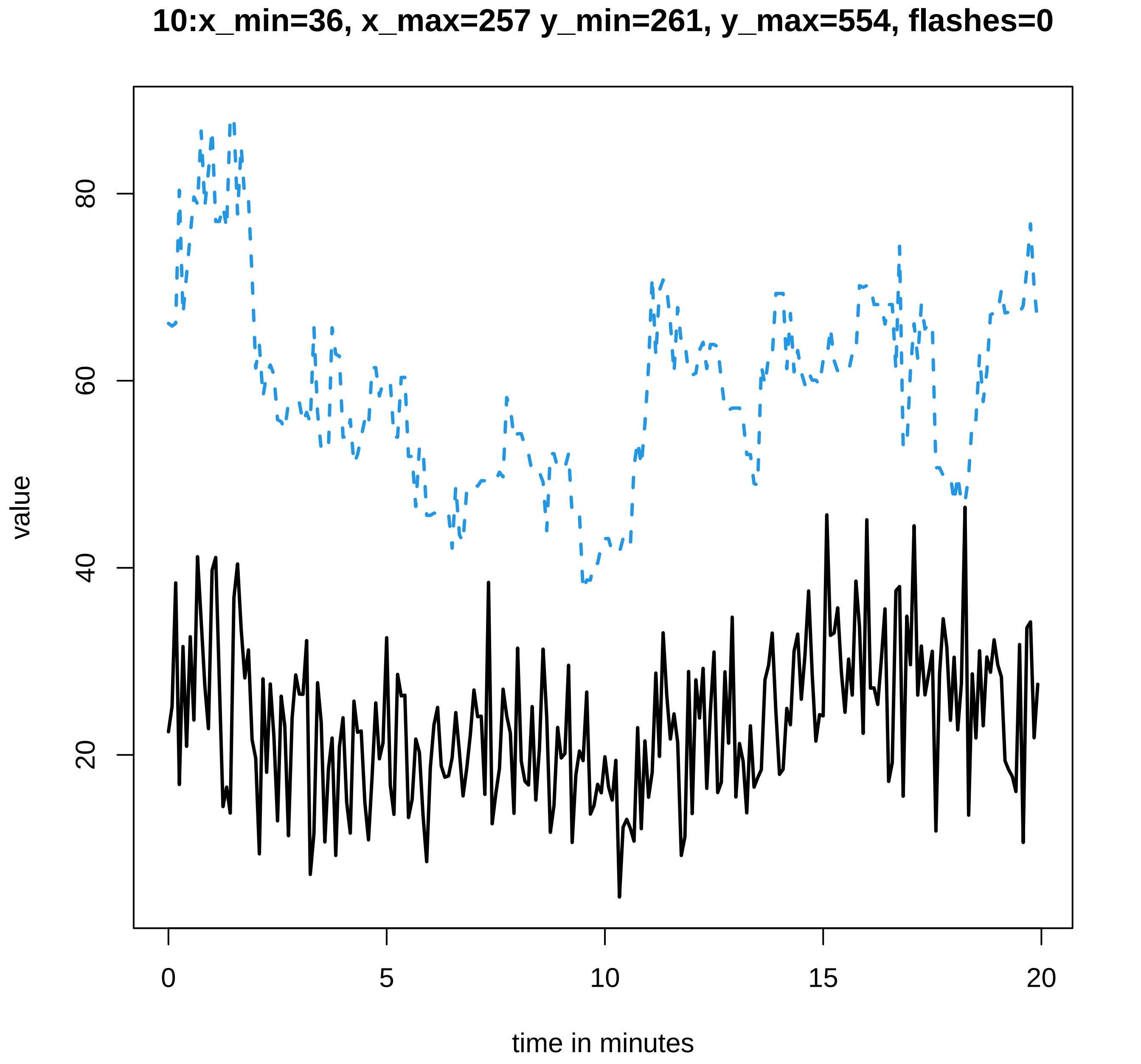
10:x_min=36, x_max=257 y_min=261, y_max=554, flashes=0
value
20
40
60
80
0
5
10
15
20
time in minutes

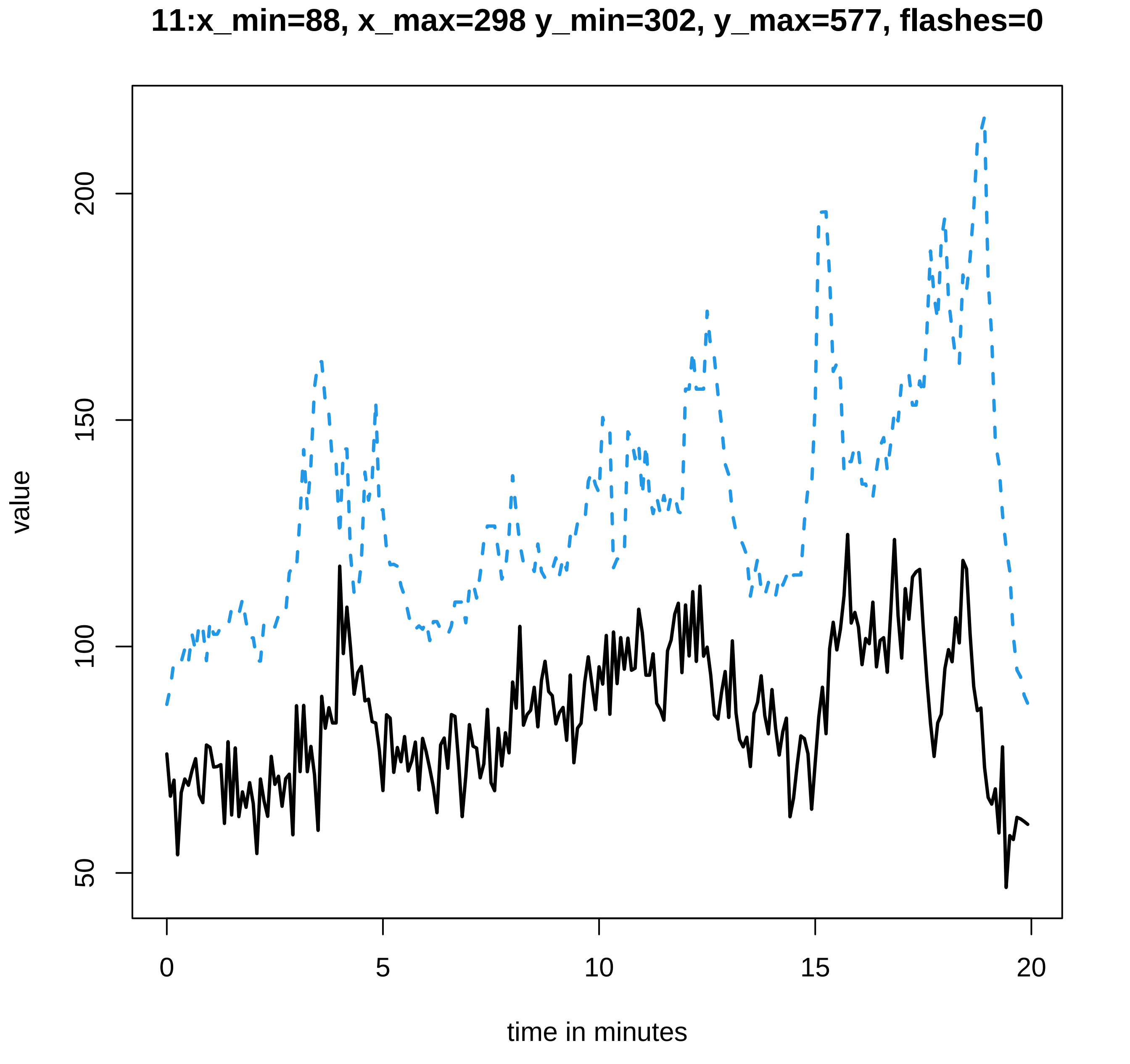

11:x_min=88, x_max=298 y_min=302, y_max=577, flashes=0
value
200
150
100
50
0
5
10
15
20
time in minutes

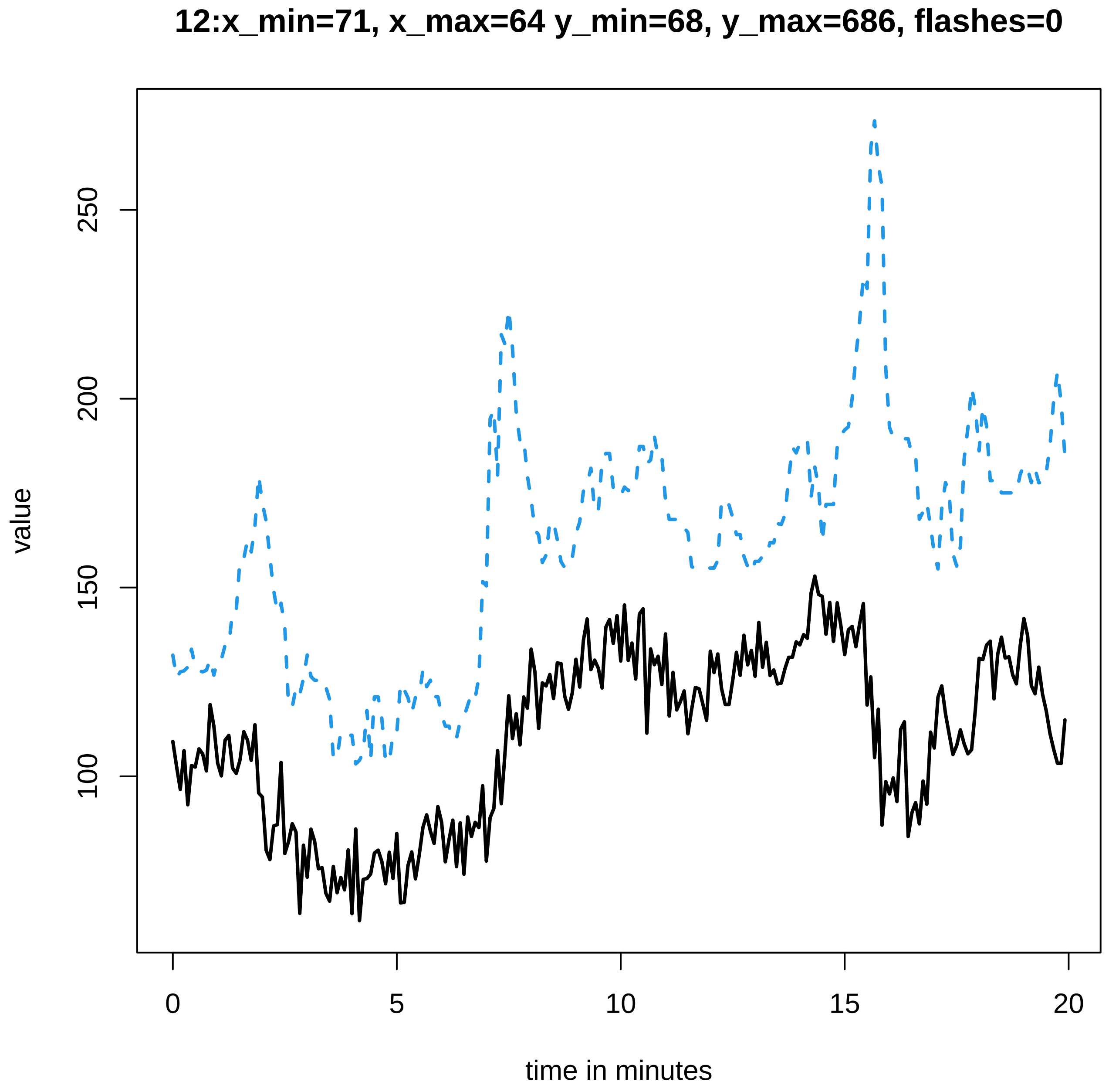
12:x_min=71, x_max=64 y_min=68, y_max=686, flashes=0
value
100
150
200
250
0
5
10
15
20
time in minutes

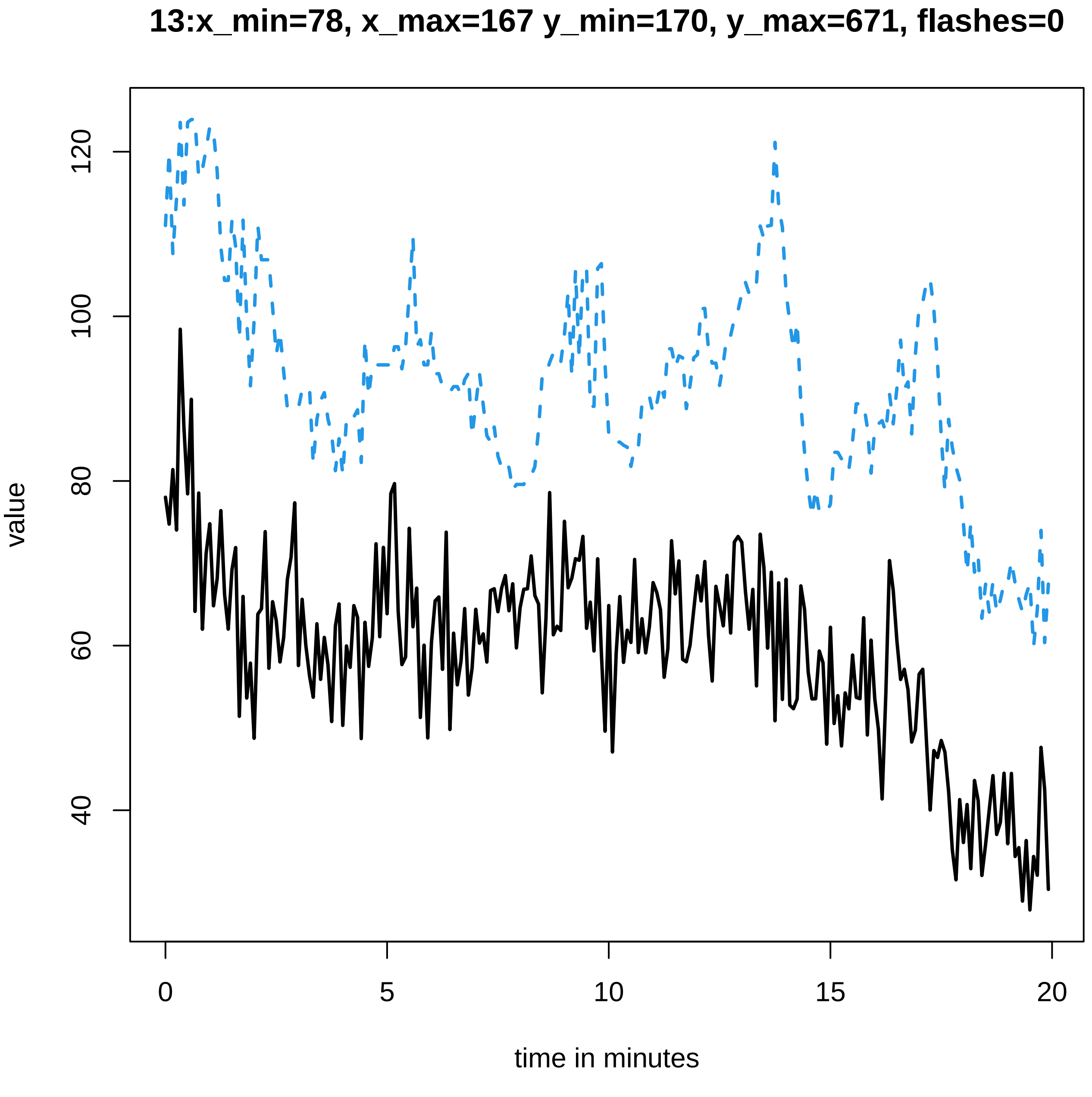
13:x_min=78, x_max=167 y_min=170, y_max=671, flashes=0
value
time in minutes
0
5
10
15
20
40
60
80
100
120

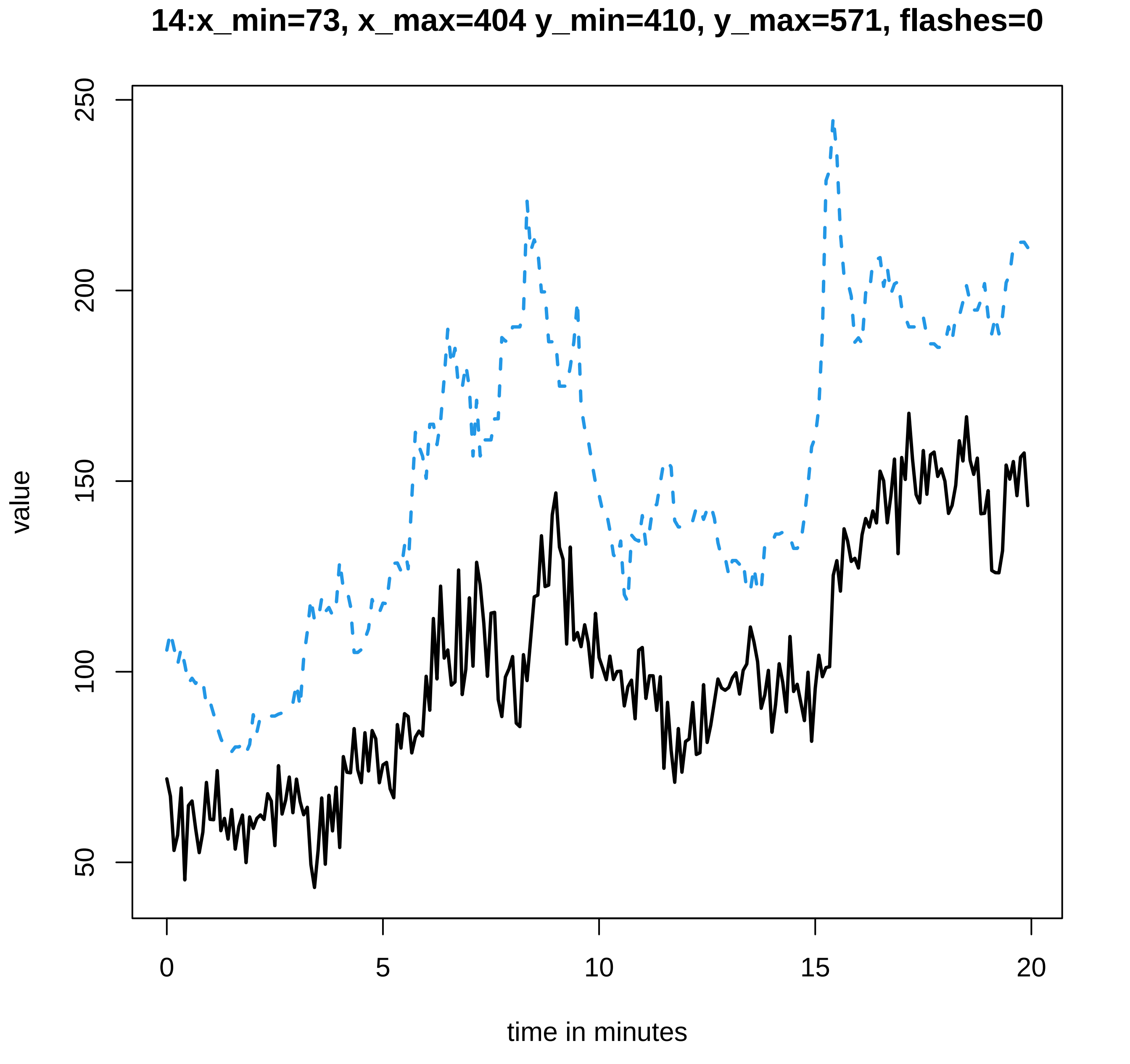
14:x_min=73, x_max=404 y_min=410, y_max=571, flashes=0
value
250
200
150
100
50
0
5
10
15
20
time in minutes

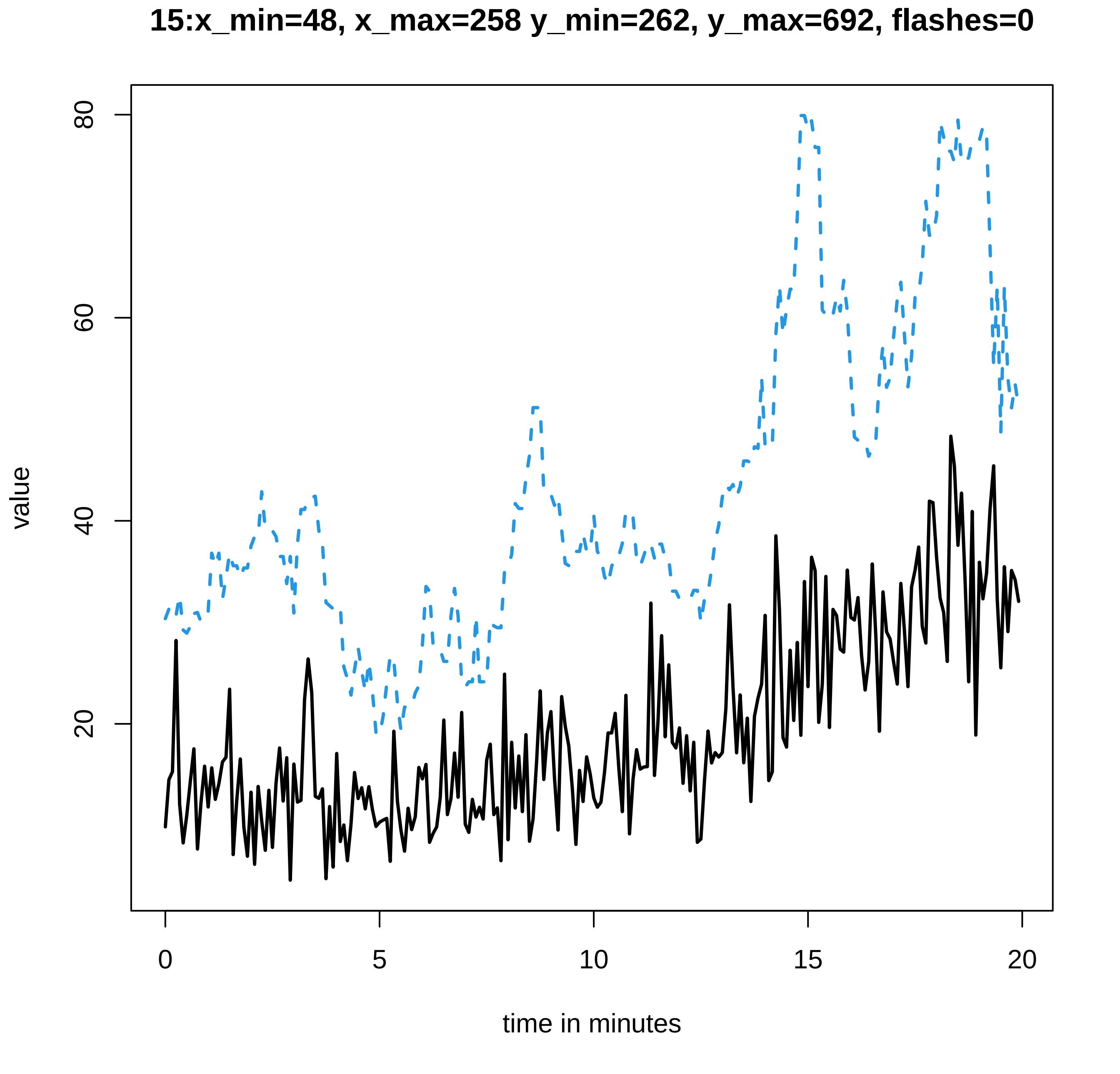
15:x_min=48, x_max=258 y_min=262, y_max=692, flashes=0
value
20
40
60
80
0
5
10
15
20
time in minutes

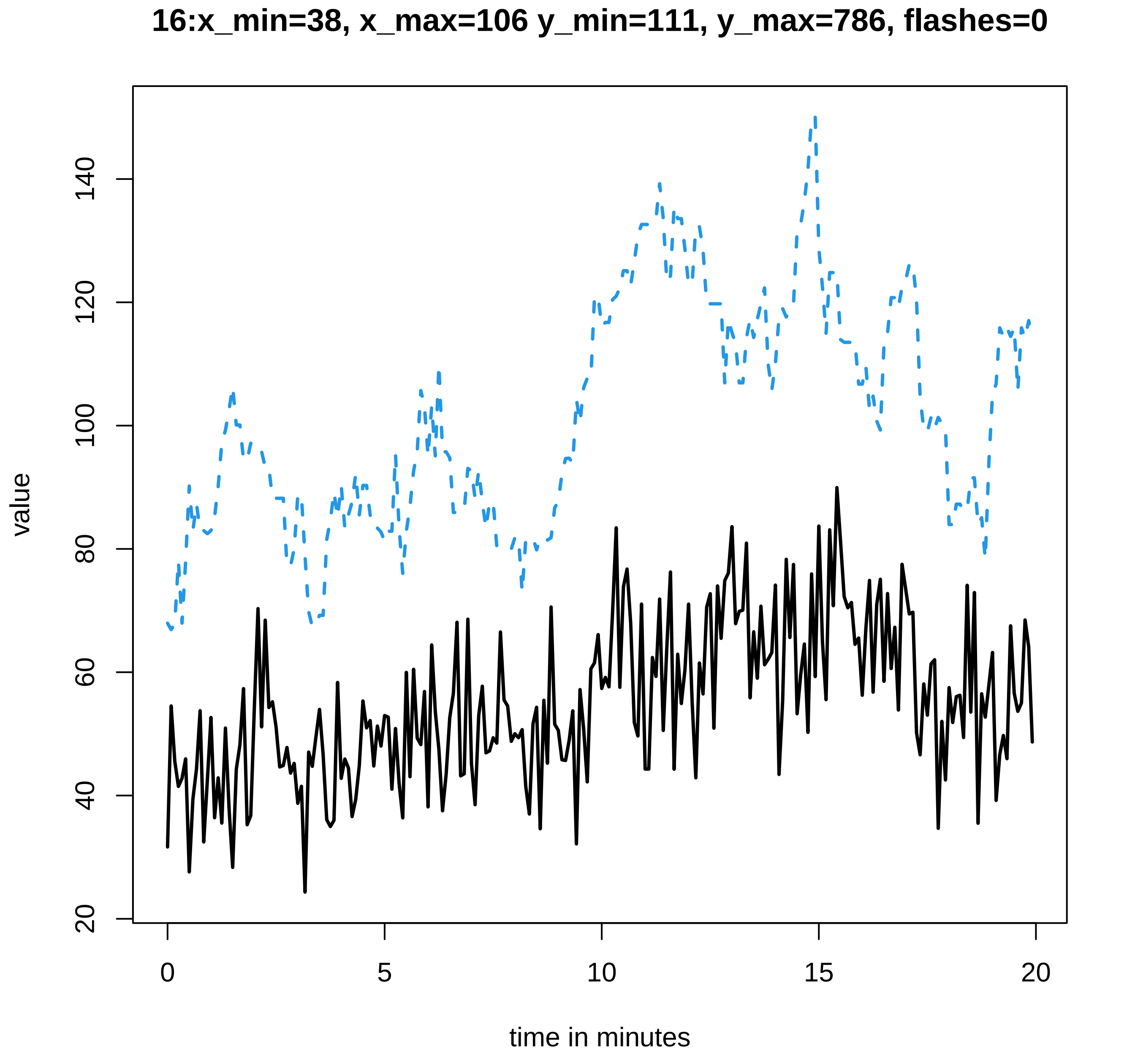

16:x_min=38, x_max=106 y_min=111, y_max=786, flashes=0
value
20
40
60
80
100
120
140
0
5
10
15
20
time in minutes

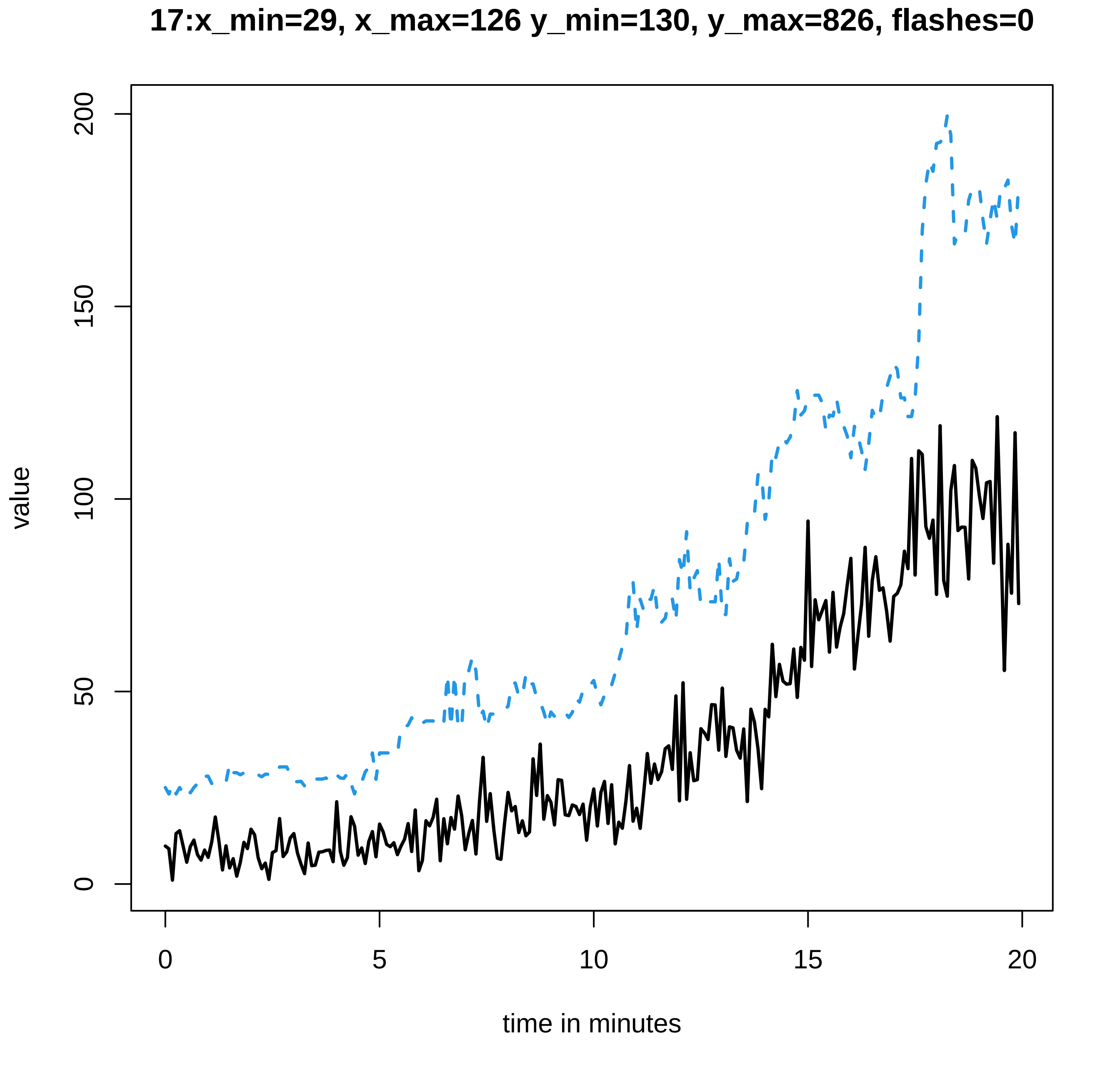
17:x_min=29, x_max=126 y_min=130, y_max=826, flashes=0
value
0
50
100
150
200
0
5
10
15
20
time in minutes

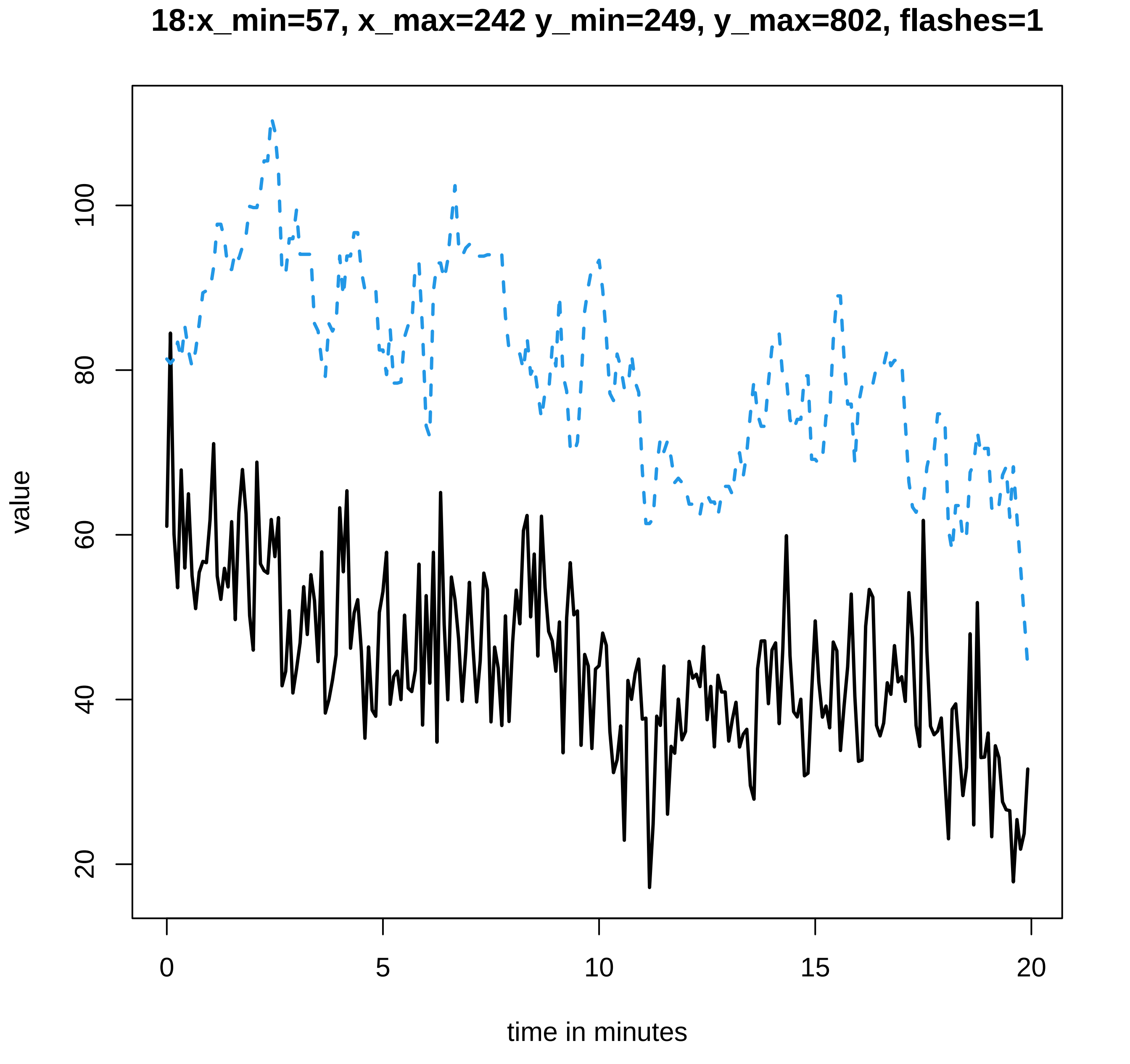
18:x_min=57, x_max=242 y_min=249, y_max=802, flashes=1
value
100
80
60
40
20
0
5
10
15
20
time in minutes

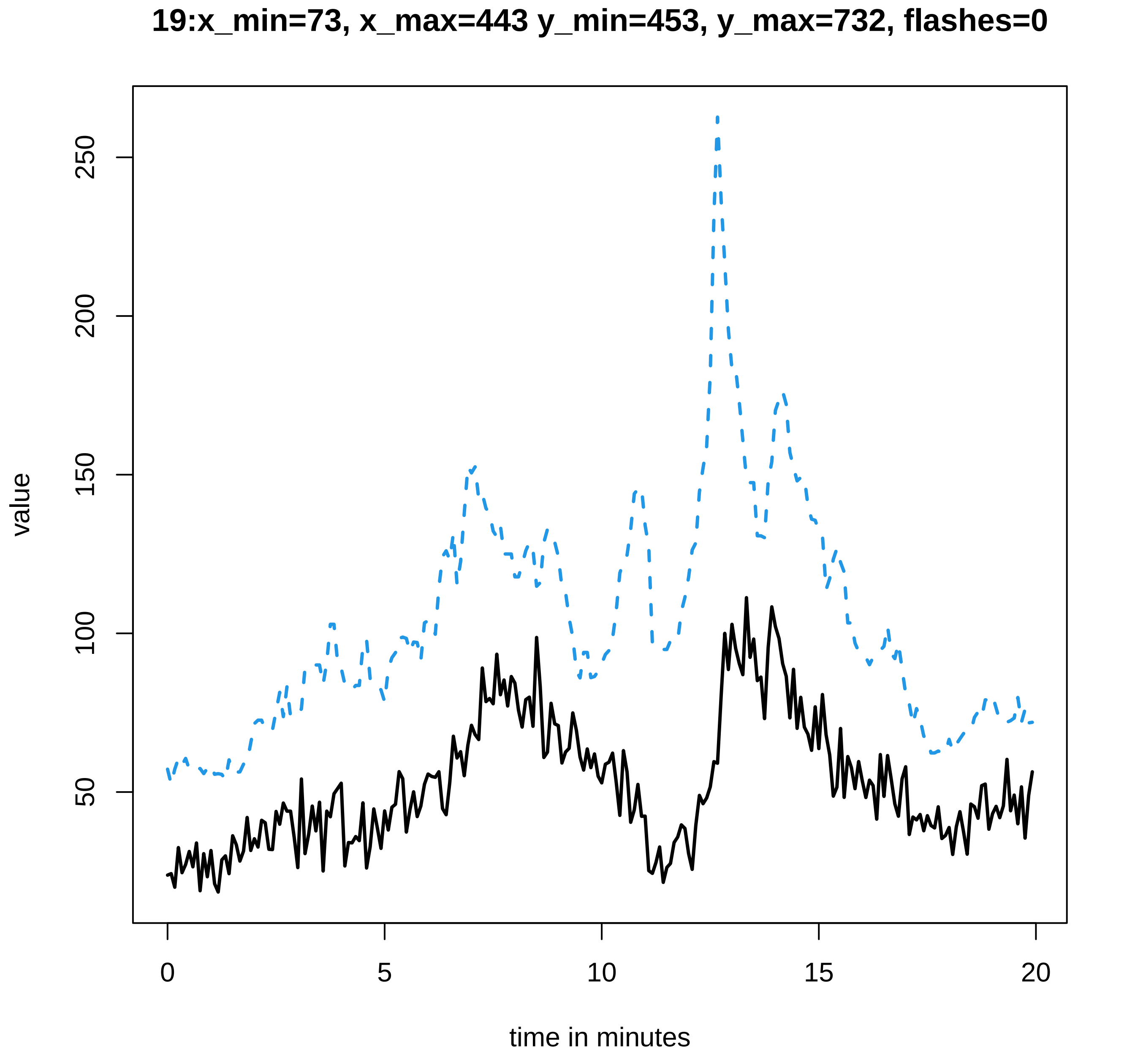
19:x_min=73, x_max=443 y_min=453, y_max=732, flashes=0
value
50
100
150
200
250
0
5
10
15
20
time in minutes

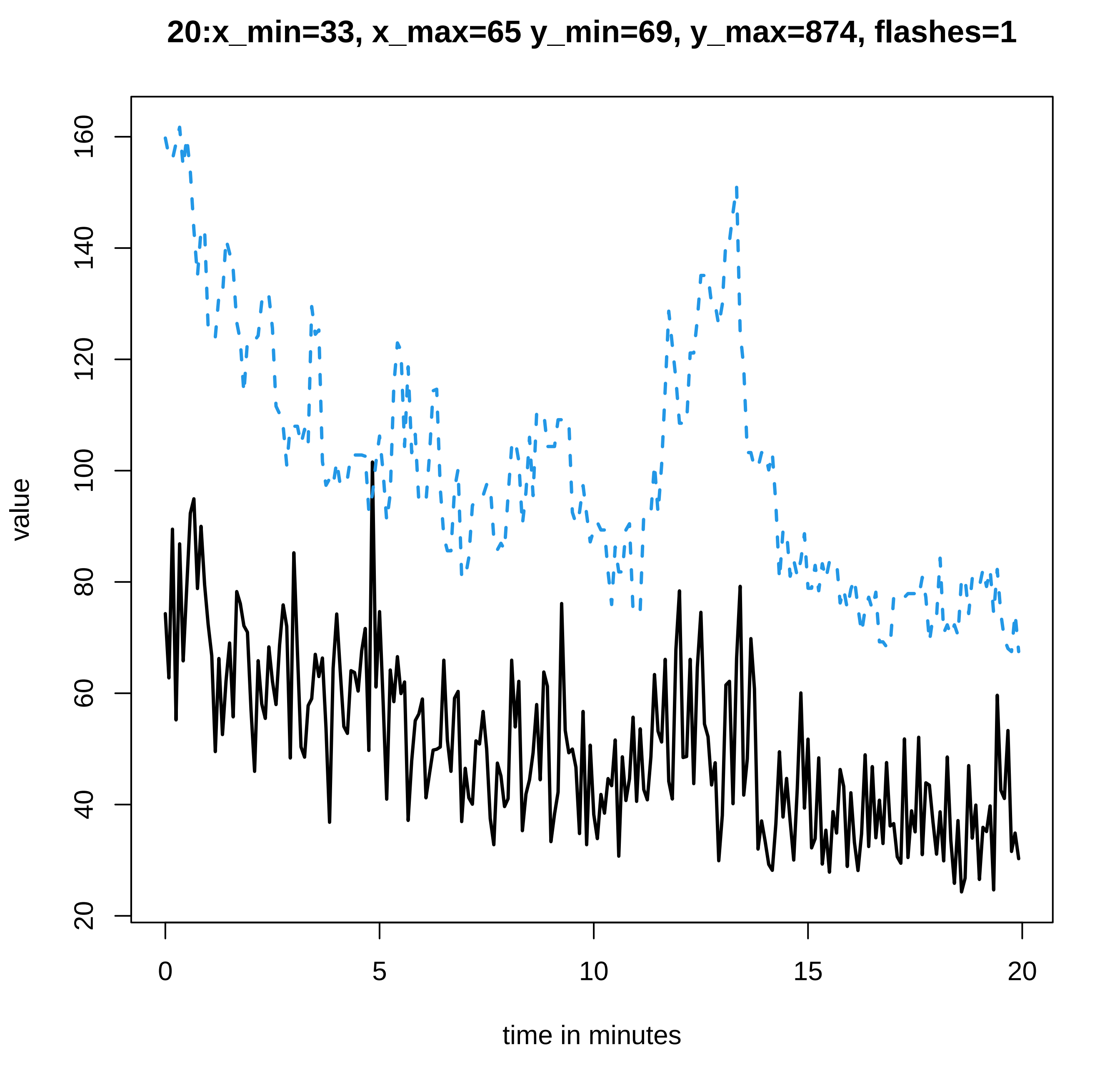
20:x_min=33, x_max=65 y_min=69, y_max=874, flashes=1
value
20
40
60
80
100
120
140
160
0
5
10
15
20
time in minutes

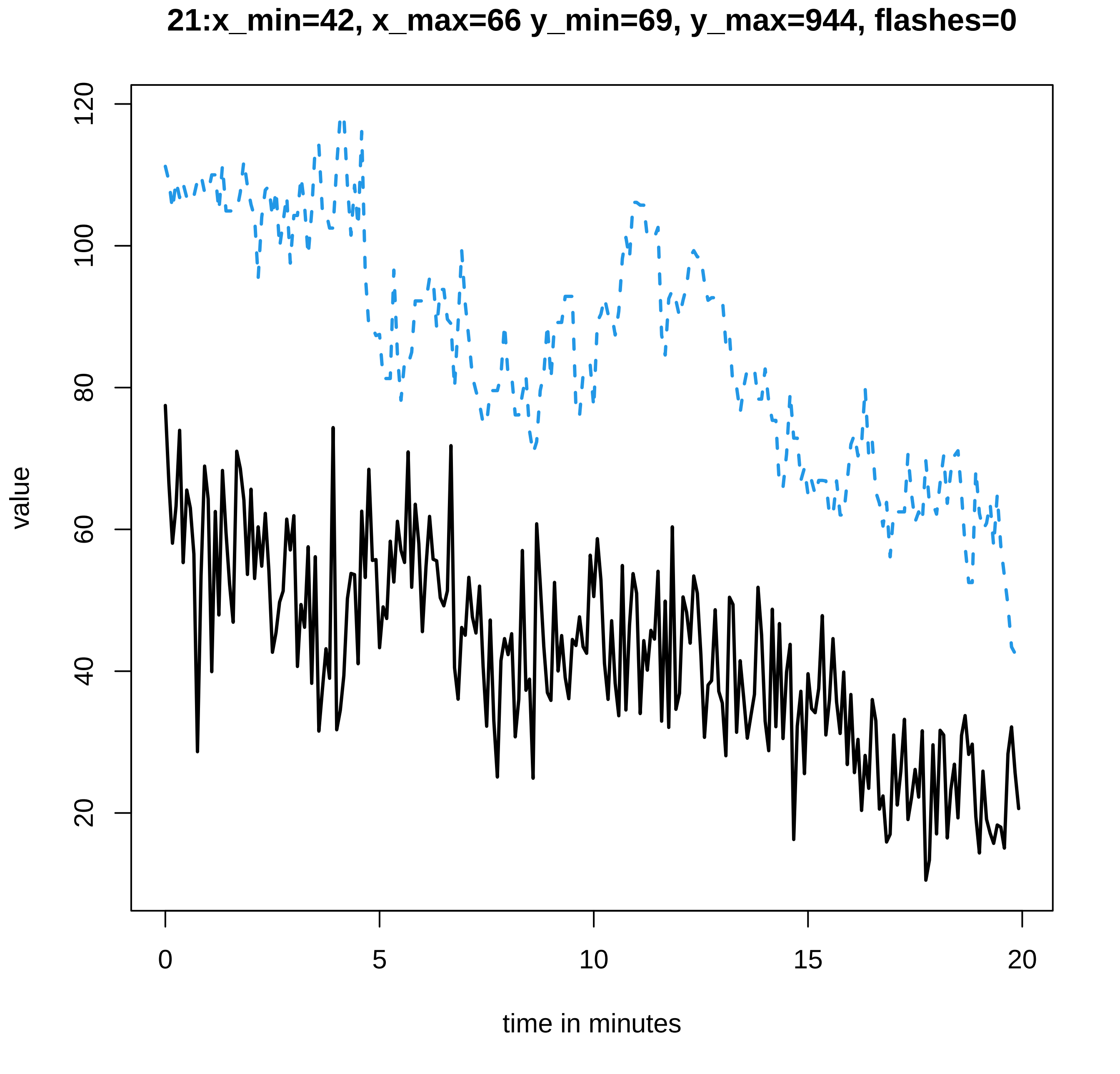
21:x_min=42, x_max=66 y_min=69, y_max=944, flashes=0
value
120
100
80
60
40
20
0
5
10
15
20
time in minutes

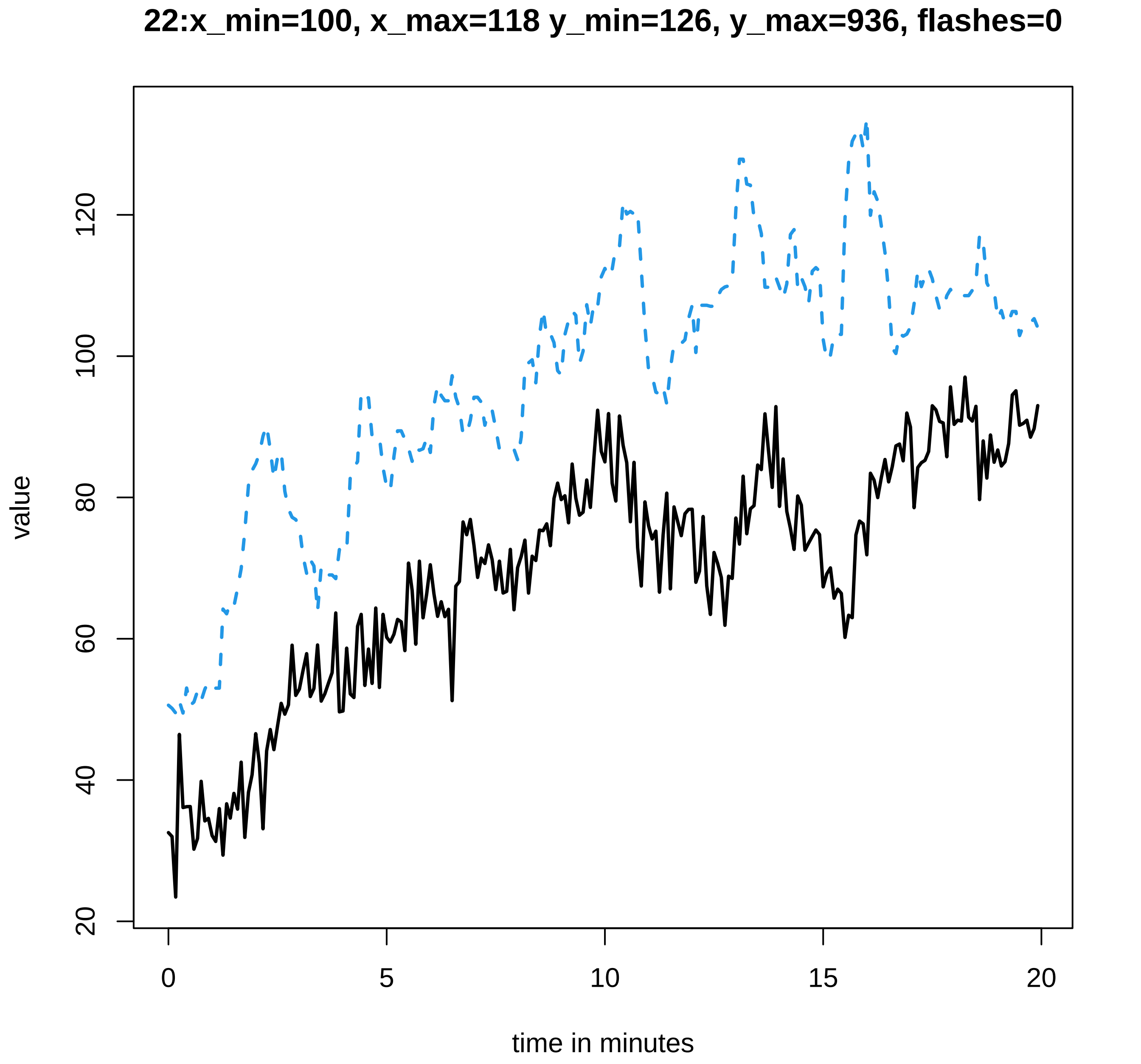
22:x_min=100, x_max=118 y_min=126, y_max=936, flashes=0
value
120
100
80
60
40
20
0
5
10
15
20
time in minutes

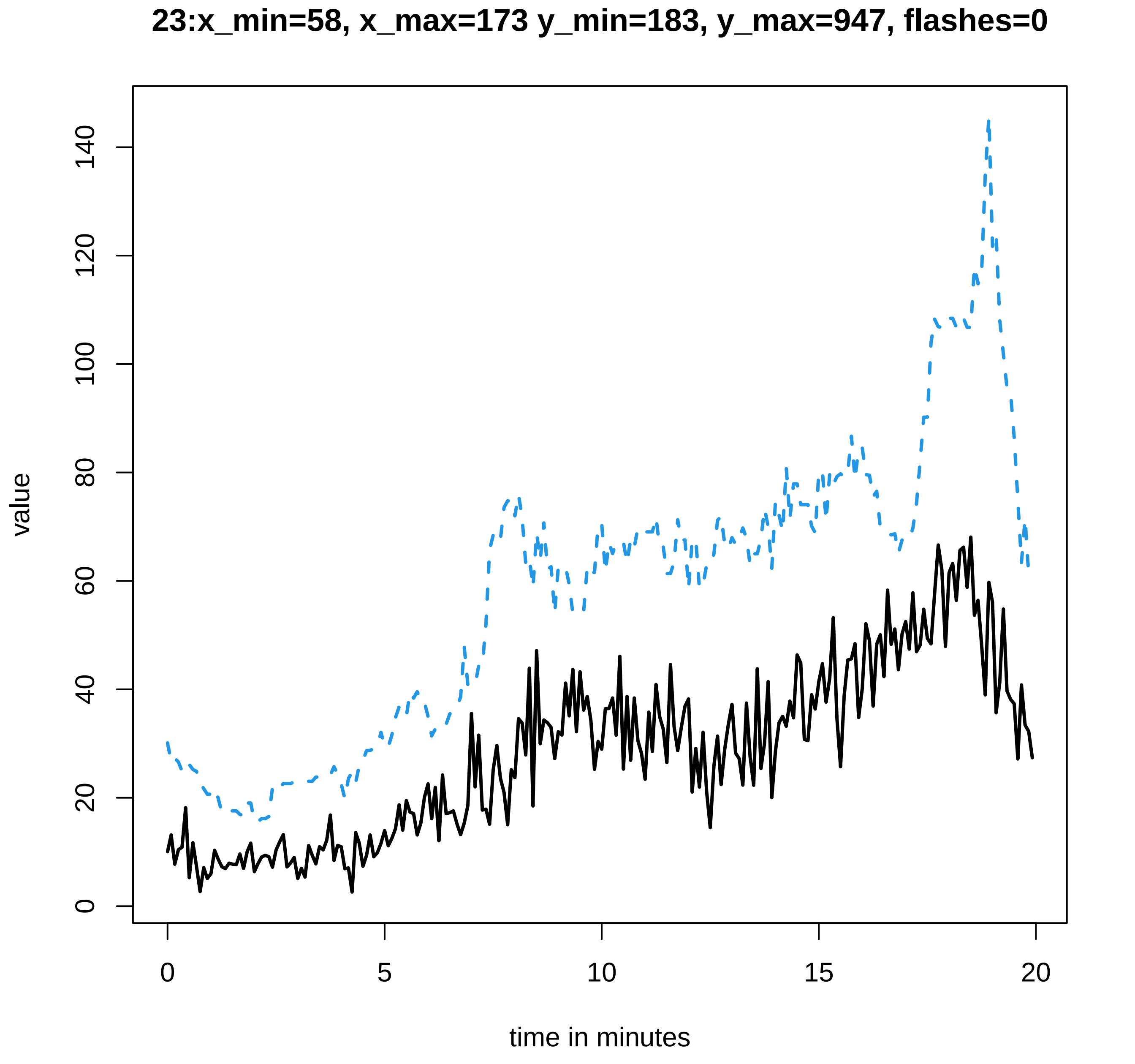
23:x_min=58, x_max=173 y_min=183, y_max=947, flashes=0
value
0
20
40
60
80
100
120
140
0
5
10
15
20
time in minutes

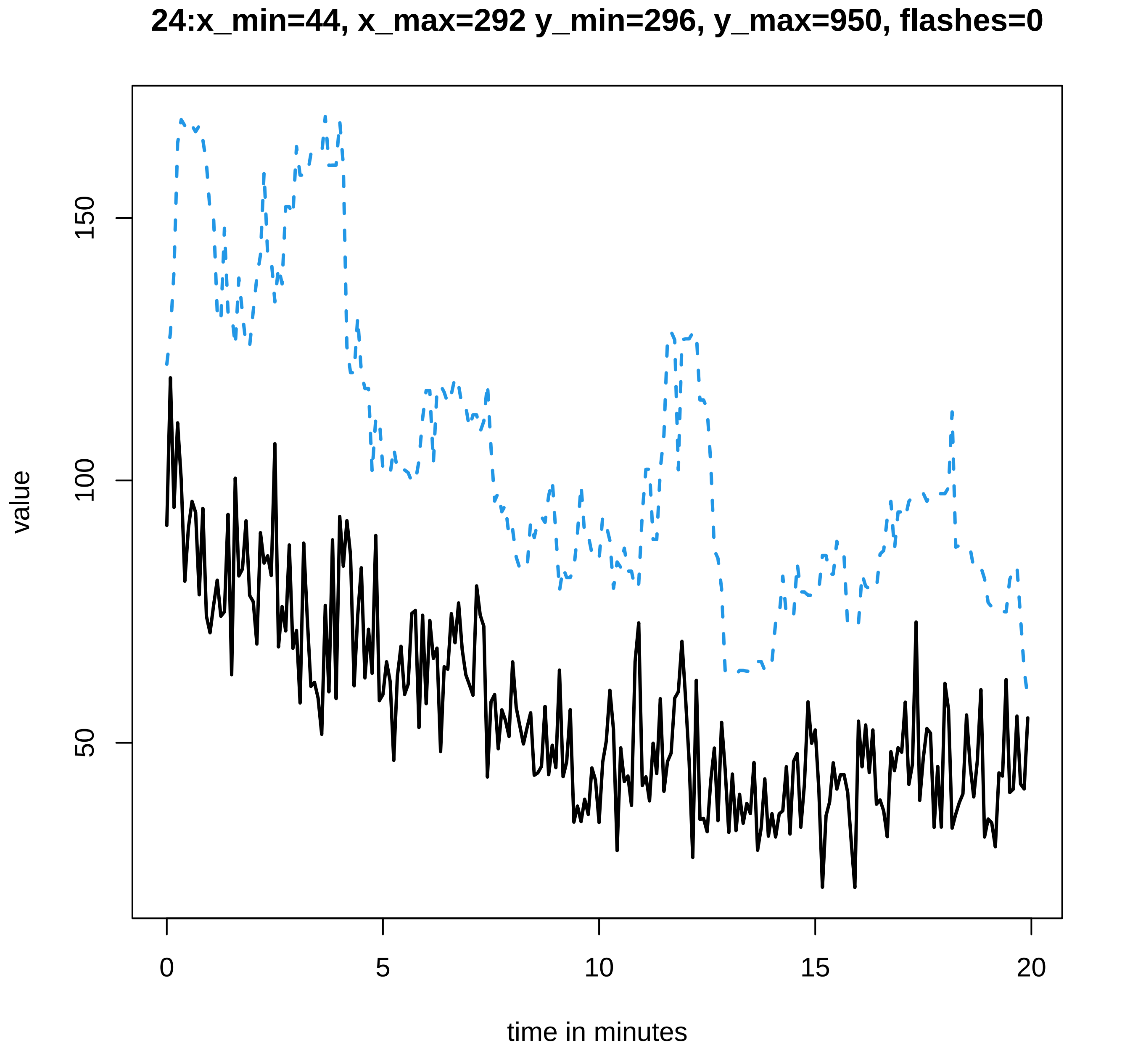
24:x_min=44, x_max=292 y_min=296, y_max=950, flashes=0
value
150
100
50
0
5
10
15
20
time in minutes

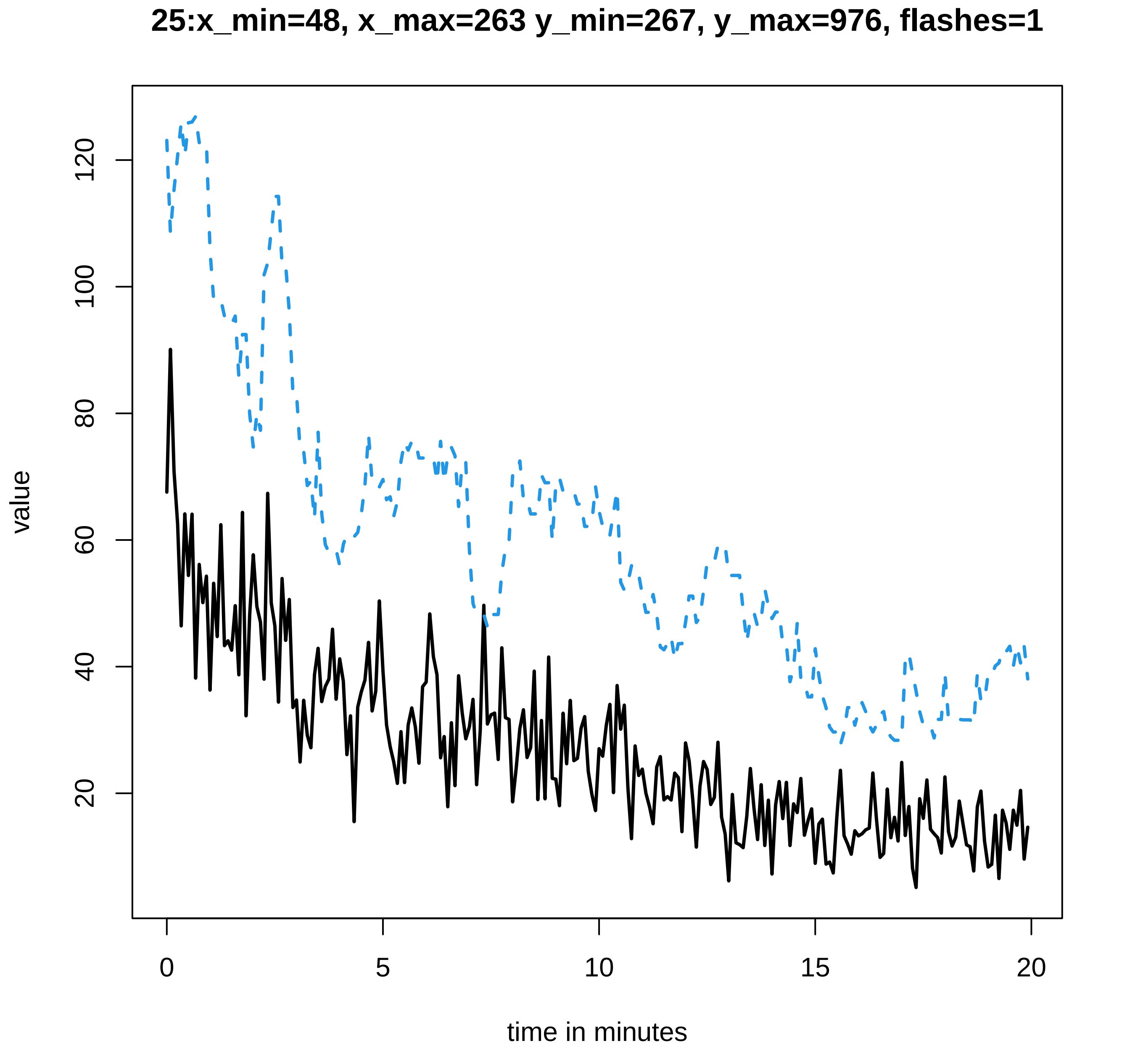
25:x_min=48, x_max=263 y_min=267, y_max=976, flashes=1
value
120
100
80
60
40
20
0
5
10
15
20
time in minutes
