## Supplementary data Calcium spiking for "LysM Receptor Proteins are Required for Ectomycorrhizal Symbiosis in Poplar": profiles-spiking-19.pdf

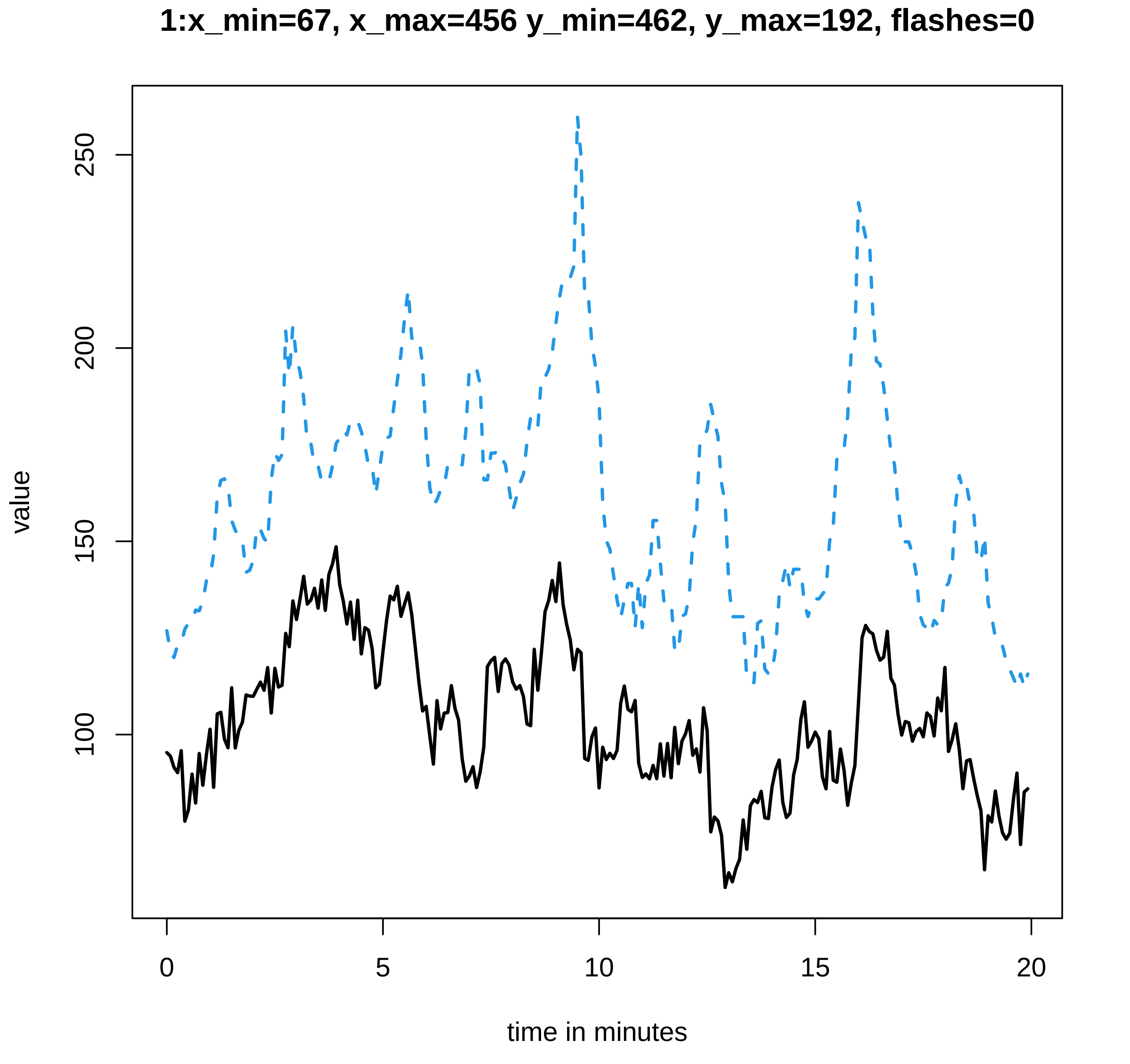
1:x_min=67, x_max=456 y_min=462, y_max=192, flashes=0
value
100
150
200
250
0
5
10
15
20
time in minutes

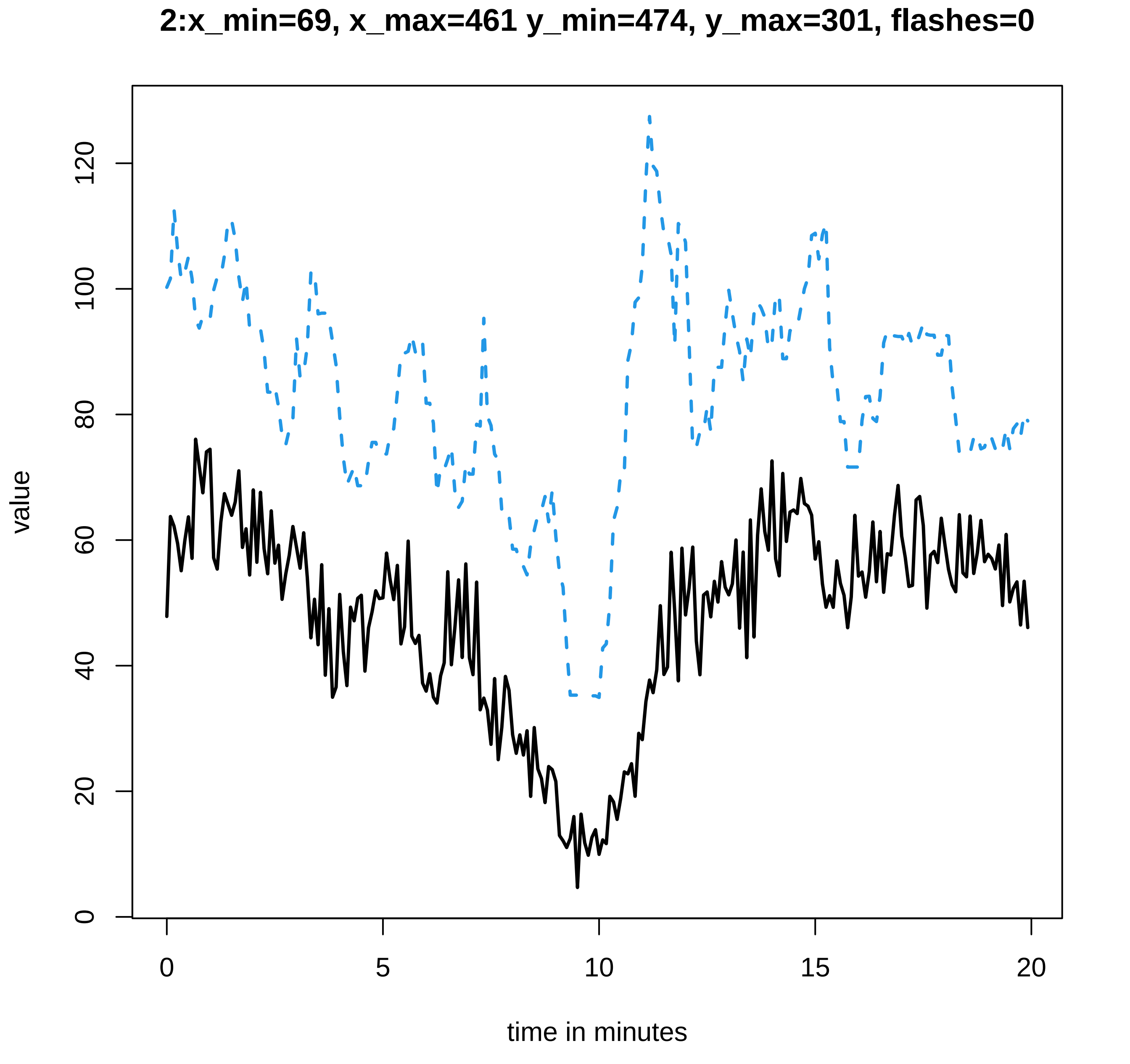

2:x_min=69, x_max=461 y_min=474, y_max=301, flashes=0
value
time in minutes
0
20
40
60
80
100
120
0
5
10
15
20

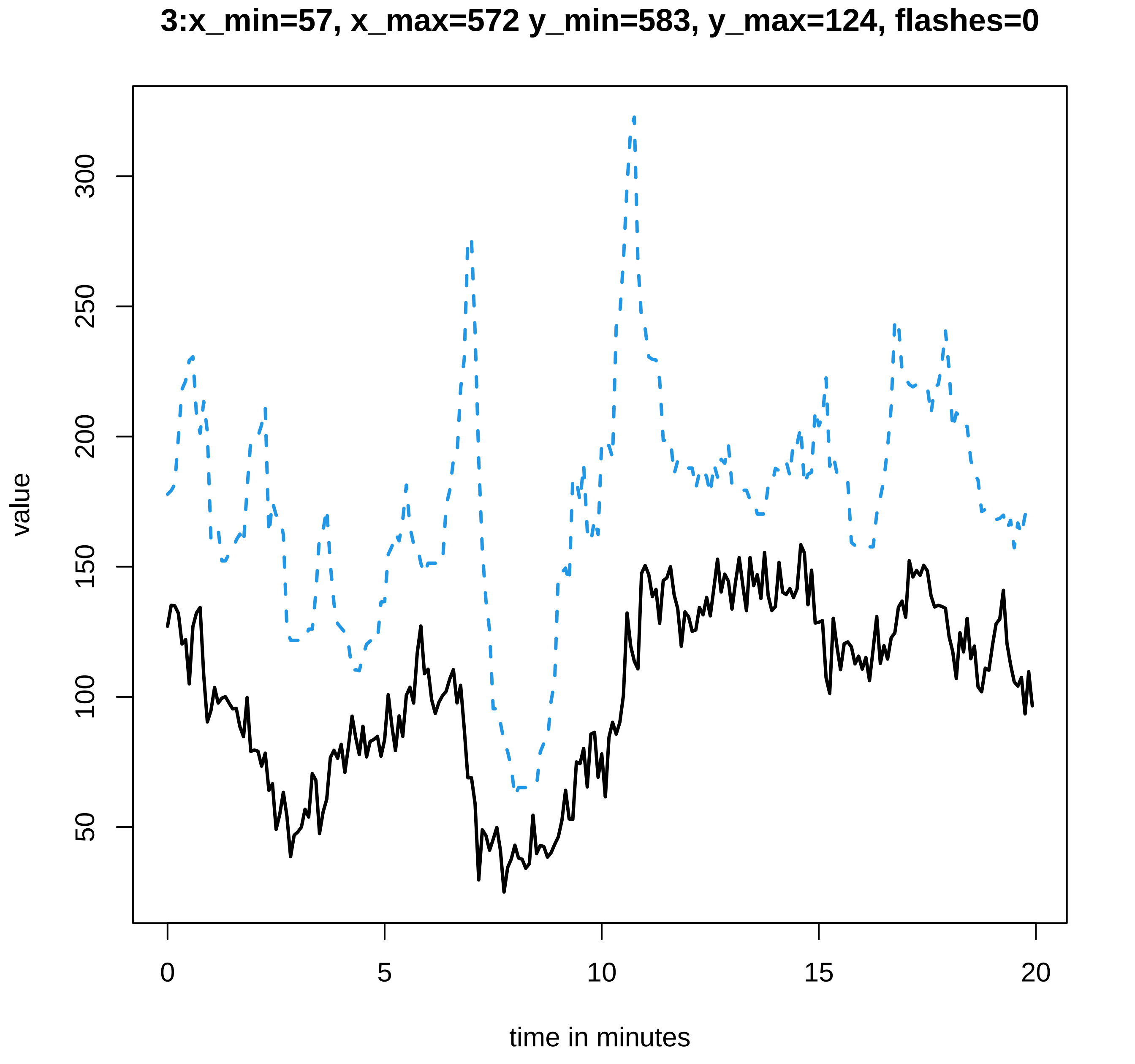
3:x_min=57, x_max=572 y_min=583, y_max=124, flashes=0
value
300
250
200
150
100
50
0
5
10
15
20
time in minutes

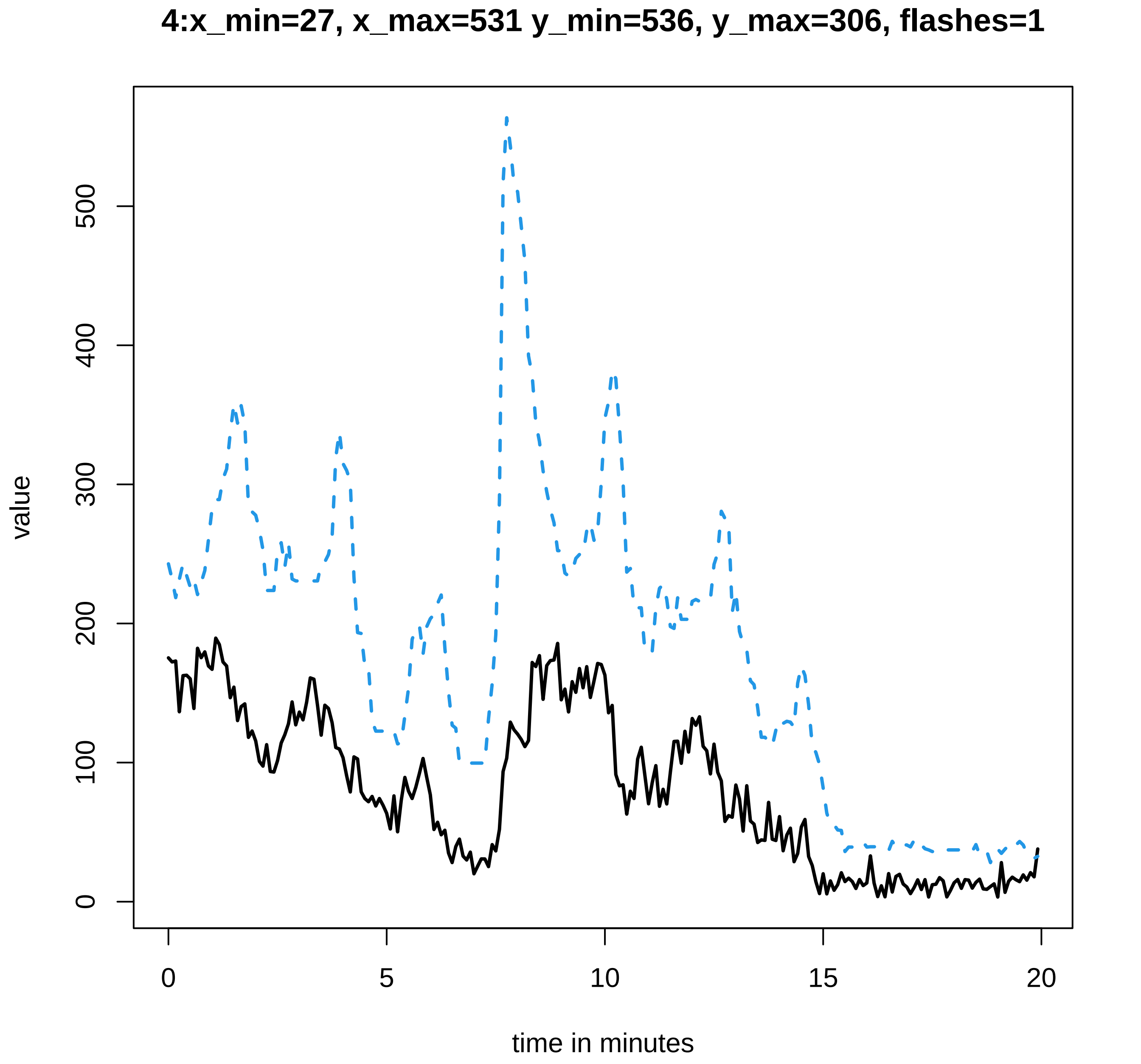

4:x_min=27, x_max=531 y_min=536, y_max=306, flashes=1
value
0
100
200
300
400
500
0
5
10
15
20
time in minutes

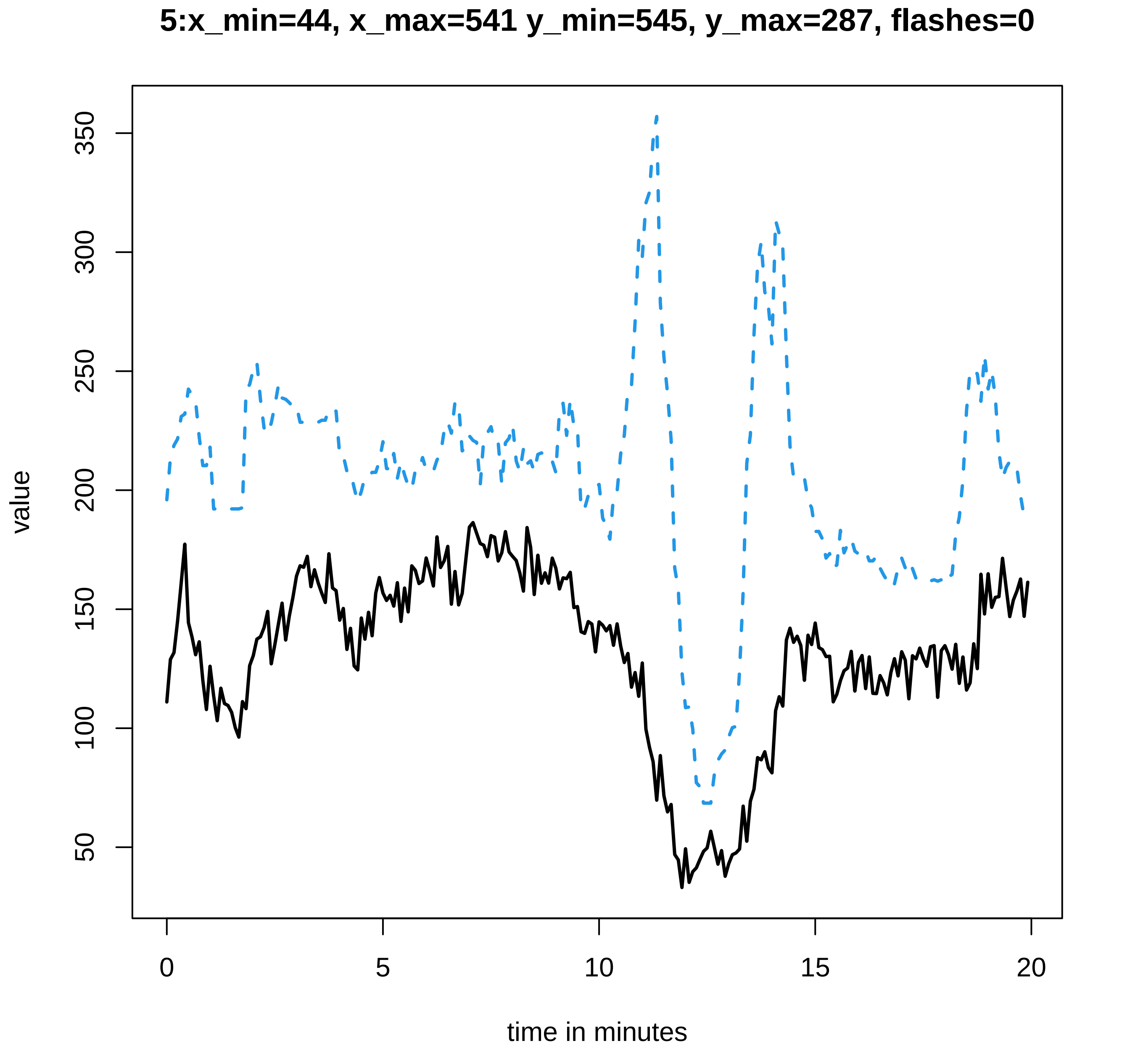
5:x_min=44, x_max=541 y_min=545, y_max=287, flashes=0
value
350
300
250
200
150
100
50
0
5
10
15
20
time in minutes

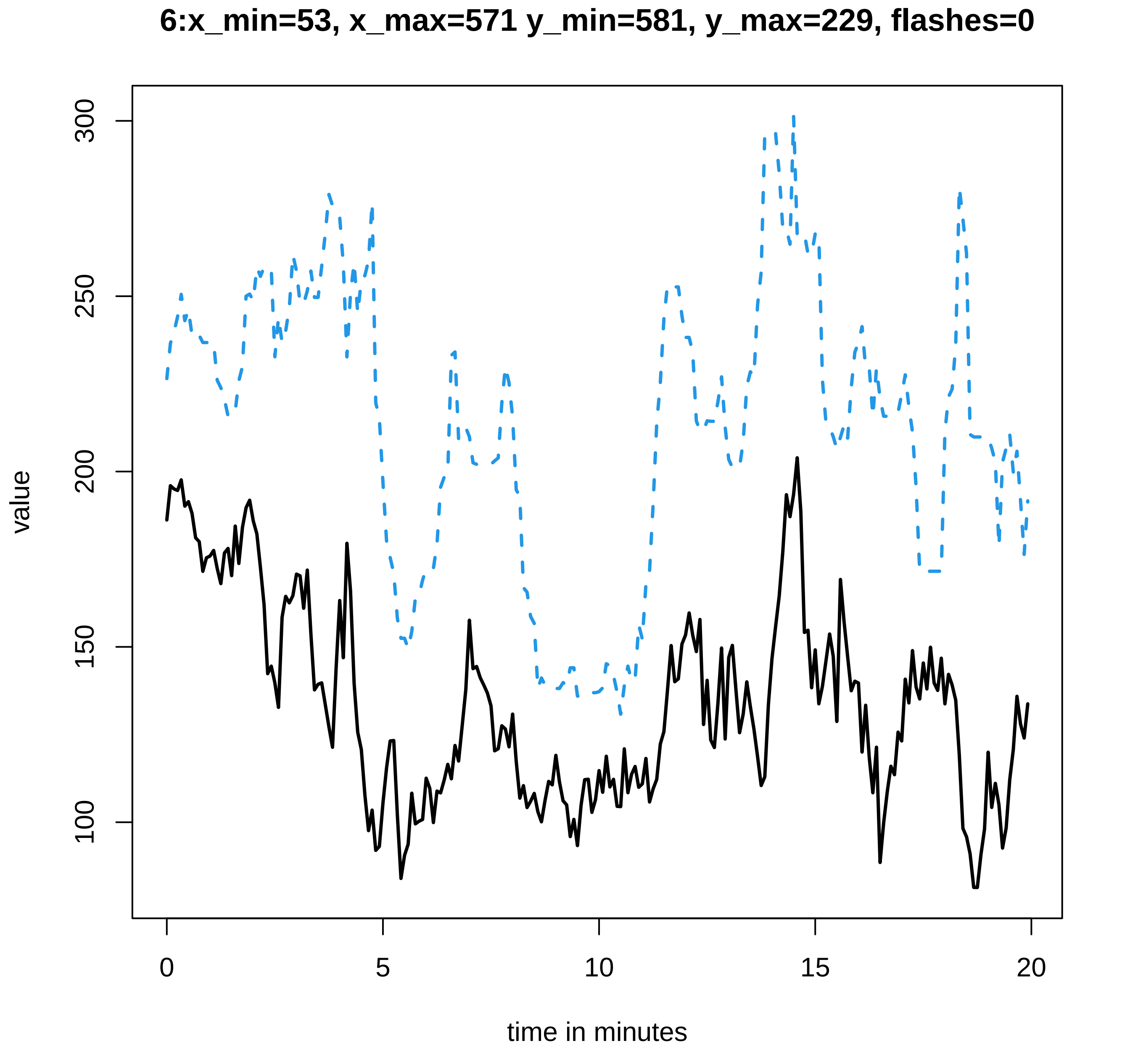
6:x_min=53, x_max=571 y_min=581, y_max=229, flashes=0
value
100
150
200
250
300
0
5
10
15
20
time in minutes

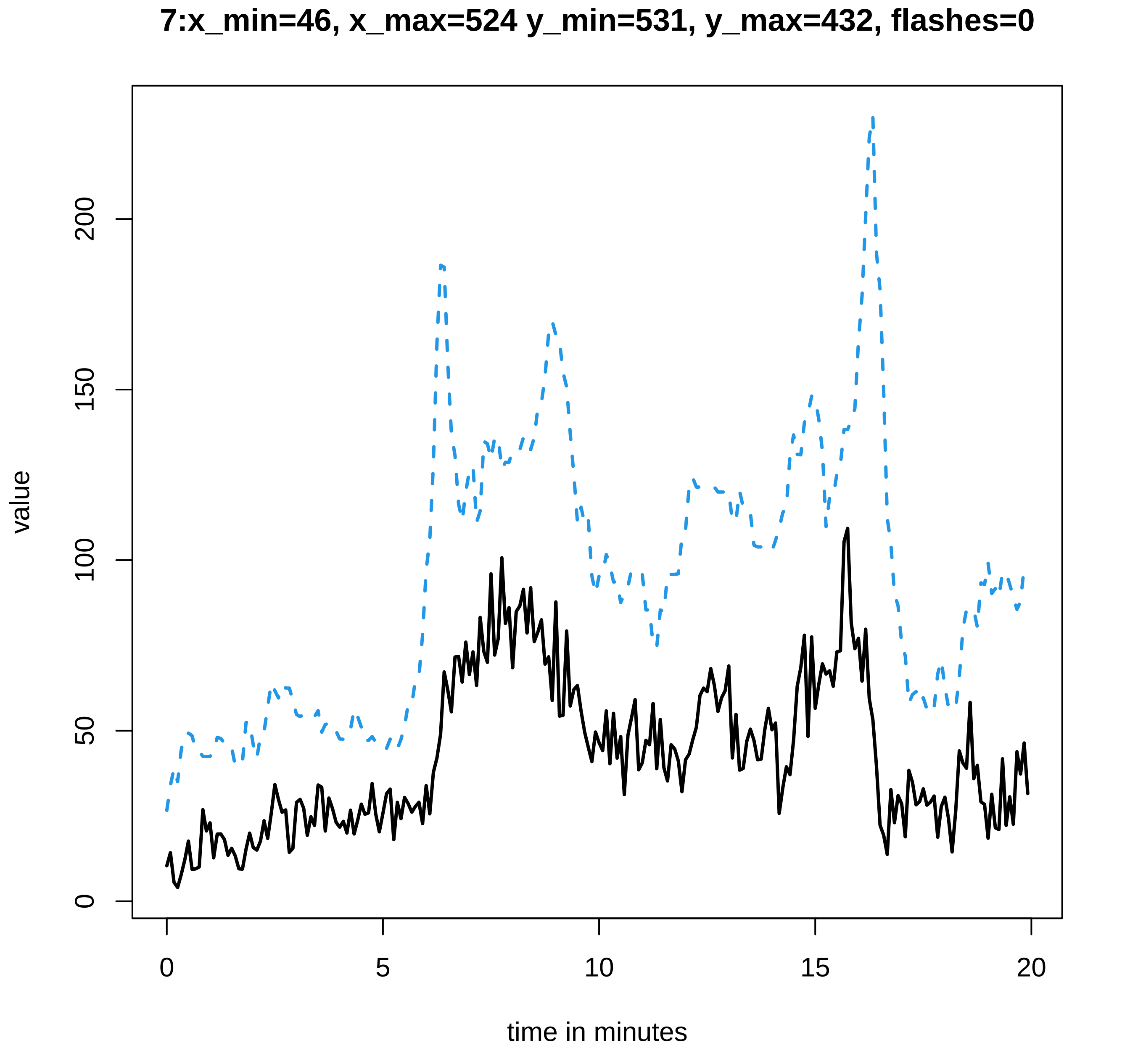
7:x_min=46, x_max=524 y_min=531, y_max=432, flashes=0
value
0
50
100
150
200
0
5
10
15
20
time in minutes

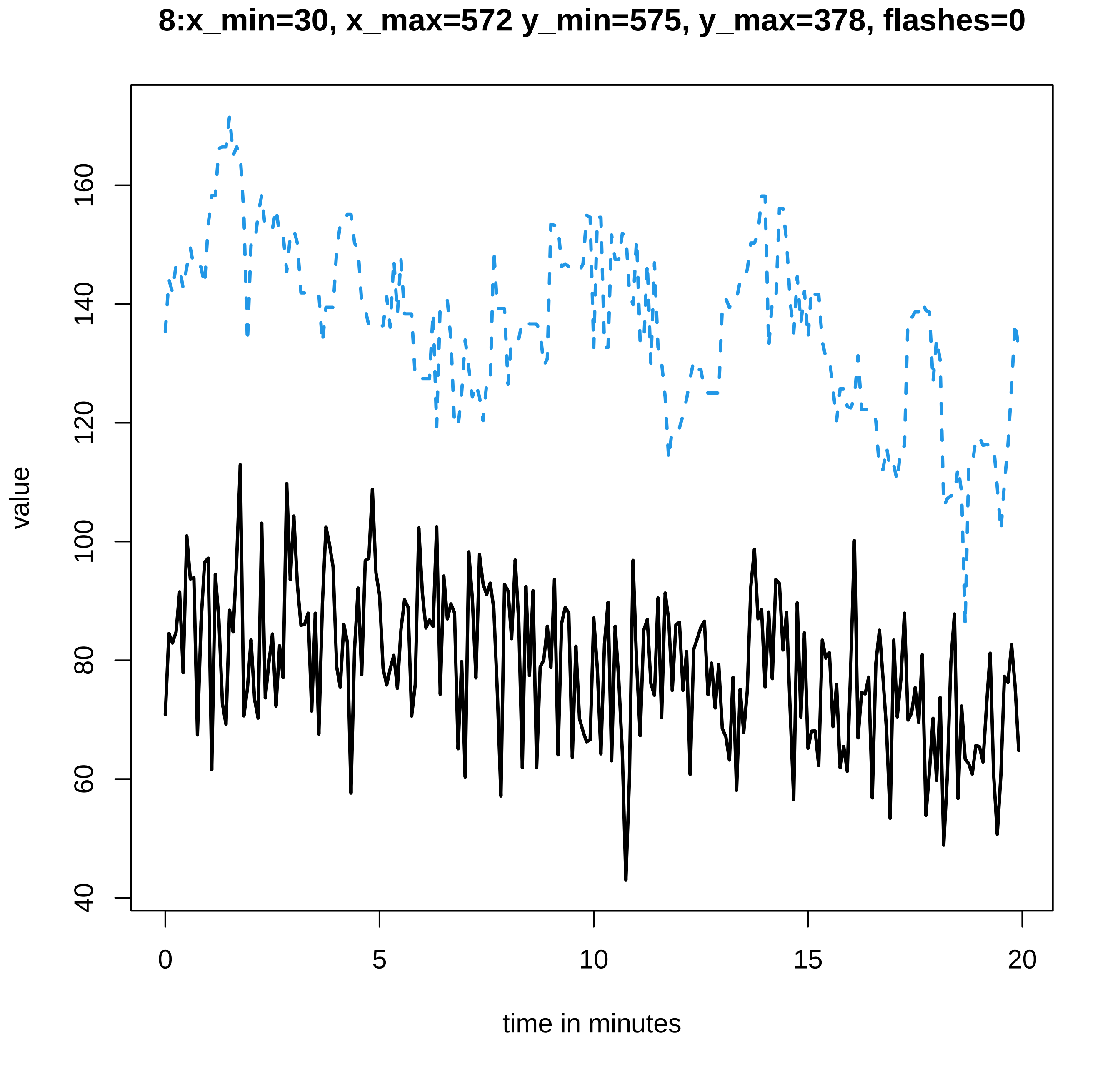
8:x_min=30, x_max=572 y_min=575, y_max=378, flashes=0
value
time in minutes
0
5
10
15
20
40
60
80
100
120
140
160

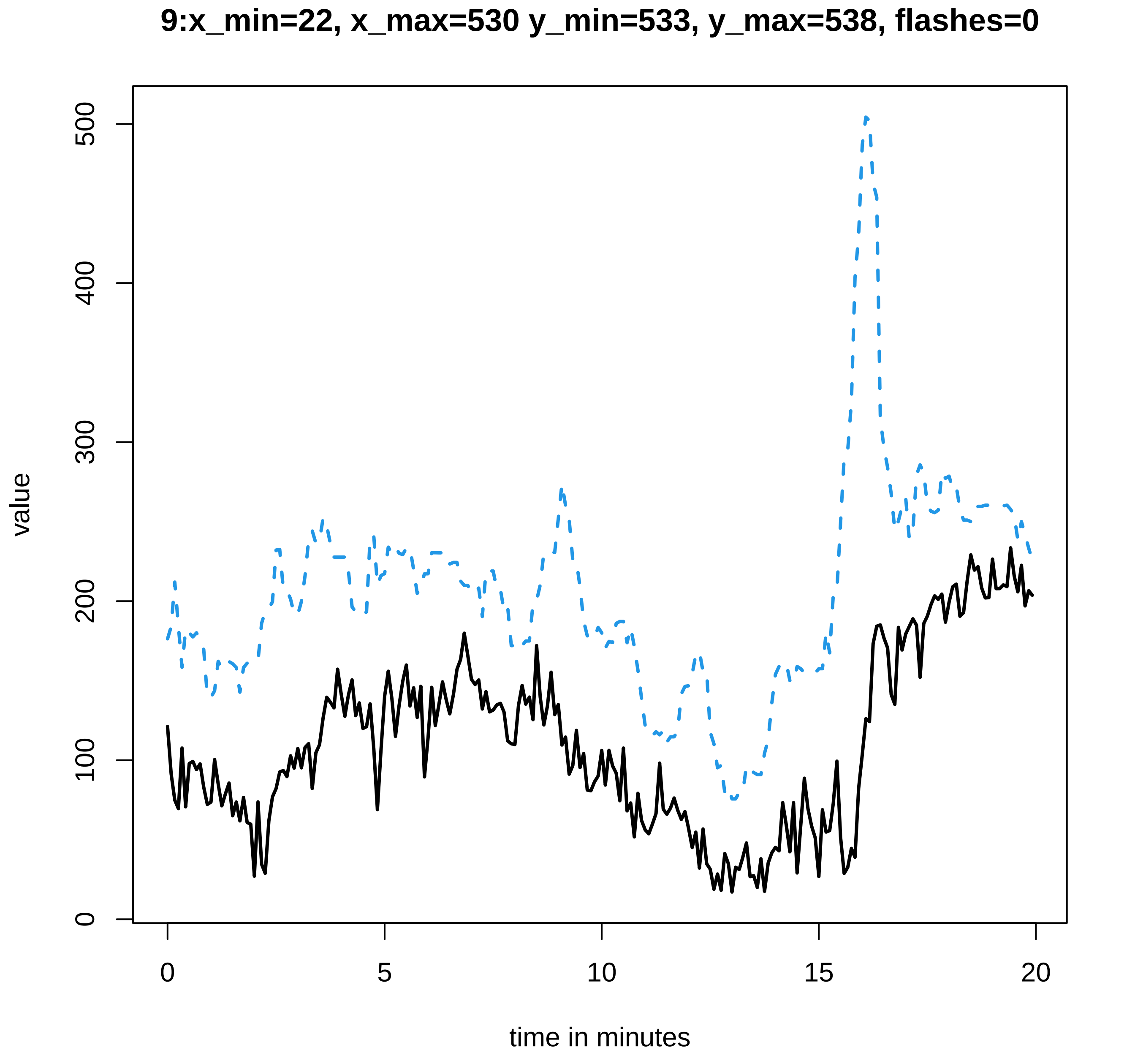
9:x_min=22, x_max=530 y_min=533, y_max=538, flashes=0
value
0
100
200
300
400
500
0
5
10
15
20
time in minutes

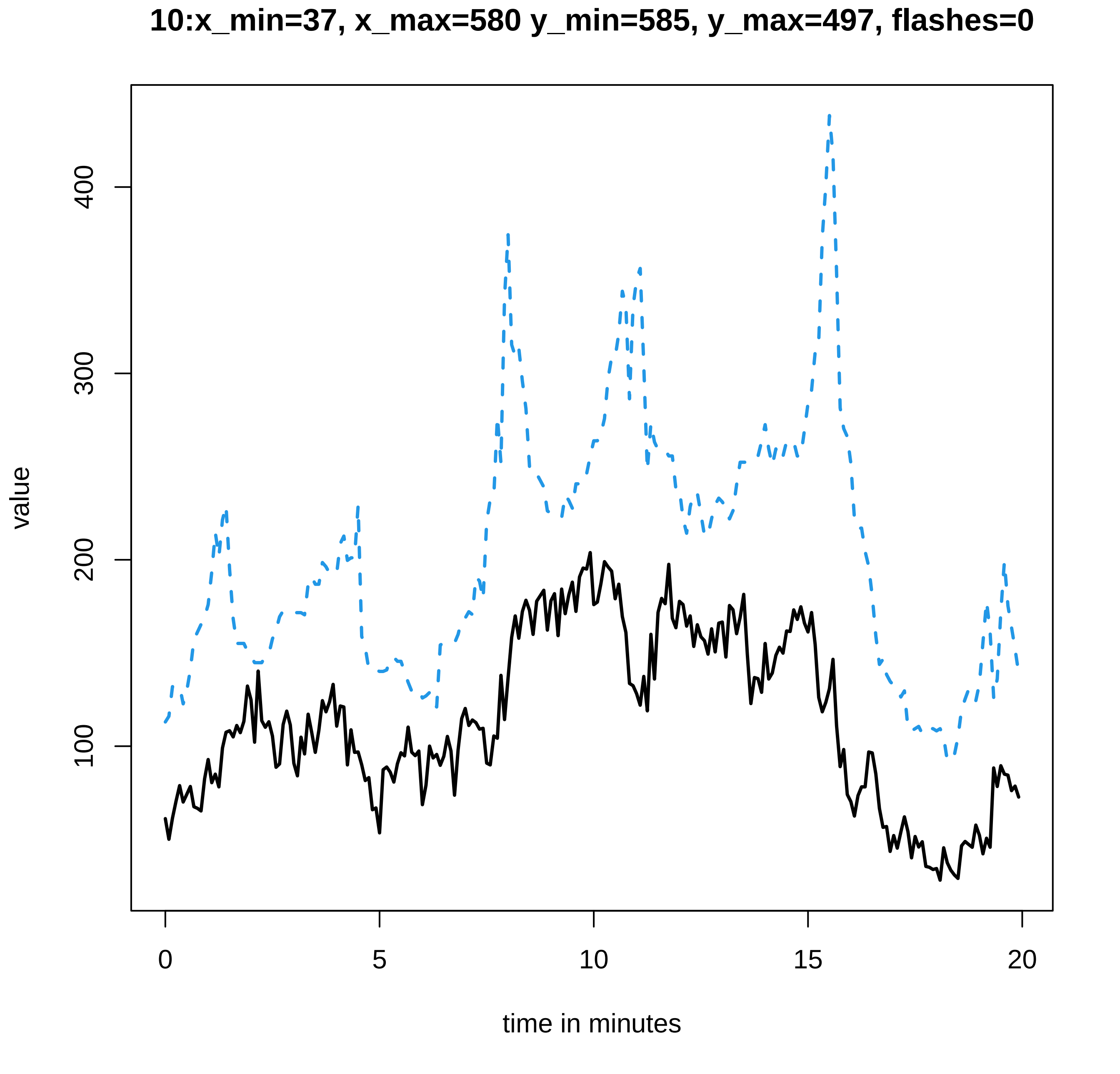
10:x_min=37, x_max=580 y_min=585, y_max=497, flashes=0
value
100
200
300
400
0
5
10
15
20
time in minutes

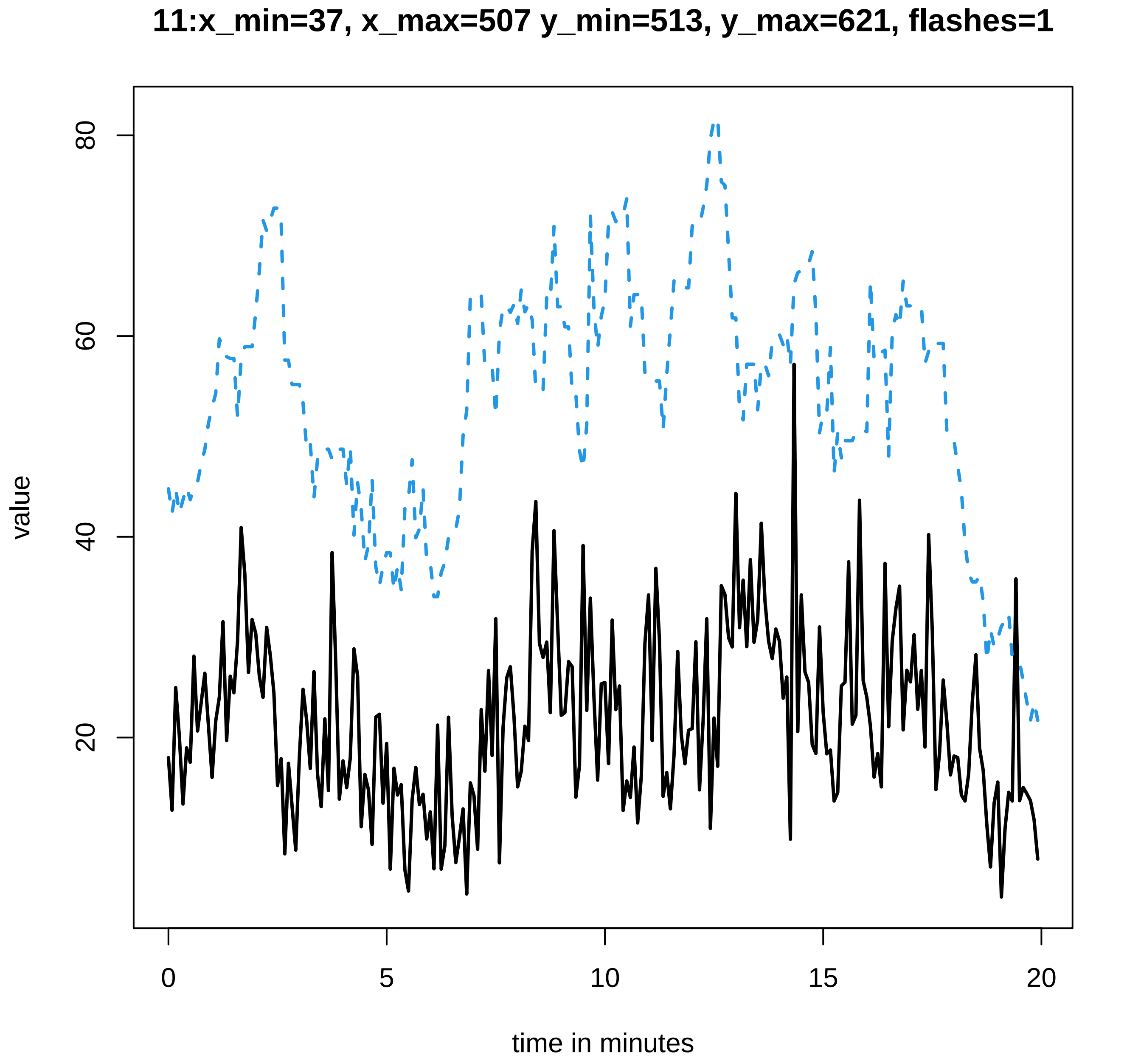

11:x_min=37, x_max=507 y_min=513, y_max=621, flashes=1
value
80
60
40
20
0
5
10
15
20
time in minutes

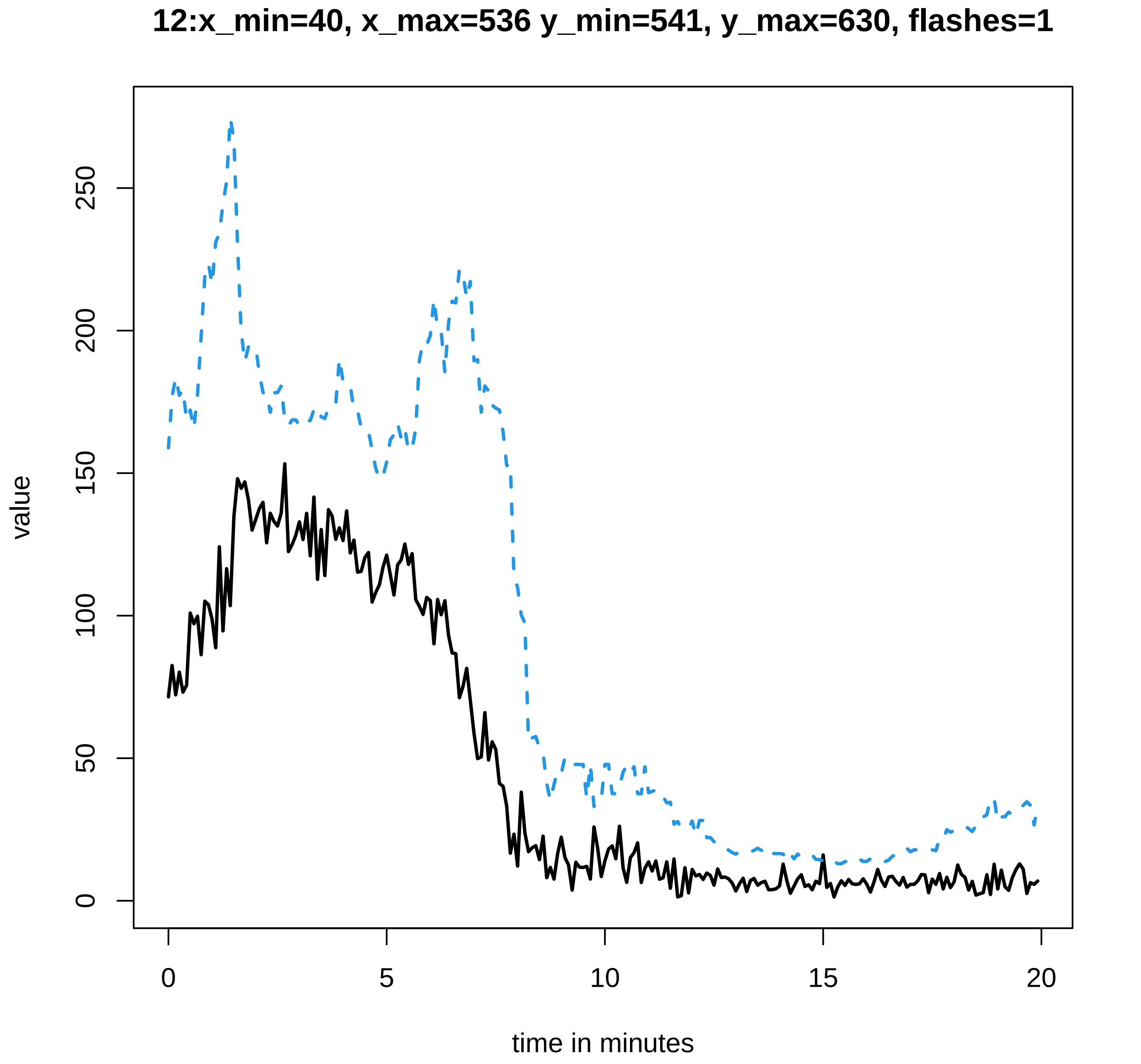
12:x_min=40, x_max=536 y_min=541, y_max=630, flashes=1
value
0
50
100
150
200
250
0
5
10
15
20
time in minutes

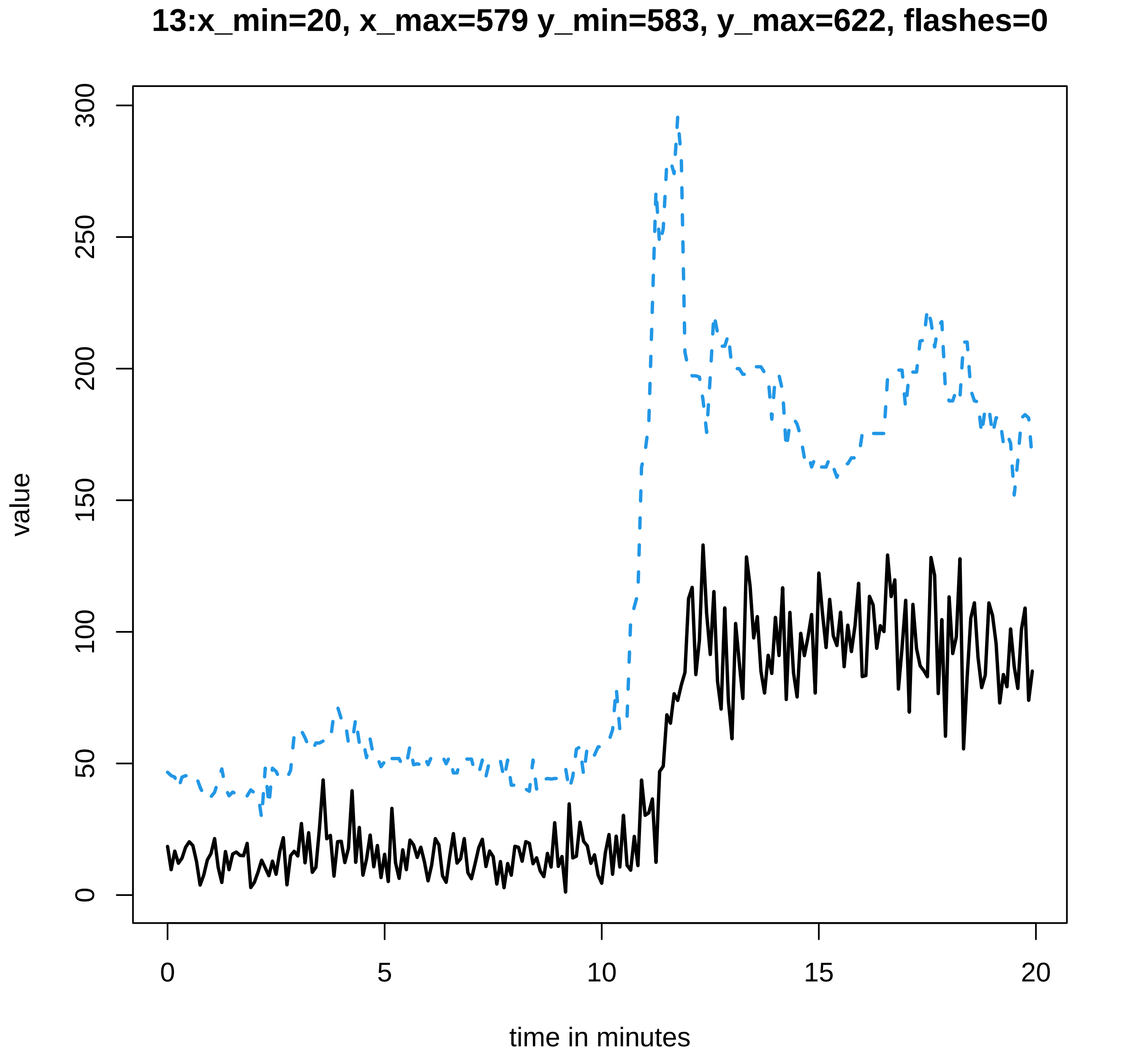
13:x_min=20, x_max=579 y_min=583, y_max=622, flashes=0
value
0
50
100
150
200
250
300
0
5
10
15
20
time in minutes

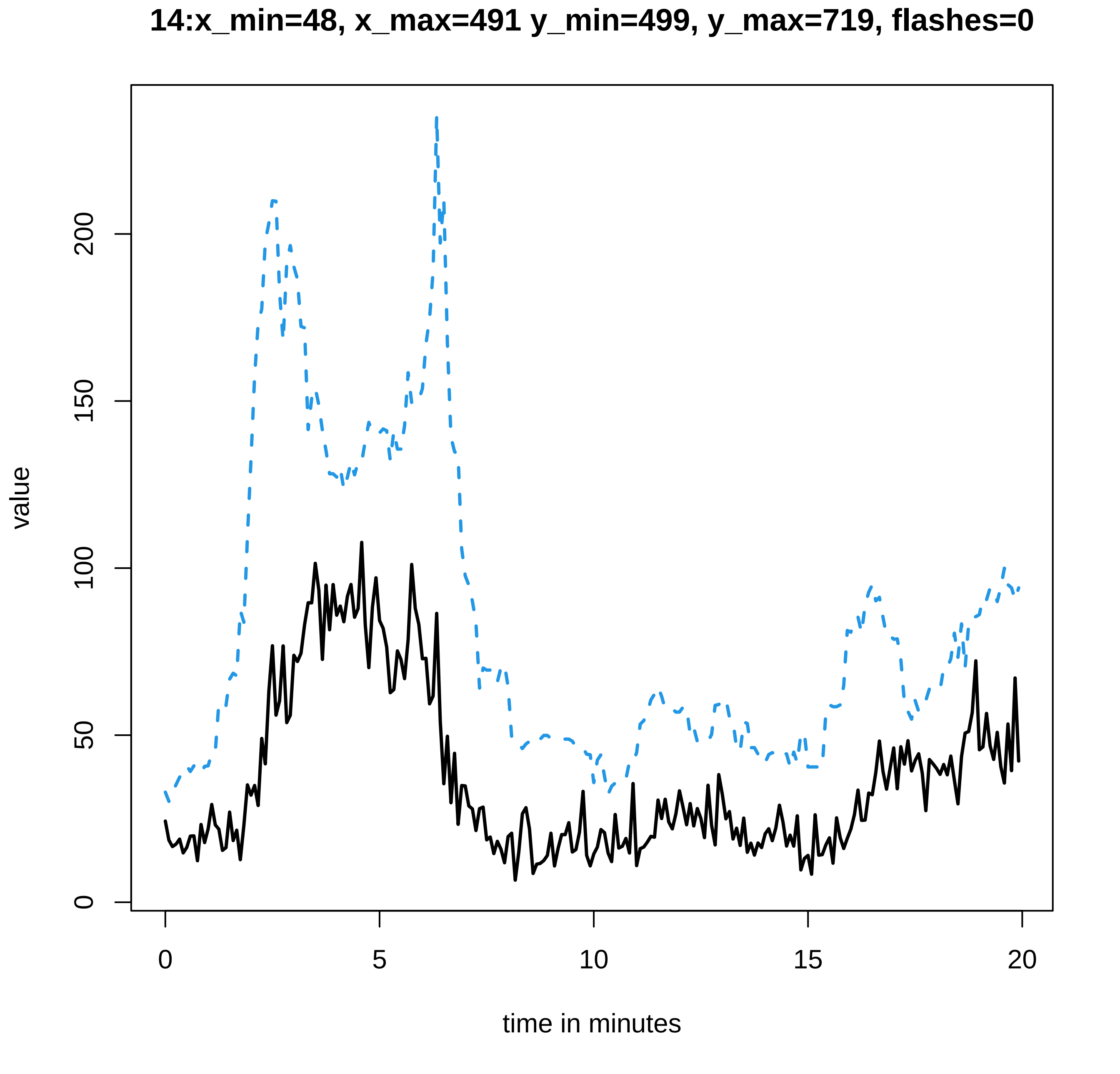
14:x_min=48, x_max=491 y_min=499, y_max=719, flashes=0
value
0
50
100
150
200
0
5
10
15
20
time in minutes

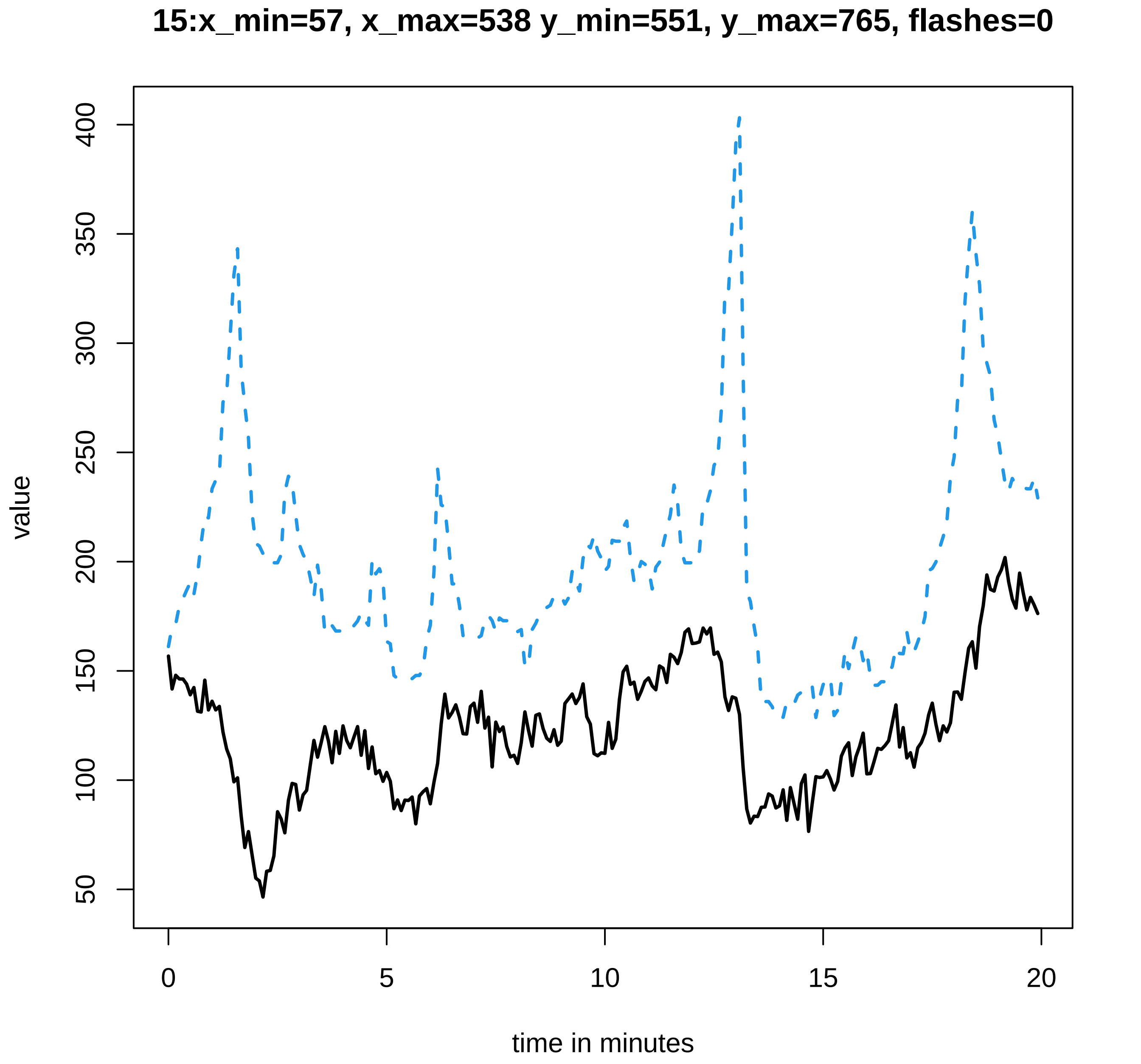

15:x_min=57, x_max=538 y_min=551, y_max=765, flashes=0
value
time in minutes
50
100
150
200
250
300
350
400
0
5
10
15
20

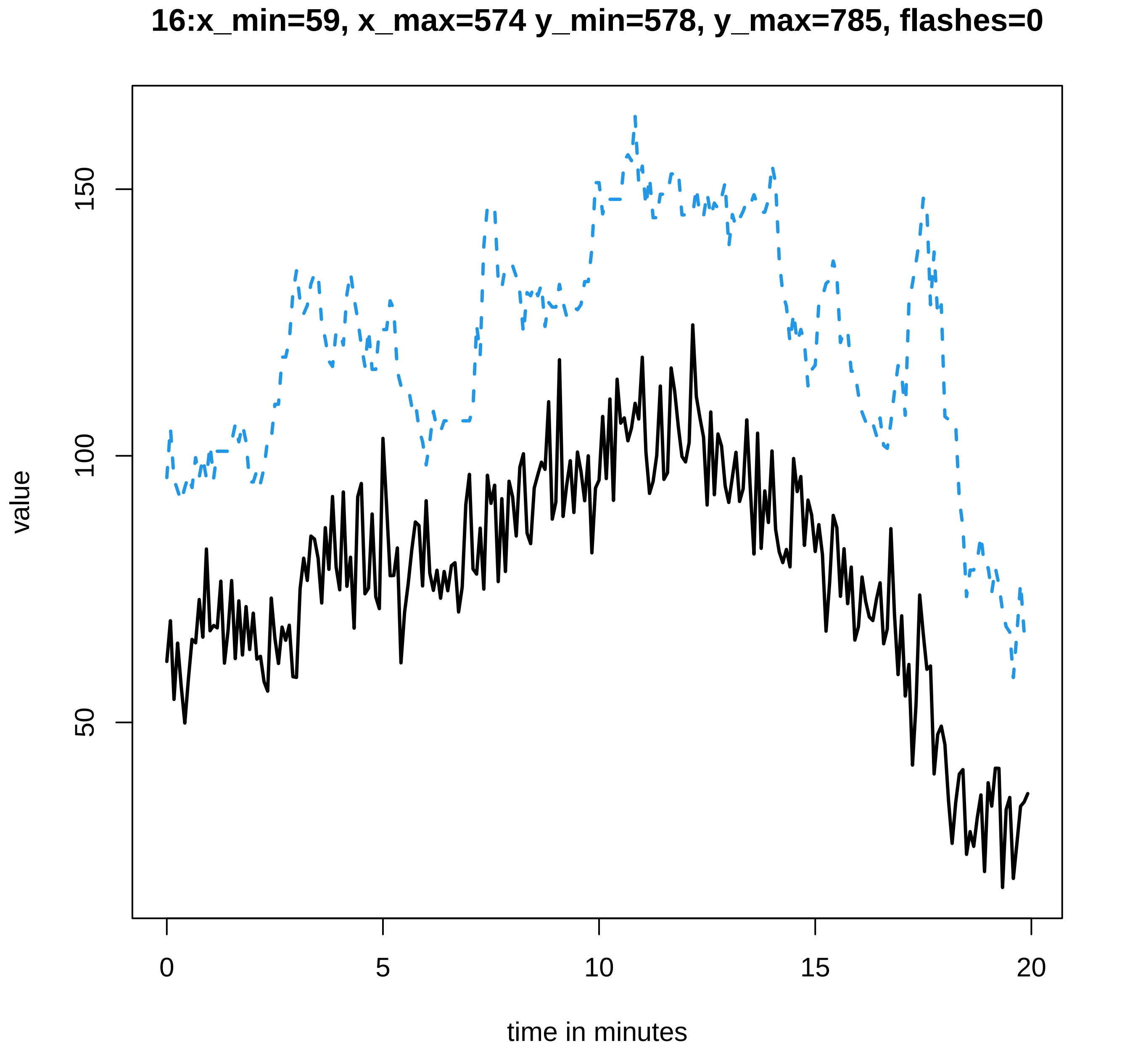
16:x_min=59, x_max=574 y_min=578, y_max=785, flashes=0
value
50
100
150
0
5
10
15
20
time in minutes

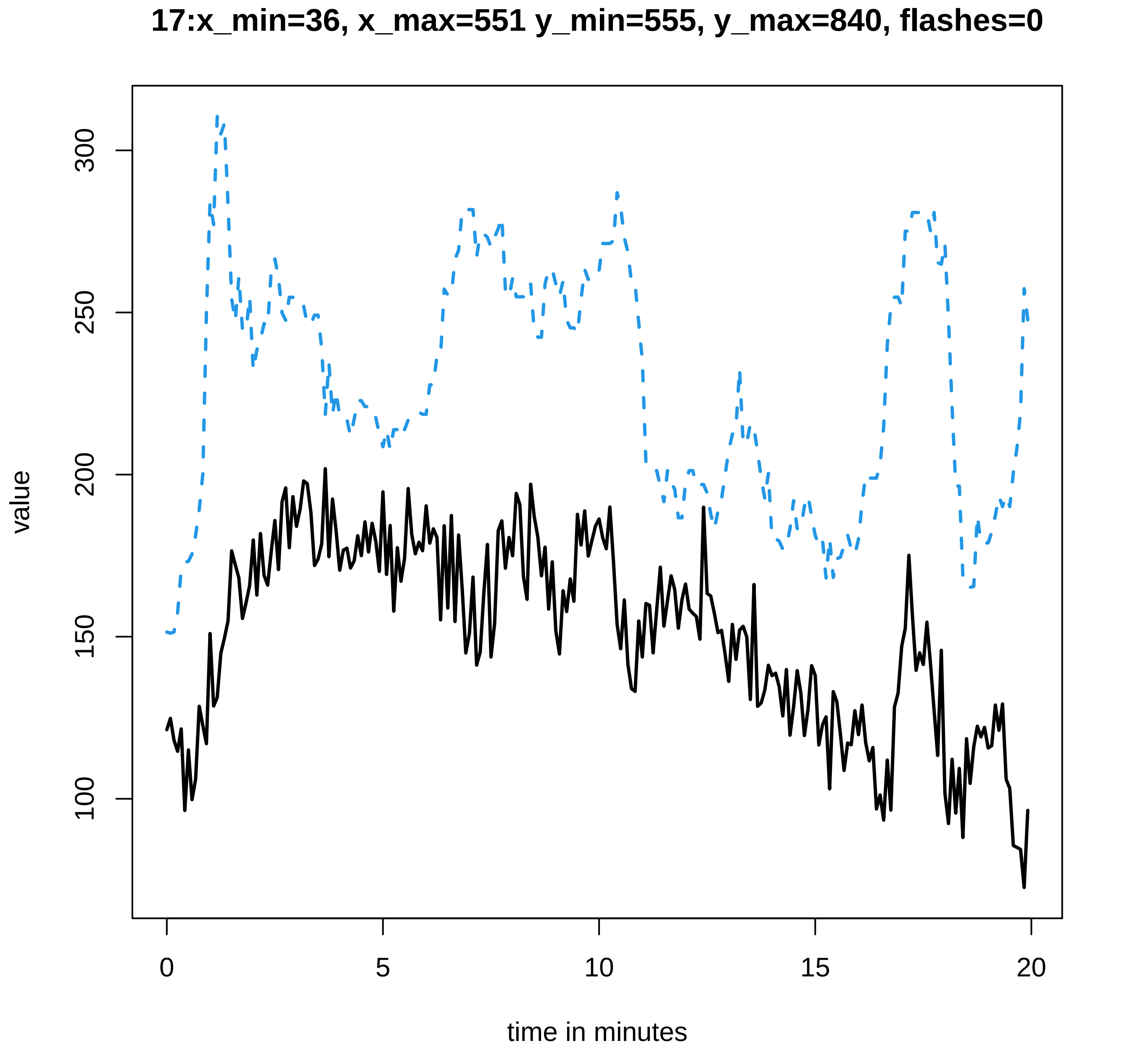
17:x_min=36, x_max=551 y_min=555, y_max=840, flashes=0
value
300
250
200
150
100
0
5
10
15
20
time in minutes

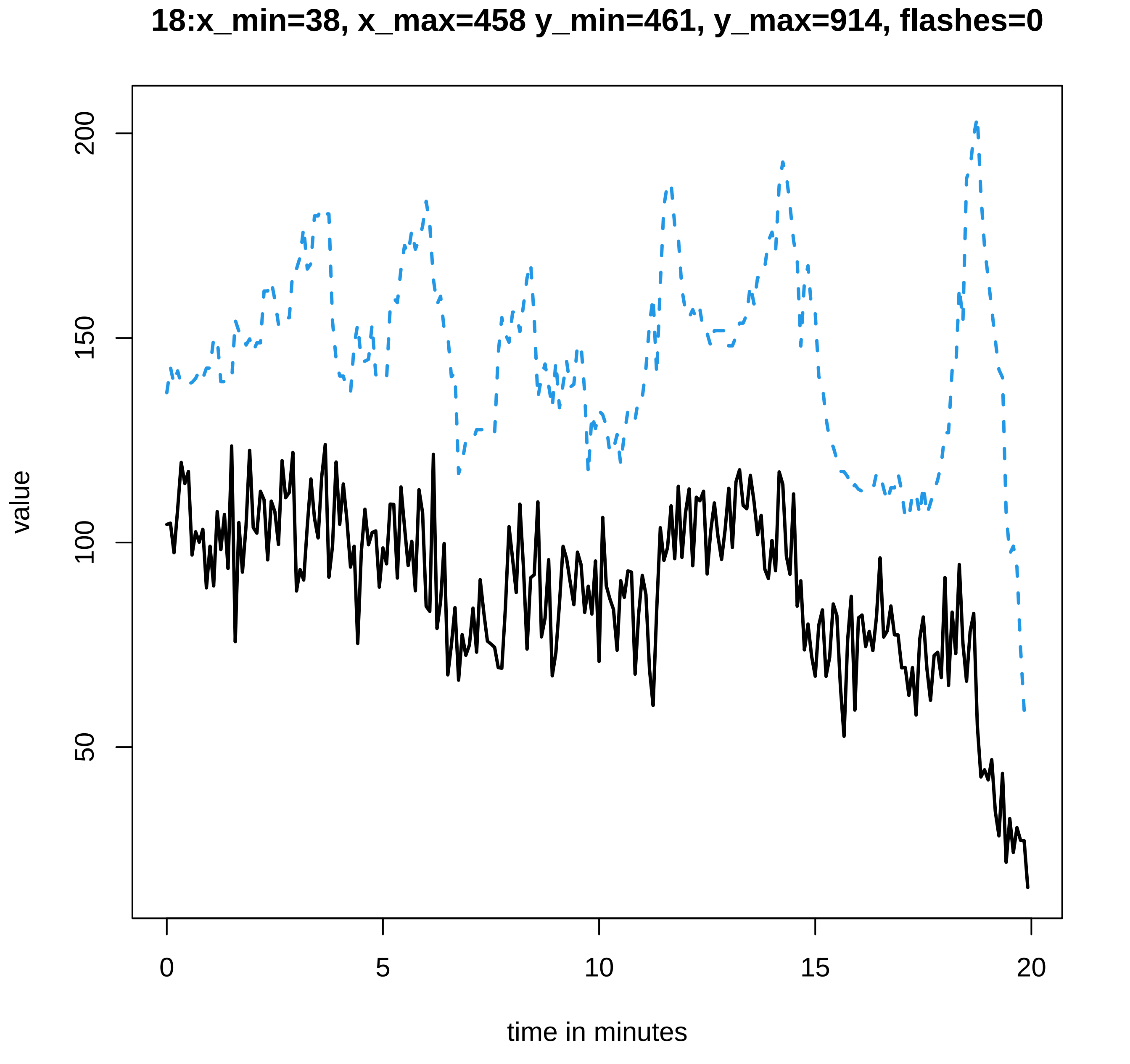

18:x_min=38, x_max=458 y_min=461, y_max=914, flashes=0
value
200
150
100
50
0
5
10
15
20
time in minutes

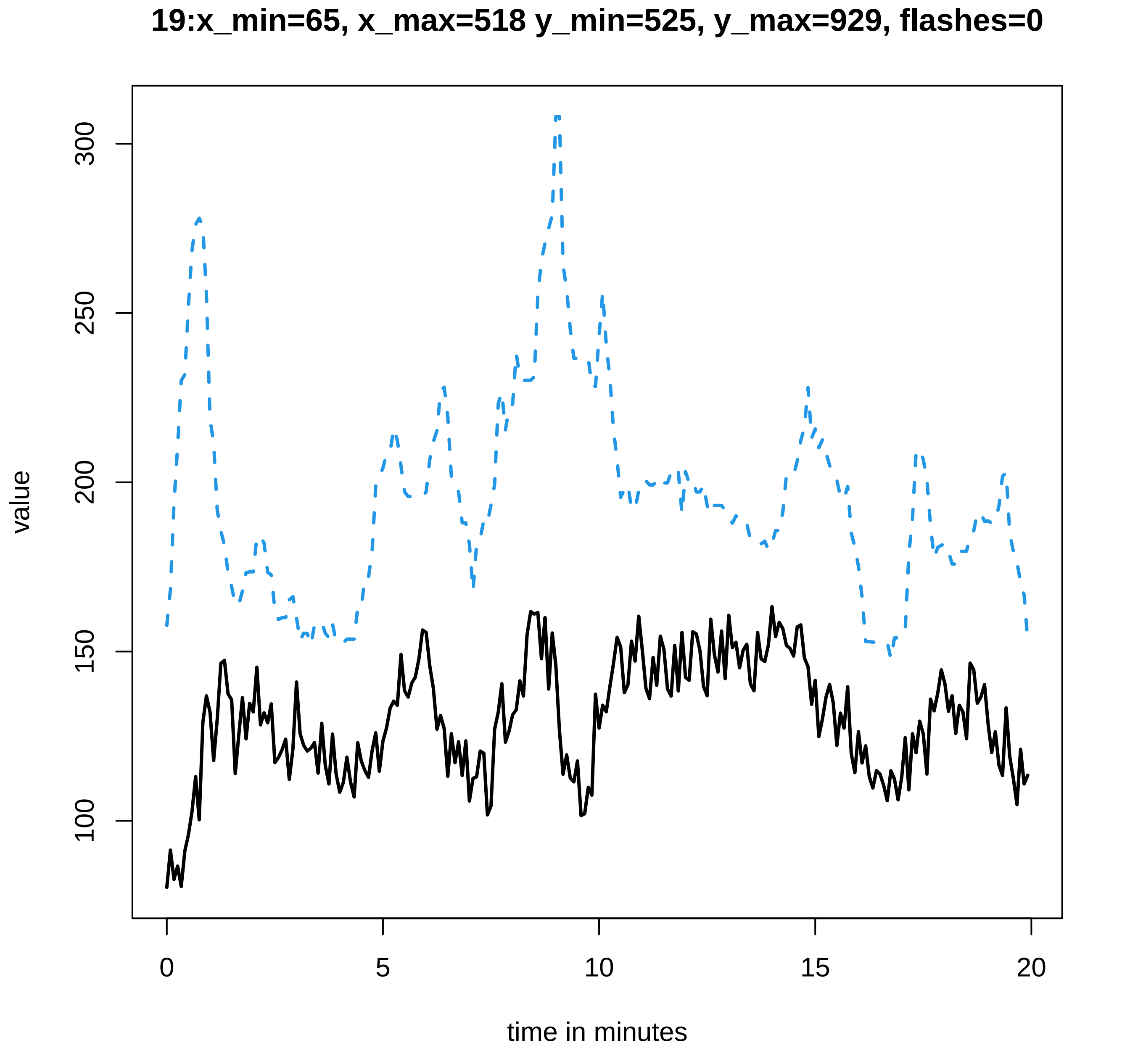

19:x_min=65, x_max=518 y_min=525, y_max=929, flashes=0
value
300
250
200
150
100
0
5
10
15
20
time in minutes

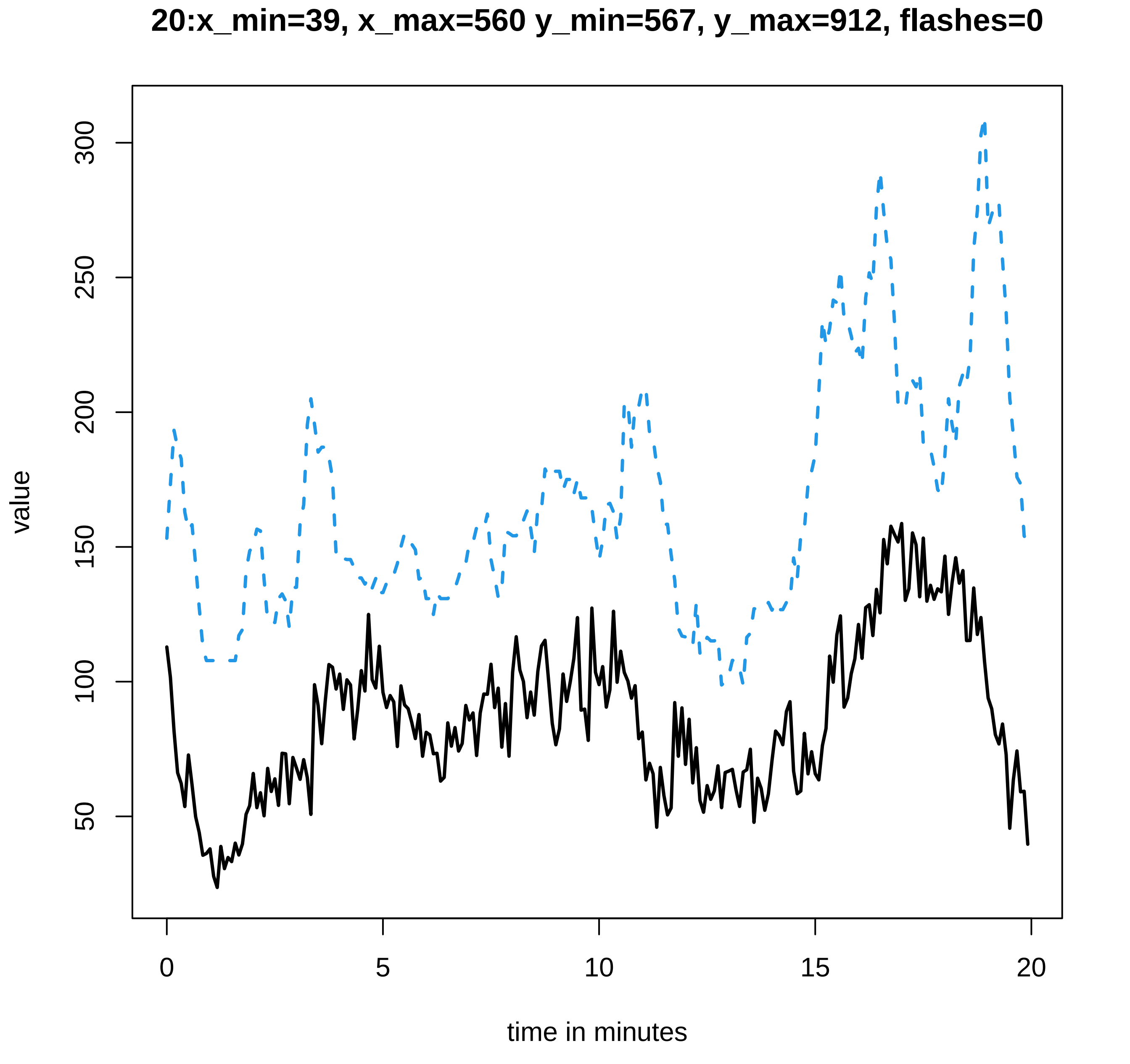
20:x_min=39, x_max=560 y_min=567, y_max=912, flashes=0
value
300
250
200
150
100
50
0
5
10
15
20
time in minutes

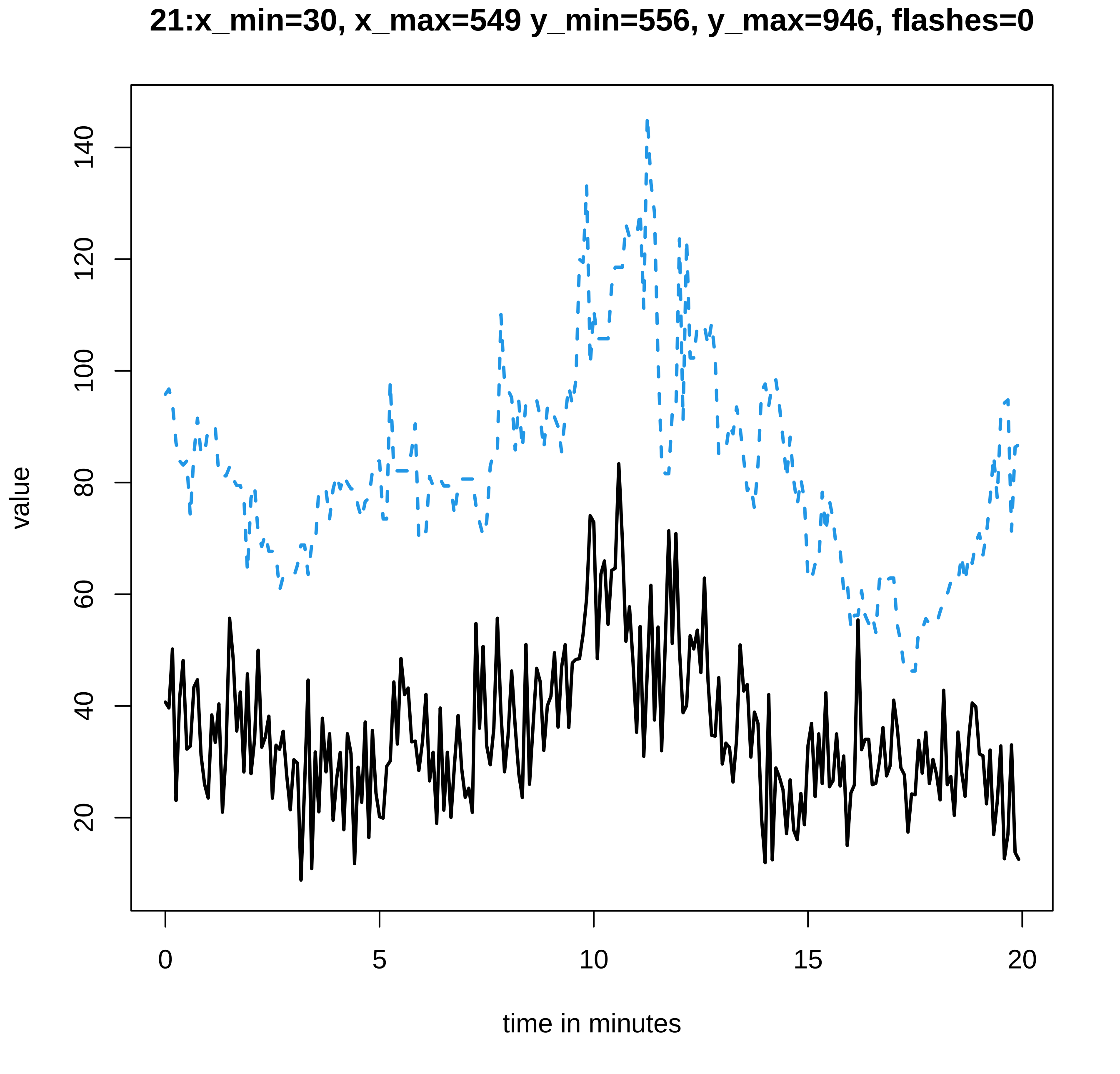
21:x_min=30, x_max=549 y_min=556, y_max=946, flashes=0
value
20
40
60
80
100
120
140
0
5
10
15
20
time in minutes

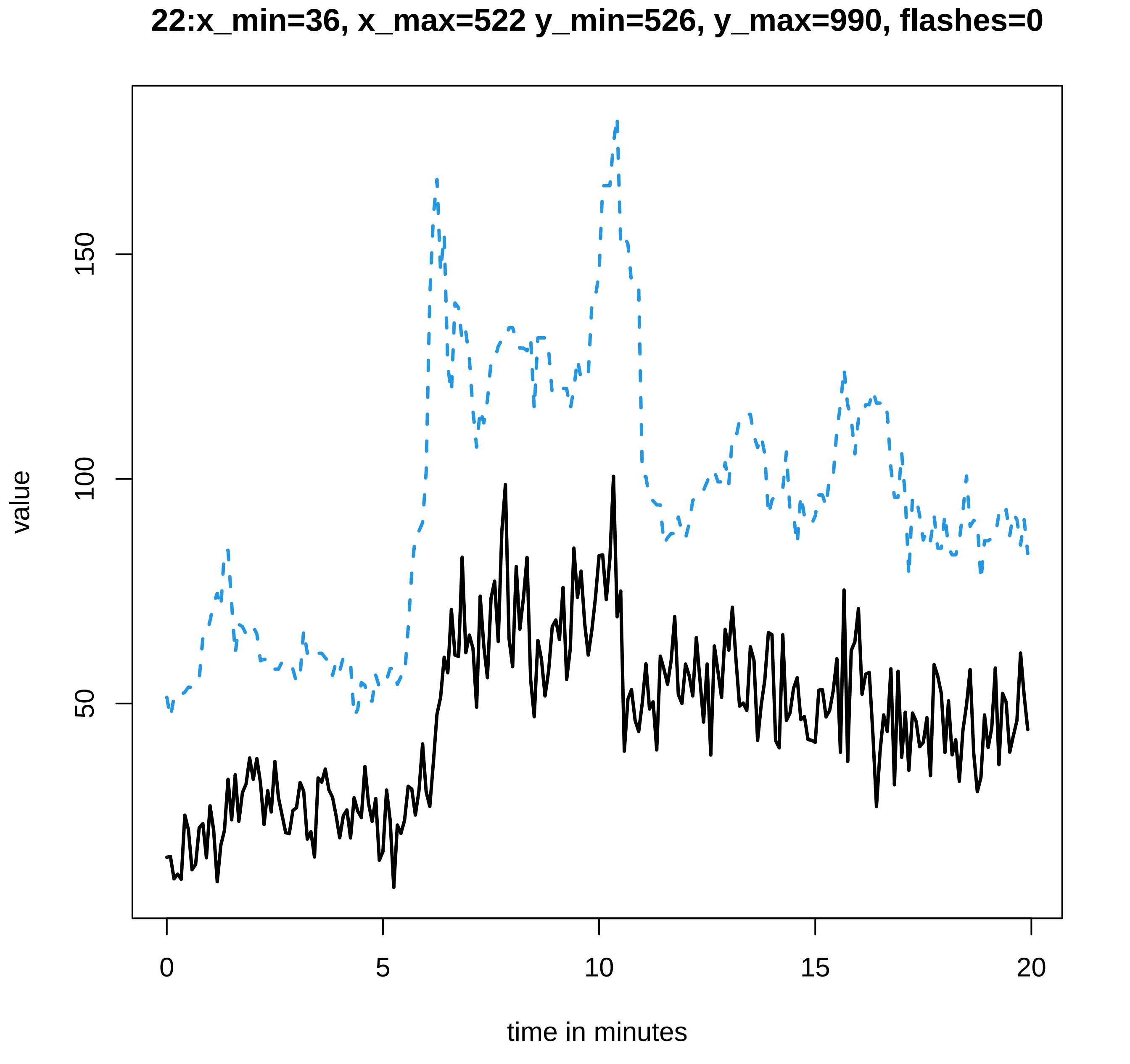

22:x_min=36, x_max=522 y_min=526, y_max=990, flashes=0
value
50
100
150
0
5
10
15
20
time in minutes
