## Supplementary figures and images for "LysM Receptor Proteins are Required for Ectomycorrhizal Symbiosis in Poplar"

### combined-summary_new.txt.pdf

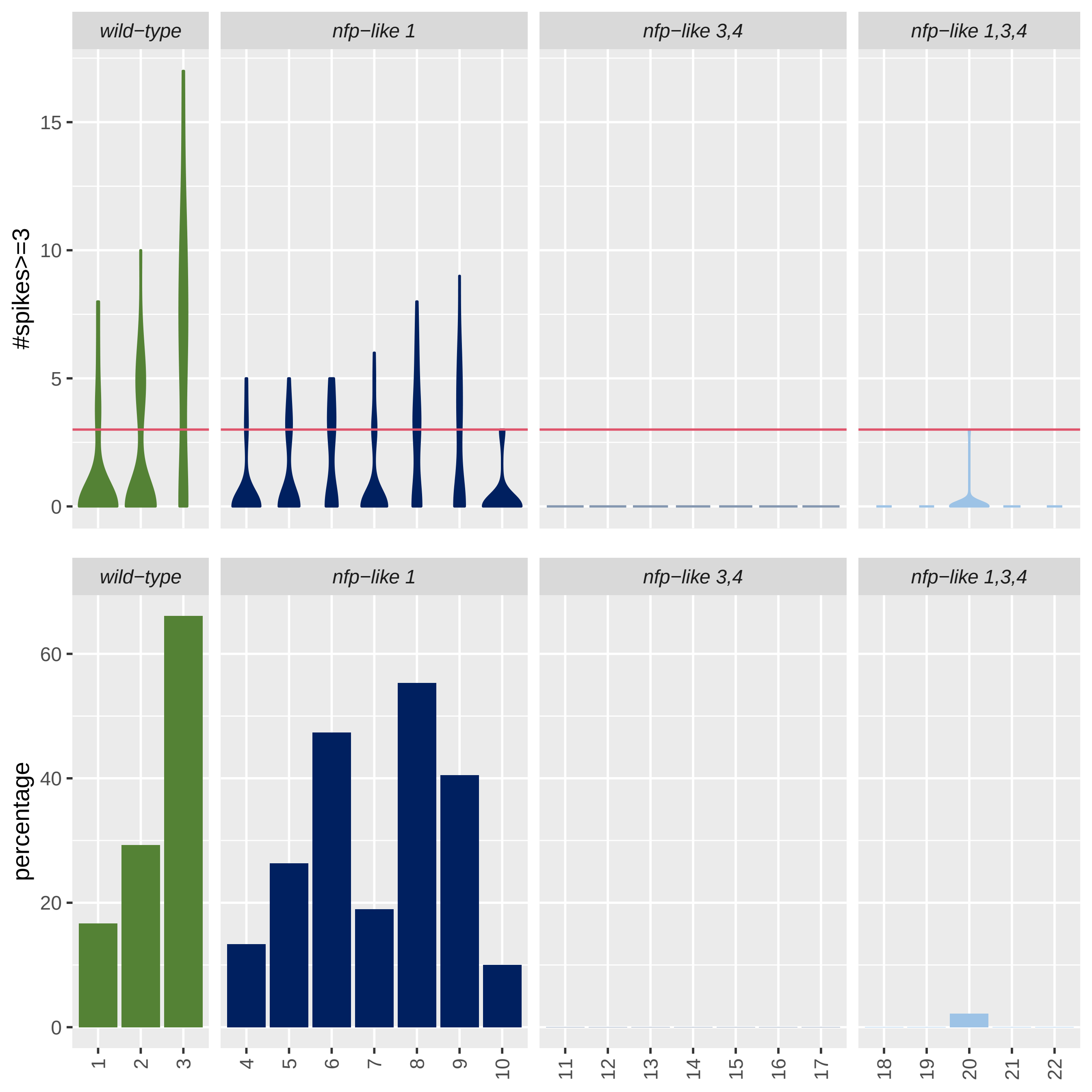

### overview-summary_new.txt.pdf

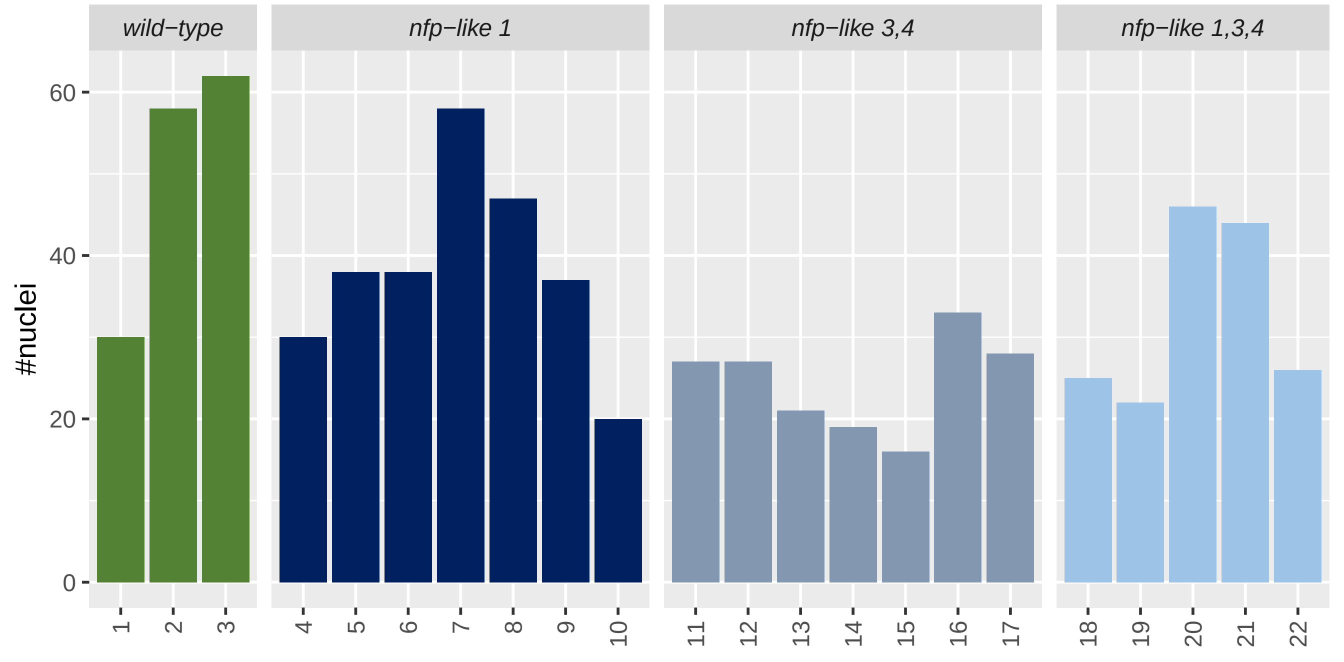

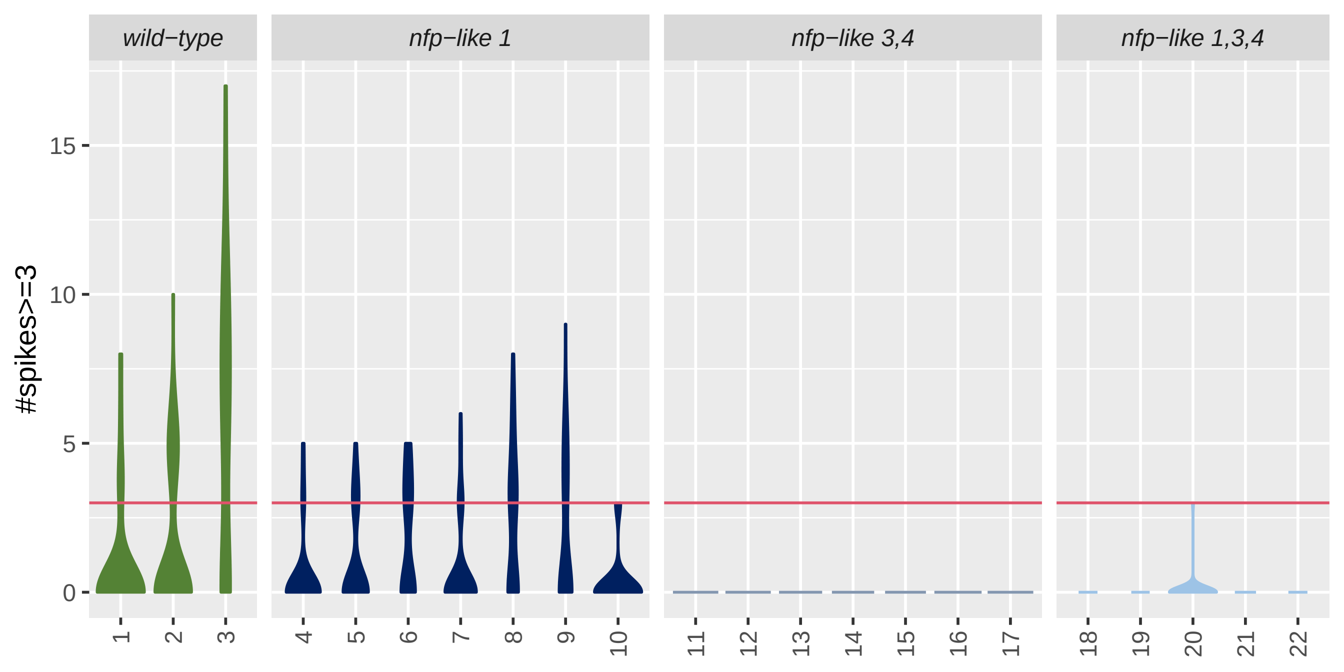

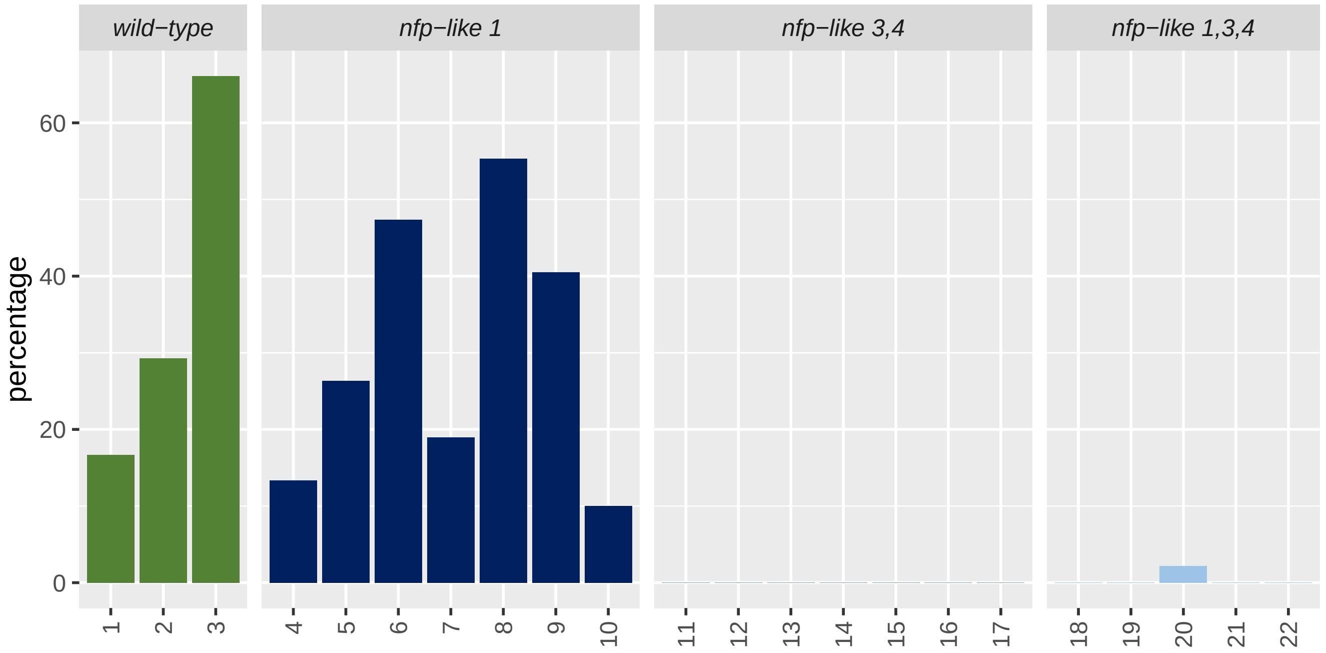
