## Supplementary data qRT-PCR & Primers for "LysM Receptor Proteins are Required for Ectomycorrhizal Symbiosis in Poplar": Supplementary information Primer.pdf

Primers used in this study:

| Nr. | Name | Sequence (5' to 3') | Purpose |
|---|---|---|---|
| 1 | Act2_RT_Fwd | CCCATTGAGCACGTTATTGT | qPCR |
| 2 | Act2_RT_Rev | TACGACCACTGGCATACAGG | qPCR |
| 3 | EF1B_RT-Fwd | CTAACCGCCTTCTCCAACAC | qPCR |
| 4 | EF1B_RT-Rev | AAGAGGACAAGAAGGCAGCA | qPCR |
| 5 | PtNIN2b_RT_Fwd | CAGGAGGAGTGACAGTTCAAAG | qPCR |
| 6 | PtNIN2b_RT_Rev | TCCTACTACACGGAGAGGAATG | qPCR |
| 7 | SS1_Tre_Alba_RT_Fwd1 | CAAGCCCCGAGCAATCTCTTTC | qPCR |
| 8 | SS1_Tre_Alba_RT_Rev1 | CCTTTCTCAGTGGGCATAGAGG | qPCR |
| 9 | pMAS_Fwd | CAGTGCGCAAGACGTGACGTAAG | genotyping PCR |
| 10 | BASTA_Rev | TGACAGCGACCACGCTCTTGAAG | genotyping PCR |
| 11 | NFPlike3_RT_Fwd | CATATACCATCGAGGCAGGCA | qPCR |
| 12 | NFPlike3_RT_Rev | TCGGGGATAAGTGTAGGGTTGAA | qPCR |
| 13 | NFP-like1_2845_Fwd | AAT TTC TTG GAC CTC GGA AA | qPCR |
| 14 | NFP-like1_2846_Rev | AGA CTC CAG GTT GCT TGC AG | qPCR |
| 15 | NFP-like4_2839_Fwd | CCG TAT AAT GCC ATT TTG TAC C | qPCR |
| 16 | NFP-like4_2840_Rev | CAT TAT AGG GTT CCA AAT CCA GA | qPCR |
| 17 | NFP3_C_for | AACAGGTCTCAGGCTCAATGAGTCCCGCATCCCGTTTAG | Cloning NFP-like3 |
| 18 | NFP3_C_rev | AACAGGTCTCACTGATCTTGCCATAACCTGAGGG | Cloning NFP-like3 |
| 19 | NFP4_C_for | AACAGGTCTCAGGCTCAATGACAGCCAAATCCCATC | Cloning NFP-like4 |
| 20 | NFP4_C_rev | AACAGGTCTCACTGATCTTGCCATTACCTGAGG | Cloning NFP-like4 |
